# Systematic histone mutagenesis reveals nucleosome-dependent maintenance of three-dimensional chromosome architecture and virulence in *Cryptococcus neoformans*

**DOI:** 10.64898/2026.05.19.726417

**Authors:** Sunhak Kwon, Tu Minh Khuong, Yu-Byeong Jang, Seong-Ryong Yu, Eui-Seong Kim, Sangyong Lim, Jong-Hyun Jung, Kyung-Tae Lee, Yong-Sun Bahn, Kwang-Woo Jung

**Author notes:** Correspondence to Y.-S. Bahn and K.-W. Jung.

## Abstract

**Background:** Genomic stability is maintained through the coordinated regulation of DNA repair, dNTP pool balance, and histone dynamics—the three pillars of the DNA damage response. Because histones constitute the fundamental 3D physical scaffold of the genome, their precise regulation is essential for the spatial organization that dictates environmental fitness. In the radiotolerant pathogen *Cryptococcus neoformans*, the Rad53-Bdr1 pathway is a central DDR mediator; however, the mechanisms linking this checkpoint to histone dynamics remain poorly understood. Because conventional one-dimensional analyses cannot capture how spatial chromatin folding shapes transcriptional reprogramming, we integrated high-throughput chromosome conformation capture (Hi-C) with transcriptomic profiling to address this gap.

**Results:** We demonstrate that *HTA1* and *HTB1*, encoding H2A and H2B, are essential for viability, whereas H3 and H4 paralogs exhibit functional redundancy. Although most core histones are regulated by Rad53, *HHT1* and variant *HTZ1* are expressed independently of the Rad53. Notably, loss of the H3 paralog *HHT2* induces growth defects under diverse stress conditions. Integrated RNA sequencing and Hi-C analyses reveal that *HHT2* deletion drives transcriptional reprogramming of stress-responsive genes, coinciding with large-scale chromatin rearrangements such as A/B compartment switching and topologically associating domain boundary shifts. Furthermore, *HHT2* loss impairs virulence factor formation and attenuates virulence.

**Conclusion:** Our findings identify core histones as essential architects of the 3D genome in *C. neoformans*. By establishing a causal link between chromatin structural collapse and transcriptional reprogramming, this study highlights 3D genome architecture as a decisive physical switch linking nucleosome-level dynamics to global transcriptional programs required for environmental survival.

## Background

In eukaryotes, genomic DNA is packaged into chromatin to enable both compaction and dynamic regulation of genome function. The fundamental chromatin subunit is the nucleosome, consisting of approximately 147 base pairs of DNA wrapped around an octamer of histone proteins: two copies each of H2A, H2B, H3, and H4 [1]. These core histones form an H3-H4 tetramer and two H2A-H2B heterodimers. Additionally, the linker histone H1 binds DNA between nucleosomes, facilitating higher-order chromatin structure formation [2].

In model yeasts, the genetic organization and functional roles of core histone complexes are well characterized. In *Saccharomyces cerevisiae*, each canonical histone is encoded by two gene copies (e.g., *HTA1*/*HTA2*), whereas *Schizosaccharomyces pombe* possesses multiple gene copies for each histone [3, 4]. Due to this redundancy, single histone gene deletions are generally non-lethal [5–7]. Eukaryotes also possess histone variants, such as H2A.Z, which are located outside canonical gene clusters and often expressed independently of the cell cycle [8]. Recent studies in pathogenic fungi, such as *Candida glabrata* and *Candida albicans*, have further demonstrated that histone dosage is pivotal for survival, stress adaptation, and full virulence [9–11]. To maintain functional integrity and respond to physiological cues, histone levels are dynamically regulated through post-translational modifications and S-phase-coupled transcriptional induction, the latter of which is orchestrated by the HIR complex and specific activators such as Spt10 and Spt21 [12].

Historically, characterizing the functional impact of factors that modulate histone architecture, such as histone deacetylases, has predominantly relied on a one-dimensional (1D) framework. These investigations have largely been confined to correlating the presence of specific genes with the linear transcriptional modulation of their downstream targets under stress [13, 14]. However, such localized analyses of histone dynamics fail to capture the broader, genome-wide transcriptional reprogramming driven by the physical clustering of distal genes or large-scale chromatin rearrangements. Because core histones serve as the fundamental physical scaffolds of the genome, their depletion or imbalance can trigger a global ‘structural collapse’ that transcends individual promoter-level control. This inherent limitation of 1D methodologies in capturing histone-dependent structural changes underscores the necessity for a multi-dimensional, systems-level approach to understand how the spatial organization of the genome dictates functional output and environmental fitness.

To overcome these limitations, recent studies have increasingly employed spatial genomics, utilizing high-throughput chromosome conformation capture (Hi-C) to reveal that 3D genome architecture—organized into hierarchical structures such as A/B compartments and topologically associating domains (TADs)—is a critical determinant of global transcriptional programs [15–17]. Hi-C analysis identified unique 3D chromatin conformations, termed “jet-like” structures, within biosynthetic gene cluster regions in *Fusarium graminearum*, where the histone acetyltransferase Gcn5 is required for the maintenance of these structures [18]. In addition, transcriptional regulation mediated by condensin, a central factor in chromosome condensation and segregation, has been characterized in yeast [19]. Recently, Polisetty et al. reported that cell cycle-dependent centromere clustering and declustering serve as fundamental drivers for global reorganization of 3D genome architecture and chromosomal interaction in *C. neoformans* [20]. However, the contribution of core histones, the fundamental building blocks of chromatin, to maintaining this spatial organization remains poorly understood. It remains unclear whether histone dosage imbalances drive transcriptional reprogramming through direct structural collapse or through indirect signaling pathways.

A comprehensive understanding of the DNA damage response (DDR) requires the elucidation of three interconnected pillars: canonical DNA repair pathways, intracellular dNTP pool regulation, and histone dynamics. Although our group has previously characterized the former two mechanisms in *Cryptococcus neoformans*, a basidiomycetous pathogen that causes fatal meningoencephalitis and exhibits remarkable radiation resistance [21, 22], the specific contribution of histone regulation under genotoxic stress has remained elusive. In this study, we systematically characterize the transcriptional and structural landscape of core histones and their variants in *C. neoformans*. Beyond characterizing their essentiality and roles in the DDR, we performed an integrated analysis of histone function and gene expression by mapping 3D physical chromosomal interactions via Hi-C in the absence of the H3 paralog *HHT2*. By integrating Hi-C and RNA-seq data, we demonstrate that structural disruption—including compartment switching and TAD boundary shifts—drives transcriptional reprogramming required for environmental stress and virulence. Collectively, this study establishes core histones as essential architects of the 3D genome and links nucleosome-level dynamics to physiological adaptation and pathogenesis in *C. neoformans*.

## Results

### *C. neoformans* core histones exhibit distinctive genomic organization with conserved structural architectures

Although the fundamental architecture of the nucleosome comprising an octamer of two copies each of histones H2A, H2B, H3, and H4 is highly conserved across eukaryotes, the genomic copy numbers of the genes encoding these core histone proteins diverge significantly among organisms. In humans, large histone gene clusters facilitate the massive synthesis of histones required for genome packaging. Conversely, yeast genomes feature a more compact and minimalist histone gene organization [23]. Specifically, while *S. cerevisiae* maintains a symmetrical system with two copies for each core histone gene, *C. neoformans* exhibits an asymmetric organization with reduced gene copy. In *C. neoformans*, histones H3 (*HHT1*, CNAG_04828; *HHT2*, CNAG_06745) and H4 (*HHF1*, CNAG_01648; *HHF2*, CNAG_07807) are encoded by two gene copies each, whereas H2A and H2B are uniquely represented by single-copy genes, *HTA1* (CNAG_06747) and *HTB1* (CNAG_06746), respectively (Fig. 1A). To complement these canonical core histones, eukaryotes such as *S. cerevisiae* and humans utilize centromere-specific H3 variants (Cse4 and CENP-A, respectively) and the H2A variant H2A.Z to functionally compensate for their canonical forms [24]. Similarly, *C. neoformans* encodes the conserved histone variants *CSE4* (CNAG_00063) and *HTZ1* (CNAG_05221). To evaluate these findings within an evolutionary context, we performed a phylogenetic analysis across diverse fungal species and metazoans (Fig. 1B). The resulting tree demonstrates that not only the core histones but also their variants, such as H2A.Z and CENP-A, are strikingly conserved across the fungal kingdom. Despite this high degree of sequence conservation, the phylogenetic distribution of these genes revealed that each organism has evolved a distinctive gene copy profile.

**Fig. 1.**
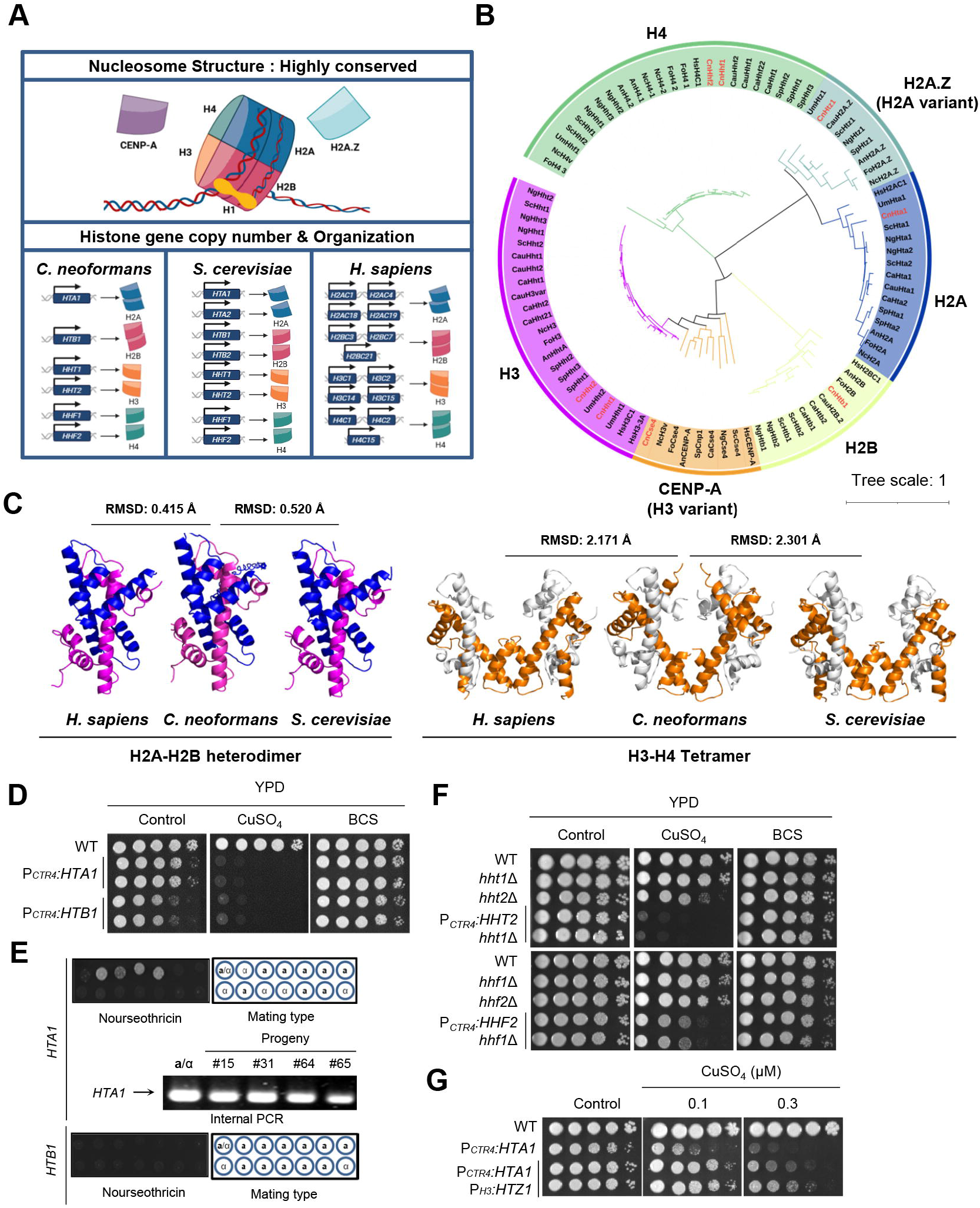
Evolutionary conservation, structural architecture, and essentiality of the *C. neoformans* histone repertoire. **A** Schematic diagram of conserved nucleosome structure and comparison of histone gene copy numbers in *C. neoformans, S. cerevisiae* and *homo sapiens*. **B** The phylogenetic analysis of core histones and histone variants. Neighbor-joining phylogenetic tree inferred from the protein sequences of core histones and histone variants from 11 eukaryotes (Cn: *Cryptococcus neoformans*; Um: *Ustilago maydis*; Sp: *Schizosaccharomyces pombe*; An: *Aspergillus nidulans*; Nc: *Neurospora crassa*; Fo: *Fusarium oxysporum*; Ca: *Candida albicans*; Cg: *Candida glabrata*; Sc: *Saccharomyces cerevisiae*; Hs: *Homo sapiens*; Cau: *Candidozyma auris*). Evolutionary analyses were carried out in MEGA11. Protein sequences were sourced from FungiDB (https://fungidb.org) and NCBI database. **C** Structural comparison of the histone complexes across species. Structures of the H2A-H2B heterodimer (left) and H3-H4 tetramer (right) from *C. neoformans, H. sapiens*, and *S. cerevisiae* were shown. Histone subunits H2A, H2B, H3, and H4 are colored in magenta, blue, orange, and white, respectively. Structural alignments and RMSD calculations were performed using PyMOL. **D** Hta1 and Htb1 were required for growth. The grown strains were serially diluted and spotted onto YPD medium containing CuSO_4_. **E** Basidiospore progeny isolated from the heterozygous *HTA1*/*hta1*Δ and *HTB1*/*htb1*Δ strains were spotted onto nourseothricin medium, followed by genetic analysis of *NAT* progeny (#15, #31, #64, and #65) using internal PCR. **F** The H3 and H4 paralogs exhibited functional redundancy in growth. The grown strains were serially diluted and spotted onto YPD medium containing the BCS (100 μM) or CuSO_4_ (20 μM). **G** Partial complementation of reduced growth through *HTZ1* overexpression in the *P*_*CTR4*_*:HTA1* strain. The grown strains were serially diluted and spotted onto YPD medium containing CuSO_4_.

Given the high degree of evolutionary conservation in histone proteins, we sought to determine whether their tertiary architectures are similarly maintained in *C. neoformans* through in silico structural analysis. We utilized AlphaFold 3 to predict the three-dimensional structures of the *C. neoformans* (Cn) H2A–H2B heterodimer and H3–H4 tetramer, subsequently comparing them with the established structures of *H. sapiens* (Hs) and *S. cerevisiae* (Sc). Structural alignment of the CnH2A–H2B heterodimer (Hta1–Htb1) revealed high structural homology, with root-mean-square deviation (RMSD) values of 0.415 Å and 0.520 Å against Hs and Sc, respectively. However, structural superposition of the H3– H4 tetrameric assemblies revealed that the Cn tetramer diverged substantially from both Hs (RMSD = 2.171 Å) and Sc (RMSD = 2.301 Å), in stark contrast to the high similarity between Sc and Hs themselves (RMSD = 0.416 Å; PDB IDs: 1ID3 and 3AV1). These results indicate that the AlphaFold-predicted Cn H3–H4 tetramer may possess unique conformational features (Fig. 1C). Consistent with the core histones, the AlphaFold-predicted structure of the Cn histone variant Htz1 exhibited high structural homology with its orthologs, demonstrating RMSD values of 0.489 Å and 1.623 Å against Hs and Sc, respectively (Additional file 2: Fig. S1A). However, a markedly different pattern was observed for the centromere-specific variant CnCse4. Structural comparisons revealed a striking similarity in the global architectures of ScCse4 and HsCENP-A. In contrast, the AlphaFold model for CnCse4 appeared structurally incomplete and lacked the extended structural features observed in its orthologs. Despite this unresolved overall conformation, the core domain aligns closely with both species, yielding low RMSD values of 0.701 Å against HsCENP-A and 0.609 Å against ScCse4 (Additional file 2: Fig. S1B).

### Systematic evaluation of histone essentiality and functional redundancy in *C. neoformans*

Given the requirement of core histones for growth and viability in yeast [5, 6], we attempted to generate individual deletion mutants for all core histone genes. However, repeated attempts to delete *HTA1* or *HTB1* were not successful. Therefore, we examined their requirement for growth using a copper-conditional expression system. Repression of either gene by CuSO_4_ treatment caused severe growth defects in P_*CTR4*_*:HTA1* and P_*CTR4*_*:HTB1* strains, whereas growth was fully rescued by bathocuproinedisulfonic acid (BCS) treatment, which increased *HTA1* and *HTB1* expression (Fig. 1D). To confirm essentiality, we analyzed meiotic progeny from heterozygous diploids (*HTA1*/*hta1*Δ and *HTB1*/*htb1*Δ). Among more than 50 basidiospores, four progeny derived from the *HTA1*/*hta1*Δ heterozygous strain exhibited nourseothricin resistance (Fig. 1E and Additional file 2: Fig. S1C). PCR analysis revealed that all four nourseothricin-resistant isolates retained the wild-type (WT) *HTA1* allele, suggesting that they are anueploid disomes rather than true deletion progeny (Fig. 1E). In contrast, no nourseothricin-resistant progeny were recovered from the *HTB1*/*htb1*Δ background (Fig. 1E and Additional file 2: Fig. S1D). These results indicate that *HTA1* and *HTB1* are essential for viability in *C. neoformans*.

Unlike *HTA1* and *HTB1, HHT1* and *HHT2* were not essential for viability: *hht1*Δ and *hht2*Δ mutants were both successfully generated, although the *hht2*Δ mutant exhibited a slight growth defect (Fig. 1F). Despite multiple attempts, we were unable to obtain *hht1*Δ *hht2*Δ double mutants, suggesting that *HHT1* and *HHT2* are synthetically lethal. Given the known functional redundancy of H3 paralogs in *S. cerevisiae* [7], we constructed P_*CTR4*_*:HHT2 hht1*Δ strains to evaluate their necessity for growth. The growth of these strains was markedly impaired in the presence of CuSO_4_–mediated repression, whereas this inhibition was fully rescued by copper chelation with BCS (Fig. 1F). These data indicate that at least one functional H3 gene is required for viability. Similarly, single deletion of *HHF1* or *HHF2* did not affect growth in *C. neoformans* (Fig. 1F), whereas an *hhf1*Δ *hhf2*Δ double mutant could not be obtained, suggesting that *HHF1* and *HHF2* are synthetically lethal. To address functional redundancy for growth, we constructed P_*CTR4*_*:HHF2 hhf1*Δ strains. These strains showed an evident growth defect in the presence of CuSO_4_, whereas it was rescued by treatment with BCS. These data indicate that *HHF1* and *HHF2* function redundantly to support viability in *C. neoformans*.

We next assessed whether histone variants are similarly required for the growth and viability. In contrast to essential *HTA1, HTZ1* was dispensable for growth: *htz1*Δ mutants exhibited no discernible growth defects. To determine whether Htz1 could functionally compensate for canonical H2A, we generated *HTZ1* overexpression strains within the *P*_*CTR4*_*:HTA1* background. Quantitative analysis confirmed robust induction of *HTZ1* in these strains (Additional file 2: Fig. S1E). Notably, overexpression of *HTZ1* rescued the growth impairments of the *P*_*CTR4*_*:HTA1* strain under both basal conditions and CuSO_4_-mediated repression, suggesting that Htz1 possesses partially overlapping functional or structural roles with canonical H2A (Fig. 1G). Next, considering that *CSE4* is essential in *S. cerevisiae* [25], we evaluated its essentiality by generating P_*CTR4*_*:CSE4* system. Repression of *CSE4* led to severe growth defects, which were fully reversed by copper chelation with BCS (Additional file 2: Fig. S1F). To further demonstrate its essentiality, we performed meiotic sporulation analysis using a *CSE4*/*cse4*Δ heterozygous strain. Notably, all nourseothricin-resistant progeny retained the *CSE4* locus, indicating that they were aneuploid segregants rather than true deletion mutants (Additional file 2: Fig. S1G-H). Collectively, these data demonstrate that *CSE4* is indispensable for the viability of *C. neoformans*. Overall, we demonstrate that *HTA1, HTB1*, and *CSE4* are indispensable for viability, whereas H3 and H4 paralogs maintain viability through functional redundancy.

### The Mec1/Te1-Rad53 axis differentially regulates core histones and variants and shapes their distinct roles in the DNA damage response

Having established the requirement of core histones for viability, we next investigated their contribution to the DDR. The Mec1/Tel1-Rad53-Bdr1 pathway is a central regulator of DNA damage repair in *C. neoformans* [21, 26]; we therefore examined whether core and variant histone genes are transcriptionally and functionally integrated into this network. To this end, we measured the transcriptional responses of all core histone and variant genes following radiation exposure in *rad53*Δ, *bdr1*Δ, and *mec1*Δ *tel1*Δ mutants, and complemented these analyses with phenotypic assays of histone mutants under diverse genotoxic stresses.

Expression of both *HTA1* and *HTB1* was strongly repressed upon γ-radiation exposure in WT and *bdr1*Δ but their radiation-induced repression was attenuated in *rad53*Δ, and the *mec1*Δ *tel1*Δ double mutant exhibited constitutively low basal expression and a failure to respond to radiation (Fig. 2A and 2B). In *S. cerevisiae*, the serine 129 (S129) residue of canonical H2A is phosphorylated upon double-strand break (DSB) induction to signal the recruitment of repair machinery [27]. We found that *C. neoformans* Hta1 also possesses this conserved S129 residue, and consistent with other yeast orthologs, it is robustly phosphorylated following radiation exposure (Additional file 2: Fig. S2A and S2B). The H3 paralogs, in contrast, exhibited a striking divergence in their dependency: *HHT1* was strongly induced upon γ-radiation exposure in WT, whereas *HHT2* was repressed. *HHT1* induction was strictly Mec1-dependent yet independent of the downstream Rad53-Bdr1 circuit, whereas *HHT2* repression required the canonical Mec1/Tel1-Rad53 signaling cascade (Fig. 2A and 2B). The H4 paralogs (*HHF1* and *HHF2*) were strongly repressed upon γ-radiation exposure in WT in a Rad53-dependent but Mec1/Tel1-independent manner (Fig. 2A and 2B). In sharp contrast to canonical histones, the variants *HTZ1* and *CSE4* were largely independent of this regulatory circuit, remaining stable and unresponsive to radiation regardless of the status of the Mec1/Tel1 or Rad53/Bdr1 pathways (Additional file 2: Fig. S2C-F).

**Fig. 2.**
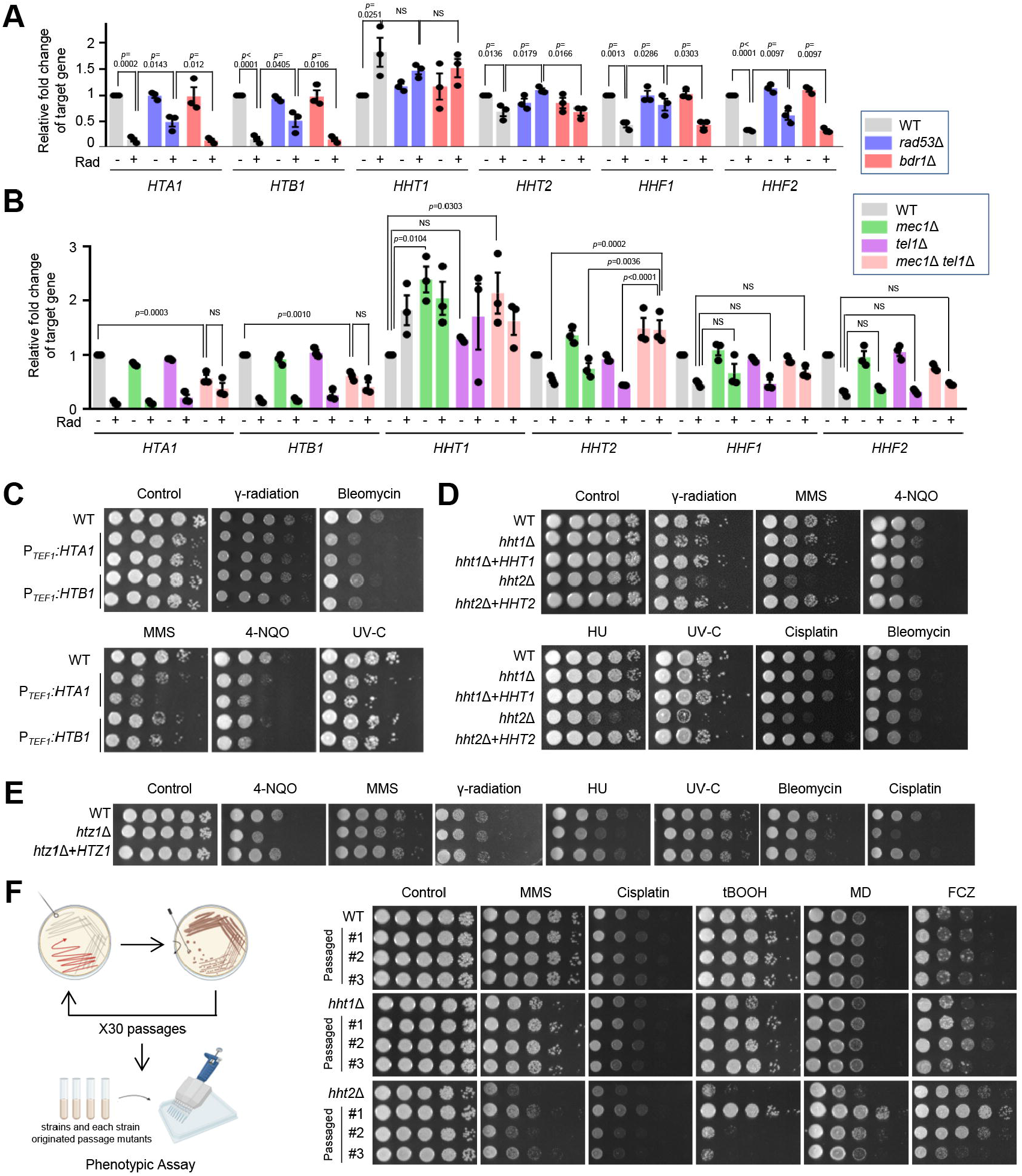
Divergent transcriptional regulation and DNA damage response of histone paralogs. **A and B** Expression levels of core histone genes in strains belonging to the DNA repair pathway under radiation exposure. Three independent biological experiments with duplicate technical replicates were performed. Error bars indicate standard errors of the means. Statistical significance of difference was determined by one-way ANOVA with Bonferroni’s multiple-comparison test (NS: not significant). **C-E** Functional roles of core histone or histone variant in DNA damage response. The diluted (1 to 10^4^) cells were spotted onto YPD media containing the indicated concentrations of chemicals and were further incubated at 30°C for 4 days. **F** Schematic representation of the microevolution experiment **G** Deletion of *HHT1* and *HHT2* promotes rapid microevolution. The phenotypic changes of original WT strain, *hht1*Δ, and *hht2*Δ mutants compared to those of passaged strains from each corresponding strains in response to diverse stress responses. The strains were spotted onto YPD medium containing 0.035 % MMS, 5 mM cisplatin, 4 μg/ml benomyl, 0.8 mM tBOOH, 0.025 mM MD, 16 μg/ml FCZ or 600 μg/ml 5-FC. The illustration was created with Biorender.

We next assessed the functional consequences of these regulatory differences using phenotypic assays. Because *HTA1* and *HTB1* are essential for viability, we employed a promoter-replacement strategy to assess the physiological consequences of their transcriptional misregulation. We replaced the native, radiation-responsive promoters of *HTA1* and *HTB1* with the constitutive *TEF1* promoter (Additional file 2: Fig. S3A) and confirmed that these strains maintained elevated histone levels even post-irradiation (Additional file 2: Fig. S3B). The resulting failure to suppress *HTA1* or *HTB1* expression caused marked susceptibility to various DNA damaging agents (Fig. 2C), underscoring the necessity of dynamic H2A-H2B dosage control for genomic maintenance. The H3 paralogs exhibited a striking functional divergence in the DDR; although the *hht1*Δ mutant showed only slight growth inhibition, the *hht2*Δ mutant displayed broad-spectrum susceptibility to methyl methanesulfonate (MMS), 4-nitroquinoline 1-oxide (4-NQO), hydroxyurea (HU), cisplatin, UV-C, and radiation, despite its intrinsic growth defect (Fig. 2D). Furthermore, *HHT1* expression was significantly upregulated in the *hht2*Δ mutant, whereas *HHT2* expression remained unchanged in the *hht1*Δ mutant (Additional file 2: Fig. S3C), suggesting that Hht2 is the major H3 isoform mediating the DDR. In contrast, the H4 paralogs appeared functionally redundant under the tested conditions, as both *hhf1*Δ and *hhf2*Δ mutants exhibited WT-level resistance to genotoxic agents (Additional file 2: Fig. S3D). *HHF1* expression was elevated in *hhf2*Δ mutants, whereas *HHF2* expression in *hhf1*Δ mutants was indistinguishable from WT (Additional file 2: Fig. S3E). Among the variants, the *htz1*Δ mutant exhibited specific susceptibility to cisplatin, HU, and 4-NQO, but WT-level resistance to bleomycin, MMS, UV-C, and γ-radiation (Fig. 2E), indicating that Htz1 contributes to the DDR in a damage type-specific manner. To circumvent the essentiality of *CSE4* and its non-responsive transcriptional pattern under radiation, we performed phenotypic analysis using the P_*CTR4*_*:CSE4* strains under BCS-mediated overexpression conditions; these strains showed a growth defect specifically in response to UV-C exposure (Additional file 2: Fig. S3F).

Given the distinct DNA damage susceptibility observed in H3 mutants, we next investigated their roles in maintaining long-term genomic stability. Quantitative assessment of spontaneous mutation rates via 5-fluoroorotic acid (5-FOA) resistance assays revealed that both *hht1*Δ and *hht2*Δ mutants maintained mutation frequencies comparable to WT (Additional file 2: Fig. S4A). Despite this stable mutation rate, we reasoned that chromatin-level perturbations might still drive microevolution through increased phenotypic plasticity. To test this hypothesis, we monitored phenotypic diversification across independent passaged lineages. Notably, lineages derived from both *hht1*Δ and *hht2*Δ mutants exhibited significantly greater phenotypic divergence from their parental strains than WT lineages, with the *hht2*Δ lineages displaying more pronounced instability (Fig. 2F and Additional file 2: Fig. S4B). These findings suggest that H3 deficiency accelerates microevolutionary changes in *C. neoformans*, likely by compromising epigenetic or structural genome integrity.

To further dissect the functional interplay between Rad53 and chromatin components, we systematically constructed double mutants of *rad53*Δ with core (*hht1*Δ, *hht2*Δ, *hhf1*Δ, *hhf2*Δ) or variant (*htz1*Δ) histones and evaluated their synergistic or epistatic effects on the DDR. *rad53*Δ *hht1*Δ, *rad53*Δ *hhf1*Δ, and *rad53*Δ *hhf2*Δ double mutants showed DNA damage resistance comparable to the *rad53*Δ single mutant. In contrast, *rad53*Δ *hht2*Δ mutants exhibited increased susceptibility to most DNA damaging agents tested, with the notable exception of bleomycin, where the double mutant phenocopied *the hht2*Δ single mutant (Additional file 2: Fig. S5A). The interaction between Rad53 and Htz1 was likewise stress-dependent: the *rad53*Δ *htz1*Δ double mutant showed enhanced sensitivity to 4-NQO compared with either single mutant, whereas its UV-C and cisplatin susceptibilities mirrored that of the *htz1*Δ single mutant (Additional file 2: Fig. S5B). These data suggest that Htz1 operates through both Rad53-dependent and Rad53-independent pathways depending on the type of genotoxic stress.

Collectively, these findings demonstrate that the Mec1/Tel1-Rad53 axis selectively regulates canonical histone dosage but not histone variants, coordinating chromatin assembly with the DDR and translating into distinct functional roles for each histone family in genomic maintenance.

### Core and variant histones are required for pleiotropic stress adaptation and full virulence in *C. neoformans*

In *C. neoformans*, histone-modifying enzymes, including histone acetyltransferases (e.g. Gcn5) and deacetylases (e.g. Hda1), are known to orchestrate stress resistance, antifungal drug tolerance, and virulence [13, 14]. However, although these modifiers are pivotal, the functional requirements of their primary substrates—the core and variant histones themselves—remain largely unexplored. To address this knowledge gap, we performed systematic phenotypic profiling of individual histones under multifaceted stress conditions. Consistent with its role in the DDR, the *hht2*Δ mutant exhibited growth defects in response to oxidative stress, osmotic stress, endoplasmic reticulum (ER) stress, heat stress, and cell wall stress (Fig. 3A and Additional file 2: Fig. S6). In contrast, the *hht1*Δ mutant displayed WT-level resistance to all environmental cues tested except for DDR. Similarly, the *hhf1*Δ and *hhf2*Δ mutants were as resistant to diverse stress as WT. Notably, both *hht2*Δ and *htz1*Δ mutants showed increased resistance to fluconazole (Fig. 3A). These findings indicate that Hht2 plays a pivotal role in diverse stress responses in *C. neoformans*.

**Fig. 3.**
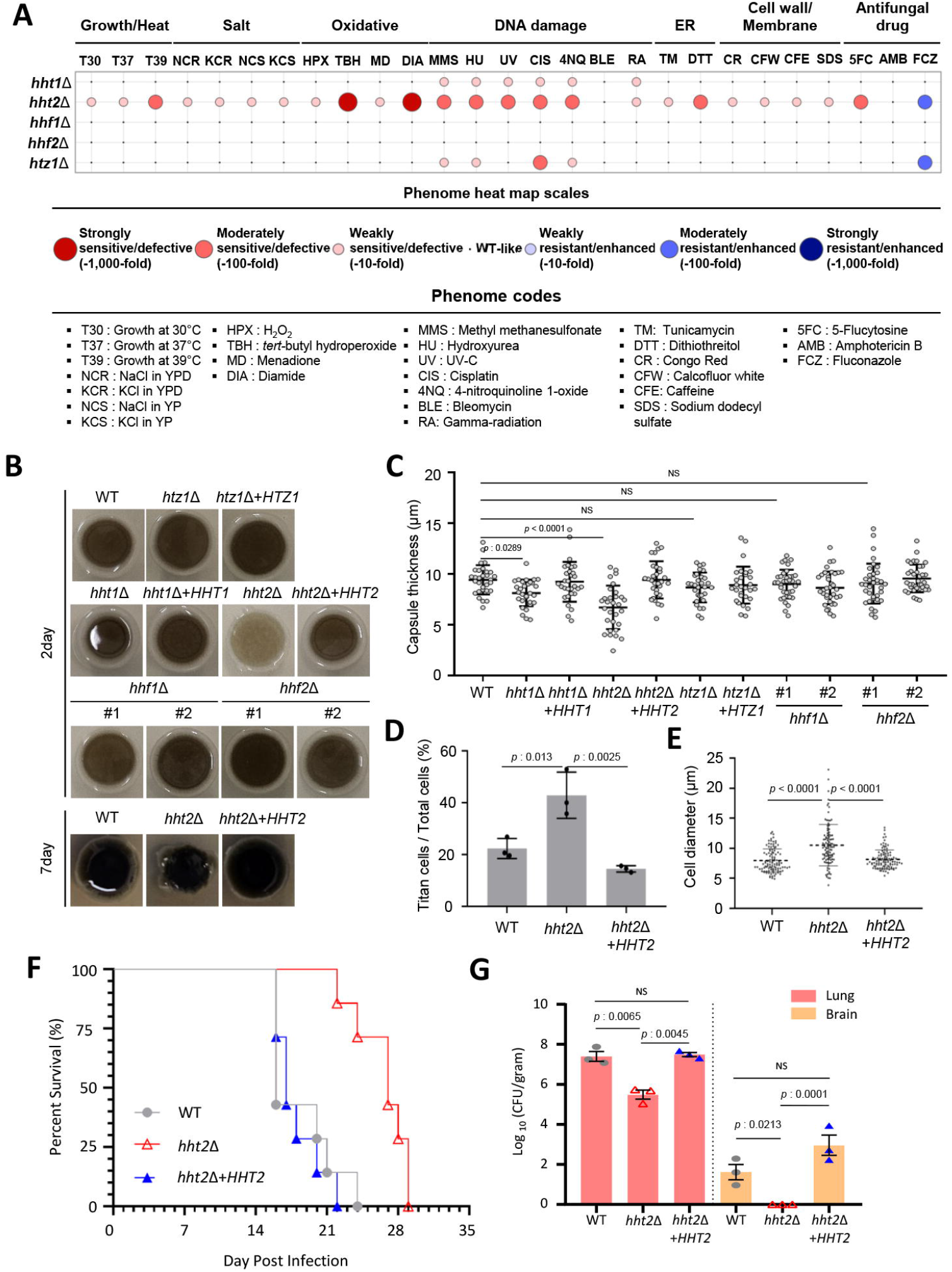
Requirement of core and variant histones for pleiotropic stress adaptation, virulence factor formation, and virulence. **A** Phenome heat map created for *C. neoformans* core histone and histone variant mutants. Phenotype scores are represented in distinct colors based on qualitative or semi-quantitative measurement of core histone and histone variant mutants under indicated growth conditions. Red and blue in the heat map represent reduction and enhancement, respectively. Phenotype strengths (strong≥ 3 log dilution factor, moderate= 2 log dilution factor and weak= 1 log dilution factor) are distinguished in gradients of red or blue. **B** The *hht2*Δ mutant showed reduced melanin production. The strains were spotted on L-DOPA medium (0.2 % glucose) and incubated at 30°C for the indicated days. **C** The *hht1*Δ and *hht2*Δ mutants showed reduced capsule production. Statistical analysis was conducted using one-way ANOVA (NS: not significant). **D and E** Deletion of *HHT2* increased the proportion of titan cells and cell body size. Quantitative measurement of titan cell percentage **(D)** and cell size **(E)** under *in vitro* titan cell inducing condition. The percentage of titan cells under *in vitro* inducing conditions was calculated from three independent biological experiments and cell size data (n = 100 cells) were representative of three independent experiments. Statistical analysis of difference was determined by one-way ANOVA with Bonferroni’s multiple-comparison test. **F** Survival curves were monitored over the experimental period. Statistical analysis was performed using the log-rank (Mantel–Cox) test: WT vs. *hht2*Δ, *P* = 0.0008; *hht2*Δ vs. *hht2*Δ+*HHT2, P* = 0.0003; WT vs. *hht2*Δ+*HHT2, P* = 0.6910. **G** Fungal burden assays were performed on day 14 post-infection, when mice were sacrificed, and fungal loads in the lung and brain were quantified. Statistical analysis was conducted using one-way ANOVA (NS: not significant).

Next, we evaluated the requirement of histones for the production of capsule and melanin, the two hallmark virulence factors of *C. neoformans* [28, 29]. On L-DOPA medium, *htz1*Δ, *hhf1*Δ, *hhf2*Δ, and *hht1*Δ mutants produced WT levels of melanin. In contrast, the *hht2*Δ mutant exhibited delayed melanization (Fig. 3B); however, because melanin levels in *hht2*Δ eventually reached WT levels after 7 days, this delay likely reflects a growth defect rather than an intrinsic biosynthetic impairment. Conversely, capsule formation was significantly reduced in both *hht1*Δ and *hht2*Δ mutants, whereas no defect in capsule formation was observed in *hhf1*Δ, *hhf2*Δ, or *htz1*Δ mutants (Fig. 3C and Additional file 2: Fig. S7A). Notably, extended incubation failed to restore capsule size in the *hht2*Δ mutant (Additional file 2: Fig. S7B), indicating that H3 paralogs are indispensable for proper capsule formation. Beyond these virulence traits, we further investigated whether histones are required for titan cell formation, a critical morphological transition for host immune evasion [30]. We found a higher proportion of titan cells and a larger cell diameter in the *hht2*Δ mutant compared with WT (Fig. 3D and 3E). These data suggest that the H3 paralog Hht2 is involved in virulence factor formation.

Given the pleiotropic defects in stress adaptation and virulence factor production— particularly in the *hht2*Δ mutant—we next assessed its virulence in a murine model of cryptococcosis. Mice infected with the *hht2*Δ mutant exhibited significantly prolonged survival compared with those infected with WT or complemented strains (Fig. 3F). Consistent with this attenuation, fungal burdens in both the lungs and brains of *hht2*Δ-infected mice were markedly reduced (Fig. 3G). Despite this reduction, all infected animals eventually succumbed to infection by day 28, indicating that *HHT2* is required for full virulence in *C. neoformans*.

### Disrupting histone maintenance induces extensive transcriptomic reprogramming in the *hht2*Δ mutant

Given the pivotal roles of Hht2 in multifaceted stress adaptation and virulence, we performed RNA sequencing (RNA-seq)-based transcriptomic analysis of the *hht2*Δ mutant to investigate how alterations in nucleosome integrity drive transcriptional reprogramming. We found that expression levels of 1,274 genes (1,053 upregulated and 221 downregulated; *P* < 0.05, |Log_2_ fold change| > 1) were altered in the *hht2*Δ mutant, compared to WT (Fig. 4A and Additional file 1: Table. S1). Kyoto Encylopedia of Genes and Genomes (KEGG) pathway enrichment analysis revealed that Hht2-dependent genes were primarily linked to metabolic processes (Fig. 4B). Specifically, genes upregulated in the *hht2*Δ mutant were enriched in the biosynthesis of secondary metabolites, carbon metabolism, and the degradation of various amino acids and fatty acids. In contrast, genes associated with the ribosome were markedly downregulated. These results suggest that the loss of *HHT2* induces a massive transcriptomic shift that simultaneously activates alternative metabolic pathways whereas suppressing the protein translation machinery (Fig. 4B).

**Fig. 4.**
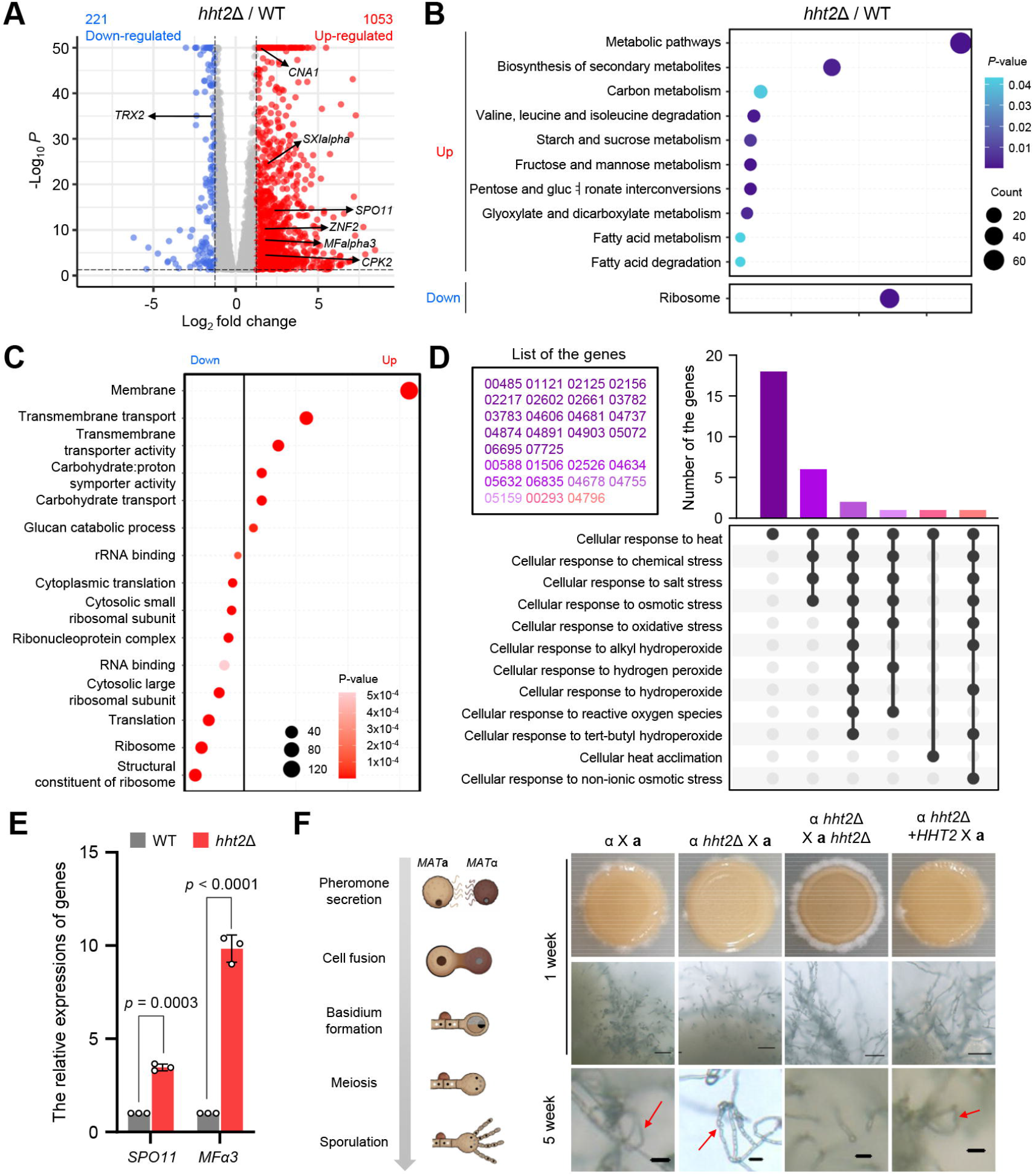
Transcriptional profiles of *C. neoformans* in WT and *hht2*Δ mutant. **A** Volcano plot showed the changes in levels of gene expression in *hht2*Δ mutant compared to the WT strain. Upregulated genes (more than 2 fold-increase, *P* < 0.05) are shown in red, whereas downregulated genes (less than 2 fold-decrease, *P* < 0.05) are shown in blue. **B** Significant differences in KEGG pathway analysis in *hht2*Δ mutant were observed at basal condition. **C** Functional categorization of DEGs (*P* < 0.05, |Log_2_FC| ≥ 1) in *hht2*Δ compared to WT. Differentially expressed genes were classified into functional categories using the DAVID tool. **D** UpSet plot analysis revealing heat-specific and multi-stress responsive genes. The plot illustrates the intersection of 29 up-regulated genes (enriched for ‘cellular response to heat’) with other environmental stress-related Gene Ontology (GO) terms retrieved from FungiDB. The vertical bar chart displays the number of genes within each specific intersection, while the connected dots below indicate the overlapping stress pathways. The inset box lists *C. neoformans* locus tags (CNAG prefix), color-coded to match the intersection bars. **E** The expression levels of genes involved in mating process were measured in WT and *hht2*Δ strains. Three independent biological experiments with duplicate technical replicates were performed. Error bars indicate standard errors of the means. Statistical significance of difference was determined by one-way ANOVA with Bonferroni’s multiple-comparison test. **F** The deletion of *HHT2* increased hyphal formation but decreased basidiospore formation. Schematic diagrams of the *C. neoformans* mating process (left) and representative images of spores produced from WT, unilateral, and bilateral crosses (right). The images were photographed after indicated weeks and red arrow indicated basidiospores. The illustration was created with Biorender.

To comprehensively profile global transcriptomic alterations and biological processes, we performed a Gene Ontology (GO) enrichment analysis using DAVID. The analysis revealed a significant downregulation of genes associated with ribosomal functions, whereas genes involved in membrane components and transmembrane transport were upregulated (Fig. 4C). Furthermore, to delineate the specific stress-responsive mechanisms directly linked to environmental adaptation and the virulence of *C. neoformans*, we interrogated our dataset with FungiDB, a fungal-specific database [31]. Consistent with our preceding phenotypic assays, this targeted analysis demonstrated that the upregulated gene set was highly enriched for the ‘cellular response to heat’ [*P* < 0.05, false discovery rate (FDR) < 0.05]. During infection, *C. neoformans* encounters a complex array of hostile conditions, including elevated mammalian host body temperature and macrophage-derived oxidative bursts [32, 33]. To test whether the 29 genes annotated to ‘cellular response to heat’ are heat-specific or broadly stress-responsive, we analyzed their overlap with other environmental stress GO terms. UpSet Plot visualization separated these genes into distinct functional groups (Fig. 4D). A subset of 19 genes exhibited highly specific roles, mapping exclusively to thermal stimuli without involvement in other tested stress pathways (Fig. 4D). Conversely, we identified a distinct group of 11 multi-stress responsive genes that mapped simultaneously to ‘cellular response to osmotic stress’, ‘cellular response to oxidative stress’, ‘response to salt stress’, and ‘cellular response to hydrogen peroxide’ (Fig. 4D). The presence of these overlapping cross-talk genes suggests that they function as core hub regulators, integrating multiple environmental cues to drive the pathogen’s overall survival and virulence network.

Furthermore, *HHT2* deletion disrupted the expression of a broad array of signaling regulators, including kinases (e.g., *YPK1*), phosphatases (e.g., *CNA1*), and transcription factors (e.g., *ZNF2*) (Additional file 3: Dataset 1). These results indicate that Hht2 governs a complex regulatory network essential for stress adaptation and developmental processes such as mating. Notably, the *hht2*Δ mutant exhibited reduced expression of *ERG1* and *ERG11* genes encoding key enzymes in the ergosterol biosynthetic pathway (Additional file 2; Fig. S8). This downregulation was consistent with the heightened susceptibility of the *hht2*Δ mutant to cell membrane stress, as noted in our preceding phenotypic analysis (Fig. 3A and Additional file 2: Fig. S6B and S6C). Paradoxically, the *hht2*Δ mutant also displayed enhanced resistance to fluconazole, suggesting that *HHT2* deletion alters cell membrane integrity in a highly complicated manner. In parallel, we observed upregulation of meiotic/mating-related genes (*SPO11* and *MFα3*) in the *hht2*Δ mutant (Fig. 4E). Supporting this, we observed accelerated basidium formation in bilateral crosses between *MAT***a** *hht2*Δ and *MAT*α *hht2*Δ mutant strains compared to WT crosses. However, these crosses failed to produce intact basidiospores (Fig. 4F), suggesting that maintaining intact nucleosomes is required for both early and late phases of the mating process. Collectively, transcriptomic alterations in the *hht2*Δ mutant correlate with its diverse phenotypic changes.

### Core histone maintenance is required for chromatin architecture and is coupled to transcriptional reprogramming

Unlike perturbations of higher-order structural organizers, the deletion of a core histone eliminates a fundamental structural unit [34]. We hypothesized that Hht2 loss leads to critical alterations in chromatin architecture, which in turn drive transcriptomic reprogramming. To investigate this link, we performed Hi-C analysis, a genome-wide chromosome conformation capture technique that maps 3D chromatin interactions (Fig. 5A). Comparative analysis of Hi-C contact matrices revealed pronounced structural differences between WT and the *hht2*Δ mutant in chromosome 10 (Chr10) (Fig. 5B). In contrast, contact matrices of other chromosomes remained unaltered in the *hht2*Δ mutant compared with WT (Fig. 5B and Additional file 2: Fig. S9A). Quantifying these observed alterations, contact probability analysis revealed a marked reduction in chromatin contact frequencies along Chr10 in the *hht2*Δ mutant (Fig. 5C). In contrast, only minor differences were observed on the other chromosomes (Additional file 2: Fig. S9B).

**Fig. 5.**
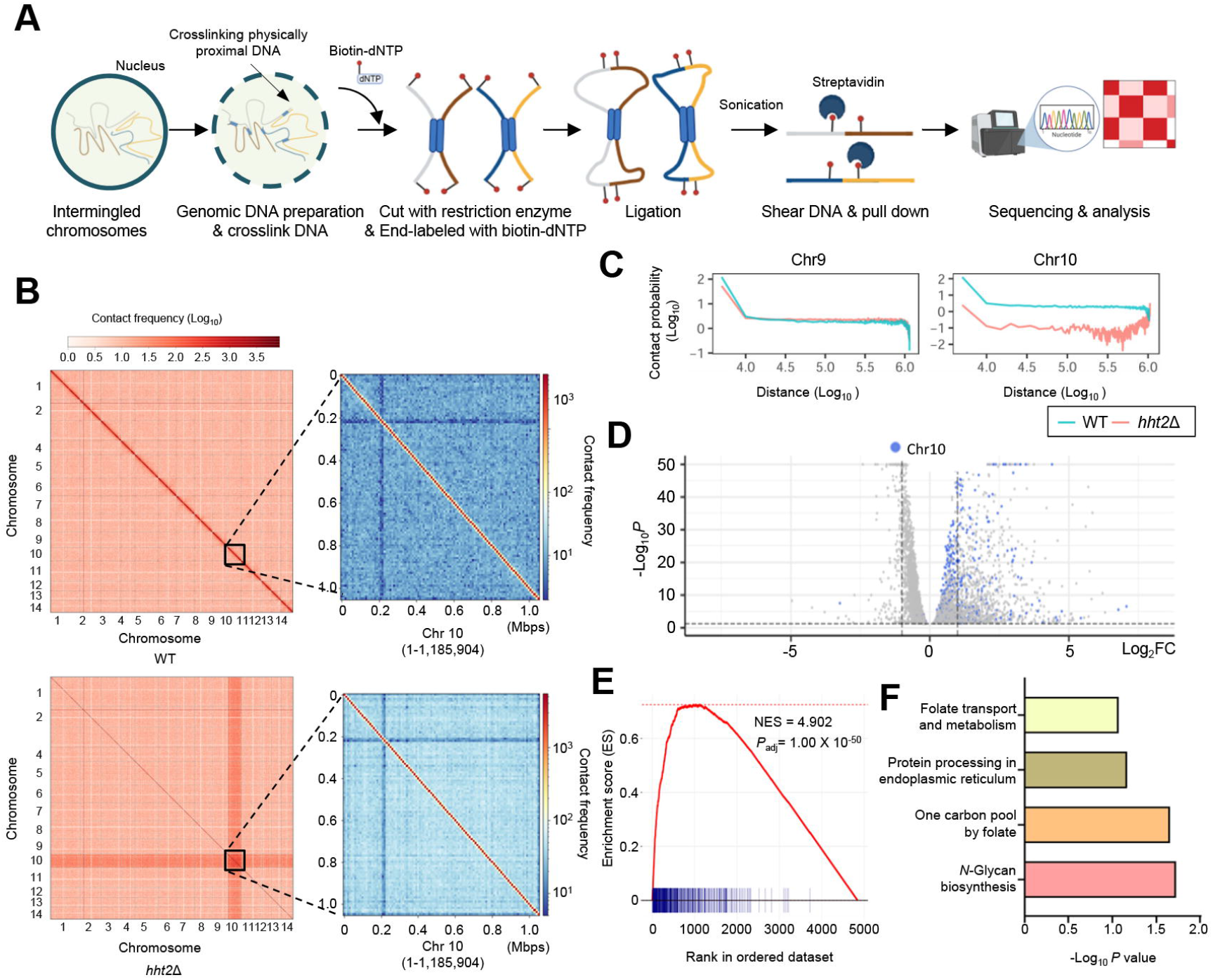
Hi-C profiles in WT and *hht2*Δ mutant revealed conformation changes of chromosome. **A** Schematic diagram of experimental processes of Hi-C. Samples were crosslinked, digested, biotin-labeled, ligated, and streptavidin-enriched to construct a 3D chromatin interaction library. Prepared libraries were subsequently sequenced and analyzed to map genome-wide spatial architecture. **B** Contact maps (10 kb-resolution) of whole genome (left) and Chromosome 10 (1,185,904 base pairs) (right) of WT (upper panels) and *hht2*Δ (down panels). **C** Average contact probability (CP) as a function of genomic distance for the chromosome 10 in WT and *hht2*Δ mutant. **D** Volcano plot illustrating the genes showing statistically significant changes in expression (*P* < 0.05) in gray, with those overlapping the chromosome 10 region highlighted in blue. **E** Positional GSEA analysis using RNA-seq DEGs and gene set in chromosome 10. All expressed genes were pre-ranked based on their differential expression test statistics (stat values) comparing the *hht2*Δ mutant to the WT. The evaluated positional gene set consists of 387 overlapping genes localized to this specific chromosome. The Normalized enrichment score (NES) and statistical significance are indicated on the plot. **F** We extracted the overlapping genes between those showing statistically significant changes in expression (*P* < 0.05) (WT vs. *hht2*Δ mutant under basal conditions) and the genes annotated in chromosome 10. Subsequently, KEGG pathway analysis was performed exclusively on this intersected gene set. The DAVID tool was utilized to conduct KEGG pathway analysis.

Integration of Hi-C and RNA-seq data demonstrated that this chromosomal reorganization was accompanied by transcriptional activation; 387 out of the 463 annotated genes on Chr10 overlapped with genes showing statistically significant expression changes (*P* < 0.05) identified in our RNA-seq analysis (Fig. 5D). To statistically validate whether this transcriptional activation reflected a coordinated response driven by chromosomal structural changes rather than stochastic events, we performed positional Gene Set Enrichment Analysis (GSEA). We defined a custom gene set comprising all genes located on Chr10 and analyzed their distribution across the ranked list of global transcriptional changes. GSEA revealed that genes on Chr10 were significantly enriched in the upregulated fraction of the transcriptome (normalized enrichment score [NES] = 4.90; adjusted p-value < 1 × 10^−50^) (Fig. 5E). Thse findings suggest that structural changes on Chr10 drive regional alterations in gene expression.

We next performed KEGG pathway analysis to characterize the functional roles of the 387 overlapping genes identified in both the Hi-C structural analysis and RNA-seq data. We found enrichment of genes involved in folate transport and metabolism and *N*-glycan biosynthesis (Fig. 5F). Folate metabolism is well known to play a critical role in the oxidative stress response [35]. This established link is consistent with our finding that the *hht2*Δ mutant showed increased susceptibility to oxidative stress, including *tert*-butyl hydroperoxide (tBOOH), menadione (MD), and hydrogen peroxide (H_2_O_2_) (Fig. 3A and Additional file 2: Fig. S6A). Furthermore, *N*-glycan biosynthesis represents a pivotal post-translational modification occurring in the ER [36]. Supporting this, *hht2*Δ mutant exhibited growth defect in response to ER stress inducers, dithiothreitol (DTT) and tunicamycin (Fig. 3A and Additional file 2: Fig. S6E). Collectively, our data suggest that *HHT2* deletion drives a profound conformational change in Chr10, which translates into transcriptional reprogramming and increased susceptibility of *C. neoformans* to oxidative stress and ER stress.

### Disrupting core histone maintenance alters physical interactions among chromosomes

In eukaryotes, genomic DNA is organized into sophisticated hierarchical 3D architecture within the nucleus [37]. This spatial organization plays a pivotal role in regulating essential genomic activities, including transcription, DNA replication, and DNA damage repair [38, 39]. Given the functional significance of this higher-order architecture, we next systematically assess the global 3D genome landscape by analyzing A/B compartments and TADs across the entire genome. As previously described [40], global chromatin regions were partitioned based on their Principal Component 1 (PC1) scores. The A compartment is characterized by positive PC1 values and represents an open, loosely packed chromatin conformation associated with active transcription, whereas the B compartment is defined by negative PC1 values and corresponds to a tightly packed, transcriptionally inactive state (Fig. 6A, left). The A/B compartment analysis revealed that the *hht2*Δ mutant exhibited an increased proportion of active A compartments compared with WT, accompanied by a decreased proportion of inactive B compartments (Fig. 6A, right).

**Fig. 6.**
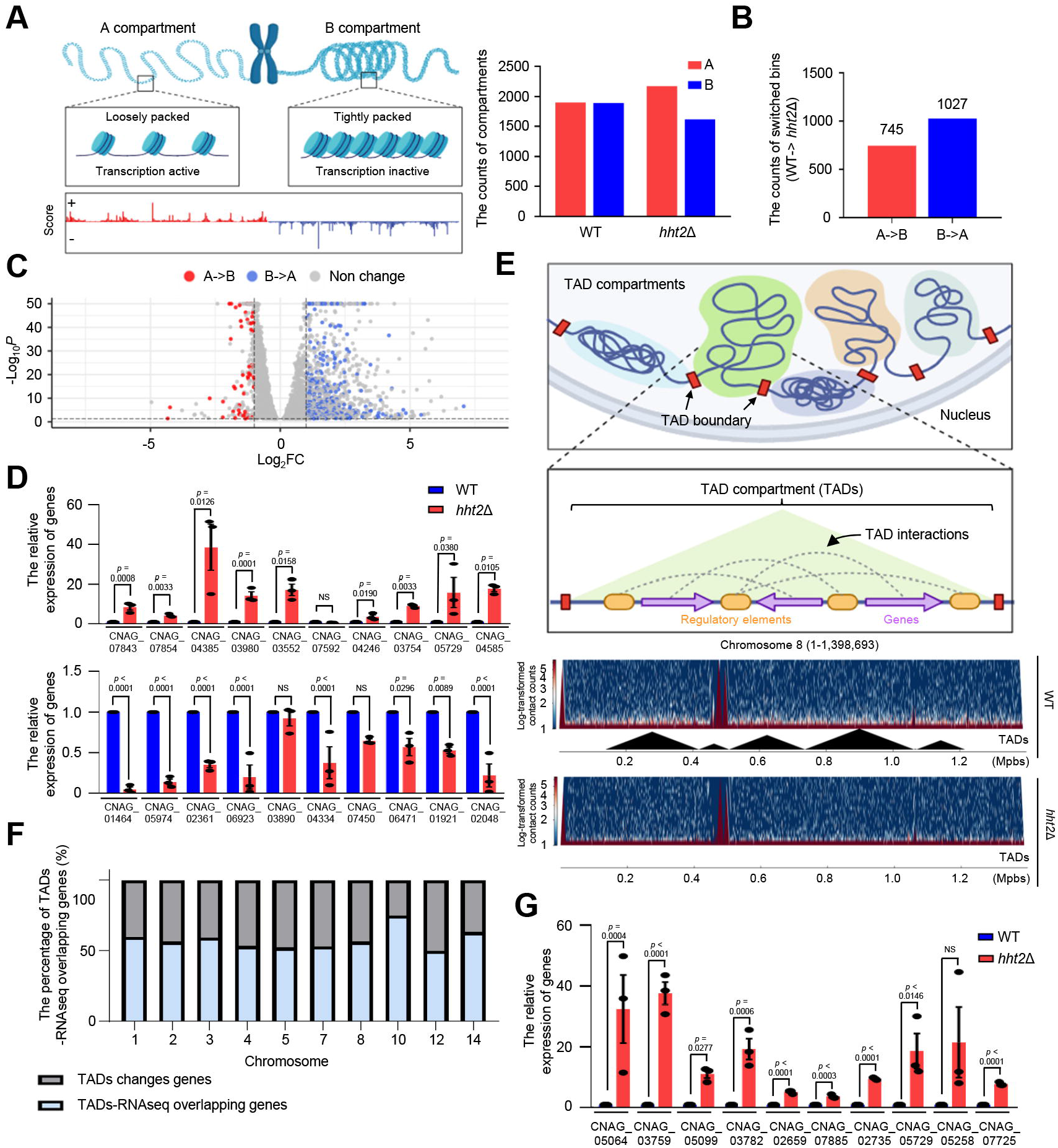
The deletion of *HHT2* led to alterations in local 3D chromatin architecture. **A** Structura and functional characterizationof of active (A) and inactive (B) chromatin compartments (left) Number of A/B compartments in the WT and the *hht2*Δ mutant (right). **B** The number of A/B compartment switched bins in the WT and the *hht2*Δ mutant. **C** Volcano plot showing statistically significant changes in expression (*P* < 0.05) (gray) and those overlapping with A/B compartment switched genes. Red dots indicated that overlapped genes between A-to-B switched genes and DEGs (*P* < 0.05, Log_2_FC ≤ −1), blue dots indicated that overlapped genes between B-to-A switched genes and DEGs (*P* < 0.05, Log_2_FC ≥ 1). **D and G** The expression levels of genes corresponded to B-to-A switched genes (Fig. D upper panel), A-to-B switched genes (Fig. D down panel) and TAD alteration genes (Fig. G) were measured in WT and *hht2*Δ strains. Three independent biological experiments with duplicate technical replicates were performed. Error bars indicate standard errors of the means. Statistical significance of difference was determined by one-way ANOVA with Bonferroni’s multiple-comparison test (NS: not significant) **E** Hierachical organization and functional architecture of Topologically Associating Domains (TADs) (Fig. E upper panel). Contact maps and corresponding TAD annotations of WT and *hht2*Δ mutants in chromosome 8, generated using the hicFindTADs tool (Fig. E down panel). **F** The graph shows the proportion of TAD-altered genes that overlap with the genes showing statistically significant changes in expression (*P* < 0.05) across individual chromosomes. The illustration was created with Biorender.

Analysis of global compartment switching between WT and the *hht2*Δ mutant showed numerous transitions. Notably, we observed a higher frequency of B-to-A switches (compartment B in WT converting to A in the *hht2*Δ mutant) compared with A-to-B switches (Fig. 6B). The direction of compartment switching was correlated with gene expression changes: genes in A-to-B regions were associated with reduced expression (log_2_FC ≤ −1), whereas genes in B-to-A regions showed increased expression in the *hht2*Δ mutant (log_2_FC ≥ 1) (Fig. 6C and Additional file 3: Dataset 2). To statistically validate the directional correlation, we performed Fisher’s exact test using a 2×2 contingency table (Additional file 2: Fig. S9C). Specifically, we compared the proportion of downregulated genes in A-to-B transitions with stable A-to-A regions. Notably, 54.17% (525 of 969 genes) within A-to-B transition regions were downregulated, a significantly higher proportion than 43.5% (577 of 1,326 genes) in stable A-to-A regions.

For the reverse direction, we performed a parallel analysis comparing the proportion of upregulated versus non-upregulated genes (unchanged or downregulated) within B-to-A transitions and stable B-to-B regions. This analysis revealed that 47.1% (627/1,331 genes) in B-to-A transition regions were upregulated, compared with 0% (0 genes) in stable B-to-B regions (Fisher’s exact test, *P* < 0.0001; Additional file 2: Fig. S9D). Collectively, these results demonstrate a significant directional concordance between A/B compartment switching and transcriptional alterations, confirming that chromatin structural changes are intrinsically linked to gene regulation. To experimentally validate whether compartment switching translates into actual gene expression changes, we performed quantitative reverse transcription-PCR (qRT-PCR) analysis. Among 10 selected genes located in B-to-A transition regions, all but one (CNAG_07572) showed expected increase in expression in the *hht2*Δ mutant compared with WT (Fig. 6D, upper panel). Similarly, we observed the expected decrease in expression for genes within A-to-B transition regions, with the exceptions of CNAG_03890 and CNAG_07450 (Fig. 6D, lower panel).

To further investigate local chromatin architecture and its potential role in gene regulation, we performed TAD analysis to achieve a finer structural resolution. TADs represent highly insulated spatial units within the nucleus that physically orchestrate and modulate local regulatory interactions [37, 41]. As illustrated in Fig. 6E (upper panel), these discrete domains compartmentalize local genes along with their corresponding regulatory elements, thereby facilitating coordinated spatial interactions within the domain boundaries. TAD analysis identified 27 TADs across the WT genome, including 5 domains on Chr8, whereas the *hht2*Δ mutant exhibited a markedly reduced number of TADs (5 in total across all chromosomes), with none detected on Chr8 (Fig. 6F and Additional file 2: Fig. S9E). Notably, although noticeable differences in the contact map were strictly restricted to Chr10, TAD alterations in *hht2*Δ mutant occurred across the entire genome.

To determine whether TADs alterations lead to transcriptional changes, we identified the genes located within altered TADs between WT and the *hht2*Δ mutant. We then assessed the extent of overlap between these genes and genes showing statistically significant expression changes (*P* < 0.05) identified via RNA-seq. Notably, the proportion of genes within altered TADs that overlapped with differentially expressed genes (DEGs; *P* < 0.05, |Log_2_FC| ≥ 1)) was 75% on Chr10 and 50% on Chr12 (Fig. 6F), indicating that TAD-level reorganization was not strictly confined to Chr10 but extended to other chromosomes that did not exhibit major contact map alterations. To validate whether the observed TAD alterations directly translated into gene expression changes, we performed qRT-PCR analysis. For 10 selected genes located within altered TADs, we confirmed significant changes in their expression levels in the *hht2*Δ mutant compared with WT (Fig. 6G). Analysis of overlapping genes associated with differential expression, TAD alterations, and A/B compartment changes showed that most DEGs overlapped with regions exhibiting either TAD or compartment changes (Additional file 2: Fig. S9F and Additional file 3: Dataset 2). Together, these results demonstrate that disrupting a core histone leads to extensive genome reorganization, accompanied by coordinated changes in chromatin architecture and gene expression.

## Discussion

In this study, we elucidated the distinct biological roles and regulatory mechanisms of histone core proteins and their variants in the pathogenic fungus *C. neoformans*. By integrating Hi-C and RNA-seq analyses, we uncovered a close association between genome structural integrity and transcriptional regulation mediated by the histone protein Hht2. Furthermore, we demonstrated that Hht2 contributes to the virulence of *C. neoformans* (Fig. 7).

**Fig. 7.**
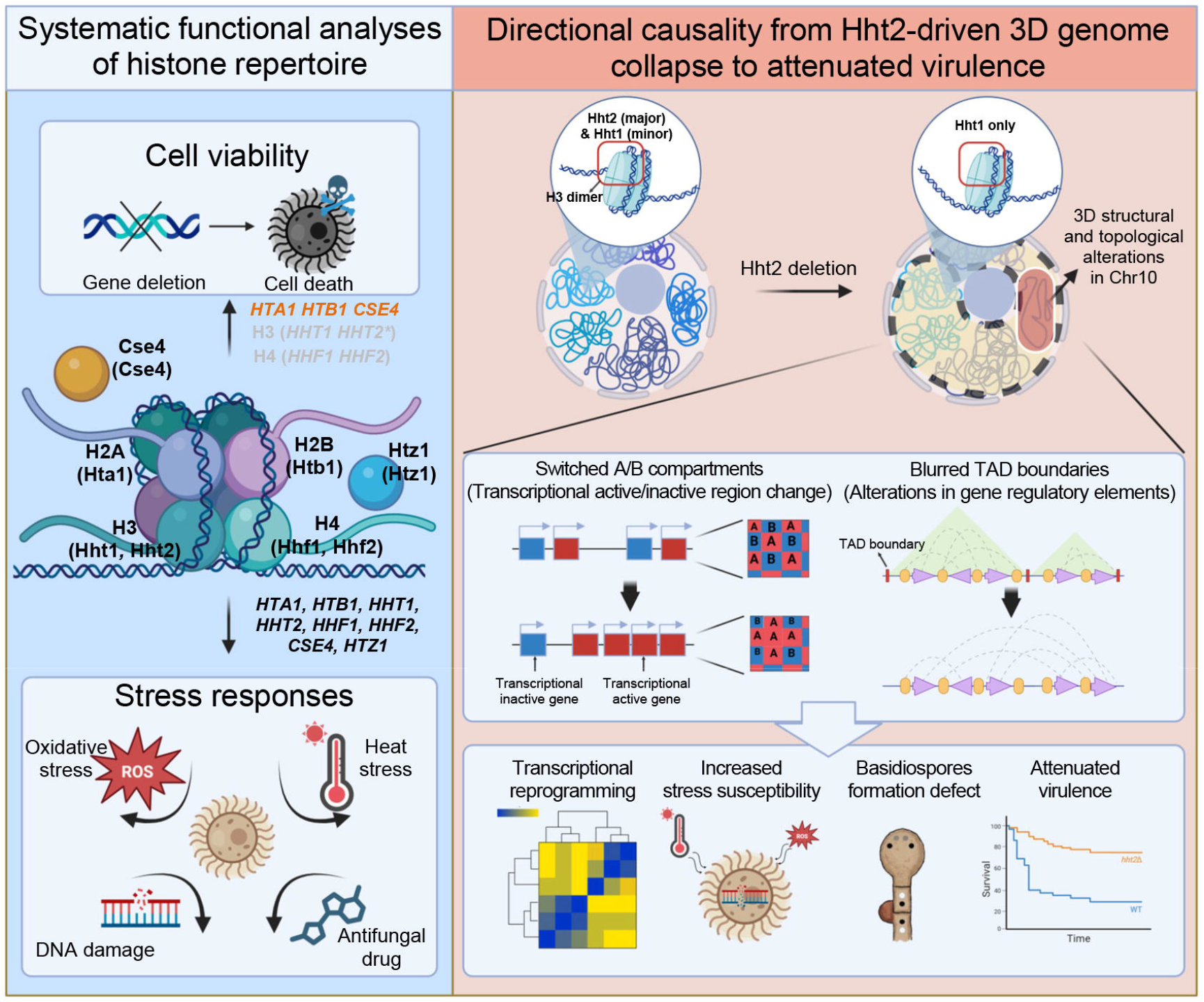
Systematic functional landscape of histones and *HHT2*-mediated 3D genome regulation of fungal pathogenesis. This schematic illustrates the functional roles of core histones and variants in cell viability and stress responses in *C. neoformans*. In the panel depicting cell viability, red text indicates essential genes (*HTA1, HTB1*, and *CSE4*), asterisks (*) denote quasi-essential gene (*HHT2*), and gray represents synthetic lethality between the paralogous gene pairs encoding H3 (*HHT1* and *HHT2*) and H4 (*HHF1* and *HHF2*). The loss of *HHT2* induces a directional cascade of 3D structural alterations in the genome, including changes in A/B compartments and topologically associating domain (TAD) boundaries. Consequently, these structural changes drive transcriptional reprogramming and result in phenotypic alterations, including altered stress susceptibility, basidiospore formation, and virulence.

Our analysis revealed that core histones in *C. neoformans* possess evolutionarily conserved yet distinct features compared with those of other fungi. We found that the *hht2*Δ mutant exhibit robust changes in DNA damage responses and virulence, whereas *hhf1*Δ and *hhf2*Δ mutants appeared dispensable for these phenotypes. This mirrors observations in *S. cerevisiae*, where H3 mutations cause growth defects, whereas individual H4 deletions in *Candida glabrata* do not lead to major phenotypic changes under DNA damage stress [9, 42]. This functional distinction is likely attributable to their post-translational modification profiles. Histone H3 modifications are closely linked to transcriptional processes, whereas H4 modifications (e.g., H4K16ac and H4K20me) mediate chromatin compaction and DNA damage signaling [43]. Thus, our findings suggest that loss of H3 impacts a broader spectrum of cellular and pathogenic phenotypes than loss of H4. Although H3 modifications by enzymes such as Gcn5 are known to modulate virulence and stress responses in various fungi [14, 44, 45], the specific modifications responsible for these phenotypes remain to be identified. Therefore, further epigenetic studies focusing on individual H3 modification sites are warranted.

A central discovery of this study is that Hht2 serves as a master structural scaffold whose absence triggers a dramatic 3D structural collapse, specifically on Chr10. Although such a chromosome-specific structural collapse driven by a core histone is unprecedented, it bears a conceptual resemblance to regulatory mechanisms observed in other fungi. In the plant pathogenic fungus *Fusarium oxysporum*, the loss of specific histone-modifying factors induces extreme structural instability and transcriptional derepression specifically within its accessory chromosomes [46]. Although *C. neoformans* lacks canonical accessory chromosomes, our results suggest that Chr10 possesses a unique chromatin architecture that is exceptionally reliant on Hht2 for its structural stability. This chromosome-specific structural collapse is functionally linked to the transcriptional landscape of the region. Comparison with our RNA-seq data revealed consistent transcriptional upregulation of genes located on Chr10. Importantly, Chr10 harbors an enrichment of genes involved in folate transport, folate metabolism, and one-carbon metabolism. These metabolic pathways are pivotal for maintaining dNTP homeostasis, which is essential for the DDR pathway [22, 47]. Based on these findings, we propose that Hht2 serves as a master regulator of the DDR and dNTP balance by modulating the structural integrity of Chr10.

We idenfied a distinctive, Rad53-independent regulatory circuit for *HHT1* that diverges from other core histone expression patterns observed in *C. neoformans*. In *S. cerevisiae*, histone genes are transcriptionally regulated as chromosomally co-localized pairs (*HTA1*-*HTB1, HHT2*-*HHF2*, and *HHT1*-*HHF1*) to facilitate coordinated control [48, 49]. In contrast, in *C. neoformans*, histone genes are distributed across different chromosomes (*HHT1*: Chr10; *HHF1*: Chr10; *HHF2*: Chr4), with only the *HHT2*-*HTA1*-*HTB1* cluster residing on a single chromosome (Chr2). Our Hi-C analysis revealed that although *HHT1* and *HHF1* are located on different chromosomes, they were positioned in close spatial proximity within the 3D genome (Additional file 2: Fig. S9G). This suggests that the distinct regulation of *HHT1* is likely driven by trans-acting factors or specific chromatin contexts rather than simple linear chromosomal positioning or spatial proximity alone.

Beyond canonical histones, our functional characterization of the variant Htz1 confirms its conserved necessity for maintaining genomic stability. In *S. cerevisiae*, Htz1 replaces the conventional H2A–H2B dimer as part of an Htz1–H2B heterodimer within the nucleosome. Htz1 is deposited at promoter regions by the Ino80/SWR complexes to regulate transcription, while its NuA4-mediated acetylation is critical for the DDR [50–52]. Consistently, our results showed that *HTZ1* overexpression rescued the growth defect caused by *HTA1* suppression, whereas *htz1*Δ mutants exhibited increased DNA damage susceptibility in a Rad53-independent manner. These results, consistent with the pleiotropic defects observed in *F. graminearum* [53], support a conserved role of Htz1 in maintaining genome stability. Although the precise mechanisms linking Htz1 to stress responses in *C. neoformans* remain elusive, the conserved presence of Ino80 and NuA4 complex components suggests that Htz1-mediated chromatin remodeling may underpin these stress-responsive phenotypes. Future characterization of the interactions between Htz1 and its regulatory complexes will further clarify the mechanistic links between chromatin dynamics and pathogenesis.

To ensure the biological authenticity of our findings, we rigorously cross-validated the observed 3D chromatin rearrangements against potential computational aritifacts and normalization biases. We carefully considered whether domain-calling algorithms, such as principle component analysis (PCA) for compartments or insulation score calculations for TADs, might have overly amplified subtle chromatin relaxation. We also considered the possibility of mathematical overestimation inherent in normalization process for Hi-C matrix balancing. To ensure our findings were not computational artifacts, we cross-validated 3D structural changes through rigorous statistical analysis and qRT-PCR, confirming that they correlate with significant alterations in transcriptional regulation. In addition, visualization of contact maps on a Log2 scale revealed distinct, large-scale structural variation in specific regions. Collectively, our validation results demonstrate that these chromatin rearrangements are not mere technical artifacts but represent biologically significant events.

Classical A/B compartment analysis in fungi has been considered challenging due to compact genome size and weak compartmentalization signals [54, 55]. However, the presence of specific histone methylation marks and repeat-rich heterochromatic regions facilitates clear spatial segregation, making this analysis feasible in species such as *F. graminearum* and *Rhizophagus irregularis* [18, 56]. Although the chromosomes of *C. neoformans* are relatively small, the distinct localization of heterochromatin marks, including H3K9me2 and 5-methylcytosine (5mC) allows for clear delination of gene density across the genome [57, 58]. Therefore, A/B compartment analysis could be applied to *C. neoformans*. The current consensus is that transcription and 3D genome architecture engage in bidirectional interactions [59, 60]. In *S. pombe*, modulation of RNA polymerase II activity induces structural alterations in genome organization, suggesting that transcriptional changes can drive reorganization of the 3D genome architecture [61, 62]. Conversely, other studies have reported that deletion of condensin leads to increased DNA damage within actively transcribed regions, supporting the notion that 3D genome reorganization can, in turn, modulate transcriptional activity [34]. Nevertheless, numerous studies have questioned the directional causality underlying this interplay [63–65]. In our study, rather than perturbing local chromatin features such as TADs or enhancers, we directly deleted the core histone gene *HHT2*, encoding a fundamental component of the nucleosome, and observed a global disruption of A/B compartments and TAD structures. Furthermore, we revealed genome-wide transcriptional reprogramming accompanying this structural disruption. These results support a model of directional causality in which the disruption of global chromatin architecture leads to subsequent transcriptional change.

By comparing our results with those of other pathogenic fungi, we demonstrate that 3D genome organization in *C. neoformans* is a decisive factor that regulates virulence. Emerging evidence suggests that 3D chromosomal reorganization provides an essential mechanistic framework for orchestrating the pathogenicity of fungi, particularly through the coordinated expression of virulence-associated effector genes. For instance, in *F. graminearum*, virulence factors are regulated by large-scale, dynamic 3D structural shifts, such as the formation of jet-like chromatin architectures, which represent a level of control beyond traditional transcriptional regulation [18]. Similarly, in the wheat pathogen *Zymoseptoria tritici*, effector genes are physically sequestered within condensed heterochromatin to prevent premature expression, only to undergo rapid 3D decondensation for explosive spatiotemporal activation upon host colonization [66]. This structural logic is further exemplified by the function of chromatin interacting domains (CIDs) as fundamental co-regulatory units [67]. Consistent with this paradigm, our study demonstrates that the loss of *HHT2* leads to global genomic structural perturbations, which leads to increased environmental susceptibility and attenuated virulence. Collectively, these findings position 3D genome architecture not merely as a scaffold, but as a decisive ‘physical switch’ indispensable for the spatiotemporal orchestration of fungal virulence.

In conclusion, our study provides an integrative framework for understanding the fundamental interplay among histone regulation, 3D genome architecture, and fungal virulence. While the current spatial analysis was centered on the *hht2*Δ mutant, the direct correlation between nucleosome occupancy and these structural shifts warrants further exploration through future ChIP-seq studies. Additionally, the specific molecular underpinnings of the unique, Chr10-specific structural collapse represent an intriguing avenue for subsequent investigation. This work establishes for the first time that histones dictate stress responses and virulence not merely through local gene control, but by orchestrating large-scale interactions across entire chromosomes in the fungal pathogen. Ultimately, a comprehensive view of histone-mediated interactions may provide key insights into the structural mechanisms that coordinate fungal virulence.

## Conclusions

In summary, this study provides a comprehensive blueprint of the regulatory interplay among histone dynamics, 3D genome architecture, and fungal pathogenesis in *C. neoformans*. We identified core histones as essential architects of the genome, whose expression is precisely orchestrated by the Rad53 checkpoint kinase. Specifically, our findings demonstrate that the H3 paralog Hht2 serves as a critical structural scaffold; its loss induces a cascade of spatial rearrangements, including A/B compartment switching and TAD boundary shifts. This integrated systems-level approach establishes a directional causality, in which chromatin structural collapse drives the transcriptional reprogramming required for environmental fitness. Beyond providing fundamental insights into fungal biology, these results highlight 3D genome organization as a master regulator of virulence. Ultimately, these findings establish a paradigm in which 3D genome architecture acts as a decisive physical switch, linking nucleosome-level structural integrity to the global transcriptional reprogramming required for eukaryotic environmental adaptation and survival.

## Methods

### Strains, growth condition, and stress susceptibility test

Strains used in this study are listed in the supplementary information (Additional file 1: Table S1). These strains were cultured on a yeast extract-peptone-dextrose (YPD) medium for stress test. Each strain was inoculated overnight at 30oC in liquid YPD medium. Next, cells from overnight culture were washed, serially diluted (1 to 10^4^), and spotted (3 μl) onto solid YPD medium containing the indicated concentration of DNA damage stress inducers. The strains were further incubated at 30°C for 2-4 days and photographed daily. For the mating assay, the V8 medium contained 5% V8 juice 0.5 g/L of KH_2_PO_4_, and 4% bacto-agar with the pH adjusted to 5 using a KOH solution was used [68]. For the melanin production assay, equivalent numbers of cells were spotted on a dopamine agar medium (1 g L-asparagine, 3 g KH_2_PO_4_, 250⍰mg MgSO_4_, 1⍰mg thiamine, and 100⍰mg _L_-DOPA per liter) with glucose [69]. For the capsule thickness measurement, strains was cultured in Littman’s medium [70]. To induce titan cell formation *in vitro*, cells were incubated in liquid YPD for 16 h at 30°C. Then, each cells were washed twice with minimal media (15mM D-Glucose, 10mM MgSO_4_, 29.4 mM KH_2_PO_4_, 13 mM glycine, 3 μM Thiamine with pH adjusted to 5.5) and suspended in minimal medium with a final concentration of 10^6^ cells/ml [71]. The 1 ml of suspension was incubated at 30°C for 3 days in a 1.5 ml eppendorf tube with cap closed. For both experiment, the diameter of the capsule and the cell body were quantitatively measured using SPOT Advanced software (version 4.6; SPOT imaging, USA) via microscopic observation [72]. Capsule thickness was calculated as the difference between the capsule and cell body diameter. Cells with a body diameter exceeding 10 μm were classified as titan cells [73]. To quantify this population, the diameters of at least 100 cells per strain were measured and the corresponding percentages were calculated.

### Construction of conditional and constitutive promoter-replacement strains

To replace the native *HTA1* (CNAG_06747), *HTB1* (CNAG_06746), *HHT2* (CNAG_06745), *HHF2* (CNAG_07807), and *CSE4* (CNAG_00063) promoters with the copper-regulated *CTR4* promoter, we constructed *HTA1, HTB1, HHT2, HHF2*, and *CSE4* promoter replacement cassettes as follows. The 5’-flanking regions of *HTA1, HTB1, HHT2, HHF2*, and *CSE4* and 5’-exon regions of *HTA1, HTB1, HHT2, HHF2*, and *CSE4* were amplified with primer pairs CTR4-L1/CTR4-L2 and CTR4-R1/CTR4-R2 using WT genomic DNA as a template. The primer pair B354/B355 was used for PCR-amplification of *NAT-CTR4* promoter using a pNAT-CTR4-2 as a template [74]. The *HTA1, HTB1, HHT2, HHF2*, and *CSE4* promoter replacement cassettes were generated by double joint PCR (DJ-PCR) primer pairs CTR4-L1/B1455 and CTR4-R2/B1454 for 5’-and 3’-flanking regions of cassettes, respectively. The promoter replacement cassettes were biolistically inserted into the native locus of *HTA1, HTB1, HHT2, HHF2*, and *CSE4* genes in the background of WT, as previously described [68, 75].

To construct the constitutive *HTA1* and *HTB1* overexpression strains, the native promoters of *HTA1* and *HTB1* were replaced with *TEF1* promoter as follows. The 5’-flanking regions of *HTA1* and *HTB1* and 5’-exon regions of *HTA1* and *HTB1* were amplified with primer pairs OE-L1/OE-L2 and OE-R1/OE-R2 using WT genomic DNA as a template. The NEO-TEF1 promoter were amplified with primer pairs J1940/J1941 using a pNEO-TEF1 as a template. The *HTA1* and *HTB1* promoter replacement cassettes were generated by DJ-PCR primer pairs OE-L1/B1887 and OE-R2/B1886 for 5’-and 3’-flanking regions of cassettes, respectively. The promoter replacement cassettes were biolistically inserted into the native locus of *HTA1* and *HTB1* genes in the background of WT. The qRT-PCR primers for each gene were used to measure overexpressions of *HTA1* and *HTB1* at basal and post-radiation exposure.

To generate *HTZ1* overexpression strains in the background of *P*_*CTR4*_*:HTA1* strain (KW1520), the native promoter of *HTZ1* was replaced with *H3* (CNAG_04828) promoter as follows. In the first round PCR, primer pairs J1801/J1836 and J1837/J1838 were used for amplification of 5’-flanking and 5’-coding regions, respectively, with WT genomic DNA as a template. Primer pairs B354/B4018 were used for amplification of NEO-H3 with a plsmid pNEO-H3 as a template. In the second round PCR, primer pairs J1801/B1887 and J1838/B1886 were used for amplification of 5’- and 3’-regions of the *P*_H3_*:HTZ1* replacement cassettes, respectively. The *P*_H3_*:HTZ1* replacement cassettes were introduced into the native promoter of *HTZ1* in *P*_*CTR4*_*:HTA1* strain. Positive transformants were screened by diagnostic PCR and then the correct genotype of the positive transformants was confirmed by Southern blot analysis. Constitutive overexpression of *HTZ1* was verified by qRT-PCR using *HTZ1* gene-specific primers (J1806/J1807). Stable transformants isolated on the YPD media containing nourseothricin (100 μg/ml) or G418 (100 μg/ml) were screened with diagnostic PCR. The correct genotype of these positive strains were verified with Southern blot analysis using gene-specific probes (Additional file 2: Fig. S10). Primers used in this study were listed in the Table S2 (Additional file 1).

### Construction of gene-deletion strains

To construct *HHF1* and *HHF2* (CNAG_01648 and CNAG_07807, respectively) genes deletion mutants, deletion cassettes were generated using DJ-PCR as follows [68]. The 5’-and 3’-flanking regions of the each gene were amplified using primer pairs L1/L2 and R1/R2, respectively, with *C. neoformans* WT genomic DNA as a template. The selectable marker *NEO* was amplified using primer pairs M13Fe/M13Re with a pJAF1 as a template. The 5’-and 3’-flanking regions of deletion cassettes were amplified using primer pairs L1/B1887 and R2/B1886, respectively, with the 1^st^ PCR product as a template. In the case of *HHT1* (CNAG_04828), *HHT2* (CNAG_06745) and *HTZ1* (CNAG_05221) deletion, the nourseothricin acetyltransferase, *NAT* was used for the selectable marker. The 5’- and 3’-flanking regions of deletion cassettes were PCR-amplified using primer pairs L1/B1455 and R2/B1454, respectively, with the 1^st^ PCR product as a template. The two PCR products were combined with gold microcarrier beads (#1652262, Bio-Rad) and introduced into the WT strain using a biolistic transformation method [68, 75].

For mating assay, to construct *HHT2* genes deletion strains, deletion cassette were introduced into KN99**a** strain through biolistic transformation. To screen for the *hht2*Δ mutants in the KN99**a** background, the *NEO* selectable marker was used. To confirm the gene disruption, Southern blot analysis was performed using a DIG-lebeled gene specific probe (Additional file 2: Fig. S10).

### Construction of heterozygous *HTA1, HTB1*, and *CSE4* deletion strains and essentiality test

To construct the heterozygous *HTA1*/*hta1*Δ, *HTB1*/*htb1*Δ, and *CSE4*/*cse4*Δ strains in the diploid *C. neoformans* WT strain AI187, 5’-and 3’-flanking regions of the each gene were amplified by primer pairs L1/L2 and R1/R2 through PCR with WT genomic DNA as a template. For nourseothricin-resistant marker, primers M13Fe and M13Re were used for PCR-amplification with pNAT as a template. Each gene cassettes with *NAT* resistant marker were generated with double-joint PCR and were introduced into AI187 strain through biolistic transformation. The transformants on the YPD media containing nourseothricin were screened with diagnostic PCR for *NAT* insertion in the target region and another intrinsic region. Next, the correct genotypes of transformants were confirmed with Southern blot analysis with gene-specific probe (Additional file 2: Fig. S10). Each mutant was spotted onto V8 agar medium and incubated on the room temperature, in the dark for 5 weeks to induce meiosis and sporulation. Following dissection of haploid basidiospores from mixed progeny onto YPD agar supplemented with adenine (20 mg/L), spores were incubated at 30°C for 3 days. Subsequently, individual spores were transferred into 96-well plates containing 100 μl of YPD medium. To assess the segregation of four meiotically inherited genetic markers (*NAT, ura5*/*URA5, ade2*/*ADE2*, and *MAT***a**/*MATα*), 3 μl aliquots of each culture were spotted onto selective media, including YPD with nourseothricin (100 μg/ml), YNB with adenine (20 mg/L), and YNB with uracil (40 mg/L) [76, 77].

### Construction of *HHT1, HHT2*, and *HTZ1* complemented strains

To demonstrate the phenotypes of the *hht1*Δ mutants in response to DNA damage stress, complemented strains were generated as follows. The *HHT1* gene fragment containing its promoter, open reading frame (ORF), and terminator was PCR-amplified using primer pair J1791/J1792 harboring SacI/SpeI restriction enzyme sites with WT genomic DNA as the template. The PCR-amplified *HHT1* gene was cloned into a plasmid pJET 1.2 vector to construct a plasmid pJET_HHT1. After confirmed with no errors, the SacI/SpeI-digested pJET_HHT1 inserts were subcloned into a plasmid pTOP-NEO to generate a plasmid pNEO-HHT1. The SalI-digested pNEO-HHT1 was transformed into *hht1*Δ mutants (KW1579 and KW1580).

To confirm the observed phenotype in the *hht2*Δ mutants, *hht2*Δ+*HHT2* complemented strains were generated as follows. The *HHT2* gene fragments including promoter, ORF, and terminator were amplified with Pfu polymerase (p410, enzynomics) using primers containing NotI and ApaI restriction enzyme site, respectively. The amplified *HHT2* gene fragments were treated with NotI and ApaI restriction enzymes. Next, the restriction enzyme-treated *HHT2* gene fragment was cloned into a plasmid pTOP-NEO and we verified DNA sequence errors. The pNEO-HHT2 plasmid was linearized with ClaI and then transformed into *hht2*Δ mutants (KW1654).

To verify the phenotype of *htz1*Δ mutants in response to DNA damage stress, *htz1*Δ +*HTZ1* complemented strains were constructed as follows. The full-length of *HTZ1* gene fragment including its promoter, ORF, and terminator was amplified using primers J1849/J1850 with WT genomic DNA as the template. The PCR-amplified *HTZ1* gene was cloned into a plasmid pJET 1.2 to generate a plasmid pJET-HTZ1. Next, we verified no DNA errors in the pJET-HTZ1, the insert of pJET-HTZ1 with XhoI and NotI restriction enzymes was subcloned into pTOP-NEO to generate a plasmid pNEO-HTZ1. The linearized pNEO-HTZ1 with MluI restriction enzyme was transformed into *htz1*Δ mutants (KW1565 and KW1566).

### Total RNA isolation, cDNA synthesis, and quantitative RT-PCR

To investigate whether expression levels of *HTA1, HTB1, HHT1, HHT2, HHF1, HHF2, HTZ1*, and *CSE4* were regulated by DNA damage pathway, total RNA was isolated form WT, *rad53*Δ, *bdr1*Δ, *mec1*Δ, *tel1*Δ, and *mec1*Δ *tel1*Δ mutants. Strains were cultured in 30 ml liquid YPD medium for 16 h at 30°C. Next, overnight culture was inoculated into 100 ml of fresh YPD medium and adjusted to OD_600_ = 0.2. Then, the cells were further incubated until OD_600_ reached approximately 0.7 ~ 0.8. The 50 ml of cells were pelleted for the zero-time sample and the remaining cells were exposed to radiation (0.5 kGy). After radiation exposure, cells were further grown at 30°C for 30 min. Total RNA was extracted from the dried cells using TRIzol reagent (EasyBlue; intron), as previously described [78]. Next, total RNA was further purified using RNeasy purification kit and cDNA was synthesized using PrimeScript 1^st^ strand cDNA synthesis kit (Takara) with the purified total RNA as a template. Relative expression of targets genes was determined using the 2^−ΔΔCt^ method with *ACT1* gene as an internal control. Statistical differences between samples were analyzed using one-way analysis of variance (ANOVA) with Bonferroni’s multiple comparison test (GraphPad Software Inc., San Diego, CA, USA). Due to high sequence similarity between two genes, we designed each gene-specific qRT-PCR primers and confirmed gene-specific expressions in WT, *hht1*Δ, and *hht2*Δ mutants (Additional file 2: Fig. S11A). Similarly, Due to high sequence similarity between them, we designed and validated gene-specific qRT-PCR primers targeting their 3’-UTR using each deletion mutant (Additional file 2: Fig. S11B).

### Protein extraction and Immunoblotting

Strains were grown in liquid YPD medium overnight at 30°C. The overnight cultures were inoculated into the liquid YPD medium and OD_600_ of culture media was adjusted to the around 0.2. The sub-culture were further incubated until OD_600_ reached approximately 0.8. The 50 ml of cell culture was collected for the zero-time sample and the remaining culture was treated for the indicated time. To harvest samples following irradiation, cultures were first collected at the zero-time point and the remaining cells were exposed to radiation at the indicated dose. After exposure, the irradiated cultures were further incubated at 30 °C for the indicated times. The proteins were extracted from the collected cells, previously described [26]. To monitor H2A phosphorylation levels, a primary anti-histone H2A (phospho S129) antibody (ab15083, Abcam) and a secondary anti-rabbit IgG horseradish peroxidase-conjugated antibody (#7074S, Santa Cruz Biotechnology) were used. To monitor H2A protein levels as the loading control, a primary anti H2A antibody (39945, Active motif) and a secondary anti-rabbit IgG horseradish peroxidase-conjugated antibody were used. The membranes were developed using the ECL system (ChemiDoc Imaging system; BioRad).

### Mutation rate assay

Each strain was cultured in liquid YPD media for 16 h. Next, 10^5^ cells from the grown strains were inoculated into fresh YPD medium and then cultured for 48 h at 25°C. After growth for 48 h, 10^8^ cells were spread onto the solid YNB media containing 5-FOA (1 g/ml) and urcil (0.05 mg/ml). The number of spontaneous colonies was counted, and the mutation rate was determined using statistic analysis [79].

### RNA-seq and data analysis

WT and *hht2*Δ mutant were incubated in YPD liquid medium overnight at 30°C. Next, the grown cells were subcultured in fresh YPD liquid medium, and then further incubated 30 °C until OD_600_ reached approximately 0.7 ~ 0.8. The WT and *hht2*Δ mutant were harvested and lyophilized. Total RNA was extracted from dried cells using Trizol reagent and purified using RNeasy purification kits followed by manufacturer’s protocol. For statistical analysis, three bioloigically independent sets were prepared for each strain. For RNAseq analysis, the purified total RNA concentration was calculated by Quant-IT RiboGreen (Invitrogen, #R11490). To assess the integrity of the total RNA, samples are run on the TapeStation RNA screentape (Agilent, #5067-5576). Only high-quality RNA preparations, with RIN greater than 7.0, were used for RNA library construction. A library was independently prepared with 0.5ug of total RNA for each sample by Illumina TruSeq Stranded Total RNA Library Prep Gold Kit (Illumina, Inc., San Diego, CA, USA, # 20020599). The first step in the workflow involves removing the rRNA in the total RNA using Ribo-Zero rRNA Removal Kit (Human/Mouse/Rat Gold) (Illumina, Inc., San Diego, CA, USA). Following this step, the remaining mRNA is fragmented into small pieces using divalent cations under elevated temperature. The cleaved RNA fragments are copied into first strand cDNA using SuperScript II reverse transcriptase (Invitrogen, #18064014) and random primers. This is followed by second strand cDNA synthesis using DNA Polymerase I, RNase H and dUTP. These cDNA fragments then go through an end repair process, the addition of a single ‘A’ base, and then ligation of the adapters. The products are then purified and enriched with PCR to create the final cDNA library. The libraries were quantified using KAPA Library Quantification kits for Illumina Sequencing platforms according to the qPCR Quantification Protocol Guide (KAPA BIOSYSTEMS, #KK4854) and qualified using the TapeStation D1000 ScreenTape (Agilent Technologies, # 5067-5582). Indexed libraries were then submitted to a Illumina NovaSeq (Illumina, Inc., San Diego, CA, USA), and the paired-end (2×100 bp) sequencing was performed by the Macrogen Incorporated.

After sequencing, data trimming and analysis were performed as described in previous studies [80]. Briefly, reads were processed to remove adapters using Cutadapt v2.4 with Python 3.7.4. The reads were then processed as previously described [81] and aligned to the *C. neoformans* H99 reference genome using Hisat2 v2.2.1 and the Hisat and bowtie2 algorithm. Genome annotation data were obtained from the NCBI FTP server. Hisat2 was executed with the options ‘-p 30’ and ‘—dta −1’ along with default settings. Aligned reads were converted and sorted using Samtools v0.1.19, applying the ‘-Sb -@ 8’ option for converting and the ‘-@ 20 –m 100000000’ option for sorting. Subread 2.0.6 was used for transcript assembly and abundance estimation. Transcript abundance was quantified using FPKM (Fragments Per Kilobase of transcript per Million mapped reads) values [82] and data matrices were generated and analyzed with the ‘isoformswitchanalyzerR.’ R package. Quality control was performed using DEBrowser [83]. Statistical analysis of gene expression was performed using DESeq2 v1.24 [84, 85]. In this study, “genes showing statistically significant changes in expression” were defined as those with a *P*-value < 0.05. Within this group, genes that met an additional stringent cutoff of an absolute |Log_2_FC| ≥ 1 were defined as “differentially expressed genes (DEGs)”. The results were visualized using the Enhanced Volcano package in R v4.1.0 [86]. The redundancy of enriched GO terms was analyzed by identifying gene-level overlaps. The curated GO dataset for *C. neoformans* was retrieved from FungiDB [87]. Intersection analysis of the gene sets associated with each GO term was performed and visualized using the UpSetR package in R v4.1.0 [88].

### Phenotypic characterization of microevolution of *C. neoformans* strains

To analysis phenotypic changes caused by the deletion of histone component during repeated passages, we followed the protocol in previous studies with minor modifications [79, 89]. The WT, *hht1*Δ, *hht2*Δ, *hht1*Δ+*HHT1*, and *hht2*Δ+*HHT2* strains were re-streaked every 3 days for 3 months (approximately more than 1,000 generations) on YPD medium and YPD medium containing nourseothricin (100 μg/ml) or G418 (100 μg/ml). Seven independent passaged strains were obtained from the each tested strain. Phenotypic changes were tested in response to diverse stresses including DNA damage, oxidative stress, and antifungal drug resistance.

### Hi-C library preparation

We performed Hi-C (High-throughput chromosome conformation capture) analysis to generate chromatin conformation capture data by following the well-established Hi-C protocol provided in the Proximo Hi-C Fungi Kit (version 4.0) from Phase Genomics (Seattle, WA) [40]. To prepare the libraries, we cultured each strain and incubated in YPD liquid medium overnight at 30°C. Then, the incubated cells were subcultured in fresh YPD liquid medium, and further incubated 30°C until OD_600_ reached approximately 0.8. Next, two samples were crosslinked using a formaldehyde solution and then harvested and lyophilized. Following the manufacturer’s instructions for the kit, intact cells from two samples were digested using the DPNII restriction enzyme, and proximity ligated with biotinylated nucleotides. This creates chimeric molecules composed of fragments from different regions of the genome that were physically proximal in vivo, but not necessarily genomically proximal. Continuing with the manufacturer’s protocol, molecules were pulled down with streptavidin beads and processed into an Illumina-compatible sequencing library. Resulting libraries were sequenced using paired-end 150 bp reads on an Illumina NovaSeq 6000.

### Hi-C data processing and analysis

The Hi-C dataset was analyzed using the HiCExplorer pipeline for contact map generation and TAD analysis, and the FAN-C framework was employed for A/B compartment analysis [90, 91]. The *C. neoformans* genome (GCF_000149245.1) was used as the reference. The contact maps of individual chromosomes were generated based on 10-kb resolution matrices and other heatmaps containing TADs boundaries were generated based on 1-kb resolution matrices. To ensure comparable read depths across samples, random downsampling was performed to match the minimum total number of valid contacts; specifically, the WT dataset was downsampled to match the depth of the *hht2*Δ sample (27,153,926 valid pairs) (Additional file 3: Dataset 3). Contact matrices were generated and normalized using the ICE method, and standard contact maps were visualized using the hicPlotMatrix module in HiCExplorer. TADs and their boundaries were identified using the hicFindTADs tool with search window parameters set to a minimum depth of 10 kb, a maximum depth of 200 kb, and a step size of 10 kb. Boundary significance was determined by applying a false discovery rate (FDR) correction for multiple testing with a threshold of 0.1. For compartment analysis, FAN-C was used to compute the first principal component (PC1) from the normalized matrix to infer A/B compartments. The positive and negative PC1 values were assigned to A and B compartments, respectively, and this assignment was biologically validated by confirming a global positive correlation with our RNA-seq expression data. All matrix processing, visualization, and plotting were performed using HiCExplorer and FAN-C tools according to the standard workflows. Gene Set Enrichment Analysis (GSEA) was performed using the fgsea package in R. All expressed genes were pre-ranked strictly by their differential expression test statistic (stat value), which intrinsically incorporates both the log2 fold-change and its associated variance. The enrichment directionality and magnitude were evaluated using the Normalized Enrichment Score (NES). Statistical significance was empirically determined via 1,000 permutations, applying a False Discovery Rate (FDR) threshold of *p*-value < 0.05.

### Virulence assay

All animal experiments were approved by the Institutional Animal Care and Use Committee (IACUC) of Jeonbuk National University (approval number: JBNU 2022-092) and were conducted in accordance with relevant national and institutional guidelines. Animal experiments were conducted at the BL3 and ABL3 facilities of the Core Facility Center for Zoonosis Research (Core-FCZR). SPF/VAF-confirmed inbred 6-week-old female BALB/cAnNCRLOri mice were purchased from Orient Bio Inc. (South Korea) and acclimatized for one week prior to experiment. For infection, the strains which we used in this experiment (WT, *hht2*Δ, and *hht2*Δ+*HHT2* for each 10 mice) were cultured in a fresh liquid YPD medium and incubated overnight at 30°C. Cells were harvested, washed, and adjusted to a concentration of 5 × 10^5^ CFU per mouse. Mice were anesthetized with isoflurane vapor (Hana Pharm. Co., Ltd., South Korea) at a flow rate of 80 cc/min, and the strains were inoculated intranasally. Following infection, mice were monitored daily for survival, and survival rates were expressed as percentages. For fungal burden analysis, lungs and brains were harvested at day 14 post-infection, weighted, homogenized, and serial dilutions were plated onto YPD agar. CFU were enumerated after incubation. Survival data were analyzed using the log-rank (Mantel-Cox) test (n = 3 mice per group). Fungal burden data were log_10_-transformed and analyzed using one-way ANOVA followed Tukey’s multiple comparisons test. Statistical analyses were performed using Graphpad Prism version 9.5.1.

### Structural Analysis and RMSD calculation

The structures of the Hta1-Htb1 heterodimer, H3-H4 tetramer, Htz1, and Cse4 from *C. neoformans* were predicted using AlphaFold3 [92]. Protein sequences were retrieved from FungiDB, while the corresponding structures for *H. sapiens* and *S. cerevisiae* were obtained from the RCSB Protein Data Bank (PDB). Structural alignment and Root Mean Square Deviation (RMSD) calculations were performed using PyMOL (The PyMOL Molecular Graphics System, Version 2.0). The RMSD values were calculated across Cα atoms to evaluate the structural divergence between the predicted *C. neoformans* models and their homologous counterpart.

## Acknowledgment

We thanks Macrogen for helping the RNAseq data submission to NCBI and PHYZEN for helping the Hi-C data submission to NCBI.

## Author’s contribution

K.-W. J. and Y.-S. B. conceptualized the study and designed the experimental framework. S.-H. K., T.-M. K., Y.-B. J., S.-R. Y., E.-S. K., S.-Y. L., J.-H. J. and K.-T. L. performed the experiments, analysed the data, and wrote the manuscript. All the authors agree with the published version of the manuscript.

## Funding

This work was supported by the Nuclear R&D program of Ministry of Science and Information and Communications Technologies (ICT) (Republic of Korea) (523510-26) to K.-W. J. This work was supported by the National Research Foundation of Korea (NRF) grant funded by the Korean government (MSIT) (RS-2022-NR072215) to K.-T. L. and (RS-2025-00555365, RS-2025-02215093, and RS-2025-18362970 to Y.-S.B.).

## Data availability

We will provide any strain and materials used in this study upon request. RNA-seq-based transcriptome profiling data for WT and *hht2*Δ mutant from this publication were deposited to Gene Expression Omnibus (GEO) under accession number GSE315909. The raw and processed Hi-C data from this publication have been deposited to the GEO under accession number GSE317247

## Declarations

### Ethics approval and consent to participate

Not applicable.

### Consent for publication

Not applicable.

### Competing interests

The authors declare that have no competing interests.

## Supplemenatary information

**Table S1** Strains used in this study

**Table S2** Primers used in this study

**Fig. S1** Structural prediction and meiotic progeny analysis for assessing the essentiality of Htz1 and Cse4. **A and B** Structure prediction of histone variants, Htz1 **(A)** and Cse4 **(B). C and D** PCR analysis to characterize the marker that segregated during meiosis (*MATα* or *MAT***a**) in progeny obtained from the sporulation of the heterozygous *HTA1*/*hta1*Δ **(A)** and *HTB1*/*htb1*Δ **(B)**. 50 basidiospore progeny were isolated from each mutant and were grown on four types of medium (YPD, +NAT, +adenine and +uracil) for genetic analysis. **E** The constitutive overexpression of *HTZ1* in the *P*_*CTR4*_*:HTA1* strain. Error bars indicate standard errors of the means. Statistical significance of difference was determined by one-way ANOVA with Bonferroni’s multiple-comparison test (***: *P* < 0.001 and NS: not significant). **F** *CSE4* was required for cell growth. The overnight-cultured WT and P_CTR4:_*CSE4* (KW1559 and KW1562) strains were spotted onto YNB (or YPD) medium containing the BCS (200 μM) or CuSO_4_ (25 μM). **G** PCR analysis to characterize the marker that segregates during meiosis (*MATα* or *MAT***a**) in progeny obtained from the sporulation of the heterozygous *CSE4*/*cse4*Δ. **H** 50 basidiospore progeny were isolated from each mutant and were grown on four types of medium (YPD, +NAT, +adenine and +uracil) for genetic analysis. Basidiospore progeny isolated from the heterozygous *CSE4*/*cse4*Δ strain were spotted onto nourseothricin medium, followed by PCR analysis of *NAT* progeny (#47, #57, #60, and #62) using internal PCR.

**Fig. S2** The expression levels of histone variant genes. **A** Sequence alignment of the phosphorylation region of Hta1 and Hta2 in *S. cerevisiae, Schizosaccharomyces pombe, Candida albicans* and Hta1 of *Aspergillus nidulans* and *C. neoformans*. **B** The phosphorylation of Hta1 in response to radiation. The WT strain was grown to the mid-logarithmic phase and exposed to the radiation (3 kGy) and further incubated for each indicated time. **C-F** The expression of *HTZ1* or *CSE4* post-radiation (0.5 kGy) exposure in strains belonging to the DNA repair pathway. Three independent biological experiments with duplicate technical replicates were performed.

**Fig. S3** Phenotypic profiling of histone mutants in response to diverse DNA damaging agents. **A** Expression levels of *HTA1, HTB1*, and *TEF1* post-radiation exposure (3 kGy). **B** The constitutive expression of *HTA1* and *HTB1* in the P_*TEF1:*_*HTA1* and P_*TEF1:*_*HTB1* strains post radiation exposure (3 kGy). **C** The compensation of *HHT1* expression in the loss of *HHT2*. The qRT-PCR analysis was used to quantify expression levels of *HHT1* and *HHT2* in WT, *hht1*Δ, *hht2*Δ mutant and their complemented strains. **D and F** The strains were spotted onto YPD medium containing MMS, 4-NQO, cisplatin, and bleomycin. For UV-C and γ-radiation exposures, the serially diluted cells were spotted and then were exposed to UV-C or γ-radiation. The two images split by a horizontal white line in each spot assay were obtained from the same plate. **E** The compensation of *HHF1* and *HHF2* expression upon loss of *HHF2* and *HHF1*, respectively. The expression levels of *HHF1* or *HHF*2 were measured in the WT, *hhf2*Δ and *hhf1*Δ mutants using qRT-PCR with gene-specific primers for *HHF1* and *HHF2* genes. For qRT-PCR analysis, three independent biological experiments with duplicate technical replicates were performed. Statistical analysis of difference was determined by one-way ANOVA with Bonferroni’s multiple-comparison test (NS: not significant).

**Fig. S4** The deletion of Histone H3 led to increased phenotypic changes. **A** Quantification of spontaneous 5-FOA-resistant rates in H3 mutants. Error bars indicate standard errors of means (NS: not significant). **B** The phenotypic changes of the original WT strain, *hht1*Δ, *hht1*Δ+*HHT1, hht2*Δ, and *hht2*Δ+*HHT2* mutants compared to those of its corresponding strains after seven independent passages (1,000 generations) in response to diverse stress responses. Each strain was cultured in liquid YPD medium at 30°C. The cultured cells were serially diluted (1 to 10^4^) and spotted on the YPD plates containing 0.035 % MMS, 5 mM cisplatin, 4 μg/ml benomyl, 0.8 mM tBOOH, 0.025 mM MD, 16 μg/ml FCZ or 600 μg/ml 5-FC. Cells were further incubated at 30°C and photographed daily for 3 days.

**Fig. S5** Hht2, unlike Hht1, Hhf1, or Hhf2, contributed to the DNA damage response through both Rad53-dependent and independent mechanisms. **A** The strains (WT, *rad53*Δ, *hht1*Δ, *hht2*Δ, *hhf1*Δ, *hhf2*Δ, *rad53*Δ *hht1*Δ, *rad53*Δ *hht2*Δ, *rad53*Δ *hhf1*Δ and *rad53*Δ *hhf2*Δ) were spotted onto YPD medium containing 0.04 % MMS, 0.04 μg/ml 4-NQO, 5 mM cisplatin, 4 μg/ml bleomycin, 10 μg/ml TBZ or 3.5 μg/ml benomyl. For UV-C and γ-radiation exposures, the serially diluted cells were spotted and then were exposed to 100 J/m^2^ UV-C or 0.5 kGy γ-radiation. The two images split by a horizontal white line in each spot assay were obtained from the same plate. **B** The grown strains were serially diluted (1 to 10^5^) cells were spotted onto YPD media containing the indicated concentrations of chemicals and were further incubated at 30°C for 4 days.

**Fig. S6** Phenotypic analysis of core histone and histone variant mutants for diverse stress responses. **A-F** The cultured cells were serially diluted (1 to 10^4^) and spotted on the YPD plates containing stress-inducing chemicals. Cells were further incubated at 30°C and photographed daily for 3 days.

**Fig. S7** Virulence factor formation in histone mutants. **A** Representative images of strains cultured on Littman medium for capsule production at 30°C for 3 days. The scale bar indicates 10 μm. **B** The strains were cultured on Littman medium for capsule production at 30°C for 3 days or for the indicated days. Statistical analysis was conducted using one-way ANOVA.

**Fig. S8** The expression levels of ergosterol synthesis genes in WT and *hht2*Δ strains. Three independent biological experiments with duplicate technical replicates were performed. Error bars indicate standard errors of the means. Statistical significance of difference was determined by one-way ANOVA with Bonferroni’s multiple-comparison test (NS: not significant).

**Fig. S9** Hht2 affected the maintenance of the 3D chromatin structure. **A** Heatmaps (10-kb resolution) showing the chromosomal interactions from chromosome 1 to chromosome 14 for WT and *hht2*Δ mutant. The panels show ICE-normalized contact matrices, with the top panels representing WT and the bottom panels representing the *hht2*Δ strain. **B** Average contact probability (CP) as a function of genomic distance in WT and *hht2*Δ mutant strains. **C** 2 × 2 contingency tables showing the number of genes used for Fisher’s exact test to evaluate expression changes in transitioning (A to B and B to A) versus stable (A to A and B to B) compartments. **D** Directional correlation between A/B compartment switching and gene expression changes. Stacked bar graphs showed the proportion of genes showing statistically significant changes in expression (*P* < 0.05) across different compartment transitions. The left panel illustrated the percentage of down-regulated genes (blue) versus non-down-regulated genes (grey) in regions undergoing A to B transitions compared to A to A regions. The right panel displays the percentage of up-regulated genes (red) versus non-up-regulated genes (grey) within B to A transition regions compared to stable B to B regions. Statistical significance was determined using Fisher’s exact test (****: *P* < 0.0001). **E** Contact maps and corresponding TAD annotations of WT and *hht2*Δ mutants across chromosome 1 to 14, generated using the hicFindTADs tool. **F** Venn diagrams described the overlap among DEGs DEGs (*P* < 0.05, |Log_2_FC| ≥ 1), genes located in TAD-altered regions and genes associated with A/B compartments changes, based on comparative analyses between the WT and *hht2*Δ mutant. **G** Calculative relative proximity between indicated genes in WT.

**Fig. S10** Genotypic analysis of the *P*_*CTR4*_*:HTA1, P*_*CTR4*_*:HTB1, P*_*CTR4*_*:CSE4, hht1*Δ, *hta1*Δ, *htb1*Δ, *hht2*Δ, *hhf1*Δ, *hhf2*Δ, *htz1*Δ, *hhf1*Δ *P*_*CTR4*_*:HHF2, P*_*CTR4*_*:HTA1 P*_H3_*:HTZ1, hht1*Δ *P*_*CTR4*_*:HHT2, P*_TEF1_*:HTA1, P*_TEF1_*:HTB1, rad53*Δ *hht1*Δ, *rad53*Δ *hht2*Δ, *rad53*Δ *hhf1*Δ and *rad53*Δ *hhf2*Δ mutants. Gene disruption strategies were illustrated in the left diagram **A, C, E, G, I, K, M, O, Q, S, U, W, Y, AA, AG, AI, AK, AM, AO** The correct gene disruptions were confirmed by Southern blot analysis using genomic DNA digested with the indicated restriction enzymes. **B, D, F, H, J, L, N, P, R, T, V, X, Z, AB, AC, AD, AE, AF, AH, AJ, AL, AN, and AP**.

**Fig. S11** Designed primers for *HHT1, HHT2, HHF1*, and *HHF2* were validated using a gene expression assay. **A and B** The cDNA was synthesized from total RNAs isolated from strains (WT, *hht1*Δ, *hht2*Δ, *hhf1*Δ, and *hhf2*Δ mutants). Independent biological experiments with duplicate technical replicates were performed two or three times. Individual dots represent each biological replicate (n = 2–3) Error bars indicate standard errors of the means. Statistical significance of difference was determined by one-way ANOVA with Bonferroni’s multiple-comparison test (NS: not significant).

### Supplementary Dataset 1

Transcriptome profiles of WT and *hht2*Δ mutants under basal conditions

### Supplementary Dataset 2

List of DEGs genes overlapping with TADs and A/B compartments

### Supplementary Dataset 3

Validated chromatin interaction pairs identified by Hi-C analysis

## Notes

### Competing Interest Statement

The authors have declared no competing interest.

