## Supplementary Table 1 for "Systematic histone mutagenesis reveals nucleosome-dependent maintenance of three-dimensional chromosome architecture and virulence in *Cryptococcus neoformans*"

**Table S1. Strains used in this study**

| **Strain** | **Genotype** | **Parent** | **Reference** |
| --- | --- | --- | --- |
| H99 | *MATα* |  | [1] |
| KW137 | *MATα CNAG_02589 (BDR1)::NAT #230* | H99 | [2] |
| KW191 | *MATα CNAG_03167 (CHK1)::NAT#234* | H99 | [3] |
| KW250 | *MATα CNAG_05216 (RAD53)::NAT#184 CNAG_03167 (CHK1)::NEO* | YSB3785 | [3] |
| YSB3785 | *MATα CNAG_05216 (RAD53)::NAT#184* | H99 | [4] |
| YSB3611 | *MATα CNAG_02233 (MEC1)::NAT#204* | H99 | [4] |
| YSB3844 | *MATα CNAG_05771 (TEL1)::NAT#225* | H99 | [4] |
| KW480 | *MATα CNAG_02233 (MEC1)::NAT#204 CNAG_05771 (TEL1)::NEO* | YSB3611 | [3] |
| KW1519 | *MATα P_CTR4:_CNAG_06747 (HTA1)-NAT* | H99 | This study |
| KW1520 | *MATα P_CTR4:_CNAG_06747 (HTA1)-NAT* | H99 | This study |
| KW1521 | *MATα P_CTR4:_CNAG_06747 (HTA1)-NAT* | H99 | This study |
| KW1522 | *MATα P_CTR4:_CNAG_06747 (HTA1)-NAT* | H99 | This study |
| KW1530 | *MATα P_CTR4:_CNAG_06746 (HTB1)-NAT* | H99 | This study |
| KW1532 | *MATα P_CTR4:_CNAG_06746 (HTB1)-NAT* | H99 | This study |
| KW1533 | *MATα P_CTR4:_CNAG_06746 (HTB1)-NAT* | H99 | This study |
| KW2167 | *MATα CNAG_06747 (HTA1)::NAT#43* | AI187 | This study |
| KW2168 | *MATα CNAG_06747 (HTA1)::NAT#43* | AI187 | This study |
| KW2169 | *MATα CNAG_06747 (HTA1)::NAT#43* | AI187 | This study |
| KW2170 | *MATα CNAG_06747 (HTA1)::NAT#43* | AI187 | This study |
| KW1929 | *MATα CNAG_06746 (HTB1)::NAT#43* | AI187 | This study |
| KW1930 | *MATα CNAG_06746 (HTB1)::NAT#43* | AI187 | This study |
| KW1931 | *MATα CNAG_06746 (HTB1)::NAT#43* | AI187 | This study |
| KW1730 | *MATα P_TEF1:_CNAG_06747 (HTA1)-NAT* | H99 | This study |
| KW1731 | *MATα P_TEF1:_CNAG_06747 (HTA1)-NAT* | H99 | This study |
| KW1947 | *MATα P_TEF1:_CNAG_06747 (HTA1)-NAT* | H99 | This study |
| KW2146 | *MATα P_TEF1:_CNAG_06747 (HTA1)-NAT* | H99 | This study |
| KW1732 | *MATα P_TEF1:_CNAG_06746 (HTB1)-NAT* | H99 | This study |
| KW1733 | *MATα P_TEF1:_CNAG_06746 (HTB1)-NAT* | H99 | This study |
| KW1734 | *MATα P_TEF1:_CNAG_06746 (HTB1)-NAT* | H99 | This study |
| KW1735 | *MATα P_TEF1:_CNAG_06746 (HTB1)-NAT* | H99 | This study |
| KW1565 | *MATα CNAG_05221 (HTZ1)::NAT#43* | H99 | This study |
| KW1566 | *MATα CNAG_05221 (HTZ1)::NAT#43* | H99 | This study |
| KW1567 | *MATα CNAG_05221 (HTZ1)::NAT#43* | H99 | This study |
| KW1568 | *MATα CNAG_05221 (HTZ1)::NAT#43* | H99 | This study |
| KW1632 | *MATα CNAG_05221 (HTZ1)::NAT#43 pNEO-CNAG_05221* | KW1566 | This study |
| KW1626 | *MATα P_CTR4:_CNAG_06747 (HTA1)-NAT P_H3:_CNAG_05221 (HTZ1)-NEO* | KW1520 | This study |
| KW1627 | *MATα P_CTR4:_CNAG_06747 (HTA1)-NAT P_H3:_CNAG_05221 (HTZ1)-NEO* | KW1520 | This study |
| KW1628 | *MATα P_CTR4:_CNAG_06747 (HTA1)-NAT P_H3:_CNAG_05221 (HTZ1)-NEO* | KW1520 | This study |
| KW1629 | *MATα P_CTR4:_CNAG_06747 (HTA1)-NAT P_H3:_CNAG_05221 (HTZ1)-NEO* | KW1520 | This study |
| KW1759 | *MATα CNAG_05216 (RAD53)::NAT#184 CNAG_05221 (HTZ1)::NEO* | YSB3785 | This study |
| KW1760 | *MATα CNAG_05216 (RAD53)::NAT#184 CNAG_05221 (HTZ1)::NEO* | YSB3785 | This study |
| KW1761 | *MATα CNAG_05216 (RAD53)::NAT#184 CNAG_05221 (HTZ1)::NEO* | YSB3785 | This study |
| KW1762 | *MATα CNAG_05216 (RAD53)::NAT#184 CNAG_05221 (HTZ1)::NEO* | YSB3785 | This study |
| KW1540 | *MATα CNAG_04828 (HHT1)::NEO* | H99 | This study |
| KW1541 | *MATα CNAG_04828 (HHT1)::NEO* | H99 | This study |
| KW1542 | *MATα CNAG_04828 (HHT1)::NEO* | H99 | This study |
| KW1543 | *MATα CNAG_04828 (HHT1)::NEO* | H99 | This study |
| KW1579 | *MATα CNAG_04828 (HHT1)::NAT* | H99 | This study |
| KW1580 | *MATα CNAG_04828 (HHT1)::NAT* | H99 | This study |
| KW1581 | *MATα CNAG_04828 (HHT1)::NAT* | H99 | This study |
| KW1667 | *MATα CNAG_04828 (HHT1)::NAT pNEO-CNAG_04828* | KW1579 | This study |
| KW1654 | *MATα CNAG_06745 (HHT2)::NAT* | H99 | This study |
| KW1655 | *MATα CNAG_06745 (HHT2)::NAT* | H99 | This study |
| KW1656 | *MATα CNAG_06745 (HHT2)::NAT* | H99 | This study |
| KW1657 | *MATα CNAG_06745 (HHT2)::NAT* | H99 | This study |
| KW1738 | *MATα CNAG_06745 (HHT2)::NAT pNEO-CNAG_06745* | KW1654 | This study |
| KW1620 | *MATα CNAG_04828 (HHT1)::NEO P_CTR4:_CNAG_06745 (HHT2)-NAT* | KW1540 | This study |
| KW1621 | *MATα CNAG_04828 (HHT1)::NEO P_CTR4:_CNAG_06745 (HHT2)-NAT* | KW1540 | This study |
| KW1622 | *MATα CNAG_04828 (HHT1)::NEO P_CTR4:_CNAG_06745 (HHT2)-NAT* | KW1540 | This study |
| KW1623 | *MATα CNAG_04828 (HHT1)::NEO P_CTR4:_CNAG_06745 (HHT2)-NAT* | KW1540 | This study |
| KW1935 | *MAT*a *CNAG_06745 (HHT2)::NEO* | KN99 | This study |
| KW1936 | *MAT*a *CNAG_06745 (HHT2)::NEO* | KN99 | This study |
| KW1937 | *MAT*a *CNAG_06745 (HHT2)::NEO* | KN99 | This study |
| KW1536 | *MATα CNAG_01648 (HHF1)::NEO* | H99 | This study |
| KW1537 | *MATα CNAG_01648 (HHF1)::NEO* | H99 | This study |
| KW1538 | *MATα CNAG_01648 (HHF1)::NEO* | H99 | This study |
| KW1544 | *MATα CNAG_07807 (HHF2)::NEO* | H99 | This study |
| KW1545 | *MATα CNAG_07807 (HHF2)::NEO* | H99 | This study |
| KW1546 | *MATα CNAG_07807 (HHF2)::NEO* | H99 | This study |
| KW1590 | *MATα CNAG_07807 (HHF2)::NEO* | H99 | This study |
| KW1591 | *MATα CNAG_07807 (HHF2)::NEO* | H99 | This study |
| KW1592 | *MATα CNAG_07807 (HHF2)::NEO* | H99 | This study |
| KW1594 | *MATα CNAG_01648 (HHF1)::NEO P_CTR4:_CNAG_07807 (HHF2)-NAT* | KW1536 | This study |
| KW1595 | *MATα CNAG_01648 (HHF1)::NEO P_CTR4:_CNAG_07807 (HHF2)-NAT* | KW1536 | This study |
| KW1596 | *MATα CNAG_01648 (HHF1)::NEO P_CTR4:_CNAG_07807 (HHF2)-NAT* | KW1536 | This study |
| KW1597 | *MATα CNAG_01648 (HHF1)::NEO P_CTR4:_CNAG_07807 (HHF2)-NAT* | KW1536 | This study |
| KW1559 | *MATα P_CTR4:_CNAG_00063 (CSE4)-NAT* | H99 | This study |
| KW1562 | *MATα P_CTR4:_CNAG_00063 (CSE4)-NAT* | H99 | This study |
| KW1563 | *MATα P_CTR4:_CNAG_00063 (CSE4)-NAT* | H99 | This study |
| KW1564 | *MATα P_CTR4:_CNAG_00063 (CSE4)-NAT* | H99 | This study |
| KW2161 | *MATα CNAG_00063 (CSE4)::NAT#43* | AI187 | This study |
| KW2162 | *MATα CNAG_00063 (CSE4)::NAT#43* | AI187 | This study |
| KW1711 | *MATα CNAG_05216 (RAD53)::NAT#184 CNAG_04828 (HHT1)::NEO* | YSB3785 | This study |
| KW1744 | *MATα CNAG_05216 (RAD53)::NAT#184 CNAG_04828 (HHT1)::NEO* | YSB3785 | This study |
| KW1745 | *MATα CNAG_05216 (RAD53)::NAT#184 CNAG_04828 (HHT1)::NEO* | YSB3785 | This study |
| KW1754 | *MATα CNAG_05216 (RAD53)::NAT#184 CNAG_06745 (HHT2)::NEO* | YSB3785 | This study |
| KW1781 | *MATα CNAG_05216 (RAD53)::NAT#184 CNAG_06745 (HHT2)::NEO* | YSB3785 | This study |
| KW1706 | *MATα CNAG_05216 (RAD53)::NAT#184 CNAG_01648 (HHF1)::NEO* | YSB3785 | This study |
| KW1707 | *MATα CNAG_05216 (RAD53)::NAT#184 CNAG_01648 (HHF1)::NEO* | YSB3785 | This study |
| KW1708 | *MATα CNAG_05216 (RAD53)::NAT#184 CNAG_01648 (HHF1)::NEO* | YSB3785 | This study |
| KW1709 | *MATα CNAG_05216 (RAD53)::NAT#184 CNAG_01648 (HHF1)::NEO* | YSB3785 | This study |
| KW1710 | *MATα CNAG_05216 (RAD53)::NAT#184 CNAG_07807 (HHF2)::NEO* | YSB3785 | This study |
| KW1713 | *MATα CNAG_05216 (RAD53)::NAT#184 CNAG_07807 (HHF2)::NEO* | YSB3785 | This study |
| KW1749 | *MATα CNAG_05216 (RAD53)::NAT#184 CNAG_07807 (HHF2)::NEO* | YSB3785 | This study |
| KW2228 | *MATα (H99) 30passaged strain* | H99 | This study |
| KW2229 | *MATα (H99) 30passaged strain* | H99 | This study |
| KW2230 | *MATα (H99) 30passaged strain* | H99 | This study |
| KW2231 | *MATα (H99) 30passaged strain* | H99 | This study |
| KW2232 | *MATα (H99) 30passaged strain* | H99 | This study |
| KW2233 | *MATα (H99) 30passaged strain* | H99 | This study |
| KW2234 | *MATα (H99) 30passaged strain* | H99 | This study |
| KW2235 | *MATα CNAG_04828::NAT 30 passaged* | KW1579 | This study |
| KW2236 | *MATα CNAG_04828::NAT 30 passaged* | KW1579 | This study |
| KW2237 | *MATα CNAG_04828::NAT 30 passaged* | KW1579 | This study |
| KW2238 | *MATα CNAG_04828::NAT 30 passaged* | KW1579 | This study |
| KW2239 | *MATα CNAG_04828::NAT 30 passaged* | KW1579 | This study |
| KW2240 | *MATα CNAG_04828::NAT 30 passaged* | KW1579 | This study |
| KW2241 | *MATα CNAG_04828::NAT 30 passaged* | KW1579 | This study |
| KW2242 | *MATα CNAG_04828::NAT pNEO-CNAG_04828 30 passaged* | KW1667 | This study |
| KW2230 | *MATα CNAG_04828::NAT pNEO-CNAG_04828 30 passaged* | KW1667 | This study |
| KW2231 | *MATα CNAG_04828::NAT pNEO-CNAG_04828 30 passaged* | KW1667 | This study |
| KW2232 | *MATα CNAG_04828::NAT pNEO-CNAG_04828 30 passaged* | KW1667 | This study |
| KW2233 | *MATα CNAG_04828::NAT pNEO-CNAG_04828 30 passaged* | KW1667 | This study |
| KW2234 | *MATα CNAG_04828::NAT pNEO-CNAG_04828 30 passaged* | KW1667 | This study |
| KW2235 | *MATα CNAG_04828::NAT pNEO-CNAG_04828 30 passaged* | KW1667 | This study |
| KW2236 | *MATα hht2Δ(CNAG_06745)::NAT 30 passaged* | KW1654 | This study |
| KW2237 | *MATα hht2Δ(CNAG_06745)::NAT 30 passaged* | KW1654 | This study |
| KW2238 | *MATα hht2Δ(CNAG_06745)::NAT 30 passaged* | KW1654 | This study |
| KW2239 | *MATα hht2Δ(CNAG_06745)::NAT 30 passaged* | KW1654 | This study |
| KW2240 | *MATα hht2Δ(CNAG_06745)::NAT 30 passaged* | KW1654 | This study |
| KW2241 | *MATα hht2Δ(CNAG_06745)::NAT 30 passaged* | KW1654 | This study |
| KW2242 | *MATα hht2Δ(CNAG_06745)::NAT 30 passaged* | KW1654 | This study |
| KW2243 | *MATα hht2Δ(CNAG_06745)::NAT pNEO-CNAG_06745 30 passaged* | KW1738 | This study |
| KW2244 | *MATα hht2Δ(CNAG_06745)::NAT pNEO-CNAG_06745 30 passaged* | KW1738 | This study |
| KW2245 | *MATα hht2Δ(CNAG_06745)::NAT pNEO-CNAG_06745 30 passaged* | KW1738 | This study |
| KW2246 | *MATα hht2Δ(CNAG_06745)::NAT pNEO-CNAG_06745 30 passaged* | KW1738 | This study |
| KW2247 | *MATα hht2Δ(CNAG_06745)::NAT pNEO-CNAG_06745 30 passaged* | KW1738 | This study |
| KW2248 | *MATα hht2Δ(CNAG_06745)::NAT pNEO-CNAG_06745 30 passaged* | KW1738 | This study |
| KW2249 | *MATα hht2Δ(CNAG_06745)::NAT pNEO-CNAG_06745 30 passaged* | KW1738 | This study |

Each *NAT-STM#* indicates the Nat^r^ marker with a unique signature tag.
