## Supplementary Table 2 for "Systematic histone mutagenesis reveals nucleosome-dependent maintenance of three-dimensional chromosome architecture and virulence in *Cryptococcus neoformans*"

**Table S2. Primers used in this study**

| Primer Name | | Sequence (5’—3’) | Comment |
| --- | --- | --- | --- |
| B79 | TGTGGATGCTGGCGGAGGATA | | Screening primer on ACT promoter |
| B1026 | GTAAAACGACGGCCAGTGAGC | | M13 forward (extended) |
| B1027 | CAGGAAACAGCTATGACCATG | | M13 reverse (extended) |
| B1454 | AAGGTGTTCCCCGACGACGAATCG | | NSL-NAT |
| B1455 | AACTCCGTCGCGAGCCCCATCAAC | | NSR-NAT |
| B1886 | TGGAAGAGATGGATGTGC | | NSL-NEO |
| B1887 | ATTGTCTGTTGTGCCCAG | | NSR-NEO |
| J1940 | GGAGCCATGAAGATCCTGAGGA | | Primer 1 for *TEF1* promoter amplification |
| J1941 | CCACAATTTTTTTCGAAACCTC | | Primer 1 for *TEF1* promoter amplification |
| J940 | TTCCCACCCTCAGCAACGCC | | Screening primer on TRP terminator |
| B354 | GCATGCAGGATTCGAGTG | | CTR4-NAT primer 1 |
| B355 | GATTGGTGAAGTCGTTGTCG | | CTR4-NAT primer 1 |
| J1710 | ACATCTCTTGCCGTCAGG | | *HTA1* –3’ screening primer for *CTR4-NAT* |
| J1680 | AGAGAATGCTGCCCAGGTG | | *HTA1*– left flanking primer 1 for *CTR4-NAT* |
| J1681 | GCCACTCGAATCCTGCATGCAGACTTGTTGGATGGAGGGGA | | *HTA1*– left flanking primer 2 for *CTR4-NAT* |
| J1682 | CGACAACGACTTCACCAATCTTCCAACAACCCTTCCACTA | | *HTA1*– right flanking primer 1 for *CTR4-NAT* |
| J1690 | GCTTGTCGTTCTTCACACAG | | *HTA1*– right flanking primer 2 for *CTR4-NAT* |
| J1127 | CTTCTTGGCACCGTCTTTG | | *HTA1*– probe primer 1 for Southern blot for *CTR4-NAT* |
| J1132 | AGCTTGCGTTTCGGTGAC | | *HTA1*– probe primer 2 for Southern blot for *CTR4-NAT* |
| J1743 | TGAAAGGAGCATCGGGCAC | | *HHT1* –5’ screening primer for deletion |
| J1744 | GCCACGAGATGGTGTTGAG | | *HHT1* – left flanking primer 1 for deletion |
| J1745 | ATTCACTGGCCGTCGTTTTACGTGATGTCGGGTGGAGATG | | *HHT1* – left flanking primer 2 for deletion |
| J1746 | CATGGTCATAGCTGTTTCCTGTTCATCTGCTCTCCGCAGG | | *HHT1* – right flanking primer 1 for deletion |
| J1747 | TACATGGCGCATTTGGCAC | | *HHT1* – right flanking primer 2 for deletion |
| J1748 | ATTTCTCGGAGAGCGACG | | *HHT1* – probe primer for Southern blot for deletion |
| J1751 | TAGCGTCTTGGATGGGTG | | *HHF1* –5’ screening primer for deletion |
| J1752 | TCAGCTCTCATTCCGGCTC | | *HHF1* – left flanking primer 1 for deletion |
| J1753 | ATTCACTGGCCGTCGTTTTACAACCCGCCGGGAATTACAC | | *HHF1* – left flanking primer 2 for deletion |
| J1754 | CATGGTCATAGCTGTTTCCTGCCGCGCTTACACAGTTTAC | | *HHF1* – right flanking primer 1 for deletion |
| J1755 | AGCCAACTTACAGAGGGA | | *HHF1* – right flanking primer 2 for deletion |
| J1756 | AAGGGCATAGACAACGTCG | | *HHF1* – probe primer for Southern blot for deletion |
| J1759 | GAGAAGCGGGTGTACCATG | | *HHF2* –5’ screening primer for deletion |
| J1760 | GCGGATGATGAGGGAAGAG | | *HHF2* – left flanking primer 1 for deletion |
| J1761 | ATTCACTGGCCGTCGTTTTACTCGGGTCTCCTCGTAGATG | | *HHF2* – left flanking primer 2 for deletion |
| J1762 | CATGGTCATAGCTGTTTCCTGTGGTAAGCCTTCTCCATCC | | *HHF2* – right flanking primer 1 for deletion |
| J1763 | ACTTATCCGCCTGCCTTC | | *HHF2* – right flanking primer 2 for deletion |
| J1764 | CGTCTATGCAGCACCAAGC | | *HHF2* – probe primer for Southern blot for deletion |
| J1796 | TCTGTCGAACAGCCAGGAG | | *CSE4* –5’ screening primer for *CTR4-NAT* |
| J1797 | TGTCCGTCTGTTCCAAGTG | | *CSE4*– left flanking primer 1 for *CTR4-NAT* |
| J1798 | GCCACTCGAATCCTGCATGCTTGGTTCGTGACTGTCGACGA | | *CSE4*– left flanking primer 2 for *CTR4-NAT* |
| J1799 | CGACAACGACTTCACCAATCCTCTTTTTCCTCCCATACTTT | | *CSE4*– right flanking primer 1 for *CTR4-NAT* |
| J1800 | AGAGAGCAAAGATGGCAGG | | *CSE4*– right flanking primer 2 for *CTR4-NAT* |
| J1736 | AGCCTTCTCTTCTGTCTGC | | *CSE4*– probe primer 1 for Southern blot for *CTR4-NAT* |
| J1740 | TTATCCGCCGAGCTAGTTG | | *CSE4*– probe primer 2 for Southern blot for *CTR4-NAT* |
| J1801 | CCTCTAAACCCAGCACACG | | *HTZ1* –5’ screening primer for deletion |
| J1802 | AGCGAAAGGGTGCGATCAG | | *HTZ1* – left flanking primer 1 for deletion |
| J1803 | ATTCACTGGCCGTCGTTTTACAGCCCAGCTTTGGATGAC | | *HTZ1* – left flanking primer 2 for deletion |
| J1804 | CATGGTCATAGCTGTTTCCTGTTTCCTGTGGGCCGTATTC | | *HTZ1* – right flanking primer 1 for deletion |
| J1805 | AGGCGCTGCAAGTAAGTC | | *HTZ1* – right flanking primer 2 for deletion |
| J1807 | ATCAAGCTCTTCGTCTCCAC | | *HTZ1* – probe primer for Southern blot for deletion |
| J1764 | CGTCTATGCAGCACCAAGC | | *HHF2*– probe primer 2 for Southern blot for *CTR4-NAT* |
| J1881 | AGTGGTCTGGTTCGGCTAG | | *HHT2* –5’ screening primer for *CTR4-NAT* |
| J1873 | AACACGTTTGGGCTGTGC | | *HHT2*– left flanking primer 1 for *CTR4-NAT* |
| J1882 | GCCACTCGAATCCTGCATGCATCTCGGAGAATAGGCATTACG | | *HHT2*– left flanking primer 2 for *CTR4-NAT* |
| J1883 | CGACAACGACTTCACCAATCAAACAAACACTCGCTCCTGG | | *HHT2*– right flanking primer 1 for *CTR4-NAT* |
| J1878 | AAGGCCAGGCTGCATCAAC | | *HHT2*– right flanking primer 2 for *CTR4-NAT* |
| J1873 | AACACGTTTGGGCTGTGC | | *HHT2*– probe primer 1 for Southern blot for *CTR4-NAT* |
| J1884 | CGAGCCTCCACTCTTCTTG | | *HHT2*– probe primer 2 for Southern blot for *CTR4-NAT* |
| J1808 | AGATGGAGACAGAGGGCAG | | *HHF2* –5’ screening primer for *CTR4-NAT* |
| J1759 | GAGAAGCGGGTGTACCATG | | *HHF2*– left flanking primer 1 for *CTR4-NAT* |
| J1809 | GCCACTCGAATCCTGCATGCACCAAACAAAGATAACGCCGAG | | *HHF2*– left flanking primer 2 for *CTR4-NAT* |
| J1810 | CGACAACGACTTCACCAATCCCGTGCCTGACCTCGTCATC | | *HHF2*– right flanking primer 1 for *CTR4-NAT* |
| J1764 | CGTCTATGCAGCACCAAGC | | *HHF2*– right flanking primer 2 for *CTR4-NAT* |
| J1759 | GAGAAGCGGGTGTACCATG | | *HHF2*– probe primer 1 for Southern blot for *CTR4-NAT* |
| J1764 | CGTCTATGCAGCACCAAGC | | *HHF2*– probe primer 2 for Southern blot for *CTR4-NAT* |
| J1835 | CCAAAGGACCCTCGACTTAC | | *HTZ1* –5’ screening primer for *H3-NEO* |
| J1836 | CACTCGAATCCTGCATGCTTATCGCTTTAAAATGGATC | | *HTZ1*– left flanking primer 2 for *H3-NEO* |
| J1837 | ACCACAACACATCTATCACACAAAACACAGACGCAACAA | | *HTZ1*– right flanking primer 1 for *H3-NEO* |
| J1838x | ACCTCAGCACATCTCTGTTC | | *HTZ1*– right flanking primer 2 for *H3-NEO* |
| J1942 | TCCTCAGGATCTTCATGGCTCCAGACTTGTTGGATGGAGGGGA | | *HTA1*– left flanking primer 2 for *TEF1-NEO* |
| J1943 | GAGGTTTCGAAAAAAATTGTGGTTCCAACAACCCTTCCACTA | | *HTA1*– right flanking primer 1 for *TEF1-NEO* |
| J1944 | TCCTCAGGATCTTCATGGCTCCGGACCCGTCGTCGTGTGGCCC | | *HTB1*– left flanking primer 2 for *TEF1-NEO* |
| J1945 | GAGGTTTCGAAAAAAATTGTGGCTGAGCTTCTCCCCACCGAGC | | *HTB1*– right flanking primer 1 for *TEF1-NEO* |
| J1127 | CTTCTTGGCACCGTCTTTG | | *HTA1* – screening primer for deletion |
| J1128 | ACAGACTTGGGAGCCATTG | | *HTA1* – left flanking primer 1 for deletion |
| J1129 | TCACTGGCCGTCGTTTTACGGGAAGAACACCACCTTGAG | | *HTA1* – left flanking primer 2 for deletion |
| J1130 | CATGGTCATAGCTGTTTCCTGACCGATGAACAGCGAAGG | | *HTA1* – right flanking primer 1 for deletion |
| J1131 | TGGCGGGCTTGTATGTTTG | | *HTA1* – right flanking primer 2 for deletion |
| J1132 | AGCTTGCGTTTCGGTGAC | | *HTA1* – probe primer for Southern blot for deletion |
| J2080 | TCGAGGCACGATACGAGAC | | *HTB1* – screening primer for deletion |
| J2081 | TTACACTCAAGGACGGCAG | | *HTB1* – left flanking primer 1 for deletion |
| J2082 | TCACTGGCCGTCGTTTTACGACCTGCTTGAGGACCTTG | | *HTB1* – left flanking primer 2 for deletion |
| J2083 | CATGGTCATAGCTGTTTCCTGGACATCTTCGAGCGAATTGC | | *HTB1* – right flanking primer 1 for deletion |
| J2084 | ACATCTACCGAGGATTGGAC | | *HTB1* – right flanking primer 2 for deletion |
| J1735 | TATTGTTGACGGCCCGACC | | *CSE4* – screening primer for deletion |
| J1736 | AGCCTTCTCTTCTGTCTGC | | *CSE4* – left flanking primer 1 for deletion |
| J1737 | ATTCACTGGCCGTCGTTTTACGTACGACTCGGGAGAATGG | | *CSE4* – left flanking primer 2 for deletion |
| J1738 | CATGGTCATAGCTGTTTCCTGAGCCATGAACGTAGGCTC | | *CSE4* – right flanking primer 1 for deletion |
| J1739 | AGCTCCTTCATCGCATACC | | *CSE4* – right flanking primer 2 for deletion |
| J1740 | TTATCCGCCGAGCTAGTTG | | *CSE4* – probe primer for Southern blot for deletion |
| J968 | TGCCCGAGACAACAAGAAG | | *HTA1* (CNAG_06747) qRT primer 1 |
| J969 | AGATGACAACAGACCCGAGG | | *HTA1* (CNAG_06747) qRT primer 2 |
| J966 | TCTTCGAGCGAATTGCCAC | | *HTB1* (CNAG_06746) qRT primer 1 |
| J967 | GGAAGGATGAGTCGGACAG | | *HTB1* (CNAG_06746) qRT primer 2 |
| J1902 | ACTTCCATCTCCACCCGAC | | *HHT1* (CNAG_04828) qRT primer 1 |
| J1903 | CACCTTTGTTCGGGCTATGC | | *HHT1* (CNAG_04828) qRT primer 2 |
| J1904 | ATCACAATGGCCCGAACG | | *HHT2* (CNAG_06745) qRT primer 1 |
| J1905 | GGTGGTGGTTTGCTTCCTG | | *HHT2* (CNAG_06745) qRT primer 2 |
| J964 | TGCCAAACTCCCATTCTCC | | *CSE4* (CNAG_00063) qRT primer 1 |
| J965 | AATGCCATGATCGCCGAAC | | *CSE4* (CNAG_00063) qRT primer 2 |
| J1806 | TATCTGACTGCCGAGGTCC | | *HTZ1* (CNAG_05221) qRT primer 1 |
| J1807 | ATCAAGCTCTTCGTCTCCAC | | *HTZ1* (CNAG_05221) qRT primer 1 |
| J1889 | CACGCTAAGAGGAAGACTG | | *HHF1* (CNAG_01628) qRT primer 1 (3’-UTR) |
| J1890 | CGAAAACGAAGAACATCCAC | | *HHF1* (CNAG_01628) qRT primer 2 (3’-UTR) |
| J1891 | ACGCCAAGAGGAAGACTGTC | | *HHF2* (CNAG_07807) qRT primer 1 (3’-UTR) |
| J1892 | CCCCAAAAAACTGTGCTCC | | *HHF2* (CNAG_07807) qRT primer 2 (3’-UTR) |
| J2627 | CACATTATGGCCGAGCTCCTTCCT | | *HTA1* internal PCR primer 1 |
| J2626 | TTACACCTCCTGAGAAGCCTTGGC | | *HTA1* internal PCR primer 2 |
| J2625 | GACAAAAGAGTTGAGGATAGCCAT | | *HTB1* internal PCR primer 1 |
| J2624 | TAACAAGGCCATGGCTATCCTCAA | | *HTB1* internal PCR primer 2 |
| J2628 | TTGGATCACGTAGGTAAGAGAAGT | | *CSE4* internal PCR l primer 1 |
| J2629 | ACTTCTCTTACCTACGTGATCCAA | | *CSE4* internal PCR l primer 2 |
| J61 | TTCCCGCCTCACTTCAATC | | *ERG1*(CNAG_06829) qRT primer 1 |
| J62 | AGGAAGACCCTGGATGGAG | | *ERG1*(CNAG_06829) qRT primer 2 |
| J59 | GGCCCTGCTCGTGAAATTG | | *ERG11*(CNAG_06829) qRT primer *1* |
| J60 | GCCTTCTTTGACTTGGCAG | | *ERG11*(CNAG_06829) qRT primer 2 |
| J1340 | ACGGTGTTGAAGACGACAG | | *HMG1*(CNAG_06534) qRT primer 1 |
| J1341 | CCGCGTGAGCGTTAAATC | | *HMG1*(CNAG_06534) qRT primer 2 |
| J2101 | CTTTGGTTGATGCCGATCC | | *SPO11* (CNAG_05472)_qRT_primer 1 |
| J2102 | CCACTGCACTCTATCCCTC | | *SPO11* (CNAG_05472)_qRT_primer 2 |
| J2103 | ACGCCTTCACTGCCATCTTC | | *MAT*alpha (CNAG_07407)_qRT_primer 1 |
| J2104 | CACAAAGGGTCATGCCAC | | *MAT*alpha (CNAG_07407)_qRT_primer 2 |
| J1791 | cgcGAGCTCGCCACGAGATGGTGTTGAG | | *HHT1* – primer 1 for complementation |
| J1792 | cgcACTAGTGGATTTCGCCGCCTATCTTG | | *HHT1* – primer 2 for complementation |
| J1793 | GCGGATGAGCCTGTGTTTG | | *HHT1* – primer 1 for sequencing |
| J1794 | GCGTACAGTCGCTACCAAG | | *HHT1* – primer 2 for sequencing |
| J1795 | CCCAACTCTGACCTAAGGCTCC | | *HHT1* – primer 3 for sequencing |
| J1816 | GCGGCCGCCTTCCCTTGGCAACACAC | | *HTZ1* – primer 1 for complementation |
| J1817 | GCGGCCGCCTCGAGCCCGAATGATGACCAATGC | | *HTZ1* – primer 2 for complementation |
| J1918 | CGCgcggccgcGCGATACCACCCAAATCAAC | | *HHT2*- primer 1 for complementation |
| J1919 | CGCgggcccTTGGGTGCGGAGAAGGATG | | *HHT2*- primer 2 for complementation |
| J1920 | CAGATATCCATCACACTGGCGCGATACCACCCAAATCAAC | | *HHT2*- primer 3 for complementation |
| J1921 | ACTATAGGGCGAATTGGGCCTTGGGTGCGGAGAAGGAT | | *HHT2*- primer 4 for complementation |
| J2632 | CGGAGGGAGAGAATGGAGATAATG | | CNAG_07843_qRT_primer_1 |
| J2633 | GGAAAGAGGAAAGGGTAAACTGAC | | CNAG_07843_qRT_primer_2 |
| J2634 | TCTTCCTCCTCATTGTCTTCAAGC | | CNAG_07854_qRT_primer_1 |
| J2635 | CAAGATAGTCTGGATGTGGGTGAG | | CNAG_07854_qRT_primer_2 |
| J2636 | AACACCCTCTCTGACCAATGAATC | | CNAG_04385_qRT_primer_1 |
| J2637 | GCTCTTCTCACTTTGGATTGTGG | | CNAG_04385_qRT_primer_2 |
| J2638 | CAATCTACTTTACCGACTCCCTTC | | CNAG_03980_qRT_primer_1 |
| J2639 | TTGGTGTCGGTAATGATGAGTAGG | | CNAG_03980_qRT_primer_2 |
| J2640 | GTTGGTGAAGGAGACAATCATTGG | | CNAG_03532_qRT_primer_1 |
| J2641 | TTAAACGGAATACCCTCATCCTGG | | CNAG_03532_qRT_primer_2 |
| J2642 | TAAGCTTTGGAGCACCTTTGCCTA | | CNAG_07592_qRT_primer_1 |
| J2643 | GCAAGGGAAAGATGATGTTCTCTG | | CNAG_07592_qRT_primer_2 |
| J2644 | GATCTATGGAAGTTGGACCCTACC | | CNAG_04246_qRT_primer_1 |
| J2645 | GAATGAGACTGCTCCGTAAAGACA | | CNAG_04246_qRT_primer_2 |
| J2646 | CATGTTCCTCGTCAACTCAAAGTG | | CNAG_03754_qRT_primer_1 |
| J2647 | CAGGAGCTTTACTGCCTTGATATG | | CNAG_03754_qRT_primer_2 |
| J2648 | GTTACTGCTTGCCTCAATGTTGTC | | CNAG_05729_qRT_primer_1 |
| J2649 | GACTGAATAAAGAAACGCCAGCAC | | CNAG_05729_qRT_primer_2 |
| J2650 | TCTATTCACAACACAGACCCTAGC | | CNAG_04585_qRT_primer_1 |
| J2651 | CCAAACCCTCAGGGGCAGCCTTCT | | CNAG_04585_qRT_primer_2 |
| J2652 | GATATTCCCGGTTTAGGCTACAAG | | CNAG_01464_qRT_primer_1 |
| J2653 | AGGGTTAGAAAGTGTAGGTACAGG | | CNAG_01464_qRT_primer_2 |
| J2654 | CTAAGGGAATGGTGTATGAGGAGG | | CNAG_05974_qRT_primer_1 |
| J2655 | GAGTCTTGATAAGGTCGAGGAAGG | | CNAG_05974_qRT_primer_2 |
| J2656 | GTTACCCGCAATCGTTTCATCATC | | CNAG_02361_qRT_primer_1 |
| J2657 | GATGATGTAGGCAAACCAATGAGC | | CNAG_02361_qRT_primer_2 |
| J2658 | TAAACTCTTACGAGCCCTTCATCC | | CNAG_06923_qRT_primer_1 |
| J2659 | GAATGTTGAGAGAGGAGACTTTGG | | CNAG_06923_qRT_primer_2 |
| J2660 | CGATAAAGTCTCGAACAAGCTCAC | | CNAG_03890_qRT_primer_1 |
| J2661 | TCGTCTCTAGGCATCTTCATGTTG | | CNAG_03890_qRT_primer_2 |
| J2662 | ATCCCTCTCGTCTTCATCATCATC | | CNAG_04334_qRT_primer_1 |
| J2663 | GGTCGGAGTTTGATATGGGATATG | | CNAG_04334_qRT_primer_2 |
| J2664 | GACCGAGATGTAGAAGTGTATGAG | | CNAG_07450_qRT_primer_1 |
| J2665 | TGTACTGTGGAAGGACGATTGTTG | | CNAG_07450_qRT_primer_2 |
| J2666 | GGTGTTTCCAAAAAAAGGCAGTCT | | CNAG_06471_qRT_primer_1 |
| J2667 | CGTCATCTGTCCGTTGGAAAGCTT | | CNAG_06471_qRT_primer_2 |
| J2668 | AATAAGTGAACCATCTGGGTATCG | | CNAG_01921_qRT_primer_1 |
| J2669 | CTGAACGAAAAGGCTTGGTCCAAT | | CNAG_01921_qRT_primer_2 |
| J2670 | ACGCTTACACTCTTCTCCTTTCTC | | CNAG_02048_qRT_primer_1 |
| J2671 | GTGTTGTATGGCGTGAATAAGGTG | | CNAG_02048_qRT_primer_2 |
| J2672 | TAACCCTAACCCTAACGCAACTAC | | CNAG_05064_qRT_primer_1 |
| J2673 | GTATCAGGAGGTAAATGGATGTCG | | CNAG_05064_qRT_primer_2 |
| J2217 | GATGAGGCCAAAGAGAATGCTTCC | | CNAG_03759_qRT_primer1 |
| J2218 | GATCATTGTTGCTCTCTTCCTTGG | | CNAG_03759_qRT_primer2 |
| J2674 | CTGGATTTGTTCATGTCTCCTTCG | | CNAG_05099_qRT_primer_1 |
| J2675 | ATCGCTGGTCATAAGATACCTTGG | | CNAG_05099_qRT_primer_2 |
| J2676 | GCTCTTCTTCTGGTGACTCTTCTG | | CNAG_03782_qRT_primer_1 |
| J2677 | GAGTCACTAGAGGAAGAGGAGGAG | | CNAG_03782_qRT_ primer_2 |
| J2678 | TATGGGTATGATGAATCCTATGAT | | CNAG_02659_qRT_ primer_1 |
| J2679 | TATGCTCCAGAACCAAGGCCTCCA | | CNAG_02659_qRT_ primer_2 |
| J2680 | GACAGAGGGCTCAACAAGAAACAC | | CNAG_07885_qRT_ primer_1 |
| J2681 | GTGCTCTTCTAGGGACTGTTCTTC | | CNAG_07885_qRT_ primer_2 |
| J2682 | ATTCATCGTCTTCATCTGCCAAGC | | CNAG_02735_qRT_ primer_1 |
| J2683 | CGCCCTCCGCAATATCTTTAACAC | | CNAG_02735_qRT_ primer_2 |
| J2684 | CACCCTATCCTGTCCCATGTCTTC | | CNAG_05258_qRT_ primer_1 |
| J2685 | CCATTGTGATTCTTCCTGTAGGAC | | CNAG_05258_qRT_ primer_2 |
| J2686 | GTATCAGCCAGTCTATCGTCGAAC | | CNAG_07725_qRT_ primer_1 |
| J2687 | CTCCTTGATCTCCTCTTCCAACTG | | CNAG_07725_qRT_ primer_2 |
