## Supplementary Figures for "Systematic histone mutagenesis reveals nucleosome-dependent maintenance of three-dimensional chromosome architecture and virulence in *Cryptococcus neoformans*"

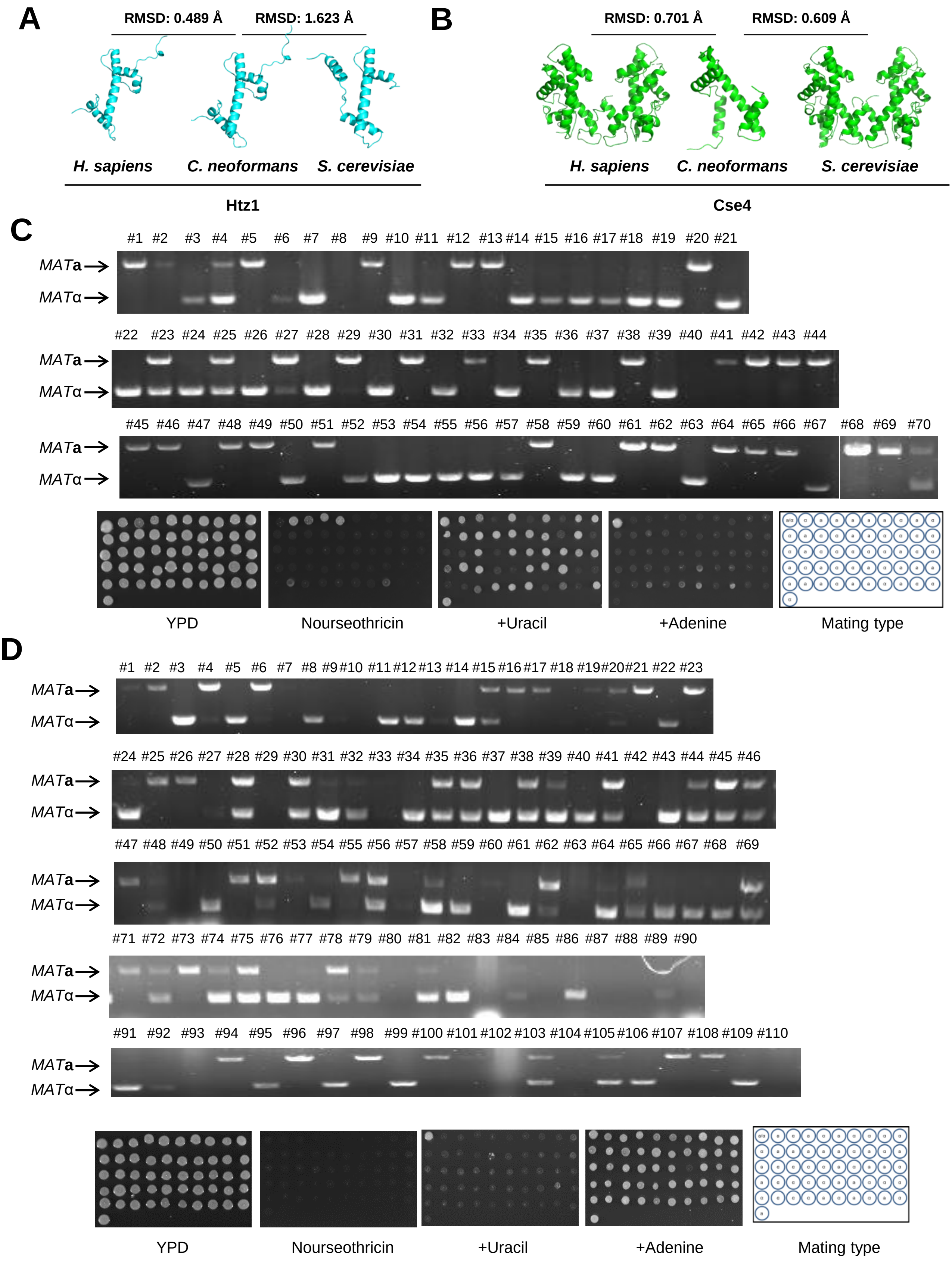
A
RMSD: 0.489 Å
RMSD: 1.623 Å
H. sapiens
C. neoformans
S. cerevisiae
Htz1
B
RMSD: 0.701 Å
RMSD: 0.609 Å
H. sapiens
C. neoformans
S. cerevisiae
Cse4
C
#1 #2 #3 #4 #5 #6 #7 #8 #9 #10 #11 #12 #13 #14 #15 #16 #17 #18 #19 #20 #21
#22 #23 #24 #25 #26 #27 #28 #29 #30 #31 #32 #33 #34 #35 #36 #37 #38 #39 #40 #41 #42 #43 #44
#45 #46 #47 #48 #49 #50 #51 #52 #53 #54 #55 #56 #57 #58 #59 #60 #61 #62 #63 #64 #65 #66 #67 #68 #69 #70
MATa
MATα
YPD
Nourseothricin
+Uracil
+Adenine
Mating type
D
#1 #2 #3 #4 #5 #6 #7 #8 #9 #10 #11 #12 #13 #14 #15 #16 #17 #18 #19 #20 #21 #22 #23
#24 #25 #26 #27 #28 #29 #30 #31 #32 #33 #34 #35 #36 #37 #38 #39 #40 #41 #42 #43 #44 #45 #46
#47 #48 #49 #50 #51 #52 #53 #54 #55 #56 #57 #58 #59 #60 #61 #62 #63 #64 #65 #66 #67 #68 #69
#71 #72 #73 #74 #75 #76 #77 #78 #79 #80 #81 #82 #83 #84 #85 #86 #87 #88 #89 #90
#91 #92 #93 #94 #95 #96 #97 #98 #99 #100 #101 #102 #103 #104 #105 #106 #107 #108 #109 #110
MATa
MATα
YPD
Nourseothricin
+Uracil
+Adenine
Mating type

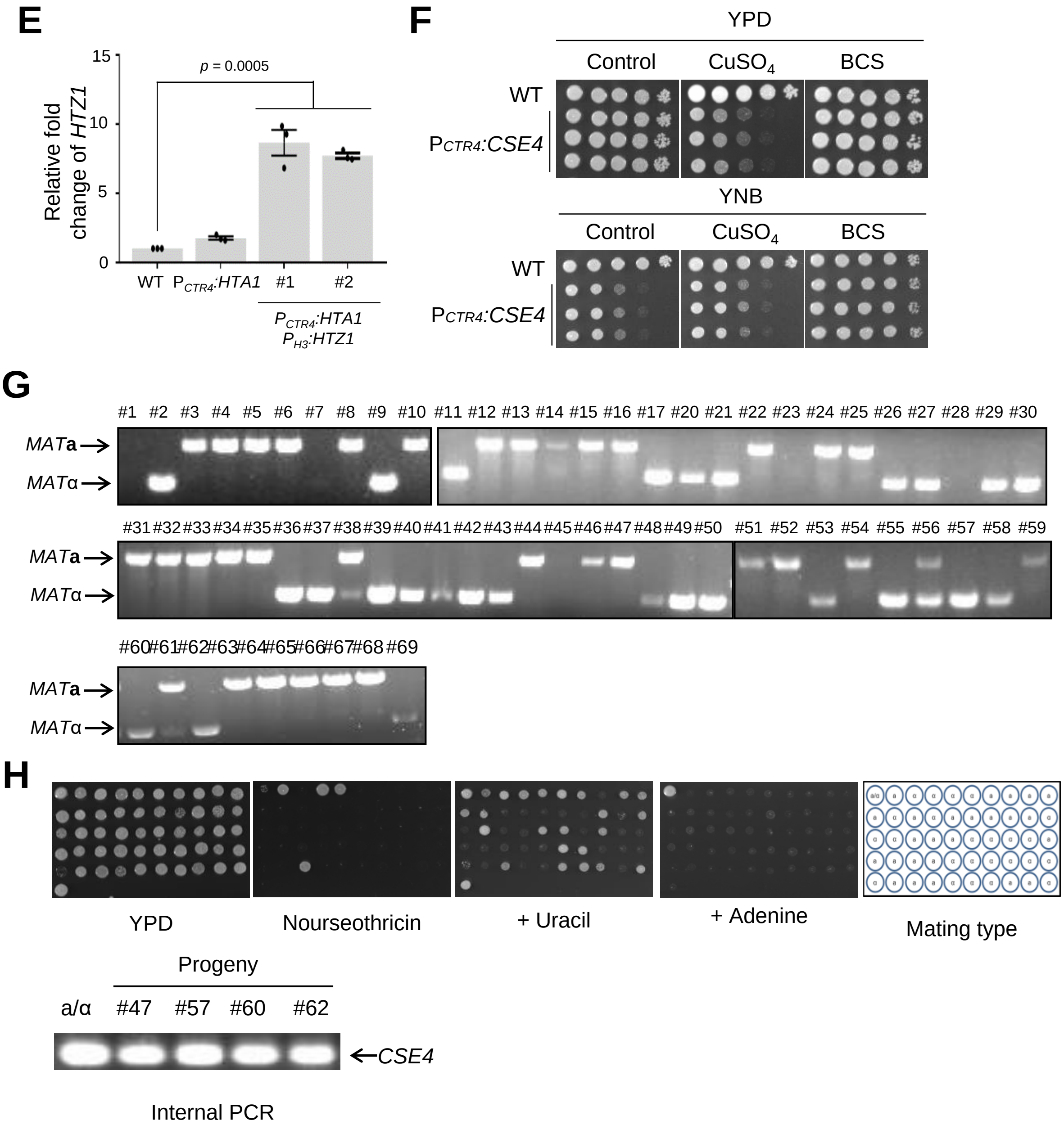


**Fig. S1** Structural prediction and meiotic progeny analysis for assessing the essentiality of Htz1 and Cse4. **A and B** Structure prediction of histone variants, Htz1 **(A)** and Cse4 **(B)**. **C and D** PCR analysis to characterize the marker that segregated during meiosis (*MATα* or *MAT***a**) in progeny obtained from the sporulation of the heterozygous *HTA1*/*hta1*Δ **(A)** and *HTB1*/*htb1*Δ **(B)**. 50 basidiospore progeny were isolated from each mutant and were grown on four types of medium (YPD, +NAT, +adenine and +uracil) for genetic analysis. **E** The constitutive overexpression of *HTZ1* in the $P_{CTR4}$:*HTA1* strain. Error bars indicate standard errors of the means. Statistical significance of difference was determined by one-way ANOVA with Bonferroni's multiple-comparison test (***: $P < 0.001$ and NS: not significant). **F** *CSE4* was required for cell growth. The overnight-cultured WT and $P_{CTR4}$:*CSE4* (KW1559 and KW1562) strains were spotted onto YNB (or YPD) medium containing the BCS (200 μM) or $CuSO_4$ (25 μM). **G** PCR analysis to characterize the marker that segregates during meiosis (*MATα* or *MAT***a**) in progeny obtained from the sporulation of the heterozygous *CSE4*/*cse4*Δ. **H** 50 basidiospore progeny were isolated from each mutant and were grown on four types of medium (YPD, +NAT, +adenine and +uracil) for genetic analysis. Basidiospore progeny isolated from the heterozygous *CSE4*/*cse4*Δ strain were spotted onto nourseothricin medium, followed by PCR analysis of *NAT* progeny (#47, #57, #60, and #62) using internal PCR.

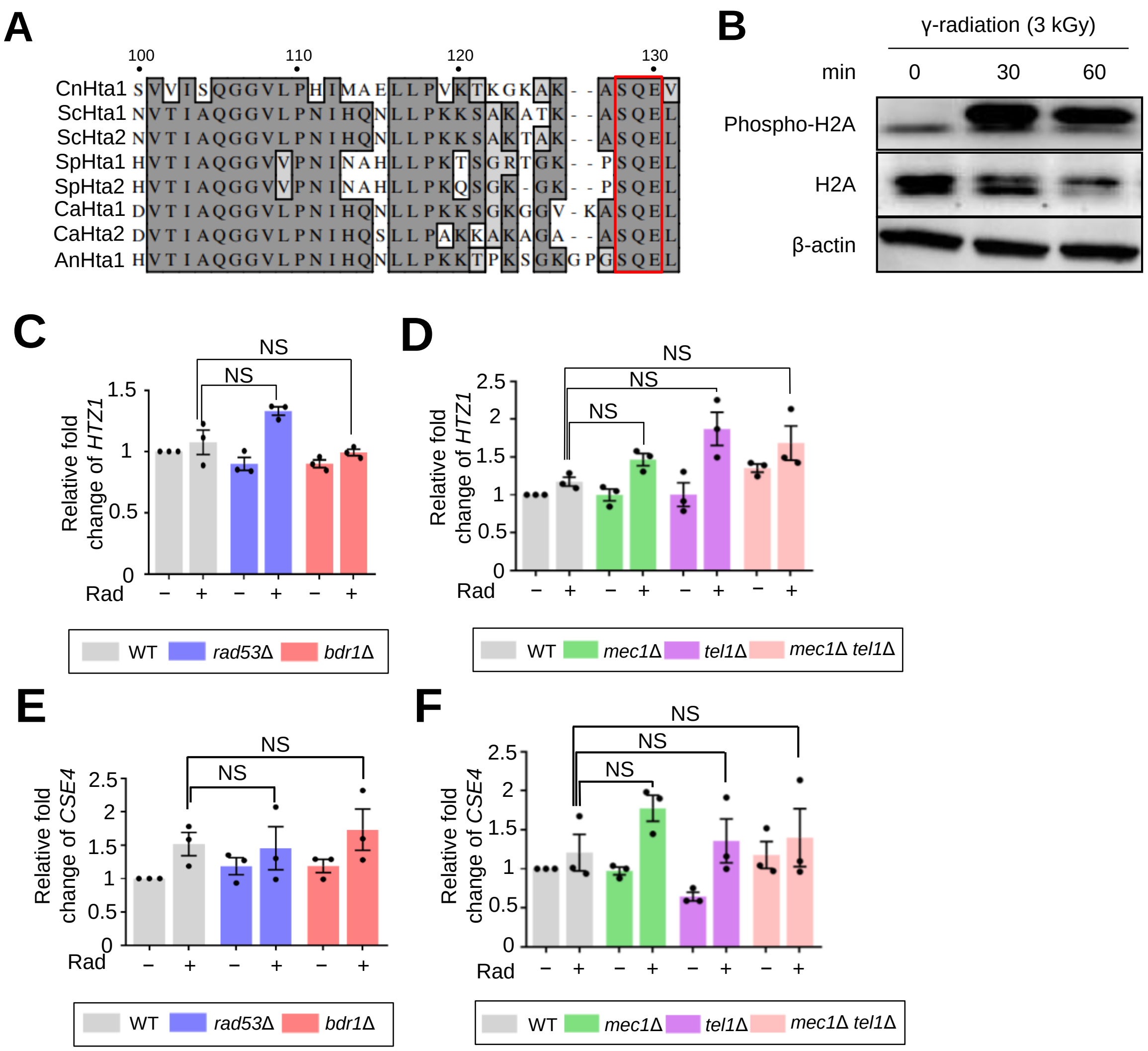


**Fig. S2** The expression levels of histone variant genes. **A** Sequence alignment of the phosphorylation region of Hta1 and Hta2 in *S. cerevisiae*, *Schizosaccharomyces pombe, Candida albicans* and Hta1 of *Aspergillus nidulans* and *C. neoformans.* **B** The phosphorylation of Hta1 in response to radiation. The WT strain was grown to the mid-logarithmic phase and exposed to the radiation (3 kGy) and further incubated for each indicated time. **C-F** The expression of *HTZ1* or *CSE4* post-radiation (0.5 kGy) exposure in strains belonging to the DNA repair pathway. Three independent biological experiments with duplicate technical replicates were performed.

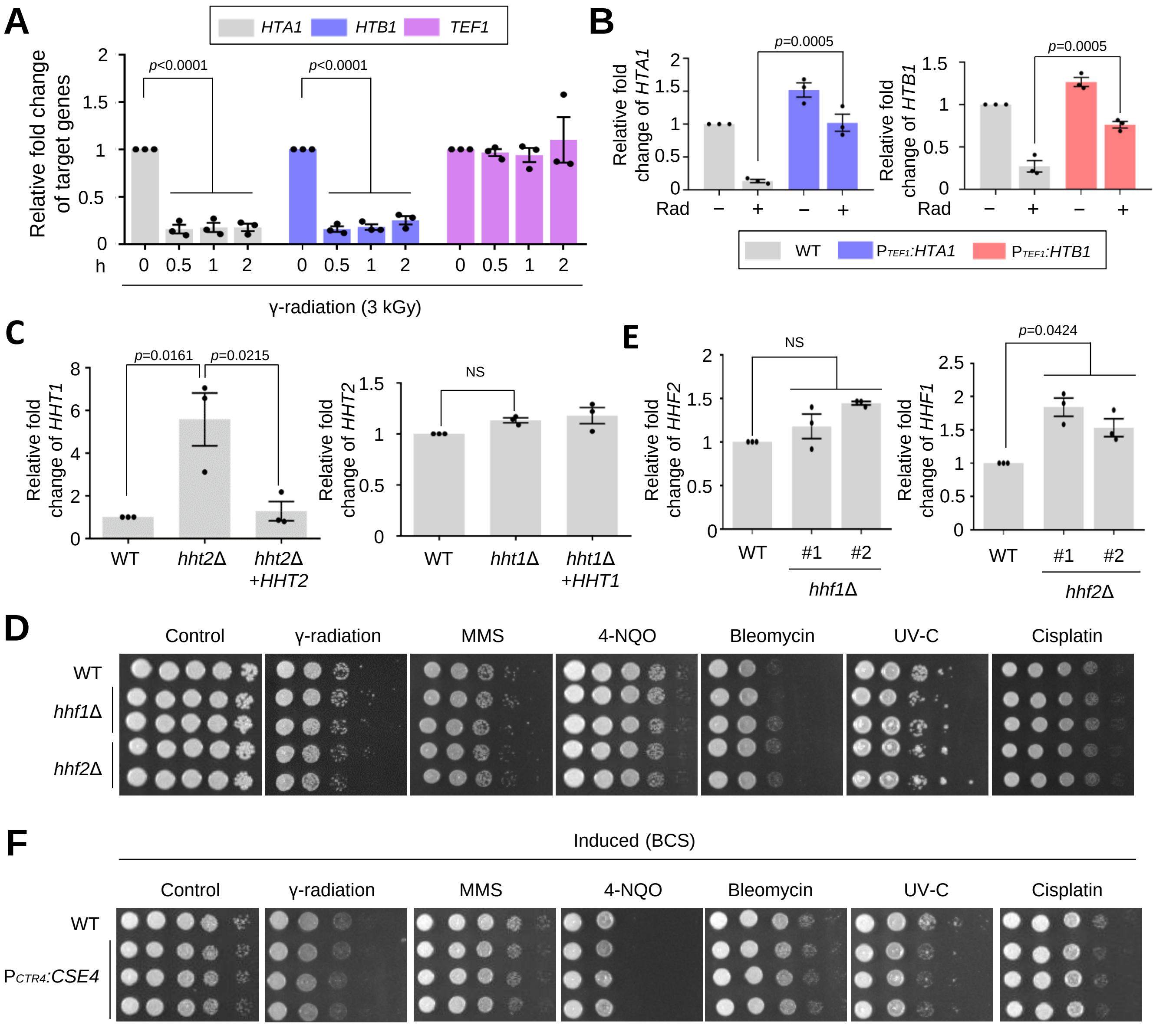


**Fig. S3** Phenotypic profiling of histone mutants in response to diverse DNA damaging agents. **A** Expression levels of *HTA1*, *HTB1*, and *TEF1* post-radiation exposure (3 kGy). **B** The constitutive expression of *HTA1* and *HTB1* in the $P_{TEF1}$:*HTA1* and $P_{TEF1}$:*HTB1* strains post radiation exposure (3 kGy). **C** The compensation of *HHT1* expression in the loss of *HHT2*. The qRT-PCR analysis was used to quantify expression levels of *HHT1* and *HHT2* in WT, *hht1*Δ, *hht2*Δ mutant and their complemented strains. **D and F** The strains were spotted onto YPD medium containing MMS, 4-NQO, cisplatin, and bleomycin. For UV-C and γ-radiation exposures, the serially diluted cells were spotted and then were exposed to UV-C or γ-radiation. The two images split by a horizontal white line in each spot assay were obtained from the same plate. **E** The compensation of *HHF1* and *HHF2* expression upon loss of *HHF2* and *HHF1*, respectively. The expression levels of *HHF1* or *HHF*2 were measured in the WT, *hhf2*Δ and *hhf1*Δ mutants using qRT-PCR with gene-specific primers for *HHF1* and *HHF2* genes. For qRT-PCR analysis, three independent biological experiments with duplicate technical replicates were performed. Statistical analysis of difference was determined by one-way ANOVA with Bonferroni's multiple-comparison test (NS: not significant).

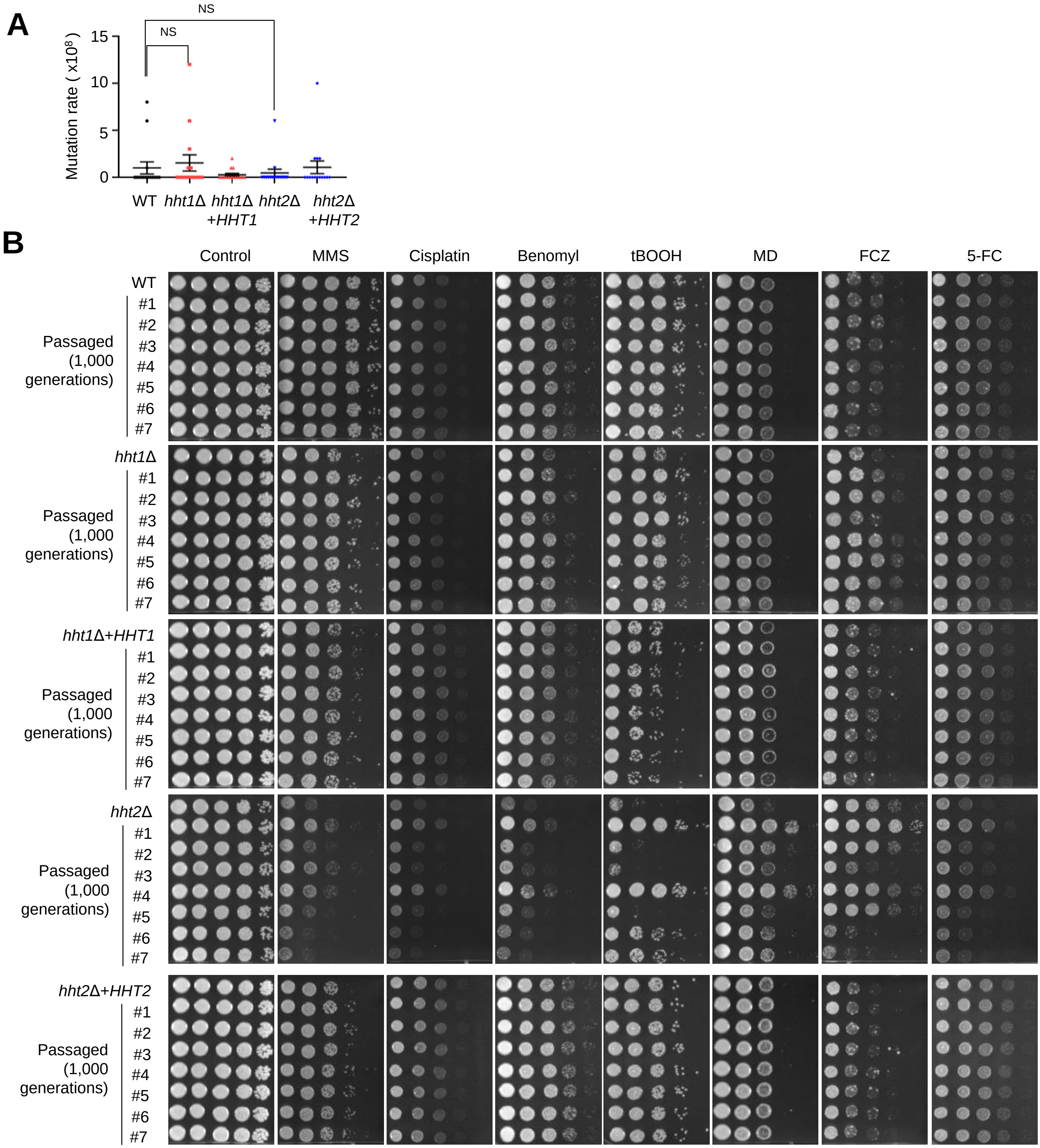


**Fig. S4** The deletion of Histone H3 led to increased phenotypic changes. **A** Quantification of spontaneous 5-FOA-resistant rates in H3 mutants. Error bars indicate standard errors of means (NS: not significant). **B** The phenotypic changes of the original WT strain, *hht1*Δ, *hht1*Δ+*HHT1*, *hht2*Δ, and *hht2*Δ+*HHT2* mutants compared to those of its corresponding strains after seven independent passages (1,000 generations) in response to diverse stress responses. Each strain was cultured in liquid YPD medium at 30°C. The cultured cells were serially diluted (1 to $10^4$) and spotted on the YPD plates containing 0.035 % MMS, 5 mM cisplatin, 4 μg/ml benomyl, 0.8 mM tBOOH, 0.025 mM MD, 16 μg/ml FCZ or 600 μg/ml 5-FC. Cells were further incubated at 30°C and photographed daily for 3 days.

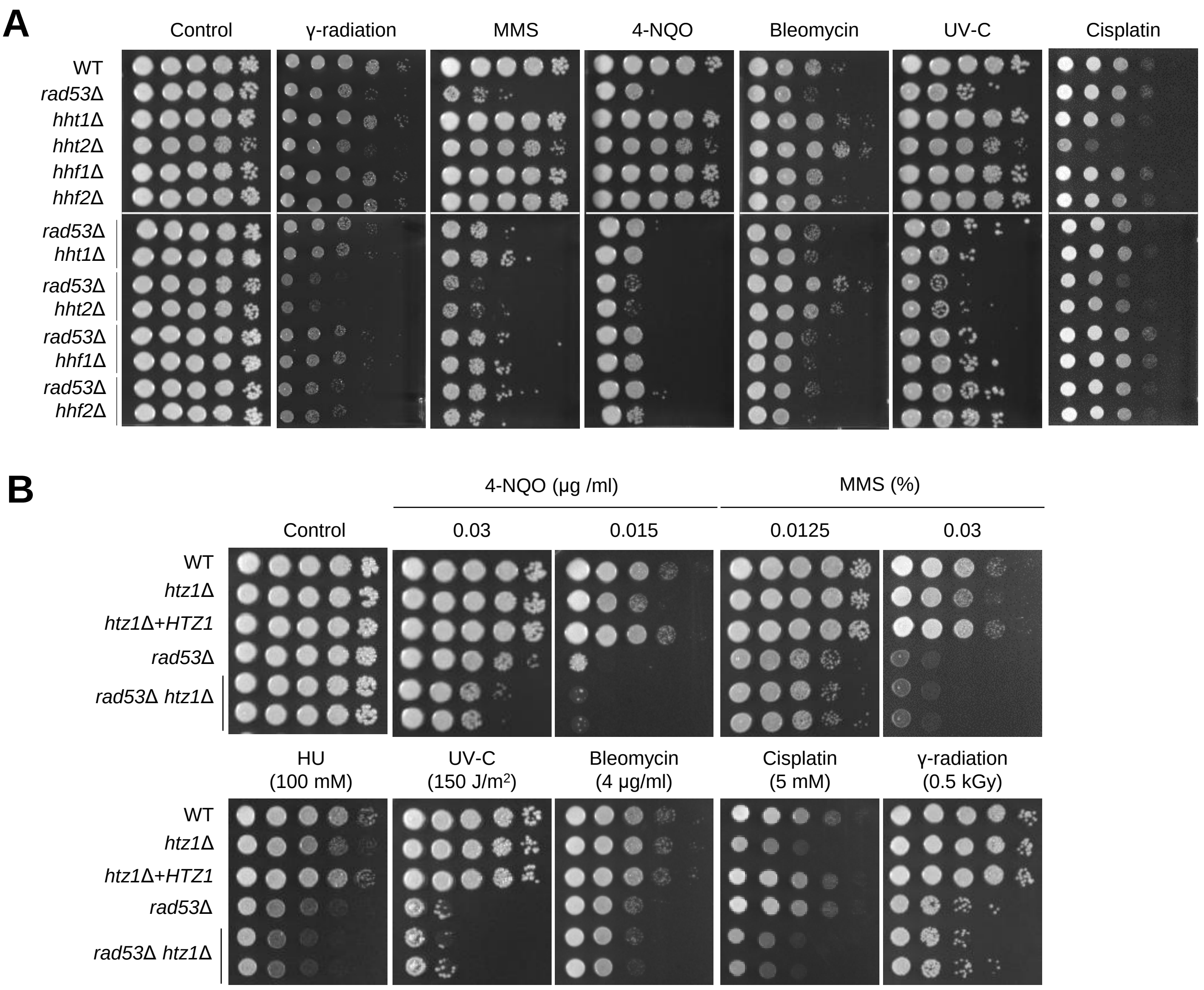


**Fig. S5** Hht2, unlike Hht1, Hhf1, or Hhf2, contributed to the DNA damage response through both Rad53-dependent and independent mechanisms. **A** The strains (WT, *rad53*Δ, *hht1*Δ, *hht2*Δ, *hhf1*Δ, *hhf2*Δ, *rad53*Δ *hht1*Δ, *rad53*Δ *hht2*Δ, *rad53*Δ *hhf1*Δ and *rad53*Δ *hhf2*Δ) were spotted onto YPD medium containing 0.04 % MMS, 0.04 μg/ml 4-NQO, 5 mM cisplatin, 4 μg/ml bleomycin, 10 μg/ml TBZ or 3.5 μg/ml benomyl. For UV-C and γ-radiation exposures, the serially diluted cells were spotted and then were exposed to 100 $J/m^2$ UV-C or 0.5 kGy γ-radiation. The two images split by a horizontal white line in each spot assay were obtained from the same plate. **B** The grown strains were serially diluted (1 to $10^5$) cells were spotted onto YPD media containing the indicated concentrations of chemicals and were further incubated at 30$^{o}$C for 4 days.

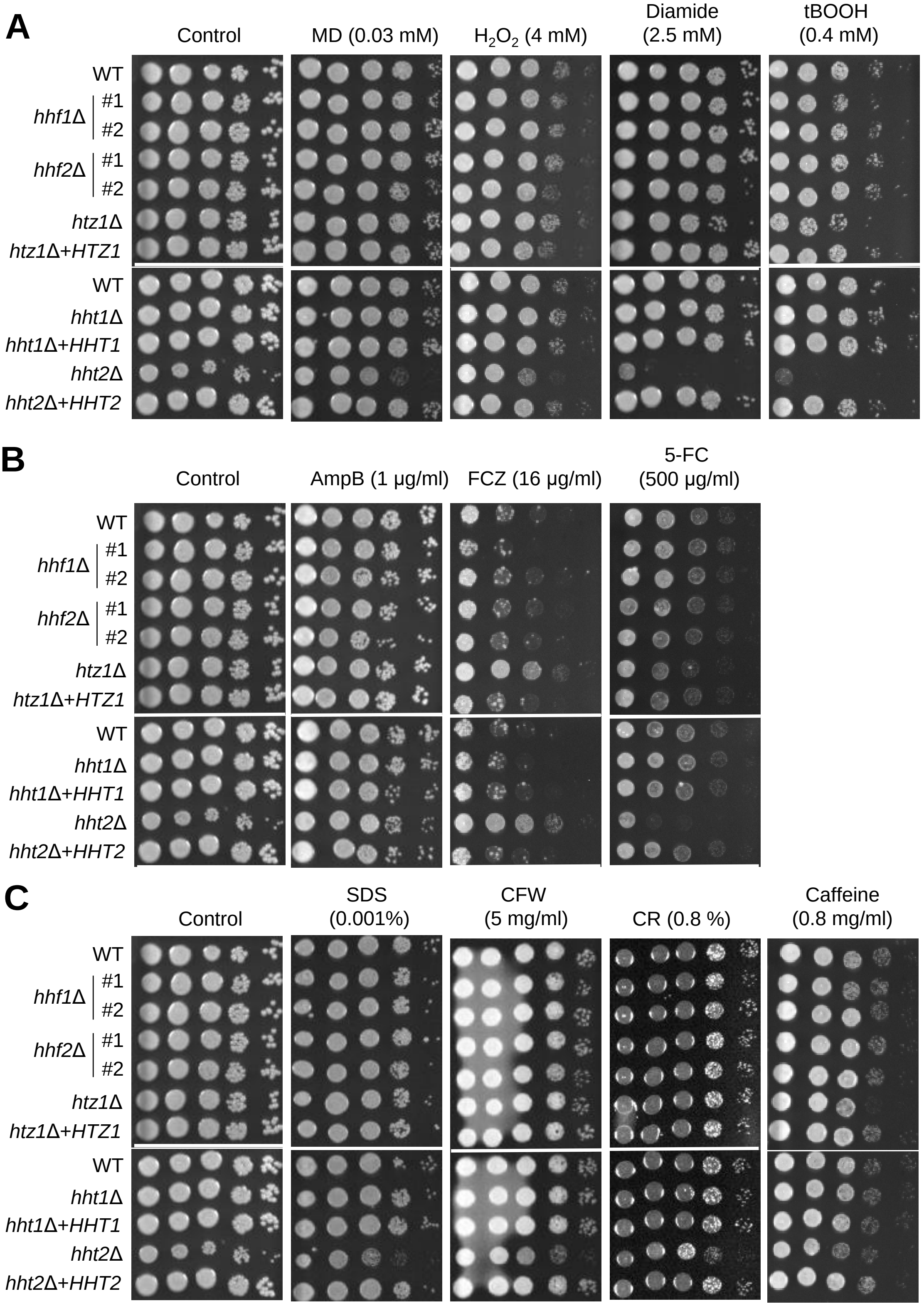
A
Control
MD (0.03 mM)
$H_2O_2$ (4 mM)
Diamide (2.5 mM)
tBOOH (0.4 mM)
WT
hhf1Δ #1
#2
hhf2Δ #1
#2
htz1Δ
htz1Δ+HTZ1
WT
hht1Δ
hht1Δ+HHT1
hht2Δ
hht2Δ+HHT2
B
Control
AmpB (1 μg/ml)
FCZ (16 μg/ml)
5-FC (500 μg/ml)
WT
hhf1Δ #1
#2
hhf2Δ #1
#2
htz1Δ
htz1Δ+HTZ1
WT
hht1Δ
hht1Δ+HHT1
hht2Δ
hht2Δ+HHT2
C
Control
SDS (0.001%)
CFW (5 mg/ml)
CR (0.8 %)
Caffeine (0.8 mg/ml)
WT
hhf1Δ #1
#2
hhf2Δ #1
#2
htz1Δ
htz1Δ+HTZ1
WT
hht1Δ
hht1Δ+HHT1
hht2Δ
hht2Δ+HHT2

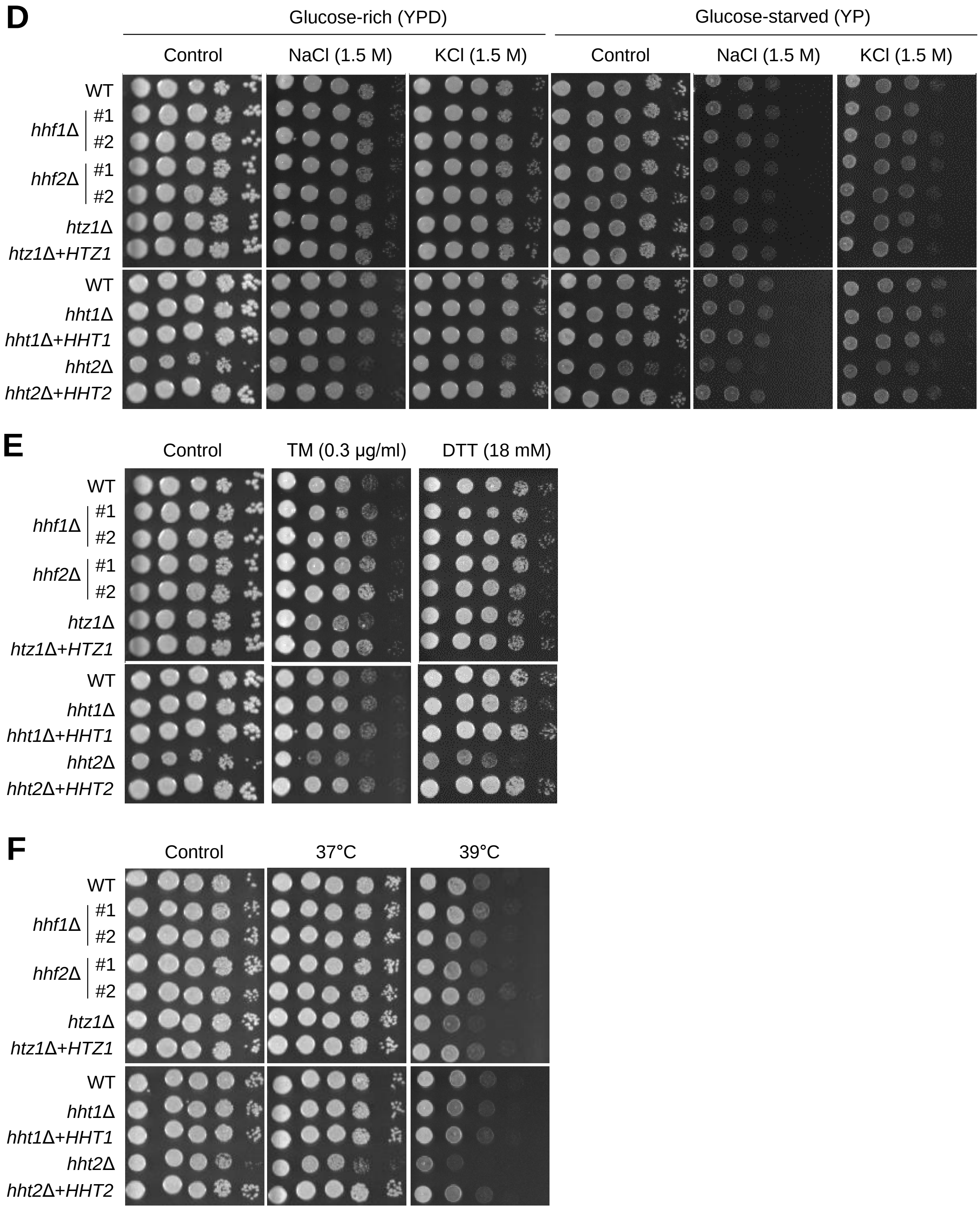


**Fig. S6** Phenotypic analysis of core histone and histone variant mutants for diverse stress responses. **A-F** The cultured cells were serially diluted (1 to $10^4$) and spotted on the YPD plates containing stress-inducing chemicals. Cells were further incubated at 30°C and photographed daily for 3 days.

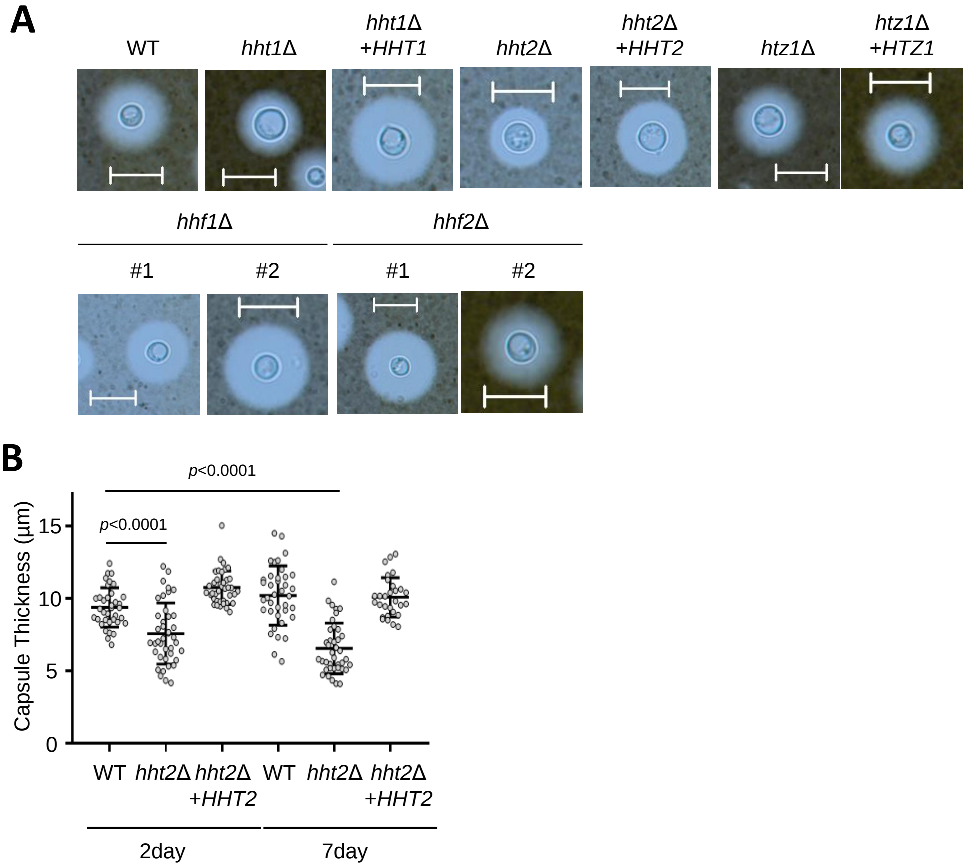


**Fig. S7** Virulence factor formation in histone mutants. **A** Representative images of strains cultured on Littman medium for capsule production at 30$^{o}$C for 3 days. The scale bar indicates 10 μm. **B** The strains were cultured on Littman medium for capsule production at 30$^{o}$C for 3 days or for the indicated days. Statistical analysis was conducted using one-way ANOVA.

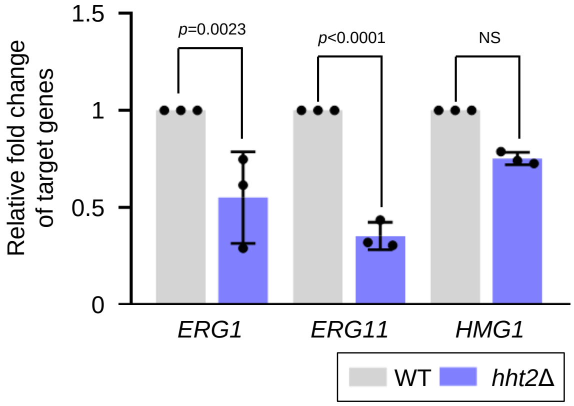


**Fig. S8** The expression levels of ergosterol synthesis genes in WT and *hht2*Δ strains. Three independent biological experiments with duplicate technical replicates were performed. Error bars indicate standard errors of the means. Statistical significance of difference was determined by one-way ANOVA with Bonferroni's multiple-comparison test (NS: not significant).

**A**

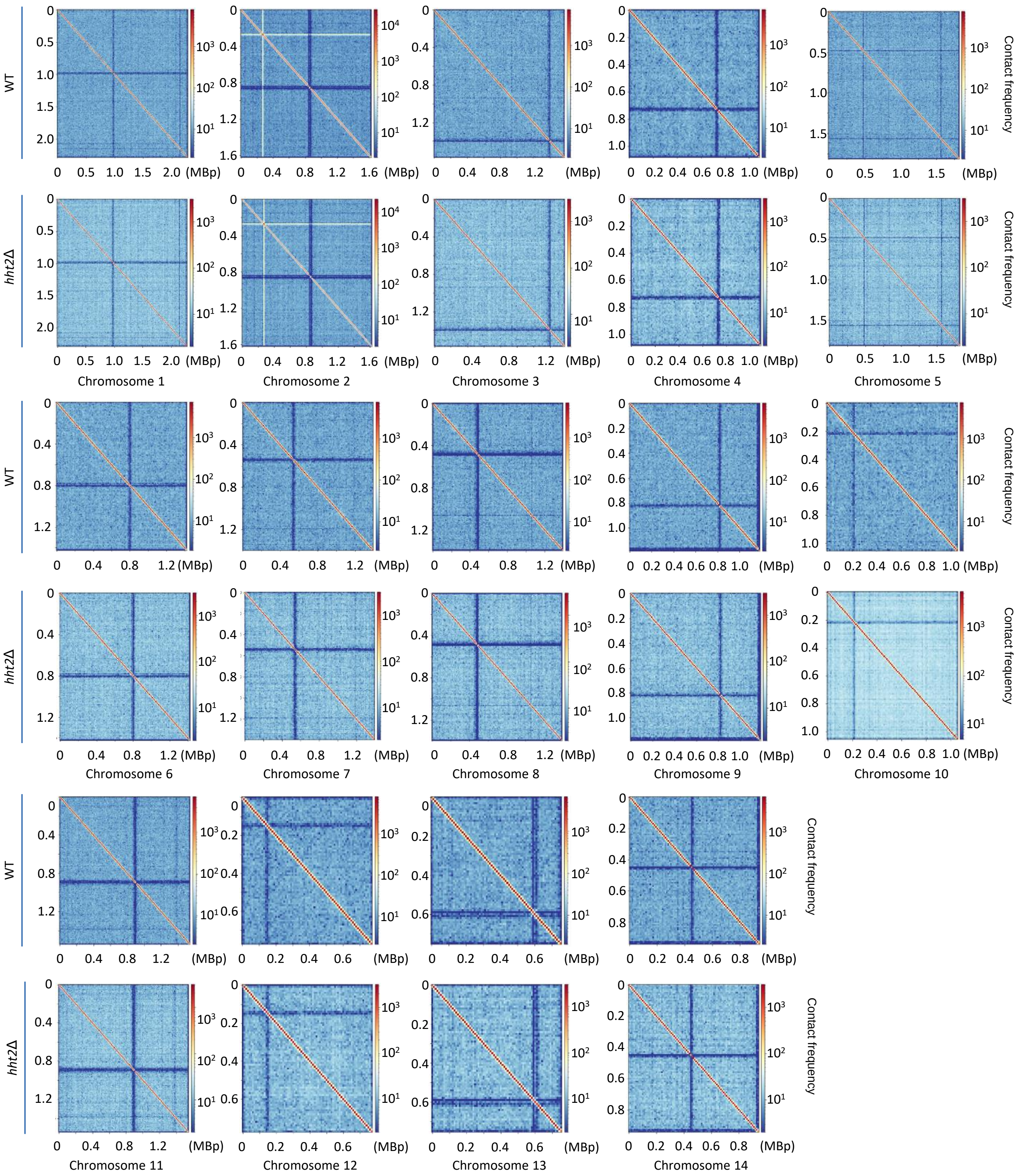

WT
hht2Δ
Chromosome 1
Chromosome 2
Chromosome 3
Chromosome 4
Chromosome 5
Chromosome 6
Chromosome 7
Chromosome 8
Chromosome 9
Chromosome 10
Chromosome 11
Chromosome 12
Chromosome 13
Chromosome 14
Contact frequency
(MBp)

**B**

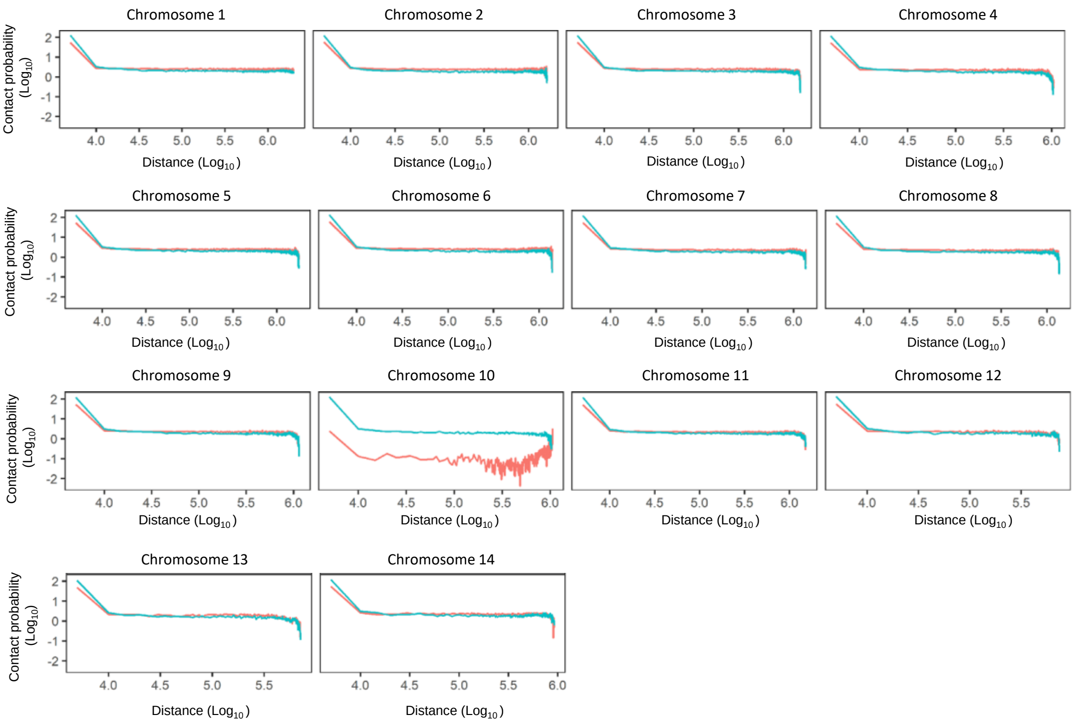


**C**

| Compartment Dynamics | Down-regulated genes | Non-Down-regulated Genes |
|---|---|---|
| A -> B | 525 | 444 |
| A -> A | 577 | 749 |
| | Up-regulated genes | Non-Up-regulated genes |
| B -> A | 627 | 703 |
| B -> B | 0 | 714 |

**D**

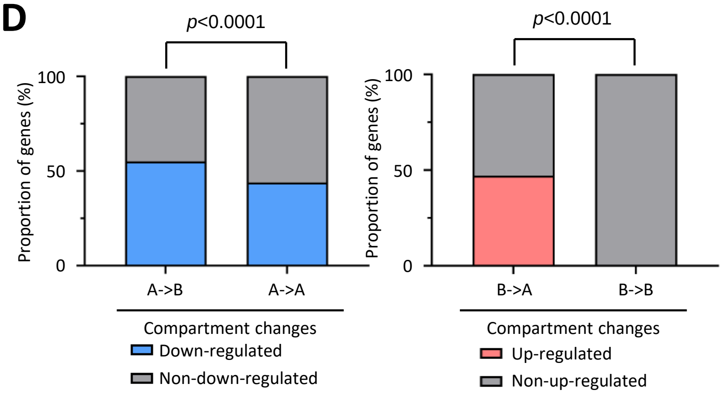


**E**

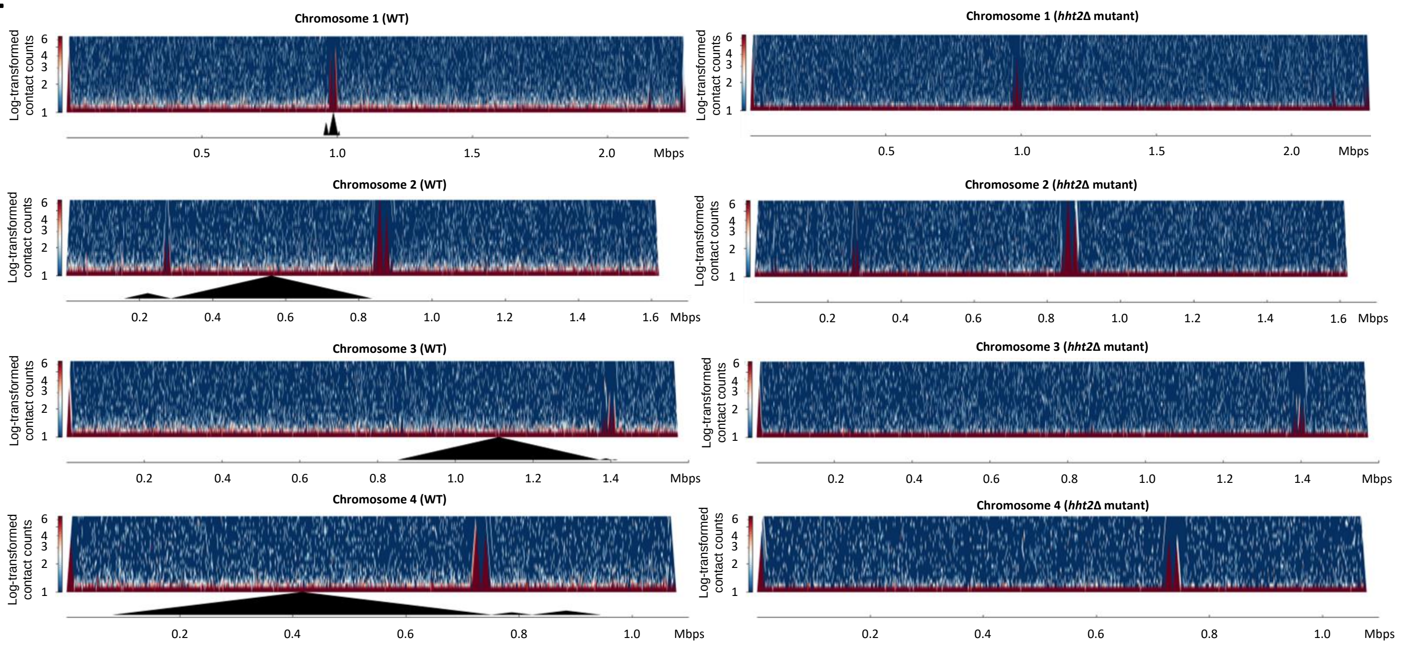

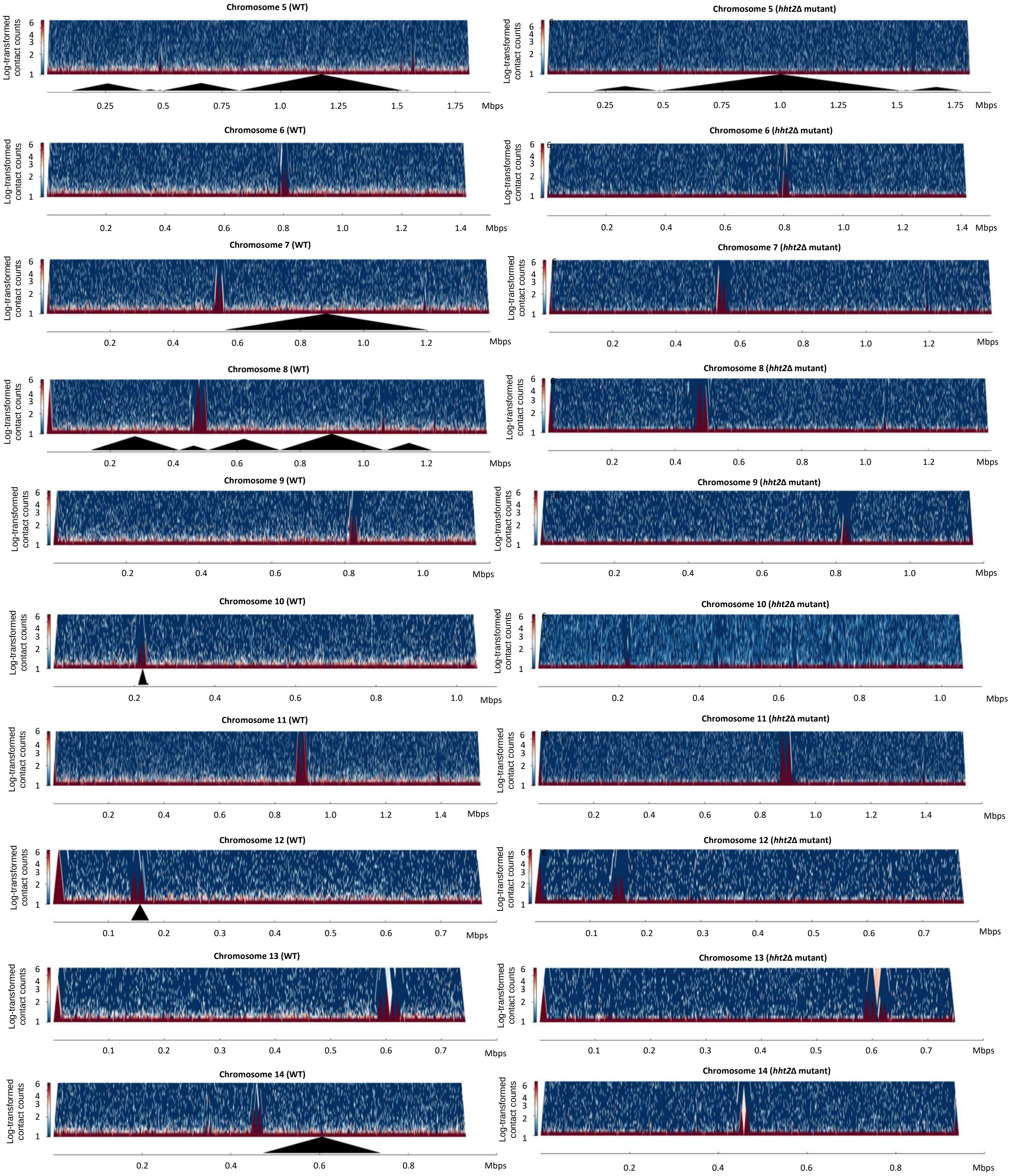
Chromosome 5 (WT)
Chromosome 5 (*hht2*Δ mutant)
Chromosome 6 (WT)
Chromosome 6 (*hht2*Δ mutant)
Chromosome 7 (WT)
Chromosome 7 (*hht2*Δ mutant)
Chromosome 8 (WT)
Chromosome 8 (*hht2*Δ mutant)
Chromosome 9 (WT)
Chromosome 9 (*hht2*Δ mutant)
Chromosome 10 (WT)
Chromosome 10 (*hht2*Δ mutant)
Chromosome 11 (WT)
Chromosome 11 (*hht2*Δ mutant)
Chromosome 12 (WT)
Chromosome 12 (*hht2*Δ mutant)
Chromosome 13 (WT)
Chromosome 13 (*hht2*Δ mutant)
Chromosome 14 (WT)
Chromosome 14 (*hht2*Δ mutant)
Log-transformed contact counts
Mbps

F

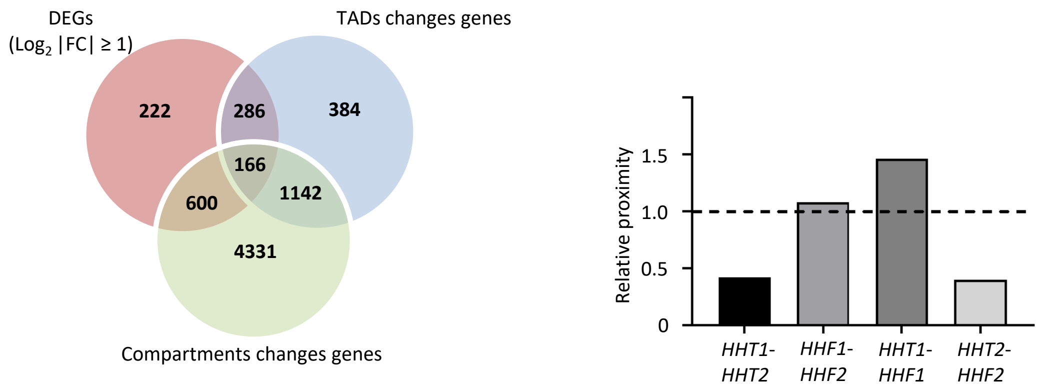


**Fig. S9** Hht2 affected the maintenance of the 3D chromatin structure. **A** Heatmaps (10-kb resolution) showing the chromosomal interactions from chromosome 1 to chromosome 14 for WT and *hht2*Δ mutant. The panels show ICE-normalized contact matrices, with the top panels representing WT and the bottom panels representing the *hht2*Δ strain. **B** Average contact probability (CP) as a function of genomic distance in WT and *hht2*Δ mutant strains. **C** 2 x 2 contingency tables showing the number of genes used for Fisher's exact test to evaluate expression changes in transitioning (A to B and B to A) versus stable (A to A and B to B) compartments. **D** Directional correlation between A/B compartment switching and gene expression changes. Stacked bar graphs showed the proportion of genes showing statistically significant changes in expression ($P$ < 0.05) across different compartment transitions. The left panel illustrated the percentage of down-regulated genes (blue) versus non-down-regulated genes (grey) in regions undergoing A to B transitions compared to A to A regions. The right panel displays the percentage of up-regulated genes (red) versus non-up-regulated genes (grey) within B to A transition regions compared to stable B to B regions. Statistical significance was determined using Fisher's exact test (****: $P$ < 0.0001). **E** Contact maps and corresponding TAD annotations of WT and *hht2*Δ mutants across chromosome 1 to 14, generated using the hicFindTADs tool. **F** Venn diagrams described the overlap among DEGs DEGs ($P$ < 0.05, $|Log_2FC| \geq 1$), genes located in TAD-altered regions and genes associated with A/B compartments changes, based on comparative analyses between the WT and *hht2*Δ mutant. **G** Calculative relative proximity between indicated genes in WT.

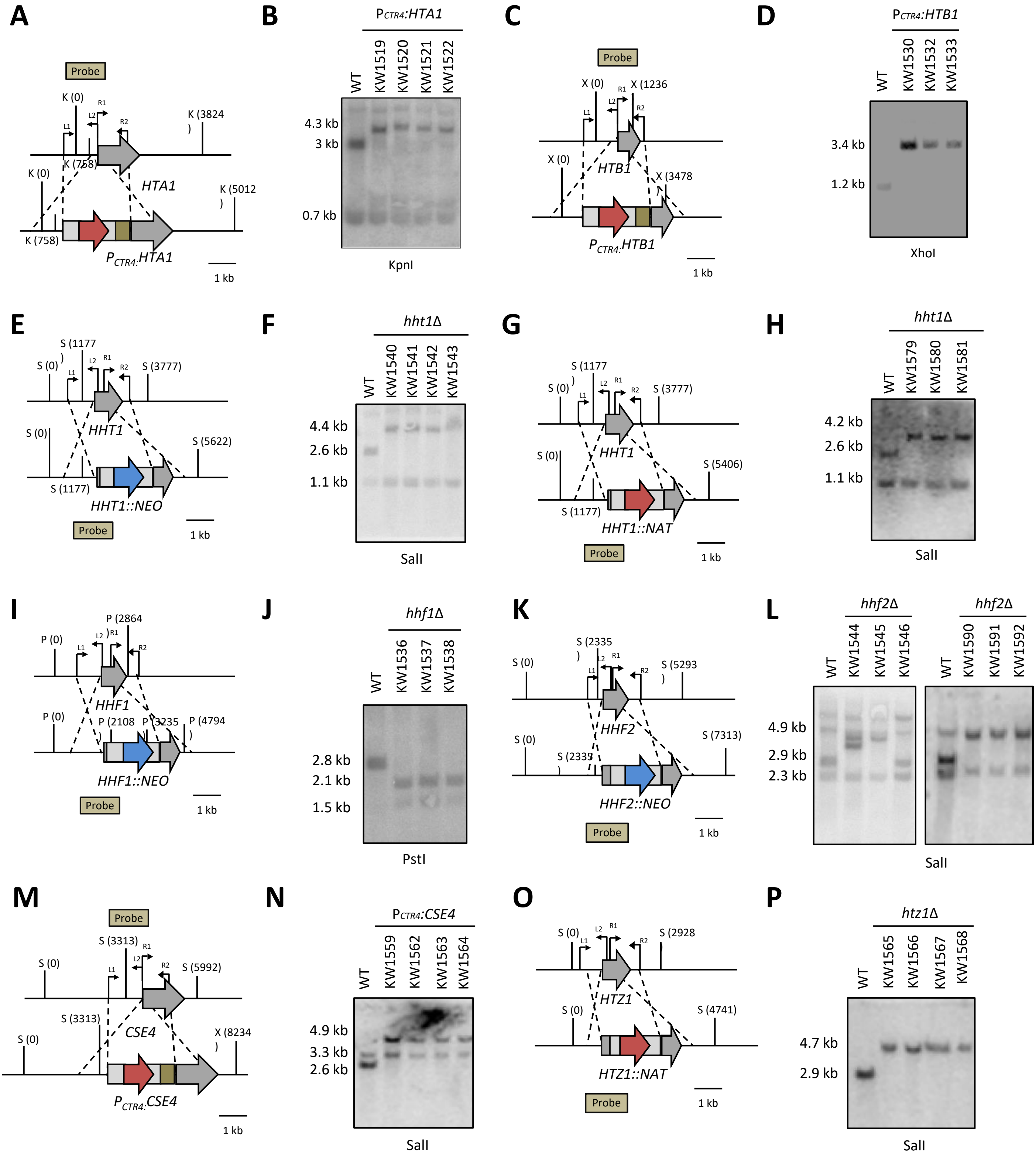
A
Probe
K (0)
L1
L2
R1
R2
K (3824
)
K (758)
K (0)
HTA1
K (5012
)
K (758)
PCTR4:HTA1
1 kb
B
PCTR4:HTA1
WT
KW1519
KW1520
KW1521
KW1522
4.3 kb
3 kb
0.7 kb
KpnI
C
Probe
X (0)
X (1236
X (0)
HTB1
X (3478
)
PCTR4:HTB1
1 kb
D
PCTR4:HTB1
WT
KW1530
KW1532
KW1533
3.4 kb
1.2 kb
XhoI
E
S (1177
)
S (0)
S (3777)
S (0)
HHT1
S (5622)
S (1177)
HHT1::NEO
Probe
1 kb
F
hht1Δ
WT
KW1540
KW1541
KW1542
KW1543
4.4 kb
2.6 kb
1.1 kb
SalI
G
S (1177
)
S (0)
S (3777)
S (0)
HHT1
S (5406)
S (1177)
HHT1::NAT
Probe
1 kb
H
hht1Δ
WT
KW1579
KW1580
KW1581
4.2 kb
2.6 kb
1.1 kb
SalI
I
P (2864
)
P (0)
P (0)
HHF1
P (2108
)
P (3235
P (4794
)
HHF1::NEO
Probe
1 kb
J
hhf1Δ
WT
KW1536
KW1537
KW1538
2.8 kb
2.1 kb
1.5 kb
PstI
K
S (2335
)
S (0)
S (5293
)
S (0)
HHF2
S (2335
)
S (7313)
HHF2::NEO
Probe
1 kb
L
hhf2Δ
hhf2Δ
WT
KW1544
KW1545
KW1546
WT
KW1590
KW1591
KW1592
4.9 kb
2.9 kb
2.3 kb
SalI
M
Probe
S (3313)
S (0)
S (5992)
S (3313)
S (0)
CSE4
X (8234
)
PCTR4:CSE4
1 kb
N
PCTR4:CSE4
WT
KW1559
KW1562
KW1563
KW1564
4.9 kb
3.3 kb
2.6 kb
SalI
O
S (0)
S (2928
S (0)
HTZ1
S (4741)
HTZ1::NAT
Probe
1 kb
P
htz1Δ
WT
KW1565
KW1566
KW1567
KW1568
4.7 kb
2.9 kb
SalI

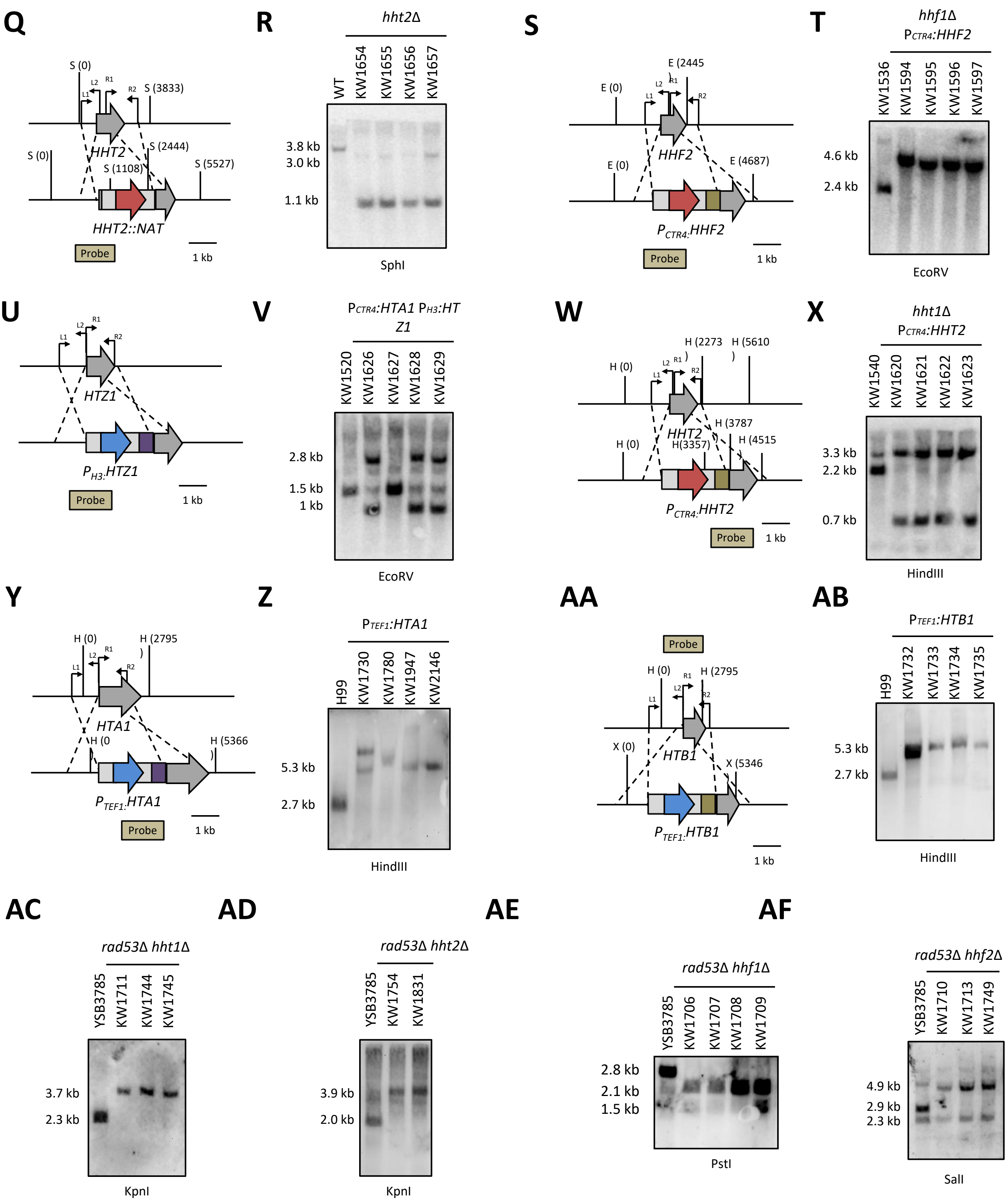
Q
S (0)
L1
L2
R1
R2
S (3833)
HHT2
S (0)
S (2444)
S (1108)
S (5527)
HHT2::NAT
Probe
1 kb
R
hht2Δ
WT
KW1654
KW1655
KW1656
KW1657
3.8 kb
3.0 kb
1.1 kb
SphI
S
E (2445)
E (0)
L1
L2
R1
R2
HHF2
E (0)
E (4687)
PCTR4:HHF2
Probe
1 kb
T
hhf1Δ
PCTR4:HHF2
KW1536
KW1594
KW1595
KW1596
KW1597
4.6 kb
2.4 kb
EcoRV
U
L1
L2
R1
R2
HTZ1
PH3:HTZ1
Probe
1 kb
V
PCTR4:HTA1 PH3:HTZ1
KW1520
KW1626
KW1627
KW1628
KW1629
2.8 kb
1.5 kb
1 kb
EcoRV
W
H (2273)
H (5610)
H (0)
L1
L2
R1
R2
HHT2
H (3787)
H (0)
H(3357)
H (4515)
PCTR4:HHT2
Probe
1 kb
X
hht1Δ
PCTR4:HHT2
KW1540
KW1620
KW1621
KW1622
KW1623
3.3 kb
2.2 kb
0.7 kb
HindIII
Y
H (0)
H (2795)
L1
L2
R1
R2
HTA1
H (0)
H (5366)
PTEF1:HTA1
Probe
1 kb
Z
PTEF1:HTA1
H99
KW1730
KW1780
KW1947
KW2146
5.3 kb
2.7 kb
HindIII
AA
Probe
H (0)
H (2795)
L1
L2
R1
R2
X (0)
HTB1
X (5346)
PTEF1:HTB1
1 kb
AB
PTEF1:HTB1
H99
KW1732
KW1733
KW1734
KW1735
5.3 kb
2.7 kb
HindIII
AC
rad53Δ hht1Δ
YSB3785
KW1711
KW1744
KW1745
3.7 kb
2.3 kb
KpnI
AD
rad53Δ hht2Δ
YSB3785
KW1754
KW1831
3.9 kb
2.0 kb
KpnI
AE
rad53Δ hhf1Δ
YSB3785
KW1706
KW1707
KW1708
KW1709
2.8 kb
2.1 kb
1.5 kb
PstI
AF
rad53Δ hhf2Δ
YSB3785
KW1710
KW1713
KW1749
4.9 kb
2.9 kb
2.3 kb
SalI

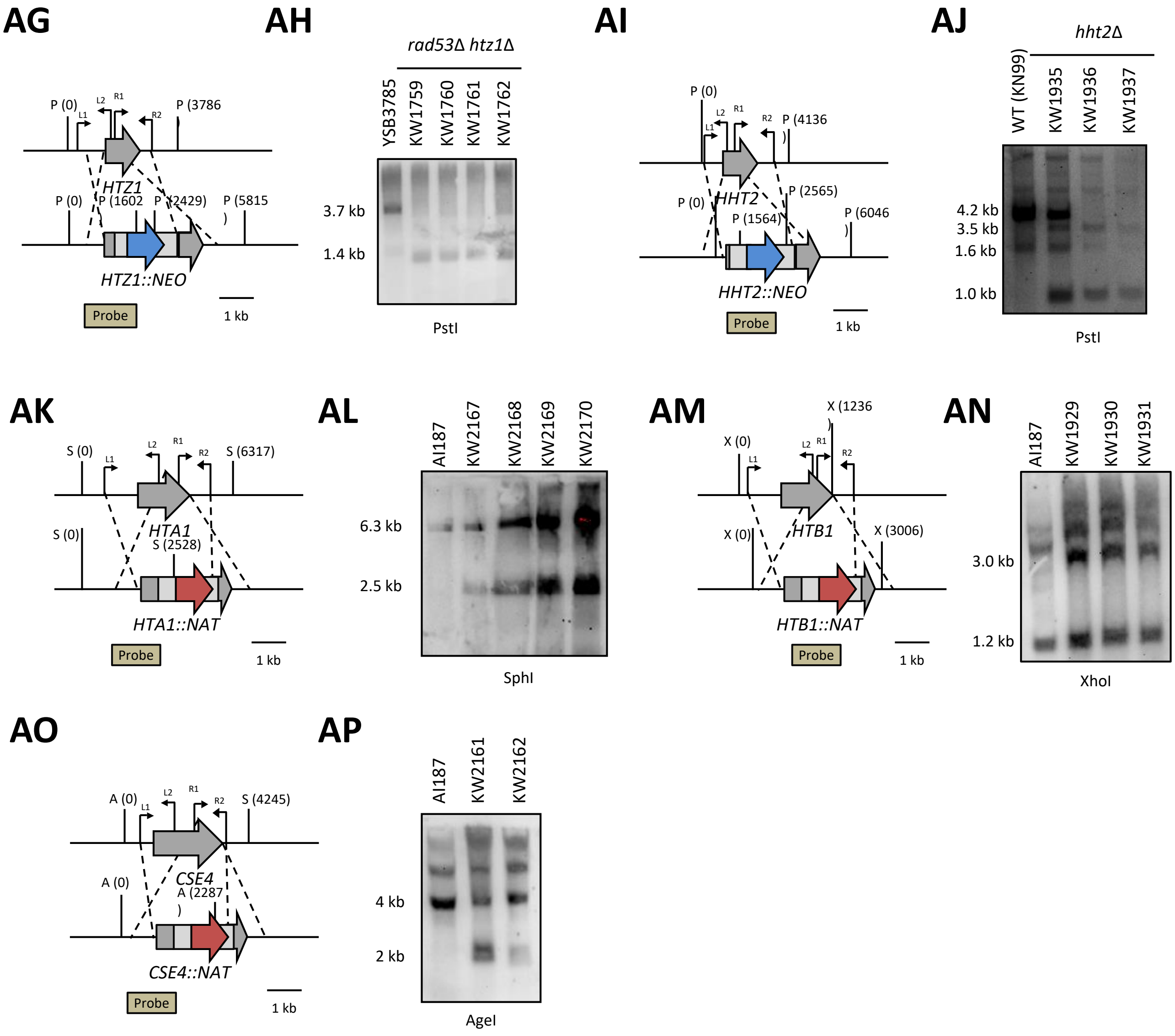


**Fig. S10** Genotypic analysis of the $P_{CTR4}$:*HTA1*, $P_{CTR4}$:*HTB1*, $P_{CTR4}$:*CSE4, hht1Δ*, *hta1Δ*, *htb1Δ*, *hht2Δ*, *hhf1Δ*, *hhf2Δ*, *htz1Δ*, *hhf1Δ* $P_{CTR4}$:*HHF2*, $P_{CTR4}$:*HTA1* $P_{H3}$:*HTZ1*, *hht1Δ* $P_{CTR4}$:*HHT2*, $P_{TEF1}$:*HTA1*, $P_{TEF1}$:*HTB1*, *rad53Δ hht1Δ*, *rad53Δ hht2Δ*, *rad53Δ hhf1Δ* and *rad53Δ hhf2Δ* mutants. Gene disruption strategies were illustrated in the left diagram **A, C, E, G, I, K, M, O, Q, S, U, W, Y, AA, AG, AI, AK, AM, AO** The correct gene disruptions were confirmed by Southern blot analysis using genomic DNA digested with the indicated restriction enzymes. **B, D, F, H, J, L, N, P, R, T, V, X, Z, AB, AC, AD, AE, AF, AH, AJ, AL, AN, and AP**.

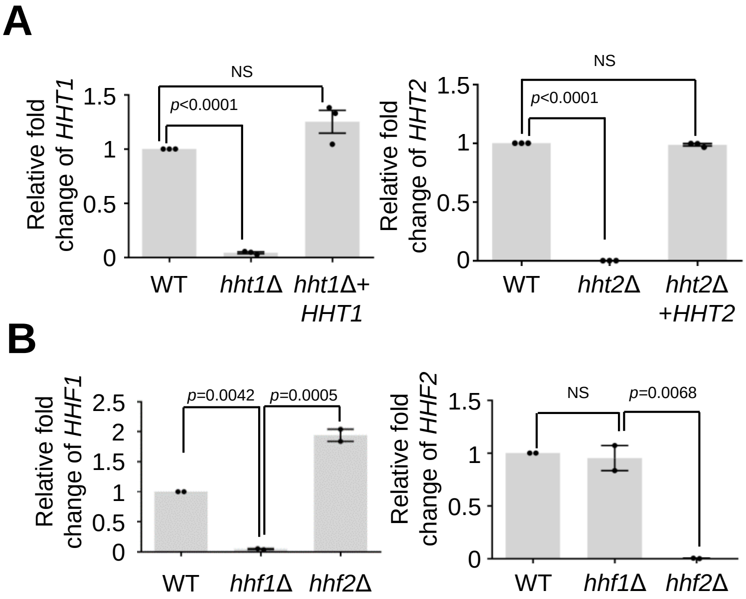


**Fig. S11** Designed primers for *HHT1*, *HHT2*, *HHF1*, and *HHF2* were validated using a gene expression assay. **A and B** The cDNA was synthesized from total RNAs isolated from strains (WT, *hht1*Δ, *hht2*Δ, *hhf1*Δ, and *hhf2*Δ mutants). Independent biological experiments with duplicate technical replicates were performed two or three times. Individual dots represent each biological replicate (n = 2–3) Error bars indicate standard errors of the means. Statistical significance of difference was determined by one-way ANOVA with Bonferroni's multiple-comparison test (NS: not significant).
