## Supplementary Dataset 1 for "Systematic histone mutagenesis reveals nucleosome-dependent maintenance of three-dimensional chromosome architecture and virulence in *Cryptococcus neoformans*"

Supplementary Dataset 1. Transcriptome profiles of WT and *hht2*Δ mutants under basal conditions

| Gene_ID | gene_name | Function | WT(H99)_1_DESeq2_normalized_cou | WT(H99)_2_DESeq2_normalized_cou | WT(H99)_3_DE | *hht2*Δ mutant_1_DESeq2_normalized_cou | *hht2*Δ mutant_2_DESeq2_normalized_cou | *hht2*Δ mutant_3_DESeq2_normalized_cou | padj | log2FoldChange | pvalue | stat | foldChange |
|---|---|---|---|---|---|---|---|---|---|---|---|---|---|
| CNAG_04869 | PNB1 | para-nitrobenzyl esterase | 441.2590452 | 524.0974819 | 521.895184 | 4818.043232 | 4833.305419 | 4657.328648 | 1.00E-50 | 3.252487983 | 0 | 42.26458437 | 9.530077743 |
| CNAG_06332 | - | hypothetical protein | 1120.856185 | 1259.961762 | 1224.671209 | 7118.94875 | 7781.460243 | 7665.87664 | 1.00E-50 | 2.631869675 | 0 | 39.06730582 | 6.198287505 |
| CNAG_05831 | - | endoribonuclease | 289.4359451 | 284.9050809 | 307.5334983 | 2631.762714 | 2566.375465 | 2642.331183 | 1.00E-50 | 3.139844158 | 2.38E-292 | 36.54214923 | 8.814288746 |
| CNAG_04631 | RIK1 | ribitol kinase | 400.3584479 | 390.4878313 | 422.7039198 | 2937.413805 | 3145.283211 | 2929.985178 | 1.00E-50 | 2.878839991 | 3.84E-274 | 35.3770948 | 7.355584516 |
| CNAG_01562 | BLP4 | pr4/barwin domain protein | 963.187364 | 965.8222163 | 725.5854329 | 5972.911176 | 5528.239829 | 5931.788289 | 1.00E-50 | 2.701169082 | 1.86E-195 | 29.82493406 | 6.503286949 |
| CNAG_04828 | H3.3 | Histone H3 | 2546.587869 | 2631.494308 | 2318.833845 | 10074.52158 | 11913.94103 | 11322.86271 | 1.00E-50 | 2.136807272 | 7.94E-191 | 29.46563203 | 4.39787705 |
| CNAG_02925 | - | D-arabinitol 2-dehydrogenase | 10220.87953 | 9634.747023 | 9716.751835 | 28094.23145 | 28836.99861 | 28778.13103 | 1.00E-50 | 1.519778671 | 5.55E-180 | 28.60667611 | 2.867470554 |
| CNAG_04851 | - | transitional endoplasmic reticulum ATPase | 13346.94877 | 13331.39914 | 13279.75974 | 35129.06279 | 37812.83056 | 36874.56056 | 1.00E-50 | 1.443270661 | 6.51E-173 | 28.03258354 | 2.71936661 |
| CNAG_10501 | - | unspecified product | 212236.7795 | 232456.525 | 238929.797 | 46437.80732 | 35447.86589 | 47200.20725 | 1.00E-50 | -2.419313906 | 3.24E-168 | -27.64455303 | 0.186945039 |
| CNAG_00515 | - | mannitol dehydrogenase, variant | 145.780625 | 139.4431383 | 120.9025635 | 1243.289479 | 1232.900902 | 1213.619982 | 1.00E-50 | 3.175203987 | 5.57E-166 | 27.45806279 | 9.032992353 |
| CNAG_02777 | PHO84 | phosphate:H symporter, variant | 3989.069171 | 3644.55089 | 3926.249108 | 940.5793641 | 1074.947441 | 994.1511485 | 1.00E-50 | -1.957975394 | 5.33E-146 | -25.73052755 | 0.257389411 |
| CNAG_04307 | URO1 | Urate oxidase | 135.2794344 | 160.9870605 | 178.478237 | 1268.93995 | 1208.619201 | 1260.224472 | 1.00E-50 | 2.966697409 | 6.06E-146 | 25.72548034 | 7.81744633 |
| CNAG_01332 | - | small subunit ribosomal protein S24e | 23359.07088 | 24527.39358 | 23827.49279 | 10202.56027 | 10185.4709 | 9921.933479 | 1.00E-50 | -1.257583239 | 3.32E-143 | -25.4795726 | 0.418244003 |
| CNAG_01257 | - | aldo-keto reductase | 896.4937876 | 757.8479487 | 772.46406 | 3282.583683 | 3224.160979 | 3643.792238 | 1.00E-50 | 2.049622685 | 7.30E-126 | 23.8636699 | 4.139976806 |
| CNAG_03400 | GRE202 | oxidoreductase | 311.5311983 | 351.6450886 | 298.9717957 | 1507.394863 | 1615.762301 | 1533.512183 | 1.00E-50 | 2.262824129 | 1.95E-124 | 23.72590472 | 4.799300438 |
| CNAG_01224 | - | large subunit ribosomal protein L18-A | 38964.88781 | 41061.65573 | 41149.84888 | 15591.36273 | 15064.04857 | 16857.09555 | 1.00E-50 | -1.365589699 | 7.07E-123 | -23.57432444 | 0.38807578 |
| CNAG_06745 | H3 | Histone H3 | 4976.280707 | 5347.972113 | 5278.613174 | 1891.861074 | 1945.370038 | 1964.325258 | 1.00E-50 | -1.442467491 | 7.29E-122 | -23.47533249 | 0.36793747 |
| CNAG_06811 | RPL22alpha | large subunit ribosomal protein L22e | 41681.17216 | 43064.67178 | 44932.57689 | 19006.59508 | 19582.24223 | 19042.99342 | 1.00E-50 | -1.185264326 | 4.14E-117 | -23.00519517 | 0.439743962 |
| CNAG_03283 | - | large subunit ribosomal protein L24e | 48568.51403 | 48515.41322 | 50756.48573 | 21236.3777 | 22640.89556 | 21286.63808 | 1.00E-50 | -1.197361608 | 4.39E-115 | -22.80194389 | 0.436072039 |
| CNAG_01951 | - | small subunit ribosomal protein S22-A, variant | 58410.15302 | 53033.11936 | 57647.89672 | 23487.17582 | 23751.82648 | 23172.81838 | 1.00E-50 | -1.279749137 | 1.62E-111 | -22.43950741 | 0.41186712 |
| CNAG_03974 | - | hypothetical protein | 885.9062025 | 823.5233614 | 805.4900282 | 2829.656316 | 2638.384745 | 2749.254172 | 1.00E-50 | 1.693664638 | 3.23E-109 | 22.20271768 | 3.234773367 |
| CNAG_06222 | - | large subunit ribosomal protein L32e | 39244.07992 | 39954.14452 | 42149.14032 | 18294.48096 | 18736.07942 | 18450.15304 | 1.00E-50 | -1.144388127 | 1.46E-108 | -22.13485228 | 0.452381511 |
| CNAG_05814 | - | small subunit ribosomal protein S10e | 47368.08381 | 47359.33595 | 51714.12206 | 21259.74634 | 21190.89638 | 20432.93646 | 1.00E-50 | -1.235052625 | 1.59E-108 | -22.13112122 | 0.424827003 |
| CNAG_01300 | - | small subunit ribosomal protein S21e | 31722.46265 | 30592.33877 | 33567.35211 | 13480.70991 | 13755.62002 | 12803.9274 | 1.00E-50 | -1.275513248 | 6.32E-108 | -22.06871196 | 0.413078178 |
| CNAG_00034 | RPL9B | large subunit ribosomal protein L9e | 63913.96002 | 62320.58899 | 66586.92903 | 29202.21984 | 30246.96544 | 29744.92528 | 1.00E-50 | -1.127752641 | 1.10E-104 | -21.72861345 | 0.45762804 |
| CNAG_04209 | - | voltage-gated potassium channel protein beta-2 subunit | 2445.975795 | 2658.322417 | 2268.64377 | 6567.453869 | 6568.171667 | 6770.311582 | 1.00E-50 | 1.418511004 | 2.47E-102 | 21.47854302 | 2.673094802 |
| CNAG_01356 | - | hypothetical protein | 362.8377576 | 327.942914 | 329.8795552 | 1374.953353 | 1501.863984 | 1402.575 | 1.00E-50 | 2.055247593 | 1.60E-101 | 21.3916238 | 4.156149623 |
| CNAG_04891 | - | hypothetical protein | 90.89628226 | 86.64839198 | 73.97607937 | 774.8474305 | 860.9927542 | 1116.476732 | 1.00E-50 | 3.450337445 | 4.11E-101 | 21.34757123 | 10.93087847 |
| CNAG_04792 | - | Phosphatidylserine decarboxylase | 2108.494926 | 2186.422928 | 2190.762898 | 5341.447819 | 5711.667326 | 5976.502942 | 1.00E-50 | 1.377872654 | 4.61E-100 | 21.23429853 | 2.598848715 |
| CNAG_04879 | - | Glycogen debranching enzyme | 2534.76569 | 2479.52135 | 1994.610187 | 7946.151559 | 7576.605886 | 7568.493173 | 1.00E-50 | 1.705345437 | 5.61E-100 | 21.22502826 | 3.261070062 |
| CNAG_04585 | - | hypothetical protein | 71.03088183 | 82.32688598 | 62.2397948 | 751.7349946 | 660.883322 | 660.8374927 | 1.00E-50 | 3.253196883 | 5.89E-99 | 21.11423022 | 9.534761703 |
| CNAG_00655 | - | large subunit acidic ribosomal protein P1 | 59442.6778 | 59921.1282 | 65759.31483 | 26756.019 | 27930.97194 | 26847.43076 | 1.00E-50 | -1.198449933 | 7.33E-99 | -21.10385928 | 0.435743203 |
| CNAG_07466 | - | U3 small nucleolar RNA-associated protein 7, variant 1 | 442315.4359 | 435261.7619 | 363444.9676 | 152310.4886 | 135643.9974 | 139878.5612 | 1.00E-50 | -1.551406053 | 7.23E-99 | -21.10452945 | 0.34117739 |
| CNAG_03739 | - | large subunit ribosomal protein L10-like | 56265.24962 | 56311.74948 | 57218.18768 | 27901.76113 | 28956.61768 | 28409.36895 | 1.00E-50 | -1.00906087 | 2.68E-98 | -21.04247285 | 0.496869582 |
| CNAG_00771 | - | large subunit ribosomal protein L29 | 38449.91881 | 36812.38008 | 38505.10394 | 17870.35094 | 18271.18228 | 17185.68765 | 1.00E-50 | -1.10874957 | 1.66E-97 | -20.95569241 | 0.463695757 |
| CNAG_01884 | - | large subunit ribosomal protein L3 | 96667.16866 | 97286.51215 | 99638.50253 | 49777.97473 | 50487.00561 | 50016.4258 | 1.00E-50 | -0.981437607 | 2.57E-96 | -20.82495799 | 0.5064748 |
| CNAG_04788 | - | hypothetical protein | 2402.672535 | 2299.536879 | 2240.880395 | 5480.839243 | 5608.114521 | 5542.569318 | 1.00E-50 | 1.245177195 | 2.30E-93 | 20.49668846 | 2.370476671 |
| CNAG_04663 | - | hypothetical protein | 354.7303818 | 435.6640196 | 456.7606565 | 1645.11589 | 1650.07109 | 1651.234169 | 1.00E-50 | 1.975408812 | 1.70E-92 | 20.39929167 | 3.932396547 |
| CNAG_00238 | - | isoleucine-tRNA ligase | 36468.24373 | 34696.30988 | 36255.9612 | 16438.26147 | 17387.92224 | 17253.37474 | 1.00E-50 | -1.088033286 | 8.57E-92 | -20.31991068 | 0.4704022 |
| CNAG_02331 | - | small subunit ribosomal protein S9 | 21887.72294 | 21280.11178 | 23392.3392 | 9821.997107 | 10300.23901 | 9669.910559 | 1.00E-50 | -1.17541187 | 7.37E-91 | -20.21400525 | 0.44275734 |
| CNAG_07863 | - | 26S proteasome regulatory subunit N1 | 13223.03481 | 12181.74858 | 12641.982 | 27709.43378 | 29288.83537 | 29983.838 | 1.00E-50 | 1.177345682 | 1.62E-90 | 20.17526335 | 2.261602968 |
| CNAG_02754 | - | small subunit ribosomal protein S12e | 49202.61797 | 47291.49984 | 52282.56903 | 22817.14377 | 23374.94461 | 22074.843 | 1.00E-50 | -1.139561297 | 7.12E-89 | -19.98716993 | 0.453897581 |
| CNAG_02144 | - | large subunit ribosomal protein L1-A | 62245.25923 | 59665.82177 | 62868.24454 | 30567.41804 | 30196.08766 | 31001.59092 | 1.00E-50 | -1.025255945 | 1.05E-88 | -19.96788627 | 0.491323127 |
| CNAG_01735 | - | hypothetical protein | 1901.880996 | 2239.217434 | 1717.319547 | 6344.417286 | 6018.801484 | 5881.10369 | 1.00E-50 | 1.624849479 | 1.72E-88 | 19.94306003 | 3.084099859 |
| CNAG_05555 | - | large subunit ribosomal protein L7Ae | 72627.72158 | 69521.08798 | 79484.56113 | 32744.98446 | 33375.45463 | 32084.06231 | 1.00E-50 | -1.190012038 | 2.04E-86 | -19.70279548 | 0.438299203 |
| CNAG_04763 | PMT2 | dolichyl-phosphate-mannose-protein mannosyltransferase | 2798.570118 | 2931.969774 | 2939.381656 | 6308.783246 | 6710.448263 | 6781.486154 | 1.00E-50 | 1.176567076 | 5.28E-86 | 19.65458731 | 2.260382736 |
| CNAG_01486 | - | large subunit ribosomal protein L15-A | 69347.55716 | 68275.56813 | 76003.67857 | 31895.09047 | 33596.29334 | 32707.94824 | 1.00E-50 | -1.136846757 | 5.62E-86 | -19.6514337 | 0.454752427 |
| CNAG_04724 | - | ubiquitin-protein ligase E3 C | 2649.092724 | 2663.702151 | 2434.987681 | 5989.713801 | 6468.121335 | 6574.942913 | 1.00E-50 | 1.281702613 | 1.25E-85 | 19.61071638 | 2.431257352 |
| CNAG_03015 | - | large subunit ribosomal protein L37-A | 26325.09494 | 24622.31257 | 25903.88772 | 11756.08069 | 12291.0491 | 11140.87887 | 1.00E-50 | -1.142756719 | 1.46E-85 | -19.6028027 | 0.452893356 |
| CNAG_02523 | - | hypothetical protein, variant | 338.3341734 | 386.088701 | 389.5694924 | 1402.352579 | 1329.101338 | 1343.778214 | 1.00E-50 | 1.857148823 | 4.09E-85 | 19.55037588 | 3.622909638 |
| CNAG_05232 | RPL2 | large subunit ribosomal protein L8 | 69678.04811 | 68077.33661 | 74129.95884 | 34605.17827 | 34369.15618 | 34674.01597 | 1.00E-50 | -1.047038417 | 4.31E-85 | -19.54777592 | 0.483960626 |
| CNAG_02928 | - | large subunit ribosomal protein L5e | 97980.83503 | 96311.43664 | 108877.0864 | 44687.06205 | 46863.33684 | 45036.29986 | 1.00E-50 | -1.165884844 | 7.63E-85 | -19.51857925 | 0.445690821 |
| CNAG_03922 | - | hypothetical protein | 39.48346716 | 30.60598741 | 38.76934616 | 478.1797353 | 471.2111616 | 502.9765107 | 1.00E-50 | 3.745856795 | 1.91E-84 | 19.47158533 | 13.4157592 |
| CNAG_12909 | - | unspecified product | 214.5756794 | 242.8069569 | 213.6390838 | 975.0589882 | 1007.56944 | 951.8620463 | 1.00E-50 | 2.117409459 | 2.93E-84 | 19.44964228 | 4.339140975 |
| CNAG_04703 | - | DNA clamp loader | 432.8375684 | 424.8682121 | 402.372455 | 1420.07415 | 1391.175961 | 1433.689439 | 1.00E-50 | 1.739111683 | 4.00E-84 | 19.43371547 | 3.338295542 |
| CNAG_04735 | MEP1 | extracellular elastinolytic metalloproteinase | 791.3028556 | 778.2500553 | 646.6032939 | 2308.726221 | 2462.578625 | 2374.039697 | 1.00E-50 | 1.67425424 | 8.87E-84 | 19.39285972 | 3.191543341 |
| CNAG_01264 | - | homoisocitrate dehydrogenase | 11368.65144 | 11522.85504 | 11809.63574 | 5079.853644 | 5476.803644 | 5587.796292 | 1.00E-50 | -1.119312247 | 4.28E-83 | -19.31175777 | 0.460313211 |
| CNAG_01990 | - | small subunit ribosomal protein S5 | 79008.33615 | 75634.44678 | 86240.55356 | 36985.92291 | 36440.13182 | 36000.69733 | 1.00E-50 | -1.154012248 | 4.78E-83 | -19.30604153 | 0.449373748 |
| CNAG_06605 | - | small subunit ribosomal protein S2 | 46748.11729 | 47765.55485 | 49862.83143 | 23021.07063 | 24639.5027 | 24170.77181 | 1.00E-50 | -1.022447386 | 9.59E-83 | -19.26999253 | 0.492280539 |
| CNAG_05497 | - | Dihydroxy-acid dehydratase | 21456.91824 | 22419.99822 | 21339.52808 | 10624.48203 | 11247.38365 | 11163.00458 | 1.00E-50 | -0.996268745 | 9.54E-83 | -19.27030562 | 0.501294828 |
| CNAG_01305 | - | methionine-tRNA ligase | 14516.89045 | 14017.15587 | 14281.71994 | 7133.101859 | 7211.443367 | 7226.381128 | 1.00E-50 | -1.004480162 | 1.92E-82 | -19.23408533 | 0.498449703 |
| CNAG_02661 | ISP4 | Identified spore protein 4 | 229.7664583 | 243.9006142 | 225.3757555 | 959.874231 | 1115.417395 | 1179.190549 | 1.00E-50 | 2.207562869 | 1.98E-82 | 19.23255503 | 4.618943404 |
| CNAG_03221 | - | large subunit ribosomal protein L29 | 14569.48096 | 15151.62758 | 17881.84365 | 6128.14049 | 6118.452172 | 5883.206813 | 1.00E-50 | -1.408084172 | 2.49E-82 | -19.22049354 | 0.376811742 |
| CNAG_05909 | - | cytochrome c1, heme protein, mitochondrial | 13804.60551 | 14109.77155 | 14867.13009 | 6166.33604 | 6854.328893 | 6534.791586 | 1.00E-50 | -1.144821315 | 3.91E-82 | -19.19721053 | 0.452245698 |
| CNAG_04874 | - | hypothetical protein | 148.0672138 | 176.0324294 | 180.5803878 | 915.4545486 | 809.8542353 | 895.0847466 | 1.00E-50 | 2.362847899 | 4.81E-82 | 19.18634919 | 5.143847599 |
| CNAG_04566 | - | beta-flanking protein | 4791.891291 | 5235.486598 | 4975.231585 | 10652.50886 | 10371.03112 | 10784.78254 | 1.00E-50 | 1.069510436 | 5.68E-82 | 19.17772404 | 2.098721066 |
| CNAG_04011 | - | large subunit ribosomal protein L37a | 40417.61028 | 38361.75004 | 43740.25425 | 18141.92136 | 18931.88479 | 18097.13524 | 1.00E-50 | -1.166795439 | 6.56E-82 | -19.17028137 | 0.445409601 |
| CNAG_04741 | - | endosome protein | 700.2011313 | 728.6636448 | 697.7457836 | 2023.147187 | 1964.698068 | 1964.467439 | 1.00E-50 | 1.470614571 | 8.97E-82 | 19.15397182 | 2.771399269 |
| CNAG_05264 | - | alpha-amylase AmyA, variant | 868.4258059 | 700.7376649 | 902.5226748 | 3117.061388 | 2851.515126 | 2771.522838 | 1.00E-50 | 1.806725593 | 2.00E-81 | 19.11217376 | 3.498473572 |
| CNAG_04142 | - | tartrate transporter | 111.8858134 | 115.7220289 | 109.1563338 | 704.7923032 | 746.1425267 | 947.9298455 | 1.00E-50 | 2.82420443 | 2.32E-81 | 19.10442311 | 7.082233597 |
| CNAG_06248 | - | protein TIF31 | 6517.522455 | 6101.179965 | 5795.950306 | 2661.729288 | 2571.637405 | 2460.555198 | 1.00E-50 | -1.274792292 | 8.97E-81 | -19.03368744 | 0.413284656 |
| CNAG_02361 | - | MFS transporter, ACS family, pantothenate transporter | 1814.605828 | 1483.31478 | 1709.103989 | 445.6405975 | 465.1187033 | 462.3604204 | 1.00E-50 | -1.883580557 | 2.02E-80 | -18.99124024 | 0.271010273 |
| CNAG_00232 | RPL30 | large subunit ribosomal protein L30e | 33449.68049 | 32314.12347 | 35870.92521 | 16361.27062 | 16203.61489 | 15692.51109 | 1.00E-50 | -1.090169133 | 3.73E-80 | -18.9589182 | 0.469706306 |
| CNAG_03127 | - | small subunit ribosomal protein S23 | 58472.34555 | 58050.75461 | 60833.46058 | 30653.84179 | 31698.72655 | 30357.69581 | 1.00E-50 | -0.951300537 | 1.54E-79 | -18.88422349 | 0.517166045 |
| CNAG_03747 | - | large subunit ribosomal protein L27Ae | 84544.66302 | 81775.80647 | 95900.45156 | 38365.79014 | 38470.58728 | 37934.60929 | 1.00E-50 | -1.20767583 | 2.86E-79 | -18.85137171 | 0.432965558 |
| CNAG_12913 | - | unspecified product | 101.3590513 | 112.4754727 | 104.880334 | 726.6041598 | 645.2626096 | 629.8759679 | 1.00E-50 | 2.642522626 | 3.24E-79 | 18.84475795 | 6.244225444 |
| CNAG_04635 | - | endopeptidase | 579.9797921 | 473.3930639 | 436.5341189 | 2097.578889 | 2011.539843 | 1792.398288 | 1.00E-50 | 1.971366506 | 1.24E-78 | 18.77351632 | 3.921393736 |
| CNAG_04731 | PCD102 | hypothetical protein | 387.3335509 | 409.7845341 | 427.9477558 | 1346.841161 | 1426.679544 | 1324.977984 | 1.00E-50 | 1.728565406 | 1.50E-78 | 18.76353307 | 3.313981172 |
| CNAG_06699 | GPD1 | glyceraldehyde-3-phosphate dehydrogenase | 118673.3621 | 112760.6162 | 114065.5128 | 57651.91379 | 60580.70848 | 60145.52302 | 1.00E-50 | -0.969277043 | 2.14E-78 | -18.7446301 | 0.51076195 |
| CNAG_05651 | - | hypothetical protein | 18062.14875 | 16977.95308 | 19418.71968 | 8356.519443 | 8207.64623 | 7908.085794 | 1.00E-50 | -1.169774181 | 1.26E-76 | -18.52639903 | 0.444490909 |
| CNAG_01152 | - | small subunit ribosomal protein S6e, variant | 73079.72862 | 71497.11395 | 77597.00866 | 35950.04296 | 38023.86287 | 37359.48362 | 1.00E-50 | -1.012325937 | 1.57E-76 | -18.51472086 | 0.495746353 |
| CNAG_03644 | CAS3 | capsule related protein | 83.86696317 | 98.48012665 | 120.8662458 | 720.3111906 | 681.6164918 | 628.7802638 | 1.00E-50 | 2.737817234 | 1.63E-76 | 18.51261435 | 6.670603241 |
| CNAG_03000 | - | small subunit ribosomal protein S19e | 53997.46335 | 51819.9741 | 57361.04382 | 26912.56634 | 27249.59767 | 26265.66775 | 1.00E-50 | -1.036326155 | 6.06E-76 | -18.44188794 | 0.487567492 |
| CNAG_00703 | - | large subunit ribosomal protein L31e | 31881.48219 | 32881.90137 | 35329.17342 | 15954.98373 | 16843.67196 | 15654.76185 | 1.00E-50 | -1.062070661 | 9.00E-76 | -18.42049217 | 0.478944151 |
| CNAG_03780 | - | small subunit ribosomal protein S16 | 75723.52114 | 73726.32753 | 82057.96281 | 38067.01791 | 38048.84925 | 38193.22336 | 1.00E-50 | -1.033633832 | 3.25E-75 | -18.35092002 | 0.488478229 |

| | | | | | | | | | | | | | |
|---|---|---|---|---|---|---|---|---|---|---|---|---|---|
| CNAG_05907 | - | pyruvate carboxylase | 11351.00915 | 11020.76348 | 10356.13682 | 5160.092842 | 5053.265356 | 5347.996734 | 1.00E-50 | -1.087853235 | 5.61E-75 | -18.32118526 | 0.470460911 |
| CNAG_04796 | CNA1 | calcineurin a catalytic subunit | 1997.457132 | 2036.617141 | 1940.126173 | 4305.453366 | 4672.252173 | 4534.561019 | 1.00E-50 | 1.162419072 | 5.94E-75 | 18.31802009 | 2.238324293 |
| CNAG_00149 | - | NADH dehydrogenase (ubiquinone) 1 alpha subcomplex 4 | 8529.89453 | 8578.263743 | 9027.260473 | 4347.612174 | 4157.94211 | 4277.805912 | 1.00E-50 | -1.047172711 | 6.44E-75 | -18.31361301 | 0.483915578 |
| CNAG_04801 | - | hypothetical protein | 2524.111938 | 2611.9594 | 2674.882496 | 5544.561651 | 5500.169051 | 5704.67322 | 1.00E-50 | 1.085678987 | 9.72E-75 | 18.29125331 | 2.122374118 |
| CNAG_01181 | - | small subunit ribosomal protein S27Ae | 73180.131 | 70553.288 | 79315.03319 | 36680.89275 | 35688.8679 | 35182.48885 | 1.00E-50 | -1.06792363 | 1.02E-74 | -18.28865452 | 0.477005025 |
| CNAG_04652 | - | enoyl reductase | 523.894727 | 530.435928 | 506.8668493 | 1563.127439 | 1689.75814 | 1506.008354 | 1.00E-50 | 1.594421694 | 2.98E-74 | 18.23013258 | 3.019734465 |
| CNAG_04021 | - | large subunit ribosomal protein L24 | 33261.72038 | 32696.61468 | 36670.75699 | 16284.97544 | 16573.65978 | 16560.6339 | 1.00E-50 | -1.06982597 | 3.33E-74 | -18.2240605 | 0.47637646 |
| CNAG_00605 | - | cytoplasmic protein | 1790.783609 | 1890.081656 | 1659.669735 | 4135.156243 | 4141.703332 | 4262.530646 | 1.00E-50 | 1.21681298 | 1.59E-73 | 18.1382435 | 2.324326888 |
| CNAG_05199 | - | chaperone DnaK | 22577.81113 | 22519.10567 | 22901.80344 | 9014.034158 | 9457.452046 | 10845.5107 | 1.00E-50 | -1.228900427 | 7.00E-73 | -18.05657932 | 0.426642494 |
| CNAG_06095 | - | large subunit ribosomal protein L13e | 79784.97479 | 77289.40706 | 84563.0896 | 41312.51922 | 41523.26874 | 41693.5419 | 1.00E-50 | -0.971874742 | 1.36E-72 | -18.01992665 | 0.509843105 |
| CNAG_04815 | BNI1 | Formin BNI1 | 589.3185144 | 615.5588338 | 608.1772173 | 1978.220527 | 1692.843979 | 1979.849112 | 1.00E-50 | 1.62558239 | 2.45E-72 | 17.98728969 | 3.085667027 |
| CNAG_04737 | - | hypothetical protein | 3488.775087 | 3561.230405 | 2781.638831 | 10274.04514 | 9772.696679 | 8785.405652 | 1.00E-50 | 1.537392434 | 7.98E-72 | 17.92175614 | 2.902693883 |
| CNAG_02209 | NOP56 | nucleolar protein 56 | 11926.83144 | 11392.54882 | 11781.95524 | 6019.394786 | 6128.951573 | 5973.009232 | 1.00E-50 | -0.969371306 | 1.36E-71 | -17.89209578 | 0.510728578 |
| CNAG_00821 | - | large subunit ribosomal protein L34e | 17200.45804 | 16960.71121 | 18894.05004 | 8700.576669 | 8512.159297 | 8366.548914 | 1.00E-50 | -1.068032868 | 1.63E-71 | -17.88188696 | 0.476968909 |
| CNAG_04755 | BCK1 | STE/STE11/BCK1 protein kinase | 1971.785412 | 2037.698327 | 1822.837914 | 4750.931705 | 4771.470759 | 4328.162847 | 1.00E-50 | 1.23310049 | 2.39E-71 | 17.86071326 | 2.350716389 |
| CNAG_02880 | - | GTP-binding protein YchF | 22584.93782 | 22000.94174 | 22049.79195 | 11539.02686 | 11843.42353 | 12153.23784 | 1.00E-50 | -0.922339243 | 9.30E-71 | -17.78459996 | 0.527652769 |
| CNAG_05425 | - | asparagine synthase (glutamine-hydrolyzing) | 14979.40966 | 14745.51932 | 15431.34799 | 7942.573571 | 8093.881341 | 7754.816477 | 1.00E-50 | -0.940035031 | 2.02E-70 | -17.74097124 | 0.521220224 |
| CNAG_02330 | - | large subunit ribosomal protein L21e | 46479.58602 | 47370.16445 | 49696.48457 | 25677.4916 | 26004.96766 | 24602.62378 | 1.00E-50 | -0.927396528 | 2.80E-70 | -17.72282211 | 0.525806351 |
| CNAG_02353 | - | betaine lipid synthase, variant | 177.1767123 | 176.0136698 | 160.3166909 | 726.6309515 | 772.0594285 | 774.1433828 | 1.00E-50 | 2.131760678 | 1.07E-69 | 17.64712641 | 4.382520009 |
| CNAG_04634 | - | hypothetical protein | 421.1646068 | 465.7815359 | 429.0171022 | 1434.19096 | 1315.185414 | 1340.526567 | 1.00E-50 | 1.621879512 | 1.81E-69 | 17.61744607 | 3.077757388 |
| CNAG_07862 | - | fumarate reductase | 1086.634095 | 989.4030501 | 1049.648141 | 2592.659313 | 2699.632383 | 2536.281986 | 1.00E-50 | 1.309367247 | 2.14E-69 | 17.60792204 | 2.478328189 |
| CNAG_05762 | - | large subunit acidic ribosomal protein P2 | 34703.939 | 35264.12082 | 38974.29685 | 16895.90466 | 18208.23806 | 17644.14994 | 1.00E-50 | -1.061796062 | 4.11E-69 | -17.57102433 | 0.47903532 |
| CNAG_00672 | - | small subunit ribosomal protein S11 | 80042.99924 | 75567.67574 | 86414.41254 | 39807.28065 | 39348.72383 | 38776.07296 | 1.00E-50 | -1.052894069 | 7.10E-69 | -17.53996371 | 0.482000294 |
| CNAG_01568 | - | T-complex protein 1 subunit alpha | 8313.81257 | 8468.328817 | 8944.039574 | 3627.539794 | 3678.305281 | 4154.567937 | 1.00E-50 | -1.181790531 | 9.05E-69 | -17.52617561 | 0.440804076 |
| CNAG_03196 | URA5 | Orotate phosphoribosyltransferase | 3462.879507 | 3307.459077 | 3508.351519 | 1555.806614 | 1534.006692 | 1549.061302 | 1.00E-50 | -1.163325006 | 1.44E-68 | -17.49958958 | 0.446482333 |
| CNAG_00640 | - | small subunit ribosomal protein S4-A | 154495.4057 | 147406.5627 | 162756.5164 | 78240.03759 | 81310.63675 | 80251.65443 | 1.00E-50 | -0.969952669 | 1.86E-68 | -17.48515838 | 0.510522812 |
| CNAG_03531 | - | 3-beta hydroxysteroid dehydrogenase/isomerase | 239.1217192 | 296.6746295 | 311.7223567 | 1048.639242 | 1239.702301 | 1137.239508 | 1.00E-50 | 2.002075489 | 2.00E-68 | 17.48098487 | 4.005758618 |
| CNAG_00370 | UBI1 | Predicted ubiquitin-ribosomal 60s subunit protein L40a fusion protein | 25449.41757 | 24430.56034 | 27243.32238 | 13002.22103 | 12009.8221 | 12227.6863 | 1.00E-50 | -1.065880695 | 7.84E-68 | -17.40291898 | 0.477680969 |
| CNAG_04677 | - | xylitol dehydrogenase | 523.9211428 | 534.7560139 | 566.5731939 | 1501.34943 | 1722.472087 | 1766.686483 | 1.00E-50 | 1.604839926 | 3.28E-67 | 17.32069493 | 3.041620006 |
| CNAG_00026 | - | kynurenine aminotransferase | 6454.767849 | 6274.736098 | 6860.209644 | 3145.894021 | 3211.738302 | 3116.591856 | 1.00E-50 | -1.063755598 | 3.69E-67 | -17.31399658 | 0.478385114 |
| CNAG_06274 | - | hypothetical protein | 13645.86554 | 13831.83838 | 14893.81016 | 7138.56654 | 7286.50437 | 6790.118001 | 1.00E-50 | -1.01339676 | 4.32E-67 | -17.30494487 | 0.495378527 |
| CNAG_04819 | - | hypothetical protein | 429.347451 | 486.2505558 | 483.3868216 | 1349.003987 | 1449.024317 | 1399.349625 | 1.00E-50 | 1.571056524 | 8.24E-67 | 17.26768146 | 2.97122225 |
| CNAG_04740 | - | hypothetical protein | 1638.936364 | 1591.64688 | 1660.693205 | 3566.962148 | 3657.829375 | 3499.953213 | 1.00E-50 | 1.117512267 | 8.61E-67 | 17.26512668 | 2.169725099 |
| CNAG_00779 | RPL27 | large subunit ribosomal protein L27e | 46494.69869 | 44921.18611 | 50404.56602 | 23664.76495 | 24267.73664 | 23791.09889 | 1.00E-50 | -0.999158886 | 9.53E-67 | -17.25925519 | 0.500291593 |
| CNAG_05534 | PLR1 | pyridoxal reductase | 805.302745 | 791.1736418 | 840.6400724 | 2078.007281 | 2314.080969 | 2499.382177 | 1.00E-50 | 1.484900247 | 1.79E-66 | 17.22267432 | 2.798978193 |
| CNAG_04642 | TSP2 | tetraspanin Tsp2 | 716.5009746 | 650.0302035 | 630.5715615 | 1927.417718 | 1999.351559 | 1770.283578 | 1.00E-50 | 1.497157553 | 4.05E-66 | 17.17559204 | 2.822859948 |
| CNAG_04738 | SAC6 | fimbrin | 6061.142738 | 5494.104079 | 5800.658004 | 11640.02819 | 12072.90934 | 12212.45801 | 1.00E-50 | 1.033888769 | 4.09E-66 | 17.17498843 | 2.047535931 |
| CNAG_07846 | - | ran-binding protein 3 | 797.124149 | 667.3071915 | 805.4538666 | 2491.995441 | 2232.957958 | 2177.658928 | 1.00E-50 | 1.58853108 | 6.23E-66 | 17.15047919 | 3.007429837 |
| CNAG_04666 | - | 26S protease regulatory subunit 8 | 5143.28365 | 4889.642727 | 5131.973815 | 9737.860269 | 10053.62744 | 9994.405822 | 1.00E-50 | 0.958456148 | 1.05E-65 | 17.12004945 | 1.9432293 |
| CNAG_04837 | MLN1 | bHLH family transcription factor | 19.5965676 | 18.73935304 | 13.15381926 | 387.6395575 | 341.8272445 | 336.6315764 | 1.00E-50 | 4.396321876 | 2.01E-65 | 17.08218135 | 21.05837016 |
| CNAG_04626 | - | hypothetical protein | 177.164649 | 128.641673 | 145.3933573 | 705.6173219 | 763.4799102 | 784.0480204 | 1.00E-50 | 2.307105707 | 3.27E-65 | 17.05395241 | 4.948892511 |
| CNAG_07778 | - | translation initiation factor 2 subunit 1 | 11728.33765 | 11156.60596 | 11674.24564 | 6055.212719 | 6141.897378 | 6100.659754 | 1.00E-50 | -0.933021852 | 3.32E-65 | -17.05297654 | 0.52376013 |
| CNAG_04847 | - | hypothetical protein | 416.4491225 | 411.9190665 | 356.5169856 | 1208.172858 | 1281.684339 | 1277.176666 | 1.00E-50 | 1.655428883 | 4.11E-65 | 17.04050716 | 3.150168246 |
| CNAG_00932 | - | hypothetical protein | 66.3678462 | 131.8588762 | 88.89586282 | 681.7437303 | 678.0369707 | 797.1079597 | 1.00E-50 | 2.906404118 | 6.02E-65 | 17.01820336 | 7.497471421 |
| CNAG_06447 | RPL17 | Large subunit ribosomal protein L17, putative | 58598.49436 | 58225.31221 | 64083.97406 | 30417.99014 | 32189.52349 | 30830.62458 | 1.00E-50 | -0.968692978 | 9.24E-65 | -16.9930667 | 0.51096877 |
| CNAG_12266 | - | unspecified product | 33765.15738 | 35999.99855 | 38530.67771 | 18724.21809 | 18479.94635 | 18278.03809 | 1.00E-50 | -0.979991041 | 4.81E-64 | -16.89607711 | 0.506982888 |
| CNAG_04835 | - | dihydrodipicolinate synthase | 59.32163533 | 43.53727469 | 70.74863135 | 529.0538265 | 447.6912715 | 475.4323039 | 1.00E-50 | 3.060683491 | 6.09E-64 | 16.88211091 | 8.343678053 |
| CNAG_05437 | - | nascent polypeptide-associated complex subunit beta | 12748.86448 | 11591.89842 | 12597.76662 | 6253.656782 | 6103.901896 | 6141.761781 | 1.00E-50 | -1.013468725 | 1.26E-63 | -16.83937343 | 0.495353817 |
| CNAG_06666 | - | starch phosphorylase | 1188.268868 | 1311.506464 | 1142.446029 | 2821.288765 | 3001.451822 | 3154.507185 | 1.00E-50 | 1.287049718 | 1.73E-63 | 16.82025295 | 2.440285115 |
| CNAG_00509 | - | translation initiation factor 3 subunit M | 9616.981641 | 8932.784733 | 9370.703337 | 4826.287885 | 4799.733567 | 4740.730951 | 1.00E-50 | -0.974264481 | 1.79E-63 | -16.81828387 | 0.50899928 |
| CNAG_03599 | - | mandelate racemase/muconate lactonizing enzyme | 466.7421248 | 429.2087954 | 572.960177 | 1592.729219 | 1634.447396 | 1696.758902 | 1.00E-50 | 1.730628502 | 2.03E-63 | 16.81081223 | 3.318723651 |
| CNAG_02217 | CHS7 | Chitin synthase | 193.6055046 | 274.0438644 | 172.0756052 | 990.4100027 | 1050.622835 | 1195.767347 | 1.00E-50 | 2.326404692 | 2.79E-63 | 16.79196257 | 5.015538798 |
| CNAG_01168 | - | H/ACA ribonucleoprotein complex subunit 4 | 15764.08304 | 14952.4008 | 16226.89993 | 7761.167611 | 7879.059696 | 8295.439079 | 1.00E-50 | -0.987274831 | 4.32E-63 | -16.76614097 | 0.504429715 |
| CNAG_03012 | CQS1 | quorum sensing-like molecule | 8165.62752 | 9389.022307 | 9270.92039 | 23695.58284 | 20955.59668 | 19529.63359 | 1.00E-50 | 1.243868956 | 4.60E-63 | 16.76237533 | 2.368328092 |
| CNAG_04448 | - | large subunit ribosomal protein L19e | 93341.42958 | 90656.03984 | 103221.7227 | 49132.78987 | 48502.24217 | 47223.27957 | 1.00E-50 | -1.003105919 | 1.50E-62 | -16.69186263 | 0.498924728 |
| CNAG_03375 | - | Succinyl-CoA synthetase alpha subunit | 12189.64105 | 11856.93174 | 13253.59228 | 6052.028906 | 6378.224302 | 6163.96783 | 1.00E-50 | -1.019923983 | 2.97E-62 | -16.651113 | 0.493142336 |
| CNAG_04697 | - | zinc finger protein | 1231.409739 | 1230.704591 | 1243.718687 | 2765.345602 | 2875.530576 | 2670.677378 | 1.00E-50 | 1.150407612 | 3.45E-62 | 16.642107 | 2.219766018 |
| CNAG_00699 | LPI15 | transmembrane receptor | 234.3739213 | 260.019813 | 189.1180691 | 890.0772601 | 937.6539576 | 906.3379362 | 1.00E-50 | 1.985831096 | 5.61E-62 | 16.61299245 | 3.960907737 |
| CNAG_04598 | - | hypothetical protein, variant | 106.0081742 | 130.7654552 | 133.6439207 | 599.9515149 | 599.5112526 | 615.3729835 | 1.00E-50 | 2.281920675 | 1.11E-61 | 16.57209207 | 4.863249725 |
| CNAG_03701 | - | 3-phosphoshikimate 1-carboxyvinyltransferase | 8996.889947 | 8384.361703 | 8676.446202 | 4495.517025 | 4270.949856 | 4493.152765 | 1.00E-50 | -0.9902502 | 1.56E-61 | -16.55142487 | 0.503390467 |
| CNAG_12640 | - | unspecified product | 896.9271024 | 831.381834 | 900.722157 | 226.9469122 | 258.1415529 | 200.3998219 | 1.00E-50 | -1.95722521 | 2.87E-61 | -16.51486574 | 0.257523285 |
| CNAG_01119 | - | POT family proton-dependent oligopeptide transporter | 12554.09905 | 13852.2548 | 13170.49558 | 6799.230301 | 7079.504927 | 7191.93005 | 1.00E-50 | -0.924281735 | 7.22E-61 | -16.45907533 | 0.526942798 |
| CNAG_04976 | - | zuotin | 6810.914678 | 6448.223134 | 7060.720929 | 3301.554081 | 3391.156961 | 3435.177193 | 1.00E-50 | -1.020132239 | 7.52E-61 | -16.45657924 | 0.493071155 |
| CNAG_03068 | - | hypothetical protein | 132.8134752 | 124.310286 | 137.9079325 | 582.6567452 | 631.5176432 | 610.9774547 | 1.00E-50 | 2.198227839 | 1.19E-60 | 16.42854993 | 4.589152786 |
| CNAG_06755 | - | threonine-tRNA ligase | 7797.735107 | 7928.569746 | 7558.846368 | 4246.509687 | 4148.514747 | 4215.594467 | 1.00E-50 | -0.899915998 | 1.34E-60 | -16.42170487 | 0.535917934 |
| CNAG_06633 | - | small subunit ribosomal protein S15 | 66710.8299 | 63584.48711 | 69227.51461 | 36874.01263 | 35895.94486 | 34913.7897 | 1.00E-50 | -0.90536148 | 2.20E-60 | -16.39137183 | 0.533898914 |
| CNAG_01695 | - | hypothetical protein | 112.9689153 | 92.00392009 | 89.94610492 | 502.3761832 | 586.7762009 | 587.505928 | 1.00E-50 | 2.49950253 | 4.58E-60 | 16.34680421 | 5.654903989 |
| CNAG_00311 | - | 3-hydroxyisobutyryl-CoA hydrolase | 221021.1963 | 236798.9054 | 244291.1454 | 125617.763 | 112021.1921 | 121270.9921 | 1.00E-50 | -0.982941419 | 5.05E-60 | -16.34087105 | 0.505947144 |
| CNAG_01480 | - | large subunit ribosomal protein L12 | 14724.67054 | 13936.3356 | 14650.6705 | 7119.890567 | 6888.823873 | 5950.903666 | 1.00E-50 | -1.133528836 | 5.31E-60 | -16.33785016 | 0.455799474 |
| CNAG_04068 | - | large subunit ribosomal protein L28e | 9432.712071 | 10752.45122 | 11014.02702 | 4967.87135 | 4653.015707 | 5043.816874 | 1.00E-50 | -1.103909001 | 7.46E-60 | -16.31706715 | 0.465254174 |
| CNAG_05465 | GIB2 | G protein beta subunit Gib2 | 27041.10371 | 25620.07682 | 28602.01345 | 14065.36716 | 14491.49566 | 13734.1514 | 1.00E-50 | -0.957994872 | 1.28E-59 | -16.28431456 | 0.514771871 |
| CNAG_13174 | - | unspecified product | 35000.57207 | 35015.26192 | 37963.34586 | 19615.41463 | 18141.80332 | 19044.02681 | 1.00E-50 | -0.942043149 | 3.07E-59 | -16.23043275 | 0.520495231 |
| CNAG_07865 | - | ferro-O2-oxidoreductase | 1912.161029 | 1863.145575 | 1915.567831 | 3796.37364 | 4104.625628 | 4022.808209 | 1.00E-50 | 1.05170385 | 3.50E-59 | 16.22249406 | 2.072976627 |
| CNAG_04906 | - | 26S protease regulatory subunit 10B | 3866.923093 | 3476.071027 | 3585.640751 | 7400.709988 | 7828.305987 | 7398.862953 | 1.00E-50 | 1.034285752 | 5.16E-59 | 16.19854464 | 2.048099425 |
| CNAG_03098 | - | hypothetical protein | 2783.221846 | 2674.98783 | 2774.637732 | 1231.255691 | 1130.279928 | 1268.173229 | 1.00E-50 | -1.196784593 | 9.09E-59 | -16.16372019 | 0.436246484 |
| CNAG_02234 | - | large subunit ribosomal protein L6e | 39761.43882 | 38227.1032 | 42630.13196 | 19710.2405 | 21618.84575 | 20588.27656 | 1.00E-50 | -0.97773872 | 1.80E-58 | -16.12141487 | 0.507775003 |
| CNAG_05556 | - | hypothetical protein | 38649.66603 | 35674.67673 | 44428.0635 | 18031.43276 | 17910.59549 | 17464.3456 | 1.00E-50 | -1.168804845 | 1.46E-57 | -15.99149325 | 0.44478966 |
| CNAG_04657 | - | Short-chain dehydrogenase | 366.3541224 | 506.7046943 | 404.5047507 | 1463.767678 | 1436.962641 | 1337.23568 | 1.00E-50 | 1.717048933 | 1.79E-57 | 15.97891624 | 3.287632257 |
| CNAG_00141 | - | monooxygenase | 229.6939763 | 254.6280368 | 259.4481204 | 821.4327409 | 883.259321 | 839.7996149 | 1.00E-50 | 1.761976755 | 3.33E-57 | 15.94025303 | 3.391625211 |
| CNAG_04259 | - | H/ACA ribonucleoprotein complex subunit 1 | 2584.86877 | 2752.521603 | 2884.434318 | 1124.542393 | 1163.942696 | 1267.059351 | 1.00E-50 | -1.22468232 | 3.41E-57 | -15.9387216 | 0.427891724 |
| CNAG_04609 | AGO1 | argonaute | 4767.243183 | 4660.126796 | 4244.71722 | 9050.714404 | 9058.223394 | 8890.932583 | 1.00E-50 | 0.966485617 | 4.33E-57 | 15.92385121 | 1.954074697 |
| CNAG_04708 | - | hypothetical protein | 446.7963161 | 444.2340483 | 484.4319617 | 1247.040078 | 1308.366886 | 1251.760438 | 1.00E-50 | 1.454502027 | 4.39E-57 | 15.92294393 | 2.740619471 |
| CNAG_03053 | - | large subunit ribosomal protein L23 | 20111.74232 | 20172.55488 | 22318.45623 | 11025.38958 | 11124.91785 | 10338.18226 | 1.00E-50 | -0.961810134 | 2.76E-56 | -15.80746376 | 0.513412336 |
| CNAG_04951 | - | 3-deoxy-7-phosphoheptulonate synthase | 9888.197721 | 11954.8293 | 11641.12321 | 5332.883119 | 5306.01314 | 5367.992308 | 1.00E-50 | -1.079342766 | 1.34E-55 | -15.70774787 | 0.473244365 |
| CNAG_06817 | UAP1 | uric acid xanthine permease | 213.4009751 | 284.7860177 | 264.7906518 | 889.9702189 | 972.1752834 | 893.0969932 | 1.00E-50 | 1.841043777 | 1.35E-55 | 15.70697793 | 3.582691391 |
| CNAG_01796 | - | hypothetical protein, variant 3 | 19.5984059 | 42.44007326 | 30.22810367 | 377.2389505 | 359.1264413 | 326.7632014 | 1.00E-50 | 3.530848516 | 1.61E-55 | 15.69608253 | 11.55822949 |
| CNAG_04904 | CHC1 | clathrin heavy chain | 13445.94483 | 12911.09826 | 11899.711 | 23870.80799 | 24546.35415 | 24460.92953 | 1.00E-50 | 0.914460155 | 3.70E-55 | 15.64317653 | 1.884863639 |

| | | | | | | | | | | | | | |
|---|---|---|---|---|---|---|---|---|---|---|---|---|---|
| CNAG_03612 | - | Threonine synthase | 3119.788162 | 3417.30931 | 3394.264945 | 1629.031084 | 1641.882987 | 1575.690141 | 1.00E-50 | -1.05024264 | 4.85E-55 | -15.62596617 | 0.482886943 |
| CNAG_06906 | TIF3 | Translation initiation factor 4B | 5034.884722 | 5125.113542 | 5628.449332 | 2636.927159 | 2664.026073 | 2597.022909 | 1.00E-50 | -1.014704673 | 9.87E-55 | -15.58055946 | 0.494929632 |
| CNAG_04257 | - | hypothetical protein | 2735.582643 | 3167.269637 | 3277.912079 | 1221.043464 | 1131.958943 | 1335.910069 | 1.00E-50 | -1.330403312 | 4.92E-54 | -15.47759304 | 0.397657059 |
| CNAG_12914 | - | unspecified product | 53.46923527 | 62.90752508 | 69.66962317 | 440.1048318 | 389.027309 | 397.8064554 | 1.00E-50 | 2.721255955 | 7.70E-54 | 15.4486676 | 6.594466531 |
| CNAG_06998 | - | protein transporter SEC61 subunit alpha | 8898.958361 | 8931.688882 | 9052.916311 | 4640.248738 | 4953.109934 | 5075.942208 | 1.00E-50 | -0.889165446 | 1.02E-53 | -15.43030689 | 0.539926359 |
| CNAG_01761 | - | hypothetical protein | 5960.401144 | 5000.127393 | 5125.148637 | 2459.427158 | 2300.802939 | 2177.462159 | 1.00E-50 | -1.229591231 | 3.00E-53 | -15.36078785 | 0.426438255 |
| CNAG_07807 | - | Histone H4 | 8952.698623 | 11298.19763 | 10990.01934 | 23772.27384 | 23278.48224 | 22388.24271 | 1.00E-50 | 1.138004049 | 3.08E-53 | 15.35907217 | 2.200763395 |
| CNAG_00490 | - | acetyl-CoA acyltransferase | 630.0837883 | 621.9899881 | 530.3335115 | 1602.679894 | 1565.230202 | 1796.666018 | 1.00E-50 | 1.463592116 | 4.28E-53 | 15.3377311 | 2.757941998 |
| CNAG_02430 | - | ABC transporter ABCC.6 | 259.9899197 | 236.325828 | 249.85539 | 777.6957269 | 852.1474655 | 876.3338036 | 1.00E-50 | 1.735927301 | 4.93E-53 | 15.32854236 | 3.330935229 |
| CNAG_04004 | - | small subunit ribosomal protein S1 | 138002.0734 | 126901.9834 | 138708.0386 | 73940.05785 | 75585.85307 | 75830.95516 | 1.00E-50 | -0.856498939 | 5.27E-53 | -15.32422334 | 0.552291206 |
| CNAG_04670 | - | hypothetical protein | 2699.203749 | 2589.33163 | 2319.796072 | 5239.332908 | 5530.395642 | 5870.015993 | 1.00E-50 | 1.113950321 | 1.18E-52 | 15.27163271 | 2.164374758 |
| CNAG_07848 | - | hypothetical protein | 971.0468821 | 1081.998926 | 1069.877044 | 2247.685533 | 2323.415902 | 2364.182193 | 1.00E-50 | 1.136813319 | 1.68E-52 | 15.24861745 | 2.198947742 |
| CNAG_05509 | HIS3 | Imidazoleglycerol-phosphate dehydratase | 5513.455727 | 5021.706808 | 5181.691721 | 2683.640548 | 2685.59261 | 2715.785292 | 1.02E-50 | -0.974828807 | 2.31E-52 | -15.22779518 | 0.508800218 |
| CNAG_04445 | - | small subunit ribosomal protein S7e | 38579.79409 | 40130.89865 | 41804.73893 | 21033.38646 | 23438.90135 | 21692.86746 | 1.07E-50 | -0.880401249 | 2.43E-52 | -15.22468146 | 0.543216328 |
| CNAG_03198 | - | small subunit ribosomal protein S8e | 111752.2731 | 110867.601 | 129792.3853 | 60518.39933 | 59631.00413 | 59163.05046 | 1.21E-50 | -0.990305 | 2.75E-52 | -15.21639108 | 0.503371346 |
| CNAG_13073 | - | unspecified product | 843738.9183 | 803874.2815 | 795955.8408 | 449465.1039 | 487550.2364 | 465693.9712 | 1.95E-50 | -0.816326492 | 4.50E-52 | -15.18417143 | 0.567886101 |
| CNAG_01976 | - | large subunit ribosomal protein L23 | 16129.57823 | 15466.31968 | 18385.28783 | 8179.152207 | 6996.563863 | 7032.183268 | 1.95E-50 | -1.185924501 | 4.49E-52 | -15.18431264 | 0.439542782 |
| CNAG_01170 | - | small subunit ribosomal protein S17 | 49640.65533 | 47077.16153 | 52110.93776 | 28199.99059 | 27484.26417 | 26651.96198 | 3.24E-50 | -0.869720584 | 7.53E-52 | -15.15041624 | 0.54725283 |
| CNAG_02811 | - | small subunit ribosomal protein S29 | 12979.04495 | 13298.48128 | 13704.73346 | 7033.562967 | 6749.119334 | 5837.647089 | 4.07E-50 | -1.042428672 | 9.49E-52 | -15.13517445 | 0.485509466 |
| CNAG_06471 | - | small subunit ribosomal protein S12 | 1174.046787 | 1174.07255 | 1346.496453 | 384.2265482 | 424.6529958 | 457.8257114 | 4.44E-50 | -1.561419464 | 1.04E-51 | -15.12905314 | 0.338817556 |
| CNAG_03819 | ERG6 | sterol 24-C-methyltransferase | 12064.72352 | 11688.87062 | 13177.88346 | 6104.180688 | 6589.575808 | 6454.802971 | 4.70E-50 | -0.963241453 | 1.11E-51 | -15.12504563 | 0.512903225 |
| CNAG_03461 | - | hypothetical protein | 70.98152194 | 95.20938128 | 88.86231184 | 448.0997714 | 510.953151 | 443.4112643 | 5.90E-50 | 2.454512031 | 1.40E-51 | 15.10973603 | 5.48127695 |
| CNAG_06679 | - | anthranilate synthase component I | 3227.12929 | 3391.452858 | 3131.988132 | 1605.672077 | 1685.03616 | 1648.929834 | 6.70E-50 | -0.99632682 | 1.60E-51 | -15.10097751 | 0.501274649 |
| CNAG_03303 | - | small subunit ribosomal protein S27 | 18007.39082 | 17601.80007 | 19054.06362 | 10162.34309 | 9151.331917 | 8908.301617 | 7.18E-50 | -0.969215279 | 1.72E-51 | -15.09610066 | 0.510783817 |
| CNAG_01598 | - | template-activating factor I | 5199.33542 | 4666.171383 | 5091.005861 | 2545.888545 | 2481.148381 | 2504.869784 | 2.28E-49 | -1.005768555 | 5.48E-51 | -15.0194264 | 0.498004763 |
| CNAG_00058 | - | T-complex protein 1 subunit epsilon | 11598.81658 | 11609.11847 | 11882.21676 | 6128.347368 | 6527.487246 | 6782.246905 | 2.87E-49 | -0.867469588 | 6.93E-51 | -15.00381351 | 0.54810736 |
| CNAG_03890 | - | protein lysine methyltransferase SET5 | 744.2366727 | 748.4440611 | 830.3316546 | 238.7099122 | 242.6279608 | 235.8547821 | 3.38E-49 | -1.714733449 | 8.21E-51 | -14.99258682 | 0.304658849 |
| CNAG_04583 | DDT1 | hypothetical protein | 1392.491382 | 1412.752701 | 1384.468931 | 2783.900377 | 2808.971645 | 2789.466904 | 4.00E-49 | 0.985511096 | 9.76E-51 | 14.98106275 | 1.980014638 |
| CNAG_04641 | - | general transcription factor 3C polypeptide 3 (transcription factor C subunit 4) | 2076.779552 | 2150.777537 | 2124.570865 | 4126.488423 | 3998.387123 | 3955.186503 | 8.62E-49 | 0.912569056 | 2.12E-50 | 14.92955002 | 1.882394559 |
| CNAG_04742 | - | DNA primase large subunit | 606.774666 | 733.9944498 | 624.1498285 | 1704.748188 | 1594.534441 | 1686.931921 | 2.27E-48 | 1.329903941 | 5.61E-50 | 14.86442558 | 2.513859363 |
| CNAG_05633 | - | ubiquinol-cytochrome c reductase subunit 9 | 5724.93768 | 5459.14669 | 6170.206253 | 2842.37619 | 3024.605647 | 2713.617356 | 3.07E-48 | -1.032067156 | 7.60E-50 | -14.84407017 | 0.489008973 |
| CNAG_05496 | - | 2-isopropylmalate synthase | 9872.989895 | 11886.94041 | 10678.1869 | 5255.832024 | 5667.488578 | 5430.145637 | 3.37E-48 | -1.002554666 | 8.39E-50 | -14.83747356 | 0.499115404 |
| CNAG_05455 | - | translation initiation factor eIF-1A | 8131.707136 | 7785.333241 | 8073.919544 | 4590.567399 | 4476.318345 | 4295.539996 | 4.19E-48 | -0.859860285 | 1.05E-49 | -14.82240825 | 0.551005916 |
| CNAG_04831 | - | hypothetical protein | 33.61923155 | 56.43026007 | 40.88833456 | 338.5689012 | 340.8368027 | 364.3163693 | 5.05E-48 | 2.99995483 | 1.27E-49 | 14.80955635 | 7.999749527 |
| CNAG_04985 | - | Nascent polypeptide-associated complex subunit alpha | 8722.771596 | 9820.515294 | 10622.61982 | 4735.1697 | 4926.528977 | 4633.075779 | 9.54E-48 | -1.043779597 | 2.41E-49 | -14.76646146 | 0.485055052 |
| CNAG_07538 | - | calcium/proton exchanger | 1119.304913 | 851.5328035 | 780.9708922 | 2676.057689 | 2784.240567 | 2710.457071 | 1.19E-47 | 1.554625916 | 3.01E-49 | 14.75142558 | 2.937575469 |
| CNAG_05395 | VAM6 | rab guanyl-nucleotide exchange factor | 129.2612967 | 110.2968983 | 119.7748157 | 543.2508808 | 514.0552043 | 501.1758109 | 1.21E-47 | 2.106030919 | 3.09E-49 | 14.74974057 | 4.305052762 |
| CNAG_01789 | - | hypothetical protein | 2796.295484 | 2930.347818 | 3052.985215 | 1478.07285 | 1377.036581 | 1461.31595 | 1.70E-47 | -1.03923202 | 4.37E-49 | -14.72636286 | 0.486586426 |
| CNAG_04823 | - | peptidyl-prolyl cis-trans isomerase H | 341.7713594 | 280.5344814 | 406.5914742 | 1133.539116 | 1148.797174 | 1226.021166 | 1.90E-47 | 1.755706707 | 4.89E-49 | 14.71868481 | 3.376916968 |
| CNAG_02710 | - | T-complex protein 1 subunit gamma | 7017.70758 | 7081.73064 | 7402.011024 | 3629.241228 | 3580.906554 | 4022.394936 | 2.11E-47 | -0.951971324 | 5.47E-49 | -14.71112937 | 0.516925643 |
| CNAG_04114 | - | small subunit ribosomal protein S0 | 40257.86047 | 41000.41211 | 45941.46266 | 21870.81851 | 22924.75528 | 23385.843 | 2.60E-47 | -0.915014119 | 6.76E-49 | -14.69676692 | 0.530338681 |
| CNAG_05800 | - | large subunit ribosomal protein L33-b | 15891.41574 | 15160.35994 | 17006.45819 | 8826.779434 | 8206.84287 | 7797.033127 | 3.57E-47 | -0.968357534 | 9.33E-49 | -14.67490918 | 0.51108759 |
| CNAG_01336 | - | hypothetical protein | 100.1418404 | 128.5861414 | 121.9027907 | 507.2386892 | 527.9579242 | 502.2537507 | 4.45E-47 | 2.123187953 | 1.17E-48 | 14.65967645 | 4.356555593 |
| CNAG_01045 | - | hypothetical protein | 1946.036376 | 1942.332784 | 2246.624087 | 794.5190358 | 771.421749 | 897.3916042 | 4.66E-47 | -1.332262155 | 1.23E-48 | -14.65615571 | 0.397145027 |
| CNAG_06475 | - | hypothetical protein | 2515.970853 | 2644.793248 | 2543.269096 | 1259.340994 | 1320.177225 | 1176.052724 | 7.79E-47 | -1.051575195 | 2.06E-48 | -14.62096509 | 0.482441127 |
| CNAG_01485 | - | D-lactate dehydrogenase (cytochrome) | 1552.441881 | 1617.945409 | 1476.821369 | 693.4624095 | 678.3419695 | 664.2823722 | 7.91E-47 | -1.205646806 | 2.11E-48 | -14.6196165 | 0.433574915 |
| CNAG_02103 | - | hypothetical protein, variant | 484.1182028 | 434.5461022 | 507.8831854 | 1199.4002 | 1305.75241 | 1302.774776 | 4.23E-46 | 1.401576375 | 1.13E-47 | 14.50472723 | 2.641900944 |
| CNAG_04621 | - | glycogen(starch) synthase, variant | 1542.014134 | 1676.717872 | 1470.88502 | 3123.104477 | 3376.59098 | 3528.705101 | 4.63E-46 | 1.082095446 | 1.24E-47 | 14.49824652 | 2.117108849 |
| CNAG_04752 | - | hypothetical protein | 820.3526356 | 874.044879 | 804.3463107 | 1830.614843 | 1812.75031 | 1812.485151 | 6.28E-46 | 1.111604142 | 1.69E-47 | 14.47697997 | 2.160857809 |
| CNAG_12692 | - | unspecified product | 1638.977883 | 1765.574157 | 1876.602818 | 742.4540239 | 641.1560906 | 756.4132265 | 7.71E-46 | -1.318156417 | 2.09E-47 | -14.46255977 | 0.401047099 |
| CNAG_05243 | XKS1 | xylulokinase | 885.7037464 | 784.6695065 | 802.2226459 | 1844.562287 | 1893.002904 | 1958.940659 | 7.91E-46 | 1.188977298 | 2.16E-47 | 14.46040206 | 2.279910668 |
| CNAG_00819 | - | small subunit ribosomal protein S30 | 18734.88063 | 17305.55558 | 20873.37063 | 9636.882766 | 9713.743906 | 9253.557708 | 9.21E-46 | -1.008515266 | 2.52E-47 | -14.44962578 | 0.497057526 |
| CNAG_06231 | - | large subunit ribosomal protein L13 | 41267.99831 | 42471.11183 | 46919.41926 | 24511.65667 | 24650.76832 | 23442.48622 | 1.41E-45 | -0.863036817 | 3.87E-47 | -14.42002994 | 0.549794044 |
| CNAG_05150 | - | ATP-binding cassette transporter | 3537.761961 | 3519.733939 | 3448.724838 | 1928.774162 | 1896.381021 | 1847.638006 | 2.42E-45 | -0.904529412 | 6.68E-47 | -14.38236525 | 0.534206927 |
| CNAG_04475 | HPP3 | Phosphoglycerate mutase-like superfamily protein | 2064.075549 | 2273.036418 | 2090.034772 | 1058.56101 | 1019.089881 | 991.7555009 | 3.15E-45 | -1.081642054 | 8.73E-47 | -14.36379156 | 0.472490735 |
| CNAG_04575 | - | G2/mitotic-specific cyclin 1/2 | 1902.798006 | 2027.943486 | 1861.179106 | 3775.428271 | 3957.971793 | 3609.954111 | 4.27E-45 | 0.954867211 | 1.19E-46 | 14.34244742 | 1.938401211 |
| CNAG_07965 | - | hypothetical protein | 3705.719922 | 3588.06973 | 3298.760897 | 6681.544116 | 6542.056773 | 6724.99744 | 5.20E-45 | 0.898107961 | 1.46E-46 | 14.32834279 | 1.863620313 |
| CNAG_06333 | - | replication factor C subunit 3/5, variant | 889.2155047 | 858.9841964 | 832.0761752 | 1885.060599 | 1831.668133 | 1982.256615 | 6.87E-45 | 1.128435869 | 1.93E-46 | 14.30876633 | 2.186215881 |
| CNAG_04678 | YPK1 | AGC/AKT protein kinase | 1538.437788 | 1388.010127 | 1516.70345 | 3106.223476 | 3018.567412 | 2989.305443 | 9.79E-45 | 1.021022826 | 2.76E-46 | 14.28377214 | 2.029357201 |
| CNAG_04702 | - | hypothetical protein | 1221.976845 | 1122.954445 | 1077.36301 | 2425.93721 | 2396.656866 | 2535.145688 | 2.62E-44 | 1.089512767 | 7.42E-46 | 14.21479398 | 2.12802156 |
| CNAG_02048 | PUT5 | Proline dehydrogenase | 2579.757 | 2298.934996 | 2114.593941 | 1021.107693 | 917.2095871 | 1005.095045 | 4.66E-44 | -1.263686508 | 1.33E-45 | -14.17404444 | 0.416478375 |
| CNAG_04798 | - | regulatory protein Cys-3 | 753.8623409 | 833.1250273 | 860.8482078 | 1832.928623 | 1956.943601 | 1778.112884 | 4.66E-44 | 1.171120978 | 1.33E-45 | 14.17372501 | 2.251865996 |
| CNAG_03869 | - | hypothetical protein | 226.1659131 | 218.0140064 | 256.2362174 | 803.6930758 | 710.4100407 | 862.9338437 | 5.81E-44 | 1.747310618 | 1.67E-45 | 14.15802833 | 3.357321316 |
| CNAG_03759 | - | conidiation-specific protein 6 | 9.031092404 | 6.856382731 | 1.37144751 | 274.3570513 | 267.6743983 | 295.2824843 | 8.20E-44 | 5.692219601 | 2.36E-45 | 14.13342699 | 51.70455941 |
| CNAG_00705 | - | RuvB-like helicase 2 | 4967.10269 | 4709.275272 | 5073.961023 | 2668.899556 | 2697.705706 | 2726.856868 | 8.64E-44 | -0.881770782 | 2.50E-45 | -14.12944359 | 0.542700904 |
| CNAG_05904 | - | small subunit ribosomal protein S14 | 38279.6471 | 37092.59669 | 42857.28016 | 20729.08622 | 22028.5792 | 21387.56706 | 1.01E-43 | -0.897839629 | 2.94E-45 | -14.11806924 | 0.536689799 |
| CNAG_04803 | - | protein transporter sec31 | 2144.457569 | 2077.518985 | 1926.239528 | 3946.62452 | 4093.353896 | 3884.133832 | 1.07E-43 | 0.940437917 | 3.11E-45 | 14.11408962 | 1.919110679 |
| CNAG_05918 | - | ATP synthase subunit beta, mitochondrial | 41182.69791 | 43962.21216 | 44771.60209 | 22306.67 | 20826.06002 | 24825.68816 | 1.16E-43 | -0.949614514 | 3.40E-45 | -14.10778518 | 0.517770791 |
| CNAG_12486 | - | unspecified product | 830.5626366 | 777.5815761 | 820.834195 | 291.7262294 | 281.474862 | 284.6623601 | 1.70E-43 | -1.519007445 | 5.00E-45 | -14.08052059 | 0.34892589 |
| CNAG_12223 | - | unspecified product | 1973.927923 | 1801.229456 | 2337.177122 | 818.7142975 | 685.1247882 | 653.281784 | 2.04E-43 | -1.518967507 | 6.04E-45 | -14.06721215 | 0.34893555 |
| CNAG_05022 | - | hypothetical protein | 32.43412052 | 38.1244306 | 13.1524087 | 331.2637547 | 276.829659 | 329.8282574 | 3.11E-43 | 3.481302683 | 9.24E-45 | 14.03710908 | 11.16802896 |
| CNAG_01612 | PSK202 | CAMK/CAMKL protein kinase, variant | 37.08341265 | 38.11827497 | 27.02091922 | 288.9131887 | 278.7205928 | 286.6977212 | 3.87E-43 | 3.047356791 | 1.15E-44 | 14.02134431 | 8.266959342 |
| CNAG_03986 | NAM9 | NAM9 protein | 3707.04356 | 3471.294629 | 4103.383201 | 1721.571769 | 1778.137475 | 1874.291266 | 6.00E-43 | -1.085763688 | 1.79E-44 | -13.98999909 | 0.471142803 |
| CNAG_04347 | - | Aspartate kinase | 3156.029747 | 3470.120535 | 3329.258927 | 1815.92261 | 1748.00351 | 1709.988603 | 7.12E-43 | -0.931916056 | 2.14E-44 | -13.97750016 | 0.524161736 |
| CNAG_12987 | - | unspecified product | 8394.519987 | 8446.855781 | 11001.06476 | 4075.888597 | 3977.62187 | 3827.153328 | 7.85E-43 | -1.244448366 | 2.37E-44 | -13.97024817 | 0.422069251 |
| CNAG_04654 | - | UNC-50 family protein | 821.4803715 | 752.3359465 | 792.6053662 | 1684.950601 | 1772.254847 | 1783.574134 | 7.95E-43 | 1.131875609 | 2.41E-44 | 13.96908426 | 2.191434577 |
| CNAG_04688 | - | acyl-CoA dehydrogenase | 255.3504018 | 278.3171324 | 292.4880294 | 766.6196482 | 873.6754666 | 921.8301573 | 8.61E-43 | 1.618662018 | 2.62E-44 | 13.96309841 | 3.070901031 |
| CNAG_01681 | FCY2 | cytosine permease | 6266.732262 | 5953.673691 | 6436.850937 | 2776.111679 | 3126.355618 | 3339.664117 | 1.09E-42 | -1.028961889 | 3.32E-44 | -13.94617935 | 0.490062653 |
| CNAG_03817 | - | cell growth-regulating nucleolar protein | 2473.566681 | 2048.002434 | 2268.081538 | 1014.892357 | 1000.079596 | 975.1281825 | 1.69E-42 | -1.199660055 | 5.17E-44 | -13.91452252 | 0.435377858 |
| CNAG_05525 | - | small subunit ribosomal protein S26 | 40361.82215 | 40838.8189 | 46208.08761 | 23773.82613 | 24012.46981 | 23559.03481 | 1.82E-42 | -0.851996142 | 5.61E-44 | -13.908739 | 0.554017656 |
| CNAG_03878 | - | hypothetical protein | 1461.414343 | 1434.878162 | 1595.08216 | 680.2526366 | 623.9714748 | 640.9677338 | 1.93E-42 | -1.223817051 | 5.97E-44 | -13.90424853 | 0.428148433 |
| CNAG_04348 | - | chaperone | 2511.199241 | 2426.123151 | 2567.74554 | 1032.69067 | 1207.145886 | 1228.179592 | 2.33E-42 | -1.129460466 | 7.22E-44 | -13.8905974 | 0.457086633 |
| CNAG_06310 | IRK7 | CAMK/CAMKL protein kinase, variant 2 | 146.8807086 | 242.7530613 | 191.2248774 | 762.4914511 | 806.5901976 | 729.9110982 | 3.97E-42 | 1.974445669 | 1.24E-43 | 13.8519676 | 3.929772156 |
| CNAG_04774 | FZC26 | nuclear protein, variant | 1289.761429 | 1436.429804 | 1305.557742 | 2660.137285 | 2760.715365 | 2616.318033 | 4.54E-42 | 0.980861276 | 1.42E-43 | 13.84214849 | 1.973643304 |
| CNAG_07002 | - | hypothetical protein, variant | 110.6408418 | 132.8996657 | 143.2147066 | 546.2586309 | 481.9255641 | 566.499257 | 7.55E-42 | 2.034190572 | 2.37E-43 | 13.80528801 | 4.095928616 |
| CNAG_02359 | - | small subunit ribosomal protein S25e | 61477.05385 | 59450.40336 | 69941.98444 | 35203.01873 | 34487.38514 | 33717.03087 | 1.12E-41 | -0.899883833 | 3.53E-43 | -13.77646993 | 0.535929883 |
| CNAG_04873 | - | hypothetical protein, variant | 94.25654424 | 100.5781957 | 75.00667491 | 458.5390817 | 439.9803687 | 394.6706425 | 1.57E-41 | 2.247854518 | 4.98E-43 | 13.7516771 | 4.749759673 |

| | | | | | | | | | | | | | |
|---|---|---|---|---|---|---|---|---|---|---|---|---|---|
| CNAG_05866 | PRM1 | plasma membrane fusion protein PRM1 | 140.9306578 | 152.2722424 | 127.2409325 | 509.3197503 | 588.4274013 | 516.7671738 | 1.80E-41 | 1.929653228 | 5.72E-43 | 13.74154064 | 3.809636183 |
| CNAG_04876 | - | endo-1,3(4)-beta-glucanase | 1141.489446 | 1105.732586 | 1276.740514 | 2458.574475 | 2471.702689 | 2647.248066 | 2.12E-41 | 1.08961373 | 6.75E-43 | 13.72961981 | 2.128170488 |
| CNAG_00284 | - | efflux protein EncT, variant | 86.19142502 | 118.9280782 | 160.2794389 | 560.7360856 | 608.1421319 | 678.4698756 | 2.40E-41 | 2.327695794 | 7.68E-43 | 13.72020877 | 5.020029333 |
| CNAG_12764 | - | unspecified product | 862.4122226 | 938.0898454 | 1050.027127 | 342.3064139 | 329.7721127 | 352.3640132 | 4.71E-41 | -1.493175399 | 1.51E-42 | -13.67106941 | 0.35522982 |
| CNAG_01752 | - | solute carrier family 25 (mitochondrial 2- oxodicarboxylate transporter), member 21 | 5821.881094 | 5447.322923 | 5722.394375 | 3247.229684 | 3124.668802 | 3233.077884 | 5.13E-41 | -0.838496907 | 1.65E-42 | -13.6645551 | 0.559225903 |
| CNAG_02084 | ERG20 | farnesyl diphosphate synthase | 4766.285636 | 4662.92953 | 4663.430584 | 2702.418649 | 2758.989614 | 2710.190807 | 5.50E-41 | -0.801730233 | 1.78E-42 | -13.65913974 | 0.57366077 |
| CNAG_07752 | GLF | UDP-galactopyranose mutase | 4892.480631 | 5223.137882 | 4935.379701 | 2876.758068 | 2786.547233 | 2943.309079 | 6.84E-41 | -0.821356322 | 2.22E-42 | -13.64297309 | 0.565909664 |
| CNAG_04589 | - | hypothetical protein | 107.1716993 | 171.6543358 | 124.0538965 | 574.9101014 | 642.8378563 | 547.8589798 | 7.02E-41 | 2.118891239 | 2.29E-42 | 13.6408805 | 4.343599961 |
| CNAG_03194 | - | saccharopine dehydrogenase, variant | 508.1686135 | 478.0080884 | 534.9149226 | 136.7094439 | 145.1468321 | 125.986899 | 7.05E-41 | -1.920234028 | 2.31E-42 | -13.6402693 | 0.264211647 |
| CNAG_00797 | - | acetyl-CoA synthetase | 9634.549056 | 9142.883379 | 8846.08118 | 4959.381447 | 5431.214949 | 5240.251919 | 7.17E-41 | -0.837013124 | 2.35E-42 | -13.63879737 | 0.559801352 |
| CNAG_05725 | - | ketol-acid reductoisomerase, mitochondrial | 35210.79862 | 34675.90129 | 34627.58452 | 20548.62235 | 22396.98513 | 20600.43219 | 7.36E-41 | -0.733313218 | 2.43E-42 | -13.63659061 | 0.601520905 |
| CNAG_01920 | UBI4 | Polyubiquitin | 8131.472952 | 8175.745824 | 7372.596753 | 14248.07567 | 13642.61387 | 13722.41276 | 9.69E-41 | 0.79838487 | 3.21E-42 | 13.61622116 | 1.739153015 |
| CNAG_05623 | - | chorismate synthase | 7972.964248 | 7863.98695 | 8372.502821 | 4245.663962 | 4759.283195 | 4558.615621 | 1.01E-40 | -0.851351352 | 3.37E-42 | -13.61265672 | 0.55426532 |
| CNAG_05847 | TRR1 | Thioredoxin reductase | 2141.106407 | 2156.757228 | 2116.702332 | 1040.616091 | 1123.519902 | 1103.829528 | 1.05E-40 | -0.988718166 | 3.50E-42 | -13.60984277 | 0.503925313 |
| CNAG_06723 | - | succinate dehydrogenase (ubiquinone) membrane anchor subunit | 6523.738975 | 6192.908282 | 6880.511401 | 3740.681211 | 3480.887748 | 3485.122146 | 2.24E-40 | -0.887765934 | 7.48E-42 | -13.55419511 | 0.540450378 |
| CNAG_05722 | - | alanine-tRNA ligase | 5101.35073 | 4787.919516 | 5236.039558 | 2450.86119 | 2672.676676 | 2787.90622 | 2.48E-40 | -0.950870531 | 8.31E-42 | -13.54648464 | 0.517320214 |
| CNAG_02833 | - | NADH dehydrogenase (ubiquinone) 1 beta subcomplex 7 | 4248.923024 | 3987.400649 | 4405.275422 | 2195.655679 | 2334.582284 | 2265.045785 | 2.54E-40 | -0.911346323 | 8.55E-42 | -13.54441472 | 0.531688688 |
| CNAG_01674 | - | MFS transporter | 46.41821543 | 37.05046651 | 39.81495113 | 252.2883868 | 344.6032788 | 330.9888955 | 7.69E-40 | 2.908314192 | 2.60E-41 | 13.46248225 | 7.507404369 |
| CNAG_03270 | - | Adenylosuccinate lyase | 7826.861999 | 7203.533679 | 7673.987206 | 3816.787479 | 4249.475342 | 4265.526877 | 8.39E-40 | -0.896356314 | 2.85E-41 | -13.45575367 | 0.537241883 |
| CNAG_07513 | - | hypothetical protein | 2546.148319 | 2574.187763 | 2528.741549 | 4450.848135 | 4419.24939 | 4604.606286 | 9.00E-40 | 0.801790644 | 3.07E-41 | 13.45027309 | 1.743263488 |
| CNAG_02054 | - | hypothetical protein | 2498.192995 | 2174.049393 | 2328.890203 | 1006.270679 | 1127.790217 | 1121.618039 | 9.53E-40 | -1.121057281 | 3.26E-41 | -13.44580283 | 0.459756769 |
| CNAG_02818 | - | glycine cleavage system T protein, variant | 639.3490775 | 649.9572336 | 631.5776848 | 1352.569132 | 1493.715435 | 1436.116526 | 1.10E-39 | 1.14235485 | 3.77E-41 | 13.43510696 | 2.207410355 |
| CNAG_02657 | - | translation initiation factor 3 subunit G | 10449.6554 | 9374.5944 | 10406.23465 | 5636.548127 | 5777.941102 | 5569.988141 | 1.13E-39 | -0.847814134 | 3.89E-41 | -13.43271375 | 0.555625943 |
| CNAG_01140 | - | ATP-dependent RNA helicase DRS1 | 6513.079592 | 5680.073071 | 6330.24073 | 3277.503915 | 3312.699968 | 3323.028038 | 1.31E-39 | -0.918006235 | 4.53E-41 | -13.4214692 | 0.52923991 |
| CNAG_01273 | - | cellular nucleic acid-binding protein | 9640.483608 | 9105.219318 | 9971.118081 | 5254.375431 | 5426.864579 | 5670.967072 | 1.61E-39 | -0.828122338 | 5.60E-41 | -13.40568963 | 0.563261849 |
| CNAG_00788 | NDH1 | NADH dehydrogenase | 6244.663726 | 6098.073163 | 6106.344024 | 3739.177971 | 3575.823052 | 3462.899084 | 2.11E-39 | -0.791010595 | 7.35E-41 | -13.38549248 | 0.577939109 |
| CNAG_05980 | - | large subunit ribosomal protein L7/L12 | 2901.341906 | 2888.360085 | 2912.273732 | 1619.824624 | 1532.367306 | 1436.911586 | 2.22E-39 | -0.938957381 | 7.79E-41 | -13.38121637 | 0.521609705 |
| CNAG_06239 | - | hypothetical protein | 174.7459626 | 95.24654953 | 107.0086865 | 597.4987573 | 627.1600629 | 643.0903706 | 2.22E-39 | 2.294821265 | 7.78E-41 | 13.38127082 | 4.906931952 |
| CNAG_00413 | OFD1 | putative oxidoreductase | 4609.728274 | 4342.936521 | 4473.57912 | 2361.436158 | 2603.736892 | 2432.678605 | 3.00E-39 | -0.875643236 | 1.06E-40 | -13.35852441 | 0.545010815 |
| CNAG_01455 | RPL39alpha | large subunit ribosomal protein L39 | 19667.99849 | 19020.81813 | 22635.15548 | 10462.64554 | 11288.83536 | 9998.480094 | 3.98E-39 | -0.965572213 | 1.40E-40 | -13.33736082 | 0.512075268 |
| CNAG_12260 | - | unspecified product | 79237.31857 | 72380.69239 | 77543.77768 | 47066.60043 | 45353.97625 | 45098.35327 | 4.04E-39 | -0.752465086 | 1.43E-40 | -13.33587311 | 0.593588445 |
| CNAG_06138 | - | NADH dehydrogenase (ubiquinone) Fe-S protein 6 | 2795.175183 | 2903.444828 | 3295.011721 | 1530.236143 | 1421.8978 | 1423.598527 | 4.55E-39 | -1.055098166 | 1.62E-40 | -13.32671809 | 0.481264474 |
| CNAG_02002 | - | hypothetical protein | 3780.660701 | 3570.430589 | 3893.386997 | 1993.323354 | 2113.77992 | 1986.406222 | 5.60E-39 | -0.899714724 | 2.00E-40 | -13.31107266 | 0.535992707 |
| CNAG_03510 | - | large subunit ribosomal protein L36e | 8583.825937 | 9100.864843 | 9980.67161 | 5189.042826 | 5178.44494 | 5077.077716 | 7.01E-39 | -0.856090198 | 2.51E-40 | -13.29394408 | 0.552447702 |
| CNAG_04189 | - | succinate dehydrogenase [ubiquinone] flavoprotein subunit, mitochondrial | 10098.28209 | 9882.029959 | 9738.710127 | 5970.492051 | 6125.590899 | 6222.695144 | 2.08E-38 | -0.713465258 | 7.48E-40 | -13.21202838 | 0.60985355 |
| CNAG_04025 | - | transaldolase | 38.32812409 | 14.4228284 | 41.9752407 | 1073.835213 | 1086.456271 | 1115.672889 | 2.75E-38 | 5.119148157 | 9.92E-40 | 13.19075918 | 34.75498836 |
| CNAG_02288 | - | solute carrier family 25 (mitochondrial citrate transporter), member 1 | 154.9074578 | 155.5027413 | 157.0692737 | 516.3317366 | 514.7631224 | 561.0425916 | 3.13E-38 | 1.756426713 | 1.13E-39 | 13.18089456 | 3.378602708 |
| CNAG_01577 | - | glutamate dehydrogenase (NADP) | 26243.4675 | 23208.82642 | 24396.01972 | 14281.82817 | 14531.22428 | 14235.88358 | 3.15E-38 | -0.794498566 | 1.14E-39 | -13.18020271 | 0.576543527 |
| CNAG_06206 | - | ATP-dependent RNA helicase DBP9 | 2047.760617 | 2099.668825 | 2135.877969 | 1056.215179 | 1121.804084 | 1098.273727 | 3.49E-38 | -0.954718158 | 1.27E-39 | -13.17214229 | 0.515942373 |
| CNAG_00715 | - | hypothetical protein | 2194.970657 | 2335.621669 | 2698.827991 | 1103.621477 | 1132.945157 | 1123.856068 | 4.31E-38 | -1.121030698 | 1.57E-39 | -13.15593861 | 0.45976524 |
| CNAG_04686 | - | hypothetical protein | 46.43175525 | 49.96911668 | 63.25060478 | 287.5591535 | 347.0548527 | 320.0726551 | 8.50E-38 | 2.578627401 | 3.11E-39 | 13.10425523 | 5.973710825 |
| CNAG_06347 | BLP2 | pr4/barwin domain protein | 150.2599478 | 153.3553835 | 122.9839965 | 593.843949 | 476.6137011 | 602.0149308 | 9.62E-38 | 1.960563389 | 3.54E-39 | 13.09459132 | 3.892139419 |
| CNAG_00602 | - | translation initiation factor 3 subunit I | 3124.34013 | 3136.140895 | 2915.522954 | 1544.193092 | 1628.114872 | 1713.264614 | 1.19E-37 | -0.924450243 | 4.40E-39 | -13.07800137 | 0.526881254 |
| CNAG_06123 | - | leucine-tRNA ligase | 4738.201419 | 4637.044671 | 4378.715548 | 2673.650847 | 2535.539724 | 2625.819099 | 1.27E-37 | -0.827310248 | 4.72E-39 | -13.07270447 | 0.563578997 |
| CNAG_04693 | SIN1 | target of rapamycin complex 2 subunit | 1343.524739 | 1552.800921 | 1508.165988 | 2828.143568 | 2900.359339 | 3143.501282 | 1.93E-37 | 0.996008125 | 7.18E-39 | 13.04067262 | 1.994473735 |
| CNAG_04820 | - | hypothetical protein | 1156.767675 | 1235.035786 | 1494.290849 | 2921.760104 | 2836.560008 | 3138.997136 | 2.33E-37 | 1.179961771 | 8.67E-39 | 13.02632578 | 2.265707732 |
| CNAG_01696 | - | chaperone DnaJ | 7534.668586 | 6281.25197 | 6478.523211 | 3265.735355 | 3451.54713 | 3549.505034 | 2.36E-37 | -0.999283938 | 8.82E-39 | -13.02501189 | 0.50024823 |
| CNAG_04880 | - | hypothetical protein | 354.620851 | 477.5813684 | 448.1687624 | 1220.041067 | 1121.911034 | 1117.473259 | 2.45E-37 | 1.420705884 | 9.17E-39 | 13.0220605 | 2.677164676 |
| CNAG_03980 | - | hypothetical protein, variant | 5.479496873 | 13.32757174 | 16.33999449 | 194.5358031 | 206.3738489 | 227.6208531 | 2.59E-37 | 4.190410619 | 9.75E-39 | 13.01736996 | 18.25741513 |
| CNAG_03476 | - | Spermidine synthase | 10633.17538 | 10566.21261 | 11491.88225 | 6249.070635 | 6687.12542 | 6410.349928 | 2.97E-37 | -0.772292941 | 1.12E-38 | -13.00670121 | 0.585486195 |
| CNAG_01153 | - | small subunit ribosomal protein S13e | 6244.583204 | 5823.34 | 6325.96896 | 3558.591718 | 3079.756446 | 3209.809737 | 5.19E-37 | -0.916997891 | 1.96E-38 | -12.96371914 | 0.529609942 |
| CNAG_00656 | - | large subunit ribosomal protein L7e | 62985.67527 | 55720.37072 | 70024.07336 | 33524.99248 | 32551.77146 | 32168.37022 | 7.76E-37 | -0.957953416 | 2.95E-38 | -12.93256479 | 0.514786664 |
| CNAG_01292 | - | pol II transcription elongation factor | 418.6816903 | 414.0272679 | 435.3493016 | 999.4306412 | 1042.610868 | 978.7575592 | 8.46E-37 | 1.237265538 | 3.22E-38 | 12.92567816 | 2.357512691 |
| CNAG_03433 | - | endo alpha-1,4 polygalactosaminidase precursor | 2852.393845 | 2922.852591 | 2986.92571 | 1674.315468 | 1577.22302 | 1622.24785 | 9.82E-37 | -0.861251791 | 3.75E-38 | -12.91402536 | 0.550474717 |
| CNAG_06113 | - | hypothetical protein | 76436.04549 | 77298.06621 | 84592.98826 | 45471.7784 | 49571.08566 | 47313.11694 | 1.02E-36 | -0.758886694 | 3.92E-38 | -12.91055927 | 0.590952184 |
| CNAG_04760 | - | cytoplasmic protein | 1366.893675 | 1285.686197 | 1637.187936 | 3261.222829 | 3073.78081 | 3114.709436 | 1.03E-36 | 1.12398844 | 3.98E-38 | 12.90954537 | 2.179486761 |
| CNAG_07637 | - | large subunit ribosomal protein L3 | 3254.022854 | 3097.437659 | 3299.394777 | 1794.91032 | 1829.996142 | 1761.028798 | 1.18E-36 | -0.857415034 | 4.56E-38 | -12.89896054 | 0.551940619 |
| CNAG_01470 | - | NADH dehydrogenase (ubiquinone) flavoprotein 2 | 4453.416304 | 4462.542775 | 4714.584371 | 2696.230402 | 2668.416493 | 2640.24885 | 1.75E-36 | -0.783208547 | 6.76E-38 | -12.8686051 | 0.581073051 |
| CNAG_03010 | - | enoyl-CoA hydratase/isomerase | 160.7436547 | 177.0275397 | 156.0100533 | 520.9327145 | 530.2917498 | 575.4805789 | 2.12E-36 | 1.706018445 | 8.24E-38 | 12.85335841 | 3.262591685 |
| CNAG_02018 | - | hypothetical protein | 94.24046178 | 96.26765594 | 94.17383408 | 406.069667 | 363.7368314 | 445.429121 | 2.59E-36 | 2.080936223 | 1.01E-37 | 12.83774854 | 4.230816818 |
| CNAG_02076 | - | leukotriene A-4 hydrolase/aminopeptidase | 7023.570458 | 6941.697562 | 7056.535964 | 3716.425233 | 4201.212966 | 4167.808427 | 3.50E-36 | -0.814046843 | 1.37E-37 | -12.81408825 | 0.568784146 |
| CNAG_01639 | - | elongation factor G, mitochondrial | 7657.718217 | 7589.25822 | 7908.630785 | 4409.958969 | 4805.889025 | 4678.480327 | 3.66E-36 | -0.752420161 | 1.44E-37 | -12.81030361 | 0.59360693 |
| CNAG_00777 | - | CCR4-NOT transcriptional complex subunit CAF120 | 1958.760144 | 1750.00452 | 1807.844738 | 3368.219134 | 3714.723562 | 3527.745378 | 3.74E-36 | 0.928053934 | 1.47E-37 | 12.80839571 | 1.902707682 |
| CNAG_05818 | CHS5 | Chitin synthase | 3555.071128 | 3484.626456 | 2991.638889 | 6170.893489 | 6379.079404 | 6482.961979 | 3.94E-36 | 0.908895629 | 1.55E-37 | 12.80411432 | 1.877607655 |
| CNAG_03944 | - | chaperone regulator | 11186.69171 | 11129.70939 | 11708.40298 | 6629.66507 | 7184.842567 | 6954.306912 | 4.33E-36 | -0.727638143 | 1.71E-37 | -12.79658452 | 0.603891745 |
| CNAG_05111 | - | hypothetical protein | 3778.460041 | 3937.794582 | 3865.720524 | 2208.968525 | 2287.17233 | 2312.729354 | 4.52E-36 | -0.78161233 | 1.79E-37 | -12.79308839 | 0.581716314 |
| CNAG_06226 | - | NADH dehydrogenase (ubiquinone) 1 alpha subcomplex 5 | 4204.468764 | 3724.531547 | 4309.273263 | 2049.305577 | 2225.033287 | 2095.21321 | 5.18E-36 | -0.958127647 | 2.06E-37 | -12.78215139 | 0.514724498 |
| CNAG_01540 | - | dehydrogenase | 722.2138157 | 694.1331925 | 701.9514405 | 1464.67968 | 1461.617875 | 1595.915938 | 6.45E-36 | 1.079042558 | 2.58E-37 | 12.76484806 | 2.112633571 |
| CNAG_01496 | PPT1 | protein phosphatase 5 | 3245.781615 | 3013.394608 | 3239.661546 | 1722.515342 | 1768.751588 | 1590.108909 | 6.63E-36 | -0.918402131 | 2.65E-37 | -12.76253097 | 0.529094699 |
| CNAG_05064 | - | hypothetical protein | 4.301300246 | 2.515281294 | 6.730717124 | 225.2218604 | 245.4885605 | 225.5587371 | 7.18E-36 | 5.840300096 | 2.88E-37 | 12.75605751 | 57.29352096 |
| CNAG_04582 | - | hypothetical protein | 60.45184182 | 96.24743695 | 53.68556232 | 363.9902379 | 382.1767857 | 402.1478035 | 8.15E-36 | 2.442825786 | 3.28E-37 | 12.74592478 | 5.437056373 |
| CNAG_12985 | - | unspecified product | 1107.66893 | 691.5487072 | 879.4511273 | 247.996289 | 239.1241104 | 238.1094115 | 8.25E-36 | -1.903893763 | 3.34E-37 | -12.74470905 | 0.267221175 |
| CNAG_01607 | TRX2 | thioredoxin | 1635.571626 | 1761.313982 | 1794.575298 | 898.2892668 | 838.8246091 | 795.2636955 | 8.46E-36 | -1.051057537 | 3.43E-37 | -12.74254989 | 0.482614265 |
| CNAG_02830 | ERG4 | delta24(24(1))-sterol reductase | 3175.903791 | 3290.265867 | 3495.594807 | 1774.620385 | 1890.378223 | 1855.376995 | 9.78E-36 | -0.866979875 | 3.98E-37 | -12.73098186 | 0.548293443 |
| CNAG_00845 | - | hypothetical protein | 2295.224171 | 2106.205026 | 2515.442031 | 1132.484502 | 1138.146771 | 1083.893377 | 1.22E-35 | -1.060335845 | 4.96E-37 | -12.71377221 | 0.479520419 |
| CNAG_02315 | - | ubiquinol-cytochrome c reductase, iron-sulfur subunit | 8852.315951 | 8679.624835 | 8951.636024 | 5618.017932 | 5201.748706 | 5284.641229 | 1.23E-35 | -0.733003564 | 5.04E-37 | -12.71256673 | 0.601650026 |
| CNAG_06799 | HEM12 | uroporphyrinogen decarboxylase | 3710.514509 | 3303.252515 | 3828.298125 | 1784.669192 | 1894.641412 | 1929.77241 | 1.45E-35 | -0.967082536 | 5.96E-37 | -12.69937369 | 0.511539469 |
| CNAG_07400 | - | aspartate-tRNA(Asn) ligase | 12496.66957 | 10819.44371 | 12288.51529 | 6248.991456 | 6829.412624 | 6522.505394 | 2.08E-35 | -0.877327451 | 8.57E-37 | -12.67094142 | 0.544374936 |
| CNAG_01437 | - | ribosomal RNA assembly protein | 3422.115377 | 3130.850813 | 3471.067702 | 1766.060335 | 1736.776476 | 1867.580251 | 2.20E-35 | -0.916216136 | 9.06E-37 | -12.66655297 | 0.529897 |
| CNAG_08028 | - | hypothetical protein | 308.9620203 | 309.5346102 | 329.7838471 | 861.1631918 | 769.8827666 | 834.3259421 | 3.40E-35 | 1.36595144 | 1.41E-36 | 12.63203024 | 2.577462504 |
| CNAG_01323 | - | ubiquinol-cytochrome c reductase subunit 7 | 12433.865 | 12296.58103 | 13716.49126 | 7778.490974 | 7600.544182 | 7473.891659 | 4.15E-35 | -0.766000112 | 1.72E-36 | -12.61605181 | 0.58804558 |
| CNAG_03641 | - | translation initiation factor 3 subunit D | 8791.574189 | 8367.190594 | 8963.352076 | 5108.895709 | 5432.969421 | 5280.19 | 5.93E-35 | -0.739064394 | 2.47E-36 | -12.58765886 | 0.599127768 |
| CNAG_01721 | HEM3 | porphobilinogen deaminase | 3502.511794 | 3014.478388 | 3130.921226 | 1682.782692 | 1658.284695 | 1688.879857 | 6.23E-35 | -0.95580906 | 2.60E-36 | -12.58354477 | 0.515552388 |
| CNAG_07363 | - | isocitrate dehydrogenase, NAD-dependent | 12529.40884 | 11058.63176 | 12109.3836 | 6668.549702 | 7095.098208 | 6849.982724 | 6.57E-35 | -0.80823551 | 2.75E-36 | -12.57910632 | 0.571079891 |
| CNAG_03713 | - | efflux protein EncT, variant | 79.08521441 | 95.17709805 | 95.22593483 | 374.1266476 | 368.2349956 | 378.9961667 | 6.94E-35 | 2.043042608 | 2.91E-36 | 12.57457869 | 4.121137526 |
| CNAG_02565 | - | homoaconitase, mitochondrial | 1918.098608 | 1946.665004 | 1885.300034 | 1013.463303 | 1055.406304 | 973.9582029 | 1.05E-34 | -0.934019482 | 4.43E-36 | -12.54141058 | 0.523398073 |
| CNAG_04855 | - | hypothetical protein | 325.2907615 | 306.3121983 | 303.1511952 | 904.8512301 | 752.5471724 | 896.4049153 | 1.48E-34 | 1.435010778 | 6.24E-36 | 12.51422004 | 2.703841865 |

| | | | | | | | | | | | | | |
|---|---|---|---|---|---|---|---|---|---|---|---|---|---|
| CNAG_03910 | ITR6 | myo-inositol transporter, putative | 678.9870503 | 641.3371985 | 559.0905933 | 1390.839523 | 1464.376984 | 1378.430955 | 1.59E-34 | 1.156764617 | 6.75E-36 | 12.50805444 | 2.229568643 |
| CNAG_01132 | - | nucleolar protein 6 | 1537.370124 | 1558.771009 | 1558.919832 | 733.9578111 | 713.7174623 | 817.3832459 | 2.20E-34 | -1.054965546 | 9.35E-36 | -12.4821007 | 0.481308716 |
| CNAG_07362 | - | nucleolin | 10243.04514 | 9532.97503 | 10349.73156 | 5627.186149 | 6066.930079 | 6097.260572 | 2.28E-34 | -0.775524165 | 9.73E-36 | -12.47891447 | 0.584176341 |
| CNAG_04707 | - | hypothetical protein | 81.45930087 | 129.62442 | 99.50993383 | 424.655601 | 436.4406762 | 438.9363913 | 2.38E-34 | 2.056782729 | 1.02E-35 | 12.47537514 | 4.160574433 |
| CNAG_02382 | - | ribosome biogenesis protein BRX1 | 3922.96617 | 3403.448039 | 3809.130879 | 1764.392075 | 1977.462559 | 1975.289599 | 2.44E-34 | -0.977890508 | 1.05E-35 | -12.47321286 | 0.507721582 |
| CNAG_00485 | - | hypothetical protein | 377.8034823 | 283.7315134 | 240.2939871 | 1031.84803 | 1011.736307 | 898.7884791 | 2.93E-34 | 1.691581569 | 1.26E-35 | 12.45836512 | 3.230106132 |
| CNAG_02656 | - | translation machinery-associated protein 20 | 2582.378291 | 2383.052501 | 2524.044783 | 1155.771079 | 1343.512195 | 1057.301349 | 5.19E-34 | -1.090908911 | 2.24E-35 | -12.41248238 | 0.469465514 |
| CNAG_04868 | - | cytoplasmic protein, variant | 282.1578648 | 272.9361251 | 343.6343072 | 808.8090756 | 801.0211928 | 894.1432551 | 5.58E-34 | 1.465056922 | 2.41E-35 | 12.406412 | 2.76074363 |
| CNAG_00116 | - | small subunit ribosomal protein S3 | 26656.03642 | 29786.47437 | 30790.41989 | 17303.95526 | 17947.86336 | 16628.29369 | 5.59E-34 | -0.764709204 | 2.42E-35 | -12.40607213 | 0.588571992 |
| CNAG_12766 | - | unspecified product | 44.05547146 | 29.50493565 | 41.93290538 | 300.1519809 | 269.2470413 | 231.5072725 | 5.80E-34 | 2.794686946 | 2.52E-35 | 12.40291235 | 6.938803663 |
| CNAG_03844 | - | nonselective cation channel protein | 196.9163461 | 224.4085359 | 199.7149046 | 575.2864761 | 603.594013 | 636.6061123 | 9.29E-34 | 1.5364522 | 4.05E-35 | 12.36481743 | 2.900802754 |
| CNAG_12216 | - | unspecified product | 900.507572 | 753.9557889 | 1045.606845 | 303.3998313 | 185.571523 | 228.1308085 | 9.90E-34 | -1.929808425 | 4.33E-35 | -12.35948633 | 0.262464021 |
| CNAG_01435 | - | serine-tRNA ligase | 14112.99203 | 13388.01649 | 13614.19315 | 8055.494271 | 8812.505713 | 8517.406845 | 1.08E-33 | -0.711288617 | 4.73E-35 | -12.35231048 | 0.610774351 |
| CNAG_03762 | - | AdoMet-dependent rRNA methyltransferase spb1 | 5994.700041 | 5545.377176 | 5795.991469 | 3128.065052 | 3340.344372 | 3486.158398 | 1.14E-33 | -0.815846786 | 5.01E-35 | -12.34775727 | 0.568074959 |
| CNAG_04501 | TRP1 | anthranilate synthase/indole-3-glycerol phosphate synthase/phosphoribosylanthranilate isomerase | 4055.249632 | 4379.519593 | 4090.767591 | 2523.465757 | 2508.85652 | 2479.265487 | 1.21E-33 | -0.752684687 | 5.36E-35 | -12.34236836 | 0.593498098 |
| CNAG_03589 | - | adrenodoxin-type ferredoxin | 1911.178974 | 1970.392721 | 2049.487905 | 1032.90029 | 1091.633049 | 1046.095693 | 1.26E-33 | -0.919254805 | 5.59E-35 | -12.33893558 | 0.528782082 |
| CNAG_07326 | - | hypothetical protein | 3286.737913 | 3125.459706 | 3625.65403 | 1821.367757 | 1792.010297 | 1655.613473 | 1.53E-33 | -0.945579747 | 6.78E-35 | -12.32334003 | 0.519220861 |
| CNAG_06344 | - | hypothetical protein | 1358.675342 | 1302.390181 | 1405.319394 | 681.1094173 | 646.4648335 | 645.3694913 | 1.81E-33 | -1.059063795 | 8.05E-35 | -12.3095528 | 0.479943407 |
| CNAG_03346 | BZP4 | bZip transcription factor, putative | 60.41444927 | 79.01440694 | 61.12841866 | 338.5533277 | 296.5167315 | 344.4588107 | 1.99E-33 | 2.275658922 | 8.86E-35 | 12.30174135 | 4.842187425 |
| CNAG_04809 | - | COP9 signalosome complex subunit 5 | 1496.366809 | 1534.471708 | 1530.540115 | 2850.859073 | 2702.772635 | 2656.345298 | 2.78E-33 | 0.832848136 | 1.24E-34 | 12.27444842 | 1.781198295 |
| CNAG_04072 | - | 60S ribosome subunit biogenesis protein Nip7 | 3717.697677 | 3666.320451 | 4122.64252 | 1974.600532 | 2199.186483 | 2095.183368 | 3.33E-33 | -0.891745997 | 1.49E-34 | -12.25963989 | 0.538961455 |
| CNAG_05899 | - | Pyrroline-5-carboxylate reductase | 2496.298719 | 2759.034963 | 2841.83797 | 1309.111445 | 1502.225484 | 1408.01278 | 3.43E-33 | -0.955351299 | 1.54E-34 | -12.25686575 | 0.515715997 |
| CNAG_04898 | - | MFS transporter | 74.401082 | 87.63192771 | 74.98179456 | 313.8719433 | 366.6982262 | 358.9584891 | 3.44E-33 | 2.12687984 | 1.55E-34 | 12.25644905 | 4.367718388 |
| CNAG_02789 | - | nitrogen permease regulator 2 | 1019.90502 | 960.2309573 | 936.5600646 | 1874.818723 | 1919.675755 | 1851.422498 | 3.93E-33 | 0.9377637 | 1.78E-34 | 12.24543025 | 1.915556661 |
| CNAG_04681 | LPI14 | transmembrane protein, putative | 98.88549798 | 101.6411877 | 78.19542904 | 378.6457532 | 361.1762836 | 450.910833 | 4.01E-33 | 2.087139536 | 1.82E-34 | 12.2435719 | 4.249047689 |
| CNAG_02923 | - | large subunit ribosomal protein L14 | 3244.755749 | 3270.879154 | 3256.775935 | 1896.901621 | 1850.681244 | 1975.22996 | 5.39E-33 | -0.787218379 | 2.45E-34 | -12.21942465 | 0.579460257 |
| CNAG_02502 | - | ATP-dependent RNA helicase DBP3 | 3417.340447 | 2899.235355 | 3409.188319 | 1658.631767 | 1684.170474 | 1671.126764 | 5.70E-33 | -0.972125273 | 2.60E-34 | -12.21457812 | 0.509754576 |
| CNAG_02338 | - | cellular nucleic acid-binding protein | 14149.21478 | 13214.57459 | 13858.40634 | 8721.810301 | 8788.339013 | 8600.670991 | 5.78E-33 | -0.674460459 | 2.64E-34 | -12.21329778 | 0.6265665 |
| CNAG_04581 | - | hypothetical protein | 618.3166169 | 670.3953073 | 577.2013528 | 1365.204543 | 1410.026233 | 1269.710343 | 9.93E-33 | 1.101657991 | 4.55E-34 | 12.16897712 | 2.146011773 |
| CNAG_00754 | - | ATP-binding cassette, sub-family E, member 1 | 10596.67288 | 9175.257148 | 9541.389394 | 5155.491435 | 5792.6381 | 5372.380839 | 1.08E-32 | -0.860994962 | 4.98E-34 | -12.16158424 | 0.550572722 |
| CNAG_02212 | - | translation initiation factor 3 subunit A | 14737.71193 | 13377.27326 | 15417.48037 | 7974.495063 | 8449.29791 | 8665.068983 | 1.40E-32 | -0.810871813 | 6.46E-34 | -12.14028544 | 0.570037283 |
| CNAG_04639 | - | hypothetical protein | 1493.993118 | 1446.13224 | 1429.239251 | 2604.757888 | 2758.951968 | 2559.759723 | 1.46E-32 | 0.843257336 | 6.74E-34 | 12.13684041 | 1.794096307 |
| CNAG_06835 | KRE61 | glucosidase | 235.5253853 | 300.9144395 | 315.9235719 | 760.437483 | 910.0536718 | 791.0706783 | 1.74E-32 | 1.518873834 | 8.05E-34 | 12.12227775 | 2.865672683 |
| CNAG_06367 | - | U3 small nucleolar RNA-associated protein 3 | 3282.088625 | 3150.240814 | 3449.74689 | 1792.538599 | 1822.207127 | 1934.166506 | 2.13E-32 | -0.848143552 | 9.87E-34 | -12.10558089 | 0.555499088 |
| CNAG_04857 | - | hypothetical protein | 250.5749608 | 214.7624175 | 199.7437299 | 827.5522273 | 653.5065793 | 645.695095 | 2.27E-32 | 1.664821928 | 1.06E-33 | 12.09992093 | 3.170745157 |
| CNAG_05507 | - | small subunit ribosomal protein S8 | 2218.052846 | 1932.749768 | 2196.635686 | 1053.881098 | 1024.266155 | 1107.14794 | 2.75E-32 | -1.010830119 | 1.28E-33 | -12.08403165 | 0.49626062 |
| CNAG_02905 | - | hypothetical protein | 2253.26058 | 2180.526309 | 2262.811598 | 1271.154647 | 1248.618602 | 1193.749137 | 3.15E-32 | -0.866341968 | 1.47E-33 | -12.07273891 | 0.548535931 |
| CNAG_04714 | - | MFS multidrug transporter protein | 610.1728487 | 710.2449513 | 629.4376912 | 1379.950581 | 1394.428105 | 1327.406824 | 3.73E-32 | 1.058209118 | 1.75E-33 | 12.05845646 | 2.082345008 |
| CNAG_04725 | - | hypothetical protein, variant | 1976.308034 | 2053.785144 | 1973.126161 | 3328.381269 | 3729.316242 | 3708.68436 | 3.89E-32 | 0.82772453 | 1.83E-33 | 12.05474375 | 1.774883744 |
| CNAG_00462 | - | electron-transferring-flavoprotein dehydrogenase | 4809.641331 | 5089.581257 | 4947.122497 | 2974.069294 | 3136.839786 | 3078.727106 | 4.05E-32 | -0.707191495 | 1.91E-33 | -12.0512241 | 0.612511359 |
| CNAG_00336 | - | hypothetical protein | 405.9329499 | 544.3330143 | 416.1973069 | 1123.989069 | 1144.361501 | 1146.308771 | 5.19E-32 | 1.30818666 | 2.45E-33 | 12.03068505 | 2.476300952 |
| CNAG_04588 | ERT1 | hypothetical protein, variant | 1898.039509 | 1898.639946 | 1970.964726 | 3243.667381 | 3354.891755 | 3299.109714 | 6.09E-32 | 0.763956721 | 2.89E-33 | 12.01717639 | 1.698141549 |
| CNAG_00613 | FCY1 | Cytosine deaminase | 1225.560284 | 1173.103525 | 1305.046961 | 568.9425482 | 587.8151133 | 604.2726566 | 6.36E-32 | -1.08944461 | 3.02E-33 | -12.01343292 | 0.469942252 |
| CNAG_00747 | - | succinyl-CoA synthetase beta subunit | 9316.000922 | 9365.964409 | 9826.126126 | 5946.402389 | 6245.534183 | 5964.037905 | 9.16E-32 | -0.666326999 | 4.36E-33 | -11.98296083 | 0.63010886 |
| CNAG_00084 | - | glutamine-tRNA ligase | 4205.763121 | 3969.098754 | 4238.954706 | 2401.23172 | 2306.098905 | 2509.21577 | 9.99E-32 | -0.79790316 | 4.77E-33 | -11.97560458 | 0.575184554 |
| CNAG_01364 | - | Guanylate kinase | 4111.263524 | 4026.204937 | 4366.91687 | 2458.835359 | 2498.505601 | 2497.020012 | 1.03E-31 | -0.761821495 | 4.92E-33 | -11.97305227 | 0.589751262 |
| CNAG_02842 | - | hypothetical protein | 3845.880747 | 3301.097158 | 3531.900599 | 1962.242915 | 1841.140786 | 1949.747598 | 1.06E-31 | -0.908214203 | 5.07E-33 | -11.97057625 | 0.532844249 |
| CNAG_06312 | - | hypothetical protein | 7.828656756 | 0.320185965 | 5.655322266 | 213.2091251 | 195.2491229 | 179.0858673 | 1.23E-31 | 5.59603207 | 5.91E-33 | 11.95782926 | 48.36971291 |
| CNAG_05115 | - | sarcosine oxidase | 3875.163383 | 3561.802704 | 3482.893965 | 2124.957426 | 2039.578056 | 2068.52381 | 1.28E-31 | -0.824697421 | 6.18E-33 | -11.95405323 | 0.564600605 |
| CNAG_00773 | - | hypothetical protein | 492.2413564 | 535.736899 | 517.4588196 | 1170.694114 | 1057.873364 | 1196.258754 | 1.47E-31 | 1.133833999 | 7.09E-33 | 11.94264001 | 2.194411365 |
| CNAG_05259 | - | hypothetical protein, variant | 34.73459662 | 44.55993376 | 44.05629008 | 238.7097652 | 275.4727373 | 232.5250391 | 1.54E-31 | 2.595366733 | 7.49E-33 | 11.93815674 | 6.043426388 |
| CNAG_04611 | UBC8 | ubiquitin-conjugating enzyme E2 H | 265.8468446 | 342.9028937 | 306.3437086 | 938.5216603 | 805.3269701 | 773.4012224 | 1.59E-31 | 1.448121487 | 7.73E-33 | 11.93546162 | 2.728525425 |
| CNAG_02781 | - | dihydrodipicolinate synthetase, variant | 40.54702782 | 42.40715905 | 40.86056037 | 253.0082951 | 222.3839051 | 255.6286976 | 1.72E-31 | 2.56614068 | 8.39E-33 | 11.9287204 | 5.922230652 |
| CNAG_01120 | - | pyruvate dehydrogenase complex dihydrolipoamide acetyltransferase | 21632.16137 | 19931.29733 | 21967.68181 | 12457.50781 | 13226.204 | 13326.61334 | 2.02E-31 | -0.719368216 | 9.88E-33 | -11.91507184 | 0.60736336 |
| CNAG_13077 | - | unspecified product | 23.0678242 | 44.54647224 | 28.07507335 | 237.4523622 | 239.9638966 | 223.5649451 | 2.18E-31 | 2.865524349 | 1.07E-32 | 11.90853916 | 7.288007023 |
| CNAG_02825 | - | Argininosuccinate lyase | 8323.490748 | 9071.782671 | 9459.233002 | 5456.061832 | 5519.209649 | 5494.445582 | 2.39E-31 | -0.720314052 | 1.18E-32 | -11.90059603 | 0.606965301 |
| CNAG_02851 | - | Threonine aldolase | 2139.984526 | 2180.469608 | 2035.703905 | 1212.013238 | 1172.705589 | 1127.141784 | 2.87E-31 | -0.870904076 | 1.41E-32 | -11.88517189 | 0.546804084 |
| CNAG_05091 | - | hypothetical protein | 2433.255984 | 2619.00605 | 2861.008943 | 1418.973719 | 1327.039402 | 1435.741945 | 3.04E-31 | -0.935700223 | 1.50E-32 | -11.88006384 | 0.522788669 |
| CNAG_06900 | - | 2,3-bisphosphoglycerate-independent phosphoglycerate mutase | 23653.67039 | 23523.44091 | 23944.94157 | 15984.87285 | 15741.45667 | 16151.93835 | 3.04E-31 | -0.586213572 | 1.50E-32 | -11.88008406 | 0.6660888 |
| CNAG_05402 | - | protein CMS1 | 2309.383005 | 2288.276987 | 2517.644976 | 1258.612917 | 1292.591258 | 1339.140599 | 3.04E-31 | -0.886631744 | 1.51E-32 | -11.87976294 | 0.540875425 |
| CNAG_00471 | - | mitochondrial protein | 1268.869743 | 1316.362679 | 1423.414675 | 647.6017149 | 587.7604232 | 669.7580442 | 4.16E-31 | -1.089289445 | 2.07E-32 | -11.85339009 | 0.469992798 |
| CNAG_00441 | IMD1 | inosine-5'-monophosphate dehydrogenase | 13607.52079 | 14148.66804 | 14264.72344 | 8648.597116 | 9323.158844 | 9185.695628 | 4.44E-31 | -0.644930335 | 2.21E-32 | -11.84772815 | 0.639523671 |
| CNAG_04832 | - | RalA-binding protein 1 | 440.8097722 | 440.9317579 | 385.2552413 | 1012.000483 | 989.9313188 | 925.5228876 | 4.86E-31 | 1.194625969 | 2.43E-32 | 11.83994767 | 2.288854839 |
| CNAG_01046 | - | N,N-dimethylglycine oxidase | 1835.039102 | 1665.508792 | 1694.405881 | 916.9639363 | 905.2820017 | 891.7947584 | 5.26E-31 | -0.953128913 | 2.64E-32 | -11.83301451 | 0.516511039 |
| CNAG_05202 | - | SET domain-containing protein 6 | 4509.50216 | 4584.293815 | 4648.502601 | 2737.460884 | 2882.377148 | 2889.989013 | 5.71E-31 | -0.706816131 | 2.87E-32 | -11.82594035 | 0.612670745 |
| CNAG_02301 | - | lupus La protein | 11313.85192 | 10401.38828 | 11990.94835 | 5692.46216 | 6422.27496 | 6610.161956 | 6.83E-31 | -0.863825637 | 3.43E-32 | -11.8107658 | 0.549493516 |
| CNAG_05159 | - | hypothetical protein, variant | 105.9183783 | 139.311084 | 111.229922 | 466.9403384 | 390.4205717 | 444.5166952 | 1.04E-30 | 1.85854599 | 5.23E-32 | 11.77535108 | 3.626419918 |
| CNAG_04861 | - | transglycosylase SLT domain-containing protein | 320.6751408 | 388.1410736 | 341.5309229 | 990.5517772 | 875.194064 | 817.8500764 | 1.07E-30 | 1.339766393 | 5.41E-32 | 11.77243703 | 2.531103308 |
| CNAG_09003 | - | small ribosomal protein subunit 3 | 29239.81228 | 24380.01121 | 26060.7531 | 15529.4537 | 14398.32521 | 15222.81578 | 1.10E-30 | -0.835507616 | 5.59E-32 | -11.76978005 | 0.560385831 |
| CNAG_05839 | - | cytochrome c oxidase subunit 6b, variant | 7245.481292 | 6848.022288 | 7470.311574 | 4459.101495 | 4498.81595 | 4211.131906 | 1.26E-30 | -0.727198924 | 6.41E-32 | -11.75815332 | 0.604075624 |
| CNAG_00007 | - | methionine-tRNA ligase | 3248.140947 | 3236.359115 | 3098.940449 | 1620.466855 | 1624.625824 | 1908.608125 | 1.32E-30 | -0.910203763 | 6.72E-32 | -11.75416325 | 0.532109932 |
| CNAG_07547 | GPM1 | hypothetical protein | 732.9697258 | 973.4885469 | 897.6087014 | 350.1905021 | 322.0418924 | 286.912107 | 1.38E-30 | -1.455707662 | 7.05E-32 | -11.75018193 | 0.364576212 |
| CNAG_01111 | - | translation initiation factor 3 subunit K | 5123.607311 | 4761.022998 | 5003.626853 | 2897.772097 | 3119.591731 | 2921.116459 | 1.88E-30 | -0.75190501 | 9.62E-32 | -11.72388818 | 0.59381893 |
| CNAG_00568 | - | hypothetical protein | 54.54535469 | 26.27958953 | 55.78302472 | 269.0295905 | 284.7181692 | 305.5166459 | 2.47E-30 | 2.651033126 | 1.27E-31 | 11.70062676 | 6.28116917 |
| CNAG_04841 | FZC43 | transcriptional regulatory protein, variant | 738.5133583 | 740.4230402 | 699.8090883 | 1394.574002 | 1432.309227 | 1439.504392 | 2.79E-30 | 0.955036459 | 1.43E-31 | 11.690026 | 1.938628625 |
| CNAG_05400 | - | minor histocompatibility antigen H13 | 2155.285042 | 2198.84127 | 2391.80626 | 1231.408264 | 1265.875562 | 1240.355363 | 2.96E-30 | -0.867424481 | 1.52E-31 | -11.68480881 | 0.548124497 |
| CNAG_07494 | - | U3 small nucleolar RNA-associated protein MPP10 | 3438.590137 | 3344.175535 | 3518.034685 | 1948.232311 | 1963.682381 | 2118.427546 | 3.16E-30 | -0.787983734 | 1.63E-31 | -11.67910617 | 0.579152933 |
| CNAG_01923 | - | ATPase GET3 | 3131.424844 | 2980.004682 | 3105.327094 | 1869.731451 | 1766.163272 | 1716.618246 | 3.25E-30 | -0.799830471 | 1.69E-31 | -11.67627431 | 0.574416673 |
| CNAG_01542 | - | taurine catabolism dioxygenase TauD | 54.53356086 | 51.02545016 | 56.84252904 | 281.7855885 | 253.4738328 | 264.6590789 | 3.57E-30 | 2.294037166 | 1.85E-31 | 11.66813039 | 4.904265779 |
| CNAG_07625 | - | plasma membrane protein | 2829.198603 | 3352.670188 | 3173.561026 | 1762.239331 | 1746.313785 | 1718.838325 | 3.70E-30 | -0.854418707 | 1.93E-31 | -11.66487798 | 0.553088133 |
| CNAG_07520 | - | endopeptidase, variant 1 | 67.42670464 | 91.93741545 | 97.34276011 | 355.4898564 | 344.070684 | 347.9548559 | 4.28E-30 | 2.021727219 | 2.23E-31 | 11.65240304 | 4.060696545 |
| CNAG_03320 | - | Mannose-1-phosphate guanylyltransferase | 218.9837612 | 158.7510942 | 156.0214374 | 589.6559953 | 569.998898 | 557.8823724 | 4.88E-30 | 1.672534244 | 2.55E-31 | 11.64105015 | 3.187740618 |
| CNAG_05917 | - | hypothetical protein | 2043.209909 | 2178.338007 | 2445.065652 | 1202.633974 | 1113.993681 | 1089.429864 | 4.94E-30 | -0.984754348 | 2.59E-31 | -11.63979797 | 0.505311757 |
| CNAG_06663 | - | ubiquinol-cytochrome c reductase subunit 6 | 5275.509428 | 5503.298479 | 5257.461678 | 3139.054598 | 3434.451368 | 2972.22084 | 5.11E-30 | -0.763649322 | 2.68E-31 | -11.63664501 | 0.589004547 |
| CNAG_00716 | - | Cytochrome c | 9946.478763 | 9488.794444 | 9836.803742 | 6420.477837 | 5840.914078 | 5875.247915 | 5.51E-30 | -0.706090786 | 2.90E-31 | -11.63003311 | 0.612978855 |
| CNAG_06927 | - | peptidyl-prolyl isomerase G (cyclophilin G) | 2285.767843 | 1971.541752 | 2149.75612 | 1087.355825 | 1135.59725 | 1138.230404 | 6.06E-30 | -0.946662233 | 3.19E-31 | -11.62177513 | 0.518831424 |
| CNAG_03693 | - | large subunit ribosomal protein L11 | 2671.332434 | 2645.960424 | 2606.242229 | 1476.583313 | 1436.624907 | 1598.878386 | 6.06E-30 | -0.827869683 | 3.20E-31 | -11.62156642 | 0.5633605 |

| | | | | | | | | | | | | | |
|---|---|---|---|---|---|---|---|---|---|---|---|---|---|
| CNAG_03629 | - | NADH dehydrogenase (quinone), G subunit | 16494.15152 | 16405.90825 | 16293.1472 | 10465.93976 | 11241.51489 | 10890.90078 | 6.34E-30 | -0.609069975 | 3.36E-31 | -11.61739215 | 0.655619207 |
| CNAG_06847 | - | small subunit ribosomal protein S28 | 7730.144568 | 8325.069278 | 8064.334484 | 4444.18537 | 5097.472094 | 4181.218696 | 6.37E-30 | -0.8289108 | 3.38E-31 | -11.61685297 | 0.562954099 |
| CNAG_04836 | FZC10 | nuclear protein | 293.7920745 | 317.0580758 | 327.642901 | 765.137093 | 734.489164 | 866.3940181 | 6.75E-30 | 1.32090002 | 3.60E-31 | 11.61165494 | 2.498219117 |
| CNAG_02327 | - | xaa-Pro dipeptidase | 5359.383479 | 4719.010746 | 5252.065008 | 2647.822219 | 2791.719014 | 3033.214378 | 7.19E-30 | -0.871561161 | 3.84E-31 | -11.606106 | 0.546555095 |
| CNAG_01898 | - | protein MAK11 | 2462.395913 | 2487.599488 | 2599.808729 | 1426.002998 | 1499.664485 | 1461.254307 | 7.21E-30 | -0.798500517 | 3.86E-31 | -11.60562859 | 0.574946445 |
| CNAG_00270 | - | mitochondrial protein | 2820.723368 | 2558.742635 | 2708.605066 | 1527.970072 | 1561.750346 | 1507.901643 | 7.28E-30 | -0.831097097 | 3.90E-31 | -11.60469658 | 0.56210163 |
| CNAG_03863 | - | protein arginine N-methyltransferase 1 | 9245.814513 | 8582.694588 | 9531.755222 | 5207.70422 | 5734.01319 | 5513.324173 | 7.38E-30 | -0.749308833 | 3.97E-31 | -11.60324927 | 0.594888489 |
| CNAG_00573 | - | NADH dehydrogenase (ubiquinone) 1 alpha subcomplex 6 | 4187.123532 | 3945.415777 | 4340.259868 | 2454.16503 | 2502.821362 | 2499.238584 | 7.53E-30 | -0.758021021 | 4.05E-31 | -11.60139497 | 0.591306884 |
| CNAG_04648 | - | sister chromatid cohesion protein pds5 | 2302.058647 | 2433.006685 | 2218.404785 | 3833.600282 | 3874.103466 | 3928.565486 | 8.49E-30 | 0.727849651 | 4.58E-31 | 11.59090739 | 1.656168717 |
| CNAG_01634 | - | large subunit GTPase 1 | 5375.846946 | 4916.190818 | 5413.119212 | 3008.271009 | 3204.962755 | 3138.692524 | 1.02E-29 | -0.763870652 | 5.54E-31 | -11.57471741 | 0.588914193 |
| CNAG_01874 | - | Glutathione S-transferase | 175.8463171 | 169.4829685 | 157.0648416 | 535.2164054 | 519.1398267 | 460.3264444 | 1.03E-29 | 1.58011093 | 5.59E-31 | 11.57383288 | 2.989928385 |
| CNAG_01040 | - | carboxypeptidase D | 70.86697691 | 55.34979212 | 74.96282779 | 319.8064587 | 274.8793248 | 336.6536267 | 1.12E-29 | 2.207267493 | 6.10E-31 | 11.56633091 | 4.617997822 |
| CNAG_07628 | - | translation initiation factor 3 subunit C | 18719.85158 | 17806.59763 | 18958.17871 | 11502.59847 | 12136.81879 | 12196.44775 | 1.20E-29 | -0.64627718 | 6.54E-31 | -11.560365 | 0.638926915 |
| CNAG_03165 | - | hypothetical protein | 2690.101551 | 2752.631791 | 2912.253252 | 1538.841367 | 1667.019649 | 1445.776063 | 1.24E-29 | -0.860673674 | 6.76E-31 | -11.55757894 | 0.550695348 |
| CNAG_00279 | - | hypothetical protein | 4493.055533 | 4223.38757 | 4492.787932 | 2752.328748 | 2606.308957 | 2639.124606 | 1.32E-29 | -0.739412865 | 7.19E-31 | -11.55221558 | 0.598983071 |
| CNAG_03226 | - | succinate dehydrogenase [ubiquinone] iron-sulfur subunit, mitochondrial | 2714.14325 | 2203.179894 | 2311.883187 | 1236.821575 | 1146.739946 | 1197.10159 | 1.38E-29 | -1.030098785 | 7.56E-31 | -11.54793239 | 0.489676618 |
| CNAG_04304 | - | T-complex protein 1 subunit zeta | 13811.79648 | 13160.71051 | 13799.75841 | 8985.691881 | 9022.115271 | 8812.695967 | 1.52E-29 | -0.619798542 | 8.34E-31 | -11.53953137 | 0.650761794 |
| CNAG_07676 | - | ATP-dependent RNA helicase DBP2-A | 8100.156543 | 7559.089376 | 7636.712234 | 4144.486176 | 4589.347373 | 4846.084034 | 1.61E-29 | -0.794161223 | 8.88E-31 | -11.53413053 | 0.576678354 |
| CNAG_04744 | - | mannose-6-phosphate isomerase, class I | 236.580447 | 261.0291897 | 190.1491344 | 752.5476351 | 640.5818885 | 636.7931198 | 1.63E-29 | 1.547134071 | 8.98E-31 | 11.53310801 | 2.922360323 |
| CNAG_00164 | - | hypothetical protein | 154.9299922 | 165.196985 | 200.7483277 | 601.5763586 | 526.7935927 | 504.6896652 | 1.70E-29 | 1.636723112 | 9.41E-31 | 11.52912238 | 3.109587282 |
| CNAG_02128 | - | translation initiation factor 3 subunit J | 9486.365974 | 8669.979022 | 9602.155856 | 5554.880119 | 5790.929201 | 5743.11434 | 1.95E-29 | -0.715729489 | 1.08E-30 | -11.51735013 | 0.608897169 |
| CNAG_03311 | ERG13 | hydroxymethylglutaryl-CoA synthase | 10463.80372 | 9719.395169 | 10493.73092 | 6414.939893 | 6631.087446 | 6392.561598 | 2.16E-29 | -0.67397154 | 1.20E-30 | -11.50808289 | 0.626778875 |
| CNAG_02440 | - | cation-transporting ATPase | 5782.349252 | 5868.593616 | 5686.201126 | 3617.752732 | 3795.856224 | 3751.463822 | 2.28E-29 | -0.650231393 | 1.27E-30 | -11.50331653 | 0.637178109 |
| CNAG_06645 | MTD1 | methylenetetrahydrofolate dehydrogenase (NAD) | 1910.093043 | 2156.729419 | 2219.011834 | 980.6483109 | 1167.564623 | 1011.723983 | 2.41E-29 | -1.007976811 | 1.35E-30 | -11.49818744 | 0.497243077 |
| CNAG_04192 | - | phosphoribosylformylglycinamidine synthase | 5978.371994 | 5740.363398 | 5414.265041 | 3398.237785 | 3625.923503 | 3295.255764 | 2.67E-29 | -0.747021488 | 1.49E-30 | -11.48931411 | 0.595832413 |
| CNAG_06106 | - | chaperone regulator | 7068.875085 | 6237.072746 | 5999.742171 | 3468.179642 | 3659.482619 | 3778.146698 | 2.76E-29 | -0.839617811 | 1.55E-30 | -11.48610864 | 0.558791581 |
| CNAG_03340 | - | flavonol synthase | 45.20343989 | 47.78542516 | 30.21254278 | 258.4606427 | 277.20648 | 213.7968261 | 3.03E-29 | 2.601004408 | 1.70E-30 | 11.47786777 | 6.06708872 |
| CNAG_03636 | - | DNA-directed RNA polymerase I and III subunit RPAC2 | 1163.714417 | 1176.297645 | 1213.372208 | 622.7956109 | 573.1575191 | 537.7101202 | 3.34E-29 | -1.051470546 | 1.88E-30 | -11.46937696 | 0.482476124 |
| CNAG_12523 | - | unspecified product | 844.9227028 | 896.1134603 | 905.1425619 | 425.8168155 | 373.8376496 | 393.3818383 | 5.21E-29 | -1.166051524 | 2.94E-30 | -11.43069614 | 0.445639332 |
| CNAG_02086 | - | Acyl-CoA-dependent ceramide synthase | 4075.018078 | 3831.209545 | 4356.226847 | 2341.285476 | 2331.133108 | 2449.27572 | 6.20E-29 | -0.799632686 | 3.51E-30 | -11.41531601 | 0.574495427 |
| CNAG_03577 | - | large subunit ribosomal protein LP0 | 22657.30207 | 21235.98977 | 21601.8896 | 10541.16382 | 13619.57615 | 11320.65592 | 6.32E-29 | -0.900268554 | 3.58E-30 | -11.41346468 | 0.535786987 |
| CNAG_00966 | - | aromatic-L-amino-acid decarboxylase | 2170.3346 | 2041.559471 | 2172.160008 | 1224.467308 | 1208.951353 | 1141.571651 | 6.51E-29 | -0.852704405 | 3.70E-30 | -11.41069254 | 0.553745738 |
| CNAG_13196 | - | unspecified product | 20149.20019 | 21825.29685 | 21594.40834 | 13295.87642 | 10576.25655 | 11761.35324 | 6.58E-29 | -0.849800313 | 3.75E-30 | -11.4095599 | 0.55486153 |
| CNAG_03080 | - | fatty acid elongase | 5393.472034 | 5229.714087 | 6090.253449 | 3135.117573 | 3273.939266 | 3268.589689 | 6.79E-29 | -0.803997986 | 3.87E-30 | -11.40672248 | 0.572759749 |
| CNAG_02751 | - | Short-chain dehydrogenase | 1001.194056 | 814.8253823 | 872.5822123 | 1877.297727 | 1839.459699 | 1789.238829 | 6.89E-29 | 1.018687264 | 3.94E-30 | 11.40517845 | 2.026074557 |
| CNAG_05693 | - | hypothetical protein | 123.3761778 | 132.8600411 | 144.2413809 | 421.3907911 | 419.0195569 | 427.9443573 | 8.33E-29 | 1.651856752 | 4.77E-30 | 11.38848446 | 3.142378039 |
| CNAG_04769 | - | hypothetical protein, variant | 869.2940415 | 903.0951415 | 865.0883526 | 1610.910734 | 1658.286843 | 1611.641416 | 8.89E-29 | 0.872578452 | 5.10E-30 | 11.38267248 | 1.830932306 |
| CNAG_00536 | - | hypothetical protein, variant | 4666.005658 | 4762.077881 | 4885.251727 | 3108.829843 | 2939.281353 | 2932.198852 | 9.14E-29 | -0.687885 | 5.26E-30 | -11.38002481 | 0.620763226 |
| CNAG_06820 | - | hypothetical protein, variant | 1255.940945 | 1257.103836 | 1258.191187 | 599.3172302 | 594.7003466 | 680.8182861 | 1.04E-28 | -1.024307593 | 6.00E-30 | -11.36849446 | 0.491646203 |
| CNAG_01799 | - | ubiquinol-cytochrome c reductase subunit 8 | 6022.913759 | 5905.263936 | 6423.05232 | 3826.352321 | 3912.287926 | 3703.770262 | 1.07E-28 | -0.697084654 | 6.20E-30 | -11.36570226 | 0.616817391 |
| CNAG_02257 | - | GTP-binding nuclear protein spi1 | 19416.9647 | 18106.12771 | 18708.65795 | 12506.69146 | 12220.47341 | 12278.61093 | 1.08E-28 | -0.619229861 | 6.26E-30 | -11.36475909 | 0.651018362 |
| CNAG_06302 | - | hypothetical protein, variant | 82.48292512 | 63.95237084 | 65.38078309 | 296.9912501 | 292.2780744 | 321.1880739 | 1.09E-28 | 2.098174009 | 6.30E-30 | 11.36427075 | 4.281671191 |
| CNAG_00822 | - | ribosome assembly protein SQT1 | 2468.17859 | 2391.724603 | 2675.477434 | 1288.143396 | 1451.35221 | 1391.332215 | 1.11E-28 | -0.882880056 | 6.44E-30 | -11.36232445 | 0.542283787 |
| CNAG_06337 | - | hypothetical protein | 19.53600579 | 26.24481689 | 29.12078171 | 182.7171226 | 194.206412 | 195.6947089 | 1.13E-28 | 2.933076955 | 6.57E-30 | 11.36060497 | 7.637375489 |
| CNAG_05976 | - | nucleolar protein 58 | 4841.066235 | 4547.704107 | 5081.433786 | 2770.893869 | 3000.552297 | 2818.983664 | 1.24E-28 | -0.768321525 | 7.23E-30 | -11.3521663 | 0.587100127 |
| CNAG_06431 | - | acyl-CoA oxidase | 597.2185989 | 552.9846388 | 526.0025176 | 1111.21266 | 1219.346504 | 1234.040698 | 1.25E-28 | 1.073880774 | 7.31E-30 | 11.35124109 | 2.105088335 |
| CNAG_04733 | - | DNA repair protein rad5 | 3016.660745 | 2889.861417 | 2924.370893 | 4725.503091 | 4728.872677 | 4776.757607 | 1.30E-28 | 0.673107372 | 7.62E-30 | 11.34762445 | 1.594503619 |
| CNAG_02900 | - | hypothetical protein | 54.60678844 | 79.02368276 | 88.81652225 | 313.1897275 | 317.2655427 | 384.2892083 | 1.37E-28 | 2.183688169 | 8.05E-30 | 11.34284121 | 4.543134978 |
| CNAG_00448 | - | V-type H -transporting ATPase subunit AC39 | 5333.905267 | 5187.686208 | 5349.174471 | 3420.124774 | 3275.679887 | 3396.217042 | 1.48E-28 | -0.668478403 | 8.73E-30 | -11.33572564 | 0.629169917 |
| CNAG_01236 | - | rRNA-processing protein EBP2 | 3731.725803 | 3774.04484 | 4139.712225 | 1959.012034 | 2023.144895 | 2337.127171 | 1.63E-28 | -0.896976181 | 9.60E-30 | -11.32738659 | 0.537011102 |
| CNAG_01060 | - | hypothetical protein | 13915.71503 | 13226.43134 | 13567.28799 | 9132.033081 | 8940.160565 | 8965.886392 | 1.66E-28 | -0.605859765 | 9.83E-30 | -11.32532961 | 0.657079681 |
| CNAG_02495 | - | cytoplasmic protein | 2399.272153 | 2299.072569 | 2533.660789 | 1292.858735 | 1361.618185 | 1407.953391 | 1.94E-28 | -0.848221745 | 1.15E-29 | -11.31157652 | 0.555468981 |
| CNAG_03742 | - | staphylococcal nuclease domain-containing protein 1 | 5428.40618 | 5145.653462 | 5316.101207 | 3223.158446 | 2961.638008 | 3341.81511 | 2.13E-28 | -0.753559886 | 1.27E-29 | -11.30320319 | 0.593138167 |
| CNAG_00433 | - | hypothetical protein | 254.0276345 | 213.6713536 | 257.2596683 | 620.4199954 | 659.6851446 | 631.206999 | 2.62E-28 | 1.383571603 | 1.56E-29 | 11.28505483 | 2.609135017 |
| CNAG_00061 | CIT1 | citrate synthase, mitochondrial | 18117.28214 | 17163.37031 | 17698.72471 | 11318.17534 | 11811.6432 | 11895.58431 | 2.86E-28 | -0.61257689 | 1.70E-29 | -11.27712909 | 0.654027458 |
| CNAG_01131 | - | hypothetical protein | 58.02295139 | 52.10196835 | 32.35084515 | 318.8619567 | 271.7261434 | 237.1074553 | 2.95E-28 | 2.524992092 | 1.76E-29 | 11.27408101 | 5.755702771 |
| CNAG_02685 | - | hypothetical protein | 109.4115728 | 144.6891672 | 114.4255476 | 466.1400844 | 453.7079709 | 375.9755139 | 2.99E-28 | 1.80352636 | 1.79E-29 | 11.27273682 | 3.490724165 |
| CNAG_03724 | - | ribosomal RNA methyltransferase Nop2 | 5954.935732 | 5112.297136 | 5655.213119 | 3160.810014 | 3336.059303 | 3217.534774 | 3.05E-28 | -0.799731526 | 1.83E-29 | -11.27073929 | 0.574456069 |
| CNAG_04748 | - | protein OS-9 | 789.9900687 | 952.6395834 | 850.171414 | 1659.97988 | 1737.699578 | 1663.797863 | 3.15E-28 | 0.950794293 | 1.89E-29 | 11.26783465 | 1.932936566 |
| CNAG_07600 | - | Beta-glucosidase | 415.0788778 | 309.5644952 | 286.1191373 | 893.6296516 | 902.8369879 | 970.7838982 | 4.13E-28 | 1.43683194 | 2.49E-29 | 11.24363773 | 2.707257171 |
| CNAG_00806 | - | hypothetical protein | 6791.384632 | 7202.457663 | 7827.53881 | 4322.061021 | 4585.080839 | 4350.995959 | 4.43E-28 | -0.73413036 | 2.67E-29 | -11.23745432 | 0.6011803 |
| CNAG_03820 | - | nucleolar protein 14 | 6689.582403 | 6113.224856 | 6768.58271 | 3734.40306 | 4052.869204 | 4022.363741 | 6.23E-28 | -0.744664723 | 3.77E-29 | -11.20700188 | 0.596806551 |
| CNAG_05367 | - | hypothetical protein | 2149.42986 | 2205.292193 | 2205.24367 | 1217.397523 | 1290.053097 | 1309.140033 | 6.66E-28 | -0.796570042 | 4.03E-29 | -11.20096549 | 0.575716298 |
| CNAG_04808 | MKT1 | XPG N-terminal domain-containing protein | 3136.932384 | 2966.362854 | 2822.021026 | 4968.378623 | 5150.761415 | 4799.001627 | 8.74E-28 | 0.725686068 | 5.32E-29 | 11.17644903 | 1.653686854 |
| CNAG_05360 | - | ATP-dependent RNA helicase MAK5 | 3203.86837 | 3073.751944 | 3413.482549 | 1698.316243 | 1925.777333 | 1875.338961 | 8.74E-28 | -0.833065293 | 5.31E-29 | -11.17660866 | 0.561335306 |
| CNAG_03421 | - | hypothetical protein | 66.18531757 | 70.38955725 | 58.98314443 | 294.0989446 | 298.5240298 | 255.9699844 | 1.11E-27 | 2.111187553 | 6.76E-29 | 11.15510845 | 4.32046787 |
| CNAG_03065 | - | ER-derived vesicles protein ERV14 | 6641.85558 | 6714.401118 | 7010.691616 | 4330.700135 | 4180.49941 | 4497.473276 | 1.20E-27 | -0.661963111 | 7.33E-29 | -11.14796835 | 0.632017709 |
| CNAG_01390 | - | adenine phosphoribosyltransferase | 5899.992658 | 4919.441978 | 5481.379774 | 3146.062387 | 3100.543787 | 3035.485926 | 1.30E-27 | -0.828571108 | 7.94E-29 | -11.14082406 | 0.563086665 |
| CNAG_06460 | - | hypothetical protein | 1135.527068 | 1101.367822 | 1153.016923 | 1939.161075 | 2132.698184 | 2202.150212 | 1.38E-27 | 0.872907042 | 8.49E-29 | 11.13484565 | 1.831349369 |
| CNAG_06331 | - | hypothetical protein | 102.4985994 | 157.6564893 | 187.9580141 | 568.8089575 | 540.7322908 | 517.9076575 | 1.58E-27 | 1.85096249 | 9.68E-29 | 11.12317839 | 3.60740772 |
| CNAG_07188 | - | hypothetical protein | 179.3336649 | 183.4653401 | 143.2170302 | 470.9670973 | 514.8205885 | 514.5291385 | 1.79E-27 | 1.553704604 | 1.10E-28 | 11.1118129 | 2.935700116 |
| CNAG_02706 | - | hypothetical protein | 35.82896201 | 29.47598399 | 19.53961005 | 201.5998976 | 201.0842357 | 191.3786237 | 1.88E-27 | 2.821424047 | 1.16E-28 | 11.10727519 | 7.068597759 |
| CNAG_06716 | - | hypothetical protein | 2192.649722 | 2192.395568 | 2474.971152 | 1309.30054 | 1222.70542 | 1257.009396 | 1.95E-27 | -0.871697462 | 1.21E-28 | -11.10351885 | 0.546503461 |
| CNAG_03456 | - | hypothetical protein | 1942.551489 | 1751.742175 | 1905.537412 | 1001.795804 | 990.6757616 | 1054.936662 | 2.06E-27 | -0.894124784 | 1.28E-28 | -11.09840192 | 0.538073521 |
| CNAG_04875 | - | hypothetical protein | 874.0498932 | 884.8308869 | 1036.752744 | 1866.34874 | 1765.18017 | 1853.589266 | 2.11E-27 | 0.957591737 | 1.31E-28 | 11.09630778 | 1.942065336 |
| CNAG_06027 | - | aryl-alcohol dehydrogenase | 43.00495325 | 13.3469481 | 29.17197182 | 679.9337235 | 670.572696 | 718.2417591 | 2.20E-27 | 4.584857059 | 1.37E-28 | 11.09216437 | 23.99824599 |
| CNAG_04614 | - | hypothetical protein | 63.95379848 | 120.9843861 | 80.31975327 | 374.0903097 | 366.4772846 | 398.8986902 | 2.31E-27 | 2.100375354 | 1.44E-28 | 11.08757333 | 4.288209393 |
| CNAG_02443 | - | NADH dehydrogenase | 2365.469894 | 2346.473097 | 2523.013481 | 1428.399297 | 1363.332425 | 1425.717001 | 2.36E-27 | -0.794218936 | 1.47E-28 | -11.08571215 | 0.576655285 |
| CNAG_00393 | - | 1,4-alpha-glucan-branching enzyme | 2855.547727 | 2728.263828 | 2242.999166 | 4868.866772 | 4881.64783 | 4954.293381 | 2.63E-27 | 0.894768217 | 1.65E-28 | 11.07562496 | 1.859311144 |
| CNAG_05356 | - | hypothetical protein | 38.20600805 | 32.72196175 | 44.04680583 | 232.6507928 | 194.6208567 | 266.5306773 | 2.71E-27 | 2.586435158 | 1.70E-28 | 11.07276434 | 6.006127738 |
| CNAG_06168 | GNO1 | S-(hydroxymethyl)glutathione dehydrogenase | 7408.997092 | 7369.462964 | 7269.874269 | 4354.722503 | 4911.171682 | 4696.218297 | 2.88E-27 | -0.674479547 | 1.81E-28 | -11.06721537 | 0.62655821 |
| CNAG_01648 | H4 | Histone H4 | 12105.50565 | 15242.59805 | 14576.44408 | 23848.11322 | 25365.77187 | 24605.30395 | 2.88E-27 | 0.80190356 | 1.81E-28 | 11.06702078 | 1.743399934 |
| CNAG_04782 | PPU1 | hypothetical protein, variant | 2048.69808 | 2258.460917 | 2147.999509 | 3494.221598 | 3552.351195 | 3703.175415 | 3.02E-27 | 0.721379009 | 1.90E-28 | 11.06271776 | 1.648757257 |
| CNAG_00722 | - | large subunit ribosomal protein L27 | 2622.214062 | 2364.813918 | 2517.71321 | 1436.921885 | 1428.036101 | 1463.467305 | 3.44E-27 | -0.809784552 | 2.17E-28 | -11.05093034 | 0.570467043 |
| CNAG_00002 | - | hypothetical protein | 924.2245349 | 945.6658517 | 906.2524885 | 458.5388216 | 423.9178264 | 361.2510616 | 3.54E-27 | -1.175058879 | 2.24E-28 | -11.04816843 | 0.442865685 |
| CNAG_03600 | STE6 | ATP-binding cassette, subfamily B (MDR/TAP), member 1 | 79.01917738 | 88.68827864 | 71.7753963 | 317.3044328 | 287.8376599 | 331.2067698 | 3.60E-27 | 1.959415226 | 2.28E-28 | 11.04633679 | 3.889043107 |
| CNAG_01253 | - | mitochondrial import inner membrane translocase subunit TIM10 | 3315.902872 | 3047.905794 | 3261.026381 | 1879.788995 | 1974.100334 | 1773.245441 | 3.62E-27 | -0.790183369 | 2.30E-28 | -11.04581628 | 0.578270588 |

| | | | | | | | | | | | | | |
|---|---|---|---|---|---|---|---|---|---|---|---|---|---|
| CNAG_00280 | - | complement component 1 Q subcomponent-binding protein, mitochondrial | 4758.082674 | 4347.297703 | 4566.389462 | 2735.144558 | 2899.647451 | 2645.808407 | 3.65E-27 | -0.739397732 | 2.32E-28 | -11.04473832 | 0.598989354 |
| CNAG_06948 | - | essential nuclear protein 1 | 4290.916879 | 3727.795894 | 4241.06944 | 2227.596082 | 2377.734471 | 2395.999465 | 3.68E-27 | -0.824483212 | 2.35E-28 | -11.04385438 | 0.564684442 |
| CNAG_12491 | - | unspecified product | 8.990126283 | 13.31389342 | 15.26015864 | 204.2065337 | 128.1622635 | 162.4041692 | 3.80E-27 | 3.764172865 | 2.42E-28 | 11.04097607 | 13.58716787 |
| CNAG_03783 | - | hypothetical protein | 13.68811807 | 28.38488373 | 18.46538514 | 198.1178358 | 172.535329 | 166.9890046 | 3.81E-27 | 3.169770318 | 2.44E-28 | 11.04055046 | 8.999035087 |
| CNAG_05147 | - | hypothetical protein | 72.03737136 | 86.534338 | 80.28830021 | 295.3155125 | 317.4160531 | 303.5826461 | 3.88E-27 | 1.92797335 | 2.49E-28 | 11.03856794 | 3.805202815 |
| CNAG_00750 | - | hypothetical protein | 1711.546575 | 1798.009412 | 1981.182927 | 876.3709266 | 1010.560572 | 971.7063871 | 4.33E-27 | -0.957596998 | 2.78E-28 | -11.02854413 | 0.514913858 |
| CNAG_04948 | TUB2 | tubulin beta | 5747.330381 | 5655.298918 | 5920.786507 | 3690.163591 | 3667.300634 | 3830.268852 | 4.72E-27 | -0.646098591 | 3.04E-28 | -11.020664 | 0.639006012 |
| CNAG_03353 | - | hypothetical protein | 2182.09076 | 2108.367796 | 2316.108167 | 1203.374541 | 1265.885637 | 1288.057926 | 4.72E-27 | -0.830067201 | 3.04E-28 | -11.0204688 | 0.56250304 |
| CNAG_05462 | - | cytochrome b5 reductase | 11887.38948 | 11598.43249 | 11934.54662 | 8062.670986 | 8020.691914 | 7998.923464 | 5.24E-27 | -0.572080025 | 3.39E-28 | -11.01089009 | 0.672646292 |
| CNAG_03871 | LTP1 | phosphotyrosine protein phosphatase | 1172.837171 | 1093.32705 | 1026.822865 | 553.4810763 | 494.6192494 | 534.3368927 | 5.48E-27 | -1.073005207 | 3.55E-28 | -11.00672571 | 0.475327835 |
| CNAG_05978 | - | glutamate-tRNA ligase | 16849.02035 | 15181.98454 | 16019.05241 | 10005.95696 | 10396.16229 | 10430.18955 | 5.65E-27 | -0.655893391 | 3.67E-28 | -11.00376244 | 0.634682341 |
| CNAG_00292 | - | zinc finger protein | 8803.038937 | 7414.771551 | 7997.16864 | 4556.272945 | 4828.286011 | 4806.155548 | 5.97E-27 | -0.787053111 | 3.88E-28 | -10.99863267 | 0.579526641 |
| CNAG_01682 | - | mitochondrial outer membrane 72K protein | 9182.768076 | 8858.485138 | 8455.806314 | 5767.448509 | 5878.087084 | 5608.784023 | 5.99E-27 | -0.634281832 | 3.90E-28 | -10.9982633 | 0.644261446 |
| CNAG_06753 | - | nuclear cap-binding protein subunit 1 | 5183.279191 | 4998.072565 | 5455.782312 | 3141.401391 | 3354.21702 | 3258.571803 | 7.05E-27 | -0.696522934 | 4.60E-28 | -10.98321003 | 0.617057598 |
| CNAG_03055 | - | bud site selection protein 20 | 1640.112815 | 1469.464672 | 1695.399057 | 837.5612134 | 855.2700979 | 826.2945077 | 7.05E-27 | -0.947820446 | 4.59E-28 | -10.98337568 | 0.518415067 |
| CNAG_03124 | - | periodic tryptophan protein 2 | 6465.272773 | 5672.552882 | 6308.935095 | 3368.571694 | 3420.528894 | 3816.964767 | 7.55E-27 | -0.814397681 | 4.94E-28 | -10.97678655 | 0.568645844 |
| CNAG_04911 | - | hypothetical protein, variant | 108.2171052 | 130.6897196 | 131.4474982 | 380.0272198 | 398.36069 | 404.606276 | 7.71E-27 | 1.664756448 | 5.06E-28 | 10.97469683 | 3.170601249 |
| CNAG_07604 | FRE5 | metalloreductase | 227.214322 | 238.4073747 | 255.115503 | 584.5668019 | 619.1013577 | 597.9197283 | 8.86E-27 | 1.306704461 | 5.82E-28 | 10.96195462 | 2.473758151 |
| CNAG_06563 | - | translation initiation factor 3 subunit F | 1620.300608 | 1518.998245 | 1606.939184 | 891.3406354 | 856.1298158 | 862.9048449 | 9.27E-27 | -0.878130748 | 6.11E-28 | -10.95753611 | 0.544071911 |
| CNAG_04280 | - | hypothetical protein | 37.02919308 | 43.46585895 | 35.52600703 | 228.8818456 | 193.8449682 | 230.1054235 | 9.27E-27 | 2.486515315 | 6.10E-28 | 10.95769948 | 5.604226696 |
| CNAG_00287 | - | cytoplasmic protein | 7424.272898 | 7467.542203 | 7998.219146 | 5047.553955 | 4861.949129 | 4930.46073 | 1.04E-26 | -0.640646549 | 6.85E-28 | -10.94727459 | 0.641425427 |
| CNAG_02412 | - | rRNA-processing protein CGR1 | 1315.543457 | 1246.408263 | 1488.442495 | 637.4467471 | 700.0048835 | 647.5874645 | 1.23E-26 | -1.044934032 | 8.13E-28 | -10.93166093 | 0.48466707 |
| CNAG_02081 | - | mannosyl-oligosaccharide alpha-1,2-mannosidase | 4197.716532 | 4165.198667 | 4239.004148 | 2694.736106 | 2746.976315 | 2709.028716 | 1.28E-26 | -0.644166893 | 8.46E-28 | -10.92805553 | 0.639862182 |
| CNAG_05664 | - | Branched-chain-amino-acid transaminase | 2426.254238 | 2534.999974 | 2741.608335 | 1375.349341 | 1497.926887 | 1511.196441 | 1.43E-26 | -0.828337513 | 9.50E-28 | -10.91760064 | 0.563177846 |
| CNAG_05260 | - | Glutamate decarboxylase | 3177.836238 | 3282.018191 | 2880.684033 | 5317.913339 | 5156.726076 | 5014.368086 | 1.47E-26 | 0.714738819 | 9.76E-28 | 10.91510876 | 1.641186075 |
| CNAG_04983 | - | hypothetical protein | 1365.88245 | 1443.523194 | 1552.483152 | 775.3282619 | 709.4307864 | 782.968139 | 1.56E-26 | -0.959185042 | 1.04E-27 | -10.90916667 | 0.514347379 |
| CNAG_01232 | PMC1 | calcium-translocating P-type ATPase, PMCA-type | 1923.689979 | 2007.419532 | 1841.94303 | 3165.056062 | 3143.487373 | 3224.728409 | 1.67E-26 | 0.70900702 | 1.11E-27 | 10.90324192 | 1.634678611 |
| CNAG_01733 | - | pre-mRNA-processing factor 19 | 5627.09641 | 5738.247012 | 5862.140252 | 3676.927114 | 3884.723271 | 3721.496005 | 1.73E-26 | -0.626055455 | 1.15E-27 | -10.89984591 | 0.647945576 |
| CNAG_03475 | - | hypothetical protein | 1140.25609 | 1322.183423 | 1063.488386 | 2281.903784 | 2148.118574 | 2342.033388 | 1.89E-26 | 0.927452155 | 1.27E-27 | 10.89119541 | 1.901914187 |
| CNAG_03931 | - | mitochondrial import receptor subunit TOM40 | 5276.769031 | 5347.144415 | 5514.473004 | 3578.144545 | 3493.942105 | 3501.680709 | 1.91E-26 | -0.625350445 | 1.29E-27 | -10.89009918 | 0.648262289 |
| CNAG_06353 | - | tRNA pseudouridine(38-40) synthase | 4559.673197 | 4364.537773 | 4609.043942 | 2754.604639 | 2955.727895 | 2822.280895 | 2.72E-26 | -0.681162282 | 1.83E-27 | -10.857724 | 0.623662629 |
| CNAG_01544 | - | histidyl-tRNA synthetase | 7272.431307 | 7180.94741 | 7421.301418 | 4681.696711 | 4880.965646 | 4928.211175 | 3.20E-26 | -0.609542778 | 2.16E-27 | -10.8428271 | 0.655404381 |
| CNAG_01451 | - | hypothetical protein | 849.4948111 | 793.8506813 | 939.1923464 | 393.0781647 | 392.0070238 | 334.6035545 | 3.29E-26 | -1.221565943 | 2.22E-27 | -10.84011542 | 0.428817015 |
| CNAG_06126 | - | Pin2-interacting protein X1 | 1642.568372 | 1669.79101 | 1690.120691 | 847.6271739 | 890.6245545 | 982.7453174 | 3.38E-26 | -0.894247396 | 2.29E-27 | -10.83750344 | 0.538027793 |
| CNAG_02914 | - | hypothetical protein | 1419.657573 | 1548.030005 | 1680.444107 | 831.3691142 | 787.0886315 | 817.398758 | 3.46E-26 | -0.947360347 | 2.34E-27 | -10.83530149 | 0.518580424 |
| CNAG_02901 | - | U6 snRNA-associated Sm-like protein LSm4 | 1921.596715 | 1828.213467 | 1886.365633 | 993.9816059 | 1042.457733 | 1117.076188 | 3.54E-26 | -0.853552997 | 2.40E-27 | -10.83298829 | 0.553420121 |
| CNAG_02586 | - | Sugar transporter | 31.21462034 | 45.61444296 | 30.20219817 | 259.7889391 | 197.3351038 | 202.5818437 | 3.54E-26 | 2.620040615 | 2.41E-27 | 10.83271334 | 6.147673793 |
| CNAG_04625 | - | cerevisin | 725.6600325 | 714.5690004 | 613.4610391 | 1523.364823 | 1371.895899 | 1333.007259 | 3.71E-26 | 1.027487128 | 2.53E-27 | 10.82818514 | 2.03847057 |
| CNAG_04548 | - | hypothetical protein | 26.4836275 | 10.08288582 | 15.26396929 | 182.4358867 | 175.2930924 | 153.7130636 | 4.47E-26 | 3.304954366 | 3.05E-27 | 10.81109584 | 9.883036483 |
| CNAG_05845 | - | protein FAM32A | 4038.799968 | 3859.199331 | 3931.857151 | 2553.136972 | 2287.140606 | 2390.442698 | 4.50E-26 | -0.725493426 | 3.08E-27 | -10.81030851 | 0.604790161 |
| CNAG_04584 | - | poly(rC)-binding protein 2/3/4 | 1886.39139 | 1947.122838 | 2049.874509 | 3559.09092 | 3425.619744 | 3144.896005 | 4.62E-26 | 0.768687417 | 3.16E-27 | 10.80773315 | 1.703719012 |
| CNAG_02765 | - | 3-hydroxyisobutyrate dehydrogenase | 99.98786753 | 79.03259957 | 104.7927028 | 349.9996564 | 330.1885534 | 334.6945507 | 5.03E-26 | 1.828966338 | 3.45E-27 | 10.79982526 | 3.552824286 |
| CNAG_02509 | - | hypothetical protein | 35.91936772 | 67.14761739 | 57.90647114 | 258.8999664 | 278.7135847 | 261.3474468 | 5.20E-26 | 2.311749509 | 3.57E-27 | 10.79659267 | 4.964847859 |
| CNAG_04136 | - | methylthioribose-1-phosphate isomerase | 2061.683268 | 1849.802014 | 2034.580829 | 1142.752685 | 1103.703518 | 1067.187062 | 5.90E-26 | -0.859519278 | 4.06E-27 | -10.78480229 | 0.551136172 |
| CNAG_04710 | - | hypothetical protein | 2769.140256 | 2851.055281 | 2895.558726 | 4534.842926 | 4576.227541 | 4343.860359 | 6.06E-26 | 0.644622619 | 4.18E-27 | 10.78213055 | 1.563330296 |
| CNAG_03483 | - | protein GET1 | 2802.207305 | 2717.131495 | 2903.763217 | 1689.902903 | 1755.010122 | 1691.044624 | 6.32E-26 | -0.729300024 | 4.37E-27 | -10.77806297 | 0.603196506 |
| CNAG_00086 | - | N2,N2-dimethylguanosine tRNA methyltransferase | 3720.088289 | 3599.563054 | 3972.333544 | 2272.062145 | 2362.253054 | 2312.723191 | 6.85E-26 | -0.71646158 | 4.74E-27 | -10.77049407 | 0.608588265 |
| CNAG_06840 | - | elongation factor 2, variant | 109713.4274 | 107226.9944 | 105211.9045 | 65427.95577 | 73166.49511 | 72996.61743 | 7.06E-26 | -0.621848809 | 4.89E-27 | -10.76762502 | 0.649837629 |
| CNAG_00334 | - | hsp75-like protein | 90531.92478 | 84771.18652 | 89306.79292 | 55003.56716 | 60241.01823 | 59382.72232 | 7.13E-26 | -0.615252578 | 4.95E-27 | -10.76650645 | 0.652815593 |
| CNAG_07557 | - | ATP-binding cassette transporter | 1986.58715 | 1606.26915 | 1609.119898 | 855.4135013 | 862.9929327 | 851.8462078 | 7.94E-26 | -1.034257296 | 5.52E-27 | -10.75647191 | 0.488267177 |
| CNAG_04650 | - | actin-like protein arp6 | 984.8665402 | 924.6743784 | 1012.231659 | 1716.681672 | 1785.916695 | 1799.227051 | 8.80E-26 | 0.844480084 | 6.13E-27 | 10.74681601 | 1.795617528 |
| CNAG_05511 | - | ATPase | 753.7380215 | 757.6811856 | 750.9976592 | 1354.607341 | 1569.506516 | 1702.441913 | 9.21E-26 | 1.017973293 | 6.44E-27 | 10.74237755 | 2.025072127 |
| CNAG_02704 | - | hypothetical protein | 1919.458909 | 2016.753756 | 2254.205239 | 1116.208368 | 1179.635121 | 1113.809648 | 9.42E-26 | -0.876083194 | 6.59E-27 | -10.74016023 | 0.544844636 |
| CNAG_00045 | - | hypothetical protein | 228.3459454 | 208.2654892 | 203.973095 | 507.2235491 | 658.1620707 | 602.2704001 | 1.07E-25 | 1.451729465 | 7.47E-27 | 10.72865872 | 2.735357625 |
| CNAG_07015 | - | hypothetical protein | 572.6750113 | 529.2732357 | 486.5481413 | 1037.255891 | 1165.904026 | 1130.837108 | 1.31E-25 | 1.054600291 | 9.16E-27 | 10.70971559 | 2.077142639 |
| CNAG_06766 | - | U3 small nucleolar RNA-associated protein 14 | 4400.761244 | 3973.444281 | 4243.254012 | 2557.704158 | 2573.555692 | 2646.864211 | 1.36E-25 | -0.713853099 | 9.58E-27 | -10.70562886 | 0.609689625 |
| CNAG_04866 | - | NAD-dependent histone deacetylase SIR2 | 651.0439537 | 777.0336136 | 750.9727382 | 1365.712266 | 1428.852892 | 1476.100693 | 1.37E-25 | 0.956292495 | 9.68E-27 | 10.70463339 | 1.940317164 |
| CNAG_02801 | TRX1 | thioredoxin, variant | 4267.767831 | 4203.987064 | 4265.668779 | 2817.761031 | 2749.554926 | 2659.093422 | 1.45E-25 | -0.646066973 | 1.02E-26 | -10.69946771 | 0.639020016 |
| CNAG_07511 | - | nucleolar protein 15 | 6529.707898 | 6466.591814 | 6969.071264 | 3740.598859 | 4123.592636 | 4318.73983 | 1.45E-25 | -0.728137967 | 1.03E-26 | -10.69892674 | 0.603682562 |
| CNAG_06891 | - | hypothetical protein, variant | 1246.673859 | 1253.901036 | 1281.659031 | 708.4591808 | 668.0865955 | 628.7072462 | 1.45E-25 | -0.930795887 | 1.03E-26 | -10.6989792 | 0.524568875 |
| CNAG_04673 | - | NAD dependent epimerase/dehydratase | 58.12840553 | 94.10286324 | 110.1316653 | 371.747697 | 432.45415 | 331.5004573 | 1.59E-25 | 2.099123897 | 1.13E-26 | 10.69068026 | 4.284491224 |
| CNAG_05050 | GCS2 | ADP-ribosylation factor GTPase-activating protein 1 | 2203.10546 | 2078.221003 | 2360.887894 | 1289.071052 | 1252.931791 | 1256.995932 | 1.82E-25 | -0.822145466 | 1.30E-26 | -10.67761797 | 0.565600199 |
| CNAG_05515 | - | hypothetical protein | 715.1678047 | 764.101452 | 620.9236279 | 1460.166494 | 1386.573108 | 1357.409039 | 1.84E-25 | 0.986935197 | 1.31E-26 | 10.67681958 | 1.981970099 |
| CNAG_04637 | MBF1 | multiprotein-bridging factor 1 | 6440.47107 | 6335.458769 | 6371.123715 | 9556.29073 | 9581.607305 | 9370.676268 | 1.84E-25 | 0.558921699 | 1.31E-26 | 10.6764266 | 1.473167729 |
| CNAG_02182 | GRE2 | D-lactaldehyde dehydrogenase | 5251.913967 | 6132.91555 | 4316.21106 | 11378.8422 | 10177.76001 | 10061.08459 | 1.88E-25 | 0.995926506 | 1.35E-26 | 10.67407694 | 1.994360903 |
| CNAG_02918 | ERG10 | Acetyl-CoA C-acetyltransferase | 5444.870908 | 5208.186193 | 6012.432474 | 3381.918642 | 3413.696312 | 3401.792694 | 2.11E-25 | -0.724495454 | 1.51E-26 | -10.66336967 | 0.605208664 |
| CNAG_04746 | - | hypothetical protein | 1778.841662 | 1618.515861 | 1626.521618 | 2853.004957 | 2931.38703 | 2784.035692 | 2.11E-25 | 0.755033813 | 1.51E-26 | 10.66338747 | 1.687671146 |
| CNAG_07893 | - | hypothetical protein | 154.8138261 | 144.7042274 | 157.0312937 | 443.1868557 | 415.448325 | 440.1929455 | 2.19E-25 | 1.496985683 | 1.57E-26 | 10.65949996 | 2.822523676 |
| CNAG_01586 | - | F-type H -transporting ATPase subunit B | 19304.95358 | 18542.5002 | 20936.41222 | 12909.0587 | 12390.37545 | 12674.94261 | 2.28E-25 | -0.645937034 | 1.64E-26 | -10.65554954 | 0.639077573 |
| CNAG_02741 | - | mitochondrial intermediate peptidase 1 | 192.1568622 | 195.3085475 | 182.6327144 | 501.4191485 | 477.5030711 | 525.6509758 | 2.29E-25 | 1.386353325 | 1.65E-26 | 10.65525097 | 2.614170656 |
| CNAG_05293 | - | glycogenin glucosyltransferase | 680.2159716 | 821.189438 | 721.1338108 | 1381.261511 | 1485.825716 | 1470.58299 | 2.32E-25 | 0.950004394 | 1.67E-26 | 10.65378904 | 1.931878542 |
| CNAG_00990 | - | F-type H -transporting ATPase subunit H, variant | 6657.023976 | 6516.179062 | 7624.886094 | 4196.791696 | 4279.719275 | 3981.337342 | 2.44E-25 | -0.755188002 | 1.76E-26 | -10.64904575 | 0.592469174 |
| CNAG_04786 | - | hypothetical protein, variant | 2229.624855 | 2089.352492 | 2293.014693 | 3611.013534 | 3624.813829 | 3635.506871 | 2.84E-25 | 0.701827744 | 2.06E-26 | 10.63465635 | 1.626564175 |
| CNAG_12094 | - | unspecified product | 2487.089827 | 3096.208431 | 3014.605531 | 1454.704264 | 1404.65284 | 1637.754739 | 3.16E-25 | -0.949676529 | 2.29E-26 | -10.62457661 | 0.517748535 |
| CNAG_06836 | - | oxidoreductase | 150.1497124 | 138.2461525 | 148.508538 | 433.0495425 | 462.2978642 | 391.5079616 | 3.19E-25 | 1.544787529 | 2.32E-26 | 10.62336976 | 2.91761097 |
| CNAG_03298 | - | hypothetical protein | 6401.20635 | 5941.922822 | 7382.756336 | 3790.486475 | 3823.390464 | 3479.58321 | 3.79E-25 | -0.846396197 | 2.76E-26 | -10.60714927 | 0.556172302 |
| CNAG_01593 | - | ribosome biogenesis protein SSF1/2 | 4227.849557 | 3578.05356 | 4226.124911 | 2281.355622 | 2265.575308 | 2330.506642 | 4.38E-25 | -0.823186842 | 3.20E-26 | -10.59344341 | 0.565192081 |
| CNAG_12392 | - | unspecified product | 28.83747879 | 30.54056408 | 13.13696646 | 191.5691248 | 199.5216606 | 163.7654274 | 4.61E-25 | 2.946533671 | 3.37E-26 | 10.58861074 | 7.708946262 |
| CNAG_06282 | - | WD-repeat protein 48 | 1477.61384 | 1521.507336 | 1351.394681 | 2416.306254 | 2522.512543 | 2531.964073 | 4.67E-25 | 0.765220815 | 3.42E-26 | 10.58719797 | 1.69963012 |
| CNAG_03110 | - | hypothetical protein | 1135.533041 | 1255.37939 | 1127.433444 | 2151.146295 | 2061.02951 | 1990.272132 | 4.73E-25 | 0.803597212 | 3.46E-26 | 10.58593085 | 1.745447801 |
| CNAG_07405 | - | hypothetical protein, variant | 44.04926407 | 66.05799335 | 45.12451939 | 229.9125294 | 265.8161061 | 246.9023328 | 4.89E-25 | 2.241131523 | 3.59E-26 | 10.58252993 | 4.727677166 |
| CNAG_07486 | - | hypothetical protein | 100.1176559 | 173.7398177 | 111.2389432 | 463.5943881 | 460.5284005 | 441.2947693 | 4.90E-25 | 1.815483374 | 3.60E-26 | 10.58225963 | 3.519775404 |
| CNAG_06611 | - | DNA-directed RNA polymerase I subunit RPA2 | 2546.318989 | 2372.327448 | 2405.753222 | 1294.390101 | 1497.97078 | 1415.735604 | 5.89E-25 | -0.81564683 | 4.34E-26 | -10.56487981 | 0.568153699 |
| CNAG_04914 | - | small nuclear ribonucleoprotein G | 4224.083825 | 4055.627945 | 4097.456948 | 6502.368889 | 6264.182188 | 6281.075965 | 6.18E-25 | 0.606621601 | 4.56E-26 | 10.56015338 | 1.522689307 |
| CNAG_05755 | - | protein MAK16 | 2669.014767 | 2445.643208 | 2864.242098 | 1460.214827 | 1550.542276 | 1532.305016 | 6.26E-25 | -0.828513128 | 4.63E-26 | -10.55880225 | 0.563109296 |
| CNAG_03532 | - | 3-beta hydroxysteroid dehydrogenase/isomerase | 14.82350581 | 17.61249364 | 9.922835446 | 127.088968 | 178.2488589 | 145.8208505 | 6.63E-25 | 3.476875534 | 4.90E-26 | 10.5532987 | 11.13381054 |
| CNAG_07836 | - | NAD binding dehydrogenase | 31.13576883 | 14.39758657 | 16.33212448 | 187.0822739 | 164.7240965 | 169.1422417 | 6.91E-25 | 3.075022366 | 5.12E-26 | 10.54926724 | 8.427018931 |

| | | | | | | | | | | | | | |
|---|---|---|---|---|---|---|---|---|---|---|---|---|---|
| CNAG_06060 | - | hypothetical protein | 1124.983279 | 1132.580376 | 932.3228091 | 2126.357436 | 1974.782579 | 1975.781274 | 8.53E-25 | 0.915270606 | 6.33E-26 | 10.52925259 | 1.88592278 |
| CNAG_04616 | - | hypothetical protein | 115.1799742 | 127.4549009 | 122.9229683 | 344.790798 | 409.7094333 | 384.6580491 | 9.03E-25 | 1.628149203 | 6.72E-26 | 10.52369066 | 3.091161868 |
| CNAG_03812 | - | histidinol-phosphate transaminase | 4375.328013 | 4780.370753 | 4871.373105 | 3006.851253 | 3007.464024 | 2807.878302 | 9.74E-25 | -0.684273568 | 7.26E-26 | -10.51640941 | 0.6223191 |
| CNAG_04030 | - | hypothetical protein, variant | 863.4257546 | 761.5531522 | 856.0888316 | 392.3171893 | 388.5625679 | 378.9406708 | 1.00E-24 | -1.114542432 | 7.49E-26 | -10.51339923 | 0.46183761 |
| CNAG_02938 | - | cytochrome c oxidase subunit 6a | 7780.380878 | 7488.029945 | 8307.474309 | 4993.802821 | 4754.952203 | 5171.34366 | 1.07E-24 | -0.675617854 | 7.96E-26 | -10.50770714 | 0.626064041 |
| CNAG_06161 | VRK1 | hypothetical protein, variant | 573.9248083 | 659.5847014 | 594.2236449 | 1147.028228 | 1196.828652 | 1213.019483 | 1.10E-24 | 0.946890927 | 8.21E-26 | 10.50480969 | 1.927713868 |
| CNAG_00854 | ERG2 | C-8 sterol isomerase | 1657.730356 | 1589.035705 | 1762.60035 | 911.5385336 | 845.7305368 | 963.8874693 | 1.11E-24 | -0.896513421 | 8.31E-26 | -10.50362822 | 0.537183382 |
| CNAG_05653 | MLS1 | malate synthase A | 311.2484459 | 322.4293363 | 353.2036521 | 724.5805407 | 717.2458197 | 784.3716923 | 1.13E-24 | 1.159034675 | 8.52E-26 | 10.50129219 | 2.233079595 |
| CNAG_04622 | - | endoplasmic reticulum protein | 507.2899759 | 449.5629897 | 439.6207135 | 922.8271497 | 1077.990036 | 1009.835033 | 1.17E-24 | 1.092796377 | 8.78E-26 | 10.49842373 | 2.132870506 |
| CNAG_02812 | ARG3 | Ornithine carbamoyltransferase | 3026.498263 | 3171.768847 | 3111.76258 | 1995.847557 | 2012.073461 | 1904.193614 | 1.24E-24 | -0.670183528 | 9.31E-26 | -10.49287601 | 0.628426739 |
| CNAG_06944 | - | hypothetical protein | 168.8750517 | 213.5908692 | 176.2403175 | 503.799775 | 500.0412601 | 493.5493834 | 1.32E-24 | 1.410956606 | 9.94E-26 | 10.48674334 | 2.659134233 |
| CNAG_01800 | - | hypothetical protein | 1261.97896 | 1311.030071 | 1562.004348 | 692.7718792 | 687.8940414 | 684.2063913 | 1.36E-24 | -1.018134916 | 1.03E-25 | -10.48372935 | 0.493754254 |
| CNAG_04108 | PKP2 | pyruvate dehydrogenase kinase | 780.6371301 | 854.0266092 | 764.4331724 | 328.3658188 | 327.2441175 | 402.1874379 | 1.45E-24 | -1.196415517 | 1.10E-25 | -10.47728286 | 0.4363581 |
| CNAG_03975 | - | ribosome biogenesis protein ERB1 | 2966.898855 | 3051.094474 | 3133.049629 | 1703.039614 | 1978.45767 | 1782.094145 | 1.52E-24 | -0.759442138 | 1.15E-25 | -10.47262632 | 0.590724708 |
| CNAG_02378 | - | H/ACA ribonucleoprotein complex subunit 2 | 5952.719405 | 5473.219454 | 5790.676009 | 3258.069697 | 3605.204698 | 3703.716726 | 1.53E-24 | -0.719902946 | 1.16E-25 | -10.47176159 | 0.607138285 |
| CNAG_04806 | - | golgi phosphoprotein 3 | 4354.88288 | 4141.832268 | 4380.053945 | 6681.340701 | 6563.503808 | 6560.830475 | 1.54E-24 | 0.605717612 | 1.17E-25 | 10.4711999 | 1.521735493 |
| CNAG_12946 | - | unspecified product | 23.0066487 | 29.45509913 | 14.19967518 | 161.6226663 | 163.8541152 | 190.0038848 | 1.64E-24 | 2.947084596 | 1.25E-25 | 10.46523756 | 7.711890653 |
| CNAG_06814 | SXI1alpha | Alpha cell-type homeodomain transcription factor | 144.4157968 | 200.6743558 | 166.6486765 | 545.434304 | 487.1018767 | 458.1025053 | 1.70E-24 | 1.532605891 | 1.30E-25 | 10.46125004 | 2.893079344 |
| CNAG_06798 | - | U3 small nucleolar RNA-associated protein 18 | 1283.84995 | 1100.986027 | 1329.563042 | 645.3599712 | 565.3565095 | 576.5498055 | 1.72E-24 | -1.071429355 | 1.32E-25 | -10.46018958 | 0.475847318 |
| CNAG_04222 | - | protein SDA1 | 4829.400905 | 4507.850349 | 4874.582881 | 2914.946327 | 2895.283325 | 3108.663647 | 1.81E-24 | -0.687804333 | 1.38E-25 | -10.45542304 | 0.620797936 |
| CNAG_02853 | - | amidophosphoribosyltransferase | 5254.54588 | 5179.075293 | 5266.010673 | 3393.639366 | 3518.984733 | 3537.180658 | 1.87E-24 | -0.602780504 | 1.44E-25 | -10.4519156 | 0.658483637 |
| CNAG_01179 | - | DNA-directed RNA polymerase I and III subunit RPAC1 | 4653.087031 | 4317.157464 | 4991.843282 | 2797.407607 | 2794.358457 | 2899.984767 | 1.95E-24 | -0.733247745 | 1.49E-25 | -10.44813646 | 0.601548204 |
| CNAG_01185 | - | Hsp70-like protein | 5028.787491 | 4192.152452 | 4278.476333 | 2632.399877 | 2594.225946 | 2553.667979 | 2.03E-24 | -0.811126828 | 1.57E-25 | -10.44368343 | 0.569936531 |
| CNAG_02684 | - | hypothetical protein | 1096.93289 | 1063.632026 | 1014.393161 | 1868.48679 | 1999.049697 | 1793.786004 | 2.03E-24 | 0.819503595 | 1.56E-25 | 10.44377812 | 1.764798653 |
| CNAG_06244 | - | hypothetical protein | 2013.570219 | 1651.551748 | 1760.528607 | 931.7184365 | 899.2068228 | 986.1199746 | 2.19E-24 | -0.961682506 | 1.69E-25 | -10.43632849 | 0.513457757 |
| CNAG_04662 | CTF4 | chromosome transmission fidelity protein 4 | 2558.894356 | 2523.515655 | 2332.513187 | 3859.175592 | 4108.745638 | 4061.801836 | 2.27E-24 | 0.683024538 | 1.76E-25 | 10.43283395 | 1.605502084 |
| CNAG_04916 | FZC16 | hypothetical protein | 20.67519466 | 28.37760496 | 23.78546821 | 164.0853953 | 162.1512985 | 170.1996312 | 2.41E-24 | 2.78999625 | 1.86E-25 | 10.4271835 | 6.916279872 |
| CNAG_04601 | - | glycine hydroxymethyltransferase | 16048.73966 | 14841.82273 | 16031.05509 | 23124.91545 | 24411.65429 | 24627.54794 | 2.53E-24 | 0.605348335 | 1.97E-25 | 10.4221027 | 1.521346034 |
| CNAG_00016 | - | U6 snRNA-associated Sm-like protein LSm5 | 2536.079513 | 2772.016454 | 2749.137883 | 1590.256645 | 1638.540607 | 1698.779619 | 2.69E-24 | -0.724700382 | 2.09E-25 | -10.41637978 | 0.605122703 |
| CNAG_00293 | RAS1 | Ras-like protein | 467.6892811 | 547.5405325 | 440.6968771 | 1045.298309 | 972.4830781 | 1085.262515 | 2.71E-24 | 1.078857122 | 2.11E-25 | 10.41530363 | 2.112362041 |
| CNAG_00580 | - | translation initiation factor IF-2 | 4298.135451 | 4243.85252 | 4328.585466 | 2629.292325 | 2857.40358 | 2815.589563 | 3.19E-24 | -0.64791852 | 2.49E-25 | -10.39963679 | 0.638200427 |
| CNAG_02742 | DCR1 | dicer-like protein 1 | 311.2504793 | 328.8877037 | 336.1582041 | 782.4191867 | 669.6734443 | 793.2380694 | 3.73E-24 | 1.187755229 | 2.91E-25 | 10.3847499 | 2.277980234 |
| CNAG_03597 | - | hypothetical protein | 58.0361672 | 81.12652261 | 45.13700687 | 281.6162736 | 260.339501 | 280.1745976 | 3.76E-24 | 2.143934463 | 2.94E-25 | 10.3837138 | 4.419657162 |
| CNAG_00862 | - | hypothetical protein | 626.4766582 | 682.7113615 | 597.0179758 | 273.9396776 | 290.2278087 | 277.9271657 | 4.82E-24 | -1.196173536 | 3.77E-25 | -10.35989713 | 0.436431296 |
| CNAG_02735 | - | hypothetical protein | 13.62802171 | 6.827437705 | 10.98022394 | 136.3709115 | 124.1977417 | 133.5250815 | 5.07E-24 | 3.744342499 | 3.97E-25 | 10.35502274 | 13.40168501 |
| CNAG_05234 | TIM54 | mitochondrial import inner membrane translocase subunit tim54 | 1241.920824 | 1123.603644 | 1291.212349 | 639.1056248 | 637.8915623 | 632.010518 | 5.14E-24 | -0.953579564 | 4.03E-25 | -10.35349103 | 0.516349723 |
| CNAG_04824 | - | mitotic spindle assembly checkpoint protein MAD1 | 657.8993758 | 560.5422248 | 640.0632435 | 1207.031512 | 1213.192912 | 1249.649142 | 6.25E-24 | 0.966571999 | 4.92E-25 | 10.33454321 | 1.954191702 |
| CNAG_01715 | - | ribosome biogenesis protein BMS1 | 7178.827347 | 6498.93694 | 6617.211998 | 4059.77268 | 4232.273539 | 4469.716447 | 7.00E-24 | -0.684980684 | 5.52E-25 | -10.32354166 | 0.622014155 |
| CNAG_02140 | - | NADH dehydrogenase (ubiquinone) 1 alpha subcomplex 12 | 5349.075423 | 5060.571073 | 5195.628365 | 3416.234705 | 3505.189114 | 3384.006518 | 7.92E-24 | -0.614126645 | 6.24E-25 | -10.31160453 | 0.653325273 |
| CNAG_01914 | - | mitochondrial protein | 1151.926907 | 1058.909976 | 1137.662463 | 558.1296601 | 604.2965912 | 487.7736235 | 8.16E-24 | -1.036741381 | 6.45E-25 | -10.3085615 | 0.487427184 |
| CNAG_06734 | - | DNA-directed RNA polymerase I, II, and III subunit rpabc5 | 1370.204286 | 1157.005228 | 1289.110713 | 630.488994 | 688.820414 | 603.1923616 | 8.49E-24 | -1.006378912 | 6.71E-25 | -10.3046309 | 0.497794118 |
| CNAG_05366 | - | translation initiation factor 2A | 12691.93487 | 12081.1297 | 12579.7476 | 8033.790475 | 8415.744174 | 8672.764426 | 8.82E-24 | -0.587797752 | 6.99E-25 | -10.30074205 | 0.665357789 |
| CNAG_02007 | ADK1 | adenylate kinase 1 | 8879.084682 | 7753.105982 | 8517.601419 | 5312.897376 | 5388.991393 | 5219.124099 | 9.94E-24 | -0.675722303 | 7.89E-25 | -10.28915571 | 0.626018717 |
| CNAG_00303 | - | high-affinity nicotinic acid transporter | 56.82403793 | 62.83179709 | 48.3159452 | 226.8019292 | 234.504314 | 255.691265 | 1.02E-23 | 2.092056995 | 8.11E-25 | 10.28640678 | 4.263555377 |
| CNAG_01776 | - | 2,4-dihydroxyhept-2-ene-1,7-dioic acid aldolase | 71.92515716 | 45.64163165 | 46.19094754 | 245.6577262 | 244.8861124 | 257.9502182 | 1.11E-23 | 2.186929789 | 8.80E-25 | 10.27858169 | 4.553354515 |
| CNAG_04878 | FZC1 | hypothetical protein, variant | 716.3247951 | 756.5596072 | 753.0913973 | 1336.167773 | 1358.950688 | 1324.127103 | 1.25E-23 | 0.837577808 | 9.94E-25 | 10.26682534 | 1.787047285 |
| CNAG_04771 | - | rab family protein | 1583.879527 | 1356.746688 | 1594.506341 | 2800.238332 | 2618.198043 | 2717.358857 | 1.26E-23 | 0.827767455 | 1.01E-24 | 10.26554439 | 1.774936553 |
| CNAG_07356 | - | succinate dehydrogenase, cytochrome b556 subunit | 1726.446562 | 1523.319538 | 1585.635397 | 918.6027824 | 821.5836368 | 888.4288172 | 1.32E-23 | -0.895440143 | 1.06E-24 | -10.26097407 | 0.537583162 |
| CNAG_07510 | - | farnesyl-diphosphate farnesyltransferase | 7911.186499 | 7774.614867 | 7935.344505 | 5016.341961 | 5452.873616 | 5373.3819 | 1.61E-23 | -0.591748419 | 1.29E-24 | -10.24166278 | 0.66353827 |
| CNAG_01287 | - | NADH-ubiquinone oxidoreductase 51 kDa subunit, mitochondrial | 8068.737963 | 7851.079651 | 7455.472224 | 4783.597972 | 5063.807425 | 5311.202656 | 1.74E-23 | -0.640177492 | 1.39E-24 | -10.23420702 | 0.641634005 |
| CNAG_04732 | PCD1 | hypothetical protein | 862.2257567 | 853.5275638 | 740.3450465 | 1484.005938 | 1525.421654 | 1560.520033 | 1.76E-23 | 0.881052994 | 1.41E-24 | 10.23295827 | 1.841719045 |
| CNAG_02814 | - | glycerol-3-phosphate dehydrogenase | 7837.588039 | 8174.268429 | 7308.317812 | 4949.424245 | 4991.337821 | 5290.113319 | 2.04E-23 | -0.629446467 | 1.64E-24 | -10.21830621 | 0.646424388 |
| CNAG_05180 | - | hypothetical protein | 187.6095451 | 249.1611015 | 258.3057632 | 601.8380534 | 593.1670687 | 592.3585254 | 2.04E-23 | 1.349245811 | 1.64E-24 | 10.21843275 | 2.547789014 |
| CNAG_03891 | - | hsp60-like protein | 22951.73291 | 21175.73533 | 23553.49304 | 12540.70611 | 14496.78321 | 14639.91031 | 2.27E-23 | -0.715304997 | 1.83E-24 | -10.20759804 | 0.609076355 |
| CNAG_02174 | - | signal recognition particle protein SRP54, variant 2 | 5666.726447 | 5455.981862 | 5698.973439 | 3439.489842 | 3822.630897 | 3650.439129 | 2.62E-23 | -0.639868834 | 2.12E-24 | -10.19357066 | 0.641771294 |
| CNAG_02156 | - | hypothetical protein | 437.3086926 | 489.3698245 | 366.0710355 | 999.5231497 | 898.3188227 | 982.0379596 | 2.64E-23 | 1.141571614 | 2.14E-24 | 10.19272338 | 2.206212283 |
| CNAG_07561 | - | 6-phosphogluconate dehydrogenase, decarboxylating 1 | 72679.45768 | 73413.95664 | 73880.57034 | 49850.60231 | 50824.42833 | 53821.87312 | 3.11E-23 | -0.524911149 | 2.52E-24 | -10.17650432 | 0.695001912 |
| CNAG_04734 | - | mitochondrial ornithine carrier protein | 1220.792497 | 1391.126731 | 1292.712923 | 2182.063514 | 2331.082893 | 2171.225984 | 3.12E-23 | 0.76131594 | 2.54E-24 | 10.17600624 | 1.69503603 |
| CNAG_00527 | - | translation initiation factor eIF-2B subunit epsilon | 6017.006253 | 5605.756418 | 5862.150214 | 3619.29214 | 3829.493039 | 3909.083094 | 3.13E-23 | -0.638039988 | 2.55E-24 | -10.17566625 | 0.642585358 |
| CNAG_04723 | - | hypothetical protein | 1874.622143 | 1909.375751 | 1850.45301 | 2900.31821 | 3015.835066 | 3078.189075 | 3.28E-23 | 0.659771506 | 2.68E-24 | 10.17081453 | 1.57983239 |
| CNAG_06767 | - | F-type H -transporting ATPase subunit delta | 6297.413225 | 6716.523771 | 7149.283218 | 4452.988729 | 3813.858179 | 3901.400231 | 3.28E-23 | -0.743776054 | 2.67E-24 | -10.1708838 | 0.597174284 |
| CNAG_01641 | - | DNA polymerase phi subunit | 9621.829483 | 8622.591857 | 9380.368743 | 5837.449653 | 6139.445806 | 5971.789751 | 3.37E-23 | -0.63799188 | 2.75E-24 | -10.16813163 | 0.642606786 |
| CNAG_03983 | - | oxidoreductase | 11074.18144 | 12337.79548 | 10576.19204 | 17787.55021 | 17240.26563 | 17806.73072 | 3.56E-23 | 0.621927384 | 2.92E-24 | 10.1624775 | 1.538929759 |
| CNAG_12075 | - | unspecified product | 23.01453035 | 31.6085278 | 26.98413922 | 151.1803866 | 199.6279614 | 169.1933103 | 3.62E-23 | 2.682823505 | 2.96E-24 | 10.16089683 | 6.421113521 |
| CNAG_06287 | GPX2 | Glutathione peroxidase | 2593.242419 | 2675.07906 | 2664.910963 | 1725.032365 | 1599.708212 | 1659.937795 | 4.18E-23 | -0.685726233 | 3.43E-24 | -10.14656655 | 0.621692796 |
| CNAG_05434 | - | NADH dehydrogenase (ubiquinone) 1 alpha subcomplex 2 | 4077.380413 | 3901.237352 | 3958.537893 | 2602.152351 | 2651.252394 | 2439.296564 | 4.22E-23 | -0.649435619 | 3.47E-24 | -10.14549397 | 0.637529666 |
| CNAG_03961 | - | hypothetical protein | 2331.57906 | 2191.363858 | 2435.572779 | 1395.715847 | 1403.058892 | 1397.958079 | 4.80E-23 | -0.74540561 | 3.95E-24 | -10.13291557 | 0.596500144 |
| CNAG_00972 | - | hypothetical protein | 2215.913043 | 2180.518234 | 2053.864586 | 1238.440971 | 1294.373134 | 1345.751026 | 5.75E-23 | -0.74911613 | 4.74E-24 | -10.11505006 | 0.594967954 |
| CNAG_05514 | - | hypothetical protein | 16.06577054 | 52.07850601 | 45.11684739 | 225.3762518 | 269.4525933 | 235.7787336 | 6.46E-23 | 2.676593169 | 5.34E-24 | 10.10341203 | 6.393443476 |
| CNAG_00249 | - | hypothetical protein | 279.6642904 | 259.9511994 | 283.8983994 | 602.3459394 | 663.958171 | 613.5303246 | 7.26E-23 | 1.17810529 | 6.00E-24 | 10.09182395 | 2.262794062 |
| CNAG_05886 | MMS2 | ubiquitin-conjugating enzyme E2 | 2826.802173 | 2874.422895 | 2894.205433 | 1883.019863 | 1831.778385 | 1835.336052 | 7.32E-23 | -0.64628167 | 6.06E-24 | -10.09096893 | 0.638924926 |
| CNAG_04365 | - | ribosome biogenesis protein MAK21 | 6777.146413 | 6173.556946 | 6724.869055 | 3605.152483 | 3984.705693 | 4287.645907 | 7.44E-23 | -0.743922932 | 6.17E-24 | -10.08915311 | 0.59711349 |
| CNAG_02259 | - | tRNA-dihydrouridine synthase 2 | 1104.024807 | 975.9772492 | 1053.436177 | 509.0509831 | 550.7848356 | 549.8442272 | 7.96E-23 | -0.977621185 | 6.61E-24 | -10.08244751 | 0.507816373 |
| CNAG_07664 | - | mitochondrial protein | 1230.439402 | 1286.237688 | 1354.162096 | 729.4594413 | 732.8234628 | 681.969314 | 8.08E-23 | -0.868602412 | 6.72E-24 | -10.08072501 | 0.547677147 |
| CNAG_04553 | - | hypothetical protein | 49.7995104 | 30.56498753 | 39.78298743 | 253.4564427 | 184.2642107 | 208.0837097 | 8.29E-23 | 2.434341117 | 6.90E-24 | 10.07812314 | 5.40517421 |
| CNAG_05543 | - | hypothetical protein | 67.34764944 | 78.98729404 | 79.20442846 | 282.2663087 | 287.216669 | 250.4551818 | 8.51E-23 | 1.855322163 | 7.10E-24 | 10.07541361 | 3.618325416 |
| CNAG_02285 | - | nucleoside diphosphate kinase | 22912.15286 | 21884.70633 | 24555.97145 | 15094.46394 | 15932.9878 | 15168.36266 | 8.68E-23 | -0.601902611 | 7.25E-24 | -10.07329777 | 0.658884452 |
| CNAG_00447 | - | T-complex protein 1 subunit beta | 11302.31362 | 10714.9508 | 11279.73659 | 7134.858529 | 7709.334344 | 7589.263384 | 9.29E-23 | -0.585339731 | 7.77E-24 | -10.06646868 | 0.666492372 |
| CNAG_03339 | - | biotin transporter, variant | 141.9898263 | 164.046418 | 137.8537262 | 411.1771234 | 393.8700464 | 425.738775 | 9.64E-23 | 1.460814123 | 8.08E-24 | 10.06266952 | 2.752636531 |
| CNAG_04695 | - | hypothetical protein | 13.65357802 | 18.68537032 | 21.63944681 | 139.1288765 | 142.3579316 | 147.9787524 | 9.90E-23 | 3.026222892 | 8.31E-24 | 10.05993327 | 8.146740146 |
| CNAG_03592 | THI20 | phosphomethylpyrimidine kinase | 4905.069126 | 4012.261746 | 4531.17542 | 2504.690863 | 2647.723232 | 2673.532425 | 1.02E-22 | -0.797631165 | 8.54E-24 | -10.05717877 | 0.575293006 |
| CNAG_02839 | - | hypothetical protein | 8.953890152 | 11.13908448 | 10.9780747 | 102.0195301 | 137.4341864 | 143.3244483 | 1.03E-22 | 3.704639115 | 8.65E-24 | 10.05592423 | 13.03789553 |
| CNAG_05336 | - | glucose transporter, variant 2 | 824.8421917 | 771.1688914 | 744.1834512 | 349.4560435 | 402.4405395 | 320.1534106 | 1.06E-22 | -1.142660919 | 8.97E-24 | -10.05238495 | 0.452923431 |
| CNAG_04860 | XPT1 | xylosylphosphotransferase | 350.9906062 | 461.3404955 | 325.5512754 | 918.6048158 | 850.9489177 | 897.663238 | 1.08E-22 | 1.216374269 | 9.15E-24 | 10.05034681 | 2.323620187 |
| CNAG_04913 | - | syntaxin 5 | 1652.734974 | 1517.237193 | 1623.295156 | 2561.020462 | 2690.692433 | 2699.614718 | 1.13E-22 | 0.71469472 | 9.52E-24 | 10.04649197 | 1.64113591 |
| CNAG_05101 | - | U3 small nucleolar RNA-associated protein 12 | 6832.088278 | 6173.587869 | 7044.792986 | 3996.727719 | 4269.384831 | 4305.432139 | 1.13E-22 | -0.689388311 | 9.54E-24 | -10.04627738 | 0.620116718 |
| CNAG_06787 | - | metacaspase | 46.34349915 | 52.07260759 | 52.56055868 | 195.4294422 | 233.8019717 | 239.0278869 | 1.22E-22 | 2.137742675 | 1.03E-23 | 10.03866151 | 4.400729436 |

| | | | | | | | | | | | | | |
|---|---|---|---|---|---|---|---|---|---|---|---|---|---|
| CNAG_07303 | - | hypothetical protein, variant | 21.85962166 | 27.31596244 | 36.5633288 | 164.5624693 | 165.453324 | 211.0158336 | 1.29E-22 | 2.66980448 | 1.09E-23 | 10.0327871 | 6.363429412 |
| CNAG_01207 | - | hypothetical protein | 347.5130055 | 465.6590544 | 418.2683033 | 899.7644802 | 902.7655161 | 877.7607157 | 1.30E-22 | 1.108255536 | 1.10E-23 | 10.03198983 | 2.155848109 |
| CNAG_00684 | - | Ribulose-phosphate 3-epimerase | 3673.442313 | 3698.677642 | 3949.959371 | 2362.375562 | 2363.099183 | 2515.830545 | 1.41E-22 | -0.660045031 | 1.20E-23 | -10.02377814 | 0.632858543 |
| CNAG_04902 | - | SEL1 protein | 1065.491103 | 1212.27934 | 1160.464937 | 1922.889168 | 1950.593709 | 2029.040722 | 1.68E-22 | 0.765102168 | 1.43E-23 | 10.00601819 | 1.699490347 |
| CNAG_00590 | - | para-aminobenzoate synthetase | 1842.327134 | 1839.00083 | 1967.394659 | 1075.002381 | 1060.570048 | 1172.56179 | 1.69E-22 | -0.787433616 | 1.44E-23 | -10.00544849 | 0.579373813 |
| CNAG_01113 | - | hypothetical protein | 1662.470946 | 1603.067051 | 1839.376377 | 993.3312198 | 958.7988918 | 940.6147662 | 1.77E-22 | -0.835613811 | 1.52E-23 | -10.00055335 | 0.560344583 |
| CNAG_06566 | - | hypothetical protein | 2633.784403 | 2162.303842 | 2525.139423 | 1401.1407 | 1368.516647 | 1287.011693 | 1.82E-22 | -0.868045463 | 1.56E-23 | -9.997890104 | 0.547888618 |
| CNAG_01271 | - | pre-mRNA-splicing factor isy1 | 2527.909145 | 2718.163931 | 2884.531772 | 1503.012508 | 1645.446219 | 1688.791724 | 1.83E-22 | -0.764321641 | 1.57E-23 | -9.997346365 | 0.588730126 |
| CNAG_01877 | GUA1 | GMP synthase [glutamine-hydrolyzing] | 7706.790958 | 7157.281999 | 8136.842948 | 4726.79538 | 5033.632222 | 4991.510353 | 1.93E-22 | -0.656608786 | 1.65E-23 | -9.991934199 | 0.634367697 |
| CNAG_00259 | - | bud site selection protein | 2178.571685 | 2015.743696 | 2214.828873 | 1282.861215 | 1290.9253 | 1291.3802 | 2.10E-22 | -0.745574105 | 1.80E-23 | -9.983418904 | 0.596430481 |
| CNAG_00976 | - | carbamoyl-phosphate synthase, small subunit | 2125.987034 | 2049.088702 | 1983.469912 | 1299.286065 | 1213.275285 | 1198.151388 | 2.46E-22 | -0.746140439 | 2.11E-23 | -9.967522947 | 0.596196397 |
| CNAG_00538 | - | hypothetical protein | 1860.916447 | 1777.58871 | 1978.022982 | 1100.767047 | 902.6231116 | 1050.50085 | 2.59E-22 | -0.89484263 | 2.23E-23 | -9.96201231 | 0.537805856 |
| CNAG_01267 | - | hypothetical protein | 772.6463361 | 925.1248016 | 798.5838349 | 404.7840641 | 411.8802566 | 396.6884715 | 2.62E-22 | -1.05729307 | 2.27E-23 | -9.960641186 | 0.480532839 |
| CNAG_02762 | - | F-type H -transporting ATPase subunit G | 12458.50644 | 12617.68784 | 13461.68064 | 8668.153855 | 9078.228423 | 8743.841352 | 2.92E-22 | -0.556176089 | 2.53E-23 | -9.949838707 | 0.680102411 |
| CNAG_00143 | - | phosphoribosylaminoimidazolesuccinocarboxamide synthase | 3832.273357 | 4237.306252 | 3893.493609 | 2427.717285 | 2695.265663 | 2438.182793 | 2.93E-22 | -0.676910266 | 2.53E-23 | -9.949550245 | 0.625503445 |
| CNAG_07488 | - | hypothetical protein | 1492.043394 | 1571.76262 | 1599.482907 | 914.7188866 | 879.4370441 | 936.1220791 | 2.97E-22 | -0.787578506 | 2.57E-23 | -9.948081723 | 0.57931563 |
| CNAG_05219 | - | hypothetical protein | 2264.988056 | 2170.861113 | 2155.166591 | 1371.625976 | 1361.667334 | 1371.297132 | 3.04E-22 | -0.698763605 | 2.64E-23 | -9.945552012 | 0.61609998 |
| CNAG_06007 | - | U3 small nucleolar RNA-associated protein 23 | 1593.520349 | 1526.552367 | 1621.87675 | 910.0244966 | 963.1757846 | 868.4627733 | 3.07E-22 | -0.805789917 | 2.67E-23 | -9.944497844 | 0.572048781 |
| CNAG_06742 | - | pre-60S factor REI1 | 3700.278628 | 3631.891224 | 3809.21681 | 2443.375116 | 2452.844633 | 2465.886947 | 3.20E-22 | -0.613346084 | 2.78E-23 | -9.940229496 | 0.653678847 |
| CNAG_07679 | DST1 | transcription elongation factor S-II | 2232.354347 | 2140.720201 | 2365.188161 | 1381.733458 | 1373.738248 | 1353.554683 | 3.27E-22 | -0.729559262 | 2.85E-23 | -9.937755691 | 0.603088127 |
| CNAG_12573 | - | unspecified product | 4844.810472 | 5838.328529 | 6305.621582 | 3161.61736 | 2572.50528 | 3180.893095 | 3.35E-22 | -0.94473661 | 2.93E-23 | -9.935163147 | 0.519524392 |
| CNAG_00386 | - | translation initiation factor 4G | 4399.295048 | 3726.658359 | 3478.673231 | 2060.243273 | 2273.380492 | 2160.674247 | 3.41E-22 | -0.8535156 | 2.98E-23 | -9.933455996 | 0.553434467 |
| CNAG_03748 | - | large subunit ribosomal protein L19 | 10881.86363 | 10098.65417 | 10122.67059 | 6927.88912 | 7048.586927 | 7147.415149 | 3.84E-22 | -0.573766911 | 3.37E-23 | -9.921250809 | 0.671860253 |
| CNAG_12265 | - | unspecified product | 116938.3664 | 112552.7826 | 113379.8832 | 77916.53126 | 84302.21449 | 78893.80299 | 4.14E-22 | -0.523517707 | 3.63E-23 | -9.913575645 | 0.695673511 |
| CNAG_04787 | - | membrane protein | 130.3491833 | 164.0406561 | 121.8779291 | 424.5757222 | 438.1240639 | 368.2199457 | 4.15E-22 | 1.552988131 | 3.64E-23 | 9.913386333 | 2.934242549 |
| CNAG_05303 | ICL1 | isocitrate lyase | 108.0993207 | 69.34689135 | 98.39834495 | 308.4298866 | 345.0324918 | 322.4852268 | 4.33E-22 | 1.809255956 | 3.81E-23 | 9.90886793 | 3.504614977 |
| CNAG_02193 | - | kinetochore protein Spc25, fungi type | 962.770075 | 1145.478207 | 1082.619054 | 1829.514963 | 1860.905325 | 1941.31846 | 4.55E-22 | 0.805497767 | 4.01E-23 | 9.903805778 | 1.747748706 |
| CNAG_04669 | - | mitochondrial matrix protein import protein | 965.0619869 | 1026.9942 | 1100.721785 | 1781.310608 | 1809.180296 | 1761.552759 | 4.57E-22 | 0.776578071 | 4.03E-23 | 9.903231329 | 1.713062835 |
| CNAG_01502 | - | major facilitator superfamily transporter | 1609.59878 | 1417.704233 | 1450.187474 | 657.592477 | 838.9339413 | 787.418115 | 5.23E-22 | -0.987845293 | 4.62E-23 | -9.889620678 | 0.504230296 |
| CNAG_01244 | - | hypothetical protein, variant | 5.425742624 | 8.981320513 | 8.842588142 | 87.69593416 | 143.667782 | 145.4488398 | 5.50E-22 | 4.143572985 | 4.87E-23 | 9.884392246 | 17.6741997 |
| CNAG_04919 | - | hypothetical protein | 72.05629684 | 109.1169666 | 93.07006065 | 324.2766051 | 315.5697169 | 299.2477434 | 5.64E-22 | 1.761307987 | 5.00E-23 | 9.881709834 | 3.390053372 |
| CNAG_05302 | - | Amine oxidase | 636.267225 | 1046.601432 | 843.2544675 | 339.2936592 | 303.0496886 | 289.1178308 | 6.03E-22 | -1.452483258 | 5.34E-23 | -9.874968177 | 0.365391947 |
| CNAG_04629 | - | Mob1 family protein | 2848.482272 | 2729.314239 | 2755.863768 | 4215.614512 | 4594.399127 | 4238.391564 | 6.26E-22 | 0.631425352 | 5.56E-23 | 9.871003645 | 1.549094711 |
| CNAG_00546 | CHS4 | Chitin synthase | 1098.08524 | 1075.471997 | 1063.425689 | 1844.359147 | 1899.780365 | 1742.740898 | 7.30E-22 | 0.745870608 | 6.49E-23 | 9.855461169 | 1.676985957 |
| CNAG_00785 | - | ATP-dependent RNA helicase eIF4A | 43058.19 | 40251.70642 | 43271.20857 | 29170.68422 | 30100.57007 | 28982.07447 | 7.65E-22 | -0.536048762 | 6.81E-23 | -9.850661107 | 0.689657151 |
| CNAG_04037 | - | hypothetical protein | 1659.065377 | 1722.634237 | 1890.582033 | 1018.988228 | 1028.689203 | 1036.052325 | 8.16E-22 | -0.789435214 | 7.28E-23 | -9.843975399 | 0.578570546 |
| CNAG_02389 | YPK101 | AGC protein kinase | 144.3064003 | 147.9123899 | 158.0786728 | 408.0875341 | 399.1015242 | 392.5463388 | 8.72E-22 | 1.40180479 | 7.79E-23 | 9.837188582 | 2.642319257 |
| CNAG_01428 | - | Translation initiation factor 5A | 24144.08471 | 22179.93818 | 25040.15111 | 16008.2223 | 16202.95515 | 15544.71192 | 8.92E-22 | -0.595382146 | 7.99E-23 | -9.834600489 | 0.661869114 |
| CNAG_06908 | - | pyridoxal biosynthesis lyase pdxS | 5982.172248 | 6426.713379 | 6770.719384 | 4353.372384 | 4026.1193 | 3960.209717 | 8.92E-22 | -0.651259623 | 7.99E-23 | -9.834636218 | 0.636724145 |
| CNAG_04604 | - | tryptophan-tRNA ligase | 1273.240071 | 1245.703293 | 1038.959132 | 2200.148955 | 2233.634279 | 2045.786511 | 9.26E-22 | 0.85005738 | 8.30E-23 | 9.830720569 | 1.802572617 |
| CNAG_03519 | - | cytoplasmic protein, variant 2 | 7.789091275 | 16.52267266 | 14.17825594 | 151.5336163 | 119.7766206 | 122.6011246 | 9.26E-22 | 3.391309894 | 8.32E-23 | 9.830562591 | 10.49266972 |
| CNAG_05600 | IGI1 | indigoidine synthase A family protein | 2261.480974 | 2191.319764 | 2155.163864 | 1326.440084 | 1302.113034 | 1424.552685 | 9.93E-22 | -0.720596145 | 8.93E-23 | -9.823417557 | 0.606846632 |
| CNAG_01186 | - | hypothetical protein | 969.6455072 | 980.6476308 | 887.4882427 | 1700.467798 | 1686.713956 | 1573.982738 | 9.96E-22 | 0.791023963 | 8.96E-23 | 9.823011868 | 1.73030212 |
| CNAG_04437 | - | nicotinamide riboside kinase | 481.5923768 | 460.8716035 | 491.34162 | 207.8212022 | 193.5617977 | 199.1160744 | 1.02E-21 | -1.272820573 | 9.20E-23 | -9.820363487 | 0.413849875 |
| CNAG_00450 | LEU2 | 3-isopropylmalate dehydrogenase | 2472.967274 | 2552.252878 | 2737.363736 | 1425.958367 | 1676.565573 | 1472.373703 | 1.06E-21 | -0.778547576 | 9.57E-23 | -9.816365942 | 0.582953382 |
| CNAG_06392 | SGF29 | SAGA-associated factor 29 | 1078.404188 | 1069.622944 | 1006.558014 | 547.2694611 | 478.2235681 | 586.4395278 | 1.16E-21 | -0.983715586 | 1.05E-22 | -9.806912963 | 0.50567572 |
| CNAG_00512 | - | mitochondrial carrier protein | 1805.951595 | 1648.307805 | 1659.25526 | 1014.35187 | 994.1811901 | 1006.079081 | 1.22E-21 | -0.777778421 | 1.10E-22 | -9.80227981 | 0.58326426 |
| CNAG_04416 | - | major facilitator superfamily transporter | 34.65090066 | 31.6189354 | 32.31267508 | 147.8034735 | 204.6989199 | 194.5880899 | 1.24E-21 | 2.474608709 | 1.12E-22 | 9.800417955 | 5.558165177 |
| CNAG_06235 | - | hypothetical protein | 1367.953323 | 1266.843755 | 1188.964197 | 698.2893841 | 686.2030704 | 715.2326851 | 1.29E-21 | -0.879656381 | 1.17E-22 | -9.79576744 | 0.543496865 |
| CNAG_00567 | - | WD40 repeat protein | 1649.53109 | 1611.631272 | 1516.370723 | 956.7700751 | 934.6686358 | 939.4708759 | 1.35E-21 | -0.770755539 | 1.22E-22 | -9.791492261 | 0.586110448 |
| CNAG_00299 | - | DNA repair protein rad5 | 2299.664521 | 2347.880187 | 1995.537236 | 3636.691475 | 3593.730073 | 3768.686295 | 1.45E-21 | 0.712737242 | 1.32E-22 | 9.783972002 | 1.638910694 |
| CNAG_03333 | - | cytoplasmic protein | 3262.78359 | 2469.306485 | 2379.1437 | 1263.169324 | 1344.298399 | 1409.112342 | 1.58E-21 | -1.030297332 | 1.44E-22 | -9.774932838 | 0.489609233 |
| CNAG_00265 | IDI1 | isopentenyl-diphosphate delta-isomerase | 2151.707583 | 1979.114158 | 2163.646764 | 1284.441945 | 1259.864652 | 1282.492886 | 1.66E-21 | -0.733919859 | 1.51E-22 | -9.770081191 | 0.601268024 |
| CNAG_05337 | - | alpha-N-arabinofuranosidase, variant | 758.3939355 | 745.3048803 | 736.6913878 | 324.5157648 | 317.7706251 | 397.7190099 | 1.66E-21 | -1.124101266 | 1.52E-22 | -9.769657357 | 0.458787737 |
| CNAG_03412 | CHI2 | Chitinase | 932.2692599 | 934.3188931 | 824.577466 | 1582.089236 | 1629.807804 | 1541.751888 | 1.68E-21 | 0.805757042 | 1.53E-22 | 9.768656251 | 1.748062832 |
| CNAG_00940 | RRP44 | exosome complex exonuclease DIS3/RRP44 | 3500.505917 | 3420.679608 | 3295.227592 | 2090.718294 | 2282.081658 | 2222.767848 | 1.70E-21 | -0.646822062 | 1.56E-22 | -9.767205271 | 0.638685648 |
| CNAG_04716 | - | KH domain-containing protein | 4065.248699 | 3614.98915 | 3901.22675 | 5910.752976 | 6146.074573 | 6107.852886 | 1.90E-21 | 0.633544623 | 1.74E-22 | 9.756025368 | 1.551371952 |
| CNAG_08000 | - | hypothetical protein | 19.47063624 | 12.22844858 | 22.70200365 | 154.341602 | 130.9834084 | 139.1935234 | 1.95E-21 | 2.980582437 | 1.79E-22 | 9.753150336 | 7.89304753 |
| CNAG_01400 | - | 3-deoxy-7-phosphoheptulonate synthase | 6082.510721 | 6107.817148 | 6720.578109 | 3742.998883 | 4303.095665 | 3729.320223 | 2.02E-21 | -0.699086671 | 1.86E-22 | -9.749191758 | 0.615962031 |
| CNAG_06127 | - | ribosome biogenesis protein NSA2 | 3888.216493 | 3582.357894 | 4038.46731 | 2476.836234 | 2423.489318 | 2410.409874 | 2.09E-21 | -0.670604449 | 1.92E-22 | -9.745735374 | 0.628243416 |
| CNAG_02160 | - | hypothetical protein, variant | 56.77546043 | 49.91825104 | 48.30037981 | 224.0075776 | 206.8519768 | 209.2442249 | 2.50E-21 | 2.046847268 | 2.30E-22 | 9.727475448 | 4.132020087 |
| CNAG_06656 | - | Diphthine synthase | 1424.212826 | 1358.484284 | 1488.552293 | 818.9653577 | 846.7087392 | 798.5079594 | 2.56E-21 | -0.81000048 | 2.36E-22 | -9.724748174 | 0.570381668 |
| CNAG_01858 | HOB2 | hypothetical protein | 30.04240248 | 40.23586206 | 50.41537513 | 185.4536972 | 243.6418224 | 189.3135159 | 2.56E-21 | 2.346976313 | 2.37E-22 | 9.724566186 | 5.087568496 |
| CNAG_04770 | - | nucleolar essential protein 1 | 2340.521141 | 2285.406833 | 2320.743576 | 3629.666094 | 3718.853608 | 3440.215284 | 2.58E-21 | 0.619792248 | 2.39E-22 | 9.723831565 | 1.536653883 |
| CNAG_04850 | - | hypothetical protein, variant | 116.3282013 | 111.318991 | 133.5614309 | 369.2276506 | 414.9783949 | 319.4126016 | 2.61E-21 | 1.598215566 | 2.42E-22 | 9.722429129 | 3.027685945 |
| CNAG_04682 | - | DNA replication complex GINS protein psf3 | 427.9649746 | 506.5810167 | 400.1663772 | 914.4724282 | 938.9896906 | 888.9060026 | 2.63E-21 | 1.024996543 | 2.44E-22 | 9.721716407 | 2.034954509 |
| CNAG_07004 | LPD1alpha | dihydrolipoyl dehydrogenase | 16331.80845 | 15076.41588 | 15880.43303 | 10764.8824 | 11121.63055 | 10858.6908 | 2.67E-21 | -0.545955364 | 2.48E-22 | -9.719766458 | 0.684937681 |
| CNAG_00727 | - | mitochondrial protein with role in iron accumulation | 1234.298903 | 903.7947654 | 930.841651 | 477.9305435 | 433.3573041 | 435.5892493 | 2.83E-21 | -1.205706576 | 2.63E-22 | -9.713869173 | 0.433556952 |
| CNAG_04810 | - | hypothetical protein | 319.4082129 | 345.0322986 | 377.7095847 | 725.3424171 | 745.774831 | 720.1320063 | 2.93E-21 | 1.057814374 | 2.73E-22 | 9.710130269 | 2.081775324 |
| CNAG_06061 | - | translation initiation factor 3 subunit E | 1492.046803 | 1536.23248 | 1617.603731 | 912.3779954 | 912.2446195 | 935.0139928 | 2.94E-21 | -0.766710277 | 2.75E-22 | -9.709477448 | 0.587756186 |
| CNAG_04795 | - | adenine nucleotide transporter | 674.2804488 | 693.0015684 | 604.9110585 | 1170.977682 | 1276.148832 | 1431.582043 | 3.15E-21 | 0.961140863 | 2.94E-22 | 9.702453717 | 1.946848828 |
| CNAG_01261 | - | myosin I binding protein | 2568.190008 | 2115.238685 | 2275.98717 | 3883.41648 | 3956.065032 | 3996.276184 | 3.18E-21 | 0.750240107 | 2.98E-22 | 9.701264959 | 1.682072753 |
| CNAG_01354 | - | hypothetical protein | 2706.599736 | 3006.894337 | 3030.664866 | 1640.000028 | 1875.798566 | 1795.382124 | 3.19E-21 | -0.734219563 | 2.99E-22 | -9.700873164 | 0.60114313 |
| CNAG_03081 | - | ATP-dependent RNA helicase MRH4, mitochondrial | 3158.319785 | 2954.17891 | 3135.201247 | 1742.73434 | 2014.681751 | 1927.483473 | 3.45E-21 | -0.718026478 | 3.24E-22 | -9.692727265 | 0.607928484 |
| CNAG_01178 | - | nuclear envelope-endoplasmic reticulum network protein | 127.9738606 | 130.6823705 | 139.9572024 | 345.5564945 | 404.4893226 | 365.8806864 | 3.47E-21 | 1.476782754 | 3.26E-22 | 9.692162912 | 2.783273638 |
| CNAG_00678 | URE7 | urease accessory protein UreG | 3979.359454 | 3789.219436 | 4329.591041 | 2587.342976 | 2549.423581 | 2553.614996 | 3.68E-21 | -0.669383066 | 3.46E-22 | -9.685833563 | 0.628775511 |
| CNAG_03263 | TUF1 | translation elongation factor Tu | 15538.82135 | 13478.6057 | 15696.93708 | 9602.073134 | 9352.43147 | 9679.722215 | 3.81E-21 | -0.658958428 | 3.59E-22 | -9.68223792 | 0.633335377 |
| CNAG_03936 | - | NAD(P)H:quinone oxidoreductase, type IV | 1930.668937 | 1827.526628 | 2002.930522 | 3123.828099 | 2990.762634 | 3120.391522 | 3.81E-21 | 0.665507007 | 3.58E-22 | 9.682337781 | 1.586125588 |
| CNAG_04600 | - | DNA topoisomerase III | 579.7118895 | 632.6463309 | 514.2723996 | 1096.442861 | 1232.347771 | 1084.34101 | 3.98E-21 | 0.969106673 | 3.75E-22 | 9.677649339 | 1.957628043 |
| CNAG_04804 | SRE1 | sterol regulatory element-binding protein | 1046.769774 | 1063.63234 | 1142.317655 | 1779.619405 | 1978.326163 | 1831.502861 | 4.02E-21 | 0.766043072 | 3.80E-22 | 9.676442941 | 1.700599092 |
| CNAG_00859 | - | hypothetical protein | 240.0996979 | 334.213281 | 262.606598 | 653.0501075 | 643.9647741 | 693.3335385 | 4.04E-21 | 1.236626422 | 3.83E-22 | 9.675643611 | 2.356468542 |
| CNAG_03861 | - | cytochrome c oxidase subunit 5a | 4461.591458 | 4312.815689 | 4531.192485 | 3044.316904 | 2634.763353 | 2699.073177 | 4.07E-21 | -0.682715141 | 3.86E-22 | -9.6747203 | 0.622991705 |
| CNAG_07595 | - | hypothetical protein | 75.44823751 | 67.15309283 | 65.36006351 | 232.7770096 | 298.6661051 | 245.9517733 | 5.07E-21 | 1.888770068 | 4.81E-22 | 9.652153337 | 3.703193841 |
| CNAG_13142 | - | unspecified product | 417.1861318 | 371.4465288 | 402.8110573 | 164.2280761 | 135.7058543 | 145.864477 | 5.12E-21 | -1.437949167 | 4.86E-22 | -9.651171295 | 0.369091606 |
| CNAG_01187 | - | ribosome production factor 2 | 3320.503 | 2799.072115 | 3159.716262 | 1844.001018 | 1910.266563 | 1792.091752 | 5.82E-21 | -0.758520969 | 5.53E-22 | -9.637858646 | 0.59110201 |
| CNAG_01475 | SMT1 | sphingolipid C9-methyltransferase | 8727.432297 | 8520.198621 | 8439.802186 | 5611.764629 | 5830.649162 | 6143.797984 | 6.15E-21 | -0.562002895 | 5.86E-22 | -9.631953804 | 0.677361129 |
| CNAG_01174 | - | hypothetical protein | 1643.374665 | 1558.155987 | 1332.223772 | 2570.430168 | 2637.215426 | 2737.307169 | 6.55E-21 | 0.79416898 | 6.24E-22 | 9.625448866 | 1.734078224 |

| | | | | | | | | | | | | |
|---|---|---|---|---|---|---|---|---|---|---|---|---|
| CNAG_04062 | - | ribosome biogenesis protein ENP2 | 2523.010223 | 2238.79166 | 2479.321016 | 1436.164657 | 1487.603459 | 1488.977047 | 7.13E-21 | -0.730718586 | 6.81E-22 | -9.616552211 | 0.602603691 |
| CNAG_04488 | - | 3-ketoacyl-CoA reductase | 2460.230855 | 2728.932638 | 2727.813283 | 1675.950731 | 1695.504709 | 1659.93959 | 7.33E-21 | -0.669212108 | 7.01E-22 | -9.613585701 | 0.628850024 |
| CNAG_03439 | - | hypothetical protein | 5047.901344 | 5020.705765 | 5196.687694 | 3425.591825 | 3597.519066 | 3409.527777 | 7.57E-21 | -0.56456304 | 7.25E-22 | -9.610125615 | 0.676160179 |
| CNAG_12920 | - | unspecified product | 12.48978312 | 21.91162348 | 22.70530681 | 172.6155354 | 128.3039554 | 133.7523636 | 7.83E-21 | 2.962643534 | 7.51E-22 | 9.606502372 | 7.795510669 |
| CNAG_06993 | - | hypothetical protein | 2729.820502 | 2684.800816 | 2727.836182 | 1819.245847 | 1623.847741 | 1706.565273 | 8.64E-21 | -0.676246993 | 8.30E-22 | -9.596170948 | 0.625791083 |
| CNAG_02935 | - | malonic semialdehyde reductase | 1021.253686 | 997.4829 | 982.0290231 | 544.9625747 | 574.1265519 | 526.5450941 | 9.17E-21 | -0.883166178 | 8.81E-22 | -9.589997535 | 0.54217625 |
| CNAG_04552 | ITR1A | putative myo-inositol transporter | 10.09662985 | 8.97256562 | 8.835149322 | 114.4791232 | 109.5883676 | 108.1031053 | 9.28E-21 | 3.654810261 | 8.93E-22 | 9.588589634 | 12.59527102 |
| CNAG_04745 | - | hypothetical protein | 1701.79843 | 1782.225046 | 1664.895485 | 2697.241897 | 2647.456546 | 2831.718943 | 9.35E-21 | 0.652583819 | 9.01E-22 | 9.58769324 | 1.571981042 |
| CNAG_04765 | - | ARP2/3 complex 21 kDa subunit | 466.4274416 | 468.9091025 | 412.9554077 | 892.5928136 | 1004.678305 | 847.9049196 | 1.11E-20 | 1.01173746 | 1.07E-21 | 9.570003259 | 2.016337945 |
| CNAG_04332 | MRPL31 | large subunit ribosomal protein L31 | 1269.001411 | 1353.030636 | 1362.728235 | 808.1524051 | 758.6925448 | 729.6909982 | 1.22E-20 | -0.810766334 | 1.18E-21 | -9.559866793 | 0.570078962 |
| CNAG_03662 | - | hypothetical protein | 1764.104116 | 1765.740255 | 1868.229249 | 1010.369166 | 1121.03142 | 1111.516287 | 1.29E-20 | -0.750634438 | 1.25E-21 | -9.553814679 | 0.594342133 |
| CNAG_07536 | - | DNA-directed RNA polymerase I subunit RPA43 | 1051.593187 | 998.5797623 | 1210.062235 | 428.7122735 | 463.5437182 | 588.6533059 | 1.31E-20 | -1.155229544 | 1.27E-21 | -9.552553811 | 0.448994742 |
| CNAG_06727 | - | hypothetical protein, variant 2 | 213.1360947 | 232.9725336 | 208.2052802 | 495.7915636 | 516.3878707 | 502.4865001 | 1.36E-20 | 1.197324734 | 1.32E-21 | 9.548438194 | 2.293140473 |
| CNAG_05371 | - | hypothetical protein | 1813.045006 | 1691.420019 | 1811.717067 | 1011.954972 | 1025.224034 | 1117.047831 | 1.39E-20 | -0.768834949 | 1.35E-21 | -9.546102439 | 0.586891229 |
| CNAG_07450 | - | hypothetical protein | 373.0642246 | 410.1068285 | 395.3530672 | 130.5769165 | 140.0397341 | 169.1029537 | 1.57E-20 | -1.438761684 | 1.52E-21 | -9.533316291 | 0.368883794 |
| CNAG_07413 | NOG2 | nucleolar GTP-binding protein 2 | 7152.246564 | 7660.369777 | 8578.295853 | 4342.991304 | 4560.00102 | 5141.351007 | 1.74E-20 | -0.751042515 | 1.69E-21 | -9.522335513 | 0.594174043 |
| CNAG_12898 | - | unspecified product | 2289.098575 | 2110.875211 | 1893.158467 | 3427.305873 | 3531.685331 | 3521.146466 | 1.91E-20 | 0.720456563 | 1.86E-21 | 9.512558724 | 1.647703393 |
| CNAG_03624 | - | pescadillo | 6816.920987 | 6158.509479 | 6781.41673 | 4159.44357 | 4439.344676 | 4390.901835 | 1.94E-20 | -0.620860692 | 1.89E-21 | -9.510641945 | 0.650282862 |
| CNAG_07804 | - | protein mgr2 | 1108.958552 | 1105.261523 | 1235.75817 | 653.9823583 | 591.2879527 | 637.5275498 | 2.00E-20 | -0.889537515 | 1.95E-21 | -9.507448091 | 0.53978713 |
| CNAG_05662 | PTP1 | Polyol transporter protein 1 | 26.43583121 | 18.68628215 | 16.31848351 | 155.0767823 | 142.3413916 | 128.2533732 | 2.26E-20 | 2.808518566 | 2.21E-21 | 9.494560563 | 7.005648313 |
| CNAG_00622 | ARG8 | Acetylornithine aminotransferase | 3089.624489 | 3238.602597 | 3338.902594 | 2110.259191 | 2176.84762 | 2108.426883 | 2.27E-20 | -0.611212172 | 2.22E-21 | -9.493971674 | 0.654646428 |
| CNAG_05764 | - | small subunit ribosomal protein S5 | 1330.985226 | 1571.686987 | 1480.047518 | 857.9124037 | 844.0971441 | 810.7253975 | 2.31E-20 | -0.81760204 | 2.27E-21 | -9.491944041 | 0.567384231 |
| CNAG_03948 | - | NAD(P)H:quinone oxidoreductase, type IV | 6866.125454 | 7044.110133 | 7500.189704 | 4836.665213 | 4412.520006 | 4888.233219 | 2.46E-20 | -0.613979707 | 2.42E-21 | -9.485341609 | 0.653391818 |
| CNAG_02736 | - | T-complex protein 1 subunit theta | 10507.01643 | 9595.508603 | 9915.767379 | 6673.383557 | 7066.739659 | 6696.714802 | 2.48E-20 | -0.570501531 | 2.43E-21 | -9.484541271 | 0.673382657 |
| CNAG_04651 | - | hypothetical protein | 960.3149782 | 1004.339139 | 837.3865018 | 1731.653351 | 1689.302856 | 1582.85744 | 2.50E-20 | 0.821621069 | 2.46E-21 | 9.483578433 | 1.767390787 |
| CNAG_03435 | - | ribonucleoprotein-associated protein | 9746.778762 | 8585.974507 | 9471.020056 | 6253.145873 | 6011.753061 | 6092.791593 | 2.56E-20 | -0.614797102 | 2.52E-21 | -9.48087706 | 0.653021727 |
| CNAG_04842 | - | hypothetical protein | 586.6940356 | 601.4187127 | 586.7405651 | 1119.115723 | 1165.842406 | 1012.258722 | 2.57E-20 | 0.879218672 | 2.54E-21 | 9.480176244 | 1.83937887 |
| CNAG_01772 | - | hypothetical protein | 3855.455458 | 3544.623184 | 3630.076538 | 2286.867765 | 2332.056603 | 2485.84676 | 2.58E-20 | -0.650338806 | 2.55E-21 | -9.479644922 | 0.637130671 |
| CNAG_04473 | - | hypothetical protein | 1419.635484 | 1512.488363 | 1447.021813 | 881.2762695 | 851.8537672 | 897.2580882 | 2.93E-20 | -0.750653843 | 2.89E-21 | -9.466483564 | 0.594334139 |
| CNAG_00510 | - | prefoldin subunit 2 | 1342.591444 | 1394.011109 | 1500.273147 | 817.4101942 | 819.9432724 | 841.7653779 | 3.12E-20 | -0.789281957 | 3.09E-21 | -9.459613744 | 0.578632011 |
| CNAG_02485 | - | ATP synthase F1, gamma subunit | 13479.10415 | 13613.22975 | 13820.04625 | 9273.659637 | 10129.73401 | 9426.586157 | 3.15E-20 | -0.520423422 | 3.12E-21 | -9.458578731 | 0.697167189 |
| CNAG_02555 | SIP402 | Putative Zn2-Cys6 zinc-finger transcription factor | 1518.791819 | 1446.831164 | 1518.449984 | 926.461097 | 885.5094781 | 859.5582303 | 3.37E-20 | -0.762793278 | 3.35E-21 | -9.451221366 | 0.589354146 |
| CNAG_00694 | CLF1 | pre-mRNA-splicing factor CLF1 | 5713.441449 | 5437.678607 | 5532.640592 | 3665.274561 | 3756.184798 | 3921.268068 | 3.38E-20 | -0.572040929 | 3.36E-21 | -9.450917265 | 0.672664521 |
| CNAG_00268 | ILV2 | acetolactate synthase, mitochondrial | 3041.630963 | 3368.854993 | 2912.376165 | 1918.763245 | 2007.776247 | 1705.538076 | 3.66E-20 | -0.742214561 | 3.65E-21 | -9.442322098 | 0.597820982 |
| CNAG_05644 | - | 2-Nitropropane dioxygenase | 366.0522797 | 389.1796717 | 391.5805317 | 759.5652683 | 759.5071932 | 745.6934624 | 3.80E-20 | 0.9665519 | 3.79E-21 | 9.438292034 | 1.954164476 |
| CNAG_04360 | - | tRNA (guanine37-N1)-methyltransferase | 1925.285721 | 2089.978688 | 1886.439063 | 1210.499086 | 1214.166149 | 1248.05628 | 3.91E-20 | -0.698932789 | 3.90E-21 | -9.435155369 | 0.616027735 |
| CNAG_00992 | - | homocitrate synthase, mitochondrial | 3348.784673 | 3472.376926 | 3561.77051 | 2353.197928 | 1959.346701 | 2044.091463 | 4.02E-20 | -0.722773477 | 4.01E-21 | -9.432305704 | 0.605931462 |
| CNAG_04577 | - | Nucleoside-diphosphate kinase | 2271.632141 | 2212.1466 | 2360.183101 | 3544.830037 | 3508.32506 | 3437.96506 | 4.07E-20 | 0.600740796 | 4.07E-21 | 9.430814856 | 1.516495058 |
| CNAG_01361 | - | importin alpha subunit | 12724.58909 | 12081.11361 | 11629.58772 | 7980.882898 | 8643.506292 | 8361.935733 | 4.13E-20 | -0.559780857 | 4.14E-21 | -9.429051348 | 0.678405204 |
| CNAG_01548 | - | ATP-dependent RNA helicase FAL1 | 6638.316242 | 6305.020161 | 6400.742876 | 4511.358396 | 4435.892094 | 4504.118499 | 4.37E-20 | -0.539653622 | 4.38E-21 | -9.422988976 | 0.687936057 |
| CNAG_00690 | - | hypothetical protein | 3159.394173 | 2875.513857 | 2812.124369 | 1909.481346 | 1744.601957 | 1846.44392 | 4.71E-20 | -0.701347851 | 4.72E-21 | -9.415155469 | 0.614997371 |
| CNAG_03264 | - | hypothetical protein | 1737.133138 | 1677.375911 | 1684.832515 | 904.4651738 | 1075.340027 | 1007.184718 | 4.94E-20 | -0.787239739 | 4.97E-21 | -9.409916015 | 0.579451678 |
| CNAG_01666 | - | U6 snRNA-associated Sm-like protein LSm2 | 1985.969865 | 1990.918764 | 1981.323152 | 1197.233155 | 1314.287794 | 1228.095317 | 5.12E-20 | -0.687032365 | 5.15E-21 | -9.406055962 | 0.621130206 |
| CNAG_02404 | - | nuclear segregation protein Bfr1 | 9809.92114 | 9193.624608 | 9663.002311 | 6193.909852 | 6673.398376 | 6717.768267 | 6.09E-20 | -0.565292453 | 6.14E-21 | -9.387630369 | 0.675818405 |
| CNAG_05105 | LIV12 | eukaryotic translation initiation factor 3 subunit I | 13244.26778 | 12028.37253 | 13347.57423 | 8471.215761 | 8827.197612 | 8861.504227 | 6.23E-20 | -0.577798375 | 6.28E-21 | -9.385185957 | 0.669985429 |
| CNAG_05197 | - | NADH dehydrogenase (ubiquinone) 1 alpha subcomplex 8 | 4475.649063 | 4209.427245 | 4830.829333 | 2988.208379 | 2837.51099 | 2864.460552 | 7.11E-20 | -0.652825919 | 7.18E-21 | -9.3710261 | 0.636033245 |
| CNAG_06543 | - | ethanolaminephosphotransferase | 3463.307245 | 3730.962863 | 3650.329404 | 2330.489886 | 2340.691525 | 2510.256985 | 7.43E-20 | -0.609560865 | 7.51E-21 | -9.366329311 | 0.655396165 |
| CNAG_03140 | - | Sugar transporter | 3701.323199 | 3430.412306 | 3439.196305 | 2293.932477 | 2392.49715 | 2236.115967 | 8.31E-20 | -0.626570639 | 8.42E-21 | -9.354305483 | 0.647714237 |
| CNAG_01384 | - | hypothetical protein | 73.14943183 | 86.50480578 | 79.20149328 | 263.3890803 | 286.3783364 | 242.7118619 | 8.49E-20 | 1.717742618 | 8.61E-21 | 9.351914148 | 3.289213416 |
| CNAG_02235 | - | hypothetical protein | 640.7256446 | 710.8261421 | 837.8184661 | 330.8158672 | 343.7335793 | 260.2533759 | 8.80E-20 | -1.244408225 | 8.93E-21 | -9.347999522 | 0.422080995 |
| CNAG_01706 | - | hypothetical protein | 30.0163405 | 48.80739476 | 34.4448828 | 207.2240736 | 178.424678 | 168.2541825 | 9.14E-20 | 2.289865888 | 9.29E-21 | 9.343860839 | 4.890106509 |
| CNAG_03696 | - | hypothetical protein | 28.81737384 | 36.97307454 | 23.78857699 | 169.549039 | 189.1945309 | 138.39025 | 9.40E-20 | 2.496500083 | 9.57E-21 | 9.340751718 | 5.643147594 |
| CNAG_03128 | - | gamma-glutamyltransferase, variant | 1942.506599 | 1738.802195 | 1679.543947 | 1018.965936 | 1013.999333 | 1095.971245 | 9.42E-20 | -0.793189349 | 9.59E-21 | -9.340449457 | 0.577066965 |
| CNAG_04376 | - | protein KRI1 | 1186.071818 | 1239.911491 | 1234.773705 | 720.9294746 | 705.2175515 | 729.6377756 | 9.78E-20 | -0.779474632 | 9.97E-21 | -9.33637991 | 0.582578905 |
| CNAG_04720 | - | GTPase activating protein | 2563.551953 | 2555.826496 | 2197.093882 | 3778.200653 | 4045.76723 | 4125.044265 | 1.00E-19 | 0.692762952 | 1.02E-20 | 9.333843587 | 1.616376131 |
| CNAG_04993 | - | hypothetical protein | 7.775490981 | 11.13517494 | 17.3617971 | 117.3755033 | 105.0141955 | 137.7769274 | 1.01E-19 | 3.343971525 | 1.03E-20 | 9.332623031 | 10.15396668 |
| CNAG_04346 | - | dihydrodipicolinate synthase | 231.9487602 | 299.7836392 | 327.5907108 | 649.9033478 | 679.4272081 | 654.5828004 | 1.04E-19 | 1.194929442 | 1.06E-20 | 9.329504364 | 2.289336353 |
| CNAG_06659 | HEX1 | beta-hexosaminidase | 618.2255638 | 646.662327 | 539.8650475 | 1178.258831 | 1071.566189 | 1299.488103 | 1.04E-19 | 0.961999387 | 1.07E-20 | 9.329334514 | 1.94800771 |
| CNAG_03584 | - | Prp8 binding protein | 2684.346438 | 2668.659948 | 2883.502767 | 1835.579388 | 1746.384237 | 1677.725062 | 1.06E-19 | -0.662650504 | 1.09E-20 | -9.326698659 | 0.631716647 |
| CNAG_03697 | - | hormone-sensitive lipase | 366.0809082 | 417.1768902 | 404.3786053 | 786.030984 | 761.17403 | 792.2779 | 1.13E-19 | 0.963814838 | 1.16E-20 | 9.320242438 | 1.950460577 |
| CNAG_05825 | - | minichromosome maintenance protein 7 (cell division control protein 47) | 4556.26821 | 4612.337562 | 4633.606772 | 3079.25375 | 3283.564118 | 2922.20332 | 1.17E-19 | -0.587334306 | 1.20E-20 | -9.316436157 | 0.66557156 |
| CNAG_03302 | - | protein LTV1 | 2840.830143 | 2864.741187 | 3200.197184 | 1650.880272 | 1858.52004 | 1903.036999 | 1.18E-19 | -0.733938677 | 1.21E-20 | -9.315651672 | 0.601260181 |
| CNAG_00028 | - | high-affinity nicotinic acid transporter, variant | 725.7560201 | 767.8742614 | 654.661498 | 346.4005272 | 368.7789629 | 352.2663535 | 1.19E-19 | -1.024359765 | 1.23E-20 | -9.314128027 | 0.491628424 |
| CNAG_07666 | - | hypothetical protein | 21.76229714 | 15.44806565 | 13.11340389 | 121.0842809 | 145.2239093 | 118.2255332 | 1.39E-19 | 2.954213705 | 1.43E-20 | 9.297994988 | 7.750093443 |
| CNAG_00302 | - | hypothetical protein | 347.3598937 | 345.0294093 | 371.3146499 | 750.3795707 | 697.3550765 | 699.0732723 | 1.45E-19 | 0.998225491 | 1.50E-20 | 9.293140694 | 1.997541521 |
| CNAG_06036 | - | hypothetical protein | 3159.53701 | 3209.470282 | 2946.510213 | 1868.882513 | 2120.806533 | 1968.560633 | 1.72E-19 | -0.660278125 | 1.78E-20 | -9.274800395 | 0.632756301 |
| CNAG_04608 | - | signal peptidase I | 1102.742147 | 1126.083078 | 1081.544559 | 1771.885337 | 1795.320799 | 1774.898569 | 1.80E-19 | 0.675306266 | 1.86E-20 | 9.270083772 | 1.596935746 |
| CNAG_02974 | - | voltage-dependent anion channel protein 2 | 18930.27045 | 20514.16529 | 20580.28747 | 14057.85201 | 14432.14828 | 14524.41014 | 1.89E-19 | -0.495701953 | 1.96E-20 | -9.264534633 | 0.70921652 |
| CNAG_06318 | - | ribosome biogenesis protein YTM1 | 2173.958251 | 2105.151814 | 2422.724323 | 1128.559795 | 1315.072097 | 1376.832257 | 1.96E-19 | -0.826517286 | 2.04E-20 | -9.26035943 | 0.563888847 |
| CNAG_05241 | - | U6 snRNA-associated Sm-like protein LSm7 | 1673.04927 | 1709.707048 | 1720.031151 | 1064.200759 | 1094.306637 | 1048.251533 | 2.02E-19 | -0.685519959 | 2.10E-20 | -9.257281763 | 0.621781691 |
| CNAG_06419 | - | hypothetical protein | 245.7560817 | 221.1565423 | 213.543422 | 498.0053052 | 582.972916 | 505.8744428 | 2.03E-19 | 1.207278766 | 2.11E-20 | 9.256750883 | 2.309016954 |
| CNAG_12827 | - | unspecified product | 1287.425426 | 1113.934415 | 1301.888833 | 672.6121694 | 604.1952834 | 700.7853988 | 2.11E-19 | -0.921093742 | 2.20E-20 | -9.252183348 | 0.528108497 |
| CNAG_06691 | - | tuftelin-interacting protein 11 | 3683.859861 | 3458.435783 | 3561.825207 | 2293.123982 | 2398.518118 | 2433.672688 | 2.15E-19 | -0.602647902 | 2.24E-20 | -9.250247939 | 0.658544163 |
| CNAG_01529 | - | solute carrier family 25 (mitochondrial phosphate transporter), member 23/24/25/41 | 1210.456458 | 1131.141115 | 1230.472865 | 617.2674235 | 707.8463917 | 679.7007237 | 2.19E-19 | -0.849841215 | 2.29E-20 | -9.247902004 | 0.5548458 |
| CNAG_04180 | - | large subunit ribosomal protein L15 | 1621.718251 | 1743.080578 | 1836.208853 | 1093.82945 | 1016.61413 | 939.5257703 | 2.23E-19 | -0.785757455 | 2.33E-20 | -9.246141484 | 0.580047336 |
| CNAG_04444 | - | splicing factor 3B subunit 1 | 8026.874969 | 8078.465341 | 8157.188458 | 5632.846341 | 5889.354516 | 5855.164517 | 2.31E-19 | -0.496807969 | 2.42E-20 | -9.241891441 | 0.708673021 |
| CNAG_04194 | - | protein transporter SEC13 | 6265.798245 | 5853.589001 | 6146.910823 | 4153.3074 | 4297.049441 | 4164.424304 | 2.51E-19 | -0.549613305 | 2.63E-20 | -9.233079521 | 0.683203227 |
| CNAG_03602 | - | U3 small nucleolar RNA-associated protein 5 | 6866.112182 | 6964.384794 | 6850.791096 | 4705.878166 | 4920.682393 | 5008.104184 | 2.55E-19 | -0.514179131 | 2.67E-20 | -9.231439415 | 0.700191217 |
| CNAG_00984 | - | glucose and ribitol dehydrogenase | 327.5741236 | 352.5714034 | 236.0039341 | 906.6558486 | 800.2343178 | 654.7891533 | 2.73E-19 | 1.352231795 | 2.87E-20 | 9.223726612 | 2.553067701 |
| CNAG_01611 | LIV8 | infection related protein of unknown function | 129.1022753 | 142.4906899 | 111.2019567 | 339.4363673 | 370.7685753 | 343.6953762 | 2.92E-19 | 1.446627831 | 3.07E-20 | 9.216473802 | 2.725701979 |
| CNAG_04743 | - | oligosaccharyltransferase complex subunit beta | 4030.192821 | 3915.53571 | 3811.642959 | 5812.716439 | 5637.059807 | 5752.647939 | 3.07E-19 | 0.533808836 | 3.23E-20 | 9.210971274 | 1.447746327 |
| CNAG_00321 | - | dRaptor | 118.5614285 | 100.5313935 | 102.6587216 | 285.5739485 | 335.4482718 | 326.9252899 | 3.07E-19 | 1.548016268 | 3.23E-20 | 9.210991489 | 2.92414787 |
| CNAG_00586 | - | hypothetical protein | 101.2182544 | 148.9534305 | 137.8238326 | 353.4601695 | 386.3063749 | 362.5396004 | 3.29E-19 | 1.497878769 | 3.47E-20 | 9.203260437 | 2.824271473 |
| CNAG_05900 | - | glycine-tRNA ligase | 18471.21468 | 18318.38052 | 18945.40859 | 11855.16276 | 13344.47121 | 13258.82778 | 3.32E-19 | -0.550707053 | 3.51E-20 | -9.202043628 | 0.682685468 |
| CNAG_05631 | - | NADH-ubiquinone oxidoreductase 49 kDa subunit, mitochondrial | 8803.293397 | 8544.966045 | 8307.56577 | 5279.359526 | 6130.022985 | 5730.867453 | 3.41E-19 | -0.597395959 | 3.60E-20 | -9.199245662 | 0.660945876 |
| CNAG_01609 | - | Cystathionine gamma-synthase | 2223.17131 | 2631.890021 | 2452.670187 | 1359.815345 | 1418.554641 | 1578.854883 | 3.47E-19 | -0.76079849 | 3.67E-20 | -9.197221533 | 0.590169598 |
| CNAG_01193 | - | hypothetical protein, variant | 88.33938943 | 105.8977885 | 113.2934688 | 307.6915169 | 335.5118844 | 282.709643 | 3.53E-19 | 1.57927142 | 3.74E-20 | 9.195264895 | 2.988189041 |

| | | | | | | | | | | | | | |
|---|---|---|---|---|---|---|---|---|---|---|---|---|---|
| CNAG_03373 | - | hypothetical protein | 1702.264908 | 1835.721934 | 1720.054739 | 1132.743911 | 1089.10007 | 1063.803418 | 3.61E-19 | -0.693688438 | 3.84E-20 | -9.192575869 | 0.618271137 |
| CNAG_03590 | CKB1 | casein kinase II subunit beta | 2707.761845 | 2901.340542 | 2873.941307 | 1792.675089 | 1804.170354 | 1941.858871 | 3.74E-19 | -0.630020598 | 3.98E-20 | -9.18870381 | 0.64616719 |
| CNAG_00108 | - | RuvB-like helicase 1 | 4466.257255 | 4076.909271 | 4446.957921 | 2863.653768 | 2910.015431 | 2891.077379 | 3.83E-19 | -0.599802016 | 4.07E-20 | -9.186097438 | 0.659844501 |
| CNAG_07855 | - | midasin | 17465.15415 | 15437.63873 | 16049.19942 | 23564.61586 | 24941.24909 | 25425.75297 | 3.99E-19 | 0.579002336 | 4.24E-20 | 9.18167565 | 1.493815876 |
| CNAG_04375 | - | hexaprenyl-diphosphate synthase | 3759.661096 | 3295.764837 | 3601.253248 | 2122.601225 | 2311.387457 | 2298.259119 | 4.05E-19 | -0.678554109 | 4.32E-20 | -9.179805918 | 0.624791136 |
| CNAG_03892 | - | chaperonin GroES | 4097.19265 | 3670.711845 | 4474.57652 | 2630.209219 | 2215.507705 | 2291.679876 | 4.16E-19 | -0.794359439 | 4.44E-20 | -9.176724504 | 0.576599128 |
| CNAG_06520 | - | hypothetical protein | 483.9663337 | 441.5440023 | 526.4840468 | 211.7093831 | 218.6389373 | 194.6961711 | 4.47E-19 | -1.234614909 | 4.77E-20 | -9.169036634 | 0.424955916 |
| CNAG_03857 | - | hypothetical protein | 299.5196903 | 295.487254 | 329.7289992 | 704.5846363 | 676.789788 | 595.9140569 | 4.49E-19 | 1.081947515 | 4.80E-20 | 9.168416036 | 2.116891777 |
| CNAG_01547 | - | wd-repeat protein | 2308.31549 | 2318.477685 | 2369.50623 | 1583.431318 | 1489.34051 | 1452.351766 | 4.69E-19 | -0.643987249 | 5.02E-20 | -9.163518621 | 0.639941863 |
| CNAG_03293 | TSR2 | pre-rRNA-processing protein TSR2 | 2681.869384 | 2340.103058 | 3006.040457 | 1559.136108 | 1563.480107 | 1483.486335 | 5.73E-19 | -0.817806449 | 6.15E-20 | -9.141741333 | 0.567303846 |
| CNAG_04513 | - | DNA-directed RNA polymerase III subunit RPC2 | 5999.55748 | 5643.48626 | 6081.832011 | 3803.779685 | 3962.333551 | 4169.940031 | 5.88E-19 | -0.586085347 | 6.31E-20 | -9.138844014 | 0.666148003 |
| CNAG_04082 | - | proline-tRNA ligase | 5995.894184 | 5254.532709 | 5424.932423 | 3565.634225 | 3771.730781 | 3589.389732 | 6.22E-19 | -0.625795048 | 6.69E-20 | -9.132579023 | 0.648062541 |
| CNAG_05802 | - | D-aspartate oxidase | 143.2239351 | 194.2009577 | 193.2554274 | 493.1113183 | 442.1615855 | 429.2549635 | 6.68E-19 | 1.349396047 | 7.18E-20 | 9.124834661 | 2.548054343 |
| CNAG_03295 | - | hypothetical protein | 273.7995222 | 279.2992036 | 272.1639892 | 585.2918575 | 551.5571912 | 606.8060927 | 7.79E-19 | 1.065018166 | 8.39E-20 | 9.108065536 | 2.092196223 |
| CNAG_06626 | - | pre-mRNA-splicing factor ATP-dependent RNA helicase DHX15/PRP43 | 6006.487931 | 5326.74294 | 6067.940556 | 3577.255448 | 3853.656695 | 3850.25239 | 8.01E-19 | -0.641164225 | 8.64E-20 | -9.104844232 | 0.641195308 |
| CNAG_04182 | - | hypothetical protein | 2338.609464 | 2184.914997 | 2405.736034 | 1476.722478 | 1408.224922 | 1496.72107 | 8.13E-19 | -0.676829297 | 8.78E-20 | -9.103085705 | 0.625538551 |
| CNAG_06474 | - | RNA-binding protein Musashi | 2707.715105 | 2897.008779 | 2832.342418 | 1636.918769 | 1698.040548 | 1931.852184 | 9.58E-19 | -0.694936933 | 1.04E-19 | -9.085137255 | 0.617736322 |
| CNAG_04943 | - | CBS domain-containing protein, variant | 797.9942225 | 829.7966736 | 792.5440305 | 1371.940598 | 1359.747452 | 1323.069863 | 9.61E-19 | 0.730090076 | 1.04E-19 | 9.084688596 | 1.658742654 |
| CNAG_04834 | - | MFS transporter | 74.41427095 | 106.9928394 | 131.4041189 | 316.8956442 | 339.694585 | 356.8187759 | 9.70E-19 | 1.686244345 | 1.05E-19 | 9.083625696 | 3.218178491 |
| CNAG_03205 | - | ATP-dependent RNA helicase ROK1 | 3340.475471 | 3004.849274 | 3311.162011 | 1905.45045 | 1975.798953 | 2162.785567 | 9.98E-19 | -0.691931774 | 1.08E-19 | -9.080320972 | 0.619024419 |
| CNAG_07753 | - | hypothetical protein | 149.0650719 | 220.0275431 | 181.5545282 | 461.6130147 | 447.2805094 | 515.6074152 | 1.06E-18 | 1.357734162 | 1.15E-19 | 9.073570775 | 2.562823567 |
| CNAG_01528 | - | hypothetical protein | 812.281106 | 810.0129597 | 892.3442108 | 426.6164428 | 372.1193835 | 467.6467907 | 1.11E-18 | -1.00593929 | 1.21E-19 | -9.068180224 | 0.49794583 |
| CNAG_07878 | - | nucleolar complex protein 2 | 2548.724178 | 2304.50704 | 2614.730318 | 1349.662458 | 1459.102702 | 1616.59692 | 1.11E-18 | -0.770676824 | 1.21E-19 | -9.067969186 | 0.586142428 |
| CNAG_01969 | - | zinc metalloprotease | 168.8456704 | 208.1946231 | 199.6548534 | 471.8272424 | 485.4146688 | 425.9851077 | 1.25E-18 | 1.249320682 | 1.36E-19 | 9.055354062 | 2.377294577 |
| CNAG_02022 | - | DNA-directed RNA polymerase II subunit RPB3 | 2176.283525 | 2151.456435 | 2064.537096 | 1423.083277 | 1307.306954 | 1361.295982 | 1.27E-18 | -0.658751038 | 1.38E-19 | -9.053730532 | 0.633426426 |
| CNAG_04104 | AKP3 | mucin-desulfating sulfatase | 1394.667663 | 1134.393467 | 1178.276602 | 679.6235424 | 649.9616517 | 674.1717635 | 1.37E-18 | -0.904439552 | 1.49E-19 | -9.045365935 | 0.534240202 |
| CNAG_01397 | - | mitochondrial import inner membrane translocase subunit | 3573.010614 | 3499.376717 | 3626.872215 | 2447.313059 | 2380.386529 | 2484.731268 | 1.53E-18 | -0.564510537 | 1.68E-19 | -9.032676964 | 0.676184787 |
| CNAG_02475 | - | Flavin-containing monooxygenase | 91.80344168 | 76.86446966 | 116.4732573 | 286.5779765 | 309.5447845 | 301.4137247 | 1.58E-18 | 1.646635179 | 1.73E-19 | 9.029301657 | 3.131025328 |
| CNAG_07469 | - | nuclear pore complex protein Nup98-Nup96 | 6973.505879 | 7025.786467 | 7009.670899 | 4389.790443 | 4707.587594 | 5082.468663 | 1.60E-18 | -0.582361128 | 1.75E-19 | -9.02779459 | 0.66786984 |
| CNAG_05596 | - | nuclear transport factor 2 | 5790.597714 | 5632.710808 | 5892.027562 | 4047.489296 | 4127.991065 | 4076.703139 | 1.82E-18 | -0.514597029 | 2.00E-19 | -9.013285789 | 0.699988425 |
| CNAG_01362 | - | cell cycle control protein cwf19 | 3902.238919 | 3682.540896 | 3746.331071 | 2465.142001 | 2527.029781 | 2645.695359 | 1.88E-18 | -0.584518457 | 2.07E-19 | -9.009675627 | 0.66687189 |
| CNAG_07626 | - | NAD-binding Rossmann fold oxidoreductase | 181.6603022 | 221.1092577 | 192.207812 | 458.431023 | 443.7875522 | 507.8897683 | 1.89E-18 | 1.230760268 | 2.08E-19 | 9.009168509 | 2.34690634 |
| CNAG_00649 | - | tryptophan synthase, beta subunit | 8963.496639 | 9803.395928 | 10153.53332 | 6374.508582 | 6939.102874 | 6311.505205 | 1.94E-18 | -0.574509878 | 2.14E-19 | -9.005937547 | 0.671514344 |
| CNAG_04586 | HOB7 | LIM-homeobox protein, variant | 548.1156085 | 566.9160824 | 565.7701276 | 964.1132623 | 989.7718017 | 972.1963054 | 2.27E-18 | 0.837398745 | 2.50E-19 | 8.988573769 | 1.786825495 |
| CNAG_01018 | - | histone-lysine N-methyltransferase Su(var)3-9 | 8450.629696 | 7637.83334 | 8327.791448 | 5344.818063 | 5571.026134 | 5700.862464 | 2.28E-18 | -0.570981213 | 2.51E-19 | -8.988214341 | 0.673158801 |
| CNAG_04175 | - | hypothetical protein, variant | 2243.976015 | 2100.854 | 2196.736857 | 1324.885833 | 1302.978623 | 1462.274538 | 2.35E-18 | -0.693081352 | 2.60E-19 | -8.984614404 | 0.61853136 |
| CNAG_02952 | - | hypothetical protein | 1646.150928 | 1628.912965 | 1668.837269 | 1074.388463 | 1032.179914 | 972.7887945 | 2.46E-18 | -0.698464352 | 2.72E-19 | -8.979439743 | 0.616227789 |
| CNAG_02609 | - | 3-demethylubiquinone-9 3-O-methyltransferase | 1599.245753 | 1376.844677 | 1533.371248 | 855.5215302 | 888.9635089 | 899.4916772 | 2.62E-18 | -0.786780409 | 2.90E-19 | -8.972506975 | 0.579636195 |
| CNAG_03001 | - | multiple RNA-binding domain-containing protein 1 | 2829.091464 | 2615.908511 | 3010.395101 | 1806.715038 | 1787.784562 | 1803.138866 | 2.63E-18 | -0.663448198 | 2.91E-19 | -8.972016783 | 0.631367455 |
| CNAG_00366 | - | hypothetical protein | 32.2687261 | 22.99882458 | 7.791103854 | 130.0027532 | 177.1324978 | 173.409831 | 2.66E-18 | 2.960271794 | 2.94E-19 | 8.970782134 | 7.782705651 |
| CNAG_03234 | - | cell wall surface anchor protein, variant | 90.58891979 | 89.74032505 | 72.82541191 | 237.2437989 | 274.9792324 | 313.3891711 | 2.69E-18 | 1.699997607 | 2.98E-19 | 8.969282263 | 3.249004196 |
| CNAG_05624 | - | hypothetical protein | 1144.083352 | 1230.210363 | 1256.076487 | 664.018353 | 711.2735252 | 741.8283663 | 2.83E-18 | -0.793622061 | 3.15E-19 | -8.963481887 | 0.57689391 |
| CNAG_00872 | - | ATP-dependent DNA helicase | 2137.730554 | 2090.046108 | 1982.432238 | 1307.803398 | 1336.679046 | 1363.494837 | 3.26E-18 | -0.647448686 | 3.63E-19 | -8.947728628 | 0.6384083 |
| CNAG_04713 | - | citrate lyase subunit beta-like protein | 1154.147624 | 1163.811353 | 1241.479988 | 1855.882781 | 1925.573269 | 1911.43801 | 3.51E-18 | 0.662307571 | 3.91E-19 | 8.939502566 | 1.582611967 |
| CNAG_01197 | - | ran GTPase-activating protein 1 | 4357.686576 | 4096.289 | 4324.330553 | 2756.9853 | 3006.667613 | 2882.189905 | 3.53E-18 | -0.579424386 | 3.94E-19 | -8.938603648 | 0.669230737 |
| CNAG_05118 | - | hypothetical protein | 1564.563197 | 1842.141123 | 1798.898976 | 1099.291113 | 986.4012917 | 930.6482335 | 3.59E-18 | -0.802093185 | 4.00E-19 | -8.936837768 | 0.573516467 |
| CNAG_04607 | - | hypothetical protein | 931.1512288 | 964.493827 | 996.2177706 | 1691.895624 | 1504.514662 | 1699.316915 | 3.65E-18 | 0.744477354 | 4.08E-19 | 8.934841022 | 1.675367223 |
| CNAG_02780 | BNA5 | Kynureninase | 3042.865341 | 3031.761294 | 3126.701055 | 2047.232268 | 2157.040435 | 2036.26274 | 3.65E-18 | -0.5753188 | 4.08E-19 | -8.934871932 | 0.67113793 |
| CNAG_03048 | IRK3 | Gluconokinase | 1831.759394 | 1772.20498 | 1772.298397 | 1189.614298 | 1029.520011 | 1078.236729 | 3.65E-18 | -0.720812969 | 4.08E-19 | -8.934702226 | 0.606755435 |
| CNAG_03677 | - | hypothetical protein | 1282.536895 | 1227.378344 | 1029.353467 | 2174.527474 | 2016.998402 | 1995.819815 | 3.72E-18 | 0.790648101 | 4.17E-19 | 8.932313223 | 1.729851388 |
| CNAG_01827 | - | hypothetical protein | 58.04282202 | 87.57754731 | 88.77489286 | 240.5410111 | 278.5633627 | 272.4868845 | 3.80E-18 | 1.747162103 | 4.26E-19 | 8.930006585 | 3.356975722 |
| CNAG_04050 | - | small nuclear ribonucleoprotein D3 | 5249.947034 | 5143.558228 | 5763.994887 | 3525.955495 | 3667.352791 | 3699.255157 | 3.98E-18 | -0.584330766 | 4.48E-19 | -8.924537502 | 0.666958654 |
| CNAG_06874 | - | HpcH/HpaI aldolase/citrate lyase | 512.2094452 | 599.7540848 | 533.019451 | 272.5233843 | 260.9176657 | 241.2967636 | 4.13E-18 | -1.103278996 | 4.65E-19 | -8.920323796 | 0.465457389 |
| CNAG_02532 | - | D-amino-acid oxidase | 520.131352 | 507.1821107 | 495.6747349 | 213.993737 | 205.6246345 | 260.0713819 | 4.22E-18 | -1.181673424 | 4.75E-19 | -8.917963946 | 0.440839858 |
| CNAG_03457 | - | arginine-tRNA ligase | 7718.448855 | 7159.429747 | 7427.72469 | 4723.692386 | 5243.284745 | 5199.070774 | 4.28E-18 | -0.572216628 | 4.83E-19 | -8.916192684 | 0.672582605 |
| CNAG_03225 | - | malate dehydrogenase, NAD-dependent | 8601.207868 | 7880.217939 | 7959.894431 | 5766.794754 | 5011.131562 | 5405.595373 | 4.66E-18 | -0.610326421 | 5.26E-19 | -8.906643124 | 0.655048475 |
| CNAG_00269 | - | Sorbitol dehydrogenase | 13.7416031 | 1.441439086 | 7.809500109 | 333.8669746 | 366.0543392 | 443.7491673 | 5.04E-18 | 5.744849099 | 5.69E-19 | 8.89788137 | 53.62556728 |
| CNAG_05376 | - | 3-dehydroshikimate dehydratase | 180.460548 | 181.2969079 | 208.1631308 | 419.399138 | 456.0160347 | 459.1291239 | 5.21E-18 | 1.213898285 | 5.89E-19 | 8.894099596 | 2.31963576 |
| CNAG_02416 | - | hypothetical protein | 1807.355702 | 1831.472304 | 1846.950373 | 1196.560348 | 1199.545954 | 1160.365356 | 5.22E-18 | -0.64062202 | 5.91E-19 | -8.893667313 | 0.641436333 |
| CNAG_02672 | - | nonsense-mediated mRNA decay protein 3 | 6045.063022 | 5392.476745 | 6458.227236 | 3750.07034 | 3772.531304 | 3964.594816 | 5.68E-18 | -0.655553445 | 6.44E-19 | -8.884191654 | 0.634831911 |
| CNAG_04380 | - | peptidase, variant | 363.6518524 | 351.4798261 | 338.2824255 | 753.55898 | 653.2748018 | 700.1574823 | 5.69E-18 | 0.98620839 | 6.45E-19 | 8.883922525 | 1.980971864 |
| CNAG_04260 | - | translation initiation factor eIF-2B subunit gamma | 2648.058924 | 2573.828362 | 2509.250455 | 1580.93139 | 1744.712369 | 1725.39855 | 5.93E-18 | -0.629846817 | 6.74E-19 | -8.879184359 | 0.646245029 |
| CNAG_07473 | - | valine-tRNA ligase | 10656.54441 | 10048.02418 | 10107.74545 | 6888.957578 | 7324.641433 | 7463.778169 | 6.02E-18 | -0.522916022 | 6.85E-19 | -8.877367915 | 0.695963706 |
| CNAG_05265 | - | hypothetical protein | 1484.983109 | 1513.589491 | 1487.543378 | 958.4198027 | 919.1808991 | 811.8605581 | 6.07E-18 | -0.753524272 | 6.91E-19 | -8.876374516 | 0.593152809 |
| CNAG_03940 | - | elongator complex protein 2 | 3943.086803 | 3630.837051 | 3944.243289 | 2459.690887 | 2323.386347 | 2653.463774 | 6.11E-18 | -0.648105728 | 6.96E-19 | -8.875486348 | 0.638117618 |
| CNAG_03261 | - | zinc finger protein | 1531.696366 | 1529.773153 | 1539.797775 | 1004.361895 | 881.1615758 | 926.1404119 | 6.19E-18 | -0.725394559 | 7.06E-19 | -8.873952084 | 0.604831608 |
| CNAG_12314 | - | unspecified product | 21.80118933 | 32.64599417 | 18.44914462 | 153.4334712 | 134.4313619 | 132.6586016 | 7.56E-18 | 2.547163526 | 8.63E-19 | 8.851506392 | 5.844839976 |
| CNAG_03596 | - | 2-oxoglutarate dehydrogenase E2 component (dihydrolipoamide succinyltransferase) | 10021.11107 | 8452.377306 | 9039.136912 | 5954.999121 | 6110.974419 | 6091.673013 | 7.67E-18 | -0.615752012 | 8.77E-19 | -8.849785375 | 0.652589639 |
| CNAG_02087 | - | Acyl-CoA-dependent ceramide synthase | 68.54515754 | 103.7253833 | 104.7605267 | 281.9649708 | 288.7651425 | 294.7339648 | 7.96E-18 | 1.635711443 | 9.11E-19 | 8.845577511 | 3.107407493 |
| CNAG_00025 | VCX1 | vacuolar calcium exchanger | 577.3898576 | 671.4087509 | 606.9992201 | 1171.3311 | 1122.594797 | 1043.334 | 8.02E-18 | 0.832779393 | 9.19E-19 | 8.844553136 | 1.781113424 |
| CNAG_02074 | - | mitochondrial processing peptidase | 3561.15259 | 3117.993185 | 3357.052659 | 2155.387569 | 2113.82321 | 2230.522845 | 8.26E-18 | -0.642758249 | 9.47E-19 | -8.841243692 | 0.640487247 |
| CNAG_02956 | - | transcription factor iws1 | 2559.277292 | 2362.686833 | 2486.821759 | 1494.53078 | 1653.275855 | 1616.615784 | 8.60E-18 | -0.652697021 | 9.87E-19 | -8.836550503 | 0.636090074 |
| CNAG_06775 | - | hypothetical protein | 1176.412855 | 944.8169873 | 1174.962801 | 569.8218019 | 602.563555 | 562.0796149 | 8.78E-18 | -0.942963441 | 1.01E-18 | -8.834141567 | 0.520163314 |
| CNAG_04363 | - | hypothetical protein | 15.94475906 | 22.95700733 | 12.04310507 | 119.671983 | 128.6687806 | 110.4757473 | 8.96E-18 | 2.850737302 | 1.03E-18 | 8.831716568 | 7.213689377 |
| CNAG_06770 | FBA1 | fructose-bisphosphate aldolase 1 | 25639.98352 | 23601.0732 | 24640.29924 | 18025.64401 | 17782.33509 | 17302.06205 | 9.00E-18 | -0.491916042 | 1.04E-18 | -8.831086585 | 0.711080086 |
| CNAG_00021 | - | S-adenosylmethionine decarboxylase proenzyme | 810.0184236 | 872.4569932 | 830.5861365 | 490.5830118 | 444.6778554 | 448.8220866 | 9.12E-18 | -0.876964522 | 1.05E-18 | -8.829494688 | 0.544511898 |
| CNAG_04721 | - | hypothetical protein | 506.2763218 | 653.112972 | 650.7002934 | 1168.951439 | 1100.108709 | 1177.503528 | 9.25E-18 | 0.915008283 | 1.07E-18 | 8.827763282 | 1.885579897 |
| CNAG_05578 | - | hypothetical protein | 1463.999226 | 1416.688188 | 1617.576299 | 897.6093327 | 931.2669059 | 865.1081556 | 9.51E-18 | -0.755047483 | 1.10E-18 | -8.82455737 | 0.592526884 |
| CNAG_05766 | SPR1 | endoglucanase C | 125.5161516 | 104.826481 | 98.39777611 | 313.1594215 | 299.0103085 | 308.1261895 | 9.57E-18 | 1.476730916 | 1.11E-18 | 8.823751748 | 2.783173632 |
| CNAG_04507 | - | hypothetical protein | 311.3507172 | 435.5001098 | 440.627255 | 803.0455532 | 879.5172317 | 864.3898824 | 9.68E-18 | 1.087861093 | 1.12E-18 | 8.822327326 | 2.125586681 |
| CNAG_03646 | - | phosphoribosylformimino-5-aminoimidazole carboxamide ribotide isomerase | 1630.902077 | 1596.5699 | 1538.764687 | 979.3560385 | 926.0237421 | 1034.881244 | 9.85E-18 | -0.711985469 | 1.14E-18 | -8.820258166 | 0.610479405 |
| CNAG_04777 | - | S-adenosylmethionine-dependent methyltransferase | 1115.640556 | 1103.498012 | 1253.202753 | 1845.770173 | 1966.165486 | 1900.334261 | 9.90E-18 | 0.702900029 | 1.15E-18 | 8.819643183 | 1.62777357 |
| CNAG_02540 | - | hypothetical protein | 2348.132737 | 3661.553039 | 3391.938112 | 978.0614344 | 1017.203303 | 1000.756803 | 1.00E-17 | -1.663528506 | 1.17E-18 | -8.817918712 | 0.315666155 |
| CNAG_02403 | - | NADH dehydrogenase (ubiquinone) 1 beta subcomplex 9 | 3368.674598 | 3391.616724 | 3358.149165 | 2357.050598 | 2389.929179 | 2284.930815 | 1.02E-17 | -0.540452341 | 1.18E-18 | -8.816430506 | 0.6875553 |
| CNAG_04990 | PRO2 | glutamate-5-semialdehyde dehydrogenase | 3531.022877 | 3588.792233 | 3648.207425 | 2484.674193 | 2566.763949 | 2431.472194 | 1.04E-17 | -0.540599168 | 1.21E-18 | -8.813559004 | 0.687485329 |
| CNAG_01768 | - | hypothetical protein | 4843.505027 | 4667.319798 | 4673.09689 | 3280.858129 | 3356.855137 | 3370.631678 | 1.12E-17 | -0.518696348 | 1.30E-18 | -8.805638992 | 0.698002279 |
| CNAG_06336 | - | glucan 1,3-beta-glucosidase | 1505.539463 | 1422.384046 | 1332.177567 | 2213.111293 | 2246.402706 | 2344.363166 | 1.12E-17 | 0.660331836 | 1.31E-18 | 8.804934784 | 1.580446102 |

| | | | | | | | | | | | | | |
|---|---|---|---|---|---|---|---|---|---|---|---|---|---|
| CNAG_04751 | - | hypothetical protein | 539.9594726 | 584.141082 | 472.6687362 | 968.7893075 | 983.7172784 | 1010.996842 | 1.20E-17 | 0.877815359 | 1.40E-18 | 8.797216162 | 1.837590571 |
| CNAG_00132 | - | hypothetical protein, variant | 45.17155466 | 54.21499827 | 68.50585595 | 197.1019216 | 219.9244008 | 210.3453314 | 1.25E-17 | 1.891195999 | 1.46E-18 | 8.792423585 | 3.709426099 |
| CNAG_05004 | - | GTP-binding protein LepA | 2033.790813 | 1896.150043 | 1967.463444 | 1296.200562 | 1220.198723 | 1263.597276 | 1.25E-17 | -0.657217414 | 1.47E-18 | -8.792263874 | 0.634100134 |
| CNAG_06873 | - | 5-oxoprolinase | 24.12891189 | 27.28493859 | 25.89895306 | 167.7210481 | 119.5047935 | 140.371388 | 1.28E-17 | 2.472160156 | 1.50E-18 | 8.789948904 | 5.548739818 |
| CNAG_01587 | - | RNA polymerase-associated protein LEO1 | 2655.096509 | 2526.455303 | 2709.693009 | 1675.944534 | 1624.730748 | 1808.635567 | 1.31E-17 | -0.642931715 | 1.54E-18 | -8.786867134 | 0.640410242 |
| CNAG_02720 | - | nuclear GTP-binding protein | 2268.404008 | 1930.66959 | 2225.492541 | 1255.584551 | 1362.565466 | 1271.411393 | 1.32E-17 | -0.740363915 | 1.55E-18 | -8.786279513 | 0.598588341 |
| CNAG_00283 | - | hypothetical protein | 138.4337608 | 130.6813797 | 156.9866082 | 380.9099713 | 345.5094819 | 357.0335933 | 1.32E-17 | 1.336106521 | 1.55E-18 | 8.785668086 | 2.524690468 |
| CNAG_02516 | HLH5 | hypothetical protein, variant 2 | 270.325448 | 313.7365133 | 274.3017672 | 575.8346452 | 586.1325034 | 605.7339367 | 1.42E-17 | 1.028484134 | 1.67E-18 | 8.777462403 | 2.039879786 |
| CNAG_06314 | - | phosphoribosylamine-glycine ligase | 5802.25137 | 5783.517654 | 5701.151642 | 3784.334055 | 4257.419698 | 3959.042505 | 1.47E-17 | -0.541958689 | 1.74E-18 | -8.773224897 | 0.686837784 |
| CNAG_01960 | - | efflux protein EncT | 48.66845636 | 72.47863595 | 56.81668752 | 220.7147226 | 228.5590675 | 197.1834065 | 1.51E-17 | 1.8595147 | 1.78E-18 | 8.770690028 | 3.628855727 |
| CNAG_01091 | - | U5 small nuclear ribonucleoprotein component | 11944.48278 | 10779.60285 | 10815.87431 | 7345.800766 | 7535.060345 | 8017.749374 | 1.52E-17 | -0.566251295 | 1.80E-18 | -8.769325722 | 0.675369393 |
| CNAG_07634 | - | hypothetical protein | 1254.484943 | 1102.427776 | 1177.520513 | 1982.13669 | 1891.89011 | 1905.90085 | 1.59E-17 | 0.693944272 | 1.88E-18 | 8.764328353 | 1.617700207 |
| CNAG_01270 | - | hypothetical protein | 9563.502286 | 8759.430542 | 8960.241837 | 6320.082905 | 6431.017782 | 6506.83406 | 1.68E-17 | -0.51819916 | 1.99E-18 | -8.757865992 | 0.698242869 |
| CNAG_00987 | - | prefoldin subunit | 2188.993588 | 2027.557643 | 1923.779131 | 1229.145533 | 1221.037786 | 1347.939334 | 1.84E-17 | -0.708271275 | 2.18E-18 | -8.747515293 | 0.612053099 |
| CNAG_06621 | - | biotin synthase | 5974.892183 | 5062.786413 | 5721.364874 | 3682.422348 | 3682.858628 | 3588.282906 | 1.90E-17 | -0.62967075 | 2.25E-18 | -8.744045941 | 0.646323902 |
| CNAG_07020 | - | hypothetical protein | 1320.165654 | 1202.252141 | 1199.622461 | 760.6723425 | 734.5630857 | 714.1245497 | 2.10E-17 | -0.768549499 | 2.49E-18 | -8.732746837 | 0.587007362 |
| CNAG_04797 | - | signal recognition particle subunit SRP19 | 2271.637512 | 2360.798808 | 2410.298101 | 3394.486954 | 3522.988001 | 3453.507315 | 2.14E-17 | 0.543253312 | 2.54E-18 | 8.730342037 | 1.45725496 |
| CNAG_07344 | - | ras guanyl-nucleotide exchange factor, variant | 175.8996382 | 236.2058227 | 238.0294822 | 469.1155927 | 523.2836244 | 561.1648974 | 2.15E-17 | 1.244426573 | 2.55E-18 | 8.729752293 | 2.369243652 |
| CNAG_07310 | - | hypothetical protein | 1949.503522 | 1621.428397 | 1744.565557 | 1073.521434 | 1046.840567 | 1051.595146 | 2.20E-17 | -0.761335383 | 2.62E-18 | -8.726701204 | 0.58995001 |
| CNAG_03007 | - | hypothetical protein | 2133.114842 | 2995.522783 | 2128.958052 | 7413.036588 | 6491.324226 | 6818.081549 | 2.37E-17 | 1.500760612 | 2.82E-18 | 8.718384439 | 2.82991871 |
| CNAG_01758 | PGU1 | Polygalacturonase | 102.2738451 | 119.863217 | 84.55546299 | 333.7156719 | 271.2715865 | 304.7716 | 2.43E-17 | 1.559615957 | 2.90E-18 | 8.715214129 | 2.947753642 |
| CNAG_03404 | - | hypothetical protein, variant | 54.56138495 | 85.43460392 | 94.09891744 | 293.3018913 | 281.9964028 | 237.2170832 | 2.44E-17 | 1.784937133 | 2.91E-18 | 8.714986925 | 3.44603447 |
| CNAG_06638 | - | malate dehydrogenase (oxaloacetate- decarboxylating) | 4743.927538 | 4298.763262 | 3749.595531 | 2490.715626 | 2734.901443 | 2623.566699 | 2.52E-17 | -0.720398415 | 3.01E-18 | -8.711004998 | 0.606929809 |
| CNAG_00067 | SSS1 | protein translocase SEC61 complex gamma subunit | 3028.885583 | 3127.63407 | 3088.328781 | 2189.751964 | 2119.056924 | 2069.563999 | 2.66E-17 | -0.550609181 | 3.18E-18 | -8.704717212 | 0.682731783 |
| CNAG_02150 | - | mRNA methyltransferase | 274.9427379 | 256.6962162 | 282.8067363 | 532.9332527 | 571.5073693 | 590.161061 | 2.76E-17 | 1.044537521 | 3.31E-18 | 8.700429556 | 2.062705019 |
| CNAG_02145 | - | hypothetical protein | 1600.724571 | 1662.320557 | 1826.61547 | 1033.035836 | 1082.227156 | 1043.808576 | 2.79E-17 | -0.703375165 | 3.34E-18 | -8.69913287 | 0.614133767 |
| CNAG_01943 | - | hypothetical protein | 3339.465668 | 3301.122894 | 3539.387984 | 2059.559364 | 2327.813923 | 2310.442335 | 2.82E-17 | -0.619664979 | 3.38E-18 | -8.697813175 | 0.650822043 |
| CNAG_01295 | - | acyl carrier protein, variant | 780.5246648 | 654.9389129 | 766.5198151 | 345.5896286 | 354.0652613 | 398.8315278 | 2.84E-17 | -1.019870527 | 3.41E-18 | -8.697017698 | 0.493160608 |
| CNAG_02496 | - | hypothetical protein | 982.8913581 | 987.8362178 | 1128.040429 | 596.3980174 | 549.9053413 | 590.8851723 | 2.92E-17 | -0.851222345 | 3.51E-18 | -8.693707901 | 0.554314886 |
| CNAG_02075 | - | hypothetical protein | 1184.98772 | 1257.176549 | 1354.167713 | 783.2567816 | 737.9997707 | 732.9931699 | 3.13E-17 | -0.768006526 | 3.77E-18 | -8.685642567 | 0.587228329 |
| CNAG_06829 | ERG1 | Squalene monooxygenase | 2053.497617 | 1833.640588 | 1833.105374 | 1086.657748 | 1124.428665 | 1248.022629 | 3.21E-17 | -0.741397463 | 3.87E-18 | -8.68261472 | 0.598159666 |
| CNAG_04315 | - | hypothetical protein | 3503.453357 | 2676.201519 | 2563.667563 | 1591.757438 | 1667.846417 | 1525.698773 | 3.23E-17 | -0.886528187 | 3.90E-18 | -8.681760711 | 0.540914251 |
| CNAG_02838 | - | small subunit ribosomal protein S17 | 1093.635155 | 986.7797614 | 1052.407106 | 618.9919706 | 612.9455426 | 572.0481111 | 3.24E-17 | -0.812772728 | 3.91E-18 | -8.681509017 | 0.569286689 |
| CNAG_01198 | - | small nuclear ribonucleoprotein F | 2843.106531 | 2610.528518 | 3155.393587 | 1860.454255 | 1768.79257 | 1696.612715 | 3.67E-17 | -0.709012772 | 4.43E-18 | -8.667251565 | 0.611738605 |
| CNAG_02681 | APH3 | Phytase | 200.2903477 | 199.6057082 | 229.4833237 | 499.1212784 | 468.8720978 | 444.8536067 | 3.71E-17 | 1.152477763 | 4.49E-18 | 8.665735984 | 2.222953489 |
| CNAG_00165 | - | methylthioadenosine phosphorylase | 1412.492259 | 1297.088858 | 1377.686728 | 871.2444933 | 793.1964806 | 801.8166978 | 4.26E-17 | -0.744678498 | 5.15E-18 | -8.649904916 | 0.596800853 |
| CNAG_07445 | - | transketolase | 23742.4094 | 22771.42278 | 22322.92195 | 15980.20438 | 17174.26401 | 16882.38563 | 4.51E-17 | -0.475635133 | 5.46E-18 | -8.643285098 | 0.719150121 |
| CNAG_04465 | - | small nuclear ribonucleoprotein, variant | 2099.302356 | 2101.92354 | 2118.909899 | 1386.451504 | 1431.597193 | 1411.229848 | 4.69E-17 | -0.594770339 | 5.69E-18 | -8.638667155 | 0.662149854 |
| CNAG_01759 | - | vacuolar membrane protein | 34.57442307 | 31.58708444 | 21.64949977 | 130.0710847 | 156.1773317 | 156.907232 | 4.69E-17 | 2.34639334 | 5.68E-18 | 8.638757092 | 5.085513097 |
| CNAG_00534 | - | hypothetical protein | 8457.78856 | 8438.341907 | 9125.482996 | 6373.112298 | 5968.66205 | 5798.603019 | 4.78E-17 | -0.535903415 | 5.80E-18 | -8.636385525 | 0.689726635 |
| CNAG_01156 | CAP2 | capsular related protein | 4493.291902 | 4750.246312 | 4572.851683 | 3269.211263 | 3320.63729 | 3308.461517 | 4.89E-17 | -0.496168153 | 5.95E-18 | -8.633469674 | 0.708987377 |
| CNAG_03061 | - | multiple drug resistance protein | 2257.993611 | 2361.481393 | 2093.338113 | 1211.121796 | 1290.887367 | 1505.553159 | 4.92E-17 | -0.759108101 | 5.99E-18 | -8.632660692 | 0.590861498 |
| CNAG_03452 | - | AFG1 family mitochondrial ATPase | 2104.951143 | 1873.537558 | 1999.445444 | 1180.089548 | 1228.828454 | 1315.744734 | 5.03E-17 | -0.698410732 | 6.13E-18 | -8.630099816 | 0.616250693 |
| CNAG_00826 | DAK101 | dihydroxyacetone kinase | 8.956947459 | 21.88878515 | 15.24048187 | 93.97059192 | 161.1145412 | 120.3702569 | 5.08E-17 | 3.073552154 | 6.20E-18 | 8.628767333 | 8.418435556 |
| CNAG_02317 | - | protein arginine N-methyltransferase 3 | 1377.625909 | 1356.353702 | 1615.403719 | 844.6586329 | 874.3181135 | 829.5828901 | 5.39E-17 | -0.787147454 | 6.58E-18 | -8.621905171 | 0.579488745 |
| CNAG_03919 | - | acetate non-utilizing protein 9, mitochondrial | 1014.237202 | 923.2064492 | 988.4120451 | 586.3306917 | 541.3086989 | 540.9526278 | 5.56E-17 | -0.826808448 | 6.80E-18 | -8.618254661 | 0.563775056 |
| CNAG_05929 | - | MFS transporter, SP family, general alpha glucoside:H symporter | 181.6148759 | 218.9310031 | 159.1786643 | 439.0350771 | 441.3100361 | 441.4113559 | 5.70E-17 | 1.228930275 | 6.98E-18 | 8.615278695 | 2.343931283 |
| CNAG_07957 | - | WD repeat and SOF domain-containing protein 1 | 1599.420314 | 1539.503765 | 1600.587305 | 941.159126 | 1007.178998 | 1033.773526 | 5.74E-17 | -0.683904579 | 7.03E-18 | -8.614391196 | 0.622478288 |
| CNAG_03416 | - | hypothetical protein | 1110.340652 | 1334.667246 | 1391.44764 | 699.0671258 | 762.198686 | 634.2674703 | 6.00E-17 | -0.88716075 | 7.35E-18 | -8.609294956 | 0.540677134 |
| CNAG_03358 | - | Phosphoglycerate kinase | 15270.98766 | 14394.66597 | 15705.76015 | 21432.12487 | 21126.03597 | 22817.99088 | 6.45E-17 | 0.511532851 | 7.92E-18 | 8.600733333 | 1.425564041 |
| CNAG_02079 | - | cytoplasmic protein | 1763.7481 | 2096.791467 | 1735.301673 | 2953.305292 | 3095.2466 | 2917.285942 | 6.47E-17 | 0.665985325 | 7.95E-18 | 8.600374278 | 1.586651547 |
| CNAG_03896 | - | hypothetical protein | 142.0683407 | 215.7154282 | 189.0013583 | 452.2864332 | 426.5354047 | 501.1828188 | 6.52E-17 | 1.322981858 | 8.02E-18 | 8.599347163 | 2.501826699 |
| CNAG_07837 | - | hypothetical protein, variant | 116.3134929 | 143.5634692 | 141.0045317 | 341.8121131 | 325.6609227 | 360.2601201 | 6.78E-17 | 1.344420331 | 8.35E-18 | 8.594729667 | 2.539281488 |
| CNAG_06713 | - | cysteine-tRNA ligase | 3383.870348 | 3358.236835 | 3533.015059 | 2343.011728 | 2451.185495 | 2284.940219 | 6.82E-17 | -0.553027722 | 8.40E-18 | -8.593968058 | 0.681588207 |
| CNAG_01380 | - | Phosphatidylserine decarboxylase | 1104.995903 | 947.2825937 | 932.2749064 | 1611.476811 | 1750.542876 | 1751.507738 | 6.88E-17 | 0.761315448 | 8.48E-18 | 8.592843513 | 1.695035452 |
| CNAG_01856 | - | hypothetical protein | 76.60012983 | 84.33869994 | 77.06029507 | 266.7591177 | 241.2792356 | 223.8711386 | 7.18E-17 | 1.608066645 | 8.86E-18 | 8.587840761 | 3.048430479 |
| CNAG_04599 | - | 3-methyl-2-oxobutanoate hydroxymethyltransferase | 2798.229012 | 2692.661797 | 2646.014839 | 3814.661719 | 4238.966572 | 4155.087877 | 7.29E-17 | 0.569876173 | 9.00E-18 | 8.586006 | 1.484396159 |
| CNAG_00316 | HAM1 | rdgB/HAM1 family non-canonical purine NTP pyrophosphatase | 1501.405434 | 1576.085831 | 1594.167915 | 895.9885537 | 903.5982422 | 1040.406576 | 7.32E-17 | -0.733778747 | 9.05E-18 | -8.585417453 | 0.601326837 |
| CNAG_05798 | - | signal recognition particle subunit SRP9 | 1340.062327 | 1270.109727 | 1268.923173 | 801.1702167 | 790.6654678 | 666.4534097 | 7.66E-17 | -0.796527372 | 9.48E-18 | -8.580067787 | 0.575733325 |
| CNAG_03392 | - | DNA supercoiling protein | 290.1958452 | 300.8644011 | 344.6389614 | 671.7642941 | 585.9677604 | 684.5035306 | 7.72E-17 | 1.039518162 | 9.57E-18 | 8.578948659 | 2.05554102 |
| CNAG_04753 | - | Gluconolactonase | 604.1196957 | 594.9314125 | 551.54914 | 1007.675582 | 1007.823486 | 1024.372921 | 7.99E-17 | 0.781403439 | 9.91E-18 | 8.574964892 | 1.718802093 |
| CNAG_06012 | - | NADH-cytochrome b5 reductase 1 | 5171.573933 | 4926.961434 | 4673.123018 | 3499.610386 | 3349.917905 | 3392.857439 | 8.73E-17 | -0.543800081 | 1.08E-17 | -8.564686791 | 0.685961694 |
| CNAG_00919 | - | carboxypeptidase D | 1635.435791 | 1583.590806 | 1353.258779 | 878.1026198 | 922.6205575 | 911.7069526 | 8.94E-17 | -0.768903772 | 1.11E-17 | -8.561818428 | 0.586863232 |
| CNAG_01653 | CIG1 | cytokine inducing-glycoprotein | 22.96696323 | 27.28405327 | 33.33796254 | 143.7923867 | 142.2801729 | 132.6840039 | 8.95E-17 | 2.331285198 | 1.11E-17 | 8.561483705 | 5.03253464 |
| CNAG_00417 | - | elongation factor 1-gamma | 59578.4346 | 56856.02246 | 58806.27127 | 42006.27006 | 43517.7264 | 44435.44996 | 9.14E-17 | -0.446771339 | 1.14E-17 | -8.558989464 | 0.733682948 |
| CNAG_04417 | - | alpha-ketoglutarate-dependent 2,4- dichlorophenoxyacetate dioxygenase | 504.9319026 | 543.6880186 | 424.3239883 | 236.7154454 | 194.402093 | 216.8577434 | 9.31E-17 | -1.20170255 | 1.16E-17 | -8.556731713 | 0.434761909 |
| CNAG_02328 | - | pre-rRNA-processing protein TSR1 | 5236.800923 | 4282.729694 | 4860.737708 | 2993.565961 | 3123.917025 | 3062.06348 | 9.48E-17 | -0.66386994 | 1.18E-17 | -8.554562991 | 0.631182915 |
| CNAG_03649 | - | hypothetical protein | 1027.181433 | 1061.03354 | 972.4774276 | 626.0366643 | 581.8618389 | 591.9976067 | 9.68E-17 | -0.782119985 | 1.21E-17 | -8.552083559 | 0.581511656 |
| CNAG_07419 | - | hypothetical protein, variant | 195.5421987 | 187.7325239 | 132.5468083 | 447.7761133 | 412.7820403 | 440.2493932 | 9.76E-17 | 1.322619622 | 1.22E-17 | 8.550933169 | 2.501198612 |
| CNAG_01997 | - | prephenate dehydrogenase (NADP) | 1597.256101 | 1857.236679 | 1705.115112 | 1027.557807 | 1125.378615 | 1063.788857 | 9.83E-17 | -0.696949682 | 1.23E-17 | -8.550049677 | 0.6168751 |
| CNAG_00400 | - | secondary thiamine-phosphate synthase enzyme, variant | 2118.020846 | 2272.10573 | 2126.393253 | 1487.702116 | 1378.919174 | 1440.098395 | 1.01E-16 | -0.612797739 | 1.27E-17 | -8.546526619 | 0.653927346 |
| CNAG_00233 | - | chaperone protein | 1667.284354 | 1808.806413 | 1868.223148 | 1181.058561 | 989.8184425 | 1068.24442 | 1.04E-16 | -0.737386038 | 1.31E-17 | -8.542882365 | 0.599825168 |
| CNAG_01794 | - | 2-hydroxyacid dehydrogenase | 221.3101633 | 232.9866539 | 246.5561094 | 521.523738 | 458.3070383 | 567.8398585 | 1.13E-16 | 1.130320618 | 1.42E-17 | 8.533342952 | 2.189073838 |
| CNAG_04736 | - | hypothetical protein | 128.0996341 | 170.5456766 | 222.0091691 | 501.669869 | 475.0496041 | 432.6010242 | 1.17E-16 | 1.422186558 | 1.47E-17 | 8.529728697 | 2.679913728 |
| CNAG_01780 | - | cytoplasmic protein | 5441.325506 | 4852.679443 | 5376.909924 | 3502.639203 | 3661.341389 | 3495.009826 | 1.18E-16 | -0.57188733 | 1.49E-17 | -8.528219356 | 0.672736141 |
| CNAG_01862 | - | hexose transporter | 13.66177938 | 20.84053875 | 37.58827315 | 149.3521762 | 134.3833566 | 150.2474308 | 1.40E-16 | 2.614028083 | 1.76E-17 | 8.508535453 | 6.122106248 |
| CNAG_06277 | - | ribosomal RNA-processing protein 12 | 1869.934251 | 1590.101284 | 1425.774569 | 927.9224579 | 933.8010561 | 897.3195797 | 1.40E-16 | -0.840792898 | 1.77E-17 | -8.50808617 | 0.558336626 |
| CNAG_04838 | - | hypothetical protein | 2216.690342 | 2151.786624 | 2032.818763 | 3104.147435 | 3220.204039 | 3421.218301 | 1.43E-16 | 0.591591476 | 1.80E-17 | 8.506105067 | 1.506908143 |
| CNAG_05833 | - | allantoate transporter | 40.41358663 | 34.82814852 | 37.61806971 | 172.6900064 | 169.0570009 | 139.5046847 | 1.54E-16 | 2.096660434 | 1.94E-17 | 8.497262874 | 4.277181518 |
| CNAG_02136 | - | small subunit ribosomal protein S6 | 2336.334557 | 2334.640935 | 2270.369044 | 1616.139253 | 1549.750014 | 1515.605367 | 1.54E-16 | -0.583650991 | 1.95E-17 | -8.496832551 | 0.667272988 |
| CNAG_04889 | - | hypothetical protein | 228.1682763 | 187.7588529 | 166.6425558 | 452.2422797 | 445.5687758 | 473.5515367 | 1.56E-16 | 1.221908684 | 1.98E-17 | 8.495012901 | 2.332551096 |
| CNAG_03885 | - | cytoplasmic protein | 2823.138915 | 2477.998111 | 2676.652539 | 1703.972542 | 1800.783262 | 1653.29239 | 1.61E-16 | -0.645382361 | 2.04E-17 | -8.491231782 | 0.639323327 |
| CNAG_06958 | - | hypothetical protein | 14.77669575 | 22.95275732 | 7.77516507 | 94.10021752 | 138.3909736 | 122.4792382 | 1.62E-16 | 3.027486935 | 2.06E-17 | 8.490525763 | 8.153881186 |
| CNAG_05070 | - | sulfite reductase (NADPH) hemoprotein, beta- component | 4782.452242 | 4007.928698 | 4054.543483 | 2546.778896 | 2891.078401 | 2434.894902 | 1.72E-16 | -0.722643645 | 2.18E-17 | -8.483933199 | 0.605985994 |
| CNAG_06441 | - | isoleucine-tRNA ligase | 2434.449466 | 2444.545298 | 2410.068546 | 1600.471397 | 1659.311064 | 1700.951467 | 1.78E-16 | -0.570671677 | 2.26E-17 | -8.479802657 | 0.673303245 |
| CNAG_06954 | - | hypothetical protein | 14.77525334 | 22.95172916 | 7.774806179 | 95.71693658 | 136.6459436 | 120.282057 | 1.83E-16 | 3.015208456 | 2.33E-17 | 8.476059349 | 8.084779662 |
| CNAG_00513 | - | nucleolar protein 53 | 1579.514431 | 1524.393269 | 1493.969161 | 878.8528369 | 948.5032247 | 1015.994892 | 2.01E-16 | -0.709279734 | 2.56E-17 | -8.46523226 | 0.611625417 |

| | | | | | | | | | | | | | |
|---|---|---|---|---|---|---|---|---|---|---|---|---|---|
| CNAG_05378 | - | 3-hydroxyacyl-CoA dehydrogenase, variant | 227.1837817 | 265.2878046 | 247.6455584 | 494.692955 | 526.5697352 | 654.2998704 | 2.10E-16 | 1.165225077 | 2.67E-17 | 8.459968958 | 2.242682013 |
| CNAG_03810 | - | large subunit ribosomal protein L13 | 2504.504995 | 2425.181596 | 2572.134872 | 1688.465596 | 1741.279653 | 1613.310361 | 2.15E-16 | -0.58862481 | 2.75E-17 | -8.4569148 | 0.664976467 |
| CNAG_00653 | - | hypothetical protein, variant 2 | 47.55723751 | 71.44527818 | 91.94777623 | 228.9120041 | 270.8831631 | 249.2182543 | 2.18E-16 | 1.822882218 | 2.79E-17 | 8.45520065 | 3.537872897 |
| CNAG_04962 | - | hypothetical protein | 3334.793381 | 3234.33929 | 3426.370415 | 2276.080315 | 2198.374029 | 2373.696865 | 2.27E-16 | -0.560920646 | 2.90E-17 | -8.4503731 | 0.677869448 |
| CNAG_05721 | MFE2 | multifunctional beta-oxidation protein | 1796.343503 | 1846.860695 | 1732.072249 | 2619.284339 | 2906.360829 | 2660.867368 | 2.31E-16 | 0.591764941 | 2.95E-17 | 8.448479197 | 1.507089339 |
| CNAG_00520 | - | hypothetical protein | 5664.503824 | 5627.314771 | 5982.654192 | 3843.510128 | 3887.285353 | 4250.952549 | 2.52E-16 | -0.543137764 | 3.22E-17 | -8.438141423 | 0.686276679 |
| CNAG_04768 | - | hypothetical protein | 509.5595161 | 511.9718799 | 473.7007385 | 921.4451987 | 862.0259478 | 872.3076001 | 2.53E-16 | 0.815395676 | 3.24E-17 | 8.437621722 | 1.759780726 |
| CNAG_04530 | - | DNA-binding protein | 2672.731399 | 2793.623122 | 2735.333839 | 1822.294151 | 1896.525895 | 1934.08096 | 2.63E-16 | -0.551909107 | 3.37E-17 | -8.432964061 | 0.682116891 |
| CNAG_02557 | - | fungal specific transcription factor | 647.5321623 | 627.9125232 | 650.3064129 | 277.014081 | 320.4527496 | 362.1684228 | 2.68E-16 | -1.019813664 | 3.44E-17 | -8.430677496 | 0.493180046 |
| CNAG_04551 | - | hypothetical protein | 18.24126211 | 3.573128165 | 16.29296154 | 116.7256071 | 117.4084253 | 105.9870665 | 2.88E-16 | 3.173342996 | 3.70E-17 | 8.422154572 | 9.021347838 |
| CNAG_02225 | EXG104 | glucan 1,3-beta-glucosidase | 30.00369793 | 55.23373204 | 26.99042734 | 184.3582949 | 158.4309972 | 183.5774869 | 2.88E-16 | 2.228988588 | 3.71E-17 | 8.421848166 | 4.688052051 |
| CNAG_03245 | ZWF1 | glucose-6-phosphate dehydrogenase | 14130.63142 | 13650.9462 | 12934.94971 | 9777.285827 | 9848.497558 | 10062.6607 | 2.97E-16 | -0.470995517 | 3.83E-17 | -8.418127219 | 0.721466585 |
| CNAG_01136 | - | hypothetical protein | 1949.70013 | 1741.034241 | 1955.696017 | 1177.046638 | 1230.603666 | 1170.365373 | 3.00E-16 | -0.674097443 | 3.86E-17 | -8.416934741 | 0.626724179 |
| CNAG_04273 | - | hypothetical protein | 2005.589285 | 1748.534307 | 1817.085485 | 1180.228366 | 967.3499297 | 1133.722907 | 3.13E-16 | -0.779554941 | 4.04E-17 | -8.411704667 | 0.582546476 |
| CNAG_04066 | - | endoribonuclease L-PSP | 4268.884764 | 3983.143147 | 3876.472641 | 2791.321544 | 2866.929643 | 2701.249763 | 3.23E-16 | -0.552567277 | 4.18E-17 | -8.407873098 | 0.681805774 |
| CNAG_05750 | ATP1 | ATP synthase subunit alpha, mitochondrial | 55366.40877 | 54237.86314 | 57593.6937 | 40234.88557 | 41741.71747 | 42441.6011 | 3.29E-16 | -0.441786206 | 4.25E-17 | -8.405877914 | 0.736222524 |
| CNAG_04719 | - | solute carrier family 41 | 607.6688975 | 650.9365569 | 588.8684869 | 1107.436661 | 1067.357408 | 1004.485859 | 3.38E-16 | 0.769537782 | 4.37E-17 | 8.402457051 | 1.704723528 |
| CNAG_07649 | - | hypothetical protein | 3938.481437 | 3659.946824 | 4023.572939 | 2748.571596 | 2612.427713 | 2540.286684 | 3.43E-16 | -0.572351395 | 4.43E-17 | -8.400824998 | 0.672519779 |
| CNAG_01189 | - | DNA-directed RNA polymerase I subunit RPA1 | 4712.635945 | 4407.651587 | 4498.176913 | 2792.721099 | 2986.763308 | 3277.345917 | 3.51E-16 | -0.604071564 | 4.55E-17 | -8.397748773 | 0.657894627 |
| CNAG_03951 | - | hypothetical protein | 3163.25646 | 3331.288566 | 4077.816174 | 2130.42338 | 2222.528036 | 2062.968146 | 4.01E-16 | -0.736271939 | 5.20E-17 | -8.382084604 | 0.600288552 |
| CNAG_07884 | BOT1 | Eukaryotic mitochondrial regulator protein, putative | 6337.156912 | 6275.944674 | 6575.628913 | 4706.761912 | 4650.706921 | 4642.872998 | 4.03E-16 | -0.470256004 | 5.23E-17 | -8.381415283 | 0.721836498 |
| CNAG_05903 | - | hypothetical protein | 483.9241606 | 496.9034257 | 523.7842279 | 858.9655367 | 919.933091 | 877.8528058 | 4.08E-16 | 0.805999072 | 5.30E-17 | 8.379811388 | 1.748356117 |
| CNAG_04602 | - | hypothetical protein | 88.31744168 | 126.2984877 | 97.32077362 | 284.2209737 | 291.3042892 | 293.6839719 | 5.00E-16 | 1.466765807 | 6.50E-17 | 8.35571619 | 2.764015693 |
| CNAG_04789 | - | hypothetical protein | 243.5046594 | 260.9965114 | 255.1110197 | 684.0251049 | 562.8681601 | 480.5397689 | 5.07E-16 | 1.172073582 | 6.60E-17 | 8.353996058 | 2.253353383 |
| CNAG_05521 | - | aldose reductase | 1659.65961 | 1526.899437 | 1324.745715 | 2591.591301 | 2478.466155 | 2392.184023 | 5.22E-16 | 0.711128097 | 6.80E-17 | 8.350388229 | 1.637083713 |
| CNAG_03675 | - | pumilio domain-containing protein | 5301.1303 | 4488.5271 | 5333.143811 | 3128.191917 | 3343.882021 | 3324.041016 | 5.33E-16 | -0.642694161 | 6.95E-17 | -8.347917141 | 0.6405157 |
| CNAG_06456 | - | hypothetical protein | 4022.626393 | 3856.040358 | 4506.621874 | 2734.464588 | 2652.930958 | 2816.676574 | 5.34E-16 | -0.609959088 | 6.97E-17 | -8.347459473 | 0.655215282 |
| CNAG_05676 | - | tyrosine aminotransferase | 174.5708283 | 180.1859972 | 162.3502084 | 364.0105291 | 399.8793733 | 445.6926672 | 5.36E-16 | 1.212247048 | 7.01E-17 | 8.346912368 | 2.316982341 |
| CNAG_12912 | - | unspecified product | 49.71710079 | 45.57990426 | 39.75829853 | 170.8168326 | 162.774472 | 178.1005942 | 5.42E-16 | 1.923887862 | 7.09E-17 | 8.345478504 | 3.794442314 |
| CNAG_12916 | - | unspecified product | 35.77289729 | 45.55575298 | 21.65871928 | 148.0928221 | 158.6031169 | 174.6345906 | 5.45E-16 | 2.234769091 | 7.13E-17 | 8.344871236 | 4.706873535 |
| CNAG_03368 | STU2 | microtubule Associated protein, variant | 2591.497911 | 2378.061195 | 2406.057343 | 3565.711308 | 3720.524228 | 3627.802298 | 5.61E-16 | 0.549571899 | 7.36E-17 | 8.341154078 | 1.463651312 |
| CNAG_06135 | - | hypothetical protein | 64.83642391 | 47.75268197 | 40.84010198 | 180.7490733 | 187.8490136 | 206.8969715 | 5.85E-16 | 1.908186443 | 7.67E-17 | 8.336229212 | 3.753369812 |
| CNAG_05930 | - | hypothetical protein | 694.0560722 | 680.0598574 | 725.3317495 | 1185.148298 | 1187.253166 | 1123.266125 | 5.88E-16 | 0.720162621 | 7.71E-17 | 8.335559198 | 1.647367716 |
| CNAG_02585 | - | hypothetical protein | 2702.487618 | 2754.048867 | 2489.263271 | 3728.272994 | 4149.264034 | 4049.625749 | 6.23E-16 | 0.570867309 | 8.18E-17 | 8.328544119 | 1.485416294 |
| CNAG_00906 | - | nucleolar pre-ribosomal-associated protein 1, variant | 5768.151293 | 4999.17929 | 5026.098547 | 3269.05307 | 3629.422412 | 3512.764479 | 6.23E-16 | -0.617250272 | 8.20E-17 | -8.328368971 | 0.651912267 |
| CNAG_01019 | SOD1 | superoxide dismutase [Cu-Zn] | 16079.71915 | 16879.98247 | 16774.12052 | 12641.52877 | 10857.6078 | 11262.80389 | 6.37E-16 | -0.531678426 | 8.39E-17 | -8.325592405 | 0.691749487 |
| CNAG_04334 | MRPS18 | hypothetical protein | 6801.356792 | 6388.880448 | 7369.851866 | 2988.664829 | 2015.127296 | 2391.768486 | 6.57E-16 | -1.490598631 | 8.66E-17 | -8.321884079 | 0.355864856 |
| CNAG_03160 | - | DNA cross-link repair 1A protein, variant | 438.4914961 | 547.5062697 | 519.5319186 | 890.132884 | 953.5977849 | 910.0287018 | 6.63E-16 | 0.857639107 | 8.75E-17 | 8.320620192 | 1.812070527 |
| CNAG_01345 | - | small subunit ribosomal protein S2 | 650.08434 | 689.3346254 | 732.4065613 | 390.1435399 | 364.4619364 | 368.8868277 | 6.84E-16 | -0.899339609 | 9.03E-17 | -8.316854789 | 0.536132089 |
| CNAG_04825 | - | hypothetical protein | 437.1845463 | 437.6427461 | 400.1300249 | 861.7072471 | 740.3596129 | 779.0352301 | 6.93E-16 | 0.886570369 | 9.16E-17 | 8.315200813 | 1.848775914 |
| CNAG_06101 | - | ADP,ATP carrier protein | 115783.5145 | 109202.01 | 114067.7539 | 84259.70214 | 84057.22576 | 86241.89589 | 6.93E-16 | -0.429043303 | 9.16E-17 | -8.315254443 | 0.742754166 |
| CNAG_01055 | - | phospholipid transporter, variant | 2587.493688 | 2761.283618 | 2608.441421 | 1805.202206 | 1847.362137 | 1857.490008 | 6.97E-16 | -0.54518696 | 9.22E-17 | -8.314415667 | 0.685302588 |
| CNAG_01121 | - | hypothetical protein | 26.51047436 | 32.68764145 | 52.5044156 | 183.662795 | 175.9543506 | 149.4575618 | 7.02E-16 | 2.189063719 | 9.30E-17 | 8.313446092 | 4.56009449 |
| CNAG_01184 | - | hypothetical protein | 823.711393 | 939.6573897 | 724.3470007 | 1461.464788 | 1522.015519 | 1379.708052 | 7.04E-16 | 0.796573955 | 9.34E-17 | 8.312902101 | 1.736971346 |
| CNAG_03682 | FRR1 | FK506-binding protein 1 | 9434.048791 | 9429.587011 | 9567.054629 | 7114.806706 | 7202.19829 | 6997.519974 | 7.29E-16 | -0.430996673 | 9.67E-17 | -8.308725772 | 0.741749178 |
| CNAG_00894 | - | hypothetical protein | 8255.827225 | 8445.888901 | 9107.360503 | 6232.969372 | 6222.292073 | 6177.121221 | 7.37E-16 | -0.485332754 | 9.80E-17 | -8.307225491 | 0.714332292 |
| CNAG_02418 | - | asparagine-tRNA ligase | 10382.36916 | 12333.15189 | 11386.40552 | 7643.130895 | 8106.988518 | 8014.440881 | 7.46E-16 | -0.535573685 | 9.92E-17 | -8.305692817 | 0.689884291 |
| CNAG_04691 | - | hypothetical protein | 4.230856027 | 11.12658498 | 18.41363538 | 133.770009 | 111.334242 | 82.96315259 | 7.62E-16 | 3.289626122 | 1.01E-16 | 8.303061311 | 9.77858775 |
| CNAG_04844 | - | mitochondrial import receptor subunit Tom22 | 9037.463698 | 8440.75297 | 8651.183843 | 11976.64911 | 12418.54982 | 11992.90958 | 7.72E-16 | 0.462163857 | 1.03E-16 | 8.301496274 | 1.377606502 |
| CNAG_05672 | - | Nucleolar protein | 2024.411283 | 1855.208148 | 1880.034477 | 1296.235668 | 1183.963152 | 1233.624959 | 7.77E-16 | -0.649130858 | 1.04E-16 | -8.300579864 | 0.637664354 |
| CNAG_06748 | - | U3 small nucleolar RNA-associated protein 7 | 3211.057703 | 3167.543557 | 3400.76881 | 2233.268438 | 2295.902952 | 2278.244853 | 7.85E-16 | -0.538140809 | 1.05E-16 | -8.299173222 | 0.688657806 |
| CNAG_04630 | YAP2 | hypothetical protein | 343.710425 | 284.7048599 | 262.597609 | 627.3362209 | 634.5083132 | 588.0811709 | 8.02E-16 | 1.040633794 | 1.07E-16 | 8.296534678 | 2.057131179 |
| CNAG_01818 | - | ATP synthase mitochondrial F1 complex assembly factor 2 | 1816.45718 | 1562.167891 | 1661.39052 | 1023.698803 | 1014.89736 | 1082.628096 | 8.46E-16 | -0.707860253 | 1.13E-16 | -8.290119365 | 0.612227497 |
| CNAG_04603 | - | replication fork protection complex subunit Csm3/Swi3 | 194.4171481 | 198.5035832 | 197.515223 | 424.1085741 | 404.0350316 | 482.3546235 | 8.93E-16 | 1.138285462 | 1.20E-16 | 8.283570301 | 2.20119272 |
| CNAG_03913 | - | hypothetical protein, variant | 288.9526045 | 313.7361976 | 288.1487744 | 568.0127487 | 573.1530645 | 592.4458761 | 9.11E-16 | 0.948145351 | 1.22E-16 | 8.281135263 | 1.929390744 |
| CNAG_04155 | - | small nuclear ribonucleoprotein E | 3766.830472 | 3509.107674 | 3772.976745 | 2555.500669 | 2615.921459 | 2526.936641 | 9.19E-16 | -0.537133993 | 1.23E-16 | -8.279915534 | 0.689138569 |
| CNAG_04759 | - | hypothetical protein | 33.39182491 | 20.8392026 | 26.96100805 | 152.5832046 | 153.6753335 | 112.9277401 | 9.22E-16 | 2.383377004 | 1.24E-16 | 8.279402427 | 5.217566214 |
| CNAG_00741 | - | ATP-dependent RNA helicase SUB2 | 13913.51564 | 13631.57779 | 14021.62316 | 10067.59356 | 10723.20062 | 9907.274104 | 9.29E-16 | -0.452781069 | 1.25E-16 | -8.278456376 | 0.730633056 |
| CNAG_12489 | - | unspecified product | 679.4076676 | 774.4433557 | 895.4787845 | 406.3951267 | 397.2200785 | 411.0747239 | 9.42E-16 | -0.966608943 | 1.27E-16 | -8.276726004 | 0.511707419 |
| CNAG_05504 | - | GDSL Lipase/Acylhydrolase | 1682.368096 | 1606.328587 | 1812.760565 | 1039.268093 | 994.1665621 | 1130.340471 | 9.80E-16 | -0.704603603 | 1.32E-16 | -8.271916035 | 0.613611062 |
| CNAG_04362 | - | ATP-dependent RNA helicase DBP5 | 2940.220375 | 3203.043681 | 3212.015372 | 2122.745022 | 2225.189639 | 2123.951517 | 9.83E-16 | -0.546978402 | 1.32E-16 | -8.271363229 | 0.684452153 |
| CNAG_03144 | - | peptide-methionine (S)-S-oxide reductase | 2129.656835 | 2041.624972 | 2263.899716 | 1441.755285 | 1363.396233 | 1425.662619 | 1.03E-15 | -0.620631334 | 1.38E-16 | -8.266074917 | 0.650386252 |
| CNAG_00598 | - | nicotinamide mononucleotide permease | 19.44602283 | 20.82139231 | 25.87847669 | 147.4787122 | 120.6306932 | 103.9916944 | 1.07E-15 | 2.516041586 | 1.45E-16 | 8.260545875 | 5.720104828 |
| CNAG_06695 | - | Macrophage activating regulator of cell wall-1 | 118.6985719 | 139.2990184 | 179.3561869 | 388.5836283 | 343.6783177 | 422.3336887 | 1.07E-15 | 1.388212908 | 1.45E-16 | 8.260428389 | 2.617542402 |
| CNAG_02120 | - | TIGR01456 family HAD hydrolase | 2510.350362 | 2445.6501 | 2513.504039 | 1721.955381 | 1755.946241 | 1666.577197 | 1.16E-15 | -0.553835172 | 1.57E-16 | -8.250825627 | 0.681206841 |
| CNAG_01049 | - | H/ACA ribonucleoprotein complex subunit 3 | 1521.197371 | 1412.407417 | 1759.359012 | 925.6311073 | 827.6338509 | 952.7618723 | 1.19E-15 | -0.810700205 | 1.61E-16 | -8.248400863 | 0.570105093 |
| CNAG_06923 | XFP2 | xylulose 5-phosphate/fructose 6-phosphate phosphoketolase | 9098.487676 | 6563.375041 | 5627.526074 | 2093.319936 | 2072.073432 | 2093.179577 | 1.19E-15 | -1.782826855 | 1.60E-16 | -8.248444481 | 0.290613403 |
| CNAG_01106 | VPH1 | V-type H -transporting ATPase subunit I | 5341.084707 | 5466.77206 | 5491.061824 | 3992.284847 | 3920.969818 | 4047.805925 | 1.42E-15 | -0.46171756 | 1.93E-16 | -8.226404184 | 0.72612128 |
| CNAG_05196 | - | nuclear cap-binding protein subunit 2 | 1051.717137 | 1084.744848 | 990.6160952 | 619.7516178 | 603.4314089 | 656.3333776 | 1.44E-15 | -0.749587497 | 1.96E-16 | -8.22461749 | 0.594773594 |
| CNAG_03733 | - | hypothetical protein | 43.90171422 | 33.75960616 | 46.12646432 | 173.3768039 | 152.3927578 | 163.7276606 | 1.51E-15 | 1.984139544 | 2.05E-16 | 8.219169363 | 3.956266316 |
| CNAG_01981 | - | sulfide:quinone oxidoreductase | 967.3310158 | 841.2879321 | 806.1141391 | 510.8233702 | 468.8419401 | 448.8451397 | 1.61E-15 | -0.890020163 | 2.18E-16 | -8.211562837 | 0.539606577 |
| CNAG_06756 | - | pre-rRNA-processing protein ESF2 | 1186.118174 | 1186.106243 | 1308.32642 | 734.934264 | 779.4688413 | 751.8354254 | 1.64E-15 | -0.716253535 | 2.22E-16 | -8.209327859 | 0.608676033 |
| CNAG_02021 | - | N-terminal acetyltransferase A complex catalytic subunit ard1 | 1566.702152 | 1471.639602 | 1515.291931 | 1002.00622 | 958.8578526 | 976.0662867 | 1.64E-15 | -0.648115187 | 2.24E-16 | -8.208356951 | 0.638113434 |
| CNAG_05015 | CAT4 | catalase 4 | 52.05869796 | 41.29522331 | 53.58726223 | 174.6293578 | 175.7749592 | 181.4769511 | 1.66E-15 | 1.844157331 | 2.26E-16 | 8.207645048 | 3.59043173 |
| CNAG_01679 | - | small subunit ribosomal protein S15 | 2236.033513 | 2406.816628 | 2378.057496 | 1603.643354 | 1585.982664 | 1623.25038 | 1.67E-15 | -0.560066423 | 2.28E-16 | -8.206343642 | 0.678270935 |
| CNAG_04508 | - | hypothetical protein | 64.94310806 | 56.38341406 | 84.47848283 | 213.4030321 | 243.2334095 | 213.8005191 | 1.75E-15 | 1.697377838 | 2.39E-16 | 8.200637458 | 3.243109729 |
| CNAG_04775 | - | hypothetical protein | 630.9249712 | 580.9350657 | 518.5107424 | 1024.076025 | 1016.46998 | 1005.523144 | 1.89E-15 | 0.80193124 | 2.59E-16 | 8.191036758 | 1.743433384 |
| CNAG_06229 | - | hypothetical protein | 2023.35549 | 1974.79149 | 1987.741706 | 1265.764877 | 1406.624786 | 1303.561483 | 1.89E-15 | -0.605989452 | 2.59E-16 | -8.191130993 | 0.657020617 |
| CNAG_02482 | - | translation machinery-associated protein 22 | 1745.300924 | 1543.855585 | 1734.931441 | 1076.71125 | 1064.114587 | 936.1857337 | 1.90E-15 | -0.723191811 | 2.60E-16 | -8.190662836 | 0.605755788 |
| CNAG_03897 | - | protein BCP1 | 1538.474314 | 1266.96862 | 1424.606376 | 844.6912869 | 810.4413331 | 859.5157565 | 1.92E-15 | -0.766590276 | 2.64E-16 | -8.188925371 | 0.587805076 |
| CNAG_07496 | - | hypothetical protein | 4.181616403 | 6.789251956 | 9.872747602 | 88.00651805 | 75.7346203 | 82.58384042 | 1.92E-15 | 3.673870104 | 2.64E-16 | 8.188944404 | 12.76277467 |
| CNAG_04846 | - | membrane protein | 1137.679409 | 1021.597849 | 965.3285192 | 1626.239261 | 1775.549416 | 1721.586079 | 2.03E-15 | 0.698135148 | 2.79E-16 | 8.182163656 | 1.622406287 |
| CNAG_05913 | - | Alpha-glucosidase | 608.9052333 | 524.5588673 | 607.6182076 | 270.0892313 | 284.212509 | 314.4585191 | 2.04E-15 | -1.01832411 | 2.81E-16 | -8.181190695 | 0.493689508 |
| CNAG_00781 | - | U6 snRNA-associated Sm-like protein LSm3 | 1609.94915 | 1529.827386 | 1754.087154 | 1082.221315 | 963.1328639 | 901.7736931 | 2.04E-15 | -0.747301982 | 2.81E-16 | -8.181211167 | 0.59571658 |
| CNAG_00517 | - | hypothetical protein | 1759.644928 | 1986.598786 | 2136.945882 | 1227.641971 | 1277.181323 | 1190.363573 | 2.06E-15 | -0.685881184 | 2.83E-16 | -8.180343971 | 0.621626027 |
| CNAG_02088 | - | hypothetical protein | 1539.889477 | 1535.170801 | 1529.151223 | 993.4254548 | 1033.971417 | 886.2150817 | 2.08E-15 | -0.675997457 | 2.87E-16 | -8.178659062 | 0.625899333 |
| CNAG_03684 | - | ATP-binding cassette, subfamily B (MDR/TAP), member 10 | 5272.018756 | 4947.454651 | 4932.250574 | 3305.678042 | 3595.792829 | 3659.255187 | 2.13E-15 | -0.536452738 | 2.93E-16 | -8.175983208 | 0.689464063 |

| | | | | | | | | | | | | | |
|---|---|---|---|---|---|---|---|---|---|---|---|---|---|
| CNAG_03772 | HXS1 | high-affinity glucose transporter | 409.4661778 | 262.2434025 | 241.3925557 | 1219.800305 | 1355.080538 | 1579.850283 | 2.13E-15 | 2.170150581 | 2.93E-16 | 8.176058687 | 4.500703676 |
| CNAG_02381 | - | hypothetical protein | 5.380260486 | 11.10477914 | 12.01255963 | 93.31181575 | 94.96573102 | 82.73475771 | 2.18E-15 | 3.24962851 | 3.01E-16 | 8.17293841 | 9.511207496 |
| CNAG_05713 | - | elongator complex protein 1 | 967.2364326 | 903.0804629 | 923.703602 | 1413.717594 | 1565.940006 | 1525.086592 | 2.20E-15 | 0.673509956 | 3.04E-16 | 8.171546487 | 1.594948627 |
| CNAG_03135 | - | hypothetical protein | 1245.300563 | 1509.599831 | 1381.201427 | 2147.759258 | 2179.137558 | 2180.108661 | 2.22E-15 | 0.639428063 | 3.08E-16 | 8.17027396 | 1.557711503 |
| CNAG_07502 | - | CTP synthase | 9310.128434 | 8464.232307 | 10114.02674 | 5403.820201 | 6429.307116 | 6181.588295 | 2.23E-15 | -0.646609394 | 3.09E-16 | -8.169620081 | 0.638779804 |
| CNAG_01919 | - | hypothetical protein | 248.1668464 | 313.7136391 | 263.634355 | 577.5455028 | 542.9482145 | 559.1594902 | 2.28E-15 | 1.01173428 | 3.16E-16 | 8.16711433 | 2.016333501 |
| CNAG_06443 | SSA1 | glucose-regulated protein | 39919.37147 | 38712.0657 | 42087.44546 | 28129.31732 | 29700.35722 | 30440.75963 | 2.33E-15 | -0.467149307 | 3.23E-16 | -8.164439771 | 0.723392574 |
| CNAG_05711 | - | hypothetical protein | 675.6379507 | 633.3684734 | 730.2527552 | 391.7411131 | 348.921629 | 337.8402307 | 2.52E-15 | -0.937096058 | 3.50E-16 | -8.154807816 | 0.522283105 |
| CNAG_01657 | - | fumarate hydratase, mitochondrial | 5508.056775 | 5517.423621 | 5689.411531 | 3951.757692 | 4061.580944 | 4173.246918 | 2.54E-15 | -0.471211192 | 3.53E-16 | -8.153754837 | 0.721358738 |
| CNAG_07838 | - | cytoplasmic protein | 558.6174065 | 541.0856188 | 592.02767 | 1007.016729 | 944.8136877 | 955.5850656 | 2.61E-15 | 0.76577649 | 3.64E-16 | 8.15008896 | 1.700284882 |
| CNAG_04308 | HAD1 | 3-hydroxyacyl-CoA dehydrogenase | 2015.141374 | 1862.772084 | 2021.83466 | 1337.490921 | 1294.42921 | 1208.127578 | 2.71E-15 | -0.635196784 | 3.78E-16 | -8.145535887 | 0.643852987 |
| CNAG_01788 | - | tRNA(His) guanylyltransferase | 3743.491735 | 3635.136425 | 3756.984776 | 2606.120092 | 2363.951463 | 2670.10207 | 2.90E-15 | -0.558734212 | 4.04E-16 | -8.137457633 | 0.678897552 |
| CNAG_06631 | - | myosin heavy chain | 2857.209065 | 3032.784286 | 2806.808394 | 1882.968411 | 1935.307434 | 2099.470062 | 2.97E-15 | -0.570393972 | 4.14E-16 | -8.134412651 | 0.673432862 |
| CNAG_01828 | - | compass component swd1 | 1545.839728 | 1592.295128 | 1743.442898 | 1058.031832 | 1064.131558 | 959.4711755 | 3.25E-15 | -0.678853622 | 4.53E-16 | -8.123452929 | 0.624661439 |
| CNAG_02686 | - | Cystathionine beta-lyase | 3984.257315 | 4770.64707 | 4478.967681 | 2929.803173 | 3086.01168 | 2977.667221 | 3.34E-15 | -0.571843224 | 4.66E-16 | -8.120027249 | 0.672756708 |
| CNAG_04399 | - | YggS family pyridoxal phosphate enzyme | 1122.81453 | 1037.390108 | 1023.654322 | 660.3077024 | 639.7050243 | 554.3178575 | 3.34E-15 | -0.795727715 | 4.67E-16 | -8.119670758 | 0.576052531 |
| CNAG_04680 | - | hypothetical protein | 180.4103873 | 199.5530503 | 173.006586 | 440.7803233 | 379.0189767 | 410.358729 | 3.79E-15 | 1.140079055 | 5.31E-16 | 8.10417399 | 2.203930997 |
| CNAG_07373 | - | Carbamoyl-phosphate synthase, large subunit | 16642.26766 | 15118.38369 | 13503.39325 | 10058.94855 | 10319.42551 | 10283.6288 | 3.97E-15 | -0.577518797 | 5.57E-16 | -8.098445665 | 0.670115277 |
| CNAG_01905 | KSP1 | serine/threonine protein kinase | 1613.274792 | 1473.7637 | 1406.549534 | 911.6264751 | 989.9714481 | 869.5521985 | 4.07E-15 | -0.713709493 | 5.71E-16 | -8.095309227 | 0.609750317 |
| CNAG_03632 | - | hypothetical protein | 1699.824262 | 1608.465912 | 1605.959304 | 1068.133101 | 1034.758303 | 1105.920585 | 4.18E-15 | -0.630444305 | 5.87E-16 | -8.091995543 | 0.645977444 |
| CNAG_03467 | - | hypothetical protein | 347.3174043 | 353.6186853 | 364.9043954 | 662.1840468 | 625.6503885 | 712.2947422 | 4.40E-15 | 0.894741697 | 6.18E-16 | 8.085712049 | 1.859276967 |
| CNAG_03213 | UVE1 | UV damage endonuclease UvdE | 1587.447516 | 1983.647828 | 1638.246531 | 2754.080761 | 2753.598145 | 2776.258117 | 4.50E-15 | 0.655286451 | 6.32E-16 | 8.082889636 | 1.574928628 |
| CNAG_01719 | - | hypothetical protein | 44.05132731 | 80.02216128 | 79.17083043 | 208.5676437 | 248.3865392 | 244.7103337 | 4.62E-15 | 1.772001219 | 6.49E-16 | 8.079693212 | 3.415273742 |
| CNAG_03459 | - | T-complex protein 1 subunit delta | 8755.54296 | 8451.29103 | 9041.261233 | 6087.360008 | 6660.529808 | 6215.986114 | 4.81E-15 | -0.48465625 | 6.77E-16 | -8.074520147 | 0.714667333 |
| CNAG_05726 | EMC3 | hypothetical protein | 1621.723058 | 1675.244747 | 1759.465804 | 1043.164085 | 1009.705529 | 1161.402768 | 4.97E-15 | -0.668804908 | 7.00E-16 | -8.07056627 | 0.629027542 |
| CNAG_04333 | - | hypothetical protein | 24.17350835 | 34.82757347 | 49.30620574 | 183.8403687 | 151.5823697 | 148.2920477 | 5.03E-15 | 2.15659137 | 7.10E-16 | 8.068860495 | 4.458601839 |
| CNAG_02919 | - | hypothetical protein | 106.8419421 | 88.66846211 | 101.5556629 | 263.2143519 | 262.7910487 | 265.9632648 | 5.45E-15 | 1.405363553 | 7.69E-16 | 8.05907655 | 2.648845234 |
| CNAG_01651 | - | glicosidase | 91.77794496 | 99.42419996 | 115.3954931 | 293.8161345 | 274.8840456 | 254.9822801 | 5.45E-15 | 1.413365745 | 7.70E-16 | 8.05890833 | 2.663578399 |
| CNAG_07410 | CID1alpha | CID1-family polyA polymerase | 3541.53437 | 3582.332978 | 3577.840664 | 2428.593755 | 2642.697126 | 2548.003603 | 5.53E-15 | -0.50540656 | 7.83E-16 | -8.056884891 | 0.704461828 |
| CNAG_06558 | - | NADPH-ferrihemoprotein reductase | 1388.350184 | 1795.775797 | 1693.323411 | 968.4319902 | 1008.036863 | 911.7516113 | 5.55E-15 | -0.770473043 | 7.85E-16 | -8.056438006 | 0.586225227 |
| CNAG_05398 | - | Phosphoserine transaminase | 823.8138361 | 804.5821794 | 701.6209912 | 409.5038761 | 463.7579259 | 399.9968302 | 5.84E-15 | -0.888500937 | 8.27E-16 | -8.050070556 | 0.540175107 |
| CNAG_05397 | - | hypothetical protein, variant | 100.9767538 | 70.37972005 | 81.3219006 | 231.9500429 | 269.0561897 | 244.8685787 | 5.94E-15 | 1.551183884 | 8.42E-16 | 8.04793554 | 2.930575253 |
| CNAG_02035 | - | triose-phosphate isomerase | 9437.548062 | 9621.349844 | 9337.784922 | 7336.678617 | 7023.622118 | 6906.496632 | 5.96E-15 | -0.432293652 | 8.46E-16 | -8.047323105 | 0.741082647 |
| CNAG_06946 | - | hypothetical protein | 1947.389066 | 1792.733286 | 1854.437029 | 1223.009639 | 1253.901776 | 1243.596812 | 5.97E-15 | -0.604314616 | 8.48E-16 | -8.047062873 | 0.657783801 |
| CNAG_01702 | - | Integral membrane protein | 407.0129454 | 561.4944622 | 483.2976409 | 920.612905 | 869.7754406 | 978.7317042 | 6.25E-15 | 0.917451085 | 8.89E-16 | 8.041342046 | 1.888775306 |
| CNAG_04715 | - | oligosaccharyltransferase complex subunit alpha (ribophorin I) | 795.7305435 | 782.4503127 | 967.3806978 | 1524.604595 | 1439.085539 | 1411.884772 | 6.30E-15 | 0.766307016 | 8.96E-16 | 8.040346857 | 1.700910248 |
| CNAG_06501 | - | 1,3-beta-glucanosyltransferase | 615.8207122 | 634.7841275 | 608.0484136 | 1107.460958 | 1041.450095 | 980.0849015 | 6.49E-15 | 0.737131582 | 9.24E-16 | 8.036494042 | 1.666858434 |
| CNAG_01538 | - | hypothetical protein | 58.09004097 | 95.14093019 | 126.0507233 | 317.2517357 | 314.7376693 | 281.5563662 | 6.61E-15 | 1.694261828 | 9.43E-16 | 8.034102334 | 3.236112648 |
| CNAG_04401 | - | alpha-soluble NSF attachment protein | 2128.335562 | 1905.866697 | 1976.013493 | 1359.276594 | 1255.565617 | 1311.325956 | 6.74E-15 | -0.6300601 | 9.62E-16 | -8.031625691 | 0.646149497 |
| CNAG_02687 | - | hypothetical protein | 3770.270097 | 3431.527001 | 3583.167206 | 2428.602711 | 2414.891837 | 2591.280516 | 6.87E-15 | -0.552455653 | 9.81E-16 | -8.0291655 | 0.681858529 |
| CNAG_12851 | - | unspecified product | 7.685214814 | 3.544424564 | 2.394219689 | 92.24960819 | 57.25064414 | 90.12183543 | 6.92E-15 | 4.300492531 | 9.89E-16 | 8.028268719 | 19.7050367 |
| CNAG_04080 | - | 3-hydroxyanthranilate 3,4-dioxygenase | 1815.547579 | 1875.6363 | 1825.641942 | 1187.967412 | 1158.096962 | 1297.94513 | 7.25E-15 | -0.61358388 | 1.04E-15 | -8.022460901 | 0.653571111 |
| CNAG_06144 | - | cytoplasmic protein | 17921.26753 | 17322.88509 | 18990.19885 | 13440.74294 | 13195.23437 | 13458.64489 | 7.51E-15 | -0.451304118 | 1.07E-15 | -8.018016842 | 0.73138142 |
| CNAG_12714 | - | unspecified product | 10782.1402 | 9251.685624 | 12981.3543 | 3811.124495 | 2617.137425 | 3572.992498 | 7.59E-15 | -1.738839029 | 1.09E-15 | -8.016628283 | 0.299610683 |
| CNAG_00101 | KEI1 | Inositolphosphorylceramide synthase subunit Kei1 domain-containing protein | 1700.045417 | 1915.46656 | 1871.467367 | 1245.658547 | 1114.946436 | 1193.648322 | 7.71E-15 | -0.641784977 | 1.10E-15 | -8.014596738 | 0.640919479 |
| CNAG_02463 | - | hypothetical protein | 29.96249168 | 42.3280763 | 13.13098703 | 191.2239637 | 155.1852779 | 128.441987 | 7.91E-15 | 2.483062614 | 1.13E-15 | 8.011318239 | 5.590830527 |
| CNAG_07977 | - | hypothetical protein | 1564.327203 | 1427.475439 | 1522.736586 | 988.7849901 | 976.1448246 | 868.450645 | 8.07E-15 | -0.68804828 | 1.16E-15 | -8.00869971 | 0.620692974 |
| CNAG_05472 | SPO11 | meiotic recombination protein SPO11, variant 2 | 27.68372682 | 47.72701126 | 52.51420844 | 167.6333764 | 195.9546733 | 156.1127412 | 8.09E-15 | 2.025036995 | 1.16E-15 | 8.008289702 | 4.070023134 |
| CNAG_01148 | FPR3 | FK506-binding protein 4 | 19807.03742 | 17488.82564 | 19450.91009 | 12890.37323 | 13708.46922 | 13660.7057 | 8.51E-15 | -0.511157288 | 1.22E-15 | -8.002043502 | 0.701659361 |
| CNAG_07730 | - | tricarboxylate transporter, variant | 43.855062 | 28.37507999 | 37.60771249 | 156.9779514 | 150.8157546 | 142.719406 | 8.66E-15 | 2.036123859 | 1.25E-15 | 7.999717242 | 4.101421055 |
| CNAG_02847 | - | thymidylate kinase | 890.539277 | 875.7592073 | 859.4116116 | 545.1144217 | 522.3833793 | 488.7791078 | 8.67E-15 | -0.770184527 | 1.25E-15 | -7.999473434 | 0.586342474 |
| CNAG_00434 | - | arginine N-methyltransferase 2 | 1014.204891 | 918.8879987 | 913.8162212 | 570.745602 | 578.4689428 | 535.399433 | 8.81E-15 | -0.773072318 | 1.27E-15 | -7.99741226 | 0.585169987 |
| CNAG_05158 | - | hypothetical protein, variant | 625.1275735 | 617.550634 | 543.0322358 | 995.1202006 | 1131.346943 | 992.2572479 | 9.41E-15 | 0.789343297 | 1.36E-15 | 7.989174028 | 1.728287581 |
| CNAG_05032 | - | hypothetical protein | 67.2143546 | 57.43949931 | 62.12911416 | 189.9495966 | 243.4323643 | 183.9279559 | 9.86E-15 | 1.709171407 | 1.42E-15 | 7.98338903 | 3.269729769 |
| CNAG_01032 | - | hypothetical protein | 59.01163052 | 47.740196 | 37.64021864 | 154.7721152 | 187.9691171 | 200.1941619 | 1.01E-14 | 1.903977722 | 1.47E-15 | 7.979846098 | 3.7424362 |
| CNAG_05431 | RIM101 | pH-response transcription factor pacC/RIM101 | 1109.670746 | 1045.283799 | 1058.06582 | 1640.302536 | 1671.088983 | 1632.832359 | 1.03E-14 | 0.606432053 | 1.49E-15 | 7.97816498 | 1.522489262 |
| CNAG_02779 | BNA6 | nicotinate-nucleotide diphosphorylase (carboxylating) | 2498.741307 | 2632.003125 | 2540.180365 | 1756.963821 | 1704.132584 | 1851.910325 | 1.03E-14 | -0.545141451 | 1.49E-15 | -7.97775362 | 0.685324205 |
| CNAG_04684 | SEC23 | protein transporter SEC23 | 1568.562581 | 1470.857517 | 1432.398386 | 2308.14451 | 2217.030272 | 2213.464342 | 1.08E-14 | 0.576618572 | 1.56E-15 | 7.971752203 | 1.491349683 |
| CNAG_04615 | - | L-aminoadipate-semialdehyde dehydrogenase | 204.8405804 | 170.5251347 | 188.9815818 | 436.0231454 | 428.3878783 | 381.6236772 | 1.09E-14 | 1.131556035 | 1.59E-15 | 7.969895148 | 2.190949202 |
| CNAG_00879 | - | Glutamate dehydrogenase | 8317.183173 | 7431.924793 | 6076.634893 | 4378.107834 | 4460.840061 | 4628.494073 | 1.11E-14 | -0.712055511 | 1.61E-15 | -7.967900662 | 0.610449767 |
| CNAG_02719 | - | hypothetical protein | 69.68479852 | 82.2187989 | 116.4440779 | 274.2742309 | 291.494029 | 244.98387 | 1.12E-14 | 1.58389897 | 1.63E-15 | 7.966771084 | 2.997789265 |
| CNAG_06338 | - | ABC transporter PMR5 | 658.9052513 | 517.4205696 | 515.315545 | 1000.642733 | 1061.393984 | 1049.873794 | 1.12E-14 | 0.863772921 | 1.63E-15 | 7.966548782 | 1.819791192 |
| CNAG_04821 | PAN3 | PAB-dependent poly(A)-specific ribonuclease subunit PAN3 | 225.9099778 | 240.4885399 | 207.1310459 | 505.2914455 | 502.5935182 | 421.6550299 | 1.15E-14 | 1.072381193 | 1.68E-15 | 7.963180891 | 2.10290138 |
| CNAG_06853 | - | hypothetical protein, variant 2 | 83.6587857 | 128.437903 | 95.1872553 | 287.442828 | 293.956521 | 266.0517622 | 1.33E-14 | 1.453274411 | 1.94E-15 | 7.944871982 | 2.738288418 |
| CNAG_04839 | - | exosome complex component MTR3, animal type | 715.0193331 | 639.1371114 | 708.2700997 | 1133.710387 | 1127.679908 | 1159.827618 | 1.35E-14 | 0.713867245 | 1.96E-15 | 7.943743261 | 1.640194887 |
| CNAG_04903 | - | hypothetical protein | 78.86995603 | 72.50104037 | 60.02247887 | 274.9773461 | 210.9830106 | 195.0884822 | 1.37E-14 | 1.677759289 | 2.00E-15 | 7.941280206 | 3.19930667 |
| CNAG_04195 | - | O-methyltransferase | 570.3850389 | 696.1641959 | 636.8359532 | 1050.39916 | 1115.701884 | 1071.037024 | 1.44E-14 | 0.752137446 | 2.10E-15 | 7.935430561 | 1.684286362 |
| CNAG_01737 | ERG25 | C-4 methyl sterol oxidase, putative | 1943.253995 | 1326.253783 | 1495.0164 | 870.3086379 | 796.5652225 | 847.3674556 | 1.46E-14 | -0.939582465 | 2.14E-15 | -7.933160609 | 0.521383754 |
| CNAG_06374 | - | malate dehydrogenase (oxaloacetate- decarboxylating)(NADP) | 722.1998636 | 707.6008649 | 613.1011126 | 339.4042676 | 400.7877599 | 346.7086311 | 1.46E-14 | -0.927655919 | 2.14E-15 | -7.932807943 | 0.525711821 |
| CNAG_02130 | NSR1 | RNA-binding protein 39 | 8164.777941 | 8630.116883 | 9323.836586 | 6007.96871 | 6277.497372 | 6373.599089 | 1.48E-14 | -0.500413442 | 2.17E-15 | -7.931334358 | 0.70690417 |
| CNAG_06537 | - | monocarboxylic acid transporter | 841.4869036 | 806.8256772 | 868.9470762 | 421.1031783 | 486.1458794 | 514.2343457 | 1.50E-14 | -0.839116207 | 2.20E-15 | -7.929263016 | 0.558985898 |
| CNAG_04580 | - | RNA helicase, variant | 1914.275893 | 1738.10183 | 2028.495857 | 2862.982392 | 2926.112215 | 2855.124057 | 1.55E-14 | 0.589826454 | 2.28E-15 | 7.925212251 | 1.505065688 |
| CNAG_00569 | - | GTP cyclohydrolase II | 3126.687171 | 2590.076979 | 2846.228863 | 1847.967649 | 1904.272223 | 1824.23876 | 1.61E-14 | -0.63514157 | 2.36E-15 | -7.920959366 | 0.643877629 |
| CNAG_04045 | - | hypothetical protein | 1827.230066 | 1783.034149 | 2020.721569 | 1148.989604 | 1175.353669 | 1286.858467 | 1.66E-14 | -0.65692953 | 2.45E-15 | -7.916372175 | 0.634226678 |
| CNAG_07381 | - | ATP-dependent RNA helicase prh1 | 1881.004852 | 1942.467893 | 2058.083153 | 1289.947597 | 1379.020038 | 1241.416464 | 1.67E-14 | -0.604612585 | 2.45E-15 | -7.915969654 | 0.657647958 |
| CNAG_00326 | - | hypothetical protein | 4751.284016 | 4504.655993 | 4689.087547 | 3348.610985 | 3437.966277 | 3358.41939 | 1.69E-14 | -0.474706253 | 2.48E-15 | -7.9144958 | 0.719613295 |
| CNAG_03335 | - | Ribose-phosphate pyrophosphokinase | 3413.055055 | 3454.105587 | 3394.412796 | 2242.536022 | 2560.774905 | 2361.518164 | 1.76E-14 | -0.533870254 | 2.59E-15 | -7.909115505 | 0.690699338 |
| CNAG_01809 | - | U4/U6 small nuclear ribonucleoprotein Prp3 | 130.2669988 | 137.1174148 | 154.8428275 | 339.4242457 | 377.7021164 | 302.81285 | 1.79E-14 | 1.258356133 | 2.64E-15 | 7.90664167 | 2.392230049 |
| CNAG_06411 | - | mutanase | 111.4314539 | 62.86128354 | 69.62337174 | 265.7818516 | 243.7845709 | 270.2767335 | 1.79E-14 | 1.667924108 | 2.64E-15 | 7.906727064 | 3.177570441 |
| CNAG_03481 | - | ribonuclease P protein component | 785.4709806 | 766.9587822 | 849.7225183 | 446.9280332 | 398.0491872 | 488.6987766 | 1.81E-14 | -0.864642456 | 2.67E-15 | -7.905305448 | 0.549182494 |
| CNAG_05307 | - | beta-catenin-like protein 1 | 2282.663788 | 2270.020692 | 2344.993068 | 1476.68246 | 1642.968707 | 1591.056331 | 1.89E-14 | -0.565721842 | 2.79E-15 | -7.900148229 | 0.675617292 |
| CNAG_06598 | - | hypothetical protein, variant | 25.29755514 | 22.99298117 | 42.90329035 | 150.8609334 | 123.8347095 | 153.5797867 | 1.89E-14 | 2.243358394 | 2.79E-15 | 7.899801494 | 4.734980202 |
| CNAG_07844 | - | hypothetical protein | 26.43650809 | 31.57121252 | 32.27061642 | 144.685834 | 116.933714 | 137.0299686 | 1.95E-14 | 2.142640565 | 2.88E-15 | 7.895971259 | 4.415695119 |
| CNAG_07512 | - | hypothetical protein | 323.9737585 | 368.6559099 | 292.4434167 | 685.036666 | 617.9636889 | 610.310676 | 1.96E-14 | 0.945274925 | 2.90E-15 | 7.894928813 | 1.925555792 |
| CNAG_02888 | - | gamma-glutamyltransferase | 283.1475039 | 298.682051 | 327.5657608 | 549.9671846 | 628.5018428 | 584.7221387 | 1.99E-14 | 0.940320455 | 2.95E-15 | 7.892831244 | 1.918954435 |
| CNAG_00140 | - | U1 small nuclear ribonucleoprotein 70kDa | 2247.525823 | 2166.569486 | 2073.09457 | 1437.048929 | 1420.345767 | 1529.975214 | 2.06E-14 | -0.57966489 | 3.06E-15 | -7.88854016 | 0.669119183 |
| CNAG_05169 | - | L-lactate dehydrogenase (cytochrome) | 735.1509018 | 774.3701193 | 640.8430349 | 413.5143416 | 390.3594391 | 377.7798851 | 2.15E-14 | -0.879724212 | 3.19E-15 | -7.883397567 | 0.543471312 |

| | | | | | | | | | | | | | |
|---|---|---|---|---|---|---|---|---|---|---|---|---|---|
| CNAG_07834 | - | hypothetical protein | 2.971373502 | 5.69694388 | 4.530975853 | 74.62631065 | 63.57021683 | 80.20825111 | 2.15E-14 | 4.288062197 | 3.20E-15 | 7.882973576 | 19.53598642 |
| CNAG_05759 | - | acetyl-CoA carboxylase/biotin carboxylase | 42781.29801 | 38009.56048 | 36806.44366 | 26661.43935 | 28384.08296 | 28198.26707 | 2.20E-14 | -0.514199959 | 3.26E-15 | -7.880343291 | 0.700181108 |
| CNAG_00135 | - | cyclophilin | 980.5545269 | 1053.490499 | 988.4538905 | 663.4997613 | 574.0943927 | 560.9436834 | 2.28E-14 | -0.764316608 | 3.39E-15 | -7.875740018 | 0.58873218 |
| CNAG_03396 | NPY1 | NAD diphosphatase | 1384.688038 | 1433.906543 | 1497.129658 | 970.1255081 | 938.183625 | 900.5980166 | 2.29E-14 | -0.634689645 | 3.41E-15 | -7.874884552 | 0.644079355 |
| CNAG_05430 | - | hypothetical protein | 427.8175212 | 421.4773709 | 408.6367415 | 755.5852651 | 703.2585621 | 777.8598995 | 2.33E-14 | 0.816336463 | 3.47E-15 | 7.872815113 | 1.76092866 |
| CNAG_00022 | - | cytochrome c heme-lyase | 1264.410622 | 1304.618705 | 1390.469788 | 833.8293346 | 856.2399371 | 858.3792556 | 2.49E-14 | -0.651191406 | 3.72E-15 | -7.86402847 | 0.636754252 |
| CNAG_05619 | - | TPR repeat-containing protein | 1853.903209 | 1595.597351 | 1836.248301 | 1078.175905 | 1077.86988 | 1170.31161 | 2.53E-14 | -0.684149936 | 3.77E-15 | -7.862233214 | 0.622372432 |
| CNAG_07908 | - | aconitate hydratase, mitochondrial | 1505.864153 | 1364.959749 | 1339.354563 | 818.9429152 | 903.6840197 | 911.6594422 | 2.54E-14 | -0.692592816 | 3.79E-15 | -7.861709319 | 0.618740847 |
| CNAG_02118 | - | hypothetical protein | 1.7826407 | 4.622700048 | 1.313255334 | 79.36601517 | 79.44193601 | 75.92784967 | 2.54E-14 | 5.264995746 | 3.80E-15 | 7.861363637 | 38.45224022 |
| CNAG_06830 | - | Histidinol dehydrogenase | 1474.35067 | 1418.778671 | 1286.052571 | 819.7427338 | 940.8298257 | 806.2736412 | 2.89E-14 | -0.719362508 | 4.32E-15 | -7.845249778 | 0.607365763 |
| CNAG_07530 | - | hypothetical protein | 206.0380436 | 209.2530844 | 164.5030736 | 485.2984438 | 404.0534026 | 411.5438032 | 2.99E-14 | 1.152013476 | 4.49E-15 | 7.840508705 | 2.222238215 |
| CNAG_03852 | - | hypothetical protein | 70.70557012 | 74.62528166 | 54.68905996 | 188.2980103 | 232.0469199 | 203.7915365 | 3.05E-14 | 1.638681365 | 4.58E-15 | 7.838041943 | 3.113810969 |
| CNAG_01623 | - | tRNA pseudouridine(55) synthase | 1063.577239 | 1134.361919 | 1229.398951 | 661.7313465 | 728.58362 | 709.637507 | 3.27E-14 | -0.722692834 | 4.91E-15 | -7.829286493 | 0.605965333 |
| CNAG_07868 | - | hypothetical protein | 343.9718957 | 444.1020532 | 466.1910614 | 849.1948059 | 755.0511915 | 810.066455 | 3.35E-14 | 0.931672277 | 5.04E-15 | 7.826022601 | 1.907485748 |
| CNAG_02677 | - | cytoplasmic protein | 5351.516901 | 5100.475021 | 5414.25878 | 3604.586382 | 3822.649868 | 3951.210739 | 3.44E-14 | -0.495224218 | 5.17E-15 | -7.822701159 | 0.709451409 |
| CNAG_07476 | - | hypothetical protein | 49.77026738 | 65.99709146 | 46.15639659 | 213.2913723 | 174.7936445 | 168.3579303 | 3.48E-14 | 1.780261232 | 5.23E-15 | 7.821194378 | 3.434883652 |
| CNAG_04190 | TOP1 | DNA topoisomerase I | 8372.521265 | 8321.972859 | 7993.000525 | 6059.408621 | 6215.420045 | 6292.537788 | 3.54E-14 | -0.426248567 | 5.34E-15 | -7.818725578 | 0.744194397 |
| CNAG_01491 | - | hypothetical protein | 1423.082175 | 1304.665587 | 1450.19841 | 913.2778349 | 858.7828237 | 893.9121931 | 3.59E-14 | -0.664487727 | 5.41E-15 | -7.816910023 | 0.63091269 |
| CNAG_02449 | - | long-chain fatty acid CoA ligase | 2024.535945 | 1944.644075 | 1972.822916 | 1350.68987 | 1391.079829 | 1343.508632 | 3.61E-14 | -0.555938438 | 5.45E-15 | -7.816122052 | 0.680214451 |
| CNAG_00820 | - | Prephenate dehydratase | 687.4972819 | 744.2845941 | 772.9542288 | 410.3399386 | 394.651631 | 445.41293 | 3.73E-14 | -0.833574606 | 5.63E-15 | -7.81196035 | 0.561137173 |
| CNAG_05229 | - | Stomatin family protein | 1008.029351 | 737.2358358 | 780.8575939 | 1434.049244 | 1605.706953 | 1579.400114 | 3.75E-14 | 0.85443026 | 5.66E-15 | 7.811236892 | 1.808044593 |
| CNAG_01949 | - | chlorophyll synthesis pathway protein BchC | 236.3942783 | 201.7724668 | 180.509952 | 386.1913446 | 544.1344637 | 587.6648928 | 3.84E-14 | 1.282183442 | 5.80E-15 | 7.808223104 | 2.43206779 |
| CNAG_06055 | - | hypothetical protein | 177.9864805 | 151.1255827 | 152.7445736 | 332.8591655 | 372.2981181 | 401.3273347 | 4.10E-14 | 1.187306111 | 6.20E-15 | 7.799876093 | 2.277271198 |
| CNAG_01180 | - | hypothetical protein | 140.845712 | 207.0653334 | 169.8032557 | 432.2563981 | 406.8110426 | 371.605625 | 4.24E-14 | 1.214432777 | 6.42E-15 | 7.795368206 | 2.320495303 |
| CNAG_05063 | SSK2 | ste/ste11/ssk protein kinase | 3506.494329 | 3550.005668 | 3444.553386 | 2439.519298 | 2514.136073 | 2617.908081 | 4.71E-14 | -0.487148462 | 7.14E-15 | -7.781943061 | 0.713433832 |
| CNAG_00090 | - | hypothetical protein, variant | 1670.814316 | 1763.595314 | 1776.566463 | 1115.566386 | 1199.583177 | 1191.40006 | 4.72E-14 | -0.586684686 | 7.17E-15 | -7.781514931 | 0.665871323 |
| CNAG_01628 | - | small subunit ribosomal protein S20 | 21735.08617 | 21758.61527 | 22765.42835 | 13383.66161 | 16969.03989 | 14457.86622 | 4.88E-14 | -0.579958343 | 7.41E-15 | -7.777357758 | 0.668983094 |
| CNAG_01117 | EF3 | elongation factor 3 | 29563.47454 | 26060.82759 | 24966.68035 | 18959.55334 | 19431.56909 | 17605.18041 | 4.94E-14 | -0.541176733 | 7.51E-15 | -7.775552108 | 0.687210158 |
| CNAG_06741 | - | ATP-dependent RNA helicase DBP10 | 2883.938963 | 2546.978525 | 3066.901895 | 1669.616027 | 1765.343834 | 1952.947272 | 4.96E-14 | -0.673252426 | 7.55E-15 | -7.774961538 | 0.627091372 |
| CNAG_04298 | NOC3 | nucleolar complex protein 3 | 4649.602286 | 4165.278645 | 4609.065765 | 2855.798595 | 3166.238228 | 3180.786398 | 5.10E-14 | -0.560693225 | 7.76E-15 | -7.771428042 | 0.677976313 |
| CNAG_07022 | - | hypothetical protein | 11.21181683 | 11.10268111 | 9.883042343 | 107.1227171 | 68.52195083 | 87.07532756 | 5.14E-14 | 3.064559143 | 7.83E-15 | 7.770247497 | 8.366122619 |
| CNAG_03352 | - | hypothetical protein | 1432.360782 | 1372.474969 | 1330.810298 | 883.6855667 | 919.2495105 | 756.3530395 | 5.22E-14 | -0.708123854 | 7.95E-15 | -7.768389445 | 0.612115645 |
| CNAG_02043 | - | hypothetical protein, variant | 90.56156917 | 107.9870562 | 70.68906257 | 228.6995385 | 261.158428 | 273.5998472 | 5.38E-14 | 1.498619395 | 8.21E-15 | 7.764324098 | 2.825721721 |
| CNAG_04565 | - | hypothetical protein | 746.6667324 | 882.5483649 | 813.8550665 | 1354.044556 | 1276.856614 | 1306.414598 | 6.02E-14 | 0.674720847 | 9.19E-15 | 7.750033236 | 1.596287871 |
| CNAG_01479 | - | hypothetical protein | 1064.264348 | 853.2209349 | 957.4886909 | 547.3570435 | 478.2682203 | 568.6523828 | 6.03E-14 | -0.86727711 | 9.21E-15 | -7.749660501 | 0.548180491 |
| CNAG_07655 | - | hypothetical protein | 5.343279813 | 8.93195591 | 7.736341648 | 82.56430473 | 67.01146605 | 75.9514774 | 6.17E-14 | 3.471067134 | 9.43E-15 | 7.746746285 | 11.08907509 |
| CNAG_02921 | - | hypothetical protein | 782.7195463 | 719.9192271 | 750.9315094 | 1214.66389 | 1196.674843 | 1188.749467 | 6.45E-14 | 0.661080254 | 9.88E-15 | 7.740739776 | 1.581266194 |
| CNAG_02447 | - | hypothetical protein | 1049.407104 | 1034.163983 | 1021.525764 | 679.0258453 | 642.2935148 | 627.5095308 | 6.45E-14 | -0.687416724 | 9.88E-15 | -7.740838962 | 0.620964749 |
| CNAG_04074 | - | coatomer beta' subunit | 7122.977408 | 6581.963687 | 7112.055695 | 5204.90568 | 5000.036663 | 4908.221668 | 6.45E-14 | -0.477602351 | 9.87E-15 | -7.74086951 | 0.718170177 |
| CNAG_06651 | - | amidohydrolase | 12.37194808 | 5.719990179 | 13.07375815 | 94.16910735 | 70.2836359 | 100.1579971 | 6.76E-14 | 3.121251658 | 1.04E-14 | 7.734681078 | 8.701424839 |
| CNAG_03543 | - | hypothetical protein, variant | 2509.155618 | 2426.251327 | 2390.891621 | 1693.152785 | 1663.610271 | 1764.213052 | 6.80E-14 | -0.531963588 | 1.04E-14 | -7.733791294 | 0.69161277 |
| CNAG_00827 | - | ribose 5-phosphate isomerase | 19.3306397 | 2.474975599 | 6.69123751 | 100.4165827 | 93.10315096 | 93.71428476 | 6.96E-14 | 3.386268733 | 1.07E-14 | 7.730687152 | 10.45606952 |
| CNAG_05300 | - | mitochondrial carrier protein | 60.23042057 | 57.42674967 | 68.49480203 | 205.9504342 | 186.89151 | 182.7508911 | 7.06E-14 | 1.615552535 | 1.09E-14 | 7.728804016 | 3.064289355 |
| CNAG_01834 | - | hypothetical protein, variant | 2529.066017 | 2486.598211 | 2702.223751 | 1837.993425 | 1599.709153 | 1748.708773 | 7.18E-14 | -0.589236008 | 1.10E-14 | -7.726584586 | 0.664694809 |
| CNAG_05458 | - | endo-1,3(4)-beta-glucanase | 162.7903143 | 127.4403502 | 134.6195358 | 361.4472318 | 315.2443426 | 331.5361181 | 7.31E-14 | 1.235483827 | 1.13E-14 | 7.724148403 | 2.35460299 |
| CNAG_03560 | - | hypothetical protein | 95.24784381 | 113.3829965 | 78.15142231 | 222.9997528 | 296.6562083 | 300.1960797 | 7.56E-14 | 1.501644471 | 1.16E-14 | 7.719853209 | 2.831652976 |
| CNAG_07810 | - | mitochondrial protein | 1264.214655 | 1239.931813 | 1155.928391 | 758.3323944 | 819.1987678 | 695.2699511 | 7.62E-14 | -0.702536783 | 1.18E-14 | -7.718677392 | 0.614490758 |
| CNAG_04833 | TEP1 | hypothetical protein | 622.7759802 | 612.1587591 | 494.0016366 | 981.1209372 | 1116.692904 | 991.1239756 | 7.76E-14 | 0.822303579 | 1.20E-14 | 7.716211791 | 1.768227102 |
| CNAG_06658 | - | rhomboid family membrane protein | 809.6171986 | 758.714031 | 844.7556695 | 1255.039544 | 1282.072677 | 1307.495923 | 8.19E-14 | 0.657433057 | 1.26E-14 | 7.70935442 | 1.577273731 |
| CNAG_04676 | - | 6-phosphofructokinase | 4066.400093 | 4055.58974 | 4066.50043 | 6103.117621 | 5642.212991 | 5443.004596 | 8.22E-14 | 0.480784396 | 1.27E-14 | 7.708658683 | 1.395502197 |
| CNAG_05657 | - | cytoplasmic protein | 635.742922 | 730.65402 | 717.8642556 | 1115.750842 | 1109.529454 | 1200.858899 | 8.47E-14 | 0.703041483 | 1.31E-14 | 7.704818287 | 1.627933179 |
| CNAG_01551 | GAT201 | GATA family transcription factor | 131.3727946 | 131.7163026 | 137.7905333 | 331.03107 | 291.9430297 | 310.4369044 | 8.78E-14 | 1.208796302 | 1.36E-14 | 7.700180587 | 2.311447031 |
| CNAG_06846 | - | hypothetical protein | 693.0997028 | 668.8712125 | 634.4010239 | 392.5196384 | 384.3635436 | 350.0321762 | 8.96E-14 | -0.840576275 | 1.39E-14 | -7.697477617 | 0.558420467 |
| CNAG_07443 | HLH4 | hypothetical protein | 7.688993376 | 7.861619987 | 9.868427751 | 70.27142454 | 89.96491848 | 71.64676367 | 8.98E-14 | 3.306260913 | 1.39E-14 | 7.697089175 | 9.891990903 |
| CNAG_06665 | - | hypothetical protein | 716.4768226 | 629.1066101 | 763.3129467 | 398.7063321 | 396.4284256 | 356.7080506 | 9.08E-14 | -0.890258765 | 1.41E-14 | -7.695459095 | 0.539517341 |
| CNAG_04776 | - | Homoserine dehydrogenase | 683.4405687 | 587.4192208 | 539.8441799 | 1162.067902 | 1023.289843 | 1035.529544 | 9.09E-14 | 0.816243584 | 1.41E-14 | 7.695216427 | 1.760815298 |
| CNAG_05884 | KAP123 | importin beta-4 subunit | 2294.113363 | 2029.765463 | 2051.747074 | 1432.428694 | 1441.958738 | 1264.767701 | 9.30E-14 | -0.639508889 | 1.45E-14 | -7.692261335 | 0.641931433 |
| CNAG_00486 | - | succinate dehydrogenase assembly factor 2, mitochondrial | 1721.940527 | 1499.692719 | 1645.382206 | 970.7321819 | 1085.714542 | 1050.438831 | 9.92E-14 | -0.663795686 | 1.54E-14 | -7.683840605 | 0.631215402 |
| CNAG_00696 | - | alpha-mannosidase | 1126.045935 | 1146.531594 | 1113.514544 | 1662.811079 | 1744.418745 | 1665.04963 | 9.92E-14 | 0.567896101 | 1.54E-14 | 7.683840827 | 1.48236025 |
| CNAG_01163 | RAD54 | DNA repair and recombination protein RAD54-like protein | 858.7584929 | 961.2246909 | 981.260144 | 1425.435135 | 1565.954831 | 1448.546977 | 1.01E-13 | 0.649462855 | 1.57E-14 | 7.681809422 | 1.568584071 |
| CNAG_00138 | - | ubiquinone biosynthesis methyltransferase | 3452.634396 | 3031.828251 | 3447.683062 | 2114.878654 | 2345.951828 | 2236.076211 | 1.02E-13 | -0.584725012 | 1.58E-14 | -7.680556603 | 0.666776419 |
| CNAG_07314 | - | hypothetical protein | 55.52916445 | 50.95653875 | 50.39755079 | 162.7480667 | 162.6946317 | 193.5663882 | 1.03E-13 | 1.72266121 | 1.61E-14 | 7.678440045 | 3.300446496 |
| CNAG_04689 | - | hypothetical protein | 157.0668545 | 174.7766393 | 170.8398613 | 363.4743722 | 370.5660247 | 349.3691579 | 1.12E-13 | 1.095820985 | 1.75E-14 | 7.667871738 | 2.137346756 |
| CNAG_02966 | - | carboxypeptidase | 61.41032222 | 69.24587493 | 69.56759833 | 189.9977405 | 208.605668 | 192.7169063 | 1.18E-13 | 1.55044933 | 1.84E-14 | 7.661001618 | 2.929083517 |
| CNAG_04653 | - | hypothetical protein | 212.1528715 | 366.5041571 | 321.2064886 | 667.0478158 | 662.9495716 | 669.0106323 | 1.19E-13 | 1.138857552 | 1.86E-14 | 7.660029497 | 2.202065759 |
| CNAG_00875 | - | rab family protein | 924.324978 | 826.2564027 | 876.4602936 | 524.0379341 | 499.052471 | 543.1050582 | 1.24E-13 | -0.761381294 | 1.94E-14 | -7.654858256 | 0.589931236 |
| CNAG_01913 | - | hypothetical protein | 144.1680972 | 118.8264259 | 135.6701981 | 305.0093446 | 296.2377016 | 369.0236771 | 1.28E-13 | 1.273225615 | 2.01E-14 | 7.650160254 | 2.41701364 |
| CNAG_03300 | - | ribosomal RNA-processing protein 1 | 2823.167124 | 2414.463239 | 2826.98536 | 1679.804949 | 1879.308463 | 1675.494919 | 1.32E-13 | -0.639914486 | 2.07E-14 | -7.646506365 | 0.641750987 |
| CNAG_07597 | - | hypothetical protein, variant | 50.95073302 | 60.64421852 | 68.49145898 | 194.9055699 | 209.5846638 | 167.317832 | 1.33E-13 | 1.663892465 | 2.08E-14 | 7.645567682 | 3.168703047 |
| CNAG_04718 | PCD101 | hypothetical protein | 145.400486 | 151.1046761 | 167.6263458 | 332.2433964 | 362.875489 | 337.1256792 | 1.33E-13 | 1.139639217 | 2.09E-14 | 7.645085911 | 2.203259181 |
| CNAG_07391 | - | hypothetical protein | 31.10958787 | 35.8839785 | 41.85117427 | 171.4313735 | 126.3850031 | 136.0859707 | 1.33E-13 | 2.001868026 | 2.09E-14 | 7.644802534 | 4.005182621 |
| CNAG_04671 | - | hypothetical protein | 579.7225234 | 699.4002203 | 660.2870713 | 1078.460973 | 1035.330141 | 1145.334636 | 1.43E-13 | 0.734997652 | 2.25E-14 | 7.635227461 | 1.664394761 |
| CNAG_00444 | - | hypothetical protein | 551.8487252 | 597.6573208 | 576.7131638 | 241.990283 | 343.9166708 | 221.3747661 | 1.46E-13 | -1.114225505 | 2.31E-14 | -7.632309501 | 0.461939076 |
| CNAG_02339 | - | hypothetical protein | 11.23795633 | 14.33731119 | 19.45942145 | 93.0587424 | 97.45334163 | 90.48207018 | 1.49E-13 | 2.65471133 | 2.34E-14 | 7.63021046 | 6.297203672 |
| CNAG_05609 | - | hypothetical protein | 1879.724534 | 1756.114424 | 1888.543705 | 1205.094473 | 1274.625641 | 1260.234044 | 1.49E-13 | -0.578409201 | 2.35E-14 | -7.629912253 | 0.669701822 |
| CNAG_01928 | - | Xaa-Pro aminopeptidase | 2185.856686 | 2327.115723 | 2476.133153 | 1555.346002 | 1623.963459 | 1621.028065 | 1.53E-13 | -0.557261181 | 2.41E-14 | -7.626369553 | 0.679591079 |
| CNAG_04709 | - | hypothetical protein | 830.6123378 | 827.6273348 | 764.8187526 | 1367.349195 | 1237.150172 | 1259.817956 | 1.56E-13 | 0.659015447 | 2.47E-14 | 7.623393071 | 1.579004679 |
| CNAG_03064 | - | nucleolar protein 16 | 1649.806986 | 1672.058129 | 2025.988518 | 1161.558942 | 1053.692834 | 1098.205175 | 1.63E-13 | -0.706122022 | 2.58E-14 | -7.618037096 | 0.612965584 |
| CNAG_13046 | - | unspecified product | 2.970506185 | 4.617694662 | 2.387789888 | 102.3880958 | 62.69111922 | 54.11404969 | 1.70E-13 | 4.753888316 | 2.69E-14 | 7.612550554 | 26.98130673 |
| CNAG_04926 | - | hypothetical protein | 74.12982752 | 53.13239283 | 29.13676896 | 214.6499643 | 206.9344338 | 186.0591297 | 1.72E-13 | 1.941764139 | 2.72E-14 | 7.610988348 | 3.84175133 |
| CNAG_03911 | - | hydrolase | 97.53592621 | 88.65904302 | 108.9909753 | 241.2541156 | 255.9033629 | 260.3934599 | 1.73E-13 | 1.35186006 | 2.74E-14 | 7.610214822 | 2.552409945 |
| CNAG_03789 | - | mRNA turnover protein 4 | 1657.823511 | 1504.009631 | 1778.628637 | 1111.015363 | 989.8717761 | 1020.494869 | 1.74E-13 | -0.678399733 | 2.75E-14 | -7.609363877 | 0.624857996 |
| CNAG_07707 | - | glycoside hydrolase family 3 domain-containing protein | 12.43447575 | 18.6504655 | 24.78966841 | 87.05749165 | 114.7947118 | 116.9034809 | 1.77E-13 | 2.538610108 | 2.81E-14 | 7.606816377 | 5.810289742 |
| CNAG_04146 | - | hypothetical protein | 3201.720816 | 3189.085628 | 3245.105507 | 2208.345517 | 2380.47347 | 2355.927112 | 1.88E-13 | -0.488050585 | 2.99E-14 | -7.598647502 | 0.712987859 |
| CNAG_06224 | MLN2 | nuclear movement protein nudC | 1189.436524 | 1159.112086 | 1038.639991 | 724.1400459 | 718.2277011 | 648.6358473 | 1.89E-13 | -0.711397367 | 3.01E-14 | -7.598042452 | 0.610728313 |
| CNAG_01946 | - | allantoate permease | 1046.975962 | 885.5792143 | 1149.349385 | 551.1526769 | 585.322551 | 611.9523621 | 1.90E-13 | -0.834173116 | 3.03E-14 | -7.597147545 | 0.560904431 |
| CNAG_06288 | - | RNA binding protein | 2787.119962 | 2628.848469 | 2847.292076 | 1978.839056 | 1948.290032 | 1899.695002 | 1.99E-13 | -0.519821233 | 3.16E-14 | -7.591492654 | 0.697458251 |

| | | | | | | | | | | | | | |
|---|---|---|---|---|---|---|---|---|---|---|---|---|---|
| CNAG_00019 | - | mitochondrial import inner membrane translocase subunit tim9 | 2638.843022 | 2600.807576 | 3029.562323 | 1819.217643 | 1891.373441 | 1540.124537 | 2.00E-13 | -0.670825303 | 3.19E-14 | -7.59042353 | 0.628147249 |
| CNAG_07572 | ELP3 | elongator complex protein 3 | 1598.246375 | 1480.278471 | 1534.499847 | 1012.097152 | 1053.805966 | 1014.912521 | 2.02E-13 | -0.598548316 | 3.21E-14 | -7.589395825 | 0.660418154 |
| CNAG_02805 | CAN1 | Putative beta-type carbonic anhydrase | 31.07667535 | 40.15430672 | 20.57900293 | 130.2479068 | 153.681319 | 123.8983868 | 2.06E-13 | 2.143172634 | 3.29E-14 | 7.586558034 | 4.417323937 |
| CNAG_00555 | - | hypothetical protein, variant | 604.3565294 | 825.4632117 | 782.9299644 | 1226.998728 | 1437.580348 | 1289.732235 | 2.09E-13 | 0.823577722 | 3.34E-14 | 7.584582834 | 1.769789434 |
| CNAG_04111 | - | lectin, mannose-binding 2 | 1121.731937 | 1184.895427 | 973.5991425 | 696.1354226 | 593.8953277 | 651.9293619 | 2.11E-13 | -0.771397933 | 3.38E-14 | -7.582966192 | 0.585849527 |
| CNAG_03310 | - | nicotinamide N-methyltransferase | 995.6182197 | 898.460111 | 977.7657516 | 590.997588 | 580.1751244 | 596.3991087 | 2.14E-13 | -0.716225008 | 3.42E-14 | -7.581386637 | 0.608688069 |
| CNAG_07608 | - | hypothetical protein | 1984.767553 | 1870.291084 | 1827.806457 | 1312.589122 | 1317.757825 | 1225.863963 | 2.15E-13 | -0.575310655 | 3.44E-14 | -7.580414045 | 0.671141719 |
| CNAG_05932 | - | peptidyl-prolyl cis-trans isomerase D | 5634.141797 | 5819.060523 | 5270.359964 | 4123.043207 | 4116.792357 | 4128.855825 | 2.27E-13 | -0.450325692 | 3.64E-14 | -7.573191307 | 0.731877606 |
| CNAG_06502 | - | solute carrier family 25 (mitochondrial carnitine/acylcarnitine transporter), member 20/29 | 1021.503266 | 1097.704121 | 1076.963205 | 640.7540903 | 668.1775537 | 710.7009118 | 2.38E-13 | -0.677354677 | 3.81E-14 | -7.567377384 | 0.625310793 |
| CNAG_01444 | RPB4 | DNA-directed RNA polymerase II subunit RPB4 | 248.4625996 | 313.1729972 | 349.369835 | 138.6528631 | 102.0651749 | 112.5512165 | 2.44E-13 | -1.384732943 | 3.92E-14 | -7.563634634 | 0.382960382 |
| CNAG_00979 | CTR4 | Copper uptake transporter | 3918.433311 | 3139.995667 | 3496.114862 | 7441.522555 | 10780.37069 | 10494.75952 | 2.53E-13 | 1.427630138 | 4.06E-14 | 7.559057065 | 2.690044684 |
| CNAG_01223 | - | hypothetical protein | 232.9635777 | 268.4919448 | 276.3814175 | 492.4935609 | 531.8643468 | 489.2865579 | 2.66E-13 | 0.947355399 | 4.28E-14 | 7.552281084 | 1.928334591 |
| CNAG_01962 | - | ubiquinone biosynthesis protein COQ4, mitochondrial | 540.185786 | 526.6668611 | 542.62901 | 282.6470437 | 302.4101857 | 301.1351486 | 3.17E-13 | -0.876901726 | 5.09E-14 | -7.529655009 | 0.544535599 |
| CNAG_00608 | - | hypothetical protein | 967.2934626 | 776.7047582 | 851.9123101 | 531.9212752 | 404.9164147 | 452.1620827 | 3.21E-13 | -0.9191785 | 5.16E-14 | -7.527762243 | 0.52881005 |
| CNAG_03410 | - | cytoplasmic protein | 2360.928256 | 2350.838439 | 2434.582285 | 1621.518256 | 1632.563018 | 1755.315277 | 3.25E-13 | -0.527719339 | 5.24E-14 | -7.525828167 | 0.693650414 |
| CNAG_04124 | - | hypothetical protein | 4065.767347 | 3959.430212 | 3752.776736 | 2959.563047 | 2677.111915 | 2748.959593 | 3.32E-13 | -0.505354019 | 5.35E-14 | -7.523131373 | 0.704487484 |
| CNAG_04845 | - | mannose-P-dolichol utilization defect 1 | 335.6828772 | 404.1998797 | 347.8645195 | 732.5068558 | 667.143821 | 619.2669071 | 3.37E-13 | 0.879382021 | 5.44E-14 | 7.520956259 | 1.839587145 |
| CNAG_12183 | - | unspecified product | 18.22468614 | 17.57164512 | 19.4748256 | 108.0470356 | 114.8719385 | 81.86002204 | 3.54E-13 | 2.472513467 | 5.71E-14 | 7.514571235 | 5.550098851 |
| CNAG_05439 | CMK1 | CAMK/CAMK1/CAMK1-CMK protein kinase | 342.7186295 | 451.5758951 | 348.9491849 | 719.8281745 | 718.9371807 | 696.8882675 | 3.62E-13 | 0.888519176 | 5.84E-14 | 7.511554642 | 1.851274945 |
| CNAG_07416 | - | exosome complex component RRP41 | 1163.956018 | 1183.94676 | 1230.524503 | 802.7912279 | 747.5209898 | 778.4479387 | 4.01E-13 | -0.635037637 | 6.48E-14 | -7.498057488 | 0.643924016 |
| CNAG_06922 | - | anon-23da protein | 1260.786844 | 1326.089263 | 1150.624232 | 834.7059048 | 780.3045619 | 770.7115795 | 4.04E-13 | -0.662983663 | 6.53E-14 | -7.49702345 | 0.631570783 |
| CNAG_00706 | - | RNA-binding protein rnp24 | 1976.774138 | 2027.59799 | 2115.696742 | 1358.429157 | 1423.847031 | 1460.033735 | 4.04E-13 | -0.543848506 | 6.54E-14 | -7.496703361 | 0.685938669 |
| CNAG_05457 | - | Pentatricopeptide repeat protein | 2441.371535 | 2265.728572 | 2385.518207 | 1436.145782 | 1560.954735 | 1718.672802 | 4.74E-13 | -0.604546177 | 7.68E-14 | -7.47564593 | 0.657678231 |
| CNAG_01883 | GAT8 | hypothetical protein | 10.13451751 | 36.90703579 | 18.43771754 | 137.0440622 | 106.5860589 | 132.4879644 | 4.92E-13 | 2.539341149 | 7.98E-14 | 7.470670433 | 5.813234671 |
| CNAG_01991 | - | cytochrome c oxidase subunit 4 | 1549.207318 | 1516.861237 | 1424.692015 | 989.535947 | 1045.20143 | 953.8793607 | 5.03E-13 | -0.603069214 | 8.16E-14 | -7.467673193 | 0.658351876 |
| CNAG_04132 | - | pre-mRNA-splicing factor CWC2, variant | 3013.634275 | 2907.86995 | 2865.468724 | 1993.531835 | 2203.672794 | 2038.459997 | 5.10E-13 | -0.510385783 | 8.27E-14 | -7.465927035 | 0.702034686 |
| CNAG_01833 | - | translation initiation factor 4E | 1246.799779 | 1227.045025 | 1262.527648 | 861.9986013 | 797.5754895 | 733.0020682 | 5.20E-13 | -0.658886201 | 8.44E-14 | -7.463293727 | 0.633367085 |
| CNAG_06953 | - | hypothetical protein | 104.5827965 | 124.1528806 | 119.6668123 | 296.004237 | 254.8098755 | 291.488379 | 5.20E-13 | 1.262649966 | 8.46E-14 | 7.463002877 | 2.399360549 |
| CNAG_01482 | - | solute carrier family 25 (mitochondrial phosphate transporter), member 23/24/25/41 | 1379.957915 | 1331.589919 | 1438.481471 | 917.1634565 | 947.7145738 | 882.8260668 | 5.33E-13 | -0.610632386 | 8.67E-14 | -7.459692761 | 0.654909568 |
| CNAG_03232 | - | lactamase | 1028.38239 | 1042.752399 | 968.2283465 | 657.9982626 | 658.7280261 | 584.2466009 | 5.44E-13 | -0.693114072 | 8.86E-14 | -7.456879028 | 0.618517332 |
| CNAG_12325 | - | unspecified product | 2.963281974 | 3.532913452 | 8.788116574 | 78.02253068 | 72.51231715 | 55.16336584 | 5.48E-13 | 3.963426464 | 8.92E-14 | 7.455990736 | 15.59948466 |
| CNAG_02864 | - | hypothetical protein | 802.655663 | 897.6294007 | 827.7166857 | 1349.330015 | 1316.567105 | 1262.071444 | 5.79E-13 | 0.621759027 | 9.44E-14 | 7.448515532 | 1.538750181 |
| CNAG_05344 | - | Hemolysin | 1591.232714 | 1558.87441 | 1455.618445 | 957.5443341 | 991.6488253 | 1080.354269 | 5.91E-13 | -0.619677577 | 9.65E-14 | -7.445641856 | 0.65081636 |
| CNAG_07889 | - | hypothetical protein | 1487.479852 | 1569.659107 | 1644.29369 | 1007.404846 | 956.2341452 | 1104.773637 | 6.08E-13 | -0.630760065 | 9.92E-14 | -7.441916048 | 0.645836075 |
| CNAG_03359 | - | cytochrome c oxidase subunit 7 | 1591.301003 | 1526.601827 | 1608.066043 | 1099.361586 | 989.028864 | 1067.069413 | 6.30E-13 | -0.598010737 | 1.03E-13 | -7.43708244 | 0.660664286 |
| CNAG_07439 | - | U3 small nucleolar RNA-associated protein 21 | 3506.467582 | 3286.09325 | 3648.188029 | 2360.101325 | 2533.997611 | 2502.479495 | 6.48E-13 | -0.513170964 | 1.06E-13 | -7.433233902 | 0.700680687 |
| CNAG_02961 | - | F-type H -transporting ATPase subunit J | 2198.556561 | 2070.732668 | 2493.127822 | 1571.041324 | 1397.891061 | 1292.53919 | 6.55E-13 | -0.682142579 | 1.07E-13 | -7.431672571 | 0.623239001 |
| CNAG_07808 | - | alanine-glyoxylate transaminase/serine- glyoxylate transaminase/serine-pyruvate transaminase | 2154.050087 | 2023.271873 | 1957.909462 | 1262.618304 | 1492.061457 | 1273.61981 | 6.59E-13 | -0.622835801 | 1.08E-13 | -7.430775061 | 0.649393207 |
| CNAG_03965 | - | ribosome recycling factor | 2288.530129 | 2186.029916 | 2577.410368 | 1551.449557 | 1593.753805 | 1599.94636 | 6.69E-13 | -0.587659158 | 1.10E-13 | -7.42879576 | 0.66542171 |
| CNAG_06794 | - | hypothetical protein | 1146.221489 | 1027.743353 | 1079.101513 | 667.2625337 | 619.8007309 | 719.5789215 | 6.82E-13 | -0.713234933 | 1.12E-13 | -7.426087892 | 0.609950921 |
| CNAG_01092 | - | hypothetical protein | 1159.161482 | 1110.682685 | 1092.998934 | 738.1593809 | 732.0347354 | 715.1853429 | 6.90E-13 | -0.636654117 | 1.13E-13 | -7.424414343 | 0.64320293 |
| CNAG_01523 | HOG1 | putative mitogen-activated protein kinase HOG1 | 8664.36547 | 8263.788118 | 7747.731559 | 5850.006687 | 6285.312234 | 5985.058082 | 7.19E-13 | -0.460870953 | 1.18E-13 | -7.418852223 | 0.72654751 |
| CNAG_05273 | - | 1,2-dihydroxy-3-keto-5-methylthiopentene dioxygenase | 2093.406063 | 1899.431921 | 2094.366006 | 1336.630724 | 1420.406961 | 1393.453141 | 7.59E-13 | -0.568497887 | 1.25E-13 | -7.411624062 | 0.674318514 |
| CNAG_01688 | - | ATP-dependent metalloprotease | 8068.832593 | 7356.639356 | 7814.864017 | 5434.379523 | 5675.436293 | 5891.763787 | 7.92E-13 | -0.466622782 | 1.30E-13 | -7.405924375 | 0.723656631 |
| CNAG_01417 | - | hypothetical protein | 59.07445531 | 70.3091664 | 57.86545433 | 199.59823 | 220.0087922 | 158.5296393 | 9.05E-13 | 1.618267926 | 1.49E-13 | 7.3880291 | 3.070062287 |
| CNAG_02467 | - | DNA repair/transcription protein MET18/MMS19 | 2043.223982 | 2004.961951 | 1919.530769 | 1346.775755 | 1429.049347 | 1395.66413 | 9.45E-13 | -0.532030664 | 1.56E-13 | -7.382174084 | 0.691580615 |
| CNAG_04638 | - | hypothetical protein | 39.21845444 | 43.39500949 | 35.47866848 | 122.8148999 | 150.8561348 | 158.060592 | 1.03E-12 | 1.862999429 | 1.70E-13 | 7.370277639 | 3.637631566 |
| CNAG_00457 | GLN1 | glutamine synthetase, variant | 10968.30149 | 10064.19686 | 9670.544213 | 7557.64112 | 7677.465339 | 7281.744824 | 1.04E-12 | -0.463086405 | 1.72E-13 | -7.368673305 | 0.725432656 |
| CNAG_05771 | TEL1 | serine/threonine-protein kinase TEL1 | 1295.318452 | 1256.433771 | 1167.926914 | 1836.372843 | 1985.993408 | 1790.531601 | 1.07E-12 | 0.578305782 | 1.77E-13 | 7.365165759 | 1.493094814 |
| CNAG_06406 | - | hypothetical protein, variant | 782.0289018 | 771.2863826 | 830.5699845 | 509.3466708 | 468.0243056 | 465.4383399 | 1.11E-12 | -0.740060082 | 1.83E-13 | -7.360367512 | 0.598714418 |
| CNAG_04640 | ACL1 | ATP-citrate synthase subunit 1 | 11840.03676 | 10073.10173 | 9231.416788 | 17876.539 | 16154.32955 | 14863.82783 | 1.14E-12 | 0.634759659 | 1.88E-13 | 7.35707675 | 1.552679065 |
| CNAG_07660 | - | pyruvate dehydrogenase (acetyl-transferring) E1 component, alpha subunit | 4263.05492 | 4057.470962 | 3765.585627 | 2943.175006 | 2900.579244 | 2612.464675 | 1.14E-12 | -0.530822368 | 1.89E-13 | -7.35638204 | 0.692160075 |
| CNAG_06242 | CFT1 | Cryptococcus ferric transporter 1 | 2758.667032 | 2140.782215 | 2258.641457 | 1425.262823 | 1455.671937 | 1573.303246 | 1.15E-12 | -0.70062526 | 1.91E-13 | -7.355264146 | 0.615305477 |
| CNAG_03084 | - | endoribonuclease L-PSP | 296.8020145 | 270.0983653 | 234.4567706 | 118.3964814 | 79.60000186 | 109.1717159 | 1.17E-12 | -1.401511581 | 1.93E-13 | -7.353436243 | 0.378532327 |
| CNAG_02522 | MRS3/4 | solute carrier family 25 (mitochondrial iron transporter), member 28/37 | 916.3944287 | 986.7210343 | 951.1304133 | 567.5867171 | 635.4635908 | 594.1835248 | 1.21E-12 | -0.682744044 | 2.01E-13 | -7.34804068 | 0.622979224 |
| CNAG_00652 | - | hypothetical protein, variant 2 | 421.9624217 | 420.3870747 | 415.0174766 | 711.9100365 | 693.7962148 | 732.3911669 | 1.23E-12 | 0.75092264 | 2.04E-13 | 7.346028227 | 1.682868723 |
| CNAG_03387 | - | hypothetical protein | 701.5700646 | 760.4752864 | 815.6088702 | 464.9244731 | 446.4648788 | 444.3410609 | 1.27E-12 | -0.764318735 | 2.12E-13 | -7.341166057 | 0.588731312 |
| CNAG_02984 | - | sarcosine oxidase | 259.6519482 | 250.7309825 | 231.2262859 | 103.5299946 | 103.0340694 | 103.618843 | 1.30E-12 | -1.27519342 | 2.15E-13 | -7.338923736 | 0.413169762 |
| CNAG_07771 | - | peptidase | 1229.262813 | 1193.648219 | 1209.221307 | 833.9646763 | 795.0079975 | 723.0035305 | 1.33E-12 | -0.642540431 | 2.22E-13 | -7.33499471 | 0.640583956 |
| CNAG_06240 | - | protein disulfide-isomerase | 1743.185916 | 1801.312003 | 1732.885227 | 1226.968597 | 1265.164506 | 1194.750981 | 1.34E-12 | -0.53293372 | 2.24E-13 | -7.333765073 | 0.691147855 |
| CNAG_05739 | - | hypothetical protein | 42.72604294 | 42.34021055 | 50.36778616 | 140.9032565 | 164.6983152 | 150.4859691 | 1.38E-12 | 1.744951752 | 2.30E-13 | 7.329876343 | 3.351836441 |
| CNAG_04696 | - | DNA clamp loader | 178.1149439 | 235.052741 | 188.9895452 | 429.670801 | 398.8790921 | 424.7976636 | 1.47E-12 | 1.043572739 | 2.45E-13 | 7.321428082 | 2.061326076 |
| CNAG_03301 | TIM13 | mitochondrial import inner membrane translocase subunit TIM13 | 1605.356259 | 1532.012207 | 1728.537485 | 1119.578576 | 1042.529395 | 1085.951547 | 1.54E-12 | -0.599263653 | 2.57E-13 | -7.315247989 | 0.660090777 |
| CNAG_03025 | - | V-type H -transporting ATPase subunit D | 2178.669528 | 2065.324504 | 2105.066347 | 1547.711998 | 1470.42119 | 1375.738713 | 1.61E-12 | -0.546948568 | 2.69E-13 | -7.309175525 | 0.684466307 |
| CNAG_12944 | - | unspecified product | 108.0177166 | 121.9777473 | 86.66583308 | 259.9985797 | 287.953659 | 247.2322242 | 1.71E-12 | 1.317172024 | 2.86E-13 | 7.300753086 | 2.491771927 |
| CNAG_06669 | - | hypothetical protein | 1215.280304 | 1231.330857 | 1209.223697 | 822.2617624 | 823.5050461 | 699.7140312 | 1.85E-12 | -0.656249135 | 3.09E-13 | -7.29056128 | 0.634525859 |
| CNAG_01471 | - | hypothetical protein | 1233.938815 | 1210.879239 | 1230.54082 | 856.5705422 | 712.9602463 | 776.2445171 | 2.07E-12 | -0.663161964 | 3.47E-13 | -7.274726316 | 0.631492732 |
| CNAG_02924 | - | histone H1/5 | 306.4476234 | 357.8628372 | 294.55214 | 607.0253295 | 559.2467082 | 591.390186 | 2.12E-12 | 0.862066809 | 3.55E-13 | 7.271580627 | 1.817640403 |
| CNAG_02752 | - | Short-chain dehydrogenase | 1150.517659 | 1077.599506 | 935.4822167 | 1815.667574 | 1766.904046 | 1548.551748 | 2.21E-12 | 0.682358933 | 3.70E-13 | 7.266064644 | 1.604761537 |
| CNAG_05939 | - | hypothetical protein, variant | 171.0208322 | 178.0052429 | 137.8425558 | 412.1200018 | 388.7593003 | 311.7893168 | 2.22E-12 | 1.177761921 | 3.72E-13 | 7.265351882 | 2.262255569 |
| CNAG_00607 | - | hypothetical protein | 1417.393507 | 1403.784343 | 1660.226796 | 920.9688858 | 907.0740566 | 1014.872479 | 2.22E-12 | -0.672460076 | 3.72E-13 | -7.265200765 | 0.627435874 |
| CNAG_00961 | - | hypothetical protein | 674.0801583 | 570.1807765 | 447.1117916 | 1147.331898 | 1052.718898 | 998.9050469 | 2.27E-12 | 0.903170547 | 3.80E-13 | 7.262332014 | 1.870171463 |
| CNAG_04292 | - | hypothetical protein | 3237.911178 | 3068.4535 | 3403.972466 | 2304.134978 | 2379.603519 | 2291.566893 | 2.29E-12 | -0.492826796 | 3.85E-13 | -7.26078331 | 0.710631332 |
| CNAG_06066 | - | nucleoside-diphosphate-sugar epimerase | 13.60977174 | 20.8044359 | 30.10530851 | 125.428308 | 100.6357866 | 102.7569721 | 2.33E-12 | 2.361138802 | 3.92E-13 | 7.258281316 | 5.137757518 |
| CNAG_03259 | - | hypothetical protein | 30.02069162 | 65.97867175 | 49.33566043 | 151.6330023 | 208.0723164 | 173.777794 | 2.38E-12 | 1.872001369 | 4.01E-13 | 7.255212277 | 3.660400146 |
| CNAG_00644 | - | sphingolipid delta-4 desaturase | 139.462503 | 126.3265427 | 127.1352753 | 283.205758 | 300.7093814 | 297.1127144 | 2.42E-12 | 1.151129989 | 4.08E-13 | 7.253041307 | 2.220877763 |
| CNAG_05461 | - | L-lactate dehydrogenase (cytochrome) | 54.57385353 | 132.7093869 | 60.05369451 | 270.2966513 | 283.6457597 | 290.243221 | 2.42E-12 | 1.765557495 | 4.09E-13 | 7.252693614 | 3.400053613 |
| CNAG_02841 | - | cytochrome P450 monooxygenase pc-2, variant | 639.1353513 | 662.7745983 | 642.1514219 | 1003.657068 | 1016.391854 | 1026.64884 | 2.55E-12 | 0.6330741 | 4.31E-13 | 7.245584214 | 1.550866067 |
| CNAG_04895 | FZC3 | nuclear protein | 1228.814355 | 1150.889117 | 1323.565705 | 1887.833768 | 1871.176842 | 1812.710438 | 2.58E-12 | 0.574102608 | 4.35E-13 | 7.244282125 | 1.488751135 |
| CNAG_00452 | - | Isovaleryl-CoA dehydrogenase | 81.09951637 | 59.58303308 | 62.12517896 | 201.0804566 | 203.3805893 | 182.8017194 | 2.59E-12 | 1.520344564 | 4.37E-13 | 7.24345688 | 2.868595532 |
| CNAG_12256 | - | unspecified product | 88970.67781 | 86436.87847 | 79626.68715 | 58693.54267 | 67478.89699 | 57968.39657 | 2.59E-12 | -0.48543338 | 4.38E-13 | -7.243116513 | 0.71428247 |
| CNAG_07849 | - | hypothetical protein | 17.0353485 | 15.41139763 | 19.46044455 | 97.8762205 | 91.27273095 | 87.21416337 | 2.62E-12 | 2.415276407 | 4.43E-13 | 7.241761047 | 5.334216597 |
| CNAG_07309 | - | mRNA surveillance protein pelota | 1519.997766 | 1389.78202 | 1455.580958 | 1002.052023 | 991.6954197 | 926.1276316 | 2.88E-12 | -0.595538373 | 4.88E-13 | -7.228626442 | 0.661797445 |
| CNAG_02712 | BUD32 | BUD32 protein kinase | 1820.337944 | 1988.770627 | 1931.226017 | 1381.907094 | 1347.947913 | 1282.464564 | 2.92E-12 | -0.532131068 | 4.95E-13 | -7.226585051 | 0.691532486 |
| CNAG_02982 | - | U3 small nucleolar RNA-associated protein 13 | 2444.856348 | 2198.94856 | 2292.763774 | 1514.842104 | 1682.661575 | 1592.174271 | 3.14E-12 | -0.550073957 | 5.33E-13 | -7.216603915 | 0.682985115 |
| CNAG_03874 | - | oxidoreductase | 146.4511744 | 118.8126441 | 137.7840579 | 298.0681096 | 302.3948076 | 304.8841595 | 3.19E-12 | 1.156809812 | 5.42E-13 | 7.214393103 | 2.229638489 |
| CNAG_06507 | - | hypothetical protein | 20.57647759 | 32.59745923 | 13.1064056 | 111.7251083 | 97.9694312 | 125.7643601 | 3.51E-12 | 2.343991015 | 5.96E-13 | 7.201491824 | 5.077051923 |

| | | | | | | | | | | | | | |
|---|---|---|---|---|---|---|---|---|---|---|---|---|---|
| CNAG_04749 | - | hypothetical protein | 1520.657925 | 1454.673269 | 1289.517043 | 2132.171228 | 2086.767723 | 2166.796344 | 3.51E-12 | 0.567376667 | 5.95E-13 | 7.201541825 | 1.481826631 |
| CNAG_05934 | LIV15 | translin domain protein | 1308.677472 | 1243.234076 | 1273.220557 | 854.9095136 | 840.7132054 | 873.8930527 | 3.53E-12 | -0.590401075 | 6.00E-13 | -7.200595403 | 0.664158243 |
| CNAG_01011 | - | hypothetical protein | 194.4110866 | 232.9105666 | 207.0950813 | 405.3169025 | 434.3731138 | 412.6543087 | 3.54E-12 | 0.968116253 | 6.02E-13 | 7.200017934 | 1.956284579 |
| CNAG_00104 | - | elongation factor 3 | 3907.982848 | 3384.131947 | 3432.830976 | 2529.07036 | 2557.274178 | 2399.286352 | 3.81E-12 | -0.534562129 | 6.48E-13 | -7.190036463 | 0.690368178 |
| CNAG_00011 | - | hypothetical protein | 750.4241867 | 714.1831263 | 736.760544 | 430.6056845 | 459.4518377 | 453.1904137 | 3.84E-12 | -0.728593774 | 6.55E-13 | -7.188616652 | 0.603491864 |
| CNAG_04605 | ERG26 | C-3 sterol dehydrogenase | 5707.091731 | 5848.420784 | 6114.053835 | 7735.691607 | 7956.472978 | 7854.238156 | 3.93E-12 | 0.39895656 | 6.69E-13 | 7.185586347 | 1.318553912 |
| CNAG_03620 | - | hypothetical protein | 1382.20543 | 1346.625785 | 1262.591483 | 915.6692355 | 901.1253342 | 821.7904829 | 3.96E-12 | -0.612851793 | 6.75E-13 | -7.18436597 | 0.653902846 |
| CNAG_04570 | LHP1 | hypothetical protein | 5977.922563 | 5504.745099 | 5582.998886 | 7502.173292 | 7862.464704 | 8032.919225 | 4.00E-12 | 0.439801506 | 6.83E-13 | 7.182760718 | 1.356417691 |
| CNAG_04865 | - | mannitol dehydrogenase | 39.28585406 | 61.66848459 | 47.19746276 | 143.0738621 | 165.4179122 | 182.4771497 | 4.18E-12 | 1.723159669 | 7.15E-13 | 7.176556354 | 3.301587014 |
| CNAG_06567 | - | hypothetical protein | 404.659064 | 569.012145 | 498.2040666 | 885.5610793 | 855.1338206 | 885.5886971 | 4.22E-12 | 0.821632014 | 7.22E-13 | 7.175242208 | 1.767404196 |
| CNAG_04783 | - | Monosaccharide transporter | 2094.185394 | 2330.614068 | 2398.54817 | 3104.058232 | 3468.69207 | 3316.956577 | 4.59E-12 | 0.520485425 | 7.86E-13 | 7.163606926 | 1.434437814 |
| CNAG_05741 | - | hypothetical protein, variant | 1632.357095 | 1882.064765 | 1795.768289 | 1273.7485 | 1127.912724 | 1159.241283 | 4.84E-12 | -0.591548757 | 8.29E-13 | -7.156302014 | 0.663630107 |
| CNAG_01881 | - | molecular chaperone GrpE | 4250.418665 | 4289.158325 | 4623.994369 | 2995.963113 | 3132.585699 | 3361.694037 | 4.89E-12 | -0.487444111 | 8.38E-13 | -7.154815884 | 0.713287645 |
| CNAG_06627 | - | pre-mRNA-processing factor 40 | 3194.586771 | 2866.97228 | 3008.363013 | 2131.340506 | 2173.428667 | 2190.521387 | 4.98E-12 | -0.49739797 | 8.54E-13 | -7.152131129 | 0.708383262 |
| CNAG_04907 | - | hypothetical protein, variant | 57.75496343 | 36.97399486 | 32.30073899 | 176.6928587 | 147.2197382 | 139.4954984 | 5.08E-12 | 1.869588515 | 8.71E-13 | 7.149447986 | 3.654283379 |
| CNAG_02883 | RAC1 | Ras-like GTP-binding protein, putative | 274.8132856 | 231.9025022 | 252.9375912 | 465.9881021 | 492.1133894 | 482.5869872 | 5.10E-12 | 0.910855884 | 8.75E-13 | 7.148880929 | 1.880160579 |
| CNAG_01137 | - | aconitate hydratase, mitochondrial | 15243.26111 | 12954.94668 | 12246.04003 | 9441.832212 | 8963.442686 | 9559.777431 | 5.19E-12 | -0.548150916 | 8.92E-13 | -7.146173404 | 0.683896108 |
| CNAG_03760 | RINT1 | hypothetical protein | 2454.180705 | 2266.795316 | 2212.804617 | 1580.292394 | 1476.381158 | 1687.594713 | 5.44E-12 | -0.562814768 | 9.35E-13 | -7.139737217 | 0.676980053 |
| CNAG_00426 | - | UBA/TS-N domain-containing protein | 216.6186965 | 279.2152569 | 224.1763634 | 485.6420313 | 439.3371722 | 501.3630937 | 5.44E-12 | 0.972878197 | 9.37E-13 | 7.139478367 | 1.96275241 |
| CNAG_04655 | - | rab family protein | 336.7794485 | 342.8311107 | 363.814278 | 662.3666092 | 604.1278961 | 579.2938851 | 5.60E-12 | 0.810402248 | 9.64E-13 | 7.13548727 | 1.753700336 |
| CNAG_00877 | - | adenylate kinase, variant | 1117.224054 | 1258.173264 | 1054.643104 | 688.2271999 | 730.2967518 | 760.6623639 | 5.67E-12 | -0.669341049 | 9.76E-13 | -7.133780478 | 0.628793823 |
| CNAG_02508 | - | hypothetical protein | 714.7777408 | 1200.877821 | 1145.030116 | 496.612747 | 487.7537864 | 527.6437157 | 5.76E-12 | -1.031705223 | 9.93E-13 | -7.131459326 | 0.489131668 |
| CNAG_03326 | CHS2 | Chitin synthase | 234.0939078 | 289.983309 | 209.2746348 | 491.829792 | 457.4771106 | 519.1099796 | 5.88E-12 | 0.988925743 | 1.02E-12 | 7.128437113 | 1.984706591 |
| CNAG_05007 | - | translation initiation factor eIF-2B subunit beta | 1841.381368 | 1986.640285 | 2083.684797 | 1304.721347 | 1353.102273 | 1421.16974 | 6.09E-12 | -0.550284762 | 1.05E-12 | -7.12368626 | 0.682885326 |
| CNAG_02290 | - | parafibromin | 2328.140997 | 2230.161671 | 2188.271392 | 1498.514585 | 1555.80464 | 1676.483418 | 6.18E-12 | -0.527807295 | 1.07E-12 | -7.121414027 | 0.693608125 |
| CNAG_01645 | - | hypothetical protein | 15.92897915 | 30.44662653 | 17.35884765 | 114.2456866 | 124.3993173 | 84.12670936 | 6.29E-12 | 2.36306353 | 1.09E-12 | 7.118917568 | 5.144616475 |
| CNAG_06916 | - | hypothetical protein | 866.0309824 | 781.0337253 | 884.9621762 | 517.8297544 | 538.820941 | 504.2855889 | 6.37E-12 | -0.714905735 | 1.10E-12 | -7.117040683 | 0.609244938 |
| CNAG_01419 | - | hypothetical protein | 1386.382658 | 1304.926299 | 1259.631479 | 2083.302237 | 1937.554239 | 1859.379972 | 6.46E-12 | 0.558643301 | 1.12E-12 | 7.115152444 | 1.472883477 |
| CNAG_06283 | LIV4 | putative myb-like mRNA polymerase I termination factor | 1198.84412 | 1094.557573 | 1178.272723 | 670.2715616 | 785.5599248 | 757.3461366 | 6.58E-12 | -0.665340186 | 1.14E-12 | -7.112472037 | 0.630540006 |
| CNAG_02878 | GEP4 | hypothetical protein | 1246.816582 | 1169.992692 | 1354.187241 | 849.5076681 | 744.8900983 | 812.8643719 | 6.79E-12 | -0.663651871 | 1.18E-12 | -7.107912183 | 0.631278328 |
| CNAG_02907 | - | tRNA (guanine-N(1)-)-methyltransferase | 1639.084354 | 1446.907691 | 1586.736554 | 1074.441028 | 975.227801 | 1050.418448 | 7.07E-12 | -0.607624954 | 1.23E-12 | -7.102360954 | 0.656276212 |
| CNAG_03591 | - | rab GDP-dissociation inhibitor | 6187.669506 | 5907.488676 | 6176.80061 | 4700.589963 | 4539.444395 | 4649.504997 | 7.63E-12 | -0.411083257 | 1.33E-12 | -7.091663604 | 0.752058474 |
| CNAG_03740 | - | hypothetical protein | 17.1900956 | 42.3428624 | 54.61911531 | 144.8433626 | 179.5347055 | 159.2936693 | 7.85E-12 | 2.078743806 | 1.36E-12 | 7.087587972 | 4.224392266 |
| CNAG_04002 | RVS161 | hypothetical protein | 2867.709924 | 2795.82723 | 2921.952135 | 2194.548329 | 1895.610057 | 1928.575894 | 8.23E-12 | -0.527761156 | 1.43E-12 | -7.08105845 | 0.693630308 |
| CNAG_05412 | - | hypothetical protein | 13.51960692 | 13.24517323 | 14.13186314 | 78.0897268 | 93.29260088 | 73.95110598 | 8.34E-12 | 2.593222357 | 1.45E-12 | 7.079119802 | 6.034450303 |
| CNAG_06763 | - | serine-tRNA ligase | 863.7841449 | 837.0321476 | 906.3043243 | 556.014325 | 529.2801066 | 561.9629216 | 8.49E-12 | -0.678231661 | 1.48E-12 | -7.076460224 | 0.624930795 |
| CNAG_01146 | - | U3 small nucleolar RNA-associated protein 22 | 3554.224746 | 3129.870578 | 3374.139074 | 2189.605868 | 2372.674186 | 2466.92912 | 8.50E-12 | -0.533042776 | 1.48E-12 | -7.076202622 | 0.691095612 |
| CNAG_02845 | - | hypothetical protein | 3103.569515 | 2836.815661 | 3187.481554 | 2153.145228 | 2095.757577 | 2230.473754 | 8.51E-12 | -0.510081737 | 1.48E-12 | -7.075920588 | 0.702182654 |
| CNAG_00811 | - | anthranilate phosphoribosyltransferase | 1755.061681 | 2137.41085 | 2178.562248 | 1262.638373 | 1414.386804 | 1278.047789 | 8.81E-12 | -0.633100328 | 1.54E-12 | -7.070997586 | 0.644789284 |
| CNAG_04579 | - | Alpha-1,2-mannosyltransferase | 420.8046834 | 394.5615272 | 451.2430757 | 718.9241158 | 739.6148967 | 692.5143577 | 9.19E-12 | 0.748885969 | 1.60E-12 | 7.065184897 | 1.680494672 |
| CNAG_00235 | AMT1 | amt family ammonium transporter | 2903.930623 | 2769.99747 | 3011.527789 | 2161.011361 | 2010.3664 | 2080.624236 | 9.37E-12 | -0.489567301 | 1.64E-12 | -7.062284954 | 0.712238683 |
| CNAG_03413 | - | alginate lyase | 653.2258733 | 780.1743463 | 650.7174406 | 1057.249838 | 1134.593487 | 1200.849588 | 9.48E-12 | 0.688683243 | 1.66E-12 | 7.060590094 | 1.611811736 |
| CNAG_01021 | - | ATP phosphoribosyltransferase | 1443.152069 | 1591.188165 | 1550.491054 | 1060.420843 | 1085.748266 | 990.5104584 | 9.50E-12 | -0.563113605 | 1.66E-12 | -7.060151629 | 0.67683984 |
| CNAG_05933 | - | hypothetical protein | 2294.441973 | 2363.771565 | 2588.10723 | 1588.003408 | 1661.898245 | 1778.625588 | 9.51E-12 | -0.542477805 | 1.67E-12 | -7.059970871 | 0.686590688 |
| CNAG_00542 | - | Salicylate hydroxylase | 139.5503061 | 139.2613561 | 161.2217032 | 352.0523772 | 343.8964396 | 281.7923271 | 9.56E-12 | 1.137087449 | 1.68E-12 | 7.059165815 | 2.199365609 |
| CNAG_00935 | - | DNA-binding protein | 2522.000817 | 2474.721849 | 2306.673307 | 1659.64547 | 1799.972687 | 1747.573198 | 9.71E-12 | -0.503694602 | 1.70E-12 | -7.056836147 | 0.705298266 |
| CNAG_04166 | - | smad nuclear-interacting protein 1, variant | 1646.31338 | 1724.825747 | 1835.189989 | 1271.448594 | 1107.213791 | 1100.426095 | 9.74E-12 | -0.596751892 | 1.71E-12 | -7.056395106 | 0.66124101 |
| CNAG_06379 | - | N-acetyltransferase 10 | 2782.472525 | 2656.860987 | 2884.615196 | 1851.073542 | 1981.074624 | 2072.81364 | 9.86E-12 | -0.510733371 | 1.73E-12 | -7.054622277 | 0.701865565 |
| CNAG_05575 | - | hypothetical protein | 1117.368297 | 1243.178859 | 1276.368844 | 819.9125371 | 781.1816265 | 798.4280616 | 9.92E-12 | -0.615384674 | 1.74E-12 | -7.053646269 | 0.652755823 |
| CNAG_03541 | PTC7 | hypothetical protein | 2277.802553 | 1896.23177 | 2264.961134 | 1470.593666 | 1432.454388 | 1269.21224 | 1.05E-11 | -0.642292188 | 1.84E-12 | -7.046242874 | 0.64069419 |
| CNAG_06167 | - | metal homeostatis protein bsd2 | 4149.422515 | 4718.187121 | 4862.039307 | 6272.54813 | 6480.665761 | 6396.569515 | 1.05E-11 | 0.46513441 | 1.85E-12 | 7.045429038 | 1.380445958 |
| CNAG_07346 | - | T-complex protein 1 subunit eta | 8980.959473 | 8651.712973 | 9027.435639 | 6818.291485 | 7000.383705 | 6840.963038 | 1.06E-11 | -0.383366301 | 1.86E-12 | -7.04465828 | 0.766646653 |
| CNAG_01020 | - | hypothetical protein | 1890.350221 | 1943.551129 | 1934.432697 | 1293.054031 | 1385.918723 | 1410.063206 | 1.06E-11 | -0.51156811 | 1.87E-12 | -7.043615558 | 0.701459585 |
| CNAG_04829 | - | hypothetical protein | 379.9206098 | 343.9320707 | 389.4039119 | 681.7422658 | 636.9017858 | 621.4590632 | 1.08E-11 | 0.787058494 | 1.90E-12 | 7.041522894 | 1.72555265 |
| CNAG_12940 | - | unspecified product | 82.47268633 | 108.0102872 | 123.8954313 | 287.6058616 | 243.6524199 | 254.943832 | 1.17E-11 | 1.31013946 | 2.06E-12 | 7.030057015 | 2.479655087 |
| CNAG_06968 | - | hypothetical protein | 749.154998 | 759.3248822 | 657.8976033 | 395.5321928 | 354.9117424 | 466.4533734 | 1.18E-11 | -0.848099018 | 2.08E-12 | -7.029046562 | 0.555516236 |
| CNAG_01671 | - | acetyl/propionyl CoA carboxylase | 61.31215636 | 52.02869684 | 41.8914331 | 135.8069471 | 208.0954535 | 173.7631538 | 1.23E-11 | 1.728552934 | 2.17E-12 | 7.022784245 | 3.313952523 |
| CNAG_07495 | - | hypothetical protein | 4.11998428 | 5.678678203 | 4.516738122 | 45.67227385 | 70.95461176 | 65.87976729 | 1.23E-11 | 3.776328649 | 2.17E-12 | 7.022841483 | 13.70213358 |
| CNAG_06238 | - | Glutathione S-transferase | 85.85799751 | 99.36038487 | 54.70982576 | 215.5704606 | 270.9402136 | 210.5890582 | 1.27E-11 | 1.531466425 | 2.25E-12 | 7.018172709 | 2.890795243 |
| CNAG_04980 | - | U6 snRNA-associated Sm-like protein LSm1 | 1598.391873 | 1631.093225 | 1620.894536 | 1126.56611 | 1184.965952 | 1085.964552 | 1.34E-11 | -0.529552656 | 2.38E-12 | -7.010170126 | 0.692769511 |
| CNAG_02013 | - | hypothetical protein, variant | 357.0356216 | 472.5491719 | 374.1697093 | 200.1390058 | 187.5846896 | 205.6894945 | 1.36E-11 | -1.036809407 | 2.41E-12 | -7.008472545 | 0.487404202 |
| CNAG_03441 | - | small subunit ribosomal protein S21 | 1714.986973 | 1500.786493 | 1685.903053 | 1081.38235 | 999.3609836 | 1146.96658 | 1.51E-11 | -0.618432849 | 2.68E-12 | -6.993773749 | 0.651378114 |
| CNAG_02484 | - | peptidyl-prolyl cis-trans isomerase, variant | 22.88590179 | 30.45243277 | 8.841884651 | 128.6161371 | 112.04494 | 92.91292244 | 1.52E-11 | 2.454539083 | 2.70E-12 | 6.992744677 | 5.481379729 |
| CNAG_02471 | - | hypothetical protein | 1164.99767 | 1074.084999 | 1120.709391 | 727.2338385 | 775.2167665 | 707.4252509 | 1.55E-11 | -0.61990576 | 2.76E-12 | -6.989412058 | 0.650713432 |
| CNAG_03008 | - | hypothetical protein | 1027.162683 | 946.9311049 | 952.2250898 | 612.786427 | 651.8418881 | 628.5831002 | 1.58E-11 | -0.64488524 | 2.81E-12 | -6.98704405 | 0.639543661 |
| CNAG_06754 | - | RNA polymerase-associated protein CTR9 | 8734.548721 | 8378.042306 | 9039.136985 | 6159.743715 | 6506.093431 | 6856.470417 | 1.59E-11 | -0.437312067 | 2.83E-12 | -6.985795643 | 0.738509269 |
| CNAG_04853 | - | derlin-2/3 | 270.1247082 | 238.3426008 | 227.3682928 | 464.5176377 | 447.1447114 | 481.431197 | 1.64E-11 | 0.905875896 | 2.92E-12 | 6.981307154 | 1.873681708 |
| CNAG_00519 | ERG3 | lathosterol oxidase | 765.6342686 | 771.2443404 | 780.4641487 | 471.1572708 | 379.0523173 | 498.6596159 | 1.66E-11 | -0.795687431 | 2.96E-12 | -6.97969253 | 0.576068617 |
| CNAG_06555 | - | aromatic amino acid aminotransferase I | 671.1370806 | 718.4320339 | 699.4319079 | 429.9056969 | 448.2686871 | 385.5423426 | 1.67E-11 | -0.741552766 | 2.97E-12 | -6.979147949 | 0.598095279 |
| CNAG_03994 | - | leucine carboxyl methyltransferase 1 | 117.2893483 | 114.4545898 | 80.27852869 | 291.5522997 | 261.0355564 | 233.9710494 | 1.71E-11 | 1.320699461 | 3.05E-12 | 6.975431377 | 2.497871846 |
| CNAG_04378 | - | NADH dehydrogenase (ubiquinone) 1 alpha subcomplex 13 | 3504.205525 | 3422.922818 | 3953.154353 | 2603.81033 | 2650.453296 | 2353.796178 | 1.74E-11 | -0.532024361 | 3.10E-12 | -6.973174774 | 0.691583636 |
| CNAG_06774 | - | tRNA (guanine-N(7)-)-methyltransferase | 1248.958684 | 1049.340946 | 1196.387165 | 706.9285117 | 681.0577062 | 789.5050857 | 1.74E-11 | -0.69864172 | 3.11E-12 | -6.972715992 | 0.616152033 |
| CNAG_07864 | - | U3 small nucleolar ribonucleoprotein IMP3 | 716.2618538 | 716.690744 | 803.1574706 | 1354.303903 | 1159.525134 | 1122.221197 | 1.83E-11 | 0.686333941 | 3.27E-12 | 6.965517423 | 1.609189179 |
| CNAG_03782 | - | hypothetical protein | 5.30367251 | 2.442265342 | 6.653322381 | 70.06180052 | 81.53499279 | 39.8781144 | 1.86E-11 | 3.852108013 | 3.32E-12 | 6.963384096 | 14.44109278 |
| CNAG_03004 | - | riboflavin synthase, alpha subunit | 2218.396265 | 2093.351895 | 2147.731675 | 1594.410551 | 1545.485709 | 1460.087547 | 1.87E-11 | -0.505311319 | 3.35E-12 | -6.962083022 | 0.704508335 |
| CNAG_00157 | - | hypothetical protein | 951.0769367 | 843.460517 | 801.8689524 | 572.4543473 | 498.2019857 | 459.9446361 | 1.93E-11 | -0.778823001 | 3.45E-12 | -6.958087596 | 0.582842101 |
| CNAG_05240 | - | signal recognition particle subunit SRP68 | 3462.134023 | 3291.485465 | 3579.957256 | 2522.083871 | 2559.01448 | 2555.755387 | 1.94E-11 | -0.451782237 | 3.47E-12 | -6.957285856 | 0.731139075 |
| CNAG_04935 | - | hypothetical protein | 548.5269841 | 574.0812145 | 611.9241193 | 350.5190572 | 316.1691602 | 341.0814427 | 2.05E-11 | -0.798261171 | 3.68E-12 | -6.949088358 | 0.575041837 |
| CNAG_04098 | PDR5-2 | ATP-binding cassette, subfamily G (WHITE), member 2, PDR | 860.9808045 | 911.6358065 | 631.5948665 | 1403.883298 | 1403.774572 | 1341.939622 | 2.05E-11 | 0.773188346 | 3.68E-12 | 6.948913603 | 1.709042585 |
| CNAG_01629 | - | multifunctional methyltransferase subunit TRM112 | 1173.322535 | 1161.357651 | 1299.813175 | 840.9711922 | 776.864994 | 775.136248 | 2.14E-11 | -0.61893556 | 3.84E-12 | -6.943049863 | 0.651151179 |
| CNAG_01627 | - | amino oxidase | 292.2725445 | 245.8898505 | 203.9518975 | 493.3691476 | 471.3150224 | 514.6971197 | 2.16E-11 | 0.982520696 | 3.89E-12 | 6.94119696 | 1.97591474 |
| CNAG_05517 | - | solute carrier family 45, member 1/2/4 | 851.6657772 | 899.7988011 | 879.9572808 | 1336.773074 | 1427.935322 | 1238.817195 | 2.19E-11 | 0.590623932 | 3.93E-12 | 6.939685109 | 1.505897872 |
| CNAG_07422 | DHA1 | delayed-type hypersensitivity antigen | 0.531883061 | 1.360410985 | 2.387777428 | 75.45579048 | 68.8968251 | 81.28726558 | 2.26E-11 | 6.207653804 | 4.06E-12 | 6.935080211 | 73.90775281 |
| CNAG_06032 | - | ADP-ribosylation factor-like 2, variant | 647.8013111 | 709.7982165 | 701.5396029 | 353.4095725 | 414.5799811 | 439.8284253 | 2.31E-11 | -0.785207768 | 4.16E-12 | -6.931533196 | 0.580268384 |
| CNAG_03657 | LDP1 | hypothetical protein | 745.3107766 | 698.3440463 | 678.4213399 | 1085.379209 | 1079.328554 | 1108.801663 | 2.31E-11 | 0.610044691 | 4.16E-12 | 6.931568494 | 1.526306489 |
| CNAG_00813 | - | hypothetical protein | 65.86848461 | 25.16325212 | 21.66297827 | 162.2484158 | 175.9822679 | 162.6292555 | 2.34E-11 | 2.150201224 | 4.22E-12 | 6.929681479 | 4.438896973 |
| CNAG_05571 | - | U3 small nucleolar RNA-associated protein 20 | 5129.545112 | 4707.204418 | 4637.933161 | 3253.556176 | 3521.62361 | 3676.985372 | 2.37E-11 | -0.485416827 | 4.28E-12 | -6.927588325 | 0.714290666 |

| | | | | | | | | | | | | | |
|---|---|---|---|---|---|---|---|---|---|---|---|---|---|
| CNAG_07735 | - | U3 small nucleolar RNA-associated protein 25 | 2305.778185 | 1975.917996 | 1976.058173 | 1400.482514 | 1465.265046 | 1387.926821 | 2.44E-11 | -0.573034954 | 4.41E-12 | -6.923410503 | 0.672201211 |
| CNAG_03477 | - | hypothetical protein | 181.4646199 | 185.5229077 | 151.6776567 | 356.4307039 | 354.1032063 | 346.042566 | 2.46E-11 | 1.015375475 | 4.44E-12 | 6.922320861 | 2.02142892 |
| CNAG_01889 | - | Glutathione S-transferase | 66.10805811 | 99.33765148 | 63.21219996 | 194.4194952 | 202.2962277 | 260.0973526 | 2.47E-11 | 1.511393256 | 4.46E-12 | 6.921924153 | 2.850852221 |
| CNAG_04692 | - | thymidylate synthase | 1366.615258 | 1491.27459 | 1383.321017 | 2146.276336 | 2034.154256 | 1942.653397 | 2.47E-11 | 0.515063816 | 4.46E-12 | 6.92179455 | 1.429057351 |
| CNAG_03851 | - | general transcription factor 3C polypeptide 5 (transcription factor C subunit 1) | 98.63786652 | 92.93137218 | 86.6338628 | 228.0841822 | 264.7634783 | 201.8149068 | 2.53E-11 | 1.31248711 | 4.58E-12 | 6.918089903 | 2.483693434 |
| CNAG_07659 | - | hypothetical protein | 490.8782712 | 556.0837291 | 456.6381869 | 886.4076178 | 824.0540424 | 796.8949068 | 2.59E-11 | 0.723894401 | 4.69E-12 | 6.91467876 | 1.651634433 |
| CNAG_00853 | - | exosome complex component RRP43 | 1185.955329 | 1063.310908 | 1095.125425 | 689.0315696 | 711.3115446 | 770.6290309 | 2.73E-11 | -0.639040753 | 4.95E-12 | -6.90701231 | 0.642139765 |
| CNAG_06876 | - | Alpha-ketoglutarate-dependent taurine dioxygenase | 504.9439607 | 427.5930855 | 457.3256919 | 271.1325249 | 234.1945266 | 235.6875071 | 2.79E-11 | -0.923251505 | 5.06E-12 | -6.903886233 | 0.527319223 |
| CNAG_00739 | - | hypothetical protein | 1988.298293 | 1836.920559 | 1867.259706 | 1314.902926 | 1400.614036 | 1269.144682 | 2.79E-11 | -0.530526616 | 5.07E-12 | -6.903685181 | 0.692301982 |
| CNAG_07446 | - | mitochondrial protein | 1336.792319 | 1385.412681 | 1324.430009 | 974.099443 | 885.5545083 | 910.5470848 | 2.81E-11 | -0.561771345 | 5.10E-12 | -6.902849843 | 0.677469854 |
| CNAG_05528 | - | hypothetical protein | 14.73789217 | 25.06703088 | 20.53102566 | 92.07299759 | 88.53570552 | 106.9528795 | 2.87E-11 | 2.26356438 | 5.22E-12 | 6.89952583 | 4.801763604 |
| CNAG_00374 | - | hypothetical protein | 291.3442376 | 376.172995 | 331.8484962 | 632.7209338 | 638.7777571 | 550.4659 | 2.91E-11 | 0.852366311 | 5.30E-12 | 6.897357543 | 1.805459817 |
| CNAG_05584 | - | methylthioribulose-1-phosphate dehydratase | 1559.787301 | 1437.217777 | 1627.241108 | 1039.361252 | 1076.242476 | 1042.654317 | 2.91E-11 | -0.566007346 | 5.30E-12 | -6.89736651 | 0.675483603 |
| CNAG_06759 | LPI1 | dehydrogenase | 620.4810355 | 562.1783264 | 525.6128471 | 342.8123533 | 288.5529567 | 213.6209863 | 2.98E-11 | -1.03363375 | 5.42E-12 | -6.894061619 | 0.488478256 |
| CNAG_03710 | ECM22 | hypothetical protein | 1489.73613 | 1413.494276 | 1546.200955 | 1064.372755 | 1005.483269 | 962.7539814 | 2.99E-11 | -0.568928272 | 5.45E-12 | -6.893453088 | 0.674117381 |
| CNAG_02564 | SEN54 | tRNA-splicing endonuclease subunit Sen54 | 1026.243709 | 1177.402477 | 1081.258989 | 752.2354504 | 676.7780776 | 714.0595478 | 3.05E-11 | -0.631047629 | 5.56E-12 | -6.890564725 | 0.645707357 |
| CNAG_00874 | - | multisite-specific tRNA:(cytosine-C5)- methyltransferase | 4114.905665 | 3968.099802 | 4221.96786 | 2883.924358 | 3225.842227 | 2927.690451 | 3.05E-11 | -0.461053066 | 5.56E-12 | -6.890418946 | 0.726455803 |
| CNAG_02421 | - | hypothetical protein | 1421.92632 | 1289.596087 | 1421.425784 | 868.0515755 | 987.4396044 | 887.260505 | 3.09E-11 | -0.607146962 | 5.65E-12 | -6.888314061 | 0.656493685 |
| CNAG_02194 | - | AGC/NDR protein kinase | 583.0815017 | 617.5099125 | 549.3969906 | 882.925247 | 994.0923941 | 922.3146495 | 3.11E-11 | 0.662808558 | 5.68E-12 | 6.887423377 | 1.583161637 |
| CNAG_06413 | - | hypothetical protein | 78.86719033 | 80.01680215 | 89.7984037 | 209.4093162 | 188.3631048 | 243.5782975 | 3.18E-11 | 1.3586371 | 5.81E-12 | 6.884219906 | 2.564428059 |
| CNAG_07717 | - | ubiquitin carboxyl-terminal hydrolase 4/11/15 | 7045.827562 | 6259.81379 | 6427.435597 | 4634.299822 | 4965.578709 | 4932.599919 | 3.19E-11 | -0.457136187 | 5.84E-12 | -6.883530768 | 0.728430791 |
| CNAG_06866 | - | hypothetical protein | 1224.683993 | 1236.750141 | 1249.752641 | 829.9985324 | 806.1955148 | 883.8530798 | 3.24E-11 | -0.573863711 | 5.93E-12 | -6.881358418 | 0.671815175 |
| CNAG_05500 | - | hypothetical protein, variant | 279.6210643 | 323.4085405 | 324.3540545 | 562.5964044 | 529.0551538 | 552.5546336 | 3.52E-11 | 0.812509292 | 6.46E-12 | 6.869201114 | 1.756263472 |
| CNAG_02294 | ADE2 | phosphoribosylaminoimidazole carboxylase | 2305.96394 | 2085.841465 | 2489.963355 | 1450.217741 | 1417.710237 | 1673.150392 | 3.54E-11 | -0.615141385 | 6.49E-12 | -6.868433071 | 0.652865909 |
| CNAG_03366 | ZNF2 | C2H2 type zinc finger transcription factor | 82.37154538 | 95.06155989 | 78.1129975 | 181.6746376 | 237.0229915 | 252.4642538 | 3.68E-11 | 1.379023269 | 6.75E-12 | 6.862879551 | 2.600922242 |
| CNAG_07393 | - | hypothetical protein | 28.73660135 | 31.56285154 | 41.81743024 | 128.8436779 | 120.5073466 | 118.329988 | 3.80E-11 | 1.856617025 | 6.98E-12 | 6.858158633 | 3.621574428 |
| CNAG_06824 | - | hypothetical protein | 224.6780534 | 221.0975915 | 223.0777317 | 413.8428622 | 473.2954415 | 394.9989982 | 3.90E-11 | 0.922691868 | 7.16E-12 | 6.85438975 | 1.895649012 |
| CNAG_02763 | - | phenylalanine-tRNA ligase, beta subunit | 4033.223525 | 4197.551281 | 4253.963986 | 2774.106701 | 3243.087146 | 3068.649711 | 3.93E-11 | -0.473655694 | 7.22E-12 | -6.853294055 | 0.720137503 |
| CNAG_05024 | - | L-xylulose reductase | 115.099976 | 148.9038994 | 148.4254778 | 309.062645 | 282.4225925 | 310.413913 | 4.12E-11 | 1.117972633 | 7.58E-12 | 6.846312315 | 2.170417572 |
| CNAG_04048 | - | 53 kda brg1-associated factor b | 2222.102742 | 2442.393787 | 2467.639664 | 1742.256764 | 1744.754858 | 1662.117198 | 4.12E-11 | -0.485063131 | 7.59E-12 | -6.846108511 | 0.714465805 |
| CNAG_00561 | HAT1 | histone acetyltransferase type B catalytic subunit | 2481.255335 | 2493.073657 | 2570.038296 | 1917.429216 | 1814.595734 | 1815.299385 | 4.29E-11 | -0.459126145 | 7.90E-12 | -6.840300285 | 0.727426735 |
| CNAG_02220 | - | hypothetical protein | 1932.378912 | 1889.720923 | 2004.803061 | 1394.327649 | 1404.028376 | 1398.982992 | 4.29E-11 | -0.488646376 | 7.90E-12 | -6.840318051 | 0.712693477 |
| CNAG_05316 | - | inositol oxygenase | 486.6065326 | 535.227583 | 504.2653696 | 268.6268154 | 292.0652324 | 308.8594324 | 4.33E-11 | -0.827858942 | 7.99E-12 | -6.838698375 | 0.563364694 |
| CNAG_05258 | SMG1 | glucose-methanol-choline oxidoreductase | 123.5920427 | 42.46626496 | 31.3028438 | 1254.094621 | 1119.136296 | 1020.546081 | 4.34E-11 | 4.093598588 | 8.02E-12 | 6.838262062 | 17.07245455 |
| CNAG_01439 | - | U4/U6 small nuclear ribonucleoprotein PRP4 | 2739.246943 | 2622.372655 | 2643.647637 | 1930.568433 | 1910.332494 | 2033.952624 | 4.47E-11 | -0.461965656 | 8.26E-12 | -6.833958399 | 0.725996423 |
| CNAG_00111 | - | splicing factor 3A subunit 1 | 3278.763261 | 3204.174007 | 3140.626361 | 2498.847296 | 2304.522974 | 2310.435359 | 4.52E-11 | -0.45135711 | 8.35E-12 | -6.83246846 | 0.731354556 |
| CNAG_02500 | - | calnexin | 9362.657816 | 8429.75511 | 8341.740781 | 6465.691542 | 6561.304879 | 6587.849853 | 4.80E-11 | -0.429683416 | 8.87E-12 | -6.823760662 | 0.742424685 |
| CNAG_07485 | - | membrane protein, variant | 1447.895656 | 1631.070392 | 1704.015318 | 1087.632858 | 1100.381407 | 1083.72698 | 4.87E-11 | -0.563222407 | 9.00E-12 | -6.821601552 | 0.676788797 |
| CNAG_04460 | - | RNA polymerase II transcription factor | 3035.933614 | 2960.704242 | 3198.163685 | 2202.953523 | 2301.983573 | 2289.310676 | 4.87E-11 | -0.452160592 | 9.01E-12 | -6.821492282 | 0.730947355 |
| CNAG_01518 | PMU1 | phosphoglycerate mutase, variant | 950.351547 | 1036.317315 | 1063.083843 | 661.082093 | 656.1138863 | 689.6070281 | 4.90E-11 | -0.618935873 | 9.08E-12 | -6.820407397 | 0.651151037 |
| CNAG_01368 | - | hypothetical protein, variant | 67.02522814 | 26.23861813 | 24.85758241 | 161.4460554 | 175.1032008 | 160.4347965 | 4.92E-11 | 2.065563288 | 9.12E-12 | 6.819744317 | 4.185973817 |
| CNAG_05881 | ALG9 | Alpha-1,2-mannosyltransferase | 2914.358622 | 2764.579911 | 2644.742525 | 2039.558827 | 2065.636942 | 1984.050121 | 5.19E-11 | -0.466411843 | 9.64E-12 | -6.811834666 | 0.723762446 |
| CNAG_05906 | - | hypothetical protein | 419.5738597 | 394.5334964 | 345.7466464 | 754.278803 | 693.0254793 | 612.6658327 | 5.20E-11 | 0.814896173 | 9.65E-12 | 6.811701369 | 1.759171545 |
| CNAG_06524 | FRE3 | hypothetical protein, variant | 12.33316253 | 8.938498475 | 11.99557411 | 61.32479835 | 62.51331254 | 105.3938915 | 5.27E-11 | 2.863281636 | 9.79E-12 | 6.809626829 | 7.276686394 |
| CNAG_04840 | - | small nuclear ribonucleoprotein B and B' | 3183.5221 | 3127.90336 | 2953.139058 | 4433.592662 | 4028.28268 | 4306.130485 | 5.33E-11 | 0.447746783 | 9.91E-12 | 6.807785444 | 1.363908425 |
| CNAG_04033 | - | alpha 1,3-glucosidase | 2570.00652 | 2866.841767 | 2537.016937 | 1962.546374 | 1933.658485 | 1704.361684 | 5.53E-11 | -0.524759211 | 1.03E-11 | -6.802498849 | 0.69507511 |
| CNAG_04395 | - | protein BFR2 | 1676.699238 | 1751.776083 | 1812.831761 | 1219.176306 | 1215.094391 | 1282.393821 | 5.55E-11 | -0.510609304 | 1.03E-11 | -6.801844121 | 0.701925926 |
| CNAG_05311 | - | arsenite-resistant protein ASR2, variant | 6343.0747 | 6472.047209 | 6714.284843 | 4907.560896 | 4998.352418 | 5166.810273 | 5.55E-11 | -0.389029142 | 1.03E-11 | -6.801652681 | 0.763643323 |
| CNAG_01513 | - | hypothetical protein | 405.0552476 | 523.3075679 | 523.3499224 | 232.7604642 | 233.305783 | 283.3257974 | 5.58E-11 | -0.970097295 | 1.04E-11 | -6.800873296 | 0.510471635 |
| CNAG_07465 | - | hypothetical protein | 909.2499051 | 866.1054451 | 859.4387916 | 588.7680701 | 578.5145368 | 519.8448821 | 5.73E-11 | -0.660013693 | 1.07E-11 | -6.797085002 | 0.63287229 |
| CNAG_05045 | - | hypothetical protein | 387.6734333 | 574.9521364 | 558.511549 | 240.5402354 | 268.7496522 | 247.9099328 | 5.79E-11 | -1.022546564 | 1.08E-11 | -6.795428021 | 0.492246699 |
| CNAG_06539 | - | monocarboxylic acid transporter, variant | 37.99106404 | 35.8556343 | 23.7650499 | 103.249306 | 161.672541 | 116.1610726 | 5.99E-11 | 1.984349317 | 1.12E-11 | 6.790422173 | 3.956841611 |
| CNAG_07746 | - | methylenetetrahydrofolate dehydrogenase (NADP) | 4990.491245 | 4538.018402 | 4085.553318 | 3231.865027 | 3259.395177 | 3178.590963 | 6.14E-11 | -0.5092503 | 1.15E-11 | -6.786772545 | 0.702587444 |
| CNAG_00305 | - | chaperone activator | 5547.714419 | 5208.238099 | 5474.006522 | 4070.914626 | 3919.237027 | 4233.168184 | 6.15E-11 | -0.424543408 | 1.15E-11 | -6.786491175 | 0.745074499 |
| CNAG_06542 | - | hypothetical protein | 1522.583813 | 1645.092337 | 1754.134463 | 1049.402303 | 1089.133229 | 1193.571818 | 6.24E-11 | -0.577949806 | 1.17E-11 | -6.784200191 | 0.669915108 |
| CNAG_01411 | - | hypothetical protein | 33.41715789 | 43.38630019 | 46.09385461 | 147.5973161 | 137.7960356 | 122.8800415 | 6.51E-11 | 1.719807207 | 1.22E-11 | 6.777981976 | 3.29392386 |
| CNAG_02240 | - | cytoplasmic protein | 3418.945888 | 3350.725744 | 3413.626375 | 2573.492259 | 2565.058527 | 2591.265446 | 6.60E-11 | -0.412857719 | 1.24E-11 | -6.775934853 | 0.751134038 |
| CNAG_04023 | - | hypothetical protein | 2040.934354 | 2108.366832 | 1967.518199 | 1310.121061 | 1375.509437 | 1551.01175 | 6.67E-11 | -0.544554107 | 1.25E-11 | -6.774325011 | 0.685603269 |
| CNAG_03554 | - | coatomer protein complex, subunit alpha (xenin) | 12498.19916 | 11941.10835 | 11910.0887 | 9326.716495 | 9572.537491 | 9635.213786 | 6.70E-11 | -0.364723588 | 1.26E-11 | -6.77355592 | 0.776617656 |
| CNAG_05072 | - | endopeptidase | 21.63765156 | 17.55060831 | 15.20790596 | 94.75537933 | 78.11242608 | 90.44505917 | 6.81E-11 | 2.290154962 | 1.28E-11 | 6.77106771 | 4.891086441 |
| CNAG_07590 | - | V-type H -transporting ATPase subunit C | 1336.703301 | 1406.902116 | 1180.513693 | 900.8796846 | 870.0530608 | 836.1988116 | 7.00E-11 | -0.604623808 | 1.31E-11 | -6.766982115 | 0.657642842 |
| CNAG_03985 | GRX5 | Grx4 family monothiol glutaredoxin | 2654.018527 | 2527.570859 | 2699.067909 | 1979.686162 | 1934.522505 | 1767.612692 | 7.40E-11 | -0.487915442 | 1.39E-11 | -6.758759876 | 0.71305465 |
| CNAG_07951 | - | WD-repeat protein JIP5 | 745.8434992 | 726.0532684 | 791.1187961 | 441.4858241 | 498.3104614 | 472.0599877 | 7.40E-11 | -0.696404838 | 1.39E-11 | -6.758820586 | 0.617108111 |
| CNAG_00559 | BZP3 | bZip transcription factor, putative | 1264.83936 | 917.8657923 | 890.4200663 | 590.1666589 | 565.4594432 | 574.247543 | 7.54E-11 | -0.845257263 | 1.42E-11 | -6.755972999 | 0.556611545 |
| CNAG_04636 | - | metacaspase-1 | 1469.245483 | 1269.398099 | 1348.125899 | 1978.020939 | 2008.306778 | 1983.66263 | 7.76E-11 | 0.530946895 | 1.46E-11 | 6.751686354 | 1.444877211 |
| CNAG_02065 | - | hypothetical protein, variant | 1226.980612 | 1209.819081 | 1222.02933 | 844.0631754 | 761.3053636 | 861.6569947 | 7.76E-11 | -0.583624544 | 1.46E-11 | -6.75154563 | 0.667285221 |
| CNAG_02202 | - | Adenylylsulfate kinase | 1388.189535 | 1390.839827 | 1443.839577 | 989.6131985 | 1017.621324 | 906.1455164 | 7.78E-11 | -0.550994317 | 1.47E-11 | -6.75112683 | 0.682549548 |
| CNAG_12988 | - | unspecified product | 1138.209085 | 1098.8533 | 1129.242268 | 770.1168508 | 785.5718173 | 698.5597999 | 7.95E-11 | -0.594291289 | 1.50E-11 | -6.747910026 | 0.662369759 |
| CNAG_01675 | - | tagatose-bisphosphate aldolase, variant | 787.8037108 | 761.580152 | 745.3273853 | 453.1709189 | 512.9849363 | 480.9409783 | 8.01E-11 | -0.681608447 | 1.51E-11 | -6.746750627 | 0.623469786 |
| CNAG_05226 | - | hypothetical protein | 756.3041523 | 716.3603186 | 791.1105398 | 478.9756862 | 491.4053518 | 406.6543011 | 8.23E-11 | -0.734558131 | 1.56E-11 | -6.742613953 | 0.601002071 |
| CNAG_08003 | - | hypothetical protein | 621.9974508 | 607.5036174 | 668.4372479 | 398.8474046 | 376.6278296 | 295.6989964 | 8.66E-11 | -0.84235784 | 1.64E-11 | -6.735144345 | 0.557731307 |
| CNAG_07304 | - | hypothetical protein | 321.6156361 | 365.4119571 | 353.1504228 | 589.7166328 | 579.0877686 | 601.4062746 | 8.93E-11 | 0.752969303 | 1.69E-11 | 6.730642171 | 1.685257801 |
| CNAG_07253 | HPI2 | cupin domain-containing protein | 1030.734729 | 978.1787843 | 993.8102815 | 677.4861506 | 628.4842999 | 681.834356 | 8.97E-11 | -0.61090791 | 1.70E-11 | -6.729872257 | 0.654784506 |
| CNAG_01255 | - | hypothetical protein | 301.740692 | 342.773521 | 271.1000197 | 600.9313242 | 517.8010614 | 543.700246 | 9.11E-11 | 0.846452979 | 1.73E-11 | 6.727510462 | 1.798074727 |
| CNAG_00419 | - | DNA polymerase beta subunit | 57.80726936 | 57.37820642 | 49.32256835 | 168.4809804 | 144.4666343 | 160.4874491 | 9.14E-11 | 1.518987237 | 1.73E-11 | 6.726975964 | 2.865897947 |
| CNAG_04344 | - | galactonate dehydratase | 474.4206585 | 441.9295194 | 416.0981008 | 703.9928554 | 755.9880755 | 731.3362916 | 9.41E-11 | 0.702295883 | 1.79E-11 | 6.72257052 | 1.627092063 |
| CNAG_06714 | - | endoplasmic reticulum vesicle protein 25 | 1771.180924 | 1751.777411 | 1695.576941 | 1279.184266 | 1247.03458 | 1225.815265 | 9.60E-11 | -0.491201641 | 1.82E-11 | -6.719703867 | 0.711432289 |
| CNAG_12770 | - | unspecified product | 1448.031731 | 780.9722897 | 875.3129863 | 269.763765 | 251.1739032 | 208.1948443 | 9.60E-11 | -2.109931725 | 1.82E-11 | -6.719486513 | 0.231657978 |
| CNAG_02602 | - | flavonol synthase | 93.91419217 | 70.3464876 | 84.47671916 | 207.1013627 | 216.2532063 | 199.4641285 | 9.72E-11 | 1.316207261 | 1.85E-11 | 6.717580112 | 2.49010618 |
| CNAG_00733 | - | hypothetical protein | 1899.660802 | 1804.620173 | 1988.78985 | 1349.957132 | 1338.452501 | 1370.112997 | 9.82E-11 | -0.503785472 | 1.87E-11 | -6.716032558 | 0.705253843 |
| CNAG_03853 | SAR1 | small copII coat GTPase | 4779.439671 | 5049.797124 | 4794.70463 | 3857.739622 | 3763.173271 | 3620.41402 | 9.98E-11 | -0.394503538 | 1.90E-11 | -6.713639345 | 0.760751122 |
| CNAG_06526 | - | hypothetical protein, variant | 2088.793965 | 2029.764464 | 1987.784021 | 1447.235203 | 1414.338093 | 1535.492089 | 1.00E-10 | -0.488786535 | 1.90E-11 | -6.713176495 | 0.712624241 |
| CNAG_00418 | - | S-adenosylmethionine synthase | 5355.005384 | 4893.638555 | 5215.92356 | 3824.174297 | 3782.105007 | 3997.820833 | 1.02E-10 | -0.429934262 | 1.93E-11 | -6.710898921 | 0.742295608 |
| CNAG_05848 | - | splicing factor U2AF 65 kDa subunit | 5234.835075 | 5252.388708 | 5254.33451 | 3918.335594 | 4192.765179 | 4069.993422 | 1.07E-10 | -0.385342066 | 2.04E-11 | -6.703246502 | 0.765597452 |
| CNAG_01464 | FHB1 | Flavohemoglobin | 7169.240134 | 2906.871331 | 4156.676392 | 675.8088866 | 683.2043127 | 838.7916013 | 1.14E-10 | -2.7152159 | 2.17E-11 | -6.694428767 | 0.152278493 |
| CNAG_05753 | ARG5 | N-acetyl-gamma-glutamyl-phosphate reductase, variant | 5742.670179 | 5251.333694 | 5304.469764 | 4177.575836 | 4086.612129 | 4070.015902 | 1.20E-10 | -0.417699819 | 2.28E-11 | -6.686610889 | 0.748617242 |
| CNAG_01392 | - | hypothetical protein | 567.3701269 | 710.8111722 | 659.950874 | 398.0079318 | 380.0526696 | 365.5385458 | 1.21E-10 | -0.774649125 | 2.31E-11 | -6.685098356 | 0.584530769 |

| | | | | | | | | | | | | | |
|---|---|---|---|---|---|---|---|---|---|---|---|---|---|
| CNAG_01574 | - | hypothetical protein | 1539.385705 | 1116.153459 | 1180.464248 | 753.6085895 | 801.895496 | 725.2366999 | 1.21E-10 | -0.767567014 | 2.32E-11 | -6.684202017 | 0.587407254 |
| CNAG_00701 | CAS31 | protein involved in gxm O-acetylation | 4142.902154 | 4043.496093 | 3892.491539 | 3112.888867 | 2902.272581 | 3083.061161 | 1.29E-10 | -0.423994001 | 2.46E-11 | -6.675748429 | 0.745358293 |
| CNAG_03426 | GMT2 | GDP-mannose transporter 2 | 979.5990809 | 1070.8205 | 1282.650428 | 623.5560464 | 734.6598482 | 673.0190102 | 1.29E-10 | -0.729819363 | 2.47E-11 | -6.675173291 | 0.602979407 |
| CNAG_05539 | - | hypothetical protein | 1059.469693 | 1134.642628 | 1005.816114 | 1550.744155 | 1514.860084 | 1601.724387 | 1.31E-10 | 0.529919757 | 2.50E-11 | 6.673256737 | 1.443848886 |
| CNAG_02761 | - | inorganic phosphate transporter PHO88 | 8808.174284 | 8569.83819 | 9396.39787 | 6976.322424 | 6863.211366 | 6808.774553 | 1.32E-10 | -0.390335428 | 2.52E-11 | -6.672004702 | 0.762952197 |
| CNAG_02166 | - | DNA-directed RNA polymerase II subunit RPB1 | 13072.65285 | 12185.67296 | 11716.01086 | 9205.26904 | 9423.289535 | 9678.511585 | 1.34E-10 | -0.400835895 | 2.56E-11 | -6.669639461 | 0.757419309 |
| CNAG_03049 | - | hypothetical protein | 1738.486866 | 1669.920732 | 1797.895084 | 1296.361729 | 1194.391555 | 1120.406896 | 1.34E-10 | -0.54314607 | 2.57E-11 | -6.669345502 | 0.686272728 |
| CNAG_01492 | - | hypothetical protein | 11789.33963 | 10866.92488 | 11208.35153 | 8779.60509 | 8728.93314 | 8818.147812 | 1.34E-10 | -0.378887673 | 2.58E-11 | -6.668773627 | 0.769030289 |
| CNAG_02733 | - | Monosaccharide transporter | 54.47091468 | 97.16084071 | 60.0069064 | 181.188832 | 229.4368652 | 200.4840788 | 1.37E-10 | 1.518288621 | 2.64E-11 | 6.665370918 | 2.864510491 |
| CNAG_01404 | - | hsp71-like protein | 7652.97563 | 6615.358137 | 7234.682062 | 4160.092826 | 5549.609837 | 4655.146083 | 1.38E-10 | -0.598294306 | 2.64E-11 | -6.665099771 | 0.660534442 |
| CNAG_07790 | - | hypothetical protein | 161.7240786 | 167.2555268 | 199.5805474 | 324.9871098 | 364.4766797 | 384.7606406 | 1.39E-10 | 1.009150555 | 2.67E-11 | 6.66347787 | 2.012725678 |
| CNAG_02055 | - | hypothetical protein | 1007.296686 | 908.1582557 | 984.169327 | 669.7871297 | 525.7418335 | 600.8433448 | 1.44E-10 | -0.707865494 | 2.77E-11 | -6.658047588 | 0.612225273 |
| CNAG_04934 | - | hypothetical protein | 520.0672439 | 597.0154808 | 520.594342 | 871.4143695 | 869.7726836 | 849.0575532 | 1.45E-10 | 0.646674956 | 2.80E-11 | 6.656697475 | 1.565555828 |
| CNAG_03060 | - | multiple drug resistance protein | 354.3281595 | 403.1294863 | 418.1787263 | 656.6416596 | 655.8806676 | 652.5146434 | 1.52E-10 | 0.725507559 | 2.92E-11 | 6.650398827 | 1.65348225 |
| CNAG_04877 | - | hypothetical protein | 7.631347585 | 9.978796691 | 6.653104827 | 56.1741769 | 62.86195045 | 63.77309095 | 1.56E-10 | 3.025811655 | 3.01E-11 | 6.646249171 | 8.144418267 |
| CNAG_02306 | - | molybdopterin binding domain-containing protein | 1021.510871 | 1098.784408 | 1031.144972 | 658.6890334 | 758.8721206 | 666.3474173 | 1.56E-10 | -0.61248238 | 3.01E-11 | -6.646300887 | 0.654070304 |
| CNAG_04961 | URA3 | orotidine 5'-phosphate decarboxylase | 1215.362713 | 1221.680111 | 1265.74103 | 805.0415662 | 853.6902028 | 886.0735107 | 1.69E-10 | -0.55606077 | 3.26E-11 | -6.634376329 | 0.680156775 |
| CNAG_03814 | - | small nuclear ribonucleoprotein Sm D2, variant | 3671.082493 | 3346.437645 | 3495.742956 | 2676.25514 | 2589.194864 | 2557.992455 | 1.70E-10 | -0.442089955 | 3.29E-11 | -6.632892985 | 0.736067534 |
| CNAG_00254 | - | NADH dehydrogenase | 765.3704237 | 607.5910146 | 687.6939848 | 424.4767406 | 406.7991398 | 377.7679134 | 1.74E-10 | -0.786789761 | 3.37E-11 | -6.629424421 | 0.579632438 |
| CNAG_00365 | - | lysophospholipid acyltransferase | 2609.695649 | 2585.739796 | 2714.001022 | 1827.766862 | 1902.580194 | 2043.926344 | 1.76E-10 | -0.46951193 | 3.41E-11 | -6.627508488 | 0.722208883 |
| CNAG_01017 | - | E3 ubiquitin-protein ligase SHPRH, variant | 2740.970371 | 2715.25366 | 2733.422892 | 3487.681641 | 3852.459057 | 3816.537011 | 1.78E-10 | 0.430767012 | 3.45E-11 | 6.626093867 | 1.347950027 |
| CNAG_03043 | - | hypothetical protein, variant 2 | 31.03251996 | 41.1977501 | 19.50409209 | 112.975566 | 138.1182845 | 109.5092365 | 1.78E-10 | 1.96583536 | 3.45E-11 | 6.625911822 | 3.906388295 |
| CNAG_01936 | - | Sugar transporter | 46.09431861 | 35.86515639 | 33.33631174 | 121.4844531 | 123.0311193 | 139.2522113 | 1.79E-10 | 1.732601044 | 3.46E-11 | 6.625547 | 3.32326432 |
| CNAG_04099 | - | 6-phosphogluconate dehydrogenase, decarboxylating | 6885.861388 | 6293.195707 | 6181.102622 | 4776.805473 | 4560.100313 | 5034.701494 | 1.81E-10 | -0.445395447 | 3.51E-11 | -6.623519483 | 0.734382992 |
| CNAG_00040 | ERG11 | cytochrome P450, family 51 (sterol 14- demethylase) | 976.7734243 | 847.7934119 | 834.9201121 | 579.4302653 | 559.522246 | 451.0921007 | 1.81E-10 | -0.757663131 | 3.51E-11 | -6.623576229 | 0.591453588 |
| CNAG_02813 | - | large subunit ribosomal protein L30 | 741.1989237 | 721.7550515 | 819.8863773 | 508.6212586 | 462.8694223 | 444.353386 | 1.81E-10 | -0.705376395 | 3.51E-11 | -6.623413737 | 0.613282464 |
| CNAG_04056 | - | hypothetical protein | 1197.795273 | 1190.421657 | 1172.98946 | 773.1309523 | 814.8682129 | 854.9853861 | 1.91E-10 | -0.559208967 | 3.72E-11 | -6.614865472 | 0.67867418 |
| CNAG_06517 | - | cytoplasmic protein | 1397.560071 | 1415.623899 | 1483.293343 | 984.9003977 | 1060.771919 | 946.0947885 | 1.99E-10 | -0.537922124 | 3.88E-11 | -6.608709366 | 0.688762201 |
| CNAG_01003 | NCP1 | NADPH-ferrihemoprotein reductase | 4123.107457 | 4043.517378 | 4081.235688 | 3073.130213 | 3217.19483 | 3144.123628 | 2.03E-10 | -0.391868213 | 3.96E-11 | -6.605656153 | 0.762142032 |
| CNAG_07925 | - | nucleoprotein TPR | 10439.37212 | 9384.380459 | 10039.49245 | 7085.143365 | 7274.612788 | 7913.3332 | 2.06E-10 | -0.438753557 | 4.00E-11 | -6.60406041 | 0.737771745 |
| CNAG_07558 | - | hypothetical protein | 12807.64669 | 11888.3362 | 12674.70405 | 9311.110261 | 9650.153604 | 9936.058343 | 2.07E-10 | -0.38653961 | 4.03E-11 | -6.602812639 | 0.764962213 |
| CNAG_04807 | FZC8 | hypothetical protein | 321.4446557 | 231.9340342 | 246.5777372 | 614.4900418 | 494.5576563 | 486.0264078 | 2.14E-10 | 0.981851318 | 4.18E-11 | 6.597700786 | 1.974998172 |
| CNAG_04813 | - | V-type ATPase, C subunit, variant | 1020.875278 | 978.4535146 | 937.557081 | 1474.561698 | 1373.375628 | 1455.201356 | 2.21E-10 | 0.536407542 | 4.31E-11 | 6.59316869 | 1.450356482 |
| CNAG_03708 | - | signal recognition particle subunit SRP72 | 2687.922302 | 2551.294151 | 2845.154618 | 1939.90095 | 2023.380519 | 1956.283621 | 2.21E-10 | -0.465414336 | 4.32E-11 | -6.592758452 | 0.724263042 |
| CNAG_02106 | - | mitochondrial intermembrane space import and assembly protein 40 | 1914.911796 | 2000.658577 | 2042.126816 | 1295.341121 | 1498.964741 | 1433.390866 | 2.23E-10 | -0.510270528 | 4.36E-11 | -6.59141246 | 0.702090773 |
| CNAG_00662 | - | carboxymethylenebutenolidase | 1588.839336 | 1376.878399 | 1451.333318 | 1042.557252 | 972.6804223 | 996.0267698 | 2.28E-10 | -0.568434664 | 4.45E-11 | -6.588218547 | 0.674348065 |
| CNAG_00192 | - | hypothetical protein | 343.6982569 | 344.9517035 | 299.8774053 | 610.9430756 | 528.102227 | 583.6355645 | 2.31E-10 | 0.78866929 | 4.51E-11 | 6.586302802 | 1.727480337 |
| CNAG_03284 | - | rRNA biogenesis protein RRP36 | 1726.740901 | 1603.121179 | 1681.674578 | 1056.377191 | 1180.619621 | 1227.985243 | 2.31E-10 | -0.548358062 | 4.51E-11 | -6.586193578 | 0.683797919 |
| CNAG_06630 | - | import inner membrane translocase subunit tim44 | 9202.737488 | 8124.88296 | 9195.902355 | 6332.548139 | 6772.655008 | 6654.460907 | 2.34E-10 | -0.440666562 | 4.58E-11 | -6.584096191 | 0.736794113 |
| CNAG_01854 | - | heparinase II/III family protein | 9246.483134 | 9581.685446 | 8313.146085 | 11732.16895 | 12299.47149 | 12411.41711 | 2.40E-10 | 0.410259495 | 4.70E-11 | 6.580098671 | 1.328924824 |
| CNAG_08018 | - | hypothetical protein | 1069.909424 | 982.7820526 | 991.9377707 | 1418.317832 | 1501.118796 | 1557.294237 | 2.47E-10 | 0.541673128 | 4.83E-11 | 6.5759815 | 1.455659702 |
| CNAG_05087 | - | hypothetical protein | 40.45182715 | 61.67386091 | 59.95163734 | 136.6855355 | 188.090457 | 162.7130647 | 2.49E-10 | 1.588414815 | 4.87E-11 | 6.574722026 | 3.007187482 |
| CNAG_06915 | - | hypothetical protein | 14.80844388 | 47.62719169 | 21.63314942 | 128.1146122 | 113.5347425 | 133.644473 | 2.61E-10 | 2.178300591 | 5.12E-11 | 6.567428098 | 4.526200803 |
| CNAG_03938 | CPR2 | Nonmating-type specific pheromone G-protein coupled receptor | 11.16249675 | 15.36474424 | 9.864976823 | 74.41078221 | 62.57504797 | 77.05020909 | 2.65E-10 | 2.591450649 | 5.19E-11 | 6.565273178 | 6.027044217 |
| CNAG_04383 | - | acetyltransferase | 824.1738217 | 858.5554848 | 886.0603476 | 588.0160562 | 558.6583738 | 533.137101 | 2.65E-10 | -0.628212201 | 5.21E-11 | -6.56490923 | 0.646977658 |
| CNAG_07529 | - | threonine ammonia-lyase, biosynthetic | 334.3368112 | 324.4884727 | 275.3626046 | 547.7058659 | 517.7916267 | 593.5343851 | 2.73E-10 | 0.813635919 | 5.36E-11 | 6.560542092 | 1.757635506 |
| CNAG_01879 | - | cathepsin A (carboxypeptidase C) | 1063.020753 | 1170.2056 | 1131.622105 | 1687.059633 | 1618.405499 | 1544.102429 | 2.75E-10 | 0.512597341 | 5.42E-11 | 6.559007997 | 1.426616279 |
| CNAG_02852 | - | 4-aminobutyrate transaminase | 1923.593018 | 1642.869171 | 1300.00358 | 1026.9066 | 986.4605105 | 819.6685573 | 2.75E-10 | -0.796997957 | 5.42E-11 | -6.559019871 | 0.575545561 |
| CNAG_01738 | - | hypothetical protein | 348.315727 | 313.7356355 | 292.4089426 | 585.2292179 | 528.1573169 | 553.6892254 | 2.78E-10 | 0.790432395 | 5.47E-11 | 6.557622199 | 1.729592767 |
| CNAG_00036 | SEC14-2 | SEC14 cytosolic factor | 203.638474 | 178.0378067 | 208.1284575 | 407.9201401 | 392.9335833 | 346.1743244 | 2.80E-10 | 0.943144078 | 5.51E-11 | 6.556490771 | 1.922713863 |
| CNAG_06641 | - | DNA-directed RNA polymerase III subunit RPC6 | 2566.567973 | 2668.695616 | 2804.642572 | 1958.594076 | 1993.172802 | 2030.632873 | 2.84E-10 | -0.441601657 | 5.60E-11 | -6.55394995 | 0.736316708 |
| CNAG_06710 | - | hypothetical protein | 1955.692587 | 1820.790521 | 2010.126058 | 1411.485843 | 1369.508598 | 1372.348208 | 2.85E-10 | -0.494120382 | 5.63E-11 | -6.553372395 | 0.709994433 |
| CNAG_00065 | - | ran-binding protein 1 | 5037.530581 | 5082.17074 | 5589.282653 | 4036.715266 | 3810.559929 | 3992.286406 | 3.00E-10 | -0.420570504 | 5.93E-11 | -6.545529586 | 0.747129119 |
| CNAG_06348 | PDR5-3 | ABC transporter PMR5 | 1405.515321 | 1247.570235 | 1276.444057 | 854.0783672 | 908.0373326 | 914.9580942 | 3.01E-10 | -0.569438694 | 5.95E-11 | -6.54487168 | 0.673878922 |
| CNAG_03349 | - | pre-mRNA-processing factor 17 | 5568.830471 | 5838.48017 | 5639.324643 | 4018.657328 | 4373.006449 | 4549.545545 | 3.17E-10 | -0.412509365 | 6.26E-11 | -6.537384434 | 0.751315429 |
| CNAG_00533 | TIM50 | mitochondrial import inner membrane translocase subunit tim50 | 4591.354796 | 4388.309031 | 4838.358132 | 3060.511335 | 3508.728598 | 3502.69566 | 3.22E-10 | -0.471994977 | 6.37E-11 | -6.534825018 | 0.720966946 |
| CNAG_04201 | - | hypothetical protein | 36.83173651 | 44.42529075 | 23.76392677 | 120.0284064 | 134.5116976 | 115.0531181 | 3.23E-10 | 1.826588685 | 6.39E-11 | 6.534284077 | 3.546973828 |
| CNAG_07076 | - | hypothetical protein | 19.3723723 | 23.99891687 | 26.89952674 | 92.84240925 | 92.0348539 | 97.13736855 | 3.30E-10 | 2.032092444 | 6.53E-11 | 6.53106802 | 4.08997619 |
| CNAG_04649 | - | peptide alpha-N-acetyltransferase | 196.6033807 | 170.4876191 | 185.7513097 | 389.2895692 | 341.9025132 | 347.1970272 | 3.31E-10 | 0.951256031 | 6.57E-11 | 6.530179078 | 1.933555307 |
| CNAG_03623 | - | malonic semialdehyde reductase | 1594.941855 | 1670.958859 | 1634.771492 | 1200.569643 | 1180.622208 | 1173.624524 | 3.33E-10 | -0.478207003 | 6.60E-11 | -6.529380303 | 0.717869246 |
| CNAG_01504 | - | hypothetical protein | 999.4268191 | 1131.109706 | 1105.76484 | 738.2027654 | 682.8283731 | 745.1061562 | 3.35E-10 | -0.594056473 | 6.64E-11 | -6.52861575 | 0.662477576 |
| CNAG_07902 | - | AAT family amino acid transporter | 1352.059114 | 1716.010876 | 1318.077011 | 982.5812595 | 945.0806527 | 888.4039911 | 3.40E-10 | -0.653779772 | 6.75E-11 | -6.52620366 | 0.635612864 |
| CNAG_05632 | - | hypothetical protein | 108.1024448 | 163.9242603 | 114.3629198 | 285.6000834 | 275.553183 | 311.4509899 | 3.40E-10 | 1.16436895 | 6.76E-11 | 6.525918695 | 2.241351551 |
| CNAG_05086 | URM1 | ubiquitin-related modifier | 866.115417 | 785.373003 | 975.5560199 | 540.4045115 | 577.6656729 | 512.0735987 | 3.49E-10 | -0.70487911 | 6.93E-11 | -6.522077413 | 0.613493894 |
| CNAG_07492 | - | hypothetical protein | 25.16084343 | 27.22017321 | 19.47672238 | 83.9080638 | 103.4056036 | 107.0278995 | 3.51E-10 | 2.03019966 | 6.98E-11 | 6.521094667 | 4.084613749 |
| CNAG_06329 | - | high-affinity nicotinic acid transporter | 65.93542262 | 46.66110646 | 63.14707282 | 150.8772325 | 167.94596 | 175.9567759 | 3.54E-10 | 1.483295295 | 7.05E-11 | 6.519647132 | 2.795866151 |
| CNAG_03182 | - | U4/U6.U5 tri-snRNP-associated protein 3 | 502.8661151 | 494.3573912 | 498.9343524 | 299.134586 | 270.4468022 | 307.7450203 | 3.60E-10 | -0.785325307 | 7.18E-11 | -6.516930619 | 0.580221111 |
| CNAG_12531 | - | unspecified product | 2.952507617 | 12.12224 | 7.718521925 | 63.48617342 | 62.83832899 | 60.53466929 | 3.66E-10 | 3.126499161 | 7.30E-11 | 6.514398958 | 8.73313209 |
| CNAG_06468 | - | nucleolar GTP-binding protein | 2638.673695 | 2187.155287 | 2623.303702 | 1810.784854 | 1585.930138 | 1645.486448 | 3.66E-10 | -0.579523543 | 7.31E-11 | -6.514236037 | 0.669184742 |
| CNAG_07903 | ROM20 | Predicted Rho guanine nucleotide exchange factor (Rho GEF) | 1024.334279 | 924.5968755 | 860.7970001 | 1376.38872 | 1398.468456 | 1444.06917 | 3.70E-10 | 0.571521158 | 7.39E-11 | 6.51254705 | 1.486089657 |
| CNAG_06119 | - | spermine transporter | 25.28367746 | 36.93716932 | 49.26081215 | 138.239647 | 127.4109727 | 120.6010042 | 3.76E-10 | 1.792553231 | 7.51E-11 | 6.510199502 | 3.464274454 |
| CNAG_07358 | - | hypothetical protein | 51.01973644 | 101.4870035 | 78.10340519 | 184.1376673 | 214.46925 | 262.3138286 | 3.82E-10 | 1.508034887 | 7.64E-11 | 6.507594364 | 2.844223598 |
| CNAG_06939 | - | hypothetical protein | 6.478971358 | 8.912434271 | 11.96783234 | 68.32598482 | 64.57919503 | 54.04190922 | 3.87E-10 | 2.880140269 | 7.74E-11 | 6.505509679 | 7.362216979 |
| CNAG_12438 | - | unspecified product | 20270.52665 | 16972.73963 | 17822.48624 | 12754.19613 | 12501.67646 | 14277.89189 | 3.92E-10 | -0.493965834 | 7.83E-11 | -6.503786013 | 0.710070495 |
| CNAG_06689 | - | cytoplasmic protein | 2522.200543 | 2668.683116 | 2700.150136 | 1974.976571 | 1995.776992 | 1970.69867 | 3.92E-10 | -0.424879252 | 7.85E-11 | -6.503498342 | 0.744901074 |
| CNAG_05797 | XRN2 | 5'-3' exoribonuclease 2 | 6804.317012 | 6889.011178 | 6976.649418 | 5065.50502 | 5527.149397 | 5472.106836 | 3.94E-10 | -0.378905853 | 7.88E-11 | -6.502847577 | 0.769020598 |
| CNAG_03389 | - | alpha-ketoglutarate catabolism dioxygenase | 3331.317843 | 3193.419678 | 3227.002575 | 2389.77199 | 2351.102592 | 2561.256456 | 3.95E-10 | -0.432768355 | 7.91E-11 | -6.502337528 | 0.740838842 |
| CNAG_01894 | - | aspartate-tRNA ligase | 2615.419549 | 2345.510657 | 2574.294841 | 1722.689295 | 1828.415151 | 1879.649903 | 4.01E-10 | -0.488491362 | 8.05E-11 | -6.499671396 | 0.712770058 |
| CNAG_04564 | - | phytanoyl-CoA dioxygenase | 276.041055 | 276.0313914 | 300.8833928 | 510.4372379 | 471.2248342 | 507.0282487 | 4.02E-10 | 0.789249431 | 8.07E-11 | 6.499238814 | 1.728175029 |
| CNAG_03244 | - | hypothetical protein | 1809.630876 | 1636.536535 | 1691.298166 | 1212.971189 | 1215.1128 | 1234.674339 | 4.09E-10 | -0.504042621 | 8.21E-11 | -6.496756822 | 0.705128149 |
| CNAG_02037 | - | large subunit ribosomal protein L22 | 722.6562195 | 787.4400703 | 874.25871 | 518.6946712 | 506.9003773 | 465.4470813 | 4.18E-10 | -0.693596351 | 8.40E-11 | -6.493271978 | 0.618310602 |
| CNAG_00793 | - | DNA-directed RNA polymerase I, II, and III subunit RPABC1 | 2136.82329 | 2318.484222 | 2307.679936 | 1675.35599 | 1479.839893 | 1680.931229 | 4.18E-10 | -0.498758854 | 8.41E-11 | -6.493072902 | 0.707715365 |
| CNAG_03808 | - | aromatic amino acid aminotransferase I | 1300.378255 | 1228.101792 | 1079.204848 | 828.4996048 | 790.6864063 | 778.4606793 | 4.35E-10 | -0.605484559 | 8.75E-11 | -6.487099663 | 0.657250592 |
| CNAG_03840 | - | pre-mRNA-splicing factor cwc26 | 1529.458485 | 1498.5905 | 1480.145742 | 1078.343021 | 1094.378079 | 1071.493014 | 4.43E-10 | -0.49016755 | 8.92E-11 | -6.484146386 | 0.71194241 |
| CNAG_01052 | - | hypothetical protein | 1488.236465 | 1326.605706 | 900.4235887 | 3299.520834 | 3247.342324 | 3019.231256 | 4.45E-10 | 1.349374909 | 8.95E-11 | 6.483673993 | 2.54801701 |
| CNAG_02898 | - | RNA-binding protein 8A | 3031.241026 | 2926.225773 | 3062.749337 | 2241.140899 | 2186.358038 | 2343.675961 | 4.51E-10 | -0.429154537 | 9.09E-11 | -6.481316366 | 0.742696901 |
| CNAG_12795 | - | unspecified product | 178.0082878 | 199.5116886 | 174.0418306 | 386.9429499 | 364.448211 | 323.9733318 | 4.67E-10 | 0.946651115 | 9.41E-11 | 6.476085088 | 1.92739346 |

| | | | | | | | | | | | | | |
|---|---|---|---|---|---|---|---|---|---|---|---|---|---|
| CNAG_02545 | IPP1 | Inorganic pyrophosphatase | 4278.477363 | 4340.885512 | 4562.171115 | 3414.868352 | 3446.637753 | 3150.840999 | 4.74E-10 | -0.412283932 | 9.56E-11 | -6.473817015 | 0.751432838 |
| CNAG_06021 | - | rab GTPase activator | 843.4714095 | 880.4020329 | 904.458852 | 1236.235333 | 1314.838899 | 1279.815924 | 5.02E-10 | 0.528370304 | 1.01E-10 | 6.464839239 | 1.442299026 |
| CNAG_06346 | BLP1 | Barwin-like protein 1 | 57.79122499 | 47.71710958 | 58.87793562 | 146.3197303 | 145.3803564 | 167.0470559 | 5.29E-10 | 1.483680728 | 1.07E-10 | 6.456899758 | 2.7966132 |
| CNAG_06917 | - | thiol-specific antioxidant protein 3 | 376.3832905 | 375.117816 | 344.646618 | 572.3958038 | 659.4663361 | 605.8969946 | 5.31E-10 | 0.730331686 | 1.07E-10 | 6.456357162 | 1.659020469 |
| CNAG_07766 | - | DNA polymerase lambda subunit | 551.5256562 | 525.9718162 | 523.7890014 | 851.9071935 | 782.4880773 | 972.0765296 | 5.32E-10 | 0.689012032 | 1.08E-10 | 6.455857717 | 1.612179109 |
| CNAG_03769 | HXK2 | Hexokinase | 12925.42786 | 11361.45324 | 11117.73094 | 8816.139211 | 9015.306739 | 8747.13103 | 5.48E-10 | -0.429462338 | 1.11E-10 | -6.451247069 | 0.742538462 |
| CNAG_04756 | - | hypothetical protein | 85.74705166 | 75.69228192 | 70.63578107 | 200.2195898 | 171.1080078 | 204.8622848 | 5.98E-10 | 1.306674874 | 1.21E-10 | 6.437952086 | 2.47370742 |
| CNAG_05342 | - | hypothetical protein | 615.956755 | 532.115654 | 540.5625338 | 331.0080418 | 354.2373683 | 315.5865905 | 6.10E-10 | -0.77036877 | 1.24E-10 | -6.434900392 | 0.586267598 |
| CNAG_12906 | - | unspecified product | 24.05033684 | 32.59963839 | 30.11100302 | 101.2820319 | 96.2002432 | 121.4010249 | 6.10E-10 | 1.872366233 | 1.24E-10 | 6.434904281 | 3.661325993 |
| CNAG_06438 | - | hypothetical protein | 6.42053282 | 5.662897869 | 4.504488824 | 50.0454972 | 52.45523569 | 53.81299699 | 6.27E-10 | 3.353887387 | 1.27E-10 | 6.430503603 | 10.22399681 |
| CNAG_00589 | - | 5-formyltetrahydrofolate cyclo-ligase | 1494.197392 | 1247.598268 | 1330.82549 | 922.6389429 | 917.5032186 | 909.4384969 | 6.29E-10 | -0.583001397 | 1.28E-10 | -6.42981591 | 0.667573505 |
| CNAG_01644 | MLN4 | hypothetical protein | 2219.658154 | 2140.780417 | 2297.018413 | 1663.670853 | 1649.87569 | 1616.575164 | 6.59E-10 | -0.449135936 | 1.34E-10 | -6.422840877 | 0.732481417 |
| CNAG_06548 | - | endoplasmic reticulum protein | 4495.578284 | 4355.964775 | 4266.814603 | 3198.390567 | 3356.903311 | 3477.139464 | 6.73E-10 | -0.402184127 | 1.37E-10 | -6.419558931 | 0.756711814 |
| CNAG_03681 | - | peptidylprolyl isomerase domain and WD-repeat protein 1 | 2240.4979 | 1918.857636 | 2130.650767 | 1527.454345 | 1476.464137 | 1441.201385 | 7.01E-10 | -0.516714912 | 1.43E-10 | -6.413061215 | 0.698961593 |
| CNAG_00006 | - | prohibitin-2 | 4455.720829 | 3765.58744 | 4132.403284 | 2930.648397 | 2989.414201 | 3070.860052 | 7.03E-10 | -0.47451961 | 1.43E-10 | -6.412649732 | 0.719706399 |
| CNAG_07740 | HAD2 | hypothetical protein | 7127.818936 | 7426.640524 | 7315.794961 | 5959.938551 | 5669.426452 | 5691.942149 | 7.03E-10 | -0.35150877 | 1.43E-10 | -6.412541784 | 0.783764009 |
| CNAG_05548 | - | hypothetical protein | 130.1674737 | 140.2911501 | 139.8994355 | 265.9317821 | 314.6216172 | 270.5916739 | 7.07E-10 | 1.038415904 | 1.44E-10 | 6.411602199 | 2.05397113 |
| CNAG_04236 | - | pyridoxal phosphate phosphatase phospho2 | 2102.868999 | 2031.951926 | 2319.353074 | 1367.701604 | 1646.492838 | 1432.337187 | 7.12E-10 | -0.553362122 | 1.45E-10 | -6.410525183 | 0.681430241 |
| CNAG_07687 | - | hypothetical protein | 29.94953492 | 29.44349772 | 57.77519198 | 118.0780887 | 132.5125605 | 174.4986495 | 7.34E-10 | 1.86688014 | 1.50E-10 | 6.405795176 | 3.647429618 |
| CNAG_05099 | - | hypothetical protein | 2.918470113 | 0.227157895 | 9.819147809 | 48.34569775 | 52.41820136 | 63.58073789 | 7.47E-10 | 3.87257332 | 1.52E-10 | 6.40286388 | 14.64740634 |
| CNAG_02486 | - | translation initiation factor 2 subunit 2 | 2102.906551 | 2150.435342 | 2105.092922 | 1624.010651 | 1561.880778 | 1565.499109 | 7.75E-10 | -0.435537469 | 1.58E-10 | -6.397123288 | 0.739418236 |
| CNAG_02667 | - | hypothetical protein | 475.5827465 | 438.6997657 | 437.4057317 | 755.5673195 | 717.0825909 | 691.4439777 | 7.77E-10 | 0.664570842 | 1.59E-10 | 6.39674859 | 1.585096685 |
| CNAG_00550 | - | DNA mismatch repair protein Msh5 | 321.5343454 | 314.806417 | 329.6830517 | 538.316142 | 519.5217112 | 570.2830816 | 7.83E-10 | 0.739888893 | 1.60E-10 | 6.395400025 | 1.670047218 |
| CNAG_01687 | - | Atypical/RIO/RIO2 protein kinase | 1249.024768 | 1105.346973 | 1126.075628 | 787.9927926 | 755.3048225 | 813.9273521 | 8.32E-10 | -0.578975357 | 1.70E-10 | -6.386113202 | 0.669439063 |
| CNAG_04667 | - | hypothetical protein | 618.1636934 | 696.1529712 | 676.2585969 | 1088.696628 | 1004.278092 | 957.925284 | 8.45E-10 | 0.600727632 | 1.73E-10 | 6.38369874 | 1.51648122 |
| CNAG_04213 | - | signal transducer | 688.1213003 | 718.7677585 | 617.6498237 | 1054.320148 | 1039.680705 | 987.8763629 | 8.55E-10 | 0.59145324 | 1.75E-10 | 6.381814982 | 1.50676376 |
| CNAG_06779 | - | small nuclear ribonucleoprotein D1 | 922.4034948 | 1106.295962 | 1044.979267 | 680.5657464 | 666.4679105 | 697.3789484 | 8.82E-10 | -0.602962403 | 1.81E-10 | -6.37692111 | 0.658400619 |
| CNAG_05562 | PBX2 | parallel beta-helix repeat protein | 2184.696617 | 2261.42237 | 2326.885509 | 1659.738685 | 1737.028626 | 1660.976233 | 8.88E-10 | -0.436597422 | 1.82E-10 | -6.375790442 | 0.738875183 |
| CNAG_06822 | - | hypothetical protein | 27.71096868 | 69.21920509 | 73.79460942 | 180.7587767 | 193.0882653 | 181.5807793 | 8.96E-10 | 1.699641206 | 1.84E-10 | 6.3743482 | 3.248201667 |
| CNAG_05540 | URE1 | urease | 9708.455851 | 9368.198259 | 9283.417649 | 7242.44005 | 7668.891984 | 7543.670637 | 8.96E-10 | -0.352330167 | 1.84E-10 | -6.374186423 | 0.783317901 |
| CNAG_00876 | FRE7 | ferric-chelate reductase | 1165.614346 | 871.5192416 | 848.8113439 | 436.5679457 | 619.9520855 | 533.1734415 | 9.07E-10 | -0.877603647 | 1.86E-10 | -6.372298259 | 0.544270728 |
| CNAG_01069 | - | hypothetical protein | 417.7656862 | 437.2608221 | 494.5718102 | 241.4134613 | 249.7764233 | 272.2231158 | 9.14E-10 | -0.838110451 | 1.88E-10 | -6.371005994 | 0.559375724 |
| CNAG_01822 | - | hypothetical protein | 690.8342087 | 607.5752145 | 683.4177899 | 412.0070701 | 391.2552769 | 425.4086581 | 9.15E-10 | -0.704896624 | 1.88E-10 | -6.370638655 | 0.613486446 |
| CNAG_01558 | - | chlorophyll synthesis pathway protein BchC | 18189.01213 | 14720.7295 | 12694.98849 | 7927.018375 | 7427.924023 | 7304.121437 | 9.40E-10 | -1.025134739 | 1.93E-10 | -6.366513896 | 0.491364407 |
| CNAG_02879 | - | chloride channel protein, nucleotide-sensitive, 1A | 1633.538983 | 1807.771008 | 1735.015448 | 1271.395662 | 1259.123752 | 1238.016409 | 9.81E-10 | -0.472605713 | 2.02E-10 | -6.359864872 | 0.720661804 |
| CNAG_01600 | - | ribosome assembly protein RRB1 | 1680.262319 | 1850.879735 | 1846.97097 | 1289.259094 | 1350.593337 | 1249.139914 | 1.02E-09 | -0.482507007 | 2.10E-10 | -6.354047479 | 0.715732796 |
| CNAG_04069 | PRP31 | U4/U6 small nuclear ribonucleoprotein PRP31 | 3507.727263 | 3405.697414 | 3763.377926 | 2752.544897 | 2698.765163 | 2554.678856 | 1.06E-09 | -0.430991284 | 2.18E-10 | -6.348323828 | 0.741751948 |
| CNAG_04404 | - | hypothetical protein, variant | 73.03851483 | 68.19039551 | 100.4094184 | 201.6643495 | 191.0615428 | 210.4383356 | 1.07E-09 | 1.305110486 | 2.21E-10 | 6.345767608 | 2.471026505 |
| CNAG_06868 | - | Phosphopyruvate hydratase | 837.4238848 | 747.4094154 | 503.3965147 | 393.2902686 | 312.6116355 | 352.2531155 | 1.08E-09 | -0.996534404 | 2.23E-10 | -6.344698409 | 0.501202528 |
| CNAG_00897 | SKN1 | putative glucosidase | 3765.043983 | 3763.569115 | 3498.089311 | 4930.789913 | 5080.73372 | 4665.917341 | 1.08E-09 | 0.397169901 | 2.23E-10 | 6.344325554 | 1.316922002 |
| CNAG_06826 | - | hypothetical protein | 475.0483965 | 514.8331984 | 591.5999207 | 302.1970059 | 273.0065276 | 327.7155418 | 1.10E-09 | -0.823659322 | 2.27E-10 | -6.342188273 | 0.565007012 |
| CNAG_06385 | - | GAF domain-containing protein, variant | 1145.391248 | 1209.82703 | 1325.412622 | 846.3884394 | 874.4286625 | 776.2636927 | 1.11E-09 | -0.575008893 | 2.29E-10 | -6.340512713 | 0.671282113 |
| CNAG_07491 | - | glutaredoxin | 655.5873614 | 766.195636 | 801.0052176 | 1077.458953 | 1166.50594 | 1172.064384 | 1.11E-09 | 0.605220556 | 2.30E-10 | 6.339966636 | 1.521211295 |
| CNAG_03246 | - | 18S rRNA biogenesis protein RCL1 | 2150.46047 | 1755.084088 | 1905.643689 | 1239.300013 | 1395.42357 | 1327.954979 | 1.14E-09 | -0.5689154 | 2.37E-10 | -6.335463613 | 0.674123395 |
| CNAG_04656 | - | arginine-tRNA-protein transferase, variant | 1466.90863 | 1317.861182 | 1376.908456 | 1932.045699 | 1927.161312 | 2010.284782 | 1.15E-09 | 0.481262472 | 2.38E-10 | 6.334789607 | 1.395964711 |
| CNAG_00142 | - | hypothetical protein | 1281.897295 | 1290.621577 | 1329.720212 | 766.7782493 | 859.6949761 | 973.7275388 | 1.16E-09 | -0.60121798 | 2.39E-10 | -6.333635504 | 0.6591972 |
| CNAG_01208 | CCH1 | calcium-channel protein CCH1 | 1946.881184 | 1940.567844 | 1829.093946 | 2646.551254 | 2484.300274 | 2719.676524 | 1.24E-09 | 0.442707331 | 2.57E-10 | 6.322569957 | 1.359152493 |
| CNAG_04285 | - | hypothetical protein | 49.59518047 | 47.67235042 | 31.22093169 | 144.427164 | 136.0560545 | 119.5789886 | 1.26E-09 | 1.636416909 | 2.62E-10 | 6.319624706 | 3.108927361 |
| CNAG_02848 | - | hypothetical protein | 204.9736806 | 269.5305742 | 264.6362364 | 440.2443035 | 514.7006154 | 428.3069928 | 1.30E-09 | 0.892355298 | 2.71E-10 | 6.314714725 | 1.856204032 |
| CNAG_07337 | - | hypothetical protein | 786.1726709 | 769.4264718 | 737.059687 | 1110.22188 | 1119.854148 | 1125.496452 | 1.32E-09 | 0.534018991 | 2.75E-10 | 6.31229215 | 1.447957233 |
| CNAG_01155 | GUT1 | glycerol kinase | 7543.021554 | 6333.027605 | 5612.73059 | 4669.414216 | 4132.20313 | 4446.399364 | 1.35E-09 | -0.572643643 | 2.80E-10 | -6.309206229 | 0.672383561 |
| CNAG_07471 | - | hypothetical protein | 19.43790168 | 42.2507224 | 29.06002817 | 114.7396194 | 113.6617288 | 108.3399993 | 1.36E-09 | 1.883078459 | 2.83E-10 | 6.307834577 | 3.688613067 |
| CNAG_00840 | - | hypothetical protein | 19.33349649 | 12.18648296 | 26.87785092 | 84.26062366 | 92.20866105 | 83.89978953 | 1.49E-09 | 2.167247838 | 3.10E-10 | 6.293912429 | 4.491657236 |
| CNAG_01099 | - | cleavage and polyadenylation specificity factor subunit 2 | 2579.298853 | 2495.241123 | 2507.148942 | 1825.478437 | 1924.188594 | 1956.237029 | 1.50E-09 | -0.425536531 | 3.12E-10 | -6.292586899 | 0.744561781 |
| CNAG_06029 | - | peptidyl-tRNA hydrolase ICT1 | 536.7371318 | 489.0441089 | 585.2341145 | 299.820247 | 349.9684759 | 281.2070803 | 1.51E-09 | -0.809353956 | 3.15E-10 | -6.290982599 | 0.570637334 |
| CNAG_03265 | - | hypothetical protein | 937.985041 | 901.9627264 | 926.8649558 | 1286.811036 | 1313.034881 | 1391.888383 | 1.53E-09 | 0.514297075 | 3.18E-10 | 6.289496814 | 1.42829806 |
| CNAG_00642 | - | hypothetical protein | 214.3734869 | 313.6920482 | 303.0245881 | 572.2024523 | 516.192199 | 482.7091249 | 1.57E-09 | 0.905518339 | 3.28E-10 | 6.284826574 | 1.873217394 |
| CNAG_00663 | - | hypothetical protein | 1172.626468 | 1058.193838 | 1030.339636 | 1524.980784 | 1597.732981 | 1588.432301 | 1.58E-09 | 0.514952182 | 3.29E-10 | 6.284312026 | 1.428946777 |
| CNAG_04730 | GPR4 | G-protein coupled receptor | 1322.074023 | 1137.940142 | 1118.864428 | 1742.986366 | 1701.174523 | 1807.096418 | 1.63E-09 | 0.537873627 | 3.40E-10 | 6.279307174 | 1.451831101 |
| CNAG_03436 | - | alanine transaminase, variant | 1776.088516 | 2273.11808 | 1976.033806 | 1337.417483 | 1429.044116 | 1396.777681 | 1.64E-09 | -0.547848681 | 3.43E-10 | -6.277827521 | 0.684039394 |
| CNAG_07321 | - | hypothetical protein, variant | 483.835821 | 520.5390714 | 498.1738772 | 802.9750227 | 771.4049586 | 749.1775326 | 1.72E-09 | 0.61446852 | 3.60E-10 | 6.270621492 | 1.530993884 |
| CNAG_01954 | - | aldo-keto reductase | 5.502725426 | 12.2623323 | 0.294544198 | 319.6100652 | 316.0826965 | 324.225059 | 1.76E-09 | 5.808893217 | 3.68E-10 | 6.267006468 | 56.05974267 |
| CNAG_06278 | TCO7 | Two-component-like sensor kinase | 5398.375701 | 5543.300727 | 5877.101458 | 4091.087307 | 4195.285522 | 4538.43512 | 1.77E-09 | -0.406293816 | 3.69E-10 | -6.266427142 | 0.754559297 |
| CNAG_03215 | - | hypothetical protein | 164.201522 | 271.6484437 | 211.3734937 | 410.645778 | 524.3637242 | 412.7322675 | 1.77E-09 | 1.045380467 | 3.71E-10 | 6.265559737 | 2.063910581 |
| CNAG_00937 | - | hypothetical protein, variant 2 | 9.922912043 | 6.752061922 | 4.51547054 | 76.005946 | 54.06894468 | 41.99981916 | 1.79E-09 | 3.154025417 | 3.75E-10 | 6.263928258 | 8.901357785 |
| CNAG_00126 | - | 2-deoxy-D-gluconate 3-dehydrogenase | 1901.268348 | 1588.31913 | 1700.052565 | 2446.518248 | 2511.177353 | 2451.058361 | 1.84E-09 | 0.497590793 | 3.86E-10 | 6.259786684 | 1.411853889 |
| CNAG_02434 | ATX1 | Copper transport protein ATX1 | 1387.060639 | 1444.693277 | 1405.484465 | 1068.322122 | 986.5166238 | 979.3706742 | 1.86E-09 | -0.497002394 | 3.90E-10 | -6.258163766 | 0.708577522 |
| CNAG_05690 | RPD304 | histone deacetylase RPD3 | 4390.49531 | 4423.813754 | 4067.411016 | 3312.890145 | 3190.391086 | 3389.445357 | 1.87E-09 | -0.395966873 | 3.93E-10 | -6.256948554 | 0.759979878 |
| CNAG_03162 | - | vacuolar protein | 299.3228007 | 259.8999714 | 278.5237697 | 429.0076011 | 562.9698097 | 534.716436 | 1.91E-09 | 0.851312942 | 4.02E-10 | 6.253386328 | 1.804142059 |
| CNAG_01103 | PRP11 | splicing factor 3a subunit 2 | 2045.577885 | 1902.662942 | 1988.830116 | 1485.461344 | 1345.308797 | 1472.22784 | 1.93E-09 | -0.479668207 | 4.07E-10 | -6.251504206 | 0.717142534 |
| CNAG_00989 | UCH2 | ubiquitin carboxyl-terminal hydrolase L5 | 667.6501557 | 651.7284078 | 737.7794568 | 451.7698057 | 414.553909 | 438.7427054 | 1.94E-09 | -0.672681331 | 4.08E-10 | -6.250934122 | 0.627339656 |
| CNAG_03957 | - | 4-nitrophenyl phosphatase | 3257.855274 | 3309.778352 | 3478.650543 | 2618.697899 | 2588.378007 | 2342.659807 | 1.96E-09 | -0.427537219 | 4.13E-10 | -6.248904596 | 0.74352996 |
| CNAG_00604 | - | hypothetical protein | 363.6549446 | 427.8836769 | 417.1179408 | 672.2478391 | 666.242351 | 623.7247527 | 1.99E-09 | 0.685378312 | 4.18E-10 | 6.247168737 | 1.608123617 |
| CNAG_01130 | - | hypothetical protein | 1099.411653 | 885.580104 | 965.0142776 | 634.6268742 | 668.2526212 | 582.0141879 | 2.03E-09 | -0.662748447 | 4.27E-10 | -6.243718255 | 0.631673762 |
| CNAG_04773 | - | hypothetical protein | 32.0702672 | 19.71866749 | 20.53837333 | 79.10221291 | 113.091331 | 100.4638087 | 2.04E-09 | 2.023296964 | 4.29E-10 | 6.243014683 | 4.06511725 |
| CNAG_01693 | - | exosome complex component RRP45 | 1530.729752 | 1569.707114 | 1670.985189 | 1153.862034 | 1190.156094 | 1059.332981 | 2.10E-09 | -0.502679387 | 4.43E-10 | -6.238075418 | 0.705794754 |
| CNAG_01637 | - | COPII-coated vesicle component Erv46 | 4104.281244 | 3839.863084 | 3582.177988 | 2761.75675 | 2949.785571 | 2909.883197 | 2.15E-09 | -0.434652099 | 4.53E-10 | -6.234472549 | 0.739872151 |
| CNAG_05694 | CKA1 | CMGC/CK2 protein kinase | 6653.70963 | 7091.537139 | 6639.683934 | 5524.084311 | 5299.390491 | 5337.79561 | 2.15E-09 | -0.34982183 | 4.54E-10 | -6.234212484 | 0.784680999 |
| CNAG_01098 | - | pre-rRNA-processing protein ipi1 | 2374.7706 | 2048.136541 | 2136.020805 | 1542.951769 | 1586.895375 | 1595.470534 | 2.17E-09 | -0.4894195 | 4.58E-10 | -6.232934307 | 0.712311655 |
| CNAG_02203 | - | karyopherin/importin that interacts with the nuclear pore complex, variant | 1945.204172 | 1808.948726 | 2081.552815 | 1326.53962 | 1372.092916 | 1456.672999 | 2.17E-09 | -0.505612285 | 4.58E-10 | -6.232793493 | 0.70436138 |
| CNAG_02664 | - | D-lactate dehydrogenase | 1461.815021 | 1576.115479 | 1454.562717 | 1115.764072 | 1052.073456 | 1075.928439 | 2.23E-09 | -0.48498043 | 4.71E-10 | -6.228606635 | 0.714506762 |
| CNAG_01456 | - | hypothetical protein, variant | 13.4984949 | 19.65867322 | 13.06141054 | 75.83146554 | 66.00683594 | 81.50067383 | 2.25E-09 | 2.288678629 | 4.75E-10 | 6.227014313 | 4.886083874 |
| CNAG_06426 | - | U6 snRNA-associated Sm-like protein LSm6 | 1001.59564 | 963.1063904 | 999.1357712 | 663.4479433 | 720.9191271 | 627.4982449 | 2.28E-09 | -0.575460295 | 4.83E-10 | -6.224594061 | 0.67107211 |
| CNAG_02745 | DCR2 | dicer-like protein 2 | 2157.724856 | 2016.884557 | 2191.43762 | 1475.204492 | 1596.415262 | 1591.012172 | 2.28E-09 | -0.464613962 | 4.84E-10 | -6.224300324 | 0.724664958 |
| CNAG_02592 | - | thioredoxin reductase GliT | 51.89905389 | 43.3832605 | 34.40819644 | 127.7120404 | 118.5677297 | 150.3057326 | 2.30E-09 | 1.610505922 | 4.87E-10 | 6.223314124 | 3.053589059 |
| CNAG_03968 | PAB2 | polyadenylate-binding protein 2 | 2474.17809 | 2385.335913 | 2338.662006 | 1845.047721 | 1738.685631 | 1819.704686 | 2.30E-09 | -0.429185982 | 4.88E-10 | -6.222763249 | 0.742680713 |
| CNAG_07586 | - | hypothetical protein | 155.8403358 | 171.5177828 | 177.1956974 | 318.1691555 | 319.5432064 | 322.7269119 | 2.47E-09 | 0.915989367 | 5.23E-10 | 6.21194662 | 1.886862595 |

| | | | | | | | | | | | | | |
|---|---|---|---|---|---|---|---|---|---|---|---|---|---|
| CNAG_06256 | - | hypothetical protein | 7.561875871 | 5.656645076 | 2.361781151 | 50.16050823 | 52.52755011 | 46.18797167 | 2.49E-09 | 3.386476541 | 5.29E-10 | 6.210392275 | 10.45757573 |
| CNAG_06872 | - | 5-oxoprolinase | 21.69251226 | 32.5678628 | 21.59857964 | 88.75943097 | 98.15209763 | 98.26304707 | 2.54E-09 | 1.926356485 | 5.40E-10 | 6.207135769 | 3.800940616 |
| CNAG_04994 | - | signal recognition particle subunit SRP14 | 1210.537173 | 1079.500922 | 1129.265027 | 802.8274268 | 769.1328154 | 789.5209337 | 2.54E-09 | -0.550025817 | 5.41E-10 | -6.206724168 | 0.683007906 |
| CNAG_06300 | - | glutamate-cysteine ligase catalytic subunit, variant | 2024.416944 | 1947.822492 | 1633.790899 | 1276.749367 | 1315.172737 | 1299.085173 | 2.55E-09 | -0.541906265 | 5.42E-10 | -6.206335741 | 0.686862742 |
| CNAG_03603 | - | ATP-dependent RNA helicase HAS1 | 1239.602387 | 1013.80349 | 1116.450434 | 735.8061883 | 712.1617194 | 777.294892 | 2.56E-09 | -0.614722837 | 5.46E-10 | -6.205307536 | 0.653055343 |
| CNAG_03342 | - | tRNA (guanine-N(7)-)-methyltransferase subunit TRM82 | 1850.426946 | 1600.990255 | 1658.24444 | 1129.598159 | 1229.801538 | 1229.117344 | 2.62E-09 | -0.525850759 | 5.58E-10 | -6.201881978 | 0.694549412 |
| CNAG_04290 | - | hypothetical protein | 4624.169448 | 4650.133926 | 5485.644606 | 3629.609834 | 3491.39753 | 3693.657082 | 2.62E-09 | -0.46419638 | 5.59E-10 | -6.201615004 | 0.724874739 |
| CNAG_01015 | - | cytoplasmic protein | 146.441946 | 138.1528599 | 148.4200636 | 269.7919244 | 307.6228831 | 284.9870237 | 2.64E-09 | 0.97773338 | 5.64E-10 | 6.200271567 | 1.969368899 |
| CNAG_04096 | - | racemase | 35.57317995 | 22.95030658 | 25.86086148 | 107.7622012 | 119.0808286 | 84.14886921 | 2.68E-09 | 1.901736632 | 5.71E-10 | 6.198240576 | 3.736627193 |
| CNAG_04973 | - | hypothetical protein | 1058.652295 | 934.0262202 | 930.9239397 | 673.621894 | 634.5518344 | 649.6605878 | 2.69E-09 | -0.594693914 | 5.75E-10 | -6.197127549 | 0.662184931 |
| CNAG_04576 | - | hypothetical protein | 22.95038418 | 51.92230088 | 30.14132513 | 128.7998978 | 132.750668 | 109.5737861 | 2.70E-09 | 1.828201269 | 5.77E-10 | 6.196611092 | 3.550940702 |
| CNAG_05734 | SLA2 | cell wall organization and biogenesis-related protein, putative | 425.4551201 | 451.5946334 | 444.8471745 | 665.0286277 | 703.3107764 | 700.2589668 | 2.78E-09 | 0.632639299 | 5.94E-10 | 6.192076587 | 1.550398736 |
| CNAG_06554 | - | FAD dependent oxidoreductase | 354.3976268 | 332.7152995 | 320.8514746 | 198.7753637 | 161.713435 | 177.9291141 | 2.80E-09 | -0.923598095 | 6.00E-10 | -6.190532366 | 0.527192556 |
| CNAG_05974 | - | hypothetical protein | 92.33649175 | 70.8763678 | 84.96515507 | 20.17738746 | 11.47651704 | 23.67980816 | 2.80E-09 | -2.203417294 | 6.00E-10 | -6.190406243 | 0.217122735 |
| CNAG_02458 | - | GTPase activating protein | 1226.357974 | 1127.145373 | 1096.456769 | 1568.495069 | 1648.583384 | 1760.440394 | 2.84E-09 | 0.513360631 | 6.07E-10 | 6.188550055 | 1.427371262 |
| CNAG_04235 | - | hypothetical protein | 1371.891411 | 1391.929146 | 1504.603849 | 1005.961806 | 1061.643763 | 898.3906884 | 2.92E-09 | -0.540839277 | 6.26E-10 | -6.183765391 | 0.68737092 |
| CNAG_02718 | - | NAD binding dehydrogenase | 1002.19193 | 1001.055725 | 922.6258486 | 1422.365411 | 1344.906182 | 1418.576542 | 2.98E-09 | 0.501380055 | 6.38E-10 | 6.180653875 | 1.41556702 |
| CNAG_01790 | - | upstream activation factor subunit UAF30 | 923.3507596 | 870.4626767 | 926.6098 | 626.9139181 | 608.6981656 | 617.4542323 | 2.99E-09 | -0.570257977 | 6.41E-10 | -6.180042498 | 0.673496346 |
| CNAG_06685 | - | U2-associated protein SR140 | 2610.840383 | 2530.800028 | 2531.683889 | 1969.56314 | 1928.483617 | 1960.692203 | 3.01E-09 | -0.404586861 | 6.46E-10 | -6.178692428 | 0.755452598 |
| CNAG_03223 | - | hypothetical protein | 490.7462965 | 444.0894949 | 283.975956 | 864.9074066 | 733.4808491 | 759.058277 | 3.08E-09 | 0.937945545 | 6.62E-10 | 6.174863329 | 1.915798123 |
| CNAG_03793 | - | hypothetical protein, variant | 1014.530947 | 1013.751145 | 1117.465218 | 732.7841764 | 743.3163867 | 647.5000861 | 3.19E-09 | -0.582696024 | 6.84E-10 | -6.169697536 | 0.667714824 |
| CNAG_04963 | - | hypothetical protein, variant | 744.2906938 | 926.6661639 | 727.5020516 | 1152.918549 | 1285.568455 | 1279.740074 | 3.23E-09 | 0.618325619 | 6.94E-10 | 6.167536767 | 1.53509253 |
| CNAG_12387 | - | unspecified product | 35.75837138 | 53.05155821 | 51.41797281 | 136.2655201 | 118.4578889 | 170.1818039 | 3.24E-09 | 1.59527664 | 6.97E-10 | 6.166874491 | 3.021524499 |
| CNAG_03360 | - | hypothetical protein | 318.0738219 | 287.9228539 | 373.3640436 | 568.7408835 | 537.642544 | 588.0445946 | 3.29E-09 | 0.775992537 | 7.07E-10 | 6.164541736 | 1.712367711 |
| CNAG_02797 | CPL1 | putative secreted protein | 49.73551893 | 77.76886507 | 25.9303015 | 196.1000345 | 158.3464264 | 161.6577137 | 3.41E-09 | 1.757567843 | 7.34E-10 | 6.158640602 | 3.381276142 |
| CNAG_03744 | - | hypothetical protein | 5.244303133 | 6.728790644 | 4.499014559 | 47.70515672 | 48.96526695 | 50.50813888 | 3.41E-09 | 3.258315846 | 7.35E-10 | 6.158394095 | 9.568652991 |
| CNAG_05654 | - | hypothetical protein | 287.705676 | 312.6174159 | 293.4407385 | 536.9695782 | 468.5830066 | 508.1747012 | 3.41E-09 | 0.746565239 | 7.36E-10 | 6.158158575 | 1.677793589 |
| CNAG_07678 | - | hypothetical protein, variant | 116.0842265 | 113.3601374 | 102.6066453 | 234.1897461 | 256.7920614 | 219.5550324 | 3.42E-09 | 1.083545161 | 7.37E-10 | 6.157878532 | 2.119237329 |
| CNAG_01578 | - | hypothetical protein | 1287.769478 | 1327.247678 | 1277.513948 | 957.771868 | 948.6185726 | 841.7571518 | 3.63E-09 | -0.518022319 | 7.83E-10 | -6.1483137 | 0.698328463 |
| CNAG_06667 | - | hypothetical protein | 513.6843768 | 653.709482 | 607.695147 | 382.5096978 | 350.728646 | 328.9170353 | 3.64E-09 | -0.756475078 | 7.87E-10 | -6.147540821 | 0.591940848 |
| CNAG_00780 | - | hypothetical protein | 152.5450359 | 255.5025898 | 233.7208032 | 408.3832231 | 441.2578755 | 441.4526817 | 3.67E-09 | 0.997644326 | 7.93E-10 | 6.146352838 | 1.996737008 |
| CNAG_01498 | AKP2 | Arylsulfatase | 441.2678281 | 518.0311933 | 568.150281 | 317.8775504 | 286.8681382 | 282.2826636 | 3.78E-09 | -0.801754654 | 8.17E-10 | -6.141535204 | 0.57365106 |
| CNAG_01075 | - | methylmalonate-semialdehyde dehydrogenase (acylating) | 91.56975825 | 66.04196817 | 91.90667185 | 196.930436 | 198.903125 | 197.2123496 | 3.79E-09 | 1.236512536 | 8.20E-10 | 6.140982513 | 2.35628253 |
| CNAG_06260 | - | Alpha-glucosidase | 507.1513033 | 530.2323439 | 488.5898702 | 743.581914 | 797.3182648 | 805.7126426 | 3.81E-09 | 0.605759941 | 8.24E-10 | 6.140311743 | 1.521780141 |
| CNAG_06610 | - | MFS transporter | 576.4316829 | 579.4510324 | 528.8579361 | 349.7494878 | 316.1756166 | 369.8790796 | 3.93E-09 | -0.717731132 | 8.52E-10 | -6.13498944 | 0.608052951 |
| CNAG_00050 | - | hypothetical protein | 531.983528 | 468.5697654 | 514.922903 | 285.8105298 | 319.7365575 | 305.5461656 | 3.95E-09 | -0.750909921 | 8.55E-10 | -6.13431845 | 0.594228654 |
| CNAG_07576 | - | hypothetical protein | 409.3864114 | 337.1845556 | 483.809314 | 229.050874 | 212.6563088 | 203.501347 | 3.95E-09 | -0.948669508 | 8.55E-10 | -6.134332358 | 0.518110057 |
| CNAG_02497 | - | thaumatin family protein, variant | 14.79763358 | 27.25889203 | 44.98040458 | 118.6708232 | 133.8019948 | 97.41003277 | 3.96E-09 | 2.018581121 | 8.59E-10 | 6.133674055 | 4.051850997 |
| CNAG_05148 | CXT1 | beta-1,2-xylosyltransferase 1 | 1108.892299 | 1075.091208 | 929.90155 | 730.4750642 | 639.6806446 | 690.7345191 | 4.00E-09 | -0.61095731 | 8.67E-10 | -6.132147302 | 0.654762086 |
| CNAG_06453 | - | benzodiazapine receptor | 2305.781682 | 2951.275363 | 1965.77007 | 6126.447641 | 4951.457805 | 5262.893383 | 4.07E-09 | 1.164652925 | 8.84E-10 | 6.129033669 | 2.241792773 |
| CNAG_03418 | - | exportin-1 | 4002.739594 | 3839.86288 | 3578.979108 | 2845.863976 | 2986.890346 | 2888.796433 | 4.10E-09 | -0.404590572 | 8.91E-10 | -6.127869519 | 0.755450655 |
| CNAG_06108 | - | hypothetical protein, variant | 958.4117357 | 925.4025466 | 960.7485125 | 648.6777618 | 659.6181672 | 660.7373354 | 4.10E-09 | -0.546821984 | 8.90E-10 | -6.127940389 | 0.684526366 |
| CNAG_01381 | - | hypothetical protein | 1295.914273 | 1235.724226 | 1305.217935 | 924.2576166 | 913.2218386 | 911.6280713 | 4.10E-09 | -0.496312796 | 8.92E-10 | -6.127539275 | 0.708916298 |
| CNAG_01510 | - | hypothetical protein | 2375.908383 | 2128.898493 | 1993.158186 | 1497.783131 | 1617.119836 | 1527.776622 | 4.11E-09 | -0.500821883 | 8.95E-10 | -6.127137265 | 0.706704067 |
| CNAG_02680 | VPS15 | VPS15 protein kinase | 2402.974276 | 2418.709803 | 2250.163982 | 1742.23785 | 1831.052887 | 1769.758851 | 4.29E-09 | -0.41952289 | 9.33E-10 | -6.120466766 | 0.747671844 |
| CNAG_07866 | - | transcription initiation factor TFIIA small subunit, variant 2 | 333.3826553 | 417.1306303 | 441.6258875 | 730.8327709 | 628.1659273 | 663.6251168 | 4.29E-09 | 0.748411204 | 9.34E-10 | 6.120304218 | 1.679941743 |
| CNAG_07322 | - | hypothetical protein | 3209.739062 | 2796.953408 | 2871.88671 | 2178.868103 | 2149.273862 | 2230.464002 | 4.29E-09 | -0.452643864 | 9.35E-10 | -6.120069028 | 0.730702544 |
| CNAG_02546 | - | transferase | 3087.246841 | 2715.108177 | 3363.405261 | 2224.013011 | 2167.377303 | 2162.786638 | 4.32E-09 | -0.499979396 | 9.42E-10 | -6.118946459 | 0.70711688 |
| CNAG_00651 | - | ATP-binding cassette, subfamily D (ALD), peroxisomal long-chain fatty acid import protein | 141.8003737 | 97.3117286 | 163.314609 | 290.2292015 | 327.587284 | 277.2573645 | 4.45E-09 | 1.139683063 | 9.71E-10 | 6.114044991 | 2.203326143 |
| CNAG_03331 | - | Phosphatidylserine decarboxylase | 2050.317666 | 2053.466978 | 1963.273083 | 1499.42325 | 1575.742568 | 1472.258972 | 4.47E-09 | -0.43142936 | 9.76E-10 | -6.11324094 | 0.741526749 |
| CNAG_01302 | - | hypothetical protein | 1111.322011 | 963.1794161 | 1181.42637 | 691.399127 | 715.6481013 | 750.6524972 | 4.49E-09 | -0.610248061 | 9.80E-10 | -6.112579947 | 0.655084055 |
| CNAG_00345 | - | porphobilinogen synthase | 2474.245425 | 2508.148 | 2399.455789 | 1887.055125 | 1856.890915 | 1906.285665 | 4.50E-09 | -0.400924076 | 9.84E-10 | -6.111961153 | 0.757373015 |
| CNAG_04179 | - | lysine-tRNA ligase | 7543.51267 | 7972.872702 | 7604.806987 | 6183.296003 | 6194.779239 | 5410.040956 | 4.52E-09 | -0.393394797 | 9.89E-10 | -6.11111036 | 0.761336 |
| CNAG_00449 | - | RNA-binding protein NOB1, variant | 2718.203091 | 2443.563569 | 2683.079059 | 1855.818753 | 1881.009584 | 2031.715353 | 4.54E-09 | -0.459446255 | 9.93E-10 | -6.110582596 | 0.727265349 |
| CNAG_00746 | CAS35 | capsular associated protein | 5328.354232 | 5652.113491 | 5738.490466 | 4405.629249 | 4376.458612 | 4459.640605 | 4.57E-09 | -0.351515456 | 1.00E-09 | -6.109349799 | 0.783760377 |
| CNAG_07953 | - | hypothetical protein | 768.1544803 | 791.7805851 | 864.7056533 | 511.6101335 | 526.7392732 | 563.026789 | 4.64E-09 | -0.613905441 | 1.02E-09 | -6.106911389 | 0.653425453 |
| CNAG_06298 | - | hypothetical protein | 6870.663979 | 1038.918347 | 1851.300423 | 111.5463635 | 107.0134774 | 101.6674329 | 4.82E-09 | -4.958329832 | 1.06E-09 | -6.100719258 | 0.032165773 |
| CNAG_04207 | - | homeobox protein cut-like | 2384.295321 | 2271.153666 | 2423.934379 | 1677.584738 | 1743.022638 | 1849.64552 | 4.87E-09 | -0.441261511 | 1.07E-09 | -6.098744615 | 0.73649033 |
| CNAG_04647 | - | Glutathione synthetase | 406.6654253 | 353.6027241 | 363.824847 | 582.534982 | 611.8703462 | 643.5691332 | 4.92E-09 | 0.694840152 | 1.08E-09 | 6.09720191 | 1.618705074 |
| CNAG_05129 | - | hypothetical protein | 21.75266567 | 36.89507224 | 36.4869207 | 96.36293986 | 105.7809761 | 126.9438029 | 5.07E-09 | 1.800126785 | 1.11E-09 | 6.092237928 | 3.482508285 |
| CNAG_02286 | - | cytoplasmic protein | 3030.047748 | 2941.293601 | 2868.695427 | 2163.273273 | 2242.466151 | 2340.334534 | 5.17E-09 | -0.405356167 | 1.14E-09 | -6.089082363 | 0.755049866 |
| CNAG_04092 | - | Sugar transporter | 15.82432581 | 14.31498134 | 20.4940143 | 83.79748782 | 85.36890341 | 61.93554255 | 5.26E-09 | 2.208543862 | 1.16E-09 | 6.086244097 | 4.622085228 |
| CNAG_06724 | - | DNA repair and recombination protein RAD52 | 260.8954401 | 271.7200192 | 305.1342729 | 438.4692684 | 498.0941064 | 523.6036875 | 5.29E-09 | 0.788185505 | 1.16E-09 | 6.085101559 | 1.726901152 |
| CNAG_06356 | - | mitochondrial protein | 1144.016897 | 1103.150492 | 1038.652282 | 728.7942868 | 804.5842455 | 748.4576311 | 5.42E-09 | -0.542279493 | 1.19E-09 | -6.081184726 | 0.686685073 |
| CNAG_01370 | - | hypothetical protein | 171.9030137 | 143.06759 | 150.1309381 | 51.27458488 | 74.70974678 | 47.07692285 | 5.49E-09 | -1.448159777 | 1.21E-09 | -6.078987064 | 0.366488599 |
| CNAG_05574 | - | phosphatidylinositol glycan, class C | 389.2908192 | 441.8794117 | 416.0585051 | 644.0731976 | 655.8418143 | 664.7315003 | 5.59E-09 | 0.640976271 | 1.23E-09 | 6.075953301 | 1.559384037 |
| CNAG_04610 | - | hypothetical protein | 7.533333765 | 2.412533912 | 0.18267641 | 47.06491338 | 39.24610003 | 54.73924867 | 5.61E-09 | 3.942946642 | 1.24E-09 | 6.075533214 | 15.37960603 |
| CNAG_02539 | - | AAT family amino acid transporter | 4036.522088 | 3524.22441 | 3681.315326 | 2585.026103 | 2873.022091 | 2847.706357 | 5.70E-09 | -0.45261817 | 1.26E-09 | -6.072770971 | 0.730715558 |
| CNAG_04388 | SOD2 | mitochondrial manganese superoxide dismutase | 2179.194587 | 1786.576922 | 1934.676299 | 2649.504165 | 3052.106749 | 2848.478138 | 5.72E-09 | 0.51902383 | 1.26E-09 | 6.07221563 | 1.43298532 |
| CNAG_06369 | - | geranylgeranyl transferase type-2 subunit alpha | 1185.034972 | 1288.421244 | 1155.961944 | 860.4202642 | 886.5162037 | 819.5212542 | 5.89E-09 | -0.515123678 | 1.30E-09 | -6.067351769 | 0.699732944 |
| CNAG_03702 | ALF1 | tubulin-folding cofactor B, variant | 916.462656 | 972.7630219 | 961.8142831 | 679.8872225 | 619.8792433 | 664.0639133 | 5.95E-09 | -0.553050103 | 1.31E-09 | -6.065735289 | 0.681577633 |
| CNAG_03319 | - | phospholipid binding protein | 340.0572971 | 229.7816309 | 266.8131025 | 490.0250911 | 555.1555842 | 509.272151 | 5.95E-09 | 0.879294177 | 1.32E-09 | 6.065503106 | 1.839475138 |
| CNAG_04864 | CIR1 | iron regulator 1, variant | 1818.384342 | 1641.08148 | 1690.444311 | 2265.044476 | 2493.959354 | 2365.588503 | 5.99E-09 | 0.45243485 | 1.32E-09 | 6.064503769 | 1.368347682 |
| CNAG_04402 | - | splicing factor U2AF 35 kDa subunit | 1898.604932 | 1989.901703 | 2065.590332 | 1355.318574 | 1530.025198 | 1482.220346 | 6.27E-09 | -0.461938905 | 1.39E-09 | -6.057053643 | 0.726009884 |
| CNAG_06071 | - | hypothetical protein | 423.4890895 | 417.869605 | 435.9938449 | 274.3425559 | 239.4271814 | 232.3338276 | 6.50E-09 | -0.790501812 | 1.44E-09 | -6.05103628 | 0.578142962 |
| CNAG_03714 | - | microtubule motor protein | 2122.096847 | 2259.471528 | 2255.631989 | 2945.363948 | 2901.825703 | 2907.372842 | 6.64E-09 | 0.384454264 | 1.47E-09 | 6.047608416 | 1.305365906 |
| CNAG_05256 | CAT2 | catalase 2 | 1250.248395 | 553.1842254 | 566.6817243 | 4324.300301 | 4325.057944 | 4152.365327 | 6.69E-09 | 2.414879741 | 1.48E-09 | 6.046273876 | 5.332750168 |
| CNAG_04785 | - | hypothetical protein | 355.3405191 | 322.3592082 | 356.3305285 | 557.7450375 | 566.1691294 | 558.1677514 | 6.71E-09 | 0.686845396 | 1.49E-09 | 6.045730254 | 1.609759759 |
| CNAG_02089 | - | hypothetical protein | 211.9049669 | 268.4228025 | 241.1865069 | 423.9655596 | 416.9441147 | 433.7554643 | 6.78E-09 | 0.806664991 | 1.51E-09 | 6.043789993 | 1.749163308 |
| CNAG_07368 | - | calcium activated cation channel protein | 1319.274688 | 1369.249117 | 1227.430288 | 947.6137842 | 948.6052575 | 910.5358897 | 7.00E-09 | -0.496034984 | 1.55E-09 | -6.038603707 | 0.709052824 |
| CNAG_09002 | - | apocytochrome b | 209761.7542 | 181104.4908 | 183375.5411 | 150164.6138 | 147038.0058 | 146514.7668 | 7.11E-09 | -0.387919069 | 1.58E-09 | -6.036103632 | 0.76423113 |
| CNAG_06006 | - | hypothetical protein | 1409.33111 | 1495.356115 | 1567.54477 | 1058.862546 | 1117.695166 | 1069.270709 | 7.45E-09 | -0.477986795 | 1.66E-09 | -6.028306281 | 0.717978827 |
| CNAG_02326 | - | V-type proton ATPase catalytic subunit A | 4309.976475 | 4364.576545 | 4170.845877 | 3457.713629 | 3331.887967 | 3365.037633 | 7.52E-09 | -0.354190699 | 1.67E-09 | -6.026680442 | 0.782308368 |
| CNAG_00994 | - | protein mago nashi | 2586.223737 | 2370.271996 | 2391.979924 | 1837.227595 | 1905.242913 | 1717.632788 | 7.52E-09 | -0.444249504 | 1.67E-09 | -6.026714955 | 0.73496655 |
| CNAG_07806 | - | hypothetical protein | 272.5059929 | 258.8011904 | 297.6689081 | 487.0780654 | 447.0578808 | 478.1861787 | 7.53E-09 | 0.756367359 | 1.68E-09 | 6.026475398 | 1.689231855 |
| CNAG_06924 | - | beta-fructofuranosidase | 718.354646 | 575.153085 | 502.2384386 | 391.1712731 | 272.9398167 | 272.3836539 | 7.53E-09 | -0.956811507 | 1.68E-09 | -6.026377821 | 0.515194284 |
| CNAG_00100 | - | stress-induced-phosphoprotein 1 | 9261.329174 | 9512.575511 | 10057.61357 | 7293.011526 | 7801.726841 | 7749.040943 | 7.56E-09 | -0.351079714 | 1.68E-09 | -6.025592507 | 0.783997134 |

| | | | | | | | | | | | | | |
|---|---|---|---|---|---|---|---|---|---|---|---|---|---|
| CNAG_05107 | - | prefoldin subunit 4 | 1555.035503 | 1341.350424 | 1556.858272 | 1037.86991 | 1084.921016 | 895.0854472 | 7.60E-09 | -0.578036143 | 1.70E-09 | -6.024558249 | 0.669875019 |
| CNAG_04957 | - | hypothetical protein | 1367.243289 | 1386.553181 | 1477.965008 | 1015.302566 | 1057.316833 | 991.5751448 | 7.65E-09 | -0.48106952 | 1.71E-09 | -6.023539689 | 0.7164463 |
| CNAG_05822 | - | protein RAI1 | 1235.013918 | 1070.887939 | 1109.016234 | 794.2492311 | 786.4068726 | 781.7573483 | 7.69E-09 | -0.547872846 | 1.72E-09 | -6.022655014 | 0.684027937 |
| CNAG_05349 | - | hypothetical protein | 5945.94102 | 5675.839187 | 5593.494614 | 4439.056264 | 4591.257995 | 4600.629192 | 7.99E-09 | -0.352141309 | 1.79E-09 | -6.016233337 | 0.783420449 |
| CNAG_01658 | - | hypothetical protein | 980.0784976 | 1050.610487 | 979.9720372 | 1293.614299 | 1607.37297 | 1516.228638 | 8.00E-09 | 0.563603629 | 1.79E-09 | 6.016084751 | 1.477956317 |
| CNAG_01169 | - | Aspartyl aminopeptidase | 707.5279165 | 811.1216041 | 832.7184023 | 510.8764681 | 516.3992819 | 525.3125962 | 8.00E-09 | -0.614257481 | 1.79E-09 | -6.015996141 | 0.653266027 |
| CNAG_07354 | MET22 | 3'(2'),5'-bisphosphate nucleotidase | 915.1694746 | 889.8251061 | 888.2420151 | 565.2879979 | 605.2545322 | 654.0307712 | 8.21E-09 | -0.576850681 | 1.84E-09 | -6.011564018 | 0.670425681 |
| CNAG_05011 | - | iron donor protein CyaY | 994.6953067 | 1010.511231 | 1084.421375 | 751.5269825 | 715.6869101 | 651.9275857 | 8.23E-09 | -0.560004603 | 1.84E-09 | -6.011238168 | 0.6783 |
| CNAG_12566 | - | unspecified product | 7.565578808 | 5.658128648 | 5.565583125 | 40.26994033 | 55.18901424 | 52.69004821 | 8.42E-09 | 3.096321236 | 1.89E-09 | 6.007284884 | 8.552352015 |
| CNAG_04467 | - | succinate-semialdehyde dehydrogenase (NADP) | 731.9208499 | 754.0549636 | 783.6842404 | 466.4313542 | 499.1491776 | 535.2597253 | 8.57E-09 | -0.611584772 | 1.92E-09 | -6.004342666 | 0.654477376 |
| CNAG_01251 | UFD4 | E3 ubiquitin-protein ligase TRIP12 | 3593.362075 | 3551.312046 | 3228.285338 | 4392.076957 | 4670.117952 | 4712.451319 | 8.74E-09 | 0.394064116 | 1.96E-09 | 6.001134494 | 1.314090027 |
| CNAG_02299 | - | hypothetical protein | 1265.589212 | 1236.795926 | 1354.239192 | 877.4831859 | 978.8591929 | 836.2008926 | 8.83E-09 | -0.534246235 | 1.98E-09 | -5.999445608 | 0.690519358 |
| CNAG_00997 | - | hypothetical protein | 982.9195947 | 1000.765897 | 999.1232975 | 590.1378134 | 759.8127409 | 568.7167605 | 8.93E-09 | -0.652564981 | 2.00E-09 | -5.99752725 | 0.636148294 |
| CNAG_03267 | - | splicing factor 3B subunit 2 | 3105.946193 | 3088.877165 | 2858.049886 | 2173.361578 | 2367.574108 | 2340.355109 | 9.29E-09 | -0.410889468 | 2.09E-09 | -5.991006102 | 0.7521595 |
| CNAG_01104 | - | ribosomal RNA-processing protein 8 | 1732.706502 | 1655.944093 | 1876.797718 | 1185.64899 | 1266.881231 | 1328.997477 | 9.34E-09 | -0.493413119 | 2.10E-09 | -5.989985733 | 0.710342584 |
| CNAG_03106 | - | hypothetical protein | 659.1284275 | 508.498171 | 650.291261 | 398.1404478 | 290.2052466 | 332.2412977 | 9.35E-09 | -0.849200963 | 2.10E-09 | -5.989665251 | 0.555092089 |
| CNAG_05248 | - | clathrin binding protein | 3115.283369 | 3015.634393 | 2876.175456 | 2281.626625 | 2302.849694 | 2352.562675 | 9.37E-09 | -0.392282933 | 2.11E-09 | -5.989237719 | 0.761922977 |
| CNAG_06887 | - | hypothetical protein | 528.5949478 | 484.7418718 | 557.5511488 | 325.6189529 | 317.9635504 | 326.6249779 | 9.46E-09 | -0.712412419 | 2.13E-09 | -5.987726163 | 0.610298767 |
| CNAG_01056 | - | hypothetical protein | 21.66271761 | 19.70420856 | 29.00947371 | 80.90264242 | 90.3535377 | 94.88862766 | 9.75E-09 | 1.920161829 | 2.19E-09 | 5.982716244 | 3.784655093 |
| CNAG_02512 | - | DNA repair protein RAD16 | 1998.38593 | 2250.852984 | 2278.020685 | 2879.15649 | 2919.090306 | 3029.433071 | 9.87E-09 | 0.42093362 | 2.22E-09 | 5.980631386 | 1.338793656 |
| CNAG_01823 | INM1 | myo-inositol-1(or 4)-monophosphatase | 2750.838416 | 2541.560671 | 2381.351114 | 1872.206512 | 1919.865195 | 1918.515665 | 9.89E-09 | -0.441855007 | 2.23E-09 | -5.980173867 | 0.736187416 |
| CNAG_00332 | SIP4 | hypothetical protein | 325.0202373 | 359.9854788 | 262.5788522 | 523.4658273 | 545.4919118 | 563.6316888 | 9.91E-09 | 0.771302763 | 2.23E-09 | 5.979762171 | 1.706810349 |
| CNAG_02899 | - | hypothetical protein | 7.622929785 | 11.04449594 | 8.775718585 | 52.9763361 | 71.80660979 | 48.54617268 | 1.00E-08 | 2.702637224 | 2.27E-09 | 5.977373277 | 6.50990831 |
| CNAG_04494 | - | U1 small nuclear ribonucleoprotein C | 1381.172183 | 1326.227652 | 1385.21016 | 1025.49506 | 990.0010444 | 953.834598 | 1.02E-08 | -0.478779601 | 2.31E-09 | -5.974139001 | 0.717584384 |
| CNAG_06644 | ERG5 | C-22 sterol desaturase | 830.6726764 | 675.4160019 | 635.5377598 | 452.5214264 | 379.1015982 | 452.0560692 | 1.04E-08 | -0.753815946 | 2.34E-09 | -5.972157791 | 0.593032902 |
| CNAG_00180 | - | ubiquitin carboxyl-terminal hydrolase L3 | 1052.984267 | 1163.373258 | 932.0432097 | 747.6121241 | 689.7652461 | 665.2407113 | 1.10E-08 | -0.59750017 | 2.49E-09 | -5.961807925 | 0.660898135 |
| CNAG_04306 | - | vesicle transporter SFT2B | 1228.132264 | 1115.072398 | 1229.483873 | 871.3696836 | 837.308941 | 802.8705875 | 1.10E-08 | -0.524772542 | 2.50E-09 | -5.961581785 | 0.695068687 |
| CNAG_05997 | - | hypothetical protein | 1424.394221 | 1429.627779 | 1509.945376 | 1138.471689 | 943.3357053 | 892.8544393 | 1.11E-08 | -0.568359586 | 2.51E-09 | -5.960592023 | 0.674383159 |
| CNAG_03399 | - | SIT4-associating protein/190 | 5961.136646 | 5609.057189 | 5819.560428 | 4593.980625 | 4492.889833 | 4678.334056 | 1.12E-08 | -0.352856755 | 2.53E-09 | -5.959311156 | 0.78303204 |
| CNAG_00492 | - | hypothetical protein | 22.95424354 | 43.3590606 | 47.1297486 | 110.4075824 | 123.0888967 | 135.9118894 | 1.18E-08 | 1.702419768 | 2.66E-09 | 5.951111962 | 3.254463578 |
| CNAG_01440 | - | hypothetical protein | 65.9234392 | 70.2575987 | 50.39239742 | 158.8542617 | 146.1577028 | 168.233548 | 1.19E-08 | 1.336714328 | 2.70E-09 | 5.949152111 | 2.525754343 |
| CNAG_03686 | - | UDP-N-acetylglucosamine transferase subunit ALG14 | 616.0595952 | 554.7440375 | 551.2466623 | 370.8012723 | 349.8721152 | 368.7970227 | 1.21E-08 | -0.678424671 | 2.74E-09 | -5.946685487 | 0.624847194 |
| CNAG_12675 | - | unspecified product | 43.80842038 | 46.59392242 | 41.84012641 | 119.792021 | 156.2369927 | 108.5714442 | 1.25E-08 | 1.543356021 | 2.83E-09 | 5.941290725 | 2.914717419 |
| CNAG_05156 | - | hypothetical protein | 1235.858739 | 1410.45601 | 1270.282581 | 1670.280891 | 2029.966239 | 2015.774313 | 1.25E-08 | 0.530782974 | 2.84E-09 | 5.940768024 | 1.444713052 |
| CNAG_00023 | - | hypothetical protein | 56.56120494 | 41.25464231 | 49.29222265 | 145.8378885 | 151.6933689 | 115.2527756 | 1.29E-08 | 1.476720482 | 2.93E-09 | 5.935236291 | 2.783153504 |
| CNAG_03327 | - | hypothetical protein | 31.06884273 | 53.00391163 | 37.58158644 | 126.3146148 | 130.0649139 | 112.9017451 | 1.29E-08 | 1.590991242 | 2.94E-09 | 5.935009911 | 3.012562646 |
| CNAG_05163 | - | exoribonuclease II | 683.9236444 | 620.5156618 | 721.7916201 | 422.1096565 | 465.5676905 | 407.7024002 | 1.31E-08 | -0.66128885 | 2.97E-09 | -5.933085236 | 0.63231316 |
| CNAG_09008 | - | ATP synthase subunit 6 | 122307.6243 | 103974.3479 | 109169.5661 | 89816.0389 | 83388.73454 | 83695.21136 | 1.34E-08 | -0.400871913 | 3.06E-09 | -5.928356092 | 0.7574004 |
| CNAG_05790 | - | OPT family small oligopeptide transporter | 1249.762696 | 1301.636918 | 1218.008941 | 1677.484321 | 1725.334125 | 1740.551066 | 1.35E-08 | 0.433507128 | 3.07E-09 | 5.927861706 | 1.350512626 |
| CNAG_01550 | - | pre-rRNA-processing protein TSR3 | 1418.441923 | 1303.598681 | 1318.046731 | 971.7344019 | 1010.747705 | 896.1390253 | 1.37E-08 | -0.50514768 | 3.13E-09 | -5.924764172 | 0.70458825 |
| CNAG_04594 | FZC27 | hypothetical protein | 352.0549681 | 449.4348218 | 468.2778372 | 676.0054951 | 705.9292472 | 692.4844591 | 1.42E-08 | 0.693455677 | 3.23E-09 | 5.919312627 | 1.617152437 |
| CNAG_01308 | - | protein NRD1 | 200.1878882 | 232.8888473 | 223.0472316 | 381.9206775 | 389.3951374 | 379.4082326 | 1.42E-08 | 0.796008344 | 3.25E-09 | 5.91852862 | 1.736290497 |
| CNAG_01567 | - | hypothetical protein | 48.37960568 | 36.92999264 | 36.51655162 | 104.022708 | 134.4749625 | 123.8521198 | 1.43E-08 | 1.572823083 | 3.26E-09 | 5.918130607 | 2.974862697 |
| CNAG_00991 | - | flap endonuclease 1 | 2030.476286 | 2096.540433 | 1899.306963 | 1501.768798 | 1535.177625 | 1503.321653 | 1.44E-08 | -0.423625236 | 3.30E-09 | -5.916080347 | 0.745548837 |
| CNAG_00626 | - | hypothetical protein | 4673.102317 | 4743.809319 | 4468.378289 | 3842.282592 | 3274.015565 | 3372.85784 | 1.47E-08 | -0.419693494 | 3.35E-09 | -5.913382819 | 0.747583434 |
| CNAG_06252 | CCD6 | Fungal Zn(2)-Cys(6) binuclear cluster domain protein | 675.8534492 | 682.9554068 | 721.8256216 | 486.0906003 | 437.8676274 | 448.7375818 | 1.49E-08 | -0.615175956 | 3.40E-09 | -5.911249544 | 0.652850265 |
| CNAG_04345 | ARO8001 | specific RNA polymerase II transcription factor | 351.7727542 | 312.6461841 | 293.4633841 | 554.7990469 | 523.8786201 | 511.5734385 | 1.49E-08 | 0.716582035 | 3.39E-09 | 5.911303677 | 1.643284227 |
| CNAG_03403 | - | mitochondrial protein | 1037.759609 | 994.339314 | 975.7075348 | 620.5141038 | 731.2652772 | 708.4593081 | 1.49E-08 | -0.561367982 | 3.42E-09 | -5.910106859 | 0.677659294 |
| CNAG_07531 | - | pyridoxal 5'-phosphate synthase, glutaminase subunit Pdx2 | 634.0249133 | 795.9823941 | 802.8253797 | 446.1872959 | 512.1522761 | 438.7982455 | 1.53E-08 | -0.691086967 | 3.50E-09 | -5.906020459 | 0.619387011 |
| CNAG_01166 | URE4 | urease accessory protein | 753.0615978 | 891.890644 | 851.9445515 | 499.0833274 | 595.8423735 | 544.2032092 | 1.55E-08 | -0.621861252 | 3.55E-09 | -5.903756911 | 0.649832025 |
| CNAG_04668 | UBC12 | ubiquitin-conjugating enzyme E2 M | 426.506087 | 371.9136535 | 360.6446865 | 671.5546654 | 682.7310776 | 563.8651367 | 1.56E-08 | 0.711818312 | 3.58E-09 | 5.902311925 | 1.637867116 |
| CNAG_05579 | - | hypothetical protein | 3786.775721 | 3522.077558 | 3784.746559 | 2867.707612 | 2866.108749 | 2894.326406 | 1.57E-08 | -0.378189723 | 3.59E-09 | -5.902123247 | 0.769402423 |
| CNAG_03926 | MAP1 | methionine aminopeptidase, type I | 6251.859421 | 5474.418102 | 6445.505649 | 4529.312871 | 4662.828961 | 4554.042856 | 1.60E-08 | -0.418680378 | 3.66E-09 | -5.898776475 | 0.748108601 |
| CNAG_05313 | - | hypothetical protein, variant 4 | 251.4855085 | 267.3668055 | 244.395237 | 450.4998723 | 457.5875529 | 398.393802 | 1.61E-08 | 0.761077253 | 3.70E-09 | 5.897133311 | 1.694755618 |
| CNAG_00637 | - | Cystathionine beta-synthase | 282.953126 | 293.2079575 | 231.6383663 | 453.4657219 | 464.4095972 | 469.3021277 | 1.63E-08 | 0.767206079 | 3.73E-09 | 5.895554577 | 1.701970556 |
| CNAG_04225 | - | hypothetical protein | 53.51123303 | 135.05144 | 135.7584006 | 518.8235274 | 539.9755589 | 596.3633317 | 1.63E-08 | 2.343329509 | 3.74E-09 | 5.895187471 | 5.074724525 |
| CNAG_03766 | - | ankyrin repeat-containing protein | 1529.42612 | 1379.055121 | 1563.271756 | 1037.054632 | 1135.84002 | 987.1736422 | 1.66E-08 | -0.517099072 | 3.81E-09 | -5.892281651 | 0.698775498 |
| CNAG_02469 | - | cytoplasmic protein | 1328.578951 | 1296.025518 | 1242.3432 | 880.5782322 | 923.5747 | 975.9660693 | 1.67E-08 | -0.492031005 | 3.83E-09 | -5.891175605 | 0.711023425 |
| CNAG_01707 | CRG3 | Putative regulator of G-protein signaling protein | 675.167408 | 552.9280988 | 504.6435173 | 915.6767211 | 1093.452132 | 865.7963494 | 1.67E-08 | 0.714675751 | 3.84E-09 | 5.89089079 | 1.641114332 |
| CNAG_04286 | - | DNA-directed RNA polymerase I, II, and III subunit RPABC3 | 1610.12769 | 1621.436 | 1759.495795 | 1287.064575 | 1195.285961 | 1084.883423 | 1.68E-08 | -0.500138037 | 3.87E-09 | -5.889620927 | 0.707039128 |
| CNAG_05696 | - | ubiquitin-conjugating enzyme E2-16 kDa | 2491.609383 | 2290.528347 | 2379.14779 | 1861.471699 | 1452.176695 | 1713.152827 | 1.72E-08 | -0.525884114 | 3.95E-09 | -5.886169995 | 0.694533355 |
| CNAG_01202 | - | hypothetical protein | 522.7965353 | 499.8058995 | 551.1699694 | 343.588685 | 317.9562141 | 326.6310319 | 1.76E-08 | -0.688386618 | 4.06E-09 | -5.881875475 | 0.620547427 |
| CNAG_07464 | MBS1 | Mbp1 and Swi4-like APSES protein 1 | 2815.226511 | 2808.765019 | 2705.533711 | 2205.427126 | 2159.675393 | 2116.12734 | 1.77E-08 | -0.377201204 | 4.08E-09 | -5.881074526 | 0.769929789 |
| CNAG_01668 | - | hypothetical protein | 7.585401068 | 7.812308413 | 8.759104621 | 49.1665911 | 51.52737352 | 52.75804248 | 1.80E-08 | 2.773238198 | 4.16E-09 | 5.8778405 | 6.836406566 |
| CNAG_04110 | - | Glutathione S-transferase | 502.4591333 | 512.9967097 | 490.7086501 | 746.7774486 | 743.766902 | 772.4292939 | 1.82E-08 | 0.573076515 | 4.20E-09 | 5.876245494 | 1.487692662 |
| CNAG_00583 | - | hypothetical protein | 4169.77068 | 3946.559205 | 3962.87261 | 2937.650949 | 3158.533395 | 3246.217429 | 1.86E-08 | -0.386076982 | 4.29E-09 | -5.872732068 | 0.765207553 |
| CNAG_07842 | - | hypothetical protein | 89.26636496 | 95.03086673 | 80.21885913 | 208.7396064 | 206.7119005 | 175.1906598 | 1.89E-08 | 1.142671462 | 4.37E-09 | 5.869524899 | 2.207894843 |
| CNAG_05647 | - | hypothetical protein | 1604.182848 | 1529.85549 | 1512.155206 | 1112.569278 | 1159.95012 | 1166.929694 | 1.99E-08 | -0.449939548 | 4.59E-09 | -5.861224242 | 0.732073523 |
| CNAG_03363 | - | hypothetical protein | 964.2106292 | 873.7248985 | 972.4585286 | 655.7217882 | 649.264796 | 635.2246755 | 2.01E-08 | -0.550467413 | 4.66E-09 | -5.859016088 | 0.682798875 |
| CNAG_05778 | - | WSC domain-containing protein, variant | 225.7584645 | 152.2397797 | 227.3033981 | 414.0846566 | 427.5344004 | 351.7489818 | 2.03E-08 | 0.963721785 | 4.70E-09 | 5.857585168 | 1.950334777 |
| CNAG_00778 | - | hypothetical protein | 477.9455999 | 474.2292052 | 482.1656103 | 698.4566471 | 719.6295497 | 762.3974859 | 2.07E-08 | 0.589416738 | 4.80E-09 | 5.854025329 | 1.50463832 |
| CNAG_06370 | BAT2 | Branched-chain amino acid aminotransferase | 3960.914124 | 4386.0893 | 4404.330501 | 2917.342771 | 3499.323255 | 3073.115517 | 2.08E-08 | -0.44127722 | 4.82E-09 | -5.853193708 | 0.736482311 |
| CNAG_03171 | SWE1 | WEE protein kinase | 1242.838259 | 1331.82824 | 1379.010224 | 1715.526215 | 1956.613277 | 1840.471307 | 2.14E-08 | 0.46455733 | 4.97E-09 | 5.848294905 | 1.379893888 |
| CNAG_06018 | - | aldehyde dehydrogenase (NAD), variant | 377.653298 | 341.3314923 | 301.7203888 | 175.2327442 | 198.0766518 | 187.9012764 | 2.19E-08 | -0.879218708 | 5.07E-09 | -5.844710201 | 0.543661772 |
| CNAG_00562 | - | hypothetical protein | 1808.369043 | 1569.734583 | 1536.702672 | 1148.353784 | 1188.395574 | 1178.059353 | 2.28E-08 | -0.499279034 | 5.29E-09 | -5.837911559 | 0.707460236 |
| CNAG_07747 | - | acyl-CoA oxidase | 516.4611058 | 526.9983826 | 474.734453 | 731.1087505 | 832.7782471 | 755.8308183 | 2.38E-08 | 0.597387246 | 5.53E-09 | 5.830396265 | 1.512974053 |
| CNAG_02113 | - | clathrin assembly protein | 1661.388819 | 1608.488239 | 1510.044482 | 1174.097412 | 1171.142886 | 1193.578795 | 2.40E-08 | -0.449042369 | 5.58E-09 | -5.82888193 | 0.732528924 |
| CNAG_03365 | - | hypothetical protein, variant | 2092.46421 | 2211.850985 | 2171.212557 | 1629.410047 | 1697.361687 | 1649.852003 | 2.42E-08 | -0.39503083 | 5.63E-09 | -5.827261765 | 0.760473125 |
| CNAG_07442 | - | phosphatidylglycerol/phosphatidylinositol transfer protein | 1997.19505 | 2245.454424 | 2202.310199 | 2957.108538 | 2888.05306 | 2761.978302 | 2.44E-08 | 0.402544554 | 5.66E-09 | 5.826406372 | 1.321837247 |
| CNAG_03147 | - | rhomboid-like protein | 1403.196045 | 1244.3452 | 1254.066638 | 975.7053592 | 862.2646426 | 903.8711961 | 2.47E-08 | -0.525273286 | 5.74E-09 | -5.824239349 | 0.694827478 |
| CNAG_01071 | - | hypothetical protein | 537.0400939 | 613.9574687 | 684.4110968 | 386.3461268 | 402.5682583 | 343.3509766 | 2.47E-08 | -0.712514144 | 5.75E-09 | -5.82386808 | 0.610255736 |
| CNAG_01613 | - | hypothetical protein | 1203.026051 | 1228.371332 | 1087.92777 | 1647.235537 | 1671.036293 | 1571.863499 | 2.53E-08 | 0.459676863 | 5.88E-09 | 5.820072317 | 1.375233757 |
| CNAG_01647 | - | translocation protein SEC66 | 1632.363306 | 1737.770984 | 1765.92429 | 1341.549103 | 1255.675921 | 1167.009815 | 2.55E-08 | -0.463295136 | 5.94E-09 | -5.818374218 | 0.725327706 |
| CNAG_04675 | - | hypothetical protein | 20.43945701 | 18.60264667 | 16.25527504 | 82.93697019 | 88.87062146 | 59.76971944 | 2.56E-08 | 2.068491785 | 5.98E-09 | 5.817343281 | 4.194479469 |
| CNAG_06449 | - | hypothetical protein, variant 2 | 62.58285812 | 84.27943743 | 90.83116869 | 155.1463314 | 216.397438 | 202.6661212 | 2.60E-08 | 1.265036358 | 6.07E-09 | 5.814817191 | 2.403332666 |
| CNAG_00151 | - | transcription elongation regulator 1 | 2864.229704 | 2808.761416 | 2689.538365 | 2013.813663 | 2138.967539 | 2237.080136 | 2.61E-08 | -0.403547864 | 6.08E-09 | -5.814560367 | 0.755996854 |
| CNAG_02771 | - | DNA repair and recombination protein RAD54B | 1575.503858 | 1534.375643 | 1325.76775 | 1986.424707 | 2065.174491 | 2241.139162 | 2.69E-08 | 0.489826797 | 6.28E-09 | 5.809158261 | 1.404276275 |

| | | | | | | | | | | | | | |
|---|---|---|---|---|---|---|---|---|---|---|---|---|---|
| CNAG_03036 | - | potassium ion transporter | 622.9050951 | 776.9406329 | 775.4086373 | 1090.037405 | 1058.580476 | 1117.683631 | 2.69E-08 | 0.572963537 | 6.28E-09 | 5.809052183 | 1.487576164 |
| CNAG_00465 | - | hypothetical protein | 746.4548298 | 730.6324299 | 530.2696359 | 1218.087018 | 1068.990198 | 1010.099915 | 2.73E-08 | 0.701414667 | 6.38E-09 | 5.806462713 | 1.626098518 |
| CNAG_12933 | - | unspecified product | 105.885521 | 31.2718503 | 41.48265961 | 8.498349641 | 7.236996345 | 4.849801383 | 2.76E-08 | -3.240133957 | 6.45E-09 | -5.804592144 | 0.105833337 |
| CNAG_00054 | - | hypothetical protein | 18.19763458 | 23.98762695 | 30.06809741 | 83.32752672 | 82.43625602 | 97.06672478 | 2.77E-08 | 1.868917332 | 6.49E-09 | 5.80371288 | 3.652583698 |
| CNAG_07979 | - | hypothetical protein | 1271.39013 | 1242.166343 | 1223.136826 | 911.0361228 | 875.2459061 | 933.7934657 | 2.82E-08 | -0.473640115 | 6.61E-09 | -5.800583062 | 0.72014528 |
| CNAG_04193 | - | hypothetical protein | 1434.854438 | 1425.304937 | 1343.658894 | 1080.810883 | 924.3563371 | 1005.985937 | 2.84E-08 | -0.496839962 | 6.65E-09 | -5.7995423 | 0.708657305 |
| CNAG_01376 | - | stress response protein NST1, variant | 3232.072888 | 2924.116979 | 3313.34534 | 2349.311944 | 2458.997864 | 2381.455512 | 2.84E-08 | -0.413463473 | 6.66E-09 | -5.799202964 | 0.75081872 |
| CNAG_01745 | - | glycerol-3-phosphate dehydrogenase (NAD()) | 3808.445313 | 2892.838938 | 2798.34011 | 2276.963123 | 2111.275148 | 1968.58855 | 2.90E-08 | -0.596174213 | 6.81E-09 | -5.795570696 | 0.661505835 |
| CNAG_01727 | - | hsp71-like protein | 95325.6708 | 85444.67092 | 91537.8825 | 73138.80422 | 70976.51222 | 73742.56034 | 2.93E-08 | -0.337596924 | 6.88E-09 | -5.793928928 | 0.791358369 |
| CNAG_01963 | - | CMP/dCMP deaminase zinc-binding protein | 146.3966874 | 114.4952189 | 151.5928322 | 262.8450831 | 292.0868624 | 267.2489661 | 3.01E-08 | 0.981949902 | 7.06E-09 | 5.789549148 | 1.975133134 |
| CNAG_01389 | - | hypothetical protein | 403.297535 | 465.569469 | 425.6594311 | 708.0804683 | 642.8220009 | 660.3392388 | 3.03E-08 | 0.62185192 | 7.11E-09 | 5.78840773 | 1.538849262 |
| CNAG_02588 | - | avenacinase | 258.5920387 | 305.0771994 | 310.4706812 | 465.7983555 | 549.9888054 | 468.2925801 | 3.04E-08 | 0.748492397 | 7.14E-09 | 5.787605098 | 1.680036291 |
| CNAG_00195 | - | exosome complex protein LRP1 | 1062.472835 | 1120.397777 | 1135.648634 | 819.2291784 | 792.4656659 | 775.0931451 | 3.04E-08 | -0.490370429 | 7.14E-09 | -5.787655781 | 0.711842301 |
| CNAG_05926 | - | U2 small nuclear ribonucleoprotein A' | 2553.638144 | 2465.088102 | 2666.005163 | 2002.306827 | 1708.420831 | 1957.351161 | 3.04E-08 | -0.454529844 | 7.16E-09 | -5.787108232 | 0.729747949 |
| CNAG_05936 | - | DNA-directed RNA polymerase III subunit RPC1 | 7097.361758 | 6913.798685 | 6669.553129 | 5222.753461 | 5575.450162 | 5659.702332 | 3.11E-08 | -0.344871525 | 7.32E-09 | -5.783440732 | 0.787378091 |
| CNAG_01843 | - | elongation factor Ts, mitochondrial | 390.7371003 | 371.5507646 | 392.2689908 | 189.9487651 | 245.5841972 | 227.8340599 | 3.12E-08 | -0.818302282 | 7.35E-09 | -5.782839379 | 0.567108906 |
| CNAG_07801 | - | aldo-keto reductase | 2812.716091 | 2419.867621 | 2572.200937 | 2004.608945 | 1896.54514 | 1897.446337 | 3.15E-08 | -0.444604465 | 7.42E-09 | -5.781163386 | 0.734785741 |
| CNAG_05789 | - | hypothetical protein | 2058.408673 | 1800.353254 | 2069.847253 | 1456.630755 | 1494.643272 | 1353.510448 | 3.15E-08 | -0.47800136 | 7.44E-09 | -5.780778048 | 0.717971578 |
| CNAG_02597 | - | hypothetical protein | 2847.978254 | 2876.657575 | 3046.737141 | 2115.778874 | 2141.508611 | 2384.712094 | 3.22E-08 | -0.416383296 | 7.60E-09 | -5.777132826 | 0.7493007 |
| CNAG_06440 | - | DNA dependent ATPase | 1581.394978 | 1633.486739 | 1586.983221 | 2068.14027 | 2126.395146 | 2195.710375 | 3.27E-08 | 0.397491254 | 7.71E-09 | 5.774605641 | 1.317215373 |
| CNAG_01616 | - | hypothetical protein | 4319.285339 | 4218.063175 | 4374.490588 | 2904.854174 | 3428.545093 | 3418.296917 | 3.41E-08 | -0.420589964 | 8.06E-09 | -5.767248242 | 0.747119041 |
| CNAG_01194 | - | hypothetical protein | 11.1133843 | 14.2663259 | 7.721744962 | 71.57445263 | 46.82957722 | 64.91074716 | 3.45E-08 | 2.486111916 | 8.15E-09 | 5.765357953 | 5.602659887 |
| CNAG_03838 | - | hypothetical protein | 35.74385754 | 53.04075282 | 54.59355372 | 152.2898852 | 136.8722875 | 113.0265659 | 3.48E-08 | 1.483309349 | 8.22E-09 | 5.763913962 | 2.795893387 |
| CNAG_00078 | - | vacuolar protein | 1705.12394 | 1629.201211 | 1517.70401 | 2054.02592 | 2352.494946 | 2347.760343 | 3.53E-08 | 0.461857109 | 8.35E-09 | 5.761145414 | 1.377313624 |
| CNAG_02260 | CRG1 | xap5-domain-containing protein | 2158.995472 | 2281.86902 | 2143.506053 | 1638.733043 | 1757.764228 | 1645.427017 | 3.56E-08 | -0.400169053 | 8.41E-09 | -5.75994779 | 0.757769484 |
| CNAG_03618 | - | NADPH2:quinone reductase | 807.2133379 | 815.7471965 | 827.6727554 | 1127.271398 | 1143.970115 | 1174.34396 | 3.57E-08 | 0.476700178 | 8.45E-09 | 5.75922096 | 1.391557167 |
| CNAG_06957 | - | hypothetical protein | 59.06758936 | 81.03441543 | 84.43185187 | 179.0540072 | 160.7593054 | 192.648871 | 3.59E-08 | 1.230992534 | 8.49E-09 | 5.758446162 | 2.34728421 |
| CNAG_04385 | - | hypothetical protein | 0.44683804 | 7.787378998 | 5.554625618 | 53.6774883 | 46.39517519 | 36.37560866 | 3.62E-08 | 3.479192483 | 8.57E-09 | 5.75691671 | 11.15170566 |
| CNAG_06690 | - | hypothetical protein | 1226.400576 | 1290.852345 | 1146.574085 | 1720.421015 | 1694.291879 | 1620.714626 | 3.62E-08 | 0.444103299 | 8.59E-09 | 5.756507972 | 1.36046826 |
| CNAG_04168 | - | histone H1/5 | 1162.440679 | 1479.43385 | 1554.964335 | 2237.430254 | 2106.652285 | 1926.024086 | 3.76E-08 | 0.565026887 | 8.93E-09 | 5.7499053 | 1.47941508 |
| CNAG_02996 | - | flavoprotein oxygenase, variant 1 | 348.3070864 | 341.7028031 | 306.2527531 | 546.9130816 | 553.2649344 | 506.0642173 | 3.76E-08 | 0.674515579 | 8.93E-09 | 5.749982362 | 1.596060765 |
| CNAG_02045 | - | acetoacetate-CoA ligase | 116.1479395 | 133.7963849 | 116.457319 | 283.5953049 | 265.3373192 | 208.6076406 | 3.84E-08 | 1.035568835 | 9.12E-09 | 5.746245206 | 2.049921743 |
| CNAG_00031 | MLR1 | hypothetical protein | 694.3915291 | 638.8025383 | 671.7267519 | 460.3888212 | 455.1982026 | 386.6418559 | 3.86E-08 | -0.638295579 | 9.17E-09 | -5.745346641 | 0.642471526 |
| CNAG_03652 | - | dynactin 6 | 539.2600101 | 546.1352347 | 622.5918283 | 373.9338311 | 355.928985 | 362.1466221 | 3.86E-08 | -0.662448302 | 9.17E-09 | -5.745368266 | 0.631805192 |
| CNAG_00501 | - | hypothetical protein | 38.14425548 | 63.82868917 | 79.07785217 | 165.1551458 | 172.3118388 | 150.6551681 | 3.86E-08 | 1.421445751 | 9.18E-09 | 5.745134383 | 2.678537977 |
| CNAG_04862 | - | glutamate synthase (NADPH/NADH), variant | 1721.358821 | 1317.901867 | 1300.185695 | 2195.24394 | 2130.763129 | 2179.025405 | 3.95E-08 | 0.568037951 | 9.39E-09 | 5.741472307 | 1.482506008 |
| CNAG_12871 | - | unspecified product | 127.4270881 | 122.3995627 | 85.23797064 | 41.311686 | 29.62646557 | 38.12158106 | 3.96E-08 | -1.643535187 | 9.42E-09 | -5.740794921 | 0.320071209 |
| CNAG_00159 | - | hypothetical protein | 444.2451384 | 329.6660693 | 449.7660617 | 241.5673951 | 227.3664359 | 196.862689 | 3.97E-08 | -0.896730034 | 9.46E-09 | -5.740140242 | 0.537102732 |
| CNAG_03395 | - | NifU-like protein c | 2829.235817 | 2641.818975 | 2936.89243 | 2123.631878 | 2162.260939 | 2162.73739 | 3.99E-08 | -0.398432926 | 9.51E-09 | -5.739225197 | 0.758681926 |
| CNAG_06148 | - | hypothetical protein | 888.9534951 | 835.1805813 | 925.777299 | 1183.193693 | 1275.103785 | 1369.647351 | 4.00E-08 | 0.515810383 | 9.55E-09 | 5.738475272 | 1.429797052 |
| CNAG_07969 | - | hypothetical protein | 12.22368794 | 5.677270233 | 9.832538175 | 65.33455931 | 43.40390467 | 58.27757081 | 4.00E-08 | 2.65810134 | 9.55E-09 | 5.738541759 | 6.312018091 |
| CNAG_05776 | - | hypothetical protein | 1375.348648 | 1334.84431 | 1375.620305 | 956.1014312 | 1001.222818 | 1042.573973 | 4.00E-08 | -0.460856525 | 9.54E-09 | -5.738643763 | 0.726554777 |
| CNAG_05151 | - | U3 small nucleolar RNA-associated protein 11 | 1193.116423 | 1083.833205 | 1223.071897 | 762.2251388 | 840.7857334 | 854.9786419 | 4.03E-08 | -0.526376424 | 9.62E-09 | -5.737361265 | 0.69429639 |
| CNAG_01238 | - | arginine biosynthesis ArgJ, mitochondrial | 1989.73516 | 2103.04687 | 2218.093324 | 1609.989371 | 1628.346394 | 1537.758592 | 4.06E-08 | -0.417625185 | 9.69E-09 | -5.736071261 | 0.74865597 |
| CNAG_04537 | - | hypothetical protein | 13.44495133 | 14.28081178 | 11.97962228 | 81.95612931 | 55.59514432 | 55.20863326 | 4.14E-08 | 2.314764173 | 9.90E-09 | 5.732517622 | 4.975233281 |
| CNAG_03497 | - | BET3 family protein | 1564.502418 | 1429.701031 | 1642.18466 | 1142.213487 | 1152.194959 | 1043.782949 | 4.16E-08 | -0.48987329 | 9.94E-09 | -5.731692824 | 0.712087637 |
| CNAG_03398 | ZIP2 | solute carrier family 39 (zinc transporter), member 1/2/3 | 12.30028103 | 14.28482043 | 15.16402735 | 59.10914442 | 79.48931157 | 57.40599898 | 4.17E-08 | 2.234324123 | 9.98E-09 | 5.731098487 | 4.705422025 |
| CNAG_00587 | - | hypothetical protein | 33.39367541 | 54.08404401 | 34.40047153 | 94.3721217 | 132.6447546 | 155.7198326 | 4.23E-08 | 1.641540015 | 1.01E-08 | 5.728809889 | 3.119909141 |
| CNAG_01549 | - | splicing factor 3a subunit 3 | 2526.758698 | 2315.360318 | 2602.02056 | 1837.191588 | 1932.837854 | 1858.574152 | 4.24E-08 | -0.419300004 | 1.02E-08 | -5.728066427 | 0.747787363 |
| CNAG_06080 | SAC1 | inositol/phosphatidylinositol phosphatase | 4478.115094 | 4346.289954 | 4337.205265 | 3416.387577 | 3564.82376 | 3539.313198 | 4.24E-08 | -0.338659167 | 1.02E-08 | -5.727898052 | 0.790775913 |
| CNAG_05223 | - | hypothetical protein | 889.4625898 | 801.5366819 | 895.6704472 | 577.048818 | 622.5728451 | 569.7418362 | 4.38E-08 | -0.563894811 | 1.05E-08 | -5.722248505 | 0.676473436 |
| CNAG_05542 | - | hypothetical protein | 5.225325734 | 4.572280125 | 7.678141072 | 40.44226354 | 44.59453842 | 50.43282628 | 4.38E-08 | 3.050466165 | 1.05E-08 | 5.722303066 | 8.284795947 |
| CNAG_05179 | - | ubiquinol-cytochrome c reductase core subunit 2 | 1912.443423 | 1943.51775 | 1645.518911 | 1364.025481 | 1348.836581 | 1309.072601 | 4.41E-08 | -0.466888775 | 1.06E-08 | -5.721239851 | 0.723523221 |
| CNAG_03692 | - | hypothetical protein | 57.89991422 | 96.04241374 | 61.04842003 | 175.1435841 | 169.5014513 | 184.9181711 | 4.50E-08 | 1.289790792 | 1.08E-08 | 5.717496922 | 2.444925987 |
| CNAG_04205 | - | bromodomain-containing protein 8, variant | 2364.555882 | 2440.291097 | 2492.205019 | 1886.283301 | 1925.078354 | 1871.883504 | 4.59E-08 | -0.375904113 | 1.10E-08 | -5.714014428 | 0.770622325 |
| CNAG_01401 | - | hypothetical protein | 14.7373313 | 32.55653579 | 25.83604017 | 74.39696943 | 91.17900727 | 109.1344826 | 4.63E-08 | 1.935307393 | 1.11E-08 | 5.712447383 | 3.824596085 |
| CNAG_06930 | - | hypothetical protein | 2182.256852 | 1952.279057 | 2359.887374 | 1504.00681 | 1638.692311 | 1567.725529 | 4.64E-08 | -0.479449915 | 1.11E-08 | -5.712314644 | 0.717251052 |
| CNAG_03738 | - | pantetheine-phosphate adenylyltransferase | 1114.80067 | 1001.926479 | 1033.296144 | 774.0961241 | 726.0213451 | 718.4866894 | 4.64E-08 | -0.520823989 | 1.12E-08 | -5.711996628 | 0.696973646 |
| CNAG_04073 | - | hypothetical protein | 3293.978026 | 3231.121672 | 3187.555323 | 2360.949862 | 2341.607395 | 2684.441135 | 4.72E-08 | -0.410151537 | 1.14E-08 | -5.709170636 | 0.752544324 |
| CNAG_01310 | - | Nucleolar protein | 1232.885321 | 1158.174798 | 1321.174941 | 902.4854684 | 908.9418294 | 833.9599318 | 4.81E-08 | -0.50534732 | 1.16E-08 | -5.705795828 | 0.704490755 |
| CNAG_06223 | MIZ1 | E3 SUMO-protein ligase PIAS1 | 1348.534961 | 1361.76471 | 1380.951776 | 996.6371357 | 1060.800897 | 957.1672968 | 4.90E-08 | -0.455907948 | 1.18E-08 | -5.702606909 | 0.729051206 |
| CNAG_04231 | - | hypothetical protein | 177.9205307 | 182.2702504 | 165.4944059 | 299.2893047 | 310.8488062 | 347.0581333 | 4.93E-08 | 0.852508507 | 1.19E-08 | 5.701410932 | 1.805637777 |
| CNAG_04215 | MET3 | sulfate adenylyltransferase | 5684.083696 | 4656.609337 | 4653.956941 | 3471.517525 | 3850.315075 | 3620.416383 | 5.01E-08 | -0.470759557 | 1.21E-08 | -5.698435413 | 0.721584594 |
| CNAG_06001 | ERG8 | phosphomevalonate kinase | 564.6965185 | 512.7235901 | 525.626318 | 325.5875619 | 296.3234562 | 370.9479251 | 5.04E-08 | -0.70736364 | 1.22E-08 | -5.69733929 | 0.612438278 |
| CNAG_04574 | - | dolichyl-phosphate mannosyltransferase polypeptide 2, regulatory subunit | 122.0225501 | 129.5377497 | 153.719455 | 325.7993118 | 259.9623559 | 240.7397972 | 5.15E-08 | 1.013049746 | 1.24E-08 | 5.69358357 | 2.018172856 |
| CNAG_03241 | - | hypothetical protein | 1175.068231 | 1302.695134 | 1157.232029 | 1598.828237 | 1767.689794 | 1658.424971 | 5.31E-08 | 0.452136932 | 1.28E-08 | 5.688254748 | 1.368065147 |
| CNAG_03181 | - | hypothetical protein, variant | 13.381729 | 8.901574998 | 9.836027825 | 57.08065898 | 51.36555326 | 59.39041495 | 5.34E-08 | 2.460841594 | 1.29E-08 | 5.687249296 | 5.505377889 |
| CNAG_06541 | - | hypothetical protein, variant | 2116.568189 | 1287.388861 | 1323.323084 | 601.5104236 | 570.4368694 | 585.50661 | 5.54E-08 | -1.445752502 | 1.34E-08 | -5.680802531 | 0.367100631 |
| CNAG_03141 | - | hypothetical protein, variant | 62.38679303 | 55.21508552 | 56.74402838 | 143.2440982 | 137.5806545 | 146.1117556 | 5.64E-08 | 1.284633288 | 1.36E-08 | 5.677734926 | 2.436201201 |
| CNAG_05216 | RAD53 | CAMK protein kinase | 1409.777698 | 1547.274138 | 1433.424151 | 1973.278045 | 2060.061163 | 1888.284582 | 5.66E-08 | 0.417006958 | 1.37E-08 | 5.677219566 | 1.335154743 |
| CNAG_12883 | - | unspecified product | 15.84747085 | 28.2377633 | 16.26205236 | 84.42580207 | 87.90546203 | 67.45548046 | 5.66E-08 | 1.998607347 | 1.37E-08 | 5.677014633 | 3.99614061 |
| CNAG_04729 | - | ATP-dependent RNA helicase DBP7 | 1682.972001 | 1664.751126 | 1696.818476 | 2131.134681 | 2332.619398 | 2246.812169 | 5.74E-08 | 0.396718456 | 1.39E-08 | 5.674460598 | 1.316509978 |
| CNAG_02816 | - | nuclear pore complex protein Nup93 | 6243.732747 | 5893.496704 | 6082.976012 | 4658.519571 | 5104.541884 | 4650.627959 | 5.79E-08 | -0.353777414 | 1.40E-08 | -5.672827207 | 0.782532506 |
| CNAG_05614 | - | YbgI/family dinuclear metal center protein | 1430.215766 | 1319.796639 | 1489.690488 | 1076.889243 | 1008.961461 | 980.4837547 | 5.83E-08 | -0.483650732 | 1.41E-08 | -5.671595178 | 0.71516561 |
| CNAG_02001 | INP5203 | inositol-polyphosphate 5-phosphatase | 965.9549275 | 838.4261584 | 933.2534844 | 1240.832943 | 1319.102536 | 1371.90723 | 6.05E-08 | 0.506646628 | 1.47E-08 | 5.665259818 | 1.420744005 |
| CNAG_06228 | - | MIF4G/MA4 domain-containing protein | 1571.466069 | 1541.672462 | 1454.573303 | 1104.046467 | 988.1696848 | 1185.76004 | 6.14E-08 | -0.493553253 | 1.49E-08 | -5.662759993 | 0.710273589 |
| CNAG_04991 | - | hypothetical protein | 1130.645141 | 1159.422373 | 1123.082314 | 1536.658428 | 1537.278691 | 1536.307625 | 6.39E-08 | 0.418718587 | 1.55E-08 | 5.65561222 | 1.336739725 |
| CNAG_04143 | - | hypothetical protein, variant | 2.849076744 | 4.551694925 | 4.474251104 | 44.10591015 | 34.05260015 | 39.48310516 | 6.58E-08 | 3.534217161 | 1.60E-08 | 5.650614394 | 11.58524911 |
| CNAG_01769 | - | solute carrier family 25 (mitochondrial aspartate/glutamate transporter), member 12/13 | 3843.814347 | 3204.254675 | 3477.618894 | 2525.939887 | 2612.491563 | 2710.035014 | 6.65E-08 | -0.439509121 | 1.62E-08 | -5.648653582 | 0.737385462 |
| CNAG_03638 | - | heterokaryon incompatibility protein Het-C | 1959.711771 | 2011.652731 | 1852.547497 | 2540.607419 | 2666.42171 | 2478.887385 | 6.71E-08 | 0.385184408 | 1.63E-08 | 5.646837188 | 1.306026715 |
| CNAG_02159 | - | hypothetical protein | 350.8883977 | 464.4991163 | 459.7544101 | 720.5563673 | 701.6087463 | 640.4090164 | 6.71E-08 | 0.680686073 | 1.63E-08 | 5.646858358 | 1.602901833 |
| CNAG_02025 | - | DNA-directed RNA polymerase III subunit RPC3 | 1081.086028 | 1113.92056 | 1096.202972 | 826.2839743 | 799.3960134 | 679.7033138 | 6.87E-08 | -0.529228883 | 1.67E-08 | -5.642740731 | 0.692925002 |
| CNAG_03313 | - | hypothetical protein, variant | 2.932438941 | 13.1713418 | 8.768309419 | 67.04616358 | 54.10121596 | 43.06669868 | 6.94E-08 | 2.821776166 | 1.69E-08 | 5.64102832 | 7.070323203 |
| CNAG_12357 | - | unspecified product | 15.85257229 | 22.89586806 | 24.74851491 | 65.03002079 | 100.2366808 | 76.20394222 | 6.97E-08 | 1.967146047 | 1.70E-08 | 5.640275319 | 3.909938857 |
| CNAG_00677 | - | E3 ubiquitin-protein ligase UHRF1 | 1120.879595 | 1265.794278 | 1166.605159 | 895.5467466 | 857.1729958 | 775.1429521 | 7.13E-08 | -0.505595123 | 1.74E-08 | -5.636212961 | 0.704369759 |
| CNAG_00092 | - | mitochondrial protein | 2426.358011 | 2353.033229 | 2370.65225 | 1828.670826 | 1911.292913 | 1827.480193 | 7.23E-08 | -0.376536332 | 1.77E-08 | -5.633582703 | 0.770284696 |
| CNAG_01553 | - | CORD and CS domain-containing protein | 1375.457456 | 1429.64728 | 1523.819548 | 1063.589751 | 1113.411086 | 990.4876529 | 7.47E-08 | -0.466042272 | 1.82E-08 | -5.627900567 | 0.723947874 |

| | | | | | | | | | | | | | |
|---|---|---|---|---|---|---|---|---|---|---|---|---|---|
| CNAG_01089 | - | hypothetical protein | 609.0445919 | 540.746894 | 554.4293234 | 402.8735565 | 320.4848947 | 341.0882486 | 7.53E-08 | -0.695494557 | 1.84E-08 | -5.626559693 | 0.617497604 |
| CNAG_06561 | - | allantoate transporter | 12.31574702 | 17.50240184 | 14.11100732 | 62.25336953 | 51.09199816 | 92.26166109 | 7.54E-08 | 2.239666 | 1.84E-08 | 5.626287335 | 4.72287712 |
| CNAG_00051 | SNT1 | putative set3c deacetylase complex subunit | 3827.717952 | 3728.936103 | 3962.845101 | 3018.703623 | 3030.867899 | 3103.015695 | 7.55E-08 | -0.347160104 | 1.85E-08 | -5.625859558 | 0.786130046 |
| CNAG_06188 | FZC15 | hypothetical protein | 82.4559776 | 131.624655 | 127.0740475 | 227.0333297 | 250.6620827 | 248.2767976 | 7.61E-08 | 1.07847994 | 1.86E-08 | 5.624485101 | 2.111809852 |
| CNAG_04204 | - | transcription elongation factor spt4 | 1087.088973 | 1256.115159 | 1243.346444 | 805.8474102 | 864.9348075 | 886.0588091 | 7.75E-08 | -0.503769311 | 1.90E-08 | -5.621282001 | 0.705261744 |
| CNAG_05162 | - | V-type proton ATPase subunit E | 698.0181998 | 760.4452031 | 654.7336417 | 499.3310855 | 452.5417857 | 462.0535534 | 7.91E-08 | -0.59455594 | 1.94E-08 | -5.617538668 | 0.662248263 |
| CNAG_12522 | - | unspecified product | 1.763828374 | 7.84168526 | 18.32453402 | 73.9869804 | 54.79038388 | 61.66739878 | 7.93E-08 | 2.839775324 | 1.94E-08 | 5.617096984 | 7.15908557 |
| CNAG_00300 | - | gamma-tubulin complex component 2 | 2139.575354 | 2321.930659 | 2011.462656 | 2907.248015 | 2781.89583 | 2914.00386 | 7.95E-08 | 0.396002772 | 1.95E-08 | 5.616511621 | 1.315857053 |
| CNAG_05858 | - | hypothetical protein | 758.6543783 | 697.0004121 | 744.2480491 | 526.5905852 | 500.9025714 | 449.8917661 | 8.08E-08 | -0.591829057 | 1.98E-08 | -5.613529132 | 0.663501183 |
| CNAG_00547 | - | hypothetical protein | 5160.467623 | 4984.30398 | 5006.016933 | 6262.322078 | 6573.778183 | 6258.98989 | 8.16E-08 | 0.318251454 | 2.00E-08 | 5.611868314 | 1.246818489 |
| CNAG_06407 | - | cytochrome c oxidase subunit 7c | 2539.473905 | 2266.845365 | 2391.951277 | 1910.52966 | 1736.118144 | 1501.224995 | 8.18E-08 | -0.49935873 | 2.01E-08 | -5.611320845 | 0.707421156 |
| CNAG_02840 | - | hypothetical protein, variant | 91.7027765 | 135.8884449 | 84.5143146 | 200.4014861 | 236.8662702 | 254.8023963 | 8.27E-08 | 1.140888083 | 2.03E-08 | 5.60929416 | 2.205167253 |
| CNAG_04627 | - | COP9 signalosome complex subunit 12 | 1336.165156 | 1325.365305 | 1354.490315 | 1848.031692 | 1881.505647 | 1700.688044 | 8.30E-08 | 0.419866303 | 2.04E-08 | 5.608710334 | 1.337803573 |
| CNAG_13106 | - | unspecified product | 322.6236222 | 166.9043749 | 160.9752873 | 97.44754149 | 75.37675023 | 49.37743454 | 8.35E-08 | -1.570224303 | 2.05E-08 | -5.607504218 | 0.336756033 |
| CNAG_02204 | - | endonuclease G, mitochondrial | 1054.169167 | 972.8503201 | 1091.895613 | 682.0710289 | 721.7154474 | 778.3567512 | 8.43E-08 | -0.530762635 | 2.07E-08 | -5.605692718 | 0.692188734 |
| CNAG_04443 | - | hypothetical protein | 1137.708192 | 1278.986703 | 1104.985489 | 1688.542719 | 1706.431148 | 1523.059624 | 8.43E-08 | 0.467829552 | 2.07E-08 | 5.605709958 | 1.383027221 |
| CNAG_12786 | - | unspecified product | 28.59327181 | 29.3491655 | 18.40777759 | 114.5993237 | 76.21546133 | 81.82238533 | 8.45E-08 | 1.838785102 | 2.08E-08 | 5.605160788 | 3.577086738 |
| CNAG_06218 | - | Amidase | 831.237935 | 867.2015819 | 894.616105 | 609.8046859 | 633.7910214 | 577.5128567 | 8.47E-08 | -0.526162781 | 2.09E-08 | -5.604774561 | 0.694399214 |
| CNAG_00700 | - | phosphoribosylaminoimidazolecarboxamide formyltransferase/IMP cyclohydrolase | 3568.419067 | 3361.53487 | 3524.546304 | 2765.782516 | 2812.669432 | 2672.318368 | 8.56E-08 | -0.357228826 | 2.11E-08 | -5.60287085 | 0.780662662 |
| CNAG_00428 | - | hypothetical protein | 29.81102844 | 39.01433657 | 26.92195835 | 92.53880448 | 113.8123658 | 92.90678923 | 8.59E-08 | 1.645294531 | 2.12E-08 | 5.602156937 | 3.128117123 |
| CNAG_07479 | - | hypothetical protein | 31.10079665 | 60.52987016 | 48.21695555 | 136.4442908 | 121.1750419 | 133.8581686 | 8.59E-08 | 1.475191371 | 2.12E-08 | 5.602056105 | 2.780205205 |
| CNAG_01948 | FZC36 | nuclear protein | 1007.527227 | 1036.351674 | 994.9144778 | 731.2334061 | 748.5052131 | 721.7937722 | 8.60E-08 | -0.480284379 | 2.12E-08 | -5.601702841 | 0.71683631 |
| CNAG_04967 | - | hypothetical protein | 72.89649231 | 67.05721557 | 67.40538215 | 169.0225756 | 176.6123188 | 138.5891674 | 8.66E-08 | 1.213694008 | 2.14E-08 | 5.600470068 | 2.319307337 |
| CNAG_02165 | - | cytoplasmic protein | 299.2296354 | 252.3355015 | 263.5827412 | 433.9056075 | 444.5054881 | 488.1077013 | 8.76E-08 | 0.730105348 | 2.16E-08 | 5.598385977 | 1.658760212 |
| CNAG_04772 | - | hypothetical protein | 782.7263366 | 790.983041 | 867.106024 | 1171.741149 | 1146.564063 | 1127.768729 | 8.98E-08 | 0.482477606 | 2.22E-08 | 5.593927464 | 1.397140981 |
| CNAG_05381 | ITR3C | myo-inositol transporter, putative | 13.43755063 | 17.48352046 | 9.853871266 | 61.64294233 | 69.80383238 | 56.27765494 | 9.01E-08 | 2.239297246 | 2.23E-08 | 5.593385169 | 4.721670104 |
| CNAG_06648 | RTF1 | RNA polymerase-associated protein RTF1 | 3237.849514 | 2921.939088 | 3024.38657 | 2195.143359 | 2377.911182 | 2436.91652 | 9.03E-08 | -0.405496618 | 2.24E-08 | -5.592742199 | 0.754976363 |
| CNAG_00010 | - | cation transporter | 1100.864148 | 1089.143771 | 1018.394756 | 817.7372337 | 742.408739 | 727.3768783 | 9.12E-08 | -0.503586308 | 2.26E-08 | -5.591049127 | 0.70535121 |
| CNAG_00352 | - | A1 cistron-splicing factor AAR2 | 833.3689829 | 790.6987577 | 743.2485099 | 542.8430076 | 537.1149277 | 554.1536099 | 9.17E-08 | -0.550248679 | 2.27E-08 | -5.589850289 | 0.682902405 |
| CNAG_06093 | SFB2 | copII vesicle coat protein | 5084.113546 | 4736.327975 | 4738.196301 | 3649.824012 | 3857.21336 | 3966.70674 | 9.17E-08 | -0.359214964 | 2.27E-08 | -5.589873491 | 0.779588674 |
| CNAG_01385 | GGT1 | Glycosylinositolphosphoceramide galactosyltransferase | 511.7360128 | 505.443409 | 400.1285735 | 721.8726628 | 783.5957889 | 719.1922894 | 9.31E-08 | 0.635519357 | 2.31E-08 | 5.587183098 | 1.553496894 |
| CNAG_13131 | - | unspecified product | 25.11146342 | 27.19512403 | 10.9585872 | 101.0830556 | 104.4819107 | 60.98884776 | 9.31E-08 | 2.080755145 | 2.31E-08 | 5.587050004 | 4.230285827 |
| CNAG_01866 | - | hypothetical protein | 99.82733416 | 119.7902033 | 116.4211957 | 228.7570706 | 241.1982169 | 206.2766675 | 9.32E-08 | 1.00014215 | 2.31E-08 | 5.58678018 | 2.000197072 |
| CNAG_04256 | LPI6 | hypothetical protein | 355.4924221 | 460.1578934 | 390.7424479 | 597.2371106 | 674.9094494 | 656.9450729 | 9.36E-08 | 0.658248434 | 2.32E-08 | 5.586046588 | 1.57816542 |
| CNAG_04843 | - | atypical/ABC1/ABC1-B protein kinase | 1068.823188 | 1133.576962 | 1180.643385 | 1521.849633 | 1511.376997 | 1595.094195 | 9.38E-08 | 0.437605392 | 2.33E-08 | 5.585514283 | 1.354354482 |
| CNAG_00117 | ERG24 | c-14 sterol reductase | 3346.636514 | 3376.600314 | 3657.80947 | 2492.449985 | 2761.777099 | 2751.097581 | 9.43E-08 | -0.390240055 | 2.34E-08 | -5.584446076 | 0.763002635 |
| CNAG_04915 | - | hypothetical protein | 5.174461179 | 2.395239349 | 2.3397226 | 28.37698407 | 43.94057871 | 45.92985879 | 9.43E-08 | 3.671585855 | 2.34E-08 | 5.584465687 | 12.74258308 |
| CNAG_06341 | - | hypothetical protein | 444.6552635 | 463.1390303 | 505.2893886 | 287.4576013 | 267.0164409 | 318.7898467 | 9.48E-08 | -0.710344179 | 2.36E-08 | -5.58345292 | 0.611174316 |
| CNAG_04454 | - | hypothetical protein | 2.860040545 | 5.630568156 | 4.479499656 | 33.23217013 | 56.41896175 | 33.04986099 | 9.49E-08 | 3.456469597 | 2.36E-08 | 5.583214593 | 10.97743889 |
| CNAG_08025 | - | hypothetical protein | 1440.785174 | 1339.209342 | 1663.457428 | 1016.797123 | 1057.274599 | 1067.037301 | 9.60E-08 | -0.516224602 | 2.39E-08 | -5.580993202 | 0.69919918 |
| CNAG_04067 | - | Haloacid dehalogenase, type II | 266.5763533 | 271.6548822 | 191.1448697 | 422.366268 | 440.3104708 | 427.132721 | 9.87E-08 | 0.809369357 | 2.46E-08 | 5.576219752 | 1.752445231 |
| CNAG_02437 | - | rRNA biogenesis protein RRP5 | 1017.768897 | 868.327809 | 903.1848749 | 573.8235867 | 693.3455862 | 610.8177212 | 1.02E-07 | -0.587664766 | 2.53E-08 | -5.571116217 | 0.665419124 |
| CNAG_03588 | LYS2 | L-aminoadipate-semialdehyde dehydrogenase | 1814.368026 | 1920.857823 | 1591.121798 | 1277.599997 | 1371.333075 | 1127.086866 | 1.02E-07 | -0.511354406 | 2.53E-08 | -5.57119868 | 0.701563499 |
| CNAG_03113 | - | trehalose synthase | 1341.027741 | 1299.605335 | 850.2778794 | 2706.379773 | 2579.601366 | 2367.672758 | 1.03E-07 | 1.117639792 | 2.57E-08 | 5.568253562 | 2.169916897 |
| CNAG_03792 | - | tRNA adenylyltransferase | 813.2709204 | 647.4617467 | 672.8387179 | 475.9115496 | 433.5317542 | 465.3743354 | 1.04E-07 | -0.650894074 | 2.60E-08 | -5.566717665 | 0.636885498 |
| CNAG_03796 | ENV7 | NAK protein kinase | 1617.059823 | 1567.564843 | 1535.623288 | 1167.870394 | 1235.027172 | 1154.747504 | 1.05E-07 | -0.423655499 | 2.62E-08 | -5.56483343 | 0.745533198 |
| CNAG_04849 | - | vacuolar protein | 437.1514431 | 517.2646489 | 421.4275227 | 792.2684403 | 723.1217508 | 651.5628047 | 1.06E-07 | 0.641091907 | 2.64E-08 | 5.563626571 | 1.55950903 |
| CNAG_05121 | - | hypothetical protein | 2784.795084 | 2479.138982 | 2797.187143 | 1929.756753 | 2102.768091 | 2057.27323 | 1.06E-07 | -0.420708829 | 2.65E-08 | -5.563196816 | 0.747057488 |
| CNAG_01239 | CDA3 | Chitin deacetylase | 546.7767094 | 518.399256 | 510.9702466 | 826.341184 | 766.1802349 | 745.8820642 | 1.06E-07 | 0.554835298 | 2.65E-08 | 5.562832929 | 1.469000919 |
| CNAG_07382 | - | hypothetical protein | 3347.746946 | 3339.950953 | 3312.333645 | 2455.116737 | 2749.731867 | 2641.201841 | 1.08E-07 | -0.365208336 | 2.69E-08 | -5.560322757 | 0.776356755 |
| CNAG_01621 | - | hypothetical protein | 3950.021905 | 2791.353939 | 2248.031827 | 1236.658594 | 1084.514519 | 1107.291959 | 1.15E-07 | -1.40775775 | 2.87E-08 | -5.549201639 | 0.376897009 |
| CNAG_02492 | - | peptidyl-prolyl cis-trans isomerase-like 3 | 1220.11168 | 1215.259842 | 1386.209395 | 963.2616105 | 832.9216617 | 909.4022502 | 1.16E-07 | -0.514126428 | 2.91E-08 | -5.546830008 | 0.700216796 |
| CNAG_02061 | - | hypothetical protein | 269.3957583 | 292.8157587 | 287.7466748 | 150.376787 | 177.4488509 | 133.5853566 | 1.17E-07 | -0.897701731 | 2.94E-08 | -5.544725382 | 0.5367411 |
| CNAG_06764 | - | Short-chain dehydrogenase | 3508.166815 | 3645.039769 | 3670.802212 | 4562.586924 | 4538.079792 | 4524.90295 | 1.18E-07 | 0.316935692 | 2.96E-08 | 5.543847726 | 1.245681888 |
| CNAG_00639 | URO3 | hypothetical protein | 833.152266 | 766.9328128 | 651.5601384 | 493.812464 | 381.649826 | 511.9537408 | 1.18E-07 | -0.715015375 | 2.97E-08 | -5.543341609 | 0.609198639 |
| CNAG_01729 | - | ATP-dependent RNA helicase DHX37/DHR1 | 2832.760122 | 2715.071435 | 2871.866387 | 2160.999523 | 2251.137312 | 2203.807152 | 1.19E-07 | -0.363467671 | 2.98E-08 | -5.542501348 | 0.777294024 |
| CNAG_02856 | - | potassium ion transporter | 1296.494663 | 1455.671231 | 1205.242094 | 1803.616612 | 1830.576043 | 1811.619154 | 1.19E-07 | 0.446017804 | 2.99E-08 | 5.541864489 | 1.362274846 |
| CNAG_04646 | - | hypothetical protein | 544.6900919 | 628.3143424 | 746.5763561 | 1044.300874 | 951.6231882 | 973.4063511 | 1.20E-07 | 0.614958913 | 3.03E-08 | 5.539933039 | 1.53151438 |
| CNAG_03391 | - | mitochondrial inner membrane protein | 1388.076665 | 1269.123878 | 1273.2643 | 953.8422229 | 1009.047493 | 865.0633717 | 1.21E-07 | -0.491033352 | 3.04E-08 | -5.539373036 | 0.711515282 |
| CNAG_02164 | - | hypothetical protein | 310.9258507 | 288.9304966 | 276.3829792 | 508.1412317 | 454.8065374 | 461.6182061 | 1.21E-07 | 0.687874138 | 3.05E-08 | 5.538550922 | 1.610908039 |
| CNAG_02552 | - | transketolase, variant 2 | 927.9789916 | 877.9830638 | 860.5368288 | 555.1485437 | 667.4660212 | 615.2244402 | 1.22E-07 | -0.551818428 | 3.07E-08 | -5.537376072 | 0.682159766 |
| CNAG_02405 | - | hypothetical protein | 142.9580044 | 155.343113 | 159.0579001 | 278.4451718 | 282.4512165 | 275.0431339 | 1.24E-07 | 0.857699185 | 3.13E-08 | 5.534259247 | 1.812145989 |
| CNAG_06800 | - | hypothetical protein | 1143.596652 | 961.9694387 | 807.2862263 | 703.3529068 | 537.8401036 | 514.3505805 | 1.24E-07 | -0.746577951 | 3.13E-08 | -5.534027478 | 0.596015622 |
| CNAG_05783 | - | transcription initiation factor TFIIB | 4922.246438 | 4723.569555 | 4812.984563 | 6168.230044 | 5920.728323 | 6123.51864 | 1.24E-07 | 0.317670266 | 3.13E-08 | 5.533919778 | 1.246316311 |
| CNAG_00507 | - | nitrilase | 12.27119124 | 9.987913127 | 16.20880633 | 65.81760407 | 53.90027254 | 61.67041735 | 1.25E-07 | 2.258527387 | 3.16E-08 | 5.532462609 | 4.785028067 |
| CNAG_07942 | - | cell division control protein 25 | 1197.966796 | 1374.607077 | 1227.411487 | 938.3013269 | 871.776516 | 944.8980258 | 1.27E-07 | -0.478721422 | 3.20E-08 | -5.530009815 | 0.717613322 |
| CNAG_03967 | - | F-actin capping protein | 2880.627593 | 2790.478002 | 2840.956492 | 2312.092346 | 2174.316795 | 2210.475803 | 1.30E-07 | -0.361203537 | 3.27E-08 | -5.526388005 | 0.77851485 |
| CNAG_06758 | - | efflux protein | 231.5452382 | 171.5907227 | 216.6494109 | 424.3520101 | 333.1135899 | 382.6897906 | 1.34E-07 | 0.865956313 | 3.39E-08 | 5.520226037 | 1.822547371 |
| CNAG_02355 | - | solute carrier family 35 (UDP-xylose/UDP-N- acetylglucosamine transporter), member B4 | 187.2972642 | 228.5275923 | 137.8446634 | 362.6371839 | 342.787898 | 354.9215824 | 1.37E-07 | 0.924656227 | 3.47E-08 | 5.515980819 | 1.898231867 |
| CNAG_07406 | - | pheromone alpha | 21.59095248 | 23.95145489 | 13.0703103 | 89.400827 | 80.03264644 | 59.77264273 | 1.38E-07 | 1.985153929 | 3.49E-08 | 5.51496365 | 3.959049016 |
| CNAG_05134 | - | ornithine-oxo-acid transaminase | 3698.518623 | 4014.573586 | 3688.961661 | 4684.697454 | 4886.578489 | 4958.933946 | 1.38E-07 | 0.334982673 | 3.49E-08 | 5.51498049 | 1.261362259 |
| CNAG_02885 | CAP64 | capsular associated protein | 2006.087182 | 2180.597968 | 2142.411361 | 1687.108271 | 1622.292812 | 1561.067814 | 1.41E-07 | -0.39277654 | 3.58E-08 | -5.510608201 | 0.761662335 |
| CNAG_04699 | - | microfibrillar-associated protein 1 | 324.7813222 | 207.1496353 | 180.5089965 | 440.2370858 | 455.8048657 | 520.1612289 | 1.42E-07 | 0.976216598 | 3.59E-08 | 5.509674966 | 1.967299485 |
| CNAG_07996 | - | hypothetical protein | 771.5568416 | 721.7894724 | 786.8964867 | 510.1200797 | 500.8588414 | 553.0109901 | 1.44E-07 | -0.558902324 | 3.66E-08 | -5.506637006 | 0.678818447 |
| CNAG_05544 | - | hypothetical protein | 48.64152135 | 62.79773254 | 103.559812 | 160.8024269 | 196.4565893 | 199.284384 | 1.45E-07 | 1.366365341 | 3.68E-08 | 5.505716906 | 2.57820207 |
| CNAG_04442 | - | hypothetical protein, variant | 1.649700862 | 6.684367476 | 0.165579422 | 37.57563311 | 36.80852064 | 39.43110729 | 1.48E-07 | 3.988232746 | 3.76E-08 | 5.501684039 | 15.87002775 |
| CNAG_03150 | - | solute carrier family 25 (mitochondrial folate transporter), member 32 | 899.0067514 | 925.4140597 | 1032.150817 | 670.5029157 | 665.6522192 | 682.9238111 | 1.50E-07 | -0.517031661 | 3.80E-08 | -5.499742757 | 0.69880815 |
| CNAG_03920 | IDP1 | isocitrate dehydrogenase, NADP-dependent | 7730.454197 | 8040.815077 | 8995.390769 | 6461.078472 | 6588.0857 | 6482.369963 | 1.50E-07 | -0.357823376 | 3.81E-08 | -5.499276364 | 0.780341009 |
| CNAG_01335 | - | hypothetical protein | 237.4426987 | 247.9675778 | 193.2563895 | 449.1932543 | 435.2241924 | 338.5385892 | 1.52E-07 | 0.836909952 | 3.86E-08 | 5.497344548 | 1.786220212 |
| CNAG_02100 | FAS2 | fatty acid synthase subunit alpha, fungi type | 20722.54345 | 19023.05048 | 17069.6485 | 14382.47391 | 15163.67922 | 14589.88749 | 1.53E-07 | -0.379920365 | 3.87E-08 | -5.496574891 | 0.768480009 |
| CNAG_01048 | - | hypothetical protein, variant | 41.56400722 | 57.35128855 | 65.23001879 | 160.0150198 | 137.6221813 | 124.0974993 | 1.56E-07 | 1.354335782 | 3.95E-08 | 5.493047172 | 2.556793742 |
| CNAG_12368 | - | unspecified product | 89.17251167 | 58.50169297 | 78.06092549 | 202.765331 | 192.1114188 | 145.3106196 | 1.59E-07 | 1.243420733 | 4.04E-08 | 5.488965106 | 2.367592403 |
| CNAG_05569 | - | hypothetical protein | 338.3340327 | 403.7100456 | 329.4301039 | 199.4368463 | 185.8987504 | 229.9913795 | 1.62E-07 | -0.817791344 | 4.13E-08 | -5.485169282 | 0.567309786 |
| CNAG_04206 | - | hypothetical protein | 438.3193741 | 503.2764507 | 450.1953176 | 802.417579 | 713.6061711 | 644.9174484 | 1.63E-07 | 0.619524567 | 4.16E-08 | 5.484129755 | 1.536368796 |
| CNAG_01445 | APG9 | autophagy-like protein 9 | 446.45234 | 489.2750836 | 454.4482502 | 643.0811269 | 734.3910852 | 718.0248925 | 1.64E-07 | 0.577969015 | 4.18E-08 | 5.482903597 | 1.492746323 |
| CNAG_01506 | - | hypothetical protein | 10.00471657 | 12.15199928 | 22.58626643 | 63.0462541 | 55.50115172 | 87.92547617 | 1.68E-07 | 2.205228602 | 4.27E-08 | 5.479208198 | 4.611476039 |
| CNAG_01147 | - | arp2/3 complex 20 kDa subunit | 1592.552717 | 1528.792877 | 1545.210961 | 1179.575499 | 1203.96441 | 1165.834453 | 1.69E-07 | -0.410721712 | 4.31E-08 | -5.477637349 | 0.752246966 |

| | | | | | | | | | | | | | |
|---|---|---|---|---|---|---|---|---|---|---|---|---|---|
| CNAG_02666 | - | hypothetical protein | 627.4728668 | 681.0727932 | 728.4690782 | 995.8663294 | 949.8698648 | 983.4049651 | 1.75E-07 | 0.508693126 | 4.46E-08 | 5.471788821 | 1.422760795 |
| CNAG_03875 | - | galactose-1-phosphate uridylyltransferase | 2120.270085 | 1942.523031 | 1861.983724 | 1525.953731 | 1490.309514 | 1455.598567 | 1.75E-07 | -0.421603183 | 4.47E-08 | -5.471221079 | 0.746594516 |
| CNAG_02753 | LIV13 | endoplasmic reticulum protein | 1361.812689 | 1375.973231 | 1157.266036 | 1767.005156 | 1923.832483 | 1739.50018 | 1.75E-07 | 0.464640794 | 4.47E-08 | 5.471076597 | 1.379973721 |
| CNAG_06473 | - | hypothetical protein | 1081.560629 | 1096.919829 | 947.1639129 | 1421.45923 | 1477.817903 | 1430.839609 | 1.77E-07 | 0.455226307 | 4.52E-08 | 5.469210359 | 1.370997848 |
| CNAG_07823 | - | solute carrier family 35 (UDP-galactose transporter), member B1 | 11.25866555 | 31.49431853 | 33.26271915 | 89.59734239 | 100.8155742 | 89.49962268 | 1.77E-07 | 1.876678099 | 4.52E-08 | 5.469230177 | 3.672285181 |
| CNAG_06208 | - | heat shock 70kDa protein 4 | 14698.11873 | 12973.30545 | 13910.71408 | 8736.502993 | 11691.97093 | 9867.30719 | 1.81E-07 | -0.473138485 | 4.63E-08 | -5.464844805 | 0.72039572 |
| CNAG_06311 | - | cytoplasmic protein, variant | 671.808372 | 760.7560763 | 683.7274488 | 1024.649095 | 1039.670636 | 955.7311125 | 1.84E-07 | 0.498811986 | 4.70E-08 | 5.462307788 | 1.413049481 |
| CNAG_06324 | - | zinc finger protein | 651.3216895 | 630.1966029 | 698.3564455 | 416.64668 | 447.4162409 | 468.652267 | 1.85E-07 | -0.587701785 | 4.72E-08 | -5.461437249 | 0.66540205 |
| CNAG_06568 | SKS1 | RAN protein kinase | 113.8351888 | 157.4259513 | 98.37509155 | 272.5361371 | 267.9252204 | 225.1807892 | 1.87E-07 | 1.040016796 | 4.80E-08 | 5.458709887 | 2.056251592 |
| CNAG_07640 | - | hypothetical protein | 548.3715838 | 486.8947908 | 519.2221923 | 322.4883764 | 362.9425617 | 306.6876488 | 1.87E-07 | -0.665493758 | 4.80E-08 | -5.458741739 | 0.63047289 |
| CNAG_03052 | PTC2 | PP2Cc protein phosphatase | 2797.5701 | 2497.425071 | 2554.084776 | 2126.114583 | 1871.505828 | 1901.894021 | 1.88E-07 | -0.427748634 | 4.80E-08 | -5.458403011 | 0.74342101 |
| CNAG_07407 | MFalpha3 | fungal mating-type pheromone | 47.27436542 | 56.23693948 | 41.84100222 | 132.530088 | 118.5882684 | 122.849135 | 1.88E-07 | 1.354822401 | 4.81E-08 | 5.458290214 | 2.557656291 |
| CNAG_05564 | - | peroxin-10 | 548.0396308 | 564.7265035 | 613.2997879 | 818.3096942 | 880.1378856 | 819.1356441 | 1.88E-07 | 0.52976408 | 4.82E-08 | 5.457841517 | 1.443693093 |
| CNAG_03660 | - | lipoate-protein ligase A | 1316.799641 | 1121.558976 | 1226.312639 | 833.1046999 | 916.7276889 | 867.2236513 | 1.88E-07 | -0.501693526 | 4.83E-08 | -5.45755782 | 0.706277222 |
| CNAG_05473 | - | hypothetical protein | 758.6390813 | 693.76589 | 713.3493826 | 483.6561085 | 481.030976 | 520.8233804 | 1.90E-07 | -0.559706126 | 4.88E-08 | -5.45565271 | 0.678440347 |
| CNAG_04764 | - | Dihydrofolate reductase | 77.73415547 | 107.9439563 | 113.1990313 | 209.2859128 | 224.835982 | 195.1375138 | 1.95E-07 | 1.065692707 | 5.00E-08 | 5.451148252 | 2.093174672 |
| CNAG_01095 | - | BEM46 family protein | 711.4883254 | 630.5107535 | 799.90908 | 1125.208976 | 1020.601633 | 1043.368973 | 1.96E-07 | 0.559198643 | 5.02E-08 | 5.450522996 | 1.47345055 |
| CNAG_13007 | - | unspecified product | 57.89905965 | 86.39033048 | 80.17620322 | 152.1935958 | 171.2474977 | 195.9232549 | 1.97E-07 | 1.201959017 | 5.07E-08 | 5.448943274 | 2.300518435 |
| CNAG_12629 | - | unspecified product | 29.81185706 | 36.87481678 | 33.2883727 | 83.72567038 | 108.5370048 | 103.8554578 | 1.98E-07 | 1.570185957 | 5.09E-08 | 5.448214948 | 2.969429864 |
| CNAG_02226 | - | hypothetical protein | 306.2293043 | 292.1384616 | 206.0817425 | 498.854252 | 456.5978685 | 459.3562593 | 2.03E-07 | 0.798479412 | 5.22E-08 | 5.443713158 | 1.739266988 |
| CNAG_12817 | - | unspecified product | 34.4382797 | 31.52969913 | 39.65812174 | 98.06709114 | 108.484752 | 95.13299366 | 2.07E-07 | 1.517402297 | 5.33E-08 | 5.439971459 | 2.862751211 |
| CNAG_00089 | - | exosome complex component RRP4 | 1688.392075 | 1738.867584 | 1711.576936 | 1281.498514 | 1363.564567 | 1309.032791 | 2.12E-07 | -0.393166492 | 5.46E-08 | -5.435783797 | 0.76145649 |
| CNAG_05013 | HRB1 | RNP domain-containing protein | 10188.52716 | 10420.85605 | 10642.06192 | 8227.00184 | 8894.633447 | 8562.786236 | 2.14E-07 | -0.2983185 | 5.51E-08 | -5.433896152 | 0.81319965 |
| CNAG_07556 | - | hypothetical protein | 763.5348529 | 810.0951306 | 839.1548608 | 580.266583 | 551.7780877 | 566.3699048 | 2.14E-07 | -0.522279436 | 5.52E-08 | -5.433538892 | 0.696270866 |
| CNAG_02062 | - | glycoside hydrolase family 2 | 638.9774056 | 563.669018 | 574.9568559 | 817.4397322 | 879.2130289 | 921.1578425 | 2.15E-07 | 0.5444532 | 5.55E-08 | 5.43265793 | 1.458467461 |
| CNAG_04556 | - | translocation protein Sec62 | 36.69766883 | 21.87063026 | 31.15809327 | 111.9009896 | 76.10259764 | 102.7075925 | 2.18E-07 | 1.69917224 | 5.61E-08 | 5.430715527 | 3.24714597 |
| CNAG_01560 | - | hypothetical protein | 18.24842193 | 33.65858758 | 40.70452587 | 99.79923983 | 96.27013472 | 99.45293204 | 2.19E-07 | 1.677749971 | 5.66E-08 | 5.42911754 | 3.199286007 |
| CNAG_04679 | - | histone deacetylase complex subunit | 2304.197033 | 2274.570482 | 2372.930381 | 3003.708296 | 2965.641925 | 2941.811101 | 2.20E-07 | 0.342905419 | 5.69E-08 | 5.428335186 | 1.268308248 |
| CNAG_04393 | - | RNA helicase | 3427.077463 | 3317.317417 | 3150.258701 | 2387.384012 | 2704.014127 | 2592.358026 | 2.24E-07 | -0.380301542 | 5.78E-08 | -5.42557037 | 0.768276994 |
| CNAG_01142 | - | endoplasmic reticulum protein | 1600.098677 | 1750.89892 | 1621.112628 | 2191.968769 | 2135.847925 | 2195.74568 | 2.26E-07 | 0.376768842 | 5.85E-08 | 5.423263809 | 1.298430543 |
| CNAG_00891 | - | ATP-binding cassette, subfamily F, member 2 | 10519.92285 | 9200.15317 | 9357.013371 | 7493.8495 | 7957.867972 | 7418.25779 | 2.29E-07 | -0.362453503 | 5.91E-08 | -5.421378694 | 0.777840629 |
| CNAG_13191 | - | unspecified product | 27.2977453 | 6.791255725 | 13.06352696 | 62.84358802 | 88.96362587 | 74.98525306 | 2.31E-07 | 2.264542521 | 5.98E-08 | 5.419475512 | 4.805020284 |
| CNAG_00664 | - | hypothetical protein | 14.72829843 | 32.54734057 | 15.21539598 | 133.2855441 | 54.28869586 | 86.15205463 | 2.32E-07 | 2.150680823 | 6.00E-08 | 5.418761022 | 4.440372854 |
| CNAG_04275 | - | Metalloendopeptidase | 794.5662832 | 739.9638233 | 568.4226843 | 484.556396 | 411.9514908 | 398.8513434 | 2.34E-07 | -0.715563455 | 6.06E-08 | -5.416996364 | 0.608967249 |
| CNAG_01581 | - | hypothetical protein | 37.93250197 | 41.18410156 | 35.43096077 | 111.5034364 | 92.59217174 | 114.9291075 | 2.37E-07 | 1.473625255 | 6.14E-08 | 5.414634918 | 2.777188794 |
| CNAG_05843 | - | translation initiation factor 4E | 430.1161367 | 491.4116725 | 438.4558256 | 694.7176645 | 691.2016918 | 650.4026398 | 2.37E-07 | 0.567725303 | 6.15E-08 | 5.414345055 | 1.482184767 |
| CNAG_05685 | - | neutral amino acid transporter | 6918.822276 | 7491.266452 | 6911.64177 | 5615.068179 | 5826.459256 | 5921.696018 | 2.38E-07 | -0.311091267 | 6.17E-08 | -5.413691412 | 0.806031839 |
| CNAG_03325 | BCH1 | ChAPs family protein | 1527.196192 | 1513.697271 | 1521.743107 | 1196.766315 | 1115.062014 | 1154.72104 | 2.38E-07 | -0.4117132 | 6.18E-08 | -5.413500066 | 0.751730164 |
| CNAG_00734 | - | dihydroorotase, homodimeric type, variant | 593.9471165 | 496.6552567 | 619.3980016 | 373.1508679 | 355.0634181 | 364.3630714 | 2.42E-07 | -0.6613835 | 6.27E-08 | -5.410947367 | 0.632271677 |
| CNAG_01114 | - | hypothetical protein | 558.0762298 | 663.4892031 | 670.6102035 | 423.7730887 | 361.0249353 | 436.464906 | 2.43E-07 | -0.646680561 | 6.32E-08 | -5.409503988 | 0.638748295 |
| CNAG_04539 | - | mitochondrial protein required for respiration | 992.3270633 | 937.2957219 | 1049.240443 | 728.9364438 | 681.1539491 | 721.7726405 | 2.44E-07 | -0.498973985 | 6.34E-08 | -5.409023919 | 0.70760984 |
| CNAG_04767 | - | hypothetical protein | 71.82423464 | 99.2792386 | 67.43636607 | 177.3995178 | 199.0603313 | 160.7474818 | 2.46E-07 | 1.163191726 | 6.40E-08 | 5.407370629 | 2.239523377 |
| CNAG_05938 | - | hypothetical protein | 34.37769739 | 34.71372091 | 15.22614183 | 86.29637223 | 92.83865088 | 105.945901 | 2.47E-07 | 1.75511495 | 6.42E-08 | 5.406605899 | 3.375532127 |
| CNAG_01127 | - | hypothetical protein | 1027.945171 | 1079.707907 | 1118.794234 | 1573.427788 | 1393.148466 | 1470.805317 | 2.47E-07 | 0.44540329 | 6.44E-08 | 5.406269574 | 1.36169471 |
| CNAG_03292 | - | hypothetical protein, variant | 27.30722682 | 14.31226408 | 10.9400859 | 81.41653255 | 56.25818909 | 88.0768822 | 2.48E-07 | 2.125146727 | 6.45E-08 | 5.405852023 | 4.362474588 |
| CNAG_12197 | - | unspecified product | 24.01526982 | 40.06442106 | 22.66554515 | 87.87443665 | 113.9270267 | 84.08330962 | 2.48E-07 | 1.709717764 | 6.45E-08 | 5.405716269 | 3.270968269 |
| CNAG_04046 | - | hypothetical protein | 833.5531107 | 837.0587215 | 885.0205181 | 596.5504718 | 626.8871647 | 600.7887591 | 2.50E-07 | -0.503148583 | 6.52E-08 | -5.403950499 | 0.705565252 |
| CNAG_07843 | - | hypothetical protein | 0.308519602 | 0.147976978 | 0.127604829 | 19.09914892 | 29.21939365 | 27.24629781 | 2.65E-07 | 7.071083416 | 6.90E-08 | 5.393818592 | 134.4646761 |
| CNAG_03806 | - | acyltransferase, variant | 667.2498657 | 804.9556374 | 834.0450382 | 1072.771351 | 1154.407173 | 1119.953573 | 2.68E-07 | 0.522503582 | 6.98E-08 | 5.391611709 | 1.436445824 |
| CNAG_07955 | - | hypothetical protein | 625.8795378 | 818.5825222 | 763.4152291 | 500.8372797 | 435.2271083 | 500.871706 | 2.70E-07 | -0.634381631 | 7.05E-08 | -5.389998578 | 0.64421688 |
| CNAG_07815 | - | hypothetical protein | 302.9319368 | 378.2888728 | 341.4084475 | 576.5887647 | 507.3723496 | 544.8462245 | 2.71E-07 | 0.657937594 | 7.06E-08 | 5.389607786 | 1.577825429 |
| CNAG_05982 | - | MFS transporter, SP family, general alpha glucoside:H symporter | 38.11631681 | 73.45083219 | 64.18786033 | 152.6715582 | 137.521548 | 157.1546816 | 2.75E-07 | 1.336464213 | 7.17E-08 | 5.386835717 | 2.5253165 |
| CNAG_05512 | - | hypothetical protein | 2421.652399 | 2273.316439 | 2302.406754 | 1803.775585 | 1836.225952 | 1837.449297 | 2.76E-07 | -0.368908538 | 7.20E-08 | -5.386166736 | 0.774368119 |
| CNAG_06421 | - | acetolactate synthase, small subunit | 581.5078976 | 730.2789006 | 733.526884 | 461.9087999 | 448.2571825 | 436.5329018 | 2.79E-07 | -0.618278617 | 7.29E-08 | -5.383996033 | 0.651447753 |
| CNAG_03356 | - | hypothetical protein | 961.9656631 | 990.0123867 | 933.0664234 | 666.5755768 | 745.1132098 | 657.4341133 | 2.81E-07 | -0.495079158 | 7.35E-08 | -5.382471152 | 0.709522747 |
| CNAG_00174 | - | hypothetical protein | 271.2114186 | 254.4429901 | 225.2149204 | 408.2999748 | 428.2080607 | 404.9844451 | 2.82E-07 | 0.709981455 | 7.37E-08 | 5.381968052 | 1.63578309 |
| CNAG_05641 | - | hypothetical protein, variant | 2.81572764 | 4.532816386 | 0.158696761 | 27.74569835 | 42.31567951 | 35.05332237 | 2.85E-07 | 4.101565535 | 7.46E-08 | 5.379754356 | 17.16699399 |
| CNAG_07791 | - | hypothetical protein | 172.24223 | 208.1377942 | 237.9267716 | 355.3940554 | 342.6819775 | 403.6672946 | 2.91E-07 | 0.81953068 | 7.61E-08 | 5.376212889 | 1.764831786 |
| CNAG_01984 | TAL1 | transaldolase | 23401.57913 | 23411.44833 | 22074.49206 | 19392.46674 | 19282.42652 | 18867.30838 | 3.00E-07 | -0.274821075 | 7.85E-08 | -5.370578346 | 0.826552822 |
| CNAG_06936 | - | Beta-glucosidase | 328.535741 | 330.951746 | 355.2516469 | 506.9798505 | 505.6503992 | 610.1692248 | 3.03E-07 | 0.663183233 | 7.95E-08 | 5.368361819 | 1.583572845 |
| CNAG_02940 | - | calcium/proton exchanger | 1302.330309 | 1380.287603 | 1371.545472 | 1849.597397 | 1773.592387 | 1745.059913 | 3.05E-07 | 0.390090324 | 7.99E-08 | 5.367442779 | 1.310475447 |
| CNAG_06005 | - | U3 small nucleolar RNA-associated protein 4 | 703.7213691 | 632.3539689 | 653.6254487 | 429.1334655 | 484.5831357 | 429.8702023 | 3.13E-07 | -0.582749158 | 8.20E-08 | -5.362585895 | 0.667690233 |
| CNAG_05537 | - | DNA repair protein rad18 | 1037.928518 | 1095.605265 | 1036.513191 | 784.1905678 | 773.5005515 | 782.8267462 | 3.17E-07 | -0.452651208 | 8.32E-08 | -5.360047086 | 0.730698825 |
| CNAG_00183 | - | alternative cyclin Pcl12, variant | 1011.491677 | 541.8276771 | 567.2127962 | 225.5039349 | 223.6562551 | 216.9789525 | 3.23E-07 | -1.692229882 | 8.48E-08 | -5.356710029 | 0.309448261 |
| CNAG_03754 | ISP1 | short-chain dehydrogenase/reductase SDR | 5.161746024 | 3.469637815 | 3.403711874 | 37.57881071 | 38.60690765 | 34.04540619 | 3.27E-07 | 3.292715378 | 8.59E-08 | 5.354212433 | 9.79954916 |
| CNAG_00288 | - | oligoribonuclease | 1162.771492 | 1112.885255 | 1185.757225 | 878.4599252 | 841.6687954 | 670.8606676 | 3.33E-07 | -0.549669467 | 8.75E-08 | -5.350882745 | 0.683176632 |
| CNAG_07997 | - | hypothetical protein | 36.88018329 | 59.45212846 | 49.27387262 | 116.5801871 | 125.5778614 | 134.9315045 | 3.35E-07 | 1.367076829 | 8.80E-08 | 5.34985725 | 2.579473864 |
| CNAG_04492 | - | hypothetical protein | 67.13985998 | 78.86829411 | 78.0395995 | 168.9169387 | 176.5441193 | 147.4377902 | 3.38E-07 | 1.123638285 | 8.88E-08 | 5.348307314 | 2.178957844 |
| CNAG_12363 | - | unspecified product | 21.62461941 | 21.82898974 | 27.92980479 | 90.80499337 | 72.86740958 | 74.03623705 | 3.39E-07 | 1.758822236 | 8.92E-08 | 5.347534155 | 3.384217368 |
| CNAG_05844 | - | mitochondrial import inner membrane translocase subunit TIM8 | 818.3303431 | 804.7353323 | 876.4634845 | 610.6681872 | 500.7773311 | 597.4293173 | 3.40E-07 | -0.564459994 | 8.96E-08 | -5.346560668 | 0.676208477 |
| CNAG_06345 | MDM10 | mitochondrial distribution and morphology protein 10 | 583.758706 | 679.6670627 | 680.2310617 | 425.2707628 | 466.4581803 | 406.5769494 | 3.42E-07 | -0.597270293 | 9.02E-08 | -5.345522419 | 0.66100345 |
| CNAG_00549 | - | hypothetical protein, variant | 74.34036654 | 113.3805686 | 154.7321544 | 245.8549992 | 259.2969892 | 251.6306939 | 3.44E-07 | 1.132118665 | 9.06E-08 | 5.344629003 | 2.191803807 |
| CNAG_03133 | ATG2602 | UDP-glucose,sterol transferase | 429.0300163 | 521.5891969 | 475.7788622 | 684.3640436 | 692.8085496 | 856.6004521 | 3.46E-07 | 0.63316359 | 9.11E-08 | 5.343578122 | 1.550962269 |
| CNAG_02372 | - | enoyl reductase | 1738.627354 | 1795.979317 | 1832.065006 | 1451.31917 | 1322.944602 | 1332.366618 | 3.50E-07 | -0.401305837 | 9.22E-08 | -5.341417808 | 0.757172628 |
| CNAG_05074 | - | hypothetical protein | 1663.092495 | 1714.273773 | 1515.56252 | 2107.059175 | 2243.764215 | 2139.148379 | 3.51E-07 | 0.392502701 | 9.25E-08 | 5.340777235 | 1.312668569 |
| CNAG_06536 | - | monocarboxylic acid transporter | 567.2575354 | 617.1627386 | 551.271295 | 377.7846708 | 404.3202231 | 385.4406126 | 3.56E-07 | -0.588260436 | 9.39E-08 | -5.33805394 | 0.665144437 |
| CNAG_06362 | - | hypothetical protein | 2307.64392 | 2277.777269 | 1935.768218 | 2898.646185 | 2869.906399 | 2976.161418 | 3.63E-07 | 0.408427178 | 9.59E-08 | 5.334238134 | 1.327238074 |
| CNAG_00790 | DPP1 | PAP2 domain-containing protein | 643.1009311 | 655.9845417 | 617.3779203 | 479.9800352 | 401.6328518 | 397.7009992 | 3.70E-07 | -0.599640511 | 9.79E-08 | -5.330618025 | 0.659918373 |
| CNAG_06865 | - | translation machinery-associated protein 16 | 1060.068593 | 1035.318221 | 1104.713173 | 692.9369853 | 824.4660281 | 776.1704426 | 3.76E-07 | -0.496549748 | 9.94E-08 | -5.327773513 | 0.708799874 |
| CNAG_06994 | - | hypothetical protein | 490.0777564 | 564.2500465 | 440.3784396 | 357.7967024 | 259.2012672 | 263.4419194 | 3.79E-07 | -0.779510754 | 1.00E-07 | -5.326201877 | 0.582564318 |
| CNAG_03606 | - | translation initiation factor aIF-2 | 9060.368751 | 8510.591828 | 9070.097836 | 6488.97168 | 7417.058236 | 7242.797361 | 3.90E-07 | -0.348822409 | 1.03E-07 | -5.321176697 | 0.785224771 |
| CNAG_04826 | - | Homocysteine S-methyltransferase | 32.07756432 | 30.4340106 | 31.15141721 | 96.72926251 | 83.1686067 | 96.10605742 | 3.98E-07 | 1.548947264 | 1.05E-07 | 5.317037753 | 2.926035482 |
| CNAG_00601 | CEL1 | glycosyl hydrolase | 48.44419028 | 57.31683876 | 46.09375182 | 143.6139744 | 117.6758998 | 118.4870495 | 3.99E-07 | 1.309342619 | 1.06E-07 | 5.316824325 | 2.478285883 |
| CNAG_05580 | - | hypothetical protein | 8.824906427 | 17.48831146 | 18.33768764 | 67.28655671 | 61.79225576 | 59.56223432 | 4.17E-07 | 2.106709442 | 1.10E-07 | 5.308646299 | 4.307077975 |
| CNAG_01378 | STE24 | STE24 endopeptidase | 1478.016282 | 1326.253429 | 1368.179845 | 1074.558353 | 1034.869875 | 992.6819843 | 4.21E-07 | -0.444178105 | 1.12E-07 | -5.30678598 | 0.735002925 |
| CNAG_01835 | - | hypothetical protein | 1332.735751 | 1497.712523 | 1412.086217 | 1759.042519 | 1985.872972 | 1999.184667 | 4.21E-07 | 0.42261075 | 1.12E-07 | 5.306636047 | 1.340350906 |
| CNAG_02332 | - | hypothetical protein | 1467.663531 | 1497.520115 | 1453.503676 | 1119.645552 | 1116.819112 | 1151.367812 | 4.27E-07 | -0.398584865 | 1.13E-07 | -5.304185363 | 0.758602029 |

| | | | | | | | | | | | | | |
|---|---|---|---|---|---|---|---|---|---|---|---|---|---|
| CNAG_03009 | ERG28 | putative ER membrane protein | 527.4781676 | 483.6856114 | 566.0890313 | 366.239731 | 324.8629336 | 329.9645571 | 4.30E-07 | -0.64476589 | 1.14E-07 | -5.302642186 | 0.639596571 |
| CNAG_06052 | GAL7 | galactose-1-phosphate uridylyltransferase | 152.3119191 | 173.6433088 | 162.2787825 | 335.6092737 | 320.503835 | 247.5086973 | 4.32E-07 | 0.874173666 | 1.15E-07 | 5.301684098 | 1.83295792 |
| CNAG_06316 | - | glycine cleavage system H protein | 668.341443 | 706.9443337 | 780.7204068 | 1129.142645 | 980.8781606 | 1007.874938 | 4.35E-07 | 0.517588934 | 1.16E-07 | 5.30047777 | 1.431560789 |
| CNAG_04327 | - | hypothetical protein | 5949.449294 | 5767.409532 | 5442.080294 | 4601.000987 | 4798.322013 | 4383.084389 | 4.53E-07 | -0.331533631 | 1.20E-07 | -5.292802834 | 0.794691252 |
| CNAG_01703 | - | hypothetical protein | 73.15568846 | 125.1731919 | 134.5088935 | 252.3087196 | 233.3136196 | 227.2835956 | 4.58E-07 | 1.086824 | 1.22E-07 | 5.290886975 | 2.124059235 |
| CNAG_02006 | - | protein N-terminal amidase | 1305.11822 | 1177.534259 | 1119.728206 | 829.2229503 | 903.7853828 | 879.4145102 | 4.71E-07 | -0.47942962 | 1.25E-07 | -5.285438304 | 0.717261142 |
| CNAG_01454 | STE12alpha | transcription factor STE12 | 2679.033819 | 2677.517047 | 2718.46485 | 3312.57812 | 3484.934627 | 3413.621095 | 4.72E-07 | 0.323174708 | 1.26E-07 | 5.284854669 | 1.251080575 |
| CNAG_04413 | - | hypothetical protein | 7.539158302 | 5.647536987 | 8.738508111 | 36.31137812 | 43.69125659 | 52.6004628 | 4.72E-07 | 2.695706881 | 1.26E-07 | 5.284855758 | 6.478711347 |
| CNAG_07552 | - | DNA repair protein Rad8 | 3287.388007 | 3236.693812 | 2883.824666 | 3896.269616 | 4351.028037 | 4087.479752 | 4.73E-07 | 0.375690044 | 1.26E-07 | 5.284590187 | 1.297459984 |
| CNAG_03856 | - | hypothetical protein | 666.0815395 | 837.2525124 | 691.2299753 | 1350.538577 | 1139.725373 | 954.7311368 | 4.73E-07 | 0.635835811 | 1.26E-07 | 5.284502938 | 1.553837689 |
| CNAG_07349 | - | hypothetical protein | 1019.192522 | 1033.127305 | 1001.31237 | 745.2659881 | 790.8264794 | 678.5504248 | 4.79E-07 | -0.479567255 | 1.28E-07 | -5.282190356 | 0.717192718 |
| CNAG_04529 | - | cytochrome c oxidase-assembly factor COX16 | 483.3566994 | 581.6025401 | 608.7226647 | 388.050998 | 304.069302 | 355.4779804 | 4.84E-07 | -0.689740892 | 1.29E-07 | -5.280125933 | 0.619965186 |
| CNAG_06241 | CFO1 | acidic laccase | 1076.153606 | 839.309991 | 1021.504928 | 686.8687219 | 622.448721 | 684.0419548 | 5.05E-07 | -0.57586506 | 1.35E-07 | -5.272023307 | 0.670883859 |
| CNAG_12024 | - | unspecified product | 540.3821752 | 548.2669934 | 560.8105788 | 325.5108409 | 385.3541373 | 380.9631199 | 5.05E-07 | -0.61167076 | 1.35E-07 | -5.272039463 | 0.654438369 |
| CNAG_07736 | - | glucan endo-1,3-alpha-glucosidase agn1 | 227.2966468 | 217.4901147 | 250.3677594 | 128.6531754 | 117.7698711 | 129.044172 | 5.14E-07 | -0.907319564 | 1.37E-07 | -5.268915312 | 0.533174777 |
| CNAG_01604 | - | TIGR01458 family HAD hydrolase, variant | 592.7554579 | 623.565951 | 496.9226502 | 415.3828094 | 306.6507696 | 331.1163216 | 5.14E-07 | -0.716291286 | 1.37E-07 | -5.268804055 | 0.608660106 |
| CNAG_01900 | NDX1 | hypothetical protein | 1370.642486 | 1332.672181 | 1269.020797 | 944.4411754 | 980.5213244 | 1042.554736 | 5.18E-07 | -0.436215608 | 1.39E-07 | -5.267042464 | 0.739070755 |
| CNAG_05220 | TLK1 | phosphatidylinositol 3-kinase | 122.1026599 | 177.9366478 | 167.595223 | 340.3554518 | 246.7727428 | 324.8485911 | 5.23E-07 | 0.951911501 | 1.40E-07 | 5.265477857 | 1.934433993 |
| CNAG_02134 | RSC8 | rsc chromatin remodeling complex subunit | 3309.340889 | 3650.22382 | 3621.581299 | 2721.369314 | 2678.904085 | 2939.788579 | 5.34E-07 | -0.358307424 | 1.43E-07 | -5.261507535 | 0.780079235 |
| CNAG_04478 | - | hypothetical protein | 2883.436097 | 2987.815901 | 3115.168726 | 3765.595968 | 3859.278291 | 3738.918149 | 5.38E-07 | 0.323407098 | 1.44E-07 | 5.260063908 | 1.251282115 |
| CNAG_07332 | - | hypothetical protein | 526.9577876 | 520.546017 | 541.8640538 | 748.2671401 | 754.9599202 | 781.3323289 | 5.40E-07 | 0.509358498 | 1.45E-07 | 5.259193135 | 1.423417125 |
| CNAG_03316 | RDI1 | rho gdp-dissociation inhibitor | 856.2479029 | 941.7483031 | 798.9198998 | 1203.584877 | 1183.622303 | 1214.331725 | 5.41E-07 | 0.457577243 | 1.45E-07 | 5.2588613 | 1.373233772 |
| CNAG_04997 | - | cytoplasmic protein | 1674.369888 | 1659.167726 | 1747.809619 | 1260.478888 | 1373.076646 | 1259.097932 | 5.43E-07 | -0.399787758 | 1.46E-07 | -5.257985228 | 0.757969784 |
| CNAG_00081 | - | pre-mRNA-processing factor 6 | 4961.557114 | 4860.213809 | 4682.752436 | 3644.366093 | 3990.082916 | 4003.337226 | 5.46E-07 | -0.332949224 | 1.47E-07 | -5.256886251 | 0.793911872 |
| CNAG_06637 | - | ubiquitin carboxyl-terminal hydrolase 22/27/51 | 2049.076934 | 1814.353778 | 1936.594601 | 1493.254288 | 1471.33616 | 1472.227575 | 5.47E-07 | -0.402311976 | 1.47E-07 | -5.256597587 | 0.756644758 |
| CNAG_03167 | CHK1 | CAMK/CAMKL/CHK1 protein kinase | 1292.518047 | 1338.055175 | 1316.982841 | 957.6961566 | 1039.241186 | 991.532284 | 5.50E-07 | -0.416874893 | 1.48E-07 | -5.255340596 | 0.74904542 |
| CNAG_04571 | - | dimethyladenosine transferase | 789.7062692 | 804.9721647 | 845.7834625 | 1100.005261 | 1094.756594 | 1176.541971 | 5.58E-07 | 0.451907096 | 1.50E-07 | 5.252644281 | 1.367847217 |
| CNAG_00453 | - | mitochondrial protein | 4793.891471 | 2456.486389 | 2963.467503 | 1139.213273 | 1102.591216 | 1129.524592 | 5.64E-07 | -1.617752164 | 1.52E-07 | -5.250719254 | 0.325842758 |
| CNAG_05386 | PRO1 | glutamate 5-kinase, variant | 470.3287798 | 543.830956 | 446.7735479 | 310.0676888 | 330.1301309 | 296.6771092 | 5.68E-07 | -0.655031772 | 1.53E-07 | -5.249145181 | 0.635061505 |
| CNAG_03924 | - | nit protein 1 | 288.0125873 | 300.3665393 | 286.7121241 | 194.2749953 | 156.6112179 | 131.3906902 | 5.88E-07 | -0.876718544 | 1.58E-07 | -5.242924664 | 0.544604744 |
| CNAG_07772 | - | hypothetical protein | 95.11224983 | 112.2306867 | 99.37508475 | 206.1695437 | 186.5786987 | 216.0674768 | 5.88E-07 | 0.976083403 | 1.58E-07 | 5.24269092 | 1.967117865 |
| CNAG_03289 | - | methionine aminopeptidase, type II | 4986.151945 | 4885.027 | 5181.808197 | 4021.937561 | 4235.934145 | 3962.310774 | 5.88E-07 | -0.316458737 | 1.58E-07 | -5.242732761 | 0.803038612 |
| CNAG_07621 | - | hypothetical protein | 169.8040856 | 181.2025582 | 191.0402507 | 346.3635996 | 342.9144254 | 273.0189313 | 5.90E-07 | 0.813584995 | 1.59E-07 | 5.241940186 | 1.757573466 |
| CNAG_03927 | - | hypothetical protein | 987.7061703 | 1039.57713 | 983.1894089 | 756.2043731 | 699.272869 | 752.8388268 | 5.94E-07 | -0.462090863 | 1.60E-07 | -5.240541355 | 0.725933418 |
| CNAG_07566 | - | hypothetical protein | 181.3102208 | 165.0356757 | 139.9244356 | 286.1693293 | 297.9867277 | 288.3664735 | 6.01E-07 | 0.829283628 | 1.62E-07 | 5.238269692 | 1.77680287 |
| CNAG_04006 | - | hypothetical protein | 10.01182669 | 17.50931524 | 24.70819253 | 71.90771742 | 64.30848384 | 68.37415395 | 6.15E-07 | 1.978421617 | 1.66E-07 | 5.234029313 | 3.94061722 |
| CNAG_04781 | - | hypothetical protein, variant 2 | 34.51858016 | 43.35229065 | 50.30548975 | 94.51276758 | 119.6244527 | 130.3957875 | 6.17E-07 | 1.418597545 | 1.67E-07 | 5.233250648 | 2.673255154 |
| CNAG_05030 | - | zinc finger protein 830, variant | 988.9203243 | 1050.365889 | 1084.444746 | 686.7356648 | 827.9517262 | 716.2667586 | 6.17E-07 | -0.501377922 | 1.67E-07 | -5.233277794 | 0.706431744 |
| CNAG_02185 | - | nuclear protein | 953.5736689 | 775.719568 | 889.291929 | 592.6294859 | 598.3619493 | 621.8629085 | 6.17E-07 | -0.547221907 | 1.66E-07 | -5.23338402 | 0.684336638 |
| CNAG_00902 | - | hypothetical protein | 360.5130187 | 348.9685736 | 401.8325001 | 231.4670395 | 238.6709058 | 209.0079917 | 6.25E-07 | -0.725544314 | 1.69E-07 | -5.230693012 | 0.604768829 |
| CNAG_07647 | - | voltage-gated chloride channel protein | 1524.188057 | 1667.93311 | 1360.956037 | 1908.461444 | 2175.643893 | 2257.803123 | 6.35E-07 | 0.463656018 | 1.72E-07 | 5.227544553 | 1.37903208 |
| CNAG_02561 | - | spermine transporter | 648.2189292 | 419.0923633 | 474.4566897 | 256.8973208 | 263.5180026 | 339.8709364 | 6.35E-07 | -0.861593013 | 1.72E-07 | -5.227458255 | 0.550344536 |
| CNAG_01097 | - | short-chain dehydrogenase/reductase SDR | 1536.598024 | 1677.405955 | 1536.628405 | 1282.438955 | 1155.600537 | 1146.987812 | 6.47E-07 | -0.420931826 | 1.75E-07 | -5.223905756 | 0.746942024 |
| CNAG_03108 | - | hypothetical protein | 439.4945902 | 495.7489429 | 508.7967639 | 727.3823059 | 666.0389155 | 750.1981315 | 6.48E-07 | 0.556838225 | 1.75E-07 | 5.223620549 | 1.471041783 |
| CNAG_03103 | - | Cullin 3 | 207.0379311 | 180.1547314 | 200.6437636 | 318.7159388 | 324.597399 | 368.1743616 | 6.62E-07 | 0.767330345 | 1.79E-07 | 5.219546718 | 1.702117162 |
| CNAG_05067 | CLR5 | hypothetical protein | 12.28852652 | 21.75500159 | 6.66381673 | 65.69399944 | 48.52845986 | 75.8500669 | 6.62E-07 | 2.257029073 | 1.79E-07 | 5.219576903 | 4.780061146 |
| CNAG_07542 | - | hypothetical protein | 126.5408384 | 136.9939213 | 106.8685288 | 230.9679363 | 235.0143897 | 232.8300083 | 6.63E-07 | 0.906661015 | 1.80E-07 | 5.219039889 | 1.874701649 |
| CNAG_01870 | - | electron transfer flavoprotein beta subunit | 2537.324521 | 2565.263957 | 2424.002979 | 1965.671635 | 1959.555725 | 2053.900759 | 6.67E-07 | -0.347540183 | 1.81E-07 | -5.217858035 | 0.785922967 |
| CNAG_07544 | - | sterol esterase | 23.99675717 | 32.56511256 | 31.14385572 | 72.72912438 | 106.969856 | 90.5975535 | 6.73E-07 | 1.625061068 | 1.83E-07 | 5.216325629 | 3.084552214 |
| CNAG_04606 | - | hypothetical protein | 79.77628827 | 67.03640444 | 42.94913948 | 151.7496588 | 140.069447 | 170.4141408 | 6.75E-07 | 1.27889856 | 1.83E-07 | 5.215544961 | 2.426536498 |
| CNAG_07360 | - | hypothetical protein | 219.9788951 | 211.3981816 | 221.9939472 | 343.4371199 | 518.467134 | 341.8308201 | 6.87E-07 | 0.868592999 | 1.86E-07 | 5.212328605 | 1.825881325 |
| CNAG_05110 | - | small subunit ribosomal protein S13 | 942.021346 | 864.0166888 | 911.70384 | 703.3658875 | 596.5898743 | 618.5738874 | 6.93E-07 | -0.518677201 | 1.88E-07 | -5.210414289 | 0.698011542 |
| CNAG_02053 | - | hypothetical protein | 13.46395775 | 13.21954081 | 22.58077689 | 58.23277053 | 76.77729954 | 60.70083647 | 6.96E-07 | 1.998947653 | 1.89E-07 | 5.209617877 | 3.997083338 |
| CNAG_07392 | - | hypothetical protein | 363.8165835 | 344.5915568 | 323.0266498 | 178.3204624 | 230.0949156 | 203.4172878 | 7.13E-07 | -0.768970467 | 1.94E-07 | -5.204963537 | 0.586836102 |
| CNAG_03464 | LAC2 | Laccase 2 | 100.8902962 | 104.7107721 | 95.11736007 | 214.1137247 | 210.9700482 | 174.1479718 | 7.22E-07 | 0.984779483 | 1.96E-07 | 5.202727243 | 1.979010797 |
| CNAG_04141 | - | vacuolar protein | 1704.55226 | 1541.730034 | 1523.905453 | 1142.131874 | 1220.348449 | 1229.082296 | 7.27E-07 | -0.424621375 | 1.98E-07 | -5.20137243 | 0.745034235 |
| CNAG_03964 | - | zinc knuckle family protein | 1708.26475 | 1658.121446 | 2026.05748 | 1204.308712 | 1390.299972 | 1315.709319 | 7.28E-07 | -0.479764196 | 1.98E-07 | -5.201035121 | 0.717094821 |
| CNAG_00929 | - | hypothetical protein | 999.2362328 | 935.1074512 | 924.5346058 | 701.7009038 | 627.6359553 | 719.5193527 | 7.28E-07 | -0.496774972 | 1.98E-07 | -5.200739072 | 0.708689229 |
| CNAG_05106 | - | hypothetical protein | 278.6939982 | 275.6487099 | 295.2003248 | 177.0413638 | 168.7660423 | 132.4858419 | 7.28E-07 | -0.845631094 | 1.98E-07 | -5.200767754 | 0.556467334 |
| CNAG_05840 | - | hypothetical protein | 800.686956 | 770.2211095 | 712.3282495 | 551.4867272 | 553.569341 | 482.0743251 | 7.31E-07 | -0.54034141 | 1.99E-07 | -5.199866715 | 0.687608169 |
| CNAG_06388 | - | hypothetical protein | 1849.919743 | 276.1098427 | 577.8797455 | 37.62700995 | 66.46927765 | 42.82347801 | 7.33E-07 | -4.243900139 | 2.00E-07 | -5.199367337 | 0.052778709 |
| CNAG_01885 | CID14 | DNA polymerase sigma subunit | 1730.301253 | 1612.838568 | 1606.0152 | 1284.689617 | 1204.761786 | 1289.026698 | 7.36E-07 | -0.405003421 | 2.01E-07 | -5.198424872 | 0.755234502 |
| CNAG_05459 | - | pi-transporter A-1 | 858.0067617 | 828.443148 | 891.4082066 | 552.0769949 | 680.4674277 | 543.1329185 | 7.39E-07 | -0.553667871 | 2.02E-07 | -5.197701051 | 0.681285841 |
| CNAG_07998 | - | hypothetical protein | 44.86313352 | 43.33391029 | 25.88035812 | 137.0245151 | 89.92517945 | 104.0064116 | 7.43E-07 | 1.544428283 | 2.03E-07 | 5.196650446 | 2.916884544 |
| CNAG_04784 | - | Monosaccharide transporter | 32.09859966 | 35.79477888 | 32.22014863 | 82.19906185 | 96.29334688 | 104.9046724 | 7.51E-07 | 1.495312405 | 2.05E-07 | 5.194585002 | 2.819251932 |
| CNAG_03242 | - | solute carrier family 25 (peroxisomal adenine nucleotide transporter), member 17 | 188.3247875 | 158.6077769 | 159.0938142 | 282.9092901 | 348.2332051 | 282.8986835 | 7.55E-07 | 0.83666119 | 2.06E-07 | 5.193562966 | 1.785912242 |
| CNAG_06693 | - | hypothetical protein | 189.8583022 | 305.0302603 | 277.422362 | 401.8186771 | 472.2261224 | 520.1903534 | 7.66E-07 | 0.839879394 | 2.10E-07 | 5.190635614 | 1.789900504 |
| CNAG_04000 | - | cytochrome c oxidase assembly protein subunit 11 | 532.140498 | 504.1247086 | 536.2790473 | 324.0157796 | 334.3727445 | 379.8139069 | 7.68E-07 | -0.615551982 | 2.10E-07 | -5.190166099 | 0.652680127 |
| CNAG_03027 | - | hypothetical protein | 3278.860462 | 3253.766098 | 3182.244906 | 2617.142206 | 2571.983299 | 2673.370729 | 7.75E-07 | -0.320398746 | 2.12E-07 | -5.188249802 | 0.800848501 |
| CNAG_07694 | - | nicotinate (nicotinamide) nucleotide adenylyltransferase, variant | 610.6252114 | 723.8280461 | 696.2500283 | 474.3979297 | 485.4235828 | 416.5866761 | 7.84E-07 | -0.576116167 | 2.15E-07 | -5.186005237 | 0.670767099 |
| CNAG_00871 | CLR3 | hypothetical protein, variant | 65.83143011 | 63.7824252 | 36.55273062 | 159.2144212 | 119.2744807 | 142.7830864 | 7.95E-07 | 1.342127096 | 2.18E-07 | 5.183427641 | 2.53524838 |
| CNAG_04384 | - | peptidyl-tRNA hydrolase | 789.2340128 | 812.2819384 | 924.4195761 | 570.8372189 | 640.7412698 | 532.0320211 | 7.98E-07 | -0.551092565 | 2.19E-07 | -5.182682372 | 0.682503067 |
| CNAG_01501 | - | hypothetical protein | 407.2413225 | 443.6781704 | 423.2301166 | 295.4621416 | 267.9805321 | 205.7634691 | 8.02E-07 | -0.74603893 | 2.20E-07 | -5.18150896 | 0.596238347 |
| CNAG_04712 | - | solute carrier family 25 (mitochondrial dicarboxylate transporter), member 10 | 1428.387484 | 1253.233644 | 1493.099876 | 1935.984948 | 1938.410464 | 1823.884426 | 8.24E-07 | 0.432532708 | 2.26E-07 | 5.176410664 | 1.349600776 |
| CNAG_02167 | - | vacuolar protein sorting-associated protein 27 | 110.3258493 | 127.3408286 | 134.5275523 | 273.4428628 | 202.8288812 | 246.105874 | 8.32E-07 | 0.944015039 | 2.29E-07 | 5.174504192 | 1.923874964 |
| CNAG_12039 | - | unspecified product | 7.663536046 | 15.35059596 | 20.45383749 | 77.87580589 | 52.05263243 | 59.56163841 | 8.37E-07 | 2.148357172 | 2.30E-07 | 5.173308489 | 4.433226803 |
| CNAG_12625 | - | unspecified product | 352.0239856 | 305.8382446 | 312.3230534 | 222.3392153 | 152.2021188 | 165.744599 | 8.43E-07 | -0.861789724 | 2.32E-07 | -5.172034968 | 0.550269502 |
| CNAG_00675 | - | carnitine/acyl carnitine carrier, variant | 2055.176361 | 2483.276003 | 2169.100409 | 1697.926085 | 1580.810555 | 1770.818745 | 8.50E-07 | -0.424410329 | 2.34E-07 | -5.170455481 | 0.745143231 |
| CNAG_06417 | - | AP-2 complex subunit sigma | 285.7185989 | 298.2310364 | 279.2779281 | 166.0058486 | 173.9370994 | 171.2149407 | 8.58E-07 | -0.772231666 | 2.36E-07 | -5.168447871 | 0.585511062 |
| CNAG_02962 | - | CMGC/DYRK/PRP4 protein kinase | 2622.64596 | 2775.358088 | 2663.939326 | 2094.850539 | 2180.410038 | 2192.678328 | 8.72E-07 | -0.332898654 | 2.40E-07 | -5.165365895 | 0.793939701 |
| CNAG_13128 | - | unspecified product | 2930.247119 | 2393.904117 | 3085.937292 | 1447.811979 | 853.2568552 | 1262.630331 | 8.85E-07 | -1.254786284 | 2.44E-07 | -5.162615168 | 0.41905564 |
| CNAG_04882 | - | hypothetical protein | 1138.825581 | 1177.736382 | 1175.320903 | 1566.25234 | 1588.203815 | 1473.095976 | 8.87E-07 | 0.391634941 | 2.44E-07 | 5.162138794 | 1.311879255 |
| CNAG_06112 | - | carbamoyl-phosphate synthase arginine-specific large chain | 7719.6794 | 7299.505016 | 6564.012307 | 5827.61596 | 5816.109495 | 5357.823146 | 8.94E-07 | -0.359753925 | 2.46E-07 | -5.160534147 | 0.77929749 |
| CNAG_06390 | - | hypothetical protein | 106.592919 | 72.47813144 | 79.14533142 | 207.2752042 | 155.406742 | 203.7908165 | 9.05E-07 | 1.120422734 | 2.50E-07 | 5.157997074 | 2.174106682 |
| CNAG_05139 | UGT1 | UDP-galactose transporter | 598.2117863 | 660.5560864 | 589.8832405 | 905.6115846 | 883.5056191 | 832.5026632 | 9.05E-07 | 0.489538764 | 2.50E-07 | 5.158000395 | 1.40399594 |
| CNAG_05691 | - | hypothetical protein | 1035.601926 | 1022.404872 | 1130.293482 | 816.9477574 | 777.816693 | 740.6885297 | 9.05E-07 | -0.465451797 | 2.50E-07 | -5.158006901 | 0.724244236 |
| CNAG_04896 | - | tyrosine-tRNA ligase | 1950.400931 | 1989.045707 | 2068.988163 | 2457.981062 | 2644.813631 | 2648.680928 | 9.09E-07 | 0.352118747 | 2.51E-07 | 5.157092437 | 1.276433827 |

| | | | | | | | | | | | | | |
|---|---|---|---|---|---|---|---|---|---|---|---|---|---|
| CNAG_05292 | TPS1 | alpha,alpha-trehalose-phosphate synthase (UDP- forming) | 6220.293112 | 5805.121178 | 5356.780763 | 4513.00326 | 4754.310079 | 4547.362706 | 9.10E-07 | -0.346897187 | 2.51E-07 | -5.156766788 | 0.786273324 |
| CNAG_05239 | - | cytoplasmic protein | 352.2910567 | 311.3156429 | 411.3641367 | 228.4057357 | 205.8054412 | 199.0168019 | 9.18E-07 | -0.78095177 | 2.53E-07 | -5.155142286 | 0.581982723 |
| CNAG_05116 | - | DNA polymerase mu subunit | 442.9901195 | 536.6434388 | 473.6390638 | 698.4746955 | 734.3392751 | 703.6598543 | 9.26E-07 | 0.542643886 | 2.56E-07 | 5.1533073 | 1.456639513 |
| CNAG_02714 | - | elongation factor 1-beta | 42531.70758 | 39949.01753 | 45849.90638 | 32884.76416 | 36358.58482 | 33775.68125 | 9.33E-07 | -0.332815657 | 2.58E-07 | -5.151913738 | 0.793985378 |
| CNAG_12040 | - | unspecified product | 30.9202907 | 32.57123605 | 32.20902435 | 90.33478635 | 85.82133197 | 93.90476316 | 9.61E-07 | 1.496554988 | 2.66E-07 | 5.146256693 | 2.821681181 |
| CNAG_07898 | - | hypothetical protein | 57.5536694 | 30.48671437 | 33.32170294 | 123.3064159 | 100.3850753 | 117.2092038 | 9.72E-07 | 1.485653174 | 2.69E-07 | 5.143951939 | 2.800439331 |
| CNAG_04289 | - | hypothetical protein | 3671.328916 | 3963.768495 | 4035.354912 | 2834.126529 | 3023.09417 | 3286.14277 | 9.74E-07 | -0.366836475 | 2.70E-07 | -5.143581949 | 0.7754811 |
| CNAG_00044 | - | ribosome assembly protein 1 | 3918.799633 | 3903.459583 | 3883.959936 | 2992.186399 | 3114.542348 | 3318.342892 | 9.76E-07 | -0.327782215 | 2.70E-07 | -5.143005939 | 0.796760364 |
| CNAG_06527 | - | MFS transporter, SP family, general alpha glucoside:H symporter, variant 2 | 20.44024555 | 28.22486993 | 13.07012268 | 61.92699602 | 90.63464974 | 73.95528345 | 9.85E-07 | 1.862522813 | 2.73E-07 | 5.141337554 | 3.636430019 |
| CNAG_07596 | - | hypothetical protein | 25.2015742 | 44.36810839 | 35.41278668 | 98.90804177 | 103.2420259 | 94.02000051 | 9.87E-07 | 1.490373802 | 2.74E-07 | 5.14082309 | 2.809617628 |
| CNAG_01650 | MRPL7 | large subunit ribosomal protein L7 | 221.2178937 | 206.6345681 | 183.3249104 | 109.9611699 | 113.5390417 | 95.82203177 | 9.90E-07 | -0.959851613 | 2.74E-07 | -5.140176295 | 0.514109789 |
| CNAG_00472 | - | hypothetical protein | 804.8348077 | 788.8096108 | 806.3382803 | 1123.467345 | 1118.115575 | 1043.450322 | 1.00E-06 | 0.438221598 | 2.78E-07 | 5.13795903 | 1.35493308 |
| CNAG_12994 | - | unspecified product | 119.7674754 | 185.4481928 | 169.7160208 | 283.8165926 | 302.3290214 | 294.993202 | 1.01E-06 | 0.880116633 | 2.79E-07 | 5.136965363 | 1.840524091 |
| CNAG_06266 | - | hypothetical protein | 282.9635004 | 321.1745832 | 233.7714229 | 481.6307254 | 428.8957522 | 467.1016065 | 1.01E-06 | 0.704591661 | 2.81E-07 | 5.136030811 | 1.629683335 |
| CNAG_04323 | - | hypothetical protein | 1628.145611 | 1831.698564 | 1626.460427 | 2159.157391 | 2469.847115 | 2173.597704 | 1.02E-06 | 0.404624364 | 2.84E-07 | 5.133457515 | 1.3237442 |
| CNAG_05350 | - | hypothetical protein, variant | 394.1641931 | 626.0522468 | 541.8886289 | 866.8372744 | 847.3658134 | 795.8065543 | 1.03E-06 | 0.670552369 | 2.86E-07 | 5.132613273 | 1.591682263 |
| CNAG_07011 | - | hypothetical protein, variant | 1242.213954 | 1252.919176 | 1145.323812 | 861.9352892 | 922.7584588 | 938.216717 | 1.06E-06 | -0.434518799 | 2.95E-07 | -5.126498703 | 0.739940515 |
| CNAG_00404 | - | RING zinc finger protein | 120.8052397 | 164.9479412 | 106.8892933 | 255.9690574 | 241.8447018 | 257.2102132 | 1.07E-06 | 0.935490767 | 2.97E-07 | 5.125656551 | 1.91254112 |
| CNAG_00810 | SSU72 | polynucleotide kinase 3'-phosphatase | 867.440054 | 903.8362952 | 885.0616759 | 651.8709612 | 657.9356728 | 645.1790544 | 1.09E-06 | -0.458120532 | 3.04E-07 | -5.12121647 | 0.727933956 |
| CNAG_07917 | - | uracil transporter FurD | 951.5891632 | 1034.201642 | 1045.001326 | 700.0251029 | 768.3696122 | 757.2806478 | 1.09E-06 | -0.460527269 | 3.05E-07 | -5.120554841 | 0.726720612 |
| CNAG_02008 | OVA1 | nuclear protein | 536.3581584 | 536.7272235 | 610.0898898 | 876.1604303 | 763.4989411 | 816.8865623 | 1.10E-06 | 0.530293196 | 3.06E-07 | 5.119553128 | 1.444222673 |
| CNAG_05267 | - | NADH dehydrogenase (ubiquinone) Fe-S protein 5 | 1522.591853 | 1556.797883 | 1631.55779 | 1276.213908 | 1182.370373 | 1119.249344 | 1.10E-06 | -0.411925399 | 3.07E-07 | -5.119169742 | 0.751619604 |
| CNAG_03407 | - | hypothetical protein | 21.64117137 | 19.69502237 | 33.23403955 | 65.73092101 | 86.03067268 | 94.82135571 | 1.10E-06 | 1.730948572 | 3.07E-07 | 5.119203063 | 3.31946001 |
| CNAG_04197 | YAK1 | CMGC/DYRK/YAK protein kinase | 1060.563382 | 1061.383321 | 1083.481337 | 1412.110457 | 1476.095906 | 1357.631282 | 1.14E-06 | 0.398665513 | 3.17E-07 | 5.112926591 | 1.318287936 |
| CNAG_05795 | - | mitogen-activated protein kinase organizer 1 | 991.2703107 | 1010.545069 | 1123.881784 | 795.9246254 | 753.6538515 | 734.017921 | 1.14E-06 | -0.468408724 | 3.18E-07 | -5.112412892 | 0.722761356 |
| CNAG_02601 | - | hypothetical protein | 20.48573518 | 29.32321126 | 25.8176114 | 93.9450576 | 85.16035661 | 63.14143754 | 1.15E-06 | 1.68894928 | 3.20E-07 | 5.111010641 | 3.224217973 |
| CNAG_03361 | - | hypothetical protein | 476.7803226 | 486.0673225 | 511.9966225 | 674.2407994 | 754.2135705 | 710.31706 | 1.15E-06 | 0.522173385 | 3.21E-07 | 5.110626598 | 1.436117094 |
| CNAG_03518 | - | Phenylacrylic acid decarboxylase | 33.38354422 | 31.57253244 | 61.98548716 | 116.7334407 | 151.0867308 | 97.53938239 | 1.16E-06 | 1.525332351 | 3.23E-07 | 5.10957384 | 2.878530205 |
| CNAG_07429 | - | asparagine-tRNA ligase | 1881.182925 | 2111.635886 | 2062.426644 | 1613.159756 | 1590.39985 | 1512.214489 | 1.18E-06 | -0.375479047 | 3.30E-07 | -5.105592877 | 0.77084941 |
| CNAG_01852 | - | CCR4-NOT transcription complex subunit 2 | 2656.55628 | 2840.017564 | 2896.391086 | 2256.023197 | 2291.694376 | 2151.654451 | 1.19E-06 | -0.340390602 | 3.33E-07 | -5.103981479 | 0.789827442 |
| CNAG_12896 | - | unspecified product | 9.950177405 | 13.19185323 | 17.25829001 | 54.50581966 | 51.29615321 | 64.87726663 | 1.19E-06 | 2.100979403 | 3.34E-07 | 5.103399636 | 4.29000522 |
| CNAG_07594 | - | hypothetical protein | 1091.974521 | 703.5449842 | 739.0023168 | 510.8072154 | 537.0940227 | 505.3932629 | 1.19E-06 | -0.723900185 | 3.34E-07 | -5.10342186 | 0.60545843 |
| CNAG_07667 | SAT4 | HAL protein kinase | 5234.829499 | 4861.344681 | 5259.663769 | 4113.000725 | 4200.533832 | 4173.212492 | 1.20E-06 | -0.313958053 | 3.35E-07 | -5.102475051 | 0.80443176 |
| CNAG_00496 | - | hypothetical protein | 865.092458 | 876.9199467 | 915.9542144 | 676.0789075 | 653.6255875 | 581.973961 | 1.20E-06 | -0.491040792 | 3.37E-07 | -5.101219253 | 0.711511612 |
| CNAG_00120 | - | hypothetical protein | 636.00843 | 552.6590139 | 632.2467763 | 373.8130414 | 393.9002825 | 445.3064748 | 1.21E-06 | -0.603500873 | 3.39E-07 | -5.100552984 | 0.658154924 |
| CNAG_07993 | - | hypothetical protein | 33.44397865 | 64.84109608 | 63.09681822 | 129.8862713 | 133.3056092 | 143.8356781 | 1.21E-06 | 1.326332762 | 3.40E-07 | 5.100046979 | 2.50764437 |
| CNAG_00728 | - | dityrosine transporter | 309.8980575 | 393.3371288 | 290.2759723 | 481.9812589 | 551.5342087 | 575.8241824 | 1.23E-06 | 0.68190065 | 3.45E-07 | 5.096980316 | 1.604251853 |
| CNAG_06686 | - | hypothetical protein | 1366.060703 | 1374.701446 | 1343.660846 | 1058.976128 | 1040.920379 | 1049.24789 | 1.23E-06 | -0.390830957 | 3.45E-07 | -5.096995266 | 0.762690187 |
| CNAG_04372 | - | DNA-directed RNA polymerase I, II, and III subunit RPABC2 | 1629.919687 | 1502.968508 | 1633.70076 | 1255.92104 | 1165.088527 | 1216.884156 | 1.23E-06 | -0.405379864 | 3.46E-07 | -5.096575618 | 0.755037464 |
| CNAG_03317 | - | AP-1 complex subunit mu-1 | 2732.423252 | 2776.475765 | 2926.249766 | 2264.576954 | 2230.411805 | 2279.277533 | 1.24E-06 | -0.331644155 | 3.49E-07 | -5.094819921 | 0.794630374 |
| CNAG_06597 | SPT8 | transcriptional activator SPT8 | 2172.966944 | 2187.080435 | 2061.406838 | 1593.570023 | 1773.319154 | 1664.276336 | 1.26E-06 | -0.36727965 | 3.54E-07 | -5.092147252 | 0.77524292 |
| CNAG_03343 | - | hypothetical protein | 1540.972735 | 1202.423576 | 1489.682695 | 987.2502515 | 977.8866062 | 1041.484064 | 1.27E-06 | -0.50997061 | 3.57E-07 | -5.090355391 | 0.702236743 |
| CNAG_02197 | - | hypothetical protein | 2306.140229 | 2342.245712 | 2334.389728 | 1771.055212 | 1969.144687 | 1748.680174 | 1.28E-06 | -0.362729661 | 3.61E-07 | -5.088582843 | 0.77769175 |
| CNAG_06412 | - | rRNA-processing protein EFG1 | 1311.200127 | 1243.316503 | 1541.881897 | 921.0315653 | 960.6473854 | 1051.440103 | 1.30E-06 | -0.497591041 | 3.66E-07 | -5.085982573 | 0.708288468 |
| CNAG_05804 | - | nuclear pore complex protein Nup188 | 4202.465189 | 3904.554285 | 3846.655993 | 3210.977679 | 3120.559463 | 3277.299093 | 1.32E-06 | -0.330542425 | 3.70E-07 | -5.083548541 | 0.795237434 |
| CNAG_02101 | - | hypothetical protein | 514.8487994 | 569.8119898 | 620.4743782 | 398.9218483 | 389.6422041 | 344.4293912 | 1.32E-06 | -0.605604764 | 3.73E-07 | -5.082456593 | 0.657195832 |
| CNAG_01211 | - | hypothetical protein | 3441.173166 | 3234.416366 | 3568.248308 | 2775.938187 | 2777.308839 | 2604.606698 | 1.36E-06 | -0.344391794 | 3.84E-07 | -5.076760973 | 0.787639957 |
| CNAG_01245 | - | hypothetical protein | 2336.483207 | 2311.015003 | 2309.871828 | 1786.630556 | 1888.871505 | 1880.726791 | 1.37E-06 | -0.340039585 | 3.86E-07 | -5.075902924 | 0.790019635 |
| CNAG_06834 | PMT1 | dolichyl-phosphate-mannose-protein mannosyltransferase | 2255.161323 | 2366.117459 | 2262.037841 | 2814.456218 | 2896.622442 | 3043.883327 | 1.37E-06 | 0.331778702 | 3.87E-07 | 5.07503681 | 1.258564105 |
| CNAG_05005 | ATG1 | ULK/ULK protein kinase | 154.7594449 | 205.9599904 | 229.3847199 | 348.4930135 | 347.9791986 | 337.239406 | 1.39E-06 | 0.796970407 | 3.93E-07 | 5.072308232 | 1.737448731 |
| CNAG_04871 | - | hypothetical protein | 281.8769415 | 356.7104025 | 278.5269199 | 507.2544845 | 509.2820355 | 449.4850748 | 1.40E-06 | 0.6647294 | 3.94E-07 | 5.071904254 | 1.585270903 |
| CNAG_03238 | - | dioxygenase subfamily protein | 49.56797696 | 44.44567996 | 51.38877402 | 139.7707622 | 126.499763 | 96.47321342 | 1.40E-06 | 1.310072538 | 3.96E-07 | 5.070943758 | 2.479540066 |
| CNAG_03063 | - | hypothetical protein, variant | 5714.673283 | 5157.620188 | 5225.570101 | 4053.757847 | 4228.963062 | 4510.661769 | 1.41E-06 | -0.347188316 | 3.98E-07 | -5.070139571 | 0.786114674 |
| CNAG_04683 | - | hypothetical protein | 2646.346562 | 2429.763016 | 2816.555028 | 3297.013504 | 3520.324707 | 3481.308974 | 1.41E-06 | 0.368294139 | 3.99E-07 | 5.069584609 | 1.290825639 |
| CNAG_03299 | - | coatomer beta subunit | 9276.367574 | 8378.085507 | 8794.980936 | 7047.140348 | 7297.997848 | 7277.213454 | 1.46E-06 | -0.306472994 | 4.13E-07 | -5.063081088 | 0.808616196 |
| CNAG_02307 | - | L-serine ammonia-lyase | 1041.375075 | 1009.462248 | 1021.574311 | 785.7967075 | 752.7977219 | 759.5171388 | 1.47E-06 | -0.434921687 | 4.15E-07 | -5.062126607 | 0.739733908 |
| CNAG_02944 | APH1 | phosphate-repressible vacuolar acid phosphatase | 143.1378386 | 250.0303157 | 167.6568493 | 318.6562158 | 347.098755 | 410.2210984 | 1.47E-06 | 0.928198175 | 4.15E-07 | 5.062010605 | 1.902897925 |
| CNAG_06315 | - | protein CWC15 | 783.1323162 | 714.2274235 | 730.4083485 | 539.0647499 | 414.4706822 | 537.4644889 | 1.47E-06 | -0.595028867 | 4.16E-07 | -5.061704509 | 0.662031208 |
| CNAG_05558 | KIN4 | CAMK/CAMKL/KIN4 protein kinase | 315.5516836 | 283.5415275 | 274.2441339 | 476.1037446 | 478.2186858 | 422.8462511 | 1.48E-06 | 0.642409034 | 4.18E-07 | 5.06059331 | 1.560933454 |
| CNAG_05920 | - | tRNA-specific adenosine deaminase 2 | 851.0067357 | 904.8523298 | 777.3998989 | 590.3251429 | 565.5518686 | 642.9067756 | 1.48E-06 | -0.509377045 | 4.20E-07 | -5.059632156 | 0.702525723 |
| CNAG_06404 | - | hypothetical protein | 2075.62455 | 1134.472648 | 1366.984806 | 631.1726193 | 529.8787199 | 501.1812456 | 1.50E-06 | -1.479756491 | 4.25E-07 | -5.057485437 | 0.358549326 |
| CNAG_05028 | - | Cysteine synthase | 4619.227204 | 3884.116959 | 4106.848541 | 3062.955229 | 3413.030824 | 3254.025544 | 1.53E-06 | -0.390220188 | 4.33E-07 | -5.053996385 | 0.763013142 |
| CNAG_04219 | - | lactoylglutathione lyase, variant | 2541.935291 | 2388.602758 | 2588.164279 | 2049.083016 | 1941.464078 | 1614.453362 | 1.54E-06 | -0.439490863 | 4.36E-07 | -5.052433905 | 0.737394795 |
| CNAG_00286 | NAB2 | putative nuclear polyadenylated mRNA-binding protein | 1376.39492 | 1190.517137 | 1312.690956 | 923.4515578 | 859.6676797 | 1025.882519 | 1.56E-06 | -0.481947915 | 4.44E-07 | -5.049259257 | 0.71601022 |
| CNAG_04181 | - | RNA-binding domain-containing protein, X-linked 2 | 491.3190158 | 540.6358501 | 475.542829 | 286.5693904 | 299.8243942 | 375.3335241 | 1.56E-06 | -0.664123518 | 4.44E-07 | -5.048968612 | 0.631071984 |
| CNAG_06694 | URO2 | hydroxyisourate hydrolase | 195.4344014 | 225.3054584 | 155.9439834 | 404.2270281 | 302.9045354 | 336.1305664 | 1.59E-06 | 0.843605641 | 4.50E-07 | 5.046361695 | 1.794529501 |
| CNAG_02344 | - | Uracil phosphoribosyltransferase | 19.17411225 | 8.920453188 | 15.15645714 | 69.79636489 | 68.92285284 | 45.39245755 | 1.59E-06 | 2.097695718 | 4.52E-07 | 5.045816766 | 4.280251941 |
| CNAG_03386 | - | solute carrier family 25 (mitochondrial carnitine/acylcarnitine transporter), member 20/29 | 2421.919053 | 2985.379691 | 2748.159557 | 2043.452777 | 2177.829972 | 2042.864108 | 1.62E-06 | -0.395323514 | 4.60E-07 | -5.04244153 | 0.760318862 |
| CNAG_06935 | - | isochorismatase hydrolase | 340.2932796 | 417.0824377 | 378.7363407 | 616.9476081 | 508.0801496 | 690.0814205 | 1.62E-06 | 0.662477764 | 4.61E-07 | 5.042069309 | 1.582798676 |
| CNAG_06565 | - | hypothetical protein | 981.6767913 | 894.1711444 | 880.8179469 | 686.9433481 | 689.8757859 | 590.8686415 | 1.62E-06 | -0.503255364 | 4.61E-07 | -5.041667926 | 0.705513032 |
| CNAG_03775 | - | hypothetical protein, variant 1 | 2963.694905 | 3293.591335 | 3221.670415 | 2559.566825 | 2396.84045 | 2592.328002 | 1.62E-06 | -0.343224975 | 4.62E-07 | -5.041426918 | 0.78827724 |
| CNAG_05524 | - | glycogen storage control protein | 350.7387187 | 369.7236612 | 416.0003873 | 565.4314171 | 601.5546019 | 550.4801876 | 1.62E-06 | 0.581030201 | 4.62E-07 | 5.041439676 | 1.495917073 |
| CNAG_05041 | - | NADH-ubiquinone oxidoreductase subunit 8 | 2438.195976 | 2774.241057 | 2573.284884 | 2131.542583 | 1981.097942 | 2026.187908 | 1.63E-06 | -0.357534264 | 4.63E-07 | -5.041053322 | 0.780497403 |
| CNAG_02690 | - | pirin | 350.523866 | 284.6453082 | 289.1769209 | 470.4653765 | 511.8778615 | 478.2707081 | 1.64E-06 | 0.644551 | 4.68E-07 | 5.038856765 | 1.56325269 |
| CNAG_00585 | - | hypothetical protein | 583.5156005 | 524.6390272 | 654.5568217 | 303.5820873 | 354.181562 | 436.3884284 | 1.67E-06 | -0.703787195 | 4.76E-07 | -5.035876657 | 0.613958397 |
| CNAG_05236 | - | hypothetical protein | 1699.793548 | 1442.62651 | 1452.456931 | 1160.913248 | 1157.370199 | 1098.143868 | 1.68E-06 | -0.443491726 | 4.80E-07 | -5.03418449 | 0.735352694 |
| CNAG_00091 | - | hypothetical protein | 1302.498535 | 1349.138084 | 909.979215 | 2837.39191 | 2324.985799 | 2170.156205 | 1.70E-06 | 1.02755287 | 4.84E-07 | 5.032364736 | 2.038563463 |
| CNAG_12488 | - | unspecified product | 104.4603439 | 121.9344839 | 119.6084463 | 205.9135764 | 202.0847565 | 252.5997731 | 1.70E-06 | 0.923339794 | 4.85E-07 | 5.032206198 | 1.896500555 |
| CNAG_06849 | - | saccharopine dehydrogenase (NAD, L-lysine forming) | 2160.247874 | 2427.307561 | 2223.497244 | 1723.573754 | 1818.991743 | 1824.111712 | 1.70E-06 | -0.358790906 | 4.86E-07 | -5.031682571 | 0.779817856 |
| CNAG_01432 | - | ribosome assembly protein 4, variant | 3301.030007 | 3066.3411 | 3375.226152 | 2589.886989 | 2563.356627 | 2638.965797 | 1.71E-06 | -0.338201228 | 4.87E-07 | -5.031247855 | 0.79102696 |
| CNAG_10502 | - | unspecified product | 30958.3911 | 23918.42878 | 12531.97345 | 6676.692076 | 4450.041858 | 6425.031121 | 1.79E-06 | -1.956576268 | 5.11E-07 | -5.022031017 | 0.257639148 |
| CNAG_01994 | - | hypothetical protein | 32.19744215 | 58.33086131 | 34.3835235 | 112.1209755 | 99.50858563 | 125.979394 | 1.80E-06 | 1.425412041 | 5.14E-07 | 5.020939299 | 2.685912006 |
| CNAG_01177 | NEM1 | nuclear envelope-endoplasmic reticulum network protein | 361.2669038 | 333.1621252 | 459.6958456 | 570.783349 | 611.853974 | 653.5516353 | 1.80E-06 | 0.653808887 | 5.15E-07 | 5.020475172 | 1.573316461 |
| CNAG_04064 | - | hypothetical protein | 887.3182434 | 916.7859856 | 999.1026812 | 729.0579956 | 589.6501123 | 661.8409988 | 1.82E-06 | -0.516723557 | 5.20E-07 | -5.018715581 | 0.698957404 |
| CNAG_01073 | - | hypothetical protein, variant | 318.5280911 | 363.9190071 | 323.0181374 | 218.2967199 | 208.4478742 | 192.3447797 | 1.82E-06 | -0.714800987 | 5.21E-07 | -5.018413894 | 0.609289175 |
| CNAG_04950 | - | hypothetical protein | 2774.364019 | 2755.978808 | 2614.902644 | 2235.047046 | 2211.468294 | 2069.511183 | 1.84E-06 | -0.337077823 | 5.26E-07 | -5.016555452 | 0.791643162 |
| CNAG_04008 | - | hypothetical protein | 26.34454396 | 22.94776881 | 43.88522882 | 134.3000227 | 85.72277898 | 76.46703145 | 1.84E-06 | 1.680651148 | 5.27E-07 | 5.016255536 | 3.205726061 |

| | | | | | | | | | | | | | |
|---|---|---|---|---|---|---|---|---|---|---|---|---|---|
| CNAG_01713 | - | hypothetical protein | 1165.601013 | 1164.790025 | 883.2251711 | 1590.524339 | 1539.911813 | 1474.158468 | 1.84E-06 | 0.504136259 | 5.28E-07 | 5.015770554 | 1.418273982 |
| CNAG_04989 | - | protein FRG1 | 1253.840158 | 1122.633265 | 1214.587054 | 878.3373914 | 942.6397221 | 851.6939281 | 1.88E-06 | -0.442070175 | 5.40E-07 | -5.011367776 | 0.736077626 |
| CNAG_05415 | - | hypothetical protein | 671.3353747 | 729.2927473 | 783.6726997 | 521.8856037 | 482.7415776 | 530.814624 | 1.89E-06 | -0.52487798 | 5.44E-07 | -5.010281872 | 0.69501789 |
| CNAG_03314 | - | YTH domain family 2 | 260.8285836 | 300.7236014 | 261.4466778 | 443.3453801 | 491.2941017 | 389.5833961 | 1.90E-06 | 0.673425773 | 5.45E-07 | 5.009771219 | 1.594855563 |
| CNAG_00761 | - | coiled-coil domain-containing protein 12 | 1069.455826 | 1034.269462 | 1158.023131 | 806.7597946 | 765.6969994 | 838.2998192 | 1.90E-06 | -0.452252266 | 5.47E-07 | -5.009042808 | 0.73090091 |
| CNAG_01379 | - | hypothetical protein | 636.2027092 | 677.5440943 | 679.19069 | 450.9997806 | 510.5172521 | 384.4314278 | 1.92E-06 | -0.583064473 | 5.51E-07 | -5.007471581 | 0.667544319 |
| CNAG_02080 | - | hypothetical protein | 1203.834066 | 1243.274723 | 1367.046773 | 945.2830082 | 961.5636542 | 969.3171376 | 1.93E-06 | -0.422607228 | 5.55E-07 | -5.006284358 | 0.746075103 |
| CNAG_05940 | ZFC3 | hypothetical protein | 663.9023073 | 612.8875439 | 516.1449152 | 412.122178 | 389.5923068 | 391.0020666 | 1.96E-06 | -0.602710715 | 5.65E-07 | -5.002925573 | 0.658515491 |
| CNAG_06578 | - | hypothetical protein | 2152.454893 | 2395.196834 | 2024.272395 | 3383.905409 | 2913.889967 | 2698.766858 | 1.98E-06 | 0.438222533 | 5.70E-07 | 5.001238893 | 1.354933957 |
| CNAG_03079 | PER1 | Protein processing in the ER protein 1, putative | 696.4265357 | 580.5942084 | 499.088559 | 361.3686551 | 399.9915731 | 379.9044424 | 2.02E-06 | -0.655937561 | 5.83E-07 | -4.99682833 | 0.63466291 |
| CNAG_04665 | - | hypothetical protein | 302.8283414 | 330.90038 | 312.6078215 | 518.9797872 | 477.245793 | 453.920384 | 2.10E-06 | 0.602275184 | 6.05E-07 | 4.989483326 | 1.518108794 |
| CNAG_12814 | - | unspecified product | 9.917661253 | 15.30748862 | 10.89018394 | 47.36430407 | 45.19980725 | 63.6763496 | 2.11E-06 | 2.175257203 | 6.08E-07 | 4.988793814 | 4.516662777 |
| CNAG_02604 | - | hypothetical protein | 96.30735524 | 119.7650689 | 114.2743296 | 191.8450505 | 216.0776796 | 220.5440231 | 2.12E-06 | 0.915758012 | 6.11E-07 | 4.987790544 | 1.886560037 |
| CNAG_00503 | - | hypothetical protein | 1053.489772 | 984.8972757 | 960.9933348 | 1291.389378 | 1404.512755 | 1337.597844 | 2.13E-06 | 0.412440463 | 6.13E-07 | 4.986984567 | 1.330935321 |
| CNAG_05478 | - | charged multivesicular body protein 2A | 1403.213083 | 1190.517924 | 1253.006684 | 949.1805029 | 984.8788737 | 913.8599818 | 2.14E-06 | -0.450435279 | 6.18E-07 | -4.985358926 | 0.731822015 |
| CNAG_04013 | - | potassium channel protein | 1041.398079 | 1099.897758 | 987.4819146 | 817.7586375 | 716.5081341 | 770.6163645 | 2.16E-06 | -0.456059163 | 6.25E-07 | -4.983442151 | 0.728974795 |
| CNAG_05896 | UBA2 | ubiquitin-like 1-activating enzyme E1 B | 1717.554126 | 1612.873839 | 1802.185046 | 1285.394189 | 1335.93828 | 1344.534325 | 2.16E-06 | -0.387893016 | 6.25E-07 | -4.983456635 | 0.764244931 |
| CNAG_05244 | - | hypothetical protein | 59.16507354 | 121.8949467 | 110.0094147 | 225.868978 | 203.957978 | 205.0590362 | 2.22E-06 | 1.113084123 | 6.42E-07 | 4.978143286 | 2.163075651 |
| CNAG_07567 | - | hypothetical protein | 1007.502792 | 921.159417 | 1095.071848 | 656.365847 | 720.0125732 | 776.1171833 | 2.22E-06 | -0.506012363 | 6.42E-07 | -4.978047572 | 0.704166079 |
| CNAG_06209 | - | hypothetical protein | 23.88403046 | 22.88024264 | 22.61564045 | 71.63252848 | 80.93042684 | 64.12980824 | 2.25E-06 | 1.666502764 | 6.50E-07 | 4.975594359 | 3.174441439 |
| CNAG_04461 | HFM1 | ATP-dependent DNA helicase HFM1/MER3 | 84.65837308 | 106.8496758 | 104.6777668 | 167.5810028 | 218.8358359 | 202.7915839 | 2.26E-06 | 0.982788735 | 6.53E-07 | 4.974987274 | 1.976281879 |
| CNAG_06486 | - | hypothetical protein | 2111.011258 | 1896.244134 | 2059.228466 | 1546.928146 | 1518.756017 | 1638.699883 | 2.26E-06 | -0.383116241 | 6.53E-07 | -4.97473284 | 0.766779547 |
| CNAG_06913 | - | L-serine ammonia-lyase | 67.153763 | 86.38367059 | 80.17098847 | 171.2342312 | 138.1543361 | 189.2949076 | 2.28E-06 | 1.081761894 | 6.59E-07 | 4.97297141 | 2.116619428 |
| CNAG_06717 | - | Acyl-CoA-dependent ceramide synthase | 240.8835836 | 243.6446909 | 215.5993077 | 393.6552676 | 368.5829714 | 361.7219529 | 2.29E-06 | 0.669705158 | 6.64E-07 | 4.971598085 | 1.590747835 |
| CNAG_03504 | - | hypothetical protein | 74.19703769 | 102.5365133 | 102.5344369 | 172.4309395 | 184.9680695 | 208.2338148 | 2.32E-06 | 1.006358449 | 6.73E-07 | 4.968899631 | 2.008834135 |
| CNAG_06947 | - | hypothetical protein | 1152.262623 | 1099.958788 | 1057.865395 | 815.2999515 | 829.5866402 | 847.1920743 | 2.36E-06 | -0.424580799 | 6.83E-07 | -4.96609949 | 0.745055189 |
| CNAG_00362 | - | hypothetical protein | 1541.238709 | 1607.401213 | 1490.849632 | 1139.041323 | 1260.945848 | 1188.02235 | 2.37E-06 | -0.385830041 | 6.87E-07 | -4.965111787 | 0.765338542 |
| CNAG_01483 | HPI1 | E3 ubiquitin ligase-like protein | 1194.278925 | 1066.606456 | 1167.653444 | 824.6162163 | 790.6957175 | 904.8826781 | 2.38E-06 | -0.460254609 | 6.89E-07 | -4.964345327 | 0.72685797 |
| CNAG_04106 | - | hypothetical protein | 189.47563 | 183.3191721 | 121.8420635 | 288.4305948 | 293.5870535 | 331.5084346 | 2.40E-06 | 0.872635603 | 6.98E-07 | 4.962027949 | 1.831004837 |
| CNAG_06983 | - | hypothetical protein | 1612.4445 | 1611.737146 | 1537.77374 | 1224.724966 | 1230.691691 | 1271.251216 | 2.44E-06 | -0.368875413 | 7.08E-07 | -4.959049322 | 0.774385899 |
| CNAG_07999 | - | hypothetical protein | 458.0223102 | 430.0525832 | 441.6340225 | 684.6658212 | 612.5736583 | 642.5981582 | 2.44E-06 | 0.528865875 | 7.08E-07 | 4.959088127 | 1.442794546 |
| CNAG_02530 | - | hypothetical protein | 3484.429914 | 3373.402437 | 3857.224968 | 2659.824979 | 2799.699557 | 2955.336461 | 2.45E-06 | -0.364303546 | 7.12E-07 | -4.958008757 | 0.776843802 |
| CNAG_01094 | - | pre-mRNA-splicing factor CWC25 | 1732.715878 | 1635.48752 | 1728.632272 | 1344.626935 | 1343.709042 | 1304.594851 | 2.45E-06 | -0.367878584 | 7.13E-07 | -4.957779796 | 0.774921145 |
| CNAG_07633 | - | hypothetical protein | 36.77437316 | 49.74192987 | 34.36779512 | 100.3300664 | 106.6374446 | 101.7698507 | 2.47E-06 | 1.353120189 | 7.18E-07 | 4.956422883 | 2.554640333 |
| CNAG_12048 | - | unspecified product | 18.12090718 | 27.14505133 | 19.43036429 | 73.33098811 | 74.80023774 | 64.09709423 | 2.47E-06 | 1.706976081 | 7.18E-07 | 4.956423751 | 3.264758057 |
| CNAG_01842 | - | hypothetical protein | 1234.061004 | 1246.463214 | 1178.366044 | 837.7563493 | 904.6184853 | 988.1289765 | 2.50E-06 | -0.437746484 | 7.26E-07 | -4.954231926 | 0.738286926 |
| CNAG_04931 | HXS2 | Putative hexose transporter | 0.271137429 | 0.130402328 | 0.112489064 | 25.33725901 | 8.517388974 | 28.16694357 | 2.53E-06 | 6.774767325 | 7.36E-07 | 4.951719037 | 109.4985022 |
| CNAG_00596 | - | hypothetical protein | 321.5364264 | 313.7315323 | 354.1796635 | 465.5578567 | 497.0203869 | 647.7902347 | 2.55E-06 | 0.688438073 | 7.43E-07 | 4.949670353 | 1.611537849 |
| CNAG_06132 | HAD8 | glycerol-1-phosphatase | 31.04702677 | 49.77391511 | 49.24187096 | 133.6578209 | 99.4916937 | 104.0650354 | 2.56E-06 | 1.367141946 | 7.45E-07 | 4.949273172 | 2.579590294 |
| CNAG_05154 | - | membrane fraction protein | 4318.330315 | 5029.313825 | 4853.337233 | 3553.3098 | 3692.440768 | 3988.88158 | 2.57E-06 | -0.352664392 | 7.48E-07 | -4.948369315 | 0.783136453 |
| CNAG_02232 | - | RNA polymerase II-associated factor 1 | 2318.894249 | 2222.660718 | 2159.518767 | 1806.189973 | 1822.464774 | 1638.794179 | 2.58E-06 | -0.362739031 | 7.53E-07 | -4.947268497 | 0.777686699 |
| CNAG_07674 | - | hypothetical protein | 310.9818674 | 345.9629672 | 281.727014 | 467.3025601 | 552.5394188 | 456.143913 | 2.60E-06 | 0.641184691 | 7.58E-07 | 4.945753427 | 1.559609331 |
| CNAG_07546 | - | peptide chain release factor 1 | 1367.347533 | 1475.974555 | 1524.909733 | 1127.460988 | 1158.27594 | 1065.943063 | 2.63E-06 | -0.397272301 | 7.67E-07 | -4.94369246 | 0.759292519 |
| CNAG_01817 | - | signal recognition particle receptor subunit alpha | 394.6220568 | 525.4776349 | 449.9155632 | 268.7145345 | 292.12605 | 308.8077316 | 2.65E-06 | -0.670313798 | 7.74E-07 | -4.941722861 | 0.628369997 |
| CNAG_05299 | - | oxidoreductase | 10.14288989 | 22.97751787 | 2.427271617 | 125.6745048 | 160.8618962 | 142.4331064 | 2.68E-06 | 3.640660049 | 7.82E-07 | 4.939863688 | 12.4723382 |
| CNAG_02176 | - | hypothetical protein | 1094.398786 | 981.7040234 | 1119.848566 | 1422.221947 | 1406.133488 | 1480.760149 | 2.69E-06 | 0.415962358 | 7.85E-07 | 4.939126433 | 1.334188358 |
| CNAG_05298 | - | hypothetical protein, variant | 62.50583673 | 97.09241746 | 61.03770334 | 145.175893 | 160.8594273 | 185.9509104 | 2.72E-06 | 1.142035031 | 7.96E-07 | 4.936388068 | 2.206921066 |
| CNAG_03159 | - | cytoplasmic protein | 649.8954832 | 510.6750135 | 609.852545 | 344.9552837 | 413.8230085 | 400.9590556 | 2.72E-06 | -0.628120883 | 7.96E-07 | -4.936294935 | 0.647018612 |
| CNAG_03822 | - | monolysocardiolipin acyltransferase | 1140.604304 | 1045.053415 | 1136.727268 | 798.9228416 | 803.6768209 | 872.6978626 | 2.73E-06 | -0.440908617 | 7.98E-07 | -4.935841934 | 0.736670503 |
| CNAG_00241 | - | protein AIR1/2 | 1955.789687 | 1934.987959 | 1934.477382 | 1515.037255 | 1493.757886 | 1604.275086 | 2.75E-06 | -0.351564707 | 8.05E-07 | -4.934237136 | 0.783733621 |
| CNAG_04728 | - | DNA-directed RNA polymerase III subunit RPC8 | 1831.322604 | 1867.311315 | 2020.989315 | 2726.086241 | 2397.16372 | 2402.300519 | 2.76E-06 | 0.380962619 | 8.08E-07 | 4.933455072 | 1.302210448 |
| CNAG_04800 | - | phenylalanine-tRNA ligase, alpha subunit | 9406.417571 | 9702.338053 | 9326.208539 | 11333.63914 | 11403.17578 | 11376.83798 | 2.79E-06 | 0.247581796 | 8.18E-07 | 4.931006759 | 1.18721547 |
| CNAG_12895 | - | unspecified product | 873.8370484 | 895.4984375 | 996.1430261 | 1326.602063 | 1148.092218 | 1473.945715 | 2.82E-06 | 0.499073687 | 8.25E-07 | 4.929297561 | 1.413305828 |
| CNAG_05192 | - | hypothetical protein | 13.72061151 | 39.17393402 | 56.81274115 | 208.1284663 | 274.8032636 | 213.6739662 | 2.84E-06 | 2.672350577 | 8.31E-07 | 4.927868914 | 6.374669636 |
| CNAG_12954 | - | unspecified product | 166.6465968 | 196.8475881 | 174.7253463 | 107.0035273 | 77.15544453 | 86.91393418 | 2.91E-06 | -1.00361893 | 8.53E-07 | -4.922941405 | 0.498747346 |
| CNAG_06498 | - | hypothetical protein | 169.0361654 | 312.708543 | 322.2757832 | 652.9364046 | 725.2181635 | 748.7683344 | 2.91E-06 | 1.390809601 | 8.53E-07 | 4.922845321 | 2.622257933 |
| CNAG_02535 | - | hypothetical protein | 317.1160956 | 276.7913468 | 316.5348612 | 202.0298384 | 180.8328649 | 141.3755368 | 2.92E-06 | -0.813518943 | 8.57E-07 | -4.922032392 | 0.568992309 |
| CNAG_00544 | - | hypothetical protein | 518.3887065 | 591.3514902 | 625.8234735 | 415.2869117 | 387.0203342 | 379.9047009 | 2.97E-06 | -0.568964918 | 8.74E-07 | -4.918131525 | 0.674100258 |
| CNAG_06221 | - | diphthamide biosynthesis protein 2 | 1315.959868 | 1426.405552 | 1438.540208 | 1034.767416 | 1095.292992 | 1091.419302 | 2.97E-06 | -0.391091194 | 8.74E-07 | -4.91813281 | 0.762552623 |
| CNAG_03423 | - | transcriptional activator | 519.9126459 | 482.8525903 | 508.8132545 | 776.5074999 | 660.8032815 | 758.0003729 | 3.05E-06 | 0.524655686 | 8.95E-07 | 4.913384342 | 1.438590204 |
| CNAG_07937 | CAS1 | O-acetyltransferase | 1654.508561 | 1679.613011 | 1661.453379 | 1389.110167 | 1270.361664 | 1168.112404 | 3.07E-06 | -0.39921711 | 9.04E-07 | -4.911440145 | 0.758269653 |
| CNAG_06439 | ALO1 | d-arabinono-1,4-lactone oxidase | 2249.989931 | 2048.15775 | 2187.211771 | 1639.520259 | 1674.908035 | 1773.022727 | 3.08E-06 | -0.365964344 | 9.07E-07 | -4.910767328 | 0.775950032 |
| CNAG_05777 | - | hypothetical protein | 347.9963987 | 449.0702005 | 487.1059884 | 271.9520126 | 251.5207264 | 271.1039573 | 3.13E-06 | -0.708524401 | 9.21E-07 | -4.90789066 | 0.611945721 |
| CNAG_00083 | CCC1 | hypothetical protein | 2382.083307 | 2471.537234 | 2480.488168 | 1892.491331 | 1855.154494 | 2073.848269 | 3.14E-06 | -0.348356701 | 9.25E-07 | -4.907023058 | 0.785478286 |
| CNAG_05308 | - | beta-catenin-like protein 1 | 860.8777716 | 823.2976807 | 882.0408635 | 1177.913686 | 1196.621864 | 1096.750435 | 3.16E-06 | 0.419998299 | 9.31E-07 | 4.905628502 | 1.337925977 |
| CNAG_12673 | - | unspecified product | 23.90486665 | 25.02813361 | 25.80194338 | 68.31222707 | 77.33664253 | 77.2641804 | 3.16E-06 | 1.589526289 | 9.33E-07 | 4.905311456 | 3.009505158 |
| CNAG_00869 | PDR5 | ATP-binding cassette transporter | 1391.302857 | 1171.044382 | 1014.194596 | 742.7166787 | 778.5948122 | 913.764621 | 3.16E-06 | -0.57121772 | 9.33E-07 | -4.905356529 | 0.673048456 |
| CNAG_12867 | - | unspecified product | 11.03389551 | 5.664600429 | 12.99238285 | 59.87800377 | 35.52102296 | 50.58012353 | 3.19E-06 | 2.346077608 | 9.42E-07 | 4.903351044 | 5.084400258 |
| CNAG_10500 | - | unspecified product | 433646.2318 | 319514.2973 | 204568.166 | 118586.4337 | 75544.08477 | 93314.00798 | 3.20E-06 | -1.752139326 | 9.43E-07 | -4.903161447 | 0.296861246 |
| CNAG_06860 | - | nuclear protein | 4382.408991 | 4311.822523 | 4298.82253 | 3527.734627 | 3659.732223 | 3598.13995 | 3.20E-06 | -0.283962924 | 9.43E-07 | -4.903191112 | 0.821331808 |
| CNAG_04486 | - | hypothetical protein | 21.66589956 | 32.548947 | 32.19008777 | 84.10165603 | 71.86015742 | 97.08320094 | 3.23E-06 | 1.554032455 | 9.54E-07 | 4.900982686 | 2.936367328 |
| CNAG_00814 | - | hypothetical protein | 739.3928108 | 629.4067752 | 707.1635818 | 905.3018671 | 1076.856809 | 983.4248426 | 3.27E-06 | 0.498340225 | 9.65E-07 | 4.898646533 | 1.41258749 |
| CNAG_01190 | - | vesicle transporter SEC22 | 1551.759477 | 1557.87924 | 1582.529199 | 1132.784756 | 1276.471527 | 1230.188345 | 3.30E-06 | -0.381777906 | 9.75E-07 | -4.896563974 | 0.76749119 |
| CNAG_00752 | - | transducin family protein | 576.3305574 | 486.9124155 | 493.6829188 | 340.438794 | 343.0263231 | 358.7650058 | 3.31E-06 | -0.594753391 | 9.79E-07 | -4.895748701 | 0.662157632 |
| CNAG_03979 | - | ATP(GTP)-binding protein Fet5 | 667.7469562 | 676.5202171 | 711.1925873 | 483.7092665 | 506.1308074 | 487.5461664 | 3.36E-06 | -0.492049195 | 9.93E-07 | -4.893094432 | 0.71101446 |
| CNAG_00801 | - | hypothetical protein | 75.18839842 | 75.62765614 | 70.58488146 | 145.3084642 | 153.0985625 | 156.1672391 | 3.36E-06 | 1.030032454 | 9.94E-07 | 4.892861928 | 2.042070187 |
| CNAG_02172 | - | ubiquitin-protein ligase | 1433.045394 | 1425.544993 | 1436.595937 | 1817.53672 | 1893.528848 | 1805.016777 | 3.37E-06 | 0.345699816 | 9.97E-07 | 4.892172223 | 1.270767252 |
| CNAG_01042 | - | hypothetical protein | 496.5725808 | 531.2615624 | 437.4137886 | 694.573297 | 726.5609701 | 697.0089416 | 3.49E-06 | 0.517243513 | 1.03E-06 | 4.884945494 | 1.431218075 |
| CNAG_07509 | - | Xaa-Pro aminopeptidase | 869.648942 | 865.0325416 | 776.3352695 | 639.4935006 | 592.3355942 | 594.1288199 | 3.53E-06 | -0.474895363 | 1.05E-06 | -4.882648428 | 0.719518974 |
| CNAG_03163 | - | hypothetical protein, variant 2 | 368.3271891 | 447.2602257 | 475.7141032 | 641.7255729 | 646.326917 | 635.9290813 | 3.54E-06 | 0.560505215 | 1.05E-06 | 4.882005085 | 1.474785579 |
| CNAG_01029 | - | impact family protein | 849.7768439 | 748.7683073 | 854.087483 | 627.8462971 | 544.8454414 | 570.8209391 | 3.63E-06 | -0.509188868 | 1.08E-06 | -4.877266349 | 0.702617362 |
| CNAG_03540 | - | aspartyl/glutamyl-tRNA(Asn/Gln) amidotransferase, B subunit | 1835.536864 | 1810.020312 | 1883.269894 | 1473.050119 | 1485.182647 | 1441.134757 | 3.63E-06 | -0.344957884 | 1.08E-06 | -4.876871769 | 0.787330961 |
| CNAG_00072 | - | protein disulfide-isomerase | 2290.901393 | 2160.198088 | 2228.818166 | 1824.100935 | 1773.261368 | 1745.319094 | 3.64E-06 | -0.337540488 | 1.08E-06 | -4.876321749 | 0.791389326 |
| CNAG_03240 | - | Mannosyl-oligosaccharide 1,2-alpha-mannosidase | 2931.332203 | 3244.210026 | 3110.921096 | 3731.254262 | 3886.843421 | 4071.895663 | 3.67E-06 | 0.317289332 | 1.09E-06 | 4.874748727 | 1.245987273 |
| CNAG_07960 | - | hypothetical protein | 488.5457723 | 581.9136835 | 579.1726675 | 808.3142516 | 787.7530462 | 758.1016677 | 3.69E-06 | 0.499072158 | 1.10E-06 | 4.87364073 | 1.41330433 |
| CNAG_03984 | YCH1 | hypothetical protein | 852.342664 | 928.6175456 | 942.6218254 | 660.4072603 | 659.6342007 | 696.2042109 | 3.69E-06 | -0.448797019 | 1.10E-06 | -4.873321056 | 0.732653511 |
| CNAG_07725 | - | HMG box factor | 0.362084733 | 4.516377287 | 2.317120719 | 27.93803829 | 19.93687702 | 45.66873396 | 3.70E-06 | 3.911573741 | 1.10E-06 | 4.872949686 | 15.04877075 |

| | | | | | | | | | | | | | |
|---|---|---|---|---|---|---|---|---|---|---|---|---|---|
| CNAG_04245 | CHI22 | chitinase, variant | 1616.692011 | 1347.757275 | 1159.246855 | 978.7247598 | 949.4225386 | 974.9054847 | 3.77E-06 | -0.522579712 | 1.12E-06 | -4.869314401 | 0.696125963 |
| CNAG_06969 | HAD7 | magnesium-dependent phosphatase-1 | 783.1114019 | 675.4802618 | 760.2283594 | 409.4761048 | 583.0140355 | 472.0617264 | 3.77E-06 | -0.615671112 | 1.12E-06 | -4.868984495 | 0.652626235 |
| CNAG_00301 | - | hypothetical protein | 657.6078264 | 612.1018986 | 481.1777745 | 899.4414235 | 833.4255256 | 867.9392973 | 3.78E-06 | 0.555430852 | 1.12E-06 | 4.868421822 | 1.469607457 |
| CNAG_12926 | - | unspecified product | 8.681743718 | 8.855715611 | 7.67653134 | 49.54814737 | 35.67684197 | 40.69876771 | 3.78E-06 | 2.415774702 | 1.13E-06 | 4.868176861 | 5.33605931 |
| CNAG_02366 | - | 4-aminobutyrate aminotransferase | 1490.945782 | 1398.438275 | 1416.181481 | 1080.706127 | 1109.93526 | 1146.911129 | 3.84E-06 | -0.383448932 | 1.14E-06 | -4.865037003 | 0.766602745 |
| CNAG_02387 | - | hypothetical protein, variant | 339.2688495 | 357.398165 | 277.2211777 | 226.2580661 | 174.7226403 | 138.0812126 | 3.89E-06 | -0.869694472 | 1.16E-06 | -4.862718673 | 0.547262735 |
| CNAG_12698 | - | unspecified product | 88.10665557 | 86.44419086 | 113.1644735 | 202.4235477 | 175.3747421 | 188.41437 | 3.93E-06 | 0.961783103 | 1.17E-06 | 4.860615483 | 1.947715692 |
| CNAG_04081 | - | hypothetical protein | 389.7922489 | 409.270494 | 429.6084153 | 247.7299083 | 254.1402468 | 294.3304329 | 4.01E-06 | -0.64109497 | 1.20E-06 | -4.856292205 | 0.641226089 |
| CNAG_01331 | - | hypothetical protein | 412.5977187 | 381.6247658 | 490.6339779 | 594.8044017 | 650.6311998 | 723.4643722 | 4.09E-06 | 0.600809528 | 1.22E-06 | 4.852299981 | 1.516567307 |
| CNAG_02211 | - | hypothetical protein | 179.0887678 | 186.5722817 | 193.1649482 | 339.3199517 | 312.571887 | 281.8538858 | 4.15E-06 | 0.724832937 | 1.24E-06 | 4.849375897 | 1.652709242 |
| CNAG_04459 | - | hypothetical protein | 9.947580209 | 12.12099562 | 19.37113262 | 61.87045709 | 61.07238767 | 44.23296899 | 4.28E-06 | 2.037661425 | 1.28E-06 | 4.843368027 | 4.105794513 |
| CNAG_06196 | - | synaptosomal-associated protein, 23kDa | 1035.561672 | 952.4137384 | 1131.341255 | 794.36439 | 744.1524406 | 751.7608747 | 4.32E-06 | -0.461605283 | 1.29E-06 | -4.841241218 | 0.726177793 |
| CNAG_13017 | - | unspecified product | 25.23467369 | 52.95647847 | 38.614945 | 125.8951961 | 102.2297338 | 89.72830191 | 4.36E-06 | 1.445249107 | 1.30E-06 | 4.839198823 | 2.723098383 |
| CNAG_02884 | - | DNA replication complex GINS protein PSF2 | 205.8194099 | 188.7307558 | 175.0818393 | 321.9902569 | 289.1278308 | 340.4645593 | 4.38E-06 | 0.726826983 | 1.31E-06 | 4.838296886 | 1.654995143 |
| CNAG_03178 | - | hypothetical protein | 712.9797033 | 606.5108132 | 611.0005094 | 462.760629 | 462.9963132 | 396.6052287 | 4.39E-06 | -0.563311566 | 1.32E-06 | -4.837462666 | 0.676746973 |
| CNAG_02311 | - | hypothetical protein | 5.214333492 | 9.917840828 | 8.731391136 | 33.10176775 | 46.43722222 | 43.91365511 | 4.39E-06 | 2.456997425 | 1.31E-06 | 4.837504086 | 5.490727924 |
| CNAG_03290 | KIC102 | ste/ste20/ysk protein kinase | 853.8186517 | 771.5910129 | 821.2660313 | 1090.666141 | 1144.00854 | 1080.05942 | 4.39E-06 | 0.42301616 | 1.31E-06 | 4.837564194 | 1.34072761 |
| CNAG_01996 | - | hypothetical protein, variant | 264.3248207 | 272.7659454 | 300.8457424 | 385.3821776 | 476.5604569 | 461.5329001 | 4.40E-06 | 0.645787442 | 1.32E-06 | 4.837266056 | 1.564593029 |
| CNAG_07483 | - | DNA polymerase zeta subunit | 1623.438007 | 1807.982392 | 1615.783327 | 2192.741688 | 2156.560767 | 2126.9592 | 4.43E-06 | 0.345075476 | 1.33E-06 | 4.835599683 | 1.270217433 |
| CNAG_13163 | - | unspecified product | 8.711390879 | 11.00626201 | 9.806828891 | 47.70505547 | 41.8278364 | 45.09997571 | 4.44E-06 | 2.233630523 | 1.33E-06 | 4.835320361 | 4.70316036 |
| CNAG_04243 | CDC24 | cell division control protein 24 | 1734.946057 | 1719.436754 | 1474.894757 | 1329.914986 | 1175.419495 | 1206.924163 | 4.45E-06 | -0.424307562 | 1.33E-06 | -4.834831481 | 0.745196311 |
| CNAG_05656 | - | hypothetical protein | 54.2798942 | 57.34946421 | 70.53472952 | 125.8447373 | 143.7490068 | 131.7846594 | 4.48E-06 | 1.124975841 | 1.34E-06 | 4.833270388 | 2.180978943 |
| CNAG_02233 | MEC1 | serine/threonine-protein kinase ATR | 2893.836826 | 2849.886952 | 2657.707141 | 3545.374631 | 3484.019329 | 3468.040416 | 4.50E-06 | 0.306469628 | 1.35E-06 | 4.832386369 | 1.236677761 |
| CNAG_02708 | - | prenylcysteine oxidase/farnesylcysteine lyase | 412.7083848 | 557.0957239 | 489.6242505 | 692.2029707 | 730.8663148 | 738.0220642 | 4.54E-06 | 0.552950187 | 1.36E-06 | 4.830565982 | 1.467082689 |
| CNAG_06174 | - | PEK/GCN2 protein kinase | 4664.980299 | 4498.224534 | 4530.240818 | 3699.73297 | 3680.381441 | 3914.501864 | 4.57E-06 | -0.293223873 | 1.37E-06 | -4.82896516 | 0.8160764 |
| CNAG_04984 | PDR16 | pleiotropic drug resistance protein | 2082.032202 | 2182.806072 | 2413.24238 | 1693.230712 | 1825.061722 | 1697.597247 | 4.60E-06 | -0.371859243 | 1.38E-06 | -4.827690231 | 0.772785943 |
| CNAG_00905 | - | MFS transporter | 73.01075918 | 100.3715814 | 95.08041239 | 166.2163415 | 169.3640388 | 205.9659259 | 4.61E-06 | 0.996625627 | 1.38E-06 | 4.827201578 | 1.995327593 |
| CNAG_02554 | - | Sugar transporter | 387.4194825 | 379.148634 | 433.8434783 | 260.2912668 | 235.1204123 | 277.7050735 | 4.61E-06 | -0.650421981 | 1.39E-06 | -4.826801775 | 0.637093939 |
| CNAG_02934 | - | hypothetical protein, variant | 58.92570758 | 78.79443361 | 43.9978826 | 159.1220686 | 144.5596239 | 118.6340271 | 4.61E-06 | 1.219968942 | 1.39E-06 | 4.826860029 | 2.329417025 |
| CNAG_06585 | - | pre-mRNA-splicing helicase BRR2 | 26305.73618 | 25120.27764 | 24640.36833 | 19677.94892 | 21228.36548 | 22075.61279 | 4.65E-06 | -0.287740887 | 1.40E-06 | -4.825028439 | 0.819183813 |
| CNAG_00307 | - | hypothetical protein | 617.3558689 | 556.949357 | 583.243513 | 424.6237274 | 403.4262073 | 402.0827165 | 4.70E-06 | -0.530434508 | 1.42E-06 | -4.822862238 | 0.692346183 |
| CNAG_06359 | - | scavenger mRNA-decapping enzyme DcpS | 203.6947659 | 284.5024678 | 201.7637207 | 397.5426571 | 362.4996783 | 394.9229309 | 4.74E-06 | 0.73131556 | 1.43E-06 | 4.821131196 | 1.660152256 |
| CNAG_03465 | LAC1 | laccase | 393.7456051 | 360.0124852 | 293.4856349 | 492.0547176 | 548.0049267 | 620.1758553 | 4.80E-06 | 0.651077664 | 1.45E-06 | 4.818533792 | 1.57034077 |
| CNAG_01944 | - | hypothetical protein | 3.935449359 | 3.438970187 | 0.145259058 | 33.0571198 | 34.4052314 | 19.8827657 | 4.86E-06 | 3.82607178 | 1.46E-06 | 4.816116886 | 14.18281282 |
| CNAG_00661 | - | hypothetical protein | 123.041252 | 138.0559686 | 107.9248859 | 244.4308476 | 210.7179659 | 220.6608023 | 4.91E-06 | 0.858507834 | 1.48E-06 | 4.81368051 | 1.813162005 |
| CNAG_02764 | PXA2 | ATP-binding cassette, subfamily D (ALD), peroxisomal long-chain fatty acid import protein | 312.0341279 | 263.0951624 | 290.2068644 | 452.6753441 | 435.8326265 | 447.1697394 | 4.93E-06 | 0.609192771 | 1.49E-06 | 4.813092402 | 1.525405462 |
| CNAG_07560 | - | hypothetical protein | 827.6539507 | 806.8897054 | 799.7548553 | 598.9560726 | 557.8015272 | 626.2537518 | 4.93E-06 | -0.4646279 | 1.49E-06 | -4.81283133 | 0.724657957 |
| CNAG_07926 | - | glycylpeptide N-tetradecanoyltransferase | 2582.852705 | 2368.171301 | 2676.684412 | 2094.189259 | 1983.709352 | 1942.945336 | 5.08E-06 | -0.357121572 | 1.53E-06 | -4.806770556 | 0.780717454 |
| CNAG_08015 | - | hypothetical protein | 2.820897899 | 4.535761152 | 4.460636953 | 56.11773611 | 26.10388462 | 19.98205819 | 5.10E-06 | 3.322135933 | 1.54E-06 | 4.805934139 | 10.00144072 |
| CNAG_03448 | - | translation initiation factor eIF-2B subunit delta | 1043.769267 | 1089.152513 | 1027.992778 | 750.6391165 | 797.6782486 | 838.2791484 | 5.10E-06 | -0.420463842 | 1.54E-06 | -4.805712369 | 0.747184358 |
| CNAG_05085 | - | deoxyhypusine hydroxylase | 609.3880352 | 614.0473796 | 696.2157883 | 440.096994 | 446.5638957 | 466.4256929 | 5.17E-06 | -0.520576182 | 1.56E-06 | -4.80319595 | 0.697093373 |
| CNAG_02580 | LPI3 | hypothetical protein | 665.3101258 | 668.9378418 | 649.3669972 | 433.8152222 | 401.5933178 | 524.0670479 | 5.17E-06 | -0.560344106 | 1.56E-06 | -4.802862718 | 0.678140397 |
| CNAG_06107 | - | G protein beta subunit-like | 1286.63857 | 1233.593513 | 1303.107186 | 959.2997918 | 950.3263859 | 1027.001852 | 5.18E-06 | -0.395838752 | 1.57E-06 | -4.80263918 | 0.760047373 |
| CNAG_07959 | - | hypothetical protein | 1438.578793 | 1585.852363 | 1550.531057 | 1188.17205 | 1114.187476 | 1237.928872 | 5.22E-06 | -0.384523612 | 1.58E-06 | -4.801069469 | 0.766031906 |
| CNAG_03894 | PDR802 | Putative Zn2-Cys6 zinc-finger transcription factor | 14.66098908 | 12.16639184 | 32.13115168 | 79.01121909 | 56.26831839 | 81.5297558 | 5.22E-06 | 1.881082575 | 1.58E-06 | 4.800737137 | 3.683513614 |
| CNAG_06000 | - | glycoprotein | 217.6089556 | 236.1052344 | 223.0401613 | 405.4964087 | 374.6927656 | 319.6540862 | 5.23E-06 | 0.684729214 | 1.59E-06 | 4.800180244 | 1.607400252 |
| CNAG_06821 | FRE2 | ferric reductase transmembrane component 4, variant | 602.2217008 | 609.6585984 | 551.2905749 | 376.182651 | 451.8539408 | 388.7846257 | 5.25E-06 | -0.551569402 | 1.59E-06 | -4.799627968 | 0.682277525 |
| CNAG_03830 | - | hypothetical protein, variant 2 | 11.1934181 | 29.28080046 | 22.60918005 | 74.94440079 | 62.43552507 | 77.18061938 | 5.30E-06 | 1.768019081 | 1.61E-06 | 4.797366892 | 3.405859877 |
| CNAG_01283 | CAP5 | beta-1,2-xylosyltransferase | 1384.858301 | 1529.82192 | 1465.227442 | 1173.436399 | 1113.367104 | 1128.07155 | 5.31E-06 | -0.37361239 | 1.61E-06 | -4.797122215 | 0.771847433 |
| CNAG_05891 | - | TDG/mug DNA glycosylase | 263.0001747 | 235.0579011 | 219.8685062 | 360.6560881 | 370.2707806 | 427.0255959 | 5.39E-06 | 0.674981275 | 1.63E-06 | 4.794153443 | 1.59657605 |
| CNAG_04567 | - | drug transporter | 152.3901319 | 218.8247358 | 191.047469 | 295.2826107 | 318.6002558 | 354.8433898 | 5.41E-06 | 0.771626202 | 1.64E-06 | 4.793138669 | 1.707193043 |
| CNAG_05667 | ITR3B | myo-inositol transporter, putative | 276.1088979 | 347.0625343 | 352.0335936 | 582.2291201 | 469.3803855 | 488.3040496 | 5.43E-06 | 0.644110308 | 1.65E-06 | 4.792195099 | 1.562775246 |
| CNAG_00743 | - | imidazoleglycerol phosphate synthase, cyclase subunit | 4673.30019 | 5290.064727 | 5190.333565 | 3683.237491 | 4406.779852 | 3857.97384 | 5.43E-06 | -0.358067735 | 1.65E-06 | -4.792205184 | 0.780208848 |
| CNAG_07853 | - | hypothetical protein | 18.06159575 | 21.77021486 | 15.16785528 | 74.49341665 | 55.53828369 | 58.5173607 | 5.44E-06 | 1.777588018 | 1.65E-06 | 4.791929017 | 3.428524943 |
| CNAG_04320 | CPS1 | polysaccharide synthase Cps1p | 692.8682727 | 838.3107978 | 731.717065 | 1210.957181 | 1009.326247 | 1002.379924 | 5.53E-06 | 0.496222128 | 1.68E-06 | 4.788482751 | 1.410515117 |
| CNAG_01942 | - | hypothetical protein | 295.6311855 | 237.2395801 | 219.8927259 | 471.7610913 | 393.5545777 | 382.859233 | 5.53E-06 | 0.714554667 | 1.68E-06 | 4.788383017 | 1.640976601 |
| CNAG_05341 | - | hypothetical protein | 79.79161931 | 70.25903762 | 59.95662334 | 132.6760787 | 180.1913152 | 139.638483 | 5.54E-06 | 1.10068498 | 1.68E-06 | 4.788008611 | 2.144564906 |
| CNAG_00504 | - | peptidyl-prolyl cis-trans isomerase-like 4 | 1187.453289 | 1210.947449 | 1308.402476 | 964.8334525 | 977.143683 | 835.0842112 | 5.55E-06 | -0.43191915 | 1.69E-06 | -4.787526218 | 0.741275046 |
| CNAG_00483 | ACT1 | actin | 3468.413405 | 3516.817703 | 3307.172975 | 4676.409452 | 4084.284765 | 4268.469609 | 5.55E-06 | 0.325227853 | 1.69E-06 | 4.787315565 | 1.252862295 |
| CNAG_04700 | - | hypothetical protein | 259.5690056 | 272.715079 | 231.5980625 | 405.2026154 | 371.9298945 | 424.8819431 | 5.58E-06 | 0.640436246 | 1.70E-06 | 4.786219711 | 1.558800443 |
| CNAG_01867 | - | mitochondrial division protein 1 | 2334.022757 | 2083.71087 | 2153.10825 | 1649.623216 | 1750.847323 | 1771.927193 | 5.65E-06 | -0.361454298 | 1.72E-06 | -4.783544914 | 0.778379545 |
| CNAG_03547 | - | hypothetical protein | 67.19644634 | 105.7174081 | 84.44277666 | 158.3975094 | 186.8452757 | 179.4819641 | 5.73E-06 | 1.016837189 | 1.75E-06 | 4.780616289 | 2.023478035 |
| CNAG_03271 | - | pre-mRNA-splicing factor prp46 | 4782.824466 | 4403.407687 | 4289.24407 | 3529.245808 | 3582.909475 | 3793.501105 | 5.79E-06 | -0.320802763 | 1.77E-06 | -4.778430805 | 0.80062426 |
| CNAG_01521 | - | metallo-beta-lactamase | 339.4582646 | 334.9287131 | 341.1192039 | 193.1943163 | 209.2971205 | 237.7207658 | 5.84E-06 | -0.681375104 | 1.78E-06 | -4.776863253 | 0.623570635 |
| CNAG_04620 | - | hypothetical protein | 955.4595031 | 961.1760371 | 967.3610197 | 1281.387152 | 1249.146874 | 1253.240819 | 5.85E-06 | 0.376751409 | 1.78E-06 | 4.7764777 | 1.298414853 |
| CNAG_01228 | - | hypothetical protein | 407.5008567 | 522.3085831 | 490.4124379 | 307.746147 | 310.2466881 | 321.0301725 | 5.85E-06 | -0.611507908 | 1.79E-06 | -4.776138142 | 0.654512247 |
| CNAG_00308 | - | gly-X carboxypeptidase | 362.153179 | 279.2656403 | 283.8510435 | 497.8670206 | 464.2746304 | 491.552107 | 5.87E-06 | 0.636056058 | 1.79E-06 | 4.775628037 | 1.554074922 |
| CNAG_06309 | - | hypothetical protein | 43.76189723 | 39.07147091 | 58.79463207 | 122.393547 | 110.8365819 | 105.1927568 | 5.96E-06 | 1.25151427 | 1.82E-06 | 4.772249161 | 2.380911952 |
| CNAG_02904 | - | hypothetical protein | 319.0615906 | 270.6422926 | 310.4563371 | 483.8870441 | 473.8615321 | 429.507841 | 6.15E-06 | 0.60752186 | 1.88E-06 | 4.766006139 | 1.52363978 |
| CNAG_06459 | - | vacuolar membrane protein | 1078.142825 | 1221.871953 | 1133.736171 | 1510.959585 | 1544.204557 | 1446.446476 | 6.22E-06 | 0.376066966 | 1.90E-06 | 4.763684729 | 1.297799006 |
| CNAG_01711 | - | glycerol-3-phosphate O- acyltransferase/dihydroxyacetone phosphate acyltransferase | 3234.44895 | 3260.199588 | 2909.273169 | 2526.855894 | 2559.076943 | 2523.522689 | 6.25E-06 | -0.320701287 | 1.91E-06 | -4.762443165 | 0.800680576 |
| CNAG_04014 | - | 26S proteasome regulatory subunit N9 | 4812.437513 | 4341.102471 | 4671.135043 | 5557.13551 | 5872.48997 | 5831.545179 | 6.38E-06 | 0.304480445 | 1.95E-06 | 4.758438542 | 1.234973808 |
| CNAG_05914 | - | MFS transporter, SP family, general alpha glucoside:H symporter | 275.8219818 | 242.5987437 | 226.2674741 | 386.4448146 | 371.9582235 | 427.0680717 | 6.38E-06 | 0.656837465 | 1.95E-06 | 4.758218166 | 1.576622715 |
| CNAG_04102 | - | hypothetical protein | 332.4799434 | 355.3361485 | 333.6663395 | 218.2782363 | 237.0252877 | 166.8984385 | 6.44E-06 | -0.73121689 | 1.97E-06 | -4.756232636 | 0.602395589 |
| CNAG_00470 | - | charged multivesicular body protein 1 | 812.1702968 | 631.3428468 | 700.5556111 | 482.896784 | 444.7439559 | 530.7862808 | 6.48E-06 | -0.573678964 | 1.99E-06 | -4.754928974 | 0.671901211 |
| CNAG_05627 | - | ATP-dependent rRNA helicase SPB4 | 1547.145133 | 1562.210355 | 1749.902825 | 1180.283143 | 1186.652164 | 1336.710765 | 6.55E-06 | -0.40726979 | 2.01E-06 | -4.752613072 | 0.754049015 |
| CNAG_01569 | - | hypothetical protein | 1005.250896 | 1035.301444 | 1013.05205 | 767.8479907 | 795.9826949 | 773.9363195 | 6.64E-06 | -0.401961516 | 2.04E-06 | -4.749796258 | 0.756828585 |
| CNAG_13199 | - | unspecified product | 5.109777188 | 0.175928197 | 4.447816883 | 22.05071044 | 37.01882975 | 34.97924178 | 6.66E-06 | 3.350648431 | 2.04E-06 | 4.74906455 | 10.20106894 |
| CNAG_07374 | - | DNA-directed RNA polymerase III subunit RPC5 | 1091.720589 | 1168.895009 | 1190.045342 | 789.4990641 | 942.6843892 | 852.7718809 | 6.67E-06 | -0.432338673 | 2.05E-06 | -4.748730317 | 0.74105952 |
| CNAG_01746 | - | E3 ubiquitin-protein ligase RNF14 | 639.1742692 | 745.6885767 | 775.3807071 | 979.3722231 | 967.0864775 | 1062.175144 | 6.68E-06 | 0.463774546 | 2.05E-06 | 4.748283468 | 1.379145382 |
| CNAG_06862 | - | vacuolar protein, variant 2 | 73.00737365 | 86.42249802 | 106.7648689 | 160.6966324 | 171.9847509 | 207.0593765 | 6.70E-06 | 1.006800209 | 2.06E-06 | 4.747805153 | 2.009449344 |
| CNAG_12932 | - | unspecified product | 3.985694527 | 5.605191363 | 5.520388303 | 41.05878425 | 23.41991742 | 35.06660477 | 6.80E-06 | 2.915086526 | 2.09E-06 | 4.744611505 | 7.54272865 |
| CNAG_03717 | - | hypothetical protein | 71.81393001 | 78.89247585 | 101.4315616 | 158.4531511 | 160.7436132 | 193.7649725 | 6.81E-06 | 1.016362254 | 2.09E-06 | 4.744222427 | 2.022812017 |
| CNAG_00766 | - | hypothetical protein | 242.1019169 | 256.5734391 | 253.9446523 | 366.0859311 | 406.6188827 | 392.7632217 | 6.86E-06 | 0.617079703 | 2.11E-06 | 4.74263279 | 1.533767391 |
| CNAG_04510 | VPS4 | vacuolar protein-sorting-associated protein 4 | 3286.87871 | 2886.40021 | 2917.773283 | 2413.997116 | 2341.658083 | 2465.77645 | 6.96E-06 | -0.348019512 | 2.14E-06 | -4.739636768 | 0.785661891 |
| CNAG_03051 | - | polyamine transporter | 404.4552726 | 423.598402 | 494.9002398 | 607.2846516 | 718.0696654 | 635.9565387 | 6.97E-06 | 0.553740657 | 2.15E-06 | 4.73917497 | 1.467886741 |
| CNAG_01636 | - | peptidyl-prolyl cis-trans isomerase-like 1 | 1508.522118 | 1453.390313 | 1504.682991 | 1170.276374 | 1128.883838 | 1205.742249 | 7.01E-06 | -0.365396271 | 2.16E-06 | -4.737962898 | 0.776255628 |

| | | | | | | | | | | | | | |
|---|---|---|---|---|---|---|---|---|---|---|---|---|---|
| CNAG_02476 | YRM101 | putative transcription factor | 156.8495845 | 162.8455318 | 140.9593446 | 271.4676937 | 239.1075748 | 277.2032085 | 7.10E-06 | 0.766242266 | 2.19E-06 | 4.735445223 | 1.70083391 |
| CNAG_00556 | CCK1 | Casein kinase I | 1179.631745 | 1124.96222 | 1160.390298 | 1595.133882 | 1526.055701 | 1426.491814 | 7.11E-06 | 0.377392235 | 2.19E-06 | 4.735133132 | 1.29899172 |
| CNAG_04171 | - | hypothetical protein | 228.122771 | 259.7842311 | 215.6085188 | 458.6623627 | 366.8117387 | 338.5180759 | 7.13E-06 | 0.712459776 | 2.20E-06 | 4.734274809 | 1.638595521 |
| CNAG_01744 | HAD1 | phosphatase | 2050.133261 | 1918.769906 | 1594.364159 | 1397.497124 | 1423.052534 | 1344.593231 | 7.17E-06 | -0.433011298 | 2.21E-06 | -4.733014584 | 0.740714098 |
| CNAG_02428 | - | GC-rich sequence DNA-binding factor | 2727.759156 | 2840.026298 | 2853.75211 | 2237.310352 | 2182.081445 | 2400.236699 | 7.25E-06 | -0.319235485 | 2.24E-06 | -4.730861948 | 0.801494494 |
| CNAG_00847 | - | hypothetical protein | 719.1329161 | 732.5441021 | 749.6042937 | 513.2547906 | 545.8065497 | 555.221273 | 7.25E-06 | -0.46296224 | 2.24E-06 | -4.730726372 | 0.725495092 |
| CNAG_07825 | - | hypothetical protein | 1844.921258 | 1986.660731 | 1863.036493 | 1519.753745 | 1425.589146 | 1568.750061 | 7.34E-06 | -0.35017982 | 2.26E-06 | -4.728256986 | 0.784486312 |
| CNAG_01347 | - | co-chaperone | 3533.378221 | 3286.121884 | 3323.02029 | 2771.260794 | 2730.797963 | 2799.930942 | 7.36E-06 | -0.304662599 | 2.27E-06 | -4.727473405 | 0.809631542 |
| CNAG_00846 | - | hypothetical protein | 602.2541969 | 560.1881569 | 651.4154266 | 347.255449 | 403.4128806 | 464.1397226 | 7.38E-06 | -0.594166331 | 2.28E-06 | -4.726954117 | 0.662427132 |
| CNAG_05846 | - | ubiquitin-conjugating enzyme E2 O | 1667.234719 | 1453.431325 | 1603.852034 | 1230.205213 | 1203.942673 | 1217.990525 | 7.48E-06 | -0.387644812 | 2.31E-06 | -4.724024769 | 0.764376425 |
| CNAG_04563 | - | fungal specific transcription factor | 810.7964196 | 976.2160565 | 895.9216208 | 1290.887876 | 1180.1343 | 1175.544682 | 7.51E-06 | 0.428684331 | 2.32E-06 | 4.72315114 | 1.346005525 |
| CNAG_06670 | - | HIV Tat-specific factor 1 | 1645.108325 | 1566.513038 | 1546.303707 | 1271.485105 | 1264.358515 | 1222.44268 | 7.52E-06 | -0.355912208 | 2.33E-06 | -4.722830299 | 0.781375428 |
| CNAG_02544 | - | DNA repair protein Swi5/Sae3 | 21.47828721 | 19.62498619 | 4.531530614 | 52.64901898 | 72.42403337 | 63.91568331 | 7.53E-06 | 2.072460761 | 2.33E-06 | 4.722365156 | 4.206034726 |
| CNAG_12420 | - | unspecified product | 105.6137169 | 109.5585182 | 87.33424589 | 46.10839677 | 40.10364327 | 40.30825016 | 7.53E-06 | -1.271167001 | 2.33E-06 | -4.722407436 | 0.414324489 |
| CNAG_01754 | - | hypothetical protein | 1187.724158 | 1052.783986 | 1032.45028 | 1446.35188 | 1437.195421 | 1467.470961 | 7.61E-06 | 0.395660922 | 2.36E-06 | 4.720060916 | 1.315545295 |
| CNAG_04704 | - | MFS transporter, SHS family, lactate transporter | 0.35539212 | 3.440808875 | 3.379057055 | 13.00152232 | 44.34129813 | 32.77982835 | 7.83E-06 | 3.876627219 | 2.42E-06 | 4.714369311 | 14.68862268 |
| CNAG_12782 | - | unspecified product | 26.43989074 | 61.57051519 | 50.31943364 | 103.1746419 | 119.5445807 | 133.7541151 | 7.87E-06 | 1.360917294 | 2.44E-06 | 4.713123983 | 2.56848437 |
| CNAG_06809 | IKS1 | IKS protein kinase | 1325.118998 | 1301.425892 | 1194.394939 | 1003.74081 | 965.8681926 | 980.4193094 | 7.96E-06 | -0.389015258 | 2.47E-06 | -4.710829782 | 0.763650673 |
| CNAG_07857 | - | hypothetical protein | 867.8105545 | 738.2152369 | 803.1484629 | 1222.508433 | 1104.270374 | 1032.378963 | 7.99E-06 | 0.463833738 | 2.48E-06 | 4.7099706 | 1.379201968 |
| CNAG_02531 | CPK2 | Calcium-dependent protein kinase 2 | 48.40772639 | 49.79965331 | 56.69034231 | 119.8642076 | 105.5111975 | 123.9064483 | 8.03E-06 | 1.16633065 | 2.49E-06 | 4.708739302 | 2.244401294 |
| CNAG_07850 | - | hypothetical protein | 0.334307757 | 3.42731199 | 0.137978055 | 18.93285015 | 31.79601062 | 32.69296337 | 8.06E-06 | 4.742537126 | 2.50E-06 | 4.707989019 | 26.76984956 |
| CNAG_00385 | - | 20S proteasome subunit beta 3 | 2235.270451 | 2248.686557 | 2243.89213 | 2828.570129 | 2758.594284 | 2755.321773 | 8.17E-06 | 0.295261342 | 2.54E-06 | 4.70523902 | 1.227107245 |
| CNAG_03186 | - | THO complex subunit 4 | 1967.519142 | 1998.557423 | 2073.088727 | 1560.170603 | 1487.679652 | 1706.381928 | 8.17E-06 | -0.360284823 | 2.54E-06 | -4.705146416 | 0.779010769 |
| CNAG_01579 | - | vacuolar membrane-associated protein IML1 | 609.9789689 | 724.1283165 | 725.2678812 | 893.6509816 | 982.7159147 | 992.2636055 | 8.45E-06 | 0.463603365 | 2.63E-06 | 4.698065324 | 1.378981751 |
| CNAG_06076 | - | cytoplasmic protein | 963.2534565 | 1010.524009 | 1022.627123 | 742.9210454 | 777.002195 | 766.1560441 | 8.45E-06 | -0.405485037 | 2.63E-06 | -4.698101401 | 0.754982424 |
| CNAG_04515 | - | hypothetical protein, variant | 182.6845413 | 238.2233107 | 218.7577613 | 358.5996022 | 325.3867879 | 354.9761763 | 8.47E-06 | 0.686574772 | 2.63E-06 | 4.697536135 | 1.609457825 |
| CNAG_07505 | - | hypothetical protein | 7.548288752 | 2.416971007 | 14.02836589 | 40.37299225 | 56.16727269 | 37.49334335 | 8.55E-06 | 2.51338073 | 2.66E-06 | 4.695390001 | 5.709564588 |
| CNAG_12434 | - | unspecified product | 13437.19832 | 15878.98142 | 18965.43854 | 9301.58117 | 7234.632141 | 9655.312596 | 8.56E-06 | -0.896802183 | 2.67E-06 | -4.694928845 | 0.537075873 |
| CNAG_04028 | - | RNA binding protein, variant | 14089.97077 | 13557.3058 | 14673.25112 | 11747.28476 | 11934.27209 | 12046.44644 | 8.56E-06 | -0.259801022 | 2.67E-06 | -4.694985335 | 0.835203103 |
| CNAG_07914 | - | protein phosphatase 2 (formerly 2A), regulatory subunit A | 6083.841915 | 5993.701587 | 5997.686104 | 4962.129748 | 5139.026417 | 5155.674654 | 8.62E-06 | -0.259779064 | 2.69E-06 | -4.693482452 | 0.835215816 |
| CNAG_01058 | - | nitric oxide synthase-interacting protein | 792.8491604 | 883.3772247 | 976.6864528 | 683.8843539 | 612.1554434 | 617.4530533 | 8.65E-06 | -0.486403712 | 2.70E-06 | -4.692671583 | 0.713802217 |
| CNAG_06400 | PMA1 | plasma-membrane proton-efflux P-type ATPase | 20701.47272 | 18893.74282 | 15846.45928 | 14740.57703 | 14722.04143 | 13649.61842 | 8.81E-06 | -0.378410838 | 2.75E-06 | -4.688882473 | 0.769284509 |
| CNAG_04827 | - | hypothetical protein | 247.9392101 | 275.9391943 | 250.7607719 | 382.485597 | 403.1159715 | 400.5372598 | 8.85E-06 | 0.603374321 | 2.76E-06 | 4.687922881 | 1.519265827 |
| CNAG_03785 | NTH1 | alpha,alpha-trehalase | 1750.575448 | 1633.486862 | 1479.306668 | 2103.205428 | 2146.248792 | 2081.433459 | 8.93E-06 | 0.365217167 | 2.79E-06 | 4.686058634 | 1.288075508 |
| CNAG_01590 | - | large subunit ribosomal protein L32 | 206.0282256 | 172.2548866 | 179.0197217 | 102.1882836 | 97.09471284 | 104.6092337 | 9.23E-06 | -0.894161699 | 2.88E-06 | -4.679109145 | 0.538059753 |
| CNAG_13013 | - | unspecified product | 251.8862643 | 262.692726 | 253.6720408 | 170.8821758 | 150.6237055 | 139.0777088 | 9.31E-06 | -0.753935239 | 2.91E-06 | -4.677351597 | 0.592983868 |
| CNAG_05314 | GLO3 | ADP-ribosylation factor GTPase-activating protein 2/3 | 2177.567823 | 2019.042672 | 2065.646745 | 1429.227816 | 1758.686338 | 1632.069483 | 9.35E-06 | -0.393403139 | 2.92E-06 | -4.676274927 | 0.761331598 |
| CNAG_01605 | - | rossman fold oxidoreductase | 54.53883344 | 134.8016418 | 113.219115 | 261.9954984 | 216.9107296 | 196.3057238 | 9.39E-06 | 1.148457751 | 2.93E-06 | 4.675387422 | 2.21676794 |
| CNAG_06072 | - | Chorismate mutase | 1114.832341 | 1009.47481 | 982.1555045 | 779.5303291 | 767.4664094 | 795.0149205 | 1.00E-05 | -0.422992182 | 3.14E-06 | -4.661408006 | 0.745876055 |
| CNAG_00745 | HRK1 | HAL protein kinase | 1787.534245 | 1596.718469 | 1716.905257 | 1343.850701 | 1257.381827 | 1372.266823 | 1.01E-05 | -0.375968871 | 3.17E-06 | -4.659733301 | 0.770587735 |
| CNAG_02782 | - | cytoplasmic protein | 1014.68878 | 1083.800173 | 1182.543047 | 779.4427362 | 880.5593088 | 793.950414 | 1.01E-05 | -0.434586684 | 3.17E-06 | -4.659305244 | 0.739905699 |
| CNAG_12552 | - | unspecified product | 781.1476405 | 859.6723828 | 912.7333024 | 620.721806 | 612.1847182 | 650.6888445 | 1.02E-05 | -0.454285191 | 3.20E-06 | -4.657806895 | 0.72987171 |
| CNAG_03294 | - | hypothetical protein | 133.5360834 | 155.2672355 | 106.8804919 | 229.2850539 | 224.5213802 | 263.7880925 | 1.03E-05 | 0.847596959 | 3.21E-06 | 4.65668082 | 1.799501068 |
| CNAG_07425 | - | hypothetical protein | 8.815782461 | 19.61613729 | 20.44672049 | 51.90833654 | 60.95263145 | 64.96272784 | 1.04E-05 | 1.883013146 | 3.25E-06 | 4.65411814 | 3.688446081 |
| CNAG_04886 | - | hypothetical protein | 310.9784403 | 365.3210404 | 221.0161411 | 562.0196025 | 505.8018682 | 441.7468874 | 1.04E-05 | 0.736911616 | 3.26E-06 | 4.65363638 | 1.666604309 |
| CNAG_04524 | - | zinc metalloprotease | 643.687187 | 650.8805925 | 634.6483276 | 840.7679623 | 931.8667518 | 856.9144906 | 1.04E-05 | 0.431752972 | 3.27E-06 | 4.652931044 | 1.348871552 |
| CNAG_02788 | - | hypothetical protein | 155.9168532 | 211.3306833 | 228.3171248 | 298.2946916 | 365.3321198 | 353.8147741 | 1.06E-05 | 0.759825776 | 3.31E-06 | 4.650348708 | 1.693286126 |
| CNAG_06129 | - | cytoplasmic protein | 902.4924841 | 1015.824353 | 861.6565618 | 704.8433753 | 669.9773416 | 694.0068981 | 1.06E-05 | -0.441070173 | 3.33E-06 | -4.649171441 | 0.736588014 |
| CNAG_05066 | - | hypothetical protein | 33.44327257 | 56.26833818 | 76.88992129 | 135.4581427 | 123.6945868 | 141.6362429 | 1.06E-05 | 1.259755759 | 3.34E-06 | 4.648707165 | 2.394551989 |
| CNAG_02698 | - | hypothetical protein | 2017.699623 | 2028.719135 | 2024.056339 | 1465.877229 | 1700.876251 | 1639.818761 | 1.07E-05 | -0.352547569 | 3.35E-06 | -4.648324273 | 0.78319987 |
| CNAG_06448 | - | Cystathionine gamma-lyase | 74.00716798 | 71.32727239 | 69.51324244 | 139.8364163 | 132.2081288 | 169.3234432 | 1.08E-05 | 1.024876548 | 3.39E-06 | 4.645947659 | 2.03478526 |
| CNAG_02224 | - | hypothetical protein | 404.3184358 | 404.1649721 | 356.3619696 | 520.0273229 | 566.9141834 | 666.8053252 | 1.09E-05 | 0.574982585 | 3.41E-06 | 4.644218106 | 1.489659482 |
| CNAG_02398 | - | hypothetical protein | 1007.528505 | 1039.581644 | 938.4361328 | 775.6990902 | 772.6914733 | 709.5964385 | 1.09E-05 | -0.418581415 | 3.42E-06 | -4.644098333 | 0.748159919 |
| CNAG_03886 | - | hypothetical protein | 2842.13494 | 2778.637553 | 2858.021944 | 2132.922504 | 2392.646374 | 2340.319629 | 1.10E-05 | -0.32011583 | 3.45E-06 | -4.641993331 | 0.801005565 |
| CNAG_01360 | - | pumilio domain-containing protein c | 1044.205682 | 1142.127913 | 869.3386204 | 1470.651615 | 1379.414227 | 1363.149413 | 1.10E-05 | 0.448773759 | 3.45E-06 | 4.642026916 | 1.364879663 |
| CNAG_03385 | PCL103 | g1/s-specific cyclin pcl1 | 128.9105447 | 163.8644098 | 103.6929387 | 271.7697097 | 240.1358101 | 219.6523766 | 1.11E-05 | 0.875699526 | 3.51E-06 | 4.638528636 | 1.834897566 |
| CNAG_12194 | - | unspecified product | 120.7128467 | 111.7904412 | 99.04726665 | 59.49079065 | 45.27836116 | 39.25068555 | 1.11E-05 | -1.221870388 | 3.51E-06 | -4.638559818 | 0.428726533 |
| CNAG_01272 | - | hypothetical protein | 113.6197712 | 109.0003843 | 87.67343544 | 214.1351147 | 195.3176128 | 174.1399166 | 1.12E-05 | 0.900671786 | 3.53E-06 | 4.637506431 | 1.866935113 |
| CNAG_04952 | - | ADP-ribosylation factor 6 | 2728.921526 | 2883.107829 | 2763.125915 | 2396.998366 | 2259.754458 | 2170.525913 | 1.14E-05 | -0.310096282 | 3.58E-06 | -4.634609794 | 0.806587928 |
| CNAG_04493 | - | ubiquitin carboxyl-terminal hydrolase 48 | 623.9103528 | 718.7140261 | 617.6123518 | 830.5517413 | 943.9151334 | 957.8300184 | 1.14E-05 | 0.464917073 | 3.58E-06 | 4.63447015 | 1.380238014 |
| CNAG_06680 | - | COP9 signalosome complex subunit 6 | 994.7013672 | 1013.743456 | 974.66528 | 689.1137288 | 745.9169327 | 807.1734938 | 1.14E-05 | -0.426717566 | 3.58E-06 | -4.634163941 | 0.743952509 |
| CNAG_03545 | - | hypothetical protein, variant 2 | 78.89991767 | 98.28668715 | 138.7099001 | 197.4227625 | 216.9869915 | 207.2804863 | 1.14E-05 | 0.966383578 | 3.61E-06 | 4.632717284 | 1.953936494 |
| CNAG_03251 | ARF2 | ADP-ribosylation factor | 12521.60625 | 11799.9989 | 12050.87269 | 10488.87513 | 10368.74348 | 9503.142766 | 1.14E-05 | -0.276260712 | 3.61E-06 | -4.632732465 | 0.825728433 |
| CNAG_06973 | - | hypothetical protein | 13.42961667 | 16.41079437 | 20.44459671 | 55.99386766 | 67.17547124 | 50.81977906 | 1.15E-05 | 1.796168416 | 3.64E-06 | 4.631157013 | 3.472966311 |
| CNAG_05924 | - | hypothetical protein | 39.0527177 | 31.53267738 | 47.08158908 | 90.03295279 | 92.66495995 | 112.6688448 | 1.16E-05 | 1.313426918 | 3.65E-06 | 4.630228935 | 2.4853119 |
| CNAG_02994 | - | hypothetical protein, variant | 5871.657373 | 6275.013132 | 5797.326649 | 6993.821339 | 7500.14082 | 7313.634566 | 1.20E-05 | 0.266360552 | 3.79E-06 | 4.622700879 | 1.202769802 |
| CNAG_03070 | - | compass component swd2 | 1096.396669 | 1150.599379 | 1186.85381 | 881.5097186 | 845.9462294 | 917.1022435 | 1.20E-05 | -0.391881417 | 3.80E-06 | -4.621804932 | 0.762135057 |
| CNAG_06190 | - | N-acetylglucosamine-6-phosphate deacetylase | 1096.294684 | 1076.26767 | 1096.225095 | 865.2315352 | 864.1438959 | 758.4651462 | 1.21E-05 | -0.409470679 | 3.83E-06 | -4.620448924 | 0.75289956 |
| CNAG_02479 | - | Monosaccharide transporter | 125.4789355 | 155.3041976 | 153.7057976 | 225.9628632 | 283.4678368 | 249.5343852 | 1.22E-05 | 0.791090775 | 3.85E-06 | 4.619258805 | 1.730382254 |
| CNAG_00676 | - | ESCRT-I complex subunit VPS28 | 744.885346 | 800.403814 | 801.8612205 | 556.8592199 | 571.6592664 | 610.705549 | 1.22E-05 | -0.447924587 | 3.86E-06 | -4.618712499 | 0.733096698 |
| CNAG_01414 | - | hypothetical protein | 2064.427291 | 1981.359742 | 2250.085884 | 1613.082395 | 1762.985157 | 1561.070349 | 1.22E-05 | -0.366605274 | 3.86E-06 | -4.61876247 | 0.775605385 |
| CNAG_05707 | - | cytoplasmic protein | 252.6464143 | 272.7425288 | 285.9179805 | 424.6534894 | 456.7252863 | 372.9368386 | 1.22E-05 | 0.614717932 | 3.88E-06 | 4.617878201 | 1.531258585 |
| CNAG_03297 | MSL1 | histone acetyltransferase type B subunit 2 | 1547.069461 | 1506.180651 | 1544.145817 | 1259.080452 | 1148.707852 | 1219.083139 | 1.22E-05 | -0.357581742 | 3.88E-06 | -4.617932006 | 0.780471718 |
| CNAG_00638 | - | GTPase | 1805.019066 | 1665.633883 | 1591.115852 | 1325.155944 | 1277.243258 | 1361.167734 | 1.23E-05 | -0.368238033 | 3.89E-06 | -4.616980606 | 0.774728096 |
| CNAG_12884 | - | unspecified product | 22.68002801 | 26.05650854 | 13.05554452 | 68.64865728 | 64.30028384 | 64.02425287 | 1.24E-05 | 1.669751984 | 3.94E-06 | 4.614748177 | 3.181598935 |
| CNAG_06550 | - | nucleoporin family protein | 2061.945583 | 1910.21651 | 1832.135956 | 1522.833112 | 1567.132452 | 1523.280132 | 1.24E-05 | -0.346935014 | 3.94E-06 | -4.614750445 | 0.786252708 |
| CNAG_05428 | TAF5 | transcription initiation factor TFIID subunit 5, variant | 2498.766828 | 2314.288254 | 2446.372049 | 1950.140214 | 1979.436895 | 1962.88534 | 1.25E-05 | -0.316670597 | 3.95E-06 | -4.613826296 | 0.802920694 |
| CNAG_01974 | - | large subunit ribosomal protein L36 | 310.3725004 | 297.2926582 | 374.0673708 | 209.6999029 | 202.3986742 | 204.504275 | 1.25E-05 | -0.689283886 | 3.98E-06 | -4.612481575 | 0.620161605 |
| CNAG_03151 | CSN4 | COP9 signalosome complex subunit 4 | 335.4475027 | 338.4486684 | 328.5972989 | 522.0137624 | 492.7701226 | 460.6039646 | 1.26E-05 | 0.543072946 | 4.00E-06 | 4.611449696 | 1.457072785 |
| CNAG_01254 | OCA101 | protein-tyrosine-phosphatase | 68.17055392 | 71.29798568 | 57.80963621 | 151.9456283 | 135.812204 | 124.1411246 | 1.27E-05 | 1.049816781 | 4.04E-06 | 4.60906768 | 2.070266912 |
| CNAG_01910 | - | hypothetical protein | 754.0479563 | 705.6308157 | 744.268158 | 454.7216558 | 531.9911109 | 579.5942856 | 1.29E-05 | -0.509722751 | 4.08E-06 | -4.607056312 | 0.7023574 |
| CNAG_04867 | - | vacuolar protein | 24.05212977 | 24.02786894 | 52.37511335 | 131.733891 | 104.0938252 | 69.9291471 | 1.29E-05 | 1.587717789 | 4.11E-06 | 4.605923529 | 3.005734935 |
| CNAG_06077 | - | mRNA export factor | 2025.770318 | 1852.056779 | 1904.622728 | 1437.128779 | 1580.950438 | 1559.888342 | 1.30E-05 | -0.352895629 | 4.12E-06 | -4.605006783 | 0.78301094 |
| CNAG_07482 | - | SCF-associated factor 1 | 737.0889304 | 785.5170752 | 625.1131116 | 1066.785357 | 961.0460502 | 969.0377401 | 1.30E-05 | 0.466014738 | 4.14E-06 | 4.604449525 | 1.381288558 |
| CNAG_05720 | - | GTPase inhibitor | 1610.207932 | 1776.541034 | 1615.625955 | 1342.321642 | 1366.175685 | 1221.37411 | 1.30E-05 | -0.362953373 | 4.15E-06 | -4.603968633 | 0.777571166 |
| CNAG_04035 | - | hypothetical protein | 955.1501855 | 1063.300071 | 1097.247062 | 802.1514792 | 816.7012333 | 715.1716969 | 1.30E-05 | -0.432107818 | 4.14E-06 | -4.604004882 | 0.741178112 |
| CNAG_05757 | - | hypothetical protein | 414.1415435 | 413.5457469 | 367.856236 | 293.1759529 | 255.0294353 | 228.9962571 | 1.32E-05 | -0.63684962 | 4.19E-06 | -4.601571438 | 0.643115774 |

| | | | | | | | | | | | | | |
|---|---|---|---|---|---|---|---|---|---|---|---|---|---|
| CNAG_04100 | - | hypothetical protein | 97.29067254 | 68.16983896 | 98.25593733 | 173.4118845 | 162.4443742 | 186.0959833 | 1.32E-05 | 0.969907513 | 4.19E-06 | 4.601586197 | 1.958715023 |
| CNAG_07817 | - | centromeric protein E | 3163.602808 | 2970.596718 | 3183.423954 | 3721.920413 | 3957.609822 | 3877.674518 | 1.32E-05 | 0.295115566 | 4.22E-06 | 4.600303236 | 1.22698326 |
| CNAG_02709 | - | small subunit ribosomal protein S19 | 350.9773946 | 311.2545868 | 317.6814324 | 229.3093826 | 207.6024749 | 171.306332 | 1.32E-05 | -0.703951821 | 4.22E-06 | -4.600071251 | 0.613888343 |
| CNAG_03999 | - | hypothetical protein | 36.77717845 | 52.95279765 | 36.4909881 | 109.9158231 | 87.33911851 | 108.3499675 | 1.33E-05 | 1.28016769 | 4.23E-06 | 4.599723987 | 2.428672046 |
| CNAG_07474 | - | AP-2 complex subunit mu-1 | 832.3148953 | 793.9754283 | 790.1663782 | 591.1496033 | 587.1795641 | 625.1457806 | 1.33E-05 | -0.436954488 | 4.24E-06 | -4.599182244 | 0.738692334 |
| CNAG_05639 | PPS1 | hypothetical protein | 1238.181656 | 1456.758492 | 1416.326046 | 1879.18 | 1672.564052 | 1883.747759 | 1.35E-05 | 0.388438461 | 4.29E-06 | 4.596636578 | 1.308975833 |
| CNAG_04540 | - | translocation protein Sec62 | 2358.633002 | 2173.147357 | 2322.657636 | 1817.811214 | 1849.180296 | 1869.621461 | 1.35E-05 | -0.323772309 | 4.31E-06 | -4.5960492 | 0.798978004 |
| CNAG_12217 | - | unspecified product | 336.5716505 | 325.5231649 | 297.6992494 | 519.758485 | 440.0266417 | 480.4931768 | 1.35E-05 | 0.571875189 | 4.31E-06 | 4.595733489 | 1.486454382 |
| CNAG_01398 | - | ATP-dependent RNA helicase DOB1 | 6525.167499 | 5957.095769 | 6334.664187 | 4790.016238 | 5297.742842 | 5312.203686 | 1.38E-05 | -0.304857816 | 4.42E-06 | -4.590505103 | 0.809521995 |
| CNAG_06469 | APT1 | Phospholipid-translocating ATPase APT1 | 2818.677058 | 2845.362376 | 2406.99086 | 2211.70706 | 2080.305948 | 2080.594912 | 1.38E-05 | -0.355846587 | 4.42E-06 | -4.590581242 | 0.781410969 |
| CNAG_00094 | - | NAD-dependent epimerase/dehydratase | 170.7425244 | 145.6477863 | 134.5681983 | 262.8796628 | 248.6928247 | 257.2929719 | 1.38E-05 | 0.756932263 | 4.43E-06 | 4.59032089 | 1.689893424 |
| CNAG_03005 | - | cytoplasmic tRNA 2-thiolation protein 2 | 113.0778446 | 133.3707673 | 154.3074804 | 64.76752673 | 72.15351568 | 54.76982234 | 1.42E-05 | -1.089205092 | 4.54E-06 | -4.584946107 | 0.470020279 |
| CNAG_06660 | - | hypothetical protein | 1026.170058 | 943.7737066 | 1000.240846 | 748.3998582 | 782.1970449 | 727.3362872 | 1.43E-05 | -0.411264439 | 4.58E-06 | -4.583241864 | 0.751964032 |
| CNAG_00160 | - | hypothetical protein | 62.50858447 | 88.51804217 | 88.65977319 | 163.3794591 | 158.236348 | 154.0399963 | 1.43E-05 | 0.982632397 | 4.58E-06 | 4.5829593 | 1.976067731 |
| CNAG_07824 | - | hypothetical protein | 173.2708754 | 218.8032903 | 165.492947 | 297.7294513 | 318.6620243 | 302.8430058 | 1.44E-05 | 0.707534184 | 4.62E-06 | 4.581392373 | 1.633010632 |
| CNAG_12407 | - | unspecified product | 61.61171565 | 100.6578725 | 61.74157666 | 29.74781458 | 20.19094993 | 25.87264804 | 1.44E-05 | -1.597504444 | 4.63E-06 | -4.580957118 | 0.330448089 |
| CNAG_01534 | - | hypothetical protein | 51.02110865 | 105.7814993 | 124.8735302 | 211.7618686 | 209.239868 | 191.7828147 | 1.45E-05 | 1.112060789 | 4.65E-06 | 4.580042245 | 2.161541881 |
| CNAG_00815 | SIT1 | MFS transporter, SIT family, siderophore-iron:H symporter | 492.3970045 | 438.4383657 | 479.7690757 | 290.5353998 | 320.6254874 | 346.503497 | 1.46E-05 | -0.57526582 | 4.67E-06 | -4.579056225 | 0.671162577 |
| CNAG_04805 | - | peptidyl-prolyl isomerase cwc27 | 1197.121199 | 1214.337752 | 1014.350263 | 1464.168904 | 1532.107017 | 1556.266168 | 1.46E-05 | 0.39541652 | 4.68E-06 | 4.57849385 | 1.315322452 |
| CNAG_03272 | - | hypothetical protein | 45.06013263 | 89.51490891 | 52.50154796 | 140.7968538 | 158.5047936 | 128.5512523 | 1.46E-05 | 1.182697462 | 4.68E-06 | 4.578552518 | 2.270008126 |
| CNAG_01357 | SDP101 | hypothetical protein | 96.14907173 | 82.12669225 | 96.13723254 | 156.7689067 | 185.0671985 | 181.7116753 | 1.47E-05 | 0.914005129 | 4.72E-06 | 4.577049135 | 1.884269246 |
| CNAG_06207 | - | hypothetical protein | 5101.867601 | 674.8029419 | 1457.763219 | 190.8979711 | 193.2464418 | 151.6439072 | 1.48E-05 | -3.783267268 | 4.76E-06 | -4.57524552 | 0.072631175 |
| CNAG_12816 | - | unspecified product | 18.14851264 | 29.30076648 | 27.91847249 | 73.1670347 | 60.59167155 | 87.0726107 | 1.48E-05 | 1.558336455 | 4.76E-06 | 4.575013452 | 2.945140489 |
| CNAG_02828 | - | hypothetical protein | 83.59402715 | 131.6052006 | 137.6893051 | 249.2199981 | 202.0562085 | 220.6358711 | 1.48E-05 | 0.920994349 | 4.76E-06 | 4.574911056 | 1.893419846 |
| CNAG_06489 | - | Adenosine kinase | 335.2541864 | 241.2149176 | 211.1385422 | 155.9719154 | 152.3659551 | 122.487747 | 1.49E-05 | -0.89200116 | 4.79E-06 | -4.573720123 | 0.538866139 |
| CNAG_12689 | - | unspecified product | 2.823352522 | 8.806821958 | 4.461829755 | 36.01581209 | 39.57089628 | 24.30677055 | 1.51E-05 | 2.80898057 | 4.85E-06 | 4.571336058 | 7.007892141 |
| CNAG_03482 | TSA1 | Thiol peroxidase | 11796.45518 | 10819.56702 | 12098.82376 | 9784.478814 | 9450.061724 | 9670.716723 | 1.51E-05 | -0.279911426 | 4.86E-06 | -4.570667965 | 0.823641583 |
| CNAG_03971 | - | hypothetical protein | 0.330020538 | 1.2686367 | 3.365108902 | 40.90474151 | 14.65465027 | 24.11163913 | 1.53E-05 | 4.271318414 | 4.92E-06 | 4.568364281 | 19.31056423 |
| CNAG_02420 | - | hypothetical protein, variant | 7.673009952 | 20.70117315 | 23.6355838 | 46.10135387 | 68.82353936 | 75.91687781 | 1.53E-05 | 1.904976717 | 4.94E-06 | 4.56731539 | 3.74502855 |
| CNAG_07458 | - | hypothetical protein, variant | 103.3044207 | 132.6715548 | 111.1040295 | 223.2441679 | 171.6597363 | 269.1505568 | 1.54E-05 | 0.92457184 | 4.96E-06 | 4.566300544 | 1.898120837 |
| CNAG_06777 | - | fructosyl amino acid oxidase | 102.2031274 | 157.4044762 | 126.0196558 | 235.6327412 | 210.6490651 | 261.5541185 | 1.54E-05 | 0.861411925 | 4.97E-06 | 4.565906217 | 1.816815507 |
| CNAG_00263 | PTP4 | serine/threonine/tyrosine-interacting protein | 702.0532271 | 657.3804964 | 710.3445303 | 937.374297 | 885.9565874 | 996.6873433 | 1.56E-05 | 0.43035975 | 5.03E-06 | 4.563401932 | 1.347569565 |
| CNAG_06564 | - | clathrin-coated vesicle protein | 2478.765145 | 2235.596592 | 2089.165713 | 1740.720496 | 1721.464529 | 1865.162667 | 1.56E-05 | -0.368379704 | 5.03E-06 | -4.563453004 | 0.774652023 |
| CNAG_05815 | KRE64 | glucosidase | 138.252113 | 171.419067 | 135.6279185 | 247.1762431 | 270.4207463 | 242.9217187 | 1.56E-05 | 0.759040335 | 5.04E-06 | 4.563160757 | 1.692364508 |
| CNAG_00377 | - | DNA topoisomerase II, variant | 5297.782521 | 4690.029232 | 4740.348399 | 4087.382448 | 3900.323321 | 4024.447008 | 1.58E-05 | -0.309803952 | 5.10E-06 | -4.560535454 | 0.806751381 |
| CNAG_00775 | - | U3 small nucleolar RNA-associated protein 15 | 1600.708373 | 1406.03201 | 1601.701568 | 1159.316694 | 1234.187476 | 1180.252135 | 1.58E-05 | -0.382633456 | 5.12E-06 | -4.559916751 | 0.767036185 |
| CNAG_12915 | - | unspecified product | 197.7111615 | 209.1599167 | 183.6030083 | 296.0370094 | 301.2380537 | 362.5995678 | 1.59E-05 | 0.688557182 | 5.14E-06 | 4.559142177 | 1.611670904 |
| CNAG_06925 | - | arsenical-resistance protein | 1288.999697 | 1272.367365 | 1282.868281 | 998.2581151 | 1053.933999 | 966.0119058 | 1.64E-05 | -0.364151552 | 5.30E-06 | -4.552640021 | 0.77692565 |
| CNAG_04233 | - | hypothetical protein | 146.4926883 | 172.5616821 | 186.7430778 | 246.0754205 | 288.424263 | 332.5734947 | 1.65E-05 | 0.766224353 | 5.32E-06 | 4.551563456 | 1.700812793 |
| CNAG_03204 | - | hypothetical protein | 21.66320989 | 38.95882596 | 29.00973988 | 83.31317025 | 67.47113767 | 97.07453242 | 1.66E-05 | 1.475956689 | 5.38E-06 | 4.549247676 | 2.781680434 |
| CNAG_04282 | MPK2 | CMGC/MAPK protein kinase | 238.6522189 | 304.977871 | 232.6675685 | 384.0147838 | 424.7454383 | 404.9828291 | 1.67E-05 | 0.632746353 | 5.39E-06 | 4.549162568 | 1.550513796 |
| CNAG_02769 | - | hypothetical protein | 845.0786893 | 722.9234502 | 826.3717731 | 549.0406093 | 633.8703813 | 541.9829279 | 1.68E-05 | -0.490078762 | 5.43E-06 | -4.547641377 | 0.711986227 |
| CNAG_02059 | - | CCR4-Not complex subunit Caf16 | 1171.057734 | 1170.001118 | 1153.835348 | 920.4559196 | 921.917552 | 899.3809572 | 1.68E-05 | -0.365956034 | 5.45E-06 | -4.546665061 | 0.775954501 |
| CNAG_03463 | - | hypothetical protein | 1652.510797 | 1596.843035 | 1480.353189 | 2051.098139 | 2022.871426 | 1952.684392 | 1.71E-05 | 0.334413329 | 5.54E-06 | 4.543317482 | 1.260864575 |
| CNAG_01282 | - | hypothetical protein | 1304.26022 | 1552.361048 | 1267.985873 | 999.7220332 | 1023.64891 | 1099.164038 | 1.74E-05 | -0.415928664 | 5.63E-06 | -4.539671019 | 0.749536862 |
| CNAG_00641 | - | transcription elongation factor spt5 | 7307.584923 | 6759.781919 | 7053.467002 | 5696.032165 | 5899.788976 | 6063.775618 | 1.75E-05 | -0.27381686 | 5.66E-06 | -4.5388023 | 0.82712836 |
| CNAG_02230 | - | phosphoketolase | 2491.073691 | 2862.7634 | 2278.090894 | 3565.815246 | 3190.721505 | 3226.027808 | 1.75E-05 | 0.372989133 | 5.68E-06 | 4.538024314 | 1.295033244 |
| CNAG_00057 | FPP1 | fructose-1,6-bisphosphatase I | 2607.818863 | 2102.293438 | 1651.141922 | 3966.104233 | 3949.239587 | 3755.401967 | 1.78E-05 | 0.859686014 | 5.77E-06 | 4.534552676 | 1.814643331 |
| CNAG_03854 | - | lysine decarboxylase | 533.9688664 | 541.0050354 | 577.033046 | 799.7682881 | 714.3071748 | 773.5970348 | 1.78E-05 | 0.454357324 | 5.79E-06 | 4.533904583 | 1.3701723 |
| CNAG_03507 | - | mitochondrial-processing peptidase subunit beta | 1551.453378 | 1408.075007 | 1240.268413 | 1144.748546 | 1011.567947 | 839.5949106 | 1.82E-05 | -0.503624801 | 5.90E-06 | -4.529999094 | 0.705332391 |
| CNAG_07541 | - | proteasome assembly chaperone 2 | 825.798761 | 830.7879566 | 747.7131997 | 1043.15051 | 1060.262754 | 1090.002098 | 1.85E-05 | 0.395601598 | 5.99E-06 | 4.52667916 | 1.3154912 |
| CNAG_00152 | - | hypothetical protein | 500.8832353 | 579.4923002 | 583.2019117 | 402.8307285 | 399.155681 | 371.0197549 | 1.85E-05 | -0.518782526 | 6.02E-06 | -4.525619006 | 0.697960586 |
| CNAG_06330 | - | hypothetical protein | 127.9319896 | 182.2491738 | 208.0367242 | 292.3347425 | 297.0423883 | 312.7368712 | 1.87E-05 | 0.787035087 | 6.07E-06 | 4.523915599 | 1.725524654 |
| CNAG_00523 | - | hypothetical protein, variant | 769.3808654 | 826.2488732 | 782.6974346 | 560.733192 | 603.6135024 | 616.2642234 | 1.91E-05 | -0.433008605 | 6.22E-06 | -4.518657762 | 0.740715481 |
| CNAG_02865 | - | hypothetical protein, variant | 50.7481719 | 64.81068052 | 51.39968869 | 115.7560905 | 136.0590443 | 109.6932177 | 1.92E-05 | 1.11088549 | 6.26E-06 | 4.517556908 | 2.159781687 |
| CNAG_00543 | - | hypothetical protein | 1069.574654 | 1133.36983 | 1215.624861 | 849.5430068 | 882.2242946 | 900.46263 | 1.93E-05 | -0.392237489 | 6.27E-06 | -4.517029647 | 0.761946977 |
| CNAG_05638 | - | hypothetical protein | 3436.439397 | 3265.625702 | 3104.420558 | 2779.897696 | 2673.795895 | 2540.210144 | 1.93E-05 | -0.31039461 | 6.27E-06 | -4.517090942 | 0.806421155 |
| CNAG_02341 | - | cyclin binding protein | 1910.6157 | 1860.822966 | 1896.226997 | 2338.975809 | 2346.303271 | 2354.537272 | 1.93E-05 | 0.297480928 | 6.28E-06 | 4.516836241 | 1.228996603 |
| CNAG_00658 | - | hypothetical protein | 205.7639856 | 189.7823668 | 150.5880142 | 289.9197633 | 295.2704865 | 321.609336 | 1.93E-05 | 0.719121768 | 6.30E-06 | 4.516037064 | 1.646179628 |
| CNAG_01930 | - | endopeptidase | 1573.208444 | 1666.864543 | 1605.099144 | 1906.874016 | 2176.485844 | 2103.604589 | 1.96E-05 | 0.337489196 | 6.40E-06 | 4.512723389 | 1.263555642 |
| CNAG_12023 | - | unspecified product | 1.584641371 | 1.274291761 | 5.48867537 | 36.54193875 | 20.91549179 | 24.14834283 | 1.97E-05 | 3.499012775 | 6.42E-06 | 4.512217629 | 11.30596926 |
| CNAG_00096 | - | hypothetical protein | 348.3595726 | 357.8642177 | 393.6092804 | 502.206533 | 517.7127627 | 604.6825064 | 1.98E-05 | 0.54869985 | 6.45E-06 | 4.511211359 | 1.462766863 |
| CNAG_01322 | - | hypothetical protein | 1272.768254 | 1361.779142 | 1411.874011 | 1017.659219 | 1076.325206 | 1080.304163 | 1.98E-05 | -0.365002859 | 6.47E-06 | -4.510479758 | 0.776467336 |
| CNAG_05263 | - | hypothetical protein | 10.02512056 | 32.47128704 | 19.42507707 | 66.92625758 | 61.59229336 | 79.32142398 | 1.99E-05 | 1.744209038 | 6.49E-06 | 4.509845264 | 3.350111326 |
| CNAG_12993 | - | unspecified product | 269818.0282 | 236555.5946 | 313610.5491 | 181750.6765 | 130649.8996 | 147604.0059 | 1.99E-05 | -0.849813211 | 6.51E-06 | -4.509160427 | 0.55485657 |
| CNAG_06997 | - | tRNA-dihydrouridine synthase 3 | 221.5315762 | 211.0614485 | 267.3786075 | 108.2116564 | 111.6891899 | 163.2936636 | 2.00E-05 | -0.886984842 | 6.52E-06 | -4.508938595 | 0.540743063 |
| CNAG_04894 | - | hypothetical protein | 43.90732908 | 88.44909653 | 61.00118281 | 150.2982003 | 144.5119351 | 132.9618474 | 2.00E-05 | 1.136898006 | 6.54E-06 | 4.50825073 | 2.199076825 |
| CNAG_05466 | - | hypothetical protein | 1.665705843 | 9.897316215 | 8.71519522 | 42.39875795 | 38.51036054 | 30.86944137 | 2.03E-05 | 2.598730592 | 6.63E-06 | 4.50511474 | 6.057533976 |
| CNAG_04337 | - | cation diffusion facilitator | 10.08310355 | 43.23455031 | 27.95185477 | 83.29053775 | 92.09761812 | 82.88315991 | 2.05E-05 | 1.672346224 | 6.69E-06 | 4.503263727 | 3.1873252 |
| CNAG_05650 | UBP5 | ubiquitin carboxyl-terminal hydrolase 7 | 11178.66762 | 10755.93893 | 10099.29938 | 8543.774187 | 9182.727099 | 9101.187432 | 2.05E-05 | -0.271257455 | 6.71E-06 | -4.502762017 | 0.828597025 |
| CNAG_02038 | - | hypothetical protein | 2648.417196 | 2998.365779 | 2974.234549 | 2010.633401 | 2243.346288 | 2475.703253 | 2.08E-05 | -0.372181922 | 6.79E-06 | -4.500021296 | 0.772613118 |
| CNAG_04370 | - | U3 small nucleolar RNA-associated protein 10 | 2141.29827 | 2027.614023 | 1761.784722 | 1518.915407 | 1609.415443 | 1483.348368 | 2.08E-05 | -0.378443159 | 6.81E-06 | -4.499675374 | 0.769267275 |
| CNAG_01338 | - | xeroderma pigmentosum group C-complementing protein | 1633.823132 | 1563.447366 | 1461.156629 | 1914.758911 | 2068.645972 | 1967.090443 | 2.08E-05 | 0.337828364 | 6.81E-06 | 4.499556586 | 1.26385273 |
| CNAG_05746 | - | Swi5-dependent recombination DNA repair protein 1, variant | 28.63763174 | 30.44114825 | 47.0535332 | 80.61576122 | 101.5810718 | 91.7644827 | 2.10E-05 | 1.357517217 | 6.89E-06 | 4.497067729 | 2.562438212 |
| CNAG_07956 | - | WD repeat and SOF domain-containing protein 1 | 796.6390095 | 790.9469567 | 778.6148648 | 1149.287971 | 1006.720679 | 1014.588185 | 2.11E-05 | 0.407612002 | 6.93E-06 | 4.495933304 | 1.326488348 |
| CNAG_02890 | - | hypothetical protein | 200.05395 | 183.3738501 | 224.0497695 | 341.5113222 | 314.2147199 | 315.0886837 | 2.13E-05 | 0.66064456 | 6.98E-06 | 4.494431358 | 1.580788723 |
| CNAG_01416 | - | hypothetical protein | 1555.046038 | 1283.208919 | 1432.156039 | 1150.901075 | 1044.334012 | 1041.515433 | 2.13E-05 | -0.416527921 | 6.99E-06 | -4.494052282 | 0.749225589 |
| CNAG_03700 | - | hypothetical protein | 1199.085582 | 1108.653418 | 1326.508423 | 881.4219261 | 929.6621451 | 947.0935863 | 2.15E-05 | -0.413795555 | 7.04E-06 | -4.492436332 | 0.750645916 |
| CNAG_04005 | - | cytoplasmic protein, variant | 1701.19184 | 1706.547111 | 1582.581978 | 1257.366445 | 1405.886448 | 1294.596259 | 2.16E-05 | -0.349990942 | 7.10E-06 | -4.490745521 | 0.784589024 |
| CNAG_04173 | - | hypothetical protein | 1341.522635 | 1312.228634 | 1292.480656 | 1069.940177 | 977.9143369 | 1053.660925 | 2.18E-05 | -0.363410437 | 7.17E-06 | -4.488663022 | 0.777324861 |
| CNAG_01115 | - | hypothetical protein | 3155.339237 | 2856.374948 | 2505.251711 | 3547.615771 | 3911.11398 | 3656.749127 | 2.20E-05 | 0.368494654 | 7.21E-06 | 4.487313427 | 1.291005059 |
| CNAG_03667 | - | hypothetical protein | 1481.969543 | 1323.20433 | 983.4974246 | 2061.635956 | 1826.271125 | 1659.627266 | 2.24E-05 | 0.534936606 | 7.35E-06 | 4.483270265 | 1.448878487 |
| CNAG_04858 | - | hypothetical protein | 58.77859001 | 46.58466104 | 50.32375276 | 129.4434546 | 121.2705841 | 95.36148373 | 2.25E-05 | 1.147528173 | 7.38E-06 | 4.482335301 | 2.21534006 |
| CNAG_05287 | - | hypothetical protein | 605.6668441 | 493.4585968 | 639.6668141 | 414.5072415 | 414.6932207 | 346.6634554 | 2.25E-05 | -0.581602641 | 7.41E-06 | -4.481512012 | 0.668221061 |
| CNAG_06058 | - | protein DGCR14 | 204.6944111 | 202.7183714 | 200.6359185 | 308.5613542 | 301.2181426 | 352.670268 | 2.26E-05 | 0.650153924 | 7.42E-06 | 4.481365634 | 1.569335623 |
| CNAG_05225 | - | bud site selection protein 31 | 1670.895702 | 1713.022848 | 1688.125544 | 1382.035845 | 1412.772942 | 1226.935908 | 2.27E-05 | -0.35037845 | 7.46E-06 | -4.480019271 | 0.784378312 |
| CNAG_08017 | - | hypothetical protein | 0.22282464 | 0.107545404 | 0.092814184 | 12.84439442 | 22.37133135 | 10.99708033 | 2.27E-05 | 6.300707362 | 7.48E-06 | 4.479587332 | 78.83188472 |

| | | | | | | | | | | | | | |
|---|---|---|---|---|---|---|---|---|---|---|---|---|---|
| CNAG_02947 | SCY1 | SCYL protein kinase | 321.5642695 | 414.8673889 | 343.541561 | 535.1085803 | 516.0142956 | 538.2108704 | 2.29E-05 | 0.545621029 | 7.55E-06 | 4.477575878 | 1.459648536 |
| CNAG_01195 | - | cation-transporting ATPase 13A3/4/5 | 3534.535516 | 3209.6396 | 3335.81132 | 2674.695728 | 2796.29644 | 2803.255549 | 2.32E-05 | -0.300409476 | 7.65E-06 | -4.474851472 | 0.81202189 |
| CNAG_07976 | - | hypothetical protein | 42.45754832 | 33.65750937 | 37.52384812 | 81.37486223 | 95.39793668 | 97.25928747 | 2.34E-05 | 1.269460004 | 7.72E-06 | 4.47266814 | 2.410713164 |
| CNAG_05270 | - | tRNA pseudouridine13 synthase | 2088.84528 | 1859.627172 | 2116.794884 | 1433.947703 | 1579.178579 | 1691.956209 | 2.35E-05 | -0.382243342 | 7.74E-06 | -4.472290946 | 0.767243625 |
| CNAG_06600 | - | vesicle-fusing ATPase | 3741.334634 | 3779.557957 | 3648.298557 | 3017.937346 | 3199.139825 | 3146.292182 | 2.36E-05 | -0.269690674 | 7.77E-06 | -4.47148627 | 0.829497378 |
| CNAG_04619 | - | prolyl oligopeptidase | 23.75555649 | 16.41726701 | 15.15875363 | 55.11564093 | 57.37906247 | 63.89935052 | 2.37E-05 | 1.680366695 | 7.81E-06 | 4.470255491 | 3.205094057 |
| CNAG_13099 | - | unspecified product | 60.09493425 | 77.73205653 | 66.30785912 | 139.9982189 | 129.6818769 | 142.8474665 | 2.38E-05 | 0.999865654 | 7.87E-06 | 4.468803611 | 1.999813766 |
| CNAG_06521 | - | hypothetical protein | 49.47985136 | 47.62089259 | 42.86797032 | 93.01560484 | 109.1823699 | 110.602467 | 2.45E-05 | 1.161598864 | 8.07E-06 | 4.463297586 | 2.237052111 |
| CNAG_12307 | - | unspecified product | 14.63387329 | 28.18773556 | 20.47433298 | 62.99394697 | 64.31170241 | 66.19380905 | 2.46E-05 | 1.6175046 | 8.12E-06 | 4.461946155 | 3.068438355 |
| CNAG_00823 | - | cadmium ion transporter | 564.3366695 | 647.5961789 | 541.901109 | 860.5127467 | 796.3254942 | 772.5644647 | 2.46E-05 | 0.45619357 | 8.12E-06 | 4.4620141 | 1.37191735 |
| CNAG_03550 | - | zinc finger HIT domain-containing protein 1 | 416.2777306 | 314.5862596 | 374.1769867 | 252.5966176 | 223.9388919 | 231.1525131 | 2.48E-05 | -0.66131398 | 8.20E-06 | -4.460021936 | 0.632302146 |
| CNAG_04371 | HSV2 | WD40 domain-containing protein | 588.9079339 | 636.8852929 | 698.58216 | 854.0450489 | 878.2931839 | 889.0581964 | 2.49E-05 | 0.432418574 | 8.21E-06 | 4.459512163 | 1.349494011 |
| CNAG_01582 | - | hypothetical protein | 46.15436492 | 58.39998359 | 68.39011145 | 107.7088374 | 143.8603488 | 125.1166844 | 2.54E-05 | 1.109409942 | 8.41E-06 | 4.454413464 | 2.157573852 |
| CNAG_07802 | - | class III aminotransferase | 29.68419399 | 27.18342859 | 22.63628057 | 98.80490239 | 88.69595326 | 48.91884971 | 2.60E-05 | 1.569161163 | 8.61E-06 | 4.449497678 | 2.967321328 |
| CNAG_06813 | CAP1alpha | O-glucosyltransferase, putative | 2342.411791 | 2397.221839 | 2487.951608 | 1971.951325 | 2045.029659 | 1920.72018 | 2.63E-05 | -0.299001885 | 8.70E-06 | -4.447099961 | 0.81281454 |
| CNAG_07909 | - | meiotic recombinase Dmc1 | 92.72725064 | 115.4087813 | 88.71811153 | 177.9667389 | 175.3956871 | 192.8043561 | 2.63E-05 | 0.868682701 | 8.70E-06 | 4.447175895 | 1.825994855 |
| CNAG_03188 | SET202 | histone-lysine N-methyltransferase, H3 lysine-36 specific | 2326.123578 | 2639.581968 | 2494.369074 | 1947.751325 | 2050.175033 | 2070.547028 | 2.65E-05 | -0.312708707 | 8.78E-06 | -4.445226363 | 0.805128684 |
| CNAG_07028 | RPN11 | 26S proteasome regulatory subunit N11 | 4307.981819 | 3763.622612 | 4350.114624 | 5087.859614 | 5146.165914 | 5419.638278 | 2.72E-05 | 0.317789394 | 9.00E-06 | 4.439826002 | 1.246419228 |
| CNAG_06011 | - | hypothetical protein | 75.03832183 | 50.91271764 | 59.91605771 | 131.4472409 | 127.1700856 | 130.6667836 | 2.75E-05 | 1.059927508 | 9.13E-06 | 4.436800619 | 2.084826762 |
| CNAG_05103 | - | transcription initiation factor TFIIF subunit beta | 900.246256 | 1006.182129 | 973.5781398 | 772.6383098 | 726.9434316 | 699.5904747 | 2.76E-05 | -0.404867741 | 9.14E-06 | -4.436482163 | 0.755305532 |
| CNAG_07540 | - | hypothetical protein | 269.0882392 | 305.0894895 | 364.7910453 | 533.8286677 | 529.1669023 | 412.9626882 | 2.79E-05 | 0.636733939 | 9.26E-06 | 4.433664534 | 1.554805309 |
| CNAG_01408 | - | PH and SEC7 domain-containing protein | 1021.979174 | 1010.729201 | 992.9613786 | 1310.927383 | 1273.272068 | 1304.303489 | 2.81E-05 | 0.347461761 | 9.33E-06 | 4.432224768 | 1.272320171 |
| CNAG_02321 | - | hypothetical protein | 20.38460289 | 18.57906753 | 25.76650023 | 51.6523734 | 71.38101871 | 70.55404579 | 2.81E-05 | 1.591848109 | 9.33E-06 | 4.432042258 | 3.014352442 |
| CNAG_03236 | - | AMME syndrome candidate protein | 44.98037333 | 74.44909305 | 42.90845934 | 110.9625332 | 143.9210005 | 115.1933981 | 2.82E-05 | 1.188215464 | 9.35E-06 | 4.431667615 | 2.27870705 |
| CNAG_04322 | - | hypothetical protein | 873.5396313 | 712.3400259 | 554.7891763 | 1206.32686 | 1085.375826 | 963.5518185 | 2.83E-05 | 0.589356903 | 9.39E-06 | 4.430816407 | 1.504575917 |
| CNAG_03555 | - | acylglycerone-phosphate reductase | 435.7535866 | 394.4833679 | 373.4065768 | 552.8563457 | 604.1059632 | 566.0186655 | 2.84E-05 | 0.503042584 | 9.42E-06 | 4.42999481 | 1.417199227 |
| CNAG_05697 | - | hypothetical protein | 807.9165034 | 797.2337606 | 893.5417596 | 658.9934368 | 613.065412 | 596.351812 | 2.85E-05 | -0.434894093 | 9.47E-06 | -4.428936421 | 0.739748057 |
| CNAG_00782 | SPS1 | STE/STE20/Fray protein kinase | 557.4307554 | 701.4646472 | 613.3308212 | 890.7549542 | 793.6195827 | 953.3398329 | 2.89E-05 | 0.480523917 | 9.62E-06 | 4.425609891 | 1.395250262 |
| CNAG_05559 | - | hypothetical protein | 2300.240261 | 2159.123158 | 2176.58137 | 1758.651355 | 1821.5933 | 1836.30731 | 2.90E-05 | -0.308627609 | 9.64E-06 | -4.425206444 | 0.807409457 |
| CNAG_03174 | - | 4'-phosphopantetheinyl transferase | 417.6633006 | 423.2367832 | 363.6109661 | 293.9277032 | 240.2991615 | 268.8632624 | 2.90E-05 | -0.601609195 | 9.66E-06 | -4.424686588 | 0.65901847 |
| CNAG_00182 | VPS29 | retrograde transporter, variant | 1177.957834 | 1117.20553 | 1063.211595 | 917.4274096 | 847.6972822 | 821.7059494 | 2.92E-05 | -0.392388509 | 9.71E-06 | -4.423538677 | 0.761867221 |
| CNAG_06652 | - | allantoate permease | 91.76626278 | 174.589698 | 119.6436333 | 234.0181145 | 246.2469308 | 242.8128987 | 2.94E-05 | 0.898143278 | 9.79E-06 | 4.421859632 | 1.863665935 |
| CNAG_00331 | - | anon-23da protein | 5.076755615 | 2.360662049 | 3.37470178 | 30.5823758 | 24.49656521 | 27.3870622 | 2.95E-05 | 3.001443033 | 9.82E-06 | 4.421041615 | 8.008005879 |
| CNAG_01039 | - | superkiller protein 3 | 5870.042223 | 5558.408261 | 5258.664896 | 4407.151987 | 4691.353211 | 4743.800352 | 2.98E-05 | -0.28510483 | 9.93E-06 | -4.418599675 | 0.820681974 |
| CNAG_12199 | - | unspecified product | 23.80742193 | 22.84570259 | 17.2924818 | 54.13159622 | 65.22880961 | 68.34569658 | 2.99E-05 | 1.577552889 | 9.98E-06 | 4.417673384 | 2.984631645 |
| CNAG_04387 | - | thioredoxin-like protein 4A | 1173.324634 | 1062.311726 | 1161.270334 | 865.9253692 | 881.3638532 | 889.3697503 | 3.00E-05 | -0.381719136 | 1.00E-05 | -4.416899784 | 0.767522455 |
| CNAG_02724 | - | hypothetical protein | 564.3342124 | 573.3282088 | 628.2081194 | 777.7506588 | 790.2786902 | 841.2817461 | 3.01E-05 | 0.432948633 | 1.00E-05 | 4.416139836 | 1.349989918 |
| CNAG_01057 | - | COP9 signalosome complex subunit 2 | 2278.14889 | 2336.866197 | 2277.890766 | 1794.415916 | 1925.981416 | 1937.319335 | 3.02E-05 | -0.30007097 | 1.01E-05 | -4.415573586 | 0.812212441 |
| CNAG_07435 | HAP2 | Transcriptional activator HAP2, putative | 85.58992424 | 71.34052522 | 74.83482239 | 154.8073963 | 140.0061912 | 152.8730483 | 3.02E-05 | 0.9397823 | 1.01E-05 | 4.415386842 | 1.918238758 |
| CNAG_06349 | - | hypothetical protein | 27.27958875 | 17.51689313 | 21.54140857 | 50.78088386 | 77.51596567 | 72.7730854 | 3.06E-05 | 1.597523441 | 1.02E-05 | 4.412522086 | 3.026233778 |
| CNAG_06589 | - | endoribonuclease L-PSP | 533.4115399 | 547.1962655 | 522.4783086 | 384.9027368 | 365.4568807 | 402.0150799 | 3.07E-05 | -0.491971951 | 1.02E-05 | -4.412011063 | 0.71105253 |
| CNAG_01791 | - | ribosome production factor 1 | 2990.530027 | 3109.398522 | 3125.709603 | 2574.373228 | 2584.969648 | 2573.458377 | 3.12E-05 | -0.269852488 | 1.04E-05 | -4.408232633 | 0.829404346 |
| CNAG_06173 | - | hypothetical protein | 509.9915137 | 537.4419154 | 454.2700333 | 349.0703576 | 374.1879959 | 323.3041045 | 3.13E-05 | -0.53511584 | 1.05E-05 | -4.407299355 | 0.690103263 |
| CNAG_03954 | - | ribosomal RNA-processing protein 17 | 641.9870262 | 582.8466612 | 682.3703425 | 478.3717102 | 469.9048146 | 420.9915767 | 3.14E-05 | -0.494631111 | 1.05E-05 | -4.406413961 | 0.709743132 |
| CNAG_02239 | - | 26S protease regulatory subunit 4 | 6794.116375 | 6497.009606 | 6721.915134 | 7848.417778 | 8207.423275 | 7915.368986 | 3.15E-05 | 0.244826734 | 1.05E-05 | 4.40578332 | 1.184950451 |
| CNAG_02099 | FAS1 | fatty acid synthase subunit beta, fungi type | 16951.73851 | 14509.69634 | 12430.61374 | 11459.27454 | 11767.78348 | 10759.87128 | 3.18E-05 | -0.384885908 | 1.06E-05 | -4.403559959 | 0.765839561 |
| CNAG_05854 | - | hypothetical protein | 592.8520732 | 537.558847 | 554.463985 | 418.427279 | 412.1064235 | 378.7905772 | 3.18E-05 | -0.494099189 | 1.07E-05 | -4.403326577 | 0.710004863 |
| CNAG_02772 | PMU101 | hypothetical protein | 1006.280601 | 898.5307513 | 972.5053481 | 775.7771233 | 657.8459928 | 721.7593325 | 3.20E-05 | -0.433294207 | 1.07E-05 | -4.40218681 | 0.74056886 |
| CNAG_06815 | FAO1 | putative fatty alcohol oxidase | 30.96146775 | 45.43001881 | 38.58839779 | 109.3874953 | 84.83519844 | 84.14143013 | 3.22E-05 | 1.279638653 | 1.08E-05 | 4.400712609 | 2.427781614 |
| CNAG_05637 | - | hypothetical protein | 57.91968925 | 101.4220408 | 88.68871845 | 222.53417 | 161.5980005 | 139.8101952 | 3.25E-05 | 1.071979348 | 1.09E-05 | 4.398620981 | 2.102315723 |
| CNAG_04441 | - | polyadenylate-binding protein, cytoplasmic and nuclear | 3507.663217 | 3299.031181 | 3114.027602 | 2692.6359 | 2758.344877 | 2716.679046 | 3.26E-05 | -0.296158229 | 1.09E-05 | -4.397701227 | 0.814418236 |
| CNAG_06726 | CSR3 | chitin synthase regulator 3 | 62.23375364 | 54.08243136 | 40.77148093 | 110.3097338 | 108.1414604 | 127.1933391 | 3.29E-05 | 1.129846879 | 1.10E-05 | 4.395948215 | 2.188355129 |
| CNAG_02524 | - | hypothetical protein | 211.7960343 | 240.4047043 | 249.6498895 | 346.6848862 | 361.6959715 | 365.0049299 | 3.29E-05 | 0.599977324 | 1.10E-05 | 4.39574224 | 1.515692743 |
| CNAG_05616 | - | ABC transporter | 1283.519708 | 1199.299542 | 1224.37813 | 1651.895734 | 1600.224461 | 1504.205001 | 3.31E-05 | 0.344023348 | 1.11E-05 | 4.394517216 | 1.269291428 |
| CNAG_00184 | - | pre-mRNA cleavage complex 2 protein Pcf11 | 2326.454919 | 2533.125427 | 2644.849253 | 3005.888619 | 3283.127645 | 3097.24764 | 3.33E-05 | 0.307786967 | 1.12E-05 | 4.393154956 | 1.237807499 |
| CNAG_04374 | - | hypothetical protein | 311.0531075 | 363.2125402 | 375.4756672 | 504.6895331 | 490.9748668 | 534.8428624 | 3.35E-05 | 0.527933659 | 1.12E-05 | 4.391880499 | 1.441862567 |
| CNAG_06707 | - | hypothetical protein | 97.53140595 | 146.6447856 | 133.4457182 | 231.8159141 | 222.8830854 | 214.0416604 | 3.38E-05 | 0.812959629 | 1.14E-05 | 4.389375107 | 1.756811774 |
| CNAG_07654 | - | hypothetical protein | 8.658019586 | 9.910940486 | 5.546928334 | 53.85334626 | 22.36279965 | 41.72113705 | 3.42E-05 | 2.384883577 | 1.15E-05 | 4.386590685 | 5.223017643 |
| CNAG_01701 | - | RTA1 domain-containing protein | 1472.67909 | 1340.452301 | 1385.414065 | 1910.341226 | 1794.282633 | 1697.379038 | 3.43E-05 | 0.348209554 | 1.15E-05 | 4.38626884 | 1.272979825 |
| CNAG_02192 | - | hypothetical protein | 18.14133364 | 33.56568922 | 14.13348066 | 44.19613563 | 72.07782965 | 109.9288419 | 3.47E-05 | 1.779389964 | 1.17E-05 | 4.38358601 | 3.432809894 |
| CNAG_12541 | - | unspecified product | 32.15723432 | 60.43230773 | 29.05921268 | 92.35273591 | 102.2632014 | 110.5249854 | 3.51E-05 | 1.316545312 | 1.18E-05 | 4.380960332 | 2.490689727 |
| CNAG_12997 | - | unspecified product | 2.80353452 | 3.453425681 | 8.679835999 | 33.67251809 | 27.9673629 | 30.71207037 | 3.56E-05 | 2.776138737 | 1.20E-05 | 4.377948963 | 6.850164993 |
| CNAG_00373 | - | glucan 1,3-beta-glucosidase | 144.018401 | 164.9563161 | 121.7872572 | 245.6947434 | 265.2722604 | 223.0022171 | 3.56E-05 | 0.758095823 | 1.20E-05 | 4.377795647 | 1.691256903 |
| CNAG_05069 | - | ubiquinol-cytochrome c reductase subunit 10 | 1195.590747 | 1150.644127 | 1291.339559 | 1021.785601 | 856.2561194 | 886.0943423 | 3.62E-05 | -0.411534862 | 1.22E-05 | -4.374022975 | 0.751823094 |
| CNAG_06495 | - | RNA exonuclease 1 | 3089.625046 | 2935.934858 | 2789.819875 | 2414.804682 | 2364.112577 | 2472.420153 | 3.63E-05 | -0.296973361 | 1.22E-05 | -4.373621871 | 0.813958215 |
| CNAG_04664 | MRPL37 | large subunit ribosomal protein L37 | 1556.71982 | 1245.696066 | 1431.270488 | 1964.03573 | 1961.715543 | 1738.462665 | 3.67E-05 | 0.40383959 | 1.24E-05 | 4.370724809 | 1.323024327 |
| CNAG_01004 | - | hypothetical protein, variant 2 | 674.0188751 | 714.4045538 | 571.7865062 | 850.8581414 | 956.0227026 | 899.0690125 | 3.70E-05 | 0.450609963 | 1.25E-05 | 4.368900591 | 1.366617933 |
| CNAG_07881 | - | hypothetical protein, variant 2 | 12.37793864 | 43.18421564 | 13.07626909 | 73.10908477 | 72.89926824 | 88.19528357 | 3.75E-05 | 1.766888455 | 1.27E-05 | 4.365787363 | 3.403191783 |
| CNAG_04131 | - | antiviral helicase SKI2 | 5926.199177 | 6038.926205 | 5706.562658 | 4681.882734 | 5204.627545 | 4975.839612 | 3.77E-05 | -0.265073515 | 1.27E-05 | -4.364831518 | 0.83215633 |
| CNAG_01756 | - | mitochondrial import inner membrane translocase subunit tim22 | 731.9451712 | 687.3473897 | 784.7600789 | 525.7456962 | 589.8850075 | 497.5831271 | 3.77E-05 | -0.466692783 | 1.27E-05 | -4.364523085 | 0.723621519 |
| CNAG_04959 | - | syntaxin 16 | 1027.444538 | 1067.619347 | 1026.924673 | 858.2891765 | 808.0469343 | 740.6893192 | 3.80E-05 | -0.390041134 | 1.29E-05 | -4.362636106 | 0.763107847 |
| CNAG_05282 | APT4 | Aminophospholipid translocase, putative | 2132.556753 | 2313.305265 | 2214.032269 | 2629.177054 | 2768.969328 | 2834.092591 | 3.83E-05 | 0.290879353 | 1.30E-05 | 4.360738756 | 1.22338573 |
| CNAG_02980 | - | membrane dipeptidase | 151.1642043 | 172.576334 | 208.0346948 | 284.4900369 | 326.5479353 | 271.8420369 | 3.83E-05 | 0.715091438 | 1.30E-05 | 4.360740991 | 1.641587258 |
| CNAG_02920 | - | hydrolase | 860.3547185 | 773.55993 | 878.6301559 | 623.0805398 | 597.5067826 | 659.5485632 | 3.91E-05 | -0.433925833 | 1.32E-05 | -4.356274997 | 0.740244703 |
| CNAG_12929 | - | unspecified product | 3.914548174 | 4.498988975 | 0.139356905 | 34.04645702 | 30.87085025 | 14.46279645 | 3.94E-05 | 3.463634618 | 1.33E-05 | 4.354731079 | 11.032093 |
| CNAG_03291 | - | hypothetical protein | 821.7222574 | 754.1093038 | 719.8099811 | 545.9650793 | 557.8546055 | 602.9269829 | 3.94E-05 | -0.444354065 | 1.33E-05 | -4.354650094 | 0.734913284 |
| CNAG_04817 | - | hypothetical protein | 642.4002497 | 560.4116537 | 552.5555174 | 850.3684987 | 785.9579797 | 781.4286862 | 3.96E-05 | 0.446311736 | 1.34E-05 | 4.353533806 | 1.362552422 |
| CNAG_07472 | - | hypothetical protein | 1121.908109 | 1062.258944 | 1061.045221 | 862.1411128 | 871.935997 | 688.5844767 | 3.98E-05 | -0.437884284 | 1.35E-05 | -4.352016075 | 0.738216411 |
| CNAG_06247 | - | hypothetical protein | 145.3065647 | 171.4728973 | 187.7930642 | 251.6418255 | 286.7381675 | 290.550179 | 3.99E-05 | 0.702438048 | 1.35E-05 | 4.351805906 | 1.627252406 |
| CNAG_07938 | - | hypothetical protein | 1306.890785 | 1365.158573 | 1070.887365 | 1699.369393 | 1659.753065 | 1611.842258 | 3.99E-05 | 0.394722593 | 1.35E-05 | 4.351528672 | 1.314689942 |
| CNAG_00704 | - | hypothetical protein, variant | 1301.015887 | 1208.991374 | 1162.55063 | 1505.343033 | 1733.193339 | 1550.794862 | 4.00E-05 | 0.367584471 | 1.36E-05 | 4.350780663 | 1.290190832 |
| CNAG_00528 | - | WD-repeat protein 23 | 683.4237849 | 728.4390515 | 723.1406381 | 894.4133381 | 980.1112107 | 955.6882823 | 4.00E-05 | 0.391646985 | 1.36E-05 | 4.350446996 | 1.311890207 |
| CNAG_12712 | - | unspecified product | 185.1168355 | 157.1971342 | 176.848953 | 101.4824473 | 90.18838229 | 99.05230736 | 4.00E-05 | -0.852666736 | 1.36E-05 | -4.350506426 | 0.553760197 |
| CNAG_12667 | - | unspecified product | 1523.82079 | 1553.597102 | 1737.11308 | 1283.942555 | 1162.477371 | 1316.744313 | 4.02E-05 | -0.370955321 | 1.36E-05 | -4.349702078 | 0.773270285 |
| CNAG_00125 | CRG1 | Putative regulator of G-protein signaling protein | 70.67392623 | 104.6470825 | 97.19356809 | 167.0852939 | 164.1866429 | 181.6903473 | 4.05E-05 | 0.90388784 | 1.37E-05 | 4.347940849 | 1.871101524 |
| CNAG_04921 | - | hypothetical protein | 30.74594875 | 26.07990376 | 16.25085851 | 73.30666843 | 69.48665739 | 64.1067638 | 4.06E-05 | 1.501232945 | 1.38E-05 | 4.347029257 | 2.830845366 |
| CNAG_06199 | - | nuclear DNA helicase II | 3453.87127 | 3098.638046 | 2968.987096 | 2445.837101 | 2510.740449 | 2696.652759 | 4.13E-05 | -0.330714401 | 1.40E-05 | -4.34356024 | 0.795142644 |

| | | | | | | | | | | | | | |
|---|---|---|---|---|---|---|---|---|---|---|---|---|---|
| CNAG_05279 | - | hypothetical protein | 9.860865112 | 11.00678831 | 11.92303761 | 58.41242018 | 29.36081788 | 44.02037777 | 4.14E-05 | 2.065079101 | 1.41E-05 | 4.342663037 | 4.184569187 |
| CNAG_04852 | NDX4 | 7,8-dihydro-8-oxoguanine triphosphatase | 672.8188962 | 664.8727607 | 605.8757157 | 889.1705223 | 886.0693857 | 830.3067738 | 4.15E-05 | 0.409235083 | 1.41E-05 | 4.342184831 | 1.327981532 |
| CNAG_03087 | - | Maltose permease | 2.852062506 | 8.830399826 | 10.82586012 | 34.94930954 | 34.93426483 | 39.49479158 | 4.22E-05 | 2.406086058 | 1.43E-05 | 4.338537334 | 5.30034423 |
| CNAG_06175 | - | 26S proteasome regulatory subunit N2 | 6294.319561 | 5920.59437 | 6064.995508 | 6901.89372 | 7852.098876 | 7745.446662 | 4.23E-05 | 0.284085256 | 1.44E-05 | 4.337649831 | 1.217637972 |
| CNAG_05194 | - | lipoyl synthase, mitochondrial | 802.2563981 | 934.0118296 | 1028.941216 | 640.0982835 | 702.8161514 | 696.2163205 | 4.25E-05 | -0.454493381 | 1.45E-05 | -4.336493295 | 0.729766393 |
| CNAG_06381 | - | hypothetical protein | 4668.335951 | 4429.21883 | 3799.791552 | 3387.609163 | 3407.806756 | 3554.833542 | 4.25E-05 | -0.332799074 | 1.45E-05 | -4.336559397 | 0.793994504 |
| CNAG_04562 | - | hypothetical protein | 26.23530525 | 33.59409542 | 26.87487414 | 59.33248159 | 80.77079718 | 89.34088743 | 4.26E-05 | 1.409146153 | 1.45E-05 | 4.33586146 | 2.65579935 |
| CNAG_12466 | - | unspecified product | 39.07427553 | 41.1788129 | 50.27200587 | 89.15659075 | 103.1411179 | 103.951627 | 4.30E-05 | 1.166876483 | 1.46E-05 | 4.33398303 | 2.245250608 |
| CNAG_13123 | - | unspecified product | 88.31217315 | 41.97549859 | 48.93569091 | 18.77468013 | 13.29764189 | 20.28921634 | 4.31E-05 | -1.812966681 | 1.47E-05 | -4.333508738 | 0.28460508 |
| CNAG_05281 | - | hypothetical protein, variant | 77.54676141 | 98.16570207 | 59.97555769 | 191.119863 | 140.8036831 | 146.3315994 | 4.31E-05 | 1.013707637 | 1.47E-05 | 4.33332021 | 2.019093383 |
| CNAG_12942 | - | unspecified product | 13.2729732 | 6.723407592 | 11.91574645 | 41.19995288 | 44.55152162 | 40.73809203 | 4.34E-05 | 2.041735636 | 1.48E-05 | 4.331624284 | 4.117405779 |
| CNAG_02292 | - | copper chaperone | 1092.94258 | 1196.915317 | 1208.188071 | 879.906774 | 967.6921154 | 882.7436892 | 4.38E-05 | -0.372938919 | 1.49E-05 | -4.329575021 | 0.772207826 |
| CNAG_12921 | - | unspecified product | 74.11707137 | 97.11940295 | 90.80329714 | 162.451601 | 174.7142084 | 152.999641 | 4.39E-05 | 0.88823979 | 1.50E-05 | 4.329048327 | 1.85091647 |
| CNAG_03377 | - | hypothetical protein | 528.8207219 | 533.2428952 | 600.2446806 | 414.5466482 | 408.66375 | 364.3757126 | 4.39E-05 | -0.501404919 | 1.50E-05 | -4.328958871 | 0.706418524 |
| CNAG_03665 | - | ATP-dependent RNA helicase DDX35 | 2759.189958 | 2623.495669 | 2626.628927 | 2144.691219 | 2195.936287 | 2281.459383 | 4.39E-05 | -0.290010952 | 1.50E-05 | -4.328624158 | 0.81789585 |
| CNAG_00412 | - | hypothetical protein | 12.24851892 | 12.11951384 | 21.48228394 | 69.250168 | 47.76780082 | 44.22617499 | 4.43E-05 | 1.814477488 | 1.51E-05 | 4.326723967 | 3.517322178 |
| CNAG_01640 | CSF1 | Cold sensitive for fermentation protein 1, putative | 5820.218467 | 5788.036187 | 5495.523014 | 6504.99943 | 7068.843268 | 7189.256568 | 4.50E-05 | 0.26440915 | 1.54E-05 | 4.323184766 | 1.201144025 |
| CNAG_02195 | - | origin recognition complex subunit 1 | 428.8146498 | 407.4140527 | 427.7480377 | 586.3850417 | 618.7514576 | 560.5228924 | 4.58E-05 | 0.467576759 | 1.57E-05 | 4.319181455 | 1.382784904 |
| CNAG_04977 | - | pre-mRNA branch site protein p14 | 1111.92453 | 1162.624403 | 1094.278629 | 1355.114116 | 1485.540854 | 1468.589284 | 4.59E-05 | 0.340491691 | 1.57E-05 | 4.318619603 | 1.266188055 |
| CNAG_07873 | - | MIPC synthase | 610.3035679 | 584.8884977 | 496.9525258 | 422.3373944 | 389.6237608 | 386.5451978 | 4.59E-05 | -0.512800452 | 1.57E-05 | -4.318349814 | 0.700860658 |
| CNAG_02308 | - | hypothetical protein | 418.9553727 | 412.5466606 | 434.9770697 | 318.0784999 | 284.3799295 | 273.3407064 | 4.62E-05 | -0.545983957 | 1.58E-05 | -4.317080804 | 0.684924107 |
| CNAG_00342 | - | hypothetical protein | 239.5773619 | 209.1839826 | 167.6539619 | 300.6345691 | 337.5777286 | 368.1959618 | 4.64E-05 | 0.693122528 | 1.59E-05 | 4.315965181 | 1.616779045 |
| CNAG_01403 | - | eclair-PA | 2239.600395 | 2234.528349 | 2232.030667 | 1888.72964 | 1797.406691 | 1861.843711 | 4.64E-05 | -0.288982049 | 1.59E-05 | -4.315821107 | 0.818479366 |
| CNAG_01797 | - | hypothetical protein | 824.2773306 | 827.392953 | 897.825715 | 658.9437301 | 608.7175482 | 668.4369036 | 4.66E-05 | -0.412972887 | 1.60E-05 | -4.314668526 | 0.751074078 |
| CNAG_03473 | - | lipoyl(octanoyl) transferase | 532.2757898 | 527.8479244 | 558.6968323 | 400.5111277 | 393.1171617 | 383.1949327 | 4.70E-05 | -0.474781846 | 1.61E-05 | -4.312911218 | 0.719575591 |
| CNAG_02695 | - | hypothetical protein, variant | 2364.457435 | 2258.228213 | 2152.07734 | 1847.425336 | 1837.965482 | 1871.836098 | 4.73E-05 | -0.301558535 | 1.62E-05 | -4.311611449 | 0.811375398 |
| CNAG_01732 | - | nuclear protein localization protein 4 | 292.0717462 | 245.8160273 | 170.8946346 | 409.219691 | 424.8313307 | 355.1329077 | 4.74E-05 | 0.732968382 | 1.63E-05 | 4.310982428 | 1.662055297 |
| CNAG_01672 | - | Ras family protein | 12.05638118 | 3.48069886 | 7.660035671 | 33.25275378 | 40.28349845 | 38.4369108 | 4.74E-05 | 2.3167745 | 1.63E-05 | 4.31068657 | 4.982170865 |
| CNAG_01276 | - | hypothetical protein | 7476.935198 | 7430.972907 | 6846.621993 | 6054.870484 | 6155.976963 | 6253.618759 | 4.78E-05 | -0.251746138 | 1.64E-05 | -4.308762814 | 0.839879268 |
| CNAG_06202 | - | hypothetical protein | 73.87742876 | 60.55646656 | 54.60562738 | 121.1343892 | 134.1750084 | 126.2613204 | 4.81E-05 | 1.010954585 | 1.65E-05 | 4.307246471 | 2.015244082 |
| CNAG_06401 | RIC8 | RIC8 guanine nucleotide exchange factor | 770.9123317 | 695.0960501 | 770.0575899 | 953.5980798 | 982.6225445 | 1046.666563 | 4.87E-05 | 0.399900445 | 1.67E-05 | 4.304532238 | 1.31941686 |
| CNAG_07789 | - | hypothetical protein | 2.712713741 | 0.145473387 | 0.125452167 | 31.00574708 | 21.98057959 | 17.60351774 | 4.90E-05 | 5.091570291 | 1.68E-05 | 4.303271353 | 34.09693823 |
| CNAG_04291 | - | glycosyl-hydrolase | 764.1794407 | 1017.149187 | 968.4134266 | 1189.422128 | 1267.323279 | 1299.787581 | 4.93E-05 | 0.435734111 | 1.69E-05 | 4.301726118 | 1.352598924 |
| CNAG_06334 | - | hypothetical protein | 410.5980492 | 405.9949534 | 341.2282665 | 268.9487595 | 277.5672218 | 214.5862645 | 4.96E-05 | -0.622336652 | 1.71E-05 | -4.300380007 | 0.649617925 |
| CNAG_02577 | - | oxidoreductase | 1106.372253 | 796.2285473 | 892.5469878 | 656.4933951 | 619.8871032 | 703.9720348 | 4.97E-05 | -0.514751473 | 1.71E-05 | -4.299979829 | 0.699913494 |
| CNAG_06214 | - | hypothetical protein | 1103.32737 | 1069.825504 | 1109.028933 | 875.3335426 | 861.5297683 | 853.8569474 | 5.01E-05 | -0.357133265 | 1.72E-05 | -4.297888695 | 0.780714373 |
| CNAG_06195 | - | hypothetical protein | 145.2287335 | 183.2413419 | 127.1228331 | 242.4157386 | 254.7630664 | 272.7596606 | 5.02E-05 | 0.74119725 | 1.73E-05 | 4.297197537 | 1.671562444 |
| CNAG_02122 | - | cytoplasmic tRNA 2-thiolation protein 1 | 1071.828111 | 1118.259307 | 1067.45612 | 808.282268 | 823.5418524 | 907.075532 | 5.03E-05 | -0.374340349 | 1.73E-05 | -4.296832488 | 0.77145807 |
| CNAG_06593 | - | rhamnogalacturonan lyase | 451.0256762 | 419.2876672 | 483.1758214 | 658.8951504 | 613.4500297 | 619.3226699 | 5.05E-05 | 0.466654058 | 1.74E-05 | 4.295908584 | 1.381900802 |
| CNAG_00921 | - | glutathione transferase | 112.4947919 | 127.2586131 | 90.87272161 | 209.3013927 | 208.3137141 | 175.2796729 | 5.05E-05 | 0.832823655 | 1.74E-05 | 4.295710046 | 1.78116807 |
| CNAG_05870 | - | hypothetical protein, variant | 16.99023232 | 34.63444327 | 25.79760491 | 64.32020834 | 81.77975386 | 68.51495751 | 5.08E-05 | 1.492514286 | 1.75E-05 | 4.294313735 | 2.813789269 |
| CNAG_02360 | - | hypothetical protein | 1243.319703 | 1090.323183 | 1180.472272 | 986.7636778 | 859.7468551 | 818.4100473 | 5.09E-05 | -0.415472218 | 1.75E-05 | -4.294137795 | 0.749774041 |
| CNAG_02413 | - | hypothetical protein, variant | 687.9900477 | 618.5949435 | 649.5736144 | 904.7419082 | 953.4478847 | 804.8273526 | 5.19E-05 | 0.4296742 | 1.79E-05 | 4.289620201 | 1.346929369 |
| CNAG_06898 | - | chitin synthase export chaperone | 1652.710004 | 1927.588375 | 1934.597282 | 2329.671583 | 2340.289531 | 2295.707839 | 5.32E-05 | 0.322353751 | 1.83E-05 | 4.28410425 | 1.250368857 |
| CNAG_02435 | BWC2 | white collar 2 protein | 3259.104625 | 3315.202907 | 3420.044168 | 2719.863299 | 2878.272402 | 2803.257743 | 5.37E-05 | -0.265950937 | 1.85E-05 | -4.281936356 | 0.83165038 |
| CNAG_06192 | - | hypothetical protein | 1576.057831 | 1306.905813 | 1387.396551 | 1098.650154 | 1131.5304 | 1081.457514 | 5.42E-05 | -0.383151383 | 1.87E-05 | -4.279709223 | 0.766760869 |
| CNAG_02033 | - | hypothetical protein | 78.80092717 | 63.46270439 | 87.12236968 | 27.29505793 | 28.02980783 | 35.77422352 | 5.47E-05 | -1.357100478 | 1.89E-05 | -4.277717713 | 0.390366058 |
| CNAG_07620 | - | hypothetical protein | 2.706353642 | 0.143212628 | 0.123508113 | 30.20237778 | 22.00216904 | 16.51959125 | 5.48E-05 | 5.070347885 | 1.89E-05 | 4.277205708 | 33.59903482 |
| CNAG_07565 | TAF9 | transcription initiation factor TFIID subunit 9B | 741.4124833 | 788.5766559 | 827.4405511 | 573.2309189 | 634.7295927 | 570.8091931 | 5.51E-05 | -0.42213776 | 1.90E-05 | -4.275820563 | 0.746317924 |
| CNAG_03401 | GAT203 | hypothetical protein | 1265.641462 | 1198.058699 | 1287.11623 | 982.7001471 | 976.2379247 | 1017.015009 | 5.52E-05 | -0.34946576 | 1.91E-05 | -4.275352151 | 0.784874689 |
| CNAG_01038 | - | hypothetical protein | 863.9739922 | 947.9806573 | 853.1074954 | 623.7534407 | 733.9489186 | 661.8233194 | 5.52E-05 | -0.41580238 | 1.91E-05 | -4.274926997 | 0.749602474 |
| CNAG_05518 | - | exocyst complex component 4 | 476.7222628 | 506.4858275 | 470.4232315 | 639.9847284 | 654.8620201 | 696.956924 | 5.52E-05 | 0.439103438 | 1.91E-05 | 4.275002478 | 1.355761528 |
| CNAG_02775 | BIM1 | BCS-inducible membrane protein | 24382.32506 | 20174.93039 | 23873.50808 | 14916.92169 | 19027.14256 | 18986.0496 | 5.53E-05 | -0.386913369 | 1.91E-05 | -4.27471023 | 0.76476406 |
| CNAG_04711 | - | hypothetical protein | 4.271992748 | 24.06445149 | 31.214927 | 150.1352471 | 151.0142517 | 146.9636589 | 5.55E-05 | 2.9094613 | 1.92E-05 | 4.273590608 | 7.513375985 |
| CNAG_04355 | - | hypothetical protein | 318.4162761 | 265.0281787 | 352.755796 | 191.7307019 | 199.8368734 | 206.6862263 | 5.56E-05 | -0.665768189 | 1.93E-05 | -4.273212029 | 0.630352972 |
| CNAG_01563 | HDA1 | histone deacetylase 6/10 | 1829.739256 | 1852.036689 | 1913.133031 | 1399.766425 | 1561.124952 | 1562.084527 | 5.58E-05 | -0.322198576 | 1.93E-05 | -4.272317099 | 0.799850028 |
| CNAG_02964 | - | hypothetical protein | 524.6158142 | 561.4380698 | 543.9900205 | 702.9934652 | 780.0451244 | 723.6915936 | 5.63E-05 | 0.422571242 | 1.95E-05 | 4.270404311 | 1.340314201 |
| CNAG_03705 | - | hypothetical protein | 1047.775323 | 1182.006892 | 1058.03642 | 1571.863864 | 1545.98767 | 1270.041409 | 5.67E-05 | 0.401248888 | 1.97E-05 | 4.268802877 | 1.320650655 |
| CNAG_03680 | ROM21 | Predicted Rho guanine nucleotide exchange factor (Rho GEF) | 929.8969041 | 1101.203059 | 1082.5176 | 1283.538155 | 1385.476849 | 1407.50021 | 5.73E-05 | 0.374311675 | 1.99E-05 | 4.266398967 | 1.296220966 |
| CNAG_06472 | - | nucleolar protein 4 | 2684.376233 | 2347.709561 | 2600.991574 | 1837.892575 | 2082.956962 | 2138.247042 | 5.76E-05 | -0.349175319 | 2.00E-05 | -4.264912659 | 0.785032714 |
| CNAG_06402 | - | 26S proteasome complex subunit DSS1 | 2631.996161 | 2609.494427 | 2768.428616 | 2235.048973 | 2282.237716 | 2119.446514 | 5.77E-05 | -0.286576251 | 2.00E-05 | -4.264675377 | 0.819845378 |
| CNAG_05283 | - | solute carrier family 25 (mitochondrial carrier protein), member 16 | 1120.993556 | 1205.56376 | 1316.928239 | 918.0474887 | 983.1894698 | 936.0150252 | 5.83E-05 | -0.375150901 | 2.02E-05 | -4.262191976 | 0.771024762 |
| CNAG_05673 | - | GPI inositol-deacylase, variant | 1028.016356 | 1305.883185 | 1182.784241 | 1521.028233 | 1620.147203 | 1485.301274 | 5.84E-05 | 0.381845796 | 2.03E-05 | 4.26143775 | 1.303007868 |
| CNAG_05531 | - | DNA excision repair protein ERCC-4 | 1963.141764 | 1959.920116 | 1806.677324 | 2292.980163 | 2375.631883 | 2455.522951 | 5.84E-05 | 0.299124049 | 2.03E-05 | 4.261481237 | 1.230397135 |
| CNAG_12069 | - | unspecified product | 8.812714581 | 24.9447556 | 18.32892266 | 60.88117633 | 48.5620121 | 62.77733292 | 5.88E-05 | 1.753341346 | 2.04E-05 | 4.259952912 | 3.371384904 |
| CNAG_03168 | - | sulfite reductase (NADPH) flavoprotein alpha- component | 1934.312948 | 1400.667039 | 1511.116392 | 1150.700377 | 1282.507102 | 1108.165116 | 5.90E-05 | -0.469618463 | 2.05E-05 | -4.258998269 | 0.722155555 |
| CNAG_01286 | - | hypothetical protein, variant | 181.5157437 | 255.4153533 | 205.9825839 | 347.6366202 | 367.0060453 | 308.5196955 | 5.91E-05 | 0.658879416 | 2.05E-05 | 4.258847472 | 1.578855803 |
| CNAG_05184 | - | glycosyl transferase family 8 protein | 0.367896414 | 9.843985308 | 2.319838834 | 29.56229835 | 26.20522075 | 34.9771185 | 5.94E-05 | 3.007097934 | 2.07E-05 | 4.257334297 | 8.039456284 |
| CNAG_12428 | - | unspecified product | 68.27497753 | 79.92830386 | 89.7105001 | 155.5337539 | 172.2401409 | 129.7876249 | 5.98E-05 | 0.934944378 | 2.08E-05 | 4.256006374 | 1.911816925 |
| CNAG_04822 | - | deoxyhypusine synthase | 553.71827 | 501.1586585 | 538.6543069 | 756.9283173 | 682.3881245 | 741.4020845 | 6.01E-05 | 0.43733151 | 2.09E-05 | 4.254792415 | 1.354097395 |
| CNAG_12179 | - | unspecified product | 19.18354885 | 21.7568735 | 18.33629686 | 55.10559058 | 55.60154024 | 62.81737615 | 6.01E-05 | 1.553563107 | 2.09E-05 | 4.254682689 | 2.935412204 |
| CNAG_00659 | - | TFIIH basal transcription factor complex TTD-A subunit | 49.63657666 | 46.61850387 | 79.00878245 | 101.2968071 | 151.6815484 | 136.1352876 | 6.09E-05 | 1.140751824 | 2.12E-05 | 4.251546403 | 2.204958991 |
| CNAG_02046 | PBP1 | poly(A)-binding protein binding protein | 1189.76611 | 1222.781473 | 1183.69835 | 995.2657694 | 867.4798694 | 959.2973 | 6.09E-05 | -0.364330818 | 2.12E-05 | -4.251416478 | 0.776829117 |
| CNAG_05743 | - | anaphase-promoting complex subunit 10 | 399.3117929 | 519.0519772 | 444.6104739 | 328.9145665 | 322.3896522 | 276.7100569 | 6.14E-05 | -0.570922107 | 2.14E-05 | -4.249695956 | 0.67318638 |
| CNAG_03485 | - | polyamine transporter 1 | 1148.122864 | 1186.334778 | 1116.679546 | 1464.968079 | 1626.240805 | 1373.234063 | 6.22E-05 | 0.356116368 | 2.17E-05 | 4.246624843 | 1.279975653 |
| CNAG_02738 | - | hypothetical protein | 2099.39887 | 2117.04574 | 1894.009001 | 1638.822196 | 1601.590423 | 1715.28294 | 6.26E-05 | -0.31756597 | 2.19E-05 | -4.245072189 | 0.802422536 |
| CNAG_03247 | - | sterol 3beta-glucosyltransferase, variant | 1641.971662 | 1541.899814 | 1452.621798 | 1985.704958 | 1908.100327 | 1950.433472 | 6.27E-05 | 0.319088089 | 2.19E-05 | 4.244779505 | 1.247541743 |
| CNAG_12060 | - | unspecified product | 91.69863403 | 150.9106739 | 107.9098149 | 225.6452094 | 179.4971844 | 242.6606933 | 6.35E-05 | 0.879616389 | 2.22E-05 | 4.241929561 | 1.839886013 |
| CNAG_05146 | - | DNA-directed RNA polymerase II subunit RPB9 | 1365.027065 | 1411.373956 | 1579.27541 | 1214.758686 | 1103.872726 | 1098.118283 | 6.48E-05 | -0.366025299 | 2.26E-05 | -4.237281387 | 0.775917248 |
| CNAG_05339 | - | hypothetical protein | 1404.561935 | 1357.482744 | 1309.560653 | 1132.242952 | 1091.854045 | 1011.542883 | 6.51E-05 | -0.347104081 | 2.28E-05 | -4.23601466 | 0.786160574 |
| CNAG_01680 | - | 3-methylcrotonyl-CoA carboxylase alpha subunit | 87.09303607 | 112.2907905 | 166.4047305 | 205.7945863 | 241.9911646 | 241.6395899 | 6.52E-05 | 0.897306242 | 2.28E-05 | 4.235716645 | 1.86258497 |
| CNAG_04044 | - | U3 small nucleolar RNA-associated protein 6 | 1989.66736 | 1827.31225 | 2122.119059 | 1531.357614 | 1578.323862 | 1642.021215 | 6.54E-05 | -0.337448989 | 2.29E-05 | -4.234843101 | 0.791439519 |
| CNAG_02548 | - | cobalamin synthesis protein, variant | 225.0054202 | 245.3981671 | 201.4941581 | 146.7189492 | 129.9087445 | 134.5760231 | 6.54E-05 | -0.728867571 | 2.29E-05 | -4.234740138 | 0.603377343 |
| CNAG_03722 | - | RAN protein binding protein | 2963.602405 | 2870.236713 | 2970.010579 | 2403.107651 | 2533.248755 | 2441.354723 | 6.54E-05 | -0.270433672 | 2.29E-05 | -4.234520413 | 0.829070291 |
| CNAG_02563 | - | DNA polymerase delta subunit 1 | 1653.430529 | 1773.343347 | 1663.620623 | 1399.148677 | 1378.22026 | 1384.488559 | 6.56E-05 | -0.30535566 | 2.30E-05 | -4.233913813 | 0.809242694 |
| CNAG_03072 | - | enolase | 36812.13192 | 37271.56911 | 37387.67661 | 30992.82309 | 33577.71977 | 32394.58111 | 6.61E-05 | -0.216443638 | 2.32E-05 | -4.232032788 | 0.860684482 |
| CNAG_05231 | - | nuclear protein | 2809.397186 | 2703.212881 | 2604.250145 | 2206.979889 | 2207.138059 | 2320.313067 | 6.65E-05 | -0.284806713 | 2.33E-05 | -4.23073477 | 0.820851576 |

| | | | | | | | | | | | | | |
|---|---|---|---|---|---|---|---|---|---|---|---|---|---|
| CNAG_00439 | - | hypothetical protein | 2016.402492 | 1805.733931 | 1831.053722 | 1571.189615 | 1538.673189 | 1420.076764 | 6.74E-05 | -0.335335068 | 2.36E-05 | -4.227669881 | 0.792600033 |
| CNAG_00624 | - | hypothetical protein | 1920.808541 | 1936.07285 | 1952.606924 | 1688.788834 | 1538.633898 | 1496.661858 | 6.83E-05 | -0.313863375 | 2.40E-05 | -4.224457669 | 0.804484553 |
| CNAG_07641 | - | Monosaccharide transporter | 28.53014952 | 37.86392755 | 18.39138737 | 91.56997092 | 61.40016494 | 76.23584951 | 6.84E-05 | 1.444464474 | 2.40E-05 | 4.224039152 | 2.721617786 |
| CNAG_01811 | - | hypothetical protein | 2390.18017 | 2236.710606 | 2334.401764 | 1875.420886 | 2008.827563 | 1826.367388 | 6.87E-05 | -0.301537054 | 2.41E-05 | -4.222919724 | 0.811387479 |
| CNAG_07756 | - | cell division control protein | 1650.898893 | 1355.4357 | 1785.055841 | 1071.988053 | 1178.046202 | 1304.51477 | 6.94E-05 | -0.447209843 | 2.44E-05 | -4.220661767 | 0.733459981 |
| CNAG_00925 | ISP7 | Identified spore protein 7 | 96.2572084 | 122.9531715 | 109.9983043 | 186.4812423 | 190.955196 | 193.9951981 | 6.95E-05 | 0.784700411 | 2.44E-05 | 4.220209694 | 1.72273454 |
| CNAG_02401 | - | G patch domain-containing protein 1 | 1454.700178 | 1367.171691 | 1255.199786 | 1046.487274 | 1064.224978 | 1105.825354 | 6.95E-05 | -0.357674118 | 2.44E-05 | -4.220249227 | 0.780421745 |
| CNAG_00563 | - | hypothetical protein | 997.0678216 | 975.0068252 | 1037.551972 | 790.4938034 | 700.9809224 | 821.6063762 | 6.96E-05 | -0.395942355 | 2.44E-05 | -4.219866889 | 0.759992794 |
| CNAG_01649 | - | hypothetical protein | 1675.875151 | 1625.935217 | 1599.765985 | 2051.045755 | 1970.183232 | 2054.781223 | 7.01E-05 | 0.295132667 | 2.46E-05 | 4.218142098 | 1.226997804 |
| CNAG_06357 | - | hypothetical protein, variant | 362.8795769 | 361.8851044 | 361.4152353 | 265.896447 | 246.442797 | 232.2665863 | 7.10E-05 | -0.560206848 | 2.50E-05 | -4.215020068 | 0.678204919 |
| CNAG_05775 | - | phosphatidylinositol glycan, class S | 317.9466712 | 349.1809621 | 310.4749412 | 439.1649024 | 481.5847162 | 482.6966814 | 7.11E-05 | 0.507694999 | 2.50E-05 | 4.214820927 | 1.4217768 |
| CNAG_06825 | - | hypothetical protein | 40.18220526 | 43.29710208 | 31.16897813 | 129.3965824 | 83.05805222 | 72.10497428 | 7.14E-05 | 1.302193689 | 2.51E-05 | 4.213614693 | 2.466035707 |
| CNAG_13060 | - | unspecified product | 5.241847545 | 16.33110788 | 10.86350129 | 51.02168251 | 32.039367 | 47.25445385 | 7.16E-05 | 2.056381197 | 2.52E-05 | 4.212903878 | 4.159416621 |
| CNAG_02208 | - | ribonucleoside-diphosphate reductase large subunit | 2373.844129 | 2360.568921 | 2193.674309 | 1913.595976 | 1916.476225 | 1908.478482 | 7.17E-05 | -0.28690887 | 2.52E-05 | -4.212643339 | 0.819656381 |
| CNAG_02423 | - | hypothetical protein | 230.3375867 | 211.3550565 | 232.5974778 | 350.7319264 | 331.4409355 | 336.187103 | 7.24E-05 | 0.580110843 | 2.55E-05 | 4.210385053 | 1.494964103 |
| CNAG_01210 | - | hypothetical protein | 6.296455153 | 8.807215285 | 4.462060494 | 26.86211282 | 29.68101765 | 41.5345046 | 7.43E-05 | 2.401287646 | 2.62E-05 | 4.204454513 | 5.282744539 |
| CNAG_05060 | - | hypothetical protein | 368.0428573 | 277.1374328 | 355.2269958 | 513.3715644 | 543.8119282 | 450.6526358 | 7.46E-05 | 0.574078705 | 2.63E-05 | 4.203309445 | 1.48872647 |
| CNAG_03784 | - | hypothetical protein | 2.803033815 | 8.789909101 | 4.451917976 | 25.33082764 | 42.38935163 | 23.17753333 | 7.82E-05 | 2.67127765 | 2.76E-05 | 4.19277238 | 6.369930578 |
| CNAG_01081 | - | hypothetical protein | 160.4354516 | 185.4595227 | 189.9389512 | 306.5472596 | 284.0705937 | 258.5548298 | 7.82E-05 | 0.652315985 | 2.76E-05 | 4.192658017 | 1.571689234 |
| CNAG_04185 | - | hypothetical protein | 68.12980856 | 73.42029361 | 47.17680581 | 133.0066841 | 150.7485275 | 108.6960877 | 7.82E-05 | 1.041419861 | 2.76E-05 | 4.192526109 | 2.058252332 |
| CNAG_01983 | RCV1 | Protein required for cytokinesis and virulence 1 | 363.4834185 | 339.5711866 | 410.6458546 | 557.7567704 | 515.1282366 | 518.2942186 | 7.87E-05 | 0.500266472 | 2.78E-05 | 4.191077494 | 1.414474797 |
| CNAG_13089 | - | unspecified product | 83.42565902 | 111.0926435 | 90.82686711 | 154.3845639 | 183.3148253 | 179.512292 | 7.96E-05 | 0.847217266 | 2.81E-05 | 4.188343566 | 1.799027532 |
| CNAG_05583 | GCS1 | ceramide glucosyltransferase | 1113.950009 | 1119.408765 | 1372.323382 | 808.1465513 | 945.2229362 | 945.971742 | 8.05E-05 | -0.433258969 | 2.84E-05 | -4.185824449 | 0.740586949 |
| CNAG_04818 | - | hypothetical protein | 13.38489267 | 16.38590302 | 18.30561123 | 48.89499221 | 59.35317449 | 45.28753868 | 8.10E-05 | 1.678085264 | 2.86E-05 | 4.184295783 | 3.200029631 |
| CNAG_02792 | - | mitochondrial chaperone BCS1 | 1850.630538 | 1679.682943 | 1760.651171 | 1369.471534 | 1471.414166 | 1439.99162 | 8.12E-05 | -0.321429921 | 2.87E-05 | -4.183711039 | 0.800276294 |
| CNAG_03239 | - | hypothetical protein, variant | 712.6678329 | 774.7818928 | 840.41246 | 999.5525112 | 1043.044982 | 1022.317144 | 8.23E-05 | 0.382461457 | 2.91E-05 | 4.180316345 | 1.303564038 |
| CNAG_01824 | - | transmembrane protein, variant | 5504.852164 | 5032.780463 | 4802.345208 | 6083.250948 | 6232.123415 | 6498.733588 | 8.28E-05 | 0.27910636 | 2.93E-05 | 4.178839123 | 1.213443015 |
| CNAG_01760 | - | hypothetical protein | 202.3271289 | 206.9962665 | 178.2716465 | 346.3540374 | 288.271927 | 287.3946028 | 8.28E-05 | 0.63734106 | 2.93E-05 | 4.178916829 | 1.555459747 |
| CNAG_05610 | - | hypothetical protein | 499.5742147 | 538.543298 | 490.4935377 | 380.3013427 | 320.5321415 | 387.5694573 | 8.32E-05 | -0.504970297 | 2.94E-05 | -4.177795755 | 0.704674886 |
| CNAG_02189 | - | Alpha-amylase | 2044.352375 | 1962.945281 | 1614.633662 | 1554.089648 | 1383.31802 | 1400.086938 | 8.33E-05 | -0.388991158 | 2.95E-05 | -4.177297664 | 0.763663429 |
| CNAG_06470 | - | hypothetical protein | 129.2207459 | 131.2475512 | 132.0643235 | 68.6998592 | 75.62427199 | 66.90149197 | 8.38E-05 | -0.922686951 | 2.97E-05 | -4.175810478 | 0.527525614 |
| CNAG_01630 | - | polyadenylation factor subunit 2 | 442.856872 | 435.4211237 | 466.1244505 | 637.0485092 | 646.3347207 | 572.7738847 | 8.41E-05 | 0.450467221 | 2.98E-05 | 4.175030089 | 1.366482724 |
| CNAG_02468 | - | AP-3 complex subunit delta-1 | 1536.598304 | 1609.566985 | 1462.07863 | 1264.510722 | 1273.880755 | 1189.143266 | 8.58E-05 | -0.321238663 | 3.04E-05 | -4.17047066 | 0.800382394 |
| CNAG_06454 | - | atypical/RIO/RIO1 protein kinase | 1542.363655 | 1372.638125 | 1588.897893 | 1140.636951 | 1254.928054 | 1154.723797 | 8.58E-05 | -0.359537395 | 3.04E-05 | -4.170159774 | 0.779414461 |
| CNAG_00738 | - | hypothetical protein | 64.69774578 | 47.70851693 | 84.34220427 | 128.1470644 | 152.4289257 | 125.227892 | 8.58E-05 | 1.037623247 | 3.04E-05 | 4.170229992 | 2.052842931 |
| CNAG_12272 | - | unspecified product | 3.933076785 | 5.575633353 | 4.437248852 | 28.87127951 | 29.89577411 | 22.02917647 | 8.67E-05 | 2.704272126 | 3.07E-05 | 4.167863827 | 6.5172897 |
| CNAG_03354 | - | hypothetical protein | 617.4714221 | 575.2862524 | 619.4999762 | 451.0911711 | 449.1933382 | 449.7748532 | 8.70E-05 | -0.44044462 | 3.09E-05 | -4.166878273 | 0.736907468 |
| CNAG_04502 | - | hypothetical protein | 1670.798939 | 1614.984429 | 1519.665924 | 1247.294311 | 1227.217569 | 1370.00113 | 8.78E-05 | -0.336491853 | 3.12E-05 | -4.164754603 | 0.791964763 |
| CNAG_00844 | - | hypothetical protein | 2.795966629 | 6.6565523 | 6.564698382 | 27.8778778 | 22.60249183 | 38.20302923 | 8.81E-05 | 2.622647879 | 3.13E-05 | 4.163778225 | 6.158794022 |
| CNAG_00597 | DIP5 | amino acid transporter, variant | 1480.527637 | 1547.069089 | 1375.705498 | 1201.461512 | 1183.28043 | 1164.696225 | 8.93E-05 | -0.326106534 | 3.17E-05 | -4.160753926 | 0.797686333 |
| CNAG_01320 | - | large subunit ribosomal protein L17 | 1488.77623 | 1513.72876 | 1636.890878 | 1295.692577 | 1131.428214 | 1257.920809 | 8.96E-05 | -0.347422015 | 3.19E-05 | -4.15976664 | 0.785987343 |
| CNAG_07589 | - | xylitol dehydrogenase | 120.7629424 | 164.9186091 | 118.5747869 | 215.9475507 | 221.9446551 | 250.5170133 | 9.09E-05 | 0.757243132 | 3.23E-05 | 4.15647888 | 1.690257598 |
| CNAG_01543 | - | AP-3 complex subunit beta | 1467.637257 | 1413.519637 | 1428.977874 | 1006.639032 | 1217.856418 | 1166.884937 | 9.25E-05 | -0.361678867 | 3.29E-05 | -4.152514879 | 0.778258392 |
| CNAG_00188 | - | endoplasmic reticulum protein | 396.5584863 | 327.4915898 | 375.2488571 | 277.6486715 | 231.7230452 | 221.1978635 | 9.32E-05 | -0.606906987 | 3.32E-05 | -4.150602931 | 0.656602894 |
| CNAG_06003 | - | hypothetical protein | 72.91070056 | 92.79923545 | 78.03362038 | 171.3465095 | 126.0057407 | 163.9265582 | 9.35E-05 | 0.91264569 | 3.33E-05 | 4.149853381 | 1.882494552 |
| CNAG_06523 | - | hypothetical protein | 166.4029436 | 225.3200558 | 249.616414 | 345.287751 | 359.2071252 | 317.3651302 | 9.37E-05 | 0.659211378 | 3.34E-05 | 4.149173185 | 1.579219138 |
| CNAG_09004 | - | cytochrome c oxidase subunit III | 194575.2162 | 173059.3086 | 165576.4791 | 154855.5056 | 146314.356 | 144914.9994 | 9.43E-05 | -0.273088882 | 3.36E-05 | -4.147812148 | 0.827545831 |
| CNAG_07559 | - | glucose-6-phosphate 1-epimerase | 1814.353376 | 1635.482756 | 1622.030359 | 1438.163971 | 1187.460508 | 1341.200047 | 9.52E-05 | -0.370009067 | 3.39E-05 | -4.145441817 | 0.773777634 |
| CNAG_05233 | - | hypothetical protein | 1234.384242 | 1020.482236 | 1066.56353 | 1496.236017 | 1413.007935 | 1438.639839 | 9.52E-05 | 0.372894825 | 3.39E-05 | 4.145363234 | 1.294948592 |
| CNAG_06279 | - | transportin-3 | 860.1016994 | 674.4516221 | 728.3151283 | 521.0246247 | 597.6366502 | 523.0957761 | 9.70E-05 | -0.479456507 | 3.46E-05 | -4.140945111 | 0.717247775 |
| CNAG_00456 | ISP6 | Identified spore protein 6 | 810.0683079 | 907.9939639 | 622.8543868 | 510.0625313 | 572.5491733 | 566.344637 | 9.72E-05 | -0.519558184 | 3.46E-05 | -4.140575398 | 0.697585431 |
| CNAG_05379 | - | regucalcin | 1812.084228 | 1613.970566 | 1738.245534 | 1376.533213 | 1343.680328 | 1438.856945 | 9.80E-05 | -0.328336734 | 3.49E-05 | -4.138601971 | 0.796454177 |
| CNAG_05453 | - | hypothetical protein, variant | 104.4660782 | 156.2858822 | 108.9782955 | 220.083094 | 202.073744 | 221.7238131 | 9.84E-05 | 0.791682308 | 3.51E-05 | 4.137383839 | 1.73109189 |
| CNAG_07330 | - | hypothetical protein | 58.97097563 | 76.68084253 | 84.37166977 | 158.9191943 | 135.7211184 | 130.8064739 | 9.97E-05 | 0.941716621 | 3.56E-05 | 4.134473512 | 1.920812398 |
| CNAG_06371 | - | Guanine deaminase | 55.24446923 | 29.41242748 | 53.47661283 | 86.60580116 | 113.5436536 | 120.4850668 | 9.98E-05 | 1.202013944 | 3.56E-05 | 4.134125927 | 2.300606023 |
| CNAG_01065 | - | hypothetical protein | 685.2103384 | 650.6945761 | 682.4301733 | 519.6364771 | 532.9295911 | 456.512006 | 9.98E-05 | -0.436172855 | 3.57E-05 | -4.134000679 | 0.739092657 |
| CNAG_01807 | UPF1 | ATP-dependent helicase | 958.4316172 | 862.9801111 | 924.52731 | 718.883928 | 663.0636962 | 732.8156879 | 0.000100398 | -0.39254014 | 3.59E-05 | -4.13257013 | 0.761787151 |
| CNAG_03086 | FZC20 | hypothetical protein, variant | 1002.927868 | 951.3379871 | 1095.110395 | 758.4798457 | 790.7950695 | 816.0783483 | 0.000101559 | -0.382197234 | 3.63E-05 | -4.1298491 | 0.767268146 |
| CNAG_05760 | - | endoplasmic reticulum protein | 7.492442848 | 9.900330687 | 8.717574623 | 41.5362632 | 41.18105524 | 25.4749891 | 0.000101807 | 2.141392573 | 3.64E-05 | 4.129209058 | 4.411877008 |
| CNAG_05172 | - | myosin-1 | 2791.065937 | 2710.903476 | 2588.379444 | 3081.276994 | 3564.34253 | 3385.865973 | 0.000102603 | 0.29485519 | 3.67E-05 | 4.127339451 | 1.226761836 |
| CNAG_01712 | - | hypothetical protein | 827.5299152 | 716.4366263 | 726.1970038 | 545.9774557 | 553.541542 | 602.9200021 | 0.000103703 | -0.432203467 | 3.71E-05 | -4.124807732 | 0.741128974 |
| CNAG_06424 | - | claudin family protein | 107.8962014 | 114.3981382 | 130.2183586 | 215.508049 | 216.0779647 | 174.2171122 | 0.000103979 | 0.770048917 | 3.72E-05 | 4.124116869 | 1.705327604 |
| CNAG_05037 | - | hypothetical protein | 16.93748015 | 22.85244995 | 28.93825683 | 55.70045664 | 60.77217383 | 72.72909504 | 0.00010437 | 1.487918466 | 3.74E-05 | 4.123173311 | 2.804839982 |
| CNAG_07927 | - | hypothetical protein, variant | 632.3929469 | 511.7336138 | 542.7420758 | 449.7100497 | 362.8250475 | 367.7039589 | 0.000105133 | -0.532397031 | 3.77E-05 | -4.121418409 | 0.691405013 |
| CNAG_04149 | - | nuclear pore complex protein Nup107 | 3611.622392 | 3397.08993 | 3305.978099 | 2706.574175 | 2915.344992 | 2961.983108 | 0.000105184 | -0.280542804 | 3.77E-05 | -4.121227363 | 0.823281205 |
| CNAG_01886 | - | hypothetical protein | 189.6876347 | 233.9419495 | 248.5737869 | 395.3752932 | 370.402388 | 295.2904004 | 0.000105241 | 0.646938912 | 3.77E-05 | 4.121024845 | 1.565842289 |
| CNAG_07000 | - | hypothetical protein | 3612.894017 | 3419.755277 | 3848.729861 | 2973.60032 | 3092.17537 | 3009.757698 | 0.000105888 | -0.277393105 | 3.80E-05 | -4.119533268 | 0.82508056 |
| CNAG_02426 | - | pre-mRNA-splicing factor syf1 | 2950.665852 | 2808.797739 | 2641.589883 | 2248.978901 | 2384.014338 | 2334.772628 | 0.000106098 | -0.285301433 | 3.81E-05 | -4.118997941 | 0.820570143 |
| CNAG_04265 | - | hypothetical protein | 3497.072826 | 3096.492177 | 2980.722216 | 2448.934936 | 2636.719937 | 2686.674927 | 0.000106138 | -0.3167574 | 3.81E-05 | -4.118832293 | 0.802872386 |
| CNAG_02082 | - | DASH complex subunit DAD1 | 399.4344494 | 564.2957754 | 510.6807312 | 352.2479052 | 298.9573762 | 348.7561278 | 0.000106282 | -0.575323574 | 3.81E-05 | -4.118443221 | 0.671135709 |
| CNAG_06236 | NGL3 | RNA exonuclease NGL2 | 1724.578909 | 1707.650867 | 1631.638594 | 1410.8453 | 1364.411379 | 1383.376125 | 0.000107317 | -0.299362125 | 3.85E-05 | -4.116130029 | 0.812611606 |
| CNAG_03381 | - | hypothetical protein | 176.8516504 | 233.9098265 | 231.5173049 | 322.5769464 | 318.4907086 | 354.9422244 | 0.000108634 | 0.620904019 | 3.90E-05 | 4.113237514 | 1.537838518 |
| CNAG_02979 | - | hypothetical protein | 95.02132011 | 116.4687445 | 85.52204312 | 168.546128 | 181.5289175 | 176.2443019 | 0.000108874 | 0.812334359 | 3.91E-05 | 4.112649119 | 1.756050531 |
| CNAG_00077 | - | solute carrier family 26 (sodium-independent sulfate anion transporter), member 11 | 2428.570241 | 2167.734663 | 2052.910127 | 1716.589853 | 1879.429579 | 1741.987666 | 0.000109565 | -0.332709611 | 3.94E-05 | -4.111111345 | 0.794043742 |
| CNAG_06079 | - | Proliferating cell nuclear antigen (PCNA) | 5138.010394 | 5270.731939 | 5086.950451 | 4466.432404 | 4461.055544 | 4462.935249 | 0.000111827 | -0.225765556 | 4.02E-05 | -4.106313072 | 0.85514113 |
| CNAG_02694 | - | hypothetical protein, variant | 54.27087686 | 62.70317612 | 76.89594038 | 138.5984773 | 123.6723448 | 119.6790767 | 0.00011248 | 0.96982098 | 4.05E-05 | 4.104809342 | 1.958597543 |
| CNAG_05077 | LPI11 | glycosyl hydrolase | 3700.420055 | 3789.222303 | 3323.071296 | 3007.877241 | 3020.560819 | 3008.642743 | 0.00011248 | -0.274031729 | 4.05E-05 | -4.104823807 | 0.82700518 |
| CNAG_04591 | - | hypothetical protein | 1.677219231 | 10.97393221 | 12.9478685 | 38.96589463 | 40.2333983 | 35.23227518 | 0.000112733 | 2.283650068 | 4.06E-05 | 4.104211044 | 4.869082915 |
| CNAG_03725 | - | dTDP-4-dehydrorhamnose reductase | 8094.621288 | 7159.533661 | 7724.263939 | 6588.842497 | 6326.735218 | 6447.917418 | 0.000113121 | -0.262840682 | 4.07E-05 | -4.103338159 | 0.83344524 |
| CNAG_03887 | - | hypothetical protein | 305.6380874 | 316.5875775 | 289.9908679 | 187.8219783 | 210.2431035 | 212.2103878 | 0.000113402 | -0.594471535 | 4.08E-05 | -4.102685678 | 0.662287009 |
| CNAG_02589 | - | hypothetical protein | 372.8765532 | 417.0715659 | 425.5945444 | 550.5351473 | 553.0847249 | 579.2976878 | 0.000114714 | 0.455552185 | 4.13E-05 | 4.099947004 | 1.371307567 |
| CNAG_05200 | BUD16 | pyridoxal kinase | 512.3985869 | 551.4634238 | 475.5957177 | 349.0047019 | 391.4433351 | 375.397967 | 0.000115435 | -0.478910387 | 4.16E-05 | -4.098417865 | 0.717519335 |
| CNAG_01878 | - | myo-inositol 2-dehydrogenase | 1422.098864 | 1298.280527 | 1427.889424 | 1118.945048 | 1088.37391 | 1121.37791 | 0.000115441 | -0.33347722 | 4.16E-05 | -4.098327458 | 0.79362137 |
| CNAG_01034 | - | hypothetical protein | 81.0617851 | 85.32377433 | 102.4877963 | 159.2784082 | 152.9229137 | 173.9269439 | 0.000116443 | 0.837655389 | 4.20E-05 | 4.096248131 | 1.787143385 |
| CNAG_02026 | - | hypothetical protein | 149.8287818 | 130.5940044 | 143.0647252 | 222.8472889 | 199.2571557 | 333.4151803 | 0.000116492 | 0.822220724 | 4.20E-05 | 4.0960723 | 1.768125554 |
| CNAG_01141 | RRP40 | exosome complex component RRP40 | 1123.114162 | 1008.447899 | 1127.133969 | 902.662543 | 845.1369312 | 776.2130867 | 0.000118288 | -0.384671925 | 4.27E-05 | -4.092448004 | 0.76595316 |
| CNAG_06405 | - | hypothetical protein | 457.9484551 | 405.2731717 | 438.4104531 | 580.8615635 | 601.4191952 | 608.1824694 | 0.000119667 | 0.444955792 | 4.32E-05 | 4.089603333 | 1.361272403 |

| | | | | | | | | | | | | | |
|---|---|---|---|---|---|---|---|---|---|---|---|---|---|
| CNAG_02955 | - | hypothetical protein | 328.9404986 | 303.738554 | 319.8133549 | 215.1882672 | 204.9983379 | 224.4162939 | 0.000119667 | -0.578550188 | 4.32E-05 | -4.089605516 | 0.669636379 |
| CNAG_03818 | SSK1 | osomolarity two-component system, response regulator SSK1 | 483.6093726 | 441.8764644 | 418.1867563 | 597.9743138 | 607.4205653 | 646.9992839 | 0.000119685 | 0.447729326 | 4.32E-05 | 4.089490426 | 1.363891921 |
| CNAG_04717 | - | hypothetical protein | 4.274632586 | 26.21631683 | 31.21908858 | 175.5739908 | 157.9644701 | 125.0538062 | 0.000120116 | 2.893527273 | 4.34E-05 | 4.088578466 | 7.430850132 |
| CNAG_04799 | - | large subunit ribosomal protein L14e | 83355.22426 | 80220.21944 | 88994.38174 | 72674.23113 | 73158.01302 | 72352.61802 | 0.000120294 | -0.226709399 | 4.35E-05 | -4.088155997 | 0.854581861 |
| CNAG_07460 | - | heat shock transcription factor | 736.7132619 | 827.2988102 | 737.9296872 | 621.6702811 | 540.5603564 | 581.8757698 | 0.000120579 | -0.415457362 | 4.36E-05 | -4.087529049 | 0.749781762 |
| CNAG_00397 | - | 2-oxoisovalerate dehydrogenase E1 component, beta subunit | 170.9919874 | 217.7569166 | 215.5277416 | 330.5749731 | 280.4309835 | 333.85811 | 0.000121386 | 0.63324227 | 4.39E-05 | 4.085902782 | 1.551046856 |
| CNAG_01367 | - | hypothetical protein | 760.3672446 | 666.0074481 | 706.0957314 | 857.7113503 | 1061.311724 | 969.0065613 | 0.000121616 | 0.421402346 | 4.40E-05 | 4.085385171 | 1.339228696 |
| CNAG_02590 | - | 3-hydroxyisobutyrate dehydrogenase | 293.4647149 | 339.4742972 | 300.8710653 | 475.2853861 | 459.1507022 | 410.6838102 | 0.000123589 | 0.513378255 | 4.47E-05 | 4.081567693 | 1.427388699 |
| CNAG_00422 | - | hypothetical protein, variant | 1979.503533 | 1947.0069 | 1900.507534 | 2237.624207 | 2437.749545 | 2576.489668 | 0.000124705 | 0.300238094 | 4.52E-05 | 4.079399467 | 1.231347611 |
| CNAG_04281 | - | hypothetical protein | 2096.019353 | 2191.415255 | 2150.979733 | 1858.419292 | 1773.283031 | 1730.877658 | 0.00012627 | -0.278956427 | 4.58E-05 | -4.07618562 | 0.824186977 |
| CNAG_05522 | - | hypothetical protein | 2911.467418 | 3154.790888 | 3247.392248 | 3911.977351 | 3716.012854 | 3687.884091 | 0.00012627 | 0.265969675 | 4.58E-05 | 4.076199309 | 1.202443973 |
| CNAG_02481 | - | vacuolar protein | 1868.339668 | 2052.398274 | 1986.732852 | 1704.339335 | 1490.277339 | 1607.628177 | 0.00012627 | -0.313564272 | 4.58E-05 | -4.076282383 | 0.804651358 |
| CNAG_01564 | - | nucleolar protein 12 | 4333.293607 | 3781.793947 | 4465.123215 | 3046.577 | 3513.104629 | 3542.600905 | 0.00012627 | -0.332565562 | 4.58E-05 | -4.076323485 | 0.794123029 |
| CNAG_01868 | - | protein KTI12 | 718.9251417 | 717.3826559 | 562.0392351 | 487.6582649 | 510.4820038 | 445.4102108 | 0.000127432 | -0.484902957 | 4.62E-05 | -4.073976554 | 0.714545132 |
| CNAG_00247 | LYS9 | alpha-aminoadipic semialdehyde synthase | 824.9091779 | 1152.875064 | 991.9068183 | 1202.485873 | 1466.684962 | 1426.331138 | 0.000127519 | 0.450591816 | 4.63E-05 | 4.073737964 | 1.366600743 |
| CNAG_03604 | - | mRNA 3'-end-processing protein yth1 | 1356.72093 | 1299.325405 | 1321.273602 | 1033.25861 | 1081.510766 | 1100.266005 | 0.000127585 | -0.32235587 | 4.63E-05 | -4.07354075 | 0.799762826 |
| CNAG_07835 | - | hypothetical protein | 14.41402887 | 9.932420984 | 10.85941288 | 37.90194291 | 45.43912301 | 40.74169431 | 0.000128451 | 1.869884812 | 4.66E-05 | 4.071885813 | 3.655033963 |
| CNAG_12781 | - | unspecified product | 54.40960613 | 113.179816 | 72.73121451 | 146.7157456 | 181.7564365 | 156.258653 | 0.000128478 | 1.002342941 | 4.67E-05 | 4.071758754 | 2.003250645 |
| CNAG_00359 | - | hypothetical protein | 212.0332041 | 397.5719664 | 294.4975164 | 477.5360081 | 506.7018032 | 472.7223814 | 0.000130151 | 0.675453504 | 4.73E-05 | 4.068666373 | 1.597098734 |
| CNAG_01351 | - | hypothetical protein | 1758.469189 | 1744.293657 | 1726.535937 | 1445.852099 | 1492.138059 | 1363.436369 | 0.000130302 | -0.297297546 | 4.74E-05 | -4.06831843 | 0.813775332 |
| CNAG_02105 | - | methylated-DNA-protein-cysteine methyltransferase, variant | 255.3370659 | 229.4029161 | 255.7882875 | 150.4798046 | 150.6344474 | 172.2272529 | 0.000131655 | -0.662350948 | 4.79E-05 | -4.065831488 | 0.631847828 |
| CNAG_03976 | - | hypothetical protein | 2138.063936 | 2176.356953 | 2329.034102 | 1687.750656 | 1928.618978 | 1815.227046 | 0.000131658 | -0.306278666 | 4.79E-05 | -4.065747691 | 0.808725123 |
| CNAG_01795 | - | hypothetical protein | 218.8047901 | 273.749597 | 245.4084918 | 360.709632 | 352.9769874 | 384.9608179 | 0.000131844 | 0.560140118 | 4.80E-05 | 4.06534038 | 1.474412409 |
| CNAG_12057 | - | unspecified product | 47.23514846 | 59.42583246 | 50.31663514 | 131.1129232 | 116.0579017 | 85.45458988 | 0.000132084 | 1.074948628 | 4.81E-05 | 4.064837824 | 2.106647056 |
| CNAG_05774 | - | hypothetical protein, variant | 859.0148897 | 788.5557153 | 689.9850725 | 644.3135611 | 535.3766993 | 524.2310949 | 0.000132331 | -0.47133705 | 4.82E-05 | -4.064323812 | 0.721295811 |
| CNAG_01704 | IRK6 | serine/threonine protein kinase | 4514.716877 | 4896.959986 | 4370.418007 | 5434.952018 | 5460.974886 | 5622.848702 | 0.000132697 | 0.246546377 | 4.83E-05 | 4.063601149 | 1.186363714 |
| CNAG_07177 | - | NADH dehydrogenase (ubiquinone) Fe-S protein 3 | 421.1904336 | 397.4560178 | 382.7847488 | 300.1553549 | 273.1718301 | 273.3069359 | 0.000134825 | -0.51966758 | 4.91E-05 | -4.059809791 | 0.697532537 |
| CNAG_03872 | UBP7 | putative ubiquitin carboxyl-terminal hydrolase | 486.0617882 | 532.3234006 | 475.7584736 | 649.3174373 | 681.6379524 | 680.3594171 | 0.000135212 | 0.414140335 | 4.93E-05 | 4.058985069 | 1.332504436 |
| CNAG_07663 | - | hypothetical protein | 13.22797695 | 8.84761181 | 7.670171938 | 31.48375394 | 46.46136467 | 37.41763515 | 0.000135212 | 2.008847241 | 4.93E-05 | 4.059054647 | 4.024605129 |
| CNAG_07861 | - | hypothetical protein | 221.002085 | 233.9083211 | 194.2667487 | 350.8303066 | 329.7637879 | 304.0731353 | 0.000135744 | 0.586254705 | 4.95E-05 | 4.057988493 | 1.501344134 |
| CNAG_07770 | FRE1 | ferric-chelate reductase | 2210.679172 | 2181.867101 | 2003.970005 | 2573.19895 | 2616.279269 | 2636.517431 | 0.000135956 | 0.276182575 | 4.96E-05 | 4.057546835 | 1.210986326 |
| CNAG_01175 | - | hypothetical protein | 1484.195365 | 1638.684051 | 1711.54571 | 1329.113841 | 1322.172239 | 1252.418427 | 0.000136889 | -0.323346576 | 4.99E-05 | -4.055870924 | 0.799213814 |
| CNAG_04316 | UTR1 | NAD kinase | 2056.41806 | 2477.935192 | 2262.949361 | 1725.895885 | 1836.243883 | 1923.98293 | 0.000137043 | -0.323627843 | 5.00E-05 | -4.055451411 | 0.799058014 |
| CNAG_00355 | - | hypothetical protein | 687.4948556 | 677.5748111 | 623.8256545 | 529.0507773 | 424.9032347 | 507.4743985 | 0.000137043 | -0.459560069 | 5.00E-05 | -4.055502858 | 0.727207978 |
| CNAG_07026 | - | hypothetical protein | 18.19440438 | 38.95347035 | 33.24387348 | 108.3296583 | 62.21292431 | 70.83408228 | 0.000137081 | 1.418305745 | 5.01E-05 | 4.055307453 | 2.672714515 |
| CNAG_05793 | - | hypothetical protein | 1472.208219 | 1298.266919 | 1328.750734 | 1138.468003 | 1084.075699 | 1030.405206 | 0.000137751 | -0.349366916 | 5.03E-05 | -4.054090315 | 0.784928465 |
| CNAG_00625 | - | ARF guanyl-nucleotide exchange factor | 2763.071792 | 2764.769747 | 2811.228486 | 3285.311552 | 3413.30472 | 3295.990535 | 0.000138213 | 0.245893865 | 5.05E-05 | 4.053228638 | 1.185827258 |
| CNAG_07417 | - | hypothetical protein | 22.68465929 | 23.92401766 | 25.7695898 | 62.9566944 | 71.36207363 | 54.22418478 | 0.000138698 | 1.402084483 | 5.07E-05 | 4.052330969 | 2.642831569 |
| CNAG_00498 | CDC14 | cell division cycle protein 14 | 207.1397422 | 240.4009484 | 257.0968187 | 373.3643866 | 332.2447302 | 352.8292175 | 0.000140343 | 0.571556813 | 5.13E-05 | 4.049495579 | 1.486126386 |
| CNAG_06140 | - | long-chain fatty acid transporter | 7622.908533 | 6892.336803 | 7745.568237 | 6318.751885 | 6245.674682 | 6271.399778 | 0.000143475 | -0.256842611 | 5.25E-05 | -4.044247717 | 0.83691754 |
| CNAG_10125 | - | unspecified product | 7.442610172 | 5.608690903 | 8.69511594 | 36.83519102 | 25.19225376 | 34.00984217 | 0.00014461 | 2.232287804 | 5.29E-05 | 4.042323965 | 4.698785155 |
| CNAG_04026 | - | chromosome transmission fidelity protein 1 | 850.398243 | 940.6612946 | 892.704829 | 1117.823404 | 1148.233364 | 1154.428899 | 0.000145066 | 0.335416822 | 5.31E-05 | 4.041506349 | 1.261741897 |
| CNAG_02350 | - | hypothetical protein | 212.905428 | 222.1029233 | 232.5959226 | 288.7818167 | 361.7828992 | 369.3586118 | 0.000148321 | 0.59695447 | 5.43E-05 | 4.036223624 | 1.512520263 |
| CNAG_04047 | - | hypothetical protein, variant | 515.3932608 | 595.9320736 | 649.5186737 | 770.7211226 | 887.0799779 | 760.370699 | 0.000149121 | 0.443342423 | 5.46E-05 | 4.034881922 | 1.35975094 |
| CNAG_02891 | - | endoplasmic reticulum protein | 1252.816399 | 1338.026933 | 1174.139946 | 1028.704263 | 1039.277265 | 896.1174 | 0.000149888 | -0.360216652 | 5.49E-05 | -4.033598371 | 0.77904758 |
| CNAG_00985 | - | hypothetical protein, variant | 402.4988761 | 422.1326974 | 342.3030515 | 262.659209 | 221.2817991 | 298.7239064 | 0.000151804 | -0.592625535 | 5.56E-05 | -4.030536838 | 0.663134981 |
| CNAG_02810 | PUF4 | pumilio-family RNA binding protein, variant | 6743.524237 | 6153.183621 | 6150.208672 | 5155.11858 | 5358.103609 | 5574.179842 | 0.000153323 | -0.259180196 | 5.62E-05 | -4.028117247 | 0.835562589 |
| CNAG_05611 | - | cysteine synthase A | 1570.417671 | 1535.264686 | 1469.538818 | 1216.967094 | 1283.38035 | 1243.498946 | 0.000154028 | -0.304975284 | 5.65E-05 | -4.026960041 | 0.809456084 |
| CNAG_05642 | FZC37 | nuclear protein | 2392.615346 | 2458.62728 | 2411.202349 | 2009.314842 | 2001.856891 | 2101.605603 | 0.000154281 | -0.263821638 | 5.66E-05 | -4.026495235 | 0.832878734 |
| CNAG_06884 | - | hypothetical protein | 461.5132351 | 432.2036421 | 479.9830317 | 652.6278477 | 575.3937518 | 661.4294842 | 0.000154675 | 0.445230637 | 5.68E-05 | 4.025817102 | 1.361531761 |
| CNAG_01436 | SIT4 | serine/threonine-protein phosphatase 6 catalytic subunit | 1026.39734 | 1098.897766 | 1168.713283 | 894.8219635 | 858.9325501 | 857.1888425 | 0.000154921 | -0.350539194 | 5.69E-05 | -4.025365209 | 0.784290921 |
| CNAG_12114 | - | unspecified product | 307.4967712 | 359.9485509 | 340.3015635 | 479.8105344 | 457.3189606 | 482.7222377 | 0.000155677 | 0.480603235 | 5.72E-05 | 4.024063123 | 1.395326974 |
| CNAG_02569 | - | hypothetical protein | 630.4923642 | 710.9564983 | 677.1086068 | 525.0924168 | 524.278317 | 480.9000505 | 0.000155677 | -0.414493346 | 5.72E-05 | -4.024122407 | 0.750282937 |
| CNAG_06560 | - | esterase | 352.4074988 | 364.0255774 | 348.637716 | 251.8129858 | 252.5146034 | 243.3266219 | 0.000156618 | -0.527364615 | 5.76E-05 | -4.022565691 | 0.693820987 |
| CNAG_03729 | - | peroxin-5, variant | 274.6807248 | 236.1551183 | 279.5048745 | 406.7742423 | 413.508971 | 351.8519272 | 0.000159378 | 0.552378595 | 5.86E-05 | 4.018373152 | 1.466501549 |
| CNAG_06899 | - | 26S proteasome regulatory subunit N7 | 5307.592094 | 4853.963259 | 5581.849404 | 6186.75038 | 6594.452764 | 6402.197019 | 0.000161397 | 0.269379123 | 5.94E-05 | 4.015326835 | 1.205289009 |
| CNAG_07563 | - | glutamate-tRNA ligase | 1407.972185 | 1232.545691 | 1286.080399 | 934.2921214 | 1034.049698 | 1103.566685 | 0.000163319 | -0.369732124 | 6.01E-05 | -4.012455101 | 0.773926184 |
| CNAG_03522 | - | hypothetical protein | 263.5657181 | 223.0068935 | 300.4893891 | 171.5930922 | 167.0594379 | 162.3246305 | 0.000163465 | -0.668708288 | 6.02E-05 | -4.012166065 | 0.62906967 |
| CNAG_01714 | - | sulfonate dioxygenase | 50.45887784 | 26.15306851 | 34.33380069 | 90.28555536 | 77.00835958 | 95.02249868 | 0.000164143 | 1.235152362 | 6.04E-05 | 4.011111847 | 2.354062072 |
| CNAG_00643 | - | hypothetical protein | 922.2032573 | 843.5616686 | 837.1131732 | 643.2867414 | 721.0077668 | 645.1792563 | 0.000164855 | -0.388860339 | 6.07E-05 | -4.010011297 | 0.763732679 |
| CNAG_02705 | - | hypothetical protein, variant 1 | 104.5496746 | 177.8243705 | 136.6683826 | 247.3432554 | 227.9666369 | 239.53067 | 0.000165063 | 0.760660697 | 6.08E-05 | 4.009634654 | 1.694266354 |
| CNAG_01676 | - | tRNA (adenine-N(1)-)-methyltransferase non- catalytic subunit TRM6 | 1203.856445 | 1261.589159 | 1287.121665 | 999.8412894 | 965.8676743 | 1055.841301 | 0.000170142 | -0.328195082 | 6.27E-05 | -4.002394155 | 0.796532382 |
| CNAG_05330 | - | MFS transporter, SP family, general alpha glucoside:H symporter | 2.794760709 | 10.90519049 | 2.320105243 | 32.90470072 | 27.10247318 | 27.46048542 | 0.000170311 | 2.608211802 | 6.28E-05 | 4.002081314 | 6.097474425 |
| CNAG_12105 | - | unspecified product | 7.589917257 | 20.61664888 | 12.99722505 | 45.85531409 | 51.50673881 | 41.93597796 | 0.000172187 | 1.799960806 | 6.35E-05 | 3.999411206 | 3.482107653 |
| CNAG_05508 | - | hypothetical protein | 254.9445287 | 258.749461 | 302.9380863 | 370.6692158 | 398.741908 | 427.1281555 | 0.000172531 | 0.53861402 | 6.36E-05 | 3.998859625 | 1.452576374 |
| CNAG_00594 | - | lipid particle protein | 873.8285104 | 1038.700072 | 998.2706803 | 1212.759345 | 1320.830866 | 1221.078001 | 0.000173541 | 0.352792067 | 6.40E-05 | 3.997399347 | 1.27702969 |
| CNAG_12304 | - | unspecified product | 13.45010345 | 32.42041209 | 9.857866836 | 54.25473779 | 50.26873988 | 76.97034709 | 0.000174788 | 1.726298574 | 6.45E-05 | 3.995626143 | 3.308778175 |
| CNAG_03044 | - | hypothetical protein | 454.0524903 | 494.3697155 | 451.0341504 | 374.2026642 | 320.6107771 | 308.862047 | 0.000175789 | -0.496341305 | 6.49E-05 | -3.994194607 | 0.708902289 |
| CNAG_04848 | - | mRNA binding protein, variant | 2398.443903 | 2472.62617 | 2577.515834 | 2081.77121 | 1671.311671 | 2131.56236 | 0.000177064 | -0.354911634 | 6.54E-05 | -3.992403279 | 0.781917535 |
| CNAG_00772 | RAD23 | UV excision repair protein Rad23 | 636.6834384 | 646.5695513 | 643.1686273 | 793.1741965 | 984.5987972 | 814.7791449 | 0.000177375 | 0.413154095 | 6.55E-05 | 3.991908309 | 1.331593834 |
| CNAG_12264 | - | unspecified product | 2.899150356 | 14.20207466 | 11.92069932 | 60.09964784 | 51.65899186 | 20.14982537 | 0.000177558 | 2.266187065 | 6.56E-05 | 3.991586194 | 4.810500703 |
| CNAG_03950 | - | transcription elongation factor spt6 | 4674.765003 | 4970.273074 | 5055.045465 | 5385.710455 | 6321.911597 | 6500.903931 | 0.000178158 | 0.293636248 | 6.59E-05 | 3.99070801 | 1.225725774 |
| CNAG_07032 | - | peroxiredoxin 5, atypical 2-Cys peroxiredoxin | 3915.281537 | 3671.856218 | 3753.871372 | 3322.397124 | 3123.181194 | 3162.958898 | 0.000179473 | -0.254620246 | 6.64E-05 | -3.988884876 | 0.838207743 |
| CNAG_06122 | HAD6 | glycerol-1-phosphatase | 2033.991155 | 1862.848813 | 1945.152566 | 1559.416262 | 1579.191789 | 1666.426206 | 0.000179578 | -0.297293461 | 6.65E-05 | -3.988588961 | 0.813777636 |
| CNAG_12934 | - | unspecified product | 43.69564626 | 45.46722434 | 50.27936364 | 97.89612792 | 116.2632382 | 82.05244059 | 0.000179578 | 1.080399175 | 6.65E-05 | 3.988650259 | 2.114621089 |
| CNAG_01112 | - | xenobiotic reductase | 1621.107212 | 1847.833388 | 1675.48664 | 2262.865658 | 2072.828184 | 2068.157431 | 0.000181903 | 0.301303502 | 6.74E-05 | 3.985458268 | 1.232257278 |
| CNAG_01162 | MAK3201 | hypothetical protein | 1191.002582 | 1238.965336 | 1265.793521 | 931.2533961 | 1028.925215 | 1013.679545 | 0.000182785 | -0.328934631 | 6.77E-05 | -3.984231414 | 0.796124171 |
| CNAG_07424 | - | DNA-directed RNA polymerase I, II, and III subunit RPABC4 | 426.9847702 | 383.4741243 | 381.7148674 | 274.3350087 | 301.7466139 | 264.4445929 | 0.000184491 | -0.519975903 | 6.84E-05 | -3.981946044 | 0.697383481 |
| CNAG_02175 | - | 5'-3' exoribonuclease 1 | 4474.788484 | 5014.254868 | 4581.452299 | 3877.193467 | 4031.503126 | 4063.267501 | 0.000184601 | -0.247662191 | 6.84E-05 | -3.981726303 | 0.842260147 |
| CNAG_06156 | FZC7 | hypothetical protein, variant | 1823.29247 | 1243.38193 | 1358.623688 | 996.5164919 | 1115.978605 | 1110.306068 | 0.000185727 | -0.4748001 | 6.89E-05 | -3.980202175 | 0.719566486 |
| CNAG_01931 | - | hypothetical protein | 9.807628287 | 10.97952362 | 10.84226029 | 44.70097645 | 35.73017195 | 32.00985711 | 0.000187677 | 1.92144448 | 6.96E-05 | 3.977640728 | 3.788021397 |
| CNAG_07592 | - | hypothetical protein | 0.270400832 | 2.314903552 | 2.27092483 | 20.21708966 | 18.48677121 | 18.58113239 | 0.000188415 | 3.793817512 | 6.99E-05 | 3.976628996 | 13.86924657 |
| CNAG_02985 | - | nuclear mRNA splicing protein | 2241.802928 | 2066.458357 | 2008.103196 | 1633.304753 | 1671.466659 | 1837.366516 | 0.000190413 | -0.312151235 | 7.07E-05 | -3.974039504 | 0.805439854 |
| CNAG_02671 | - | pre-mRNA-splicing factor cef1 | 9708.512028 | 9115.049176 | 9348.487259 | 7676.001215 | 8004.445261 | 8441.70788 | 0.000190743 | -0.239409474 | 7.08E-05 | -3.973549625 | 0.847091975 |
| CNAG_03843 | ARK1 | NAK protein kinase, variant | 260.5511096 | 219.9585044 | 193.2185944 | 316.9875216 | 345.2965137 | 369.3735206 | 0.000192078 | 0.600475016 | 7.13E-05 | 3.971810257 | 1.516215707 |
| CNAG_03028 | - | hypothetical protein | 201.2151203 | 180.1489366 | 240.0107755 | 332.8784193 | 319.4174182 | 304.031567 | 0.000192295 | 0.60503189 | 7.14E-05 | 3.971463466 | 1.521012374 |
| CNAG_07573 | - | DNA ligase 4, variant | 926.1842848 | 877.1373605 | 876.7170769 | 1121.726422 | 1111.958098 | 1183.259652 | 0.000192653 | 0.334849218 | 7.16E-05 | 3.970941828 | 1.261245584 |

| | | | | | | | | | | | | | |
|---|---|---|---|---|---|---|---|---|---|---|---|---|---|
| CNAG_04148 | - | NEK protein kinase | 1534.558168 | 1446.014462 | 1466.451626 | 1822.963577 | 1817.540328 | 1837.203746 | 0.000193831 | 0.285166525 | 7.21E-05 | 3.969410531 | 1.218550908 |
| CNAG_07436 | - | hypothetical protein | 6.295495251 | 6.675604215 | 8.69378069 | 39.35438382 | 27.88907025 | 27.54141912 | 0.000197186 | 2.20196984 | 7.33E-05 | 3.965241682 | 4.601071386 |
| CNAG_04335 | - | phosphatidylinositol 4-kinase | 2335.223773 | 2295.894791 | 1960.15664 | 1773.471664 | 1751.691197 | 1824.092964 | 0.000197718 | -0.316348567 | 7.35E-05 | -3.96452004 | 0.803099938 |
| CNAG_12515 | - | unspecified product | 24.08378377 | 61.5199188 | 48.16453443 | 101.050775 | 112.7246201 | 95.22915739 | 0.000199195 | 1.198761891 | 7.41E-05 | 3.962666396 | 2.295425949 |
| CNAG_01821 | - | hypothetical protein | 669.2733916 | 636.8643625 | 554.71003 | 939.233719 | 801.4358168 | 790.3579668 | 0.00019947 | 0.429386999 | 7.42E-05 | 3.96225833 | 1.346661259 |
| CNAG_06096 | - | tricarboxylate carrier | 8445.12015 | 8916.755406 | 8548.65648 | 7534.46429 | 7770.750241 | 7222.829933 | 0.000199492 | -0.216948372 | 7.43E-05 | -3.962154165 | 0.86038342 |
| CNAG_06420 | - | solute carrier family 25 (mitochondrial citrate transporter), member 1 | 3220.380996 | 2867.020418 | 2942.303126 | 2339.984273 | 2505.610847 | 2606.722379 | 0.000200239 | -0.2930374 | 7.46E-05 | -3.961184019 | 0.816181888 |
| CNAG_05389 | - | sphinganine-1-phosphate aldolase | 1074.188658 | 1103.201507 | 1086.65068 | 837.1122277 | 887.4377446 | 899.3279912 | 0.000200448 | -0.330116468 | 7.47E-05 | -3.960855938 | 0.795472263 |
| CNAG_12093 | - | unspecified product | 337.2513597 | 377.9662066 | 350.7422678 | 222.8788561 | 182.4060515 | 286.4525264 | 0.000202529 | -0.639027847 | 7.55E-05 | -3.958232803 | 0.64214551 |
| CNAG_02151 | - | amidohydrolase | 952.717649 | 997.5837762 | 925.6358834 | 761.6889963 | 723.472819 | 790.5274923 | 0.000202529 | -0.352706841 | 7.55E-05 | -3.958243244 | 0.783113411 |
| CNAG_06722 | - | hypothetical protein | 104.2753069 | 97.15964898 | 101.4609094 | 157.490155 | 184.1449261 | 177.3435885 | 0.00020263 | 0.762853622 | 7.56E-05 | 3.958035114 | 1.69684363 |
| CNAG_12877 | - | unspecified product | 195.513018 | 253.2990873 | 250.7096648 | 385.1352273 | 323.5833442 | 350.6140709 | 0.000204151 | 0.585754721 | 7.62E-05 | 3.956170601 | 1.500823914 |
| CNAG_05640 | - | metal iron transporter | 1562.909618 | 2097.796789 | 2031.64395 | 2433.952113 | 2455.87324 | 2511.025919 | 0.000204227 | 0.364816137 | 7.62E-05 | 3.956002781 | 1.287717507 |
| CNAG_04503 | - | hypothetical protein | 1053.206358 | 1040.7605 | 1204.945547 | 881.5669709 | 886.5731758 | 825.0254969 | 0.00020531 | -0.362633394 | 7.66E-05 | -3.954660721 | 0.777743645 |
| CNAG_04331 | - | hypothetical protein | 41.35624587 | 45.44846476 | 38.60163188 | 128.4754625 | 90.029829 | 68.84417517 | 0.000205316 | 1.190765727 | 7.67E-05 | 3.954575552 | 2.2827387 |
| CNAG_05618 | - | cleft lip and palate associated transmembrane protein | 1145.44912 | 1196.936122 | 1175.167045 | 943.8103612 | 957.3007056 | 955.9637034 | 0.000205379 | -0.315521598 | 7.67E-05 | -3.954424022 | 0.803560416 |
| CNAG_01615 | - | aldo-keto reductase | 891.8732173 | 826.3249354 | 848.8206072 | 688.5823039 | 580.1970901 | 693.9447513 | 0.000206728 | -0.402659587 | 7.72E-05 | -3.952780022 | 0.75646247 |
| CNAG_13118 | - | unspecified product | 7.377299316 | 6.647778471 | 0.147951007 | 25.45183706 | 36.17241502 | 23.12944023 | 0.000208541 | 2.676431494 | 7.80E-05 | 3.95061267 | 6.392727041 |
| CNAG_01348 | - | cyanate hydratase, variant | 1166.783057 | 1263.865591 | 1049.523257 | 1519.545151 | 1469.087594 | 1437.568202 | 0.000213544 | 0.332677669 | 7.98E-05 | 3.944856915 | 1.259348581 |
| CNAG_13194 | - | unspecified product | 0.271293397 | 0.130475854 | 2.27140743 | 16.80656743 | 31.26487312 | 11.09902424 | 0.000213569 | 4.840532957 | 7.99E-05 | 3.944750442 | 28.65138464 |
| CNAG_03037 | - | hypothetical protein, variant | 1278.519977 | 1205.618119 | 1370.271631 | 1094.928431 | 995.204879 | 970.4551861 | 0.000215842 | -0.348520239 | 8.08E-05 | -3.942135055 | 0.785389252 |
| CNAG_03449 | - | DNA repair protein, variant | 370.166777 | 458.7964184 | 463.7351817 | 269.529176 | 290.4176365 | 335.3596314 | 0.00021625 | -0.544545868 | 8.09E-05 | -3.941603935 | 0.685607184 |
| CNAG_02702 | CLC-A | putative voltage-gated chloride channel | 2323.818428 | 2580.359384 | 2585.003268 | 2048.991226 | 2115.747341 | 2103.8527 | 0.000216282 | -0.271919264 | 8.10E-05 | -3.94149011 | 0.828217009 |
| CNAG_05486 | - | hypothetical protein | 2920.372604 | 2898.218843 | 2731.166005 | 2384.464936 | 2325.300989 | 2493.485105 | 0.000217265 | -0.262466123 | 8.14E-05 | -3.940323934 | 0.833661651 |
| CNAG_00730 | AFR1 | ATP-binding cassette transporter | 10413.81363 | 10101.94083 | 9390.117386 | 8163.993226 | 8893.803357 | 8533.900567 | 0.000217366 | -0.240193015 | 8.14E-05 | -3.940134967 | 0.846632036 |
| CNAG_05182 | - | hypothetical protein | 612.1534272 | 651.9231877 | 597.3259263 | 890.091872 | 772.0773042 | 798.0980504 | 0.000220629 | 0.388511879 | 8.27E-05 | 3.936480663 | 1.309042447 |
| CNAG_05728 | - | hypothetical protein | 2.716119997 | 2.336615425 | 4.408613997 | 25.88405818 | 19.25071231 | 20.82293584 | 0.000223964 | 2.984589485 | 8.40E-05 | 3.932798681 | 7.915000738 |
| CNAG_03466 | RDP1 | RNA-dependent RNA polymerase 1 | 1780.635627 | 1799.216833 | 1647.65255 | 1425.58676 | 1420.48452 | 1472.162485 | 0.000224517 | -0.290734665 | 8.42E-05 | -3.932127254 | 0.817485664 |
| CNAG_00137 | - | hypothetical protein | 669.5538446 | 831.8535069 | 859.6073372 | 1095.468141 | 1009.321321 | 1037.866032 | 0.000225153 | 0.398556358 | 8.45E-05 | 3.931213249 | 1.318188197 |
| CNAG_07622 | - | COP9 signalosome complex subunit 1 | 1877.631652 | 1926.374706 | 1906.762307 | 1534.51296 | 1561.947763 | 1658.642534 | 0.000225153 | -0.279473017 | 8.45E-05 | -3.931261724 | 0.823891911 |
| CNAG_00133 | - | hypothetical protein | 134.7595506 | 188.5922249 | 127.1133117 | 263.7353385 | 232.2338324 | 244.0098092 | 0.000225153 | 0.70525934 | 8.45E-05 | 3.931364077 | 1.630437728 |
| CNAG_02725 | - | 20S proteasome subunit beta 2 | 1988.846982 | 1922.237928 | 2032.712799 | 2401.218703 | 2373.870378 | 2434.470192 | 0.000225864 | 0.263258151 | 8.48E-05 | 3.930377058 | 1.20018612 |
| CNAG_03091 | - | hypothetical protein | 7.523701869 | 15.24091698 | 8.731595473 | 34.74656832 | 35.705693 | 46.07419361 | 0.000226235 | 1.925998482 | 8.50E-05 | 3.929903868 | 3.799997533 |
| CNAG_12945 | - | unspecified product | 0.305964016 | 5.538448517 | 2.28962511 | 17.43124528 | 31.05308247 | 18.68230581 | 0.000227827 | 3.224901053 | 8.56E-05 | 3.928139565 | 9.349576726 |
| CNAG_07792 | - | hypothetical protein | 6.357375435 | 14.16601005 | 8.724523692 | 37.30641441 | 33.96777166 | 40.63344086 | 0.0002284 | 1.961376055 | 8.58E-05 | 3.927456592 | 3.894332468 |
| CNAG_01363 | - | XPA-binding protein 1 | 913.1226356 | 1035.276014 | 976.7993495 | 780.3795835 | 752.8233617 | 786.1090555 | 0.000229251 | -0.349740199 | 8.62E-05 | -3.926484071 | 0.784725399 |
| CNAG_03017 | - | transcription initiation factor TFIIF subunit alpha | 2122.827398 | 2040.622717 | 2213.867439 | 1667.560134 | 1719.785746 | 1864.010754 | 0.000229661 | -0.295511344 | 8.64E-05 | -3.925975429 | 0.814783492 |
| CNAG_06591 | SET302 | histone deacetylase complex subunit, putative | 3458.844926 | 3633.04455 | 3779.431715 | 3082.612726 | 3156.88193 | 3027.523295 | 0.000229666 | -0.245665021 | 8.64E-05 | -3.925892907 | 0.843426923 |
| CNAG_06056 | - | hypothetical protein | 448.2464069 | 508.3536126 | 439.3256022 | 331.9544989 | 352.6182088 | 334.3385297 | 0.000232151 | -0.470664513 | 8.74E-05 | -3.923223133 | 0.721632134 |
| CNAG_06170 | - | S-formylglutathione hydrolase | 1572.815502 | 1631.132951 | 1510.069064 | 1317.451069 | 1292.836165 | 1285.686074 | 0.000232631 | -0.290410353 | 8.76E-05 | -3.922648171 | 0.817669451 |
| CNAG_00358 | - | tRNA dimethylallyltransferase | 1077.635554 | 1033.20174 | 1069.579256 | 827.8004718 | 886.6061039 | 837.2003657 | 0.000233278 | -0.33358502 | 8.79E-05 | -3.921900687 | 0.793562072 |
| CNAG_00070 | - | hypothetical protein | 124.1137331 | 139.0830203 | 91.94712979 | 205.2509823 | 203.9000325 | 195.177856 | 0.000233741 | 0.753920138 | 8.81E-05 | 3.921345146 | 1.686368867 |
| CNAG_04027 | - | hypothetical protein | 600.1166338 | 115.4805727 | 158.9227718 | 26.01461622 | 40.67377445 | 43.86654381 | 0.00023478 | -3.02414682 | 8.85E-05 | -3.92019804 | 0.122925249 |
| CNAG_07578 | - | hypothetical protein | 51.79789714 | 46.55717819 | 53.4789653 | 88.18624377 | 118.7909415 | 102.9513847 | 0.000236461 | 1.022084299 | 8.91E-05 | 3.918400814 | 2.030850864 |
| CNAG_02995 | - | RNA-binding protein with serine-rich domain 1 | 3320.608969 | 2584.766683 | 2773.802028 | 2130.567981 | 2251.964815 | 2444.642036 | 0.000237528 | -0.362634124 | 8.96E-05 | -3.917236539 | 0.777743251 |
| CNAG_00603 | - | pre-rRNA-processing protein PNO1 | 1502.703339 | 1337.100113 | 1648.585264 | 1130.51038 | 1159.958676 | 1244.573591 | 0.00023855 | -0.360701553 | 9.00E-05 | -3.916123725 | 0.77878578 |
| CNAG_12762 | - | unspecified product | 7.626123401 | 18.52181984 | 22.53347809 | 57.85196113 | 54.8930612 | 42.03703887 | 0.000239509 | 1.688878842 | 9.04E-05 | 3.915077351 | 3.224060557 |
| CNAG_00660 | - | histone deacetylase HOS3 | 113.7172326 | 119.7790971 | 136.6094977 | 220.1542413 | 174.2959923 | 223.8952182 | 0.000240064 | 0.730689575 | 9.06E-05 | 3.914440329 | 1.659432073 |
| CNAG_05788 | - | V-type ATPase, G subunit | 1533.196534 | 1669.918611 | 1641.197803 | 1410.16096 | 1289.358059 | 1252.424257 | 0.000241159 | -0.308598009 | 9.11E-05 | -3.913264707 | 0.807426023 |
| CNAG_03900 | - | f-actin-capping protein subunit beta | 1538.9766 | 1648.355301 | 1498.338154 | 1252.772567 | 1298.888728 | 1306.756434 | 0.00024236 | -0.295519816 | 9.15E-05 | -3.911986858 | 0.814778707 |
| CNAG_06368 | - | hypothetical protein | 781.1336861 | 841.3683469 | 869.0426959 | 584.0557548 | 709.8334409 | 616.314056 | 0.000244287 | -0.399166685 | 9.23E-05 | -3.909996885 | 0.758296156 |
| CNAG_03719 | - | Putative transporter of the major facilitator superfamily (MFS) | 204.8657504 | 288.8083648 | 243.2858439 | 330.892725 | 421.4018023 | 372.8106749 | 0.000244858 | 0.597376081 | 9.25E-05 | 3.909354358 | 1.512962344 |
| CNAG_00917 | - | alkylbase DNA N-glycosylase | 918.8757613 | 966.3455565 | 918.159368 | 702.4382373 | 761.5094607 | 762.7846804 | 0.000249043 | -0.347682876 | 9.42E-05 | -3.905179674 | 0.785845237 |
| CNAG_12930 | - | unspecified product | 12.12944241 | 13.12913441 | 10.85759842 | 36.2773258 | 44.5599299 | 40.73242009 | 0.00024971 | 1.762594749 | 9.44E-05 | 3.904454905 | 3.393078362 |
| CNAG_06956 | - | hypothetical protein | 606.5996892 | 865.2328713 | 852.1520584 | 1117.292181 | 996.3512903 | 1111.057829 | 0.000249916 | 0.459055762 | 9.45E-05 | 3.904177985 | 1.374641826 |
| CNAG_04535 | - | Tartrate dehydrogenase | 43.63565552 | 44.36841025 | 40.71426642 | 100.5082395 | 76.91131254 | 96.21102783 | 0.000251807 | 1.084035485 | 9.53E-05 | 3.902275766 | 2.119957709 |
| CNAG_02494 | - | carboxypeptidase D | 728.9019802 | 700.4588436 | 750.8560768 | 859.263285 | 1025.01531 | 975.6636697 | 0.000256244 | 0.376933253 | 9.70E-05 | 3.897968621 | 1.298578522 |
| CNAG_06594 | - | oxysterol binding protein | 5714.744898 | 5171.653546 | 5266.115135 | 4612.744762 | 4658.591873 | 4501.816737 | 0.00025978 | -0.245712017 | 9.84E-05 | -3.894570135 | 0.843399448 |
| CNAG_02138 | - | DNA replication ATP-dependent helicase dna2 | 259.5623324 | 264.1101413 | 275.2446105 | 421.6758792 | 355.4824875 | 378.4038724 | 0.000260191 | 0.517094776 | 9.86E-05 | 3.894108841 | 1.43107053 |
| CNAG_12633 | - | unspecified product | 7.391492452 | 2.371309975 | 7.617932995 | 25.39590935 | 30.71460677 | 29.60335474 | 0.000260993 | 2.374598566 | 9.89E-05 | 3.893284418 | 5.185915022 |
| CNAG_01816 | - | hypothetical protein, variant | 16.90234847 | 29.22747082 | 16.22334919 | 66.42637852 | 63.53754387 | 45.42660927 | 0.000261262 | 1.498821581 | 9.90E-05 | 3.892955855 | 2.826117759 |
| CNAG_04172 | - | transcription factor C subunit 6 | 766.2726253 | 840.4411613 | 701.8603398 | 1037.826491 | 976.5489333 | 984.5988575 | 0.000261454 | 0.362823507 | 9.91E-05 | 3.892699836 | 1.285940157 |
| CNAG_07619 | - | ATP-binding cassette, subfamily G (WHITE), member 2, PDR, variant | 609.3924355 | 692.5890719 | 582.2470513 | 437.749862 | 521.7784078 | 436.50731 | 0.000261534 | -0.448639043 | 9.92E-05 | -3.892547173 | 0.732733742 |
| CNAG_03096 | - | UDP-glucose 4-epimerase | 1200.145 | 1019.250181 | 1172.996495 | 875.2769383 | 932.3157998 | 857.211161 | 0.000262217 | -0.364225775 | 9.95E-05 | -3.89183725 | 0.77688568 |
| CNAG_13085 | - | unspecified product | 403.158407 | 624.7016577 | 752.5525607 | 361.3623152 | 382.6928876 | 405.3908338 | 0.000263723 | -0.645976956 | 0.000100092 | -3.890369585 | 0.639059889 |
| CNAG_05763 | - | hypothetical protein | 332.853072 | 440.4498918 | 429.6134151 | 283.7254305 | 265.3870526 | 284.3727565 | 0.000264552 | -0.543763826 | 0.000100439 | -3.889529539 | 0.685978932 |
| CNAG_01764 | - | cytoplasmic protein | 1170.976487 | 1073.07093 | 1082.399525 | 850.3423025 | 934.0581155 | 872.726876 | 0.000270824 | -0.33977135 | 0.000102853 | -3.883761554 | 0.790166534 |
| CNAG_03260 | - | large subunit ribosomal protein L6 | 305.4846694 | 259.5922535 | 283.5524002 | 193.4261189 | 204.2539534 | 164.596972 | 0.000271013 | -0.610471299 | 0.000102958 | -3.883513567 | 0.654982697 |
| CNAG_12924 | - | unspecified product | 602.5401673 | 707.726544 | 736.759981 | 528.2202994 | 425.7447967 | 540.7450609 | 0.000272916 | -0.468285078 | 0.000103714 | -3.881734643 | 0.722823302 |
| CNAG_07584 | - | hypothetical protein | 92.79681721 | 134.7657505 | 114.2572231 | 185.643402 | 190.0543788 | 200.6357074 | 0.000274597 | 0.744100569 | 0.000104386 | 3.880163045 | 1.674929729 |
| CNAG_01173 | LAG1 | LAG1 family transcription factor, putative | 814.0343359 | 756.4572207 | 681.6014161 | 1026.178733 | 954.1192776 | 963.5063399 | 0.000275655 | 0.371372236 | 0.000104822 | 3.879149506 | 1.293582652 |
| CNAG_05422 | LIV11 | virulence related protein of unknown function | 2191.821773 | 2303.486661 | 2382.37671 | 1936.188312 | 1950.136603 | 1907.369238 | 0.00027584 | -0.262395395 | 0.000104926 | -3.87890784 | 0.833702522 |
| CNAG_07984 | - | hypothetical protein | 0.257293718 | 1.230682093 | 2.263754784 | 16.89951716 | 17.62981706 | 19.60711676 | 0.000275973 | 4.101017859 | 0.000105011 | 3.878713078 | 17.1604783 |
| CNAG_00411 | - | nucleoporin p58/p45 | 2588.741794 | 2389.735378 | 2691.633053 | 2036.489437 | 2170.957693 | 2164.903655 | 0.000276114 | -0.283277518 | 0.000105098 | -3.878510205 | 0.821722105 |
| CNAG_00973 | - | hypothetical protein | 805.408875 | 722.8971333 | 735.7884759 | 624.0405458 | 576.0062377 | 524.2209873 | 0.000276813 | -0.407556815 | 0.000105398 | -3.877817568 | 0.753899011 |
| CNAG_00904 | - | aflatoxin efflux pump AFLT | 337.7475959 | 243.4177057 | 227.1308432 | 174.7175906 | 136.7076441 | 182.2303311 | 0.000277153 | -0.729953349 | 0.000105561 | -3.877440698 | 0.60292341 |
| CNAG_06340 | - | pre-rRNA-processing protein TSR4 | 2569.078247 | 2865.891749 | 3090.45481 | 2252.857483 | 2484.975237 | 2240.455338 | 0.000280434 | -0.304249838 | 0.000106845 | -3.874497289 | 0.809863214 |
| CNAG_06136 | - | hypothetical protein | 802.6169435 | 1042.958757 | 845.8199295 | 1127.130365 | 1189.658243 | 1229.865441 | 0.000280982 | 0.384715126 | 0.000107088 | 3.873943445 | 1.305601958 |
| CNAG_00144 | - | hypothetical protein, variant | 177.8405648 | 183.3063458 | 185.679693 | 265.7114502 | 266.7318369 | 295.0107186 | 0.000281159 | 0.584855918 | 0.00010719 | 3.873712176 | 1.499889188 |
| CNAG_01164 | - | glutamine-fructose-6-phosphate transaminase (isomerizing) | 5268.909651 | 4518.863757 | 4336.317162 | 5831.174004 | 5743.909167 | 5874.851847 | 0.000284457 | 0.289314005 | 0.000108482 | 3.870792679 | 1.222059056 |
| CNAG_12756 | - | unspecified product | 12.15507441 | 13.14201024 | 14.03826838 | 40.27958547 | 46.27088955 | 40.78163305 | 0.000285889 | 1.676576708 | 0.000109063 | 3.869490563 | 3.196685264 |
| CNAG_12331 | - | unspecified product | 4.979306699 | 1.247778195 | 1.213130262 | 15.84734312 | 28.41701227 | 17.566283 | 0.000286059 | 3.303732532 | 0.000109163 | 3.869267194 | 9.874669977 |
| CNAG_05238 | - | hypothetical protein, variant | 5.098677351 | 7.713333814 | 4.443562675 | 29.6194964 | 28.93932709 | 24.21112364 | 0.000286135 | 2.325550206 | 0.000109227 | 3.869124343 | 5.012569052 |
| CNAG_04761 | RAS2 | Ras family protein | 170.8348681 | 170.3942531 | 182.470311 | 246.1616499 | 276.3383118 | 276.1767086 | 0.000287051 | 0.597007548 | 0.000109611 | 3.868267244 | 1.512575911 |
| CNAG_07963 | - | hypothetical protein | 6.303531412 | 7.746190138 | 8.697788581 | 44.309435 | 29.66007414 | 22.16092951 | 0.000287133 | 2.162453185 | 0.000109678 | 3.868120006 | 4.476754439 |
| CNAG_02488 | - | methionyl-tRNA formyltransferase | 61.2879374 | 70.25175959 | 91.80425593 | 126.3800222 | 126.1143763 | 174.8075384 | 0.000289655 | 0.925531788 | 0.000110747 | 3.865752191 | 1.89938424 |
| CNAG_04358 | ATM1 | iron-sulfur clusters transporter ATM1, mitochondrial, variant | 1045.994854 | 936.2432474 | 961.8824497 | 726.5467 | 772.6880962 | 816.0505141 | 0.000289655 | -0.362030899 | 0.000110739 | -3.865770518 | 0.778068512 |

| | | | | | | | | | | | | | |
|---|---|---|---|---|---|---|---|---|---|---|---|---|---|
| CNAG_04266 | - | YEATS domain-containing protein 4 | 1408.121358 | 1315.515684 | 1442.818286 | 1062.79985 | 1139.307826 | 1172.410815 | 0.000289655 | -0.319870099 | 0.000110691 | -3.8658758 | 0.80114201 |
| CNAG_03486 | - | peptidyl-prolyl cis-trans isomerase B | 2014.309521 | 2153.701658 | 2118.981468 | 1834.309119 | 1711.161921 | 1719.762317 | 0.000294235 | -0.270703044 | 0.000112535 | -3.861843735 | 0.828915506 |
| CNAG_01989 | - | hypothetical protein | 244.0762641 | 304.7376206 | 334.6344382 | 203.5617707 | 195.5459535 | 184.5319196 | 0.000295797 | -0.61446355 | 0.000113168 | -3.860472691 | 0.653172723 |
| CNAG_01123 | - | DRAP deaminase | 1314.613564 | 1209.902861 | 1230.62872 | 1097.323517 | 949.4673512 | 929.3928882 | 0.000298368 | -0.350693814 | 0.000114188 | -3.858279931 | 0.78420687 |
| CNAG_04038 | - | MFS quinate transporter QutD | 303.0373548 | 231.6217719 | 273.9352938 | 162.912525 | 160.9621272 | 197.723315 | 0.000298753 | -0.652947321 | 0.000114372 | -3.857886704 | 0.635979725 |
| CNAG_00960 | - | ATP-dependent RNA helicase DHX8/PRP22 | 4460.615661 | 4228.877219 | 4142.080918 | 3597.81693 | 3810.730413 | 3553.745904 | 0.000301328 | -0.242820462 | 0.000115394 | -3.855710002 | 0.845091546 |
| CNAG_06377 | - | solute carrier family 25 (mitochondrial phosphate transporter), member 3 | 45851.72997 | 47263.67577 | 48146.04547 | 38058.08593 | 42833.87264 | 41684.39201 | 0.000306175 | -0.219935773 | 0.000117288 | -3.851726843 | 0.85860366 |
| CNAG_04701 | - | hypothetical protein | 313.291709 | 350.2559474 | 322.1884483 | 441.5148008 | 519.6734409 | 431.7681766 | 0.000306319 | 0.482864448 | 0.00011738 | 3.851533484 | 1.397515659 |
| CNAG_07394 | - | hypothetical protein | 22.8467549 | 35.77630179 | 48.0936849 | 75.9188138 | 74.41451749 | 104.8205791 | 0.000309699 | 1.254635717 | 0.000118713 | 3.848767805 | 2.386068924 |
| CNAG_04211 | - | nuclear RNA export factor 1/2 | 1083.799219 | 996.7372816 | 1011.086567 | 1282.050292 | 1348.390539 | 1258.842429 | 0.000313095 | 0.315778714 | 0.000120054 | 3.846016734 | 1.244683307 |
| CNAG_01090 | - | alpha/beta hydrolase fold protein | 327.1921888 | 321.1882398 | 289.1553007 | 465.9051814 | 462.6226134 | 400.7142295 | 0.000313822 | 0.486709221 | 0.000120371 | 3.84536985 | 1.401244996 |
| CNAG_06785 | - | kynurenine 3-monooxygenase | 186.0204574 | 197.3004132 | 193.1496231 | 299.3620621 | 303.9559719 | 263.0222669 | 0.000314786 | 0.571438369 | 0.000120779 | 3.844539803 | 1.486004381 |
| CNAG_04246 | - | hypothetical protein | 4.978422368 | 2.326352808 | 0.119645562 | 26.03256327 | 14.78699111 | 20.77116289 | 0.000318818 | 3.306355443 | 0.000122365 | 3.841339178 | 9.892639082 |
| CNAG_05665 | - | hypothetical protein | 1873.987625 | 1812.154145 | 1641.267468 | 1431.790713 | 1440.320569 | 1503.242526 | 0.000319141 | -0.299304478 | 0.000122528 | -3.841012589 | 0.812644077 |
| CNAG_00650 | - | GRAM domain-containing protein | 275.6836186 | 223.1948797 | 185.7748465 | 337.3214302 | 392.927487 | 328.4812452 | 0.000319835 | 0.613946999 | 0.000122833 | 3.840401303 | 1.530440544 |
| CNAG_04229 | - | phytanoyl-CoA dioxygenase | 708.6844365 | 735.7887396 | 734.7037208 | 579.5868734 | 571.7141778 | 555.2373143 | 0.000319943 | -0.367706121 | 0.000122914 | -3.840240693 | 0.775013786 |
| CNAG_03235 | - | THO complex subunit 1 | 2065.596137 | 2094.443795 | 1969.711303 | 1652.817251 | 1667.172324 | 1796.294562 | 0.000325261 | -0.27571436 | 0.000124997 | -3.836113606 | 0.826041197 |
| CNAG_13019 | - | unspecified product | 23.9319474 | 39.9978088 | 30.05085771 | 69.00179645 | 80.78197315 | 69.68040158 | 0.000326112 | 1.230785012 | 0.000125363 | 3.835393429 | 2.346946593 |
| CNAG_01343 | - | ATP-dependent Clp protease ATP-binding subunit ClpX | 181.4131897 | 229.5555746 | 184.6527682 | 286.7061188 | 316.9057764 | 296.2266199 | 0.000329734 | 0.581872163 | 0.000126796 | 3.832599748 | 1.496790351 |
| CNAG_05855 | - | hypothetical protein | 1455.487021 | 1078.481038 | 1029.134068 | 904.0942868 | 918.4725668 | 799.5560212 | 0.000334726 | -0.459175785 | 0.000128757 | -3.828824796 | 0.727401706 |
| CNAG_07493 | - | hypothetical protein | 209.4351551 | 104.8977858 | 70.75273796 | 490.3648383 | 399.8957264 | 435.7268112 | 0.000336245 | 1.771487355 | 0.000129382 | 3.827632024 | 3.414057495 |
| CNAG_05546 | - | hypothetical protein | 674.7690879 | 643.1791037 | 719.7297143 | 547.714756 | 492.2810935 | 521.9193978 | 0.000337373 | -0.399618339 | 0.000129857 | -3.826729438 | 0.758058799 |
| CNAG_01921 | - | hypothetical protein | 40.70024349 | 51.47307039 | 63.61010225 | 16.44251444 | 13.33201295 | 24.62046673 | 0.000339089 | -1.553084297 | 0.000130559 | -3.825401612 | 0.340780739 |
| CNAG_12059 | - | unspecified product | 124.3720481 | 115.0900844 | 104.4146448 | 64.11302202 | 65.26830724 | 55.81679764 | 0.000342693 | -0.914040482 | 0.000131989 | -3.822718204 | 0.530696713 |
| CNAG_07490 | - | hypothetical protein | 5.117979417 | 9.850331404 | 4.450957568 | 35.36607898 | 26.17862682 | 25.32633639 | 0.000344257 | 2.243090729 | 0.000132633 | 3.82151758 | 4.734101799 |
| CNAG_06064 | PTP1 | Putative protein tyrosine phosphatase | 812.4185252 | 784.2483048 | 684.6559713 | 609.186862 | 548.341748 | 590.7400717 | 0.000346416 | -0.399255101 | 0.000133507 | -3.819897894 | 0.758249685 |
| CNAG_03406 | SPP101 | compass component spp1 | 702.8998404 | 777.7783408 | 746.4393996 | 577.9849268 | 584.6527692 | 582.9680609 | 0.00034739 | -0.367092736 | 0.000133925 | -3.819126924 | 0.775343366 |
| CNAG_06389 | - | hypothetical protein | 9.95226772 | 20.66505418 | 23.60042396 | 46.34626878 | 48.62761934 | 63.79953022 | 0.000347616 | 1.580733215 | 0.000134054 | 3.818888334 | 2.991218326 |
| CNAG_03145 | - | hypothetical protein | 126.7128012 | 198.3181664 | 177.1473354 | 283.9137243 | 248.5827797 | 275.0597422 | 0.000347796 | 0.672393427 | 0.000134166 | 3.81868265 | 1.593714744 |
| CNAG_12117 | - | unspecified product | 61.40107085 | 90.70039577 | 120.5564369 | 178.1799668 | 155.4882849 | 172.8686813 | 0.000350356 | 0.883581244 | 0.000135197 | 3.816794816 | 1.844949393 |
| CNAG_02537 | - | hypothetical protein | 0.16521652 | 0.080077893 | 0.069146833 | 8.865746165 | 9.727473805 | 12.97529364 | 0.000350974 | 5.700817433 | 0.000135478 | 3.816281365 | 52.01361601 |
| CNAG_03761 | - | mRNA 3'-end-processing protein rna14 | 1429.14222 | 1353.215465 | 1434.303171 | 1214.788449 | 1135.835881 | 1091.44979 | 0.000353088 | -0.308990258 | 0.000136337 | -3.81472081 | 0.807206525 |
| CNAG_05723 | - | Alcohol dehydrogenase | 267.8045508 | 352.3365329 | 247.6027854 | 423.7743115 | 411.6332544 | 438.2788023 | 0.000354415 | 0.540863825 | 0.000136893 | 3.813716202 | 1.454843356 |
| CNAG_06733 | RPN14 | 19S proteasome regulatory particle base assembly chaperone, putative | 784.8829239 | 671.4048307 | 768.9852446 | 985.6211683 | 1048.303995 | 896.9723233 | 0.000356048 | 0.381772041 | 0.000137567 | 3.8125024 | 1.302941256 |
| CNAG_02161 | ASP1 | Inositol heptakisphosphate kinase | 2976.475458 | 2939.188662 | 2972.15716 | 2459.930502 | 2556.525321 | 2592.2968 | 0.000356637 | -0.239672998 | 0.000137838 | -3.812015854 | 0.846937258 |
| CNAG_06534 | HMG1 | hydroxymethylglutaryl-CoA reductase (NADPH) | 1572.630778 | 1450.14557 | 1323.465711 | 1222.537183 | 1134.945977 | 1135.841677 | 0.000358212 | -0.331134727 | 0.000138491 | -3.810848648 | 0.794911014 |
| CNAG_02407 | - | Phospholipase D | 1362.899944 | 1393.166308 | 1095.411435 | 1735.193434 | 1591.526125 | 1645.137106 | 0.000363056 | 0.353884261 | 0.000140408 | 3.807449038 | 1.277996833 |
| CNAG_05017 | - | GabA permease, variant | 191.3949148 | 214.2613962 | 250.3577306 | 129.4543609 | 161.1839339 | 111.3586401 | 0.000364393 | -0.723417499 | 0.000140969 | -3.806461405 | 0.605661034 |
| CNAG_03269 | - | Aldehyde dehydrogenase | 1338.10576 | 1402.711764 | 1302.10437 | 1051.149424 | 1146.245384 | 1116.92577 | 0.000365151 | -0.301979871 | 0.000141307 | -3.805869324 | 0.811138473 |
| CNAG_00521 | - | hypothetical protein | 176.5807842 | 155.3309069 | 164.3655513 | 261.2173679 | 234.7537922 | 268.3806812 | 0.000366107 | 0.608785036 | 0.000141722 | 3.805144214 | 1.524974413 |
| CNAG_12161 | - | unspecified product | 149.0764902 | 136.7246143 | 156.5719987 | 81.15882166 | 92.04474264 | 85.72584566 | 0.000366666 | -0.789186225 | 0.000141983 | -3.804688597 | 0.578670408 |
| CNAG_06608 | - | hypothetical protein | 964.5727062 | 1167.776289 | 1106.884137 | 916.7065875 | 830.4560664 | 785.0836676 | 0.000366697 | -0.36946362 | 0.000142085 | -3.804510951 | 0.774070235 |
| CNAG_01987 | - | carbohydrate binding protein | 1707.252532 | 1971.578523 | 1873.692593 | 1508.075549 | 1571.479746 | 1518.81827 | 0.000366697 | -0.286768351 | 0.000142047 | -3.804576387 | 0.81973622 |
| CNAG_04394 | - | hypothetical protein | 16.86707013 | 15.3390474 | 26.77994148 | 46.26892425 | 53.8949314 | 62.7613564 | 0.000368923 | 1.482344274 | 0.000142992 | 3.802934473 | 2.794023731 |
| CNAG_00148 | - | hypothetical protein | 933.2468812 | 979.452813 | 891.6633519 | 1197.265268 | 1298.446454 | 1092.38048 | 0.000369565 | 0.340607975 | 0.000143286 | 3.802425859 | 1.266290116 |
| CNAG_05888 | - | hypothetical protein | 2242.986611 | 1928.609349 | 2220.258901 | 1820.283849 | 1764.676791 | 1577.737554 | 0.000372338 | -0.324278835 | 0.000144407 | -3.800496205 | 0.798697534 |
| CNAG_12224 | - | unspecified product | 12.0139789 | 6.685599924 | 5.530106808 | 40.888914 | 26.04170069 | 32.97505628 | 0.000372832 | 2.074719852 | 0.000144644 | 3.80009006 | 4.212626041 |
| CNAG_02948 | - | peptide chain release factor 1 | 11860.72632 | 11551.11423 | 11593.37874 | 9624.841927 | 10398.09765 | 10472.20245 | 0.000374858 | -0.214384001 | 0.000145475 | -3.798668594 | 0.8619141 |
| CNAG_05499 | - | SHO1 osmosensor | 3466.982996 | 3450.986047 | 3762.357766 | 2943.245121 | 3100.815965 | 3048.591984 | 0.000375622 | -0.247550286 | 0.000145818 | -3.798086093 | 0.842325481 |
| CNAG_03046 | - | hypothetical protein | 103.0961463 | 95.00449127 | 101.4521454 | 171.7691663 | 159.7970492 | 172.9083864 | 0.000377934 | 0.732149139 | 0.000146761 | 3.79648648 | 1.661111758 |
| CNAG_00391 | - | hypothetical protein | 186.5557058 | 183.0479597 | 208.8218557 | 131.2101144 | 102.2439926 | 124.5345997 | 0.000381905 | -0.707826012 | 0.00014835 | -3.79381527 | 0.612242028 |
| CNAG_12212 | - | unspecified product | 298.4157514 | 241.2923317 | 264.3699943 | 190.3735088 | 178.2961273 | 165.6680701 | 0.000382219 | -0.60735755 | 0.000148519 | -3.793533279 | 0.656397864 |
| CNAG_05326 | - | hypothetical protein | 44.70172573 | 38.98589926 | 30.08713074 | 76.70751073 | 99.02244329 | 74.20517808 | 0.000382471 | 1.136758999 | 0.000148664 | 3.793291438 | 2.198864949 |
| CNAG_04198 | - | hypothetical protein | 448.4893408 | 325.599779 | 380.8409224 | 537.336367 | 555.7397007 | 544.914659 | 0.000383124 | 0.488280009 | 0.000148964 | 3.79278983 | 1.402771485 |
| CNAG_06410 | - | ATP-dependent Clp endopeptidase, proteolytic subunit ClpP | 1598.292717 | 1284.32075 | 1521.726536 | 1107.1611 | 1245.460792 | 1103.674449 | 0.00038429 | -0.366262008 | 0.000149464 | -3.7919578 | 0.77578995 |
| CNAG_12894 | - | unspecified product | 5278.161213 | 5025.147776 | 6072.245479 | 4613.517474 | 4573.171984 | 4452.982745 | 0.000386327 | -0.279590342 | 0.000150304 | -3.790566803 | 0.823824912 |
| CNAG_05864 | - | hypothetical protein | 394.8930332 | 390.1271383 | 338.2169501 | 533.53913 | 540.2382064 | 477.3037732 | 0.000389218 | 0.45093389 | 0.000151476 | 3.788636925 | 1.366924813 |
| CNAG_07461 | - | hypothetical protein | 1.480678838 | 0.120185799 | 1.194800937 | 25.53780801 | 11.28735414 | 15.3243668 | 0.000389324 | 4.656380569 | 0.000151565 | 3.788491449 | 25.21797573 |
| CNAG_12222 | - | unspecified product | 21.35987028 | 15.30977716 | 13.00990325 | 47.34382526 | 54.07615823 | 43.08156906 | 0.000389985 | 1.538924439 | 0.00015187 | 3.787991921 | 2.905777905 |
| CNAG_01557 | CAM1 | Calmodulin | 9499.521626 | 9289.575747 | 8993.384166 | 8213.935757 | 8100.206788 | 8085.38799 | 0.000391783 | -0.202633083 | 0.000152618 | -3.786770822 | 0.868963159 |
| CNAG_06816 | - | hypothetical protein | 99.64986428 | 108.9607976 | 94.02379961 | 177.2387276 | 142.3540861 | 201.5847528 | 0.00039191 | 0.774369846 | 0.000152715 | 3.786611963 | 1.710442784 |
| CNAG_03100 | - | cytoplasmic protein | 626.2132874 | 675.6420195 | 702.8495261 | 853.2424886 | 876.5508017 | 851.3790483 | 0.000392926 | 0.350125827 | 0.000153159 | 3.785890424 | 1.274671795 |
| CNAG_01025 | - | hypothetical protein | 141.6382 | 146.6708072 | 141.9724186 | 216.7205379 | 232.3614308 | 219.6201349 | 0.000399372 | 0.624897236 | 0.000155721 | 3.781764522 | 1.542100977 |
| CNAG_02526 | - | hypothetical protein | 34.10146599 | 21.78278594 | 13.05383975 | 65.43544022 | 59.8970298 | 61.83296871 | 0.000402627 | 1.427572267 | 0.000157039 | 3.779665873 | 2.689936781 |
| CNAG_03480 | - | NAD epimerase domain-containing protein | 566.5011869 | 487.1555945 | 532.2504907 | 689.8092718 | 691.0654156 | 726.9675593 | 0.000403247 | 0.394267391 | 0.00015733 | 3.779205233 | 1.314275194 |
| CNAG_06606 | RHO104 | rho family GTPase | 686.7559036 | 639.0187954 | 567.4985251 | 879.109173 | 857.6043801 | 773.7333653 | 0.000406925 | 0.39173052 | 0.000158864 | 3.776787679 | 1.31196617 |
| CNAG_07610 | - | hypothetical protein | 5.010501801 | 2.33750445 | 1.221122626 | 35.21957801 | 19.24477491 | 12.2456105 | 0.000406925 | 3.02739383 | 0.000158847 | 3.776813644 | 8.15335499 |
| CNAG_02152 | - | hypothetical protein | 445.249714 | 462.3528957 | 510.8941996 | 603.3138854 | 751.7465567 | 599.4256525 | 0.000407061 | 0.447261042 | 0.000158967 | 3.776626086 | 1.363449288 |
| CNAG_03933 | - | Transmembrane protein | 395.4075599 | 336.0940268 | 346.5197783 | 261.9722705 | 262.0258804 | 236.6968971 | 0.000410435 | -0.520768367 | 0.000160335 | -3.774490697 | 0.697000518 |
| CNAG_12892 | - | unspecified product | 48.46348321 | 80.88664208 | 56.71722844 | 133.1695121 | 124.6237751 | 104.2392648 | 0.000411359 | 0.958778539 | 0.000160746 | 3.773852467 | 1.943663591 |
| CNAG_03908 | - | wd-repeat protein | 1215.546475 | 1261.603632 | 1320.174974 | 1012.285885 | 991.7551196 | 1086.913628 | 0.000413334 | -0.312396774 | 0.000161568 | -3.772579936 | 0.805302784 |
| CNAG_04051 | - | ATP-dependent RNA helicase A | 2134.432884 | 2010.440267 | 2007.024728 | 1588.924666 | 1726.729165 | 1791.857656 | 0.00041364 | -0.284364903 | 0.000161738 | -3.772317249 | 0.821102992 |
| CNAG_02396 | - | hypothetical protein | 1116.188029 | 1073.071794 | 1114.372449 | 867.4981475 | 923.6925074 | 894.9081631 | 0.000415445 | -0.314867254 | 0.000162495 | -3.771153112 | 0.803924959 |
| CNAG_04623 | - | hypothetical protein | 13.60232997 | 51.83153891 | 41.75971554 | 74.19169676 | 88.38748618 | 113.6539479 | 0.000415539 | 1.377727017 | 0.000162582 | 3.771018722 | 2.59858638 |
| CNAG_05969 | - | O-sialoglycoprotein endopeptidase | 15.77715088 | 30.31312579 | 22.58664005 | 45.23029942 | 62.572574 | 75.96463431 | 0.000416548 | 1.428702743 | 0.000163028 | 3.770335963 | 2.692045405 |
| CNAG_07414 | PAN6 | pantoate-beta-alanine ligase | 112.4997624 | 87.54638751 | 141.8925651 | 211.6546187 | 191.7760329 | 183.0059606 | 0.000417267 | 0.766505873 | 0.00016336 | 3.769827378 | 1.701144713 |
| CNAG_02858 | ADE12 | adenylosuccinate synthetase | 17604.84381 | 15233.84796 | 18884.62562 | 13348.11215 | 15125.80625 | 14237.92712 | 0.000417581 | -0.292440127 | 0.000163585 | -3.7694839 | 0.816519855 |
| CNAG_07372 | OXK1 | 6-phosphofructo-2-kinase/fructose-2, 6- bisphosphatase | 771.6019534 | 704.5899312 | 742.1698568 | 604.5500842 | 522.4417942 | 585.1752431 | 0.000417581 | -0.389212789 | 0.000163544 | -3.76954696 | 0.763546122 |
| CNAG_01808 | - | solute carrier family 25, member 34/35 | 1348.387605 | 1131.289889 | 1196.509359 | 976.4945861 | 1000.438029 | 955.99663 | 0.000418712 | -0.342367411 | 0.000164079 | -3.768731262 | 0.788745945 |
| CNAG_00808 | - | myosin regulatory light chain cdc4 | 2663.535531 | 2605.200439 | 2864.396368 | 2288.763303 | 2396.13537 | 2086.170891 | 0.000420078 | -0.280096484 | 0.000164666 | -3.767840476 | 0.82353594 |
| CNAG_04872 | - | mitochondrial protein | 801.306874 | 849.0851013 | 808.4576226 | 1007.281185 | 1044.725002 | 1045.639675 | 0.000422623 | 0.317918606 | 0.000165767 | 3.766176323 | 1.246530866 |
| CNAG_06436 | - | hypothetical protein | 238.141049 | 272.4406664 | 298.4028593 | 184.0826606 | 193.8977188 | 160.1436442 | 0.000422623 | -0.604556345 | 0.000165754 | -3.766196241 | 0.657673596 |
| CNAG_05407 | - | hypothetical protein | 641.3958712 | 724.09771 | 674.0921964 | 806.3519291 | 951.6883473 | 905.7306182 | 0.000425139 | 0.369782499 | 0.000166806 | 3.764615377 | 1.29215801 |
| CNAG_03153 | SEC14-1 | SEC14 cytosolic factor | 7677.868914 | 7567.850463 | 7684.822125 | 6743.73273 | 6664.90016 | 6773.163768 | 0.000428608 | -0.199486736 | 0.000168219 | -3.762506404 | 0.870860332 |
| CNAG_05551 | - | hypothetical protein | 1060.528893 | 1129.204475 | 1037.75399 | 1334.981861 | 1401.871991 | 1282.178197 | 0.000431346 | 0.301326224 | 0.000169347 | 3.760835821 | 1.232276686 |
| CNAG_01391 | - | cell division cycle protein 37 | 2613.508519 | 2320.905463 | 2366.549172 | 2821.364825 | 2965.5972 | 3239.245757 | 0.000431533 | 0.290591949 | 0.000169473 | 3.760649807 | 1.223142039 |
| CNAG_00726 | - | hypothetical protein | 10.82069671 | 5.598006613 | 4.454432322 | 32.8004689 | 30.64644734 | 25.34521337 | 0.00043478 | 2.178144749 | 0.000170801 | 3.75869686 | 4.525711904 |

| | | | | | | | | | | | | |
|---|---|---|---|---|---|---|---|---|---|---|---|---|---|
| CNAG_00748 | - | hypothetical protein | 356.5648992 | 421.3683085 | 431.9792487 | 516.1595789 | 585.1096164 | 556.0239016 | 0.000437949 | 0.441106246 | 0.000172099 | 3.756801225 | 1.357644959 |
| CNAG_07327 | - | hypothetical protein | 138.1716638 | 104.4054093 | 113.9582579 | 43.522989 | 54.79587009 | 81.12788443 | 0.00044001 | -1.011886653 | 0.000173016 | -3.755470199 | 0.495897325 |
| CNAG_04632 | - | uracil permease | 48.39294482 | 56.21718988 | 63.04593379 | 103.1809822 | 123.051917 | 99.73532056 | 0.00044001 | 0.945341781 | 0.000173008 | 3.755482213 | 1.925645026 |
| CNAG_01036 | - | geranylgeranyl transferase type-2 subunit beta | 598.7164037 | 523.5923949 | 564.0658679 | 444.9595425 | 418.1359925 | 409.8308829 | 0.000443876 | -0.422558158 | 0.000174591 | -3.753200899 | 0.74610048 |
| CNAG_04888 | - | hypothetical protein | 184.9309175 | 214.5316491 | 224.039121 | 292.904889 | 318.5844907 | 310.6394703 | 0.000444468 | 0.549816271 | 0.000174878 | 3.752789075 | 1.463899255 |
| CNAG_01585 | - | hypothetical protein | 92.55931047 | 86.36771244 | 73.7853138 | 179.267284 | 149.5387818 | 124.2966881 | 0.000445044 | 0.833237969 | 0.000175213 | 3.752308891 | 1.78167966 |
| CNAG_03567 | CBK1 | RAM signaling network protein kinase, putative | 3443.531148 | 3381.997556 | 3199.345625 | 2874.825023 | 2842.013625 | 2889.833034 | 0.000445044 | -0.235291964 | 0.000175178 | -3.752359648 | 0.849513062 |
| CNAG_05411 | LPI9 | Endoglucanase | 810.6035496 | 798.4718571 | 795.6571827 | 993.2899895 | 997.2453211 | 1043.394816 | 0.000445385 | 0.320679355 | 0.000175402 | 3.752039127 | 1.248918517 |
| CNAG_03402 | - | hypothetical protein | 1149.336739 | 1225.12467 | 1284.054695 | 1503.859214 | 1586.459999 | 1454.25074 | 0.000446694 | 0.297386632 | 0.000176026 | 3.751148386 | 1.228916277 |
| CNAG_02766 | - | hypothetical protein | 245.5930758 | 244.7476496 | 297.5905414 | 389.5879805 | 358.1760005 | 388.3416831 | 0.000446694 | 0.514758035 | 0.000176009 | 3.751172659 | 1.428754493 |
| CNAG_02991 | - | Cofilin | 4225.947104 | 4111.437182 | 4186.844187 | 3825.260698 | 3470.800608 | 3315.099168 | 0.000448351 | -0.254436168 | 0.000176734 | -3.75014204 | 0.8383147 |
| CNAG_00029 | - | D-3-phosphoglycerate dehydrogenase | 61.25015292 | 83.08866453 | 71.61588747 | 133.6475626 | 111.3533216 | 156.0385273 | 0.000449248 | 0.881911162 | 0.000177143 | 3.749563211 | 1.842814892 |
| CNAG_00867 | ITR3A | myo-inositol transporter, putative | 15.64761519 | 14.23852402 | 17.23590287 | 30.25713471 | 50.54472019 | 63.66517193 | 0.000459097 | 1.620927711 | 0.000181082 | 3.744043551 | 3.075727544 |
| CNAG_05042 | - | carnitine acetyltransferase | 553.7084125 | 571.1086995 | 485.3786356 | 747.5745022 | 674.61879 | 714.7962109 | 0.000459685 | 0.393222781 | 0.00018137 | 3.743644374 | 1.313323913 |
| CNAG_01556 | - | cytoplasmic protein | 1236.458769 | 1143.133475 | 1263.654931 | 984.2921001 | 1000.437945 | 981.5065785 | 0.000462274 | -0.312353635 | 0.000182448 | -3.742155811 | 0.805326864 |
| CNAG_05325 | - | hypothetical protein | 61.19175471 | 61.62849996 | 76.89242914 | 125.9000708 | 136.7949125 | 108.6776575 | 0.000462417 | 0.882410116 | 0.000182561 | 3.742000149 | 1.843452338 |
| CNAG_13178 | - | unspecified product | 59.98831795 | 49.82015847 | 71.56332498 | 94.24508255 | 145.6545369 | 116.3017513 | 0.000463627 | 0.971361135 | 0.000183095 | 3.741266282 | 1.960689568 |
| CNAG_12565 | - | unspecified product | 2.717242315 | 1.256768122 | 5.463817873 | 21.64412995 | 31.95909802 | 12.24433366 | 0.000464523 | 2.982103876 | 0.000183506 | 3.740703399 | 7.901375782 |
| CNAG_02363 | - | SCR1 protein | 140.4481305 | 146.6556485 | 121.7571666 | 240.4267254 | 235.0170242 | 183.1582932 | 0.000466122 | 0.677891186 | 0.000184195 | 3.739761702 | 1.599799589 |
| CNAG_07535 | - | small subunit ribosomal protein S16 | 642.1641666 | 653.9405034 | 747.4268516 | 482.8962299 | 553.6508919 | 523.0285412 | 0.000467 | -0.405742536 | 0.000184599 | -3.739210986 | 0.754847683 |
| CNAG_12710 | - | unspecified product | 20.63986676 | 72.2712823 | 54.56289114 | 95.20833935 | 118.6684745 | 133.7545678 | 0.000469655 | 1.233471202 | 0.000185706 | 3.73770759 | 2.351320501 |
| CNAG_06343 | - | hypothetical protein, variant | 435.4155832 | 440.5878131 | 480.8324981 | 339.8123019 | 335.3360983 | 332.1061638 | 0.000469781 | -0.444964266 | 0.000185813 | -3.737562579 | 0.734602512 |
| CNAG_00764 | - | ATP-dependent RNA helicase DBP4 | 1684.804239 | 1507.303096 | 1650.781982 | 1141.290159 | 1445.638598 | 1259.069631 | 0.000471021 | -0.348370516 | 0.000186361 | -3.736821893 | 0.785470764 |
| CNAG_05817 | GMT1 | GDP-mannose transporter 1 | 6820.835676 | 7336.175522 | 7472.571961 | 6265.841771 | 6412.200514 | 6165.936112 | 0.000471916 | -0.214000693 | 0.00018863 | -3.736188963 | 0.862143131 |
| CNAG_06355 | - | hypothetical protein | 364.2018763 | 426.4565623 | 375.3157549 | 306.4651943 | 272.3309301 | 252.2539218 | 0.000471916 | -0.503733972 | 0.000186826 | -3.736194166 | 0.705279019 |
| CNAG_05733 | - | cytoplasmic protein | 190.5800979 | 163.9558741 | 178.2226905 | 297.9911035 | 264.1633705 | 248.5738759 | 0.000473262 | 0.591639878 | 0.000187421 | 3.735394949 | 1.506958699 |
| CNAG_00809 | - | ATP-dependent RNA helicase DED1 | 2541.786144 | 2155.904036 | 2079.576116 | 1954.179289 | 1773.24221 | 1722.032424 | 0.000475681 | -0.330096957 | 0.000188437 | -3.734034291 | 0.795483021 |
| CNAG_05699 | - | hypothetical protein | 273.4824308 | 256.5704964 | 259.2648234 | 386.4756415 | 353.7856688 | 378.373382 | 0.000478923 | 0.48803227 | 0.000189779 | 3.732246701 | 1.402530622 |
| CNAG_12834 | - | unspecified product | 13.38888738 | 22.78303493 | 16.19318982 | 34.22402628 | 58.4501456 | 59.40939546 | 0.000479706 | 1.552234326 | 0.000190148 | 3.731757521 | 2.932709814 |
| CNAG_12182 | - | unspecified product | 4.048793121 | 16.30102946 | 10.84468555 | 36.41117878 | 33.92909482 | 44.98586671 | 0.000481377 | 1.901723414 | 0.00019087 | 3.730804116 | 3.736592959 |
| CNAG_02016 | - | hypothetical protein | 2.713297163 | 4.473259083 | 3.349926552 | 18.28846718 | 29.24687412 | 16.53294459 | 0.000482117 | 2.792545935 | 0.000191222 | 3.730339507 | 6.928513869 |
| CNAG_03916 | - | glucose-6-phosphate isomerase | 6938.678876 | 6822.26795 | 7067.331691 | 5720.939908 | 6313.037247 | 6036.029205 | 0.00048226 | -0.220404984 | 0.000191337 | -3.73018731 | 0.85832446 |
| CNAG_06115 | APH4 | Acid phosphatase | 224.6412497 | 258.7126413 | 286.933381 | 361.4341085 | 363.3367048 | 384.9922386 | 0.000487741 | 0.513907115 | 0.000193572 | 3.727261182 | 1.427912044 |
| CNAG_12819 | - | unspecified product | 0.361671647 | 10.89717968 | 4.443821384 | 25.4331112 | 24.43325482 | 34.95071586 | 0.000489281 | 2.558368203 | 0.000194243 | 3.726389054 | 5.890410602 |
| CNAG_06516 | - | hypothetical protein | 219.3336535 | 232.5805625 | 247.2439515 | 127.7718435 | 158.4890026 | 168.8711155 | 0.000489315 | -0.638839006 | 0.000194316 | -3.72629402 | 0.642229568 |
| CNAG_05716 | - | hypothetical protein | 545.6663075 | 602.3768947 | 630.3276526 | 774.6779413 | 780.8031323 | 758.1284759 | 0.000490555 | 0.364688558 | 0.000194868 | 3.725577876 | 1.287603638 |
| CNAG_06522 | - | hypothetical protein | 188.5500129 | 251.1576443 | 275.1965908 | 367.0574171 | 340.8911682 | 361.6936302 | 0.000495627 | 0.566520642 | 0.000196943 | 3.722905354 | 1.480947648 |
| CNAG_05096 | - | histone deacetylase 1/2, variant 2 | 968.7745437 | 813.4113031 | 790.2190052 | 710.4294627 | 631.1682164 | 628.5358576 | 0.000496665 | -0.400695284 | 0.000197417 | -3.722299653 | 0.757493134 |
| CNAG_00371 | - | methylglutaconyl-CoA hydratase | 268.9642507 | 317.9254237 | 298.7101422 | 381.496002 | 451.4702436 | 416.1396041 | 0.000502324 | 0.484331222 | 0.000199727 | 3.719361387 | 1.398937221 |
| CNAG_13044 | - | unspecified product | 182.6824988 | 228.5483596 | 254.9409663 | 353.8989239 | 293.3238324 | 361.6077854 | 0.000505298 | 0.586417446 | 0.000200971 | 3.717792388 | 1.5015135 |
| CNAG_00708 | - | pre-mRNA-splicing factor slt11 | 2267.69012 | 2304.575458 | 2416.502928 | 1900.324964 | 2099.435671 | 1859.668443 | 0.000506585 | -0.26970024 | 0.000201545 | -3.717071851 | 0.829491878 |
| CNAG_05608 | - | hypothetical protein | 326.1069374 | 345.9624188 | 337.1038378 | 500.1755509 | 490.2180237 | 412.9613668 | 0.000507667 | 0.460782555 | 0.000202038 | 3.716455029 | 1.37628815 |
| CNAG_05826 | - | translation elongation factor Tu | 10016.87319 | 9759.356043 | 10168.5804 | 8359.366577 | 8952.450948 | 8891.342992 | 0.000507757 | -0.20799225 | 0.000202136 | -3.716332305 | 0.865741216 |
| CNAG_05501 | - | hypothetical protein, variant | 44.94778474 | 58.36765659 | 68.35669129 | 127.033585 | 101.1339767 | 105.2499303 | 0.000511326 | 0.955414188 | 0.000203619 | 3.714484094 | 1.939136267 |
| CNAG_02017 | - | hypothetical protein | 4.944772558 | 1.23757914 | 0.112052936 | 25.34698857 | 13.03734777 | 17.51323035 | 0.000512708 | 3.442529234 | 0.000204232 | 3.713723803 | 10.87187776 |
| CNAG_04687 | - | stearoyl-CoA desaturase (delta-9 desaturase) | 1724.82921 | 1491.290746 | 1524.057132 | 1937.350515 | 2067.777647 | 1896.088573 | 0.000516918 | 0.300186187 | 0.000205972 | 3.71157702 | 1.231303309 |
| CNAG_02358 | - | MRP family ATP-binding protein | 163.2184474 | 159.3563063 | 192.7878082 | 111.7055789 | 84.96120037 | 113.4065151 | 0.000517254 | -0.754012969 | 0.000206169 | -3.711335109 | 0.592951919 |
| CNAG_04039 | - | iron sulfur cluster assembly protein 2, mitochondrial | 982.0418313 | 1054.729991 | 1130.321299 | 823.1266362 | 847.7419517 | 874.9018709 | 0.000517594 | -0.330622839 | 0.000206494 | -3.710936378 | 0.79519311 |
| CNAG_00178 | - | DNA repair protein REV1 | 962.7222177 | 1382.360891 | 1253.154697 | 1481.977831 | 1615.780484 | 1692.769741 | 0.000517594 | 0.398881831 | 0.000206442 | 3.711000021 | 1.318485614 |
| CNAG_07404 | - | hypothetical protein | 25.10875621 | 31.47103873 | 44.8841897 | 84.17763293 | 71.01917396 | 73.01148135 | 0.000517594 | 1.174037153 | 0.000206389 | 3.711065654 | 2.256422382 |
| CNAG_05120 | - | Nicotinate phosphoribosyltransferase | 3186.612028 | 3151.414835 | 2911.409729 | 2520.613521 | 2714.402703 | 2633.39384 | 0.000518375 | -0.248588109 | 0.000206869 | -3.710477359 | 0.84171976 |
| CNAG_01820 | PYK1 | pyruvate kinase, variant | 11830.23217 | 10695.63805 | 10340.3264 | 9415.575478 | 9669.231054 | 9077.911374 | 0.000518933 | -0.238585674 | 0.000207155 | -3.710127508 | 0.847575814 |
| CNAG_02683 | - | hypothetical protein | 464.9190975 | 430.0149532 | 403.2528485 | 594.9589119 | 538.3003678 | 628.1076754 | 0.000520444 | 0.425185288 | 0.000207822 | 3.709313878 | 1.342744943 |
| CNAG_04522 | - | hypothetical protein | 616.4386329 | 639.9007415 | 667.4863716 | 518.1479744 | 515.6861419 | 433.2079106 | 0.000521522 | -0.406067785 | 0.000208316 | -3.708712448 | 0.754677525 |
| CNAG_04474 | - | monocarboxylic acid transporter | 576.6797673 | 607.537484 | 556.6390832 | 465.2046387 | 432.7942628 | 438.6712691 | 0.00052189 | -0.396342451 | 0.000208526 | -3.708456547 | 0.759782058 |
| CNAG_00828 | SIP401 | pathway-specific nitrogen regulator | 119.4557548 | 89.69657545 | 138.7088562 | 175.3421825 | 196.1124215 | 217.2016512 | 0.000526728 | 0.748670078 | 0.000210524 | 3.706041094 | 1.680243215 |
| CNAG_02870 | MET7 | folylpolyglutamate synthase | 766.3211491 | 712.3510717 | 890.513449 | 975.3154461 | 1156.181471 | 1006.819934 | 0.000529622 | 0.389718273 | 0.000211745 | 3.704575071 | 1.310137538 |
| CNAG_03598 | - | Ubiquitin | 1377.770782 | 1339.19127 | 1363.93134 | 1140.009018 | 1033.121556 | 1166.848812 | 0.000531299 | -0.303991438 | 0.000212481 | -3.703696106 | 0.810008281 |
| CNAG_01476 | - | hypothetical protein | 1435.028734 | 1482.454652 | 1423.666083 | 1136.776195 | 1121.124609 | 1290.036551 | 0.000533129 | -0.306001986 | 0.000213278 | -3.702746926 | 0.808880235 |
| CNAG_06428 | OTU1 | ubiquitin thioesterase OTU1 | 815.0706758 | 917.8636147 | 937.3026461 | 729.0553151 | 704.5473148 | 686.2349811 | 0.000533943 | -0.347799756 | 0.000213669 | -3.702282382 | 0.785781575 |
| CNAG_06599 | - | hypothetical protein | 503.5079358 | 415.0125965 | 535.4072551 | 622.7993374 | 680.8036124 | 683.6593287 | 0.000535114 | 0.433948124 | 0.000214203 | 3.701649283 | 1.350925507 |
| CNAG_12861 | - | unspecified product | 1.46971149 | 1.221351094 | 1.190451484 | 13.59476338 | 16.78652422 | 18.49100705 | 0.000536233 | 3.954521413 | 0.000214716 | 3.701041945 | 15.50349322 |
| CNAG_05380 | FZC44 | hypothetical protein | 618.6281108 | 555.9174308 | 616.3017812 | 480.7882079 | 462.1667408 | 416.5290963 | 0.000541147 | -0.413481523 | 0.00021675 | -3.69864957 | 0.750809327 |
| CNAG_03827 | - | hypothetical protein | 395.0162017 | 456.8887721 | 403.2348222 | 540.3314199 | 599.7695748 | 550.5192803 | 0.000543823 | 0.414858572 | 0.000217954 | 3.697242606 | 1.33316798 |
| CNAG_00493 | - | hypothetical protein | 151.9911914 | 80.05773278 | 140.88071 | 194.0362121 | 216.8246917 | 247.1208222 | 0.000543823 | 0.806955637 | 0.000217922 | 3.697280344 | 1.749515731 |
| CNAG_07477 | - | hypothetical protein, variant | 410.1340275 | 453.6521771 | 359.5558924 | 574.7666554 | 596.3324052 | 511.7338866 | 0.00054415 | 0.445247666 | 0.000218152 | 3.697012372 | 1.361547832 |
| CNAG_04009 | - | aminopeptidase 2 | 2081.944415 | 2012.59971 | 2049.672414 | 1758.768758 | 1779.374244 | 1650.95222 | 0.000551533 | -0.259289073 | 0.000221179 | -3.693511137 | 0.835499533 |
| CNAG_03415 | - | hypothetical protein | 349.3981915 | 609.4499533 | 520.2569479 | 326.4526518 | 363.8471353 | 286.7246653 | 0.000553252 | -0.613515878 | 0.000221936 | -3.692642323 | 0.653601918 |
| CNAG_06220 | - | allergen | 2100.25652 | 3214.063475 | 2258.977383 | 5313.729568 | 4760.073131 | 3803.35067 | 0.000555884 | 0.860732253 | 0.00022306 | 3.691358282 | 1.815959783 |
| CNAG_03309 | - | pre-mRNA-splicing factor CWC21 | 693.2274167 | 621.5878612 | 567.3397838 | 483.0422121 | 450.0112638 | 491.9218168 | 0.000556131 | -0.416973268 | 0.000223227 | -3.691167881 | 0.748994345 |
| CNAG_00474 | - | hypothetical protein | 98.64636343 | 138.0244308 | 137.6707473 | 194.9328579 | 199.5104912 | 216.1691381 | 0.000558343 | 0.693087143 | 0.000224183 | 3.690080858 | 1.61673939 |
| CNAG_02824 | - | hypothetical protein | 1662.728046 | 1675.331138 | 1595.384156 | 1372.685972 | 1396.371701 | 1378.923928 | 0.000563245 | -0.265630469 | 0.00022622 | -3.68777921 | 0.831835137 |
| CNAG_03573 | - | kynurenine 3-monooxygenase | 228.1200164 | 267.3082413 | 272.0267776 | 359.8888968 | 384.143391 | 351.7915029 | 0.00058029 | 0.497404444 | 0.000233136 | 3.680106806 | 1.411671534 |
| CNAG_05799 | CDA1 | Chitin deacetylase | 2288.817698 | 2077.369453 | 1686.234577 | 2619.236567 | 2530.879748 | 2608.73969 | 0.00058651 | 0.342972068 | 0.000235707 | 3.677309675 | 1.268366843 |
| CNAG_03104 | - | hypothetical protein | 796.1836479 | 740.1551179 | 825.3181517 | 630.9760579 | 658.0497936 | 528.6906067 | 0.000593176 | -0.394517123 | 0.000238531 | -3.674269639 | 0.760743959 |
| CNAG_12002 | - | unspecified product | 2.836272378 | 14.13045981 | 6.588745508 | 34.24596691 | 38.60237891 | 26.49946043 | 0.000593176 | 2.144186534 | 0.000238477 | 3.674327117 | 4.420429443 |
| CNAG_03925 | - | mitochondrial inner membrane protease subunit 2 | 637.4797087 | 615.1903394 | 763.39227 | 511.8153233 | 508.7187956 | 513.0408289 | 0.00059479 | -0.410964594 | 0.000239253 | -3.673497991 | 0.752120334 |
| CNAG_01552 | - | blocked early in transport 1 | 841.0763168 | 957.8910671 | 915.0893689 | 1135.751546 | 1092.947087 | 1191.030075 | 0.000599318 | 0.318628111 | 0.000241147 | 3.671483075 | 1.24714405 |
| CNAG_01324 | - | hypothetical protein | 243.1158235 | 229.6285738 | 221.9540322 | 328.7713733 | 350.5104128 | 317.3847326 | 0.000602946 | 0.506338522 | 0.0002426 | 3.669863052 | 1.420440619 |
| CNAG_06296 | - | hypothetical protein, variant | 22.5462502 | 18.52740778 | 16.19788861 | 50.45726999 | 50.42543497 | 50.74905764 | 0.000605856 | 1.405825435 | 0.000243926 | 3.668554655 | 2.649693403 |
| CNAG_03056 | - | hypothetical protein | 117.1099784 | 119.739225 | 113.1906102 | 183.2997309 | 194.4195551 | 181.8785037 | 0.00060962 | 0.661816811 | 0.000245516 | 3.666892844 | 1.582073703 |
| CNAG_01423 | - | hypothetical protein | 3695.763379 | 3561.943772 | 3474.481653 | 2977.49265 | 3103.392097 | 3160.692266 | 0.00060983 | -0.231038979 | 0.000245674 | -3.666727377 | 0.852021075 |
| CNAG_05001 | - | hypothetical protein | 704.9981005 | 621.6390362 | 660.0575841 | 508.7083715 | 545.0359013 | 475.35096 | 0.000616711 | -0.393406462 | 0.000248522 | -3.663778229 | 0.761329844 |
| CNAG_05016 | - | hypothetical protein | 20.49259553 | 29.32780843 | 44.87753181 | 66.51567477 | 62.24320349 | 101.382957 | 0.000617074 | 1.285803639 | 0.000248743 | 3.66355014 | 2.438178311 |
| CNAG_00798 | - | hypothetical protein | 37.81719942 | 37.91797783 | 38.56651855 | 98.37545142 | 93.73904826 | 57.78195826 | 0.000627627 | 1.131128322 | 0.000253074 | 3.659128876 | 2.190299752 |
| CNAG_01256 | - | hypothetical protein | 483.5131849 | 403.1034988 | 362.7639967 | 528.5674613 | 626.5572356 | 589.3089291 | 0.000628709 | 0.466036632 | 0.000253664 | 3.658531926 | 1.381309521 |
| CNAG_05097 | CKY1 | YjeF family protein | 1130.004672 | 1085.915152 | 896.9067738 | 863.7599862 | 824.4644265 | 699.6530964 | 0.000628709 | -0.398135998 | 0.000253647 | -3.658548894 | 0.75883809 |

| | | | | | | | | | | | | | |
|---|---|---|---|---|---|---|---|---|---|---|---|---|---|
| CNAG_05830 | - | cytoplasmic protein, variant | 2845.757876 | 2912.255905 | 3069.176289 | 2490.317864 | 2576.380745 | 2511.280934 | 0.000631822 | -0.235405008 | 0.000254997 | -3.657188202 | 0.849446501 |
| CNAG_04089 | - | coatomer zeta subunit | 2191.742414 | 2193.592413 | 2181.915996 | 1963.546839 | 1806.050416 | 1784.162165 | 0.000631837 | -0.256744317 | 0.000255081 | -3.657104375 | 0.836974563 |
| CNAG_01751 | - | hypothetical protein, variant | 375.133758 | 409.5033251 | 332.9000367 | 653.1674846 | 467.5328504 | 490.6173158 | 0.000633917 | 0.514225048 | 0.000255997 | 3.65618421 | 1.428226754 |
| CNAG_03092 | - | hypothetical protein | 428.1459936 | 337.2455657 | 423.2254882 | 265.7254649 | 300.0155287 | 277.7289277 | 0.00063457 | -0.513054232 | 0.000256339 | -3.655842445 | 0.700737383 |
| CNAG_03443 | - | hypothetical protein | 0.347644799 | 7.700371541 | 6.548576993 | 33.10416505 | 26.29972975 | 18.79385638 | 0.000637216 | 2.554349284 | 0.000257485 | 3.654697401 | 5.874024503 |
| CNAG_03595 | - | hypothetical protein | 3537.94148 | 3168.669432 | 2881.571437 | 2594.560879 | 2628.078071 | 2716.648573 | 0.000637682 | -0.287704134 | 0.000257752 | -3.65443244 | 0.819204682 |
| CNAG_06051 | GAL1 | galactokinase | 1041.898105 | 1221.822744 | 999.3920026 | 1472.158913 | 1334.480762 | 1337.662082 | 0.000639948 | 0.330998137 | 0.000258746 | 3.653444696 | 1.257883348 |
| CNAG_05995 | - | hypothetical protein | 921.0189197 | 889.8357591 | 753.9944219 | 650.3260416 | 676.0897097 | 678.4318225 | 0.000647153 | -0.370149278 | 0.000261738 | -3.650492465 | 0.773702436 |
| CNAG_07854 | - | hypothetical protein | 2.622434311 | 0.114690511 | 0.09896651 | 20.47260817 | 15.88243102 | 12.09711099 | 0.000647569 | 4.536999943 | 0.000261985 | 3.650250134 | 23.21523442 |
| CNAG_05377 | ITR3 | myo-inositol transporter, putative | 30.8126711 | 27.175487 | 37.45476194 | 75.48790112 | 71.99382032 | 65.29407112 | 0.000648109 | 1.150704087 | 0.000262283 | 3.649958333 | 2.220222228 |
| CNAG_03731 | - | hypothetical protein, variant | 806.7233798 | 830.5850464 | 758.2164831 | 619.9951663 | 665.7918132 | 620.7288039 | 0.000649985 | -0.345525025 | 0.000263121 | -3.649138539 | 0.787021512 |
| CNAG_04284 | - | hypothetical protein | 1300.90512 | 1593.361643 | 1459.906037 | 1159.384606 | 1134.077003 | 1231.244358 | 0.000650598 | -0.318744432 | 0.000263449 | -3.648818778 | 0.801767347 |
| CNAG_05687 | - | hypothetical protein | 928.6214275 | 1066.685346 | 923.6538183 | 1215.910344 | 1260.395309 | 1185.573051 | 0.000653791 | 0.312687275 | 0.000264822 | 3.647483078 | 1.242019027 |
| CNAG_05975 | - | ribonuclease Z | 3719.152924 | 3701.998189 | 3622.706594 | 2930.720894 | 3160.313149 | 3337.177065 | 0.000659042 | -0.243330313 | 0.000267029 | -3.645349133 | 0.844792942 |
| CNAG_03846 | - | vacuolar membrane protein | 840.7095344 | 881.2673334 | 954.3519357 | 757.9267224 | 702.8186292 | 656.2988571 | 0.000660862 | -0.354101204 | 0.000267847 | -3.644562211 | 0.782356898 |
| CNAG_01950 | - | hypothetical protein | 533.7074095 | 629.1080607 | 723.9064642 | 501.8151671 | 474.2313282 | 402.1491633 | 0.000662011 | -0.46806194 | 0.000268394 | -3.644037921 | 0.722935108 |
| CNAG_03763 | - | IQ domain-containing GTPase activating protein | 4156.061896 | 3717.234936 | 3541.797621 | 4482.303534 | 4700.248428 | 4611.509956 | 0.000664152 | 0.257443311 | 0.000269343 | 3.643129811 | 1.195358458 |
| CNAG_06823 | - | hypothetical protein | 2.743953737 | 2.34927525 | 8.636639942 | 23.17655325 | 24.58817882 | 23.03128344 | 0.000664742 | 2.528107551 | 0.000269663 | 3.642824072 | 5.768145477 |
| CNAG_01001 | - | hypothetical protein | 75.37731273 | 115.4271768 | 124.861928 | 214.997542 | 194.4870647 | 147.6487761 | 0.000669261 | 0.807350628 | 0.000271578 | 3.641002796 | 1.749994791 |
| CNAG_07780 | - | geranylgeranyl diphosphate synthase, type III | 694.4882862 | 602.2584624 | 655.7853155 | 493.1147656 | 519.9841462 | 497.5037232 | 0.000673201 | -0.386499053 | 0.000273259 | -3.639413865 | 0.764983718 |
| CNAG_12597 | - | unspecified product | 11.10397486 | 22.79487408 | 20.4301103 | 61.8468212 | 61.05831187 | 33.35702133 | 0.000676779 | 1.534080518 | 0.000274794 | 3.637971032 | 2.896037971 |
| CNAG_07369 | - | hypothetical protein | 8.789383522 | 13.18865751 | 29.92482028 | 56.17569998 | 42.4485786 | 57.26086539 | 0.000682924 | 1.61743871 | 0.000277373 | 3.635564182 | 3.068298219 |
| CNAG_04900 | - | WD40 repeat protein Ciao1 | 1744.773062 | 1718.584718 | 1709.596395 | 2121.074176 | 2179.016575 | 1959.412606 | 0.000687458 | 0.260060149 | 0.000279298 | 3.633780774 | 1.197528631 |
| CNAG_01107 | - | phosphatidylinositol glycan, class A | 1570.634064 | 1785.202148 | 1764.900439 | 1448.219804 | 1407.555054 | 1410.018672 | 0.000693679 | -0.278073711 | 0.000281995 | -3.631302025 | 0.824691412 |
| CNAG_00320 | - | anaphase-promoting complex subunit 3 | 592.3519851 | 650.8513228 | 664.4592865 | 809.657854 | 807.4992278 | 826.9266149 | 0.000693679 | 0.342578789 | 0.000281985 | 3.631310725 | 1.268021132 |
| CNAG_07571 | - | hypothetical protein | 425.3021465 | 409.5573444 | 440.5220474 | 599.715974 | 586.7731573 | 521.7330736 | 0.000701 | 0.405704748 | 0.000285057 | 3.628514574 | 1.324735878 |
| CNAG_02922 | - | hypothetical protein | 1918.492783 | 1982.390042 | 1855.603423 | 1653.703831 | 1585.235532 | 1638.687553 | 0.000707654 | -0.254108437 | 0.000287935 | -3.625919196 | 0.838505158 |
| CNAG_03960 | - | Gly-Xaa carboxypeptidase | 401.7504286 | 306.1733634 | 309.4236853 | 477.4330925 | 461.6287319 | 499.3445643 | 0.000707654 | 0.484095931 | 0.000287891 | 3.625958979 | 1.398709086 |
| CNAG_07686 | - | topoisomerase 1-associated factor 1 | 2303.740961 | 2226.975068 | 2019.864727 | 1843.573512 | 1822.456152 | 1838.52669 | 0.000712159 | -0.266081449 | 0.000289855 | -3.624201769 | 0.831575149 |
| CNAG_07042 | - | hypothetical protein | 0.152038788 | 0.073762007 | 0.063701092 | 11.56033398 | 9.772803451 | 6.598682598 | 0.000716655 | 5.542628608 | 0.000291773 | 3.62249715 | 46.61197082 |
| CNAG_00878 | - | hypothetical protein | 329.214535 | 370.493726 | 374.1944702 | 257.2601172 | 268.9459357 | 252.1972791 | 0.000719161 | -0.480612525 | 0.000292881 | -3.621516708 | 0.716673281 |
| CNAG_07653 | - | hypothetical protein | 23.97158111 | 30.41037979 | 48.06927602 | 76.86433265 | 65.70517951 | 94.89862393 | 0.000719455 | 1.203642026 | 0.000293088 | 3.621333439 | 2.303203723 |
| CNAG_12568 | - | unspecified product | 21.8180942 | 63.73834208 | 76.85704909 | 114.9694948 | 128.1903897 | 120.6703358 | 0.000720841 | 1.164846202 | 0.000293741 | 3.620757953 | 2.242093126 |
| CNAG_03038 | - | hypothetical protein | 1705.588613 | 2723.832547 | 2532.937593 | 3730.069089 | 3728.29926 | 3783.159058 | 0.00072324 | 0.678317124 | 0.000294807 | 3.619820842 | 1.600271979 |
| CNAG_05443 | - | hypothetical protein | 120.8139933 | 196.0902118 | 122.8496707 | 249.6412167 | 223.5891944 | 247.2888527 | 0.000723606 | 0.70220194 | 0.000295045 | 3.619612228 | 1.626986117 |
| CNAG_04094 | - | hypothetical protein | 236.2278387 | 261.9170448 | 254.982443 | 362.3098691 | 364.2601973 | 334.0586918 | 0.000724803 | 0.482182701 | 0.000295621 | 3.619107246 | 1.396855417 |
| CNAG_05732 | - | hypothetical protein | 286.99721 | 298.28903 | 272.9418438 | 203.5690082 | 203.3510234 | 195.5885673 | 0.000725827 | -0.525286893 | 0.000296127 | -3.618664442 | 0.694820925 |
| CNAG_01814 | - | hypothetical protein | 254.8286697 | 262.994767 | 237.9535904 | 370.1376811 | 331.3299725 | 365.0389633 | 0.000729223 | 0.486593725 | 0.000297602 | 3.617378457 | 1.401132823 |
| CNAG_06781 | - | hypothetical protein | 1026.085679 | 847.9353412 | 930.9449644 | 730.5291105 | 761.5121233 | 721.7569169 | 0.000729787 | -0.357730789 | 0.000297921 | -3.617100783 | 0.78039109 |
| CNAG_00713 | - | V-type H -transporting ATPase 54 kDa subunit | 2008.235979 | 1742.194124 | 1813.996977 | 1533.006365 | 1559.391021 | 1523.25886 | 0.000730768 | -0.285393462 | 0.000298411 | -3.616675358 | 0.820517801 |
| CNAG_06984 | - | hypothetical protein | 927.0136325 | 929.7377497 | 891.5115288 | 752.3912901 | 721.7827984 | 753.904293 | 0.000735733 | -0.318527342 | 0.000300528 | -3.614844108 | 0.801888002 |
| CNAG_04381 | - | transcription initiation factor TFIIH subunit 2 | 803.1161151 | 729.3692645 | 741.1285774 | 634.1544309 | 569.0733406 | 584.0960009 | 0.00073719 | -0.362587107 | 0.000301214 | -3.614253989 | 0.777768598 |
| CNAG_02169 | - | hypothetical protein | 36.86035964 | 60.50652262 | 70.47477286 | 111.903072 | 124.7720421 | 97.55803423 | 0.000740544 | 0.985542821 | 0.000302674 | 3.613000337 | 1.980058179 |
| CNAG_04859 | - | rho family protein | 375.3310098 | 483.8548459 | 514.0714719 | 641.6835848 | 585.7686482 | 665.8624555 | 0.000741504 | 0.448547106 | 0.000303157 | 3.612587158 | 1.364665252 |
| CNAG_02701 | - | hypothetical protein | 2459.588073 | 1264.214502 | 1144.652174 | 3811.888453 | 4688.83221 | 4617.647878 | 0.000741638 | 1.411859181 | 0.000303303 | 3.61246274 | 2.660798354 |
| CNAG_00570 | PKR1 | cAMP-dependent protein kinase regulator | 3531.18494 | 3489.768332 | 3610.95621 | 3017.999869 | 3156.899131 | 3059.694662 | 0.000749524 | -0.218817759 | 0.000306632 | -3.609631714 | 0.859269291 |
| CNAG_06297 | - | hypothetical protein | 1787.760918 | 196.453972 | 426.6099987 | 101.4603747 | 85.45535212 | 65.03310166 | 0.000759895 | -3.286913862 | 0.000310955 | -3.605979724 | 0.102456693 |
| CNAG_04156 | - | Homoserine kinase | 1178.157813 | 1263.719234 | 1180.521187 | 936.7298399 | 1033.256126 | 1007.014767 | 0.000760776 | -0.297992122 | 0.000311408 | -3.605620255 | 0.81338364 |
| CNAG_12869 | - | unspecified product | 75.30177427 | 66.6253238 | 72.30986399 | 24.9437372 | 20.18356289 | 46.69527523 | 0.000761219 | -1.2577997 | 0.000311682 | -3.605391798 | 0.418181255 |
| CNAG_02591 | - | hypothetical protein | 143.8429099 | 139.1040955 | 90.89419056 | 230.3826065 | 205.5779814 | 186.3960089 | 0.000765021 | 0.724308525 | 0.000313333 | 3.604020185 | 1.6521086 |
| CNAG_12555 | - | unspecified product | 61.89560549 | 38.65573235 | 34.07709491 | 15.75021679 | 18.67629978 | 12.54218918 | 0.000769212 | -1.580718822 | 0.000315143 | -3.602523497 | 0.334315275 |
| CNAG_07459 | - | hypothetical protein | 77.58030486 | 97.11434777 | 101.4198482 | 157.7306418 | 141.6164411 | 169.5043952 | 0.000769985 | 0.749578704 | 0.000315554 | 3.602184758 | 1.681301784 |
| CNAG_02441 | - | hypothetical protein | 10.84808086 | 9.868937304 | 2.330552174 | 38.52333542 | 27.8946721 | 26.4595059 | 0.000770147 | 2.100790477 | 0.000315714 | 3.602052976 | 4.289443465 |
| CNAG_12542 | - | unspecified product | 0.145855173 | 0.070794075 | 0.061141582 | 8.102629683 | 5.18994985 | 13.97250441 | 0.000783468 | 5.544177385 | 0.000321271 | 3.597515891 | 46.66203706 |
| CNAG_12025 | - | unspecified product | 7.263348044 | 0.145383486 | 0.125374863 | 31.00767253 | 13.8377082 | 21.88403459 | 0.000786612 | 3.195045667 | 0.000322752 | 3.596318872 | 9.158083169 |
| CNAG_02505 | - | cardiolipin-specific phospholipase | 118.3357565 | 149.8212047 | 105.774168 | 202.062102 | 198.6615782 | 197.3998062 | 0.000786612 | 0.668285786 | 0.000322697 | 3.596363194 | 1.589183573 |
| CNAG_06590 | - | hypothetical protein, variant | 61.27012223 | 92.74409623 | 71.62554619 | 117.7161861 | 157.5766883 | 132.984618 | 0.000793977 | 0.843922236 | 0.000325871 | 3.593815231 | 1.794923348 |
| CNAG_12840 | - | unspecified product | 37.95979611 | 57.25278442 | 58.77336041 | 94.60397834 | 99.53661208 | 106.2215895 | 0.000794783 | 0.961980662 | 0.000326299 | 3.593473733 | 1.947982426 |
| CNAG_01093 | - | hypothetical protein | 3354.591599 | 3398.042622 | 4424.29007 | 2410.07939 | 1299.239713 | 1633.460139 | 0.000797422 | -1.080005048 | 0.00032748 | -3.592532594 | 0.473027168 |
| CNAG_06153 | - | 26S protease regulatory subunit 6A-B | 6031.527046 | 5602.739564 | 5789.834574 | 6574.291751 | 6888.527756 | 6979.47384 | 0.000805527 | 0.214968552 | 0.000330906 | 3.589820035 | 1.160678614 |
| CNAG_05807 | - | hypothetical protein | 295.6802642 | 282.4237865 | 290.1794367 | 398.789298 | 440.2593794 | 370.7298659 | 0.000809437 | 0.462893513 | 0.000332611 | 3.588480015 | 1.378303415 |
| CNAG_06380 | - | transcription initiation factor TFIIA large subunit | 1067.570238 | 1147.531944 | 1128.374349 | 1341.140795 | 1486.446945 | 1313.272344 | 0.000809524 | 0.293759291 | 0.000332746 | 3.58837459 | 1.225830317 |
| CNAG_03011 | - | glycerate-and formate-dehydrogenase | 103.1533452 | 125.0675809 | 94.0354319 | 175.5723234 | 164.0660315 | 189.5113601 | 0.000812877 | 0.699296328 | 0.000334223 | 3.587219165 | 1.623712637 |
| CNAG_00712 | - | hypothetical protein | 552.2344086 | 604.3093306 | 573.6798869 | 448.0152739 | 453.5473211 | 443.1025477 | 0.000816084 | -0.378312062 | 0.000335642 | -3.586114225 | 0.769337181 |
| CNAG_01743 | - | hypothetical protein | 401.9273241 | 401.9841134 | 374.4478377 | 490.4298999 | 601.6030769 | 508.3591566 | 0.000823265 | 0.427848167 | 0.000338696 | 3.583750306 | 1.345225626 |
| CNAG_00829 | - | E3 ubiquitin ligase complex SCF subunit sconC | 3660.797492 | 3622.287902 | 3709.073494 | 3279.608592 | 3036.917403 | 3197.342467 | 0.000831438 | -0.223726169 | 0.00034216 | -3.581093003 | 0.856350809 |
| CNAG_04188 | - | O-methyltransferase | 1342.660428 | 1268.071145 | 1224.256337 | 1111.293353 | 1059.100479 | 941.619966 | 0.000832254 | -0.316998616 | 0.000342597 | -3.580759371 | 0.802738159 |
| CNAG_01514 | - | hypothetical protein | 0.20211901 | 1.199526586 | 0.084333653 | 18.16909629 | 14.19161687 | 6.687975839 | 0.000832374 | 5.128017679 | 0.000342748 | 3.580643997 | 34.96931616 |
| CNAG_07591 | - | mitochondrial metalloendopeptidase OMA1 | 1175.665092 | 1083.847561 | 1074.948396 | 929.868828 | 893.4500239 | 902.68218 | 0.000835387 | -0.306157699 | 0.000344091 | -3.579622382 | 0.808792935 |
| CNAG_01873 | - | hypothetical protein | 0.273935613 | 1.239650376 | 4.386833327 | 17.63847761 | 30.33363674 | 8.955910418 | 0.000838042 | 3.442512693 | 0.000345287 | 3.578715786 | 10.8717531 |
| CNAG_01664 | - | CMGC/CDK/CDC2 protein kinase | 1157.30604 | 955.8343595 | 1024.956792 | 1454.318264 | 1318.124688 | 1249.999162 | 0.000841746 | 0.342581238 | 0.000347019 | 3.577407694 | 1.268023284 |
| CNAG_12538 | - | unspecified product | 29.62175516 | 20.74303355 | 35.31750458 | 82.93339006 | 58.86072033 | 60.86246409 | 0.000841746 | 1.235150873 | 0.000346944 | 3.577463848 | 2.354059642 |
| CNAG_00162 | AOX1 | alternative oxidase, mitochondrial | 5957.090751 | 1873.788755 | 2752.375288 | 1029.399722 | 992.1379705 | 978.5926112 | 0.000845912 | -1.839550805 | 0.000348839 | -3.576039182 | 0.279408767 |
| CNAG_02886 | - | 20S proteasome subunit beta 4 | 1211.172902 | 1157.289176 | 1339.491406 | 1523.320045 | 1555.357599 | 1510.837001 | 0.000845982 | 0.292001585 | 0.000348971 | 3.575940206 | 1.224337737 |
| CNAG_03842 | - | hypothetical protein | 10.81115756 | 4.525803733 | 6.569207674 | 27.82753671 | 30.6669272 | 27.48529943 | 0.000848806 | 2.052944606 | 0.00035024 | 3.574991088 | 4.149520416 |
| CNAG_01507 | - | protein CGI121 | 631.7322234 | 712.0731348 | 745.3204598 | 578.1141048 | 539.7874275 | 501.9944041 | 0.000852843 | -0.382852089 | 0.00035201 | -3.573671992 | 0.766919953 |
| CNAG_03633 | - | hypothetical protein | 374.0039182 | 401.9904084 | 413.8613995 | 473.2051932 | 555.7309996 | 589.2222657 | 0.000856055 | 0.429754971 | 0.00035344 | 3.572610467 | 1.34700478 |
| CNAG_06145 | - | hypothetical protein, variant | 615.5844855 | 626.0599342 | 560.0081533 | 809.0177843 | 759.1848067 | 750.3780851 | 0.000857653 | 0.349036119 | 0.000354205 | 3.572044951 | 1.273709363 |
| CNAG_07919 | - | hypothetical protein | 207.2144822 | 255.496193 | 302.9087839 | 430.365071 | 325.2006416 | 389.4391135 | 0.000857793 | 0.567937031 | 0.000354367 | 3.571924763 | 1.482402306 |
| CNAG_01736 | - | DASH complex subunit dad4 | 686.5669195 | 739.0238125 | 775.1929156 | 538.9755268 | 635.6619809 | 539.7252891 | 0.000858163 | -0.377111297 | 0.00035473 | -3.571657111 | 0.769977772 |
| CNAG_00751 | - | hypothetical protein | 9.638330373 | 3.442070397 | 5.499594812 | 33.00960975 | 23.55156979 | 23.12401139 | 0.000858163 | 2.198837113 | 0.000354646 | 3.571719226 | 4.591091272 |
| CNAG_13096 | - | unspecified product | 53.18043305 | 57.85914463 | 43.63367014 | 25.26738157 | 17.70641727 | 18.07478773 | 0.000867704 | -1.365026688 | 0.00035878 | -3.568683488 | 0.388227256 |
| CNAG_00344 | - | hypothetical protein | 700.9025181 | 796.2544862 | 658.1325119 | 851.50263 | 962.8372644 | 975.638686 | 0.000870407 | 0.357623501 | 0.000360004 | 3.567790878 | 1.281313497 |
| CNAG_05359 | - | hypothetical protein | 651.7210219 | 909.1192039 | 811.4667577 | 596.6321083 | 611.3865268 | 594.0967469 | 0.000870619 | -0.410638256 | 0.000360197 | -3.567649838 | 0.752290483 |
| CNAG_01222 | - | hypothetical protein | 667.6356111 | 586.0893392 | 626.9971926 | 459.5922236 | 496.6849528 | 500.7952724 | 0.000872116 | -0.384714735 | 0.000360923 | -3.567122205 | 0.765930431 |
| CNAG_06740 | - | septation protein imp2 | 1003.314683 | 1017.187 | 1049.450429 | 1292.222543 | 1185.191748 | 1337.570233 | 0.000882428 | 0.298481112 | 0.000365406 | 3.563884724 | 1.229848931 |
| CNAG_01825 | - | hypothetical protein | 362.7394145 | 315.6141254 | 319.8632989 | 243.2944782 | 251.7117455 | 214.5098748 | 0.000882428 | -0.508287186 | 0.000365369 | -3.563911416 | 0.703056634 |

| | | | | | | | | | | | | | |
|---|---|---|---|---|---|---|---|---|---|---|---|---|---|
| CNAG_02893 | - | hypothetical protein, variant | 296.889395 | 523.2855084 | 494.5921743 | 314.1518003 | 304.2784539 | 224.6202673 | 0.00088842 | -0.654481537 | 0.000367996 | -3.562031398 | 0.63530376 |
| CNAG_06596 | - | hypothetical protein | 164.8487797 | 140.2408247 | 141.9832423 | 224.5290489 | 225.3702565 | 230.6911704 | 0.000889587 | 0.5939289 | 0.000368588 | 3.56160946 | 1.509351582 |
| CNAG_03045 | - | hypothetical protein | 1207.147091 | 1065.547719 | 1080.279377 | 908.0254894 | 927.9929982 | 890.4856886 | 0.000896247 | -0.31452827 | 0.000371457 | -3.559573937 | 0.804113876 |
| CNAG_01947 | - | 2,4-dienoyl-CoA reductase | 176.4068923 | 141.3041966 | 93.04795998 | 204.1149646 | 247.125815 | 241.701481 | 0.000899053 | 0.739353246 | 0.00037273 | 3.558675239 | 1.669427274 |
| CNAG_06731 | - | hypothetical protein | 0.294107249 | 3.400168611 | 5.45443709 | 11.61923781 | 25.65862097 | 25.05868049 | 0.000905872 | 2.916832927 | 0.000375667 | 3.556612655 | 7.551864748 |
| CNAG_07426 | - | proteasome maturation protein | 1364.090432 | 1291.948651 | 1393.903681 | 1647.069275 | 1644.166986 | 1627.40797 | 0.000918461 | 0.265336787 | 0.000381 | 3.552906159 | 1.201916595 |
| CNAG_02137 | - | hypothetical protein | 999.784243 | 1048.396943 | 958.838847 | 1204.158699 | 1214.587802 | 1302.05018 | 0.000928022 | 0.293046312 | 0.00038508 | 3.550103447 | 1.225224662 |
| CNAG_12173 | - | unspecified product | 17.05445737 | 53.92111819 | 31.13707496 | 96.09778657 | 85.0175604 | 64.314886 | 0.000936762 | 1.26678615 | 0.000388821 | 3.547557472 | 2.406249348 |
| CNAG_04927 | YFH701 | hypothetical protein | 58.51200369 | 29.37259597 | 33.28060943 | 92.61896791 | 75.21602771 | 95.05798634 | 0.000937429 | 1.109470643 | 0.000389441 | 3.547137606 | 2.157664632 |
| CNAG_06867 | - | hypothetical protein | 255.0397128 | 358.8100957 | 300.856213 | 408.8193622 | 436.6930588 | 448.280967 | 0.000937429 | 0.486997084 | 0.000389403 | 3.54716313 | 1.401524617 |
| CNAG_00147 | - | pre-mRNA-processing-splicing factor 8 | 27485.14151 | 25474.76783 | 23891.76107 | 21419.85997 | 22655.08299 | 22589.62614 | 0.000937429 | -0.220686056 | 0.000389215 | -3.547290772 | 0.858157253 |
| CNAG_03664 | NIC1 | high-affinity nickel-transporter | 1679.118742 | 1618.283357 | 1812.869708 | 1453.675807 | 1434.317144 | 1385.6111 | 0.000937999 | -0.273677265 | 0.000389793 | -3.54690001 | 0.827208396 |
| CNAG_05304 | - | hypothetical protein | 39.16807728 | 69.06593093 | 59.86904113 | 135.04505 | 99.40019178 | 96.45390288 | 0.000939447 | 0.969234459 | 0.000390509 | 3.546416202 | 1.957801446 |
| CNAG_03893 | SDP102 | MAP kinase phosphatase | 238.4574476 | 239.2970519 | 203.8561176 | 309.9705497 | 305.4558851 | 379.3005791 | 0.000941229 | 0.531462358 | 0.000391364 | 3.545839502 | 1.445393547 |
| CNAG_06259 | - | MFS transporter, SP family, general alpha glucoside:H symporter | 44.94372409 | 80.83683946 | 42.8945665 | 117.4843 | 97.6437413 | 118.4140423 | 0.00094265 | 0.987668703 | 0.000392071 | 3.545364341 | 1.982978044 |
| CNAG_06290 | HXT2 | Low-affinity glucose transporter HXT2 | 313.2888334 | 221.9013031 | 216.4635624 | 174.0166055 | 164.4969693 | 144.6082987 | 0.000942758 | -0.654478217 | 0.000392231 | -3.54525678 | 0.635305222 |
| CNAG_04147 | - | ATP-dependent rRNA helicase RRP3 | 1530.77841 | 1438.356721 | 1616.644364 | 1230.202543 | 1254.879148 | 1317.831746 | 0.000946952 | -0.285780443 | 0.000394091 | -3.54400945 | 0.820297739 |
| CNAG_05353 | - | nuclear pore complex protein Nup133 | 3340.805449 | 3301.201448 | 3225.993682 | 2834.347409 | 2790.250951 | 2937.542505 | 0.000948821 | -0.219879665 | 0.000394985 | -3.543412169 | 0.858637052 |
| CNAG_06806 | ETF1alpha | electron transfer flavoprotein alpha subunit | 3127.01275 | 2961.804495 | 2795.165834 | 2655.442817 | 2433.988311 | 2449.136819 | 0.000955526 | -0.252405712 | 0.000397893 | -3.54147755 | 0.839495378 |
| CNAG_06002 | - | tRNA (5-methylaminomethyl-2-thiouridylate)- methyltransferase | 972.5399408 | 884.5594177 | 1029.004825 | 729.6938775 | 734.6971792 | 827.1231719 | 0.000956125 | -0.348838724 | 0.000398259 | -3.541234941 | 0.785215891 |
| CNAG_01861 | - | 26S proteasome non-ATPase regulatory subunit 10 | 1228.707578 | 1239.149242 | 1369.355699 | 1562.22822 | 1576.899716 | 1555.242004 | 0.000962852 | 0.275560481 | 0.000401179 | 3.539307221 | 1.210464258 |
| CNAG_05535 | FHL1 | Pre-rRNA-processing protein FHL1, putative | 1032.019904 | 891.0345487 | 1015.168373 | 755.4010503 | 830.5587979 | 762.8301862 | 0.000964407 | -0.34005696 | 0.000401944 | -3.538803961 | 0.79001012 |
| CNAG_07377 | TRA1 | transformation/transcription domain-associated protein | 14110.92328 | 13336.41783 | 12840.09012 | 11212.57508 | 11786.80057 | 12165.18927 | 0.000964449 | -0.211639317 | 0.000402079 | -3.538715138 | 0.863555426 |
| CNAG_00679 | - | hypothetical protein | 34.35616376 | 49.66340275 | 32.20330614 | 82.36410528 | 80.58135291 | 78.56472675 | 0.00096575 | 1.047594217 | 0.00040274 | 3.538282049 | 2.067079988 |
| CNAG_00088 | - | prohibitin PHB1 | 5339.96995 | 4890.464203 | 5389.784675 | 4246.847835 | 4572.349965 | 4601.680028 | 0.000966464 | -0.23467528 | 0.000403156 | -3.538009519 | 0.849876267 |
| CNAG_02149 | - | hypothetical protein | 52.2699769 | 53.72253385 | 70.06275722 | 21.90654067 | 23.77603172 | 30.1748762 | 0.000968258 | -1.257042692 | 0.000404022 | -3.537442436 | 0.41840074 |
| CNAG_00999 | - | hypothetical protein | 1232.832407 | 1101.091356 | 1096.278895 | 880.6964403 | 884.7893576 | 998.0792871 | 0.000980473 | -0.327587124 | 0.000409239 | -3.534053072 | 0.796868114 |
| CNAG_05582 | - | hypothetical protein | 33.42777591 | 56.25494471 | 89.59785322 | 118.0614861 | 129.0027383 | 119.6167436 | 0.000987311 | 1.025238437 | 0.000412213 | 3.532138447 | 2.035295734 |
| CNAG_04312 | MAN1 | mannose-6-phosphate isomerase | 1473.426665 | 1351.049238 | 1307.451055 | 1182.855768 | 1123.765443 | 1103.638029 | 0.000988034 | -0.293091218 | 0.000412636 | -3.531867496 | 0.816151442 |
| CNAG_06350 | - | hypothetical protein | 1187.605541 | 1368.220704 | 1342.577914 | 990.4080777 | 1050.444221 | 1133.524668 | 0.000988342 | -0.311158818 | 0.000412885 | -3.531707773 | 0.805994099 |
| CNAG_02659 | - | hypothetical protein | 0.22541254 | 0.108773863 | 2.245624187 | 12.82895952 | 12.27460484 | 20.56531945 | 0.000992168 | 4.444066804 | 0.000414605 | 3.53060852 | 21.76694154 |
| CNAG_12372 | - | unspecified product | 82.17386854 | 73.03303711 | 57.54109994 | 36.09930529 | 32.42164663 | 31.37682645 | 0.000993242 | -1.113965235 | 0.000415175 | -3.530245079 | 0.46202242 |
| CNAG_04925 | - | nuclear protein | 1553.20623 | 1466.469067 | 1419.539078 | 1785.583259 | 1782.163115 | 1802.797749 | 0.0009952 | 0.259347742 | 0.000416115 | 3.529646652 | 1.196937434 |
| CNAG_00836 | - | 2-hydroxyacid dehydrogenase | 441.1511537 | 431.9492963 | 420.1162975 | 306.2444244 | 336.2474467 | 330.9628936 | 0.000996831 | -0.425910182 | 0.000416919 | -3.529136126 | 0.744368968 |
| CNAG_05079 | - | hypothetical protein | 7.448528706 | 10.93815453 | 8.697789929 | 27.65911653 | 32.35228598 | 32.94595256 | 0.001002783 | 1.849611883 | 0.000419531 | 3.527483057 | 3.604032156 |
| CNAG_03621 | CPA2 | Cyclophilin A | 17923.74977 | 17140.87218 | 19394.43113 | 16119.03837 | 15814.95158 | 15584.53939 | 0.001003317 | -0.212289451 | 0.000419876 | -3.527264973 | 0.863166363 |
| CNAG_05228 | - | U3 small nucleolar ribonucleoprotein Lcp5 | 704.1572784 | 727.2432111 | 989.3919378 | 632.5297398 | 629.5275939 | 513.1723868 | 0.00100783 | -0.46378569 | 0.000421888 | -3.525999267 | 0.725081118 |
| CNAG_01588 | - | plasma membrane proteolipid 3 | 4278.400774 | 4234.206695 | 3629.161773 | 3647.787437 | 3291.36893 | 3218.508668 | 0.001017759 | -0.272636724 | 0.000426169 | -3.52332471 | 0.827805234 |
| CNAG_02115 | NHP6B02 | Non-histone chromosomal protein 6, putative | 740.2148633 | 757.3507374 | 754.979006 | 612.2946015 | 611.4179178 | 585.204646 | 0.001018037 | -0.33137322 | 0.00042641 | -3.52317507 | 0.794779618 |
| CNAG_12878 | - | unspecified product | 21.48851719 | 19.62941289 | 24.68881136 | 73.74066973 | 32.59832991 | 67.19847243 | 0.001019664 | 1.411549471 | 0.000427215 | 3.522674629 | 2.660227209 |
| CNAG_04389 | - | hypothetical protein | 469.2985232 | 467.5450634 | 525.6199181 | 373.2926957 | 364.6608405 | 374.2639413 | 0.001025051 | -0.410443623 | 0.000429598 | -3.521200518 | 0.752391981 |
| CNAG_05274 | - | ste/ste20/ysk protein kinase | 1112.852562 | 1189.424433 | 1321.201599 | 929.7223838 | 984.0385791 | 1023.645593 | 0.001026839 | -0.318053539 | 0.000430473 | -3.520660994 | 0.802151398 |
| CNAG_12891 | - | unspecified product | 16.80430148 | 20.64100972 | 17.24196827 | 43.24442152 | 48.73081471 | 49.60223471 | 0.001031233 | 1.386810272 | 0.00043244 | 3.519451265 | 2.614998777 |
| CNAG_04184 | FZC47 | hypothetical protein | 644.7886331 | 606.7193064 | 659.1310054 | 783.1173277 | 849.8476212 | 803.6475058 | 0.00103808 | 0.335493224 | 0.000435439 | 3.517618123 | 1.261808718 |
| CNAG_01452 | MAT3 | mat3 pheromone repeat protein | 46.02431537 | 57.25050634 | 52.41147667 | 108.9916717 | 90.78339901 | 96.34663812 | 0.001045684 | 0.909113585 | 0.000438756 | 3.515603442 | 1.877891337 |
| CNAG_03651 | - | syntaxin 7 | 1181.501221 | 1074.161058 | 1102.66059 | 954.0391342 | 839.9000553 | 930.411193 | 0.001047013 | -0.317429469 | 0.000439441 | -3.515188897 | 0.802498461 |
| CNAG_03949 | PHO13 | 4-nitrophenyl phosphatase | 1827.424262 | 1770.19773 | 1960.045858 | 1480.789534 | 1631.897842 | 1548.780493 | 0.001056113 | -0.269870919 | 0.00044339 | -3.512812944 | 0.82939375 |
| CNAG_07323 | - | cactin | 2301.529831 | 2325.039937 | 2284.305051 | 1956.415393 | 1951.844978 | 2040.536806 | 0.001063953 | -0.231654624 | 0.000446941 | -3.510692618 | 0.851657567 |
| CNAG_03661 | - | hypothetical protein, variant | 972.6889343 | 970.7457392 | 1196.379555 | 815.3553381 | 808.0282778 | 859.3614811 | 0.001063953 | -0.353781945 | 0.000446878 | -3.510730053 | 0.782530048 |
| CNAG_00615 | - | hypothetical protein | 22.55639622 | 15.32948063 | 19.37354347 | 32.54736369 | 59.28032522 | 63.79515225 | 0.001070364 | 1.442948388 | 0.000449765 | 3.509017836 | 2.71875922 |
| CNAG_07964 | - | hypothetical protein | 1112.933254 | 1038.722879 | 833.0747745 | 1282.081434 | 1430.456381 | 1196.717891 | 0.001073769 | 0.374125243 | 0.000451327 | 3.508095681 | 1.296053473 |
| CNAG_01734 | - | pre-mRNA-splicing factor ATP-dependent RNA helicase DHX16 | 5535.049603 | 5514.258121 | 5520.993041 | 4712.327482 | 4992.438726 | 4903.651431 | 0.001082943 | -0.197139419 | 0.000455315 | -3.505754618 | 0.872278407 |
| CNAG_01252 | - | thiosulfate/3-mercaptopyruvate sulfurtransferase | 103.009402 | 72.44428568 | 98.23224161 | 173.5732465 | 145.978306 | 148.5775463 | 0.00108744 | 0.7550108 | 0.000457338 | 3.504574169 | 1.687644226 |
| CNAG_02834 | - | sterol 3-beta-glucosyltransferase | 738.2251614 | 786.57878 | 706.0971294 | 938.0805715 | 904.0572725 | 961.2493039 | 0.001089349 | 0.315104334 | 0.000458407 | 3.503952468 | 1.244101622 |
| CNAG_03663 | - | L-lactate dehydrogenase | 2090.105496 | 1987.826448 | 2061.396962 | 1768.124976 | 1812.178633 | 1578.82828 | 0.001089349 | -0.26660954 | 0.000458316 | -3.504005225 | 0.831270811 |
| CNAG_04466 | - | hypothetical protein | 414.9516646 | 520.4517376 | 491.711547 | 616.6300347 | 681.7251408 | 601.6257694 | 0.00108986 | 0.398903127 | 0.000458755 | 3.503750245 | 1.318505077 |
| CNAG_06446 | - | mitochondrial splicing suppressor | 759.8451059 | 657.1955863 | 677.1440161 | 563.269489 | 569.1707648 | 496.4553407 | 0.001093999 | -0.378408326 | 0.000460631 | -3.502663516 | 0.769285848 |
| CNAG_01341 | ISP3 | mannose-6-phosphate isomerase, variant | 695.0845149 | 801.6423387 | 640.0192919 | 1095.756803 | 910.9843 | 839.2798505 | 0.001106646 | 0.398560469 | 0.000466091 | 3.499522945 | 1.318191954 |
| CNAG_05692 | LCB3 | sphingosine-1-phosphate phosphatase | 1845.087508 | 2128.906755 | 2158.405291 | 1610.743488 | 1729.320512 | 1769.665512 | 0.001109357 | -0.278105502 | 0.000467369 | -3.498793315 | 0.824673239 |
| CNAG_02951 | - | hypothetical protein | 1098.719509 | 1070.926697 | 1154.877929 | 955.6101594 | 864.9515298 | 903.7894463 | 0.001110623 | -0.302719488 | 0.000468038 | -3.498411859 | 0.810722738 |
| CNAG_12907 | - | unspecified product | 0.140781169 | 0.068356703 | 0.059039394 | 9.880513031 | 6.120516952 | 9.746541068 | 0.001114888 | 5.428056156 | 0.000469971 | 3.497312479 | 43.05342648 |
| CNAG_04022 | - | translation initiation factor | 712.4420462 | 690.7073768 | 563.2509232 | 786.9118134 | 889.5240049 | 912.3396084 | 0.001121605 | 0.381604032 | 0.00047294 | 3.495632535 | 1.302789532 |
| CNAG_06149 | - | hypothetical protein, variant | 207.1835373 | 287.7277755 | 264.5710956 | 366.1830221 | 367.6970247 | 352.8867813 | 0.001127637 | 0.503240317 | 0.000475621 | 3.494123508 | 1.417393479 |
| CNAG_03158 | CMT1 | alpha-1,3-mannosyltransferase CMT1 | 959.8880723 | 1161.304332 | 1031.209301 | 849.6514454 | 818.3785891 | 876.0080539 | 0.001128697 | -0.32359657 | 0.000476205 | -3.49379559 | 0.799075336 |
| CNAG_01313 | - | hypothetical protein | 304.9564065 | 273.810644 | 272.071132 | 401.188097 | 432.4983257 | 354.1072654 | 0.00113825 | 0.467666557 | 0.000480375 | 3.491467449 | 1.382870975 |
| CNAG_00356 | - | survival factor 1 | 4800.484267 | 4534.893602 | 4716.878764 | 4203.435576 | 4028.038571 | 4042.190286 | 0.001138456 | -0.210580804 | 0.000480601 | -3.491341716 | 0.864189254 |
| CNAG_04366 | - | hypothetical protein | 827.7538209 | 827.3842105 | 810.4477961 | 615.257664 | 646.7451673 | 712.7715169 | 0.001150456 | -0.335734991 | 0.000485807 | -3.488462206 | 0.79238035 |
| CNAG_06230 | DRP1 | Putative transporter | 202.1489411 | 184.3549054 | 127.1437537 | 257.9706845 | 275.4846939 | 264.0131597 | 0.001150466 | 0.618521714 | 0.000485952 | 3.488382576 | 1.535301198 |
| CNAG_12037 | - | unspecified product | 34.14246873 | 22.86368783 | 20.4835381 | 64.51007306 | 43.95523363 | 83.67027624 | 0.001156451 | 1.306323121 | 0.000488622 | 3.486917765 | 2.473104362 |
| CNAG_00791 | HLH1 | hypothetical protein | 332.7798873 | 471.5700548 | 340.1725218 | 282.2391678 | 272.3585371 | 244.4853138 | 0.001157422 | -0.532008839 | 0.000489173 | -3.486616097 | 0.691591077 |
| CNAG_13108 | - | unspecified product | 0.340297815 | 5.567216123 | 8.648592115 | 32.33319908 | 21.82132122 | 20.92422274 | 0.001163127 | 2.476242161 | 0.000491726 | 3.485223407 | 5.564461819 |
| CNAG_03888 | - | hypothetical protein | 5690.34523 | 5788.978456 | 5541.266644 | 4623.530588 | 4890.60643 | 5248.86051 | 0.001179634 | -0.220409563 | 0.000498849 | -3.481373822 | 0.858321735 |
| CNAG_06455 | - | hypothetical protein | 44.74141257 | 45.41707476 | 36.45938924 | 76.60589709 | 101.5774706 | 76.44120172 | 0.001180932 | 1.00182051 | 0.000499542 | 3.481001945 | 2.002525356 |
| CNAG_00139 | - | hypothetical protein | 2767.618447 | 2499.735301 | 2367.634672 | 3260.588561 | 2912.901217 | 3122.774121 | 0.001183978 | 0.268595946 | 0.000500975 | 3.480234524 | 1.204634887 |
| CNAG_07847 | - | hypothetical protein | 5.023222562 | 5.54599319 | 1.224407074 | 31.73947477 | 14.70222771 | 20.84978984 | 0.00118433 | 2.54905249 | 0.000501269 | 3.480077621 | 5.852497813 |
| CNAG_05964 | - | Poly(3-hydroxybutyrate) depolymerase | 104.2750095 | 112.1773711 | 85.52130206 | 213.6162615 | 132.7851556 | 172.9356171 | 0.001185229 | 0.766135273 | 0.000501794 | 3.47979707 | 1.700707778 |
| CNAG_02998 | - | nuclear protein | 2766.250058 | 2688.14682 | 2730.069019 | 2069.080117 | 2399.571949 | 2432.407209 | 0.00118869 | -0.261628079 | 0.000503404 | -3.478938441 | 0.834146056 |
| CNAG_13158 | - | unspecified product | 29.06256144 | 9.690696873 | 11.6792775 | 1.87363168 | 2.317496977 | 2.419862438 | 0.001211923 | -3.312622834 | 0.000513391 | -3.4736705 | 0.100647076 |
| CNAG_01731 | - | hypothetical protein | 408.9771047 | 454.7318448 | 390.4490011 | 563.8054445 | 591.9974543 | 512.8479805 | 0.001217052 | 0.397257498 | 0.000515713 | 3.472459545 | 1.317001965 |
| CNAG_02897 | KCS1 | inositol hexaphosphate kinase 1 | 1543.883044 | 1421.241019 | 1432.335086 | 1656.976818 | 1798.556107 | 2019.137535 | 0.001220183 | 0.30069804 | 0.000517189 | 3.471692423 | 1.23174024 |
| CNAG_00063 | CSE4 | Histone H3 | 1022.786216 | 995.4944516 | 1020.532236 | 818.5019286 | 866.780959 | 817.2086534 | 0.001230881 | -0.295583996 | 0.000521873 | -3.469270766 | 0.814742462 |
| CNAG_01246 | - | hypothetical protein, variant 2 | 541.0893013 | 703.6014485 | 669.7943296 | 818.9940119 | 830.8126866 | 843.5699392 | 0.001234081 | 0.366594139 | 0.00052338 | 3.468495914 | 1.289305491 |
| CNAG_03390 | - | AAA family ATPase | 1576.322731 | 1522.375268 | 1594.282859 | 1261.324114 | 1364.488867 | 1328.955053 | 0.001235231 | -0.262735483 | 0.000524019 | -3.468168168 | 0.833506016 |
| CNAG_07515 | - | hypothetical protein | 274.2214823 | 261.7614315 | 318.6702611 | 213.7827376 | 204.226369 | 175.6746929 | 0.00124483 | -0.54269718 | 0.000528396 | -3.465932713 | 0.686486293 |
| CNAG_02960 | - | RNA exonuclease 4 | 441.9733863 | 321.0824828 | 372.097645 | 241.5356979 | 253.3184518 | 300.9283434 | 0.00124483 | -0.530558424 | 0.000528304 | -3.465979168 | 0.692286719 |
| CNAG_02153 | TUP1 | glucose repression regulatory protein TUP1 | 1399.251016 | 1645.26303 | 1512.30173 | 1886.046417 | 1881.39616 | 1776.206676 | 0.00124815 | 0.268275714 | 0.000529957 | 3.465139295 | 1.204367526 |
| CNAG_07585 | - | hypothetical protein | 0.233614081 | 0.112663999 | 2.250386642 | 17.92420126 | 23.23312734 | 5.663721759 | 0.001259604 | 4.474131256 | 0.000534974 | 3.462604518 | 22.22530414 |

| | | | | | | | | | | | | |
|---|---|---|---|---|---|---|---|---|---|---|---|---|
| CNAG_03020 | - | hypothetical protein | 333.1513753 | 340.6148083 | 402.0912257 | 474.2135417 | 462.4406078 | 532.6092091 | 0.001263529 | 0.434330461 | 0.000536795 | 3.461689828 | 1.351283571 |
| CNAG_07481 | - | hypothetical protein | 692.3061038 | 689.481294 | 700.5832569 | 594.5001405 | 526.8153143 | 530.8181358 | 0.001264585 | -0.350031936 | 0.000537398 | -3.461387682 | 0.78456673 |
| CNAG_02143 | - | hypothetical protein | 347.1015663 | 594.4239934 | 555.4131603 | 352.2190396 | 364.6855154 | 302.2487109 | 0.001265206 | -0.568876292 | 0.000537817 | -3.461178143 | 0.674141669 |
| CNAG_01463 | - | hypothetical protein | 476.3199016 | 521.3417716 | 495.8230083 | 414.6937214 | 368.0964449 | 358.7763333 | 0.001284944 | -0.403078228 | 0.000546364 | -3.456931372 | 0.756242991 |
| CNAG_02679 | - | mitochondrial distribution and morphology protein 31 | 435.3139453 | 406.1324304 | 441.3987304 | 321.9134931 | 286.0693517 | 347.559432 | 0.00128642 | -0.441657413 | 0.000547149 | -3.456544598 | 0.736288252 |
| CNAG_03918 | SOG2 | ram signaling network protein | 786.0838568 | 832.9025942 | 717.8443817 | 941.0830776 | 966.1881631 | 1042.248873 | 0.001286626 | 0.322012121 | 0.000547393 | 3.456424125 | 1.250072805 |
| CNAG_07901 | FZC29 | hypothetical protein | 1594.199424 | 1688.399207 | 1694.645955 | 1945.074802 | 1941.701345 | 2082.514481 | 0.001287492 | 0.247302559 | 0.000547919 | 3.456165289 | 1.186985704 |
| CNAG_03382 | - | hypothetical protein | 929.7955425 | 1063.459688 | 946.0400798 | 1144.148228 | 1212.884551 | 1325.318549 | 0.001288121 | 0.310730475 | 0.000548344 | 3.455956442 | 1.240335556 |
| CNAG_00390 | YMR1 | myotubularin | 677.6489385 | 828.5834705 | 756.2004505 | 883.3870681 | 1081.988583 | 961.2872474 | 0.001288782 | 0.356823792 | 0.000548783 | 3.455740764 | 1.280603441 |
| CNAG_06549 | CET1 | mRNA-capping enzyme subunit beta | 715.6204592 | 624.9228326 | 804.9918976 | 545.2828599 | 494.8258585 | 592.8965161 | 0.001295945 | -0.409644547 | 0.000551991 | -3.454169024 | 0.752808829 |
| CNAG_00511 | - | hypothetical protein | 95.07565861 | 143.3183761 | 87.66718285 | 192.8716186 | 158.8052355 | 198.3674922 | 0.001298542 | 0.740745045 | 0.000553256 | 3.453551853 | 1.671038583 |
| CNAG_00046 | - | 26S proteasome regulatory subunit N8 | 5669.295641 | 5609.057207 | 5326.922986 | 5049.468395 | 4781.05789 | 4719.401692 | 0.001305472 | -0.205831072 | 0.000556368 | -3.452038725 | 0.867039081 |
| CNAG_03154 | - | hypothetical protein, variant | 375.0966473 | 412.7110776 | 347.7985074 | 520.2656346 | 489.2036847 | 504.9792667 | 0.001320183 | 0.401298922 | 0.000562799 | 3.4489366 | 1.320696457 |
| CNAG_07518 | - | ribonuclease P/MRP protein subunit POP5 | 566.3998545 | 631.3088556 | 769.758933 | 469.6886503 | 507.0198127 | 510.8002887 | 0.001325388 | -0.418988222 | 0.000565341 | -3.447719199 | 0.747948986 |
| CNAG_01268 | - | hypothetical protein | 4.029674621 | 9.901502533 | 16.10038 | 36.55279163 | 27.75027914 | 41.67671942 | 0.001325388 | 1.828471132 | 0.000565329 | 3.447725218 | 3.551604985 |
| CNAG_03823 | HRT1 | RING-box protein 1 | 896.7920876 | 976.0639835 | 990.6468019 | 802.2385953 | 776.155294 | 762.8162179 | 0.001332786 | -0.305491335 | 0.000568822 | -3.446060857 | 0.809166594 |
| CNAG_00053 | - | hypothetical protein | 308.089113 | 339.2131621 | 305.9986706 | 237.1026622 | 247.4308473 | 193.4542034 | 0.001332786 | -0.506855589 | 0.000568667 | -3.446134603 | 0.703754629 |
| CNAG_05421 | - | hypothetical protein | 908.4529753 | 1041.730607 | 934.1706054 | 778.0472534 | 743.3261403 | 816.0435354 | 0.001338437 | -0.317420391 | 0.000571398 | -3.44483998 | 0.802503511 |
| CNAG_02427 | - | hypothetical protein | 491.8831886 | 489.2773253 | 521.5676648 | 758.7075273 | 644.4627897 | 592.8179218 | 0.001341347 | 0.394392658 | 0.000572804 | 3.444175388 | 1.314389315 |
| CNAG_01182 | - | cytoplasmic protein | 1416.262258 | 1419.951479 | 1245.621356 | 1115.052726 | 1178.177965 | 1079.224021 | 0.001359584 | -0.290169616 | 0.000580758 | -3.440445322 | 0.817805904 |
| CNAG_05702 | - | hypothetical protein | 1268.998385 | 1133.400659 | 1060.050941 | 934.4827798 | 976.3201323 | 898.2768041 | 0.001359679 | -0.317351128 | 0.000580964 | -3.440349055 | 0.80254204 |
| CNAG_07564 | - | hypothetical protein | 510.6828927 | 652.945412 | 568.5168639 | 806.7575094 | 739.3518204 | 718.1993891 | 0.001365241 | 0.373226477 | 0.000583507 | 3.439166901 | 1.295246314 |
| CNAG_00239 | YAP1 | hypothetical protein | 87.87328641 | 74.54739746 | 83.32026894 | 144.5686301 | 120.856042 | 151.7396486 | 0.001371309 | 0.750753146 | 0.000586269 | 3.437888752 | 1.682671024 |
| CNAG_01424 | - | hypothetical protein | 1178.401225 | 1017.251151 | 1210.463801 | 1356.678373 | 1460.502384 | 1438.638597 | 0.001382451 | 0.3051638 | 0.000591201 | 3.435619483 | 1.235558911 |
| CNAG_02929 | - | transcription initiation factor TFIID subunit 1 | 3327.034949 | 3320.69841 | 3090.667652 | 3709.424603 | 3781.553771 | 3957.599062 | 0.001385038 | 0.218304921 | 0.000592476 | 3.435035417 | 1.163365899 |
| CNAG_04103 | - | hypothetical protein | 233.9965513 | 93.8166032 | 101.343547 | 71.00541369 | 68.61399717 | 70.23820416 | 0.001390889 | -1.049704549 | 0.000595319 | -3.433738281 | 0.483067082 |
| CNAG_00770 | MSH2 | DNA mismatch repair protein Msh2 | 2036.675314 | 2006.242955 | 1920.762174 | 2310.867321 | 2364.390656 | 2388.965112 | 0.001390889 | 0.229038996 | 0.00059525 | 3.433769418 | 1.172053964 |
| CNAG_05173 | - | DNA-3-methyladenine glycosylase II | 553.3316488 | 552.6394125 | 557.6767813 | 452.7723494 | 409.487255 | 437.5302017 | 0.001395731 | -0.37178331 | 0.000597562 | -3.432718756 | 0.772826618 |
| CNAG_06688 | - | mitofusin | 2204.469139 | 2064.304534 | 1942.006132 | 1713.555926 | 1825.107952 | 1710.878388 | 0.00140101 | -0.258354804 | 0.000599993 | -3.431617647 | 0.836040766 |
| CNAG_00473 | OST2 | oligosaccharyltransferase complex subunit epsilon | 935.2172314 | 1053.548435 | 845.7062774 | 725.0391977 | 782.2263137 | 758.362342 | 0.001408309 | -0.337417284 | 0.000603463 | -3.430053268 | 0.791456913 |
| CNAG_04364 | - | dolichyl-diphosphooligosaccharide-protein glycosyltransferase | 2524.484435 | 2524.333848 | 2241.691636 | 2106.679361 | 2121.814697 | 1954.030388 | 0.001408309 | -0.253084038 | 0.0006034 | -3.430081366 | 0.839100757 |
| CNAG_07920 | - | hypothetical protein | 3.768266454 | 0.117365824 | 2.256057622 | 16.13802675 | 13.12478376 | 18.50082018 | 0.001412783 | 3.214785358 | 0.000605552 | 3.429115282 | 9.284249915 |
| CNAG_02906 | - | ubiquitin-conjugating enzyme (huntingtin interacting protein 2) | 2078.340568 | 1787.468088 | 1977.1407 | 1639.657886 | 1633.562396 | 1626.492776 | 0.001418012 | -0.270200046 | 0.000607967 | -3.428035128 | 0.829204559 |
| CNAG_02238 | - | ubiquitin-conjugating enzyme E2 G2 | 518.2511543 | 474.0382555 | 535.2429781 | 393.5337089 | 401.8023237 | 382.0599293 | 0.001419917 | -0.392525932 | 0.000608957 | -3.427593407 | 0.761794654 |
| CNAG_04452 | RSM18 | small subunit ribosomal protein S18 | 927.003836 | 858.6891792 | 959.7054524 | 741.4775458 | 788.3103082 | 675.1748925 | 0.001430872 | -0.331990114 | 0.00061383 | -3.425428667 | 0.794439844 |
| CNAG_06386 | - | hypothetical protein | 918.6939105 | 752.0654409 | 885.0565927 | 664.3684129 | 692.5031003 | 668.4560314 | 0.001456353 | -0.352292774 | 0.000624939 | -3.420553223 | 0.783338203 |
| CNAG_06010 | - | aldehyde dehydrogenase (NAD), variant | 1592.556357 | 1442.648852 | 1455.674089 | 1199.838372 | 1229.000217 | 1311.156731 | 0.001463874 | -0.279293027 | 0.000628345 | -3.419074532 | 0.823994706 |
| CNAG_04814 | - | hypothetical protein | 24.89011535 | 21.75256967 | 21.50639935 | 43.77299888 | 52.07935547 | 66.06076064 | 0.00146734 | 1.271187581 | 0.000630012 | 3.418353633 | 2.413601639 |
| CNAG_02493 | - | hypothetical protein | 871.3572782 | 1350.82375 | 1213.483188 | 747.4037581 | 826.96385 | 966.9906393 | 0.001482602 | -0.448906261 | 0.000636746 | -3.415459125 | 0.732598036 |
| CNAG_05394 | - | cation:cation antiporter | 428.6980159 | 391.2264969 | 373.3831318 | 518.5787665 | 504.7046841 | 564.8443456 | 0.001492073 | 0.396664426 | 0.000640995 | 3.413647132 | 1.316460675 |
| CNAG_07432 | - | hypothetical protein | 93.78244965 | 74.59608919 | 108.8595445 | 196.4837329 | 148.566752 | 132.0730342 | 0.001494386 | 0.775283245 | 0.000642172 | 3.413147495 | 1.711526044 |
| CNAG_04116 | - | hypothetical protein, variant | 9.821558305 | 12.05208705 | 14.01534407 | 41.32111121 | 48.20875539 | 23.36646625 | 0.001503362 | 1.662515966 | 0.000646396 | 3.411360381 | 3.165681177 |
| CNAG_12919 | - | unspecified product | 53.17403966 | 55.22972773 | 107.7058387 | 114.5526856 | 141.8665137 | 159.3714549 | 0.001503362 | 0.929460504 | 0.000646246 | 3.411423836 | 1.904563651 |
| CNAG_12825 | - | unspecified product | 72.66096528 | 61.60352411 | 57.77105092 | 132.4417332 | 98.43383579 | 113.0030256 | 0.00150699 | 0.836804465 | 0.00064814 | 3.410625635 | 1.786089611 |
| CNAG_03647 | - | hypothetical protein | 402.7040491 | 402.8805745 | 428.5996794 | 319.6220306 | 303.4081342 | 310.9950192 | 0.00150855 | -0.417865249 | 0.000648995 | -3.410266134 | 0.748531405 |
| CNAG_06833 | - | hypothetical protein | 2.808675658 | 3.45605008 | 13.94498874 | 19.48870966 | 46.86896517 | 23.19145057 | 0.001509253 | 2.272048231 | 0.000649666 | 3.40998437 | 4.830083838 |
| CNAG_00250 | - | hypothetical protein | 158.0716231 | 134.4952415 | 105.5804578 | 72.60111121 | 86.04460125 | 73.53687641 | 0.001509253 | -0.797069188 | 0.000649663 | -3.409985709 | 0.575517144 |
| CNAG_03905 | - | hypothetical protein | 266.8679118 | 190.7346534 | 253.6423493 | 148.943812 | 162.8024341 | 161.1576866 | 0.001525913 | -0.606732818 | 0.000657024 | -3.406911683 | 0.656682167 |
| CNAG_07335 | - | cystinosin | 104.5165881 | 174.5738039 | 129.2061793 | 182.904438 | 218.4833527 | 273.676693 | 0.001526858 | 0.712632699 | 0.000657618 | 3.406665149 | 1.638791936 |
| CNAG_12798 | - | unspecified product | 112.5644524 | 167.0045312 | 96.21141335 | 200.4254 | 229.9270146 | 188.6011862 | 0.001537097 | 0.704868969 | 0.000662215 | 3.404762949 | 1.629996615 |
| CNAG_06237 | - | YeeE/YedE family protein | 1432.758203 | 1441.565541 | 1565.458744 | 1192.83762 | 1278.222945 | 1254.574394 | 0.001537454 | -0.268425316 | 0.000662557 | -3.404622158 | 0.830225232 |
| CNAG_00769 | PBS2 | STE/STE7 protein kinase | 534.9147935 | 508.6211505 | 370.2658123 | 657.2415585 | 683.4397244 | 601.6358799 | 0.001540673 | 0.443797616 | 0.000664132 | 3.403973471 | 1.36018003 |
| CNAG_02265 | - | exportin-T | 2540.81169 | 2370.311219 | 2309.921569 | 2042.028637 | 2115.778838 | 2033.920624 | 0.001547356 | -0.23739928 | 0.000667201 | -3.402713753 | 0.848273101 |
| CNAG_02310 | - | casein kinase II subunit beta | 1109.342166 | 1130.201569 | 1361.69477 | 986.6594533 | 980.5929077 | 943.7831612 | 0.001559655 | -0.322932061 | 0.000672695 | -3.400472371 | 0.799443477 |
| CNAG_04618 | - | hypothetical protein | 210.4036463 | 203.7495143 | 166.5500213 | 282.8590432 | 338.6496831 | 246.4449377 | 0.001561465 | 0.565222767 | 0.000673857 | 3.400000441 | 1.479615959 |
| CNAG_07699 | - | zinc finger family protein | 547.7946202 | 510.8031912 | 477.8908579 | 664.9341043 | 652.2362313 | 663.7426985 | 0.001561465 | 0.352102597 | 0.000673744 | 3.40004647 | 1.276419538 |
| CNAG_00039 | ZFC6 | hypothetical protein | 573.2536866 | 605.4185616 | 607.7909492 | 461.2313082 | 501.9254517 | 442.0260933 | 0.001563755 | -0.361602789 | 0.000675037 | -3.399522376 | 0.778299433 |
| CNAG_02717 | - | aryl-alcohol dehydrogenase | 1536.72853 | 1715.163658 | 1599.638921 | 1368.03466 | 1374.80454 | 1370.036399 | 0.001570536 | -0.252911816 | 0.000678155 | -3.398261641 | 0.839200932 |
| CNAG_04113 | PTP3 | hypothetical protein | 3250.907041 | 3197.78012 | 3206.789557 | 2822.699137 | 2867.936504 | 2768.83982 | 0.001584696 | -0.206191363 | 0.000684463 | -3.395728061 | 0.866822578 |
| CNAG_05385 | - | hypothetical protein | 1174.552528 | 1118.321966 | 1141.04511 | 833.8982426 | 944.373769 | 1015.829314 | 0.001586593 | -0.313572277 | 0.000685476 | -3.395323159 | 0.804646893 |
| CNAG_00566 | - | DNA ligase 1 | 1424.513545 | 1473.831225 | 1344.781342 | 1150.046885 | 1185.892277 | 1229.016847 | 0.001589637 | -0.265887586 | 0.000686986 | -3.394721029 | 0.831686901 |
| CNAG_03479 | - | hypothetical protein | 1353.517201 | 1320.991854 | 1162.554643 | 1534.945457 | 1582.080412 | 1576.318336 | 0.001589694 | 0.275772375 | 0.000687205 | 3.394633803 | 1.210642056 |
| CNAG_02660 | - | hypothetical protein | 16.82410202 | 25.97542671 | 14.07665539 | 60.31497731 | 46.02046058 | 39.85932342 | 0.001596197 | 1.385036061 | 0.000690211 | 3.39343847 | 2.611784856 |
| CNAG_12041 | - | unspecified product | 9.617096703 | 6.638225586 | 2.310556967 | 23.02349026 | 24.49251969 | 28.46289715 | 0.00160173 | 2.125699626 | 0.000692799 | 3.392413231 | 4.364146785 |
| CNAG_02047 | - | hypothetical protein | 699.0847236 | 607.6150516 | 582.2642082 | 494.7286091 | 490.6244436 | 489.7208668 | 0.001605947 | -0.373798733 | 0.000694819 | -3.3916155 | 0.771747745 |
| CNAG_12951 | - | unspecified product | 11.02130912 | 23.78575571 | 8.753135733 | 42.69282521 | 35.55210768 | 49.46572946 | 0.001620226 | 1.573280469 | 0.000701392 | 3.38903436 | 2.975805984 |
| CNAG_00717 | - | hypothetical protein | 249.0766261 | 269.4749764 | 281.6216023 | 414.6789261 | 339.920417 | 361.7829129 | 0.001620226 | 0.464575103 | 0.000701272 | 3.389081414 | 1.379910887 |
| CNAG_00167 | - | hypothetical protein | 321.4038472 | 342.7109671 | 327.4997621 | 418.0988183 | 447.0049099 | 460.5200799 | 0.001624683 | 0.40551044 | 0.00070352 | 3.38820362 | 1.32455747 |
| CNAG_12605 | - | unspecified product | 0.310885427 | 2.339653457 | 8.621025352 | 17.40083101 | 31.02413701 | 17.6238367 | 0.001637411 | 2.684205964 | 0.000709232 | 3.385985407 | 6.427269487 |
| CNAG_00525 | - | dynein light intermediate chain 1, cytosolic | 1011.25404 | 1125.818461 | 1119.695741 | 889.3488812 | 876.1981396 | 924.8443652 | 0.001646869 | -0.290391385 | 0.000713529 | -3.384327216 | 0.817680202 |
| CNAG_12043 | - | unspecified product | 2.814776281 | 9.862689628 | 11.84794662 | 33.5835844 | 27.0244801 | 27.51756254 | 0.001650933 | 1.970345537 | 0.000715492 | 3.383573012 | 3.91861962 |
| CNAG_12697 | - | unspecified product | 34.55980357 | 73.36174842 | 67.30415331 | 109.451511 | 100.2331045 | 130.5201154 | 0.001651061 | 0.949453918 | 0.000715749 | 3.383474355 | 1.931141554 |
| CNAG_05057 | - | hypothetical protein, variant | 271.1034596 | 269.4473414 | 221.9878011 | 370.1263677 | 335.655499 | 361.7257249 | 0.001663201 | 0.471294409 | 0.000721215 | 3.381384743 | 1.386352768 |
| CNAG_12201 | - | unspecified product | 21.58180645 | 32.48954304 | 34.25471932 | 73.27302323 | 67.70220693 | 54.29760113 | 0.001667045 | 1.142503773 | 0.000723085 | 3.380673054 | 2.207638228 |
| CNAG_05230 | - | sentrin/sumo-specific protease | 761.280544 | 806.9041721 | 799.7671331 | 602.8304099 | 664.932114 | 650.6570737 | 0.001681493 | -0.319444704 | 0.000729557 | -3.378223601 | 0.80137827 |
| CNAG_06609 | - | hypothetical protein | 3.928459865 | 11.93809287 | 2.3102008 | 22.19212128 | 27.20312483 | 26.31474353 | 0.001688046 | 2.217966139 | 0.000732607 | 3.377076515 | 4.652370972 |
| CNAG_02205 | - | pH-response regulator protein palC, variant | 294.5854689 | 306.1135982 | 323.2130718 | 447.9824724 | 459.2133402 | 365.2551147 | 0.001690789 | 0.446134355 | 0.00073421 | 3.376475121 | 1.362384905 |
| CNAG_12537 | - | unspecified product | 34.53254473 | 71.19202516 | 60.91957085 | 101.6227588 | 111.6934581 | 106.3037328 | 0.001690789 | 0.933419309 | 0.000734092 | 3.376519281 | 1.909797017 |
| CNAG_01426 | - | vacuolar protein sorting 26, variant 2 | 2414.750873 | 2397.219307 | 2238.4759 | 2072.454188 | 2025.182396 | 1978.418363 | 0.001699495 | -0.229742039 | 0.000738198 | -3.374984705 | 0.852787361 |
| CNAG_12022 | - | unspecified product | 59.82527939 | 122.4210548 | 118.0619755 | 50.81483372 | 50.54710957 | 50.23208477 | 0.001702483 | -1.008153413 | 0.000739704 | -3.374423777 | 0.497182212 |
| CNAG_01893 | - | glutathione S-transferase Gst3 | 302.5930366 | 269.4941809 | 247.571639 | 450.6305745 | 353.7299556 | 358.5029654 | 0.001707251 | 0.489762887 | 0.000741984 | 3.373576634 | 1.404214068 |
| CNAG_02663 | - | ribonucleoside-diphosphate reductase subunit M2 | 976.3769121 | 899.7715085 | 949.2136015 | 1181.688779 | 1246.63841 | 1082.389381 | 0.001734348 | 0.297551185 | 0.000753972 | 3.369161558 | 1.229056455 |
| CNAG_01730 | STE7 | STE/STE7/MEK1 protein kinase | 432.5386032 | 615.2241852 | 542.9067795 | 692.1419338 | 716.1291688 | 710.3506752 | 0.001740528 | 0.400677892 | 0.000756871 | 3.36810363 | 1.320128065 |
| CNAG_04032 | - | AFG3 family protein | 8455.541555 | 7980.520317 | 8193.524146 | 6760.725939 | 7326.516221 | 7491.405511 | 0.001743351 | -0.206330028 | 0.000758312 | -3.36757934 | 0.866739267 |
| CNAG_02348 | - | hypothetical protein | 749.4735575 | 735.7994196 | 695.2937984 | 597.5216116 | 557.0087228 | 598.4770916 | 0.001759032 | -0.32980714 | 0.000765348 | -3.365032039 | 0.795642838 |
| CNAG_01837 | - | vacuolar protein sorting-associated protein VPS35 | 3800.854424 | 3684.763191 | 3463.836017 | 3150.334515 | 3110.265303 | 3260.600081 | 0.001763607 | -0.216731544 | 0.000767554 | -3.364237866 | 0.860512739 |
| CNAG_01215 | - | hypothetical protein | 695.5913354 | 533.3710061 | 677.0661624 | 490.8236832 | 487.1668802 | 478.6387444 | 0.001763827 | -0.405216236 | 0.000767864 | -3.364126128 | 0.755123104 |

| | | | | | | | | | | | | | |
|---|---|---|---|---|---|---|---|---|---|---|---|---|---|
| CNAG_01350 | RMD5 | putative ubiquitin ligase | 615.6284783 | 659.4470967 | 587.7280747 | 883.1091958 | 770.3719382 | 738.2224708 | 0.001776019 | 0.346673634 | 0.000773389 | 3.362147054 | 1.271625307 |
| CNAG_00539 | - | membrane transporter | 2154.214 | 884.5859103 | 911.8098063 | 492.543748 | 411.6629982 | 508.8809239 | 0.001777372 | -1.503751054 | 0.000774196 | -3.361859328 | 0.352635334 |
| CNAG_00313 | - | hypothetical protein | 334.2684876 | 362.1046341 | 356.2789901 | 474.3121067 | 453.8452276 | 466.1246271 | 0.001784139 | 0.390136366 | 0.000777361 | 3.360732455 | 1.31051727 |
| CNAG_06696 | - | protein phosphatase PP2A regulatory subunit B | 5433.448384 | 5398.971907 | 5317.311285 | 4718.600045 | 4897.561008 | 4726.04029 | 0.001809781 | -0.186525707 | 0.000788754 | -3.356711359 | 0.878719304 |
| CNAG_00756 | - | hypothetical protein | 525.3642559 | 565.5308458 | 535.2969063 | 433.2701392 | 429.398161 | 416.4629189 | 0.001812833 | -0.361514088 | 0.000790749 | -3.356012989 | 0.778347286 |
| CNAG_08002 | - | hypothetical protein | 3299.905467 | 3175.151319 | 3111.889025 | 2742.49877 | 2718.660364 | 2879.806097 | 0.001812833 | -0.216198587 | 0.000790731 | -3.356018988 | 0.860830687 |
| CNAG_01023 | - | cohesin complex subunit SCC1 | 768.0275758 | 674.4254003 | 687.8078524 | 623.370779 | 542.3600489 | 482.0567991 | 0.001812833 | -0.386146706 | 0.000790356 | -3.356150441 | 0.765170571 |
| CNAG_04310 | - | hypothetical protein | 728.6541777 | 989.858726 | 748.6228239 | 511.52385 | 740.9823038 | 555.3148936 | 0.001815796 | -0.462968829 | 0.000792263 | -3.355483899 | 0.725491779 |
| CNAG_02446 | - | response regulator receiver protein | 231.462429 | 205.9653835 | 241.0961331 | 336.6929514 | 339.2913219 | 287.4898025 | 0.001821 | 0.490952574 | 0.000794755 | 3.35461488 | 1.405372501 |
| CNAG_05768 | - | hypothetical protein | 318.9430679 | 293.1807465 | 257.1765284 | 368.2307978 | 472.292081 | 381.8049744 | 0.00182488 | 0.476562137 | 0.000796895 | 3.353871072 | 1.391424025 |
| CNAG_02790 | - | galactinol synthase | 792.1692772 | 566.714354 | 547.1004251 | 451.7972785 | 455.1986455 | 493.0295466 | 0.00182488 | -0.46184037 | 0.00079674 | -3.353924688 | 0.726059472 |
| CNAG_00017 | - | hypothetical protein | 89.22073312 | 101.4443902 | 118.4517413 | 155.9426705 | 192.0086713 | 153.0834624 | 0.001831261 | 0.683616679 | 0.000799905 | 3.352827642 | 1.606161183 |
| CNAG_00024 | - | hypothetical protein, variant 2 | 4863.581536 | 4815.004851 | 4776.615451 | 4387.130403 | 4177.26452 | 4216.477497 | 0.001840314 | -0.192826621 | 0.000804084 | -3.351385183 | 0.874889901 |
| CNAG_07328 | - | hypothetical protein | 387.9174514 | 414.8722227 | 353.1305647 | 522.5742665 | 477.9350051 | 532.6833558 | 0.001840658 | 0.393138433 | 0.000804459 | 3.351255967 | 1.313247132 |
| CNAG_01033 | - | exosome complex component CSL4 | 901.2779526 | 836.0471199 | 861.6381299 | 706.4514864 | 728.7544354 | 690.6577895 | 0.001849161 | -0.306213248 | 0.000808401 | -3.349902105 | 0.808761794 |
| CNAG_02303 | - | pre-mRNA-splicing factor ATP-dependent RNA helicase prp16 | 4749.096367 | 4488.560693 | 4617.703553 | 3835.15202 | 4057.392333 | 4201.999584 | 0.001849433 | -0.211614971 | 0.000808746 | -3.349784083 | 0.863569999 |
| CNAG_07527 | - | alpha-1,6-mannosyltransferase | 9.796537124 | 13.10205886 | 9.781127937 | 37.31317286 | 17.03877274 | 54.6468735 | 0.001860299 | 1.788191499 | 0.000813725 | 3.348083596 | 3.453816652 |
| CNAG_12928 | - | unspecified product | 0.204672472 | 0.098915542 | 2.233251996 | 12.09435772 | 13.26369401 | 14.14152268 | 0.00186548 | 4.269408943 | 0.000816219 | 3.347235417 | 19.28502276 |
| CNAG_04981 | CAT1 | Catalase 1 | 4827.024077 | 5042.122561 | 3624.907866 | 3443.721973 | 3045.435813 | 2715.753707 | 0.001884078 | -0.566744553 | 0.000824586 | -3.344407225 | 0.675138523 |
| CNAG_06016 | CAP6 | alpha-1,3-mannosyltransferase | 1359.269307 | 1499.708289 | 1549.465189 | 1234.162258 | 1259.235718 | 1213.52422 | 0.001910393 | -0.264925858 | 0.000836337 | -3.340480162 | 0.832241504 |
| CNAG_01929 | - | hypothetical protein | 51.7843877 | 54.04310795 | 54.53343689 | 85.03043284 | 93.40433248 | 116.0875184 | 0.001926949 | 0.871947452 | 0.000843819 | 3.338005931 | 1.830131676 |
| CNAG_03214 | - | tRNA (uracil-O(2)-)-methyltransferase | 310.8190507 | 283.5087626 | 299.770131 | 394.8302431 | 450.6208177 | 375.1766655 | 0.001931456 | 0.434760084 | 0.000846029 | 3.337279133 | 1.351686033 |
| CNAG_05358 | - | hypothetical protein | 8.51937899 | 9.838552733 | 1.244932033 | 30.44227055 | 29.83225251 | 20.98779355 | 0.001936106 | 2.119639391 | 0.000848303 | 3.336533349 | 4.345853046 |
| CNAG_05992 | - | hypothetical protein | 13.38418126 | 23.84402498 | 19.36185192 | 43.19014675 | 38.93562035 | 63.7349706 | 0.001937394 | 1.375562553 | 0.000849103 | 3.336271003 | 2.594690665 |
| CNAG_06165 | - | sterility protein Ste20 | 2243.452492 | 2417.799886 | 2341.986457 | 2772.459801 | 2682.633261 | 2750.894277 | 0.001959656 | 0.213876237 | 0.0008591 | 3.333016143 | 1.159800156 |
| CNAG_03513 | - | ribosomal RNA-processing protein 7 | 1729.348716 | 1684.00587 | 1912.038183 | 1395.933691 | 1428.228918 | 1605.315621 | 0.00196658 | -0.281492068 | 0.000862375 | -3.331957262 | 0.822739681 |
| CNAG_00171 | - | peroxin-2 | 643.827801 | 801.6619822 | 804.1488542 | 913.0665839 | 951.5487154 | 1010.054726 | 0.001971231 | 0.339968818 | 0.000864896 | 3.331144754 | 1.265729237 |
| CNAG_04506 | - | hypothetical protein | 28.69656838 | 52.9503147 | 61.92924467 | 79.55762574 | 100.4992254 | 106.1510446 | 0.001971231 | 1.001351783 | 0.000864809 | 3.331172927 | 2.001874847 |
| CNAG_02333 | - | hypothetical protein | 660.7982338 | 713.1234316 | 631.3132812 | 507.8831348 | 543.2770199 | 549.6280418 | 0.00197502 | -0.340556919 | 0.0008668 | -3.330532802 | 0.789736393 |
| CNAG_00059 | - | hypothetical protein | 429.62272 | 442.7489935 | 482.9718094 | 334.3155436 | 343.1101266 | 360.9090285 | 0.001986649 | -0.39894318 | 0.000872146 | -3.328820468 | 0.758413642 |
| CNAG_00514 | - | hypothetical protein | 883.65364 | 753.1094038 | 796.5880559 | 656.6424084 | 644.165951 | 655.1134773 | 0.001989609 | -0.330810263 | 0.000873689 | -3.328328255 | 0.795089811 |
| CNAG_00687 | - | hypothetical protein, variant | 769.6455794 | 746.7261678 | 606.9721368 | 860.9037246 | 899.7880214 | 983.3776196 | 0.002001215 | 0.355454479 | 0.000879274 | 3.326552742 | 1.279388551 |
| CNAG_02096 | - | peroxin-16 | 76.10534643 | 64.8123017 | 49.2778513 | 123.7077755 | 110.6843949 | 106.4021103 | 0.002001215 | 0.827361457 | 0.00087912 | 3.326601463 | 1.774437127 |
| CNAG_03040 | - | transketolase, variant | 2931.084628 | 2755.052069 | 2218.384279 | 3420.14384 | 3255.435075 | 3141.69489 | 0.002009344 | 0.297279597 | 0.000883091 | 3.325345417 | 1.228825106 |
| CNAG_05765 | UBC6 | ubiquitin-conjugating enzyme E2 J2 | 2489.338008 | 2104.227431 | 2228.849688 | 1912.834599 | 1835.355872 | 1980.593693 | 0.002029482 | -0.268059853 | 0.000892189 | -3.322486841 | 0.830435571 |
| CNAG_00177 | - | hypothetical protein | 60.13959627 | 115.2597663 | 55.70489001 | 127.1457188 | 138.3107446 | 161.6295179 | 0.002029849 | 0.877276622 | 0.000892598 | 3.322358844 | 1.836904498 |
| CNAG_03642 | - | hypothetical protein | 196.4789571 | 186.5595226 | 215.4962622 | 265.5740801 | 311.748544 | 279.6125416 | 0.002042933 | 0.502490149 | 0.000898601 | 3.320488319 | 1.416656659 |
| CNAG_02119 | - | hypothetical protein | 628.6102656 | 766.0992149 | 743.3741573 | 897.5948565 | 938.6773943 | 870.2913654 | 0.002043548 | 0.325392418 | 0.000899121 | 3.32032685 | 1.253005214 |
| CNAG_01690 | - | MFS transporter, variant 2 | 415.8760329 | 409.5064228 | 358.4651149 | 540.5118113 | 505.5904486 | 512.7736977 | 0.002051554 | 0.382433339 | 0.000902894 | 3.319157765 | 1.303538632 |
| CNAG_13056 | - | unspecified product | 26.01427217 | 15.3413065 | 23.61438536 | 64.9631176 | 49.4514135 | 44.28675986 | 0.002062336 | 1.27773947 | 0.000907891 | 3.317616315 | 2.424587752 |
| CNAG_12681 | - | unspecified product | 0.205563092 | 0.099339495 | 2.23379342 | 8.65991467 | 15.09647793 | 16.26367751 | 0.002072663 | 4.26919477 | 0.000912691 | 3.316143125 | 19.28216004 |
| CNAG_05345 | - | amino acid transporter | 7.479946805 | 10.95744488 | 13.98505587 | 33.31649752 | 34.94877168 | 29.77521693 | 0.002074058 | 1.663132599 | 0.000913558 | 3.315877713 | 3.167034533 |
| CNAG_07523 | - | hypothetical protein | 270.0634736 | 296.3850597 | 291.1645284 | 396.5148004 | 364.0971111 | 400.5744833 | 0.00207938 | 0.423638385 | 0.000916156 | 3.315083861 | 1.341305981 |
| CNAG_06791 | - | hypothetical protein | 808.7395361 | 742.2005451 | 558.8860319 | 555.4722943 | 509.5392057 | 525.2682691 | 0.002080438 | -0.422959361 | 0.000916876 | -3.31486425 | 0.745893024 |
| CNAG_03834 | - | C4-hydroxylase | 194.0405632 | 162.8739101 | 176.0878973 | 228.8756888 | 260.7172331 | 300.480579 | 0.002083432 | 0.551026872 | 0.00091845 | 3.31438476 | 1.465128165 |
| CNAG_03230 | - | yippee-like, putative | 56.34704412 | 32.61695787 | 52.39868243 | 94.7032636 | 132.0629695 | 66.6900371 | 0.002089092 | 1.04235926 | 0.0009212 | 3.313548761 | 2.059592983 |
| CNAG_04944 | - | hypothetical protein, variant | 1152.329796 | 1017.089578 | 1122.899545 | 917.4097285 | 861.5059552 | 928.1798678 | 0.00208953 | -0.298050738 | 0.000921649 | -3.313412681 | 0.813350593 |
| CNAG_05862 | - | DNA excision repair protein ERCC-1, variant | 500.0476707 | 517.2622426 | 531.1605464 | 660.2375192 | 624.5647931 | 696.9893224 | 0.002094543 | 0.340078715 | 0.000924116 | 3.312665068 | 1.265825657 |
| CNAG_04356 | - | TPR repeat-containing protein | 1224.846098 | 1209.908722 | 1265.805922 | 963.9730172 | 1111.808397 | 1002.603423 | 0.002140582 | -0.281203435 | 0.00094469 | -3.306501154 | 0.822904299 |
| CNAG_04287 | - | mitochondrial import inner membrane translocase subunit | 1507.532171 | 1580.538184 | 1675.307148 | 1346.251571 | 1407.639692 | 1263.516567 | 0.002167771 | -0.26110019 | 0.000957218 | -3.302808304 | 0.834451329 |
| CNAG_08021 | - | hypothetical protein | 54.11973208 | 40.13834099 | 70.44741965 | 103.2869941 | 103.8570384 | 100.7844442 | 0.002167771 | 0.883329943 | 0.000957094 | 3.302844771 | 1.844628053 |
| CNAG_07816 | - | 2-hydroxyacid dehydrogenase | 1.443913806 | 2.281120478 | 1.180207904 | 10.29889362 | 14.12546417 | 17.36464254 | 0.00217047 | 3.343794846 | 0.000958675 | 3.302381783 | 10.15272325 |
| CNAG_04892 | - | hypothetical protein, variant 4 | 191.9394369 | 240.3459553 | 234.6995655 | 310.8593272 | 307.2527346 | 320.6385923 | 0.002176295 | 0.48021188 | 0.000961514 | 3.301552414 | 1.394948519 |
| CNAG_03637 | YKU80 | ATP-dependent DNA helicase 2 subunit 2 | 1274.250884 | 1449.166703 | 1261.71203 | 1633.850375 | 1520.725083 | 1713.918776 | 0.002181434 | 0.274092439 | 0.000964051 | 3.300813163 | 1.209233154 |
| CNAG_04127 | - | pre-mRNA-splicing factor cwc24 | 657.1794795 | 609.769518 | 612.092402 | 524.4069895 | 485.4340736 | 485.290851 | 0.002181805 | -0.345490777 | 0.000964481 | -3.300687979 | 0.787040195 |
| CNAG_00835 | - | hypothetical protein | 483.4281122 | 561.2069932 | 560.8388668 | 415.3353027 | 370.6168387 | 447.4603801 | 0.002186296 | -0.395167074 | 0.000967 | -3.299955863 | 0.760401312 |
| CNAG_02713 | - | hypothetical protein | 560.4749858 | 620.4948729 | 632.3010948 | 442.4550538 | 493.2599522 | 494.1200065 | 0.002186296 | -0.357536123 | 0.000966989 | -3.299959129 | 0.780496397 |
| CNAG_05071 | - | sulfite reductase (NADPH) hemoprotein beta- component | 79.66309993 | 73.42123425 | 63.10477565 | 113.139551 | 128.896355 | 125.1883166 | 0.002186644 | 0.752112811 | 0.000967421 | 3.299833801 | 1.684257602 |
| CNAG_12451 | - | unspecified product | 0.222463055 | 0.107373724 | 3.298364871 | 18.87210322 | 14.11309071 | 9.929941517 | 0.002206982 | 3.755864116 | 0.000976689 | 3.297157091 | 13.50914175 |
| CNAG_13132 | - | unspecified product | 166.2389252 | 215.5389402 | 179.2993712 | 260.982228 | 251.9745887 | 309.383601 | 0.002229111 | 0.536226865 | 0.000986754 | 3.29427649 | 1.450174856 |
| CNAG_06328 | - | hypothetical protein | 408.8097778 | 536.3819588 | 530.9483117 | 359.9906928 | 379.3609594 | 373.1610741 | 0.002234534 | -0.422232798 | 0.000989427 | -3.29351593 | 0.746268762 |
| CNAG_05700 | - | endoplasmic reticulum protein | 1007.581082 | 917.9620854 | 1034.364733 | 804.5126694 | 843.4927498 | 780.5893145 | 0.002236751 | -0.301251821 | 0.000990682 | -3.293159577 | 0.811547914 |
| CNAG_00145 | - | actin-2 | 1391.779395 | 1359.656241 | 1305.313467 | 1155.586083 | 1181.627748 | 1084.773192 | 0.002247669 | -0.261088602 | 0.000995792 | -3.291712817 | 0.834458032 |
| CNAG_07609 | - | ATP-dependent RNA helicase DDX51/DBP6 | 1677.788458 | 1464.221797 | 1550.575703 | 1230.154953 | 1341.183353 | 1348.919317 | 0.002256632 | -0.275251212 | 0.001000039 | -3.290515827 | 0.826306423 |
| CNAG_00786 | - | hypothetical protein | 757.5415485 | 671.1950988 | 671.8260857 | 564.8080522 | 519.027088 | 585.1359025 | 0.002279716 | -0.347090759 | 0.001010547 | -3.287573808 | 0.786167834 |
| CNAG_02549 | - | hypothetical protein | 541.1059838 | 759.5830794 | 644.2293977 | 838.4707608 | 846.3432289 | 848.0238378 | 0.002291051 | 0.367400641 | 0.001015851 | 3.286099462 | 1.290026446 |
| CNAG_06450 | - | hypothetical protein, variant | 2443.895698 | 2238.882319 | 2278.979394 | 1947.832327 | 1989.812453 | 2050.530397 | 0.002296695 | -0.23285549 | 0.001018634 | -3.285328848 | 0.850948962 |
| CNAG_03095 | - | hypothetical protein | 4.927016881 | 0.125344354 | 1.199916381 | 32.31749924 | 5.846627705 | 14.28520014 | 0.002303724 | 3.332078978 | 0.001022033 | 3.28439025 | 10.07060867 |
| CNAG_07928 | - | hypothetical protein | 40.38510723 | 37.53532564 | 40.3315318 | 10.99638942 | 20.50824821 | 13.6023633 | 0.00230433 | -1.460254162 | 0.001022865 | -3.284160997 | 0.363429098 |
| CNAG_00136 | - | ubiquitin-activating enzyme E1 | 6582.51322 | 6799.599306 | 6192.895209 | 5598.030225 | 5791.145457 | 5911.65809 | 0.00230433 | -0.193076772 | 0.001022661 | -3.284217184 | 0.874738215 |
| CNAG_06592 | UBC7 | ubiquitin-conjugating enzyme E2 G1 | 1447.760325 | 1279.977927 | 1355.412136 | 1168.833816 | 1139.314181 | 1109.179211 | 0.002306905 | -0.272389692 | 0.001024571 | -3.283691199 | 0.827946991 |
| CNAG_06434 | - | regulatory subunit for Cdc7 protein kinase | 2765.236617 | 2952.1117 | 3100.097738 | 2565.08681 | 2622.978446 | 2442.479851 | 0.002306905 | -0.223959469 | 0.001024436 | -3.283728508 | 0.856212339 |
| CNAG_06026 | - | aspartate aminotransferase | 2704.38655 | 2660.135702 | 2634.10824 | 2266.94452 | 2284.80037 | 2415.753684 | 0.002325105 | -0.214746064 | 0.001032938 | -3.281398131 | 0.861697818 |
| CNAG_03115 | FZC46 | hypothetical protein | 575.602918 | 608.6565665 | 622.7117785 | 434.6501005 | 473.3788243 | 515.1688592 | 0.00233407 | -0.359090036 | 0.001037206 | -3.28023517 | 0.779656184 |
| CNAG_07859 | - | peptidyl-prolyl cis-trans isomerase-like 2 | 749.8118205 | 688.5877968 | 697.5467467 | 875.7655689 | 929.1881148 | 858.0850471 | 0.002347675 | 0.303305759 | 0.001043538 | 3.278517668 | 1.233968665 |
| CNAG_06423 | - | hypothetical protein | 696.800239 | 602.252066 | 622.7573349 | 483.7564132 | 488.0081178 | 541.8238669 | 0.002355076 | -0.360320866 | 0.001047403 | -3.277474119 | 0.778991307 |
| CNAG_04128 | - | GTPase inhibitor | 895.5842266 | 960.9720556 | 924.5600374 | 801.5070582 | 744.2207352 | 751.7055984 | 0.002355076 | -0.290363135 | 0.00104712 | -3.277550483 | 0.817696214 |
| CNAG_04613 | - | ATP-binding cassette, subfamily F, member 3 | 5623.987932 | 5073.741522 | 5641.573465 | 6506.714372 | 6243.283335 | 6363.367092 | 0.002361448 | 0.210588283 | 0.001050526 | 3.276633712 | 1.157159939 |
| CNAG_03952 | - | hypothetical protein | 2152.163647 | 2286.261188 | 2445.283681 | 2044.463149 | 1947.537189 | 1901.82902 | 0.00237181 | -0.239402921 | 0.001055715 | -3.275242117 | 0.847095822 |
| CNAG_02067 | - | vacuolar protein | 1626.650862 | 1822.90602 | 1709.479019 | 1349.219412 | 1522.354488 | 1488.801121 | 0.00237181 | -0.257353752 | 0.001055506 | -3.27529793 | 0.836621076 |
| CNAG_06034 | - | Allantoin permease | 852.3704409 | 901.7223815 | 891.4875483 | 711.8789208 | 686.3991618 | 770.5030549 | 0.002377049 | -0.301920154 | 0.001058337 | -3.274541236 | 0.811172049 |
| CNAG_04863 | VPS25 | ESCRT-II complex subunit VPS25 | 367.0508436 | 433.2069286 | 443.6990997 | 577.1303033 | 561.7413028 | 503.9724473 | 0.002379715 | 0.385895528 | 0.001059814 | 3.274147073 | 1.306670628 |
| CNAG_02123 | - | methionyl-tRNA formyltransferase | 889.7536996 | 1015.868063 | 925.6243207 | 867.8342373 | 711.3962194 | 658.5608519 | 0.00238816 | -0.354011384 | 0.001063867 | -3.2730684 | 0.782405608 |
| CNAG_00236 | - | 8-Amino-7-oxononanoate synthase | 491.6522444 | 587.0676053 | 600.2811289 | 416.8149778 | 415.5353107 | 466.3394881 | 0.002388241 | -0.386108896 | 0.001064195 | -3.272981396 | 0.765190625 |
| CNAG_05979 | - | phosphoglycerate dehydrogenase | 621.0269699 | 607.5906164 | 592.8960897 | 444.7861937 | 514.8662838 | 486.3703837 | 0.002389965 | -0.349266183 | 0.001065255 | -3.272699915 | 0.784983273 |
| CNAG_06088 | - | aspartate aminotransferase, mitochondrial, variant | 6742.486984 | 6425.793823 | 6857.237623 | 5515.466346 | 6002.493737 | 6058.194147 | 0.002404428 | -0.203756427 | 0.001071995 | -3.270916648 | 0.86828681 |
| CNAG_04101 | - | hypothetical protein | 1838.564514 | 1412.474144 | 1244.598989 | 1134.414295 | 1229.034995 | 1134.749535 | 0.002405387 | -0.378243582 | 0.001072716 | -3.270726453 | 0.7693737 |

| | | | | | | | | | | | | | |
|---|---|---|---|---|---|---|---|---|---|---|---|---|---|
| CNAG_05617 | GPI13 | phosphatidylinositol glycan, class O, variant | 402.2237798 | 331.7392706 | 287.944897 | 219.7767295 | 288.090961 | 190.1670389 | 0.0024103 | -0.565769398 | 0.001075202 | -3.270071903 | 0.675595021 |
| CNAG_13155 | - | unspecified product | 30.32849613 | 47.10639582 | 48.77284817 | 18.20345945 | 18.69621933 | 11.43625289 | 0.002420992 | -1.42545814 | 0.001080267 | -3.268742139 | 0.37230112 |
| CNAG_02302 | - | hypothetical protein | 70.64139909 | 103.5502632 | 108.8557447 | 139.5214295 | 171.2362541 | 155.1964917 | 0.00244025 | 0.71224382 | 0.001089158 | 3.266422066 | 1.638350258 |
| CNAG_00759 | - | hypothetical protein | 5.122596616 | 10.91484448 | 8.680654666 | 28.65311238 | 23.47073058 | 31.79073057 | 0.002441514 | 1.82079435 | 0.00109002 | 3.266197995 | 3.532756592 |
| CNAG_00522 | - | C2 domain-containing protein, variant | 1202.836958 | 1155.060652 | 894.9282989 | 1409.786887 | 1381.979034 | 1344.314024 | 0.002445448 | 0.331519614 | 0.001092075 | 3.265664627 | 1.258338105 |
| CNAG_03364 | - | hypothetical protein | 924.7108749 | 931.9036631 | 920.2937527 | 747.6816176 | 707.0806461 | 820.4453905 | 0.002449382 | -0.30227265 | 0.001094131 | -3.265131927 | 0.810973878 |
| CNAG_02817 | SAV1 | GTP-binding protein ypt2 | 4196.692 | 3985.371033 | 3722.99169 | 3393.921766 | 3388.875324 | 3563.668438 | 0.002451184 | -0.217837535 | 0.001095235 | -3.264846147 | 0.859853311 |
| CNAG_05460 | - | 2-dehydropantoate 2-reductase | 375.085757 | 365.3745916 | 396.7848361 | 478.8297191 | 489.2164161 | 522.6867401 | 0.002463556 | 0.375048583 | 0.001101064 | 3.263342343 | 1.296883226 |
| CNAG_07944 | - | Urea carboxylase | 688.866512 | 742.2369692 | 695.2784937 | 557.7426354 | 563.9465928 | 600.6729969 | 0.002467792 | -0.31989831 | 0.001103259 | -3.262778079 | 0.801126344 |
| CNAG_06881 | - | SBDS family rRNA metabolism protein | 1879.798016 | 1738.915046 | 1611.411683 | 1554.165717 | 1456.735414 | 1355.676255 | 0.00247276 | -0.27579528 | 0.001105782 | -3.262130579 | 0.825994866 |
| CNAG_04138 | - | hypothetical protein | 0.238183609 | 1.220151636 | 4.360699345 | 13.60766781 | 12.23207503 | 20.60844401 | 0.002474724 | 3.155373917 | 0.001107265 | 3.261750719 | 8.909681853 |
| CNAG_02125 | - | hypothetical protein, variant | 5.441375127 | 29.41014402 | 39.68563103 | 135.3664795 | 100.4156759 | 141.2872406 | 0.002474724 | 2.353311601 | 0.001107222 | 3.261761688 | 5.109958595 |
| CNAG_03050 | - | hypothetical protein | 224.4810503 | 247.8766626 | 196.3934733 | 282.5881211 | 307.2335261 | 368.1992359 | 0.002477223 | 0.506359248 | 0.001108685 | 3.261387222 | 1.420461026 |
| CNAG_05408 | - | hypothetical protein | 2544.446266 | 2530.853603 | 2634.08077 | 2131.47732 | 2312.464787 | 2254.811908 | 0.002478731 | -0.218220383 | 0.001109693 | -3.261137345 | 0.859625162 |
| CNAG_01911 | SEC6 | exocyst complex component | 1690.671346 | 1634.387226 | 1481.306027 | 1409.370355 | 1277.265527 | 1351.163986 | 0.002480196 | -0.266407368 | 0.001110622 | -3.260892466 | 0.83138731 |
| CNAG_12780 | - | unspecified product | 548.8137402 | 673.201169 | 577.9705168 | 497.1625209 | 468.1993898 | 440.9217569 | 0.002500765 | -0.370201656 | 0.001120443 | -3.258395133 | 0.773674347 |
| CNAG_06139 | - | hypothetical protein | 48.43521344 | 70.16036736 | 70.4933205 | 122.9405814 | 114.1985661 | 98.70106761 | 0.002500765 | 0.814756874 | 0.001120367 | 3.258414294 | 1.759001697 |
| CNAG_05029 | - | meiotic Sister-Chromatid recombination aldehyde dehydrogenase, variant | 584.9552875 | 664.6228029 | 589.7079673 | 499.4500589 | 496.6972648 | 466.4333861 | 0.002507968 | -0.345120861 | 0.001123976 | -3.257501566 | 0.787242023 |
| CNAG_03868 | - | hypothetical protein | 505.2247563 | 478.2421646 | 403.1224385 | 350.7297476 | 287.7346138 | 385.2770428 | 0.0025121 | -0.454404611 | 0.001126135 | -3.25695694 | 0.729811298 |
| CNAG_02567 | - | hypothetical protein | 292.5293207 | 220.8566324 | 245.1989261 | 179.4506651 | 185.2759965 | 160.1111693 | 0.002521395 | -0.547195334 | 0.00113061 | -3.255831137 | 0.684349243 |
| CNAG_06204 | - | high-affinity nicotinic acid transporter | 454.0648095 | 469.6427479 | 441.4564026 | 364.020432 | 369.9212922 | 319.9424564 | 0.002523608 | -0.389919706 | 0.00113191 | -3.255504678 | 0.763172078 |
| CNAG_07318 | - | AP-1 complex subunit gamma-1 | 2287.234792 | 1914.576623 | 1857.764179 | 1714.39798 | 1684.451681 | 1635.399083 | 0.002533326 | -0.283375347 | 0.001136888 | -3.254258551 | 0.821666386 |
| CNAG_03203 | - | DNA polymerase epsilon p12 subunit | 2428.963351 | 2764.603689 | 2764.171818 | 2214.776354 | 2268.416777 | 2355.819178 | 0.002533326 | -0.233408394 | 0.00113677 | -3.254287818 | 0.850622904 |
| CNAG_12078 | - | unspecified product | 38.88588745 | 45.3554437 | 24.76429361 | 74.5235088 | 71.02136301 | 75.19550463 | 0.002535907 | 1.025816844 | 0.001138356 | 3.253891964 | 2.03611189 |
| CNAG_02453 | PTP5 | hypothetical protein | 1617.426027 | 1535.418349 | 1479.257812 | 1844.724464 | 1921.959204 | 1772.886461 | 0.002556769 | 0.242279405 | 0.001148033 | 3.251486388 | 1.182860061 |
| CNAG_01749 | - | hypothetical protein | 494.3008934 | 564.653632 | 592.9877353 | 683.5176634 | 722.159298 | 701.5017119 | 0.002566329 | 0.337110654 | 0.001152639 | 3.250348037 | 1.263224147 |
| CNAG_03650 | - | hypothetical protein | 8.660219368 | 17.35641319 | 8.726848153 | 35.62658745 | 50.06288298 | 22.25899798 | 0.002576903 | 1.683546038 | 0.001157703 | 3.249101268 | 3.21216508 |
| CNAG_04509 | - | hypothetical protein | 7.336792929 | 6.629619999 | 4.429721087 | 23.93257114 | 25.43960111 | 21.99310249 | 0.002579163 | 2.005514926 | 0.001159033 | 3.248774573 | 4.015319884 |
| CNAG_02377 | - | Aldehyde dehydrogenase | 2441.693359 | 2415.576965 | 2580.746454 | 2101.944124 | 2154.572402 | 2203.731262 | 0.002585652 | -0.218787815 | 0.001162265 | -3.24798237 | 0.859287126 |
| CNAG_04992 | - | hypothetical protein | 347.567057 | 308.0597472 | 291.1073262 | 230.8205849 | 230.0953267 | 234.3952912 | 0.002588319 | -0.462470103 | 0.00116378 | -3.247611707 | 0.725742618 |
| CNAG_06087 | - | kinetochore protein Spc7/SPC105 | 1686.436799 | 1695.968558 | 1810.876757 | 1900.556533 | 2214.404841 | 2145.813255 | 0.002596704 | 0.25451143 | 0.001167867 | 3.246614005 | 1.192931687 |
| CNAG_07840 | - | hypothetical protein | 0.265178112 | 4.442212067 | 3.326484445 | 24.5343435 | 9.433362635 | 18.5657871 | 0.002608473 | 2.854149165 | 0.001173478 | 3.245249295 | 7.230769389 |
| CNAG_12593 | - | unspecified product | 127.8589052 | 31.34968632 | 50.00429623 | 13.17036322 | 13.28834053 | 2.641078678 | 0.002609255 | -2.915768516 | 0.001174149 | -3.245086654 | 0.132515359 |
| CNAG_05523 | - | hypothetical protein | 19.15759135 | 31.32890313 | 14.0918013 | 59.28332532 | 45.03531296 | 52.98147896 | 0.002618071 | 1.272620047 | 0.001178436 | 3.244048798 | 2.415999318 |
| CNAG_07683 | - | hypothetical protein | 605.4975391 | 492.3224623 | 446.8537811 | 357.5771202 | 422.5646042 | 375.411953 | 0.00264541 | -0.435217383 | 0.001191064 | -3.241011335 | 0.739582307 |
| CNAG_04499 | - | clathrin light chain | 2800.175777 | 2757.118965 | 2847.381237 | 2460.016387 | 2457.336499 | 2462.415225 | 0.002652735 | -0.202989328 | 0.001194686 | -3.240145679 | 0.868748612 |
| CNAG_03902 | RDS2 | Regulator of drug sensitivity 2, putative | 1347.352754 | 1335.9162 | 1175.236727 | 1147.910731 | 1047.857873 | 992.6552938 | 0.002654893 | -0.290885721 | 0.001195982 | -3.239836559 | 0.817400074 |
| CNAG_12900 | - | unspecified product | 127.4882454 | 108.9931833 | 100.4243358 | 218.9235506 | 183.1601354 | 141.0382869 | 0.002660346 | 0.678254842 | 0.001199089 | 3.239096827 | 1.600202896 |
| CNAG_07367 | - | amino acid transporter, variant | 29.70849672 | 40.01827904 | 38.53526933 | 65.67558863 | 93.91489511 | 62.0783431 | 0.002660346 | 1.024964362 | 0.0011988 | 3.239165361 | 2.034909116 |
| CNAG_13037 | - | unspecified product | 34.40289688 | 22.58205593 | 34.93068103 | 7.959158091 | 14.47120006 | 6.957722561 | 0.002681941 | -1.754913531 | 0.00120915 | -3.236712949 | 0.296290951 |
| CNAG_05689 | DAM1 | pre-mRNA-splicing factor | 1718.766778 | 1590.278605 | 1739.311358 | 1508.244854 | 1407.563321 | 1356.759927 | 0.002686125 | -0.256499506 | 0.001211364 | -3.236190765 | 0.837116601 |
| CNAG_02393 | - | hypothetical protein | 2267.912493 | 2303.603282 | 2202.297794 | 2530.271761 | 2658.516503 | 2759.731204 | 0.002690492 | 0.21572467 | 0.001214319 | 3.235495273 | 1.161287086 |
| CNAG_02678 | - | hypothetical protein | 518.7420062 | 578.6315987 | 522.6614066 | 746.0538547 | 685.8797542 | 638.3117147 | 0.002690492 | 0.33991885 | 0.001213996 | 3.235571223 | 1.265685399 |
| CNAG_02102 | - | hypothetical protein | 26.00443614 | 22.80802589 | 17.26736848 | 57.66377523 | 53.89875209 | 43.18738218 | 0.002690492 | 1.219343478 | 0.001213811 | 3.235614808 | 2.328407352 |
| CNAG_00389 | - | mitochondrial protein | 1009.911569 | 936.2626499 | 1005.591844 | 774.089971 | 865.9526924 | 795.0046865 | 0.002700349 | -0.29310334 | 0.001219098 | -3.23437386 | 0.816144584 |
| CNAG_04199 | - | hypothetical protein | 69.3749923 | 94.89808973 | 79.06475366 | 119.2565342 | 135.7180403 | 149.5090442 | 0.002702451 | 0.721684924 | 0.001220377 | 3.234074361 | 1.649106902 |
| CNAG_00304 | - | hypothetical protein | 577.9325142 | 575.3041948 | 648.2753016 | 493.2405912 | 466.4614099 | 470.847364 | 0.002708997 | -0.348348382 | 0.001223994 | -3.23322897 | 0.785482815 |
| CNAG_04220 | - | ATP-dependent DNA helicase II subunit 1 | 1159.766801 | 1122.792396 | 1246.716482 | 1454.088763 | 1367.217112 | 1454.190516 | 0.002708997 | 0.262106816 | 0.001223716 | 3.233293978 | 1.1992287 |
| CNAG_01603 | - | hypothetical protein | 527.1390828 | 728.3828587 | 718.8305995 | 905.5537648 | 880.8784991 | 804.8140918 | 0.002718469 | 0.377780962 | 0.001228606 | 3.232154588 | 1.299341774 |
| CNAG_03610 | - | mRNA guanylyltransferase | 1082.668463 | 1076.433717 | 1044.142324 | 1198.560356 | 1458.890983 | 1291.035748 | 0.002725894 | 0.286327291 | 0.001232294 | 3.231297904 | 1.219531726 |
| CNAG_05621 | - | folylpolyglutamate synthase | 686.3797412 | 714.1811471 | 575.9048759 | 536.8142137 | 523.4178769 | 499.7382877 | 0.002726048 | -0.357047418 | 0.001232697 | -3.231204512 | 0.780760831 |
| CNAG_03669 | - | hypothetical protein | 1573.124037 | 1648.513969 | 1536.836655 | 1837.661095 | 1903.798286 | 1883.855187 | 0.002728044 | 0.226268637 | 0.001233933 | 3.230918122 | 1.169805467 |
| CNAG_05660 | - | hypothetical protein | 726.0452169 | 653.9558453 | 657.9582752 | 522.6891661 | 576.1028653 | 534.1281711 | 0.002747778 | -0.334826128 | 0.001243194 | -3.228780202 | 0.792879688 |
| CNAG_01441 | - | hypothetical protein, variant | 106.7258865 | 133.7098582 | 128.0867874 | 204.4997589 | 219.5810409 | 153.2401867 | 0.002754245 | 0.635361551 | 0.001246457 | 3.228030555 | 1.553326978 |
| CNAG_01084 | UBC4 | ubiquitin-conjugating enzyme 4 | 117.0666858 | 103.6243693 | 118.4869166 | 211.8373165 | 160.5486985 | 161.9844843 | 0.002762699 | 0.638781946 | 0.00125062 | 3.227076592 | 1.557014031 |
| CNAG_00529 | - | sulfite transporter | 308.6124158 | 354.5439854 | 339.215018 | 451.7443287 | 439.2076092 | 433.9565708 | 0.002764397 | 0.387602753 | 0.001251726 | 3.226823582 | 1.308217804 |
| CNAG_05772 | - | hypothetical protein | 99.71459136 | 120.7969039 | 125.9293976 | 203.1163369 | 170.1040381 | 163.098537 | 0.002767166 | 0.625115911 | 0.001253318 | 3.226459948 | 1.542334737 |
| CNAG_01053 | - | nuclear pore complex protein Nup160 | 3388.564906 | 3030.797414 | 3031.909532 | 2584.430088 | 2829.14216 | 2716.651616 | 0.002774958 | -0.232862966 | 0.001257186 | -3.225577996 | 0.850944553 |
| CNAG_12947 | - | unspecified product | 6.119309206 | 1.250466064 | 4.40102021 | 19.19432243 | 23.8376228 | 16.50958257 | 0.002782477 | 2.379682098 | 0.001260932 | 3.224726218 | 5.204220529 |
| CNAG_12047 | - | unspecified product | 5.006152917 | 3.4075280l5 | 5.462407518 | 21.65822481 | 20.1606897 | 19.74959258 | 0.002795689 | 2.190025975 | 0.001267261 | 3.223292481 | 4.563137019 |
| CNAG_00443 | - | hypothetical protein | 211.5418861 | 180.1012896 | 198.4630672 | 274.2634775 | 272.7497795 | 287.322564 | 0.00279765 | 0.484322556 | 0.001268491 | 3.223014516 | 1.398928819 |
| CNAG_00599 | - | anion transporter | 635.4021441 | 599.1570319 | 606.8949097 | 735.6022054 | 817.9452628 | 757.0375996 | 0.002872051 | 0.311999463 | 0.001302927 | 3.215334321 | 1.24142703 |
| CNAG_05591 | - | hypothetical protein, variant | 268.9846629 | 311.4849742 | 339.1780161 | 414.3368161 | 461.8278378 | 379.6356832 | 0.002872051 | 0.434828206 | 0.001302915 | 3.21533695 | 1.351749859 |
| CNAG_00682 | - | kinesin | 3496.366819 | 3527.56141 | 3376.458233 | 3949.935397 | 4080.894922 | 3947.677525 | 0.002880167 | 0.188487755 | 0.001306961 | 3.214446974 | 1.139568584 |
| CNAG_03640 | - | hypothetical protein | 1094.166769 | 1203.405476 | 1206.082812 | 1017.872992 | 957.2881409 | 964.8464638 | 0.002882776 | -0.268643668 | 0.001308497 | -3.214109724 | 0.830099587 |
| CNAG_00685 | - | pre-mRNA-splicing ATP-dependent RNA helicase PRP28 | 1019.628408 | 995.6463241 | 1007.876524 | 1093.395775 | 1436.520434 | 1271.014688 | 0.002889453 | 0.315359978 | 0.00131188 | 3.213368146 | 1.244322095 |
| CNAG_00982 | - | hypothetical protein | 1236.57724 | 1256.239887 | 1326.588101 | 1089.431148 | 1053.038101 | 1090.26435 | 0.002890581 | -0.255589256 | 0.001312745 | -3.213178798 | 0.837644936 |
| CNAG_05089 | - | hypothetical protein, variant | 5.155955835 | 13.06353886 | 9.754360513 | 26.81710362 | 40.42938629 | 25.40543969 | 0.002893521 | 1.739410661 | 0.001314434 | 3.2128096 | 3.338987427 |
| CNAG_06512 | - | pirin | 209.1750308 | 166.1158538 | 186.7441174 | 261.7804332 | 302.3039129 | 249.7054962 | 0.002924143 | 0.5171612 | 0.001328701 | 3.209707415 | 1.431136419 |
| CNAG_05892 | SHB17 | Phosphoglycerate mutase | 2820.001533 | 2722.645442 | 2880.427407 | 2218.566192 | 2487.550564 | 2541.197034 | 0.002924842 | -0.232448489 | 0.001329376 | -3.209561498 | 0.851189059 |
| CNAG_04890 | - | hypothetical protein | 4.901663772 | 0.118969316 | 2.257970959 | 12.70483358 | 17.6729318 | 17.44721979 | 0.00292527 | 2.91946518 | 0.001329928 | 3.209442185 | 7.565655998 |
| CNAG_04521 | - | oxidoreductase | 6.328780019 | 14.14256114 | 11.87594712 | 32.50575341 | 30.48061523 | 34.08422924 | 0.002933194 | 1.617750509 | 0.001333889 | 3.208587166 | 3.068961419 |
| CNAG_03494 | - | hypothetical protein | 869.7971459 | 856.4897224 | 832.8608894 | 691.6437902 | 705.4389024 | 720.5832733 | 0.002933531 | -0.288354489 | 0.0013344 | -3.208476899 | 0.818835475 |
| CNAG_04758 | AMT2 | amt family ammonium transporter | 159.0731068 | 177.8307673 | 121.7817021 | 244.1663021 | 216.6606936 | 226.2948347 | 0.002936902 | 0.571616654 | 0.001336292 | 3.208069477 | 1.486188029 |
| CNAG_07361 | - | hypothetical protein | 1739.844201 | 1664.619772 | 1853.406479 | 1464.514766 | 1563.762933 | 1449.992579 | 0.002945008 | -0.247403299 | 0.00134034 | -3.207199535 | 0.842411304 |
| CNAG_01128 | - | Hydroxyacylglutathione hydrolase | 1496.871974 | 1429.706086 | 1405.557434 | 1262.247629 | 1208.308842 | 1220.169186 | 0.002960419 | -0.246105364 | 0.001347716 | -3.2056206 | 0.843169529 |
| CNAG_04272 | CMK2 | CAMK/CAMK1 protein kinase | 2449.826341 | 2289.544244 | 2580.73146 | 2064.589401 | 2146.829257 | 2097.185653 | 0.002963403 | -0.230181659 | 0.001349436 | -3.205253458 | 0.852527538 |
| CNAG_03021 | - | hypothetical protein | 6.556491781 | 37.85314283 | 40.62498658 | 77.92881866 | 80.82275541 | 60.91638013 | 0.002968956 | 1.374359706 | 0.00135269 | 3.204560254 | 2.592528243 |
| CNAG_01070 | - | class II aldolase/adducin family protein | 29.79400383 | 46.48807124 | 43.87843041 | 103.0519453 | 113.8761946 | 44.65215028 | 0.002968956 | 1.124917463 | 0.001352522 | 3.204595928 | 2.180890692 |
| CNAG_03417 | - | hypothetical protein | 220.9723247 | 253.2367427 | 186.8066964 | 302.2465332 | 328.9379533 | 301.8322054 | 0.002973466 | 0.483927118 | 0.001355108 | 3.204046128 | 1.39854543 |
| CNAG_03839 | - | CCR4-NOT transcription complex subunit 4 | 971.2777826 | 910.3538176 | 833.9620689 | 776.5989547 | 694.1491064 | 741.6986497 | 0.002989494 | -0.310989951 | 0.001362777 | -3.202420976 | 0.806088446 |
| CNAG_01297 | - | hypothetical protein, variant 2 | 747.1364417 | 747.6336003 | 680.3756092 | 578.79088 | 637.3897029 | 526.4240258 | 0.00299554 | -0.335815887 | 0.001365899 | -3.201761757 | 0.79233592 |
| CNAG_13183 | - | unspecified product | 1.592458139 | 4.506872546 | 10.75857615 | 43.21586037 | 15.49524846 | 17.71985676 | 0.003011784 | 2.307720176 | 0.001373674 | 3.200126293 | 4.95100078 |
| CNAG_13097 | - | unspecified product | 117.9785247 | 65.7204348 | 74.57849062 | 35.07942385 | 49.78527161 | 46.85249028 | 0.003018982 | -0.995269323 | 0.001377325 | -3.199361136 | 0.501642219 |
| CNAG_02030 | - | glyoxal oxidase | 2017.76964 | 1537.637496 | 1379.09564 | 2207.600268 | 2107.390072 | 2044.830959 | 0.003034354 | 0.349814315 | 0.001384709 | 3.197819601 | 1.274396593 |
| CNAG_05526 | - | ino eighty subunit 2 | 626.0105543 | 827.2769435 | 760.2862915 | 599.8524485 | 599.3492461 | 544.1680187 | 0.003042974 | -0.359522704 | 0.001389014 | -3.196924226 | 0.779422398 |
| CNAG_00537 | - | Carnitine O-acetyltransferase | 264.2113956 | 279.1622067 | 289.0828196 | 360.557897 | 394.4610388 | 365.1244733 | 0.00305142 | 0.412337479 | 0.001393242 | 3.196047413 | 1.330840318 |

| | | | | | | | | | | | | | |
|---|---|---|---|---|---|---|---|---|---|---|---|---|---|
| CNAG_05756 | - | mitochondrial protein | 1363.86386 | 1383.378964 | 1469.490172 | 1216.31436 | 1127.182932 | 1225.686247 | 0.003067129 | -0.255559297 | 0.001400789 | -3.194488397 | 0.837662331 |
| CNAG_04633 | - | hypothetical protein | 12.11759727 | 18.43958306 | 8.737515442 | 51.17067006 | 33.87973244 | 28.80418768 | 0.003098911 | 1.546094573 | 0.001415683 | 3.191434337 | 2.920255447 |
| CNAG_01309 | - | arf/Sar family protein | 562.7404175 | 568.8217668 | 609.9054048 | 451.1108381 | 493.3038213 | 440.9008737 | 0.003115076 | -0.34484651 | 0.001423448 | -3.189853777 | 0.787391744 |
| CNAG_00248 | - | ESCRT-II complex subunit VPS36 | 332.0455612 | 426.7030484 | 397.860954 | 512.4131347 | 498.7030817 | 513.8534635 | 0.003116012 | 0.384698362 | 0.001424256 | 3.189689707 | 1.305586787 |
| CNAG_03220 | - | hypothetical protein | 823.1369431 | 815.5647644 | 859.4785701 | 689.3554777 | 639.8071062 | 719.4470047 | 0.003130181 | -0.301864382 | 0.001431115 | -3.18830105 | 0.811203407 |
| CNAG_07894 | - | hypothetical protein | 0.123388374 | 0.059988009 | 0.05182 | 9.993499131 | 5.247238613 | 6.542412676 | 0.003131013 | 5.164842011 | 0.001431877 | 3.188146981 | 35.8733858 |
| CNAG_12294 | - | unspecified product | 51.57237302 | 30.42735294 | 40.68157613 | 80.73659146 | 76.17391119 | 81.85660678 | 0.003137939 | 0.953089278 | 0.001435811 | 3.187353581 | 1.936013856 |
| CNAG_01007 | - | C2 domain-containing protein | 720.7341613 | 807.028207 | 690.1092122 | 1009.891432 | 928.2567261 | 851.4811972 | 0.003137939 | 0.315904123 | 0.001435641 | 3.187387978 | 1.244791507 |
| CNAG_04561 | - | monooxygenase | 455.4345533 | 328.8163815 | 351.0067509 | 502.2042355 | 463.2086635 | 626.8311674 | 0.003154302 | 0.47241624 | 0.001443684 | 3.18577194 | 1.387431207 |
| CNAG_05104 | FRK101 | CAMK protein kinase, variant | 32.03693036 | 40.03693006 | 45.96075836 | 67.18362709 | 84.14724961 | 77.43657929 | 0.003172975 | 0.947563748 | 0.001453005 | 3.183909376 | 1.928613094 |
| CNAG_12038 | - | unspecified product | 23.65194249 | 17.44866627 | 18.30211514 | 52.19476968 | 43.38456073 | 44.18800475 | 0.003172975 | 1.245021297 | 0.001452778 | 3.183954526 | 2.370220531 |
| CNAG_12243 | - | unspecified product | 9.828363187 | 15.24856566 | 16.12799708 | 37.16568334 | 37.45990531 | 36.37530196 | 0.00318034 | 1.445205297 | 0.001456766 | 3.183160977 | 2.723015693 |
| CNAG_02882 | - | hypothetical protein | 661.3045158 | 807.0387787 | 787.0934382 | 988.7803058 | 929.0980741 | 909.1518976 | 0.00318525 | 0.311090946 | 0.001459404 | 3.182637181 | 1.240645504 |
| CNAG_01927 | - | hypothetical protein | 455.2194018 | 423.3966131 | 477.6532841 | 339.7803027 | 344.8340003 | 367.5588873 | 0.003185678 | -0.383443618 | 0.001459989 | -3.182521029 | 0.766605568 |
| CNAG_06342 | - | SUMO activating enzyme | 922.3097001 | 884.5092492 | 855.2655523 | 762.5786427 | 729.5834726 | 711.7503859 | 0.003189206 | -0.288513064 | 0.001461995 | -3.182123399 | 0.818745477 |
| CNAG_02542 | IRK2 | fructosamine kinase | 3590.44957 | 912.7604863 | 1179.508781 | 516.4837102 | 420.1948793 | 525.6359619 | 0.003198446 | -1.980216449 | 0.001466622 | -3.181208217 | 0.253451841 |
| CNAG_07703 | - | hypothetical protein | 296.8962997 | 297.5029553 | 308.3004595 | 340.752754 | 513.7911506 | 411.7320294 | 0.003212189 | 0.472687795 | 0.001473708 | 3.179811692 | 1.387692385 |
| CNAG_01231 | - | agmatinase | 711.4943171 | 844.7539592 | 811.6346677 | 916.0848498 | 963.5761727 | 1111.00924 | 0.003212189 | 0.323258777 | 0.00147342 | 3.179868277 | 1.25115348 |
| CNAG_07611 | - | hypothetical protein | 0.277833449 | 6.571718795 | 1.207936555 | 29.55400999 | 18.45618369 | 7.885442417 | 0.003244356 | 2.938308078 | 0.001488862 | 3.176845752 | 7.665118405 |
| CNAG_09007 | - | NADH dehydrogenase subunit 6 | 50802.27877 | 37459.04721 | 36065.30752 | 28453.09154 | 28641.94739 | 29274.02212 | 0.00324441 | -0.542102512 | 0.001489283 | -3.176763777 | 0.686769316 |
| CNAG_10124 | - | unspecified product | 2.788049852 | 13.01628093 | 4.444554495 | 27.09507051 | 33.44098054 | 19.91054139 | 0.003276266 | 2.045175116 | 0.001504306 | 3.173851612 | 4.127233655 |
| CNAG_05315 | - | taurine catabolism dioxygenase TauD | 90.44205073 | 126.1559702 | 118.4876137 | 222.8943245 | 157.9366837 | 158.67813 | 0.003277913 | 0.671064055 | 0.001506263 | 3.173474182 | 1.592246891 |
| CNAG_12776 | - | unspecified product | 189.0428601 | 220.6784851 | 205.712869 | 145.2182165 | 118.6629276 | 155.525462 | 0.003277913 | -0.569070184 | 0.001506155 | -3.17349496 | 0.674051074 |
| CNAG_12918 | - | unspecified product | 29.56318842 | 29.2654357 | 24.71577625 | 58.87318945 | 77.52591386 | 45.5100456 | 0.003277913 | 1.118542658 | 0.001505832 | 3.173557264 | 2.171275298 |
| CNAG_02896 | ERG130 | hydroxymethylglutaryl-CoA synthase | 4131.330216 | 3621.275542 | 4223.068364 | 3292.684325 | 3505.361472 | 3500.408583 | 0.00328907 | -0.233784441 | 0.001511791 | -3.172410543 | 0.850401212 |
| CNAG_12262 | - | unspecified product | 0.164661057 | 1.176880724 | 0.068917526 | 9.73640074 | 10.65101302 | 9.803615992 | 0.003315139 | 4.687908229 | 0.001524178 | 3.170040202 | 25.77513769 |
| CNAG_03654 | - | ATP-dependent DNA helicase | 1235.674744 | 1271.440498 | 1295.790019 | 1508.496486 | 1462.10025 | 1565.197167 | 0.003331447 | 0.239651697 | 0.001532083 | 3.168536831 | 1.180707575 |
| CNAG_00888 | CNB1 | calcineurin subunit B | 1606.657349 | 1632.21788 | 1469.568541 | 1386.803333 | 1234.119295 | 1358.912125 | 0.003348681 | -0.25766857 | 0.001540418 | -3.166959521 | 0.836438532 |
| CNAG_01281 | - | hypothetical protein | 329.3557532 | 395.2950686 | 444.5082847 | 303.3093982 | 308.6781938 | 241.1914346 | 0.00336308 | -0.470170907 | 0.001547452 | -3.165634266 | 0.721879076 |
| CNAG_12904 | - | unspecified product | 23.72874544 | 21.74394247 | 23.61255013 | 53.57829584 | 56.54920082 | 43.19270193 | 0.003364687 | 1.166025761 | 0.001548602 | 3.165418123 | 2.243927029 |
| CNAG_12575 | - | unspecified product | 1.503337066 | 2.313862463 | 5.434724652 | 15.97367572 | 9.424201463 | 28.15729424 | 0.003367188 | 2.686651939 | 0.001550165 | 3.165124723 | 6.438175653 |
| CNAG_09009 | - | Cytochrome c oxidase subunit I | 468966.6562 | 413129.5936 | 408010.1242 | 362287.657 | 365905.1654 | 398144.8419 | 0.003398966 | -0.211525544 | 0.00156521 | -3.162313517 | 0.86362353 |
| CNAG_06538 | - | monocarboxylic acid transporter | 470.6502075 | 596.7181976 | 553.3959361 | 405.1479166 | 443.2447916 | 413.1324534 | 0.003413844 | -0.3749885 | 0.001572478 | -3.160964322 | 0.771111559 |
| CNAG_00064 | - | COPII-coated vesicle protein | 1997.875524 | 1956.566511 | 1904.661164 | 1757.284337 | 1643.036955 | 1669.780416 | 0.003427164 | -0.223705193 | 0.001579032 | -3.15975259 | 0.85636326 |
| CNAG_06780 | - | hypothetical protein | 884.4420083 | 1218.592301 | 1162.47733 | 1373.913749 | 1324.982211 | 1441.922365 | 0.003432481 | 0.32838092 | 0.001581901 | 3.159223638 | 1.255603469 |
| CNAG_12348 | - | unspecified product | 114.0260677 | 127.9134677 | 91.69445327 | 49.86219938 | 65.27282567 | 77.81650919 | 0.003438763 | -0.811985266 | 0.001585216 | -3.158613501 | 0.569597506 |
| CNAG_06009 | - | cyclohydrolase | 61.04715786 | 70.11962874 | 39.70118895 | 97.64960686 | 93.31856657 | 122.7530868 | 0.003438907 | 0.862115132 | 0.001585702 | 3.158524143 | 1.817701286 |
| CNAG_06875 | - | allantoate permease | 236.7788626 | 266.9625762 | 211.1286404 | 141.0727529 | 142.8384266 | 200.9421745 | 0.003446816 | -0.576040822 | 0.00158977 | -3.157777237 | 0.670802131 |
| CNAG_04456 | - | hypothetical protein | 555.502635 | 515.9969501 | 477.7458118 | 451.3696598 | 389.6809044 | 344.4033412 | 0.003449129 | -0.401301632 | 0.001591258 | -3.157504522 | 0.757174835 |
| CNAG_03876 | - | Ras family protein | 862.0638931 | 946.0475705 | 943.8657119 | 1211.414428 | 1053.216819 | 1114.512108 | 0.003451417 | 0.281410038 | 0.001592735 | 3.157234026 | 1.215382175 |
| CNAG_07453 | - | hypothetical protein | 2102.841961 | 2154.712422 | 1747.938796 | 1416.029212 | 1412.588114 | 1921.618867 | 0.003452726 | -0.352520064 | 0.001594182 | -3.156969139 | 0.783214802 |
| CNAG_06354 | - | hypothetical protein, variant | 210.7004305 | 310.3236401 | 270.9736081 | 406.0746942 | 354.654216 | 351.8175265 | 0.003452726 | 0.478158937 | 0.001594155 | 3.156974105 | 1.392964931 |
| CNAG_12585 | - | unspecified product | 26.11360078 | 27.12280819 | 28.93921008 | 70.27643688 | 46.63856393 | 59.66406825 | 0.003454516 | 1.096592987 | 0.00159543 | 3.156740891 | 2.13849078 |
| CNAG_07765 | - | hypothetical protein | 4.936633267 | 4.442582186 | 0.110250144 | 19.39987073 | 16.68568328 | 15.36622334 | 0.003467776 | 2.629569385 | 0.001601978 | 3.155546359 | 6.188412584 |
| CNAG_06273 | - | hypothetical protein | 742.6743182 | 803.6931844 | 880.757983 | 587.1868302 | 658.858753 | 702.7795671 | 0.003470657 | -0.33178125 | 0.001603733 | -3.155227001 | 0.794554866 |
| CNAG_01956 | - | hypothetical protein | 713.2482931 | 602.3073803 | 743.1867248 | 530.4864331 | 544.9778196 | 560.733807 | 0.003472906 | -0.348862211 | 0.001605196 | -3.154960963 | 0.785203108 |
| CNAG_01472 | - | endoplasmic reticulum protein | 957.43789 | 929.7932742 | 1014.10722 | 785.8147674 | 801.1814357 | 829.3674808 | 0.003474066 | -0.279438779 | 0.001606156 | -3.154786471 | 0.823911464 |
| CNAG_00732 | CCD3 | hypothetical protein | 527.4972742 | 500.9023179 | 445.7703085 | 384.9887941 | 363.7765354 | 399.7550803 | 0.003493607 | -0.376351128 | 0.001615617 | -3.153072369 | 0.770383587 |
| CNAG_06068 | - | chromatin structure-remodeling complex subunit RSC1/2 | 1993.477058 | 1994.384398 | 1927.150155 | 2239.227579 | 2337.657926 | 2332.357002 | 0.003496226 | 0.208914832 | 0.001617255 | 3.152776614 | 1.155818472 |
| CNAG_00487 | - | hypothetical protein | 32.10795327 | 50.77053518 | 52.35768917 | 78.15877785 | 93.63127531 | 83.03512156 | 0.003498245 | 0.908259077 | 0.001618616 | 3.152530997 | 1.876779392 |
| CNAG_05742 | STP1 | putative site-2 protease | 729.6868017 | 703.5206727 | 747.5031025 | 570.1819179 | 604.5327496 | 608.46018 | 0.003531837 | -0.305606671 | 0.00163459 | -3.149662741 | 0.809101908 |
| CNAG_07941 | - | hypothetical protein, variant | 5324.899993 | 5877.285743 | 5163.768338 | 4732.596716 | 4970.885945 | 4707.180605 | 0.003541237 | -0.198324659 | 0.001639373 | -3.148808922 | 0.871562084 |
| CNAG_03338 | SAF2 | hypothetical protein, variant | 854.676356 | 837.1274251 | 909.5914826 | 736.8760836 | 677.7749823 | 731.6846997 | 0.003543128 | -0.293497246 | 0.001640682 | -3.14857582 | 0.815921778 |
| CNAG_01355 | - | thioredoxin | 1775.926566 | 1654.893667 | 1602.865094 | 1389.779601 | 1431.733603 | 1479.90621 | 0.003561065 | -0.24228482 | 0.001649422 | -3.14702258 | 0.845405369 |
| CNAG_07990 | - | hypothetical protein | 311.1606013 | 204.7740265 | 278.1923477 | 183.3115419 | 160.9582516 | 197.7273147 | 0.003563759 | -0.571094356 | 0.001651105 | -3.146724348 | 0.67310601 |
| CNAG_05456 | - | hypothetical protein | 907.2580171 | 951.2935586 | 938.4209757 | 748.4322932 | 792.5848395 | 796.0682878 | 0.00357043 | -0.27440098 | 0.001654632 | -3.146100352 | 0.826793539 |
| CNAG_07773 | - | hypothetical protein | 204.7562731 | 243.591673 | 249.6161724 | 348.4271876 | 312.4005625 | 304.0933425 | 0.003575973 | 0.455217707 | 0.001658074 | 3.14549251 | 1.370989676 |
| CNAG_01316 | RPA32 | replication factor A2 | 1253.134585 | 1217.580953 | 1243.543212 | 1479.713884 | 1491.49422 | 1446.475652 | 0.003575973 | 0.23475599 | 0.001657885 | 3.145525975 | 1.176707698 |
| CNAG_02933 | - | quinone oxidoreductase | 471.6326146 | 479.3794034 | 506.4569214 | 363.1203678 | 426.9445345 | 337.7150518 | 0.003596482 | -0.386359379 | 0.001668023 | -3.143742254 | 0.765057784 |
| CNAG_12349 | - | unspecified product | 15.63934245 | 19.56334926 | 20.399562 | 44.96310656 | 39.01063287 | 48.47526295 | 0.003597555 | 1.264084213 | 0.001669399 | 3.143500867 | 2.401747045 |
| CNAG_06155 | - | hypothetical protein | 2099.540458 | 2099.879677 | 2179.772767 | 1801.519896 | 1875.981993 | 1882.900144 | 0.003597555 | -0.213611857 | 0.001669348 | -3.143509828 | 0.862375527 |
| CNAG_03694 | CFT3 | iron ion transporter | 1015.904147 | 1149.495975 | 1044.023079 | 880.7841264 | 911.6187597 | 899.3301715 | 0.003609984 | -0.268787406 | 0.001676048 | -3.14233735 | 0.830016887 |
| CNAG_00318 | - | arp2/3 complex 16 kda subunit | 1929.068624 | 1988.888348 | 2013.410515 | 1718.2937 | 1766.453482 | 1678.671045 | 0.003609984 | -0.215322555 | 0.001675683 | -3.142401072 | 0.861353558 |
| CNAG_04690 | - | acylpyruvate hydrolase, variant | 1366.08567 | 1400.554739 | 1210.435916 | 1138.474426 | 1101.348958 | 1111.375959 | 0.003628511 | -0.262114241 | 0.001685093 | -3.140761291 | 0.833865011 |
| CNAG_06532 | - | hypothetical protein | 404.2022607 | 386.8995904 | 371.2337358 | 479.5714372 | 490.0559493 | 541.5348169 | 0.003631385 | 0.363947004 | 0.001686871 | 3.140452417 | 1.286941972 |
| CNAG_01216 | - | 6-phosphogluconolactonase | 785.50332 | 713.1685528 | 616.4311168 | 574.1673845 | 577.79266 | 524.1804307 | 0.003634458 | -0.351366056 | 0.001688743 | -3.140127578 | 0.783841544 |
| CNAG_11003 | - | unspecified product | 133.4791702 | 131.617773 | 157.8993753 | 182.1764296 | 223.7433973 | 223.9802442 | 0.003652297 | 0.561420281 | 0.001697477 | 3.138616028 | 1.475721296 |
| CNAG_02422 | - | hypothetical protein | 607.0294275 | 580.6805892 | 622.7093278 | 546.3806011 | 452.6324773 | 409.8965899 | 0.003669033 | -0.378083431 | 0.001705704 | -3.137198862 | 0.769459111 |
| CNAG_01765 | - | hypothetical protein, variant | 1110.83026 | 1288.651085 | 1278.718265 | 1476.601274 | 1451.779252 | 1506.372347 | 0.003682436 | 0.255477921 | 0.001712384 | 3.136052684 | 1.193731124 |
| CNAG_05123 | - | hypothetical protein | 584.2661918 | 802.663034 | 637.8527284 | 900.8341583 | 834.1950713 | 868.0187359 | 0.003726802 | 0.34939554 | 0.00173347 | 3.132461721 | 1.274026724 |
| CNAG_06015 | - | hypothetical protein | 14.71626371 | 41.07984667 | 45.93611559 | 74.55509576 | 80.72549008 | 67.53103966 | 0.003727433 | 1.130883898 | 0.001734674 | 3.132257918 | 2.189928698 |
| CNAG_00535 | - | large subunit ribosomal protein L16 | 1569.350986 | 1656.994183 | 1518.60989 | 1426.534355 | 1249.648217 | 1353.375072 | 0.003727433 | -0.250533432 | 0.001734321 | -3.132317616 | 0.840585554 |
| CNAG_06361 | - | 26S proteasome regulatory subunit N6 | 5598.243199 | 4970.296625 | 5204.352925 | 6117.512337 | 6258.871373 | 5961.520391 | 0.003735915 | 0.201456242 | 0.001739078 | 3.1315135 | 1.149858425 |
| CNAG_05552 | - | transcription activator snf2l1 | 8670.434431 | 8536.452382 | 8190.352479 | 7201.28295 | 7744.016597 | 7716.7783 | 0.003766091 | -0.179687025 | 0.001753585 | -3.129073417 | 0.882894509 |
| CNAG_04541 | - | cleavage stimulation factor subunit 2 | 434.3108504 | 504.0638702 | 438.2691909 | 364.7840342 | 377.6968569 | 324.3803867 | 0.003793679 | -0.384359896 | 0.001766894 | -3.126851138 | 0.766118839 |
| CNAG_01298 | - | lysophospholipase NTE1 | 3486.936109 | 3173.118591 | 3004.298051 | 3642.458699 | 3796.22916 | 3998.656645 | 0.003796764 | 0.227531524 | 0.001768794 | 3.126535044 | 1.170829924 |
| CNAG_06617 | - | hypothetical protein | 1591.889464 | 1742.261085 | 1773.5465 | 2030.749118 | 1970.161782 | 2029.281028 | 0.003803293 | 0.224825652 | 0.0017723 | 3.125952756 | 1.168636012 |
| CNAG_13006 | - | unspecified product | 54.23698124 | 75.53458893 | 73.69348913 | 101.3535487 | 122.8737993 | 121.8196348 | 0.003829217 | 0.753668433 | 0.001784848 | 3.123877425 | 1.686074675 |
| CNAG_00429 | - | tRNA-dihydrouridine synthase 1 | 500.5370516 | 397.5736381 | 443.5706578 | 339.8089067 | 335.3340364 | 352.0406814 | 0.003850226 | -0.401288537 | 0.001795111 | -3.122189968 | 0.757181707 |
| CNAG_06530 | - | hypothetical protein | 251.4548107 | 273.803052 | 321.0343986 | 346.3900663 | 384.8864608 | 433.7655721 | 0.003881743 | 0.445771597 | 0.001810279 | 3.119712005 | 1.362042383 |
| CNAG_00986 | UBA4 | adenylyltransferase and sulfurtransferase | 990.1912262 | 957.834567 | 1194.256801 | 851.2177038 | 780.3789076 | 903.7280729 | 0.003884938 | -0.325243293 | 0.001812244 | -3.119392485 | 0.798163774 |
| CNAG_12936 | - | unspecified product | 6.304517985 | 9.875895608 | 13.96577803 | 30.97431731 | 31.45156519 | 27.55956976 | 0.003901056 | 1.63711416 | 0.001820239 | 3.118095389 | 3.110430261 |
| CNAG_03105 | - | hypothetical protein | 38.94174994 | 53.92841039 | 27.96015899 | 79.17996665 | 75.32315118 | 78.55222422 | 0.003923769 | 0.961859111 | 0.001831316 | 3.116306853 | 1.94781831 |
| CNAG_07657 | - | U3 small nucleolar RNA-associated protein 19 | 409.4511551 | 345.8078278 | 351.8763845 | 310.4455038 | 256.7804674 | 257.7642776 | 0.00392536 | -0.440861421 | 0.001832538 | -3.116110227 | 0.736694603 |
| CNAG_03821 | - | phosphatidylinositol 3-kinase | 739.3055749 | 710.1095873 | 668.7684283 | 847.6970642 | 933.5278041 | 843.6590942 | 0.003932261 | 0.293735951 | 0.00183624 | 3.115515061 | 1.225810486 |
| CNAG_06761 | - | siderophore-iron transporter Str1 | 91.35558459 | 92.77701892 | 74.8323445 | 129.4662188 | 129.5518824 | 157.264268 | 0.003943819 | 0.676570573 | 0.001842118 | 3.11457224 | 1.598335836 |
| CNAG_03734 | - | hypothetical protein, variant 2 | 199.9575512 | 224.1638398 | 175.0632747 | 302.4903261 | 249.3077527 | 299.5032334 | 0.003944328 | 0.492992934 | 0.001842838 | 3.11445706 | 1.407361483 |

| | | | | | | | | | | | | | |
|---|---|---|---|---|---|---|---|---|---|---|---|---|---|
| CNAG_05166 | - | helicase swr1 | 7710.508827 | 7522.581146 | 7072.71714 | 6141.985079 | 6752.068929 | 6783.126368 | 0.003980801 | -0.196172964 | 0.001860851 | -3.111586279 | 0.872862938 |
| CNAG_02131 | ISA1 | iron-sulfur cluster assembly protein | 324.6034869 | 374.8127275 | 377.4042868 | 283.8525546 | 275.8498595 | 256.6416576 | 0.003980801 | -0.416084382 | 0.001860405 | -3.111657054 | 0.749455964 |
| CNAG_02110 | - | peptide alpha-N-acetyltransferase | 1474.520581 | 1251.958951 | 1246.671816 | 1083.132071 | 1129.851496 | 1096.95034 | 0.00398132 | -0.279453981 | 0.00186158 | -3.111470686 | 0.823902782 |
| CNAG_03231 | - | hypothetical protein, variant | 628.3791681 | 764.8840707 | 883.8899878 | 591.2166997 | 593.26996 | 599.6150891 | 0.00401157 | -0.366327505 | 0.001876704 | -3.109080834 | 0.775754731 |
| CNAG_02032 | - | nuclear export factor | 3129.432106 | 3011.389607 | 2990.310985 | 2646.005425 | 2626.376861 | 2765.456658 | 0.00401157 | -0.199275392 | 0.001876596 | -3.10909782 | 0.870987916 |
| CNAG_13204 | - | unspecified product | 157.1847626 | 134.5981293 | 142.7944618 | 100.0858446 | 97.24939211 | 83.53362269 | 0.004016094 | -0.644880676 | 0.00187931 | -3.108670714 | 0.639545685 |
| CNAG_05334 | - | hypothetical protein, variant | 4.044371177 | 19.47990171 | 12.95249307 | 35.61855081 | 37.52254516 | 34.16928549 | 0.00404769 | 1.587106261 | 0.00189459 | 3.10627727 | 3.004461138 |
| CNAG_05972 | - | hypothetical protein | 31.97831868 | 33.59535538 | 42.75345832 | 65.75898414 | 69.31270059 | 74.06091356 | 0.00406161 | 0.957281938 | 0.001901601 | 3.105184848 | 1.941648348 |
| CNAG_13018 | - | unspecified product | 213.8489165 | 329.3999158 | 286.7431567 | 191.8564867 | 190.3827229 | 198.8788781 | 0.00406613 | -0.529621815 | 0.001904214 | -3.104778737 | 0.692736303 |
| CNAG_12618 | - | unspecified product | 14.7587884 | 38.99696674 | 61.86790869 | 96.68783438 | 94.57055924 | 67.6655041 | 0.00407485 | 1.171230671 | 0.001908795 | 3.104067874 | 2.252037219 |
| CNAG_03442 | - | phosphatidylinositol glycan, class T | 831.4529565 | 978.1783121 | 934.137089 | 740.6834524 | 695.8641273 | 807.1270597 | 0.004096499 | -0.305389684 | 0.001919437 | -3.102422706 | 0.809223609 |
| CNAG_04498 | - | alpha-1,2-mannosidase | 1383.709344 | 1424.306091 | 1430.059908 | 1171.079369 | 1270.498737 | 1191.309563 | 0.004135103 | -0.237526471 | 0.00193803 | -3.099568186 | 0.848198319 |
| CNAG_00242 | - | hypothetical protein | 7.445505484 | 6.678206353 | 13.96316421 | 40.98906774 | 18.9247974 | 30.78537267 | 0.004158463 | 1.743591262 | 0.001949486 | 3.097821821 | 3.348677084 |
| CNAG_02262 | - | ATP-binding cassette transporter | 3442.350146 | 3447.697785 | 3002.088894 | 2872.503014 | 2892.935467 | 2838.775979 | 0.004168684 | -0.216541126 | 0.001954787 | -3.097016988 | 0.860626324 |
| CNAG_07974 | - | hypothetical protein | 216.8224995 | 194.9319684 | 207.8405014 | 168.0369337 | 112.5856438 | 137.860014 | 0.004209549 | -0.580579455 | 0.001974463 | -3.094046838 | 0.668695144 |
| CNAG_05704 | - | ESCRT-II complex subunit VPS22 | 213.6504344 | 253.034186 | 256.8638158 | 178.7330642 | 143.6834555 | 186.6054583 | 0.00421473 | -0.524382909 | 0.001977408 | -3.093604668 | 0.695256432 |
| CNAG_12051 | - | unspecified product | 11.11605041 | 29.19533787 | 23.6083466 | 53.61228664 | 61.00664716 | 39.91470511 | 0.004216215 | 1.266945836 | 0.001978619 | 3.093422949 | 2.4065157 |
| CNAG_06019 | - | hypothetical protein, variant | 1448.995699 | 1380.147151 | 1388.480432 | 1128.228223 | 1128.911728 | 1297.785395 | 0.004254222 | -0.262097401 | 0.001996975 | -3.090681776 | 0.833874744 |
| CNAG_06492 | - | cytoplasmic protein | 718.1181422 | 756.2924744 | 818.9196099 | 563.0789961 | 620.0353065 | 671.6866161 | 0.004264762 | -0.321068354 | 0.002002443 | -3.089869684 | 0.800476884 |
| CNAG_01196 | - | tyrosyl-DNA phosphodiesterase 1 | 2525.533838 | 2224.865428 | 2123.312461 | 1842.695362 | 1951.859453 | 2038.305311 | 0.004284607 | -0.252651388 | 0.002012285 | -3.088413219 | 0.839352433 |
| CNAG_00505 | FZC28 | hypothetical protein | 747.5504959 | 787.6555938 | 719.9512179 | 922.4720964 | 866.9062075 | 992.292872 | 0.004292684 | 0.288459296 | 0.002016602 | 3.087776253 | 1.221335275 |
| CNAG_08026 | - | hypothetical protein | 252.2467541 | 181.2042111 | 187.8493637 | 294.531305 | 281.3343695 | 313.9261247 | 0.004300369 | 0.502068047 | 0.002021263 | 3.087090198 | 1.416242236 |
| CNAG_12141 | - | unspecified product | 29.82642741 | 65.75160516 | 44.9581365 | 82.06188545 | 97.94918067 | 87.46515108 | 0.004300369 | 0.934090844 | 0.00202082 | 3.08715523 | 1.910686181 |
| CNAG_12574 | - | unspecified product | 28.52330174 | 34.65669349 | 41.68862372 | 73.0495303 | 56.13315719 | 77.3148324 | 0.00434266 | 0.985488527 | 0.002041671 | 3.084102766 | 1.979983663 |
| CNAG_06372 | - | hypothetical protein | 114.6793644 | 97.15784899 | 112.083045 | 172.5131035 | 162.382704 | 159.7070233 | 0.004352457 | 0.594248277 | 0.002046808 | 3.083355001 | 1.509685752 |
| CNAG_02859 | POS5 | NADH kinase, variant 2 | 399.5901681 | 406.2829248 | 403.2014503 | 484.1904035 | 528.0746793 | 538.2540317 | 0.004355286 | 0.344469463 | 0.002048671 | 3.083084375 | 1.269683983 |
| CNAG_03523 | - | solute carrier family 25, member 38, variant | 633.968065 | 647.4631321 | 663.2528102 | 558.6968754 | 475.8829118 | 534.088226 | 0.004371291 | -0.325298109 | 0.002056733 | -3.081915443 | 0.798133448 |
| CNAG_12852 | - | unspecified product | 47.13000375 | 43.31863158 | 61.92548949 | 99.55590847 | 86.48230202 | 87.51092378 | 0.004400557 | 0.833348426 | 0.00207104 | 3.079851271 | 1.781816076 |
| CNAG_00922 | - | hypothetical protein | 814.9077259 | 753.1058224 | 827.4835992 | 658.2120461 | 638.1226307 | 677.2809782 | 0.004410398 | -0.294562545 | 0.00207621 | -3.079108594 | 0.815319517 |
| CNAG_13087 | - | unspecified product | 17.95901058 | 20.6479168 | 22.52833749 | 37.50071655 | 48.70120392 | 53.96609406 | 0.004444322 | 1.210999073 | 0.002092722 | 3.076747845 | 2.314978947 |
| CNAG_00500 | PTC1 | Protein phosphatase | 554.8578114 | 591.5482961 | 490.7007227 | 712.4533658 | 662.5327451 | 693.72674 | 0.004449662 | 0.323515166 | 0.002095781 | 3.076312503 | 1.251375849 |
| CNAG_10067 | - | unspecified product | 0.18521499 | 2.256896189 | 0.077388533 | 7.045775462 | 10.5795531 | 17.25796708 | 0.004483435 | 4.076960063 | 0.002112235 | 3.073980137 | 16.87668989 |
| CNAG_08019 | - | hypothetical protein | 244.1767786 | 209.1665345 | 205.9565619 | 313.2718471 | 311.6231057 | 285.2279978 | 0.00450381 | 0.45040634 | 0.002122384 | 3.072549834 | 1.366425061 |
| CNAG_00588 | - | hypothetical protein | 25.9705567 | 24.92387082 | 11.96429935 | 40.66839196 | 64.61658881 | 43.15049082 | 0.004513225 | 1.277346825 | 0.002127923 | 3.071771824 | 2.423927965 |
| CNAG_03992 | - | GTP-binding protein 1, variant | 794.33107 | 919.072816 | 822.3201707 | 1065.757376 | 1053.340882 | 982.4525028 | 0.004513225 | 0.275599454 | 0.002127819 | 3.071786496 | 1.210496958 |
| CNAG_02490 | MRE11 | DNA repair protein MRE11 | 966.8564356 | 1055.790558 | 1034.395134 | 827.0598135 | 811.4858336 | 913.6958634 | 0.004533129 | -0.274879713 | 0.002137861 | -3.07038063 | 0.826519227 |
| CNAG_05324 | - | Sugar transporter | 2.716440367 | 7.65848995 | 4.40877613 | 26.73013463 | 16.53613453 | 17.61103233 | 0.004539223 | 2.170098424 | 0.00214129 | 3.069902088 | 4.500540966 |
| CNAG_04330 | - | hypothetical protein | 28.3691235 | 16.43141036 | 31.03933572 | 61.46088422 | 68.79324995 | 41.09341305 | 0.004564572 | 1.162892076 | 0.002153805 | 3.068161116 | 2.239058273 |
| CNAG_02715 | ALG12 | alpha-1,6-mannosyltransferase | 754.619095 | 810.2999649 | 834.0104546 | 944.9681577 | 1005.053837 | 973.5144787 | 0.004577008 | 0.270026959 | 0.002160231 | 3.067270688 | 1.205830361 |
| CNAG_12193 | - | unspecified product | 0.118430101 | 0.05759837 | 0.049758096 | 10.91307181 | 6.184103792 | 3.354926199 | 0.00458035 | 5.090114319 | 0.002162368 | 3.066975171 | 34.06254484 |
| CNAG_12803 | - | unspecified product | 87.02145233 | 56.07566127 | 96.73471946 | 53.48989325 | 36.70352887 | 34.74034562 | 0.004594765 | -0.975422809 | 0.002169734 | -3.065958488 | 0.508590772 |
| CNAG_12615 | - | unspecified product | 33.10105933 | 32.51595042 | 38.51056066 | 62.60910441 | 68.47746974 | 71.84017622 | 0.004595457 | 0.949120591 | 0.002170623 | 3.065836113 | 1.930695424 |
| CNAG_02431 | - | TatD DNase | 838.3367179 | 860.795753 | 848.8446248 | 706.4694657 | 719.2620071 | 706.1709779 | 0.004603195 | -0.27186259 | 0.00217484 | -3.065255718 | 0.828249545 |
| CNAG_05450 | - | hypothetical protein | 346.0341514 | 397.6703041 | 440.4713179 | 493.5982237 | 539.3383843 | 513.8760104 | 0.004633163 | 0.370134473 | 0.002189564 | 3.063237221 | 1.292473296 |
| CNAG_00364 | - | tRNA threonylcarbamoyladenosine biosynthesis protein | 1228.478728 | 1315.491777 | 1421.478047 | 1109.628865 | 1153.158882 | 1095.846423 | 0.004648828 | -0.254960939 | 0.002197535 | -3.062149772 | 0.838009823 |
| CNAG_01899 | - | prefoldin alpha subunit | 662.8846433 | 498.9025846 | 665.3057525 | 543.275994 | 388.6675694 | 429.8360894 | 0.004652153 | -0.440652123 | 0.002199675 | -3.06185842 | 0.736801486 |
| CNAG_05467 | - | hypothetical protein | 6.15230542 | 4.479405358 | 5.467461613 | 24.14223347 | 19.22770593 | 18.69777026 | 0.004678446 | 2.007723914 | 0.002213249 | 3.060016318 | 4.021472665 |
| CNAG_00264 | - | nuclease I, variant | 811.7999416 | 901.8376385 | 775.4197285 | 1058.771217 | 940.1994836 | 1072.252215 | 0.004678446 | 0.289338587 | 0.002212948 | 3.060057081 | 1.222079879 |
| CNAG_07913 | - | hypothetical protein | 8.810926426 | 23.87715601 | 32.05976316 | 42.19598782 | 59.20351462 | 57.33676655 | 0.004700995 | 1.312746501 | 0.002224491 | 3.058498658 | 2.484140031 |
| CNAG_01739 | - | hypothetical protein, variant | 357.6106553 | 410.5438594 | 359.5026425 | 510.9396539 | 488.3739196 | 461.7622423 | 0.004710774 | 0.359080236 | 0.002229693 | 3.057798647 | 1.282607934 |
| CNAG_01413 | PUB1 | nucleolysin TIA-1/TIAR | 5412.706143 | 5326.917177 | 5888.97358 | 6065.220193 | 6908.450519 | 6417.773816 | 0.004713699 | 0.206266273 | 0.002231653 | 3.057535322 | 1.153698519 |
| CNAG_12660 | - | unspecified product | 19.29112381 | 38.89817666 | 36.37631632 | 69.99035857 | 63.26361879 | 61.95792465 | 0.004753146 | 1.039817961 | 0.00225091 | 3.054959409 | 2.055968216 |
| CNAG_12044 | - | unspecified product | 101.5953203 | 140.8792948 | 137.3666668 | 52.89508054 | 82.59556165 | 90.0263997 | 0.004816686 | -0.769054107 | 0.002282176 | -3.050819634 | 0.586802081 |
| CNAG_03503 | YOR1 | ATP-binding cassette transporter protein YOR1, variant | 1896.543696 | 1846.790191 | 1822.637337 | 2046.134626 | 2209.136002 | 2300.109162 | 0.004816686 | 0.221168923 | 0.002282119 | 3.050827225 | 1.165677678 |
| CNAG_07874 | - | Sugar transporter | 5.039981343 | 5.554433995 | 6.53092237 | 25.72447213 | 12.87351587 | 28.37600814 | 0.004818496 | 2.008077874 | 0.002283623 | 3.050629403 | 4.022459439 |
| CNAG_12897 | - | unspecified product | 0.153548226 | 1.16987787 | 0.064325477 | 8.930538652 | 7.003910698 | 11.88833705 | 0.004832266 | 4.581736208 | 0.002290738 | 3.049695065 | 23.94638887 |
| CNAG_05061 | - | hypothetical protein, variant | 59.25558088 | 68.69793989 | 61.70860244 | 29.80257279 | 21.96326302 | 42.27037227 | 0.004848484 | -1.044962645 | 0.002299019 | -3.048611134 | 0.484657457 |
| CNAG_01342 | - | hypothetical protein | 973.8928974 | 1074.113724 | 1084.501629 | 810.6354953 | 906.4681246 | 901.5230824 | 0.004849489 | -0.273667654 | 0.002300087 | -3.048471511 | 0.827213908 |
| CNAG_01939 | - | hypothetical protein | 41.12557782 | 21.82949238 | 38.51520117 | 67.41177298 | 85.20581269 | 56.55883874 | 0.004881545 | 1.038600121 | 0.002315888 | 3.046413948 | 2.054233418 |
| CNAG_03730 | - | DNA-directed RNA polymerase II subunit RPB11 | 971.3794842 | 922.238321 | 935.2320483 | 838.1331505 | 778.7499242 | 745.0721496 | 0.004890716 | -0.275839041 | 0.002320835 | -3.045772266 | 0.825969812 |
| CNAG_12581 | - | unspecified product | 9.705727196 | 7.737382418 | 10.79744928 | 38.57225621 | 19.84634874 | 27.52465041 | 0.004894134 | 1.676418383 | 0.002323055 | 3.045484794 | 3.196334471 |
| CNAG_04303 | - | E3 ubiquitin-protein ligase bre1 | 6422.664517 | 6992.480752 | 7160.102597 | 5891.443233 | 6105.118269 | 6284.662946 | 0.004906007 | -0.185484489 | 0.00232929 | -3.044678694 | 0.87935372 |
| CNAG_01525 | - | hypothetical protein | 131.8640656 | 141.0499085 | 173.5893742 | 102.4073544 | 95.48063854 | 93.47683161 | 0.004920761 | -0.633000081 | 0.002337497 | -3.043620673 | 0.644834089 |
| CNAG_05348 | CDC42 | cell division control protein 42 | 1138.68834 | 1083.989703 | 1121.969081 | 1410.577708 | 1342.262769 | 1260.012259 | 0.004920761 | 0.248108177 | 0.002336937 | 3.043692714 | 1.187648715 |
| CNAG_01952 | - | aryl-alcohol dehydrogenase, variant | 577.8890129 | 549.4630818 | 619.4980225 | 460.4711208 | 471.676444 | 475.2575394 | 0.004925127 | -0.327160997 | 0.002340172 | -3.043276491 | 0.797103519 |
| CNAG_01918 | - | cytoskeletal regulatory protein binding protein | 669.3342678 | 702.5525915 | 693.2594822 | 833.7275812 | 834.2081116 | 854.7051635 | 0.004938111 | 0.273512955 | 0.002346945 | 3.04240684 | 1.208747542 |
| CNAG_03583 | - | hypothetical protein | 951.7588066 | 1061.203259 | 1160.173923 | 845.7163451 | 904.7288093 | 882.6800791 | 0.004957491 | -0.284816293 | 0.002356761 | -3.0411504 | 0.820846126 |
| CNAG_05597 | - | transcription initiation factor TFIID subunit 9/adenylate kinase | 768.1119125 | 622.8002802 | 830.595557 | 644.366824 | 580.3413839 | 492.0637878 | 0.004966696 | -0.387982558 | 0.002361743 | -3.040514505 | 0.7641975 |
| CNAG_05659 | - | hypothetical protein | 4.874776735 | 1.217238932 | 1.186921738 | 17.9289313 | 14.07403157 | 11.01921507 | 0.004987189 | 2.789505813 | 0.002372097 | 3.03919701 | 6.913929116 |
| CNAG_12603 | - | unspecified product | 17.71588671 | 8.834356513 | 9.774050183 | 31.60291832 | 30.4374204 | 37.35657522 | 0.005002383 | 1.510282231 | 0.002379934 | 3.038203176 | 2.848657612 |
| CNAG_00541 | - | dimethylaniline monooxygenase | 147.413927 | 158.4707735 | 137.7118425 | 209.6764639 | 214.1430432 | 212.9761664 | 0.005006611 | 0.507817966 | 0.002382558 | 3.037871194 | 1.42189799 |
| CNAG_12671 | - | unspecified product | 35.53480648 | 46.47787397 | 50.22360886 | 82.25935688 | 84.03294765 | 75.32676077 | 0.005021961 | 0.864716688 | 0.002390476 | 3.036871227 | 1.820982033 |
| CNAG_12873 | - | unspecified product | 4.128571653 | 36.61123964 | 10.9004494 | 57.83241142 | 38.91453314 | 57.25028256 | 0.005044469 | 1.596970551 | 0.002401805 | 3.03544561 | 3.025074245 |
| CNAG_13008 | - | unspecified product | 1093.571132 | 848.9774808 | 756.1565767 | 700.1779928 | 747.7415276 | 658.5183694 | 0.005116654 | -0.373869747 | 0.002436799 | -3.031080927 | 0.771709758 |
| CNAG_05787 | HPI4 | symplekin | 5013.088901 | 5127.449285 | 4937.661502 | 4433.74756 | 4524.051298 | 4590.566118 | 0.005125849 | -0.16947882 | 0.002441804 | -3.030461364 | 0.889163838 |
| CNAG_03546 | - | hypothetical protein | 34.50123848 | 62.60396282 | 65.13441561 | 95.38724937 | 107.4132775 | 91.9717837 | 0.005128087 | 0.849863063 | 0.002443497 | 3.030252116 | 1.802329844 |
| CNAG_04300 | - | hypothetical protein | 544.4607205 | 636.7951543 | 582.3629925 | 733.3738075 | 710.8425651 | 751.4376003 | 0.00514743 | 0.301797732 | 0.002453342 | 3.029037487 | 1.23267949 |
| CNAG_12305 | - | unspecified product | 6.311354641 | 18.36615527 | 1.260935752 | 21.83591285 | 35.91761926 | 39.42556074 | 0.005150349 | 1.916500098 | 0.002455363 | 3.02878881 | 3.775061369 |
| CNAG_03508 | - | hypothetical protein | 639.7726548 | 649.606823 | 622.7663763 | 523.5670759 | 533.8102407 | 509.6984028 | 0.005150582 | -0.302602227 | 0.002456102 | -3.028697797 | 0.810788636 |
| CNAG_01288 | - | solute carrier family 25, member 33/36 | 1641.818125 | 1639.833983 | 1854.455819 | 1453.644988 | 1517.172995 | 1425.565143 | 0.005152003 | -0.239714094 | 0.002457409 | -3.028537085 | 0.846913132 |
| CNAG_00688 | - | hypothetical protein | 714.5088184 | 670.1434556 | 754.9456322 | 559.2864833 | 574.3065655 | 613.9831899 | 0.005154136 | -0.307728907 | 0.002459056 | -3.028334678 | 0.807912576 |
| CNAG_03132 | FZC5 | hypothetical protein | 20.47929225 | 42.13342154 | 41.6928622 | 67.3852626 | 76.37085676 | 66.40594412 | 0.005180508 | 1.00726193 | 0.002472271 | 3.026714827 | 2.010092553 |
| CNAG_03557 | - | transcription initiation factor TFIIA small subunit | 831.1638862 | 715.4138301 | 791.237005 | 662.1686517 | 606.1822511 | 642.88899 | 0.005196699 | -0.306863879 | 0.002480632 | -3.025694022 | 0.808397139 |
| CNAG_03170 | - | hypothetical protein | 575.6118361 | 594.6742594 | 622.7156959 | 507.295351 | 482.8835243 | 463.094504 | 0.005207371 | -0.317930599 | 0.002486362 | -3.024996223 | 0.802219757 |
| CNAG_07636 | CSR2 | protoplast regeneration and killer toxin resistance protein | 1564.942438 | 1609.735989 | 1554.953943 | 1931.982595 | 1804.539116 | 1811.729008 | 0.005210713 | 0.215453731 | 0.00248923 | 3.024647526 | 1.161069016 |
| CNAG_05941 | - | hypothetical protein | 28.41660982 | 28.19707079 | 31.06033099 | 58.07372873 | 56.32155247 | 62.95630415 | 0.005210713 | 1.001301571 | 0.002488609 | 3.024723045 | 2.001805175 |
| CNAG_03195 | - | hypothetical protein | 109.1802692 | 172.4465185 | 164.3081705 | 247.3883898 | 209.7503302 | 213.015068 | 0.005219268 | 0.57446041 | 0.002493955 | 3.024073946 | 1.489120406 |

| | | | | | | | | | | | | | |
|---|---|---|---|---|---|---|---|---|---|---|---|---|---|
| CNAG_01059 | - | vacuolar membrane protein, variant | 245.4543827 | 201.6850375 | 266.6590447 | 324.8204943 | 298.4937652 | 374.9081414 | 0.005222087 | 0.468430627 | 0.00249594 | 3.023833252 | 1.383603556 |
| CNAG_05100 | - | ENTH domain-containing protein c | 1374.417249 | 1451.244716 | 1534.535782 | 1132.843686 | 1336.953999 | 1216.844139 | 0.005270285 | -0.258022863 | 0.00251962 | -3.020975209 | 0.836233148 |
| CNAG_04158 | HIRA | Nucleosome assembly complex protein | 2381.054311 | 2577.157755 | 2702.305005 | 1999.853726 | 2226.176038 | 2345.791451 | 0.005293122 | -0.236440208 | 0.002532477 | -3.019433744 | 0.848837202 |
| CNAG_12130 | - | unspecified product | 24.84784259 | 16.40128855 | 26.77174125 | 43.06805932 | 51.2702337 | 54.04082275 | 0.005293122 | 1.148517599 | 0.0025324 | 3.019442899 | 2.2168599 |
| CNAG_06784 | - | hypothetical protein, variant | 1649.287027 | 2199.055195 | 2152.130031 | 2443.251852 | 2385.931144 | 2524.366492 | 0.005293122 | 0.279109847 | 0.002532077 | 3.019481575 | 1.213445948 |
| CNAG_12143 | - | unspecified product | 11.89228956 | 3.442157364 | 7.610649951 | 21.28945215 | 24.45132351 | 30.64305805 | 0.005297684 | 1.81844895 | 0.002535306 | 3.019095452 | 3.527018028 |
| CNAG_03609 | - | hypothetical protein | 534.5627047 | 496.6364302 | 511.8244056 | 375.5332349 | 353.3553649 | 466.2535854 | 0.005331844 | -0.383266206 | 0.002552305 | -3.01707041 | 0.766699845 |
| CNAG_03870 | - | delta8-fatty-acid desaturase | 869.8745511 | 907.1160803 | 877.6448452 | 750.8684894 | 707.9822519 | 766.0816661 | 0.005345035 | -0.270065563 | 0.002559273 | -3.016243965 | 0.829281859 |
| CNAG_12293 | - | unspecified product | 127.8161759 | 210.0814608 | 149.4595183 | 236.9272314 | 222.6516743 | 270.5415415 | 0.005351715 | 0.570077271 | 0.002563124 | 3.015787948 | 1.484603084 |
| CNAG_08029 | - | hypothetical protein | 452.1675016 | 490.2950316 | 463.9918786 | 543.2754094 | 627.3150282 | 611.5451932 | 0.005402845 | 0.327124083 | 0.002588272 | 3.012826043 | 1.254510098 |
| CNAG_12207 | - | unspecified product | 0.114623122 | 0.055762415 | 0.048173806 | 6.522800585 | 8.981117263 | 4.411035708 | 0.005408116 | 5.011519215 | 0.002591458 | 3.012452747 | 32.25652712 |
| CNAG_01642 | - | DNA mismatch repair protein MSH3 | 2985.975468 | 2652.736396 | 2722.724764 | 3182.528689 | 3296.822839 | 3284.879189 | 0.005519385 | 0.208094718 | 0.002645449 | 3.00618851 | 1.155161623 |
| CNAG_06576 | CAR1 | cAMP-regulated gene 1 | 5505.632406 | 6318.847788 | 4627.344168 | 4426.031272 | 3607.797454 | 3172.058417 | 0.005535752 | -0.568212372 | 0.00265397 | -3.005210596 | 0.674451977 |
| CNAG_04400 | - | small subunit ribosomal protein S10 | 265.0994999 | 320.9290633 | 342.1539026 | 183.8386641 | 225.7917193 | 259.8107931 | 0.005542507 | -0.489485135 | 0.002657885 | -3.004762232 | 0.712279249 |
| CNAG_06505 | - | hypothetical protein | 730.9060634 | 712.1545179 | 796.5340572 | 615.4063941 | 634.7445699 | 595.1863211 | 0.005551886 | -0.294898347 | 0.002663061 | -3.004170436 | 0.815129765 |
| CNAG_06902 | - | hypothetical protein | 1019.510676 | 902.9986917 | 932.1577815 | 1137.243583 | 1157.683893 | 1156.68722 | 0.005555267 | 0.258415039 | 0.002665361 | 3.003907733 | 1.196163864 |
| CNAG_02163 | - | hypothetical protein | 28.19958724 | 15.31599861 | 15.13118721 | 54.63957295 | 49.59543083 | 36.58270282 | 0.005625796 | 1.243678588 | 0.002699887 | 2.999989701 | 2.368015605 |
| CNAG_07124 | - | hypothetical protein | 14.3591746 | 15.23356463 | 9.784842708 | 38.9227706 | 35.73542919 | 29.84300434 | 0.005635885 | 1.409695838 | 0.002705418 | 2.999366389 | 2.656811437 |
| CNAG_05309 | - | hypothetical protein | 1111.87219 | 1219.667932 | 986.599967 | 1370.019018 | 1246.404321 | 1460.7757 | 0.005696478 | 0.283308635 | 0.0027352 | 2.996029497 | 1.21698268 |
| CNAG_01010 | - | hypothetical protein | 1744.592151 | 1792.811991 | 1938.723866 | 1594.564399 | 1638.806188 | 1489.976171 | 0.005714934 | -0.228871935 | 0.002744759 | -2.994965453 | 0.853301841 |
| CNAG_05449 | MTN1 | Copper-detoxifying metallothionein 1 | 2552.670537 | 1847.988026 | 2247.083114 | 2794.328164 | 2747.395234 | 2682.067809 | 0.005753614 | 0.290409538 | 0.002764039 | 2.992829729 | 1.222987398 |
| CNAG_04433 | YAK103 | CMGC/DYRK/DYRK2 protein kinase | 565.1676245 | 444.048712 | 417.137379 | 564.2895244 | 681.7334626 | 639.2906624 | 0.005753667 | 0.385605534 | 0.002764768 | 2.992749302 | 1.306408003 |
| CNAG_07880 | - | TRIAD3, E3 ubiquitin-protein ligase RNF216 | 112.4663945 | 130.4617135 | 112.122718 | 141.5218847 | 200.5319656 | 201.7085408 | 0.005755918 | 0.599823718 | 0.002766552 | 2.992552402 | 1.515531374 |
| CNAG_09006 | ND5 | NADH:ubiquinone oxidoreductase subunit 5 | 234467.1087 | 205720.7269 | 205179.309 | 245977.1996 | 241615.4738 | 258269.9637 | 0.005770439 | 0.193072082 | 0.002774236 | 2.991705661 | 1.143195452 |
| CNAG_12080 | - | unspecified product | 12.23084487 | 24.90367832 | 21.47073285 | 40.76950507 | 62.93397401 | 36.58091313 | 0.005776794 | 1.26143857 | 0.002777997 | 2.991292064 | 2.397346709 |
| CNAG_05902 | - | AP-3 complex subunit sigma | 383.9966417 | 386.7026236 | 375.3311767 | 297.0301495 | 308.6680367 | 288.8024945 | 0.005801202 | -0.372611954 | 0.002790442 | -2.989926866 | 0.772382855 |
| CNAG_01599 | - | hypothetical protein | 98.32634001 | 96.01246381 | 80.15611087 | 140.4630626 | 136.4716066 | 150.7127078 | 0.005801759 | 0.621670572 | 0.002791419 | 2.989819989 | 1.53865584 |
| CNAG_03013 | OPT1 | OPT family small oligopeptide transporter | 296.5408858 | 363.9598778 | 306.0238233 | 232.3391683 | 261.2524812 | 231.1003384 | 0.005804226 | -0.429056755 | 0.002793315 | -2.989612604 | 0.742747241 |
| CNAG_06888 | - | cytoplasmic protein | 538.4693113 | 503.2667636 | 515.1753646 | 656.3545764 | 651.3803674 | 632.7094299 | 0.005841178 | 0.301971874 | 0.002811811 | 2.987595831 | 1.23282829 |
| CNAG_07698 | - | hypothetical protein | 15.58810928 | 14.21250672 | 19.31911598 | 29.60006108 | 40.89723811 | 51.62535907 | 0.005845727 | 1.306492785 | 0.002815429 | 2.987202803 | 2.473395222 |
| CNAG_07879 | - | TRIAD3, E3 ubiquitin-protein ligase RNF216 | 194.0963251 | 204.7963333 | 182.4909163 | 273.5521153 | 252.8424862 | 275.1290056 | 0.005845727 | 0.44910213 | 0.002815248 | 2.98722248 | 1.365190358 |
| CNAG_04538 | - | ubiquitin-like modifier-activating enzyme ATG7 | 505.9048236 | 576.466417 | 530.1093596 | 694.5453401 | 661.7017614 | 653.8052031 | 0.005886275 | 0.303087745 | 0.002835677 | 2.985011505 | 1.233782207 |
| CNAG_05448 | - | hypothetical protein | 2.755594352 | 7.689541614 | 9.695971684 | 26.46990962 | 19.13195908 | 23.06015518 | 0.005941031 | 1.904262667 | 0.002863506 | 2.982022914 | 3.743175437 |
| CNAG_04870 | - | hypothetical protein | 9.872838089 | 11.01290377 | 26.69346484 | 40.2389827 | 41.7872789 | 40.79987889 | 0.005941031 | 1.379525603 | 0.002863321 | 2.98204261 | 2.601828018 |
| CNAG_05363 | - | pre-mRNA-splicing factor syf2 | 2263.068214 | 2299.211687 | 2496.481827 | 1916.636929 | 2095.086484 | 2116.016264 | 0.005944216 | -0.219394444 | 0.002865767 | -2.981781259 | 0.858925886 |
| CNAG_02572 | - | hypothetical protein | 12.9710791 | 1.274593519 | 4.431836474 | 16.38396239 | 25.42364198 | 31.65402502 | 0.005949452 | 2.062272692 | 0.002869018 | 2.981434135 | 4.176437048 |
| CNAG_01101 | - | hypothetical protein | 1111.467185 | 993.3797247 | 1046.143716 | 889.3928212 | 890.9080374 | 880.463025 | 0.00596682 | -0.260172099 | 0.002878122 | -2.98046387 | 0.834988308 |
| CNAG_05114 | - | peroxisomal copper amine oxidase | 93.88255536 | 134.7123918 | 104.6575928 | 156.647903 | 168.4359306 | 182.8829891 | 0.005968648 | 0.598521937 | 0.002879732 | 2.980292535 | 1.514164487 |
| CNAG_07114 | - | V-type proton ATPase proteolipid subunit 2 | 1637.092318 | 1725.956955 | 1569.810266 | 1449.036348 | 1405.850613 | 1425.532521 | 0.005974962 | -0.219518108 | 0.002883508 | -2.979891165 | 0.858852264 |
| CNAG_01386 | - | phosphatidylinositol glycan, class P | 91.08357897 | 46.60555557 | 57.7728996 | 113.3224162 | 122.9039944 | 107.5168452 | 0.00602369 | 0.80023449 | 0.00290776 | 2.977324594 | 1.741384141 |
| CNAG_01982 | - | hypothetical protein | 72.49981105 | 49.77049176 | 47.11775857 | 98.54483939 | 92.49945845 | 104.0538616 | 0.006043395 | 0.791148205 | 0.00291801 | 2.976245664 | 1.730451138 |
| CNAG_12853 | - | unspecified product | 166.6097564 | 201.1120696 | 117.3776901 | 95.20224851 | 95.42002395 | 117.7865347 | 0.006044927 | -0.670428743 | 0.002919488 | -2.976090388 | 0.628319934 |
| CNAG_12899 | - | unspecified product | 4.044286566 | 12.04344661 | 20.32492967 | 36.44445991 | 29.48371964 | 41.73030328 | 0.00605803 | 1.544418402 | 0.002926556 | 2.9753488 | 2.916864566 |
| CNAG_03282 | - | hypothetical protein | 380.8777017 | 549.2830724 | 546.9162639 | 355.3067776 | 330.9247366 | 414.1363541 | 0.006121726 | -0.438207466 | 0.002958074 | -2.972061524 | 0.73805106 |
| CNAG_12087 | - | unspecified product | 16527.19365 | 14635.92827 | 14376.94196 | 17557.59401 | 18151.87508 | 16947.68831 | 0.006126666 | 0.193642489 | 0.002961209 | 2.971736252 | 1.143647533 |
| CNAG_03330 | - | RecQ-mediated genome instability protein 1 | 298.0831588 | 336.2289803 | 321.0873062 | 391.5923167 | 402.9264711 | 462.6719734 | 0.006133999 | 0.379455469 | 0.00296775 | 2.971058754 | 1.300850769 |
| CNAG_12586 | - | unspecified product | 16.766047 | 22.74648671 | 15.11336066 | 38.47444904 | 35.48643306 | 54.949274 | 0.006133999 | 1.247274603 | 0.002967328 | 2.971102384 | 2.373925405 |
| CNAG_02091 | NAP1 | Nucleosome assembly protein 1 | 11071.3825 | 10872.35039 | 11188.12486 | 9805.514253 | 9783.925134 | 10347.83715 | 0.006133999 | -0.16163703 | 0.002967054 | -2.971130774 | 0.894010059 |
| CNAG_05242 | - | hypothetical protein | 3429.521393 | 3315.211662 | 3124.708222 | 2854.580195 | 2873.944864 | 2989.710689 | 0.006133999 | -0.194351265 | 0.002965727 | -2.971268113 | 0.873965803 |
| CNAG_02040 | - | hypothetical protein | 36.78150203 | 65.78690597 | 53.45833198 | 103.4971212 | 83.82434532 | 91.91941039 | 0.00613867 | 0.833757614 | 0.00297076 | 2.970747457 | 1.782321521 |
| CNAG_05875 | - | cytochrome c heme-lyase | 2003.831322 | 2109.556097 | 2128.585804 | 1764.939278 | 1799.187169 | 1897.303883 | 0.00616221 | -0.207658122 | 0.002982904 | -2.96949427 | 0.865941745 |
| CNAG_07775 | - | hypothetical protein | 196.2200962 | 149.9345449 | 138.8156095 | 214.9494421 | 229.6237182 | 266.098731 | 0.00616387 | 0.537832488 | 0.00298446 | 2.969333972 | 1.451789702 |
| CNAG_04361 | - | hypothetical protein | 1563.779644 | 1689.432982 | 1431.291641 | 1931.196487 | 1976.32453 | 1712.990617 | 0.006188852 | 0.247951513 | 0.002999579 | 2.967781032 | 1.187519754 |
| CNAG_01315 | - | protein transporter | 1038.332731 | 1118.417583 | 1045.200343 | 1188.455425 | 1295.695652 | 1364.219258 | 0.006188852 | 0.250342399 | 0.002999553 | 2.967783707 | 1.189489387 |
| CNAG_02697 | - | hypothetical protein | 352.536832 | 375.9121749 | 376.3612237 | 246.2203637 | 319.1153926 | 276.5870535 | 0.006188852 | -0.406532889 | 0.002999446 | -2.967794676 | 0.754434267 |
| CNAG_01266 | - | ATP-dependent protease La | 1512.278496 | 1408.27301 | 1263.864706 | 1596.340967 | 1689.019812 | 1746.141253 | 0.006188852 | 0.250936838 | 0.002998508 | 2.967890838 | 1.189979597 |
| CNAG_03193 | - | hypothetical protein | 42.23136557 | 34.6419628 | 21.5640982 | 58.63608492 | 59.70155824 | 82.72057363 | 0.006192406 | 1.021551511 | 0.003002058 | 2.967527125 | 2.030101008 |
| CNAG_12369 | - | unspecified product | 9.74145456 | 10.94547634 | 12.920132 | 30.92385573 | 28.73211466 | 30.81543474 | 0.006195857 | 1.490195861 | 0.003004488 | 2.967278416 | 2.809271113 |
| CNAG_12367 | - | unspecified product | 90.7909736 | 106.3905552 | 84.18173631 | 68.34059541 | 44.44598007 | 50.22308691 | 0.006227981 | -0.80360434 | 0.003020825 | -2.965610771 | 0.57291605 |
| CNAG_12649 | - | unspecified product | 15.85388908 | 44.25623761 | 39.56871453 | 69.05022612 | 68.47568811 | 69.65785828 | 0.006235265 | 1.059345116 | 0.00302512 | 2.965173749 | 2.083985321 |
| CNAG_00351 | - | hypothetical protein | 294.005855 | 219.885729 | 378.2591404 | 232.5517687 | 166.0667568 | 207.7738672 | 0.006261197 | -0.574968607 | 0.003038466 | -2.963819386 | 0.671300859 |
| CNAG_01330 | - | hypothetical protein | 3287.144073 | 3261.358256 | 3339.023331 | 2838.213821 | 3018.910246 | 2948.654728 | 0.006302247 | -0.182733082 | 0.003059156 | -2.961730288 | 0.881032362 |
| CNAG_07376 | - | hypothetical protein | 2066.813621 | 1954.457813 | 2169.087634 | 1745.484442 | 1753.455301 | 1870.661899 | 0.006334013 | -0.220820747 | 0.003075349 | -2.960104277 | 0.858077139 |
| CNAG_05191 | - | hypothetical protein | 84.53704942 | 76.7439999 | 130.0979938 | 148.9800209 | 152.0428114 | 167.3278248 | 0.006381114 | 0.671403051 | 0.003098998 | 2.957743621 | 1.592621072 |
| CNAG_01050 | - | hypothetical protein | 1234.275403 | 1299.323643 | 1340.457414 | 1097.973285 | 1038.338606 | 1166.814767 | 0.006386146 | -0.245495968 | 0.003103001 | -2.95734563 | 0.84352576 |
| CNAG_01840 | TUB1 | tubulin beta chain | 11028.06663 | 9989.950702 | 10350.96972 | 9792.41158 | 9017.947813 | 9060.122206 | 0.006386146 | -0.186251487 | 0.003102776 | -2.957367968 | 0.878886342 |
| CNAG_00384 | - | RAD51-like protein 2 | 142.9002642 | 172.4973341 | 186.6819104 | 256.5962341 | 222.6537753 | 234.0955965 | 0.006415058 | 0.495286468 | 0.003117833 | 2.955875148 | 1.409600624 |
| CNAG_12289 | - | unspecified product | 9.52016751 | 3.409923414 | 2.290781421 | 22.47737675 | 16.52833772 | 22.97016692 | 0.006433533 | 2.082144124 | 0.003127598 | 2.954910512 | 4.234360567 |
| CNAG_12419 | - | unspecified product | 26.20332917 | 38.91071534 | 35.32861279 | 73.95367645 | 51.77607321 | 72.90303188 | 0.006442412 | 0.971678176 | 0.003132701 | 2.954407486 | 1.961120489 |
| CNAG_01005 | - | glutaredoxin | 1108.983047 | 984.7064512 | 906.4853158 | 944.9380198 | 731.19926 | 716.2724216 | 0.00650294 | -0.341924033 | 0.003162927 | -2.95144322 | 0.788988385 |
| CNAG_03279 | CCD4 | hypothetical protein | 322.5843728 | 386.8139213 | 301.9496854 | 400.0676065 | 454.7732916 | 483.7858456 | 0.006586828 | 0.391765917 | 0.003204533 | 2.947404884 | 1.31199836 |
| CNAG_12152 | - | unspecified product | 0.117830818 | 0.057309428 | 0.049508769 | 13.58403411 | 5.262154913 | 1.205988879 | 0.006610913 | 5.090160862 | 0.003217058 | 2.946198547 | 34.06364377 |
| CNAG_05737 | - | oligopeptide transporter 8 | 1055.416846 | 1096.683936 | 874.5293112 | 830.9889418 | 870.2402477 | 795.027052 | 0.006617623 | -0.292848992 | 0.003221132 | -2.945807159 | 0.816288483 |
| CNAG_03921 | - | hypothetical protein | 504.1238871 | 449.2318515 | 435.0856883 | 349.8956351 | 395.8479977 | 348.7590944 | 0.006619285 | -0.361818358 | 0.003222749 | -2.945651888 | 0.778183147 |
| CNAG_00627 | - | specific transcriptional repressor | 26.06626971 | 27.09969556 | 25.74795211 | 57.46455946 | 58.21019544 | 46.52476653 | 0.006656125 | 1.030217559 | 0.003241498 | 2.943856925 | 2.042332212 |
| CNAG_02773 | RRP6 | exosome complex exonuclease Rrp6 | 4067.201828 | 3998.329188 | 4227.361866 | 3334.666609 | 3794.373695 | 3700.219063 | 0.006703858 | -0.198427447 | 0.003265562 | -2.941566967 | 0.87149999 |
| CNAG_02823 | - | bloom syndrome protein | 1617.546875 | 1792.873878 | 1673.329549 | 1900.637415 | 1982.258993 | 2079.197604 | 0.006733497 | 0.215413698 | 0.003280822 | 2.940122776 | 1.161036798 |
| CNAG_01958 | - | HET-C2 protein | 2174.074775 | 1876.883968 | 1960.108722 | 1838.285533 | 1707.754573 | 1562.17123 | 0.006783168 | -0.250834805 | 0.003305852 | -2.937767162 | 0.840409977 |
| CNAG_05289 | - | hypothetical protein | 91.43145652 | 85.30721568 | 103.5317529 | 125.3526746 | 186.989055 | 133.1345088 | 0.006786012 | 0.654126882 | 0.003308067 | 2.937559482 | 1.573663285 |
| CNAG_04968 | - | hypothetical protein, variant | 443.6473862 | 486.8744659 | 455.3183118 | 345.199664 | 385.4624146 | 372.0186271 | 0.006786566 | -0.343649246 | 0.003309166 | -2.937456506 | 0.788045456 |
| CNAG_04055 | - | cell cycle checkpoint protein | 454.5127353 | 517.2029544 | 452.2806675 | 655.0012307 | 589.2446499 | 560.5964812 | 0.006792516 | 0.327596258 | 0.003312897 | 2.937107113 | 1.254920749 |
| CNAG_12977 | - | unspecified product | 12.13870415 | 6.725408101 | 24.56873538 | 47.75905079 | 32.05287559 | 37.50285728 | 0.006793232 | 1.448417652 | 0.003314075 | 2.936996802 | 2.72908561 |
| CNAG_06864 | - | hypothetical protein | 1083.413822 | 959.9752024 | 989.6331221 | 827.8472243 | 863.3158297 | 862.6821306 | 0.006828717 | -0.263887538 | 0.003332221 | -2.935303057 | 0.832840691 |
| CNAG_01172 | PBX1 | parallel beta-helix repeat protein | 1843.770123 | 1807.896825 | 1896.089784 | 1417.659072 | 1632.751601 | 1697.457019 | 0.006837291 | -0.240118471 | 0.003337239 | -2.934836091 | 0.846675782 |
| CNAG_04628 | - | translation initiation factor 6 | 745.3485378 | 736.0423142 | 943.805193 | 1007.278425 | 1226.140238 | 903.7074209 | 0.006838627 | 0.355452442 | 0.003338727 | 2.934697821 | 1.279386745 |
| CNAG_03227 | - | hypothetical protein | 585.1950316 | 366.3304109 | 333.8578001 | 309.4313721 | 321.5731229 | 273.3563533 | 0.006846475 | -0.525146196 | 0.003343394 | -2.934264282 | 0.69488869 |
| CNAG_06773 | SEC24 | protein transporter SEC24 | 1214.194552 | 1044.035456 | 1147.441126 | 852.6102132 | 1005.690791 | 975.9062136 | 0.006865533 | -0.281836382 | 0.003353539 | -2.933323835 | 0.822543349 |

| | | | | | | | | | | | | | |
|---|---|---|---|---|---|---|---|---|---|---|---|---|---|
| CNAG_01597 | - | hypothetical protein | 772.7517325 | 665.8415513 | 708.0725676 | 576.4564531 | 558.7460769 | 619.5286831 | 0.00687707 | -0.306854457 | 0.003360014 | -2.932724949 | 0.808402418 |
| CNAG_05429 | - | hypothetical protein | 92.47179217 | 83.11531506 | 79.06740366 | 131.9232402 | 139.1976021 | 126.4153687 | 0.006885238 | 0.625070687 | 0.003364846 | 2.932278738 | 1.54228639 |
| CNAG_05545 | - | bis(5'-adenosyl)-triphosphatase | 401.5387164 | 378.1496502 | 437.1144199 | 338.4096181 | 297.3514327 | 312.100814 | 0.006897584 | -0.376201377 | 0.003372564 | -2.931567177 | 0.770463556 |
| CNAG_13075 | - | unspecified product | 1.471926329 | 4.422498622 | 2.255285899 | 13.58378143 | 21.3544712 | 9.969761888 | 0.006897584 | 2.605388286 | 0.003372262 | 2.931595019 | 6.085552656 |
| CNAG_05598 | - | hypothetical protein | 1053.378108 | 1376.756102 | 1141.078615 | 988.9944029 | 954.6826348 | 1012.549679 | 0.006901889 | -0.287165033 | 0.003375512 | -2.93129584 | 0.819510857 |
| CNAG_04790 | EZA1 | hypothetical protein | 610.9235974 | 598.0757714 | 629.2659357 | 815.2655683 | 774.7426467 | 690.5233238 | 0.006916907 | 0.295586423 | 0.003383701 | 2.930543039 | 1.22738378 |
| CNAG_00834 | - | phosphatidylserine decarboxylase, variant | 1715.699821 | 1329.420744 | 1045.178003 | 942.1410762 | 880.378622 | 868.3934551 | 0.006936531 | -0.620271111 | 0.003394148 | -2.929585149 | 0.650548665 |
| CNAG_00278 | - | methionine aminopeptidase, type I | 1424.526145 | 1425.381865 | 1362.906573 | 1186.672377 | 1254.094961 | 1207.94754 | 0.006957781 | -0.222702734 | 0.003405396 | -2.92855684 | 0.856958512 |
| CNAG_07912 | - | hypothetical protein | 14.4262972 | 21.64501907 | 11.92080396 | 42.78912427 | 39.16470808 | 33.18017291 | 0.007012477 | 1.295663256 | 0.003433023 | 2.926044106 | 2.454898295 |
| CNAG_03249 | - | THO complex subunit 4 | 3186.684474 | 3096.513187 | 3158.797161 | 2916.198639 | 2803.238191 | 2632.32098 | 0.00701411 | -0.192278771 | 0.003434678 | -2.925894101 | 0.875222195 |
| CNAG_04405 | - | cyclin H | 936.3406101 | 920.0517632 | 859.544688 | 791.4005873 | 783.9815349 | 695.1397716 | 0.007021367 | -0.274245208 | 0.00343909 | -2.925494768 | 0.826882815 |
| CNAG_00015 | - | hypothetical protein | 94.98744284 | 118.5929792 | 104.6338519 | 167.0582513 | 154.5908067 | 154.1627577 | 0.007033435 | 0.573584681 | 0.003445859 | 2.924882846 | 1.48821677 |
| CNAG_06503 | - | uridine permease, variant | 3.88134334 | 7.667850686 | 6.523174695 | 14.02480692 | 21.02316507 | 27.27045242 | 0.007041257 | 1.940965517 | 0.003450551 | 2.924459351 | 3.83962527 |
| CNAG_04339 | - | exocyst complex component EXO84, variant | 2157.921769 | 2333.611826 | 2046.52683 | 1969.761293 | 1819.84998 | 1900.677215 | 0.007042308 | -0.215185642 | 0.003451926 | -2.924335353 | 0.861435305 |
| CNAG_05538 | JJJ1 | DnaJ-like cochaperone | 2407.760679 | 2379.991518 | 2241.678211 | 1991.421828 | 2040.710731 | 2141.538801 | 0.007062619 | -0.202338479 | 0.003462744 | -2.923361311 | 0.869140622 |
| CNAG_07395 | - | hypothetical protein | 15.66849566 | 17.44840883 | 27.80028029 | 54.64932387 | 44.27317905 | 38.75422833 | 0.00706839 | 1.189355246 | 0.003466437 | 2.923029437 | 2.280508023 |
| CNAG_06466 | - | jumonji domain containing 5 | 555.6826115 | 530.0583857 | 570.4666316 | 462.9081469 | 455.3046571 | 426.461963 | 0.007085227 | -0.316900035 | 0.003475559 | -2.922211056 | 0.802793012 |
| CNAG_07910 | - | hypothetical protein | 0.144996848 | 1.164387963 | 0.060786097 | 6.370189027 | 11.65041814 | 7.644052126 | 0.007104074 | 4.464864139 | 0.003485672 | 2.921306071 | 22.08299796 |
| CNAG_07972 | - | hypothetical protein | 570.7354252 | 625.7611328 | 443.6981102 | 478.6217629 | 359.3610433 | 405.3776848 | 0.007111897 | -0.413498568 | 0.003490379 | -2.920885644 | 0.750800456 |
| CNAG_00807 | - | hypothetical protein | 623.6655713 | 588.3571694 | 521.6253694 | 782.6024808 | 738.5187342 | 661.650887 | 0.007128666 | 0.317137525 | 0.003499479 | 2.920074313 | 1.245856171 |
| CNAG_07689 | - | pre-mRNA-splicing factor cwc22 | 1029.839781 | 1173.150899 | 942.7579478 | 890.9825075 | 928.9316944 | 758.460745 | 0.007138082 | -0.302186787 | 0.003504973 | -2.919585402 | 0.811022145 |
| CNAG_03324 | - | hypothetical protein | 608.1270955 | 615.0778681 | 505.5200857 | 487.0641973 | 461.3164937 | 433.1400889 | 0.007138698 | -0.338415189 | 0.003505019 | -2.919403513 | 0.790909655 |
| CNAG_02744 | - | hypothetical protein | 1008.209442 | 1386.662228 | 1322.437883 | 1515.517296 | 1502.683873 | 1586.270588 | 0.007138698 | 0.294867046 | 0.003506894 | 2.919414638 | 1.226771917 |
| CNAG_07549 | - | hypothetical protein | 22.76520248 | 34.64818251 | 40.62234751 | 98.95160144 | 47.36166193 | 58.7247082 | 0.007159177 | 1.06654515 | 0.003517954 | 2.918433019 | 2.094411828 |
| CNAG_06757 | - | hypothetical protein, variant 2 | 1577.690392 | 1820.771209 | 1828.887249 | 1441.138472 | 1551.678958 | 1502.135093 | 0.007162783 | -0.232233514 | 0.003520601 | -2.918198565 | 0.851315903 |
| CNAG_04414 | - | multidrug resistance protein fnx1 | 493.5343267 | 431.9761881 | 390.3266094 | 335.9192579 | 325.829156 | 361.9950487 | 0.007167057 | -0.379476136 | 0.003523577 | -2.917935096 | 0.768716673 |
| CNAG_07437 | ERG27 | 3-keto sterol reductase | 818.445759 | 755.2763082 | 871.1860641 | 692.5097866 | 695.9884089 | 637.3918529 | 0.007174468 | -0.287805802 | 0.003528097 | -2.917535332 | 0.819146954 |
| CNAG_06540 | - | dihydroxy-acid dehydratase, variant | 140.1084024 | 88.57476325 | 83.3947518 | 193.0640471 | 133.6587134 | 180.644883 | 0.007186851 | 0.685872297 | 0.003535064 | 2.916920064 | 1.608674341 |
| CNAG_06697 | MPS1 | TTK protein kinase, variant | 519.8887056 | 539.8806216 | 566.3343727 | 647.6652319 | 704.0565636 | 661.567797 | 0.007233769 | 0.29380924 | 0.003559025 | 2.914812324 | 1.225872758 |
| CNAG_01064 | - | hypothetical protein | 287.5267822 | 295.3209208 | 310.4012747 | 383.935548 | 396.1111151 | 381.7842635 | 0.007240814 | 0.363757888 | 0.003563375 | 2.914431022 | 1.286773284 |
| CNAG_02097 | - | hypothetical protein | 878.34332 | 976.1701903 | 831.9392546 | 1051.589614 | 1131.002792 | 1065.679705 | 0.007243161 | 0.259406267 | 0.003565415 | 2.914252434 | 1.19698599 |
| CNAG_02473 | - | endoplasmic reticulum protein | 225.3754044 | 178.9949326 | 145.2396419 | 261.8487508 | 272.8473065 | 254.0941215 | 0.007248288 | 0.506228138 | 0.003568824 | 2.913954085 | 1.420331942 |
| CNAG_00863 | - | Flavin-containing monooxygenase | 131.0872435 | 138.0227345 | 132.3547934 | 184.6930826 | 181.2585733 | 215.0619457 | 0.007262501 | 0.517612228 | 0.003576708 | 2.913265087 | 1.431583904 |
| CNAG_03139 | CHO2 | Phosphatidylethanolamine N-methyltransferase | 1826.50245 | 1881.240088 | 1787.443099 | 2074.233581 | 2191.042997 | 2100.365377 | 0.007272959 | 0.197156395 | 0.003582747 | 2.912738281 | 1.146436458 |
| CNAG_12448 | - | unspecified product | 27.14702969 | 15.33939124 | 27.83619254 | 52.7694696 | 49.46236063 | 49.71951777 | 0.007292643 | 1.103262363 | 0.003593334 | 2.911816694 | 2.148399608 |
| CNAG_03287 | - | 28 kda golgi snare protein | 1070.864184 | 1219.564324 | 1230.603784 | 966.3852588 | 998.7287027 | 1024.747127 | 0.007294781 | -0.250769716 | 0.003595278 | -2.911647742 | 0.840447894 |
| CNAG_05877 | - | CDK-activating kinase assembly factor MAT1 | 669.0493615 | 712.0972624 | 704.8598031 | 611.6383017 | 557.042989 | 557.4373909 | 0.007304965 | -0.288436329 | 0.00360119 | -2.911134471 | 0.818789026 |
| CNAG_05224 | - | hypothetical protein | 125.2431931 | 131.5570116 | 125.9557159 | 176.1341153 | 185.6773379 | 190.7340205 | 0.007311719 | 0.517227458 | 0.003605412 | 2.910768354 | 1.431202149 |
| CNAG_02112 | - | ubiquitin-conjugating enzyme E2 W | 616.3679484 | 566.7044053 | 612.0681153 | 478.378908 | 492.3880481 | 497.450074 | 0.007320767 | -0.305909879 | 0.003610768 | -2.910304503 | 0.808931879 |
| CNAG_00623 | EGCrP1 | fungus-specific glucosylceramide | 1048.458348 | 968.5930282 | 1028.003085 | 893.3577063 | 837.3948846 | 849.3815767 | 0.007326196 | -0.254699391 | 0.00361434 | -2.909995443 | 0.838161761 |
| CNAG_04937 | - | peroxin-12 | 118.4055066 | 158.4518956 | 151.5184324 | 212.1180974 | 210.7161132 | 195.2793109 | 0.007336467 | 0.516618273 | 0.003620303 | 2.909480188 | 1.430597944 |
| CNAG_05440 | - | pyridoxamine 5'-phosphate oxidase | 1059.124846 | 1124.778373 | 1186.869777 | 883.0332693 | 916.7419581 | 1032.460296 | 0.007355937 | -0.26598058 | 0.003630809 | -2.908574259 | 0.831633293 |
| CNAG_12237 | - | unspecified product | 7.270772618 | 3.40873372 | 5.463711869 | 25.03034403 | 18.34284211 | 16.54016547 | 0.007427667 | 1.958573136 | 0.003667122 | 2.905461325 | 3.886773765 |
| CNAG_00698 | - | hypothetical protein | 1488.862697 | 1508.38527 | 1711.546488 | 1305.729533 | 1388.648343 | 1343.387719 | 0.007428792 | -0.237300083 | 0.003668584 | -2.905336545 | 0.848331429 |
| CNAG_06391 | - | hypothetical protein | 120.4130001 | 81.04914396 | 107.8172932 | 141.7819459 | 165.9237452 | 171.7813503 | 0.007460781 | 0.614416243 | 0.003685292 | 2.903914121 | 1.530938409 |
| CNAG_06463 | - | hypothetical protein, variant | 8.874798633 | 38.88758551 | 38.48188853 | 74.09310453 | 64.17872253 | 57.56862591 | 0.00747732 | 1.203040463 | 0.003694375 | 2.903143339 | 2.302243552 |
| CNAG_05684 | - | hypothetical protein | 835.0490119 | 764.006937 | 806.3012906 | 991.7874639 | 976.5474738 | 939.1399681 | 0.007527867 | 0.25737192 | 0.003720269 | 2.900955376 | 1.195299308 |
| CNAG_01027 | - | succinate-semialdehyde dehydrogenase (NADP) | 1932.324722 | 1792.774899 | 1561.319304 | 1466.848119 | 1586.204906 | 1420.039529 | 0.007536078 | -0.256537132 | 0.003725247 | -2.900536322 | 0.837094769 |
| CNAG_02183 | - | anaphase-promoting complex subunit 1 | 847.7557561 | 936.1910416 | 901.0976859 | 745.3801758 | 793.4948575 | 722.8529544 | 0.007572206 | -0.263041335 | 0.00374403 | -2.898959707 | 0.833329331 |
| CNAG_02436 | - | hypothetical protein | 579.2817051 | 517.277115 | 515.1930982 | 682.8321529 | 663.4334683 | 658.2357401 | 0.007574579 | 0.298605203 | 0.003746128 | 2.898784037 | 1.229954719 |
| CNAG_12820 | - | unspecified product | 82.9167117 | 35.64509231 | 60.594615 | 34.72935141 | 31.6725451 | 11.5337473 | 0.007613874 | -1.239090124 | 0.003766492 | -2.897083721 | 0.423639752 |
| CNAG_01815 | - | hypothetical protein | 299.1478852 | 308.2210724 | 280.5899023 | 406.6628846 | 372.7367884 | 378.4588673 | 0.007617458 | 0.367141198 | 0.003769195 | 2.896858658 | 1.289794478 |
| CNAG_12821 | - | unspecified product | 128.7328645 | 144.441383 | 111.0811251 | 191.8721045 | 203.9189841 | 168.6895308 | 0.007646766 | 0.538400708 | 0.003784631 | 2.895576221 | 1.452361616 |
| CNAG_01217 | - | protein MPE1 | 2243.204979 | 2283.043364 | 2360.006611 | 2007.814385 | 2071.80279 | 2018.354239 | 0.007684768 | -0.19106719 | 0.003804378 | -2.893942534 | 0.875957519 |
| CNAG_03643 | - | hypothetical protein | 107.6301298 | 94.96794807 | 93.98408935 | 147.4535878 | 148.5924402 | 149.6787715 | 0.007693473 | 0.579159364 | 0.003809626 | 2.893509599 | 1.493978477 |
| CNAG_03458 | - | hypothetical protein | 1256.299763 | 1164.675432 | 1178.397924 | 996.7489326 | 1011.652204 | 1070.245468 | 0.007721751 | -0.241136092 | 0.003824572 | -2.892279754 | 0.846078781 |
| CNAG_03678 | - | hypothetical protein | 761.1332705 | 669.0812729 | 740.0444291 | 619.3709336 | 608.8556559 | 559.6870677 | 0.007739832 | -0.296837204 | 0.003834473 | -2.891467477 | 0.814035037 |
| CNAG_06907 | - | hypothetical protein, variant | 11.95303098 | 8.795216238 | 8.684011706 | 25.29171397 | 21.66130935 | 35.03099738 | 0.007758672 | 1.569591789 | 0.003844754 | 2.890625963 | 2.968207167 |
| CNAG_04059 | - | hypothetical protein | 63.51230199 | 77.70328848 | 82.20333123 | 123.4767468 | 123.6616217 | 109.7929866 | 0.007807981 | 0.662379305 | 0.003870142 | 2.888556716 | 1.58269066 |
| CNAG_00098 | - | palmitoyl-protein thioesterase | 380.8783181 | 395.4818607 | 354.1754037 | 458.5382867 | 466.748761 | 521.5536152 | 0.007901322 | 0.341151216 | 0.003917373 | 2.884739779 | 1.266767023 |
| CNAG_00648 | - | esterase/lipase | 852.1 | 770.2905771 | 678.287097 | 613.0208102 | 629.5313775 | 639.5485357 | 0.007909658 | -0.305604715 | 0.003922472 | -2.88433026 | 0.809103005 |
| CNAG_00695 | - | hypothetical protein | 295.7883832 | 352.3776805 | 341.3319778 | 413.4473782 | 416.7371267 | 453.8534379 | 0.00793126 | 0.361250972 | 0.003934153 | 2.883393793 | 1.284539248 |
| CNAG_07402 | - | pantothenate transporter | 370.5410088 | 447.1911796 | 446.8930335 | 553.6785704 | 502.9243038 | 559.3525815 | 0.007956227 | 0.339303011 | 0.003947509 | 2.882326171 | 1.265145234 |
| CNAG_00455 | UBP16 | ubiquitin carboxyl-terminal hydrolase Ubp16 | 764.8239329 | 835.987981 | 827.4901057 | 606.6856709 | 764.272227 | 607.4458208 | 0.00795779 | -0.310544302 | 0.003949256 | -2.882186716 | 0.806337486 |
| CNAG_03491 | NDX3 | cleavage and polyadenylation specificity factor subunit 5 | 1.425214049 | 3.331694782 | 0.085194742 | 18.15212638 | 5.963632561 | 13.07953741 | 0.007987339 | 3.159500505 | 0.003964896 | 2.8809412 | 8.935202993 |
| CNAG_02911 | - | hypothetical protein | 283.9982963 | 289.9253946 | 282.7054667 | 427.8933761 | 371.0455264 | 334.1603969 | 0.007996778 | 0.391370265 | 0.003971535 | 2.880413852 | 1.3116386 |
| CNAG_01741 | - | hypothetical protein, variant | 461.5459143 | 539.8244418 | 488.5190772 | 610.3636865 | 605.6203392 | 642.6285942 | 0.007996778 | 0.305048371 | 0.003971155 | 2.880443995 | 1.235460059 |
| CNAG_04476 | - | hypothetical protein | 11.09284784 | 35.547379 | 11.96002356 | 45.60057529 | 61.99462737 | 38.77869584 | 0.008017572 | 1.326022246 | 0.003982841 | 2.879517583 | 2.5071047 |
| CNAG_12365 | - | unspecified product | 13.44435071 | 36.67163589 | 22.56825553 | 59.97433578 | 58.25388822 | 42.14110539 | 0.00804019 | 1.150876166 | 0.003995059 | 2.878551652 | 2.220487063 |
| CNAG_04791 | - | hypothetical protein | 1891.880273 | 1731.548195 | 1962.300066 | 2211.307812 | 2103.827722 | 2251.292527 | 0.008129987 | 0.217885937 | 0.00404067 | 2.874969108 | 1.163028086 |
| CNAG_02507 | - | translation initiation factor 3 subunit H | 293.9436525 | 258.5399312 | 292.0822738 | 255.3783559 | 171.2906592 | 178.9937948 | 0.008135693 | -0.496620064 | 0.004044499 | -2.874670024 | 0.708765328 |
| CNAG_05003 | - | hypothetical protein | 285.4026319 | 412.6610919 | 362.6729637 | 507.9386304 | 452.1013849 | 447.3120342 | 0.008192529 | 0.395062278 | 0.004073755 | 2.872393391 | 1.314999525 |
| CNAG_01150 | - | omega-6 fatty acid desaturase (delta-12 desaturase) | 3946.635131 | 3399.265077 | 3099.145369 | 3104.497043 | 2901.508279 | 2925.384407 | 0.008205916 | -0.241668656 | 0.004081414 | -2.871799845 | 0.845766512 |
| CNAG_07885 | - | hypothetical protein | 0.140649945 | 1.161561372 | 0.058985004 | 9.005305348 | 7.963857874 | 7.634873014 | 0.00822512 | 4.404376495 | 0.00409197 | 2.870983442 | 21.17626862 |
| CNAG_01847 | - | hypothetical protein | 66.7992567 | 48.71451979 | 66.21812455 | 101.6360187 | 102.0714274 | 98.61535922 | 0.008240045 | 0.725375102 | 0.004100401 | 2.870332711 | 1.653330447 |
| CNAG_00532 | - | hypothetical protein | 1298.823191 | 1386.745521 | 1583.679567 | 1671.125351 | 1691.584743 | 1737.287255 | 0.008282291 | 0.241835138 | 0.004122435 | 2.868637879 | 1.182495865 |
| CNAG_03816 | PRE3 | 20S proteasome subunit beta 1 | 2650.876781 | 2509.415064 | 2516.892565 | 2792.512419 | 3093.299447 | 2999.56869 | 0.008298679 | 0.195505969 | 0.004131605 | 2.867934928 | 1.145125698 |
| CNAG_02767 | - | transcription initiation factor TFIIE subunit alpha | 793.8809511 | 752.0055608 | 769.9263398 | 644.2191596 | 690.870512 | 588.5670172 | 0.008302693 | -0.283427448 | 0.004135631 | -2.86762675 | 0.821636713 |
| CNAG_07701 | CTR1 | putative copper ion transporter | 1682.815514 | 1936.074372 | 2105.051735 | 1470.603125 | 1738.896429 | 1618.709055 | 0.008302693 | -0.260952747 | 0.004134926 | -2.867680673 | 0.834536614 |
| CNAG_00414 | MAK32 | Maintenance of killer protein 32, putative | 496.0773134 | 510.57677 | 451.0908854 | 438.1773804 | 354.2677532 | 356.552293 | 0.008320679 | -0.358797782 | 0.004145607 | -2.866864352 | 0.779814139 |
| CNAG_04451 | - | ribonuclease P/MRP protein subunit POP1 | 1482.991014 | 1477.13797 | 1623.054559 | 1269.919536 | 1361.050311 | 1332.271561 | 0.008356872 | -0.225169551 | 0.004164659 | -2.865421818 | 0.855494479 |
| CNAG_07830 | - | hypothetical protein | 139.3887176 | 198.2471071 | 145.191301 | 211.8379263 | 242.6729467 | 239.5777674 | 0.008362295 | 0.507931737 | 0.004168383 | 2.865129822 | 1.422010125 |
| CNAG_05136 | - | cation-transporting ATPase 13A1 | 3671.087532 | 3229.029622 | 2975.429221 | 2754.133221 | 2989.592517 | 2761.085315 | 0.008377848 | -0.231459212 | 0.004177159 | -2.864463815 | 0.851772931 |
| CNAG_06616 | - | hypothetical protein | 1014.579491 | 930.8824346 | 986.4121736 | 881.7396506 | 786.4785268 | 802.7696847 | 0.008386558 | -0.262519789 | 0.00418355 | -2.863979583 | 0.833630641 |
| CNAG_12835 | - | unspecified product | 3.825398665 | 5.514105183 | 4.390098114 | 14.22137371 | 18.45311709 | 19.67237309 | 0.008386558 | 2.073406202 | 0.004183257 | 2.864001748 | 4.208791969 |
| CNAG_12459 | - | unspecified product | 68.50167183 | 70.89864965 | 62.81920527 | 32.10822925 | 28.92072571 | 47.80232436 | 0.00842371 | -0.916878502 | 0.004203871 | -2.862444367 | 0.529653771 |
| CNAG_04936 | - | condensin complex subunit 2 | 1958.170482 | 1922.086082 | 1824.692231 | 1643.561086 | 1627.52441 | 1727.453872 | 0.00842137 | -0.206088155 | 0.004203261 | -2.862490352 | 0.866884592 |

| | | | | | | | | | | | | | |
|---|---|---|---|---|---|---|---|---|---|---|---|---|---|
| CNAG_12843 | - | unspecified product | 15.59088702 | 14.21372402 | 20.37478166 | 51.72893609 | 27.55500446 | 41.88721117 | 0.008469023 | 1.265019736 | 0.004227789 | 2.860645929 | 2.403304976 |
| CNAG_02284 | - | 2-oxoisovalerate dehydrogenase E1 component, alpha subunit | 210.356518 | 175.7939367 | 194.1985153 | 318.3089512 | 271.0434637 | 227.615942 | 0.008469705 | 0.478970491 | 0.004229164 | 2.860542854 | 1.393748731 |
| CNAG_05666 | - | hypothetical protein | 2.747264283 | 5.558741721 | 9.689383656 | 35.81985877 | 12.85905204 | 18.74739258 | 0.008492742 | 2.008619548 | 0.004241704 | 2.859603936 | 4.023969996 |
| CNAG_02707 | - | hypothetical protein | 12.21886695 | 18.50688984 | 27.79089072 | 55.53412967 | 37.20481202 | 41.99166239 | 0.008511363 | 1.207843177 | 0.004252044 | 2.858831655 | 2.309920464 |
| CNAG_03898 | - | phosphoadenosine phosphosulfate reductase | 1522.566229 | 1311.270221 | 1594.239637 | 1291.051546 | 1215.193774 | 1242.373349 | 0.008515494 | -0.256175913 | 0.004255147 | -2.85860017 | 0.837304386 |
| CNAG_00581 | - | saccharopepsin | 7745.787661 | 7953.627064 | 7058.94373 | 8456.979247 | 8809.444507 | 8617.009574 | 0.008521276 | 0.170669649 | 0.004259077 | 2.858307294 | 1.125580819 |
| CNAG_12056 | - | unspecified product | 27.16371574 | 13.20523961 | 31.01012124 | 47.03624782 | 53.86363245 | 55.17863093 | 0.008560352 | 1.108839008 | 0.004279653 | 2.856777785 | 2.15672018 |
| CNAG_07731 | - | hypothetical protein | 35.60711748 | 48.66261226 | 65.09601365 | 76.41006223 | 82.99536175 | 111.5962951 | 0.008622091 | 0.847565929 | 0.004311572 | 2.854418304 | 1.799462365 |
| CNAG_02072 | - | hypothetical protein | 442.8012118 | 489.1922706 | 407.514726 | 655.1489302 | 564.2429881 | 506.2525344 | 0.008624601 | 0.35178596 | 0.00431388 | 2.854248271 | 1.276139425 |
| CNAG_00109 | - | large subunit ribosomal protein L34, variant | 375.6617099 | 312.4465265 | 349.7115711 | 280.7756056 | 256.8311199 | 261.0438894 | 0.008636859 | -0.394282561 | 0.004321066 | -2.853719538 | 0.760867655 |
| CNAG_03614 | - | hypothetical protein | 13.27770724 | 12.06717691 | 21.40942561 | 38.71354818 | 30.27678656 | 45.07344633 | 0.00863899 | 1.281246929 | 0.004323187 | 2.853563591 | 2.430489547 |
| CNAG_05269 | - | 20S proteasome subunit alpha 7 | 3865.222141 | 3816.258416 | 4222.009854 | 3541.081187 | 3568.338313 | 3484.880755 | 0.008698426 | -0.183675788 | 0.004353993 | -2.851306843 | 0.880456853 |
| CNAG_01442 | - | cysteine desulfurase IscS | 1328.77002 | 1315.492517 | 1283.981962 | 1124.440227 | 1105.665965 | 1172.382521 | 0.008702316 | -0.222275916 | 0.004357003 | -2.85108714 | 0.857212079 |
| CNAG_13009 | - | unspecified product | 101.5952733 | 126.969204 | 139.4867971 | 75.81751186 | 86.083146 | 74.61333234 | 0.008719087 | -0.654055583 | 0.004366464 | -2.850397334 | 0.635491361 |
| CNAG_06022 | - | trans-aconitate 3-methyltransferase | 431.0252308 | 348.1919908 | 431.9618009 | 477.1360874 | 534.1187123 | 558.1989257 | 0.008739668 | 0.358699548 | 0.004377838 | 2.849569916 | 1.282269533 |
| CNAG_05785 | STB4 | putative transcription factor | 6.209730561 | 3.432724689 | 11.80188904 | 20.55216318 | 22.71319067 | 27.37050307 | 0.008761602 | 1.769626516 | 0.004389895 | 2.848694932 | 3.409656762 |
| CNAG_05645 | - | COP9 signalosome complex subunit 3 | 1105.721866 | 1138.756173 | 1031.253725 | 892.4293949 | 945.2676367 | 960.3530984 | 0.008819758 | -0.242198574 | 0.004420111 | -2.846511695 | 0.84545591 |
| CNAG_00146 | PBY101 | 5'/3'-nucleotidase SurE | 324.8704655 | 324.4167435 | 347.7209172 | 425.9698434 | 447.0339666 | 405.1464131 | 0.008903749 | 0.344564941 | 0.004463291 | 2.843415036 | 1.269768013 |
| CNAG_05135 | PMR1 | Ca2 -transporting ATPase | 815.2794316 | 870.6080952 | 786.0705146 | 923.0529749 | 1118.175439 | 975.7686458 | 0.008935269 | 0.272408701 | 0.004480182 | 2.842211044 | 1.207822707 |
| CNAG_02347 | - | hypothetical protein | 276.5119773 | 268.193231 | 266.5426835 | 198.9020271 | 196.4410455 | 218.7890317 | 0.008966431 | -0.419603029 | 0.004496902 | -2.841023323 | 0.747630314 |
| CNAG_04612 | - | hypothetical protein, variant 2 | 1416.58276 | 1353.332348 | 1353.395582 | 1693.837581 | 1542.290868 | 1608.54915 | 0.009032756 | 0.217239113 | 0.004531269 | 2.838594583 | 1.162506766 |
| CNAG_00707 | - | protein sym1 | 1492.492026 | 1351.214165 | 1502.670184 | 1735.786762 | 1635.451697 | 1763.926954 | 0.009095296 | 0.22484272 | 0.004563753 | 2.83631418 | 1.168649838 |
| CNAG_07444 | - | hypothetical protein | 45.06417358 | 104.5123922 | 81.17112043 | 139.1872417 | 125.3077719 | 125.2611232 | 0.009124613 | 0.74571224 | 0.004580692 | 2.835130865 | 1.676801881 |
| CNAG_12716 | - | unspecified product | 3.897856692 | 5.555653076 | 9.685635824 | 29.93429303 | 18.26881659 | 16.58902496 | 0.009124613 | 1.87495559 | 0.004580289 | 2.835158988 | 3.667903264 |
| CNAG_07522 | - | hypothetical protein | 731.1384146 | 746.7040778 | 648.5181373 | 889.8591895 | 852.3137193 | 844.7603592 | 0.009149814 | 0.267693432 | 0.00459446 | 2.834171964 | 1.203881533 |
| CNAG_12751 | - | unspecified product | 6.223725948 | 9.824700619 | 7.604916746 | 25.51866833 | 27.18844818 | 18.80126142 | 0.009185352 | 1.655319851 | 0.004613427 | 2.832855263 | 3.149930179 |
| CNAG_13026 | - | unspecified product | 32.01895229 | 48.56775634 | 36.42435289 | 90.56202594 | 54.29157398 | 75.21974312 | 0.009233843 | 0.903555725 | 0.004638909 | 2.831093936 | 1.870670838 |
| CNAG_01522 | - | phenylalanine-tRNA ligase | 71.77514056 | 102.4609923 | 107.7782316 | 176.8096573 | 129.4554919 | 136.4375546 | 0.009246042 | 0.641256182 | 0.004646167 | 2.830593887 | 1.559686617 |
| CNAG_00354 | - | vacuolar protein 8 | 2678.985653 | 2863.843229 | 2693.922258 | 3015.908068 | 3164.826943 | 3261.54387 | 0.009275206 | 0.18229463 | 0.004661955 | 2.829508588 | 1.13468719 |
| CNAG_07981 | - | hypothetical protein | 43.41014038 | 37.85829742 | 25.81100396 | 81.12126875 | 64.05901681 | 57.64964269 | 0.009305093 | 0.928557573 | 0.004678113 | 2.828401255 | 1.903372026 |
| CNAG_00892 | - | hypothetical protein | 420.4091415 | 301.8845346 | 366.9453592 | 495.3540348 | 503.1223452 | 440.6953061 | 0.009313432 | 0.386998528 | 0.004683443 | 2.828036778 | 1.307670014 |
| CNAG_02223 | - | signal sequence binding protein | 5620.241388 | 5339.728933 | 5261.86863 | 4757.51457 | 4698.246978 | 5065.68636 | 0.009320127 | -0.175066613 | 0.004687947 | -2.827728993 | 0.885726622 |
| CNAG_00665 | - | DNA topoisomerase 2-associated protein PAT1 | 2926.317431 | 2971.513893 | 2931.651512 | 2639.048308 | 2718.747733 | 2535.713668 | 0.009322754 | -0.176963742 | 0.004690408 | -2.827561013 | 0.884562666 |
| CNAG_12053 | - | unspecified product | 8.369022501 | 4.466822692 | 2.284190833 | 20.03380464 | 19.29247551 | 17.58278921 | 0.009378019 | 1.983157063 | 0.004719357 | 2.825590365 | 3.953573 |
| CNAG_06535 | - | ribosome biogenesis protein UTP30 | 348.8930111 | 309.2002171 | 348.6257295 | 276.1293641 | 246.4686493 | 254.3799361 | 0.009397468 | -0.388742126 | 0.004730292 | -2.824848848 | 0.763795261 |
| CNAG_13074 | - | unspecified product | 1.3785892 | 0.081532092 | 1.154186329 | 9.71571142 | 12.48329194 | 7.691513955 | 0.009443104 | 3.789325544 | 0.004754417 | 2.823218437 | 13.82613051 |
| CNAG_07428 | - | hypothetical protein | 1677.95397 | 1746.434781 | 1617.793339 | 1470.040285 | 1329.859828 | 1550.914189 | 0.00945021 | -0.22752043 | 0.004759148 | -2.822899533 | 0.854101581 |
| CNAG_12492 | - | unspecified product | 183.3304058 | 256.1299837 | 195.1189356 | 157.725025 | 154.2098928 | 138.9969401 | 0.009526664 | -0.511627529 | 0.004798814 | -2.820237357 | 0.701430696 |
| CNAG_04518 | ZFC5 | hypothetical protein | 622.3399342 | 713.1080308 | 608.931042 | 542.2756747 | 532.9231028 | 527.4476517 | 0.009560543 | -0.293897227 | 0.004817047 | -2.81902029 | 0.815695599 |
| CNAG_02199 | - | hypothetical protein | 32.01269783 | 51.76442224 | 31.12609486 | 84.14791774 | 79.8044583 | 54.42811974 | 0.00957134 | 0.915018156 | 0.004823656 | 2.818580169 | 1.885592801 |
| CNAG_02351 | CHI4 | endochitinase | 830.6085754 | 1095.718125 | 873.5340639 | 1077.202706 | 1257.885079 | 1133.391209 | 0.009577101 | 0.295184052 | 0.004827729 | 2.818309219 | 1.227041507 |
| CNAG_05405 | - | hypothetical protein | 140.6614866 | 212.2921757 | 197.3660179 | 287.0099885 | 231.2057286 | 264.0335525 | 0.009631978 | 0.496715808 | 0.004856568 | 2.816396527 | 1.410997868 |
| CNAG_02367 | - | GTP-binding protein ypt3 | 4906.946548 | 5625.187134 | 5378.082636 | 4759.112279 | 4832.004244 | 4631.684805 | 0.009675812 | -0.17643432 | 0.004879852 | -2.814859814 | 0.884887331 |
| CNAG_02057 | URE6 | urease accessory protein | 932.9005893 | 867.3300899 | 993.8339172 | 790.5601939 | 738.1604169 | 817.1366521 | 0.009685518 | -0.267218876 | 0.004885929 | -2.814459781 | 0.83091979 |
| CNAG_04261 | - | hypothetical protein | 94.84915521 | 94.93457252 | 85.46157976 | 128.6132126 | 160.0344785 | 129.7834293 | 0.009693543 | 0.595698772 | 0.004891161 | 2.81411574 | 1.511204363 |
| CNAG_01539 | - | Myo-inositol-1-phosphate synthase | 8265.10282 | 7504.286968 | 7025.813443 | 6680.676986 | 6855.564591 | 6681.031806 | 0.00969895 | -0.188777608 | 0.004895074 | -2.813858717 | 0.877348782 |
| CNAG_01446 | HSP12 | heat shock protein, 12Kda | 7665.563512 | 10534.05943 | 6634.635783 | 14150.10841 | 11517.0406 | 12394.75177 | 0.009705052 | 0.603108589 | 0.004899339 | 2.813578731 | 1.518986016 |
| CNAG_04012 | FZC18 | hypothetical protein | 455.3135873 | 496.5710769 | 452.1375429 | 392.8702082 | 382.8439441 | 357.637766 | 0.009746587 | -0.325821918 | 0.004922687 | -2.812049879 | 0.797843717 |
| CNAG_05374 | - | chromosome transmission fidelity protein 18 | 1723.44641 | 1654.898456 | 1632.717132 | 1400.679372 | 1460.215032 | 1517.631847 | 0.009746587 | -0.209926192 | 0.004922613 | -2.812054737 | 0.864581462 |
| CNAG_01290 | - | hypothetical protein | 30.84031573 | 48.54517212 | 34.29296569 | 72.16574202 | 68.41340822 | 67.52039177 | 0.009787466 | 0.872574832 | 0.004945724 | 2.810547822 | 1.830927712 |
| CNAG_12620 | - | unspecified product | 36.48686886 | 28.22457434 | 35.31732284 | 63.54276774 | 67.67987741 | 57.58693331 | 0.009787466 | 0.901294319 | 0.004944654 | 2.810617445 | 1.867740882 |
| CNAG_07534 | TRS130 | Trafficking protein particle complex II-specific subunit 130 | 2498.870268 | 2528.66868 | 2336.598696 | 2138.558621 | 2149.394888 | 2237.020009 | 0.009805681 | -0.189398383 | 0.004956126 | -2.809871659 | 0.87697135 |
| CNAG_07899 | - | hypothetical protein | 3.808516795 | 4.444576256 | 4.382627963 | 12.5900807 | 17.58630451 | 19.64022655 | 0.009891643 | 2.11089513 | 0.005000782 | 2.806983414 | 4.319592237 |
| CNAG_05444 | - | NADPH dehydrogenase | 706.450212 | 814.3982313 | 715.5657326 | 619.3011747 | 622.6403707 | 625.1199289 | 0.009896779 | -0.274918727 | 0.005004587 | -2.806738385 | 0.826496877 |
| CNAG_06511 | - | hypothetical protein | 1253.856901 | 1078.497448 | 1094.160747 | 982.8204603 | 899.4496164 | 1005.862266 | 0.009928221 | -0.262690042 | 0.005025336 | -2.805405208 | 0.833532269 |
| CNAG_02740 | RPP1 | hypothetical protein | 457.9569689 | 446.164594 | 478.8913314 | 570.6851344 | 620.4364761 | 537.289138 | 0.009928221 | 0.306980459 | 0.005025013 | 2.805425905 | 1.237115722 |
| CNAG_00076 | NDX6 | A/G-specific adenine glycosylase | 1151.189722 | 1160.288853 | 984.3603217 | 936.8820843 | 971.177099 | 874.9590559 | 0.009928221 | -0.259831996 | 0.005023089 | -2.805549342 | 0.835185173 |
| CNAG_05895 | - | protein CFT1 | 3513.632561 | 3359.409907 | 3399.824074 | 2886.430873 | 2992.113803 | 3230.5777 | 0.009928221 | -0.18872308 | 0.005022314 | -2.805599067 | 0.877381943 |
| CNAG_02515 | - | hypothetical protein | 559.5932598 | 628.1780601 | 572.7683 | 751.3615043 | 727.2808913 | 679.3849103 | 0.009953887 | 0.279458631 | 0.005039543 | 2.804495267 | 1.213739345 |
| CNAG_06351 | SBH1 | protein transporter SEC61 subunit beta | 775.3420188 | 773.5772866 | 903.1538398 | 665.1856465 | 657.1012603 | 714.9998437 | 0.00996753 | -0.282970189 | 0.005047667 | -2.803975938 | 0.82189717 |
| CNAG_02716 | - | hypothetical protein | 441.474176 | 363.2452529 | 391.4809828 | 491.2497645 | 483.1022239 | 563.7135215 | 0.009981235 | 0.347332864 | 0.005055827 | 2.80345513 | 1.272206501 |
| CNAG_07570 | - | prenylated SNARE protein Ykt6p | 54.20832415 | 77.65532486 | 69.43507347 | 96.66476284 | 117.6885484 | 112.9864669 | 0.009981711 | 0.686197138 | 0.005057287 | 2.803362019 | 1.609036595 |
| CNAG_12876 | - | unspecified product | 370.286035 | 442.7387911 | 519.1592338 | 331.2071516 | 344.8538257 | 356.4685762 | 0.010005976 | -0.383315262 | 0.005070803 | -2.802501212 | 0.766673775 |
| CNAG_07630 | - | U3 small nucleolar ribonucleoprotein IMP4 | 435.3910347 | 416.9187989 | 459.5289404 | 402.4329877 | 307.6619959 | 309.9468948 | 0.01001463 | -0.379230169 | 0.005077634 | -2.802066915 | 0.768847743 |
| CNAG_01631 | - | mRNA cap guanine-N7 methyltransferase | 378.4178697 | 335.1903599 | 312.5981522 | 457.2014615 | 432.2702303 | 431.7525111 | 0.01001463 | 0.34937249 | 0.00507658 | 2.802133915 | 1.274006369 |
| CNAG_07839 | - | large subunit ribosomal protein L11, variant | 65876.3813 | 66143.56332 | 70158.71493 | 61698.8454 | 62425.21222 | 61046.71302 | 0.01001604 | -0.14213518 | 0.005079572 | -2.801943821 | 0.906177026 |
| CNAG_04029 | - | Cytochrome P450 | 110.1613899 | 131.5405256 | 129.1301417 | 166.7364075 | 178.7632376 | 191.7998768 | 0.010038338 | 0.517352005 | 0.005092106 | 2.801148645 | 1.431325708 |
| CNAG_04544 | - | hypothetical protein | 271.9163143 | 280.0291434 | 276.1329646 | 216.1216579 | 212.0191603 | 204.4357584 | 0.010099917 | -0.403108695 | 0.005124576 | -2.799096858 | 0.756227021 |
| CNAG_02774 | MAL13 | Fungal Zn(2)-Cys(6) binuclear cluster domain protein | 821.2081372 | 949.2520782 | 920.3971765 | 1055.4986 | 1061.906473 | 1104.48727 | 0.010110826 | 0.244781461 | 0.005131346 | 2.798670557 | 1.184913267 |
| CNAG_07949 | - | hypothetical protein | 11.92034345 | 11.96446302 | 1.247089807 | 21.22074129 | 31.60783902 | 26.38412114 | 0.010122948 | 1.747362363 | 0.005138734 | 2.798205918 | 3.357441734 |
| CNAG_03528 | - | AP-2 complex subunit alpha | 2802.49701 | 2926.219175 | 2609.617594 | 2332.287016 | 2410.743304 | 2592.243519 | 0.01012937 | -0.199781731 | 0.005143231 | -2.797923372 | 0.870682281 |
| CNAG_06499 | DPP101 | hypothetical protein | 433.4997551 | 489.199278 | 440.541629 | 569.1335038 | 653.3114014 | 510.6945866 | 0.010134902 | 0.331667174 | 0.005147278 | 2.797669345 | 1.258466815 |
| CNAG_04698 | - | cytoplasmic protein | 1512.399567 | 1464.325951 | 1605.040311 | 1886.858052 | 1832.208612 | 1673.014984 | 0.010152724 | 0.219326764 | 0.005157569 | 2.797024091 | 1.164190188 |
| CNAG_01592 | STE14 | protein-S-isoprenylcysteine O-methyltransferase | 133.4469817 | 129.4560528 | 150.4432258 | 207.4530273 | 241.1832342 | 158.827905 | 0.010210776 | 0.540665717 | 0.005188306 | 2.795103744 | 1.454643593 |
| CNAG_06967 | APH2 | Phytase | 2434.62493 | 2277.682963 | 2392.012948 | 2040.473402 | 2108.012993 | 2140.444019 | 0.010215373 | -0.191371843 | 0.00519189 | -2.794880543 | 0.875772563 |
| CNAG_03394 | - | hypothetical protein | 240.9082363 | 294.1841503 | 268.8338257 | 336.3753849 | 336.4946266 | 388.3000223 | 0.010229081 | 0.388054389 | 0.005200106 | 2.794369294 | 1.308627405 |
| CNAG_07749 | - | 3-ketodihydrosphingosine reductase tsc10 | 505.5901616 | 525.7343478 | 594.941471 | 377.773421 | 451.8761019 | 468.5373828 | 0.010232808 | -0.340762928 | 0.00520325 | -2.79417384 | 0.789623631 |
| CNAG_04129 | - | hypothetical protein | 45.05358969 | 92.72070892 | 90.7118608 | 117.0147592 | 125.3383213 | 137.3317436 | 0.010240151 | 0.726341569 | 0.005208234 | 2.793864233 | 1.654438391 |
| CNAG_12992 | - | unspecified product | 35.40020006 | 42.1268758 | 30.04764956 | 80.3150656 | 68.46301007 | 53.2763999 | 0.01034527 | 0.895033607 | 0.005262962 | 2.790482043 | 1.859653205 |
| CNAG_12684 | - | unspecified product | 44.55280284 | 83.68033154 | 74.4385326 | 36.89833787 | 35.92827206 | 36.8585872 | 0.010413835 | -0.922681543 | 0.005299115 | -2.788265191 | 0.527527591 |
| CNAG_02041 | - | hypothetical protein | 919.842692 | 469.8024671 | 527.8554348 | 366.815488 | 319.5681321 | 359.9713036 | 0.010429121 | -0.893307391 | 0.005308167 | -2.787712267 | 0.538378465 |
| CNAG_00914 | KRE6 | glucosidase | 3336.443735 | 3689.13809 | 3364.725196 | 3887.651286 | 4023.092284 | 3921.033357 | 0.010429653 | 0.172726047 | 0.005309711 | 2.787618027 | 1.127186351 |
| CNAG_07904 | - | hypothetical protein | 287.7080359 | 187.5218441 | 220.6876775 | 183.5225872 | 141.9825371 | 161.1567915 | 0.010459577 | -0.533654096 | 0.005326223 | -2.786611889 | 0.690802833 |
| CNAG_04262 | - | E3 ubiquitin-protein ligase NRDP1, variant | 391.5361769 | 447.213103 | 471.4097503 | 541.8688517 | 511.5273771 | 614.7783777 | 0.010486875 | 0.33328484 | 0.005341404 | 2.785689339 | 1.259878701 |
| CNAG_03474 | - | efflux protein EncT | 21.50206057 | 22.83824494 | 36.31541932 | 63.92208675 | 43.16353523 | 57.43118872 | 0.010512394 | 1.029759215 | 0.005355685 | 2.784823579 | 2.041683467 |
| CNAG_02365 | - | Amidase | 378.3100138 | 263.0931953 | 293.3992605 | 451.9038029 | 388.2226563 | 417.2793394 | 0.010528107 | 0.410882954 | 0.005364976 | 2.784261497 | 1.329499241 |
| CNAG_12676 | - | unspecified product | 43.92901679 | 109.8979029 | 87.55939775 | 155.7210711 | 130.4740255 | 123.1130533 | 0.010613746 | 0.759362078 | 0.005409913 | 2.781555226 | 1.692741973 |

| | | | | | | | | | | | | | |
|---|---|---|---|---|---|---|---|---|---|---|---|---|---|
| CNAG_03228 | - | universal stress protein | 877.5574355 | 418.0856435 | 451.0983769 | 295.8645828 | 302.3643599 | 293.4160788 | 0.010614496 | -0.990832408 | 0.005411591 | -2.781454507 | 0.503187362 |
| CNAG_06321 | - | hypothetical protein, variant | 15.51162133 | 14.17888282 | 12.95882611 | 40.51053886 | 43.7448802 | 21.19205469 | 0.010633372 | 1.326870477 | 0.005422513 | 2.780800001 | 2.508579183 |
| CNAG_07883 | - | hypothetical protein | 825.5554323 | 871.5775415 | 891.4823257 | 744.6661063 | 760.7075241 | 686.2332172 | 0.010676682 | -0.255550425 | 0.005447207 | -2.779324572 | 0.837667482 |
| CNAG_06674 | - | hypothetical protein | 685.6578943 | 713.3206815 | 696.4596086 | 852.4462724 | 812.5968787 | 854.7124667 | 0.010676682 | 0.251010618 | 0.005446848 | 2.779345966 | 1.190040455 |
| CNAG_02503 | GPX2 | glutathione peroxidase, variant | 1722.276827 | 1649.512608 | 1645.507992 | 1347.688075 | 1459.353788 | 1549.802637 | 0.010700925 | -0.219222276 | 0.005460882 | -2.778510046 | 0.859028394 |
| CNAG_01832 | - | hypothetical protein | 603.9408363 | 659.4287077 | 609.024781 | 770.742639 | 765.2264012 | 734.8669155 | 0.010714598 | 0.263343995 | 0.005469168 | 2.778017444 | 1.200257537 |
| CNAG_12284 | - | unspecified product | 0.230678295 | 3.353109391 | 4.354967716 | 11.09130078 | 19.57621888 | 11.01423516 | 0.010724275 | 2.536273656 | 0.005475417 | 2.777646365 | 5.80088756 |
| CNAG_13181 | - | unspecified product | 73.48650633 | 93.51382055 | 89.42402506 | 57.30270132 | 50.61625729 | 46.88398237 | 0.010764669 | -0.751425258 | 0.005497355 | -2.77634669 | 0.594016431 |
| CNAG_03262 | INM101 | myo-inositol-1(or 4)-monophosphatase | 2045.658826 | 1745.45995 | 1915.30373 | 1615.518103 | 1669.826797 | 1663.098462 | 0.01082159 | -0.221638416 | 0.005527746 | -2.774553992 | 0.857590949 |
| CNAG_02875 | - | hypothetical protein | 920.1073375 | 912.5543352 | 982.1238351 | 815.4964169 | 793.4290587 | 794.9752121 | 0.010821711 | -0.243727686 | 0.005529129 | -2.774472612 | 0.844560286 |
| CNAG_07639 | - | Triacylglycerol lipase | 1023.13972 | 1078.561588 | 1060.109249 | 1290.662249 | 1204.190307 | 1229.989648 | 0.01086823 | 0.221090887 | 0.005554224 | 2.772999296 | 1.165614627 |
| CNAG_07652 | - | hypothetical protein | 1.38988136 | 0.085611256 | 2.215795287 | 11.40030758 | 6.940224024 | 13.00707002 | 0.010881794 | 3.305450127 | 0.005562485 | 2.77251563 | 9.886433225 |
| CNAG_06378 | - | hypothetical protein | 627.0234575 | 653.9398258 | 683.5110213 | 543.809552 | 571.8048828 | 519.7029277 | 0.010895709 | -0.280809486 | 0.005570928 | -2.772021922 | 0.823129036 |
| CNAG_00322 | - | hypothetical protein | 254.8922495 | 303.879504 | 275.2359992 | 359.0292876 | 341.6491256 | 387.2336382 | 0.010957973 | 0.371271547 | 0.005604102 | 2.770088764 | 1.293492373 |
| CNAG_00080 | - | hypothetical protein | 42.61714963 | 56.21210306 | 81.05894509 | 88.07461258 | 108.1876783 | 111.792818 | 0.010997797 | 0.763299528 | 0.005625812 | 2.768829212 | 1.69736817 |
| CNAG_04802 | - | ribosomal RNA-processing protein 9 | 1083.742507 | 936.416251 | 1035.580744 | 1159.708952 | 1217.178916 | 1294.286659 | 0.011006706 | 0.249272054 | 0.005631713 | 2.768487604 | 1.188607224 |
| CNAG_07809 | - | hypothetical protein | 25.8723172 | 15.29947912 | 17.23027141 | 35.99420512 | 46.13032184 | 46.29812697 | 0.011038408 | 1.139024172 | 0.005649282 | 2.767472473 | 2.202320095 |
| CNAG_03445 | - | hypothetical protein | 268.903843 | 287.7848414 | 322.1024344 | 404.3420117 | 385.7432386 | 354.0988021 | 0.011134785 | 0.366951926 | 0.005699966 | 2.76455981 | 1.289625277 |
| CNAG_03973 | - | hydantoinase | 1315.931006 | 1347.789414 | 1261.594236 | 1018.423535 | 1125.533731 | 1215.63413 | 0.011154966 | -0.239372998 | 0.005711659 | -2.76389117 | 0.847113392 |
| CNAG_02760 | - | hypothetical protein | 1243.386697 | 1117.26575 | 1090.970862 | 926.6516027 | 1050.570875 | 955.9651995 | 0.011168746 | -0.250973898 | 0.005721442 | -2.763332653 | 0.840328956 |
| CNAG_04750 | - | hypothetical protein | 124.1227417 | 157.3292514 | 116.4039361 | 191.0340216 | 196.0598119 | 183.0533021 | 0.011168746 | 0.507561565 | 0.005720311 | 2.763397182 | 1.421645307 |
| CNAG_06629 | - | hypothetical protein | 27.92474089 | 3.490279184 | 4.487311907 | 34.75483239 | 38.39131892 | 38.50992775 | 0.011181127 | 1.672264555 | 0.00572915 | 2.76289325 | 3.187144776 |
| CNAG_02551 | DAK102 | dihydroxyacetone kinase 1 | 118.4066005 | 85.07153406 | 88.45306585 | 68.24484401 | 54.88224367 | 56.85350188 | 0.011250409 | -0.725428801 | 0.005766023 | -2.760798532 | 0.604817253 |
| CNAG_12320 | - | unspecified product | 151.754843 | 191.563163 | 168.4098495 | 124.2340597 | 114.464158 | 121.1705167 | 0.011286339 | -0.520764992 | 0.005789951 | -2.75944569 | 0.697002148 |
| CNAG_00285 | - | hypothetical protein | 783.6449037 | 756.4204691 | 675.180018 | 849.1770641 | 964.5766409 | 882.5027484 | 0.011286339 | 0.268237103 | 0.005788884 | 2.759505926 | 1.204335294 |
| CNAG_00442 | - | cyclin | 711.3854781 | 723.050591 | 788.1394314 | 829.6646933 | 938.6530378 | 927.9598346 | 0.011286339 | 0.262550587 | 0.005788309 | 2.759538391 | 1.199597638 |
| CNAG_01519 | EEP1 | endonuclease/exonuclease/phosphatase | 279.1929659 | 365.080769 | 373.0957982 | 262.0340432 | 288.0509294 | 206.7888004 | 0.011286339 | -0.442629475 | 0.005786655 | -2.759631727 | 0.735792321 |
| CNAG_01953 | - | hypothetical protein | 241.8894539 | 358.5393527 | 323.0120786 | 233.9881653 | 232.7141167 | 221.1050633 | 0.011321392 | -0.440399973 | 0.005809316 | -2.758354506 | 0.736930274 |
| CNAG_00175 | - | hypothetical protein | 92.46134451 | 98.11065821 | 59.9496989 | 147.0160302 | 124.3788087 | 124.2047671 | 0.011357796 | 0.651234203 | 0.00583077 | 2.757149424 | 1.570511168 |
| CNAG_04472 | - | membrane protein | 25.97218432 | 21.72912603 | 22.54273693 | 48.80023313 | 48.63485522 | 44.23976628 | 0.011357796 | 1.026024042 | 0.005830522 | 2.757163357 | 2.036404335 |
| CNAG_00720 | - | DNA repair protein RAD51 | 1089.839908 | 1342.496132 | 1284.050747 | 1442.289194 | 1456.953206 | 1534.109094 | 0.011421111 | 0.24020871 | 0.005866064 | 2.755175643 | 1.181163524 |
| CNAG_02340 | - | hypothetical protein | 742.72756 | 888.7497003 | 833.9021525 | 661.270437 | 738.3140012 | 656.2489571 | 0.011421111 | -0.275991912 | 0.005865803 | -2.755190189 | 0.825882295 |
| CNAG_03202 | CAC1 | Adenylate cyclase | 3013.782582 | 2850.847991 | 2705.595182 | 2449.842951 | 2585.028895 | 2565.643197 | 0.011525962 | -0.188580105 | 0.005921324 | -2.752106616 | 0.877468898 |
| CNAG_03120 | AGS1 | alpha-1,3-glucan synthase | 10710.59412 | 9791.759865 | 8264.08515 | 11375.67116 | 11782.75101 | 10745.2262 | 0.011546389 | 0.221480535 | 0.005933228 | 2.751448871 | 1.165929483 |
| CNAG_04294 | - | hypothetical protein | 99.64108509 | 117.5362627 | 114.2062962 | 172.5204762 | 166.7403294 | 145.3756309 | 0.011580887 | 0.534705082 | 0.00595237 | 2.750393722 | 1.44864599 |
| CNAG_12335 | - | unspecified product | 2.756624556 | 9.809697765 | 8.645924841 | 30.67448837 | 18.22838065 | 18.76705896 | 0.01165594 | 1.784206251 | 0.005992369 | 2.748198674 | 3.444289122 |
| CNAG_13184 | - | unspecified product | 0.271793585 | 4.447718863 | 6.486875447 | 15.95457465 | 8.515833925 | 28.16959459 | 0.011662077 | 2.339103353 | 0.005996948 | 2.747948226 | 5.059880641 |
| CNAG_01037 | - | DNA mismatch repair protein MLH3 | 385.6454736 | 428.8859847 | 429.8377214 | 506.8248048 | 491.7142504 | 567.0730304 | 0.011680594 | 0.317844116 | 0.006007896 | 2.747350127 | 1.246466505 |
| CNAG_01546 | - | S-adenosylmethionine-dependent methyltransferase | 382.9071255 | 408.239425 | 406.2302061 | 365.8473665 | 300.83227 | 261.1552779 | 0.011696052 | -0.384205465 | 0.006017275 | -2.746838513 | 0.766200852 |
| CNAG_01365 | - | hypothetical protein | 272.2876079 | 258.70676 | 273.0904059 | 326.9764897 | 333.0327495 | 397.1468128 | 0.011791906 | 0.379877623 | 0.006068029 | 2.744082471 | 1.301231473 |
| CNAG_02029 | WSP1 | wiskott-Aldrich syndrome protein | 312.7960914 | 343.5438688 | 296.4467045 | 266.8278018 | 241.3273928 | 228.8868963 | 0.011846691 | -0.386476546 | 0.006100561 | -2.742326795 | 0.764995653 |
| CNAG_04090 | ATF1 | activating transcription factor | 867.4509871 | 837.1081995 | 803.0196426 | 689.318162 | 731.3692452 | 709.4884837 | 0.011846691 | -0.251153938 | 0.00610049 | -2.742330655 | 0.840224094 |
| CNAG_01199 | - | nuclear pore complex protein Nup62 | 1643.000213 | 1314.606485 | 1343.839362 | 1579.09968 | 2038.602394 | 1699.599997 | 0.011846691 | 0.289485255 | 0.006098136 | 2.742457372 | 1.222204124 |
| CNAG_07305 | - | hypothetical protein | 111.49219 | 154.1889391 | 182.3749574 | 230.0946113 | 236.6981163 | 188.7123031 | 0.01188995 | 0.534249168 | 0.00612429 | 2.741051548 | 1.448188267 |
| CNAG_12150 | - | unspecified product | 7.207310639 | 3.388204742 | 2.274615252 | 14.2288762 | 13.92005903 | 24.99805421 | 0.011967196 | 2.071043479 | 0.006165539 | 2.738845228 | 4.201904809 |
| CNAG_00378 | - | hypothetical protein | 107.3711313 | 134.4699987 | 127.8337366 | 82.92504824 | 93.04759394 | 65.80735987 | 0.011975795 | -0.631504627 | 0.006171432 | -2.738531114 | 0.645502851 |
| CNAG_05388 | - | formamidopyrimidine-DNA glycosylase | 1008.627696 | 950.2066676 | 805.2083005 | 777.3220824 | 787.4161241 | 743.9428158 | 0.012009894 | -0.275569962 | 0.00619047 | -2.737518156 | 0.826123879 |
| CNAG_01509 | - | hypothetical protein | 746.0935363 | 796.1208242 | 739.0191568 | 594.2856238 | 612.2475929 | 698.2993758 | 0.012027487 | -0.27542052 | 0.006201008 | -2.736958707 | 0.826209458 |
| CNAG_12813 | - | unspecified product | 0.132601705 | 0.064423749 | 1.134551593 | 6.430262608 | 8.915892552 | 7.617600397 | 0.012040004 | 4.272763037 | 0.006208931 | 2.736538591 | 19.32991029 |
| CNAG_02968 | PLC2 | Phosphatidylinositol-specific phospholipase C, putative | 325.1436744 | 257.425857 | 215.4664815 | 191.8934182 | 206.0126905 | 183.381124 | 0.012067622 | -0.474628935 | 0.006226121 | -2.73562885 | 0.719651863 |
| CNAG_07333 | - | ubiquitin carboxyl-terminal hydrolase 25/28 | 2711.575786 | 2608.51734 | 2425.202637 | 2830.662346 | 3024.248067 | 3086.135345 | 0.012067622 | 0.191887017 | 0.006225398 | 2.735667071 | 1.142256788 |
| CNAG_06655 | GPI8 | GPI-anchor transamidase | 357.7826499 | 508.5464774 | 456.5054794 | 556.6897477 | 568.5914832 | 569.3799853 | 0.012073538 | 0.344010202 | 0.006230647 | 2.735389654 | 1.269279862 |
| CNAG_01500 | - | Taurine dioxygenase | 3.860793879 | 7.653621156 | 5.459920731 | 13.25800025 | 25.62139521 | 18.66961407 | 0.01207528 | 1.886656834 | 0.006233021 | 2.735264295 | 3.697773438 |
| CNAG_03032 | - | dynactin 2 | 1472.332078 | 1328.466026 | 1398.07806 | 1232.676213 | 1130.631548 | 1258.968457 | 0.012092771 | -0.229233956 | 0.006243526 | -2.734709969 | 0.853087745 |
| CNAG_12548 | - | unspecified product | 8.564084681 | 8.799051728 | 13.95030721 | 26.92554808 | 25.23245849 | 29.66866135 | 0.012101174 | 1.448501012 | 0.006249342 | 2.734403428 | 2.729243304 |
| CNAG_05321 | - | transposase subfamily, variant | 262.7030961 | 296.1777384 | 301.6925532 | 214.4811503 | 218.9090912 | 228.8001515 | 0.012169969 | -0.395024467 | 0.006286356 | -2.732458662 | 0.760476479 |
| CNAG_05708 | UBP14 | ubiquitin carboxyl-terminal hydrolase 5/13 | 1084.528014 | 885.6741096 | 998.1403557 | 774.0705479 | 822.7484856 | 881.5078223 | 0.012171045 | -0.276465145 | 0.006288398 | -2.732351659 | 0.825611433 |
| CNAG_12263 | - | unspecified product | 2.757941231 | 12.98254773 | 5.486632886 | 23.93371633 | 24.53694569 | 19.84461349 | 0.012197934 | 1.78564711 | 0.00630378 | 2.73154673 | 3.447730745 |
| CNAG_00837 | - | hypothetical protein | 331.4936817 | 333.929614 | 348.6348089 | 263.570571 | 240.3945986 | 287.5856891 | 0.012240025 | -0.372340263 | 0.006327027 | -2.730333569 | 0.772528326 |
| CNAG_07357 | - | hypothetical protein | 6.35309728 | 18.40851576 | 13.99981527 | 33.20096051 | 44.72602208 | 22.24391106 | 0.012257032 | 1.406737102 | 0.006337315 | 2.729797966 | 2.651368326 |
| CNAG_02728 | - | peroxin-13 | 229.1012809 | 210.2492337 | 245.336245 | 302.2479792 | 317.6681337 | 288.562178 | 0.012293951 | 0.391350181 | 0.006357905 | 2.72872838 | 1.311620342 |
| CNAG_00372 | - | chromatin structure-remodeling complex protein RSC7 | 2268.049193 | 2576.183909 | 2664.01634 | 2761.399986 | 2961.2905 | 3007.312264 | 0.012430531 | 0.202556807 | 0.006430056 | 2.725004747 | 1.150735933 |
| CNAG_08004 | - | hypothetical protein | 26.04783912 | 18.55690417 | 33.13301872 | 46.97310629 | 56.47556526 | 54.12408061 | 0.012439782 | 1.001919566 | 0.006436361 | 2.724681179 | 2.002662855 |
| CNAG_05874 | - | hypothetical protein | 921.40248 | 126.3929826 | 252.7803618 | 99.30037592 | 89.86341739 | 97.13024212 | 0.012462982 | -2.211577425 | 0.006449886 | -2.723987952 | 0.215898118 |
| CNAG_02012 | - | TPR repeat-containing protein | 659.3022705 | 461.2167958 | 589.6519181 | 433.9599247 | 457.0122318 | 457.5014513 | 0.01260738 | -0.36020016 | 0.006526155 | -2.720103174 | 0.779056486 |
| CNAG_02730 | - | sorting nexin-41 | 413.6217898 | 462.2562768 | 405.3567005 | 508.3181462 | 508.9714088 | 598.1257096 | 0.0126325 | 0.319970641 | 0.006540701 | 2.719366914 | 1.248305146 |
| CNAG_05927 | - | ATP synthase mitochondrial F1 complex assembly factor 1 | 772.7468421 | 618.4870436 | 765.6000293 | 619.404924 | 501.7111959 | 613.9843895 | 0.012654049 | -0.330392412 | 0.006553403 | -2.718725173 | 0.795320128 |
| CNAG_05729 | - | hypothetical protein | 1.363983309 | 1.172321606 | 0.065918203 | 8.040881213 | 9.754138872 | 9.786600956 | 0.012663807 | 3.685757653 | 0.006560004 | 2.718392155 | 12.86837209 |
| CNAG_06249 | - | taurine catabolism dioxygenase TauD, variant | 96.22348285 | 111.1310402 | 139.7300062 | 160.5296197 | 152.7311963 | 206.0656716 | 0.012720869 | 0.567471896 | 0.006591116 | 2.716826465 | 1.481924446 |
| CNAG_01891 | - | RAD57 protein | 323.5872399 | 305.006386 | 279.5336691 | 416.0161081 | 358.8577186 | 399.5074686 | 0.012749056 | 0.355867512 | 0.006607277 | 2.716015768 | 1.279754884 |
| CNAG_03945 | - | hypothetical protein | 99.33826565 | 119.5051266 | 162.8386793 | 93.18156405 | 75.59363756 | 75.73706964 | 0.012756677 | -0.660056256 | 0.006612785 | -2.715739894 | 0.632853619 |
| CNAG_13095 | - | unspecified product | 65.81465206 | 79.8419179 | 80.07884306 | 113.9586737 | 107.9485264 | 129.5708402 | 0.012814549 | 0.625241777 | 0.006644349 | 2.714162856 | 1.542469301 |
| CNAG_01002 | - | hypothetical protein | 321.4822278 | 373.9432611 | 393.5557727 | 435.9444029 | 464.2130155 | 486.0686702 | 0.012825018 | 0.334731289 | 0.006651343 | 2.713814336 | 1.261142491 |
| CNAG_01841 | GLN3 | GATA family transcription factor | 32.96858263 | 31.38870769 | 24.70897676 | 61.36133901 | 61.65197132 | 46.58055935 | 0.012839188 | 0.933989323 | 0.00666026 | 2.713370477 | 1.910551733 |
| CNAG_02316 | - | hypothetical protein | 238.3334827 | 204.8531261 | 205.9440814 | 269.3897372 | 336.8167473 | 271.9292068 | 0.012845809 | 0.421103682 | 0.006665263 | 2.713121655 | 1.33895148 |
| CNAG_00048 | - | Sugar transporter | 50.39052037 | 34.69505417 | 40.67000922 | 70.38282069 | 64.79656064 | 88.37527238 | 0.012872274 | 0.824154414 | 0.006680567 | 2.712361596 | 1.770497018 |
| CNAG_01622 | - | CTD kinase subunit beta | 761.676888 | 843.7063699 | 928.8886383 | 992.4432529 | 1068.039209 | 996.858351 | 0.012978336 | 0.256236585 | 0.006738782 | 2.709484676 | 1.194359032 |
| CNAG_12290 | - | unspecified product | 13.42365954 | 21.74296774 | 33.11203376 | 54.39941728 | 68.08752208 | 30.11364328 | 0.012978336 | 1.166157508 | 0.006738137 | 2.70951643 | 2.244131954 |
| CNAG_02455 | - | choline transporter | 2652.004468 | 3261.227794 | 2719.455242 | 2490.347333 | 2576.399862 | 2460.218807 | 0.012987979 | -0.21206542 | 0.006745375 | -2.709160265 | 0.863300411 |
| CNAG_02850 | - | glucan endo-1,3-alpha-glucosidase agn1 | 7852.88253 | 6962.3561 | 6757.058078 | 6383.385303 | 6356.113168 | 6477.849933 | 0.012989523 | -0.182474751 | 0.006747763 | -2.709042813 | 0.881190135 |
| CNAG_04534 | - | hypothetical protein, variant | 258.0056426 | 346.627744 | 243.1617889 | 219.2471833 | 225.0144717 | 179.0035053 | 0.01317779 | -0.4571121 | 0.006848782 | -2.704109015 | 0.728442953 |
| CNAG_00887 | - | hypothetical protein | 2.601143298 | 2.284036419 | 3.299028063 | 17.13871377 | 13.19496246 | 8.864078127 | 0.01317779 | 2.424780845 | 0.006848628 | 2.704116482 | 5.36947429 |
| CNAG_02873 | - | signal peptidase I | 181.283461 | 203.6972148 | 179.2811315 | 247.7228315 | 279.8023545 | 236.4053998 | 0.013195795 | 0.421681711 | 0.006861362 | 2.703499168 | 1.33948805 |
| CNAG_01041 | - | hypothetical protein | 140.3474352 | 142.3137284 | 124.9127121 | 191.794711 | 184.7401378 | 195.2019178 | 0.013195795 | 0.474500095 | 0.006861138 | 2.70351003 | 1.389436685 |
| CNAG_02915 | PDK1 | AGC/PDK1 protein kinase | 1712.054619 | 1517.161731 | 1826.845926 | 1864.790759 | 2102.244489 | 2034.819622 | 0.013263321 | 0.23124958 | 0.006898093 | 2.701724328 | 1.173851232 |
| CNAG_00508 | VPS17 | vacuolar protein sorting-associated protein vps17 | 284.1256454 | 379.2425442 | 299.792569 | 433.0846449 | 416.7768777 | 400.6872418 | 0.013327903 | 0.362632606 | 0.006933309 | 2.700030652 | 1.28577001 |
| CNAG_00464 | - | pre-mRNA-splicing factor rse1 | 10544.58613 | 9419.987347 | 9434.908219 | 8416.208458 | 8994.731895 | 8934.626849 | 0.013370471 | -0.173957744 | 0.006957086 | -2.69889147 | 0.886407661 |

| | | | | | | | | | | | | | |
|---|---|---|---|---|---|---|---|---|---|---|---|---|---|
| CNAG_00399 | - | transformer-2-beta isoform 3 | 810.7419512 | 995.56301 | 937.4626428 | 1159.969553 | 1089.513785 | 1051.284728 | 0.013371078 | 0.252327173 | 0.006959035 | 2.698798265 | 1.191126942 |
| CNAG_00369 | - | hypothetical protein | 962.1888339 | 941.6746674 | 1144.157621 | 845.7858558 | 784.7131949 | 914.8017909 | 0.01338362 | -0.27621396 | 0.006968831 | -2.698330054 | 0.825755192 |
| CNAG_02487 | PHS1 | 3-hydroxy acyl-CoA dehydratase | 87.91358139 | 84.21238948 | 101.3839958 | 161.9031471 | 121.6776627 | 131.9784623 | 0.01338362 | 0.587207578 | 0.006967856 | 2.698376616 | 1.502336071 |
| CNAG_12417 | - | unspecified product | 47.58429465 | 62.13723836 | 41.54756081 | 22.80736535 | 15.93254463 | 35.56676017 | 0.013420444 | -1.071771389 | 0.006991282 | -2.697259216 | 0.475734518 |
| CNAG_00735 | - | aldehyde dehydrogenase family 7 member A1 | 662.0840048 | 794.9702544 | 673.9740607 | 546.8046408 | 526.7795987 | 660.5277268 | 0.013420444 | -0.311826259 | 0.006990451 | -2.697298827 | 0.805621304 |
| CNAG_00275 | - | hypothetical protein | 332.9948396 | 372.8011884 | 268.9187065 | 452.6212878 | 406.3674773 | 411.7734652 | 0.01345501 | 0.370240953 | 0.007010933 | 2.696324511 | 1.292568692 |
| CNAG_00066 | - | membrane protein | 1639.406765 | 1549.33948 | 1673.203935 | 1461.527463 | 1449.028616 | 1346.755285 | 0.013548488 | -0.207372794 | 0.007061295 | -2.693939638 | 0.866113023 |
| CNAG_00461 | - | protein kinase C substrate 80K-H | 4219.046973 | 4348.473349 | 4266.855661 | 3817.353707 | 3950.474066 | 3897.833775 | 0.013551784 | -0.152935272 | 0.007064668 | -2.693780459 | 0.899418664 |
| CNAG_01580 | SCP1 | sterol regulatory element-binding protein cleavage-activating protein | 1450.603635 | 1716.35236 | 1551.753889 | 1752.749267 | 1861.51911 | 1911.574492 | 0.013575619 | 0.213353475 | 0.007078751 | 2.693116579 | 1.159379977 |
| CNAG_07876 | - | hypothetical protein | 243.1444339 | 268.3369551 | 240.0581809 | 338.9091622 | 319.2750327 | 320.729526 | 0.013612538 | 0.36475128 | 0.007099664 | 2.692132935 | 1.287659618 |
| CNAG_12616 | - | unspecified product | 46.98612259 | 26.14504824 | 54.44186684 | 82.34784187 | 85.84660078 | 64.34321998 | 0.013628169 | 0.856861994 | 0.007109481 | 2.691672095 | 1.811094711 |
| CNAG_12101 | - | unspecified product | 43.61234282 | 32.19019082 | 27.68446999 | 14.29809351 | 19.68472701 | 10.29323158 | 0.013672512 | -1.26294593 | 0.007134283 | -2.690510322 | 0.41669222 |
| CNAG_02148 | - | ubiquitin-conjugating enzyme E2 35, variant | 557.8503781 | 446.086333 | 537.3893307 | 463.8584373 | 388.8051401 | 368.7837653 | 0.013688027 | -0.353441238 | 0.00714405 | -2.690053797 | 0.782714872 |
| CNAG_00273 | - | hypothetical protein | 92.57704658 | 109.9588551 | 86.53720217 | 143.5728661 | 157.3582066 | 128.7190536 | 0.013808571 | 0.561831125 | 0.007208651 | 2.687048389 | 1.476141605 |
| CNAG_01478 | - | monooxygenase | 406.1308364 | 368.4488988 | 398.7693726 | 331.4248881 | 269.6864832 | 324.2505786 | 0.013854372 | -0.358402925 | 0.007234253 | -2.685864005 | 0.780027599 |
| CNAG_02547 | - | nuclear pore complex protein Nup205 | 8959.946621 | 8388.846627 | 7723.275122 | 6966.982549 | 7599.963297 | 7730.083196 | 0.01387165 | -0.184680435 | 0.007244968 | -2.685369395 | 0.879843945 |
| CNAG_06544 | - | non-histone chromosomal protein 6 | 4808.814508 | 5053.123761 | 5286.347135 | 4717.939296 | 4480.04066 | 4457.382858 | 0.013906661 | -0.164813366 | 0.007264952 | -2.684448743 | 0.892043911 |
| CNAG_12083 | - | unspecified product | 196.100708 | 188.541712 | 259.9621364 | 163.1965514 | 138.5675797 | 161.1045802 | 0.013939779 | -0.49245485 | 0.007285658 | -2.683497215 | 0.710814566 |
| CNAG_02787 | - | dephospho-CoA kinase | 1037.904944 | 916.8933343 | 1015.190605 | 709.330169 | 884.9435162 | 879.2895 | 0.013939779 | -0.279717644 | 0.007284496 | -2.683550542 | 0.823752222 |
| CNAG_07796 | - | uroporphyrinogen-III C-methyltransferase | 161.0666245 | 157.2838964 | 222.6007342 | 135.2009094 | 91.84611059 | 139.9600652 | 0.014058218 | -0.580256337 | 0.007349277 | -2.680588743 | 0.668844927 |
| CNAG_07869 | - | hypothetical protein | 7.177596361 | 2.310891856 | 2.267486531 | 12.61097584 | 22.16234745 | 14.29415396 | 0.014197979 | 2.106604953 | 0.007424074 | 2.677198013 | 4.30676604 |
| CNAG_00812 | IRR1 | cohesin complex subunit SA-1/2 | 3145.811947 | 2985.555959 | 3133.198385 | 2677.895297 | 2730.7673 | 2880.893698 | 0.014360669 | -0.176074344 | 0.007510898 | -2.673300292 | 0.885108152 |
| CNAG_05784 | - | hypothetical protein, variant | 374.9774684 | 351.3438237 | 329.6523606 | 475.1157202 | 517.0402949 | 378.640475 | 0.014360997 | 0.362527314 | 0.007512823 | 2.673214327 | 1.285676174 |
| CNAG_03422 | CTS1 | calcineurin temperature suppressor Cts1 | 92.57740929 | 96.02472271 | 98.21855126 | 113.5092979 | 160.8451195 | 158.4486012 | 0.014399151 | 0.576418803 | 0.007534542 | 2.672245887 | 1.491143191 |
| CNAG_06048 | - | hypothetical protein | 1146.704469 | 1208.823374 | 1233.823702 | 1073.93693 | 999.5568717 | 1045.849695 | 0.014400452 | -0.217628487 | 0.007536981 | -2.672137272 | 0.859977913 |
| CNAG_00191 | - | etoposide-induced protein | 369.1950925 | 393.3070333 | 340.3136397 | 459.4037266 | 452.0897854 | 473.8957293 | 0.014410741 | 0.313895196 | 0.007544126 | 2.671819321 | 1.243059364 |
| CNAG_05568 | PRY1 | septum formation protein Maf | 600.0418322 | 602.197177 | 566.2555932 | 533.8813985 | 421.5107021 | 490.7876333 | 0.014412993 | -0.306966061 | 0.007547065 | -2.671688607 | 0.808339885 |
| CNAG_12960 | - | unspecified product | 75.3855572 | 64.52896204 | 62.83640173 | 40.07173116 | 41.17572588 | 35.77473723 | 0.014443602 | -0.808943611 | 0.007564856 | -2.670898296 | 0.570799663 |
| CNAG_04196 | - | actin binding protein | 5097.086819 | 4824.71854 | 4705.188928 | 4394.870291 | 4392.939308 | 4441.8104 | 0.014465366 | -0.160471634 | 0.007578022 | -2.670314553 | 0.894732524 |
| CNAG_07440 | - | nuclear distribution protein PAC1 | 187.2072759 | 206.9841605 | 236.7820451 | 299.2988303 | 290.9068092 | 256.415269 | 0.014473808 | 0.410672555 | 0.007584212 | 2.670040418 | 1.329305365 |
| CNAG_05513 | - | septum-promoting GTP-binding protein 1 | 941.35414 | 842.6887179 | 1007.814927 | 1146.664854 | 1054.076208 | 1157.765193 | 0.014515272 | 0.251224596 | 0.007607711 | 2.669001485 | 1.190216972 |
| CNAG_04238 | - | peroxisomal 2,4-dienoyl-CoA reductase | 244.3396212 | 232.8802688 | 292.2226423 | 330.2170451 | 344.3631219 | 334.0351374 | 0.014554584 | 0.376649237 | 0.007630092 | 2.668014678 | 1.298322903 |
| CNAG_03776 | - | hypothetical protein | 196.7083784 | 271.5956834 | 307.1384237 | 366.2135978 | 330.461374 | 352.8725063 | 0.01461872 | 0.422928387 | 0.007665501 | 2.666458774 | 1.340646042 |
| CNAG_04899 | - | cytoplasmic protein, variant | 1700.41653 | 1661.494171 | 1850.312567 | 1973.835183 | 1956.33211 | 2139.136903 | 0.014632527 | 0.204316942 | 0.007674527 | 2.666063168 | 1.152140725 |
| CNAG_12354 | - | unspecified product | 7.267112446 | 7.657170475 | 2.289009764 | 24.19726617 | 16.54059055 | 17.60770441 | 0.014664604 | 1.818600957 | 0.007693142 | 2.66524865 | 3.527389665 |
| CNAG_06649 | - | Haloacid dehalogenase, type II | 181.2105773 | 182.1796208 | 164.362169 | 228.9992337 | 275.5657829 | 218.6627287 | 0.014699619 | 0.436762168 | 0.007713306 | 2.664368321 | 1.353563122 |
| CNAG_12013 | - | unspecified product | 3.910657968 | 7.688043431 | 10.7442488 | 23.95933244 | 20.03898833 | 21.98251203 | 0.014708207 | 1.664008396 | 0.007719608 | 2.664093586 | 3.168957685 |
| CNAG_03199 | - | FAD dependent oxidoreductase | 28.22894423 | 58.89836109 | 54.144889 | 25.2965625 | 25.64679127 | 16.97035981 | 0.014714347 | -1.098215033 | 0.007726425 | -2.663796672 | 0.467094048 |
| CNAG_04162 | PKA2 | AGC/PKA protein kinase | 473.164964 | 495.6936777 | 502.3561801 | 632.2950826 | 621.2162928 | 555.073659 | 0.014714347 | 0.283358895 | 0.007725983 | 2.663815892 | 1.217025077 |
| CNAG_04295 | - | hypothetical protein | 823.0932891 | 787.5607058 | 789.1377074 | 698.7676937 | 645.0171525 | 696.1426158 | 0.014723609 | -0.249856914 | 0.007733086 | -2.663506742 | 0.840979819 |
| CNAG_02141 | - | hypothetical protein | 190.372138 | 223.9942515 | 249.3699633 | 158.4107504 | 170.6388513 | 161.1379555 | 0.014769228 | -0.451950538 | 0.00775885 | -2.662387521 | 0.731053788 |
| CNAG_00606 | - | transcription initiation factor TFIID subunit 7 | 1061.661179 | 1402.675463 | 1382.0141 | 1094.882806 | 1050.443928 | 1082.503177 | 0.014820101 | -0.267407663 | 0.007787385 | -2.661151769 | 0.830811066 |
| CNAG_05648 | - | glutaminyl cyclase | 1452.6094 | 1525.568791 | 1447.150408 | 1254.3914 | 1207.40962 | 1388.8132152 | 0.014832074 | -0.215689787 | 0.007795487 | -2.66080162 | 0.861134332 |
| CNAG_01294 | IPK1 | inositol-pentakisphosphate 2-kinase, variant | 440.5294243 | 503.2101484 | 488.4949813 | 612.0350964 | 618.6585664 | 541.7529714 | 0.01483784 | 0.292618206 | 0.007802142 | 2.660514294 | 1.224861142 |
| CNAG_13182 | - | unspecified product | 3.763650778 | 4.421287848 | 2.254502328 | 13.59199404 | 14.95792642 | 14.23626262 | 0.01483784 | 2.173469689 | 0.007801949 | 2.660522619 | 4.511070051 |
| CNAG_13192 | - | unspecified product | 11.97866435 | 7.740391909 | 13.9575073 | 25.21945236 | 26.10272721 | 31.84417873 | 0.014853485 | 1.386921679 | 0.007812182 | 2.660081169 | 2.61520072 |
| CNAG_04945 | - | hypothetical protein | 168.3470624 | 167.0945932 | 146.2440716 | 218.1915049 | 247.0588201 | 199.7938967 | 0.014876619 | 0.449750782 | 0.007827983 | 2.659400577 | 1.365804301 |
| CNAG_01860 | - | ornithine cyclodeaminase | 741.2790822 | 642.1555131 | 719.7818748 | 574.9193441 | 545.7967615 | 626.1654494 | 0.014876619 | -0.284223678 | 0.007827792 | -2.659408796 | 0.821183373 |
| CNAG_03709 | - | hypothetical protein | 271.2879067 | 340.5048932 | 317.8730489 | 433.9703219 | 395.1906106 | 370.7569917 | 0.014895039 | 0.35395339 | 0.007839494 | 2.658905534 | 1.278058072 |
| CNAG_02983 | - | hypothetical protein | 1255.472372 | 1226.198013 | 1288.315295 | 1407.206215 | 1470.766415 | 1511.925703 | 0.014926659 | 0.204022752 | 0.007857959 | 2.658112791 | 1.151905808 |
| CNAG_01265 | - | charged multivesicular body protein 6 | 671.6543731 | 672.4080429 | 716.6976347 | 774.4247942 | 816.0732308 | 901.253656 | 0.014939322 | 0.259576417 | 0.00786645 | 2.657748815 | 1.19712717 |
| CNAG_01026 | - | gamma-glutamyltransferase | 261.657626 | 236.0600915 | 208.1045098 | 332.7729386 | 308.9540582 | 293.0183981 | 0.014971435 | 0.388789965 | 0.007885187 | 2.65694686 | 1.309294795 |
| CNAG_12755 | - | unspecified product | 5.1311185 | 14.10142431 | 10.79256458 | 31.94151956 | 26.13979178 | 23.19803479 | 0.014983122 | 1.451313057 | 0.007893172 | 2.656605621 | 2.734568226 |
| CNAG_12925 | - | unspecified product | 1.644077135 | 19.41711481 | 8.698231198 | 33.46842558 | 26.96783106 | 29.71569524 | 0.015003399 | 1.64352934 | 0.007905686 | 2.656071447 | 3.124292096 |
| CNAG_02298 | - | niemann-Pick C1 protein | 2151.147593 | 2315.41797 | 2072.198886 | 2526.484544 | 2392.761331 | 2604.323959 | 0.015048386 | 0.187563258 | 0.007931229 | 2.654983487 | 1.138838568 |
| CNAG_12328 | - | unspecified product | 35.35226543 | 41.03361642 | 24.73789627 | 65.13316291 | 76.48798833 | 48.85691745 | 0.015059687 | 0.903598934 | 0.007939024 | 2.654652095 | 1.870726865 |
| CNAG_02126 | - | hypothetical protein | 1149.118859 | 1367.141512 | 1282.888667 | 1167.401897 | 1021.951628 | 1064.753521 | 0.015154843 | -0.238348763 | 0.007992889 | -2.652370045 | 0.84771501 |
| CNAG_05842 | - | Cytochrome P450 | 207.0813025 | 282.2817339 | 225.1396412 | 302.1431459 | 316.7325674 | 331.7423934 | 0.015154843 | 0.396537788 | 0.007991576 | 2.652425504 | 1.316345123 |
| CNAG_06671 | YKL1 | TKL protein kinase | 2134.749621 | 2077.3543 | 2086.03577 | 2313.899945 | 2511.92654 | 2381.2411 | 0.015174113 | 0.178888585 | 0.008004905 | 2.651862844 | 1.132011477 |
| CNAG_12278 | - | unspecified product | 5.117000308 | 8.787628507 | 14.9864954 | 21.18921127 | 25.28260627 | 31.77659738 | 0.015179479 | 1.49716206 | 0.008009589 | 2.651665311 | 2.822868765 |
| CNAG_06901 | ALG7 | UDP-N-acetylglucosamine-dolichyl-phosphate N- acetylglucosaminephosphotransferase | 280.4988912 | 313.5725742 | 270.9906121 | 355.050982 | 374.5238883 | 377.3079887 | 0.015194038 | 0.343181884 | 0.008019127 | 2.651263426 | 1.268551318 |
| CNAG_06272 | - | hypothetical protein | 357.1614466 | 341.5036313 | 377.4156264 | 319.0736847 | 259.3832156 | 266.6147887 | 0.015239261 | -0.364422498 | 0.008044856 | -2.650181428 | 0.776779753 |
| CNAG_07399 | - | hypothetical protein | 19.0419461 | 24.87110458 | 14.05566312 | 41.75130021 | 40.82101177 | 41.94045866 | 0.015245887 | 1.083795588 | 0.008050216 | 2.649956406 | 2.119605224 |
| CNAG_06004 | - | transcription initiation factor TFIIH subunit 3 | 198.8119359 | 220.9488563 | 211.2454483 | 262.4123283 | 265.7660918 | 310.572856 | 0.0153202 | 0.397469769 | 0.008091326 | 2.648235011 | 1.317195756 |
| CNAG_01171 | - | hypothetical protein | 422.7789839 | 466.4776998 | 574.6080753 | 338.0922389 | 448.5665877 | 349.9042466 | 0.015448724 | -0.380844474 | 0.008161092 | -2.645331518 | 0.767987922 |
| CNAG_06687 | - | hypothetical protein | 4040.504656 | 3733.395624 | 4000.30534 | 4369.38417 | 4304.197196 | 4840.145515 | 0.015587523 | 0.18338196 | 0.008236319 | 2.64222555 | 1.135542703 |
| CNAG_12635 | - | unspecified product | 2.565026982 | 2.267419187 | 1.170491985 | 16.45124826 | 12.36457677 | 5.613566321 | 0.01559198 | 2.707020289 | 0.008242481 | 2.641972227 | 6.529716198 |
| CNAG_00636 | CDC7 | CDC7 protein kinase | 1398.870551 | 1395.232557 | 1397.016598 | 1200.702038 | 1192.79021 | 1285.597052 | 0.01559198 | -0.20358321 | 0.008240704 | -2.642045264 | 0.868391066 |
| CNAG_01096 | - | hypothetical protein | 156.7038451 | 132.71134 | 167.4889956 | 247.4364783 | 199.350881 | 194.2274276 | 0.015624706 | 0.476302981 | 0.00826169 | 2.641183745 | 1.391174103 |
| CNAG_05655 | CML1 | EF-hand calcium-binding protein | 700.5266867 | 730.4058789 | 676.1067945 | 538.2116566 | 623.5744462 | 607.3301638 | 0.015679295 | -0.267473256 | 0.008292469 | -2.63992373 | 0.830773293 |
| CNAG_03217 | - | hypothetical protein | 659.880728 | 632.5279603 | 600.5059707 | 774.6220375 | 807.5608976 | 709.3839646 | 0.015683857 | 0.260715924 | 0.008296797 | 2.639746876 | 1.19807309 |
| CNAG_02000 | - | Short-chain dehydrogenase | 691.3972135 | 736.9574732 | 480.1117116 | 807.3184208 | 790.2168212 | 841.3351839 | 0.015757426 | 0.339988995 | 0.008337639 | 2.638082119 | 1.265746939 |
| CNAG_12683 | - | unspecified product | 169.4108367 | 232.5113319 | 219.5520972 | 150.6707185 | 129.9187831 | 167.7006144 | 0.01578083 | -0.489052938 | 0.00835195 | -2.637500525 | 0.712492662 |
| CNAG_01977 | - | hypothetical protein | 30.72089471 | 27.13748854 | 37.40945254 | 61.25684848 | 50.99915695 | 64.07138208 | 0.015825169 | 0.881177934 | 0.008377349 | 2.636470492 | 1.841878548 |
| CNAG_01430 | - | hypothetical protein | 574.8551269 | 689.5900917 | 672.9753859 | 790.955422 | 820.4843548 | 742.6660397 | 0.015832405 | 0.266817342 | 0.008383112 | 2.636237136 | 1.203150686 |
| CNAG_00540 | - | pantothenate transporter | 7.469857927 | 8.822619905 | 19.23609003 | 29.22401499 | 36.77432738 | 25.4363903 | 0.015899668 | 1.388063903 | 0.008420669 | 2.63472009 | 2.617272071 |
| CNAG_08022 | PHO85 | CMGC/CDK/CDK5 protein kinase, variant | 1500.556361 | 1596.702454 | 1632.678868 | 1414.033882 | 1409.346482 | 1346.732542 | 0.015921734 | -0.196631904 | 0.0084343 | -2.634170995 | 0.872585313 |
| CNAG_01771 | - | hypothetical protein | 432.9867867 | 416.8841057 | 388.1680375 | 308.6137695 | 344.9178316 | 344.2369814 | 0.015927088 | -0.326786508 | 0.008439081 | -2.633978587 | 0.797310455 |
| CNAG_12738 | - | unspecified product | 2.650180135 | 7.604915926 | 1.198446113 | 14.34524152 | 10.35841481 | 22.79848303 | 0.015943635 | 2.191996021 | 0.008449795 | 2.633547747 | 4.569372386 |
| CNAG_02215 | HAP3 | transcriptional activator, variant | 311.6584198 | 305.944827 | 349.6583815 | 219.7826385 | 288.0961524 | 237.7491303 | 0.015945421 | -0.390992242 | 0.008452689 | -2.633431458 | 0.762604927 |
| CNAG_03183 | FZC24 | nuclear protein | 1572.896262 | 1580.558984 | 1652.940955 | 1303.337778 | 1389.475544 | 1505.384178 | 0.016060317 | -0.210083699 | 0.008515557 | -2.630914039 | 0.864487076 |
| CNAG_02596 | - | hypothetical protein | 8.495610644 | 7.70429452 | 8.658879791 | 17.99380724 | 34.40972442 | 18.80437395 | 0.016107209 | 1.568295404 | 0.008542387 | 2.629844736 | 2.965541176 |
| CNAG_12238 | - | unspecified product | 84.57097405 | 133.5999937 | 100.383054 | 143.3553976 | 169.4028109 | 159.6671496 | 0.016175858 | 0.5576192 | 0.00858077 | 2.628320198 | 1.471838319 |
| CNAG_02392 | - | hypothetical protein | 165.0976648 | 210.1803359 | 221.8625252 | 235.0219386 | 298.7888173 | 280.6801729 | 0.016178029 | 0.433213446 | 0.008583897 | 2.628196274 | 1.350237738 |
| CNAG_03849 | ASG1 | nuclear protein | 1298.648861 | 1280.049807 | 1241.421005 | 1472.656935 | 1572.631517 | 1405.451941 | 0.016204669 | 0.204873861 | 0.008600011 | 2.627558276 | 1.152585569 |
| CNAG_04120 | FCP1 | RNA polymerase II subunit A domain phosphatase | 1241.180978 | 1328.349286 | 1112.330108 | 1015.423578 | 977.9499278 | 1143.468315 | 0.016212432 | -0.246198534 | 0.008610888 | -2.627128217 | 0.843115078 |
| CNAG_06765 | LMP1 | Low mating performance protein | 1089.691784 | 1095.82774 | 1172.060124 | 1260.11664 | 1318.962549 | 1346.514672 | 0.016336998 | 0.210291849 | 0.00867423 | 2.624633434 | 1.156922199 |

| | | | | | | | | | | | | | |
|---|---|---|---|---|---|---|---|---|---|---|---|---|---|
| CNAG_05970 | PAK1 | ste/ste20/paka protein kinase | 777.8733106 | 828.570944 | 719.9576997 | 910.7370796 | 956.7400289 | 905.8295787 | 0.016359632 | 0.238854295 | 0.008688245 | 2.624083619 | 1.180055158 |
| CNAG_05119 | - | GABA permease | 228.0691649 | 133.6787419 | 174.8030435 | 102.1339433 | 143.1219338 | 116.7785166 | 0.016402528 | -0.585303691 | 0.008713029 | -2.623113298 | 0.666509022 |
| CNAG_04949 | - | hypothetical protein | 885.106978 | 935.1434615 | 930.9588025 | 796.0315125 | 827.1452911 | 739.5147771 | 0.016434455 | -0.234605216 | 0.008731996 | -2.622372398 | 0.849917541 |
| CNAG_00388 | - | 1-phosphatidylinositol-4-phosphate 5-kinase | 433.3954294 | 440.7324722 | 398.960308 | 582.6258682 | 529.7474936 | 478.4864146 | 0.016472546 | 0.307009283 | 0.008754246 | 2.621505064 | 1.237140439 |
| CNAG_02364 | FZC19 | hypothetical protein | 423.9043152 | 500.8730749 | 481.9523917 | 388.1773388 | 410.5294875 | 332.162617 | 0.016508654 | -0.328094521 | 0.008775451 | -2.620680296 | 0.796587905 |
| CNAG_04325 | - | extradiol ring-cleavage dioxygenase | 506.4923482 | 460.0039016 | 431.9050723 | 367.8458342 | 374.1934658 | 395.2931415 | 0.016545778 | -0.313814851 | 0.008797206 | -2.619836014 | 0.804511612 |
| CNAG_01801 | - | hypothetical protein | 15.24399113 | 5.577396084 | 0.145472468 | 16.32447425 | 34.40205426 | 25.25956258 | 0.016602691 | 1.887502828 | 0.008829493 | 2.618586397 | 3.699942442 |
| CNAG_01304 | - | polynucleotide 5'-hydroxyl-kinase GRC3 | 1091.677014 | 1049.373541 | 1026.970486 | 883.9060229 | 984.173165 | 858.2962752 | 0.016605854 | -0.231957741 | 0.008833203 | -2.618443057 | 0.851478649 |
| CNAG_00635 | - | cytoplasmic protein | 2383.474206 | 2310.079543 | 2338.780206 | 2614.357276 | 2713.721874 | 2653.227821 | 0.016637482 | 0.167246171 | 0.008852059 | 2.617715441 | 1.122913012 |
| CNAG_06360 | - | Alcohol dehydrogenase | 825.6359641 | 929.6688433 | 962.9130235 | 739.1110923 | 737.3229759 | 825.9824804 | 0.016680306 | -0.253444237 | 0.00887688 | -2.616759717 | 0.838891285 |
| CNAG_11008 | - | unspecified product | 23.6213661 | 18.50309738 | 20.40383722 | 40.84667896 | 46.98790956 | 40.89447333 | 0.016730105 | 1.030410649 | 0.008905425 | 2.615663584 | 2.042605577 |
| CNAG_06159 | - | hypothetical protein, variant | 13.34792672 | 27.00427936 | 18.28609677 | 42.52873045 | 33.69467974 | 51.7179517 | 0.016749629 | 1.107006921 | 0.008917863 | 2.615186938 | 2.153983085 |
| CNAG_00383 | APT3 | Aminophospholipid translocase, putative | 3007.944262 | 2924.090662 | 2616.033748 | 2514.495565 | 2500.458228 | 2580.067797 | 0.016828635 | -0.185739054 | 0.008961983 | -2.61350098 | 0.879198571 |
| CNAG_01660 | GCN3 | translation initiation factor eif-2b subunit alpha | 644.5005675 | 643.1832548 | 674.9925392 | 572.6968328 | 529.442993 | 552.9559882 | 0.016891126 | -0.261726748 | 0.008997325 | -2.612155783 | 0.834089008 |
| CNAG_05237 | - | hypothetical protein | 420.5268134 | 367.5486384 | 421.3019716 | 546.7771433 | 523.7633258 | 454.0691254 | 0.016982713 | 0.318345596 | 0.009048184 | 2.61022822 | 1.246899852 |
| CNAG_07916 | - | hypothetical protein | 0.087003557 | 0.042411767 | 0.036649734 | 4.026313072 | 7.217606137 | 3.306550816 | 0.01709196 | 4.648603149 | 0.009108476 | 2.607955621 | 25.08239402 |
| CNAG_04440 | - | thiamine pyrophosphokinase, variant | 382.9415759 | 423.3089944 | 391.3412981 | 321.9694056 | 321.5836014 | 327.6020821 | 0.017118857 | -0.318448005 | 0.0091249 | -2.607338875 | 0.801932101 |
| CNAG_07886 | - | hypothetical protein | 75.1537513 | 108.8253803 | 69.51013695 | 143.8269708 | 148.8027203 | 103.2982477 | 0.017132832 | 0.6308354 | 0.009134442 | 2.606981032 | 1.548461379 |
| CNAG_00161 | - | auxin-induced protein | 459.3145623 | 319.9834383 | 290.1224711 | 254.896741 | 257.6796919 | 286.5112959 | 0.017233349 | -0.438715297 | 0.009190138 | -2.604898889 | 0.73779131 |
| CNAG_04223 | - | hypothetical protein | 0.153126723 | 2.232386182 | 0.064151137 | 10.67796433 | 6.087473946 | 9.776451206 | 0.017338625 | 3.634065139 | 0.009248396 | 2.602732961 | 12.41545423 |
| CNAG_06100 | - | hypothetical protein | 312.1980611 | 416.9823755 | 381.8555257 | 453.1013879 | 529.0811363 | 440.6977322 | 0.017370658 | 0.344195197 | 0.009267603 | 2.602021534 | 1.26944263 |
| CNAG_00274 | - | hypothetical protein | 210.3222412 | 202.6390441 | 167.5900303 | 261.8036632 | 284.0979984 | 236.432887 | 0.017382782 | 0.416353777 | 0.009276195 | 2.601703747 | 1.334550387 |
| CNAG_05730 | - | acyl-protein thioesterase 1 | 1143.232721 | 1235.753891 | 1261.54668 | 1084.041539 | 1070.349955 | 1020.354771 | 0.01743004 | -0.212288795 | 0.009303542 | -2.600693924 | 0.863166754 |
| CNAG_03917 | NUP75 | nuclear pore complex protein Nup85 | 2289.846946 | 2197.942633 | 2243.786815 | 1996.163729 | 2069.236397 | 1959.527354 | 0.01743195 | -0.175168134 | 0.00930669 | -2.600577837 | 0.885664297 |
| CNAG_03995 | - | hypothetical protein | 43.66016398 | 51.86686643 | 59.79661224 | 109.228758 | 68.98534481 | 82.00494051 | 0.017538689 | 0.744847618 | 0.009365818 | 2.598404166 | 1.675797257 |
| CNAG_04438 | - | hypothetical protein | 273.1691542 | 282.222225 | 310.2121813 | 234.067107 | 218.8959064 | 223.2812681 | 0.017667754 | -0.373005989 | 0.009436898 | -2.595807284 | 0.772171928 |
| CNAG_03143 | - | hypothetical protein | 3351.826113 | 4434.702441 | 3138.750674 | 5571.629596 | 5132.480566 | 4440.629277 | 0.0176982 | 0.457612286 | 0.009455321 | 2.59513703 | 1.373267128 |
| CNAG_04739 | - | cytochrome c oxidase assembly protein Cox19 | 248.9059514 | 196.2982213 | 259.1954176 | 392.3580581 | 275.1312482 | 296.3550195 | 0.017802389 | 0.437937898 | 0.009513158 | 2.593040448 | 1.354666664 |
| CNAG_12644 | - | unspecified product | 5.036829771 | 4.487970669 | 10.73111052 | 22.36802041 | 24.60915893 | 15.50601058 | 0.017858201 | 1.665961644 | 0.009545164 | 2.591885132 | 3.173251007 |
| CNAG_12292 | - | unspecified product | 19.1328976 | 22.79559753 | 29.9318155 | 45.53758896 | 41.54132036 | 54.0241795 | 0.017864784 | 0.985185563 | 0.009550864 | 2.591679725 | 1.979567913 |
| CNAG_01863 | SNF2 | ATP-dependent helicase STH1/SNF2 | 3992.341955 | 3845.295661 | 3409.48103 | 3202.440183 | 3362.183808 | 3427.110824 | 0.01791001 | -0.185958947 | 0.009581604 | -2.590573926 | 0.879064575 |
| CNAG_02056 | - | hypothetical protein | 102.7678014 | 155.8721526 | 110.8689475 | 83.7121076 | 87.81253081 | 72.41965847 | 0.01791001 | -0.614358685 | 0.009579993 | -2.590631794 | 0.653220202 |
| CNAG_02822 | - | hypothetical protein | 727.8203005 | 871.6702992 | 853.1994229 | 983.953866 | 980.8479361 | 962.4393657 | 0.01791001 | 0.24043387 | 0.009577239 | 2.590730751 | 1.181347881 |
| CNAG_01512 | - | hypothetical protein, variant | 71.46230626 | 70.14832819 | 55.63462665 | 122.2122251 | 111.6123138 | 81.0974709 | 0.01793904 | 0.664586698 | 0.009599325 | 2.58993788 | 1.585114106 |
| CNAG_01673 | - | hypothetical protein, variant | 1392.170052 | 1505.217175 | 1508.001773 | 1611.858841 | 1723.506102 | 1763.929987 | 0.017985656 | 0.195537762 | 0.009626466 | 2.588965781 | 1.145150934 |
| CNAG_04003 | - | pumilio 2 | 502.4238513 | 571.0932163 | 575.9262354 | 714.8278128 | 670.3306288 | 627.2209177 | 0.01804451 | 0.272317745 | 0.00966017 | 2.587762004 | 1.207746561 |
| CNAG_13124 | - | unspecified product | 10.90462457 | 12.02172493 | 16.09106323 | 27.52122687 | 33.15266912 | 31.93718115 | 0.018184213 | 1.24464213 | 0.009737181 | 2.585025455 | 2.369597675 |
| CNAG_03002 | - | hypothetical protein | 157.7102833 | 45.39870153 | 45.97912923 | 48.68699451 | 43.7012506 | 25.94586397 | 0.018212115 | -1.111315995 | 0.009754346 | -2.584418139 | 0.462871617 |
| CNAG_13207 | - | unspecified product | 1.526110564 | 6.582427519 | 7.550797971 | 15.85194836 | 19.32074829 | 18.6334473 | 0.018236411 | 1.879985223 | 0.009769586 | 2.583879731 | 3.680712902 |
| CNAG_00014 | - | hypothetical protein | 486.0861644 | 575.3809594 | 548.2131127 | 621.929381 | 656.5344447 | 687.0329383 | 0.018260015 | 0.27302737 | 0.00978446 | 2.583354938 | 1.208340767 |
| CNAG_10090 | - | unspecified product | 6.360764304 | 16.29353904 | 18.21822651 | 39.76496949 | 39.32155355 | 22.25679051 | 0.01826099 | 1.308018174 | 0.009787213 | 2.583257915 | 2.476011773 |
| CNAG_12118 | - | unspecified product | 86.202492 | 99.98210396 | 86.29787863 | 57.19732543 | 45.32019632 | 68.89512353 | 0.018266271 | -0.686923672 | 0.009792274 | -2.583079558 | 0.621177004 |
| CNAG_04018 | - | hypothetical protein | 262.0558711 | 178.8713703 | 192.9965867 | 162.4353667 | 150.7320846 | 144.5242388 | 0.018421448 | -0.487585498 | 0.009877711 | -2.580081112 | 0.713217745 |
| CNAG_05712 | RIB4 | 6,7-dimethyl-8-ribityllumazine synthase | 190.711017 | 193.0243617 | 254.8786042 | 251.371786 | 323.8329048 | 291.8128781 | 0.018445674 | 0.425689353 | 0.009892954 | 2.579548595 | 1.343214168 |
| CNAG_12793 | - | unspecified product | 1.310753354 | 0.058166456 | 0.050248287 | 8.253229814 | 4.337013421 | 7.588904414 | 0.018489999 | 4.129429061 | 0.009918985 | 2.578640873 | 17.50177161 |
| CNAG_05620 | - | hypothetical protein | 468.2656703 | 425.6514982 | 315.8644671 | 569.4140375 | 511.6263895 | 488.4227359 | 0.018526624 | 0.360998419 | 0.009940894 | 2.577878504 | 1.284314402 |
| CNAG_12481 | - | unspecified product | 28.9181423 | 32.12969661 | 30.78452538 | 14.39393894 | 14.42322153 | 11.35038112 | 0.018552238 | -1.221372368 | 0.009956904 | -2.577322379 | 0.428874556 |
| CNAG_05176 | HOB3 | hypothetical protein | 456.3159593 | 395.4092074 | 442.4925226 | 375.8325816 | 363.0453308 | 292.225448 | 0.018575579 | -0.344144008 | 0.009971699 | -2.57680915 | 0.787775247 |
| CNAG_07767 | - | hypothetical protein | 59.99108083 | 37.69898352 | 53.12111451 | 25.24705799 | 8.949264152 | 35.55074861 | 0.01857623 | -1.147486813 | 0.009974317 | -2.576718395 | 0.451410908 |
| CNAG_01014 | ZFC4 | hypothetical protein | 470.6310792 | 425.6669108 | 368.0681367 | 544.3262071 | 520.2416668 | 517.248085 | 0.018630237 | 0.308554402 | 0.010005591 | 2.575636035 | 1.23846612 |
| CNAG_05885 | - | hypothetical protein | 2.6048389 | 3.349339101 | 3.300726543 | 8.565456464 | 16.84676024 | 14.1979779 | 0.018668467 | 2.233046339 | 0.010028402 | 2.574848425 | 4.701256313 |
| CNAG_07805 | - | insulysin | 1322.895797 | 1265.942948 | 1228.540523 | 1153.335301 | 1090.151059 | 1099.152521 | 0.018719013 | -0.206305969 | 0.010057841 | -2.573834377 | 0.866753721 |
| CNAG_03613 | - | hypothetical protein | 216.3010035 | 219.9102894 | 238.9405902 | 325.0603534 | 292.5580783 | 267.5293995 | 0.018880233 | 0.378111566 | 0.010146771 | 2.570787014 | 1.299639561 |
| CNAG_12647 | - | unspecified product | 150.7156724 | 209.8668788 | 193.9588941 | 112.3013582 | 119.6057863 | 158.7535001 | 0.0188814 | -0.518646952 | 0.010149704 | -2.570686917 | 0.698026178 |
| CNAG_05413 | - | phosphatidylinositol glycan, class Q | 461.4071222 | 483.8018761 | 388.3333122 | 512.1347863 | 531.4009278 | 641.3807885 | 0.018928511 | 0.323689215 | 0.01017734 | 2.569745009 | 1.251526826 |
| CNAG_01527 | - | hypothetical protein | 4.893580577 | 3.362465944 | 2.255579876 | 12.72866791 | 21.35192673 | 8.900171714 | 0.018929921 | 2.170653138 | 0.010180409 | 2.569640528 | 4.502271751 |
| CNAG_07758 | - | hypothetical protein | 585.2800707 | 647.5799154 | 623.9339282 | 727.8318158 | 720.3042582 | 781.4021954 | 0.018950063 | 0.24955783 | 0.010193556 | 2.569193375 | 1.188842692 |
| CNAG_06073 | - | hypothetical protein, variant | 61.25296132 | 87.37546549 | 96.0253578 | 134.4315169 | 119.2021203 | 120.8584684 | 0.018961875 | 0.597958006 | 0.010202225 | 2.56889878 | 1.513572735 |
| CNAG_12600 | - | unspecified product | 182.2450238 | 145.6153415 | 150.4950089 | 219.780677 | 247.069714 | 194.2655666 | 0.018975651 | 0.45218531 | 0.010214272 | 2.5684898 | 1.368111023 |
| CNAG_12732 | - | unspecified product | 27.30011435 | 42.07363305 | 26.84149484 | 72.54101705 | 47.45510908 | 59.72695783 | 0.018975651 | 0.892836052 | 0.010213414 | 2.568518922 | 1.856822683 |
| CNAG_06728 | - | kinesin | 2580.665921 | 2678.435335 | 2523.217885 | 2360.445965 | 2234.760829 | 2395.76592 | 0.019117221 | -0.169796775 | 0.010292811 | -2.565833879 | 0.888967897 |
| CNAG_04590 | - | hypothetical protein | 144.1173599 | 182.1981984 | 216.4927272 | 239.9294527 | 232.1156787 | 277.2501883 | 0.019145452 | 0.450201865 | 0.010310348 | 2.565243289 | 1.36623141 |
| CNAG_05674 | - | solute carrier family 35, member E1 | 162.735689 | 198.3386533 | 211.2025996 | 306.6133115 | 250.2689519 | 225.3746814 | 0.019228441 | 0.437101881 | 0.010357389 | 2.563663567 | 1.353881885 |
| CNAG_12800 | - | unspecified product | 53.12221057 | 44.03971744 | 41.5084113 | 22.07981088 | 15.08927995 | 32.24591926 | 0.019538847 | -1.015057943 | 0.010526974 | -2.558020972 | 0.494808455 |
| CNAG_12784 | - | unspecified product | 179.2262612 | 133.5743204 | 137.5309237 | 101.5654032 | 94.58110958 | 117.7551814 | 0.01984752 | -0.541213234 | 0.010695702 | -2.552486601 | 0.687192771 |
| CNAG_02491 | - | nuclear protein | 751.9675034 | 770.3146516 | 797.6393832 | 631.7018386 | 665.7910537 | 687.2433418 | 0.01985223 | -0.240372219 | 0.010700665 | -2.552325015 | 0.846526878 |
| CNAG_01565 | - | biotin-[acetyl-CoA-carboxylase] ligase | 1905.702413 | 1988.872585 | 1916.385405 | 1657.552998 | 1889.915981 | 1532.195646 | 0.019871535 | -0.209381238 | 0.010713497 | -2.551907462 | 0.864908105 |
| CNAG_05396 | - | LRP16 family protein | 638.6468861 | 616.2675826 | 676.0446453 | 502.4411641 | 559.7182451 | 561.8049455 | 0.019899156 | -0.266318129 | 0.010730818 | -2.551344543 | 0.831438737 |
| CNAG_01908 | HEM4 | uroporphyrinogen-III synthase | 764.8042486 | 796.1554832 | 794.451748 | 732.3714114 | 646.7665955 | 621.8484573 | 0.020018161 | -0.250927635 | 0.010797438 | -2.549187028 | 0.840355903 |
| CNAG_12870 | - | unspecified product | 48.39083533 | 41.82899211 | 30.916441 | 23.85356725 | 14.27359246 | 20.18522025 | 0.020037771 | -1.077031196 | 0.010810462 | -2.548766621 | 0.474003233 |
| CNAG_05587 | - | hypothetical protein | 101.9530093 | 103.5922679 | 130.1374538 | 155.9302484 | 157.1657135 | 166.3137585 | 0.020144377 | 0.501806985 | 0.010870436 | 2.546836468 | 1.415985984 |
| CNAG_05392 | ZAP104 | specific RNA polymerase II transcription factor | 91.55208745 | 118.6154898 | 128.0207007 | 145.631846 | 163.2394275 | 179.5540963 | 0.020148026 | 0.516649172 | 0.010874866 | 2.546694289 | 1.430628584 |
| CNAG_01108 | DAL2 | Allantoicase | 433.2995487 | 382.5987287 | 380.8208312 | 469.3963483 | 503.0285112 | 513.8539053 | 0.020294199 | 0.297814858 | 0.010956241 | 2.544091414 | 1.229281102 |
| CNAG_03177 | - | hypothetical protein, variant | 54.00031565 | 56.14122374 | 44.96240984 | 108.4411261 | 82.13746589 | 67.74676151 | 0.020334865 | 0.723867063 | 0.010980678 | 2.543313114 | 1.651603136 |
| CNAG_03405 | - | hypothetical protein | 712.5627178 | 798.409812 | 759.3724496 | 863.2003664 | 936.9111663 | 891.3842364 | 0.020354996 | 0.230659879 | 0.01099652 | 2.542809381 | 1.173371519 |
| CNAG_06732 | - | hypothetical protein | 1538.001412 | 1485.836923 | 1425.921389 | 1714.721854 | 1674.287978 | 1726.218751 | 0.020354996 | 0.18602171 | 0.010994345 | 2.542878522 | 1.137622347 |
| CNAG_00190 | - | hypothetical protein | 455.5810885 | 449.3710396 | 430.9396189 | 577.0238415 | 620.494684 | 472.996605 | 0.020446599 | 0.307655002 | 0.011048504 | 2.541160971 | 1.23769428 |
| CNAG_00294 | - | splicing factor 45 | 1735.22662 | 1716.33406 | 1866.219715 | 1457.450027 | 1614.659898 | 1617.546343 | 0.020598402 | -0.197063855 | 0.011133048 | -2.538494735 | 0.872324095 |
| CNAG_02506 | RIB3 | 3,4-dihydroxy-2-butanone-4-phosphate synthase | 693.457653 | 618.4521264 | 706.9698104 | 563.2769613 | 606.3440139 | 519.7284188 | 0.020640965 | -0.272871323 | 0.011160132 | -2.537644377 | 0.827670635 |
| CNAG_06219 | - | amyloid beta protein binding protein 1 | 1421.126207 | 1621.409612 | 1388.526536 | 1291.797395 | 1323.960007 | 1267.903651 | 0.020654876 | -0.205501772 | 0.011168615 | -2.537378425 | 0.867237008 |
| CNAG_13014 | - | unspecified product | 4.975022499 | 8.69385622 | 2.279725102 | 9.145599524 | 16.60526925 | 28.22991866 | 0.020753024 | 1.909042608 | 0.011224221 | 2.535639515 | 3.755597903 |
| CNAG_01912 | - | NAD synthetase | 8024.581528 | 6938.687161 | 7419.290361 | 6553.800568 | 6834.011477 | 6746.509766 | 0.020783039 | -0.16867788 | 0.011242992 | -2.535054224 | 0.889657611 |
| CNAG_03493 | - | 1-alkyl-2-acetylglycerophosphocholine esterase | 233.8469143 | 235.0146145 | 280.4988055 | 360.0896315 | 321.0109458 | 296.3922232 | 0.02085638 | 0.369881114 | 0.011285214 | 2.533740899 | 1.292246338 |
| CNAG_06643 | - | ribonuclease III | 157.7694306 | 136.970428 | 136.6243256 | 200.3609818 | 205.5394735 | 184.2155754 | 0.021040619 | 0.441509193 | 0.011387474 | 2.53057807 | 1.358024204 |
| CNAG_07315 | LIV6 | T-cell immuno-modulatory protein homolog | 1926.703248 | 1984.565078 | 1847.0915 | 1648.981014 | 1787.193696 | 1700.84385 | 0.021131379 | -0.180091699 | 0.011441755 | -2.528909413 | 0.882646893 |
| CNAG_06752 | - | hypothetical protein, variant | 313.5033532 | 474.0991198 | 480.9863605 | 521.579302 | 507.2238943 | 635.8013627 | 0.021131379 | 0.378789528 | 0.011439207 | 2.528987587 | 1.300250442 |
| CNAG_05919 | - | hypothetical protein | 1276.534208 | 1372.682469 | 1310.72781 | 1435.166219 | 1602.800153 | 1544.150508 | 0.02116091 | 0.195873232 | 0.011460329 | 2.528340055 | 1.145417247 |

| | | | | | | | | | | | | |
|---|---|---|---|---|---|---|---|---|---|---|---|---|
| CNAG_01698 | - | hypothetical protein | 488.3388901 | 447.2862641 | 569.4823545 | 590.0356759 | 638.4694668 | 633.7717977 | 0.021167911 | 0.291450048 | 0.011466706 | 2.528144771 | 1.223869767 |
| CNAG_03367 | URK1 | Uridine kinase | 1941.745779 | 1817.58161 | 1679.677368 | 1476.117529 | 1683.69482 | 1600.923875 | 0.021187049 | -0.207675608 | 0.01147966 | -2.52774835 | 0.86593125 |
| CNAG_03582 | RIM20 | pH-response regulator protein palA/RIM20 | 1714.20626 | 1719.554742 | 1790.522931 | 1539.298695 | 1607.770344 | 1509.916977 | 0.021221429 | -0.18146759 | 0.011503471 | -2.527020744 | 0.881805518 |
| CNAG_06914 | TUB4 | tubulin gamma chain | 1275.120442 | 1339.177662 | 1296.769781 | 1163.430326 | 1015.006053 | 1213.418463 | 0.021221429 | -0.220228498 | 0.011503449 | -2.527021416 | 0.858429465 |
| CNAG_00035 | - | hypothetical protein | 22.65404086 | 34.56890143 | 29.9832251 | 48.49314291 | 61.71088647 | 50.90716676 | 0.021260136 | 0.889267042 | 0.011529645 | 2.526222447 | 1.85223486 |
| CNAG_00476 | - | hypothetical protein | 22.6103078 | 18.55262687 | 37.34686889 | 48.62844407 | 52.0716188 | 51.93089502 | 0.021260136 | 0.955231213 | 0.011527353 | 2.526292295 | 1.938890345 |
| CNAG_00855 | - | ligase | 739.1287852 | 777.8373331 | 773.123993 | 609.8710483 | 694.3249043 | 652.8654872 | 0.021276255 | -0.242727753 | 0.011540985 | -2.525877093 | 0.845145854 |
| CNAG_05153 | GAT5 | hypothetical protein | 414.6905569 | 443.9190744 | 340.3514478 | 482.6755282 | 504.7484223 | 514.9698997 | 0.021288044 | 0.31083112 | 0.011549979 | 2.525603383 | 1.240422087 |
| CNAG_03379 | - | N-acetyltransferase 5 | 649.192925 | 638.8944174 | 711.2269786 | 568.7534898 | 538.0648085 | 582.8982753 | 0.021392715 | -0.258556234 | 0.011609382 | -2.523800424 | 0.835924046 |
| CNAG_01085 | - | hypothetical protein | 281.451691 | 333.8781305 | 336.8842657 | 252.7075997 | 224.8634867 | 269.8261 | 0.021512 | -0.365163805 | 0.011681996 | -2.521607546 | 0.776380719 |
| CNAG_02402 | - | hypothetical protein | 1375.731464 | 1363.013563 | 1355.518296 | 1474.067866 | 1617.427164 | 1639.590746 | 0.021512 | 0.19339868 | 0.011680132 | 2.521663672 | 1.143454279 |
| CNAG_05049 | PIP201 | hypothetical protein | 818.7655955 | 703.732049 | 929.9370293 | 932.4322304 | 1065.490521 | 995.7181216 | 0.021512 | 0.27210337 | 0.011677393 | 2.521746206 | 1.207567111 |
| CNAG_12218 | - | unspecified product | 28.26888707 | 17.47052207 | 27.82943167 | 39.80797482 | 53.92121007 | 50.78712499 | 0.021566468 | 0.962813197 | 0.011714208 | 2.52063865 | 1.949106871 |
| CNAG_07797 | CLR6 | transcriptional regulator, variant | 1138.812037 | 1604.148721 | 1644.323184 | 1198.323351 | 1255.794018 | 1195.766343 | 0.021580471 | -0.279921957 | 0.011724449 | -2.520331092 | 0.823635571 |
| CNAG_03175 | - | hypothetical protein | 750.7387986 | 800.4183665 | 690.0026208 | 644.2597346 | 639.9101564 | 626.2393777 | 0.021592713 | -0.245546456 | 0.011733737 | -2.520052379 | 0.843496241 |
| CNAG_02963 | - | anaphase-promoting complex subunit 8 | 831.4093503 | 853.2919294 | 927.7235728 | 774.2870971 | 715.7625932 | 754.989931 | 0.021646629 | -0.234337209 | 0.011768323 | -2.519016251 | 0.850075444 |
| CNAG_02757 | - | hypothetical protein | 203.5700539 | 241.428053 | 260.2444523 | 304.5534609 | 295.966359 | 319.5479199 | 0.021646629 | 0.368201888 | 0.011766303 | 2.519076666 | 1.290743102 |
| CNAG_02085 | HST302 | histone deacetylase | 155.7858374 | 210.1654823 | 221.8488613 | 268.8469459 | 264.9924556 | 256.3382195 | 0.02165816 | 0.412862724 | 0.011777236 | 2.518749651 | 1.331324928 |
| CNAG_06966 | AKP1 | alkaline phosphatase D | 12.14582574 | 18.45834128 | 19.30813842 | 34.57333205 | 55.22924583 | 23.41368747 | 0.021753299 | 1.177203908 | 0.011831627 | 2.517126709 | 2.261380731 |
| CNAG_01924 | - | hypothetical protein | 667.5306205 | 569.9099886 | 522.575351 | 510.4659479 | 512.3085191 | 391.0470815 | 0.02176775 | -0.331230848 | 0.011842145 | -2.516813635 | 0.794858054 |
| CNAG_02869 | - | hypothetical protein | 42.14587936 | 28.20811925 | 26.83966017 | 54.77246249 | 60.69201085 | 61.90128047 | 0.021915293 | 0.849861907 | 0.011925088 | 2.51435337 | 1.8023284 |
| CNAG_05451 | - | hypothetical protein | 175.7428077 | 272.6428645 | 293.2763749 | 354.567222 | 330.5250703 | 320.7411651 | 0.021941106 | 0.426166184 | 0.011941813 | 2.513859099 | 1.343658193 |
| CNAG_05131 | - | hypothetical protein | 652.598786 | 633.4746665 | 614.2508689 | 529.0165772 | 530.3411487 | 552.9275362 | 0.022126982 | -0.252543703 | 0.012045681 | -2.510803242 | 0.839415086 |
| CNAG_05122 | - | homoserine O-acetyltransferase | 3071.310935 | 3583.47718 | 3849.787188 | 3163.64804 | 3160.359202 | 2907.644823 | 0.022177588 | -0.201275851 | 0.012075939 | -2.509917445 | 0.86978103 |
| CNAG_05823 | - | hypothetical protein | 212.3120033 | 226.1046883 | 194.0684138 | 142.7545313 | 183.7097249 | 147.8486673 | 0.022247116 | -0.436304766 | 0.012116514 | -2.508732669 | 0.739025082 |
| CNAG_06024 | - | peroxin-14 | 378.269673 | 403.9451608 | 399.8485178 | 320.4274644 | 295.6315909 | 343.0884579 | 0.022329712 | -0.315526873 | 0.012164226 | -2.507344019 | 0.803557477 |
| CNAG_05968 | CDC420 | cell division control protein 42 | 114.2094037 | 125.8713913 | 107.6585175 | 81.40835304 | 70.41985552 | 85.5985682 | 0.022361325 | -0.566529631 | 0.012184178 | -2.50676473 | 0.675239108 |
| CNAG_04077 | - | hypothetical protein | 18.79860847 | 9.886967034 | 8.707024487 | 39.18440411 | 26.91987604 | 23.27334934 | 0.022456094 | 1.313035397 | 0.012238558 | 2.50519015 | 2.484637525 |
| CNAG_01606 | - | hypothetical protein | 0.087372237 | 0.042590335 | 0.036803911 | 4.904814548 | 3.494811479 | 6.464580002 | 0.02246058 | 4.543850961 | 0.012243745 | 2.505040273 | 23.32574025 |
| CNAG_03815 | - | hypothetical protein | 596.9942414 | 682.0550052 | 668.7126181 | 839.3643073 | 752.212621 | 747.0986399 | 0.022477905 | 0.250027174 | 0.012255934 | 2.504688315 | 1.189229514 |
| CNAG_05994 | - | multidrug transporter | 1863.643094 | 1276.629924 | 978.008261 | 946.8843333 | 808.7421155 | 660.9426941 | 0.02248925 | -0.787091013 | 0.012264866 | -2.504430606 | 0.579511416 |
| CNAG_00420 | - | dolichyl-phosphate mannosyltransferase polypeptide 3 | 431.60759 | 340.4625982 | 347.6450997 | 298.6240383 | 285.3104425 | 305.3926013 | 0.022541951 | -0.348291163 | 0.012296361 | -2.503523235 | 0.785513969 |
| CNAG_06640 | - | cathepsin A (carboxypeptidase C) | 3380.469399 | 3119.154141 | 3133.226026 | 2711.310075 | 2900.71814 | 3015.216601 | 0.022560383 | -0.174687865 | 0.01230917 | -2.503154793 | 0.885959181 |
| CNAG_01619 | - | hypothetical protein | 528.1105968 | 629.2389471 | 581.2800689 | 720.1759652 | 673.7204425 | 703.7427976 | 0.022607357 | 0.255710528 | 0.012340321 | 2.502260152 | 1.193923606 |
| CNAG_01303 | - | hypothetical protein, variant | 923.9269617 | 991.3006807 | 1034.488073 | 1056.868554 | 1218.975175 | 1207.72573 | 0.022607357 | 0.224651649 | 0.012338832 | 2.502302872 | 1.168495072 |
| CNAG_03179 | - | hypothetical protein | 6.131860124 | 6.594520932 | 5.459558347 | 24.22875496 | 11.14160696 | 20.80821812 | 0.022608447 | 1.679618646 | 0.012343676 | 2.502163906 | 3.203432622 |
| CNAG_05967 | - | hypothetical protein | 70.17103947 | 56.18056247 | 44.98874194 | 115.3863047 | 87.28059376 | 77.69583332 | 0.022731652 | 0.698964132 | 0.012413719 | 2.500160095 | 1.623338802 |
| CNAG_12890 | - | unspecified product | 6.088904872 | 2.320366479 | 6.493242769 | 23.55895361 | 13.00875395 | 14.33032092 | 0.022782037 | 1.818994189 | 0.012444017 | 2.499296444 | 3.528351249 |
| CNAG_07716 | - | hypothetical protein | 48.42538883 | 85.13360803 | 67.30591525 | 107.8484903 | 98.47843681 | 110.766694 | 0.022788499 | 0.642197252 | 0.012450329 | 2.499116738 | 1.560704332 |
| CNAG_04054 | - | translation initiation factor SUI1 | 1441.159171 | 1381.366775 | 1558.101365 | 1764.631468 | 1643.225323 | 1667.407738 | 0.022841883 | 0.196703047 | 0.012482284 | 2.498208249 | 1.146076262 |
| CNAG_02936 | - | CCR4-NOT transcription complex subunit 1 | 3564.740773 | 2935.990553 | 2738.686392 | 2801.778752 | 2734.245824 | 2410.336382 | 0.023130873 | -0.233514694 | 0.012643032 | -2.493669182 | 0.850560231 |
| CNAG_07718 | CIN1 | Intersectin 1 | 1882.421358 | 1734.727905 | 1549.671437 | 2069.6899 | 2140.200872 | 1855.098123 | 0.023149661 | 0.215870976 | 0.012656128 | 2.49330164 | 1.16140486 |
| CNAG_07365 | - | oxidoreductase | 513.451855 | 425.5799314 | 442.547217 | 376.4653416 | 374.2040081 | 379.7795722 | 0.023189998 | -0.305529695 | 0.012681012 | -2.492604182 | 0.809145079 |
| CNAG_02228 | - | hypothetical protein, variant | 296.3558122 | 328.4035833 | 235.7342921 | 229.3845604 | 227.5767143 | 204.4681016 | 0.02322609 | -0.39479291 | 0.012703585 | -2.491972568 | 0.760598548 |
| CNAG_05252 | - | chaperone regulator | 1108.157932 | 1189.408898 | 1142.135281 | 1067.003947 | 996.1506796 | 939.3370761 | 0.023227276 | -0.211048993 | 0.01270707 | -2.491875128 | 0.863908849 |
| CNAG_12838 | - | unspecified product | 7.304403188 | 5.552632384 | 8.632204244 | 15.65606266 | 27.32825886 | 18.72774316 | 0.023328277 | 1.567987834 | 0.012765174 | 2.49025426 | 2.964909016 |
| CNAG_00491 | - | hypothetical protein | 435.1631369 | 341.5616138 | 376.4076478 | 318.1367442 | 294.7981809 | 309.8499597 | 0.023410426 | -0.338959239 | 0.012812985 | -2.488925408 | 0.790611454 |
| CNAG_00923 | - | membrane protein | 288.7326977 | 359.858739 | 290.1911064 | 412.8259861 | 378.7408853 | 398.4274597 | 0.023450383 | 0.328224549 | 0.012837717 | 2.488239701 | 1.255467384 |
| CNAG_02538 | - | hypothetical protein | 4.95643733 | 8.681461004 | 1.207324037 | 16.77289382 | 12.11226944 | 21.79896354 | 0.02347106 | 1.903149616 | 0.012851903 | 2.487846949 | 3.740288661 |
| CNAG_05201 | - | DNA mismatch repair protein Msh4 | 196.4233324 | 211.2449044 | 191.0041102 | 284.5384721 | 230.284426 | 276.2401325 | 0.023516323 | 0.387069884 | 0.012879559 | 2.487082333 | 1.307734694 |
| CNAG_03307 | - | hypothetical protein | 13.18866827 | 16.2762182 | 11.88403989 | 34.91695869 | 34.0196689 | 24.39025692 | 0.023567105 | 1.186948806 | 0.01291025 | 2.486235511 | 2.276707268 |
| CNAG_07982 | - | hypothetical protein | 628.2032103 | 641.0343019 | 718.6717439 | 607.0541199 | 569.200553 | 467.6177732 | 0.023586101 | -0.289756585 | 0.012923536 | -2.485869464 | 0.818040069 |
| CNAG_01691 | - | hypothetical protein, variant | 1504.791351 | 1167.955975 | 1183.771117 | 1106.56303 | 1084.111802 | 1081.401282 | 0.023589125 | -0.252876622 | 0.012928073 | -2.485744536 | 0.839221404 |
| CNAG_00931 | - | protein transporter | 372.4154628 | 404.99888 | 379.6054749 | 350.2251794 | 282.6735424 | 298.7853492 | 0.023705336 | -0.32735071 | 0.012994658 | -2.483915667 | 0.796998708 |
| CNAG_02770 | - | transcription initiation factor TFIID subunit 11 | 651.656077 | 715.3663067 | 803.9749717 | 636.5266174 | 589.8301559 | 602.9266654 | 0.023705885 | -0.262574567 | 0.012997854 | -2.483828094 | 0.833598989 |
| CNAG_12067 | - | unspecified product | 75.71387369 | 82.80645097 | 81.98983659 | 54.17709225 | 50.6516058 | 47.95683566 | 0.023747455 | -0.671386513 | 0.013023546 | -2.483124763 | 0.627902946 |
| CNAG_02871 | - | GPI mannosyltransferase 2 | 148.7778595 | 200.4693536 | 209.0591953 | 264.2481746 | 219.0761534 | 275.0661165 | 0.02386342 | 0.430452053 | 0.013090058 | 2.481309716 | 1.347655784 |
| CNAG_03548 | - | hypothetical protein | 17.56180748 | 5.595359177 | 6.569539926 | 27.82383052 | 29.76455912 | 21.02403829 | 0.023878857 | 1.440400341 | 0.013101442 | 2.480999885 | 2.713961662 |
| CNAG_02170 | - | hypothetical protein | 61.50254811 | 162.6882673 | 148.2857596 | 164.2505662 | 229.1235903 | 194.075465 | 0.023947852 | 0.649387821 | 0.013150994 | 2.479653966 | 1.568502492 |
| CNAG_02114 | - | hypothetical protein | 1672.301247 | 1646.364898 | 1551.765438 | 1772.964838 | 1931.4119 | 1880.53509 | 0.023947852 | 0.182119137 | 0.013149331 | 2.479699047 | 1.134549172 |
| CNAG_00267 | - | hypothetical protein | 122.8917417 | 128.3202007 | 129.128691 | 169.8987928 | 187.4675669 | 167.5362003 | 0.023947852 | 0.449362257 | 0.013148261 | 2.479728086 | 1.365436533 |
| CNAG_11011 | - | unspecified product | 49.42222803 | 52.94271526 | 57.68461847 | 77.99640135 | 68.093218 | 117.0793186 | 0.023947852 | 0.715150919 | 0.013146361 | 2.47977961 | 1.641654941 |
| CNAG_02036 | CAS4 | putative sugar transporter | 456.6304873 | 529.9958436 | 535.2654589 | 413.0251372 | 424.2577158 | 425.2872464 | 0.023968252 | -0.283136825 | 0.013165123 | -2.479271009 | 0.821802244 |
| CNAG_05973 | - | serine carboxypeptidase | 54.29319607 | 90.56744036 | 90.70200695 | 113.0918963 | 129.7358332 | 117.5174279 | 0.024019885 | 0.60348449 | 0.013196418 | 2.478424125 | 1.519381848 |
| CNAG_01149 | - | hypothetical protein | 2594.818037 | 2634.344718 | 2287.643725 | 2705.330307 | 2956.984383 | 2965.136493 | 0.024031458 | 0.183875043 | 0.01320571 | 2.478172981 | 1.135930873 |
| CNAG_03268 | - | hypothetical protein | 2684.646946 | 2466.298561 | 2110.643137 | 3100.306009 | 2854.315811 | 2612.228813 | 0.024120024 | 0.223164027 | 0.013257324 | 2.476780947 | 1.16729081 |
| CNAG_02748 | - | UTP-glucose-1-phosphate uridylyltransferase | 13095.24279 | 12799.94603 | 12803.88093 | 14237.3572 | 14305.63996 | 14087.90932 | 0.024255493 | 0.124244176 | 0.013334746 | 2.474701821 | 1.089936568 |
| CNAG_02349 | - | hypothetical protein | 245.5796244 | 311.3971287 | 274.1659546 | 391.1883047 | 356.3898457 | 321.9397093 | 0.024273274 | 0.349325851 | 0.013347485 | 2.474360735 | 1.273965184 |
| CNAG_04512 | - | hypothetical protein | 567.398074 | 518.2560211 | 622.686994 | 479.2382749 | 495.0322841 | 446.4417619 | 0.024304591 | -0.281437853 | 0.013367674 | -2.473820784 | 0.822770599 |
| CNAG_01495 | - | myo-inositol 2-dehydrogenase | 769.8027612 | 808.1613217 | 909.688327 | 933.9946028 | 1005.019544 | 1014.566715 | 0.024333636 | 0.232633061 | 0.01338662 | 2.47331471 | 1.174977443 |
| CNAG_05718 | - | multidrug resistance protein fnx1 | 234.9289173 | 241.4257869 | 232.5755396 | 308.4849708 | 280.370163 | 321.7548427 | 0.024438586 | 0.347644239 | 0.01344734 | 2.471697095 | 1.27248111 |
| CNAG_03787 | - | tubulin alpha-1A chain | 8493.020656 | 8398.575803 | 8810.995647 | 7675.334709 | 8089.063652 | 7879.977694 | 0.024450262 | -0.135829145 | 0.013456751 | -2.471446977 | 0.910146604 |
| CNAG_06285 | - | hypothetical protein | 127.5577428 | 134.7756577 | 131.2691153 | 165.096125 | 199.5952839 | 176.3949361 | 0.024501329 | 0.446859154 | 0.013487848 | 2.47062152 | 1.363069528 |
| CNAG_05468 | APN1 | AP endonuclease 1 | 943.4857993 | 980.4053036 | 973.6269557 | 848.9660753 | 878.9013118 | 798.3376423 | 0.02452621 | -0.213566217 | 0.01350454 | -2.470179144 | 0.862402809 |
| CNAG_05744 | - | 2-dehydropantoate 2-reductase | 331.7680152 | 308.2501839 | 314.6827333 | 405.7594408 | 389.1198205 | 395.1157455 | 0.024642148 | 0.30376093 | 0.013571387 | 2.468412395 | 1.234358044 |
| CNAG_08023 | - | hypothetical protein | 121.8827837 | 172.4040976 | 146.2029609 | 215.270718 | 186.3862675 | 204.1082604 | 0.02471735 | 0.448882867 | 0.013615822 | 2.467242216 | 1.36498289 |
| CNAG_01421 | - | hypothetical protein | 1073.176149 | 1016.064232 | 836.2417339 | 1152.795326 | 1236.249739 | 1115.676688 | 0.024768227 | 0.244765647 | 0.013646873 | 2.466426517 | 1.184900279 |
| CNAG_01723 | - | hypothetical protein | 1852.18393 | 1980.33192 | 1750.133471 | 2072.656983 | 2166.862151 | 2132.551374 | 0.024768321 | 0.175628492 | 0.013649949 | 2.466345788 | 1.129456331 |
| CNAG_04694 | - | periodic tryptophan protein 1 | 6210.073127 | 6106.885008 | 6767.637774 | 5343.473553 | 5728.186447 | 6090.343681 | 0.024785746 | -0.168556946 | 0.013662579 | -2.466014548 | 0.88973219 |
| CNAG_02658 | - | hypothetical protein | 285.1740151 | 286.7059547 | 310.3891575 | 430.9979181 | 364.1021504 | 333.064292 | 0.024835253 | 0.339934725 | 0.013695934 | 2.465141015 | 1.265699326 |
| CNAG_12628 | - | unspecified product | 80.91530924 | 88.46547188 | 92.85962908 | 127.9954639 | 127.8774971 | 124.1979769 | 0.024835253 | 0.524842353 | 0.013695457 | 2.465153487 | 1.438776352 |
| CNAG_12823 | - | unspecified product | 65.73775241 | 61.59865378 | 81.10023204 | 118.1420359 | 115.061129 | 87.71966982 | 0.024930596 | 0.612486612 | 0.013751557 | 2.463688487 | 1.52889212 |
| CNAG_01803 | - | hypothetical protein | 212.9037172 | 47.58858223 | 59.79592203 | 31.8494589 | 30.52170784 | 36.9970612 | 0.025003006 | -1.73452435 | 0.013794551 | -2.462569301 | 0.300508072 |
| CNAG_06782 | MNN2 | putative alpha-1,2-mannosyltransferase | 1128.109443 | 1288.531262 | 1298.867345 | 1096.472156 | 1072.049715 | 1084.706913 | 0.025087695 | -0.206418281 | 0.013844339 | -2.461277101 | 0.866686249 |
| CNAG_10127 | - | unspecified product | 1.304524408 | 0.056125323 | 0.048486978 | 8.27900151 | 2.517525749 | 8.631256198 | 0.025121352 | 4.0521912 | 0.013869048 | 2.460637345 | 16.58941606 |
| CNAG_07276 | - | hypothetical protein | 62.21810728 | 44.43686141 | 82.12571568 | 86.455659 | 118.6721065 | 100.8339886 | 0.025121352 | 0.676298311 | 0.013867082 | 2.460688208 | 1.59803423 |
| CNAG_12297 | - | unspecified product | 10.98747064 | 24.81616204 | 14.02673025 | 29.71082086 | 45.45223757 | 33.15602868 | 0.025238043 | 1.143679032 | 0.013936553 | 2.458894596 | 2.209437363 |

| | | | | | | | | | | | | | |
|---|---|---|---|---|---|---|---|---|---|---|---|---|---|
| CNAG_04117 | - | hypothetical protein | 368.0254054 | 330.918636 | 394.6210121 | 396.0456775 | 505.7533045 | 481.6406487 | 0.025373047 | 0.323859532 | 0.014014201 | 2.456899182 | 1.251674583 |
| CNAG_01726 | - | FACT complex subunit spt16 | 6231.238758 | 6458.156295 | 6062.8486 | 6537.58881 | 7292.217955 | 7202.638343 | 0.025379572 | 0.150463776 | 0.014020904 | 2.456727382 | 1.109926217 |
| CNAG_05144 | CAN2 | Beta-type carbonic anhydrase | 2892.338285 | 2676.322639 | 3329.27726 | 1853.2811 | 2654.986214 | 2128.375119 | 0.025548321 | -0.439571359 | 0.014117249 | -2.454266054 | 0.737353652 |
| CNAG_06294 | - | hypothetical protein | 288.4823388 | 439.2220897 | 291.1550448 | 198.5913647 | 312.3124853 | 239.9880717 | 0.025561612 | -0.45626729 | 0.014127715 | -2.453999583 | 0.728869638 |
| CNAG_12280 | - | unspecified product | 14.54003622 | 29.17103082 | 25.7020221 | 35.02185575 | 50.44320946 | 51.82170311 | 0.025568158 | 0.986679998 | 0.014137577 | 2.453748617 | 1.981619538 |
| CNAG_02977 | - | metal resistance protein YCF1 | 2955.364955 | 2753.878703 | 2546.710146 | 2545.721537 | 2466.8431 | 2344.775074 | 0.025568158 | -0.181852377 | 0.014135368 | -2.45380481 | 0.88157036 |
| CNAG_06839 | - | Methylene-fatty-acyl-phospholipid synthase | 1181.774405 | 1304.694452 | 1283.956256 | 1119.834232 | 1162.702998 | 1017.060597 | 0.02561198 | -0.20724706 | 0.014164935 | -2.453053288 | 0.86618851 |
| CNAG_02793 | - | kynurenine 3-monooxygenase | 1756.310785 | 1927.456336 | 1871.586755 | 1695.025029 | 1681.927567 | 1589.859414 | 0.025663153 | -0.176668568 | 0.014196371 | -2.45225577 | 0.884743665 |
| CNAG_03034 | - | RNA exonuclease 1 | 424.0099855 | 366.4696958 | 427.6887454 | 494.3908862 | 500.4127198 | 506.1180345 | 0.025733367 | 0.28579284 | 0.014238355 | 2.451193105 | 1.219080031 |
| CNAG_04043 | - | hypothetical protein | 257.0073824 | 214.5620343 | 242.1578441 | 314.7375817 | 308.9594113 | 297.4380915 | 0.025795289 | 0.350226857 | 0.014278916 | 2.450169042 | 1.274761061 |
| CNAG_00074 | - | protein PNS1 | 3124.725358 | 3038.300929 | 2716.282632 | 2701.347504 | 2707.521354 | 2533.499794 | 0.025795289 | -0.176219261 | 0.01427595 | -2.450243852 | 0.885019249 |
| CNAG_00451 | - | cytoplasmic protein | 295.3153314 | 279.0405934 | 333.6587009 | 226.1316588 | 244.8339122 | 248.7683075 | 0.025807854 | -0.352729275 | 0.014289023 | -2.449914278 | 0.783101233 |
| CNAG_00349 | - | hypothetical protein | 36.56471829 | 38.93525981 | 42.75399949 | 63.34275622 | 55.23120461 | 81.70660026 | 0.025851347 | 0.763930192 | 0.01431626 | 2.449228476 | 1.698110324 |
| CNAG_01214 | HOC2 | alpha-1,6-mannosyltransferase | 182.3312712 | 142.4228589 | 191.9979657 | 242.4404442 | 217.4474666 | 237.4072432 | 0.025864374 | 0.418399581 | 0.014326633 | 2.44896761 | 1.33644418 |
| CNAG_01044 | - | dihydroceramidase | 1438.836137 | 983.6420989 | 854.2750965 | 757.6820132 | 687.1351902 | 750.6614222 | 0.026008078 | -0.594592034 | 0.014409409 | -2.446891805 | 0.662231695 |
| CNAG_02221 | ZIP3 | solute carrier family 39 (zinc transporter), member 9 | 105.3551585 | 112.1388989 | 98.24518758 | 162.4082313 | 157.2586036 | 129.8823408 | 0.026087041 | 0.493057098 | 0.014456343 | 2.445719475 | 1.407424076 |
| CNAG_07762 | - | hypothetical protein | 7.422215935 | 13.0428587 | 11.84551359 | 30.26887029 | 18.96716563 | 30.7387122 | 0.026158028 | 1.318284715 | 0.014498875 | 2.444660004 | 2.49369447 |
| CNAG_06097 | - | hypothetical protein | 564.3718468 | 706.8140187 | 676.1742801 | 722.2469183 | 790.2302058 | 849.0826162 | 0.026173139 | 0.264516017 | 0.014510447 | 2.444372218 | 1.201233002 |
| CNAG_10048 | - | unspecified product | 6.930060468 | 12.6949386 | 9.49418517 | 2.882859741 | 0 | 2.340843984 | 0.026286962 | -2.766337867 | 0.014576761 | -2.442726967 | 0.146976981 |
| CNAG_07593 | YAP4 | hypothetical protein | 1707.070934 | 1557.958785 | 1595.390787 | 1370.329732 | 1476.647871 | 1461.028215 | 0.02639857 | -0.189624227 | 0.014645098 | -2.441038398 | 0.876834077 |
| CNAG_12777 | - | unspecified product | 2.576458377 | 2.272684363 | 3.287646668 | 7.800854452 | 10.50787049 | 17.32741997 | 0.02639857 | 2.265224966 | 0.014643412 | 2.441079957 | 4.807293763 |
| CNAG_01853 | - | splicing factor 3B subunit 5 | 1333.589572 | 1372.634272 | 1564.378074 | 1241.981886 | 1278.246405 | 1223.496066 | 0.02645214 | -0.205459378 | 0.014681277 | -2.440147234 | 0.867262493 |
| CNAG_05403 | - | ribonuclease P protein subunit RPR2, variant | 319.8792972 | 356.4997422 | 330.5443655 | 294.1278664 | 278.5008162 | 226.723292 | 0.02645214 | -0.347700942 | 0.014678913 | -2.440205395 | 0.785835397 |
| CNAG_05912 | - | hypothetical protein | 838.6780699 | 990.1523136 | 845.7849769 | 1070.354575 | 1085.260385 | 998.0145209 | 0.026496836 | 0.223700704 | 0.014709319 | 2.439457834 | 1.167725118 |
| CNAG_07934 | - | hypothetical protein | 5.977472668 | 2.285987722 | 2.24613557 | 11.11468983 | 12.27164766 | 15.26201118 | 0.026617107 | 2.060995721 | 0.014779336 | 2.437741552 | 4.172741997 |
| CNAG_02394 | - | hypothetical protein | 680.5943248 | 608.7457908 | 670.7287335 | 582.8576347 | 576.1254341 | 485.3459087 | 0.026826882 | -0.26896383 | 0.014899092 | -2.434822614 | 0.829915392 |
| CNAG_00939 | - | glucan 1,3-beta-glucosidase | 2.661889574 | 7.614469834 | 2.268328682 | 11.75816111 | 5.836639332 | 32.39318074 | 0.026828305 | 2.116950307 | 0.014903158 | 2.434723869 | 4.337760221 |
| CNAG_05165 | - | hypothetical protein | 180.1972132 | 206.9606102 | 228.2481727 | 275.0577153 | 270.152982 | 258.5820173 | 0.026865345 | 0.372167792 | 0.014927014 | 2.434144995 | 1.294296178 |
| CNAG_01383 | - | hypothetical protein | 83.82429952 | 75.37780963 | 97.9005524 | 62.05775327 | 59.35029467 | 40.29880322 | 0.026866389 | -0.679588472 | 0.014930875 | -2.434051396 | 0.624343343 |
| CNAG_01625 | - | hypothetical protein | 837.2153196 | 904.9486633 | 832.8786634 | 770.4104146 | 692.4502226 | 760.5214394 | 0.02696067 | -0.226330175 | 0.014986564 | -2.432703589 | 0.854806524 |
| CNAG_06281 | - | hypothetical protein | 0.082148916 | 0.040059538 | 0.034618693 | 2.303090431 | 9.110962669 | 2.239140684 | 0.02700069 | 4.54038389 | 0.015012107 | 2.432086863 | 23.26975136 |
| CNAG_04462 | - | F-box protein 9 | 253.5830016 | 243.6072891 | 251.7585093 | 284.8367657 | 332.2943251 | 341.7275999 | 0.027050158 | 0.342001771 | 0.015042914 | 2.431344264 | 1.267514078 |
| CNAG_05505 | - | serine/threonine-protein phosphatase 2A activator 2 | 831.1212232 | 702.482431 | 716.6404246 | 647.3783944 | 609.6621098 | 652.8453785 | 0.027068902 | -0.252989891 | 0.015056643 | -2.43101375 | 0.839155517 |
| CNAG_03786 | LYSO1 | lysophospholipase | 384.4218826 | 450.3674939 | 366.9749174 | 482.6933245 | 501.2992735 | 495.0261545 | 0.02741567 | 0.286103099 | 0.015252875 | 2.426318571 | 1.219342229 |
| CNAG_05250 | - | telomere maintenance protein | 1203.772569 | 1162.496349 | 1081.387516 | 980.4331506 | 999.599711 | 1043.5952 | 0.027449658 | -0.204660696 | 0.015275137 | -2.42578928 | 0.867742745 |
| CNAG_03397 | - | hypothetical protein | 138.2185762 | 204.6808254 | 146.2505022 | 227.5968859 | 253.112845 | 194.2871833 | 0.027478018 | 0.451984688 | 0.015297629 | 2.425255195 | 1.367920786 |
| CNAG_12006 | - | unspecified product | 6.245753928 | 9.838700788 | 11.82438271 | 17.09992742 | 39.76295209 | 16.67609692 | 0.027478018 | 1.453539803 | 0.01529741 | 2.425260384 | 2.738792189 |
| CNAG_05436 | - | hypothetical protein | 178.9188869 | 197.2293805 | 177.1363331 | 245.454067 | 259.0334721 | 222.0011262 | 0.027579189 | 0.380307265 | 0.015357321 | 2.423841157 | 1.301619044 |
| CNAG_12550 | - | unspecified product | 86.58836964 | 64.8542354 | 83.27036766 | 107.5625247 | 104.4303668 | 142.7838665 | 0.027698814 | 0.583381282 | 0.015427316 | 2.422189202 | 1.498356875 |
| CNAG_07744 | PIK1 | 1-phosphatidylinositol 4-kinase | 1738.572705 | 1537.500857 | 1604.98748 | 1468.522634 | 1434.340633 | 1421.094051 | 0.027798318 | -0.190298585 | 0.015486131 | -2.420806196 | 0.876424315 |
| CNAG_07320 | - | hypothetical protein | 1519.170068 | 1446.99531 | 1623.063749 | 1376.679723 | 1388.662619 | 1294.565211 | 0.027855561 | -0.19273707 | 0.015521422 | -2.419978569 | 0.874944208 |
| CNAG_03414 | - | hypothetical protein | 478.7365053 | 497.7235685 | 534.1982845 | 422.4115203 | 411.2872313 | 431.9307577 | 0.027870229 | -0.270730772 | 0.015532999 | -2.419707435 | 0.828899575 |
| CNAG_00266 | YFH7 | hypothetical protein | 386.5179675 | 472.817086 | 403.0885671 | 367.257961 | 309.4112334 | 348.6872554 | 0.027895466 | -0.314532473 | 0.015550471 | -2.41929857 | 0.804111533 |
| CNAG_03152 | - | calcium-binding protein | 728.8457805 | 670.292598 | 808.3758532 | 902.2803718 | 850.5470451 | 879.1592478 | 0.027900524 | 0.238517026 | 0.015556697 | 2.419152965 | 1.17977932 |
| CNAG_07093 | - | hypothetical protein | 463.6563734 | 436.4286185 | 381.9176553 | 566.9887594 | 514.1785549 | 498.4240695 | 0.028042269 | 0.286652908 | 0.015639155 | 2.417229453 | 1.219807006 |
| CNAG_07508 | - | hypothetical protein | 297.9091353 | 290.9829794 | 266.7231716 | 383.3043713 | 365.8915128 | 329.7041444 | 0.028077191 | 0.321089318 | 0.01566206 | 2.416696736 | 1.249273466 |
| CNAG_01903 | - | hypothetical protein | 331.6333095 | 376.9941793 | 383.8175865 | 283.0053002 | 335.5587963 | 251.1342787 | 0.028147595 | -0.342111384 | 0.01570477 | -2.415705213 | 0.788885932 |
| CNAG_13061 | - | unspecified product | 205.4871568 | 239.0504212 | 228.135886 | 174.8595524 | 168.0003354 | 177.7381802 | 0.028166326 | -0.387364814 | 0.015718661 | -2.415383254 | 0.764524789 |
| CNAG_01722 | - | vacuolar protein sorting-associated protein vps13 | 14038.69792 | 13084.38384 | 11600.98644 | 14511.32281 | 14441.9152 | 14957.11321 | 0.0281671 | 0.166010057 | 0.015722532 | 2.415293571 | 1.121951301 |
| CNAG_07489 | - | hypothetical protein | 250.1277501 | 257.594527 | 261.5419415 | 332.611682 | 318.3945237 | 324.059111 | 0.028170281 | 0.322050674 | 0.015727748 | 2.415172765 | 1.250106211 |
| CNAG_05177 | - | DNA polymerase kappa subunit | 1090.671454 | 1196.946943 | 1151.728668 | 1022.54888 | 1056.602187 | 934.9022194 | 0.028196956 | -0.205448175 | 0.015746083 | -2.414748371 | 0.867269227 |
| CNAG_00970 | - | phosphatidate cytidylyltransferase | 3650.399991 | 3373.501747 | 3169.587202 | 3661.85367 | 3926.477186 | 4025.333805 | 0.028200534 | 0.172694069 | 0.015751526 | 2.414622489 | 1.127161366 |
| CNAG_02093 | - | hypothetical protein | 1127.896319 | 1011.725568 | 1159.153975 | 1019.529345 | 891.7056473 | 942.6214967 | 0.028260025 | -0.224852756 | 0.015788205 | -2.413775079 | 0.855682353 |
| CNAG_00154 | - | oxidoreductase | 332.7946806 | 278.0892937 | 232.6615462 | 386.4045655 | 425.63908 | 304.2637239 | 0.02828203 | 0.389008993 | 0.015803953 | 2.413411787 | 1.309493585 |
| CNAG_13103 | - | unspecified product | 24.52119213 | 12.0416202 | 6.608260548 | 33.98353291 | 32.16950283 | 34.16279247 | 0.028418386 | 1.219927415 | 0.015883618 | 2.411578816 | 2.329349975 |
| CNAG_02575 | - | Rab small monomeric GTPase | 2546.55823 | 2101.013663 | 2217.138437 | 2141.058756 | 2036.429882 | 1799.748062 | 0.028435066 | -0.216076194 | 0.015896414 | -2.411285174 | 0.86090372 |
| CNAG_00988 | - | importin subunit beta-1 | 1394.046801 | 1338.092597 | 1159.265108 | 1157.992486 | 1201.531742 | 981.5848691 | 0.028476398 | -0.236217982 | 0.015922997 | -2.410675753 | 0.848967963 |
| CNAG_01443 | - | hypothetical protein | 384.0781088 | 384.5903613 | 407.2983542 | 301.6473681 | 356.2273456 | 302.1253944 | 0.02863601 | -0.308591419 | 0.016015744 | -2.408556573 | 0.807429712 |
| CNAG_02603 | - | early growth response protein 1 | 33.01599368 | 13.23764118 | 46.92008724 | 62.84800675 | 49.22337398 | 69.53415008 | 0.028643503 | 0.950418671 | 0.016023432 | 2.408381385 | 1.93243337 |
| CNAG_03847 | - | hypothetical protein | 1481.637391 | 1286.46125 | 1343.708027 | 1165.658202 | 1230.804451 | 1215.694985 | 0.028686988 | -0.202984133 | 0.016051262 | -2.40774788 | 0.868751739 |
| CNAG_07342 | - | cytoplasmic protein | 1488.870739 | 1577.308322 | 1574.041403 | 1357.161761 | 1376.555537 | 1414.391286 | 0.028882076 | -0.176755872 | 0.016161946 | -2.405192565 | 0.884690127 |
| CNAG_06934 | - | hexose transporter protein, variant 3 | 220.9995963 | 238.2060374 | 254.9260398 | 288.0724182 | 290.7609397 | 338.3502084 | 0.02889535 | 0.348572868 | 0.016174904 | 2.404944916 | 1.27330044 |
| CNAG_02599 | - | 2,4-dihydroxyhept-2-ene-1,7-dioic acid aldolase | 77.35623303 | 53.06058284 | 99.17506955 | 119.4963329 | 119.2889193 | 114.1942467 | 0.029073226 | 0.603368459 | 0.016278024 | 2.402621543 | 1.519259654 |
| CNAG_01812 | - | large subunit ribosomal protein L24e | 1596.228553 | 1535.331496 | 1717.966694 | 1192.667306 | 1580.269952 | 1398.886477 | 0.029127253 | -0.23315564 | 0.016311831 | -2.401862676 | 0.850771942 |
| CNAG_13133 | - | unspecified product | 27.3291765 | 31.42761813 | 45.88662427 | 47.42707618 | 62.38839747 | 75.04421565 | 0.029245888 | 0.821927822 | 0.016385411 | 2.400215748 | 1.767766619 |
| CNAG_03450 | - | ATP-binding cassette transporter | 3326.790205 | 3264.573581 | 3039.401024 | 2941.837855 | 3014.62142 | 2743.326284 | 0.029245888 | -0.16206263 | 0.016382439 | -2.400282138 | 0.893746362 |
| CNAG_04520 | - | D-aminoacylase | 204.4347504 | 144.5945224 | 196.280973 | 259.6045486 | 244.2783608 | 230.8403566 | 0.029269989 | 0.41357737 | 0.016402489 | 2.399834434 | 1.33198457 |
| CNAG_04353 | CLR1 | hypothetical protein, variant | 296.8144533 | 292.088743 | 310.3953631 | 424.698309 | 357.1587766 | 351.8906913 | 0.029307516 | 0.318794518 | 0.016427097 | 2.399285586 | 1.247287909 |
| CNAG_02231 | - | cytosolic Fe-S cluster assembly factor NBP35 | 680.4093805 | 573.1657057 | 543.9012027 | 451.8066078 | 507.0774651 | 530.6988415 | 0.029313871 | -0.287216706 | 0.016434239 | -2.399126442 | 0.819481505 |
| CNAG_06577 | - | hypothetical protein | 972.5155729 | 1070.773373 | 756.1921407 | 865.474749 | 706.2043902 | 740.6232175 | 0.029346876 | -0.290167367 | 0.016456326 | -2.398634624 | 0.817807179 |
| CNAG_04093 | YRM103 | putative transcription factor | 8.470669049 | 10.87116144 | 4.430718458 | 18.89639314 | 35.38539136 | 12.30991546 | 0.029426251 | 1.547613282 | 0.016504429 | 2.397565513 | 2.923331186 |
| CNAG_01369 | - | hypothetical protein, variant | 47.2528984 | 53.01215448 | 92.73188417 | 112.6959707 | 95.89129888 | 101.9517547 | 0.029447354 | 0.666382152 | 0.016519861 | 2.397223108 | 1.58708803 |
| CNAG_00900 | - | hypothetical protein | 217.9098472 | 147.6922259 | 215.2511164 | 121.6638657 | 144.7397233 | 155.497824 | 0.029736378 | -0.480041524 | 0.016685634 | -2.393562599 | 0.716956988 |
| CNAG_03250 | - | hypothetical protein | 557.1949289 | 525.9046434 | 610.0282264 | 677.2908475 | 701.4259336 | 653.8375357 | 0.029817689 | 0.2477822 | 0.0167349 | 2.392480871 | 1.187380396 |
| CNAG_00975 | - | hypothetical protein | 387.8947577 | 398.7254877 | 398.9180891 | 517.9254325 | 509.1079294 | 430.7752476 | 0.029824254 | 0.284241753 | 0.016742227 | 2.392320251 | 1.217770063 |
| CNAG_04377 | - | vacuolar protein | 426.4363971 | 431.061803 | 461.8115941 | 543.482544 | 542.6948145 | 511.7379827 | 0.029857598 | 0.260780296 | 0.016764591 | 2.391830317 | 1.198126548 |
| CNAG_03078 | NPP1 | type I phosphodiesterase/nucleotide pyrophosphatase | 974.066666 | 956.8462126 | 1041.94865 | 1141.876079 | 1108.425323 | 1208.834478 | 0.029883138 | 0.20328488 | 0.016786229 | 2.391356829 | 1.151316812 |
| CNAG_05962 | - | hypothetical protein, variant 2 | 20.3097697 | 26.010952 | 32.062732 | 59.24729354 | 42.36475481 | 45.37977076 | 0.029883138 | 0.889809817 | 0.016785186 | 2.39137965 | 1.852931845 |
| CNAG_04241 | - | cytoplasmic protein, variant | 950.628343 | 979.4003507 | 789.3072877 | 1043.008275 | 1073.99301 | 1101.165681 | 0.029903361 | 0.228308742 | 0.016804893 | 2.390948877 | 1.171460852 |
| CNAG_03942 | - | THO complex subunit 2 | 8001.334017 | 7784.424043 | 7737.101817 | 6546.690852 | 7199.734994 | 7604.593822 | 0.029903361 | -0.155163285 | 0.016803227 | -2.390985276 | 0.898030728 |
| CNAG_00194 | - | hypothetical protein | 1781.902454 | 1750.801697 | 1795.867317 | 1467.567435 | 1667.309027 | 1615.334788 | 0.030115985 | -0.180854614 | 0.016928059 | -2.388266546 | 0.882180262 |
| CNAG_04202 | - | cytosolic Fe-S cluster assembly factor NAR1 | 1909.539393 | 1170.148848 | 1360.744523 | 1001.997325 | 1093.546436 | 996.0377505 | 0.030126792 | -0.540301029 | 0.016937813 | -2.388054861 | 0.687627416 |
| CNAG_03958 | - | hypothetical protein | 727.6487236 | 760.7006974 | 683.6862881 | 892.1894327 | 802.1924186 | 868.0465553 | 0.030140711 | 0.224735143 | 0.016949319 | 2.387805282 | 1.168562698 |
| CNAG_01139 | - | hypothetical protein | 24.68625696 | 12.08475555 | 19.32503033 | 25.4945797 | 38.2016306 | 55.97802713 | 0.030297733 | 1.084224121 | 0.017041319 | 2.385815059 | 2.120234917 |
| CNAG_04036 | - | hypothetical protein | 70.45951436 | 65.93108251 | 100.2488583 | 129.763033 | 133.2278215 | 96.62069654 | 0.030300414 | 0.591057668 | 0.017046527 | 2.385702674 | 1.506350677 |
| CNAG_05168 | - | hypothetical protein, variant | 34.99500417 | 15.31433733 | 16.1872962 | 44.85468433 | 43.38926812 | 44.18443783 | 0.030317128 | 0.993364821 | 0.017059632 | 2.385420013 | 1.990822809 |
| CNAG_06919 | NOP1 | rRNA 2'-O-methyltransferase fibrillarin | 920.1416193 | 920.105569 | 1003.45159 | 866.9478602 | 802.907846 | 809.4090679 | 0.030562563 | -0.21412357 | 0.017201472 | -2.382372841 | 0.862069704 |

| | | | | | | | | | | | | | |
|---|---|---|---|---|---|---|---|---|---|---|---|---|---|
| CNAG_04161 | - | hypothetical protein | 978.4899776 | 916.9059342 | 1087.668744 | 934.7371394 | 779.5420204 | 837.1681305 | 0.030624393 | -0.240830018 | 0.017240011 | -2.381548692 | 0.846258299 |
| CNAG_04908 | CLR4 | hypothetical protein, variant | 55.26119885 | 81.8620503 | 41.81535608 | 77.80301092 | 112.6349954 | 98.57767858 | 0.030664553 | 0.683829503 | 0.017266364 | 2.380986091 | 1.606398139 |
| CNAG_03229 | YOX101 | specific transcriptional repressor | 4.845321532 | 2.280375747 | 2.241322703 | 12.01520306 | 15.96267775 | 8.853865803 | 0.0308029 | 2.111813953 | 0.017348024 | 2.379247482 | 4.322344172 |
| CNAG_13001 | - | unspecified product | 40.96067866 | 71.83233744 | 58.50319539 | 43.55336622 | 20.24281468 | 29.10711555 | 0.030913145 | -0.88988334 | 0.017413889 | -2.377850404 | 0.539657755 |
| CNAG_12135 | - | unspecified product | 19.14963135 | 24.93577801 | 32.05210646 | 51.17381675 | 37.9716041 | 54.05432264 | 0.031075437 | 0.906457524 | 0.017509106 | 2.375838895 | 1.874437242 |
| CNAG_00889 | - | hypothetical protein, variant | 19.94309347 | 13.08224745 | 5.535694118 | 28.35987316 | 42.12962756 | 21.1236979 | 0.031166395 | 1.281405342 | 0.017564161 | 2.374680195 | 2.430756437 |
| CNAG_05634 | - | protein phosphatase 2 (formerly 2A), regulatory subunit B' | 503.3100843 | 409.5621853 | 430.9400981 | 536.3721571 | 623.0880523 | 506.2331223 | 0.031210346 | 0.294827793 | 0.017592741 | 2.37407994 | 1.226738539 |
| CNAG_07324 | - | hypothetical protein | 440.7225827 | 675.5098629 | 583.407895 | 712.3790375 | 722.1355763 | 681.5712698 | 0.031239191 | 0.303194335 | 0.017612815 | 2.373658845 | 1.233873365 |
| CNAG_12433 | - | unspecified product | 0.198987573 | 5.435196907 | 1.170133913 | 12.12879082 | 6.8834034 | 15.18531168 | 0.03124663 | 2.451908329 | 0.017620825 | 2.373490939 | 5.471393544 |
| CNAG_04107 | - | hypothetical protein | 157.4780695 | 193.6737281 | 136.5266979 | 125.8693023 | 108.4103198 | 120.0405281 | 0.031348828 | -0.475258836 | 0.017682285 | -2.372204826 | 0.719337721 |
| CNAG_01770 | - | mitochondrion organization and biogenesis-related protein, putative | 909.4444781 | 890.9466362 | 761.4774882 | 782.9184502 | 733.0682328 | 670.7049522 | 0.031361526 | -0.243727827 | 0.017693278 | -2.371975211 | 0.844560204 |
| CNAG_04396 | - | hypothetical protein | 179.0933923 | 257.4919746 | 211.2558377 | 311.0365111 | 272.6735403 | 268.5928321 | 0.031506422 | 0.381584102 | 0.017778871 | 2.370191564 | 1.302771534 |
| CNAG_12665 | - | unspecified product | 7.394959593 | 9.84457548 | 13.93097053 | 22.05045571 | 19.01702917 | 32.83239516 | 0.031507104 | 1.317429837 | 0.017786951 | 2.370023571 | 2.492217253 |
| CNAG_01314 | PPP1 | vacuolar protein, variant | 1477.197889 | 1387.775374 | 1159.368613 | 1587.112361 | 1575.135705 | 1556.380569 | 0.031507104 | 0.21460684 | 0.017786697 | 2.370028846 | 1.160387645 |
| CNAG_01395 | - | Ribose-5-phosphate isomerase | 1145.445192 | 1110.802791 | 1119.744693 | 1023.358553 | 999.6209186 | 961.5070432 | 0.03150906 | -0.193085288 | 0.017791903 | -2.369920655 | 0.874733052 |
| CNAG_01250 | TRL1 | tRNA ligase | 1292.796064 | 1246.658072 | 1249.940838 | 1347.181703 | 1538.97892 | 1483.084968 | 0.031937653 | 0.19045003 | 0.018037813 | 2.364841011 | 1.141119617 |
| CNAG_04277 | - | hypothetical protein | 1773.798988 | 1831.60189 | 1911.025389 | 1610.101242 | 1585.256105 | 1738.52334 | 0.03196215 | -0.175961282 | 0.018055551 | -2.364476951 | 0.88517752 |
| CNAG_00916 | - | transcription factor | 2125.093541 | 1911.354985 | 1923.868775 | 1595.16315 | 1873.473055 | 1796.292675 | 0.032044128 | -0.19469191 | 0.018105774 | -2.363447867 | 0.873759469 |
| CNAG_03275 | - | glutaryl-CoA dehydrogenase, variant | 910.6630856 | 844.6847277 | 854.2023606 | 735.2612192 | 783.1439843 | 754.9949962 | 0.032166296 | -0.214555196 | 0.018178729 | -2.361957431 | 0.861811828 |
| CNAG_06938 | - | hypothetical protein | 9.860276916 | 15.26963993 | 27.73397587 | 41.12329965 | 28.47346499 | 42.93669598 | 0.032225216 | 1.107654266 | 0.018215964 | 2.361198777 | 2.154949807 |
| CNAG_07814 | - | hypothetical protein | 1002.265786 | 1153.997963 | 1294.660437 | 1439.362556 | 1401.808131 | 1250.056645 | 0.032316308 | 0.230886619 | 0.018271402 | 2.360071734 | 1.173555945 |
| CNAG_03114 | - | hypothetical protein | 364.6026955 | 474.0095949 | 301.9931598 | 542.9951289 | 479.7096252 | 445.1590469 | 0.032345894 | 0.350405792 | 0.018292079 | 2.359652125 | 1.274919178 |
| CNAG_01241 | - | enzyme regulator | 1975.593515 | 1783.126502 | 1704.202028 | 1717.671906 | 1545.56679 | 1562.100712 | 0.032398992 | -0.194733196 | 0.018326063 | -2.358963394 | 0.873734465 |
| CNAG_05002 | - | hypothetical protein | 1.377962516 | 1.178823669 | 2.209939726 | 11.45702372 | 11.56202078 | 4.494800317 | 0.03248861 | 2.683897926 | 0.018380122 | 2.357858015 | 6.425897309 |
| CNAG_04593 | - | hypothetical protein | 13.30098709 | 15.27453724 | 25.63612931 | 50.12674683 | 34.67983988 | 28.86114925 | 0.032522191 | 1.063979688 | 0.018403692 | 2.357394339 | 2.090690765 |
| CNAG_02445 | - | Phosphoacetylglucosamine mutase | 1352.415058 | 1368.404205 | 1435.473134 | 1539.521249 | 1562.158074 | 1647.367207 | 0.03254053 | 0.177273138 | 0.018418043 | 2.357104898 | 1.130744625 |
| CNAG_01481 | - | hypothetical protein | 77.45668143 | 113.1044013 | 67.38394449 | 138.2933974 | 104.3218164 | 145.1049678 | 0.032585318 | 0.577672754 | 0.018447373 | 2.356513992 | 1.492439816 |
| CNAG_12142 | - | unspecified product | 6.202541717 | 7.691640161 | 11.79739254 | 23.09258808 | 16.4247459 | 25.2122627 | 0.032607103 | 1.400298079 | 0.018463687 | 2.356185651 | 2.639561132 |
| CNAG_06718 | - | hypothetical protein | 0.189051731 | 3.318870426 | 2.223598338 | 18.27695047 | 7.820479356 | 5.597064712 | 0.032637362 | 2.573828371 | 0.018484807 | 2.355760987 | 5.953872681 |
| CNAG_05873 | - | hypothetical protein | 144.6301879 | 91.76716032 | 72.7481597 | 162.395144 | 149.4110883 | 148.6183522 | 0.032689968 | 0.563554244 | 0.018522585 | 2.355002417 | 1.477905726 |
| CNAG_12984 | - | unspecified product | 6.223612087 | 10.88372281 | 10.76044749 | 26.35726108 | 17.28411064 | 24.17724976 | 0.032689968 | 1.352044833 | 0.018520353 | 2.355047204 | 2.552736865 |
| CNAG_00329 | - | NADH dehydrogenase (ubiquinone) 1 beta subcomplex 8 | 490.4712812 | 487.0117548 | 642.8712615 | 412.925885 | 480.4070443 | 431.9854272 | 0.032772167 | -0.307078001 | 0.018573163 | -2.353988961 | 0.808277167 |
| CNAG_07487 | - | F-type H -transporting ATPase subunit D | 11411.23791 | 11092.17235 | 12100.99015 | 10449.18228 | 10619.78383 | 10808.56372 | 0.032894502 | -0.13389402 | 0.018646511 | -2.352523514 | 0.911368227 |
| CNAG_00499 | - | solute carrier family 25 (mitochondrial carnitine/acylcarnitine transporter), member 20/29 | 865.0061147 | 707.9009825 | 786.996081 | 659.7712227 | 655.4030507 | 698.3447715 | 0.032958738 | -0.244928653 | 0.018686949 | -2.351717749 | 0.843857527 |
| CNAG_12232 | - | unspecified product | 22.26591784 | 15.23068119 | 6.60787514 | 33.16383549 | 28.60152816 | 37.40153398 | 0.03297745 | 1.172823353 | 0.018701585 | 2.351426484 | 2.254524758 |
| CNAG_12288 | - | unspecified product | 2.674121485 | 8.679021829 | 4.387209649 | 19.33523357 | 14.83605037 | 14.31924564 | 0.033079936 | 1.735747314 | 0.018767784 | 2.350111583 | 3.330519698 |
| CNAG_02877 | FZC51 | hypothetical protein | 488.9665157 | 434.1599247 | 410.5827206 | 367.1367831 | 388.084302 | 346.5279269 | 0.033079936 | -0.290505007 | 0.018765306 | -2.350160731 | 0.817615807 |
| CNAG_06134 | BZP1 | hypothetical protein | 4207.26538 | 3841.053533 | 4055.691051 | 3433.568232 | 3749.508226 | 3774.576234 | 0.033093626 | -0.15934469 | 0.018779592 | -2.349877462 | 0.895431707 |
| CNAG_00502 | - | large subunit ribosomal protein L35 | 137.0720854 | 104.4220404 | 92.77613638 | 98.19355637 | 45.20210596 | 69.0444086 | 0.033113964 | -0.667004658 | 0.018795177 | -2.349568661 | 0.629812957 |
| CNAG_05294 | - | F-box and WD-40 domain-containing protein CDC4 | 397.204624 | 437.4495864 | 349.9275244 | 486.6368542 | 518.6256217 | 454.0333951 | 0.033121753 | 0.286705994 | 0.018803643 | 2.349401017 | 1.219851892 |
| CNAG_03440 | - | hypothetical protein | 383.1498776 | 380.3971592 | 349.8957924 | 453.9378397 | 431.3278034 | 478.318992 | 0.033269697 | 0.277528931 | 0.018895758 | 2.347581127 | 1.212116974 |
| CNAG_01677 | - | ER to Golgi transporter Yif1 | 451.8372951 | 443.87786 | 500.0597454 | 388.944833 | 362.0726786 | 410.8062962 | 0.033269697 | -0.280382409 | 0.01889225 | -2.347650285 | 0.82337274 |
| CNAG_00124 | CAS32 | Capsule structure designer protein | 336.7436549 | 443.9419253 | 444.7441118 | 429.4373742 | 615.4393746 | 519.4394931 | 0.033331444 | 0.339218702 | 0.018934898 | 2.346810194 | 1.265071303 |
| CNAG_06376 | - | vacuolar membrane protein | 2658.921273 | 2730.168835 | 2671.442475 | 2313.631066 | 2409.905231 | 2576.685891 | 0.033338164 | -0.158149972 | 0.018942786 | -2.346654982 | 0.896173535 |
| CNAG_05719 | - | hypothetical protein | 728.7339761 | 873.6752946 | 811.5215361 | 722.1596971 | 695.1188667 | 659.5689211 | 0.033649021 | -0.230992681 | 0.019123525 | -2.343114275 | 0.852048417 |
| CNAG_02806 | - | solute carrier family 30 (zinc transporter), member 1, variant | 1193.095293 | 1037.531455 | 955.5656638 | 941.6225743 | 976.3988751 | 792.8643086 | 0.033673545 | -0.249494718 | 0.019145686 | -2.342682145 | 0.841190978 |
| CNAG_07708 | - | hypothetical protein | 54.95006419 | 37.89589834 | 37.48832786 | 72.82676502 | 73.60740965 | 66.48777352 | 0.033673545 | 0.704643682 | 0.019144198 | 2.342711148 | 1.6297421 |
| CNAG_00564 | - | hypothetical protein | 379.8246545 | 479.4461147 | 402.145653 | 486.4962948 | 535.8330421 | 524.9855299 | 0.033750229 | 0.279868774 | 0.019193408 | 2.341753095 | 1.214084448 |
| CNAG_01358 | - | hypothetical protein | 26.07632339 | 34.56328341 | 27.86585571 | 56.62112688 | 56.42110018 | 43.26601578 | 0.033794464 | 0.818657712 | 0.019222691 | 2.341184004 | 1.763764217 |
| CNAG_05701 | TPM1 | tropomyosin | 5666.051467 | 5551.00054 | 5951.892277 | 6621.108385 | 6332.11372 | 6166.932208 | 0.034078523 | 0.139848545 | 0.019388428 | 2.337977256 | 1.101789443 |
| CNAG_04024 | ITR5 | myo-inositol transporter | 176.630132 | 203.690618 | 188.8496097 | 217.8902754 | 248.5779037 | 279.4830116 | 0.034225608 | 0.377336918 | 0.019476289 | 2.336286989 | 1.298941914 |
| CNAG_12233 | - | unspecified product | 9.721295534 | 14.11808857 | 12.90999458 | 26.8506173 | 29.67351073 | 25.39155572 | 0.034356459 | 1.178438484 | 0.019554946 | 2.334779431 | 2.263316721 |
| CNAG_12530 | - | unspecified product | 7.603673798 | 31.26083671 | 22.5122251 | 41.67629159 | 37.22571772 | 49.56866654 | 0.03437179 | 1.070914238 | 0.01956787 | 2.334532247 | 2.100764203 |
| CNAG_06719 | FZC49 | hypothetical protein | 412.5352093 | 498.8791579 | 479.9510562 | 519.904612 | 596.2350198 | 574.9448138 | 0.034475918 | 0.267448835 | 0.01963136 | 2.333319939 | 1.203677442 |
| CNAG_06518 | - | hypothetical protein | 4.848070919 | 0.105956863 | 4.34624572 | 8.594706317 | 15.95699482 | 13.11821322 | 0.034723419 | 2.15001901 | 0.019780773 | 2.330480437 | 4.438336372 |
| CNAG_12414 | - | unspecified product | 148.0820327 | 151.7945772 | 141.7791986 | 114.192426 | 111.1091908 | 99.01578176 | 0.034723419 | -0.456704325 | 0.019776562 | -2.330560204 | 0.728648875 |
| CNAG_06063 | - | cytochrome c oxidase assembly protein subunit 15 | 983.1154968 | 943.8034245 | 1005.596666 | 701.5364441 | 832.2556919 | 955.7952449 | 0.034728206 | -0.251765544 | 0.019791982 | -2.330268178 | 0.839867971 |
| CNAG_07574 | - | hypothetical protein | 78.24246879 | 112.8421299 | 89.50526817 | 65.08033108 | 54.01733565 | 66.73696847 | 0.034728206 | -0.606280601 | 0.01978869 | -2.330330498 | 0.656888038 |
| CNAG_03631 | - | hypothetical protein | 949.3683046 | 803.8730453 | 813.790473 | 971.3766318 | 995.4742776 | 1065.608231 | 0.034834666 | 0.225055174 | 0.019856908 | 2.329040727 | 1.168821948 |
| CNAG_00841 | - | hypothetical protein | 332.8169995 | 366.2559852 | 401.9216563 | 277.5013086 | 275.7976441 | 335.2659862 | 0.034857465 | -0.323998472 | 0.019874161 | -2.328715154 | 0.798852762 |
| CNAG_02269 | - | autophagy-like protein 18 Atg18 | 195.3127492 | 225.2375429 | 211.2383542 | 238.1084896 | 282.2645499 | 297.2914627 | 0.034900332 | 0.357977565 | 0.019902864 | 2.328174048 | 1.281627994 |
| CNAG_03348 | - | hypothetical protein | 1839.068843 | 1688.340851 | 1805.463155 | 1531.462767 | 1673.348853 | 1563.192981 | 0.034902494 | -0.177346907 | 0.019908359 | -2.328070534 | 0.884327766 |
| CNAG_00361 | RVS167 | hypothetical protein | 2920.452698 | 2746.380307 | 2946.566449 | 2585.338148 | 2618.663497 | 2631.134339 | 0.034946708 | -0.152158717 | 0.019937846 | -2.327515486 | 0.899902922 |
| CNAG_12571 | - | unspecified product | 13.41284434 | 29.18980006 | 30.99269488 | 52.0148123 | 56.59097646 | 33.36932413 | 0.034979459 | 0.95570694 | 0.019960802 | 2.327083858 | 1.939529797 |
| CNAG_11009 | - | unspecified product | 44.20240705 | 11.062645 | 18.32934921 | 52.7389406 | 42.36495988 | 57.34522174 | 0.035183955 | 1.029297844 | 0.020081793 | 2.324816143 | 2.041030646 |
| CNAG_07760 | - | hypothetical protein | 28.52067194 | 41.05960923 | 44.85928763 | 49.72552992 | 80.80272302 | 65.29796491 | 0.035542686 | 0.765005036 | 0.020290885 | 2.320925131 | 1.69937593 |
| CNAG_05297 | - | hypothetical protein | 374.7689612 | 394.2622775 | 395.5848373 | 322.0074328 | 294.7740769 | 343.0807236 | 0.035592353 | -0.293132399 | 0.020323586 | -2.320319767 | 0.816128145 |
| CNAG_04504 | - | hypothetical protein | 186.1621781 | 132.5076565 | 130.1039137 | 111.0269517 | 95.44476045 | 114.4545049 | 0.035606013 | -0.501632019 | 0.020335733 | -2.320095098 | 0.706307334 |
| CNAG_05038 | - | hypothetical protein | 11.17306238 | 42.04537762 | 34.22520091 | 54.87327344 | 74.02354335 | 43.31110988 | 0.035675866 | 0.962537737 | 0.020379986 | 2.319277666 | 1.948734755 |
| CNAG_03122 | - | hypothetical protein | 49.24421842 | 44.31133706 | 39.61078599 | 67.97810269 | 79.74273552 | 67.59592099 | 0.035746754 | 0.67464905 | 0.020424846 | 2.318450586 | 1.596208432 |
| CNAG_02999 | - | hypothetical protein | 739.5218177 | 928.753622 | 887.3237467 | 958.9045015 | 1044.719835 | 1015.705212 | 0.03589992 | 0.22588543 | 0.020516745 | 2.316761177 | 1.169494786 |
| CNAG_07867 | - | hypothetical protein, variant | 1262.214889 | 884.7521858 | 1014.284749 | 1230.582353 | 1289.664173 | 1306.534965 | 0.035951915 | 0.258771147 | 0.02055085 | 2.316135891 | 1.196459155 |
| CNAG_02117 | - | NAD-binding Rossmann fold oxidoreductase | 679.5824206 | 707.8174003 | 714.4692458 | 599.8678766 | 622.6942372 | 600.6919678 | 0.036224204 | -0.220824853 | 0.02071092 | -2.313213204 | 0.858074697 |
| CNAG_06635 | - | CCR4-NOT transcription complex subunit 3 | 1955.81974 | 1821.921007 | 1855.601458 | 1483.840994 | 1833.844608 | 1653.108253 | 0.036325725 | -0.19619527 | 0.0207734 | -2.312077735 | 0.872849442 |
| CNAG_01110 | - | co-chaperone | 780.043427 | 841.4065253 | 833.9144928 | 724.4529377 | 691.6345264 | 727.2188101 | 0.036444722 | -0.211082762 | 0.020845901 | -2.310763881 | 0.863888628 |
| CNAG_06364 | - | hypothetical protein | 1973.495923 | 1712.137891 | 1775.708269 | 2109.32807 | 2054.681706 | 2079.268589 | 0.036581066 | 0.177316374 | 0.020928354 | 2.309274489 | 1.130778513 |
| CNAG_01289 | - | hypothetical protein | 97.31847131 | 123.9650586 | 117.388565 | 136.199913 | 161.5299448 | 172.915703 | 0.036626612 | 0.462375156 | 0.020958885 | 2.308724307 | 1.377808283 |
| CNAG_04398 | ARO80 | specific RNA polymerase II transcription factor | 1271.848207 | 1382.363819 | 1249.958966 | 1408.694516 | 1457.768963 | 1651.732601 | 0.036688156 | 0.196163854 | 0.020998582 | 2.308009972 | 1.145648008 |
| CNAG_12564 | - | unspecified product | 20.43573278 | 44.22283665 | 36.3718367 | 63.56130365 | 80.93430356 | 40.09699094 | 0.036917159 | 0.85819128 | 0.02113416 | 2.305579131 | 1.812764206 |
| CNAG_01859 | - | hypothetical protein | 679.8670212 | 687.5036413 | 804.0999152 | 830.5285144 | 884.2758914 | 853.6359222 | 0.036946821 | 0.226938897 | 0.021155653 | 2.305195029 | 1.170349072 |
| CNAG_06035 | - | Alcohol dehydrogenase | 209.2430697 | 231.6836965 | 192.0875404 | 292.2824845 | 279.6622067 | 244.2340378 | 0.03700555 | 0.352033083 | 0.0211938 | 2.304514127 | 1.276358037 |
| CNAG_04227 | - | hypothetical protein | 2954.150826 | 2377.94839 | 2779.120626 | 2237.309582 | 2363.311902 | 2535.619374 | 0.037156176 | -0.201004719 | 0.021284604 | -2.302897627 | 0.869944507 |
| CNAG_01493 | - | hypothetical protein, variant | 670.8598319 | 317.8821023 | 306.1275541 | 237.57291 | 247.2266557 | 221.2542697 | 0.03718008 | -0.893802973 | 0.021302837 | -2.302573756 | 0.538193558 |
| CNAG_12506 | - | unspecified product | 22.53922565 | 49.35188418 | 67.81673437 | 16.44188517 | 32.71576559 | 20.2519768 | 0.037433982 | -1.035441164 | 0.021452885 | -2.299917659 | 0.487866672 |
| CNAG_02776 | - | hypothetical protein | 81.96156021 | 95.90402578 | 53.54805846 | 102.0379275 | 136.7707441 | 109.7915234 | 0.037508095 | 0.585124393 | 0.021504519 | 2.299007399 | 1.500168332 |
| CNAG_01633 | - | anaphase-promoting complex subunit 5 | 865.3954222 | 803.8562056 | 848.9444592 | 927.7491591 | 984.2837662 | 1021.221816 | 0.037508095 | 0.205228612 | 0.021504148 | 2.299013931 | 1.152869018 |
| CNAG_00186 | - | mitotic spindle organizing protein 1 | 206.4187062 | 191.7173738 | 195.0927496 | 139.6970537 | 179.4410765 | 131.2370827 | 0.037541356 | -0.414802034 | 0.021528173 | -2.298591038 | 0.750122419 |

| | | | | | | | | | | | | | |
|---|---|---|---|---|---|---|---|---|---|---|---|---|---|
| CNAG_04328 | - | ubiquitin-conjugating enzyme E2 I | 1119.818544 | 1126.963444 | 1175.175313 | 1040.487512 | 1044.512766 | 939.3376501 | 0.037572174 | -0.193403668 | 0.021550434 | -2.298199559 | 0.874540033 |
| CNAG_07870 | - | hypothetical protein | 0.082979192 | 0.040461953 | 0.034966173 | 0.612840207 | 6.297215407 | 7.501616998 | 0.037608041 | 4.542678806 | 0.021575599 | 2.297757435 | 23.30679634 |
| CNAG_12324 | - | unspecified product | 51.55619084 | 46.45652052 | 38.55844885 | 63.92742126 | 70.92243029 | 85.11971205 | 0.037721726 | 0.676185316 | 0.021645426 | 2.296532977 | 1.597909073 |
| CNAG_12001 | - | unspecified product | 21.47534628 | 30.28526571 | 31.02038098 | 56.72017298 | 56.48080505 | 37.78174679 | 0.037772455 | 0.846859491 | 0.021679148 | 2.295942879 | 1.798581445 |
| CNAG_03031 | - | protein FRA10AC1 | 204.6535818 | 217.7440548 | 241.0520678 | 314.9350322 | 263.9893105 | 273.0262917 | 0.037888779 | 0.345572292 | 0.021750537 | 2.294696263 | 1.270654929 |
| CNAG_07751 | - | siderophore iron transporter mirB | 186.8267822 | 217.515331 | 216.3857203 | 163.9925169 | 166.3493189 | 150.0543541 | 0.038016778 | -0.384768204 | 0.021828659 | -2.293336159 | 0.765902045 |
| CNAG_00983 | - | hypothetical protein | 566.4180658 | 536.625636 | 482.1490759 | 594.6400869 | 692.8834725 | 621.6273897 | 0.038032638 | 0.252235989 | 0.02184241 | 2.293097186 | 1.19105166 |
| CNAG_05068 | - | GTP-binding protein ypt1 | 5202.2853 | 5108.098343 | 5333.295753 | 4960.003751 | 4773.337702 | 4678.299126 | 0.038071127 | -0.133732028 | 0.021869164 | -2.292632639 | 0.911470565 |
| CNAG_12097 | - | unspecified product | 80.63739407 | 105.4276219 | 112.8548251 | 71.32556495 | 63.56319899 | 68.988641 | 0.038090173 | -0.570497246 | 0.021884755 | -2.29236213 | 0.673384657 |
| CNAG_00107 | - | hypothetical protein | 420.5061976 | 425.6308877 | 362.7173427 | 445.9256004 | 551.4734191 | 489.4952341 | 0.038185312 | 0.283793661 | 0.021944081 | 2.291334398 | 1.21739189 |
| CNAG_05284 | - | hypothetical protein | 454.6171945 | 603.3474737 | 533.2895797 | 672.7448934 | 641.8602294 | 613.8789237 | 0.038223069 | 0.262416548 | 0.021970447 | 2.290878419 | 1.19948619 |
| CNAG_04523 | - | glyceraldehyde-3-phosphate dehydrogenase, type I | 12.11594857 | 20.56175373 | 17.18359502 | 32.21837517 | 44.59820769 | 26.63392927 | 0.038241973 | 1.043264418 | 0.021985982 | 2.290609964 | 2.060885593 |
| CNAG_12096 | - | unspecified product | 4.48440354 | 3.196023636 | 4.200288783 | 0.647333892 | 0 | 0.046501109 | 0.03827608 | -4.333685804 | 0.022010265 | -2.290190684 | 0.049594165 |
| CNAG_00799 | - | cellulase | 4952.412068 | 4461.713282 | 4195.518723 | 5167.200877 | 5376.485353 | 4885.817531 | 0.038328307 | 0.165346534 | 0.022044978 | 2.289592014 | 1.121435413 |
| CNAG_05679 | - | pre-mRNA-splicing factor ini1 | 2176.547092 | 2025.565893 | 2065.700794 | 1898.909987 | 1894.107557 | 1881.790843 | 0.038535784 | -0.158971272 | 0.022169017 | -2.287459499 | 0.895663505 |
| CNAG_13195 | - | unspecified product | 3.821928874 | 8.681347142 | 4.388564415 | 14.23377915 | 15.73656435 | 18.59858989 | 0.038815842 | 1.635587293 | 0.02233961 | 2.284543492 | 3.107140098 |
| CNAG_05547 | - | hypothetical protein | 60.94754854 | 51.8882558 | 60.87766581 | 85.90650521 | 99.59355252 | 78.76465261 | 0.038815842 | 0.599444226 | 0.022336502 | 2.284596437 | 1.515132775 |
| CNAG_06184 | - | hypothetical protein, variant | 496.5923849 | 610.9058228 | 552.4849625 | 650.0073987 | 648.7266836 | 682.6234008 | 0.0390058 | 0.241676386 | 0.022453699 | 2.282604105 | 1.182365752 |
| CNAG_02722 | - | aldose reductase | 3175.098413 | 3495.101109 | 3123.645032 | 2830.427718 | 3135.408131 | 2913.140649 | 0.039058156 | -0.156378269 | 0.022488608 | -2.28201241 | 0.897274758 |
| CNAG_02052 | - | dimeric dihydrodiol dehydrogenase | 15.58551037 | 26.96678614 | 15.09760018 | 35.33155836 | 36.45116455 | 40.79527406 | 0.039178704 | 0.974043493 | 0.0225628 | 2.280757514 | 1.964338408 |
| CNAG_12927 | - | unspecified product | 8.637602209 | 17.3383148 | 16.09892104 | 24.18124092 | 44.76821201 | 24.39291988 | 0.03922829 | 1.151679532 | 0.022596146 | 2.280194651 | 2.221723888 |
| CNAG_02094 | - | ARP2/3 complex 34 kDa subunit | 1907.894001 | 1671.108799 | 1751.09469 | 1643.681755 | 1555.94904 | 1558.75332 | 0.039262579 | -0.17988427 | 0.022620692 | -2.279780804 | 0.882773808 |
| CNAG_06684 | - | pre-rRNA-processing protein IPI3 | 43.64255453 | 42.23227295 | 70.37303607 | 75.67641266 | 93.56859515 | 80.88568554 | 0.039308297 | 0.662002313 | 0.022656632 | 2.279175543 | 1.582277139 |
| CNAG_01061 | FRK102 | serine/threonine protein kinase | 200.906585 | 170.3589146 | 164.3569822 | 214.8747636 | 251.27595 | 235.2201995 | 0.039308297 | 0.374675957 | 0.022652346 | 2.279247688 | 1.296548305 |
| CNAG_03977 | - | hypothetical protein | 1597.462444 | 1646.282275 | 1704.138172 | 1452.897684 | 1407.556114 | 1569.772188 | 0.039472371 | -0.174936335 | 0.022756022 | -2.277506078 | 0.885806608 |
| CNAG_02511 | CPK1 | Calcium-dependent protein kinase 1 | 350.6165326 | 391.1760634 | 412.7447155 | 506.2706478 | 445.1242979 | 458.4291829 | 0.039570235 | 0.274406097 | 0.022817273 | 2.276480392 | 1.209496083 |
| CNAG_02334 | - | hypothetical protein | 2.557183855 | 4.378688818 | 0.081276517 | 7.004355716 | 17.90689236 | 7.742396301 | 0.039655317 | 2.35598042 | 0.022871176 | 2.275579724 | 5.119420175 |
| CNAG_00909 | - | phosphopantothenoylcysteine decarboxylase | 216.8263833 | 187.4224077 | 185.525886 | 142.0626643 | 138.6242797 | 169.8826612 | 0.039741993 | -0.404590447 | 0.02292602 | -2.274665231 | 0.75545072 |
| CNAG_02587 | - | alfa-L-rhamnosidase | 734.6837782 | 937.2546132 | 966.0869005 | 694.6897212 | 743.4038989 | 800.4513857 | 0.039809738 | -0.251701517 | 0.022969961 | -2.2739339 | 0.839905245 |
| CNAG_07774 | - | hypothetical protein | 30.85956463 | 31.47225766 | 60.74696658 | 72.09787538 | 76.29411942 | 59.89381124 | 0.039830918 | 0.758452299 | 0.022987046 | 2.273649877 | 1.691674848 |
| CNAG_02598 | CHI21 | Chitinase | 114.7666965 | 115.4338323 | 145.0539954 | 160.4795468 | 176.1902944 | 174.1282881 | 0.039847373 | 0.431971027 | 0.023001408 | 2.273411263 | 1.349075441 |
| CNAG_01583 | - | vacuolar-sorting protein SNF7 | 1116.268684 | 1119.402678 | 1103.746614 | 1082.666276 | 960.7771002 | 874.984224 | 0.039997135 | -0.210265729 | 0.023092741 | -2.27189686 | 0.864378007 |
| CNAG_00409 | - | hypothetical protein | 1051.151908 | 1073.189649 | 1136.861973 | 1212.641759 | 1233.527354 | 1298.774498 | 0.040041815 | 0.1841418 | 0.023123427 | 2.271389214 | 1.136140929 |
| CNAG_00691 | - | hypothetical protein | 602.7377688 | 666.9426213 | 575.9796474 | 764.5652324 | 682.295011 | 738.1596183 | 0.040119134 | 0.230318193 | 0.023172976 | 2.270570739 | 1.173093652 |
| CNAG_07537 | - | mitochondrial GTPase 1, variant | 735.6841253 | 776.7861246 | 827.4850724 | 690.210117 | 670.9555226 | 676.1742752 | 0.040204264 | -0.214651452 | 0.023227057 | -2.26967915 | 0.86175433 |
| CNAG_13016 | - | unspecified product | 64.50864391 | 34.50964026 | 42.58965984 | 25.24731894 | 29.14451733 | 23.5508941 | 0.040264911 | -0.880324191 | 0.023267012 | -2.269021608 | 0.543245344 |
| CNAG_12285 | - | unspecified product | 68.14499927 | 100.1992589 | 71.60098894 | 132.1659906 | 116.6453815 | 103.214294 | 0.04037506 | 0.540726254 | 0.023335591 | 2.267895257 | 1.454704632 |
| CNAG_07798 | - | amidohydrolase 3 | 43.54147764 | 43.25508243 | 50.20685888 | 83.96670381 | 80.57935368 | 56.67205891 | 0.040427324 | 0.671614482 | 0.023370735 | 2.267319169 | 1.592854492 |
| CNAG_00633 | - | hypothetical protein | 2265.296031 | 2136.530815 | 2145.686087 | 1810.793929 | 2001.084403 | 2069.350439 | 0.040470445 | -0.170267002 | 0.023400606 | -2.266830117 | 0.888678197 |
| CNAG_12095 | - | unspecified product | 8.716051966 | 25.89942086 | 20.36775763 | 42.74236306 | 32.90778225 | 36.44871344 | 0.040679861 | 1.041394834 | 0.02352666 | 2.264772234 | 2.058216627 |
| CNAG_07434 | HAD3 | HAD hydrolase, family IIID | 695.7941334 | 648.5866272 | 660.0987708 | 629.6503319 | 551.0195802 | 537.4656345 | 0.040852043 | -0.237751782 | 0.023631228 | -2.263072372 | 0.848065863 |
| CNAG_03572 | OPS1 | opsin 1 | 800.6823469 | 681.9761243 | 614.3115995 | 641.2851641 | 564.8070436 | 558.5572105 | 0.040864635 | -0.26462431 | 0.023643502 | -2.262873273 | 0.832415475 |
| CNAG_11007 | - | unspecified product | 43.46643622 | 47.48939771 | 34.29969972 | 67.28903042 | 56.95134234 | 80.65743039 | 0.040871183 | 0.696922854 | 0.023652282 | 2.26273091 | 1.621043555 |
| CNAG_00410 | - | hypothetical protein | 1433.772206 | 1290.747936 | 1263.747536 | 1223.401545 | 1155.716502 | 1154.660701 | 0.040904955 | -0.190042117 | 0.023676821 | -2.262333264 | 0.876580131 |
| CNAG_05137 | - | ER lumen protein retaining receptor | 107.7901557 | 115.417822 | 139.7249425 | 175.5523945 | 167.5356699 | 152.0414848 | 0.040950821 | 0.437203729 | 0.02370837 | 2.261822533 | 1.353977467 |
| CNAG_05816 | - | hypothetical protein | 935.2088975 | 856.5559181 | 913.9036326 | 792.9250539 | 762.3624091 | 816.0154085 | 0.04099426 | -0.205731876 | 0.023738525 | -2.261334924 | 0.867098698 |
| CNAG_02323 | - | hypothetical protein | 2391.435414 | 2426.309567 | 2160.655585 | 2151.114642 | 2029.483691 | 2120.461995 | 0.041049631 | -0.162227513 | 0.023775602 | -2.260736133 | 0.893644223 |
| CNAG_01012 | - | hypothetical protein | 266.5334232 | 274.8600233 | 321.0151664 | 330.0104938 | 405.7263419 | 349.6299314 | 0.041069851 | 0.316712599 | 0.023792328 | 2.260466263 | 1.245489275 |
| CNAG_07521 | - | hypothetical protein | 139.4431625 | 196.1390341 | 199.4571096 | 251.0218335 | 251.2619295 | 209.8212582 | 0.041171545 | 0.398585545 | 0.023856268 | 2.259436143 | 1.318214865 |
| CNAG_07412 | - | hypothetical protein | 0.14901533 | 3.282781205 | 0.062449956 | 9.829466798 | 7.0168943 | 7.652447721 | 0.04121936 | 2.930145038 | 0.023889007 | 2.258909622 | 7.621870194 |
| CNAG_12903 | - | unspecified product | 5.342003579 | 42.02118899 | 32.09603843 | 61.45825849 | 60.82621829 | 45.45699103 | 0.041464178 | 1.060730952 | 0.024035957 | 2.256554007 | 2.085988135 |
| CNAG_00928 | - | peptidase | 876.9104689 | 884.5320277 | 907.4992731 | 725.8538315 | 770.1551272 | 837.0661546 | 0.041564102 | -0.208813549 | 0.024098957 | -2.255547941 | 0.865248506 |
| CNAG_05181 | BWC1 | white collar 1 protein | 549.9884279 | 485.9986994 | 412.864421 | 584.6663803 | 648.0515005 | 543.975775 | 0.041564165 | 0.279976428 | 0.024104069 | 2.255466403 | 1.214175046 |
| CNAG_02375 | FIG4 | polyphosphoinositide phosphatase | 1801.981484 | 1829.534344 | 1875.92263 | 1978.424104 | 2167.756258 | 2062.626778 | 0.041610328 | 0.157839458 | 0.024135921 | 2.254958706 | 1.115615171 |
| CNAG_00911 | - | phytanoyl-CoA dioxygenase | 1.394088896 | 0.087144259 | 4.313716034 | 8.787268148 | 13.35940021 | 7.714490684 | 0.041649384 | 2.466552849 | 0.024163661 | 2.254517016 | 5.527215449 |
| CNAG_00577 | - | hypothetical protein | 663.2265495 | 694.8797066 | 689.9453433 | 566.3476669 | 560.4995499 | 639.4555748 | 0.041959478 | -0.228065656 | 0.024348692 | -2.251582121 | 0.853778859 |
| CNAG_05585 | - | arf/Sar family protein, variant | 672.5816394 | 694.8976423 | 718.7256673 | 626.4291654 | 585.5387463 | 599.5769089 | 0.042275573 | -0.218487567 | 0.024537281 | -2.2486106 | 0.859465976 |
| CNAG_00920 | - | YjeF family protein | 726.2141926 | 725.0543844 | 693.1764434 | 606.8530208 | 613.1608526 | 647.2667803 | 0.042323045 | -0.215493848 | 0.024570003 | -2.248097031 | 0.861251295 |
| CNAG_01922 | - | alternative cyclin Pho80 | 204.6198238 | 210.2057953 | 229.3331555 | 315.7989685 | 267.4968692 | 245.3560475 | 0.042425785 | 0.347806394 | 0.024634828 | 2.247081355 | 1.272624141 |
| CNAG_05138 | - | glucan 1,3-beta-glucosidase | 3631.510428 | 3472.511885 | 3100.210974 | 3066.337373 | 3257.032128 | 2828.830749 | 0.042436443 | -0.172491637 | 0.024646199 | -2.246903437 | 0.887308911 |
| CNAG_04237 | - | hypothetical protein | 1114.097015 | 1268.063444 | 1265.817514 | 1098.836007 | 1087.608459 | 1050.311401 | 0.042449923 | -0.187770899 | 0.024659212 | -2.246699915 | 0.877961208 |
| CNAG_00996 | PMT4 | dolichyl-phosphate-mannose-protein mannosyltransferase | 2559.539201 | 2266.936279 | 2464.532818 | 2224.260432 | 2025.121559 | 2279.181127 | 0.042499179 | -0.174710938 | 0.024698204 | -2.246090619 | 0.885945011 |
| CNAG_04457 | FZC30 | hypothetical protein | 17.90996702 | 22.74903608 | 22.50394035 | 42.54824333 | 51.48389971 | 26.74053834 | 0.042499179 | 0.948765118 | 0.02469602 | 2.246124727 | 1.930219769 |
| CNAG_12157 | - | unspecified product | 7.160443072 | 4.43497418 | 1.198141117 | 20.32297459 | 15.80987889 | 8.925409007 | 0.042598573 | 1.824478947 | 0.024761168 | 2.245108499 | 3.541790669 |
| CNAG_03639 | - | hypothetical protein | 204.416682 | 257.3525058 | 247.3155149 | 172.4052431 | 201.7743946 | 182.2079178 | 0.04267545 | -0.365872942 | 0.024811066 | -2.244331724 | 0.775999193 |
| CNAG_13078 | - | unspecified product | 0.077058031 | 0.037591038 | 0.032487053 | 6.741802662 | 2.596374344 | 3.290062118 | 0.043022172 | 4.302676047 | 0.0250179 | 2.241126221 | 19.73488281 |
| CNAG_00936 | - | lipid particle protein | 162.3586414 | 148.7598694 | 107.9072703 | 189.4003164 | 171.6851974 | 211.7649791 | 0.043063913 | 0.438472458 | 0.025047431 | 2.24067042 | 1.355168699 |
| CNAG_03019 | - | long-chain acyl-CoA synthetase | 1627.729546 | 1601.024332 | 1551.676774 | 1317.356005 | 1455.080795 | 1503.171616 | 0.043132295 | -0.176046713 | 0.025092472 | -2.239976138 | 0.885125105 |
| CNAG_01531 | - | DNA helicase II/ATP-dependent DNA helicase PcrA | 899.2816827 | 874.9457621 | 891.6046857 | 997.0611393 | 1093.047354 | 989.1271423 | 0.043166397 | 0.193364771 | 0.025117582 | 2.239589542 | 1.143427403 |
| CNAG_04079 | - | HAT1-interacting factor 1 | 792.7752466 | 733.7337357 | 813.6340031 | 692.551416 | 696.8784591 | 648.4595774 | 0.043401716 | -0.215229701 | 0.025259809 | -2.237406115 | 0.861408998 |
| CNAG_07375 | - | guanyl-nucleotide exchange factor | 1638.42046 | 1610.803381 | 1504.841515 | 1850.966756 | 1735.497878 | 1799.509325 | 0.043504545 | 0.164985223 | 0.025324968 | 2.236409361 | 1.121154595 |
| CNAG_02827 | RUB1 | ubiquitin-like protein Nedd8 | 1049.809269 | 1110.788011 | 1177.28331 | 885.365042 | 1091.193606 | 931.5452542 | 0.043575151 | -0.213981032 | 0.025371391 | -2.235700584 | 0.86215488 |
| CNAG_01235 | - | DNA-directed RNA polymerase II subunit RPB2, variant | 11649.4145 | 11287.17025 | 11325.72237 | 10344.88071 | 10769.89403 | 10788.57773 | 0.043618863 | -0.118366413 | 0.025402169 | -2.235231284 | 0.921230184 |
| CNAG_06157 | - | hypothetical protein | 2313.132779 | 2032.057822 | 2194.72658 | 1876.238271 | 1976.918185 | 2024.966276 | 0.043908493 | -0.170050872 | 0.025576201 | -2.232586866 | 0.888811339 |
| CNAG_01277 | - | hypothetical protein | 892.4071761 | 1110.789803 | 857.545762 | 1233.979252 | 1133.511648 | 1030.252006 | 0.043931866 | 0.234169807 | 0.025595181 | 2.232299413 | 1.176229686 |
| CNAG_00047 | PKP1 | pyruvate dehydrogenase kinase | 529.9623231 | 479.4477768 | 523.5593965 | 481.8002217 | 424.2446694 | 376.5536333 | 0.043943119 | -0.272956694 | 0.025607103 | -2.232118941 | 0.827621659 |
| CNAG_06396 | - | hypothetical protein | 55.01605285 | 46.46632096 | 39.62462959 | 71.89754655 | 81.43684131 | 69.82618468 | 0.044088041 | 0.642775566 | 0.025696937 | 2.230761411 | 1.561330076 |
| CNAG_07548 | - | cytoplasmic protein | 639.856537 | 660.4071881 | 661.1497451 | 564.0896419 | 613.2812295 | 508.6309692 | 0.044191584 | -0.233469979 | 0.025762684 | -2.229770479 | 0.850586593 |
| CNAG_02655 | - | hypothetical protein | 1292.594113 | 1262.720316 | 1272.247477 | 1226.60698 | 1065.962265 | 1106.925868 | 0.044204626 | -0.186816706 | 0.025775686 | -2.22957478 | 0.87854208 |
| CNAG_13047 | - | unspecified product | 0.127526584 | 2.211443909 | 0.053539578 | 9.085080431 | 8.006763614 | 3.368050634 | 0.044327068 | 3.311631285 | 0.025852495 | 2.228420391 | 9.928882044 |
| CNAG_01204 | - | ATP synthase F1, delta subunit | 3267.192962 | 3053.418655 | 2906.097297 | 2804.064649 | 2896.451009 | 2678.921136 | 0.044437564 | -0.154665114 | 0.025922365 | -2.227372862 | 0.898340876 |
| CNAG_02095 | - | cyclin | 888.7296079 | 837.2386606 | 834.0319436 | 1001.834695 | 944.5278772 | 1016.802696 | 0.044463425 | 0.195175091 | 0.02594288 | 2.227065748 | 1.144863097 |
| CNAG_01317 | - | hypothetical protein | 1283.296291 | 1263.811189 | 1338.344004 | 1098.705764 | 1192.010713 | 1180.150475 | 0.044591157 | -0.178012377 | 0.026022853 | -2.225870564 | 0.883919948 |
| CNAG_06205 | BLP3 | pr4/barwin domain protein | 191.2584941 | 192.7368211 | 175.9219615 | 178.4038376 | 125.6556962 | 112.4174691 | 0.044703652 | -0.442239985 | 0.026093962 | -2.224810504 | 0.735990992 |
| CNAG_02729 | - | hypothetical protein | 202.422776 | 260.7813949 | 258.1242471 | 264.5235254 | 342.7531876 | 319.5651091 | 0.04473832 | 0.346840509 | 0.026119662 | 2.224428002 | 1.271772403 |
| CNAG_00012 | - | hypothetical protein | 1551.93233 | 1550.421255 | 1736.101506 | 1428.780534 | 1450.747294 | 1458.810386 | 0.044756516 | -0.17313189 | 0.02613575 | -2.224188709 | 0.88691522 |
| CNAG_12181 | - | unspecified product | 27.18831923 | 22.82587985 | 35.24352033 | 47.78512232 | 55.59112477 | 46.50246544 | 0.044785135 | 0.800750639 | 0.026157931 | 2.223859014 | 1.742007263 |

| | | | | | | | | | | | | | |
|---|---|---|---|---|---|---|---|---|---|---|---|---|---|
| CNAG_05999 | - | peptide alpha-N-acetyltransferase | 609.4867258 | 547.3790934 | 721.8268471 | 502.4898062 | 518.2569004 | 558.452703 | 0.044801955 | -0.266181177 | 0.026173227 | -2.223631805 | 0.831517668 |
| CNAG_02190 | - | hypothetical protein | 586.1356353 | 534.4408012 | 686.6452461 | 507.2445686 | 533.8629381 | 484.171874 | 0.044823583 | -0.260901259 | 0.026191335 | -2.223362963 | 0.834566399 |
| CNAG_00445 | NHP6B01 | nonhistone chromosomal protein | 1840.157883 | 1568.770307 | 1728.680471 | 1563.495844 | 1578.443767 | 1417.814863 | 0.044984691 | -0.188336479 | 0.026290967 | -2.221886641 | 0.877617088 |
| CNAG_07352 | - | hypothetical protein | 643.1290497 | 512.9011918 | 604.6067948 | 533.8646534 | 496.7194126 | 448.6872543 | 0.045050837 | -0.268384555 | 0.026335127 | -2.221233841 | 0.830248689 |
| CNAG_12561 | - | unspecified product | 13.33607105 | 24.8693112 | 24.60748135 | 44.21630056 | 39.93738135 | 34.33989597 | 0.04512952 | 0.92578076 | 0.026386634 | 2.220473631 | 1.899712054 |
| CNAG_01457 | BSP1 | hypothetical protein | 735.9234633 | 861.9487042 | 782.8378954 | 936.4658869 | 923.8959388 | 904.7312295 | 0.045380332 | 0.20187539 | 0.026555447 | 2.217990992 | 1.150192544 |
| CNAG_05409 | - | replication factor C subunit 2/4 | 742.4825 | 720.7320194 | 655.8731485 | 562.3907918 | 566.5165965 | 680.4956133 | 0.045380332 | -0.242658606 | 0.026551586 | -2.218047618 | 0.845186362 |
| CNAG_05189 | - | hypothetical protein | 1059.389766 | 1243.35664 | 1135.828631 | 1251.50628 | 1374.206346 | 1328.78637 | 0.045380332 | 0.187155826 | 0.026550487 | 2.21806373 | 1.138516994 |
| CNAG_03824 | - | solute carrier family 25 (mitochondrial phosphate transporter), member 3 | 3839.270785 | 3650.250034 | 2869.906086 | 3109.938355 | 3047.334531 | 2956.463575 | 0.045380332 | -0.20010818 | 0.026548824 | -2.218088129 | 0.870485288 |
| CNAG_07670 | - | hypothetical protein | 11.98382126 | 13.05558404 | 12.90809505 | 25.20496254 | 32.37601077 | 22.14856194 | 0.045435294 | 1.118633241 | 0.026593157 | 2.217438268 | 2.171411631 |
| CNAG_05387 | - | galactose transporter | 623.4457195 | 322.4521808 | 259.4569497 | 717.3519964 | 719.7204358 | 786.6937953 | 0.045462471 | 0.866006344 | 0.026614615 | 2.217124058 | 1.822610576 |
| CNAG_10039 | - | unspecified product | 13.28589111 | 9.936569964 | 29.83290928 | 48.5536427 | 30.26175336 | 31.01096163 | 0.045544242 | 1.078931384 | 0.026673609 | 2.216261342 | 2.112470777 |
| CNAG_12632 | - | unspecified product | 17.83230913 | 22.70367994 | 15.0874636 | 37.87412004 | 35.6095071 | 33.17407401 | 0.045544242 | 0.94516072 | 0.0266685 | 2.216335983 | 1.925403369 |
| CNAG_00079 | - | hypothetical protein | 76.13048901 | 53.03843743 | 89.59472817 | 102.1712024 | 108.9031589 | 114.1172121 | 0.045623968 | 0.555031091 | 0.026725873 | 2.215498414 | 1.469200296 |
| CNAG_02191 | - | hypothetical protein | 928.1564544 | 839.3080654 | 852.0759584 | 775.0426443 | 790.9106503 | 731.7177487 | 0.045656113 | -0.205593279 | 0.026750279 | -2.215142594 | 0.867182003 |
| CNAG_06490 | SNF102 | CAMK/CAMKL protein kinase | 1253.981169 | 1180.83144 | 1129.374658 | 1022.46747 | 1028.051651 | 1110.180374 | 0.045842739 | -0.188085022 | 0.02687082 | -2.213389264 | 0.877770067 |
| CNAG_01571 | - | hypothetical protein | 1111.736069 | 1243.283955 | 1228.500369 | 1001.404096 | 1064.314871 | 1112.404376 | 0.045842739 | -0.187905555 | 0.026869939 | -2.213402051 | 0.877879266 |
| CNAG_03189 | - | DIL and Ankyrin domain-containing protein | 1943.06909 | 1944.732269 | 1931.32334 | 1680.892697 | 1751.779279 | 1843.989764 | 0.045861076 | -0.15681314 | 0.026892769 | -2.21307074 | 0.897004333 |
| CNAG_06795 | - | hypothetical protein | 126.21526 | 120.7442851 | 90.82714197 | 167.8188368 | 153.7033735 | 144.2571687 | 0.045861076 | 0.450510659 | 0.026890399 | 2.213105117 | 1.366523869 |
| CNAG_12970 | - | unspecified product | 1.470319213 | 8.636973037 | 2.254525874 | 13.5917471 | 14.04548616 | 13.1705406 | 0.045877537 | 1.852872779 | 0.026913627 | 2.212768261 | 3.612187493 |
| CNAG_05023 | CAS91 | maltose O-acetyltransferase | 935.5806445 | 1026.827783 | 1015.30029 | 1184.783242 | 1156.797732 | 1081.291484 | 0.045877537 | 0.186063587 | 0.026913143 | 2.212775271 | 1.137655369 |
| CNAG_02024 | - | hypothetical protein | 956.2994546 | 977.1719322 | 928.8654173 | 823.2313762 | 840.8988628 | 872.6279862 | 0.045967629 | -0.189605587 | 0.026972091 | -2.211921471 | 0.876845406 |
| CNAG_02798 | - | hypothetical protein | 872.1420422 | 770.3849906 | 876.5539964 | 695.5313237 | 735.6694298 | 762.7205807 | 0.045971429 | -0.215819756 | 0.026979935 | -2.21180799 | 0.861056758 |
| CNAG_03735 | CAP4 | beta-1,2-xylosyltransferase | 809.0286154 | 785.3709728 | 685.754884 | 705.1046875 | 622.6009274 | 638.4542432 | 0.046082851 | -0.22899679 | 0.027050954 | -2.210781748 | 0.853227997 |
| CNAG_12332 | - | unspecified product | 34.06892457 | 19.31227355 | 12.84797681 | 8.944416922 | 9.256759293 | 9.064112871 | 0.046092912 | -1.339177669 | 0.027062489 | -2.210615297 | 0.395245881 |
| CNAG_07888 | - | hypothetical protein | 33565.93088 | 31436.32119 | 33858.84746 | 36593.76206 | 35455.5949 | 36648.57927 | 0.046168741 | 0.12128681 | 0.027112648 | 2.209892147 | 1.087704607 |
| CNAG_13134 | - | unspecified product | 58.61627271 | 59.35896447 | 49.20929006 | 80.38645881 | 104.9100106 | 71.05644036 | 0.04619481 | 0.603800525 | 0.027133598 | 2.209590454 | 1.519714717 |
| CNAG_01971 | - | hypothetical protein | 4.958400591 | 2.319409347 | 8.587837969 | 21.01455635 | 22.09788893 | 5.723839541 | 0.046213125 | 1.758956125 | 0.027149999 | 2.209354413 | 3.384531454 |
| CNAG_06313 | - | phosphoglucomutase, variant | 7912.707203 | 7550.689761 | 7132.511214 | 8537.995153 | 8221.10745 | 8254.012163 | 0.046227496 | 0.131330768 | 0.027164087 | 2.209151755 | 1.095303564 |
| CNAG_13058 | - | unspecified product | 5.003257612 | 0.145409756 | 12.80158834 | 20.82509664 | 17.45108887 | 18.67421758 | 0.046245524 | 1.674576461 | 0.027185975 | 2.208837076 | 3.192256241 |
| CNAG_06023 | - | hypothetical protein | 66.32844859 | 60.29186166 | 75.53828667 | 44.82038686 | 39.39060577 | 46.75655472 | 0.046245524 | -0.655394415 | 0.027185139 | -2.208849088 | 0.634901893 |
| CNAG_02068 | - | stromal membrane-associated protein | 330.4117912 | 356.5397068 | 352.9258508 | 309.704757 | 249.0086121 | 293.1628761 | 0.046454862 | -0.301319296 | 0.02731471 | -2.206990688 | 0.811509958 |
| CNAG_12968 | - | unspecified product | 35.24618705 | 37.7805768 | 22.59165493 | 56.51290827 | 52.82258714 | 53.12029647 | 0.046655018 | 0.761770815 | 0.027438096 | 2.205228046 | 1.695570552 |
| CNAG_01122 | - | hypothetical protein | 41.20805369 | 46.43916249 | 44.89720301 | 73.63681831 | 57.79054904 | 78.51437575 | 0.046810633 | 0.652554058 | 0.02753533 | 2.203843815 | 1.571948614 |
| CNAG_01846 | - | flavoprotein | 2979.421814 | 2340.113592 | 1955.934128 | 1966.554293 | 1807.716384 | 1737.604918 | 0.046820854 | -0.416845107 | 0.02754706 | -2.203677114 | 0.749060885 |
| CNAG_01438 | SWI6 | Cell-cycle box factor subunit SWI6, putative | 195.2783391 | 240.2571265 | 173.9870028 | 304.1451368 | 265.8283799 | 227.6225846 | 0.046899643 | 0.374435108 | 0.027599142 | 2.202937665 | 1.296331872 |
| CNAG_13129 | - | unspecified product | 38.23074292 | 43.93434625 | 33.01221887 | 26.31834688 | 23.11558589 | 10.34134775 | 0.046975207 | -0.965317461 | 0.027649347 | -2.202226025 | 0.512165698 |
| CNAG_05991 | - | glycosyl hydrolase family 88, variant | 129.8367417 | 104.7133025 | 149.3164022 | 181.7310181 | 183.1154656 | 157.6110993 | 0.047060891 | 0.431346876 | 0.027705526 | 2.201431 | 1.348491919 |
| CNAG_12765 | - | unspecified product | 15.4589673 | 13.09314028 | 16.09998525 | 33.24690052 | 28.64498565 | 29.80272602 | 0.04719756 | 1.016774662 | 0.027791749 | 2.200213521 | 2.023390339 |
| CNAG_12767 | - | unspecified product | 155.7061422 | 198.2980439 | 199.4668493 | 225.0364616 | 251.2347361 | 239.669145 | 0.047274503 | 0.358664309 | 0.027842829 | 2.199493808 | 1.282238213 |
| CNAG_12196 | - | unspecified product | 9.721469271 | 21.52585176 | 3.399505955 | 33.49769073 | 19.82136304 | 30.7816669 | 0.047820422 | 1.303287677 | 0.028170194 | 2.19490806 | 2.467906397 |
| CNAG_07800 | - | syntaxin-binding protein 1 | 630.4392087 | 638.8382763 | 590.8019277 | 542.3330342 | 463.8013285 | 578.3949962 | 0.047839598 | -0.245900348 | 0.028187332 | -2.19466925 | 0.843289357 |
| CNAG_06838 | - | mitochondrial RNA helicase | 1470.07973 | 1341.420457 | 1512.162452 | 1229.473982 | 1235.916945 | 1374.377523 | 0.048028628 | -0.187268225 | 0.028304574 | -2.193038928 | 0.878267166 |
| CNAG_07162 | - | origin recognition complex subunit 2 | 326.0868092 | 333.0457714 | 404.1778351 | 470.4786533 | 433.1194825 | 406.3000699 | 0.048071196 | 0.284896047 | 0.028335531 | 2.192609432 | 1.218322475 |
| CNAG_12729 | - | unspecified product | 0.176648753 | 1.184280722 | 4.310824601 | 11.40113931 | 6.940577915 | 10.89064626 | 0.048087454 | 2.41846777 | 0.028350986 | 2.192395155 | 5.346029394 |
| CNAG_03136 | - | hypothetical protein | 1533.162093 | 1468.5305 | 1538.851336 | 1307.314661 | 1361.038507 | 1417.699864 | 0.048115163 | -0.167455706 | 0.028373198 | -2.192087373 | 0.890411601 |
| CNAG_12271 | - | unspecified product | 28.78718425 | 25.73422709 | 25.4925681 | 11.23111979 | 8.277780644 | 17.81210155 | 0.048158405 | -1.130382517 | 0.028404579 | -2.191652907 | 0.456794594 |
| CNAG_00978 | - | NADH dehydrogenase (ubiquinone) 1 alpha subcomplex 9 | 115.192341 | 108.6779397 | 87.46005953 | 72.83216511 | 72.24619739 | 74.52498637 | 0.048162711 | -0.523148561 | 0.028413 | -2.191536381 | 0.695851537 |
| CNAG_00407 | - | glyoxal oxidase | 3042.395276 | 1385.632301 | 1320.34371 | 1095.276667 | 1080.432383 | 955.1029682 | 0.04827163 | -0.895656538 | 0.02848315 | -2.190566876 | 0.537502534 |
| CNAG_02267 | - | cell cycle arrest protein BUB3 | 705.0913976 | 581.8588078 | 703.7691161 | 611.7092897 | 532.8799659 | 549.648234 | 0.048737292 | -0.249348891 | 0.02876387 | -2.186707686 | 0.841276009 |
| CNAG_03841 | - | hypothetical protein | 871.0294584 | 814.5396916 | 901.0833216 | 807.0648518 | 740.8183293 | 716.1826344 | 0.048819311 | -0.208075494 | 0.028824199 | -2.185882551 | 0.865691264 |
| CNAG_07799 | - | ABC transporter | 800.2796759 | 1108.630183 | 661.4233186 | 1136.500747 | 1178.436855 | 1325.227596 | 0.048819311 | 0.489414724 | 0.028821773 | 2.185915708 | 1.403875233 |
| CNAG_05602 | PUT2 | 1-pyrroline-5-carboxylate dehydrogenase | 16933.38204 | 15440.67324 | 14898.36811 | 13707.80727 | 14965.41016 | 14686.35969 | 0.0488634 | -0.140322417 | 0.028856198 | -2.185445501 | 0.907316363 |
| CNAG_06059 | - | cytoplasmic protein | 47.05412794 | 58.24795063 | 48.12023695 | 75.76472038 | 85.73288743 | 74.2692939 | 0.048905729 | 0.59834691 | 0.028887167 | 2.185022904 | 1.5139808 |
| CNAG_12734 | - | unspecified product | 114.2921806 | 73.44398045 | 55.68737441 | 128.8962025 | 96.52842497 | 146.1391265 | 0.048937856 | 0.596429061 | 0.02891212 | 2.184682697 | 1.511969525 |
| CNAG_06802 | - | hypothetical protein | 0.077847441 | 0.037973933 | 0.03281771 | 4.954613184 | 0 | 8.533393225 | 0.048961054 | 4.430318803 | 0.028931803 | 2.184414503 | 21.56050109 |
| CNAG_00187 | UBP16 | ubiquitin carboxyl-terminal hydrolase 1 | 1787.762134 | 1859.584882 | 1723.387253 | 1563.387564 | 1609.452116 | 1688.574756 | 0.048962702 | -0.158657868 | 0.028938756 | -2.184319806 | 0.895858095 |
| CNAG_05910 | - | hypothetical protein | 187.3596195 | 299.5042781 | 266.6644765 | 341.2648831 | 335.7417455 | 297.4991843 | 0.049054736 | 0.357999089 | 0.028999142 | 2.183498189 | 1.281647115 |
| CNAG_05055 | - | DNA/RNA-binding protein KIN17 | 841.8966662 | 787.629345 | 970.3454328 | 749.3179327 | 731.3141342 | 773.837593 | 0.049183927 | -0.221195777 | 0.029081521 | -2.182379704 | 0.85785411 |
| CNAG_04151 | - | oxysterol-binding protein | 3899.034889 | 3956.27828 | 3623.822782 | 3501.407925 | 3457.054194 | 3596.953277 | 0.049238759 | -0.136107275 | 0.029119955 | -2.181858805 | 0.909971159 |
| CNAG_05254 | - | solute carrier family 35 (UDP-galactose transporter), member B1 | 1170.074769 | 1215.307693 | 1264.754119 | 1088.692665 | 1033.201173 | 1135.718445 | 0.049459116 | -0.179433365 | 0.029256314 | -2.18001548 | 0.883049756 |
| CNAG_00762 | DPH1 | diphthamide biosynthesis protein 1 | 907.1495497 | 749.951278 | 938.3760111 | 759.4689626 | 759.8286718 | 722.8326312 | 0.049481218 | -0.227900656 | 0.02927543 | -2.179757654 | 0.85387651 |
| CNAG_03628 | - | hypothetical protein | 1241.30672 | 1428.549277 | 1223.226206 | 1104.951021 | 1079.762657 | 1246.696967 | 0.049710655 | -0.197063525 | 0.029417247 | -2.177849449 | 0.872324295 |
| CNAG_11016 | - | unspecified product | 83.94264922 | 29.40915027 | 58.77076346 | 101.0072363 | 80.27697054 | 98.53820975 | 0.049860028 | 0.696145802 | 0.02951173 | 2.17658253 | 1.620170677 |
