## Supplementary Dataset 2 for "Systematic histone mutagenesis reveals nucleosome-dependent maintenance of three-dimensional chromosome architecture and virulence in *Cryptococcus neoformans*"

Supplementary Dataset 2. List of DEGs genes overlapping with TADs and A/B compartments

| Gene_ID | gene_nam | Function | WT(H99)_1_DESeq2_normalized_coun | WT(H99)_2_DESeq2_normalized_coun | WT(H99)_3_DESeq2_normalized_coun | *hht2*Δ mutant_1_DESeq2_normalized_cou | *hht2*Δ mutant_2_DESeq2_normalized_cou | *hht2*Δ mutant_3_DESeq2_normalized_cou | padj | log2FoldC | pvalue | stat | foldChang | log10padj |
|---|---|---|---|---|---|---|---|---|---|---|---|---|---|---|
| CNAG_048 | PNB1 | para-nitro | 441.2590452 | 524.0974819 | 521.895184 | 4818.043232 | 4833.305419 | 4657.328648 | 1.00E-50 | 3.252488 | 0 | 42.26458 | 9.530078 | |
| CNAG_063 | - | hypothetic | 1120.856185 | 1259.961762 | 1224.671209 | 7118.94875 | 7781.460243 | 7665.87664 | 1.00E-50 | 2.63187 | 0 | 39.06731 | 6.198288 | |
| CNAG_058 | - | endoribon | 289.4359451 | 284.9050809 | 307.5334983 | 2631.762714 | 2566.375465 | 2642.331183 | 1.00E-50 | 3.139844 | 2.38E-292 | 36.54215 | 8.814289 | 288.188 |
| CNAG_015 | BLP4 | pr4/barwir | 963.187364 | 965.8222163 | 725.5854329 | 5972.911176 | 5528.239829 | 5931.788289 | 1.00E-50 | 2.701169 | 1.86E-195 | 29.82493 | 6.503287 | 191.5171 |
| CNAG_029 | - | D-arabinit | 10220.87953 | 9634.747023 | 9716.751835 | 28094.23145 | 28836.99861 | 28778.13103 | 1.00E-50 | 1.519779 | 5.55E-180 | 28.60668 | 2.867471 | 176.1876 |
| CNAG_048 | - | transitiona | 13346.94877 | 13331.39914 | 13279.75974 | 35129.06279 | 37812.83056 | 36874.56056 | 1.00E-50 | 1.443271 | 6.51E-173 | 28.03258 | 2.719367 | 169.176 |
| CNAG_005 | - | mannitol c | 145.780625 | 139.4431383 | 120.9025635 | 1243.289479 | 1232.900902 | 1213.619982 | 1.00E-50 | 3.175204 | 5.57E-166 | 27.45806 | 9.032992 | 162.3412 |
| CNAG_027 | PHO84 | phosphate | 3989.069171 | 3644.55089 | 3926.249108 | 940.5793641 | 1074.947441 | 994.1511485 | 1.00E-50 | -1.95798 | 5.33E-146 | -25.7305 | 0.257389 | 142.4018 |
| CNAG_043 | URO1 | Urate oxic | 135.2794344 | 160.9870605 | 178.478237 | 1268.93995 | 1208.619201 | 1260.224472 | 1.00E-50 | 2.966697 | 6.06E-146 | 25.72548 | 7.817446 | 142.3831 |
| CNAG_013 | - | small subu | 23359.07088 | 24527.39358 | 23827.49279 | 10202.56027 | 10185.4709 | 9921.933479 | 1.00E-50 | -1.25758 | 3.32E-143 | -25.4796 | 0.418244 | 139.6795 |
| CNAG_012 | - | aldo-keto | 896.4937876 | 757.8479487 | 772.46406 | 3282.583683 | 3224.160979 | 3643.792238 | 1.00E-50 | 2.049623 | 7.30E-126 | 23.86367 | 4.139977 | 122.3693 |
| CNAG_012 | - | large subu | 38964.88781 | 41061.65573 | 41149.84888 | 15591.36273 | 15064.04857 | 16857.09555 | 1.00E-50 | -1.36559 | 7.07E-123 | -23.5743 | 0.388076 | 119.4415 |
| CNAG_068 | RPL22alph | large subu | 41681.17216 | 43064.67178 | 44932.57689 | 19006.59508 | 19582.24223 | 19042.99342 | 1.00E-50 | -1.18526 | 4.14E-117 | -23.0052 | 0.439744 | 113.7255 |
| CNAG_032 | - | large subu | 48568.51403 | 48515.41322 | 50756.48573 | 21236.3777 | 22640.89556 | 21286.63808 | 1.00E-50 | -1.19736 | 4.39E-115 | -22.8019 | 0.436072 | 111.7235 |
| CNAG_019 | - | small subu | 58410.15302 | 53033.11936 | 57647.89672 | 23487.17582 | 23751.82648 | 23172.81838 | 1.00E-50 | -1.27975 | 1.62E-111 | -22.4395 | 0.411867 | 108.1782 |
| CNAG_039 | - | hypothetic | 885.9062025 | 823.5233614 | 805.4900282 | 2829.656316 | 2638.384745 | 2749.254172 | 1.00E-50 | 1.693665 | 3.23E-109 | 22.20272 | 3.234773 | 105.8994 |
| CNAG_062 | - | large subu | 39244.07992 | 39954.14452 | 42149.14032 | 18294.48096 | 18736.07942 | 18450.15304 | 1.00E-50 | -1.14439 | 1.46E-108 | -22.1349 | 0.452382 | 105.2649 |
| CNAG_058 | - | small subu | 47368.08381 | 47359.33595 | 51714.12206 | 21259.74634 | 21190.89638 | 20432.93646 | 1.00E-50 | -1.23505 | 1.59E-108 | -22.1311 | 0.424827 | 105.2482 |
| CNAG_000 | RPL9B | large subu | 63913.96002 | 62320.58899 | 66586.92903 | 29202.21984 | 30246.96544 | 29744.92528 | 1.00E-50 | -1.12775 | 1.10E-104 | -21.7286 | 0.457628 | 101.443 |
| CNAG_013 | - | hypothetic | 362.8377576 | 327.942914 | 329.8795552 | 1374.953353 | 1501.863984 | 1402.575 | 1.00E-50 | 2.055248 | 1.60E-101 | 21.39162 | 4.15615 | 98.3143 |
| CNAG_047 | - | Phosphati | 2108.494926 | 2186.422928 | 2190.762898 | 5341.447819 | 5711.667326 | 5976.502942 | 1.00E-50 | 1.377873 | 4.61E-100 | 21.2343 | 2.598849 | 96.88592 |
| CNAG_045 | - | hypothetic | 71.03088183 | 82.32688598 | 62.2397948 | 751.7349946 | 660.883322 | 660.8374927 | 1.00E-50 | 3.253197 | 5.89E-99 | 21.11423 | 9.534762 | 95.8083 |
| CNAG_006 | - | large subu | 59442.6778 | 59921.1282 | 65759.31483 | 26756.019 | 27930.97194 | 26847.43076 | 1.00E-50 | -1.19845 | 7.33E-99 | -21.1039 | 0.435743 | 95.74017 |
| CNAG_037 | - | large subu | 56265.24962 | 56311.74948 | 57218.18768 | 27901.76113 | 28956.61768 | 28409.36895 | 1.00E-50 | -1.00906 | 2.68E-98 | -21.0425 | 0.49687 | 95.19007 |
| CNAG_007 | - | large subu | 38449.91881 | 36812.38008 | 38505.10394 | 17870.35094 | 18271.18228 | 17185.68765 | 1.00E-50 | -1.10875 | 1.66E-97 | -20.9557 | 0.463696 | 94.40945 |
| CNAG_018 | - | large subu | 96667.16866 | 97286.51215 | 99638.50253 | 49777.97473 | 50487.00561 | 50016.4258 | 1.00E-50 | -0.98144 | 2.57E-96 | -20.825 | 0.506475 | 93.23288 |
| CNAG_046 | - | hypothetic | 354.7303818 | 435.6640196 | 456.7606565 | 1645.11589 | 1650.07109 | 1651.234169 | 1.00E-50 | 1.975409 | 1.70E-92 | 20.39929 | 3.932397 | 89.43699 |
| CNAG_002 | - | isoleucine | 36468.24373 | 34696.30988 | 36255.9612 | 16438.26147 | 17387.92224 | 17253.37474 | 1.00E-50 | -1.08803 | 8.57E-92 | -20.3199 | 0.470402 | 88.74469 |
| CNAG_023 | - | small subu | 21887.72294 | 21280.11178 | 23392.3392 | 9821.997107 | 10300.23901 | 9669.910559 | 1.00E-50 | -1.17541 | 7.37E-91 | -20.214 | 0.442757 | 87.82127 |
| CNAG_027 | - | small subu | 49202.61797 | 47291.49984 | 52282.56903 | 22817.14377 | 23374.94461 | 22074.843 | 1.00E-50 | -1.13956 | 7.12E-89 | -19.9872 | 0.453898 | 85.8574 |
| CNAG_021 | - | large subu | 62245.25923 | 59665.82177 | 62868.24454 | 30567.41804 | 30196.08766 | 31001.59092 | 1.00E-50 | -1.02526 | 1.05E-88 | -19.9679 | 0.491323 | 85.6999 |
| CNAG_017 | - | hypothetic | 1901.880996 | 2239.217434 | 1717.319547 | 6344.417286 | 6018.801484 | 5881.10369 | 1.00E-50 | 1.624849 | 1.72E-88 | 19.94306 | 3.0841 | 85.49418 |
| CNAG_047 | PMT2 | dolichyl-p | 2798.570118 | 2931.969774 | 2939.381656 | 6308.783246 | 6710.448263 | 6781.486154 | 1.00E-50 | 1.176567 | 5.28E-86 | 19.65459 | 2.260383 | 83.02676 |
| CNAG_047 | - | ubiquitin-| | 2649.092724 | 2663.702151 | 2434.987681 | 5989.713801 | 6468.121335 | 6574.942913 | 1.00E-50 | 1.281703 | 1.25E-85 | 19.61072 | 2.431257 | 82.67021 |
| CNAG_025 | - | hypothetic | 338.3341734 | 386.088701 | 389.5694924 | 1402.352579 | 1329.101338 | 1343.778214 | 1.00E-50 | 1.857149 | 4.09E-85 | 19.55038 | 3.62291 | 82.17349 |
| CNAG_029 | - | large subu | 97980.83503 | 96311.43664 | 108877.0864 | 44687.06205 | 46863.33684 | 45036.29986 | 1.00E-50 | -1.16588 | 7.63E-85 | -19.5186 | 0.445691 | 81.92007 |
| CNAG_039 | - | hypothetic | 39.48346716 | 30.60598741 | 38.76934616 | 478.1797353 | 471.2111616 | 502.9765107 | 1.00E-50 | 3.745857 | 1.91E-84 | 19.47159 | 13.41576 | 81.52942 |
| CNAG_047 | - | DNA clam | 432.8375684 | 424.8682121 | 402.372455 | 1420.07415 | 1391.175961 | 1433.689439 | 1.00E-50 | 1.739112 | 4.00E-84 | 19.43372 | 3.338296 | 81.22474 |
| CNAG_047 | MEP1 | extracellul | 791.3028556 | 778.2500553 | 646.6032939 | 2308.726221 | 2462.578625 | 2374.039697 | 1.00E-50 | 1.674254 | 8.87E-84 | 19.39286 | 3.191543 | 80.88719 |
| CNAG_012 | - | homoisoci | 11368.65144 | 11522.85504 | 11809.63574 | 5079.853644 | 5476.803644 | 5587.796292 | 1.00E-50 | -1.11931 | 4.28E-83 | -19.3118 | 0.460313 | 80.21144 |
| CNAG_066 | - | small subu | 46748.11729 | 47765.55485 | 49862.83143 | 23021.07063 | 24639.5027 | 24170.77181 | 1.00E-50 | -1.02245 | 9.59E-83 | -19.27 | 0.492281 | 79.88288 |
| CNAG_054 | - | Dihydroxy | 21456.91824 | 22419.99822 | 21339.52808 | 10624.48203 | 11247.38365 | 11163.00458 | 1.00E-50 | -0.99627 | 9.54E-83 | -19.2703 | 0.501295 | 79.88288 |
| CNAG_026 | ISP4 | Identified | 229.7664583 | 243.9006142 | 225.3757555 | 959.874231 | 1115.417395 | 1179.190549 | 1.00E-50 | 2.207563 | 1.98E-82 | 19.23256 | 4.618943 | 79.58327 |
| CNAG_059 | - | cytochrom | 13804.60551 | 14109.77155 | 14867.13009 | 6166.33604 | 6854.328893 | 6534.791586 | 1.00E-50 | -1.14482 | 3.91E-82 | -19.1972 | 0.452246 | 79.30132 |
| CNAG_048 | - | hypothetic | 148.0672138 | 176.0324294 | 180.5803878 | 915.4545486 | 809.8542353 | 895.0847466 | 1.00E-50 | 2.362848 | 4.81E-82 | 19.18635 | 5.143848 | 79.21728 |
| CNAG_045 | - | beta-flank | 4791.891291 | 5235.486598 | 4975.231585 | 10652.50886 | 10371.03112 | 10784.78254 | 1.00E-50 | 1.06951 | 5.68E-82 | 19.17772 | 2.098721 | 79.15186 |
| CNAG_040 | - | large subu | 40417.61028 | 38361.75004 | 43740.25425 | 18141.92136 | 18931.88479 | 18097.13524 | 1.00E-50 | -1.1668 | 6.56E-82 | -19.1703 | 0.44541 | 79.09625 |
| CNAG_047 | - | endosome | 700.2011313 | 728.6636448 | 697.7457836 | 2023.147187 | 1964.698068 | 1964.467439 | 1.00E-50 | 1.470615 | 8.97E-82 | 19.15397 | 2.771399 | 78.96659 |
| CNAG_041 | - | tartrate tra | 111.8858134 | 115.7220289 | 109.1563338 | 704.7923032 | 746.1425267 | 947.9298455 | 1.00E-50 | 2.824204 | 2.32E-81 | 19.10442 | 7.082234 | 78.56642 |
| CNAG_062 | - | protein TII | 6517.522455 | 6101.179965 | 5795.950306 | 2661.729288 | 2571.637405 | 2460.555198 | 1.00E-50 | -1.27479 | 8.97E-81 | -19.0337 | 0.413285 | 77.98518 |
| CNAG_023 | - | MFS trans | 1814.605828 | 1483.31478 | 1709.103989 | 445.6405975 | 465.1187033 | 462.3604204 | 1.00E-50 | -1.88358 | 2.02E-80 | -18.9912 | 0.27101 | 77.6398 |
| CNAG_002 | RPL30 | large subu | 33449.68049 | 32314.12347 | 35870.92521 | 16361.27062 | 16203.61489 | 15692.51109 | 1.00E-50 | -1.09017 | 3.73E-80 | -18.9589 | 0.469706 | 77.3787 |
| CNAG_031 | - | small subu | 58472.34555 | 58050.75461 | 60833.46058 | 30653.84179 | 31698.72655 | 30357.69581 | 1.00E-50 | -0.9513 | 1.54E-79 | -18.8842 | 0.517166 | 76.76909 |
| CNAG_046 | - | endopepti | 579.9797921 | 473.3930639 | 436.5341189 | 2097.578889 | 2011.539843 | 1792.398288 | 1.00E-50 | 1.971367 | 1.24E-78 | 18.77352 | 3.921394 | 75.87853 |
| CNAG_047 | PCD102 | hypothetic | 387.3335509 | 409.7845341 | 427.9477558 | 1346.841161 | 1426.679544 | 1324.977984 | 1.00E-50 | 1.728565 | 1.50E-78 | 18.76353 | 3.313981 | 75.80253 |
| CNAG_056 | - | hypothetic | 18062.14875 | 16977.95308 | 19418.71968 | 8356.519443 | 8207.64623 | 7908.085794 | 1.00E-50 | -1.16977 | 1.26E-76 | -18.5264 | 0.444491 | 73.88786 |
| CNAG_011 | - | small subu | 73079.72862 | 71497.11395 | 77597.00866 | 35950.04296 | 38023.86287 | 37359.48362 | 1.00E-50 | -1.01233 | 1.57E-76 | -18.5147 | 0.495746 | 73.79905 |
| CNAG_030 | - | small subu | 53997.46335 | 51819.9741 | 57361.04382 | 26912.56634 | 27249.59767 | 26265.66775 | 1.00E-50 | -1.03633 | 6.06E-76 | -18.4419 | 0.487567 | 73.22346 |
| CNAG_007 | - | large subu | 31881.48219 | 32881.90137 | 35329.17342 | 15954.98373 | 16843.67196 | 15654.76185 | 1.00E-50 | -1.06207 | 9.00E-76 | -18.4205 | 0.478944 | 73.05689 |
| CNAG_001 | - | NADH del | 8529.89453 | 8578.263743 | 9027.260473 | 4347.612174 | 4157.94211 | 4277.805912 | 1.00E-50 | -1.04717 | 6.44E-75 | -18.3136 | 0.483916 | 72.22204 |
| CNAG_040 | - | large subu | 33261.72038 | 32696.61468 | 36670.75699 | 16284.97544 | 16573.65978 | 16560.6339 | 1.00E-50 | -1.06983 | 3.33E-74 | -18.2241 | 0.476376 | 71.52872 |
| CNAG_006 | - | cytoplasm | 1790.783609 | 1890.081656 | 1659.669735 | 4135.156243 | 4141.703332 | 4262.530646 | 1.00E-50 | 1.216813 | 1.59E-73 | 18.13824 | 2.324327 | 70.85376 |
| CNAG_048 | BNI1 | Formin BN | 589.3185144 | 615.5588338 | 608.1772173 | 1978.220527 | 1692.843979 | 1979.849112 | 1.00E-50 | 1.625582 | 2.45E-72 | 17.98729 | 3.085667 | 69.67978 |
| CNAG_047 | - | hypothetic | 3488.775087 | 3561.230405 | 2781.638831 | 10274.04514 | 9772.696679 | 8785.405652 | 1.00E-50 | 1.537392 | 7.98E-72 | 17.92176 | 2.902694 | 69.1717 |
| CNAG_022 | NOP56 | nucleolar | 11926.83144 | 11392.54882 | 11781.95524 | 6019.394786 | 6128.951573 | 5973.009232 | 1.00E-50 | -0.96937 | 1.36E-71 | -17.8921 | 0.510729 | 68.94478 |
| CNAG_008 | - | large subu | 17200.45804 | 16960.71121 | 18894.05004 | 8700.576669 | 8512.159297 | 8366.548914 | 1.00E-50 | -1.06803 | 1.63E-71 | -17.8819 | 0.476969 | 68.86964 |

| | | | | | | | | | | | | | | |
|---|---|---|---|---|---|---|---|---|---|---|---|---|---|---|
| CNAG_028 | - | GTP-bindi | 22584.93782 | 22000.94174 | 22049.79195 | 11539.02686 | 11843.42353 | 12153.23784 | 1.00E-50 | -0.92234 | 9.30E-71 | -17.7846 | 0.527653 | 68.12249 |
| CNAG_054 | - | asparagin | 14979.40966 | 14745.51932 | 15431.34799 | 7942.573571 | 8093.881341 | 7754.816477 | 1.00E-50 | -0.94004 | 2.02E-70 | -17.741 | 0.52122 | 67.78915 |
| CNAG_023 | - | large subu | 46479.58602 | 47370.16445 | 49696.48457 | 25677.4916 | 26004.96766 | 24602.62378 | 1.00E-50 | -0.9274 | 2.80E-70 | -17.7228 | 0.525806 | 67.65318 |
| CNAG_006 | - | small subu | 80042.99924 | 75567.67574 | 86414.41254 | 39807.28065 | 39348.72383 | 38776.07296 | 1.00E-50 | -1.05289 | 7.10E-69 | -17.54 | 0.482 | 66.2691 |
| CNAG_031 | URA5 | Orotate pl | 3462.879507 | 3307.459077 | 3508.351519 | 1555.806614 | 1534.006692 | 1549.061302 | 1.00E-50 | -1.16333 | 1.44E-68 | -17.4996 | 0.446482 | 65.96888 |
| CNAG_006 | - | small subu | 154495.4057 | 147406.5627 | 162756.5164 | 78240.03759 | 81310.63675 | 80251.65443 | 1.00E-50 | -0.96995 | 1.86E-68 | -17.4852 | 0.510523 | 65.86282 |
| CNAG_035 | - | 3-beta hy | 239.1217192 | 296.6746295 | 311.7223567 | 1048.639242 | 1239.702301 | 1137.239508 | 1.00E-50 | 2.002075 | 2.00E-68 | 17.48098 | 4.005759 | 65.83493 |
| CNAG_003 | UBI1 | Predicted | 25449.41757 | 24430.56034 | 27243.32238 | 13002.22103 | 12009.8221 | 12227.6863 | 1.00E-50 | -1.06588 | 7.84E-68 | -17.4029 | 0.477681 | 65.24551 |
| CNAG_046 | - | xylitol deh | 523.9211428 | 534.7560139 | 566.5731939 | 1501.34943 | 1722.472087 | 1766.686483 | 1.00E-50 | 1.60484 | 3.28E-67 | 17.32069 | 3.04162 | 64.62731 |
| CNAG_000 | - | kynurenin | 6454.767849 | 6274.736098 | 6860.209644 | 3145.894021 | 3211.738302 | 3116.591856 | 1.00E-50 | -1.06376 | 3.69E-67 | -17.314 | 0.478385 | 64.58056 |
| CNAG_055 | PLR1 | pyridoxal | 805.302745 | 791.1736418 | 840.6400724 | 2078.007281 | 2314.080969 | 2499.382177 | 1.00E-50 | 1.4849 | 1.79E-66 | 17.22267 | 2.798978 | 63.91189 |
| CNAG_078 | - | ran-bindir | 797.124149 | 667.3071915 | 805.4538666 | 2491.995441 | 2232.957958 | 2177.658928 | 1.00E-50 | 1.588531 | 6.23E-66 | 17.15048 | 3.00743 | 63.38193 |
| CNAG_046 | - | 26S prote | 5143.28365 | 4889.642727 | 5131.973815 | 9737.860269 | 10053.62744 | 9994.405822 | 1.00E-50 | 0.958456 | 1.05E-65 | 17.12005 | 1.943229 | 63.15823 |
| CNAG_046 | - | hypothetic | 177.164649 | 128.641673 | 145.3933573 | 705.6173219 | 763.4799102 | 784.0480204 | 1.00E-50 | 2.307106 | 3.27E-65 | 17.05395 | 4.948893 | 62.67302 |
| CNAG_048 | - | hypothetic | 416.4491225 | 411.9190665 | 356.5169856 | 1208.172858 | 1281.684339 | 1277.176666 | 1.00E-50 | 1.655429 | 4.11E-65 | 17.04051 | 3.150168 | 62.57998 |
| CNAG_009 | - | hypothetic | 66.3678462 | 131.8588762 | 88.89586282 | 681.7437303 | 678.0369707 | 797.1079597 | 1.00E-50 | 2.906404 | 6.02E-65 | 17.0182 | 7.497471 | 62.41784 |
| CNAG_064 | RPL17 | Large sub | 58598.49436 | 58225.31221 | 64083.97406 | 30417.99014 | 32189.52349 | 30830.62458 | 1.00E-50 | -0.96869 | 9.24E-65 | -16.9931 | 0.510969 | 62.23491 |
| CNAG_122 | - | unspecifie | 33765.15738 | 35999.99855 | 38530.67771 | 18724.21809 | 18479.94635 | 18278.03809 | 1.00E-50 | -0.97999 | 4.81E-64 | -16.8961 | 0.506983 | 61.52203 |
| CNAG_054 | - | nascent p | 12748.86448 | 11591.89842 | 12597.76662 | 6253.656782 | 6103.901896 | 6141.761781 | 1.00E-50 | -1.01347 | 1.26E-63 | -16.8394 | 0.495354 | 61.11177 |
| CNAG_005 | - | translatior | 9616.981641 | 8932.784733 | 9370.703337 | 4826.287885 | 4799.733567 | 4740.730951 | 1.00E-50 | -0.97426 | 1.79E-63 | -16.8183 | 0.508999 | 60.96358 |
| CNAG_022 | CHS7 | Chitin syn | 193.6055046 | 274.0438644 | 172.0756052 | 990.4100027 | 1050.622835 | 1195.767347 | 1.00E-50 | 2.326405 | 2.79E-63 | 16.79196 | 5.015539 | 60.77719 |
| CNAG_011 | - | H/ACA rib | 15764.08304 | 14952.4008 | 16226.89993 | 7761.167611 | 7879.059696 | 8295.439079 | 1.00E-50 | -0.98727 | 4.32E-63 | -16.7661 | 0.50443 | 60.59152 |
| CNAG_030 | CQS1 | quorum se | 8165.62752 | 9389.022507 | 9270.92039 | 23695.58284 | 20955.59668 | 19529.63359 | 1.00E-50 | 1.243869 | 4.60E-63 | 16.76238 | 2.368328 | 60.56714 |
| CNAG_044 | - | large subu | 93341.42958 | 90656.03984 | 103221.7227 | 49132.78987 | 48502.24217 | 47223.27957 | 1.00E-50 | -1.00311 | 1.50E-62 | -16.6919 | 0.498925 | 60.0562 |
| CNAG_033 | - | Succinyl-C | 12189.64105 | 11856.93174 | 13253.59228 | 6052.028906 | 6378.224302 | 6163.96783 | 1.00E-50 | -1.01992 | 2.97E-62 | -16.6511 | 0.493142 | 59.76319 |
| CNAG_006 | LPI15 | transmem | 234.3739213 | 260.019813 | 189.1180691 | 890.0772601 | 937.6539576 | 906.3379362 | 1.00E-50 | 1.985831 | 5.61E-62 | 16.61299 | 3.960908 | 59.49297 |
| CNAG_037 | - | 3-phosph | 8996.889947 | 8384.361703 | 8676.446202 | 4495.517025 | 4270.949856 | 4493.152765 | 1.00E-50 | -0.99025 | 1.56E-61 | -16.5514 | 0.50339 | 59.05402 |
| CNAG_126 | - | unspecifie | 896.9271024 | 831.381834 | 900.722157 | 226.9469122 | 258.1415529 | 200.3998219 | 1.00E-50 | -1.95723 | 2.87E-61 | -16.5149 | 0.257523 | 58.79355 |
| CNAG_011 | - | POT famil | 12554.09905 | 13852.2548 | 13170.49558 | 6799.230301 | 7079.504927 | 7191.93005 | 1.00E-50 | -0.92428 | 7.22E-61 | -16.4591 | 0.526943 | 58.39558 |
| CNAG_030 | - | hypothetic | 132.8134752 | 124.310286 | 137.9079325 | 582.6567452 | 631.5176432 | 610.9774547 | 1.00E-50 | 2.198228 | 1.19E-60 | 16.42855 | 4.589153 | 58.18265 |
| CNAG_067 | - | threonine | 7797.735107 | 7928.569746 | 7558.846368 | 4246.509687 | 4148.514747 | 4215.594467 | 1.00E-50 | -0.89992 | 1.34E-60 | -16.4217 | 0.535918 | 58.13655 |
| CNAG_066 | - | small subu | 66710.8299 | 63584.48711 | 69227.51461 | 36874.01263 | 35895.94486 | 34913.7897 | 1.00E-50 | -0.90536 | 2.20E-60 | -16.3914 | 0.533899 | 57.92251 |
| CNAG_003 | - | 3-hydroxy | 221021.1963 | 236798.9054 | 244291.1454 | 125617.763 | 112021.1921 | 121270.9921 | 1.00E-50 | -0.98294 | 5.05E-60 | -16.3409 | 0.505947 | 57.56795 |
| CNAG_014 | - | large subu | 14724.67054 | 13936.3356 | 14650.6705 | 7119.890567 | 6888.823873 | 5950.903666 | 1.00E-50 | -1.13353 | 5.31E-60 | -16.3379 | 0.455799 | 57.54926 |
| CNAG_040 | - | large subu | 9432.712071 | 10752.45122 | 11014.02702 | 4967.87135 | 4653.015707 | 5043.816874 | 1.00E-50 | -1.10391 | 7.46E-60 | -16.3171 | 0.465254 | 57.40415 |
| CNAG_054 | GIB2 | G protein | 27041.10371 | 25620.07682 | 28602.01345 | 14065.36716 | 14491.49566 | 13734.1514 | 1.00E-50 | -0.95799 | 1.28E-59 | -16.2843 | 0.514772 | 57.17421 |
| CNAG_131 | - | unspecifie | 35000.57207 | 35015.26192 | 37963.34586 | 19615.41463 | 18141.80332 | 19044.02681 | 1.00E-50 | -0.94204 | 3.07E-59 | -16.2304 | 0.520495 | 56.79513 |
| CNAG_078 | - | ferro-O2-c | 1912.161029 | 1863.145575 | 1915.567831 | 3796.37364 | 4104.625628 | 4022.808209 | 1.00E-50 | 1.051704 | 3.50E-59 | 16.22249 | 2.072977 | 56.74173 |
| CNAG_049 | - | 26S prote | 3866.923093 | 3476.071027 | 3585.640751 | 7400.709988 | 7828.305987 | 7398.862953 | 1.00E-50 | 1.034286 | 5.16E-59 | 16.19854 | 2.048099 | 56.57522 |
| CNAG_030 | - | hypothetic | 2783.221846 | 2674.98783 | 2774.637732 | 1231.255691 | 1130.279928 | 1268.173229 | 1.00E-50 | -1.19678 | 9.09E-59 | -16.1637 | 0.436246 | 56.3323 |
| CNAG_022 | - | large subu | 39761.43882 | 38227.1032 | 42630.13196 | 19710.2405 | 21618.84575 | 20588.27656 | 1.00E-50 | -0.97774 | 1.80E-58 | -16.1214 | 0.507775 | 56.03729 |
| CNAG_042 | - | H/ACA rib | 2584.86877 | 2752.521603 | 2884.434318 | 1124.542393 | 1163.942696 | 1267.059351 | 1.00E-50 | -1.22468 | 3.41E-57 | -15.9387 | 0.427892 | 54.77116 |
| CNAG_030 | - | large subu | 20111.74232 | 20172.55488 | 22318.45623 | 11025.38958 | 11124.91785 | 10338.18226 | 1.00E-50 | -0.96181 | 2.76E-56 | -15.8075 | 0.513412 | 53.87059 |
| CNAG_049 | - | 3-deoxy-7 | 9888.197721 | 11954.8293 | 11641.12321 | 5332.883119 | 5306.01314 | 5367.992308 | 1.00E-50 | -1.07934 | 1.34E-55 | -15.7077 | 0.473244 | 53.18804 |
| CNAG_049 | CHC1 | clathrin he | 13445.94483 | 12911.09826 | 11899.711 | 23870.80799 | 24546.35415 | 24460.92953 | 1.00E-50 | 0.91446 | 3.70E-55 | 15.64318 | 1.884864 | 52.75432 |
| CNAG_036 | - | Threonine | 3119.788162 | 3417.30931 | 3394.264945 | 1629.031084 | 1641.882987 | 1575.690141 | 1.00E-50 | -1.05024 | 4.85E-55 | -15.626 | 0.482887 | 52.6395 |
| CNAG_069 | TIF3 | Translatior | 5034.884722 | 5125.113542 | 5628.449332 | 2636.927159 | 2664.026073 | 2597.022909 | 1.00E-50 | -1.0147 | 9.87E-55 | -15.5806 | 0.49493 | 52.33306 |
| CNAG_042 | - | hypothetic | 2735.582643 | 3167.269637 | 3277.912079 | 1221.043464 | 1131.958943 | 1335.910069 | 1.00E-50 | -1.3304 | 4.92E-54 | -15.4776 | 0.397657 | 51.63826 |
| CNAG_069 | - | protein tra | 8898.958361 | 8931.688882 | 9052.916311 | 4640.248738 | 4953.309948 | 5075.942208 | 1.00E-50 | -0.88917 | 1.02E-53 | -15.4303 | 0.539926 | 51.32452 |
| CNAG_078 | - | Histone H | 8952.698623 | 11298.19763 | 10990.01934 | 23772.27384 | 23278.48224 | 22388.24271 | 1.00E-50 | 1.138004 | 3.08E-53 | 15.35907 | 2.200763 | 50.85114 |
| CNAG_004 | - | acetyl-Co/ | 630.0837883 | 621.9899881 | 530.3335115 | 1602.679894 | 1565.230202 | 1796.666018 | 1.00E-50 | 1.463592 | 4.28E-53 | 15.33773 | 2.757942 | 50.71071 |
| CNAG_024 | - | ABC trans | 259.9899197 | 236.325828 | 249.85539 | 777.6957269 | 852.1474655 | 876.3338036 | 1.00E-50 | 1.735927 | 4.93E-53 | 15.32854 | 3.330935 | 50.65167 |
| CNAG_040 | - | small subu | 138002.0734 | 126901.9834 | 138708.0386 | 73940.05785 | 75585.85307 | 75830.95516 | 1.00E-50 | -0.8565 | 5.27E-53 | -15.3242 | 0.552291 | 50.62519 |
| CNAG_046 | - | hypothetic | 2699.203749 | 2589.33163 | 2319.796072 | 5239.332908 | 5530.395642 | 5870.015993 | 1.00E-50 | 1.11395 | 1.18E-52 | 15.27163 | 2.164375 | 50.27669 |
| CNAG_078 | - | hypothetic | 971.0468821 | 1081.998926 | 1069.877044 | 2247.685533 | 2323.415902 | 2364.182193 | 1.00E-50 | 1.136813 | 1.68E-52 | 15.24862 | 2.198948 | 50.12587 |
| CNAG_055 | HIS3 | Imidazole | 5513.455727 | 5021.706808 | 5181.691721 | 2683.640548 | 2685.59261 | 2715.785292 | 1.02E-50 | -0.97483 | 2.31E-52 | -15.2278 | 0.5088 | 49.98984 |
| CNAG_044 | - | small subu | 38579.79409 | 40130.89865 | 41804.73893 | 21033.38646 | 23438.90135 | 21692.86746 | 1.07E-50 | -0.8804 | 2.43E-52 | -15.2247 | 0.543216 | 49.9715 |
| CNAG_031 | - | small subu | 111752.2731 | 110867.601 | 129792.3853 | 60518.39933 | 59631.00413 | 59163.05046 | 1.21E-50 | -0.99031 | 2.75E-52 | -15.2164 | 0.503371 | 49.9188 |
| CNAG_130 | - | unspecifie | 843738.9183 | 803874.2815 | 795955.8408 | 449465.1039 | 487550.2364 | 465693.9712 | 1.95E-50 | -0.81633 | 4.50E-52 | -15.1842 | 0.567886 | 49.70981 |
| CNAG_011 | - | small subu | 49640.65533 | 47077.16153 | 52110.93776 | 28199.99059 | 27484.26417 | 26651.96198 | 3.24E-50 | -0.86972 | 7.53E-52 | -15.1504 | 0.547253 | 49.4888 |
| CNAG_028 | - | small subu | 12979.04495 | 13298.48128 | 13704.73346 | 7033.562967 | 6749.119334 | 5837.647089 | 4.07E-50 | -1.04243 | 9.49E-52 | -15.1352 | 0.485509 | 49.39041 |
| CNAG_064 | - | small subu | 1174.046787 | 1174.07255 | 1346.496453 | 384.2265482 | 424.6529958 | 457.8257114 | 4.44E-50 | -1.56142 | 1.04E-51 | -15.1291 | 0.338818 | 49.35227 |
| CNAG_038 | ERG6 | sterol 24-( | 12064.72352 | 11688.87062 | 13177.88346 | 6104.180688 | 6589.575808 | 6454.802971 | 4.70E-50 | -0.96324 | 1.11E-51 | -15.125 | 0.512903 | 49.32809 |
| CNAG_034 | - | hypothetic | 70.98152194 | 95.20938128 | 88.86231184 | 448.0997714 | 510.953151 | 443.4112643 | 5.90E-50 | 2.454512 | 1.40E-51 | 15.10974 | 5.481277 | 49.22938 |
| CNAG_066 | - | anthranila | 3227.12929 | 3391.452858 | 3131.988132 | 1605.672077 | 1685.03616 | 1648.929834 | 6.70E-50 | -0.99633 | 1.60E-51 | -15.101 | 0.501275 | 49.17391 |
| CNAG_033 | - | small subu | 18007.39082 | 17601.80007 | 19054.06362 | 10162.34309 | 9151.331917 | 8908.301617 | 7.18E-50 | -0.96922 | 1.72E-51 | -15.0961 | 0.510784 | 49.14401 |
| CNAG_015 | - | template-a | 5199.33542 | 4666.171383 | 5091.005861 | 2545.888545 | 2481.148381 | 2504.869784 | 2.28E-49 | -1.00577 | 5.48E-51 | -15.0194 | 0.498005 | 48.64262 |
| CNAG_000 | - | T-complex | 11598.81658 | 11609.11847 | 11882.21676 | 6128.347368 | 6527.487246 | 6782.246905 | 2.87E-49 | -0.86747 | 6.93E-51 | -15.0038 | 0.548107 | 48.54258 |

| | | | | | | | | | | | | | | |
|---|---|---|---|---|---|---|---|---|---|---|---|---|---|---|
| CNAG_038 | - | protein lys | 744.2366727 | 748.4440611 | 830.3316546 | 238.7099122 | 242.6279608 | 235.8547821 | 3.38E-49 | -1.71473 | 8.21E-51 | -14.9926 | 0.304659 | 48.47132 |
| CNAG_045 | DDT1 | hypothetic | 1392.491382 | 1412.752701 | 1384.468931 | 2783.900377 | 2808.971645 | 2789.466904 | 4.00E-49 | 0.985511 | 9.76E-51 | 14.98106 | 1.980015 | 48.39816 |
| CNAG_047 | - | DNA prim | 606.774666 | 733.9944498 | 624.1498285 | 1704.748918 | 1594.534441 | 1686.931921 | 2.27E-48 | 1.329904 | 5.61E-50 | 14.86443 | 2.513859 | 47.6432 |
| CNAG_054 | - | 2-isopropy | 9872.989895 | 11886.94041 | 10678.1869 | 5255.832024 | 5667.488578 | 5430.145637 | 3.37E-48 | -1.00255 | 8.39E-50 | -14.8375 | 0.499115 | 47.47287 |
| CNAG_049 | - | Nascent p | 8722.771596 | 9820.515294 | 10622.61982 | 4735.1697 | 4926.528977 | 4633.075779 | 9.54E-48 | -1.04378 | 2.41E-49 | -14.7665 | 0.485055 | 47.02065 |
| CNAG_075 | - | calcium/pr | 1119.304913 | 851.5328035 | 780.9708922 | 2676.057689 | 2784.240567 | 2710.457071 | 1.19E-47 | 1.554626 | 3.01E-49 | 14.75143 | 2.937575 | 46.92593 |
| CNAG_017 | - | hypothetic | 2796.295484 | 2930.347818 | 3052.985215 | 1478.07285 | 1377.036581 | 1461.31595 | 1.70E-47 | -1.03923 | 4.37E-49 | -14.7264 | 0.486586 | 46.76893 |
| CNAG_041 | - | small subu | 40257.86047 | 41000.41211 | 45941.46266 | 21870.81851 | 22924.75528 | 23385.843 | 2.60E-47 | -0.91501 | 6.76E-49 | -14.6968 | 0.530339 | 46.58513 |
| CNAG_058 | - | large subu | 15891.41574 | 15160.35994 | 17006.45819 | 8826.779434 | 8206.84287 | 7797.033127 | 3.57E-47 | -0.96836 | 9.33E-49 | -14.6749 | 0.511088 | 46.44711 |
| CNAG_010 | - | hypothetic | 1946.036376 | 1942.332784 | 2246.624087 | 794.5190358 | 771.421749 | 897.3916042 | 4.66E-47 | -1.33226 | 1.23E-48 | -14.6562 | 0.397145 | 46.33116 |
| CNAG_064 | - | hypothetic | 2515.970853 | 2644.793248 | 2543.269096 | 1259.340994 | 1320.177225 | 1176.052724 | 7.79E-47 | -1.05158 | 2.06E-48 | -14.621 | 0.482441 | 46.10841 |
| CNAG_047 | - | hypothetic | 820.3526356 | 874.044879 | 804.3463107 | 1830.614843 | 1812.75031 | 1812.485151 | 6.28E-46 | 1.111604 | 1.69E-47 | 14.47698 | 2.160858 | 45.20231 |
| CNAG_126 | - | unspecifie | 1638.977883 | 1765.574157 | 1876.602818 | 742.4540239 | 641.1560906 | 756.4132265 | 7.71E-46 | -1.31816 | 2.09E-47 | -14.4626 | 0.401047 | 45.11322 |
| CNAG_052 | XKS1 | xylulokina | 885.7037464 | 784.6695065 | 802.2226459 | 1844.562287 | 1893.002904 | 1958.940659 | 7.91E-46 | 1.188977 | 2.16E-47 | 14.4604 | 2.279911 | 45.10156 |
| CNAG_008 | - | small subu | 18734.88063 | 17305.55558 | 20873.37063 | 9636.882766 | 9713.743906 | 9253.557708 | 9.21E-46 | -1.00852 | 2.52E-47 | -14.4496 | 0.497058 | 45.03553 |
| CNAG_062 | - | large subu | 41267.99831 | 42471.11183 | 46919.41926 | 24511.65667 | 24650.76832 | 23442.48622 | 1.41E-45 | -0.86304 | 3.87E-47 | -14.42 | 0.549794 | 44.85104 |
| CNAG_051 | - | ATP-bindi | 3537.761961 | 3519.733939 | 3448.724838 | 1928.774162 | 1896.381021 | 1847.638006 | 2.42E-45 | -0.90453 | 6.68E-47 | -14.3824 | 0.534207 | 44.61628 |
| CNAG_044 | HPP3 | Phosphog | 2064.075549 | 2273.036418 | 2090.034772 | 1058.56101 | 1019.089881 | 991.7555009 | 3.15E-45 | -1.08164 | 8.73E-47 | -14.3638 | 0.472491 | 44.5017 |
| CNAG_063 | - | replicatior | 889.2155047 | 858.9841964 | 832.0761752 | 1885.060599 | 1831.668133 | 1982.256615 | 6.87E-45 | 1.128436 | 1.93E-46 | 14.30877 | 2.186216 | 44.16315 |
| CNAG_046 | YPK1 | AGC/AKT | 1538.437788 | 1388.010127 | 1516.70345 | 3106.223476 | 3018.567412 | 2989.305443 | 9.79E-45 | 1.021023 | 2.76E-46 | 14.28377 | 2.029357 | 44.0091 |
| CNAG_020 | PUT5 | Proline de | 2579.757 | 2298.934996 | 2114.593941 | 1021.107693 | 917.2095871 | 1005.095045 | 4.66E-44 | -1.26369 | 1.33E-45 | -14.174 | 0.416478 | 43.33146 |
| CNAG_038 | - | hypothetic | 226.1659131 | 218.0140064 | 256.2362174 | 803.6930758 | 710.4100407 | 862.9338437 | 5.81E-44 | 1.747311 | 1.67E-45 | 14.15803 | 3.357321 | 43.23615 |
| CNAG_007 | - | RuvB-like | 4967.10269 | 4709.275272 | 5073.961023 | 2668.899556 | 2697.705706 | 2726.856868 | 8.64E-44 | -0.88177 | 2.50E-45 | -14.1294 | 0.542701 | 43.06338 |
| CNAG_059 | - | ATP synth | 41182.69791 | 43962.21216 | 44771.60209 | 22306.67 | 20826.06002 | 24825.68816 | 1.16E-43 | -0.94961 | 3.40E-45 | -14.1078 | 0.517771 | 42.93538 |
| CNAG_124 | - | unspecifie | 830.5626366 | 777.5815761 | 820.834195 | 291.7262294 | 281.474862 | 284.6623601 | 1.70E-43 | -1.51901 | 5.00E-45 | -14.0805 | 0.348926 | 42.76947 |
| CNAG_122 | - | unspecifie | 1973.927923 | 1801.229456 | 2337.177122 | 818.7142975 | 685.1247882 | 653.281784 | 2.04E-43 | -1.51897 | 6.04E-45 | -14.0672 | 0.348936 | 42.68952 |
| CNAG_050 | - | hypothetic | 32.43412052 | 38.1244306 | 13.1524087 | 331.2637547 | 276.829659 | 329.8282574 | 3.11E-43 | 3.481303 | 9.24E-45 | 14.03711 | 11.16803 | 42.50668 |
| CNAG_039 | NAM9 | NAM9 prc | 3707.04356 | 3471.294629 | 4103.383201 | 1721.571769 | 1778.137475 | 1874.291266 | 6.00E-43 | -1.08576 | 1.79E-44 | -13.99 | 0.471143 | 42.22208 |
| CNAG_043 | - | Aspartate | 3156.029747 | 3470.120535 | 3329.258927 | 1815.92261 | 1748.00351 | 1709.988603 | 7.12E-43 | -0.93192 | 2.14E-44 | -13.9775 | 0.524162 | 42.14756 |
| CNAG_046 | - | acyl-CoA | 255.3504018 | 278.3171324 | 292.4880294 | 766.6196482 | 873.6754666 | 921.8301573 | 8.61E-43 | 1.618662 | 2.62E-44 | 13.9631 | 3.070901 | 42.065 |
| CNAG_016 | FCY2 | cytosine p | 6266.732262 | 5953.673691 | 6436.850937 | 2776.111679 | 3126.355618 | 3339.664117 | 1.09E-42 | -1.02896 | 3.32E-44 | -13.9462 | 0.490063 | 41.96368 |
| CNAG_038 | - | cell growt | 2473.566681 | 2048.002434 | 2268.081538 | 1014.892357 | 1000.079596 | 975.1281825 | 1.69E-42 | -1.19966 | 5.17E-44 | -13.9145 | 0.435378 | 41.77292 |
| CNAG_055 | - | small subu | 40361.82215 | 40838.8189 | 46208.08761 | 23773.82613 | 24012.46981 | 23559.03481 | 1.82E-42 | -0.852 | 5.61E-44 | -13.9087 | 0.554018 | 41.73952 |
| CNAG_038 | - | hypothetic | 1461.414343 | 1434.878162 | 1595.08216 | 680.2526366 | 623.9714748 | 640.9677338 | 1.93E-42 | -1.22382 | 5.97E-44 | -13.9042 | 0.428148 | 41.71398 |
| CNAG_043 | - | chaperone | 2511.199241 | 2426.123151 | 2567.74554 | 1032.69067 | 1207.145886 | 1228.179592 | 2.33E-42 | -1.12946 | 7.22E-44 | -13.8906 | 0.457087 | 41.63288 |
| CNAG_047 | FZC26 | nuclear pr | 1289.761429 | 1436.429804 | 1305.557742 | 2660.137285 | 2760.715365 | 2616.318033 | 4.54E-42 | 0.980861 | 1.42E-43 | 13.84215 | 1.973643 | 41.34302 |
| CNAG_023 | - | small subu | 61477.05385 | 59450.40336 | 69941.98444 | 35203.01873 | 34487.38514 | 33717.03087 | 1.12E-41 | -0.89988 | 3.53E-43 | -13.7765 | 0.53593 | 40.95046 |
| CNAG_048 | - | hypothetic | 94.25654424 | 100.5781957 | 75.00667491 | 458.5390187 | 439.9803687 | 394.6706425 | 1.57E-41 | 2.247855 | 4.98E-43 | 13.75168 | 4.74976 | 40.80317 |
| CNAG_058 | PRM1 | plasma m | 140.9306578 | 152.2722424 | 127.2409325 | 509.3197503 | 588.4274013 | 516.7671738 | 1.80E-41 | 1.929653 | 5.72E-43 | 13.74154 | 3.809636 | 40.74401 |
| CNAG_002 | - | efflux prot | 86.19142502 | 118.9280782 | 160.2794389 | 560.7360856 | 608.1421319 | 678.4698756 | 2.40E-41 | 2.327696 | 7.68E-43 | 13.72021 | 5.020029 | 40.61946 |
| CNAG_127 | - | unspecifie | 862.4122226 | 938.0898454 | 1050.027127 | 342.3064139 | 329.7721127 | 352.3640132 | 4.71E-41 | -1.49318 | 1.51E-42 | -13.6711 | 0.35523 | 40.3273 |
| CNAG_017 | - | solute car | 5821.881094 | 5447.322923 | 5722.394375 | 3247.229684 | 3124.668802 | 3233.077884 | 5.13E-41 | -0.8385 | 1.65E-42 | -13.6646 | 0.559226 | 40.29007 |
| CNAG_020 | ERG20 | farnesyl d | 4766.285636 | 4662.92953 | 4663.430584 | 2702.418649 | 2758.989614 | 2710.190807 | 5.50E-41 | -0.80173 | 1.78E-42 | -13.6591 | 0.573661 | 40.25941 |
| CNAG_077 | GLF | UDP-galac | 4892.480631 | 5223.137882 | 4935.379701 | 2876.758068 | 2786.547233 | 2943.309079 | 6.84E-41 | -0.82136 | 2.22E-42 | -13.643 | 0.56591 | 40.16469 |
| CNAG_045 | - | hypothetic | 107.1716993 | 171.6543358 | 124.0538965 | 574.9101014 | 642.8378563 | 547.8589798 | 7.02E-41 | 2.118891 | 2.29E-42 | 13.64088 | 4.3436 | 40.15386 |
| CNAG_007 | - | acetyl-CoA | 9634.549056 | 9142.883753 | 8846.08118 | 4959.381447 | 5431.214949 | 5240.251919 | 7.17E-41 | -0.83701 | 2.35E-42 | -13.6388 | 0.559801 | 40.1447 |
| CNAG_057 | - | ketol-acid | 35210.79862 | 34675.90129 | 34627.58452 | 20548.62235 | 22396.98513 | 20600.43219 | 7.36E-41 | -0.73331 | 2.43E-42 | -13.6366 | 0.601521 | 40.13317 |
| CNAG_019 | UBI4 | Polyubiqu | 8131.472952 | 8175.745824 | 7372.596753 | 14248.07567 | 13642.61387 | 13722.41276 | 9.69E-41 | 0.798385 | 3.21E-42 | 13.61622 | 1.739153 | 40.01359 |
| CNAG_056 | - | chorismat | 7972.964248 | 7863.98695 | 8372.502821 | 4245.663962 | 4759.283195 | 4558.615621 | 1.01E-40 | -0.85135 | 3.37E-42 | -13.6127 | 0.554265 | 39.994 |
| CNAG_058 | TRR1 | Thioredox | 2141.106407 | 2156.757228 | 2116.702332 | 1040.616091 | 1123.519902 | 1103.829528 | 1.05E-40 | -0.98872 | 3.50E-42 | -13.6098 | 0.503925 | 39.97887 |
| CNAG_067 | - | succinate | 6523.738975 | 6192.908282 | 6880.511401 | 3740.681211 | 3480.887748 | 3485.122146 | 2.24E-40 | -0.88777 | 7.48E-42 | -13.5542 | 0.54045 | 39.65045 |
| CNAG_028 | - | NADH del | 4248.923024 | 3987.400649 | 4405.275422 | 2195.655679 | 2334.582284 | 2265.045785 | 2.54E-40 | -0.91135 | 8.55E-42 | -13.5444 | 0.531689 | 39.59575 |
| CNAG_016 | - | MFS trans | 46.41821543 | 37.05046651 | 39.81495113 | 252.2883868 | 344.6032788 | 330.9888955 | 7.69E-40 | 2.908314 | 2.60E-41 | 13.46248 | 7.507404 | 39.11422 |
| CNAG_075 | - | hypothetic | 2546.148319 | 2574.187763 | 2528.741549 | 4450.848135 | 4419.24939 | 4604.606286 | 9.00E-40 | 0.801791 | 3.07E-41 | 13.45027 | 1.743263 | 39.04561 |
| CNAG_020 | - | hypothetic | 2498.192995 | 2174.049393 | 2328.890203 | 1006.270679 | 1127.790217 | 1121.618039 | 9.53E-40 | -1.12106 | 3.26E-41 | -13.4458 | 0.459757 | 39.02091 |
| CNAG_028 | - | glycine cle | 639.3490775 | 649.9572336 | 631.5776848 | 1352.569132 | 1493.715435 | 1436.116526 | 1.10E-39 | 1.142355 | 3.77E-41 | 13.43511 | 2.20741 | 38.95968 |
| CNAG_026 | - | translatior | 10449.6554 | 9374.5944 | 10406.23465 | 5636.548127 | 5777.941102 | 5569.988141 | 1.13E-39 | -0.84781 | 3.89E-41 | -13.4327 | 0.555626 | 38.94719 |
| CNAG_011 | - | ATP-depe | 6513.079592 | 5680.073071 | 6330.24073 | 3277.503915 | 3312.699968 | 3323.028038 | 1.31E-39 | -0.91801 | 4.53E-41 | -13.4215 | 0.52924 | 38.88279 |
| CNAG_012 | - | cellular nu | 9640.483608 | 9105.219318 | 9971.118081 | 5254.375431 | 5426.864579 | 5670.967072 | 1.61E-39 | -0.82812 | 5.60E-41 | -13.4057 | 0.563262 | 38.7919 |
| CNAG_062 | - | hypothetic | 174.7459626 | 95.24654953 | 107.0086865 | 597.4987573 | 627.1600629 | 643.0903706 | 2.22E-39 | 2.294821 | 7.78E-41 | 13.38127 | 4.906932 | 38.65332 |
| CNAG_004 | OFD1 | putative o | 4609.728274 | 4342.936521 | 4473.57912 | 2361.436158 | 2603.736892 | 2432.678605 | 3.00E-39 | -0.87564 | 1.06E-40 | -13.3585 | 0.545011 | 38.52234 |
| CNAG_122 | - | unspecifie | 79237.31857 | 72380.69239 | 77543.77768 | 47066.60043 | 45353.97625 | 45098.35327 | 4.04E-39 | -0.75247 | 1.43E-40 | -13.3359 | 0.593588 | 38.39332 |
| CNAG_061 | - | NADH del | 2795.175183 | 2903.444828 | 3295.011721 | 1530.236143 | 1421.8978 | 1423.598527 | 4.55E-39 | -1.0551 | 1.62E-40 | -13.3267 | 0.481264 | 38.34151 |
| CNAG_035 | - | large subu | 8583.825937 | 9100.864843 | 9980.67161 | 5189.042826 | 5178.44494 | 5077.077716 | 7.01E-39 | -0.85609 | 2.51E-40 | -13.2939 | 0.552448 | 38.15398 |
| CNAG_041 | - | succinate | 10098.28209 | 9882.029959 | 9738.710127 | 5970.492051 | 6125.590899 | 6222.695144 | 2.08E-38 | -0.71347 | 7.48E-40 | -13.212 | 0.609854 | 37.68132 |
| CNAG_022 | - | solute car | 154.9074578 | 155.5027413 | 157.0692737 | 516.3317366 | 514.7631224 | 561.0425916 | 3.13E-38 | 1.756427 | 1.13E-39 | 13.18089 | 3.378603 | 37.50482 |
| CNAG_015 | - | glutamate | 26243.4675 | 23208.82642 | 24396.01972 | 14281.82817 | 14531.22428 | 14235.88358 | 3.15E-38 | -0.7945 | 1.14E-39 | -13.1802 | 0.576544 | 37.5023 |
| CNAG_062 | - | ATP-depe | 2047.760617 | 2099.668825 | 2135.877969 | 1056.215179 | 1121.804084 | 1098.273727 | 3.49E-38 | -0.95472 | 1.27E-39 | -13.1721 | 0.515942 | 37.45737 |

| | | | | | | | | | | | | | | |
|---|---|---|---|---|---|---|---|---|---|---|---|---|---|---|
| CNAG_007 | - | hypothetic | 2194.970657 | 2335.621669 | 2698.827991 | 1103.621477 | 1132.945157 | 1123.856068 | 4.31E-38 | -1.12103 | 1.57E-39 | -13.1559 | 0.459765 | 37.36566 |
| CNAG_046 | - | hypothetic | 46.43175525 | 49.96911668 | 63.25060478 | 287.5591535 | 347.0548527 | 320.0726551 | 8.50E-38 | 2.578627 | 3.11E-39 | 13.10426 | 5.973711 | 37.07071 |
| CNAG_063 | BLP2 | pr4/barwir | 150.2599478 | 153.3553835 | 122.9839965 | 593.843949 | 476.6137011 | 602.0149308 | 9.62E-38 | 1.960563 | 3.54E-39 | 13.09459 | 3.892139 | 37.01686 |
| CNAG_006 | - | translatior | 3124.34013 | 3136.140895 | 2915.522954 | 1544.193092 | 1628.114872 | 1713.264614 | 1.19E-37 | -0.92445 | 4.40E-39 | -13.078 | 0.526881 | 36.92347 |
| CNAG_016 | - | chaperone | 7534.668586 | 6281.25197 | 6478.523211 | 3265.735355 | 3451.54713 | 3549.505034 | 2.36E-37 | -0.99928 | 8.82E-39 | -13.025 | 0.500248 | 36.62708 |
| CNAG_039 | - | hypothetic | 5.479496873 | 13.32757174 | 16.33999449 | 194.5358031 | 206.3738489 | 227.6208531 | 2.59E-37 | 4.190411 | 9.75E-39 | 13.01737 | 18.25742 | 36.58645 |
| CNAG_011 | - | small subu | 6244.583204 | 5823.34 | 6325.96896 | 3558.591718 | 3079.756446 | 3209.809737 | 5.19E-37 | -0.917 | 1.96E-38 | -12.9637 | 0.52961 | 36.2848 |
| CNAG_006 | - | large subu | 62985.67527 | 55720.37072 | 70024.07336 | 33524.99248 | 32551.77146 | 32168.37022 | 7.76E-37 | -0.95795 | 2.95E-38 | -12.9326 | 0.514787 | 36.10998 |
| CNAG_034 | - | endo alph | 2852.393845 | 2922.852591 | 2986.92571 | 1674.315468 | 1577.22302 | 1622.24785 | 9.82E-37 | -0.86125 | 3.75E-38 | -12.914 | 0.550475 | 36.00809 |
| CNAG_061 | - | hypothetic | 76436.04549 | 77298.06621 | 84592.98826 | 45471.7784 | 49571.08556 | 47313.11694 | 1.02E-36 | -0.75889 | 3.92E-38 | -12.9106 | 0.590952 | 35.98993 |
| CNAG_076 | - | large subu | 3254.022854 | 3097.437659 | 3299.394777 | 1794.91032 | 1829.996142 | 1761.028798 | 1.18E-36 | -0.85742 | 4.56E-38 | -12.899 | 0.551941 | 35.92729 |
| CNAG_030 | - | enoyl-CoA | 160.7436547 | 177.0275397 | 156.0100533 | 520.9327145 | 530.2917498 | 575.4805789 | 2.12E-36 | 1.706018 | 8.24E-38 | 12.85336 | 3.262592 | 35.6735 |
| CNAG_016 | - | elongatior | 7657.718217 | 7589.25822 | 7908.630785 | 4409.958969 | 4805.889025 | 4678.480327 | 3.66E-36 | -0.75242 | 1.44E-37 | -12.8103 | 0.593607 | 35.43621 |
| CNAG_058 | CHS5 | Chitin syn | 3555.071128 | 3484.626456 | 2991.638889 | 6170.893489 | 6379.079404 | 6482.961979 | 3.94E-36 | 0.908896 | 1.55E-37 | 12.80411 | 1.877608 | 35.40427 |
| CNAG_039 | - | chaperone | 11186.69171 | 11129.70939 | 11708.40298 | 6629.66507 | 7184.842657 | 6954.306912 | 4.33E-36 | -0.72764 | 1.71E-37 | -12.7966 | 0.603892 | 35.3635 |
| CNAG_051 | - | hypothetic | 3778.460041 | 3937.794582 | 3865.720524 | 2208.968525 | 2287.17233 | 2312.729354 | 4.52E-36 | -0.78161 | 1.79E-37 | -12.7931 | 0.581716 | 35.34529 |
| CNAG_062 | - | NADH deł | 4204.468764 | 3724.531547 | 4309.273263 | 2049.305577 | 2225.033287 | 2095.21321 | 5.18E-36 | -0.95813 | 2.06E-37 | -12.7822 | 0.514724 | 35.28552 |
| CNAG_015 | - | dehydrog | 722.2138157 | 694.1331925 | 701.9514405 | 1464.67968 | 1461.617875 | 1595.915938 | 6.45E-36 | 1.079043 | 2.58E-37 | 12.76485 | 2.112634 | 35.19028 |
| CNAG_014 | PPT1 | protein ph | 3245.781615 | 3013.394608 | 3239.661546 | 1722.515342 | 1768.751588 | 1590.108909 | 6.63E-36 | -0.9184 | 2.65E-37 | -12.7625 | 0.529095 | 35.17869 |
| CNAG_129 | - | unspecifie | 1107.66893 | 691.5487072 | 879.4511273 | 247.996289 | 239.1241104 | 238.1094115 | 8.25E-36 | -1.90389 | 3.34E-37 | -12.7447 | 0.267221 | 35.08333 |
| CNAG_028 | ERG4 | delta24(24 | 3175.903791 | 3290.265867 | 3495.594807 | 1774.620385 | 1890.378223 | 1855.376995 | 9.78E-36 | -0.86698 | 3.98E-37 | -12.731 | 0.548293 | 35.00954 |
| CNAG_023 | - | ubiquinol- | 8852.315951 | 8679.624835 | 8951.636024 | 5618.017932 | 5201.748706 | 5284.641229 | 1.23E-35 | -0.733 | 5.04E-37 | -12.7126 | 0.60165 | 34.90978 |
| CNAG_067 | HEM12 | uroporphy | 3710.514509 | 3303.252515 | 3828.298125 | 1784.669192 | 1894.641412 | 1929.77241 | 1.45E-35 | -0.96708 | 5.96E-37 | -12.6994 | 0.511539 | 34.83783 |
| CNAG_014 | - | ribosomal | 3422.115377 | 3130.850813 | 3471.067702 | 1766.060335 | 1736.776476 | 1867.580251 | 2.20E-35 | -0.91622 | 9.06E-37 | -12.6666 | 0.529897 | 34.65851 |
| CNAG_013 | - | ubiquinol- | 12433.865 | 12296.58103 | 13716.49126 | 7778.490974 | 7600.544182 | 7473.891659 | 4.15E-35 | -0.766 | 1.72E-36 | -12.6161 | 0.588046 | 34.38211 |
| CNAG_036 | - | translatior | 8791.574189 | 8367.190594 | 8963.352076 | 5108.895709 | 5432.969421 | 5280.19 | 5.93E-35 | -0.73906 | 2.47E-36 | -12.5877 | 0.599128 | 34.22702 |
| CNAG_073 | - | isocitrate | 12529.40884 | 11058.63176 | 12109.3836 | 6668.549702 | 7095.098208 | 6849.982724 | 6.57E-35 | -0.80824 | 2.75E-36 | -12.5791 | 0.57108 | 34.18253 |
| CNAG_039 | ITR6 | myo-inosi | 678.9870503 | 641.3371985 | 559.0905933 | 1390.839523 | 1464.376984 | 1378.430955 | 1.59E-34 | 1.156765 | 6.75E-36 | 12.50805 | 2.229569 | 33.79808 |
| CNAG_073 | - | nucleolin | 10243.04514 | 9532.97503 | 10349.73156 | 5627.186149 | 6066.930079 | 6097.260572 | 2.28E-34 | -0.77552 | 9.73E-36 | -12.4789 | 0.584176 | 33.64146 |
| CNAG_004 | - | hypothetic | 377.8034823 | 283.7315134 | 240.2939871 | 1031.84803 | 1011.736307 | 898.7884791 | 2.93E-34 | 1.691582 | 1.26E-35 | 12.45837 | 3.230106 | 33.5332 |
| CNAG_048 | - | cytoplasm | 282.1578648 | 272.9361251 | 343.6343072 | 808.8090756 | 801.0211928 | 894.1432551 | 5.58E-34 | 1.465057 | 2.41E-35 | 12.40641 | 2.760744 | 33.25335 |
| CNAG_127 | - | unspecifie | 44.05547146 | 29.50493565 | 41.93290538 | 300.1519809 | 269.2470413 | 231.5072725 | 5.80E-34 | 2.794687 | 2.52E-35 | 12.40291 | 6.938804 | 33.23683 |
| CNAG_038 | - | nonselecti | 196.9163461 | 224.4085359 | 199.7149046 | 575.2864761 | 603.594013 | 636.6061123 | 9.29E-34 | 1.536452 | 4.05E-35 | 12.36482 | 2.900803 | 33.03184 |
| CNAG_014 | - | serine-tRN | 14112.99203 | 13388.01649 | 13614.19315 | 8055.494271 | 8812.505713 | 8517.406845 | 1.08E-33 | -0.71129 | 4.73E-35 | -12.3523 | 0.610774 | 32.96671 |
| CNAG_037 | - | AdoMet-d | 5994.700041 | 5545.377176 | 5795.991469 | 3128.065052 | 3340.344372 | 3486.158398 | 1.14E-33 | -0.81585 | 5.01E-35 | -12.3478 | 0.568075 | 32.94334 |
| CNAG_045 | TRP1 | anthranila | 4055.249632 | 4379.519593 | 4090.767591 | 2523.465757 | 2508.85652 | 2479.265487 | 1.21E-33 | -0.75268 | 5.36E-35 | -12.3424 | 0.593498 | 32.91546 |
| CNAG_035 | - | adrenodo | 1911.178974 | 1970.392721 | 2049.487905 | 1032.90029 | 1091.633049 | 1046.095693 | 1.26E-33 | -0.91925 | 5.59E-35 | -12.3389 | 0.528782 | 32.89814 |
| CNAG_073 | - | hypothetic | 3286.737913 | 3125.459706 | 3625.65403 | 1821.367757 | 1792.010297 | 1655.613473 | 1.53E-33 | -0.94558 | 6.78E-35 | -12.3233 | 0.519221 | 32.81528 |
| CNAG_063 | - | hypothetic | 1358.675342 | 1302.390181 | 1405.319394 | 681.1094173 | 646.4648335 | 645.3694913 | 1.81E-33 | -1.05906 | 8.05E-35 | -12.3096 | 0.479943 | 32.74225 |
| CNAG_048 | - | COP9 sigr | 1496.366809 | 1534.471708 | 1530.540115 | 2850.859073 | 2702.772635 | 2656.345298 | 2.78E-33 | 0.832848 | 1.24E-34 | 12.27445 | 1.781198 | 32.556 |
| CNAG_040 | - | 60S ribosc | 3717.697677 | 3666.320451 | 4122.64252 | 1974.600532 | 2199.186483 | 2095.183368 | 3.33E-33 | -0.89175 | 1.49E-34 | -12.2596 | 0.538961 | 32.47778 |
| CNAG_048 | - | MFS trans | 74.401082 | 87.63192771 | 74.98719456 | 313.8719433 | 366.6982262 | 358.9584891 | 3.44E-33 | 2.12688 | 1.55E-34 | 12.25645 | 4.367718 | 32.46304 |
| CNAG_027 | - | nitrogen p | 1019.90502 | 960.2309573 | 936.5600646 | 1874.818723 | 1919.675755 | 1851.422498 | 3.93E-33 | 0.937764 | 1.78E-34 | 12.24543 | 1.915557 | 32.40521 |
| CNAG_029 | - | large subu | 3244.755749 | 3270.879154 | 3256.775935 | 1896.901621 | 1850.681244 | 1975.22996 | 5.39E-33 | -0.78722 | 2.45E-34 | -12.2194 | 0.57946 | 32.26848 |
| CNAG_025 | - | ATP-deper | 3417.340447 | 2899.235355 | 3409.188319 | 1658.631767 | 1684.170474 | 1671.126764 | 5.70E-33 | -0.97213 | 2.60E-34 | -12.2146 | 0.509755 | 32.24376 |
| CNAG_023 | - | cellular nu | 14149.21478 | 13214.57459 | 13858.40634 | 8721.810301 | 8788.339013 | 8600.670991 | 5.78E-33 | -0.67446 | 2.64E-34 | -12.2133 | 0.626567 | 32.23809 |
| CNAG_007 | - | ATP-bindi | 10596.67288 | 9175.257148 | 9541.389394 | 5155.491435 | 5792.6381 | 5372.380839 | 1.08E-32 | -0.86099 | 4.98E-34 | -12.1616 | 0.550573 | 31.96487 |
| CNAG_046 | - | hypothetic | 1493.993118 | 1446.13224 | 1429.239251 | 2604.757888 | 2758.951968 | 2559.759723 | 1.46E-32 | 0.843257 | 6.74E-34 | 12.13684 | 1.794096 | 31.83574 |
| CNAG_068 | KRE61 | glucosidas | 235.5253853 | 300.9144395 | 315.9235719 | 760.437483 | 910.0536718 | 791.0706783 | 1.74E-32 | 1.518874 | 8.05E-34 | 12.12228 | 2.865673 | 31.75966 |
| CNAG_063 | - | U3 small r | 3282.088625 | 3150.240814 | 3449.74689 | 1792.538599 | 1822.207127 | 1934.166506 | 2.13E-32 | -0.84814 | 9.87E-34 | -12.1056 | 0.555499 | 31.67237 |
| CNAG_055 | - | small subu | 2218.052846 | 1932.749768 | 2196.635686 | 1053.881098 | 1024.266155 | 1107.14794 | 2.75E-32 | -1.01083 | 1.28E-33 | -12.084 | 0.496261 | 31.5607 |
| CNAG_029 | - | hypothetic | 2253.26058 | 2180.526309 | 2262.811598 | 1271.154647 | 1248.618602 | 1193.749137 | 3.15E-32 | -0.86634 | 1.47E-33 | -12.0727 | 0.548536 | 31.5022 |
| CNAG_003 | - | hypothetic | 405.9329499 | 544.3330143 | 416.1973069 | 1123.989069 | 1144.361501 | 1146.308771 | 5.19E-32 | 1.308187 | 2.45E-33 | 12.03069 | 2.476301 | 31.2851 |
| CNAG_045 | ERT1 | hypothetic | 1898.039509 | 1898.639946 | 1970.964726 | 3243.667381 | 3354.891755 | 3299.109714 | 6.09E-32 | 0.763957 | 2.89E-33 | 12.01718 | 1.698142 | 31.2152 |
| CNAG_007 | - | succinyl-C | 9316.000922 | 9365.964409 | 9826.126126 | 5946.402389 | 6245.534183 | 5964.037905 | 9.16E-32 | -0.66633 | 4.36E-33 | -11.983 | 0.630109 | 31.0379 |
| CNAG_028 | - | hypothetic | 3845.880747 | 3301.097158 | 3531.900599 | 1962.242915 | 1841.140786 | 1949.747598 | 1.06E-31 | -0.90821 | 5.07E-33 | -11.9706 | 0.532844 | 30.97636 |
| CNAG_063 | - | hypothetic | 7.828656756 | 0.320185965 | 5.655322266 | 213.2091251 | 195.2491229 | 179.0858673 | 1.23E-31 | 5.596032 | 5.91E-33 | 11.95783 | 48.36971 | 30.91078 |
| CNAG_051 | - | sarcosine | 3875.163383 | 3561.802704 | 3482.893965 | 2124.957426 | 2039.578056 | 2068.52381 | 1.28E-31 | -0.8247 | 6.18E-33 | -11.9541 | 0.564601 | 30.89214 |
| CNAG_007 | - | hypothetic | 492.2413564 | 535.736899 | 517.4588196 | 1170.694114 | 1057.873364 | 1196.258754 | 1.47E-31 | 1.133834 | 7.09E-33 | 11.94264 | 2.194411 | 30.8336 |
| CNAG_052 | - | hypothetic | 34.73459662 | 44.55993376 | 44.05629008 | 238.7097652 | 275.4727373 | 232.5250391 | 1.54E-31 | 2.595367 | 7.49E-33 | 11.93816 | 6.043426 | 30.81129 |
| CNAG_027 | - | dihydrodi | 40.54702782 | 42.40715905 | 40.86056037 | 253.0088051 | 222.3839051 | 255.6286976 | 1.72E-31 | 2.566141 | 8.39E-33 | 11.92872 | 5.922231 | 30.76423 |
| CNAG_011 | - | pyruvate c | 21632.16137 | 19931.29733 | 21967.68181 | 12457.50781 | 13226.204 | 13326.61334 | 2.02E-31 | -0.71937 | 9.88E-33 | -11.9151 | 0.607363 | 30.69416 |
| CNAG_130 | - | unspecifie | 23.0678242 | 44.54647224 | 28.07507335 | 237.4523622 | 239.9638966 | 223.5649451 | 2.18E-31 | 2.865524 | 1.07E-32 | 11.90854 | 7.288007 | 30.66121 |
| CNAG_028 | - | Argininosu | 8323.490748 | 9071.782671 | 9459.233002 | 5456.061832 | 5519.209649 | 5494.445582 | 2.39E-31 | -0.72031 | 1.18E-32 | -11.9006 | 0.606965 | 30.62094 |
| CNAG_028 | - | Threonine | 2139.984526 | 2180.469608 | 2035.703905 | 1212.013238 | 1172.705589 | 1127.141784 | 2.87E-31 | -0.8709 | 1.41E-32 | -11.8852 | 0.546804 | 30.5418 |
| CNAG_054 | - | protein CN | 2309.383005 | 2288.276987 | 2517.644976 | 1258.612917 | 1292.591258 | 1339.140599 | 3.04E-31 | -0.88663 | 1.51E-32 | -11.8798 | 0.540875 | 30.51691 |
| CNAG_004 | - | mitochonc | 1268.869743 | 1316.362679 | 1423.414675 | 647.6071149 | 587.7604232 | 669.7580442 | 4.16E-31 | -1.08929 | 2.07E-32 | -11.8534 | 0.469993 | 30.38111 |
| CNAG_004 | IMD1 | inosine-5' | 13607.52079 | 14148.66804 | 14264.72344 | 8648.597116 | 9323.158844 | 9185.695628 | 4.44E-31 | -0.64493 | 2.21E-32 | -11.8477 | 0.639524 | 30.35283 |

| | | | | | | | | | | | | | | |
|---|---|---|---|---|---|---|---|---|---|---|---|---|---|---|
| CNAG_01( | - | N,N-dime | 1835.039102 | 1665.508792 | 1694.405881 | 916.9639363 | 905.2820017 | 891.7947584 | 5.26E-31 | -0.95313 | 2.64E-32 | -11.833 | 0.516511 | 30.27876 |
| CNAG_052 | - | SET doma | 4509.50216 | 4584.293815 | 4648.502601 | 2737.460884 | 2882.377148 | 2889.989013 | 5.71E-31 | -0.70682 | 2.87E-32 | -11.8259 | 0.612671 | 30.24322 |
| CNAG_023 | - | lupus La p | 11313.85192 | 10401.38828 | 11990.94835 | 5692.46216 | 6422.27496 | 6610.161956 | 6.83E-31 | -0.86383 | 3.43E-32 | -11.8108 | 0.549494 | 30.16584 |
| CNAG_051 | - | hypotheti | 105.9183783 | 139.311084 | 111.229922 | 466.9403384 | 390.4205717 | 444.5166952 | 1.04E-30 | 1.858546 | 5.23E-32 | 11.77535 | 3.62642 | 29.98423 |
| CNAG_00( | - | methionin | 3248.140947 | 3236.359115 | 3098.940449 | 1620.466855 | 1624.625824 | 1908.608125 | 1.32E-30 | -0.9102 | 6.72E-32 | -11.7542 | 0.53211 | 29.87938 |
| CNAG_011 | - | translatior | 5123.607311 | 4761.022998 | 5003.626853 | 2897.772097 | 3119.591731 | 2921.116459 | 1.88E-30 | -0.75191 | 9.62E-32 | -11.7239 | 0.593819 | 29.72601 |
| CNAG_005 | - | hypotheti | 54.54535469 | 26.27958953 | 55.78302472 | 269.0295905 | 284.7181692 | 305.5166459 | 2.47E-30 | 2.651033 | 1.27E-31 | 11.70063 | 6.281169 | 29.60787 |
| CNAG_048 | FZC43 | transcripti | 738.5133583 | 740.4230402 | 699.8090883 | 1394.574002 | 1432.309227 | 1439.504392 | 2.79E-30 | 0.955036 | 1.43E-31 | 11.69003 | 1.938629 | 29.55468 |
| CNAG_054 | - | minor hist | 2155.285042 | 2198.84127 | 2391.80626 | 1231.408264 | 1265.875562 | 1240.355363 | 2.96E-30 | -0.86742 | 1.52E-31 | -11.6848 | 0.548124 | 29.52903 |
| CNAG_074 | - | U3 small r | 3438.590137 | 3344.175535 | 3518.034685 | 1948.232311 | 1963.682381 | 2118.427546 | 3.16E-30 | -0.78798 | 1.63E-31 | -11.6791 | 0.579153 | 29.50092 |
| CNAG_019 | - | ATPase GI | 3131.424844 | 2980.004682 | 3105.327094 | 1869.731451 | 1766.163272 | 1716.618246 | 3.25E-30 | -0.79983 | 1.69E-31 | -11.6763 | 0.574417 | 29.48748 |
| CNAG_076 | - | plasma m | 2829.198603 | 3352.670188 | 3173.561026 | 1762.239331 | 1746.313785 | 1718.838325 | 3.70E-30 | -0.85442 | 1.93E-31 | -11.6649 | 0.553088 | 29.43134 |
| CNAG_075 | - | endopepti | 67.42670464 | 91.93741545 | 97.34276011 | 355.4898564 | 344.070684 | 347.9548559 | 4.28E-30 | 2.021727 | 2.23E-31 | 11.6524 | 4.060697 | 29.36874 |
| CNAG_033 | - | Mannose- | 218.9837612 | 158.7510942 | 156.0214374 | 589.6559953 | 569.998898 | 557.8823724 | 4.88E-30 | 1.672534 | 2.55E-31 | 11.64105 | 3.187741 | 29.31191 |
| CNAG_059 | - | hypotheti | 2043.209909 | 2178.338007 | 2445.065652 | 1202.633974 | 1113.993681 | 1089.429864 | 4.94E-30 | -0.98475 | 2.59E-31 | -11.6398 | 0.505312 | 29.30655 |
| CNAG_007 | - | Cytochron | 9946.478763 | 9488.794444 | 9836.803742 | 6420.477837 | 5840.914078 | 5875.247915 | 5.51E-30 | -0.70609 | 2.90E-31 | -11.63 | 0.612979 | 29.25887 |
| CNAG_069 | - | peptidyl-p | 2285.767843 | 1971.541752 | 2149.75612 | 1087.355825 | 1135.59725 | 1138.230404 | 6.06E-30 | -0.94666 | 3.19E-31 | -11.6218 | 0.518831 | 29.21788 |
| CNAG_036 | - | large subu | 2671.332434 | 2645.960424 | 2606.242229 | 1476.583313 | 1436.624907 | 1598.878386 | 6.06E-30 | -0.82787 | 3.20E-31 | -11.6216 | 0.563361 | 29.21782 |
| CNAG_068 | - | small subu | 7730.144568 | 8325.069278 | 8064.334484 | 4444.18537 | 5097.472094 | 4181.218696 | 6.37E-30 | -0.82891 | 3.38E-31 | -11.6169 | 0.562954 | 29.19586 |
| CNAG_023 | - | xaa-Pro di | 5359.383479 | 4719.010746 | 5252.065008 | 2647.822219 | 2791.719014 | 3033.214378 | 7.19E-30 | -0.87156 | 3.84E-31 | -11.6061 | 0.546555 | 29.14327 |
| CNAG_018 | - | protein M | 2462.395913 | 2487.599488 | 2599.808729 | 1426.002998 | 1499.664485 | 1461.254307 | 7.21E-30 | -0.7985 | 3.86E-31 | -11.6056 | 0.574946 | 29.14183 |
| CNAG_038 | - | protein ar | 9245.814513 | 8582.694588 | 9531.755222 | 5207.70422 | 5734.01319 | 5513.324173 | 7.38E-30 | -0.74931 | 3.97E-31 | -11.6032 | 0.594888 | 29.13173 |
| CNAG_005 | - | NADH del | 4187.123532 | 3945.415777 | 4340.259868 | 2454.16503 | 2502.821362 | 2499.238584 | 7.53E-30 | -0.75802 | 4.05E-31 | -11.6014 | 0.591307 | 29.12331 |
| CNAG_016 | - | large subu | 5375.846946 | 4916.190818 | 5413.119212 | 3008.271009 | 3204.962755 | 3138.692524 | 1.02E-29 | -0.76387 | 5.54E-31 | -11.5747 | 0.588914 | 28.99003 |
| CNAG_018 | - | Glutathior | 175.8463171 | 169.4829685 | 157.0648416 | 535.2164054 | 519.1398267 | 460.3264444 | 1.03E-29 | 1.580111 | 5.59E-31 | 11.57383 | 2.989928 | 28.98653 |
| CNAG_01( | - | carboxype | 70.86697691 | 55.34979212 | 74.96282779 | 319.8064587 | 274.8793248 | 336.6536267 | 1.12E-29 | 2.207267 | 6.10E-31 | 11.56633 | 4.617998 | 28.94953 |
| CNAG_043 | - | T-complex | 13811.79648 | 13160.71051 | 13799.75841 | 8985.691881 | 9022.115271 | 8812.695967 | 1.52E-29 | -0.6198 | 8.34E-31 | -11.5395 | 0.650762 | 28.81893 |
| CNAG_047 | - | mannose- | 236.580447 | 261.0291897 | 190.1491344 | 752.5476351 | 640.5818885 | 636.7931198 | 1.63E-29 | 1.547134 | 8.98E-31 | 11.53311 | 2.92236 | 28.78843 |
| CNAG_021 | - | translatior | 9486.365974 | 8669.979022 | 9602.155856 | 5554.880119 | 5790.929201 | 5743.11434 | 1.95E-29 | -0.71573 | 1.08E-30 | -11.5174 | 0.608897 | 28.71089 |
| CNAG_033 | ERG13 | hydroxym | 10463.80372 | 9719.395169 | 10493.73092 | 6414.939893 | 6631.087446 | 6392.561598 | 2.16E-29 | -0.67397 | 1.20E-30 | -11.5081 | 0.626779 | 28.66517 |
| CNAG_066 | MTD1 | methylene | 1910.093043 | 2156.729419 | 2219.011834 | 980.6483109 | 1167.564623 | 1011.723983 | 2.41E-29 | -1.00798 | 1.35E-30 | -11.4982 | 0.497243 | 28.61727 |
| CNAG_041 | - | phosphori | 5978.371994 | 5740.363398 | 5414.265041 | 3398.237785 | 3625.923503 | 3295.255764 | 2.67E-29 | -0.74702 | 1.49E-30 | -11.4893 | 0.595832 | 28.5736 |
| CNAG_061 | - | chaperone | 7068.875085 | 6237.072746 | 5999.742171 | 3468.179642 | 3659.482619 | 3778.146698 | 2.76E-29 | -0.83962 | 1.55E-30 | -11.4861 | 0.558792 | 28.55843 |
| CNAG_036 | - | DNA-direc | 1163.714417 | 1176.297645 | 1213.372208 | 622.7956109 | 573.1575191 | 537.7101202 | 3.34E-29 | -1.05147 | 1.88E-30 | -11.4694 | 0.482476 | 28.47629 |
| CNAG_125 | - | unspecifie | 844.9227028 | 896.1134603 | 905.1425619 | 425.8168155 | 373.8376496 | 393.3818383 | 5.21E-29 | -1.16605 | 2.94E-30 | -11.4307 | 0.445639 | 28.28344 |
| CNAG_035 | - | large subu | 22657.30207 | 21235.98977 | 21601.8896 | 10541.16382 | 13619.57615 | 11320.65592 | 6.32E-29 | -0.90027 | 3.58E-30 | -11.4135 | 0.535787 | 28.19919 |
| CNAG_131 | - | unspecifie | 20149.20019 | 21825.29685 | 21594.40834 | 13295.87642 | 10576.25655 | 11761.35324 | 6.58E-29 | -0.8498 | 3.75E-30 | -11.4096 | 0.554862 | 28.18156 |
| CNAG_027 | - | Short-chai | 1001.194056 | 814.8253823 | 872.5822123 | 1877.297727 | 1839.459699 | 1789.238829 | 6.89E-29 | 1.018687 | 3.94E-30 | 11.40518 | 2.026075 | 28.16155 |
| CNAG_056 | - | hypotheti | 123.3761778 | 132.8600411 | 144.2413809 | 421.3907911 | 419.0195569 | 427.9443573 | 8.33E-29 | 1.651857 | 4.77E-30 | 11.38848 | 3.142378 | 28.07923 |
| CNAG_005 | - | hypotheti | 4666.005658 | 4762.077881 | 4885.251727 | 3108.829843 | 2939.281353 | 2932.198852 | 9.14E-29 | -0.68789 | 5.26E-30 | -11.38 | 0.620763 | 28.03893 |
| CNAG_017 | - | ubiquinol- | 6022.913759 | 5905.263936 | 6423.05232 | 3826.352321 | 3912.287926 | 3703.770262 | 1.07E-28 | -0.69708 | 6.20E-30 | -11.3657 | 0.616817 | 27.96949 |
| CNAG_022 | - | GTP-bindi | 19416.9647 | 18106.12771 | 18708.65795 | 12506.69146 | 12220.47341 | 12278.61093 | 1.08E-28 | -0.61923 | 6.26E-30 | -11.3648 | 0.651018 | 27.96572 |
| CNAG_063 | - | hypotheti | 82.48292512 | 63.95237084 | 65.38078309 | 296.9912501 | 292.2780744 | 321.1880739 | 1.09E-28 | 2.098174 | 6.30E-30 | 11.36427 | 4.281671 | 27.9642 |
| CNAG_059 | - | nucleolar | 4841.066235 | 4547.704107 | 5081.433786 | 2770.893603 | 3000.552297 | 2818.983664 | 1.24E-28 | -0.76832 | 7.23E-30 | -11.3522 | 0.5871 | 27.90677 |
| CNAG_064 | - | acyl-CoA ( | 597.2185989 | 552.9846388 | 526.0025176 | 1111.21266 | 1219.346504 | 1234.040698 | 1.25E-28 | 1.073881 | 7.31E-30 | 11.35124 | 2.105088 | 27.90308 |
| CNAG_029 | - | hypotheti | 54.60678844 | 79.02368276 | 88.81652225 | 313.1897275 | 317.2655457 | 384.2892083 | 1.37E-28 | 2.183688 | 8.05E-30 | 11.34284 | 4.543135 | 27.86318 |
| CNAG_004 | - | V-type H | 5333.905267 | 5187.686208 | 5349.174417 | 3420.124774 | 3275.679887 | 3396.217042 | 1.48E-28 | -0.66848 | 8.73E-30 | -11.3357 | 0.62917 | 27.82877 |
| CNAG_012 | - | rRNA-proc | 3731.725803 | 3774.04484 | 4139.712225 | 1959.012034 | 2023.144895 | 2337.127171 | 1.63E-28 | -0.89698 | 9.60E-30 | -11.3274 | 0.537011 | 27.78832 |
| CNAG_01( | - | hypotheti | 13915.71503 | 13226.43134 | 13567.28799 | 9132.033081 | 8940.160565 | 8965.886392 | 1.66E-28 | -0.60586 | 9.83E-30 | -11.3253 | 0.65708 | 27.77902 |
| CNAG_024 | - | cytoplasm | 2399.272153 | 2299.072569 | 2533.660789 | 1292.858785 | 1361.618185 | 1407.953391 | 1.94E-28 | -0.84822 | 1.15E-29 | -11.3116 | 0.555469 | 27.7118 |
| CNAG_037 | - | staphyloco | 5428.40618 | 5145.653462 | 5316.101207 | 3223.158446 | 2961.638008 | 3341.81511 | 2.13E-28 | -0.75356 | 1.27E-29 | -11.3032 | 0.593138 | 27.67126 |
| CNAG_004 | - | hypotheti | 254.0276345 | 213.6713536 | 257.2596683 | 620.4199954 | 659.6851446 | 631.206999 | 2.62E-28 | 1.383572 | 1.56E-29 | 11.28505 | 2.609135 | 27.58244 |
| CNAG_00( | CIT1 | citrate syr | 18117.28214 | 17163.37031 | 17698.72471 | 11318.17534 | 11811.6432 | 11895.58431 | 2.86E-28 | -0.61258 | 1.70E-29 | -11.2771 | 0.654027 | 27.5442 |
| CNAG_026 | - | hypotheti | 109.4115728 | 144.6891672 | 114.4255476 | 466.1400844 | 453.7079709 | 375.9755139 | 2.99E-28 | 1.803526 | 1.79E-29 | 11.27274 | 3.490724 | 27.52431 |
| CNAG_037 | - | ribosomal | 5954.935732 | 5112.297136 | 5655.213119 | 3160.810014 | 3336.059303 | 3217.534774 | 3.05E-28 | -0.79973 | 1.83E-29 | -11.2707 | 0.574456 | 27.51534 |
| CNAG_047 | - | protein O | 789.9900687 | 952.6395834 | 850.171414 | 1659.97988 | 1737.699578 | 1663.797863 | 3.15E-28 | 0.950794 | 1.89E-29 | 11.26783 | 1.932937 | 27.50189 |
| CNAG_076 | - | Beta-gluc | 415.0788778 | 309.5644952 | 286.1191373 | 893.6296516 | 902.8369879 | 970.7838982 | 4.13E-28 | 1.436832 | 2.49E-29 | 11.24364 | 2.707257 | 27.38357 |
| CNAG_008 | - | hypotheti | 6791.384632 | 7202.457663 | 7827.53881 | 4322.061021 | 4585.080839 | 4350.995959 | 4.43E-28 | -0.73413 | 2.67E-29 | -11.2375 | 0.60118 | 27.35403 |
| CNAG_038 | - | nucleolar | 6689.582403 | 6113.224856 | 6768.58271 | 3734.40306 | 4052.869204 | 4022.363741 | 6.23E-28 | -0.74466 | 3.77E-29 | -11.207 | 0.596807 | 27.20533 |
| CNAG_053 | - | hypotheti | 2149.42986 | 2205.292193 | 2205.24367 | 1217.397523 | 1290.053097 | 1309.140033 | 6.66E-28 | -0.79657 | 4.03E-29 | -11.201 | 0.575716 | 27.17661 |
| CNAG_048 | MKT1 | XPG N-ter | 3136.932384 | 2966.362854 | 2822.021026 | 4968.378623 | 5150.761415 | 4799.001627 | 8.74E-28 | 0.725686 | 5.32E-29 | 11.17645 | 1.653687 | 27.05829 |
| CNAG_053 | - | ATP-depe | 3203.86837 | 3073.751944 | 3413.482549 | 1698.316243 | 1925.777333 | 1875.338961 | 8.74E-28 | -0.83307 | 5.31E-29 | -11.1766 | 0.561335 | 27.05829 |
| CNAG_034 | - | hypotheti | 66.18531757 | 70.38955725 | 58.98314443 | 294.0989446 | 298.5240298 | 255.9699844 | 1.11E-27 | 2.111188 | 6.76E-29 | 11.15511 | 4.320468 | 26.95486 |
| CNAG_013 | - | adenine p | 5899.992658 | 4919.441978 | 5481.379774 | 3146.062387 | 3100.543787 | 3035.485926 | 1.30E-27 | -0.82857 | 7.94E-29 | -11.1408 | 0.563087 | 26.88689 |
| CNAG_063 | - | hypotheti | 102.4985994 | 157.6564893 | 187.9580141 | 568.8089575 | 540.7322908 | 517.9076575 | 1.58E-27 | 1.850962 | 9.68E-29 | 11.12318 | 3.607408 | 26.80263 |
| CNAG_071 | - | hypotheti | 179.3336649 | 183.4653401 | 143.2170302 | 470.9670973 | 514.8205885 | 514.5291385 | 1.79E-27 | 1.553705 | 1.10E-28 | 11.11181 | 2.9357 | 26.74818 |
| CNAG_067 | - | hypotheti | 2192.649722 | 2192.395568 | 2474.971152 | 1309.30054 | 1222.70542 | 1257.009396 | 1.95E-27 | -0.8717 | 1.21E-28 | -11.1035 | 0.546503 | 26.70957 |
| CNAG_034 | - | hypotheti | 1942.551489 | 1751.742175 | 1905.537412 | 1001.795804 | 990.6757616 | 1054.936662 | 2.06E-27 | -0.89412 | 1.28E-28 | -11.0984 | 0.538074 | 26.68556 |

| | | | | | | | | | | | | | |
|---|---|---|---|---|---|---|---|---|---|---|---|---|---|
| CNAG_048 | - | hypothetic | 874.0498932 | 884.8308869 | 1036.752744 | 1866.34874 | 1765.18017 | 1853.589266 | 2.11E-27 | 0.957592 | 1.31E-28 | 11.09631 | 1.942065 | 26.67625 |
| CNAG_060 | - | aryl-alcoh | 43.00495325 | 13.3469481 | 29.17197182 | 679.9337235 | 670.572696 | 718.2417591 | 2.20E-27 | 4.584857 | 1.37E-28 | 11.09216 | 23.99825 | 26.65698 |
| CNAG_046 | - | hypothetic | 63.95379848 | 120.9843861 | 80.31975327 | 374.0903097 | 366.4772846 | 398.8986902 | 2.31E-27 | 2.100375 | 1.44E-28 | 11.08757 | 4.288209 | 26.63554 |
| CNAG_024 | - | NADH deh | 2365.469894 | 2346.473097 | 2523.013481 | 1428.399297 | 1363.332425 | 1425.717001 | 2.36E-27 | -0.79422 | 1.47E-28 | -11.0857 | 0.576655 | 26.62736 |
| CNAG_003 | - | 1,4-alpha- | 2855.547727 | 2728.263828 | 2242.999166 | 4868.866772 | 4881.64783 | 4954.293381 | 2.63E-27 | 0.894768 | 1.65E-28 | 11.07562 | 1.859311 | 26.57927 |
| CNAG_053 | - | hypothetic | 38.20600805 | 32.72196175 | 44.04680583 | 232.6507928 | 194.6208567 | 266.5306773 | 2.71E-27 | 2.586435 | 1.70E-28 | 11.07276 | 6.006128 | 26.56625 |
| CNAG_061 | GNO1 | S-(hydroxy | 7408.997092 | 7369.462964 | 7269.874269 | 4354.722503 | 4911.171682 | 4696.218297 | 2.88E-27 | -0.67448 | 1.81E-28 | -11.0672 | 0.626558 | 26.54021 |
| CNAG_047 | PPU1 | hypothetic | 2048.69808 | 2258.460917 | 2147.999509 | 3494.221598 | 3552.351195 | 3703.175415 | 3.02E-27 | 0.721379 | 1.90E-28 | 11.06272 | 1.648757 | 26.52011 |
| CNAG_007 | - | large subu | 2622.214062 | 2364.813918 | 2517.71321 | 1436.921885 | 1428.036101 | 1463.467305 | 3.44E-27 | -0.80978 | 2.17E-28 | -11.0509 | 0.570467 | 26.46389 |
| CNAG_012 | - | mitochond | 3315.902872 | 3047.905794 | 3261.026381 | 1879.788995 | 1974.100334 | 1773.245441 | 3.62E-27 | -0.79018 | 2.30E-28 | -11.0458 | 0.578271 | 26.44167 |
| CNAG_002 | - | compleme | 4758.082674 | 4347.297703 | 4566.389462 | 2735.144558 | 2899.647451 | 2645.808407 | 3.65E-27 | -0.7394 | 2.32E-28 | -11.0447 | 0.598989 | 26.43729 |
| CNAG_069 | - | essential r | 4290.916879 | 3727.795894 | 4241.06944 | 2227.596082 | 2377.734471 | 2395.999465 | 3.68E-27 | -0.82448 | 2.35E-28 | -11.0439 | 0.564684 | 26.43385 |
| CNAG_007 | - | hypothetic | 1711.546575 | 1798.009412 | 1981.182927 | 876.3709266 | 1010.560572 | 971.7063871 | 4.33E-27 | -0.9576 | 2.78E-28 | -11.0285 | 0.514914 | 26.36319 |
| CNAG_049 | TUB2 | tubulin be | 5747.330381 | 5655.298918 | 5920.786507 | 3690.163591 | 3667.300634 | 3830.268852 | 4.72E-27 | -0.6461 | 3.04E-28 | -11.0207 | 0.639006 | 26.32598 |
| CNAG_033 | - | hypothetic | 2182.09076 | 2108.367796 | 2316.108167 | 1203.374541 | 1265.885637 | 1288.057926 | 4.72E-27 | -0.83007 | 3.04E-28 | -11.0205 | 0.562503 | 26.32586 |
| CNAG_002 | - | zinc finger | 8803.038937 | 7414.771551 | 7997.16864 | 4556.272945 | 4828.286011 | 4806.155548 | 5.97E-27 | -0.78705 | 3.88E-28 | -10.9986 | 0.579527 | 26.22388 |
| CNAG_030 | - | bud site s | 1640.112815 | 1469.464672 | 1695.399057 | 837.5612134 | 855.2700979 | 826.2945077 | 7.05E-27 | -0.94782 | 4.59E-28 | -10.9834 | 0.518415 | 26.15211 |
| CNAG_031 | - | periodic tr | 6465.272773 | 5672.552882 | 6308.935095 | 3368.571694 | 3420.528894 | 3816.964767 | 7.55E-27 | -0.8144 | 4.94E-28 | -10.9768 | 0.568646 | 26.12204 |
| CNAG_049 | - | hypothetic | 108.2171052 | 130.6897196 | 131.4474982 | 380.0272198 | 398.36069 | 404.606276 | 7.71E-27 | 1.664756 | 5.06E-28 | 10.9747 | 3.170601 | 26.11281 |
| CNAG_065 | - | translation | 1620.300608 | 1518.998245 | 1606.939184 | 891.3406354 | 856.1298158 | 862.9048449 | 9.27E-27 | -0.87813 | 6.11E-28 | -10.9575 | 0.544072 | 26.03283 |
| CNAG_042 | - | hypothetic | 37.02919308 | 43.46585895 | 35.52600703 | 228.8818456 | 193.8449682 | 230.1054235 | 9.27E-27 | 2.486515 | 6.10E-28 | 10.9577 | 5.604227 | 26.03283 |
| CNAG_002 | - | cytoplasm | 7424.272898 | 7467.542203 | 7998.219146 | 5047.553955 | 4861.949129 | 4930.46073 | 1.04E-26 | -0.64065 | 6.85E-28 | -10.9473 | 0.641425 | 25.98442 |
| CNAG_024 | - | rRNA-proc | 1315.543457 | 1246.408263 | 1488.442495 | 637.4467471 | 700.0048835 | 647.5874645 | 1.23E-26 | -1.04493 | 8.13E-28 | -10.9317 | 0.484667 | 25.91043 |
| CNAG_020 | - | mannosyl- | 4197.716532 | 4165.198667 | 4239.004148 | 2694.736106 | 2746.976315 | 2709.028716 | 1.28E-26 | -0.64417 | 8.46E-28 | -10.9281 | 0.639862 | 25.89398 |
| CNAG_056 | - | Branched- | 2426.254238 | 2534.999974 | 2741.608335 | 1375.349341 | 1497.926887 | 1511.196441 | 1.43E-26 | -0.82834 | 9.50E-28 | -10.9176 | 0.563178 | 25.84477 |
| CNAG_052 | - | Glutamate | 3177.836238 | 3282.018191 | 2880.684033 | 5317.913339 | 5156.726076 | 5014.368086 | 1.47E-26 | 0.714739 | 9.76E-28 | 10.91511 | 1.641186 | 25.83366 |
| CNAG_049 | - | hypothetic | 1365.88245 | 1443.523194 | 1552.483152 | 775.3282619 | 709.4307864 | 782.968139 | 1.56E-26 | -0.95919 | 1.04E-27 | -10.9092 | 0.514347 | 25.80606 |
| CNAG_012 | PMC1 | calcium-tr | 1923.689979 | 2007.419532 | 1841.94303 | 3165.056062 | 3143.487373 | 3224.728409 | 1.67E-26 | 0.709007 | 1.11E-27 | 10.90324 | 1.634679 | 25.77856 |
| CNAG_017 | - | pre-mRNA | 5627.09641 | 5738.247012 | 5862.140252 | 3676.927114 | 3884.723271 | 3721.496005 | 1.73E-26 | -0.62606 | 1.15E-27 | -10.8998 | 0.647946 | 25.76315 |
| CNAG_034 | - | hypothetic | 1140.25609 | 1322.183423 | 1063.488386 | 2281.903784 | 2148.118574 | 2342.033388 | 1.89E-26 | 0.927452 | 1.27E-27 | 10.8912 | 1.901914 | 25.72267 |
| CNAG_063 | - | tRNA pseu | 4559.673197 | 4364.537773 | 4609.043942 | 2754.604639 | 2955.727895 | 2822.280895 | 2.72E-26 | -0.68116 | 1.83E-27 | -10.8577 | 0.623663 | 25.56485 |
| CNAG_015 | - | histidyl-tR | 7272.431307 | 7180.94741 | 7421.301418 | 4681.696711 | 4880.965646 | 4928.211175 | 3.20E-26 | -0.60954 | 2.16E-27 | -10.8428 | 0.655404 | 25.49486 |
| CNAG_014 | - | hypothetic | 849.4948111 | 793.8506813 | 939.1923464 | 393.0781647 | 392.0070238 | 334.6035545 | 3.29E-26 | -1.22157 | 2.22E-27 | -10.8401 | 0.428817 | 25.48277 |
| CNAG_029 | - | U6 snRNA | 1921.596715 | 1828.213467 | 1886.365633 | 993.9816059 | 1042.457733 | 1117.076188 | 3.54E-26 | -0.85355 | 2.40E-27 | -10.833 | 0.55342 | 25.4513 |
| CNAG_025 | - | Sugar tran | 31.21462034 | 45.61444296 | 30.20219817 | 259.7889391 | 197.3351038 | 202.5818437 | 3.54E-26 | 2.620041 | 2.41E-27 | 10.83271 | 6.147674 | 25.45077 |
| CNAG_046 | - | cerevisin | 725.6600325 | 714.5690004 | 613.4610391 | 1523.364823 | 1371.895899 | 1333.007259 | 3.71E-26 | 1.027487 | 2.53E-27 | 10.82819 | 2.038471 | 25.43007 |
| CNAG_058 | - | protein FA | 4038.799968 | 3859.199331 | 3931.857151 | 2553.136972 | 2287.140606 | 2390.442698 | 4.50E-26 | -0.72549 | 3.08E-27 | -10.8103 | 0.60479 | 25.34692 |
| CNAG_045 | - | poly(rC)-b | 1886.39139 | 1947.122838 | 2049.874509 | 3559.09092 | 3425.619744 | 3144.896005 | 4.62E-26 | 0.768687 | 3.16E-27 | 10.80773 | 1.703719 | 25.33551 |
| CNAG_025 | - | hypothetic | 35.91936772 | 67.14761739 | 57.90647114 | 258.8999664 | 278.7135847 | 261.3474468 | 5.20E-26 | 2.31175 | 3.57E-27 | 10.79659 | 4.964848 | 25.28435 |
| CNAG_041 | - | methylthio | 2061.683268 | 1849.802014 | 2034.580829 | 1142.752685 | 1103.703518 | 1067.187062 | 5.90E-26 | -0.85952 | 4.06E-27 | -10.7848 | 0.551136 | 25.2294 |
| CNAG_034 | - | protein GE | 2802.207305 | 2717.131495 | 2903.763217 | 1689.902903 | 1755.010122 | 1691.044624 | 6.32E-26 | -0.7293 | 4.37E-27 | -10.7781 | 0.603197 | 25.19911 |
| CNAG_000 | - | N2,N2-din | 3720.088289 | 3599.563504 | 3972.333544 | 2272.062145 | 2362.253054 | 2312.723191 | 6.85E-26 | -0.71646 | 4.74E-27 | -10.7705 | 0.608588 | 25.16416 |
| CNAG_068 | - | elongation | 109713.4274 | 107226.9944 | 105211.9045 | 65427.95577 | 73166.49511 | 72996.61743 | 7.06E-26 | -0.62185 | 4.89E-27 | -10.7676 | 0.649838 | 25.1514 |
| CNAG_075 | - | ATP-bindi | 1986.58715 | 1606.26915 | 1609.119898 | 855.4135013 | 862.9929327 | 851.8462078 | 7.94E-26 | -1.03426 | 5.52E-27 | -10.7565 | 0.488267 | 25.10035 |
| CNAG_055 | - | ATPase | 753.7380215 | 757.6811856 | 750.9976592 | 1354.607341 | 1569.506516 | 1702.441913 | 9.21E-26 | 1.017973 | 6.44E-27 | 10.74238 | 2.025072 | 25.03551 |
| CNAG_000 | - | hypothetic | 228.3459454 | 208.2654892 | 203.973095 | 507.2235491 | 658.1620707 | 602.2704001 | 1.07E-25 | 1.451729 | 7.47E-27 | 10.72866 | 2.735358 | 24.97252 |
| CNAG_070 | - | hypothetic | 572.6750113 | 529.2732357 | 486.5481413 | 1037.255891 | 1165.904026 | 1130.837108 | 1.31E-25 | 1.0546 | 9.16E-27 | 10.70972 | 2.077143 | 24.88434 |
| CNAG_067 | - | U3 small r | 4400.761244 | 3973.444281 | 4243.254012 | 2557.704158 | 2573.555692 | 2646.864211 | 1.36E-25 | -0.71385 | 9.58E-27 | -10.7056 | 0.60969 | 24.86593 |
| CNAG_048 | - | NAD-depe | 651.0439537 | 777.0336136 | 750.9727382 | 1365.712266 | 1428.852892 | 1476.100693 | 1.37E-25 | 0.956292 | 9.68E-27 | 10.70463 | 1.940317 | 24.86201 |
| CNAG_028 | TRX1 | thioredoxi | 4267.767831 | 4203.987064 | 4265.668779 | 2817.761031 | 2749.554926 | 2659.093422 | 1.45E-25 | -0.64607 | 1.02E-26 | -10.6995 | 0.63902 | 24.83855 |
| CNAG_068 | - | hypothetic | 1246.673859 | 1253.901036 | 1281.659031 | 708.4591808 | 668.0865955 | 628.7072462 | 1.45E-25 | -0.9308 | 1.03E-26 | -10.699 | 0.524569 | 24.83751 |
| CNAG_046 | - | NAD depe | 58.12840553 | 94.10286324 | 110.1316653 | 371.747697 | 432.45415 | 331.5004573 | 1.59E-25 | 2.099124 | 1.13E-26 | 10.69068 | 4.284491 | 24.79963 |
| CNAG_050 | GCS2 | ADP-ribos | 2203.10546 | 2078.221003 | 2360.887894 | 1289.071052 | 1252.931791 | 1256.995932 | 1.82E-25 | -0.82215 | 1.30E-26 | -10.6776 | 0.5656 | 24.73924 |
| CNAG_055 | - | hypothetic | 715.1678047 | 764.101452 | 620.9236279 | 1460.166494 | 1386.573108 | 1357.409039 | 1.84E-25 | 0.986935 | 1.31E-26 | 10.67682 | 1.98197 | 24.73626 |
| CNAG_046 | MBF1 | multiprote | 6440.47107 | 6335.458769 | 6371.123715 | 9556.29073 | 9581.607305 | 9370.676268 | 1.84E-25 | 0.558922 | 1.31E-26 | 10.67643 | 1.473168 | 24.73516 |
| CNAG_021 | GRE2 | D-lactalde | 5251.913967 | 6132.91555 | 4316.21106 | 11378.8422 | 10177.76001 | 10061.08459 | 1.88E-25 | 0.995927 | 1.35E-26 | 10.67408 | 1.994361 | 24.72492 |
| CNAG_015 | - | F-type H - | 19304.95358 | 18542.5002 | 20936.41222 | 12909.0587 | 12390.37545 | 12674.94261 | 2.28E-25 | -0.64594 | 1.64E-26 | -10.6555 | 0.639078 | 24.64132 |
| CNAG_052 | - | glycogeni | 680.2159716 | 821.189438 | 721.1338108 | 1381.261511 | 1485.825716 | 1470.58299 | 2.32E-25 | 0.950004 | 1.67E-26 | 10.65379 | 1.931879 | 24.63458 |
| CNAG_009 | - | F-type H - | 6657.023976 | 6516.179062 | 7624.886094 | 4196.791696 | 4279.719275 | 3981.337342 | 2.44E-25 | -0.75519 | 1.76E-26 | -10.649 | 0.592469 | 24.61318 |
| CNAG_047 | - | hypothetic | 2229.624855 | 2089.352492 | 2293.014693 | 3611.013534 | 3624.813829 | 3635.506871 | 2.84E-25 | 0.701828 | 2.06E-26 | 10.63466 | 1.626564 | 24.54683 |
| CNAG_120 | - | unspecifie | 2487.089827 | 3096.208431 | 3014.605531 | 1454.704264 | 1404.65284 | 1637.754739 | 3.16E-25 | -0.94968 | 2.29E-26 | -10.6246 | 0.517749 | 24.50063 |
| CNAG_068 | - | oxidoredu | 150.1497124 | 138.2461525 | 148.508538 | 433.0495425 | 462.2978642 | 391.5079616 | 3.19E-25 | 1.544788 | 2.32E-26 | 10.62337 | 2.917611 | 24.49574 |
| CNAG_015 | - | ribosome | 4227.849557 | 3578.05356 | 4226.124911 | 2281.355622 | 2265.575308 | 2330.506642 | 4.38E-25 | -0.82319 | 3.20E-26 | -10.5934 | 0.565192 | 24.35812 |
| CNAG_031 | - | hypothetic | 1135.533041 | 1255.37939 | 1127.433444 | 2151.146295 | 2061.02951 | 1990.272132 | 4.73E-25 | 0.803597 | 3.46E-26 | 10.58593 | 1.745448 | 24.32544 |
| CNAG_074 | - | hypothetic | 100.1176559 | 173.7398177 | 111.2389432 | 463.5943881 | 460.5284005 | 441.2947693 | 4.90E-25 | 1.815483 | 3.60E-26 | 10.58226 | 3.519775 | 24.30986 |
| CNAG_066 | - | DNA-direc | 2546.318989 | 2372.327448 | 2405.753222 | 1294.390101 | 1497.97078 | 1415.735604 | 5.89E-25 | -0.81565 | 4.34E-26 | -10.5649 | 0.568154 | 24.23007 |
| CNAG_057 | - | protein M | 2669.014767 | 2445.643208 | 2864.242098 | 1460.214827 | 1550.542276 | 1532.305016 | 6.26E-25 | -0.82851 | 4.63E-26 | -10.5588 | 0.563109 | 24.20339 |
| CNAG_035 | - | 3-beta hy | 14.82350581 | 17.61249364 | 9.922835446 | 127.088968 | 178.2488589 | 145.8208505 | 6.63E-25 | 3.476876 | 4.90E-26 | 10.5533 | 11.13381 | 24.17865 |

| | | | | | | | | | | | | | | |
|---|---|---|---|---|---|---|---|---|---|---|---|---|---|---|
| CNAG_06( | - | hypothetic | 1124.983279 | 1132.580376 | 932.3228091 | 2126.357436 | 1974.782579 | 1975.781274 | 8.53E-25 | 0.915271 | 6.33E-26 | 10.52925 | 1.885923 | 24.06903 |
| CNAG_04( | - | hypothetic | 115.1799742 | 127.4549009 | 122.9229683 | 344.790798 | 409.7094333 | 384.6580491 | 9.03E-25 | 1.628149 | 6.72E-26 | 10.52369 | 3.091162 | 24.04409 |
| CNAG_03{ | - | histidinol- | 4375.328013 | 4780.370753 | 4871.373105 | 3006.851253 | 3007.464024 | 2807.878302 | 9.74E-25 | -0.68427 | 7.26E-26 | -10.5164 | 0.622319 | 24.01124 |
| CNAG_04( | - | hypothetic | 863.4257546 | 761.5531522 | 856.0888316 | 392.3171893 | 388.5625679 | 378.9406708 | 1.00E-24 | -1.11454 | 7.49E-26 | -10.5134 | 0.461838 | 23.99808 |
| CNAG_02! | - | cytochrom | 7780.380878 | 7488.029945 | 8307.474309 | 4993.802821 | 4754.952203 | 5171.34366 | 1.07E-24 | -0.67562 | 7.96E-26 | -10.5077 | 0.626064 | 23.97258 |
| CNAG_06' | VRK1 | hypothetic | 573.9248083 | 659.5847014 | 594.2236449 | 1147.028228 | 1196.828652 | 1213.019483 | 1.10E-24 | 0.946891 | 8.21E-26 | 10.50481 | 1.927714 | 23.95995 |
| CNAG_00{ | ERG2 | C-8 sterol | 1657.730356 | 1589.035705 | 1762.60035 | 911.5385336 | 845.7305368 | 963.8874693 | 1.11E-24 | -0.89651 | 8.31E-26 | -10.5036 | 0.537183 | 23.95522 |
| CNAG_02{ | ARG3 | Ornithine | 3026.498263 | 3171.768847 | 3111.76258 | 1995.847557 | 2012.073461 | 1904.193614 | 1.24E-24 | -0.67018 | 9.31E-26 | -10.4929 | 0.628427 | 23.90788 |
| CNAG_01{ | - | hypothetic | 1261.97896 | 1311.030071 | 1562.004348 | 692.7718792 | 687.8940414 | 684.2063913 | 1.36E-24 | -1.01813 | 1.03E-25 | -10.4837 | 0.493754 | 23.86725 |
| CNAG_04' | PKP2 | pyruvate ( | 780.6371301 | 854.0266092 | 764.4331724 | 328.3658188 | 327.2441175 | 402.1874379 | 1.45E-24 | -1.19642 | 1.10E-25 | -10.4773 | 0.436358 | 23.83834 |
| CNAG_02: | - | H/ACA rib | 5952.719405 | 5473.219454 | 5790.676009 | 3258.069697 | 3605.204698 | 3703.716726 | 1.53E-24 | -0.7199 | 1.16E-25 | -10.4718 | 0.607138 | 23.8144 |
| CNAG_04{ | - | golgi pho: | 4354.88288 | 4141.832268 | 4380.053945 | 6681.340701 | 6563.503808 | 6560.830475 | 1.54E-24 | 0.605718 | 1.17E-25 | 10.4712 | 1.521735 | 23.81252 |
| CNAG_06; | - | U3 small r | 1283.84995 | 1100.986027 | 1329.563042 | 645.3599712 | 565.3565095 | 576.5498055 | 1.72E-24 | -1.07143 | 1.32E-25 | -10.4602 | 0.475847 | 23.76411 |
| CNAG_04: | - | protein SE | 4829.400905 | 4507.850349 | 4874.582881 | 2914.946327 | 2895.283325 | 3108.663647 | 1.81E-24 | -0.6878 | 1.38E-25 | -10.4554 | 0.620798 | 23.74296 |
| CNAG_02{ | - | amidopho | 5254.54588 | 5179.075293 | 5266.010673 | 3393.639366 | 3518.984733 | 3537.180658 | 1.87E-24 | -0.60278 | 1.44E-25 | -10.4519 | 0.658484 | 23.72759 |
| CNAG_01' | - | Hsp70-like | 5028.787491 | 4192.152452 | 4278.476333 | 2632.399877 | 2594.225946 | 2553.667979 | 2.03E-24 | -0.81113 | 1.57E-25 | -10.4437 | 0.569937 | 23.69197 |
| CNAG_02( | - | hypothetic | 1096.93289 | 1063.632026 | 1014.393161 | 1868.48679 | 1999.049697 | 1793.786004 | 2.03E-24 | 0.819504 | 1.56E-25 | 10.44378 | 1.764799 | 23.69197 |
| CNAG_04( | CTF4 | chromoso | 2558.894356 | 2523.515655 | 2332.513187 | 3859.175592 | 4108.745638 | 4061.801836 | 2.27E-24 | 0.683025 | 1.76E-25 | 10.43283 | 1.605502 | 23.64372 |
| CNAG_00( | - | U6 snRNA | 2536.079513 | 2772.016454 | 2749.137883 | 1590.256645 | 1638.540607 | 1698.779619 | 2.69E-24 | -0.7247 | 2.09E-25 | -10.4164 | 0.605123 | 23.5706 |
| CNAG_00! | - | translation | 4298.135451 | 4243.85252 | 4328.585466 | 2629.292325 | 2857.40358 | 2815.589563 | 3.19E-24 | -0.64792 | 2.49E-25 | -10.3996 | 0.6382 | 23.4956 |
| CNAG_00{ | - | hypothetic | 626.4766582 | 682.7113615 | 597.0179758 | 273.9396776 | 290.2278087 | 277.9271657 | 4.82E-24 | -1.19617 | 3.77E-25 | -10.3599 | 0.436431 | 23.31686 |
| CNAG_05: | TIM54 | mitochon( | 1241.920824 | 1123.603644 | 1291.212349 | 639.1056248 | 637.8915623 | 632.010518 | 5.14E-24 | -0.95358 | 4.03E-25 | -10.3535 | 0.51635 | 23.28914 |
| CNAG_02' | - | NADH del | 5349.075423 | 5060.571073 | 5195.628365 | 3416.234705 | 3505.189114 | 3384.006518 | 7.92E-24 | -0.61413 | 6.24E-25 | -10.3116 | 0.653325 | 23.10147 |
| CNAG_01! | - | mitochon( | 1151.926907 | 1058.909976 | 1137.662463 | 558.1296601 | 604.2965912 | 487.7736235 | 8.16E-24 | -1.03674 | 6.45E-25 | -10.3086 | 0.487427 | 23.08839 |
| CNAG_06; | - | DNA-direc | 1370.204286 | 1157.005228 | 1289.110713 | 630.488994 | 688.820414 | 603.1923616 | 8.49E-24 | -1.00638 | 6.71E-25 | -10.3046 | 0.497794 | 23.0713 |
| CNAG_05: | - | translation | 12691.93487 | 12081.1297 | 12579.7476 | 8033.790475 | 8415.744174 | 8672.764426 | 8.82E-24 | -0.5878 | 6.99E-25 | -10.3007 | 0.665358 | 23.05441 |
| CNAG_00: | - | high-affini | 56.82403793 | 62.83179709 | 48.3159452 | 226.8019292 | 234.504314 | 255.691265 | 1.02E-23 | 2.092057 | 8.11E-25 | 10.28641 | 4.263555 | 22.99107 |
| CNAG_01; | - | 2,4-dihydr | 71.92515716 | 45.64163165 | 46.19094754 | 245.6577262 | 244.8861124 | 257.9502182 | 1.11E-23 | 2.18693 | 8.80E-25 | 10.27858 | 4.553355 | 22.95647 |
| CNAG_07! | - | farnesyl-d | 7911.186499 | 7774.614867 | 7935.344505 | 5016.341961 | 5452.873616 | 5373.3819 | 1.61E-23 | -0.59175 | 1.29E-24 | -10.2417 | 0.663538 | 22.79308 |
| CNAG_01: | - | NADH-ubi | 8068.737963 | 7851.079651 | 7455.472224 | 4783.597972 | 5063.807425 | 5311.202656 | 1.74E-23 | -0.64018 | 1.39E-24 | -10.2342 | 0.641634 | 22.76028 |
| CNAG_04; | PCD1 | hypothetic | 862.2257567 | 853.5275638 | 740.3450465 | 1484.005938 | 1525.421654 | 1560.520033 | 1.76E-23 | 0.881053 | 1.41E-24 | 10.23296 | 1.841719 | 22.75534 |
| CNAG_02{ | - | glycerol-3 | 7837.588039 | 8174.268429 | 7308.317812 | 4949.424245 | 4991.337821 | 5290.113319 | 2.04E-23 | -0.62945 | 1.64E-24 | -10.2183 | 0.646424 | 22.69098 |
| CNAG_05' | - | hypothetic | 187.6095451 | 249.1611015 | 258.3057632 | 601.8380534 | 593.1670687 | 592.3585254 | 2.04E-23 | 1.349246 | 1.64E-24 | 10.21843 | 2.547789 | 22.69098 |
| CNAG_03{ | - | hsp60-like | 22951.73291 | 21175.73533 | 23553.49304 | 12540.70611 | 14496.78321 | 14639.91031 | 2.27E-23 | -0.7153 | 1.83E-24 | -10.2076 | 0.609076 | 22.6437 |
| CNAG_02' | - | hypothetic | 437.3086926 | 489.3698245 | 366.0710355 | 999.5231497 | 898.3188227 | 982.0379596 | 2.64E-23 | 1.141572 | 2.14E-24 | 10.19272 | 2.206212 | 22.57849 |
| CNAG_07! | - | 6-phospho | 72679.45768 | 73413.95664 | 73880.57034 | 49850.60231 | 50824.42833 | 53821.87312 | 3.11E-23 | -0.52491 | 2.52E-24 | -10.1765 | 0.695002 | 22.50673 |
| CNAG_04; | - | hypothetic | 1874.622143 | 1909.375751 | 1850.45301 | 2900.31821 | 3015.835066 | 3078.189075 | 3.28E-23 | 0.659772 | 2.68E-24 | 10.17081 | 1.579832 | 22.48396 |
| CNAG_06; | - | F-type H - | 6297.413225 | 6716.523771 | 7149.283218 | 4452.988729 | 3813.858179 | 3901.400231 | 3.28E-23 | -0.74378 | 2.67E-24 | -10.1709 | 0.597174 | 22.48396 |
| CNAG_01( | - | DNA poly | 9621.829483 | 8622.591857 | 9380.368743 | 5837.449653 | 6139.445806 | 5971.789751 | 3.37E-23 | -0.63799 | 2.75E-24 | -10.1681 | 0.642607 | 22.47265 |
| CNAG_03! | - | oxidoredu | 11074.18144 | 12337.79548 | 10576.19204 | 17787.55021 | 17240.26563 | 17806.73072 | 3.56E-23 | 0.621927 | 2.92E-24 | 10.16248 | 1.53893 | 22.4481 |
| CNAG_05< | - | NADH del | 4077.380413 | 3901.237352 | 3958.537893 | 2602.152351 | 2651.252394 | 2439.296564 | 4.22E-23 | -0.64944 | 3.47E-24 | -10.1455 | 0.63753 | 22.37443 |
| CNAG_00! | - | hypothetic | 2215.913043 | 2180.518234 | 2053.864586 | 1238.440971 | 1294.373134 | 1345.751026 | 5.75E-23 | -0.74912 | 4.74E-24 | -10.1151 | 0.594968 | 22.2405 |
| CNAG_05! | - | hypothetic | 16.06577054 | 52.07850601 | 45.11684739 | 225.3762518 | 269.4525933 | 235.7787336 | 6.46E-23 | 2.676593 | 5.34E-24 | 10.10341 | 6.393443 | 22.18956 |
| CNAG_00: | - | hypothetic | 279.6642904 | 259.9511994 | 283.8983994 | 602.3459394 | 663.958171 | 613.5303246 | 7.26E-23 | 1.178105 | 6.00E-24 | 10.09182 | 2.262794 | 22.13889 |
| CNAG_02: | - | tRNA-dihy | 1104.024807 | 975.9772492 | 1053.436177 | 509.0509831 | 550.7848356 | 549.8442272 | 7.96E-23 | -0.97762 | 6.61E-24 | -10.0824 | 0.507816 | 22.09934 |
| CNAG_07( | - | mitochon( | 1230.439402 | 1286.237868 | 1354.162096 | 729.4594413 | 732.8234268 | 681.969314 | 8.08E-23 | -0.8686 | 6.72E-24 | -10.0807 | 0.547677 | 22.09236 |
| CNAG_02: | - | nucleoside | 22912.15286 | 21884.70633 | 24555.97145 | 15094.46394 | 15932.9878 | 15168.36266 | 8.68E-23 | -0.6019 | 7.25E-24 | -10.0733 | 0.658884 | 22.06146 |
| CNAG_00< | - | T-complex | 11302.31362 | 10714.9508 | 11279.73659 | 7134.858529 | 7709.334344 | 7589.263384 | 9.29E-23 | -0.58534 | 7.77E-24 | -10.0665 | 0.666492 | 22.03193 |
| CNAG_03! | THI20 | phosphom | 4905.069126 | 4012.261746 | 4531.17542 | 2504.690863 | 2647.723232 | 2673.532425 | 1.02E-22 | -0.79763 | 8.54E-24 | -10.0572 | 0.575293 | 21.99284 |
| CNAG_05' | - | U3 small r | 6832.088278 | 6173.587869 | 7044.792986 | 3996.727719 | 4269.384831 | 4305.432139 | 1.13E-22 | -0.68939 | 9.54E-24 | -10.0463 | 0.620117 | 21.94794 |
| CNAG_01: | - | hypothetic | 347.5130055 | 465.6590544 | 418.2683033 | 899.7644802 | 902.7655161 | 877.7607157 | 1.30E-22 | 1.108256 | 1.10E-23 | 10.03199 | 2.155848 | 21.88691 |
| CNAG_00( | - | Ribulose-p | 3673.442313 | 3698.677642 | 3949.959371 | 2362.375562 | 2363.099183 | 2515.830545 | 1.41E-22 | -0.66005 | 1.20E-23 | -10.0238 | 0.632859 | 21.85143 |
| CNAG_04! | - | SEL1 prote | 1065.491103 | 1212.27934 | 1160.464937 | 1922.889168 | 1950.593709 | 2029.040722 | 1.68E-22 | 0.765102 | 1.43E-23 | 10.00602 | 1.69949 | 21.77405 |
| CNAG_01' | - | hypothetic | 1662.470946 | 1603.067051 | 1839.376377 | 993.3312198 | 958.7988918 | 940.6147662 | 1.77E-22 | -0.83561 | 1.52E-23 | -10.0006 | 0.560345 | 21.75132 |
| CNAG_01: | - | pre-mRNA | 2527.909145 | 2718.163931 | 2884.531772 | 1503.012508 | 1645.446219 | 1688.791724 | 1.83E-22 | -0.76432 | 1.57E-23 | -9.99735 | 0.58873 | 21.73849 |
| CNAG_00: | - | bud site s | 2178.571685 | 2015.743696 | 2214.828873 | 1282.861215 | 1290.9253 | 1291.3802 | 2.10E-22 | -0.74557 | 1.80E-23 | -9.98342 | 0.59643 | 21.67871 |
| CNAG_00! | - | hypothetic | 1860.916447 | 1777.58871 | 1978.022982 | 1100.767047 | 902.6231116 | 1050.50085 | 2.59E-22 | -0.89484 | 2.23E-23 | -9.96201 | 0.537806 | 21.58631 |
| CNAG_02; | - | F-type H - | 12458.50644 | 12617.68784 | 13461.68064 | 8668.153855 | 9078.228423 | 8743.841352 | 2.92E-22 | -0.55618 | 2.53E-23 | -9.94984 | 0.680102 | 21.53438 |
| CNAG_00' | - | phosphori | 3832.273357 | 4237.306252 | 3893.493609 | 2427.717285 | 2695.265663 | 2438.182793 | 2.93E-22 | -0.67691 | 2.53E-23 | -9.94955 | 0.625503 | 21.53374 |
| CNAG_07< | - | hypothetic | 1492.043394 | 1571.76262 | 1599.482907 | 914.7188866 | 879.4370441 | 936.1220791 | 2.97E-22 | -0.78758 | 2.57E-23 | -9.94808 | 0.579316 | 21.52794 |
| CNAG_07( | DST1 | transcripti | 2232.354347 | 2140.720201 | 2365.188161 | 1381.733458 | 1373.738248 | 1353.554683 | 3.27E-22 | -0.72956 | 2.85E-23 | -9.93776 | 0.603088 | 21.48535 |
| CNAG_12! | - | unspecifie | 4844.810472 | 5838.328529 | 6305.621582 | 3161.61736 | 2572.50528 | 3180.893095 | 3.35E-22 | -0.94474 | 2.93E-23 | -9.93516 | 0.519524 | 21.47466 |
| CNAG_00: | - | translation | 4399.295048 | 3726.658359 | 3478.673231 | 2060.243273 | 2273.380492 | 2160.674247 | 3.41E-22 | -0.85352 | 2.98E-23 | -9.93346 | 0.553434 | 21.46783 |
| CNAG_12: | - | unspecifie | 116938.3664 | 112552.7826 | 113379.8832 | 77916.53126 | 84302.21449 | 78893.80299 | 4.14E-22 | -0.52352 | 3.63E-23 | -9.91358 | 0.695674 | 21.38251 |
| CNAG_04; | - | membrane | 130.3491833 | 164.0406561 | 121.8779291 | 424.5757222 | 438.1240639 | 368.2199457 | 4.15E-22 | 1.552988 | 3.64E-23 | 9.913386 | 2.934243 | 21.38229 |
| CNAG_02' | - | kinetocho | 962.770075 | 1145.478207 | 1082.619054 | 1829.514963 | 1860.905325 | 1941.31846 | 4.55E-22 | 0.805498 | 4.01E-23 | 9.903806 | 1.747749 | 21.34185 |
| CNAG_04( | - | mitochon( | 965.0619869 | 1026.9942 | 1100.721785 | 1781.310608 | 1809.180296 | 1761.552759 | 4.57E-22 | 0.776578 | 4.03E-23 | 9.903231 | 1.713063 | 21.33996 |
| CNAG_01: | - | hypothetic | 5.425742624 | 8.981320513 | 8.842588142 | 87.69593416 | 143.667782 | 145.4488398 | 5.50E-22 | 4.143573 | 4.87E-23 | 9.884392 | 17.6742 | 21.2594 |

| | | | | | | | | | | | | | | |
|---|---|---|---|---|---|---|---|---|---|---|---|---|---|---|
| CNAG_005 | CHS4 | Chitin syn | 1098.08524 | 1075.471997 | 1063.425689 | 1844.359147 | 1899.780365 | 1742.740898 | 7.30E-22 | 0.745871 | 6.49E-23 | 9.855461 | 1.676986 | 21.13654 |
| CNAG_007 | - | ATP-depe | 43058.19 | 40251.70642 | 43271.20857 | 29170.68422 | 30100.57007 | 28982.07447 | 7.65E-22 | -0.53605 | 6.81E-23 | -9.85066 | 0.689657 | 21.11638 |
| CNAG_023 | YPK101 | AGC prote | 144.3064003 | 147.9123899 | 158.0786728 | 408.0875341 | 399.1015242 | 392.5463388 | 8.72E-22 | 1.401805 | 7.79E-23 | 9.837189 | 2.642319 | 21.05939 |
| CNAG_069 | - | pyridoxal | 5982.172248 | 6426.713379 | 6770.719384 | 4353.372384 | 4026.1193 | 3960.209717 | 8.92E-22 | -0.65126 | 7.99E-23 | -9.83464 | 0.636724 | 21.04941 |
| CNAG_035 | - | cytoplasm | 7.789091275 | 16.52267266 | 14.17825594 | 151.5336163 | 119.7766206 | 122.6011246 | 9.26E-22 | 3.39131 | 8.32E-23 | 9.830563 | 10.49267 | 21.03318 |
| CNAG_056 | IGI1 | indigoidin | 2261.480974 | 2191.319764 | 2155.163864 | 1326.440084 | 1302.113034 | 1424.552685 | 9.93E-22 | -0.7206 | 8.93E-23 | -9.82342 | 0.606847 | 21.00297 |
| CNAG_011 | - | hypothetic | 969.6455072 | 980.6476308 | 887.4882427 | 1700.467798 | 1686.713956 | 1573.982738 | 9.96E-22 | 0.791024 | 8.96E-23 | 9.823012 | 1.730302 | 21.00181 |
| CNAG_044 | - | nicotinam | 481.5923768 | 460.8716035 | 491.34162 | 207.8212022 | 193.5617977 | 199.1160744 | 1.02E-21 | -1.27282 | 9.20E-23 | -9.82036 | 0.41385 | 20.99098 |
| CNAG_004 | LEU2 | 3-isopropy | 2472.967274 | 2552.252878 | 2737.363736 | 1425.958367 | 1676.565573 | 1472.373703 | 1.06E-21 | -0.77855 | 9.57E-23 | -9.81637 | 0.582953 | 20.97435 |
| CNAG_063 | SGF29 | SAGA-ass | 1078.404188 | 1069.622944 | 1006.558014 | 547.2694611 | 478.2235681 | 586.4395278 | 1.16E-21 | -0.98372 | 1.05E-22 | -9.80691 | 0.505676 | 20.93425 |
| CNAG_005 | - | mitochonc | 1805.951595 | 1648.307805 | 1659.25526 | 1014.35187 | 994.1811901 | 1006.079081 | 1.22E-21 | -0.77778 | 1.10E-22 | -9.80228 | 0.583264 | 20.91491 |
| CNAG_044 | - | major faci | 34.65090066 | 31.6189354 | 32.31267508 | 147.8034735 | 204.6989199 | 194.5880899 | 1.24E-21 | 2.474609 | 1.12E-22 | 9.800418 | 5.558165 | 20.90749 |
| CNAG_062 | - | hypothetic | 1367.953323 | 1266.843755 | 1188.964197 | 698.2893841 | 686.2030704 | 715.2326851 | 1.29E-21 | -0.87966 | 1.17E-22 | -9.79577 | 0.543497 | 20.88808 |
| CNAG_005 | - | WD40 rep | 1649.53109 | 1611.631272 | 1516.370723 | 956.7700751 | 934.6686358 | 939.4708759 | 1.35E-21 | -0.77076 | 1.22E-22 | -9.79149 | 0.58611 | 20.8703 |
| CNAG_002 | - | DNA repa | 2299.664521 | 2347.880187 | 1995.537236 | 3636.691475 | 3593.730073 | 3768.686295 | 1.45E-21 | 0.712737 | 1.32E-22 | 9.783972 | 1.638911 | 20.83858 |
| CNAG_033 | - | cytoplasm | 3262.78359 | 2469.306485 | 2379.1437 | 1263.169324 | 1344.298399 | 1409.112342 | 1.58E-21 | -1.0303 | 1.44E-22 | -9.77493 | 0.489609 | 20.80038 |
| CNAG_002 | IDI1 | isopenten | 2151.707583 | 1979.114158 | 2163.646764 | 1284.441945 | 1259.864652 | 1282.492886 | 1.66E-21 | -0.73392 | 1.51E-22 | -9.77008 | 0.601268 | 20.78016 |
| CNAG_034 | CHI2 | Chitinase | 932.2692599 | 934.3188931 | 824.577466 | 1582.089236 | 1629.807804 | 1541.751888 | 1.68E-21 | 0.805757 | 1.53E-22 | 9.768656 | 1.748063 | 20.77522 |
| CNAG_009 | RRP44 | exosome | 3500.505917 | 3420.679608 | 3295.227592 | 2090.718294 | 2282.081658 | 2222.767848 | 1.70E-21 | -0.64682 | 1.56E-22 | -9.76721 | 0.638686 | 20.76958 |
| CNAG_047 | - | KH domai | 4065.248699 | 3614.98915 | 3901.22675 | 5910.752976 | 6146.074573 | 6107.852886 | 1.90E-21 | 0.633545 | 1.74E-22 | 9.756025 | 1.551372 | 20.72227 |
| CNAG_014 | - | 3-deoxy-7 | 6082.510721 | 6107.817148 | 6720.578109 | 3742.998883 | 4303.095665 | 3729.320223 | 2.02E-21 | -0.69909 | 1.86E-22 | -9.74919 | 0.615962 | 20.69418 |
| CNAG_021 | - | hypothetic | 56.77546043 | 49.91825104 | 48.30037981 | 224.0075776 | 206.8519768 | 209.2442249 | 2.50E-21 | 2.046847 | 2.30E-22 | 9.727475 | 4.13202 | 20.60254 |
| CNAG_048 | - | hypothetic | 116.3282013 | 111.318991 | 133.5614309 | 369.2276506 | 414.9783949 | 319.4126016 | 2.61E-21 | 1.598216 | 2.42E-22 | 9.722429 | 3.027686 | 20.5833 |
| CNAG_070 | LPD1alpha | dihydrolip | 16331.80845 | 15076.41588 | 15880.43303 | 10764.8824 | 11121.63055 | 10858.6908 | 2.67E-21 | -0.54596 | 2.48E-22 | -9.71977 | 0.684938 | 20.57309 |
| CNAG_007 | - | mitochonc | 1234.298903 | 903.7947654 | 930.841651 | 477.9305435 | 433.3573041 | 435.5892493 | 2.83E-21 | -1.20571 | 2.63E-22 | -9.71387 | 0.433557 | 20.54851 |
| CNAG_048 | - | hypothetic | 319.4082129 | 345.0322986 | 377.7095847 | 725.3424171 | 745.774831 | 720.1320063 | 2.93E-21 | 1.057814 | 2.73E-22 | 9.71013 | 2.081775 | 20.53315 |
| CNAG_060 | - | translatior | 1492.046803 | 1536.23248 | 1617.603731 | 912.3779954 | 912.2446195 | 935.0139928 | 2.94E-21 | -0.76671 | 2.75E-22 | -9.70948 | 0.587756 | 20.53094 |
| CNAG_047 | - | adenine n | 674.2804488 | 693.0015684 | 604.9110585 | 1170.977682 | 1276.148832 | 1431.582043 | 3.15E-21 | 0.961141 | 2.94E-22 | 9.702454 | 1.946849 | 20.50159 |
| CNAG_012 | - | myosin I b | 2568.190008 | 2115.238685 | 2275.98717 | 3883.41648 | 3956.065032 | 3996.276184 | 3.18E-21 | 0.75024 | 2.98E-22 | 9.701265 | 1.682073 | 20.4971 |
| CNAG_013 | - | hypothetic | 2706.599736 | 3006.894337 | 3030.664866 | 1640.000028 | 1875.798566 | 1795.382124 | 3.19E-21 | -0.73422 | 2.99E-22 | -9.70087 | 0.601143 | 20.496 |
| CNAG_011 | - | nuclear er | 127.9738606 | 130.6823705 | 139.9572024 | 345.5564945 | 404.4893226 | 365.8806864 | 3.47E-21 | 1.476783 | 3.26E-22 | 9.692163 | 2.783274 | 20.46006 |
| CNAG_039 | - | NAD(P)H:c | 1930.668937 | 1827.526628 | 2002.930522 | 3123.828099 | 2990.762634 | 3120.391522 | 3.81E-21 | 0.665507 | 3.58E-22 | 9.682338 | 1.586126 | 20.41956 |
| CNAG_048 | SRE1 | sterol regu | 1046.769774 | 1063.63234 | 1142.317655 | 1779.619405 | 1978.326163 | 1831.502861 | 4.02E-21 | 0.766043 | 3.80E-22 | 9.676443 | 1.700599 | 20.39607 |
| CNAG_038 | - | cytochron | 4461.591458 | 4312.815689 | 4531.192485 | 3044.316904 | 2634.763353 | 2699.073177 | 4.07E-21 | -0.68272 | 3.86E-22 | -9.67472 | 0.622992 | 20.38988 |
| CNAG_075 | - | hypothetic | 75.44823751 | 67.15309283 | 65.36006351 | 232.7770096 | 298.6661051 | 245.9517733 | 5.07E-21 | 1.88877 | 4.81E-22 | 9.652153 | 3.703194 | 20.29474 |
| CNAG_011 | - | ribosome | 3320.503 | 2799.072115 | 3159.716262 | 1844.001018 | 1910.266563 | 1792.091752 | 5.82E-21 | -0.75852 | 5.53E-22 | -9.63786 | 0.591102 | 20.23535 |
| CNAG_014 | SMT1 | sphingolip | 8727.432297 | 8520.198621 | 8439.802186 | 5611.764629 | 5830.649162 | 6143.797984 | 6.15E-21 | -0.562 | 5.86E-22 | -9.63195 | 0.677361 | 20.21093 |
| CNAG_011 | - | hypothetic | 1643.374665 | 1558.155987 | 1332.223772 | 2570.430168 | 2637.215426 | 2737.307169 | 6.55E-21 | 0.794169 | 6.24E-22 | 9.625449 | 1.734078 | 20.184 |
| CNAG_040 | - | ribosome | 2523.010223 | 2238.79166 | 2479.321016 | 1436.164657 | 1487.603459 | 1488.977047 | 7.13E-21 | -0.73072 | 6.81E-22 | -9.61655 | 0.602604 | 20.14699 |
| CNAG_034 | - | hypothetic | 5047.901344 | 5020.705765 | 5196.687694 | 3425.591825 | 3597.519066 | 3409.527777 | 7.57E-21 | -0.56456 | 7.25E-22 | -9.61013 | 0.67616 | 20.12098 |
| CNAG_129 | - | unspecifie | 12.48978312 | 21.91162348 | 22.70530681 | 172.6155354 | 128.3039554 | 133.7523636 | 7.83E-21 | 2.962644 | 7.51E-22 | 9.606502 | 7.795511 | 20.10626 |
| CNAG_029 | - | malonic se | 1021.253686 | 997.4829 | 982.0290231 | 544.9625747 | 574.1265519 | 526.5450941 | 9.17E-21 | -0.88317 | 8.81E-22 | -9.59 | 0.542176 | 20.03783 |
| CNAG_047 | - | ARP2/3 cc | 466.4274416 | 468.9091025 | 412.9554077 | 892.5928136 | 1004.678305 | 847.9049196 | 1.11E-20 | 1.011737 | 1.07E-21 | 9.570003 | 2.016338 | 19.95541 |
| CNAG_036 | - | hypothetic | 1764.104116 | 1765.740255 | 1868.229249 | 1010.369166 | 1121.03142 | 1111.516287 | 1.29E-20 | -0.75063 | 1.25E-21 | -9.55381 | 0.594342 | 19.88856 |
| CNAG_075 | - | DNA-direc | 1051.593187 | 998.5797623 | 1210.062235 | 428.7122735 | 463.5437182 | 588.6533059 | 1.31E-20 | -1.15523 | 1.27E-21 | -9.55255 | 0.448995 | 19.88382 |
| CNAG_067 | - | hypothetic | 213.1360947 | 232.9725336 | 208.2052802 | 495.7915636 | 516.3878707 | 502.4865001 | 1.36E-20 | 1.197325 | 1.32E-21 | 9.548438 | 2.29314 | 19.86712 |
| CNAG_053 | - | hypothetic | 1813.045006 | 1691.420019 | 1811.717067 | 1011.954972 | 1025.224034 | 1117.047831 | 1.39E-20 | -0.76883 | 1.35E-21 | -9.5461 | 0.586891 | 19.85787 |
| CNAG_074 | - | hypothetic | 373.0642246 | 410.1068285 | 395.3530672 | 130.5769165 | 140.0397341 | 169.1029537 | 1.57E-20 | -1.43876 | 1.52E-21 | -9.53332 | 0.368884 | 19.80488 |
| CNAG_074 | NOG2 | nucleolar | 7152.246564 | 7660.369777 | 8578.295853 | 4342.991304 | 4560.00102 | 5141.351007 | 1.74E-20 | -0.75104 | 1.69E-21 | -9.52234 | 0.594174 | 19.75949 |
| CNAG_128 | - | unspecifie | 2289.098575 | 2110.875211 | 1893.158467 | 3427.305873 | 3531.685331 | 3521.146466 | 1.91E-20 | 0.720457 | 1.86E-21 | 9.512559 | 1.647703 | 19.71919 |
| CNAG_036 | - | pescadillo | 6816.920987 | 6158.509479 | 6781.41673 | 4159.44357 | 4439.344676 | 4390.901835 | 1.94E-20 | -0.62086 | 1.89E-21 | -9.51064 | 0.650283 | 19.71173 |
| CNAG_078 | - | protein m | 1108.958552 | 1105.261523 | 1235.75817 | 653.9823583 | 591.2879527 | 637.5275498 | 2.00E-20 | -0.88954 | 1.95E-21 | -9.50745 | 0.539787 | 19.69894 |
| CNAG_056 | PTP1 | Polyol trar | 26.43583121 | 18.68628215 | 16.31848351 | 155.0767823 | 142.3413916 | 128.2533732 | 2.26E-20 | 2.808519 | 2.21E-21 | 9.494561 | 7.005648 | 19.64573 |
| CNAG_039 | - | NAD(P)H:c | 6866.125454 | 7044.110133 | 7500.189704 | 4836.665213 | 4412.520006 | 4888.233219 | 2.46E-20 | -0.61398 | 2.42E-21 | -9.48534 | 0.653392 | 19.60895 |
| CNAG_048 | - | hypothetic | 586.6940356 | 601.4187127 | 586.7405651 | 1119.115723 | 1165.842406 | 1012.258722 | 2.57E-20 | 0.879219 | 2.54E-21 | 9.480176 | 1.839379 | 19.5896 |
| CNAG_017 | - | hypothetic | 3855.455458 | 3544.623184 | 3630.076538 | 2286.867765 | 2332.056603 | 2485.84676 | 2.58E-20 | -0.65034 | 2.55E-21 | -9.47964 | 0.637131 | 19.58792 |
| CNAG_044 | - | hypothetic | 1419.635484 | 1512.488163 | 1447.021813 | 881.2762695 | 851.8537672 | 897.2580882 | 2.93E-20 | -0.75065 | 2.89E-21 | -9.46648 | 0.594334 | 19.53372 |
| CNAG_005 | - | prefoldin | 1342.591444 | 1394.011109 | 1500.273147 | 817.4101942 | 819.9432724 | 841.7653779 | 3.12E-20 | -0.78928 | 3.09E-21 | -9.45961 | 0.578632 | 19.50572 |
| CNAG_006 | CLF1 | pre-mRNA | 5713.441449 | 5437.678607 | 5532.640592 | 3665.274561 | 3756.184798 | 3921.268068 | 3.38E-20 | -0.57204 | 3.36E-21 | -9.45092 | 0.672665 | 19.47122 |
| CNAG_043 | - | tRNA (gua | 1925.285721 | 2089.978688 | 1886.439063 | 1210.499086 | 1214.166149 | 1248.05628 | 3.91E-20 | -0.69893 | 3.90E-21 | -9.43516 | 0.616028 | 19.40746 |
| CNAG_009 | - | homocitra | 3348.784673 | 3472.376926 | 3561.77051 | 2353.197928 | 1959.346701 | 2044.091463 | 4.02E-20 | -0.72277 | 4.01E-21 | -9.43231 | 0.605931 | 19.39619 |
| CNAG_045 | - | Nucleosid | 2271.632141 | 2212.1466 | 2360.183101 | 3544.830037 | 3508.32506 | 3437.96506 | 4.07E-20 | 0.600741 | 4.07E-21 | 9.430815 | 1.516495 | 19.39055 |
| CNAG_015 | - | ATP-depe | 6638.316242 | 6305.020161 | 6400.742876 | 4511.358396 | 4435.892094 | 4504.118499 | 4.37E-20 | -0.53965 | 4.38E-21 | -9.42299 | 0.687936 | 19.35922 |
| CNAG_006 | - | hypothetic | 3159.394173 | 2875.513857 | 2812.124369 | 1909.481346 | 1744.601957 | 1846.44392 | 4.71E-20 | -0.70135 | 4.72E-21 | -9.41516 | 0.614997 | 19.32735 |
| CNAG_016 | - | U6 snRNA | 1985.969865 | 1990.918764 | 1981.323152 | 1197.233155 | 1314.287794 | 1228.095317 | 5.12E-20 | -0.68703 | 5.15E-21 | -9.40606 | 0.62113 | 19.2908 |
| CNAG_024 | - | nuclear se | 9809.92114 | 9193.624608 | 9663.002311 | 6193.909852 | 6673.398376 | 6717.768267 | 6.09E-20 | -0.56529 | 6.14E-21 | -9.38763 | 0.675818 | 19.2153 |
| CNAG_051 | LIV12 | eukaryotic | 13244.26778 | 12028.37253 | 13347.57423 | 8471.215761 | 8827.197612 | 8861.504227 | 6.23E-20 | -0.5778 | 6.28E-21 | -9.38519 | 0.669985 | 19.20575 |
| CNAG_051 | - | NADH del | 4475.649063 | 4209.427245 | 4830.829333 | 2988.208379 | 2837.51099 | 2864.460552 | 7.11E-20 | -0.65283 | 7.18E-21 | -9.37103 | 0.636033 | 19.14796 |

| | | | | | | | | | | | | | | |
|---|---|---|---|---|---|---|---|---|---|---|---|---|---|---|
| CNAG_031 | - | Sugar trar | 3701.323199 | 3430.412306 | 3439.196305 | 2293.932477 | 2392.49715 | 2236.115967 | 8.31E-20 | -0.62657 | 8.42E-21 | -9.35431 | 0.647714 | 19.08027 |
| CNAG_013 | - | hypothetic | 73.14943183 | 86.50480578 | 79.20149328 | 263.3890803 | 286.3783364 | 242.7118619 | 8.49E-20 | 1.717743 | 8.61E-21 | 9.351914 | 3.289213 | 19.07097 |
| CNAG_022 | - | hypothetic | 640.7256446 | 710.8261421 | 837.8184661 | 330.8158672 | 343.7335793 | 260.2533759 | 8.80E-20 | -1.24441 | 8.93E-21 | -9.348 | 0.422081 | 19.05542 |
| CNAG_017 | - | hypothetic | 30.0163405 | 48.80739476 | 34.4448828 | 207.2240736 | 178.424678 | 168.2541825 | 9.14E-20 | 2.289866 | 9.29E-21 | 9.343861 | 4.890107 | 19.03895 |
| CNAG_031 | - | gamma-gl | 1942.506599 | 1738.802195 | 1679.543947 | 1018.965936 | 1013.999333 | 1095.971245 | 9.42E-20 | -0.79319 | 9.59E-21 | -9.34045 | 0.577067 | 19.026 |
| CNAG_043 | - | protein KF | 1186.071818 | 1239.911491 | 1234.773705 | 720.9294746 | 705.2175515 | 729.6377756 | 9.78E-20 | -0.77947 | 9.97E-21 | -9.33638 | 0.582579 | 19.00983 |
| CNAG_047 | - | GTPase ac | 2563.551953 | 2555.826496 | 2197.093882 | 3778.200653 | 4045.76723 | 4125.044265 | 1.00E-19 | 0.692763 | 1.02E-20 | 9.333844 | 1.616376 | 18.99995 |
| CNAG_043 | - | dihydrodip | 231.9487602 | 299.7836392 | 327.5907108 | 649.9033478 | 679.4272081 | 654.5828004 | 1.04E-19 | 1.194929 | 1.06E-20 | 9.329504 | 2.289336 | 18.98321 |
| CNAG_06( | HEX1 | beta-hexo | 618.2255638 | 646.662327 | 539.8650475 | 1178.258831 | 1071.566189 | 1299.488103 | 1.04E-19 | 0.961999 | 1.07E-20 | 9.329335 | 1.948008 | 18.98303 |
| CNAG_035 | - | Prp8 bind | 2684.346438 | 2668.659948 | 2883.502767 | 1835.579388 | 1746.384237 | 1677.725062 | 1.06E-19 | -0.66265 | 1.09E-20 | -9.3267 | 0.631717 | 18.97275 |
| CNAG_058 | - | minichron | 4556.26821 | 4612.337562 | 4633.606772 | 3079.25375 | 3283.564118 | 2922.20332 | 1.17E-19 | -0.58733 | 1.20E-20 | -9.31644 | 0.665572 | 18.93177 |
| CNAG_033 | - | protein LT | 2840.830143 | 2864.741187 | 3200.197184 | 1650.880272 | 1858.52004 | 1903.036999 | 1.18E-19 | -0.73394 | 1.21E-20 | -9.31565 | 0.60126 | 18.92907 |
| CNAG_00( | - | high-affini | 725.7560201 | 767.8742614 | 654.661498 | 346.4005272 | 368.7789629 | 352.2663535 | 1.19E-19 | -1.02436 | 1.23E-20 | -9.31413 | 0.491628 | 18.92335 |
| CNAG_076 | - | hypothetic | 21.76229714 | 15.44806565 | 13.11340389 | 121.0842809 | 145.2239093 | 118.2255332 | 1.39E-19 | 2.954214 | 1.43E-20 | 9.297995 | 7.750093 | 18.85793 |
| CNAG_003 | - | hypothetic | 347.3598937 | 345.0294093 | 371.3146499 | 750.3795707 | 697.3550765 | 699.0732723 | 1.45E-19 | 0.998225 | 1.50E-20 | 9.293141 | 1.997542 | 18.83862 |
| CNAG_04( | - | signal pep | 1102.742147 | 1126.083078 | 1081.544559 | 1771.885337 | 1795.320799 | 1774.898569 | 1.80E-19 | 0.675306 | 1.86E-20 | 9.270084 | 1.596936 | 18.74565 |
| CNAG_029 | - | voltage-de | 18930.27045 | 20514.16529 | 20580.28747 | 14057.85201 | 14432.14828 | 14524.41014 | 1.89E-19 | -0.4957 | 1.96E-20 | -9.26453 | 0.709217 | 18.72358 |
| CNAG_063 | - | ribosome | 2173.958251 | 2105.151814 | 2422.724323 | 1128.559795 | 1315.072097 | 1376.832257 | 1.96E-19 | -0.82652 | 2.04E-20 | -9.26036 | 0.563889 | 18.7071 |
| CNAG_052 | - | U6 snRNA | 1673.04927 | 1709.707048 | 1720.031151 | 1064.200759 | 1094.306637 | 1048.251533 | 2.02E-19 | -0.68552 | 2.10E-20 | -9.25728 | 0.621782 | 18.69509 |
| CNAG_06( | - | tuftelin-in | 3683.859861 | 3458.435783 | 3561.825207 | 2293.123982 | 2398.518118 | 2433.672688 | 2.15E-19 | -0.60265 | 2.24E-20 | -9.25025 | 0.658544 | 18.66803 |
| CNAG_015 | - | solute car | 1210.456458 | 1131.141115 | 1230.472865 | 617.2674235 | 707.8463917 | 679.7007237 | 2.19E-19 | -0.84984 | 2.29E-20 | -9.2479 | 0.554846 | 18.65901 |
| CNAG_044 | - | splicing fa | 8026.874969 | 8078.465341 | 8157.188458 | 5632.846341 | 5889.354516 | 5855.164517 | 2.31E-19 | -0.49681 | 2.42E-20 | -9.24189 | 0.708673 | 18.63562 |
| CNAG_036 | - | U3 small r | 6866.112182 | 6964.384794 | 6850.791096 | 4705.878166 | 4920.682393 | 5008.104184 | 2.55E-19 | -0.51418 | 2.67E-20 | -9.23144 | 0.700191 | 18.59422 |
| CNAG_01( | LIV8 | infection r | 129.1022753 | 142.4906899 | 111.2019567 | 339.4363673 | 370.7685753 | 343.6953762 | 2.92E-19 | 1.446628 | 3.07E-20 | 9.216474 | 2.725702 | 18.53459 |
| CNAG_047 | - | oligosacch | 4030.192821 | 3915.53571 | 3811.642959 | 5812.716439 | 5637.059807 | 5752.647939 | 3.07E-19 | 0.533809 | 3.23E-20 | 9.210971 | 1.447746 | 18.51333 |
| CNAG_05( | - | NADH-ubi | 8803.293397 | 8544.966045 | 8307.56577 | 5279.359526 | 6130.022985 | 5730.867453 | 3.41E-19 | -0.5974 | 3.60E-20 | -9.19925 | 0.660946 | 18.46742 |
| CNAG_01( | - | Cystathior | 2223.17131 | 2631.890021 | 2452.670187 | 1359.815345 | 1418.554641 | 1578.854883 | 3.47E-19 | -0.7608 | 3.67E-20 | -9.19722 | 0.59017 | 18.45974 |
| CNAG_011 | - | hypothetic | 88.33938943 | 105.8977885 | 113.2934688 | 307.6915169 | 335.5118844 | 282.709643 | 3.53E-19 | 1.579271 | 3.74E-20 | 9.195265 | 2.988189 | 18.45233 |
| CNAG_033 | - | hypothetic | 1702.264908 | 1835.721934 | 1720.054739 | 1132.743911 | 1089.10007 | 1063.803418 | 3.61E-19 | -0.69369 | 3.84E-20 | -9.19258 | 0.618271 | 18.44197 |
| CNAG_035 | CKB1 | casein kin | 2707.761845 | 2901.340542 | 2873.941307 | 1792.675089 | 1804.170354 | 1941.858871 | 3.74E-19 | -0.63002 | 3.98E-20 | -9.1887 | 0.646167 | 18.42684 |
| CNAG_001 | - | RuvB-like | 4466.257255 | 4076.909271 | 4446.957921 | 2863.653768 | 2910.015431 | 2891.077379 | 3.83E-19 | -0.5998 | 4.07E-20 | -9.1861 | 0.659845 | 18.41682 |
| CNAG_043 | - | hexapreny | 3759.661096 | 3295.764837 | 3601.253248 | 2122.601225 | 2311.387457 | 2298.259119 | 4.05E-19 | -0.67855 | 4.32E-20 | -9.17981 | 0.624791 | 18.39243 |
| CNAG_038 | - | chaperoni | 4097.19265 | 3670.711845 | 4474.57652 | 2630.209219 | 2215.507705 | 2291.679876 | 4.16E-19 | -0.79436 | 4.44E-20 | -9.17672 | 0.576599 | 18.3805 |
| CNAG_065 | - | hypothetic | 483.9663337 | 441.5440023 | 526.4840468 | 211.7093831 | 218.6389373 | 194.6961711 | 4.47E-19 | -1.23461 | 4.77E-20 | -9.16904 | 0.424956 | 18.35002 |
| CNAG_038 | - | hypothetic | 299.5196903 | 295.487254 | 329.7289992 | 704.5846363 | 676.789788 | 595.9140569 | 4.49E-19 | 1.081948 | 4.80E-20 | 9.168416 | 2.116892 | 18.34801 |
| CNAG_015 | - | wd-repeat | 2308.31549 | 2318.477685 | 2369.50623 | 1583.431318 | 1489.34051 | 1452.351766 | 4.69E-19 | -0.64399 | 5.02E-20 | -9.16352 | 0.639942 | 18.32879 |
| CNAG_045 | - | DNA-direc | 5999.55748 | 5643.48626 | 6081.832011 | 3803.779685 | 3962.333551 | 4169.940031 | 5.88E-19 | -0.58609 | 6.31E-20 | -9.13884 | 0.666148 | 18.23057 |
| CNAG_058 | - | D-asparta | 143.2239351 | 194.2009577 | 193.2554274 | 493.1113183 | 442.1615855 | 429.2549635 | 6.68E-19 | 1.349396 | 7.18E-20 | 9.124835 | 2.548054 | 18.17534 |
| CNAG_032 | - | hypothetic | 273.7995222 | 279.2992036 | 272.1639892 | 585.2918575 | 551.5571912 | 606.8060927 | 7.79E-19 | 1.065018 | 8.39E-20 | 9.108066 | 2.092196 | 18.10866 |
| CNAG_064 | - | RNA-bind | 2707.715105 | 2897.008779 | 2832.342418 | 1636.918769 | 1698.040548 | 1931.852184 | 9.58E-19 | -0.69494 | 1.04E-19 | -9.08514 | 0.617736 | 18.01849 |
| CNAG_049 | - | CBS doma | 797.9942225 | 829.7966736 | 792.5440305 | 1371.940598 | 1359.747452 | 1323.069863 | 9.61E-19 | 0.73009 | 1.04E-19 | 9.084689 | 1.658743 | 18.01719 |
| CNAG_032 | - | ATP-depe | 3340.475471 | 3004.849274 | 3311.162011 | 1905.45045 | 1975.798953 | 2162.785567 | 9.98E-19 | -0.69193 | 1.08E-19 | -9.08032 | 0.619024 | 18.00073 |
| CNAG_015 | - | hypothetic | 812.281106 | 810.0129597 | 892.3442108 | 426.6164428 | 372.1193835 | 467.6467907 | 1.11E-18 | -1.00594 | 1.21E-19 | -9.06818 | 0.497946 | 17.9533 |
| CNAG_019 | - | zinc metal | 168.8456704 | 208.1946231 | 199.6548534 | 471.8272424 | 485.4146688 | 425.9851077 | 1.25E-18 | 1.249321 | 1.36E-19 | 9.055354 | 2.377295 | 17.90319 |
| CNAG_02( | - | DNA-direc | 2176.283525 | 2151.456435 | 2064.537096 | 1423.083277 | 1307.306954 | 1361.295982 | 1.27E-18 | -0.65875 | 1.38E-19 | -9.05373 | 0.633426 | 17.89722 |
| CNAG_041 | AKP3 | mucin-des | 1394.667663 | 1134.393467 | 1178.276602 | 679.6235424 | 649.9616517 | 674.1717635 | 1.37E-18 | -0.90444 | 1.49E-19 | -9.04537 | 0.53424 | 17.86444 |
| CNAG_024 | - | Flavin-con | 91.80344168 | 76.86446966 | 116.4732573 | 286.5779765 | 309.5447845 | 301.4137247 | 1.58E-18 | 1.646635 | 1.73E-19 | 9.029302 | 3.131025 | 17.80161 |
| CNAG_074 | - | nuclear pc | 6973.505879 | 7025.786467 | 7009.670899 | 4389.790443 | 4707.587594 | 5082.468663 | 1.60E-18 | -0.58236 | 1.75E-19 | -9.02779 | 0.66787 | 17.79611 |
| CNAG_07( | - | NAD-bind | 181.6603022 | 221.1092577 | 192.207812 | 458.431023 | 443.7875522 | 507.8897683 | 1.89E-18 | 1.23076 | 2.08E-19 | 9.009169 | 2.346906 | 17.72373 |
| CNAG_045 | HOB7 | LIM-home | 548.1156085 | 566.9160824 | 506.7701276 | 964.1132623 | 989.7718017 | 972.1963054 | 2.27E-18 | 0.837399 | 2.50E-19 | 8.988574 | 1.786825 | 17.64324 |
| CNAG_01( | - | histone-ly | 8450.629696 | 7637.83334 | 8327.791448 | 5344.818063 | 5571.026134 | 5700.862464 | 2.28E-18 | -0.57098 | 2.51E-19 | -8.98821 | 0.673159 | 17.6423 |
| CNAG_041 | - | hypothetic | 2243.976015 | 2100.854 | 2196.736857 | 1324.885833 | 1302.978623 | 1462.274538 | 2.35E-18 | -0.69308 | 2.60E-19 | -8.98461 | 0.618531 | 17.62856 |
| CNAG_029 | - | hypothetic | 1646.150928 | 1628.912965 | 1668.837269 | 1074.388463 | 1032.179914 | 972.7887945 | 2.46E-18 | -0.69846 | 2.72E-19 | -8.97944 | 0.616228 | 17.60861 |
| CNAG_02( | - | 3-demethy | 1599.245753 | 1376.844677 | 1533.371248 | 855.5215302 | 888.9635089 | 899.4916772 | 2.62E-18 | -0.78678 | 2.90E-19 | -8.97251 | 0.579636 | 17.58174 |
| CNAG_03( | - | multiple R | 2829.091464 | 2615.908511 | 3010.395101 | 1806.715038 | 1787.784562 | 1803.138866 | 2.63E-18 | -0.66345 | 2.91E-19 | -8.97202 | 0.631367 | 17.58028 |
| CNAG_05( | - | hypothetic | 1144.083352 | 1230.210363 | 1256.076487 | 664.018353 | 711.2735252 | 741.8283663 | 2.83E-18 | -0.79362 | 3.15E-19 | -8.96348 | 0.576894 | 17.54807 |
| CNAG_011 | - | ran GTPas | 4357.686576 | 4096.289 | 4324.330553 | 2756.9853 | 3006.667613 | 2882.189905 | 3.53E-18 | -0.57942 | 3.94E-19 | -8.9386 | 0.669231 | 17.45161 |
| CNAG_051 | - | hypothetic | 1564.563197 | 1842.141123 | 1798.898976 | 1099.291113 | 986.4012917 | 930.6482335 | 3.59E-18 | -0.80209 | 4.00E-19 | -8.93684 | 0.573516 | 17.44515 |
| CNAG_04( | - | hypothetic | 931.1512288 | 964.493827 | 996.2177706 | 1691.895624 | 1504.514662 | 1699.316915 | 3.65E-18 | 0.744477 | 4.08E-19 | 8.934841 | 1.675367 | 17.43826 |
| CNAG_03( | - | hypothetic | 1282.536895 | 1227.378344 | 1029.353467 | 2174.527474 | 2016.998402 | 1995.819815 | 3.72E-18 | 0.790648 | 4.17E-19 | 8.932313 | 1.729851 | 17.42928 |
| CNAG_04( | - | small nucl | 5249.947034 | 5143.558228 | 5763.994887 | 3525.955495 | 3667.352791 | 3699.255157 | 3.98E-18 | -0.58433 | 4.48E-19 | -8.92454 | 0.666959 | 17.3997 |
| CNAG_068 | - | HpcH/Hpa | 512.2094452 | 599.7540848 | 533.019451 | 272.5233843 | 260.9176657 | 241.2967636 | 4.13E-18 | -1.10328 | 4.65E-19 | -8.92032 | 0.465457 | 17.38364 |
| CNAG_025 | - | D-amino-a | 520.131352 | 507.1821107 | 495.6747349 | 213.993737 | 205.6246345 | 260.0713819 | 4.22E-18 | -1.18167 | 4.75E-19 | -8.91796 | 0.44084 | 17.37486 |
| CNAG_034 | - | arginine-t | 7718.448855 | 7159.429747 | 7427.72469 | 4723.692386 | 5243.284745 | 5199.070774 | 4.28E-18 | -0.57222 | 4.83E-19 | -8.91619 | 0.672583 | 17.36839 |
| CNAG_053 | - | 3-dehydrc | 180.460548 | 181.2969079 | 208.1631308 | 419.399138 | 456.0160347 | 459.1291239 | 5.21E-18 | 1.213898 | 5.89E-19 | 8.8941 | 2.319636 | 17.2833 |
| CNAG_024 | - | hypothetic | 1807.355702 | 1831.472304 | 1846.950373 | 1196.560348 | 1199.545954 | 1160.365356 | 5.22E-18 | -0.64062 | 5.91E-19 | -8.89367 | 0.641436 | 17.28208 |
| CNAG_02( | - | nonsense- | 6045.063022 | 5392.476745 | 6458.227236 | 3750.07034 | 3772.531304 | 3964.594816 | 5.68E-18 | -0.65555 | 6.44E-19 | -8.88419 | 0.634832 | 17.24552 |
| CNAG_043 | - | peptidase, | 363.6518524 | 351.4798261 | 338.2824255 | 753.55898 | 653.2748018 | 700.1574823 | 5.69E-18 | 0.986208 | 6.45E-19 | 8.883923 | 1.980972 | 17.24493 |

| | | | | | | | | | | | | | | |
|---|---|---|---|---|---|---|---|---|---|---|---|---|---|---|
| CNAG_074 | - | valine-tRN | 10656.54441 | 10048.02418 | 10107.74545 | 6888.957578 | 7324.641433 | 7463.778169 | 6.02E-18 | -0.52292 | 6.85E-19 | -8.87737 | 0.695964 | 17.22028 |
| CNAG_039 | - | elongator | 3943.086803 | 3630.837051 | 3954.243289 | 2459.690887 | 2323.386347 | 2653.463774 | 6.11E-18 | -0.64811 | 6.96E-19 | -8.87549 | 0.638118 | 17.21386 |
| CNAG_123 | - | unspecifie | 21.80118933 | 32.64599417 | 18.44914462 | 153.4334712 | 134.4313619 | 132.6586016 | 7.56E-18 | 2.547164 | 8.63E-19 | 8.851506 | 5.84484 | 17.12134 |
| CNAG_035 | - | 2-oxogluta | 10021.11107 | 8452.377306 | 9039.136912 | 5954.999121 | 6110.974419 | 6091.673013 | 7.67E-18 | -0.61575 | 8.77E-19 | -8.84979 | 0.65259 | 17.11511 |
| CNAG_020 | - | Acyl-CoA- | 68.54515754 | 103.7253833 | 104.7605267 | 281.9649708 | 288.7651425 | 294.7339648 | 7.96E-18 | 1.635711 | 9.11E-19 | 8.845578 | 3.107407 | 17.0992 |
| CNAG_029 | - | transcripti | 2559.277292 | 2362.686833 | 2486.821759 | 1494.53078 | 1653.275855 | 1616.615784 | 8.60E-18 | -0.6527 | 9.87E-19 | -8.83655 | 0.63609 | 17.06549 |
| CNAG_067 | - | hypothetic | 1176.412855 | 944.8169873 | 1174.962801 | 569.8218019 | 602.563555 | 562.0796149 | 8.78E-18 | -0.94296 | 1.01E-18 | -8.83414 | 0.520163 | 17.0566 |
| CNAG_043 | - | hypothetic | 15.94475906 | 22.95700733 | 12.04310507 | 119.671983 | 128.6687806 | 110.4757473 | 8.96E-18 | 2.850737 | 1.03E-18 | 8.831717 | 7.213689 | 17.04764 |
| CNAG_067 | FBA1 | fructose-b | 25639.98352 | 23601.0732 | 24640.29924 | 18025.64401 | 17782.33509 | 17302.06205 | 9.00E-18 | -0.49192 | 1.04E-18 | -8.83109 | 0.71108 | 17.04565 |
| CNAG_000 | - | S-adenosy | 810.0184236 | 872.4569932 | 830.5861365 | 490.5830118 | 444.6778554 | 448.8220866 | 9.12E-18 | -0.87696 | 1.05E-18 | -8.82949 | 0.544512 | 17.03993 |
| CNAG_047 | - | hypothetic | 506.2763218 | 653.112972 | 650.7002934 | 1168.951439 | 1100.108709 | 1177.503528 | 9.25E-18 | 0.915008 | 1.07E-18 | 8.827763 | 1.88558 | 17.03367 |
| CNAG_055 | - | hypothetic | 1463.999226 | 1416.688188 | 1617.576299 | 897.6093327 | 931.2669059 | 865.1081556 | 9.51E-18 | -0.75505 | 1.10E-18 | -8.82456 | 0.592527 | 17.02169 |
| CNAG_057 | SPR1 | endogluca | 125.5161516 | 104.826481 | 98.39777611 | 313.1594215 | 299.0103085 | 308.1261895 | 9.57E-18 | 1.476731 | 1.11E-18 | 8.823752 | 2.783174 | 17.01902 |
| CNAG_045 | - | hypothetic | 311.3507172 | 435.5001098 | 440.627255 | 803.0455532 | 879.5172317 | 864.3898824 | 9.68E-18 | 1.087861 | 1.12E-18 | 8.822327 | 2.125587 | 17.01395 |
| CNAG_024 | - | NADH del | 3368.674598 | 3391.616724 | 3358.149165 | 2357.050598 | 2389.929178 | 2284.930815 | 1.02E-17 | -0.54045 | 1.18E-18 | -8.81643 | 0.687555 | 16.99291 |
| CNAG_049 | PRO2 | glutamate | 3531.022877 | 3588.792233 | 3648.207425 | 2484.674193 | 2566.763949 | 2431.472194 | 1.04E-17 | -0.5406 | 1.21E-18 | -8.81356 | 0.687485 | 16.98224 |
| CNAG_017 | - | hypothetic | 4843.505027 | 4667.319798 | 4673.09689 | 3280.858129 | 3356.855137 | 3370.631678 | 1.12E-17 | -0.5187 | 1.30E-18 | -8.80564 | 0.698002 | 16.95201 |
| CNAG_047 | - | hypothetic | 539.9594726 | 584.141082 | 472.6687362 | 968.7893075 | 983.7172784 | 1010.996842 | 1.20E-17 | 0.877815 | 1.40E-18 | 8.797216 | 1.837591 | 16.92032 |
| CNAG_001 | - | hypothetic | 45.17155466 | 54.21499827 | 68.50585595 | 197.1019216 | 219.9244008 | 210.3453314 | 1.25E-17 | 1.891196 | 1.46E-18 | 8.792424 | 3.709426 | 16.90224 |
| CNAG_050 | - | GTP-bindi | 2033.790813 | 1896.150043 | 1967.463444 | 1296.200562 | 1220.198723 | 1263.597276 | 1.25E-17 | -0.65722 | 1.47E-18 | -8.79226 | 0.6341 | 16.90207 |
| CNAG_068 | - | 5-oxoproli | 24.12891189 | 27.28493859 | 25.89895306 | 167.7210481 | 119.5047935 | 140.371388 | 1.28E-17 | 2.47216 | 1.50E-18 | 8.789949 | 5.54874 | 16.89357 |
| CNAG_015 | - | RNA polyr | 2655.096509 | 2526.455303 | 2709.693009 | 1675.944534 | 1624.730748 | 1808.635567 | 1.31E-17 | -0.64293 | 1.54E-18 | -8.78687 | 0.64041 | 16.88212 |
| CNAG_027 | - | nuclear G | 2268.404008 | 1930.66959 | 2225.492541 | 1255.584551 | 1362.565466 | 1271.411393 | 1.32E-17 | -0.74036 | 1.55E-18 | -8.78628 | 0.598588 | 16.8803 |
| CNAG_002 | - | hypothetic | 138.4337608 | 130.6813797 | 156.9866082 | 380.9099713 | 345.5094819 | 357.0335933 | 1.32E-17 | 1.336107 | 1.55E-18 | 8.785668 | 2.52469 | 16.87839 |
| CNAG_063 | - | phosphori | 5802.25137 | 5783.517654 | 5701.151642 | 3784.334055 | 4257.419698 | 3959.042505 | 1.47E-17 | -0.54196 | 1.74E-18 | -8.77322 | 0.686838 | 16.83124 |
| CNAG_076 | - | hypothetic | 1254.484943 | 1102.427776 | 1177.520513 | 1982.13669 | 1891.89011 | 1905.90085 | 1.59E-17 | 0.693944 | 1.88E-18 | 8.764328 | 1.6177 | 16.79828 |
| CNAG_012 | - | hypothetic | 9563.502286 | 8759.430542 | 8960.241837 | 6320.082905 | 6431.017782 | 6506.83406 | 1.68E-17 | -0.5182 | 1.99E-18 | -8.75787 | 0.698243 | 16.77383 |
| CNAG_066 | - | biotin syn | 5974.892183 | 5062.786413 | 5721.364874 | 3682.422348 | 3682.858628 | 3588.282906 | 1.90E-17 | -0.62967 | 2.25E-18 | -8.74405 | 0.646324 | 16.72154 |
| CNAG_070 | - | hypothetic | 1320.165654 | 1202.252141 | 1199.622461 | 760.6723425 | 734.5630857 | 714.1245497 | 2.10E-17 | -0.76855 | 2.49E-18 | -8.73275 | 0.587007 | 16.67856 |
| CNAG_073 | - | hypothetic | 1949.503352 | 1621.428397 | 1744.565557 | 1073.521434 | 1046.804567 | 1051.595146 | 2.20E-17 | -0.76134 | 2.62E-18 | -8.7267 | 0.58995 | 16.65668 |
| CNAG_030 | - | hypothetic | 2133.114842 | 2995.522783 | 2128.958052 | 7413.036588 | 6491.324226 | 6818.081549 | 2.37E-17 | 1.500761 | 2.82E-18 | 8.718384 | 2.829919 | 16.62522 |
| CNAG_066 | - | malate de | 4743.927538 | 4298.763262 | 3749.595531 | 2490.715626 | 2734.901443 | 2623.566699 | 2.52E-17 | -0.7204 | 3.01E-18 | -8.711 | 0.60693 | 16.59826 |
| CNAG_000 | SSS1 | protein tra | 3028.885583 | 3127.63407 | 3088.328781 | 2189.751964 | 2119.056924 | 2069.563999 | 2.66E-17 | -0.55061 | 3.18E-18 | -8.70472 | 0.682732 | 16.57462 |
| CNAG_021 | - | hypothetic | 1600.724571 | 1662.320557 | 1826.61547 | 1033.035836 | 1082.227156 | 1043.808576 | 2.79E-17 | -0.70338 | 3.34E-18 | -8.69913 | 0.614134 | 16.55413 |
| CNAG_019 | - | hypothetic | 3339.465668 | 3301.122894 | 3539.387984 | 2059.559364 | 2327.813923 | 2310.442335 | 2.82E-17 | -0.61966 | 3.38E-18 | -8.69781 | 0.650822 | 16.54953 |
| CNAG_024 | - | hypothetic | 982.8913581 | 987.8362178 | 1128.040429 | 596.3980174 | 549.9053413 | 590.8851723 | 2.92E-17 | -0.85122 | 3.51E-18 | -8.69371 | 0.554315 | 16.53471 |
| CNAG_068 | ERG1 | Squalene | 2053.497617 | 1833.640588 | 1833.105374 | 1086.657748 | 1124.428665 | 1248.022629 | 3.21E-17 | -0.7414 | 3.87E-18 | -8.68261 | 0.59816 | 16.49319 |
| CNAG_011 | - | small nucl | 2843.106531 | 2610.528478 | 3155.393587 | 1860.454255 | 1768.79257 | 1696.612715 | 3.67E-17 | -0.70901 | 4.43E-18 | -8.66725 | 0.611739 | 16.43588 |
| CNAG_026 | APH3 | Phytase | 200.2903477 | 199.6057082 | 229.4833237 | 499.1212784 | 468.8720978 | 444.8536067 | 3.71E-17 | 1.152478 | 4.49E-18 | 8.665736 | 2.222953 | 16.43054 |
| CNAG_074 | - | transketol | 23742.4094 | 22771.42278 | 22322.92195 | 15980.20438 | 17174.26401 | 16882.38563 | 4.51E-17 | -0.47564 | 5.46E-18 | -8.64329 | 0.71915 | 16.34593 |
| CNAG_044 | - | small nucl | 2099.302356 | 2101.92354 | 2118.909899 | 1386.451504 | 1431.597193 | 1411.229848 | 4.69E-17 | -0.59477 | 5.69E-18 | -8.63867 | 0.66215 | 16.32925 |
| CNAG_005 | - | hypothetic | 8457.78856 | 8438.341907 | 9125.482996 | 6373.112298 | 5968.66205 | 5798.603019 | 4.78E-17 | -0.5359 | 5.80E-18 | -8.63639 | 0.689727 | 16.32102 |
| CNAG_030 | - | multiple d | 2257.993611 | 2361.481393 | 2093.338113 | 1211.121796 | 1290.887367 | 1505.553159 | 4.92E-17 | -0.75911 | 5.99E-18 | -8.63266 | 0.590861 | 16.30774 |
| CNAG_008 | DAK101 | dihydroxy | 8.956947459 | 21.88878515 | 15.24048187 | 93.97059192 | 161.1145412 | 120.3702569 | 5.08E-17 | 3.073552 | 6.20E-18 | 8.628767 | 8.418436 | 16.29383 |
| CNAG_023 | - | protein ar | 1377.625909 | 1356.353702 | 1615.403719 | 844.6586329 | 874.3181135 | 829.5828901 | 5.39E-17 | -0.78715 | 6.58E-18 | -8.62191 | 0.579489 | 16.26822 |
| CNAG_039 | - | acetate nc | 1014.237202 | 923.2064492 | 988.4120451 | 586.3306917 | 541.3086991 | 540.9526278 | 5.56E-17 | -0.82681 | 6.80E-18 | -8.61825 | 0.563775 | 16.25481 |
| CNAG_059 | - | MFS trans | 181.6148759 | 218.9310031 | 159.1786643 | 439.0350771 | 441.3100361 | 441.4113559 | 5.70E-17 | 1.22893 | 6.98E-18 | 8.615279 | 2.343931 | 16.24396 |
| CNAG_079 | - | WD repea | 1599.420314 | 1539.503765 | 1600.587305 | 941.159126 | 1007.178998 | 1033.773526 | 5.74E-17 | -0.6839 | 7.03E-18 | -8.61439 | 0.622478 | 16.24103 |
| CNAG_034 | - | hypothetic | 1110.340652 | 1334.667246 | 1391.44764 | 699.0671258 | 762.198686 | 634.2674703 | 6.00E-17 | -0.88716 | 7.35E-18 | -8.60929 | 0.540677 | 16.22215 |
| CNAG_033 | - | Phosphog | 15270.98766 | 14394.66597 | 15705.76015 | 21432.12487 | 21126.03597 | 22817.99088 | 6.45E-17 | 0.511533 | 7.92E-18 | 8.600733 | 1.425564 | 16.19016 |
| CNAG_020 | - | cytoplasm | 1763.7481 | 2096.791467 | 1735.301673 | 2953.305292 | 3095.2466 | 2917.285942 | 6.47E-17 | 0.665985 | 7.95E-18 | 8.600374 | 1.586652 | 16.18924 |
| CNAG_038 | - | hypothetic | 142.0683407 | 215.7154282 | 189.0013583 | 452.2864332 | 426.5354047 | 501.1828188 | 6.52E-17 | 1.322982 | 8.02E-18 | 8.599347 | 2.501827 | 16.18578 |
| CNAG_078 | - | hypothetic | 116.3134929 | 143.5634692 | 141.0045317 | 341.8121131 | 325.6609227 | 360.2601201 | 6.78E-17 | 1.34442 | 8.35E-18 | 8.59473 | 2.539281 | 16.16875 |
| CNAG_067 | - | cysteine-t | 3383.870348 | 3358.236835 | 3533.015059 | 2343.011728 | 2451.185495 | 2284.940219 | 6.82E-17 | -0.55303 | 8.40E-18 | -8.59397 | 0.681588 | 16.1663 |
| CNAG_018 | - | hypothetic | 76.60012983 | 84.33869994 | 77.06029507 | 266.7591177 | 241.2792356 | 223.8711386 | 7.18E-17 | 1.608067 | 8.86E-18 | 8.587841 | 3.04843 | 16.14399 |
| CNAG_003 | HAM1 | rdgB/HAM | 1501.405434 | 1576.085831 | 1594.167915 | 895.9885537 | 903.5982422 | 1040.406576 | 7.32E-17 | -0.73378 | 9.05E-18 | -8.58542 | 0.601327 | 16.1357 |
| CNAG_033 | - | DNA supe | 290.1958452 | 300.8644011 | 344.6389614 | 671.7642941 | 585.9677604 | 684.5035306 | 7.72E-17 | 1.039518 | 9.57E-18 | 8.578949 | 2.055541 | 16.11212 |
| CNAG_047 | - | Gluconola | 604.1196957 | 594.9314125 | 551.54914 | 1007.675582 | 1007.823486 | 1024.372921 | 7.99E-17 | 0.781403 | 9.91E-18 | 8.574965 | 1.718802 | 16.09751 |
| CNAG_009 | - | carboxype | 1635.435791 | 1583.590806 | 1353.258779 | 878.1026198 | 922.6205575 | 911.7069526 | 8.94E-17 | -0.7689 | 1.11E-17 | -8.56182 | 0.586863 | 16.0488 |
| CNAG_016 | CIG1 | cytokine i | 22.96696323 | 27.28405327 | 33.33796254 | 143.7923867 | 142.2801729 | 132.6840039 | 8.95E-17 | 2.331285 | 1.11E-17 | 8.561484 | 5.032535 | 16.04796 |
| CNAG_044 | - | alpha-ketc | 504.9319026 | 543.6880186 | 424.3239883 | 236.7154454 | 194.402093 | 216.8577434 | 9.31E-17 | -1.2017 | 1.16E-17 | -8.55673 | 0.434762 | 16.03092 |
| CNAG_023 | - | pre-rRNA- | 5236.800923 | 4282.729694 | 4860.737708 | 2993.565961 | 3123.917025 | 3062.06348 | 9.48E-17 | -0.66387 | 1.18E-17 | -8.55456 | 0.631183 | 16.02318 |
| CNAG_036 | - | hypothetic | 1027.181433 | 1061.03354 | 972.4774276 | 626.0366643 | 581.8618389 | 591.9976067 | 9.68E-17 | -0.78212 | 1.21E-17 | -8.55208 | 0.581512 | 16.01427 |
| CNAG_019 | - | prephenat | 1597.256101 | 1857.236679 | 1705.115112 | 1027.557807 | 1125.378615 | 1063.788857 | 9.83E-17 | -0.69695 | 1.23E-17 | -8.55005 | 0.616875 | 16.00746 |
| CNAG_004 | - | secondary | 2118.020846 | 2272.10573 | 2126.393253 | 1487.702116 | 1378.919174 | 1440.098395 | 1.01E-16 | -0.6128 | 1.27E-17 | -8.54653 | 0.653927 | 15.99463 |
| CNAG_002 | - | chaperone | 1667.284354 | 1808.806413 | 1868.223148 | 1181.058561 | 989.8184425 | 1068.24442 | 1.04E-16 | -0.73739 | 1.31E-17 | -8.54288 | 0.599825 | 15.98135 |
| CNAG_017 | - | 2-hydroxy | 221.3101633 | 232.9866539 | 246.5561094 | 521.523738 | 458.3070383 | 567.8398585 | 1.13E-16 | 1.130321 | 1.42E-17 | 8.533343 | 2.189074 | 15.94593 |
| CNAG_047 | - | hypothetic | 128.0996341 | 170.5456766 | 222.0091691 | 501.669869 | 475.0496041 | 432.6010242 | 1.17E-16 | 1.422187 | 1.47E-17 | 8.529729 | 2.679914 | 15.93278 |

| | | | | | | | | | | | | | | |
|---|---|---|---|---|---|---|---|---|---|---|---|---|---|---|
| CNAG_017 | - | cytoplasm | 5441.325506 | 4852.679443 | 5376.909924 | 3502.639203 | 3661.341389 | 3495.009826 | 1.18E-16 | -0.57189 | 1.49E-17 | -8.52822 | 0.672736 | 15.92754 |
| CNAG_062 | - | ribosomal | 1869.934251 | 1590.101284 | 1425.774569 | 927.9224579 | 933.8010561 | 897.3195797 | 1.40E-16 | -0.84079 | 1.77E-17 | -8.50809 | 0.558337 | 15.8529 |
| CNAG_058 | - | allantoate | 40.41358663 | 34.82814852 | 37.61806971 | 172.6900064 | 169.0570009 | 139.5046847 | 1.54E-16 | 2.09666 | 1.94E-17 | 8.497263 | 4.277182 | 15.81323 |
| CNAG_064 | - | isoleucine | 2434.449466 | 2444.545298 | 2410.068546 | 1600.471397 | 1659.311064 | 1700.951467 | 1.78E-16 | -0.57067 | 2.26E-17 | -8.4798 | 0.673303 | 15.75051 |
| CNAG_005 | - | nucleolar | 1579.514431 | 1524.393269 | 1493.969161 | 878.8528369 | 948.5032247 | 1015.994892 | 2.01E-16 | -0.70928 | 2.56E-17 | -8.46523 | 0.611625 | 15.697 |
| CNAG_053 | - | 3-hydroxy | 227.1837817 | 265.2878046 | 247.6455584 | 494.692955 | 526.5697352 | 654.2998704 | 2.10E-16 | 1.165225 | 2.67E-17 | 8.459969 | 2.242682 | 15.67781 |
| CNAG_038 | - | large subu | 2504.504995 | 2425.181596 | 2572.134872 | 1688.465596 | 1741.279653 | 1613.310361 | 2.15E-16 | -0.58862 | 2.75E-17 | -8.45691 | 0.664976 | 15.66686 |
| CNAG_006 | - | hypothetic | 47.55723751 | 71.44527818 | 91.94777623 | 228.9120041 | 270.8831631 | 249.2182543 | 2.18E-16 | 1.822882 | 2.79E-17 | 8.455201 | 3.537873 | 15.66089 |
| CNAG_049 | - | hypothetic | 3334.793381 | 3234.33929 | 3426.370415 | 2276.080315 | 2198.374029 | 2373.696865 | 2.27E-16 | -0.56092 | 2.90E-17 | -8.45037 | 0.677869 | 15.64334 |
| CNAG_057 | MFE2 | multifunct | 1796.343503 | 1846.860695 | 1732.072249 | 2619.284339 | 2906.360829 | 2660.867368 | 2.31E-16 | 0.591765 | 2.95E-17 | 8.448479 | 1.507089 | 15.63671 |
| CNAG_045 | - | DNA-bind | 2672.731399 | 2793.623122 | 2735.333839 | 1822.294151 | 1896.525895 | 1934.08096 | 2.63E-16 | -0.55191 | 3.37E-17 | -8.43296 | 0.682117 | 15.58031 |
| CNAG_022 | EXG104 | glucan 1,3 | 30.00369793 | 55.23373204 | 26.99042734 | 184.3582949 | 158.4309972 | 183.5774869 | 2.88E-16 | 2.228989 | 3.71E-17 | 8.421848 | 4.688052 | 15.5403 |
| CNAG_032 | ZWF1 | glucose-6 | 14130.63142 | 13650.9462 | 12934.94971 | 9777.285827 | 9848.497558 | 10062.6607 | 2.97E-16 | -0.471 | 3.83E-17 | -8.41813 | 0.721467 | 15.52692 |
| CNAG_040 | - | endoribon | 4268.884764 | 3983.143147 | 3876.472641 | 2791.321544 | 2866.929643 | 2701.249763 | 3.23E-16 | -0.55257 | 4.18E-17 | -8.40787 | 0.681806 | 15.49017 |
| CNAG_057 | ATP1 | ATP synth | 55366.40877 | 54237.86314 | 57593.6937 | 40234.88557 | 41741.71747 | 42441.6011 | 3.29E-16 | -0.44179 | 4.25E-17 | -8.40588 | 0.736223 | 15.4832 |
| CNAG_047 | - | solute car | 607.6688975 | 650.9365569 | 588.8684869 | 1107.436661 | 1067.357408 | 1004.485859 | 3.38E-16 | 0.769538 | 4.37E-17 | 8.402457 | 1.704724 | 15.47095 |
| CNAG_076 | - | hypothetic | 3938.481437 | 3659.946824 | 4023.572939 | 2748.571596 | 2612.427713 | 2540.286684 | 3.43E-16 | -0.57235 | 4.43E-17 | -8.40082 | 0.67252 | 15.46533 |
| CNAG_011 | - | DNA-direc | 4712.635945 | 4407.651587 | 4498.176913 | 2792.721099 | 2986.763308 | 3277.345917 | 3.51E-16 | -0.60407 | 4.55E-17 | -8.39775 | 0.657895 | 15.45436 |
| CNAG_078 | BOT1 | Eukaryotic | 6337.156912 | 6275.944674 | 6575.628913 | 4706.761912 | 4650.706921 | 4642.872998 | 4.03E-16 | -0.47026 | 5.23E-17 | -8.38142 | 0.721836 | 15.39484 |
| CNAG_047 | - | hypothetic | 243.5046594 | 260.9965114 | 255.1110197 | 684.0251049 | 562.8681601 | 480.5397689 | 5.07E-16 | 1.172074 | 6.60E-17 | 8.353996 | 2.253353 | 15.29504 |
| CNAG_055 | - | aldose red | 1659.65961 | 1526.899437 | 1324.745715 | 2591.591301 | 2478.466155 | 2392.184023 | 5.22E-16 | 0.711128 | 6.80E-17 | 8.350388 | 1.637084 | 15.28218 |
| CNAG_036 | - | pumilio do | 5301.1303 | 4488.5271 | 5333.143811 | 3128.191917 | 3343.882021 | 3324.041016 | 5.33E-16 | -0.64269 | 6.95E-17 | -8.34792 | 0.640516 | 15.2735 |
| CNAG_064 | - | hypothetic | 4022.626393 | 3856.040358 | 4506.621874 | 2734.464588 | 2652.930958 | 2816.676574 | 5.34E-16 | -0.60996 | 6.97E-17 | -8.34746 | 0.655215 | 15.27222 |
| CNAG_056 | - | tyrosine a | 174.5708283 | 180.1859972 | 162.3502084 | 364.0105291 | 399.8793733 | 445.6926672 | 5.36E-16 | 1.212247 | 7.01E-17 | 8.346912 | 2.316982 | 15.27062 |
| CNAG_129 | - | unspecifie | 35.77289729 | 45.55575298 | 21.65871928 | 148.0928221 | 158.6031169 | 174.6345906 | 5.45E-16 | 2.234769 | 7.13E-17 | 8.344871 | 4.706874 | 15.26393 |
| CNAG_033 | STU2 | microtubu | 2591.497911 | 2378.061195 | 2406.057343 | 3565.711308 | 3720.524228 | 3627.802298 | 5.61E-16 | 0.549572 | 7.36E-17 | 8.341154 | 1.463651 | 15.25067 |
| CNAG_059 | - | hypothetic | 694.0560722 | 680.0598574 | 725.3317495 | 1185.148298 | 1187.253166 | 1123.266125 | 5.88E-16 | 0.720163 | 7.71E-17 | 8.335559 | 1.647368 | 15.23094 |
| CNAG_025 | - | hypothetic | 2702.487618 | 2754.048867 | 2489.263271 | 3728.272994 | 4149.264034 | 4049.625749 | 6.23E-16 | 0.570867 | 8.18E-17 | 8.328544 | 1.485416 | 15.2056 |
| CNAG_009 | - | nucleolar | 5768.151293 | 4999.17929 | 5026.098547 | 3269.05307 | 3629.422412 | 3512.764479 | 6.23E-16 | -0.61725 | 8.20E-17 | -8.32837 | 0.651912 | 15.20536 |
| CNAG_010 | SOD1 | superoxid | 16079.71915 | 16879.98247 | 16774.12052 | 12641.52877 | 10857.6078 | 11262.80389 | 6.37E-16 | -0.53168 | 8.39E-17 | -8.32559 | 0.691749 | 15.19558 |
| CNAG_043 | MRPS18 | hypothetic | 6801.356792 | 6388.880448 | 7369.851866 | 2988.664829 | 2015.127296 | 2391.768486 | 6.57E-16 | -1.4906 | 8.66E-17 | -8.32188 | 0.355865 | 15.18239 |
| CNAG_031 | - | DNA cros | 438.4914961 | 547.5062697 | 519.5319186 | 890.132884 | 953.5977849 | 910.0287018 | 6.63E-16 | 0.857639 | 8.75E-17 | 8.32062 | 1.812071 | 15.17816 |
| CNAG_061 | - | ADP,ATP c | 115783.5145 | 109202.01 | 114067.7539 | 84259.70214 | 84057.22576 | 86241.89589 | 6.93E-16 | -0.42904 | 9.16E-17 | -8.31525 | 0.742754 | 15.15952 |
| CNAG_010 | - | phospholi | 2587.493688 | 2761.283618 | 2608.441421 | 1805.202206 | 1847.362137 | 1857.490008 | 6.97E-16 | -0.54519 | 9.22E-17 | -8.31442 | 0.685303 | 15.15704 |
| CNAG_036 | FRR1 | FK506-bin | 9434.048791 | 9429.587011 | 9567.054629 | 7114.806706 | 7202.19829 | 6997.519974 | 7.29E-16 | -0.431 | 9.67E-17 | -8.30873 | 0.741749 | 15.13741 |
| CNAG_008 | - | hypothetic | 8255.827225 | 8445.888901 | 9107.360503 | 6232.969372 | 6222.292073 | 6177.121221 | 7.37E-16 | -0.48533 | 9.80E-17 | -8.30723 | 0.714332 | 15.13232 |
| CNAG_024 | - | asparagin | 10382.36916 | 12333.15189 | 11386.40552 | 7643.130895 | 8106.988518 | 8014.440881 | 7.46E-16 | -0.53557 | 9.92E-17 | -8.30569 | 0.689884 | 15.12712 |
| CNAG_048 | - | mitochon | 9037.463698 | 8440.75297 | 8651.183843 | 11976.64911 | 12418.54982 | 11992.90958 | 7.72E-16 | 0.462164 | 1.03E-16 | 8.301496 | 1.377607 | 15.11257 |
| CNAG_056 | - | Nucleolar | 2024.411283 | 1855.208148 | 1880.034477 | 1296.235668 | 1183.963152 | 1233.624959 | 7.77E-16 | -0.64913 | 1.04E-16 | -8.30058 | 0.637664 | 15.10961 |
| CNAG_007 | - | ATP-depe | 13913.51564 | 13631.57779 | 14021.62316 | 10067.59356 | 10723.20062 | 9907.274104 | 9.29E-16 | -0.45278 | 1.25E-16 | -8.27846 | 0.730633 | 15.03201 |
| CNAG_124 | - | unspecifie | 679.4076676 | 774.4433557 | 895.4787845 | 406.3951267 | 397.2200785 | 411.0747239 | 9.42E-16 | -0.96661 | 1.27E-16 | -8.27673 | 0.511707 | 15.02609 |
| CNAG_043 | - | ATP-depe | 2940.220375 | 3203.043681 | 3212.015372 | 2122.745022 | 2225.189639 | 2123.951517 | 9.83E-16 | -0.54698 | 1.32E-16 | -8.27136 | 0.684452 | 15.00734 |
| CNAG_031 | - | peptide-m | 2129.656835 | 2041.624972 | 2263.899716 | 1441.755285 | 1363.396233 | 1425.662619 | 1.03E-15 | -0.62063 | 1.38E-16 | -8.26607 | 0.650386 | 14.98847 |
| CNAG_066 | 44986 | Macropha | 118.6985719 | 139.2990184 | 179.3561869 | 388.5836283 | 343.6783177 | 422.3336887 | 1.07E-15 | 1.388213 | 1.45E-16 | 8.260428 | 2.617542 | 14.96871 |
| CNAG_021 | - | TIGR01456 | 2510.350362 | 2445.6501 | 2513.504039 | 1721.955381 | 1755.946241 | 1666.577197 | 1.16E-15 | -0.55384 | 1.57E-16 | -8.25083 | 0.681207 | 14.93418 |
| CNAG_010 | - | H/ACA rib | 1521.197371 | 1412.407417 | 1759.359012 | 925.6338073 | 827.6338509 | 952.7618723 | 1.19E-15 | -0.8107 | 1.61E-16 | -8.2484 | 0.570105 | 14.92615 |
| CNAG_069 | XFP2 | xylulose 5 | 9098.487676 | 6563.375041 | 5627.526074 | 2093.319936 | 2072.073432 | 2093.179577 | 1.19E-15 | -1.78283 | 1.60E-16 | -8.24844 | 0.290613 | 14.92615 |
| CNAG_051 | - | nuclear ca | 1051.717137 | 1084.744848 | 990.6160952 | 619.7516178 | 603.4314089 | 656.3333776 | 1.44E-15 | -0.74959 | 1.96E-16 | -8.22462 | 0.594774 | 14.84064 |
| CNAG_067 | - | pre-rRNA- | 1186.118174 | 1186.106243 | 1308.32642 | 734.934264 | 779.4688413 | 751.8354254 | 1.64E-15 | -0.71625 | 2.22E-16 | -8.20933 | 0.608676 | 14.78646 |
| CNAG_020 | - | N-termina | 1566.702152 | 1471.639602 | 1515.291931 | 1002.00622 | 958.8578526 | 976.0662867 | 1.64E-15 | -0.64812 | 2.24E-16 | -8.20854 | 0.638113 | 14.78399 |
| CNAG_050 | CAT4 | catalase 4 | 52.05869796 | 41.29522331 | 53.58726223 | 174.6293578 | 175.7749352 | 181.4769511 | 1.66E-15 | 1.844157 | 2.26E-16 | 8.207645 | 3.590432 | 14.78115 |
| CNAG_016 | - | small subu | 2236.033513 | 2406.816628 | 2378.057496 | 1603.643354 | 1585.982664 | 1623.25038 | 1.67E-15 | -0.56007 | 2.28E-16 | -8.20634 | 0.678271 | 14.77684 |
| CNAG_045 | - | hypothetic | 64.94310806 | 56.38341406 | 84.47848283 | 213.4030321 | 243.2334095 | 213.8005191 | 1.75E-15 | 1.697378 | 2.39E-16 | 8.200637 | 3.24311 | 14.7566 |
| CNAG_062 | - | hypothetic | 2023.35549 | 1974.79149 | 1987.741706 | 1265.764877 | 1406.624786 | 1303.561483 | 1.89E-15 | -0.60599 | 2.59E-16 | -8.19113 | 0.657021 | 14.72271 |
| CNAG_024 | - | translation | 1745.300924 | 1543.855585 | 1734.931441 | 1076.71125 | 1064.114587 | 936.1857337 | 1.90E-15 | -0.72319 | 2.60E-16 | -8.19066 | 0.605756 | 14.72175 |
| CNAG_048 | - | membrane | 1137.679409 | 1021.597849 | 965.3285192 | 1626.239261 | 1775.549416 | 1721.586079 | 2.03E-15 | 0.698135 | 2.79E-16 | 8.182164 | 1.622406 | 14.69226 |
| CNAG_059 | - | Alpha-glu | 608.9052333 | 524.5588672 | 607.6182076 | 270.0892313 | 284.212509 | 314.4585191 | 2.04E-15 | -1.01832 | 2.81E-16 | -8.18119 | 0.49369 | 14.68952 |
| CNAG_007 | - | U6 snRNA | 1609.94915 | 1529.827386 | 1754.087154 | 1082.221315 | 963.1328639 | 901.7736931 | 2.04E-15 | -0.7473 | 2.81E-16 | -8.18121 | 0.595717 | 14.68952 |
| CNAG_020 | - | hypothetic | 1539.889477 | 1535.170801 | 1529.151223 | 993.4254548 | 1033.971417 | 886.2150817 | 2.08E-15 | -0.676 | 2.87E-16 | -8.17866 | 0.625899 | 14.68117 |
| CNAG_036 | - | ATP-bindi | 5272.018756 | 4947.454651 | 4932.250574 | 3305.678042 | 3595.792829 | 3659.255187 | 2.13E-15 | -0.53645 | 2.93E-16 | -8.17598 | 0.689464 | 14.67229 |
| CNAG_037 | HXS1 | high-affini | 409.4661778 | 262.2434025 | 241.3925557 | 1219.800305 | 1355.080538 | 1579.850283 | 2.13E-15 | 2.170151 | 2.93E-16 | 8.176059 | 4.500704 | 14.67229 |
| CNAG_057 | - | elongator | 967.2364326 | 903.0804629 | 923.703602 | 1413.717594 | 1565.940006 | 1525.086592 | 2.20E-15 | 0.67351 | 3.04E-16 | 8.171546 | 1.594949 | 14.65708 |
| CNAG_019 | - | hypothetic | 248.1668464 | 313.7136391 | 263.634355 | 577.5455028 | 542.9482145 | 559.1594902 | 2.28E-15 | 1.011734 | 3.16E-16 | 8.167114 | 2.016334 | 14.64228 |
| CNAG_016 | - | fumarate l | 5508.056775 | 5517.423621 | 5689.411531 | 3951.757692 | 4061.580944 | 4173.246918 | 2.54E-15 | -0.47121 | 3.53E-16 | -8.15375 | 0.721359 | 14.59539 |
| CNAG_043 | HAD1 | 3-hydroxy | 2015.141374 | 1862.772084 | 2021.83466 | 1337.490921 | 1294.42921 | 1208.127578 | 2.71E-15 | -0.6352 | 3.78E-16 | -8.14554 | 0.643853 | 14.56664 |
| CNAG_017 | - | tRNA(His) | 3743.491735 | 3635.136425 | 3756.984776 | 2606.120092 | 2363.951463 | 2670.10207 | 2.90E-15 | -0.55873 | 4.04E-16 | -8.13746 | 0.678898 | 14.53803 |
| CNAG_066 | - | myosin he | 2857.209065 | 3032.784286 | 2806.808394 | 1882.968411 | 1935.307434 | 2099.470062 | 2.97E-15 | -0.57039 | 4.14E-16 | -8.13441 | 0.673433 | 14.5275 |
| CNAG_026 | - | Cystathior | 3984.257315 | 4770.64707 | 4478.967681 | 2929.803173 | 3086.01168 | 2977.667221 | 3.34E-15 | -0.57184 | 4.66E-16 | -8.12003 | 0.672757 | 14.47674 |

| | | | | | | | | | | | | | | |
|---|---|---|---|---|---|---|---|---|---|---|---|---|---|---|
| CNAG_04( | - | hypothetic | 180.4103873 | 199.5530503 | 173.006586 | 440.7803233 | 379.0189767 | 410.358729 | 3.79E-15 | 1.140079 | 5.31E-16 | 8.104174 | 2.203931 | 14.42082 |
| CNAG_01( | KSP1 | serine/thre | 1613.274792 | 1473.7637 | 1406.549534 | 911.6264751 | 989.9714481 | 869.5521985 | 4.07E-15 | -0.71371 | 5.71E-16 | -8.09531 | 0.60975 | 14.38993 |
| CNAG_03 | - | hypothetic | 347.3174043 | 353.6186853 | 364.9043954 | 662.1840468 | 625.6503885 | 712.2947422 | 4.40E-15 | 0.894742 | 6.18E-16 | 8.085712 | 1.859277 | 14.35646 |
| CNAG_03 | UVE1 | UV damag | 1587.447516 | 1983.647828 | 1638.246531 | 2754.080761 | 2753.598758 | 2776.258117 | 4.50E-15 | 0.655286 | 6.32E-16 | 8.08289 | 1.574929 | 14.34678 |
| CNAG_03 | - | T-complex | 8755.54296 | 8451.29103 | 9041.261233 | 6087.360008 | 6660.529808 | 6215.986114 | 4.81E-15 | -0.48466 | 6.77E-16 | -8.07452 | 0.714667 | 14.31774 |
| CNAG_05 | EMC3 | hypothetic | 1621.723058 | 1675.244747 | 1759.465804 | 1043.164085 | 1009.705529 | 1161.402768 | 4.97E-15 | -0.6688 | 7.00E-16 | -8.07057 | 0.629028 | 14.30405 |
| CNAG_02 | - | hypothetic | 106.8419421 | 88.66846211 | 101.5556629 | 263.2143519 | 262.7910487 | 265.9632648 | 5.45E-15 | 1.405364 | 7.69E-16 | 8.059077 | 2.648845 | 14.26395 |
| CNAG_01( | - | glicosidas | 91.77794496 | 99.42419996 | 115.3954931 | 293.8161345 | 274.8840456 | 254.9822801 | 5.45E-15 | 1.413366 | 7.70E-16 | 8.058908 | 2.663578 | 14.26373 |
| CNAG_07 | CID1alpha | CID1-fami | 3541.53437 | 3582.332978 | 3577.840664 | 2428.593755 | 2642.697126 | 2548.003603 | 5.53E-15 | -0.50541 | 7.83E-16 | -8.05688 | 0.704462 | 14.25692 |
| CNAG_02( | - | triose-pho | 9437.548062 | 9621.349844 | 9337.784922 | 7336.678617 | 7023.622118 | 6906.496632 | 5.96E-15 | -0.43229 | 8.46E-16 | -8.04732 | 0.741083 | 14.22448 |
| CNAG_06 | - | hypothetic | 1947.389066 | 1792.733286 | 1854.437029 | 1223.009639 | 1253.901776 | 1243.596812 | 5.97E-15 | -0.60431 | 8.48E-16 | -8.04706 | 0.657784 | 14.22393 |
| CNAG_01 | - | Integral m | 407.0129454 | 561.4944622 | 483.2976409 | 920.612905 | 869.7754406 | 978.7317042 | 6.25E-15 | 0.917451 | 8.89E-16 | 8.041342 | 1.888775 | 14.20402 |
| CNAG_04 | - | oligosacch | 795.7305435 | 782.4503127 | 967.3806978 | 1524.604595 | 1439.085539 | 1411.884772 | 6.30E-15 | 0.766307 | 8.96E-16 | 8.040347 | 1.70091 | 14.20086 |
| CNAG_01 | - | hypothetic | 58.09004097 | 95.14093019 | 126.0507233 | 317.2517357 | 314.7376693 | 281.5563662 | 6.61E-15 | 1.694262 | 9.43E-16 | 8.034102 | 3.236113 | 14.17948 |
| CNAG_04 | - | alpha-solu | 2128.335562 | 1905.866697 | 1976.013493 | 1359.276594 | 1255.565617 | 1311.325956 | 6.74E-15 | -0.63006 | 9.62E-16 | -8.03163 | 0.646149 | 14.17108 |
| CNAG_12 | - | unspecifie | 7.685214814 | 3.544424564 | 2.394219689 | 92.24960819 | 57.25064414 | 90.12183543 | 6.92E-15 | 4.300493 | 9.89E-16 | 8.028269 | 19.70504 | 14.15994 |
| CNAG_12 | - | unspecifie | 10782.1402 | 9251.685624 | 12981.3543 | 3811.124495 | 2617.137425 | 3572.992498 | 7.59E-15 | -1.73884 | 1.09E-15 | -8.01663 | 0.299611 | 14.11989 |
| CNAG_02 | - | hypothetic | 29.96249168 | 42.3280763 | 13.13098703 | 191.2239637 | 155.1852779 | 128.441987 | 7.91E-15 | 2.483063 | 1.13E-15 | 8.011318 | 5.590831 | 14.10187 |
| CNAG_07 | - | hypothetic | 1564.327203 | 1427.475439 | 1522.736586 | 988.7849901 | 976.1448246 | 868.450645 | 8.07E-15 | -0.68805 | 1.16E-15 | -8.0087 | 0.620693 | 14.09299 |
| CNAG_07 | - | tricarboxyl | 43.855062 | 28.37507999 | 37.60771249 | 156.9779514 | 150.8157546 | 142.719406 | 8.66E-15 | 2.036124 | 1.25E-15 | 7.999717 | 4.101421 | 14.0624 |
| CNAG_02 | - | thymidylat | 890.539277 | 875.7592073 | 859.4116116 | 545.1144217 | 522.3833793 | 488.7791078 | 8.67E-15 | -0.77018 | 1.25E-15 | -7.99947 | 0.586342 | 14.06191 |
| CNAG_05 | - | hypothetic | 625.1275735 | 617.550634 | 543.0322358 | 995.1202006 | 1131.346943 | 992.2572479 | 9.41E-15 | 0.789343 | 1.36E-15 | 7.989174 | 1.728288 | 14.02634 |
| CNAG_05( | - | hypothetic | 67.2143546 | 57.43949931 | 62.12911416 | 189.9495966 | 243.4323643 | 183.9279559 | 9.86E-15 | 1.709171 | 1.42E-15 | 7.983389 | 3.26973 | 14.00634 |
| CNAG_05 | RIM101 | pH-respor | 1109.670746 | 1045.283799 | 1058.06582 | 1640.302536 | 1671.088983 | 1632.832359 | 1.03E-14 | 0.606432 | 1.49E-15 | 7.978165 | 1.522489 | 13.98869 |
| CNAG_04( | SEC23 | protein tra | 1568.562581 | 1470.857517 | 1432.398386 | 2308.14451 | 2217.030272 | 2213.464342 | 1.08E-14 | 0.576619 | 1.56E-15 | 7.971752 | 1.49135 | 13.96687 |
| CNAG_04( | - | L-aminoac | 204.8405804 | 170.5251347 | 188.9815818 | 436.0231454 | 428.3878783 | 381.6236772 | 1.09E-14 | 1.131556 | 1.59E-15 | 7.969895 | 2.190949 | 13.96071 |
| CNAG_02 | - | hypothetic | 69.68479852 | 82.2187989 | 116.4440779 | 274.2742309 | 291.494029 | 244.98387 | 1.12E-14 | 1.583899 | 1.63E-15 | 7.966771 | 2.997789 | 13.95046 |
| CNAG_06 | - | ABC trans | 658.9052513 | 517.4205696 | 515.315545 | 1000.642733 | 1061.393984 | 1049.873794 | 1.12E-14 | 0.863773 | 1.63E-15 | 7.966549 | 1.819791 | 13.95005 |
| CNAG_04 | - | hypothetic | 78.86995603 | 72.50104037 | 60.02247887 | 274.9773461 | 210.9830106 | 195.0884822 | 1.37E-14 | 1.677759 | 2.00E-15 | 7.94128 | 3.199307 | 13.86288 |
| CNAG_04 | - | O-methylt | 570.3850389 | 696.1641959 | 636.8359532 | 1050.39916 | 1115.701884 | 1071.037024 | 1.44E-14 | 0.752137 | 2.10E-15 | 7.935431 | 1.684286 | 13.84276 |
| CNAG_01 | ERG25 | C-4 methy | 1943.253995 | 1326.253783 | 1495.0164 | 870.3086379 | 796.5652225 | 847.3674556 | 1.46E-14 | -0.93958 | 2.14E-15 | -7.93316 | 0.521384 | 13.83519 |
| CNAG_02 | NSR1 | RNA-bind | 8164.777941 | 8630.116883 | 9323.836586 | 6007.96871 | 6277.497372 | 6373.599089 | 1.48E-14 | -0.50041 | 2.17E-15 | -7.93133 | 0.706904 | 13.82952 |
| CNAG_06 | - | monocarb | 841.4869036 | 806.8256772 | 868.9470762 | 421.1031783 | 486.1458794 | 514.2343457 | 1.50E-14 | -0.83912 | 2.20E-15 | -7.92926 | 0.558986 | 13.82264 |
| CNAG_00 | - | GTP cyclo | 3126.687171 | 2590.076979 | 2846.228863 | 1847.967649 | 1904.272223 | 1824.23876 | 1.61E-14 | -0.63514 | 2.36E-15 | -7.92096 | 0.643878 | 13.79434 |
| CNAG_04( | - | hypothetic | 1827.230066 | 1783.034149 | 2020.721569 | 1148.989604 | 1175.353669 | 1286.858467 | 1.66E-14 | -0.65693 | 2.45E-15 | -7.91637 | 0.634227 | 13.77868 |
| CNAG_07 | - | ATP-depe | 1881.004852 | 1942.467893 | 2058.083153 | 1289.947597 | 1379.020038 | 1241.416464 | 1.67E-14 | -0.60461 | 2.45E-15 | -7.91597 | 0.657648 | 13.77764 |
| CNAG_00 | - | hypothetic | 4751.284016 | 4504.655993 | 4689.087547 | 3348.610985 | 3437.966277 | 3358.41939 | 1.69E-14 | -0.47471 | 2.48E-15 | -7.9145 | 0.719613 | 13.77285 |
| CNAG_03 | - | Ribose-ph | 3413.055055 | 3454.105587 | 3394.412796 | 2242.536022 | 2560.774905 | 2361.518164 | 1.76E-14 | -0.53387 | 2.59E-15 | -7.90912 | 0.690699 | 13.75444 |
| CNAG_06 | - | mutanase | 111.4314539 | 62.86128354 | 69.62337174 | 265.7818516 | 243.7845709 | 270.2767335 | 1.79E-14 | 1.667924 | 2.64E-15 | 7.906727 | 3.17757 | 13.74653 |
| CNAG_03 | - | ribonuclea | 785.4709806 | 766.9587822 | 849.7225183 | 446.9280332 | 398.0491872 | 488.6987766 | 1.81E-14 | -0.86464 | 2.67E-15 | -7.90531 | 0.549182 | 13.74223 |
| CNAG_05 | - | beta-cater | 2282.663788 | 2270.020692 | 2344.993068 | 1476.68246 | 1642.968707 | 1591.056331 | 1.89E-14 | -0.56572 | 2.79E-15 | -7.90015 | 0.675617 | 13.72462 |
| CNAG_06 | - | hypothetic | 25.29755514 | 22.99298117 | 42.90329035 | 150.8609334 | 123.8347095 | 153.5797867 | 1.89E-14 | 2.243358 | 2.79E-15 | 7.899801 | 4.73498 | 13.72377 |
| CNAG_07 | - | hypothetic | 323.9737585 | 368.6559099 | 292.4434167 | 685.036666 | 617.9636889 | 610.310676 | 1.96E-14 | 0.945275 | 2.90E-15 | 7.894929 | 1.925556 | 13.70751 |
| CNAG_05 | - | acetyl-Co/ | 42781.29801 | 38009.56048 | 36806.44366 | 26661.43935 | 28384.08296 | 28198.26707 | 2.20E-14 | -0.5142 | 3.26E-15 | -7.88034 | 0.700181 | 13.65856 |
| CNAG_00 | - | cyclophilir | 980.5545269 | 1053.490499 | 988.4538905 | 663.4997613 | 574.0943927 | 560.9436834 | 2.28E-14 | -0.76432 | 3.39E-15 | -7.87574 | 0.588732 | 13.64292 |
| CNAG_03 | NPY1 | NAD diph | 1384.688038 | 1433.906543 | 1497.129658 | 970.125081 | 938.183625 | 900.5980166 | 2.29E-14 | -0.63469 | 3.41E-15 | -7.87488 | 0.644079 | 13.6403 |
| CNAG_00( | - | cytochron | 1264.410622 | 1304.618705 | 1390.469788 | 833.8293346 | 856.2399371 | 858.3792556 | 2.49E-14 | -0.65119 | 3.72E-15 | -7.86403 | 0.636754 | 13.60333 |
| CNAG_05( | - | TPR repea | 1853.903209 | 1595.597351 | 1836.248301 | 1078.175905 | 1077.86988 | 1170.31161 | 2.53E-14 | -0.68415 | 3.77E-15 | -7.86223 | 0.622372 | 13.59746 |
| CNAG_06 | - | Histidinol | 1474.35067 | 1418.778671 | 1286.052571 | 819.7427338 | 940.8298257 | 806.2736412 | 2.89E-14 | -0.71936 | 4.32E-15 | -7.84525 | 0.607366 | 13.53968 |
| CNAG_07 | - | hypothetic | 206.0380436 | 209.2530844 | 164.5030736 | 485.2984428 | 404.0534026 | 411.5438032 | 2.99E-14 | 1.152013 | 4.49E-15 | 7.840509 | 2.222238 | 13.52363 |
| CNAG_03 | - | hypothetic | 70.70557012 | 74.62528166 | 54.68905996 | 188.2980103 | 232.0469199 | 203.7915365 | 3.05E-14 | 1.638681 | 4.58E-15 | 7.838042 | 3.113811 | 13.51546 |
| CNAG_01( | - | tRNA pseu | 1063.577239 | 1134.361919 | 1229.398951 | 661.7313465 | 728.58362 | 709.637507 | 3.27E-14 | -0.72269 | 4.91E-15 | -7.82929 | 0.605965 | 13.48555 |
| CNAG_02( | - | cytoplasm | 5351.516901 | 5100.475021 | 5414.25878 | 3604.586382 | 3822.649868 | 3951.210739 | 3.44E-14 | -0.49522 | 5.17E-15 | -7.8227 | 0.709451 | 13.46352 |
| CNAG_07 | - | hypothetic | 49.77026738 | 65.99709146 | 46.15639659 | 213.2913723 | 174.7936445 | 168.3579303 | 3.48E-14 | 1.780261 | 5.23E-15 | 7.821194 | 3.434884 | 13.45867 |
| CNAG_04 | TOP1 | DNA topo | 8372.521265 | 8321.972859 | 7993.000525 | 6059.408621 | 6215.420045 | 6292.537788 | 3.54E-14 | -0.42625 | 5.34E-15 | -7.81873 | 0.744194 | 13.45051 |
| CNAG_01 | - | hypothetic | 1423.082175 | 1304.665587 | 1450.19841 | 913.2778349 | 858.7828237 | 893.9121931 | 3.59E-14 | -0.66449 | 5.41E-15 | -7.81691 | 0.630913 | 13.4446 |
| CNAG_02 | - | long-chair | 2024.535945 | 1944.644075 | 1972.822916 | 1350.68987 | 1391.079829 | 1343.508632 | 3.61E-14 | -0.55594 | 5.45E-15 | -7.81612 | 0.680214 | 13.44223 |
| CNAG_00 | - | Prephenat | 687.4972819 | 744.2845941 | 772.9542288 | 410.3399386 | 394.651631 | 445.41293 | 3.73E-14 | -0.83357 | 5.63E-15 | -7.81196 | 0.561137 | 13.42824 |
| CNAG_05 | - | Stomatin | 1008.029351 | 737.2358358 | 780.8575939 | 1434.049244 | 1605.706953 | 1579.400114 | 3.75E-14 | 0.85443 | 5.66E-15 | 7.811237 | 1.808045 | 13.4261 |
| CNAG_01 | - | chlorophy | 236.3942783 | 201.7724668 | 180.509952 | 386.1913446 | 544.1344637 | 587.6648928 | 3.84E-14 | 1.282183 | 5.80E-15 | 7.808223 | 2.432068 | 13.41606 |
| CNAG_06( | - | hypothetic | 177.9864805 | 151.1255827 | 152.7445736 | 332.8591655 | 372.2981181 | 401.3273347 | 4.10E-14 | 1.187306 | 6.20E-15 | 7.799876 | 2.277271 | 13.38767 |
| CNAG_01( | - | small subu | 21735.08617 | 21758.61527 | 22765.42835 | 13383.66161 | 16969.03989 | 14457.86622 | 4.88E-14 | -0.57996 | 7.41E-15 | -7.77736 | 0.668983 | 13.31169 |
| CNAG_01 | EF3 | elongation | 29563.47454 | 26060.82759 | 24966.68035 | 18959.55334 | 19431.56909 | 17605.18041 | 4.94E-14 | -0.54118 | 7.51E-15 | -7.77555 | 0.68721 | 13.30584 |
| CNAG_04 | NOC3 | nucleolar | 4649.602286 | 4165.278645 | 4609.065765 | 2855.798595 | 3166.238228 | 3180.786398 | 5.10E-14 | -0.56069 | 7.76E-15 | -7.77143 | 0.677976 | 13.29239 |
| CNAG_03 | - | hypothetic | 1432.360782 | 1372.474969 | 1330.810298 | 883.6855667 | 919.2495105 | 756.3530395 | 5.22E-14 | -0.70812 | 7.95E-15 | -7.76839 | 0.612116 | 13.28267 |
| CNAG_02( | - | hypothetic | 90.56156917 | 107.9870562 | 70.68906257 | 228.6995385 | 261.158428 | 273.5998472 | 5.38E-14 | 1.498619 | 8.21E-15 | 7.764324 | 2.825722 | 13.26908 |
| CNAG_01 | - | hypothetic | 1064.264348 | 853.2209349 | 957.4886909 | 547.3570435 | 478.2682203 | 568.6523828 | 6.03E-14 | -0.86728 | 9.21E-15 | -7.74966 | 0.54818 | 13.21959 |
| CNAG_02 | - | hypothetic | 782.7195463 | 719.9192271 | 750.9315094 | 1214.66389 | 1196.674843 | 1188.749467 | 6.45E-14 | 0.66108 | 9.88E-15 | 7.74074 | 1.581266 | 13.19048 |

| | | | | | | | | | | | | | | |
|---|---|---|---|---|---|---|---|---|---|---|---|---|---|---|
| CNAG_024 | - | hypothetic | 1049.407104 | 1034.163983 | 1021.525764 | 679.0258453 | 642.2935148 | 627.5095308 | 6.45E-14 | -0.68742 | 9.88E-15 | -7.74084 | 0.620965 | 13.19048 |
| CNAG_066 | - | amidohyd | 12.37194808 | 5.719990179 | 13.07375815 | 94.16910735 | 70.2836359 | 100.1579971 | 6.76E-14 | 3.121252 | 1.04E-14 | 7.734681 | 8.701425 | 13.17014 |
| CNAG_035 | - | hypothetic | 2509.155618 | 2426.251327 | 2390.891621 | 1693.152785 | 1663.610271 | 1764.213052 | 6.80E-14 | -0.53196 | 1.04E-14 | -7.73379 | 0.691613 | 13.16745 |
| CNAG_008 | - | ribose 5-p | 19.3306397 | 2.474975599 | 6.69123751 | 100.4165827 | 93.10315096 | 93.71428476 | 6.96E-14 | 3.386269 | 1.07E-14 | 7.730687 | 10.45607 | 13.1572 |
| CNAG_018 | - | hypothetic | 2529.066017 | 2486.598211 | 2702.223751 | 1837.993425 | 1599.709153 | 1748.708773 | 7.18E-14 | -0.58924 | 1.10E-14 | -7.72658 | 0.664695 | 13.14389 |
| CNAG_054 | - | endo-1,3(4 | 162.7903143 | 127.4403502 | 134.6195358 | 361.4472318 | 315.2443426 | 331.5361181 | 7.31E-14 | 1.235484 | 1.13E-14 | 7.724148 | 2.354603 | 13.13593 |
| CNAG_066 | - | rhomboid | 809.6171986 | 758.714031 | 844.7556695 | 1255.039544 | 1282.072677 | 1307.495923 | 8.19E-14 | 0.657433 | 1.26E-14 | 7.709354 | 1.577274 | 13.08692 |
| CNAG_046 | - | 6-phospho | 4066.400093 | 4055.58974 | 4066.50043 | 6103.117621 | 5642.212991 | 5443.004596 | 8.22E-14 | 0.480784 | 1.27E-14 | 7.708659 | 1.395502 | 13.0849 |
| CNAG_056 | - | cytoplasm | 635.742922 | 730.65402 | 717.8642556 | 1115.750842 | 1109.529454 | 1200.858899 | 8.47E-14 | 0.703041 | 1.31E-14 | 7.704818 | 1.627933 | 13.07218 |
| CNAG_068 | - | hypothetic | 693.0997028 | 668.8712125 | 634.4010239 | 392.5196384 | 384.3635436 | 350.0321762 | 8.96E-14 | -0.84058 | 1.39E-14 | -7.69748 | 0.55842 | 13.04791 |
| CNAG_074 | HLH4 | hypothetic | 7.688993376 | 7.861619987 | 9.868427751 | 70.27142454 | 89.96491848 | 71.64676367 | 8.98E-14 | 3.306261 | 1.39E-14 | 7.697089 | 9.891991 | 13.04693 |
| CNAG_058 | KAP123 | importin b | 2294.113363 | 2029.765463 | 2051.747074 | 1432.428694 | 1441.958738 | 1264.767701 | 9.30E-14 | -0.63951 | 1.45E-14 | -7.69226 | 0.641931 | 13.03156 |
| CNAG_004 | - | succinate | 1721.940527 | 1499.692719 | 1645.382206 | 970.7321819 | 1085.714542 | 1050.438831 | 9.92E-14 | -0.6638 | 1.54E-14 | -7.68384 | 0.631215 | 13.00366 |
| CNAG_006 | - | alpha-mar | 1126.045935 | 1146.531594 | 1113.514544 | 1662.811079 | 1744.418745 | 1665.04963 | 9.92E-14 | 0.567896 | 1.54E-14 | 7.683841 | 1.48236 | 13.00366 |
| CNAG_011 | RAD54 | DNA repa | 858.7584929 | 961.2246909 | 981.260144 | 1425.435135 | 1565.954831 | 1448.546977 | 1.01E-13 | 0.649463 | 1.57E-14 | 7.681809 | 1.568584 | 12.99712 |
| CNAG_001 | - | ubiquinon | 3452.634396 | 3031.828251 | 3447.683062 | 2114.878654 | 2345.951828 | 2236.076211 | 1.02E-13 | -0.58473 | 1.58E-14 | -7.68056 | 0.666776 | 12.99321 |
| CNAG_073 | - | hypothetic | 55.52916445 | 50.95653875 | 50.39755079 | 162.7480667 | 162.6946317 | 193.5663882 | 1.03E-13 | 1.722661 | 1.61E-14 | 7.67844 | 3.300446 | 12.98637 |
| CNAG_029 | - | carboxype | 61.41032222 | 69.24587493 | 69.56759833 | 189.9977405 | 208.605668 | 192.7169063 | 1.18E-13 | 1.550449 | 1.84E-14 | 7.661002 | 2.929084 | 12.92801 |
| CNAG_008 | - | rab family | 924.324978 | 826.2564027 | 876.4602936 | 524.0379341 | 499.052471 | 543.1050582 | 1.24E-13 | -0.76138 | 1.94E-14 | -7.65486 | 0.589931 | 12.90792 |
| CNAG_019 | - | hypothetic | 144.1680972 | 118.8264259 | 135.6701981 | 305.0093446 | 296.2377016 | 369.0236771 | 1.28E-13 | 1.273226 | 2.01E-14 | 7.65016 | 2.417014 | 12.89239 |
| CNAG_075 | - | hypothetic | 50.95073302 | 60.64421852 | 68.49145898 | 194.9055699 | 209.5846638 | 167.317832 | 1.33E-13 | 1.663892 | 2.08E-14 | 7.645568 | 3.168703 | 12.87756 |
| CNAG_047 | PCD101 | hypothetic | 145.400486 | 151.1046761 | 167.6263458 | 332.2433964 | 362.875489 | 337.1256792 | 1.33E-13 | 1.139639 | 2.09E-14 | 7.645086 | 2.203259 | 12.87627 |
| CNAG_046 | - | hypothetic | 579.7225234 | 699.4002203 | 660.2870713 | 1078.460973 | 1035.330141 | 1145.334636 | 1.43E-13 | 0.734998 | 2.25E-14 | 7.635227 | 1.664395 | 12.84369 |
| CNAG_023 | - | hypothetic | 11.23795633 | 14.33731119 | 19.45942145 | 93.0587424 | 97.45334163 | 90.48207018 | 1.49E-13 | 2.654711 | 2.34E-14 | 7.63021 | 6.297204 | 12.82746 |
| CNAG_056 | - | hypothetic | 1879.724534 | 1756.114424 | 1888.543705 | 1205.094473 | 1274.625641 | 1260.234044 | 1.49E-13 | -0.57841 | 2.35E-14 | -7.62991 | 0.669702 | 12.82679 |
| CNAG_019 | - | Xaa-Pro a | 2185.856686 | 2327.115723 | 2476.133153 | 1555.346002 | 1623.963459 | 1621.028065 | 1.53E-13 | -0.55726 | 2.41E-14 | -7.62637 | 0.679591 | 12.81519 |
| CNAG_049 | - | hypothetic | 74.12982752 | 53.13239283 | 29.13676896 | 214.6499643 | 206.9344338 | 186.0591297 | 1.72E-13 | 1.941764 | 2.72E-14 | 7.610988 | 3.841751 | 12.76479 |
| CNAG_037 | - | mRNA tur | 1657.823511 | 1504.009631 | 1778.628637 | 1111.015363 | 989.8717761 | 1020.494869 | 1.74E-13 | -0.6784 | 2.75E-14 | -7.60936 | 0.624858 | 12.76 |
| CNAG_077 | - | glycoside | 12.43447575 | 18.6504655 | 24.78966841 | 87.05749165 | 114.7947118 | 116.9034809 | 1.77E-13 | 2.53861 | 2.81E-14 | 7.606816 | 5.81029 | 12.75178 |
| CNAG_041 | - | hypothetic | 3201.720816 | 3189.085628 | 3245.105507 | 2208.345517 | 2380.47347 | 2355.927112 | 1.88E-13 | -0.48805 | 2.99E-14 | -7.59865 | 0.712988 | 12.72469 |
| CNAG_062 | MLN2 | nuclear m | 1189.436524 | 1159.112086 | 1038.639991 | 724.1400459 | 718.2277011 | 648.6358473 | 1.89E-13 | -0.7114 | 3.01E-14 | -7.59804 | 0.610728 | 12.72299 |
| CNAG_019 | - | allantoate | 1046.975962 | 885.5792143 | 1149.349385 | 551.1526769 | 585.322551 | 611.9523621 | 1.90E-13 | -0.83417 | 3.03E-14 | -7.59715 | 0.560904 | 12.72033 |
| CNAG_000 | - | mitochonc | 2638.843022 | 2600.807576 | 3029.562323 | 1819.217643 | 1891.373441 | 1540.124537 | 2.00E-13 | -0.67083 | 3.19E-14 | -7.59042 | 0.628147 | 12.69844 |
| CNAG_075 | ELP3 | elongator | 1598.246375 | 1480.278471 | 1534.499847 | 1012.097152 | 1053.805966 | 1014.912521 | 2.02E-13 | -0.59855 | 3.21E-14 | -7.5894 | 0.660418 | 12.69533 |
| CNAG_028 | CAN1 | Putative b | 31.07667535 | 40.15430672 | 20.57900293 | 130.2479068 | 153.681319 | 123.8983868 | 2.06E-13 | 2.143173 | 3.29E-14 | 7.586558 | 4.417324 | 12.68616 |
| CNAG_041 | - | lectin, mar | 1121.731937 | 1184.895427 | 973.5991425 | 696.1354226 | 593.8953277 | 651.9293619 | 2.11E-13 | -0.7714 | 3.38E-14 | -7.58297 | 0.58585 | 12.67479 |
| CNAG_033 | - | nicotinam | 995.6182197 | 898.460111 | 977.7657516 | 590.997588 | 580.1751244 | 596.3991087 | 2.14E-13 | -0.71623 | 3.42E-14 | -7.58139 | 0.608688 | 12.66983 |
| CNAG_076 | - | hypothetic | 1984.767553 | 1870.291084 | 1827.806457 | 1312.589122 | 1317.757825 | 1225.863963 | 2.15E-13 | -0.57531 | 3.44E-14 | -7.58041 | 0.671142 | 12.66691 |
| CNAG_059 | - | peptidyl-p | 5634.141797 | 5819.060523 | 5270.359964 | 4123.043207 | 4116.792357 | 4128.855825 | 2.27E-13 | -0.45033 | 3.64E-14 | -7.57319 | 0.731878 | 12.64307 |
| CNAG_014 | RPB4 | DNA-direc | 248.4625996 | 313.1729972 | 349.369835 | 138.6528631 | 102.0651749 | 112.5512165 | 2.44E-13 | -1.38473 | 3.92E-14 | -7.56363 | 0.38296 | 12.61179 |
| CNAG_012 | - | hypothetic | 232.9635777 | 268.4919448 | 276.3814175 | 492.4935609 | 531.8643468 | 489.2865579 | 2.66E-13 | 0.947355 | 4.28E-14 | 7.552281 | 1.928335 | 12.57456 |
| CNAG_006 | - | hypothetic | 967.2934626 | 776.7047582 | 851.9123101 | 531.9212752 | 404.9164147 | 452.1620827 | 3.21E-13 | -0.91918 | 5.16E-14 | -7.52776 | 0.52881 | 12.49356 |
| CNAG_034 | - | cytoplasm | 2360.928256 | 2350.838439 | 2434.582285 | 1621.518256 | 1632.563018 | 1755.315277 | 3.25E-13 | -0.52772 | 5.24E-14 | -7.52583 | 0.69365 | 12.48746 |
| CNAG_048 | - | mannose- | 335.6828772 | 404.1998797 | 347.8645195 | 732.5068558 | 667.143821 | 619.2669071 | 3.37E-13 | 0.879382 | 5.44E-14 | 7.520956 | 1.839587 | 12.47193 |
| CNAG_121 | - | unspecifie | 18.22468614 | 17.57164512 | 19.4748256 | 108.0470356 | 114.8719385 | 81.86002204 | 3.54E-13 | 2.472513 | 5.71E-14 | 7.514571 | 5.550099 | 12.45105 |
| CNAG_054 | CMK1 | CAMK/CA | 342.7186295 | 451.5758951 | 348.9491849 | 719.8281807 | 718.9371807 | 696.8882675 | 3.62E-13 | 0.888519 | 5.84E-14 | 7.511555 | 1.851275 | 12.44137 |
| CNAG_074 | - | exosome | 1163.956018 | 1183.94676 | 1230.524503 | 802.7912279 | 747.5209898 | 778.4479387 | 4.01E-13 | -0.63504 | 6.48E-14 | -7.49806 | 0.643924 | 12.39695 |
| CNAG_069 | - | anon-23da | 1260.786844 | 1326.089263 | 1150.624232 | 834.7059048 | 780.3045619 | 770.7115795 | 4.04E-13 | -0.66298 | 6.53E-14 | -7.49702 | 0.631571 | 12.39385 |
| CNAG_007 | - | RNA-bind | 1976.774138 | 2027.59799 | 2115.696742 | 1358.429157 | 1423.847031 | 1460.033735 | 4.04E-13 | -0.54385 | 6.54E-14 | -7.4967 | 0.685939 | 12.39312 |
| CNAG_018 | GAT8 | hypothetic | 10.13451751 | 36.90703579 | 18.43771754 | 137.0440622 | 106.5860589 | 132.4879644 | 4.92E-13 | 2.539341 | 7.98E-14 | 7.47067 | 5.813235 | 12.30771 |
| CNAG_018 | - | translatior | 1246.799779 | 1227.045025 | 1262.527648 | 861.9986013 | 797.5754895 | 733.0020682 | 5.20E-13 | -0.65889 | 8.44E-14 | -7.46329 | 0.633367 | 12.28435 |
| CNAG_028 | - | hypothetic | 802.655663 | 897.6294007 | 827.7166857 | 1349.330015 | 1316.567105 | 1262.071444 | 5.79E-13 | 0.621759 | 9.44E-14 | 7.448516 | 1.53875 | 12.2373 |
| CNAG_053 | - | Hemolysir | 1591.232714 | 1558.87441 | 1455.618445 | 957.5443341 | 991.6488253 | 1080.354269 | 5.91E-13 | -0.61968 | 9.65E-14 | -7.44564 | 0.650816 | 12.22816 |
| CNAG_078 | - | hypothetic | 1487.479852 | 1569.659107 | 1644.29369 | 1007.404846 | 956.2341452 | 1104.773637 | 6.08E-13 | -0.63076 | 9.92E-14 | -7.44192 | 0.645836 | 12.21623 |
| CNAG_033 | - | cytochron | 1591.301003 | 1526.601827 | 1608.066043 | 1099.361586 | 989.028864 | 1067.069413 | 6.30E-13 | -0.59801 | 1.03E-13 | -7.43708 | 0.660664 | 12.20067 |
| CNAG_074 | - | U3 small r | 3506.467582 | 3286.09325 | 3648.188029 | 2360.101325 | 2533.997611 | 2502.479495 | 6.48E-13 | -0.51317 | 1.06E-13 | -7.43323 | 0.700681 | 12.18835 |
| CNAG_029 | - | F-type H - | 2198.556561 | 2070.732668 | 2493.127822 | 1571.041324 | 1397.891061 | 1292.53919 | 6.55E-13 | -0.68214 | 1.07E-13 | -7.43167 | 0.623239 | 12.18355 |
| CNAG_067 | - | hypothetic | 1146.221489 | 1027.743353 | 1079.101513 | 667.2625337 | 619.8007309 | 719.5789215 | 6.82E-13 | -0.71323 | 1.12E-13 | -7.42609 | 0.609951 | 12.16618 |
| CNAG_052 | - | 1,2-dihydr | 2093.406063 | 1899.431921 | 2094.366006 | 1336.630724 | 1420.406961 | 1393.453141 | 7.59E-13 | -0.5685 | 1.25E-13 | -7.41162 | 0.674319 | 12.11973 |
| CNAG_016 | - | ATP-deper | 8068.832593 | 7356.639356 | 7814.864017 | 5434.379523 | 5675.436293 | 5891.763787 | 7.92E-13 | -0.46662 | 1.30E-13 | -7.40592 | 0.723657 | 12.10139 |
| CNAG_014 | - | hypothetic | 59.07445531 | 70.3091664 | 57.86545433 | 199.59823 | 220.0087922 | 158.5296393 | 9.05E-13 | 1.618268 | 1.49E-13 | 7.388029 | 3.070062 | 12.04321 |
| CNAG_024 | - | DNA repa | 2043.223982 | 2004.961951 | 1919.530769 | 1346.775755 | 1429.049347 | 1395.66413 | 9.45E-13 | -0.53203 | 1.56E-13 | -7.38217 | 0.691581 | 12.02442 |
| CNAG_046 | - | hypothetic | 39.21845444 | 43.39500949 | 35.47866848 | 122.8148999 | 150.8561348 | 158.060592 | 1.03E-12 | 1.862999 | 1.70E-13 | 7.370278 | 3.637632 | 11.98596 |
| CNAG_064 | - | hypothetic | 782.0289018 | 771.2863826 | 830.5699845 | 509.3466708 | 468.0243056 | 465.4383399 | 1.11E-12 | -0.74006 | 1.83E-13 | -7.36037 | 0.598714 | 11.95466 |
| CNAG_076 | - | pyruvate | 4263.05492 | 4057.470962 | 3765.585627 | 2943.175006 | 2900.579244 | 2612.464675 | 1.14E-12 | -0.53082 | 1.89E-13 | -7.35638 | 0.69216 | 11.94233 |
| CNAG_030 | - | endoribon | 296.8020145 | 270.0983653 | 234.4567706 | 118.3964814 | 79.60000186 | 109.1717159 | 1.17E-12 | -1.40151 | 1.93E-13 | -7.35344 | 0.378532 | 11.9334 |
| CNAG_025 | MRS3/4 | solute car | 916.3944287 | 986.7210343 | 951.1304133 | 567.5867171 | 635.4635908 | 594.1835248 | 1.21E-12 | -0.68274 | 2.01E-13 | -7.34804 | 0.622979 | 11.91618 |
| CNAG_006 | - | hypothetic | 421.9624217 | 420.3870747 | 415.0174766 | 711.9100365 | 693.7962148 | 732.3911669 | 1.23E-12 | 0.750923 | 2.04E-13 | 7.346028 | 1.682869 | 11.90997 |

| | | | | | | | | | | | | | |
|---|---|---|---|---|---|---|---|---|---|---|---|---|---|
| CNAG_033 | - | hypothetic | 701.5700646 | 760.4752864 | 815.6088702 | 464.9244731 | 446.4648788 | 444.3410609 | 1.27E-12 | -0.76432 | 2.12E-13 | -7.34117 | 0.588731 | 11.8945 |
| CNAG_057 | - | hypothetic | 42.72604294 | 42.34021055 | 50.36778616 | 140.9032565 | 164.6983152 | 150.4859691 | 1.38E-12 | 1.744952 | 2.30E-13 | 7.329876 | 3.351836 | 11.85917 |
| CNAG_046 | - | DNA clam | 178.1149439 | 235.052741 | 188.9895452 | 429.670801 | 398.8790921 | 424.7976636 | 1.47E-12 | 1.043573 | 2.45E-13 | 7.321428 | 2.061326 | 11.83212 |
| CNAG_033 | TIM13 | mitochonc | 1605.356259 | 1532.012207 | 1728.537485 | 1119.529576 | 1042.529395 | 1085.951547 | 1.54E-12 | -0.59926 | 2.57E-13 | -7.31525 | 0.660091 | 11.81244 |
| CNAG_129 | - | unspecifie | 108.0177166 | 121.9777473 | 86.66583308 | 259.9985797 | 287.953659 | 247.2322242 | 1.71E-12 | 1.317172 | 2.86E-13 | 7.300753 | 2.491772 | 11.76624 |
| CNAG_066 | - | hypothetic | 1215.280304 | 1231.330857 | 1209.223697 | 822.2617624 | 823.5050461 | 699.7140312 | 1.85E-12 | -0.65625 | 3.09E-13 | -7.29056 | 0.634526 | 11.73368 |
| CNAG_029 | - | histone H | 306.4476234 | 357.8628372 | 294.55214 | 607.0253295 | 559.2467082 | 591.390186 | 2.12E-12 | 0.862067 | 3.55E-13 | 7.271581 | 1.81764 | 11.67321 |
| CNAG_027 | - | Short-chai | 1150.517659 | 1077.599506 | 935.4822167 | 1815.667574 | 1766.904046 | 1548.551748 | 2.21E-12 | 0.682359 | 3.70E-13 | 7.266065 | 1.604762 | 11.65579 |
| CNAG_006 | - | hypothetic | 1417.393507 | 1403.784343 | 1660.226796 | 920.9688858 | 907.0740566 | 1014.872479 | 2.22E-12 | -0.67246 | 3.72E-13 | -7.2652 | 0.627436 | 11.65365 |
| CNAG_009 | - | hypothetic | 674.0801583 | 570.1807765 | 447.1117916 | 1147.331898 | 1052.718898 | 998.9050469 | 2.27E-12 | 0.903171 | 3.80E-13 | 7.262332 | 1.870171 | 11.64475 |
| CNAG_042 | - | hypothetic | 3237.911178 | 3068.4535 | 3403.972466 | 2304.134978 | 2379.603519 | 2291.566893 | 2.29E-12 | -0.49283 | 3.85E-13 | -7.26078 | 0.710631 | 11.64009 |
| CNAG_060 | - | nucleoside | 13.60977174 | 20.8044359 | 30.10530851 | 125.428308 | 100.6357866 | 102.7569721 | 2.33E-12 | 2.361139 | 3.92E-13 | 7.258281 | 5.137758 | 11.63237 |
| CNAG_006 | - | sphingolip | 139.462503 | 126.3265427 | 127.1352753 | 283.205758 | 300.7093814 | 297.1127144 | 2.42E-12 | 1.15113 | 4.08E-13 | 7.253041 | 2.220878 | 11.61619 |
| CNAG_054 | - | L-lactate c | 54.57385353 | 132.7093869 | 60.05369451 | 270.2966513 | 283.6457597 | 290.243221 | 2.42E-12 | 1.765557 | 4.09E-13 | 7.252694 | 3.400054 | 11.61539 |
| CNAG_028 | - | cytochror | 639.1353513 | 662.7745983 | 642.1514219 | 1003.657068 | 1016.391854 | 1026.64884 | 2.55E-12 | 0.633074 | 4.31E-13 | 7.245584 | 1.550866 | 11.59291 |
| CNAG_048 | FZC3 | nuclear pr | 1228.814355 | 1150.889117 | 1323.565705 | 1887.833768 | 1871.176842 | 1812.710438 | 2.58E-12 | 0.574103 | 4.35E-13 | 7.244282 | 1.488751 | 11.58905 |
| CNAG_004 | - | Isovaleryl- | 81.09951637 | 59.58303308 | 62.12517896 | 201.0804566 | 203.3805893 | 182.8017194 | 2.59E-12 | 1.520345 | 4.37E-13 | 7.243457 | 2.868596 | 11.58672 |
| CNAG_122 | - | unspecifie | 88970.67781 | 86436.87847 | 79626.68715 | 58693.54267 | 67478.89699 | 57968.39657 | 2.59E-12 | -0.48543 | 4.38E-13 | -7.24312 | 0.714282 | 11.58595 |
| CNAG_078 | - | hypothetic | 17.0353485 | 15.41139763 | 19.46044455 | 97.8762205 | 91.27273095 | 87.21416337 | 2.62E-12 | 2.415276 | 4.43E-13 | 7.241761 | 5.334217 | 11.58192 |
| CNAG_073 | - | mRNA sur | 1519.997766 | 1389.78202 | 1455.580958 | 1002.052023 | 991.6954197 | 926.1276316 | 2.88E-12 | -0.59554 | 4.88E-13 | -7.22863 | 0.661797 | 11.5402 |
| CNAG_029 | - | U3 small r | 2444.856348 | 2198.94856 | 2292.763774 | 1514.842104 | 1682.661575 | 1592.174271 | 3.14E-12 | -0.55007 | 5.33E-13 | -7.2166 | 0.682985 | 11.50242 |
| CNAG_038 | - | oxidoredu | 146.4511744 | 118.8126441 | 137.7840579 | 298.0681096 | 302.3948076 | 304.8841595 | 3.19E-12 | 1.15681 | 5.42E-13 | 7.214393 | 2.229638 | 11.49567 |
| CNAG_065 | - | hypothetic | 20.57647759 | 32.59745923 | 13.1064056 | 111.7251083 | 97.9694312 | 125.7643601 | 3.51E-12 | 2.343991 | 5.96E-13 | 7.201492 | 5.077052 | 11.45516 |
| CNAG_047 | - | hypothetic | 1520.657925 | 1454.673269 | 1289.517043 | 2132.171228 | 2086.767723 | 2166.796344 | 3.51E-12 | 0.567377 | 5.95E-13 | 7.201542 | 1.481827 | 11.45516 |
| CNAG_010 | - | hypothetic | 194.4110866 | 232.9105666 | 207.0950813 | 405.3169025 | 434.3731138 | 412.6543087 | 3.54E-12 | 0.968116 | 6.02E-13 | 7.200018 | 1.956285 | 11.45109 |
| CNAG_001 | - | elongatior | 3907.982848 | 3384.131947 | 3432.830976 | 2529.07036 | 2557.274178 | 2399.286352 | 3.81E-12 | -0.53456 | 6.48E-13 | -7.19004 | 0.690368 | 11.41963 |
| CNAG_000 | - | hypothetic | 750.4241867 | 714.1831263 | 736.760544 | 430.6056845 | 459.4518377 | 453.1904137 | 3.84E-12 | -0.72859 | 6.55E-13 | -7.18862 | 0.603492 | 11.41543 |
| CNAG_036 | - | hypothetic | 1382.20543 | 1346.625785 | 1262.591483 | 915.6692355 | 901.1253342 | 821.7904829 | 3.96E-12 | -0.61285 | 6.75E-13 | -7.18437 | 0.653903 | 11.40253 |
| CNAG_045 | LHP1 | hypothetic | 5977.922563 | 5504.745099 | 5582.998886 | 7502.173292 | 7862.464704 | 8032.919225 | 4.00E-12 | 0.439802 | 6.83E-13 | 7.182761 | 1.356418 | 11.39774 |
| CNAG_065 | - | hypothetic | 404.659064 | 569.012145 | 498.2040666 | 885.5610793 | 855.1338206 | 885.5886971 | 4.22E-12 | 0.821632 | 7.22E-13 | 7.175242 | 1.767404 | 11.37448 |
| CNAG_047 | - | Monosacc | 2094.185394 | 2330.614068 | 2398.54817 | 3104.058232 | 3468.69207 | 3316.956577 | 4.59E-12 | 0.520485 | 7.86E-13 | 7.163607 | 1.434438 | 11.33789 |
| CNAG_018 | - | molecular | 4250.418665 | 4289.158325 | 4623.994369 | 2995.963113 | 3132.585699 | 3361.694037 | 4.89E-12 | -0.48744 | 8.38E-13 | -7.15482 | 0.713288 | 11.31066 |
| CNAG_028 | RAC1 | Ras-like G | 274.8132856 | 231.9025022 | 252.9375912 | 465.9881021 | 492.1133894 | 482.5869872 | 5.10E-12 | 0.910856 | 8.75E-13 | 7.148881 | 1.880161 | 11.29281 |
| CNAG_004 | - | UBA/TS-N | 216.6186965 | 279.2152569 | 224.1763634 | 485.6420313 | 439.3371722 | 501.3630937 | 5.44E-12 | 0.972878 | 9.37E-13 | 7.139478 | 1.962752 | 11.26401 |
| CNAG_008 | - | adenylate | 1117.224054 | 1258.173264 | 1054.643104 | 688.2271999 | 730.2967518 | 760.6623639 | 5.67E-12 | -0.66934 | 9.76E-13 | -7.13378 | 0.628794 | 11.24663 |
| CNAG_025 | - | hypothetic | 714.7777408 | 1200.877821 | 1145.030116 | 496.612747 | 487.7537864 | 527.6437157 | 5.76E-12 | -1.03171 | 9.93E-13 | -7.13146 | 0.489132 | 11.23961 |
| CNAG_033 | CHS2 | Chitin syn | 234.0939078 | 289.983309 | 209.2746348 | 491.829792 | 457.4771106 | 519.1099796 | 5.88E-12 | 0.988926 | 1.02E-12 | 7.128437 | 1.984707 | 11.23038 |
| CNAG_022 | - | parafibron | 2328.140997 | 2230.161671 | 2188.271392 | 1498.514585 | 1555.80464 | 1676.483418 | 6.18E-12 | -0.52781 | 1.07E-12 | -7.12141 | 0.693608 | 11.20885 |
| CNAG_016 | - | hypothetic | 15.92897915 | 30.44662653 | 17.35884765 | 114.2456866 | 124.3993173 | 84.12670936 | 6.29E-12 | 2.363064 | 1.09E-12 | 7.118918 | 5.144616 | 11.20129 |
| CNAG_069 | - | hypothetic | 866.0309824 | 781.0337253 | 884.9621762 | 517.8297544 | 538.820941 | 504.2855889 | 6.37E-12 | -0.71491 | 1.10E-12 | -7.11704 | 0.609245 | 11.19569 |
| CNAG_014 | - | hypothetic | 1386.382658 | 1304.926299 | 1259.631479 | 2083.302237 | 1937.554239 | 1859.379972 | 6.46E-12 | 0.558643 | 1.12E-12 | 7.115152 | 1.472883 | 11.19005 |
| CNAG_028 | GEP4 | hypothetic | 1246.816582 | 1169.992692 | 1354.187241 | 849.5076681 | 744.8900983 | 812.8643719 | 6.79E-12 | -0.66365 | 1.18E-12 | -7.10791 | 0.631278 | 11.16787 |
| CNAG_029 | - | tRNA (gua | 1639.084354 | 1446.907691 | 1586.736554 | 1074.441028 | 975.227801 | 1050.418448 | 7.07E-12 | -0.60762 | 1.23E-12 | -7.10236 | 0.656276 | 11.15072 |
| CNAG_035 | - | rab GDP-c | 6187.669506 | 5907.488676 | 6176.80061 | 4700.589963 | 4539.444395 | 4649.504997 | 7.63E-12 | -0.41108 | 1.33E-12 | -7.09166 | 0.752058 | 11.11742 |
| CNAG_037 | - | hypothetic | 17.1900956 | 42.3428624 | 54.61911531 | 144.8433626 | 179.5347055 | 159.2936693 | 7.85E-12 | 2.078744 | 1.36E-12 | 7.087588 | 4.224392 | 11.10494 |
| CNAG_011 | - | U3 small r | 3554.224746 | 3129.870578 | 3374.139074 | 2189.605868 | 2372.674186 | 2466.92912 | 8.50E-12 | -0.53304 | 1.48E-12 | -7.0762 | 0.691096 | 11.07047 |
| CNAG_045 | - | Alpha-1,2- | 420.8046834 | 394.5615272 | 451.2430757 | 718.9241158 | 739.6148967 | 692.5143577 | 9.19E-12 | 0.748886 | 1.60E-12 | 7.065185 | 1.680495 | 11.03689 |
| CNAG_002 | AMT1 | amt family | 2903.930623 | 2769.99747 | 3011.527789 | 2161.011361 | 2010.3664 | 2080.624236 | 9.37E-12 | -0.48957 | 1.64E-12 | -7.06228 | 0.712239 | 11.02813 |
| CNAG_034 | - | alginate ly | 653.2258733 | 780.1743463 | 650.7174406 | 1057.249838 | 1134.593487 | 1200.849588 | 9.48E-12 | 0.688683 | 1.66E-12 | 7.06059 | 1.611812 | 11.02314 |
| CNAG_010 | - | ATP phos | 1443.152069 | 1591.188165 | 1550.491054 | 1060.420843 | 1085.748266 | 990.5104584 | 9.50E-12 | -0.56311 | 1.66E-12 | -7.06015 | 0.67684 | 11.02207 |
| CNAG_009 | - | DNA-bind | 2522.000817 | 2474.721849 | 2306.673307 | 1659.64547 | 1799.972687 | 1747.573198 | 9.71E-12 | -0.50369 | 1.70E-12 | -7.05684 | 0.705298 | 11.01262 |
| CNAG_041 | - | smad nucl | 1646.31338 | 1724.825747 | 1835.189989 | 1271.448594 | 1107.213791 | 1100.426095 | 9.74E-12 | -0.59675 | 1.71E-12 | -7.0564 | 0.661241 | 11.01154 |
| CNAG_063 | - | N-acetyltr | 2782.472525 | 2656.860987 | 2884.615196 | 1851.073542 | 1981.074624 | 2072.81364 | 9.86E-12 | -0.51073 | 1.73E-12 | -7.05462 | 0.701866 | 11.00631 |
| CNAG_055 | - | hypothetic | 1117.368297 | 1243.178859 | 1276.368844 | 819.9125371 | 781.1816265 | 798.4280616 | 9.92E-12 | -0.61538 | 1.74E-12 | -7.05365 | 0.652756 | 11.00356 |
| CNAG_035 | PTC7 | hypothetic | 2277.802553 | 1896.23177 | 2264.961134 | 1470.593666 | 1432.454388 | 1269.21224 | 1.05E-11 | -0.64229 | 1.84E-12 | -7.04624 | 0.640694 | 10.98075 |
| CNAG_073 | - | T-complex | 8980.959473 | 8651.712973 | 9027.435639 | 6818.291485 | 7000.383705 | 6840.963038 | 1.06E-11 | -0.38337 | 1.86E-12 | -7.04466 | 0.766647 | 10.97641 |
| CNAG_010 | - | hypothetic | 1890.350221 | 1943.551129 | 1934.432697 | 1293.054031 | 1385.918723 | 1410.063206 | 1.06E-11 | -0.51157 | 1.87E-12 | -7.04362 | 0.70146 | 10.97346 |
| CNAG_069 | - | hypothetic | 749.154998 | 759.3248822 | 657.8976033 | 395.5321928 | 354.9117424 | 466.4533734 | 1.18E-11 | -0.8481 | 2.08E-12 | -7.02905 | 0.555516 | 10.92898 |
| CNAG_016 | - | acetyl/pro | 61.31215636 | 52.02869684 | 41.8914331 | 135.8069471 | 208.0954535 | 173.7631538 | 1.23E-11 | 1.728553 | 2.17E-12 | 7.022784 | 3.313953 | 10.9101 |
| CNAG_062 | - | Glutathior | 85.85799751 | 99.36038487 | 54.70982576 | 215.5704606 | 270.9402136 | 210.5890582 | 1.27E-11 | 1.531466 | 2.25E-12 | 7.018173 | 2.890795 | 10.89606 |
| CNAG_030 | - | hypothetic | 1027.162683 | 946.9311049 | 952.2250898 | 612.786427 | 651.8418881 | 628.5831002 | 1.58E-11 | -0.64489 | 2.81E-12 | -6.98704 | 0.639544 | 10.80133 |
| CNAG_067 | - | RNA polyr | 8734.548721 | 8378.042306 | 9039.136985 | 6159.743715 | 6506.093431 | 6856.470417 | 1.59E-11 | -0.43731 | 2.83E-12 | -6.9858 | 0.738509 | 10.79777 |
| CNAG_065 | - | aromatic a | 671.1370806 | 718.4320339 | 699.4319079 | 429.9056969 | 448.2686871 | 385.5423426 | 1.67E-11 | -0.74155 | 2.97E-12 | -6.97915 | 0.598095 | 10.7781 |
| CNAG_039 | - | leucine ca | 117.2893483 | 114.4545898 | 80.27852869 | 291.5522997 | 261.0355564 | 233.9710494 | 1.71E-11 | 1.320699 | 3.05E-12 | 6.975431 | 2.497872 | 10.76692 |
| CNAG_067 | - | tRNA (gua | 1248.958684 | 1049.340946 | 1196.387165 | 706.9285117 | 681.0577062 | 789.5050857 | 1.74E-11 | -0.69864 | 3.11E-12 | -6.97272 | 0.616152 | 10.75912 |
| CNAG_078 | - | U3 small r | 716.2618538 | 716.690744 | 803.1574706 | 1354.303903 | 1159.525134 | 1122.221197 | 1.83E-11 | 0.686334 | 3.27E-12 | 6.965517 | 1.609189 | 10.7372 |
| CNAG_030 | - | riboflavin | 2218.396265 | 2093.351895 | 2147.731675 | 1594.410551 | 1545.485709 | 1460.087547 | 1.87E-11 | -0.50531 | 3.35E-12 | -6.96208 | 0.704508 | 10.7272 |
| CNAG_052 | - | signal rec | 3462.134023 | 3291.485465 | 3579.957256 | 2522.083871 | 2559.01448 | 2555.755387 | 1.94E-11 | -0.45178 | 3.47E-12 | -6.95729 | 0.731139 | 10.71301 |

| | | | | | | | | | | | | | | |
|---|---|---|---|---|---|---|---|---|---|---|---|---|---|---|
| CNAG_01€ | - | multifunct | 1173.322535 | 1161.357651 | 1299.813175 | 840.9711922 | 776.864994 | 775.136248 | 2.14E-11 | -0.61894 | 3.84E-12 | -6.94305 | 0.651151 | 10.67007 |
| CNAG_01€ | - | amino oxi | 292.2725445 | 245.8898505 | 203.9518975 | 493.3691476 | 471.3150224 | 514.6971197 | 2.16E-11 | 0.982521 | 3.89E-12 | 6.941197 | 1.975915 | 10.66467 |
| CNAG_05! | - | solute car | 851.6657772 | 899.7988011 | 879.9572808 | 1336.773074 | 1427.935322 | 1238.817195 | 2.19E-11 | 0.590624 | 3.93E-12 | 6.939685 | 1.505898 | 10.66031 |
| CNAG_06( | - | ADP-ribos | 647.8013111 | 709.7982165 | 701.5396029 | 353.4095725 | 414.5799811 | 439.8284253 | 2.31E-11 | -0.78521 | 4.16E-12 | -6.93153 | 0.580268 | 10.63615 |
| CNAG_03( | LDP1 | hypothetic | 745.3107766 | 698.3440463 | 678.4213399 | 1085.379209 | 1079.328554 | 1108.801663 | 2.31E-11 | 0.610045 | 4.16E-12 | 6.931568 | 1.526306 | 10.63615 |
| CNAG_05! | - | U3 small r | 5129.545112 | 4707.204418 | 4637.933161 | 3253.556176 | 3521.62361 | 3676.985372 | 2.37E-11 | -0.48542 | 4.28E-12 | -6.92759 | 0.714291 | 10.62463 |
| CNAG_01{ | - | Glutathior | 66.10805811 | 99.33765148 | 63.21219996 | 194.4194952 | 202.2962277 | 260.0973526 | 2.47E-11 | 1.511393 | 4.46E-12 | 6.921924 | 2.850852 | 10.60813 |
| CNAG_03{ | - | general tr | 98.63786652 | 92.93137218 | 86.6338628 | 228.0841822 | 264.7634783 | 201.8149068 | 2.53E-11 | 1.312487 | 4.58E-12 | 6.91809 | 2.483693 | 10.59697 |
| CNAG_06{ | - | Alpha-ket | 504.9439607 | 427.5930855 | 457.3256919 | 271.1325249 | 234.1945266 | 235.6875071 | 2.79E-11 | -0.92325 | 5.06E-12 | -6.90389 | 0.527319 | 10.55436 |
| CNAG_00 | - | hypothetic | 1988.298293 | 1836.920559 | 1867.259706 | 1314.902926 | 1400.614036 | 1269.144682 | 2.79E-11 | -0.53053 | 5.07E-12 | -6.90369 | 0.692302 | 10.55403 |
| CNAG_07 | - | mitochonc | 1336.792319 | 1385.412681 | 1324.430009 | 974.099443 | 885.5545083 | 910.5470848 | 2.81E-11 | -0.56177 | 5.10E-12 | -6.90285 | 0.67747 | 10.55177 |
| CNAG_00: | - | hypothetic | 291.3442376 | 376.172995 | 331.8484962 | 632.7209338 | 638.7777571 | 550.4659 | 2.91E-11 | 0.852366 | 5.30E-12 | 6.897358 | 1.80546 | 10.53585 |
| CNAG_03 | ECM22 | hypothetic | 1489.73613 | 1413.494276 | 1546.200955 | 1064.372755 | 1005.483269 | 962.7539814 | 2.99E-11 | -0.56893 | 5.45E-12 | -6.89345 | 0.674117 | 10.52451 |
| CNAG_00{ | - | multisite-s | 4114.905665 | 3968.099802 | 4221.96786 | 2883.924358 | 3225.842227 | 2927.690451 | 3.05E-11 | -0.46105 | 5.56E-12 | -6.89042 | 0.726456 | 10.51582 |
| CNAG_02 | - | AGC/NDR | 583.0815017 | 617.5099125 | 549.3969906 | 882.925247 | 994.0923941 | 922.3146495 | 3.11E-11 | 0.662809 | 5.68E-12 | 6.887423 | 1.583162 | 10.50726 |
| CNAG_06 | - | hypothetic | 78.86719033 | 80.01680215 | 89.7984037 | 209.4093162 | 188.3631048 | 243.5782975 | 3.18E-11 | 1.358637 | 5.81E-12 | 6.88422 | 2.564428 | 10.49778 |
| CNAG_06{ | - | hypothetic | 1224.683993 | 1236.750141 | 1249.752641 | 829.9985324 | 806.1955148 | 883.8530798 | 3.24E-11 | -0.57386 | 5.93E-12 | -6.88136 | 0.671815 | 10.48963 |
| CNAG_05! | - | hypothetic | 279.6210643 | 323.4085405 | 324.3540545 | 562.5964044 | 529.0551538 | 552.5546336 | 3.52E-11 | 0.812509 | 6.46E-12 | 6.869201 | 1.756263 | 10.45288 |
| CNAG_02 | ADE2 | phosphori | 2305.96394 | 2085.841465 | 2489.963355 | 1450.217741 | 1417.710237 | 1673.150392 | 3.54E-11 | -0.61514 | 6.49E-12 | -6.86843 | 0.652866 | 10.45083 |
| CNAG_03: | ZNF2 | C2H2 type | 82.37154538 | 95.06155989 | 78.1129975 | 181.6746376 | 237.0229915 | 252.4642538 | 3.68E-11 | 1.379023 | 6.75E-12 | 6.86288 | 2.600922 | 10.43422 |
| CNAG_06{ | - | hypothetic | 224.6780534 | 221.0975915 | 223.0777317 | 413.8428622 | 473.2954415 | 394.9989982 | 3.90E-11 | 0.922692 | 7.16E-12 | 6.85439 | 1.895649 | 10.409 |
| CNAG_02 | - | phenylalar | 4033.223525 | 4197.551281 | 4253.963986 | 2774.106701 | 3243.087146 | 3068.649711 | 3.93E-11 | -0.47366 | 7.22E-12 | -6.85329 | 0.720138 | 10.40596 |
| CNAG_05( | - | L-xylulose | 115.099976 | 148.9038994 | 148.4254778 | 309.062645 | 282.4225925 | 310.413913 | 4.12E-11 | 1.117973 | 7.58E-12 | 6.846312 | 2.170418 | 10.38505 |
| CNAG_02 | - | hypothetic | 1932.378912 | 1889.720923 | 2004.803061 | 1394.327649 | 1404.028376 | 1398.982992 | 4.29E-11 | -0.48865 | 7.90E-12 | -6.84032 | 0.712693 | 10.36768 |
| CNAG_05 | SMG1 | glucose-m | 123.5920427 | 42.46626496 | 31.3028438 | 1254.094621 | 1119.136296 | 1020.546081 | 4.34E-11 | 4.093599 | 8.02E-12 | 6.838262 | 17.07245 | 10.36208 |
| CNAG_01 | - | U4/U6 sm | 2739.246943 | 2622.372655 | 2643.647637 | 1930.568433 | 1910.332494 | 2033.952624 | 4.47E-11 | -0.46197 | 8.26E-12 | -6.83396 | 0.725996 | 10.34933 |
| CNAG_00 | - | splicing fa | 3278.763261 | 3204.174007 | 3140.626361 | 2498.847296 | 2304.522974 | 2310.435359 | 4.52E-11 | -0.45136 | 8.35E-12 | -6.83247 | 0.731355 | 10.3451 |
| CNAG_02! | - | calnexin | 9362.657816 | 8429.75511 | 8341.740781 | 6465.691542 | 6561.304879 | 6587.849853 | 4.80E-11 | -0.42968 | 8.87E-12 | -6.82376 | 0.742425 | 10.31903 |
| CNAG_07 | - | membrane | 1447.895656 | 1631.070392 | 1704.015318 | 1087.632858 | 1100.381407 | 1083.72698 | 4.87E-11 | -0.56322 | 9.00E-12 | -6.8216 | 0.676789 | 10.31279 |
| CNAG_04 | - | RNA polyr | 3035.933614 | 2960.704242 | 3198.163685 | 2202.953523 | 2301.983573 | 2289.310676 | 4.87E-11 | -0.45216 | 9.01E-12 | -6.82149 | 0.730947 | 10.31275 |
| CNAG_05{ | ALG9 | Alpha-1,2- | 2914.358622 | 2764.579911 | 2644.742525 | 2039.558827 | 2065.636942 | 1984.050121 | 5.19E-11 | -0.46641 | 9.64E-12 | -6.81183 | 0.723762 | 10.28442 |
| CNAG_04( | - | alpha 1,3- | 2570.00652 | 2866.841767 | 2537.016937 | 1962.546374 | 1933.658485 | 1704.361684 | 5.53E-11 | -0.52476 | 1.03E-11 | -6.8025 | 0.695075 | 10.25739 |
| CNAG_05: | - | arsenite-r | 6343.0747 | 6472.047209 | 6714.284843 | 4907.560896 | 4998.352418 | 5166.810273 | 5.55E-11 | -0.38903 | 1.03E-11 | -6.80165 | 0.763643 | 10.25541 |
| CNAG_01! | - | hypothetic | 405.0552476 | 523.3075679 | 523.3499224 | 232.7604642 | 233.305783 | 283.3257974 | 5.58E-11 | -0.9701 | 1.04E-11 | -6.80087 | 0.510472 | 10.25335 |
| CNAG_07 | - | hypothetic | 909.2499051 | 866.1054451 | 859.4387916 | 588.7680701 | 578.5145368 | 519.8448821 | 5.73E-11 | -0.66001 | 1.07E-11 | -6.79709 | 0.632872 | 10.24221 |
| CNAG_05( | - | hypothetic | 387.6734333 | 574.9521364 | 558.511549 | 240.5402354 | 268.7496522 | 247.9099328 | 5.79E-11 | -1.02255 | 1.08E-11 | -6.79543 | 0.492247 | 10.2375 |
| CNAG_06! | - | monocarb | 37.99106404 | 35.8556343 | 23.7650499 | 103.249306 | 161.672541 | 116.1610726 | 5.99E-11 | 1.984349 | 1.12E-11 | 6.790422 | 3.956842 | 10.22271 |
| CNAG_00: | - | chaperone | 5547.714419 | 5208.238099 | 5474.006522 | 4070.914626 | 3919.237027 | 4233.168184 | 6.15E-11 | -0.42454 | 1.15E-11 | -6.78649 | 0.745074 | 10.21145 |
| CNAG_06! | - | hypothetic | 1522.583813 | 1645.092337 | 1754.134463 | 1049.402303 | 1089.133229 | 1193.571818 | 6.24E-11 | -0.57795 | 1.17E-11 | -6.7842 | 0.669915 | 10.20484 |
| CNAG_02 | - | cytoplasm | 3418.945888 | 3350.725744 | 3413.626375 | 2573.492259 | 2565.058527 | 2591.265446 | 6.60E-11 | -0.41286 | 1.24E-11 | -6.77593 | 0.751134 | 10.18056 |
| CNAG_03! | - | coatomer | 12498.19916 | 11941.10835 | 11910.0887 | 9326.716495 | 9572.537491 | 9635.213786 | 6.70E-11 | -0.36472 | 1.26E-11 | -6.77356 | 0.776618 | 10.17398 |
| CNAG_03! | GRX5 | Grx4 famil | 2654.018527 | 2527.570859 | 2699.067909 | 1979.686162 | 1934.522505 | 1767.612692 | 7.40E-11 | -0.48792 | 1.39E-11 | -6.75876 | 0.713055 | 10.13072 |
| CNAG_07! | - | WD-repea | 745.8434992 | 726.0532684 | 791.1187961 | 441.4858241 | 498.3104614 | 472.0599877 | 7.40E-11 | -0.6964 | 1.39E-11 | -6.75882 | 0.617108 | 10.13072 |
| CNAG_00! | BZP3 | bZip trans | 1264.83936 | 917.8657923 | 890.4200663 | 590.1666589 | 565.4594432 | 574.247543 | 7.54E-11 | -0.84526 | 1.42E-11 | -6.75597 | 0.556612 | 10.12265 |
| CNAG_04( | - | metacaspa | 1469.245483 | 1269.398099 | 1348.125899 | 1978.020939 | 2008.306778 | 1983.66263 | 7.76E-11 | 0.530947 | 1.46E-11 | 6.751686 | 1.444877 | 10.11009 |
| CNAG_02( | - | hypothetic | 1226.980612 | 1209.819081 | 1222.02933 | 844.0631754 | 761.3053636 | 861.6569947 | 7.76E-11 | -0.58362 | 1.46E-11 | -6.75155 | 0.667285 | 10.10995 |
| CNAG_01( | - | tagatose-t | 787.8037108 | 761.580152 | 745.3273853 | 453.1709189 | 512.9849363 | 480.9409783 | 8.01E-11 | -0.68161 | 1.51E-11 | -6.74675 | 0.62347 | 10.09645 |
| CNAG_05 | - | hypothetic | 756.3041523 | 716.3603186 | 791.1105398 | 478.9756862 | 491.4053518 | 406.6543011 | 8.23E-11 | -0.73456 | 1.56E-11 | -6.74261 | 0.601002 | 10.08436 |
| CNAG_01 | - | hypothetic | 301.740692 | 342.773521 | 271.1000197 | 600.9313242 | 517.8010614 | 543.700246 | 9.11E-11 | 0.846453 | 1.73E-11 | 6.72751 | 1.798075 | 10.04036 |
| CNAG_00 | - | DNA polyr | 57.80726936 | 57.37820642 | 49.32256835 | 168.4809804 | 144.4666343 | 160.4874491 | 9.14E-11 | 1.518987 | 1.73E-11 | 6.726976 | 2.865898 | 10.03905 |
| CNAG_04: | - | galactonat | 474.4206585 | 441.9295194 | 416.0981008 | 703.9928554 | 755.9880755 | 731.3362916 | 9.41E-11 | 0.702296 | 1.79E-11 | 6.722571 | 1.627092 | 10.02619 |
| CNAG_06 | - | endoplasn | 1771.180924 | 1751.777411 | 1695.576941 | 1279.184266 | 1247.03458 | 1225.815265 | 9.60E-11 | -0.4912 | 1.82E-11 | -6.7197 | 0.711432 | 10.01792 |
| CNAG_12 | - | unspecifie | 1448.031731 | 780.9722897 | 875.3129863 | 269.763765 | 251.1739032 | 208.1948443 | 9.60E-11 | -2.10993 | 1.82E-11 | -6.71949 | 0.231658 | 10.01756 |
| CNAG_00 | - | hypothetic | 1899.660802 | 1804.620173 | 1988.78985 | 1349.957132 | 1338.452501 | 1370.112997 | 9.82E-11 | -0.50379 | 1.87E-11 | -6.71603 | 0.705254 | 10.00782 |
| CNAG_03{ | SAR1 | small copl | 4779.439671 | 5049.797124 | 4794.70463 | 3857.739622 | 3763.173271 | 3620.41402 | 9.98E-11 | -0.3945 | 1.90E-11 | -6.71364 | 0.760751 | 10.00097 |
| CNAG_06! | - | hypothetic | 2088.793965 | 2029.764464 | 1987.784021 | 1447.235203 | 1414.338093 | 1535.492089 | 1.00E-10 | -0.48879 | 1.90E-11 | -6.71318 | 0.712624 | 9.999874 |
| CNAG_00 | - | S-adenosy | 5355.005384 | 4893.638565 | 5215.92356 | 3824.174297 | 3782.105007 | 3997.820833 | 1.02E-10 | -0.42993 | 1.93E-11 | -6.7109 | 0.742296 | 9.993372 |
| CNAG_05{ | - | splicing fa | 5234.835075 | 5252.388708 | 5254.33451 | 3918.335594 | 4192.765179 | 4069.993422 | 1.07E-10 | -0.38534 | 2.04E-11 | -6.70325 | 0.765597 | 9.970884 |
| CNAG_01 | FHB1 | Flavohemc | 7169.240134 | 2906.871331 | 4156.676392 | 675.8088866 | 683.2043127 | 838.7916013 | 1.14E-10 | -2.71522 | 2.17E-11 | -6.69443 | 0.152278 | 9.944961 |
| CNAG_05 | ARG5 | N-acetyl-g | 5742.670179 | 5251.333694 | 5304.469764 | 4177.575836 | 4086.612129 | 4070.015902 | 1.20E-10 | -0.4177 | 2.28E-11 | -6.68661 | 0.748617 | 9.922036 |
| CNAG_01: | - | hypothetic | 567.3701269 | 710.8111722 | 659.950874 | 398.0079318 | 380.0526696 | 365.5385458 | 1.21E-10 | -0.77465 | 2.31E-11 | -6.6851 | 0.584531 | 9.917827 |
| CNAG_01! | - | hypothetic | 1539.385705 | 1116.153459 | 1180.464248 | 753.6085895 | 801.895496 | 725.2366999 | 1.21E-10 | -0.76757 | 2.32E-11 | -6.6842 | 0.587407 | 9.915447 |
| CNAG_00 | CAS31 | protein in | 4142.902154 | 4043.496093 | 3892.491539 | 3112.888867 | 2902.272581 | 3083.061161 | 1.29E-10 | -0.42399 | 2.46E-11 | -6.67575 | 0.745358 | 9.890672 |
| CNAG_03 | GMT2 | GDP-man | 979.5990809 | 1070.8205 | 1282.650428 | 623.5560464 | 734.6598482 | 673.0190102 | 1.29E-10 | -0.72982 | 2.47E-11 | -6.67517 | 0.602979 | 9.889246 |
| CNAG_05! | - | hypothetic | 1059.469693 | 1134.642628 | 1005.816114 | 1550.744155 | 1514.860084 | 1601.724387 | 1.31E-10 | 0.52992 | 2.50E-11 | 6.673257 | 1.443849 | 9.883848 |
| CNAG_02 | - | inorganic | 8808.174284 | 8569.83819 | 9396.39787 | 6976.322424 | 6863.211366 | 6808.774553 | 1.32E-10 | -0.39034 | 2.52E-11 | -6.672 | 0.762952 | 9.880419 |
| CNAG_02 | - | DNA-direc | 13072.65285 | 12185.67296 | 11716.01086 | 9205.26904 | 9423.289535 | 9678.511585 | 1.34E-10 | -0.40084 | 2.56E-11 | -6.66964 | 0.757419 | 9.873696 |
| CNAG_03( | - | hypothetic | 1738.486866 | 1669.920732 | 1797.895084 | 1296.361729 | 1194.391555 | 1120.406896 | 1.34E-10 | -0.54315 | 2.57E-11 | -6.66935 | 0.686273 | 9.873102 |

| | | | | | | | | | | | | | | |
|---|---|---|---|---|---|---|---|---|---|---|---|---|---|---|
| CNAG_014 | - | hypothetic | 11789.33963 | 10866.92488 | 11208.35153 | 8779.60509 | 8728.93314 | 8818.147812 | 1.34E-10 | -0.37889 | 2.58E-11 | -6.66877 | 0.76903 | 9.871687 |
| CNAG_027 | - | Monosacc | 54.47091468 | 97.16084071 | 60.0069064 | 181.188832 | 229.4368652 | 200.4840788 | 1.37E-10 | 1.518289 | 2.64E-11 | 6.665371 | 2.86451 | 9.861898 |
| CNAG_014 | - | hsp71-like | 7652.97563 | 6615.358137 | 7234.682062 | 4160.092826 | 5549.609837 | 4655.146083 | 1.38E-10 | -0.59829 | 2.64E-11 | -6.6651 | 0.660534 | 9.861372 |
| CNAG_020 | - | hypothetic | 1007.296686 | 908.1582557 | 984.169327 | 669.7871297 | 525.7418335 | 600.8433448 | 1.44E-10 | -0.70787 | 2.77E-11 | -6.65805 | 0.612225 | 9.84108 |
| CNAG_030 | - | multiple d | 354.3281595 | 403.1294863 | 418.1787263 | 656.6416596 | 655.8806676 | 652.5146434 | 1.52E-10 | 0.725508 | 2.92E-11 | 6.650399 | 1.653482 | 9.819048 |
| CNAG_023 | - | molybdop | 1021.510871 | 1098.784408 | 1031.144972 | 658.6890334 | 758.8721206 | 666.3474173 | 1.56E-10 | -0.61248 | 3.01E-11 | -6.6463 | 0.65407 | 9.807356 |
| CNAG_038 | - | small nucl | 3671.082493 | 3346.437645 | 3495.742956 | 2676.25514 | 2589.194864 | 2557.992455 | 1.70E-10 | -0.44209 | 3.29E-11 | -6.63289 | 0.736068 | 9.768555 |
| CNAG_002 | - | NADH del | 765.3704237 | 607.5910146 | 687.6939848 | 424.4767406 | 406.7991398 | 377.7679134 | 1.74E-10 | -0.78679 | 3.37E-11 | -6.62942 | 0.579632 | 9.758622 |
| CNAG_003 | - | lysophosp | 2609.695649 | 2585.739796 | 2714.001022 | 1827.766862 | 1902.580194 | 2043.926344 | 1.76E-10 | -0.46951 | 3.41E-11 | -6.62751 | 0.722209 | 9.753261 |
| CNAG_010 | - | E3 ubiquit | 2740.970371 | 2715.25366 | 2733.422892 | 3487.681641 | 3852.459057 | 3816.537011 | 1.78E-10 | 0.430767 | 3.45E-11 | 6.626094 | 1.34795 | 9.749375 |
| CNAG_030 | - | hypothetic | 31.03251996 | 41.1977501 | 19.50409209 | 112.975566 | 138.1182845 | 109.5092365 | 1.78E-10 | 1.965835 | 3.45E-11 | 6.625912 | 3.906388 | 9.749113 |
| CNAG_019 | - | Sugar trar | 46.09431861 | 35.86515639 | 33.33631174 | 121.4844531 | 123.0311193 | 139.2522113 | 1.79E-10 | 1.732601 | 3.46E-11 | 6.625547 | 3.323264 | 9.748314 |
| CNAG_000 | ERG11 | cytochrom | 976.7734243 | 847.7934119 | 834.9201121 | 579.4302653 | 559.522246 | 451.0921007 | 1.81E-10 | -0.75766 | 3.51E-11 | -6.62358 | 0.591454 | 9.742901 |
| CNAG_028 | - | large subu | 741.1989237 | 721.7550515 | 819.8863773 | 508.6212586 | 462.8694223 | 444.353386 | 1.81E-10 | -0.70538 | 3.51E-11 | -6.62341 | 0.613282 | 9.742863 |
| CNAG_010 | NCP1 | NADPH-fe | 4123.107457 | 4043.517378 | 4081.235688 | 3073.130213 | 3217.19483 | 3144.123628 | 2.03E-10 | -0.39187 | 3.96E-11 | -6.60566 | 0.762142 | 9.691552 |
| CNAG_079 | - | nucleopro | 10439.37212 | 9384.380459 | 10039.49245 | 7085.143365 | 7274.612788 | 7913.3332 | 2.06E-10 | -0.43875 | 4.00E-11 | -6.60406 | 0.737772 | 9.687147 |
| CNAG_075 | - | hypothetic | 12807.64669 | 11888.3362 | 12674.70405 | 9311.110261 | 9650.153024 | 9936.058343 | 2.07E-10 | -0.38654 | 4.03E-11 | -6.60281 | 0.764962 | 9.683762 |
| CNAG_048 | FZC8 | hypothetic | 321.4446557 | 231.9340342 | 246.5777372 | 614.4900418 | 494.5576563 | 486.0264078 | 2.14E-10 | 0.981851 | 4.18E-11 | 6.597701 | 1.974998 | 9.669059 |
| CNAG_048 | - | V-type AT | 1020.875278 | 978.4535146 | 937.557081 | 1474.561698 | 1373.375628 | 1455.201356 | 2.21E-10 | 0.536408 | 4.31E-11 | 6.593169 | 1.450356 | 9.656064 |
| CNAG_037 | - | signal rec | 2687.922302 | 2551.294151 | 2845.154618 | 1939.90095 | 2023.380519 | 1956.283621 | 2.21E-10 | -0.46541 | 4.32E-11 | -6.59276 | 0.724263 | 9.655135 |
| CNAG_001 | - | hypothetic | 343.6982569 | 344.9517035 | 299.8774053 | 610.9430756 | 528.102227 | 583.6355645 | 2.31E-10 | 0.788669 | 4.51E-11 | 6.586303 | 1.72748 | 9.637067 |
| CNAG_032 | - | rRNA bioç | 1726.740901 | 1603.121179 | 1681.674578 | 1056.377191 | 1180.619621 | 1227.985243 | 2.31E-10 | -0.54836 | 4.51E-11 | -6.58619 | 0.683798 | 9.637019 |
| CNAG_066 | - | import inr | 9202.737488 | 8124.88296 | 9195.902355 | 6332.548139 | 6772.655008 | 6654.460907 | 2.34E-10 | -0.44067 | 4.58E-11 | -6.5841 | 0.736794 | 9.631159 |
| CNAG_018 | - | heparinas | 9246.483134 | 9581.685446 | 8313.146085 | 11732.16895 | 12299.47149 | 12411.41711 | 2.40E-10 | 0.410259 | 4.70E-11 | 6.580099 | 1.328925 | 9.61975 |
| CNAG_080 | - | hypothetic | 1069.909424 | 982.7820526 | 991.9377707 | 1418.317832 | 1501.118796 | 1557.294237 | 2.47E-10 | 0.541673 | 4.83E-11 | 6.575982 | 1.45566 | 9.607998 |
| CNAG_069 | - | hypothetic | 14.80844388 | 47.62719169 | 21.63314942 | 128.1146122 | 113.5347425 | 133.644473 | 2.61E-10 | 2.178301 | 5.12E-11 | 6.567428 | 4.526201 | 9.583585 |
| CNAG_039 | CPR2 | Nonmatin | 11.16249675 | 15.36474424 | 9.864976823 | 74.41078221 | 62.57504797 | 77.05020909 | 2.65E-10 | 2.591451 | 5.19E-11 | 6.565273 | 6.027044 | 9.577574 |
| CNAG_043 | - | acetyltrans | 824.1738217 | 858.5554848 | 886.0603476 | 588.0160562 | 558.6583738 | 533.137101 | 2.65E-10 | -0.62821 | 5.21E-11 | -6.56491 | 0.646978 | 9.576783 |
| CNAG_075 | - | threonine | 334.3368112 | 324.4884727 | 275.3626046 | 547.7058659 | 517.7916267 | 593.5343851 | 2.73E-10 | 0.813636 | 5.36E-11 | 6.560542 | 1.757636 | 9.564329 |
| CNAG_018 | - | cathepsin | 1063.020753 | 1170.2056 | 1131.622105 | 1687.059633 | 1618.405499 | 1544.102429 | 2.75E-10 | 0.512597 | 5.42E-11 | 6.559008 | 1.426616 | 9.5604 |
| CNAG_028 | - | 4-aminob | 1923.593018 | 1642.869171 | 1300.00358 | 1026.9066 | 986.4605105 | 819.6685873 | 2.75E-10 | -0.797 | 5.42E-11 | -6.55902 | 0.575546 | 9.5604 |
| CNAG_017 | - | hypothetic | 348.315727 | 313.7356355 | 292.4089426 | 585.2292179 | 528.1573169 | 553.6892254 | 2.78E-10 | 0.790432 | 5.47E-11 | 6.557622 | 1.729593 | 9.556635 |
| CNAG_000 | SEC14-2 | SEC14 cyt | 203.638474 | 178.0378067 | 208.1284575 | 407.9201401 | 392.9335833 | 346.1743244 | 2.80E-10 | 0.943144 | 5.51E-11 | 6.556491 | 1.922714 | 9.55361 |
| CNAG_066 | - | DNA-direc | 2566.567973 | 2668.695616 | 2804.642572 | 1958.594076 | 1993.172802 | 2030.632873 | 2.84E-10 | -0.4416 | 5.60E-11 | -6.55395 | 0.736317 | 9.546484 |
| CNAG_067 | - | hypothetic | 1955.692587 | 1820.790521 | 2010.126058 | 1411.485843 | 1369.508598 | 1372.348208 | 2.85E-10 | -0.49412 | 5.63E-11 | -6.55337 | 0.709994 | 9.545072 |
| CNAG_000 | - | ran-bindir | 5037.530581 | 5082.17074 | 5559.282653 | 4036.715266 | 3810.559929 | 3992.286406 | 3.00E-10 | -0.42057 | 5.93E-11 | -6.54553 | 0.747129 | 9.522535 |
| CNAG_063 | PDR5-3 | ABC trans | 1405.515321 | 1247.570235 | 1276.444057 | 854.0783672 | 908.0373326 | 914.9580942 | 3.01E-10 | -0.56944 | 5.95E-11 | -6.54487 | 0.673879 | 9.520891 |
| CNAG_033 | - | pre-mRNA | 5568.830471 | 5838.48017 | 5639.324643 | 4018.657328 | 4373.006449 | 4549.545545 | 3.17E-10 | -0.41251 | 6.26E-11 | -6.53738 | 0.751315 | 9.499414 |
| CNAG_005 | TIM50 | mitochond | 4591.354796 | 4388.309031 | 4838.358132 | 3060.511335 | 3508.728598 | 3502.69566 | 3.22E-10 | -0.47199 | 6.37E-11 | -6.53483 | 0.720967 | 9.492254 |
| CNAG_042 | - | hypothetic | 36.83173651 | 44.42529075 | 23.76392677 | 120.0284064 | 134.5116976 | 115.0531181 | 3.23E-10 | 1.826589 | 6.39E-11 | 6.534284 | 3.546974 | 9.490953 |
| CNAG_070 | - | hypothetic | 19.3723723 | 23.99891687 | 26.89952674 | 92.84240925 | 92.0348539 | 97.13736855 | 3.30E-10 | 2.032092 | 6.53E-11 | 6.531068 | 4.089976 | 9.481891 |
| CNAG_036 | - | malonic s | 1594.941855 | 1670.958859 | 1634.771492 | 1200.569643 | 1180.622208 | 1173.624524 | 3.33E-10 | -0.47821 | 6.60E-11 | -6.52938 | 0.717869 | 9.477533 |
| CNAG_079 | - | AAT famil | 1352.059114 | 1716.010876 | 1318.077011 | 982.5812595 | 945.0806527 | 888.4039911 | 3.40E-10 | -0.65378 | 6.75E-11 | -6.5262 | 0.635613 | 9.468859 |
| CNAG_063 | - | high-affini | 65.93542262 | 46.66110646 | 63.14707282 | 150.8772325 | 167.94596 | 175.9567759 | 3.54E-10 | 1.483295 | 7.05E-11 | 6.519647 | 2.795866 | 9.450934 |
| CNAG_031 | - | U4/U6.U5 | 502.8661151 | 494.3573912 | 498.9343524 | 299.134586 | 270.4468022 | 307.7450203 | 3.60E-10 | -0.78533 | 7.18E-11 | -6.51693 | 0.580221 | 9.443337 |
| CNAG_064 | - | nucleolar | 2638.673695 | 2187.155287 | 2623.303702 | 1810.784854 | 1585.930138 | 1645.486448 | 3.66E-10 | -0.57952 | 7.31E-11 | -6.51424 | 0.669185 | 9.436072 |
| CNAG_079 | ROM20 | Predicted | 1024.334279 | 924.5968755 | 860.7970001 | 1376.38872 | 1398.468456 | 1444.06917 | 3.70E-10 | 0.571521 | 7.39E-11 | 6.512547 | 1.48609 | 9.431453 |
| CNAG_061 | - | spermine | 25.28367746 | 36.93716932 | 49.26081215 | 138.239647 | 127.4109727 | 120.6010042 | 3.76E-10 | 1.792553 | 7.51E-11 | 6.5102 | 3.464274 | 9.42493 |
| CNAG_066 | - | cytoplasm | 2522.200543 | 2668.683116 | 2700.150136 | 1974.976571 | 1995.776992 | 1970.69867 | 3.92E-10 | -0.42488 | 7.85E-11 | -6.5035 | 0.744901 | 9.406626 |
| CNAG_033 | - | alpha-ketc | 3331.317843 | 3193.419678 | 3227.002575 | 2389.77199 | 2351.102592 | 2561.256456 | 3.95E-10 | -0.43277 | 7.91E-11 | -6.50234 | 0.740839 | 9.403803 |
| CNAG_032 | - | hypothetic | 1809.630876 | 1636.536535 | 1691.298166 | 1212.971189 | 1215.1128 | 1234.674339 | 4.09E-10 | -0.50404 | 8.21E-11 | -6.49676 | 0.705128 | 9.388487 |
| CNAG_020 | - | large subu | 722.6562195 | 787.4400703 | 874.25871 | 518.6946712 | 506.9003773 | 465.4470813 | 4.18E-10 | -0.6936 | 8.40E-11 | -6.49327 | 0.618311 | 9.378698 |
| CNAG_010 | - | hypothetic | 1488.236465 | 1326.605706 | 900.4235887 | 3299.520834 | 3247.342324 | 3019.231256 | 4.45E-10 | 1.349375 | 8.95E-11 | 6.483674 | 2.548017 | 9.352091 |
| CNAG_028 | - | RNA-bind | 3031.241026 | 2926.225773 | 3062.749337 | 2241.140899 | 2186.358038 | 2343.675961 | 4.51E-10 | -0.42915 | 9.09E-11 | -6.48132 | 0.742697 | 9.345566 |
| CNAG_127 | - | unspecifie | 178.0082878 | 199.5116886 | 174.0418306 | 386.9429499 | 364.448211 | 323.9733318 | 4.67E-10 | 0.946651 | 9.41E-11 | 6.476085 | 1.927393 | 9.330774 |
| CNAG_025 | IPP1 | Inorganic | 4278.477363 | 4340.885512 | 4562.171115 | 3414.868352 | 3446.637753 | 3150.840999 | 4.74E-10 | -0.41228 | 9.56E-11 | -6.47382 | 0.751433 | 9.324514 |
| CNAG_060 | - | rab GTPas | 843.4714095 | 880.4020329 | 904.458852 | 1236.235333 | 1314.838899 | 1279.815924 | 5.02E-10 | 0.52837 | 1.01E-10 | 6.464839 | 1.442299 | 9.298976 |
| CNAG_069 | - | thiol-spec | 376.3832905 | 375.117816 | 344.646618 | 572.3958038 | 659.4663361 | 605.8969946 | 5.31E-10 | 0.730332 | 1.07E-10 | 6.456357 | 1.65902 | 9.275156 |
| CNAG_077 | - | DNA poly | 551.5256562 | 525.9718162 | 523.7890014 | 851.9071935 | 782.4880773 | 972.0765296 | 5.32E-10 | 0.689012 | 1.08E-10 | 6.455858 | 1.612179 | 9.273986 |
| CNAG_037 | HXK2 | Hexokinas | 12925.42786 | 11361.45324 | 11117.73094 | 8816.139211 | 9015.306739 | 8747.13103 | 5.48E-10 | -0.42946 | 1.11E-10 | -6.45125 | 0.742538 | 9.261029 |
| CNAG_129 | - | unspecifie | 24.05033684 | 32.59963839 | 30.11100302 | 101.2820319 | 96.2002432 | 121.4010249 | 6.10E-10 | 1.872366 | 1.24E-10 | 6.434904 | 3.661326 | 9.215019 |
| CNAG_064 | - | hypothetic | 6.42053282 | 5.662897869 | 4.504488824 | 50.0454972 | 52.45523569 | 53.81299699 | 6.27E-10 | 3.353887 | 1.27E-10 | 6.430504 | 10.224 | 9.202714 |
| CNAG_016 | MLN4 | hypothetic | 2219.658154 | 2140.780417 | 2297.018413 | 1663.670853 | 1649.87569 | 1616.575164 | 6.59E-10 | -0.44914 | 1.34E-10 | -6.42284 | 0.732481 | 9.181354 |
| CNAG_065 | - | endoplasn | 4495.578284 | 4355.964775 | 4266.814603 | 3198.390567 | 3356.903311 | 3477.139464 | 6.73E-10 | -0.40218 | 1.37E-10 | -6.41956 | 0.756712 | 9.17225 |
| CNAG_036 | - | peptidylpr | 2240.4979 | 1918.857636 | 2130.650767 | 1527.454345 | 1476.464137 | 1441.201385 | 7.01E-10 | -0.51671 | 1.43E-10 | -6.41306 | 0.698962 | 9.153984 |
| CNAG_000 | - | prohibitin | 4455.720829 | 3765.58744 | 4132.403284 | 2930.648397 | 2989.414201 | 3070.860052 | 7.03E-10 | -0.47452 | 1.43E-10 | -6.41265 | 0.719706 | 9.153072 |
| CNAG_077 | HAD2 | hypothetic | 7127.818936 | 7426.640524 | 7315.794961 | 5959.938551 | 5669.426452 | 5691.942149 | 7.03E-10 | -0.35151 | 1.43E-10 | -6.41254 | 0.783764 | 9.153025 |
| CNAG_042 | - | pyridoxal | 2102.868999 | 2031.951926 | 2319.353074 | 1367.701604 | 1646.492838 | 1432.337187 | 7.12E-10 | -0.55336 | 1.45E-10 | -6.41053 | 0.68143 | 9.1478 |

| | | | | | | | | | | | | | | |
|---|---|---|---|---|---|---|---|---|---|---|---|---|---|---|
| CNAG_05( | - | hypothetic | 2.918470113 | 0.227157895 | 9.819147809 | 48.34569775 | 52.41820136 | 63.58073789 | 7.47E-10 | 3.872573 | 1.52E-10 | 6.402864 | 14.64741 | 9.126507 |
| CNAG_01( | - | Atypical/R | 1249.024768 | 1105.346973 | 1126.075628 | 787.9927926 | 755.3048225 | 813.9273521 | 8.32E-10 | -0.57898 | 1.70E-10 | -6.38611 | 0.669439 | 9.079939 |
| CNAG_04( | - | hypothetic | 618.1636934 | 696.1529712 | 676.2585969 | 1088.696628 | 1004.278092 | 957.925284 | 8.45E-10 | 0.600728 | 1.73E-10 | 6.383699 | 1.516481 | 9.073346 |
| CNAG_05! | PBX2 | parallel be | 2184.696617 | 2261.42237 | 2326.885509 | 1659.738685 | 1737.028626 | 1660.976233 | 8.88E-10 | -0.4366 | 1.82E-10 | -6.37579 | 0.738875 | 9.051697 |
| CNAG_01( | - | hypothetic | 417.7656862 | 437.2608221 | 494.5718102 | 241.4134613 | 249.7764233 | 272.2231158 | 9.14E-10 | -0.83811 | 1.88E-10 | -6.37101 | 0.559376 | 9.039175 |
| CNAG_01( | - | hypothetic | 690.8342087 | 607.5752145 | 683.4177899 | 412.0070701 | 391.2552769 | 425.4086581 | 9.15E-10 | -0.7049 | 1.88E-10 | -6.37064 | 0.613486 | 9.038393 |
| CNAG_02( | - | chloride c | 1633.538983 | 1807.771008 | 1735.015448 | 1271.395662 | 1259.123752 | 1238.016409 | 9.81E-10 | -0.47261 | 2.02E-10 | -6.35986 | 0.720662 | 9.008423 |
| CNAG_01( | - | ribosome | 1680.262319 | 1850.879735 | 1846.97097 | 1289.259094 | 1350.593337 | 1249.139914 | 1.02E-09 | -0.48251 | 2.10E-10 | -6.35405 | 0.715733 | 8.99224 |
| CNAG_04( | PRP31 | U4/U6 sm | 3507.727263 | 3405.697414 | 3763.377926 | 2752.544897 | 2698.765163 | 2554.678856 | 1.06E-09 | -0.43099 | 2.18E-10 | -6.34832 | 0.741752 | 8.976336 |
| CNAG_04 | - | hypothetic | 73.03851483 | 68.19039551 | 100.4094184 | 201.6643495 | 191.0615428 | 210.4383356 | 1.07E-09 | 1.30511 | 2.21E-10 | 6.345768 | 2.471027 | 8.96938 |
| CNAG_06( | - | Phosphop | 837.4238848 | 747.4094154 | 503.3965147 | 393.2902686 | 312.6116355 | 352.2531155 | 1.08E-09 | -0.99653 | 2.23E-10 | -6.3447 | 0.501203 | 8.966621 |
| CNAG_00( | SKN1 | putative g | 3765.043983 | 3763.569115 | 3498.089311 | 4930.789913 | 5080.73372 | 4665.917341 | 1.08E-09 | 0.39717 | 2.23E-10 | 6.344326 | 1.316922 | 8.965826 |
| CNAG_06( | - | hypothetic | 475.0483965 | 514.8331984 | 591.5999207 | 302.1970059 | 273.0065276 | 327.7155418 | 1.10E-09 | -0.82366 | 2.27E-10 | -6.34219 | 0.565007 | 8.960055 |
| CNAG_06: | - | GAF doma | 1145.391248 | 1209.82703 | 1325.412622 | 846.3884394 | 874.4286625 | 776.2636927 | 1.11E-09 | -0.57501 | 2.29E-10 | -6.34051 | 0.671282 | 8.955587 |
| CNAG_00 | - | hypothetic | 1281.897295 | 1290.621577 | 1329.720212 | 766.7782493 | 859.6949761 | 973.7275388 | 1.16E-09 | -0.60122 | 2.39E-10 | -6.33364 | 0.659197 | 8.937235 |
| CNAG_01: | CCH1 | calcium-cl | 1946.881184 | 1940.567844 | 1829.093946 | 2646.551254 | 2484.300274 | 2719.676524 | 1.24E-09 | 0.442707 | 2.57E-10 | 6.32257 | 1.359152 | 8.906354 |
| CNAG_04: | - | hypothetic | 49.59518047 | 47.67235042 | 31.22093169 | 144.427164 | 136.0560545 | 119.5789886 | 1.26E-09 | 1.636417 | 2.62E-10 | 6.319625 | 3.108927 | 8.898331 |
| CNAG_02( | - | hypothetic | 204.9736806 | 269.5305742 | 264.6362364 | 440.2443035 | 514.7006154 | 428.3069928 | 1.30E-09 | 0.892355 | 2.71E-10 | 6.314715 | 1.856204 | 8.884794 |
| CNAG_07: | - | hypothetic | 786.1726709 | 769.4264718 | 737.059687 | 1110.22188 | 1119.854148 | 1125.496452 | 1.32E-09 | 0.534019 | 2.75E-10 | 6.312292 | 1.447957 | 8.878247 |
| CNAG_07 | - | hypothetic | 19.43790168 | 42.2507224 | 29.06002817 | 114.7396194 | 113.6617288 | 108.3399993 | 1.36E-09 | 1.883078 | 2.83E-10 | 6.307835 | 3.688613 | 8.866249 |
| CNAG_06( | - | peptidyl-tl | 536.7371318 | 489.0441089 | 585.2341145 | 299.820247 | 349.9684759 | 281.2070803 | 1.51E-09 | -0.80935 | 3.15E-10 | -6.29098 | 0.570637 | 8.8198 |
| CNAG_00( | - | hypothetic | 214.3734869 | 313.6920482 | 303.0245881 | 572.2024523 | 516.192199 | 482.7091249 | 1.57E-09 | 0.905518 | 3.28E-10 | 6.284827 | 1.873217 | 8.803092 |
| CNAG_04: | GPR4 | G-protein | 1322.074023 | 1137.940142 | 1118.864428 | 1742.986366 | 1701.174523 | 1807.096418 | 1.63E-09 | 0.537874 | 3.40E-10 | 6.279307 | 1.451831 | 8.788178 |
| CNAG_07: | - | hypothetic | 483.835821 | 520.5390714 | 498.1738772 | 802.9750227 | 771.4049586 | 749.1775326 | 1.72E-09 | 0.614469 | 3.60E-10 | 6.270621 | 1.530994 | 8.764441 |
| CNAG_03: | - | hypothetic | 164.201522 | 271.6484437 | 211.3734937 | 410.645778 | 524.3637242 | 412.7322675 | 1.77E-09 | 1.04538 | 3.71E-10 | 6.26556 | 2.063911 | 8.751088 |
| CNAG_00( | - | hypothetic | 9.922912043 | 6.752061922 | 4.51547054 | 76.005946 | 54.06894468 | 41.99981916 | 1.79E-09 | 3.154025 | 3.75E-10 | 6.263928 | 8.901358 | 8.746794 |
| CNAG_00 | - | 2-deoxy-D | 1901.268348 | 1588.31913 | 1700.052565 | 2446.518248 | 2511.177353 | 2451.058361 | 1.84E-09 | 0.497591 | 3.86E-10 | 6.259787 | 1.411854 | 8.73551 |
| CNAG_02 | ATX1 | Copper tra | 1387.060639 | 1444.693277 | 1405.484465 | 1068.322122 | 986.5166238 | 979.3706742 | 1.86E-09 | -0.497 | 3.90E-10 | -6.25816 | 0.708578 | 8.731244 |
| CNAG_05( | RPD304 | histone de | 4390.49531 | 4423.813754 | 4067.411016 | 3312.890145 | 3190.391086 | 3389.445357 | 1.87E-09 | -0.39597 | 3.93E-10 | -6.25695 | 0.75998 | 8.728114 |
| CNAG_01 | PRP11 | splicing fa | 2045.577885 | 1902.662942 | 1988.830116 | 1485.461344 | 1345.308797 | 1472.22784 | 1.93E-09 | -0.47967 | 4.07E-10 | -6.2515 | 0.717143 | 8.71347 |
| CNAG_00( | UCH2 | ubiquitin | 667.6501557 | 651.7284078 | 737.7794568 | 451.7698057 | 414.553909 | 438.7427054 | 1.94E-09 | -0.67268 | 4.08E-10 | -6.25093 | 0.62734 | 8.712137 |
| CNAG_00( | - | hypothetic | 363.6549446 | 427.8836769 | 417.1179408 | 672.2478391 | 666.242351 | 623.7247527 | 1.99E-09 | 0.685378 | 4.18E-10 | 6.247169 | 1.608124 | 8.702172 |
| CNAG_04: | - | hypothetic | 32.0702672 | 19.71866749 | 20.53837333 | 79.10221291 | 113.091331 | 100.4638087 | 2.04E-09 | 2.023297 | 4.29E-10 | 6.243015 | 4.065117 | 8.691133 |
| CNAG_01( | - | exosome | 1530.729752 | 1569.707114 | 1670.985189 | 1153.862034 | 1190.156094 | 1059.332981 | 2.10E-09 | -0.50268 | 4.43E-10 | -6.23808 | 0.705795 | 8.67767 |
| CNAG_01( | - | COPII-coa | 4104.281244 | 3839.863084 | 3582.177988 | 2761.75675 | 2949.785571 | 2909.883197 | 2.15E-09 | -0.43465 | 4.53E-10 | -6.23447 | 0.739872 | 8.667924 |
| CNAG_05( | CKA1 | CMGC/CK | 6653.70963 | 7091.537139 | 6639.683934 | 5524.084311 | 5299.390491 | 5337.79561 | 2.15E-09 | -0.34982 | 4.54E-10 | -6.23421 | 0.784681 | 8.667454 |
| CNAG_01( | - | pre-rRNA- | 2374.7706 | 2048.136541 | 2136.020805 | 1542.951769 | 1586.895375 | 1595.470534 | 2.17E-09 | -0.48942 | 4.58E-10 | -6.23293 | 0.712312 | 8.66416 |
| CNAG_02( | - | D-lactate | 1461.815021 | 1576.115479 | 1454.562717 | 1115.764072 | 1052.073456 | 1075.928439 | 2.23E-09 | -0.48498 | 4.71E-10 | -6.22861 | 0.714507 | 8.652664 |
| CNAG_06 | - | U6 snRNA | 1001.59564 | 963.1063904 | 999.1357712 | 663.4479433 | 720.9191271 | 627.4982449 | 2.28E-09 | -0.57546 | 4.83E-10 | -6.22459 | 0.671072 | 8.642047 |
| CNAG_02: | DCR2 | dicer-like | 2157.724856 | 2016.884557 | 2191.43762 | 1475.204492 | 1596.415262 | 1591.012172 | 2.28E-09 | -0.46461 | 4.84E-10 | -6.2243 | 0.724665 | 8.641484 |
| CNAG_02! | - | thioredoxi | 51.89905389 | 43.3832605 | 34.40819644 | 127.7120404 | 118.5677297 | 150.3057326 | 2.30E-09 | 1.610506 | 4.87E-10 | 6.223314 | 3.053589 | 8.639003 |
| CNAG_03( | PAB2 | polyadeny | 2474.17809 | 2385.335913 | 2338.662006 | 1845.047721 | 1738.685683 | 1819.704686 | 2.30E-09 | -0.42919 | 4.88E-10 | -6.22276 | 0.742681 | 8.637727 |
| CNAG_06( | - | 5-oxoproli | 21.69251226 | 32.5678628 | 21.59857964 | 88.75943097 | 98.15209763 | 98.26304707 | 2.54E-09 | 1.926356 | 5.40E-10 | 6.207136 | 3.800941 | 8.595255 |
| CNAG_03( | - | ATP-depe | 1239.602387 | 1013.80349 | 1116.450434 | 735.8061883 | 712.1617194 | 777.294892 | 2.56E-09 | -0.61472 | 5.46E-10 | -6.20531 | 0.653055 | 8.590953 |
| CNAG_04: | - | hypothetic | 4624.169448 | 4650.133926 | 5485.644606 | 3629.609834 | 3491.39753 | 3693.657082 | 2.62E-09 | -0.4642 | 5.59E-10 | -6.20162 | 0.724875 | 8.581256 |
| CNAG_01( | - | cytoplasm | 146.441946 | 138.1528599 | 148.4200636 | 269.7919244 | 307.6228831 | 284.9870237 | 2.64E-09 | 0.977733 | 5.64E-10 | 6.200272 | 1.969369 | 8.577798 |
| CNAG_04( | - | racemase | 35.57317995 | 22.95030658 | 25.86086148 | 107.7622012 | 119.0808286 | 84.14886921 | 2.68E-09 | 1.901737 | 5.71E-10 | 6.198241 | 3.736627 | 8.572443 |
| CNAG_06! | - | FAD depe | 354.3976268 | 332.7152995 | 320.8514746 | 198.7753637 | 161.713435 | 177.9291141 | 2.80E-09 | -0.9236 | 6.00E-10 | -6.19053 | 0.527193 | 8.552184 |
| CNAG_05! | - | hypothetic | 92.33649175 | 70.8763678 | 84.96515507 | 20.17738746 | 11.47651704 | 23.67980816 | 2.80E-09 | -2.20342 | 6.00E-10 | -6.19041 | 0.217123 | 8.552084 |
| CNAG_02 | - | GTPase ac | 1226.357974 | 1127.145373 | 1096.456769 | 1568.495069 | 1648.583384 | 1760.440394 | 2.84E-09 | 0.513361 | 6.07E-10 | 6.18855 | 1.427371 | 8.547218 |
| CNAG_04: | - | hypothetic | 1371.891411 | 1391.929146 | 1504.603849 | 1005.961806 | 1061.643763 | 898.3906884 | 2.92E-09 | -0.54084 | 6.26E-10 | -6.18377 | 0.687371 | 8.534291 |
| CNAG_02: | - | NAD bind | 1002.19193 | 1001.055725 | 922.6258486 | 1422.365411 | 1344.906182 | 1418.576542 | 2.98E-09 | 0.50138 | 6.38E-10 | 6.180654 | 1.415567 | 8.525976 |
| CNAG_01: | - | upstream | 923.3507596 | 870.4626767 | 926.6098 | 626.9139181 | 608.6981656 | 617.4542323 | 2.99E-09 | -0.57026 | 6.41E-10 | -6.18004 | 0.673496 | 8.524542 |
| CNAG_06( | - | U2-associ | 2610.840383 | 2530.800028 | 2531.683889 | 1969.56314 | 1928.483617 | 1960.692203 | 3.01E-09 | -0.40459 | 6.46E-10 | -6.17869 | 0.755453 | 8.521075 |
| CNAG_03: | - | hypothetic | 1014.530947 | 1013.751145 | 1117.465218 | 732.7841764 | 743.3163867 | 647.5000861 | 3.19E-09 | -0.5827 | 6.84E-10 | -6.1697 | 0.667715 | 8.496847 |
| CNAG_04! | - | hypothetic | 744.2906938 | 926.6661639 | 727.5020516 | 1152.918549 | 1285.568455 | 1279.740074 | 3.23E-09 | 0.618326 | 6.94E-10 | 6.167537 | 1.535093 | 8.49116 |
| CNAG_12: | - | unspecifie | 35.75837138 | 53.05155821 | 51.41797281 | 136.2655201 | 118.4578889 | 170.1818039 | 3.24E-09 | 1.595277 | 6.97E-10 | 6.166874 | 3.021524 | 8.489589 |
| CNAG_03: | - | hypothetic | 318.0738219 | 287.9228539 | 373.3640436 | 568.7408835 | 537.642544 | 588.0445946 | 3.29E-09 | 0.775993 | 7.07E-10 | 6.164542 | 1.712368 | 8.483432 |
| CNAG_02: | CPL1 | putative se | 49.73551893 | 77.76886507 | 25.9303015 | 196.1000345 | 158.3464264 | 161.6577137 | 3.41E-09 | 1.757568 | 7.34E-10 | 6.158641 | 3.381276 | 8.467491 |
| CNAG_03: | - | hypothetic | 5.244303133 | 6.728790644 | 4.499014559 | 47.70515672 | 48.96526695 | 50.50813888 | 3.41E-09 | 3.258316 | 7.35E-10 | 6.158394 | 9.568653 | 8.467061 |
| CNAG_07( | - | hypothetic | 116.0842265 | 113.3601374 | 102.6066453 | 234.1897461 | 256.7920614 | 219.5550324 | 3.42E-09 | 1.083545 | 7.37E-10 | 6.157879 | 2.119237 | 8.46614 |
| CNAG_01! | - | hypothetic | 1287.769478 | 1327.247678 | 1277.513948 | 957.771868 | 948.6185726 | 841.7571518 | 3.63E-09 | -0.51802 | 7.83E-10 | -6.14831 | 0.698328 | 8.440183 |
| CNAG_06( | - | hypothetic | 513.6843768 | 653.709482 | 607.695147 | 382.5096978 | 350.728646 | 328.9170353 | 3.64E-09 | -0.75648 | 7.87E-10 | -6.14754 | 0.591941 | 8.438313 |
| CNAG_00: | - | hypothetic | 152.5450359 | 255.5025898 | 233.7208032 | 408.3832231 | 441.2578755 | 441.4526817 | 3.67E-09 | 0.997644 | 7.93E-10 | 6.146353 | 1.996737 | 8.435307 |
| CNAG_06: | - | Alpha-glu | 507.1513033 | 530.2323439 | 488.5898702 | 743.581914 | 797.3182648 | 805.7126426 | 3.81E-09 | 0.60576 | 8.24E-10 | 6.140312 | 1.52178 | 8.419518 |
| CNAG_06( | - | MFS trans | 576.4316829 | 579.4510324 | 528.8579361 | 349.7494878 | 316.1756166 | 369.8790796 | 3.93E-09 | -0.71773 | 8.52E-10 | -6.13499 | 0.608053 | 8.405217 |
| CNAG_00( | - | hypothetic | 531.983528 | 468.5697654 | 514.922903 | 285.8105298 | 319.7365575 | 305.5461656 | 3.95E-09 | -0.75091 | 8.55E-10 | -6.13432 | 0.594229 | 8.403874 |
| CNAG_02 | - | thaumatin | 14.79763358 | 27.25889203 | 44.98040458 | 118.6708232 | 133.8019948 | 97.41003277 | 3.96E-09 | 2.018581 | 8.59E-10 | 6.133674 | 4.051851 | 8.402359 |

| | | | | | | | | | | | | | | |
|---|---|---|---|---|---|---|---|---|---|---|---|---|---|---|
| CNAG_064 | - | benzodiaz | 2305.781682 | 2951.275363 | 1965.77007 | 6126.447641 | 4951.457805 | 5262.893383 | 4.07E-09 | 1.164653 | 8.84E-10 | 6.129034 | 2.241793 | 8.390178 |
| CNAG_034 | - | exportin-1 | 4002.739594 | 3839.86288 | 3578.979108 | 2845.863976 | 2986.890346 | 2888.796433 | 4.10E-09 | -0.40459 | 8.91E-10 | -6.12787 | 0.755451 | 8.387489 |
| CNAG_061 | - | hypothetic | 958.4117357 | 925.4025466 | 960.7485125 | 648.6777618 | 659.6181672 | 660.7373354 | 4.10E-09 | -0.54682 | 8.90E-10 | -6.12794 | 0.684526 | 8.387489 |
| CNAG_015 | - | hypothetic | 2375.908383 | 2128.898493 | 1993.158186 | 1497.783131 | 1617.119836 | 1527.776622 | 4.11E-09 | -0.50082 | 8.95E-10 | -6.12714 | 0.706704 | 8.385979 |
| CNAG_026 | VPS15 | VPS15 prc | 2402.974276 | 2418.709803 | 2250.163982 | 1742.23785 | 1831.052887 | 1769.758851 | 4.29E-09 | -0.41952 | 9.33E-10 | -6.12047 | 0.747672 | 8.368031 |
| CNAG_078 | - | transcripti | 333.3826553 | 417.1306303 | 441.6258875 | 730.8327709 | 628.1659273 | 663.6251168 | 4.29E-09 | 0.748411 | 9.34E-10 | 6.120304 | 1.679942 | 8.367832 |
| CNAG_073 | - | hypothetic | 3209.739062 | 2796.953408 | 2871.88671 | 2178.868103 | 2149.273862 | 2230.464002 | 4.29E-09 | -0.45264 | 9.35E-10 | -6.12007 | 0.730703 | 8.367434 |
| CNAG_025 | - | transferas | 3087.246841 | 2715.108177 | 3363.405261 | 2224.013011 | 2167.377303 | 2162.786638 | 4.32E-09 | -0.49998 | 9.42E-10 | -6.11895 | 0.707117 | 8.364618 |
| CNAG_006 | - | ATP-bindi | 141.8003737 | 97.3117286 | 163.314609 | 290.2292015 | 327.587284 | 277.2573645 | 4.45E-09 | 1.139683 | 9.71E-10 | 6.114045 | 2.203326 | 8.351509 |
| CNAG_003 | - | porphobili | 2474.245425 | 2508.148 | 2399.455789 | 1887.055125 | 1856.890915 | 1906.285665 | 4.50E-09 | -0.40092 | 9.84E-10 | -6.11196 | 0.757373 | 8.346565 |
| CNAG_004 | - | RNA-bind | 2718.203091 | 2443.563569 | 2683.079059 | 1855.818753 | 1881.009584 | 2031.715353 | 4.54E-09 | -0.45945 | 9.93E-10 | -6.11058 | 0.727265 | 8.343297 |
| CNAG_007 | CAS35 | capsular a | 5328.354232 | 5652.113491 | 5738.490466 | 4405.629249 | 4376.458612 | 4459.640605 | 4.57E-09 | -0.35152 | 1.00E-09 | -6.10935 | 0.78376 | 8.340185 |
| CNAG_051 | - | hypothetic | 21.75266567 | 36.89507224 | 36.4869207 | 96.36293986 | 105.7809761 | 126.9438029 | 5.07E-09 | 1.800127 | 1.11E-09 | 6.092238 | 3.482508 | 8.294896 |
| CNAG_067 | - | DNA repa | 260.8954401 | 271.7200192 | 305.1342729 | 438.4692684 | 498.0941064 | 523.6036875 | 5.29E-09 | 0.788186 | 1.16E-09 | 6.085102 | 1.726901 | 8.276265 |
| CNAG_063 | - | mitochonc | 1144.016897 | 1103.150492 | 1038.652282 | 728.7942868 | 804.5842455 | 748.4576311 | 5.42E-09 | -0.54228 | 1.19E-09 | -6.08118 | 0.686685 | 8.265892 |
| CNAG_055 | - | phosphati | 389.2908192 | 441.8794117 | 416.0585051 | 644.0731976 | 655.8418143 | 664.7315003 | 5.59E-09 | 0.640976 | 1.23E-09 | 6.075953 | 1.559384 | 8.252207 |
| CNAG_043 | SOD2 | mitochonc | 2179.194587 | 1786.576922 | 1934.676299 | 2649.504165 | 3052.106749 | 2848.478138 | 5.72E-09 | 0.519024 | 1.26E-09 | 6.072216 | 1.432985 | 8.242815 |
| CNAG_063 | - | geranylge | 1185.034972 | 1288.421244 | 1155.961944 | 860.4202642 | 886.5162037 | 819.5212542 | 5.89E-09 | -0.51512 | 1.30E-09 | -6.06735 | 0.699733 | 8.229902 |
| CNAG_037 | ALF1 | tubulin-fo | 916.462656 | 972.7630219 | 961.8142831 | 679.8872225 | 619.8792433 | 664.0639133 | 5.95E-09 | -0.55305 | 1.31E-09 | -6.06574 | 0.681578 | 8.225773 |
| CNAG_033 | - | phospholi | 340.0572971 | 229.7816309 | 266.8131025 | 490.0250911 | 555.1555842 | 509.272151 | 5.95E-09 | 0.879294 | 1.32E-09 | 6.065503 | 1.839475 | 8.225386 |
| CNAG_048 | CIR1 | iron regul | 1818.384342 | 1641.08148 | 1690.444311 | 2265.044476 | 2493.959354 | 2365.588503 | 5.99E-09 | 0.452435 | 1.32E-09 | 6.064504 | 1.368348 | 8.222925 |
| CNAG_044 | - | splicing fa | 1898.604932 | 1989.901703 | 2065.590332 | 1355.318574 | 1530.025198 | 1482.220346 | 6.27E-09 | -0.46194 | 1.39E-09 | -6.05705 | 0.72601 | 8.203047 |
| CNAG_060 | - | hypothetic | 423.4890895 | 417.869605 | 435.9938449 | 274.3425559 | 239.4271814 | 232.3338276 | 6.50E-09 | -0.7905 | 1.44E-09 | -6.05104 | 0.578143 | 8.187055 |
| CNAG_052 | CAT2 | catalase 2 | 1250.248395 | 553.1842254 | 566.6817243 | 4324.300301 | 4325.057944 | 4152.365327 | 6.69E-09 | 2.41488 | 1.48E-09 | 6.046274 | 5.33275 | 8.174698 |
| CNAG_047 | - | hypothetic | 355.3405191 | 322.3592082 | 356.3305285 | 557.7450375 | 566.1691294 | 558.1677514 | 6.71E-09 | 0.686845 | 1.49E-09 | 6.04573 | 1.60976 | 8.173472 |
| CNAG_020 | - | hypothetic | 211.9049669 | 268.4228025 | 241.1865069 | 423.9655596 | 416.9441147 | 433.7554643 | 6.78E-09 | 0.806665 | 1.51E-09 | 6.04379 | 1.749163 | 8.168485 |
| CNAG_073 | - | calcium ac | 1319.274688 | 1369.249117 | 1227.430288 | 947.6137842 | 948.6052575 | 910.5358897 | 7.00E-09 | -0.49603 | 1.55E-09 | -6.0386 | 0.709053 | 8.154762 |
| CNAG_023 | - | V-type prc | 4309.976475 | 4364.576545 | 4170.845877 | 3457.713629 | 3331.887967 | 3365.037633 | 7.52E-09 | -0.35419 | 1.67E-09 | -6.02668 | 0.782308 | 8.123662 |
| CNAG_009 | - | protein m | 2586.223737 | 2370.271996 | 2391.979924 | 1837.227595 | 1905.242913 | 1717.632788 | 7.52E-09 | -0.44425 | 1.67E-09 | -6.02671 | 0.734967 | 8.123662 |
| CNAG_078 | - | hypothetic | 272.5059929 | 258.8011904 | 297.6689081 | 487.0780654 | 447.0578808 | 478.1861787 | 7.53E-09 | 0.756367 | 1.68E-09 | 6.026475 | 1.689232 | 8.12335 |
| CNAG_069 | - | beta-fruct | 718.354646 | 575.153085 | 502.2384386 | 391.1712731 | 272.9398167 | 272.3836539 | 7.53E-09 | -0.95681 | 1.68E-09 | -6.02638 | 0.515194 | 8.123326 |
| CNAG_051 | - | prefoldin | 1555.035503 | 1341.350424 | 1556.858272 | 1037.86991 | 1084.921016 | 895.0854472 | 7.60E-09 | -0.57804 | 1.70E-09 | -6.02456 | 0.669875 | 8.118915 |
| CNAG_058 | - | protein R/ | 1235.013918 | 1070.887939 | 1109.016234 | 794.2492311 | 786.4068736 | 781.7573483 | 7.69E-09 | -0.54787 | 1.72E-09 | -6.02266 | 0.684028 | 8.114281 |
| CNAG_053 | - | hypothetic | 5945.94102 | 5675.839187 | 5593.494614 | 4439.056264 | 4591.257995 | 4600.629192 | 7.99E-09 | -0.35214 | 1.79E-09 | -6.01623 | 0.78342 | 8.097291 |
| CNAG_016 | - | hypothetic | 980.0784976 | 1050.610487 | 927.9720372 | 1293.614299 | 1607.372597 | 1516.228638 | 8.00E-09 | 0.563604 | 1.79E-09 | 6.016085 | 1.477956 | 8.09713 |
| CNAG_011 | - | Aspartyl a | 707.5279165 | 811.1216041 | 832.7184023 | 510.8764681 | 516.3992819 | 525.3125962 | 8.00E-09 | -0.61426 | 1.79E-09 | -6.016 | 0.653266 | 8.097129 |
| CNAG_073 | MET22 | 3'(2'),5'-bi: | 915.1694746 | 889.8251061 | 888.2420151 | 565.2879979 | 605.2545322 | 654.0307712 | 8.21E-09 | -0.57685 | 1.84E-09 | -6.01156 | 0.670426 | 8.085486 |
| CNAG_050 | - | iron dono | 994.6953067 | 1010.511231 | 1084.421375 | 751.5269825 | 715.6869101 | 651.9275857 | 8.23E-09 | -0.56 | 1.84E-09 | -6.01124 | 0.6783 | 8.08485 |
| CNAG_044 | - | succinate- | 731.9208499 | 754.0549636 | 783.6842404 | 466.4313542 | 499.1491776 | 535.2597253 | 8.57E-09 | -0.61158 | 1.92E-09 | -6.00434 | 0.654477 | 8.066858 |
| CNAG_022 | - | hypothetic | 1265.589212 | 1236.795926 | 1354.239192 | 877.4831859 | 978.8591929 | 836.2008926 | 8.83E-09 | -0.53425 | 1.98E-09 | -5.99945 | 0.690519 | 8.05423 |
| CNAG_032 | - | splicing fa | 3105.946193 | 3088.877165 | 2858.049886 | 2173.361578 | 2367.574108 | 2340.355109 | 9.29E-09 | -0.41089 | 2.09E-09 | -5.99101 | 0.75216 | 8.032147 |
| CNAG_011 | - | ribosomal | 1732.706502 | 1655.944093 | 1876.797718 | 1185.64899 | 1266.881231 | 1328.997477 | 9.34E-09 | -0.49341 | 2.10E-09 | -5.98999 | 0.710343 | 8.029658 |
| CNAG_031 | - | hypothetic | 659.1284275 | 508.498171 | 650.291261 | 398.1404478 | 290.2052466 | 332.2412977 | 9.35E-09 | -0.8492 | 2.10E-09 | -5.98967 | 0.555092 | 8.029039 |
| CNAG_052 | - | clathrin bi | 3115.283369 | 3015.634393 | 2876.175456 | 2281.626625 | 2302.849694 | 2352.562675 | 9.37E-09 | -0.39228 | 2.11E-09 | -5.98924 | 0.761923 | 8.028133 |
| CNAG_068 | - | hypothetic | 528.5949478 | 484.7418718 | 557.5511488 | 325.6189529 | 317.9635504 | 326.6249779 | 9.46E-09 | -0.71241 | 2.13E-09 | -5.98773 | 0.610299 | 8.024333 |
| CNAG_010 | - | hypothetic | 21.66271761 | 19.70420856 | 29.00947371 | 80.90624242 | 90.3535377 | 94.88862766 | 9.75E-09 | 1.920162 | 2.19E-09 | 5.982716 | 3.784655 | 8.011201 |
| CNAG_025 | - | DNA repa | 1998.38593 | 2250.852984 | 2278.020685 | 2879.15649 | 2919.090306 | 3029.433071 | 9.87E-09 | 0.420934 | 2.22E-09 | 5.980631 | 1.338794 | 8.005876 |
| CNAG_003 | SIP4 | hypothetic | 325.0202373 | 359.9854788 | 262.5788522 | 523.4658273 | 545.4919118 | 563.6316888 | 9.91E-09 | 0.771303 | 2.23E-09 | 5.979762 | 1.70681 | 8.004029 |
| CNAG_028 | - | hypothetic | 7.622929785 | 11.04449594 | 8.775718585 | 52.9763361 | 71.80660979 | 48.54617268 | 1.00E-08 | 2.702637 | 2.27E-09 | 5.977373 | 6.509908 | 7.997897 |
| CNAG_066 | ERG5 | C-22 sterc | 830.6726764 | 675.4160019 | 635.5377598 | 452.5214264 | 379.1015982 | 452.0560692 | 1.04E-08 | -0.75382 | 2.34E-09 | -5.97216 | 0.593033 | 7.984473 |
| CNAG_001 | - | ubiquitin | 1052.984267 | 1163.373258 | 932.0432097 | 747.6121241 | 689.7652461 | 665.2407113 | 1.10E-08 | -0.5975 | 2.49E-09 | -5.96181 | 0.660898 | 7.957171 |
| CNAG_043 | - | vesicle tra | 1228.132264 | 1115.072398 | 1229.483873 | 871.3696836 | 837.308941 | 802.8705875 | 1.10E-08 | -0.52477 | 2.50E-09 | -5.96158 | 0.695069 | 7.956804 |
| CNAG_059 | - | hypothetic | 1424.394221 | 1429.627779 | 1509.945376 | 1138.471689 | 943.3357053 | 892.8544393 | 1.11E-08 | -0.56836 | 2.51E-09 | -5.96059 | 0.674383 | 7.954408 |
| CNAG_004 | - | hypothetic | 22.95424354 | 43.3590606 | 47.1297486 | 110.4075824 | 123.0888967 | 135.9118894 | 1.18E-08 | 1.70242 | 2.66E-09 | 5.951112 | 3.254464 | 7.929698 |
| CNAG_014 | - | hypothetic | 65.9234392 | 70.2575987 | 50.39239742 | 158.8542617 | 146.1577028 | 168.233548 | 1.19E-08 | 1.336714 | 2.70E-09 | 5.949152 | 2.525754 | 7.924732 |
| CNAG_036 | - | UDP-N-ac | 616.0595952 | 554.7440375 | 551.2466623 | 370.8012723 | 349.8721152 | 368.7970227 | 1.21E-08 | -0.67842 | 2.74E-09 | -5.94669 | 0.624847 | 7.918423 |
| CNAG_126 | - | unspecifie | 43.80842038 | 46.59392242 | 41.84012641 | 119.792021 | 156.2369927 | 108.5714442 | 1.25E-08 | 1.543356 | 2.83E-09 | 5.941291 | 2.914717 | 7.904356 |
| CNAG_051 | - | hypothetic | 1235.858739 | 1410.45601 | 1270.282581 | 1670.280891 | 2029.966239 | 2015.774313 | 1.25E-08 | 0.530783 | 2.84E-09 | 5.940768 | 1.444713 | 7.903205 |
| CNAG_000 | - | hypothetic | 56.56120494 | 41.25464231 | 49.29222265 | 145.8378885 | 151.6933689 | 115.2527756 | 1.29E-08 | 1.47672 | 2.93E-09 | 5.935236 | 2.783154 | 7.888789 |
| CNAG_033 | - | hypothetic | 31.06884273 | 53.00391163 | 37.58158644 | 126.3146148 | 130.0649139 | 112.9017451 | 1.29E-08 | 1.590991 | 2.94E-09 | 5.93501 | 3.012563 | 7.888423 |
| CNAG_051 | - | exoribonu | 683.9236444 | 620.5156618 | 721.7916201 | 422.1096565 | 465.5676905 | 407.7024002 | 1.31E-08 | -0.66129 | 2.97E-09 | -5.93309 | 0.632313 | 7.883562 |
| CNAG_045 | FZC27 | hypothetic | 352.0549681 | 449.4348218 | 468.2778372 | 676.0054951 | 705.9292472 | 692.4844591 | 1.42E-08 | 0.693456 | 3.23E-09 | 5.919313 | 1.617152 | 7.848089 |
| CNAG_013 | - | protein NI | 200.1878882 | 232.8888473 | 223.0472316 | 381.9206775 | 389.3951374 | 379.4082326 | 1.42E-08 | 0.796008 | 3.25E-09 | 5.918529 | 1.73629 | 7.846251 |
| CNAG_015 | - | hypothetic | 48.37960568 | 36.92999264 | 36.51655162 | 104.022708 | 134.4749625 | 123.8521198 | 1.43E-08 | 1.572823 | 3.26E-09 | 5.918131 | 2.974863 | 7.845433 |
| CNAG_009 | - | flap endoi | 2030.476286 | 2096.540433 | 1899.306963 | 1501.768798 | 1535.177625 | 1503.321653 | 1.44E-08 | -0.42363 | 3.30E-09 | -5.91608 | 0.745549 | 7.840254 |
| CNAG_062 | CCD6 | Fungal Zn | 675.8534492 | 682.9554068 | 721.8256216 | 486.0906003 | 437.8676272 | 448.7375818 | 1.49E-08 | -0.61518 | 3.40E-09 | -5.91125 | 0.65285 | 7.828207 |
| CNAG_043 | ARO8001 | specific RI | 351.7727542 | 312.6461841 | 293.4633841 | 554.7990469 | 523.8786201 | 511.5734385 | 1.49E-08 | 0.716582 | 3.39E-09 | 5.911304 | 1.643284 | 7.828207 |
| CNAG_075 | - | pyridoxal | 634.0249133 | 795.9823941 | 802.8253797 | 446.1872959 | 512.1522761 | 438.7982455 | 1.53E-08 | -0.69109 | 3.50E-09 | -5.90602 | 0.619387 | 7.814887 |

| | | | | | | | | | | | | | |
|---|---|---|---|---|---|---|---|---|---|---|---|---|---|
| CNAG_011 | URE4 | urease acc | 753.0615978 | 891.890644 | 851.9445515 | 499.0833274 | 595.8423735 | 544.2032092 | 1.55E-08 | -0.62186 | 3.55E-09 | -5.90376 | 0.649832 | 7.809156 |
| CNAG_046 | UBC12 | ubiquitin-( | 426.506087 | 371.9136535 | 360.6446865 | 671.5546654 | 682.7310776 | 563.8651367 | 1.56E-08 | 0.711818 | 3.58E-09 | 5.902312 | 1.637867 | 7.805582 |
| CNAG_055 | - | hypothetic | 3786.775721 | 3522.077558 | 3784.746559 | 2867.707612 | 2866.108749 | 2894.326406 | 1.57E-08 | -0.37819 | 3.59E-09 | -5.90212 | 0.769402 | 7.805317 |
| CNAG_039 | MAP1 | methionin | 6251.859421 | 5474.418102 | 6445.505649 | 4529.312871 | 4662.828961 | 4554.042876 | 1.60E-08 | -0.41868 | 3.66E-09 | -5.89878 | 0.748109 | 7.796738 |
| CNAG_053 | - | hypothetic | 251.4855085 | 267.3668055 | 244.395237 | 450.4998723 | 457.5875529 | 398.393802 | 1.61E-08 | 0.761077 | 3.70E-09 | 5.897133 | 1.694756 | 7.792646 |
| CNAG_006 | - | Cystathior | 282.953126 | 293.2079575 | 231.6383663 | 453.4657219 | 464.4095972 | 469.3021277 | 1.63E-08 | 0.767206 | 3.73E-09 | 5.895555 | 1.701971 | 7.788724 |
| CNAG_042 | - | hypothetic | 53.51123303 | 135.05144 | 135.7584006 | 518.8235274 | 539.9755589 | 596.3633317 | 1.63E-08 | 2.34333 | 3.74E-09 | 5.895187 | 5.074725 | 7.787989 |
| CNAG_037 | - | ankyrin re | 1529.42612 | 1379.055121 | 1563.271756 | 1037.054632 | 1135.84002 | 987.1736422 | 1.66E-08 | -0.5171 | 3.81E-09 | -5.89228 | 0.698775 | 7.780579 |
| CNAG_024 | - | cytoplasm | 1328.578951 | 1296.025518 | 1242.3432 | 880.5782322 | 923.5747 | 975.9660693 | 1.67E-08 | -0.49203 | 3.83E-09 | -5.89118 | 0.711023 | 7.777902 |
| CNAG_017 | CRG3 | Putative re | 675.167408 | 552.9280988 | 504.6435173 | 915.6767211 | 1093.452132 | 865.7963494 | 1.67E-08 | 0.714676 | 3.84E-09 | 5.890891 | 1.641114 | 7.777384 |
| CNAG_042 | - | DNA-direc | 1610.12769 | 1621.436 | 1759.495795 | 1287.064575 | 1195.285961 | 1084.883423 | 1.68E-08 | -0.50014 | 3.87E-09 | -5.88962 | 0.707039 | 7.774277 |
| CNAG_056 | - | ubiquitin-( | 2491.609383 | 2290.528347 | 2379.14779 | 1861.471699 | 1452.176695 | 1713.152827 | 1.72E-08 | -0.52588 | 3.95E-09 | -5.88617 | 0.694533 | 7.765441 |
| CNAG_074 | MBS1 | Mbp1 and | 2815.226511 | 2808.765019 | 2705.533711 | 2205.427126 | 2159.675393 | 2116.12734 | 1.77E-08 | -0.3772 | 4.08E-09 | -5.88107 | 0.76993 | 7.752525 |
| CNAG_041 | - | Glutathior | 502.4591333 | 512.9967097 | 490.7086501 | 746.7774486 | 743.766902 | 772.4292939 | 1.82E-08 | 0.573077 | 4.20E-09 | 5.876245 | 1.487693 | 7.740317 |
| CNAG_078 | - | hypothetic | 89.26636496 | 95.03086673 | 80.21885913 | 208.7396064 | 206.7119005 | 175.1906598 | 1.89E-08 | 1.142671 | 4.37E-09 | 5.869525 | 2.207895 | 7.723163 |
| CNAG_056 | - | hypothetic | 1604.182848 | 1529.85549 | 1512.155206 | 1112.569278 | 1159.95012 | 1166.929694 | 1.99E-08 | -0.44994 | 4.59E-09 | -5.86122 | 0.732074 | 7.701665 |
| CNAG_033 | - | hypothetic | 964.2106292 | 873.7248985 | 972.4585286 | 655.7217882 | 649.264796 | 635.2246755 | 2.01E-08 | -0.55047 | 4.66E-09 | -5.85902 | 0.682799 | 7.69612 |
| CNAG_057 | - | WSC dom | 225.7584645 | 152.2397797 | 227.3033981 | 414.0846566 | 427.5344004 | 351.7489818 | 2.03E-08 | 0.963722 | 4.70E-09 | 5.857585 | 1.950335 | 7.692608 |
| CNAG_063 | BAT2 | Branched- | 3960.914124 | 4386.0893 | 4404.330501 | 2917.342771 | 3499.323255 | 3073.115517 | 2.08E-08 | -0.44128 | 4.82E-09 | -5.85319 | 0.736482 | 7.68159 |
| CNAG_031 | SWE1 | WEE prote | 1242.838259 | 1331.82824 | 1379.010224 | 1715.526215 | 1956.613277 | 1840.471307 | 2.14E-08 | 0.464557 | 4.97E-09 | 5.848295 | 1.379894 | 7.669026 |
| CNAG_077 | - | acyl-CoA ( | 516.4611058 | 526.9983826 | 474.734453 | 731.1087505 | 832.7782471 | 755.8308183 | 2.38E-08 | 0.597387 | 5.53E-09 | 5.830396 | 1.512974 | 7.623058 |
| CNAG_021 | - | clathrin as | 1661.388819 | 1608.488239 | 1510.044482 | 1174.097412 | 1171.142886 | 1193.578795 | 2.40E-08 | -0.44904 | 5.58E-09 | -5.82888 | 0.732529 | 7.619346 |
| CNAG_033 | - | hypothetic | 2092.46421 | 2211.850985 | 2171.212557 | 1629.410047 | 1697.361687 | 1649.852003 | 2.42E-08 | -0.39503 | 5.63E-09 | -5.82726 | 0.760473 | 7.615359 |
| CNAG_074 | - | phosphati | 1997.19505 | 2245.454424 | 2202.310199 | 2957.108538 | 2888.05306 | 2761.978302 | 2.44E-08 | 0.402545 | 5.66E-09 | 5.826406 | 1.321837 | 7.613362 |
| CNAG_031 | - | rhomboid- | 1403.196045 | 1244.3452 | 1254.066638 | 975.7053592 | 862.2646426 | 903.8711961 | 2.47E-08 | -0.52527 | 5.74E-09 | -5.82424 | 0.694827 | 7.607954 |
| CNAG_010 | - | hypothetic | 537.0400939 | 613.9574687 | 684.4110968 | 386.3461268 | 402.5682583 | 343.3509766 | 2.47E-08 | -0.71251 | 5.75E-09 | -5.82387 | 0.610256 | 7.607217 |
| CNAG_016 | - | hypothetic | 1203.026051 | 1228.371332 | 1087.92777 | 1647.235537 | 1671.036293 | 1571.863499 | 2.53E-08 | 0.459677 | 5.88E-09 | 5.820072 | 1.375234 | 7.597579 |
| CNAG_046 | - | hypothetic | 20.43945701 | 18.60264667 | 16.25527504 | 82.93697019 | 88.87072146 | 59.76971944 | 2.56E-08 | 2.068492 | 5.98E-09 | 5.817343 | 4.194479 | 7.590945 |
| CNAG_027 | - | DNA repa | 1575.503858 | 1534.375643 | 1325.76775 | 1986.424707 | 2065.174491 | 2241.139162 | 2.69E-08 | 0.489827 | 6.28E-09 | 5.809158 | 1.404276 | 7.570382 |
| CNAG_030 | - | potassium | 622.9050951 | 776.9406329 | 775.4086373 | 1090.037405 | 1058.580476 | 1117.683631 | 2.69E-08 | 0.572964 | 6.28E-09 | 5.809052 | 1.487576 | 7.570334 |
| CNAG_004 | - | hypothetic | 746.4548298 | 730.6324299 | 530.2696359 | 1218.087018 | 1068.990198 | 1010.099915 | 2.73E-08 | 0.701415 | 6.38E-09 | 5.806463 | 1.626099 | 7.563846 |
| CNAG_079 | - | hypothetic | 1271.39013 | 1242.166343 | 1223.136826 | 911.0361228 | 875.2459061 | 933.7934657 | 2.82E-08 | -0.47364 | 6.61E-09 | -5.80058 | 0.720145 | 7.54929 |
| CNAG_013 | - | stress resp | 3232.072888 | 2924.116979 | 3313.34534 | 2349.311944 | 2458.997864 | 2381.455512 | 2.84E-08 | -0.41346 | 6.66E-09 | -5.7992 | 0.750819 | 7.546169 |
| CNAG_017 | - | glycerol-3 | 3808.445313 | 2892.838938 | 2798.34011 | 2276.963123 | 2111.275148 | 1968.58855 | 2.90E-08 | -0.59617 | 6.81E-09 | -5.79557 | 0.661506 | 7.536992 |
| CNAG_017 | - | hsp71-like | 95325.6708 | 85444.67092 | 91537.8825 | 73138.80422 | 70976.51222 | 73742.56034 | 2.93E-08 | -0.3376 | 6.88E-09 | -5.79393 | 0.791358 | 7.53297 |
| CNAG_013 | - | hypothetic | 403.297535 | 465.569469 | 425.6594311 | 708.0804683 | 642.8220009 | 660.3392388 | 3.03E-08 | 0.621852 | 7.11E-09 | 5.788408 | 1.538849 | 7.519143 |
| CNAG_025 | - | avenacina | 258.5920387 | 305.0771994 | 310.4706812 | 465.7983555 | 549.9888054 | 468.2925801 | 3.04E-08 | 0.748492 | 7.14E-09 | 5.787605 | 1.680036 | 7.51752 |
| CNAG_001 | - | exosome ( | 1062.472835 | 1120.397777 | 1135.648634 | 819.2291784 | 792.4656659 | 775.0931451 | 3.04E-08 | -0.49037 | 7.14E-09 | -5.78766 | 0.711842 | 7.51752 |
| CNAG_059 | - | U2 small r | 2553.638144 | 2465.088102 | 2666.005163 | 2002.306827 | 1708.420831 | 1957.351161 | 3.04E-08 | -0.45453 | 7.16E-09 | -5.78711 | 0.729748 | 7.516461 |
| CNAG_059 | - | DNA-direc | 7097.361758 | 6913.798685 | 6669.553129 | 5222.753461 | 5575.450162 | 5659.702332 | 3.11E-08 | -0.34487 | 7.32E-09 | -5.78344 | 0.787378 | 7.507211 |
| CNAG_064 | - | DNA depe | 1581.394978 | 1633.486739 | 1586.983221 | 2068.14027 | 2126.395146 | 2195.710375 | 3.27E-08 | 0.397491 | 7.71E-09 | 5.774606 | 1.317215 | 7.485533 |
| CNAG_016 | - | hypothetic | 4319.285339 | 4218.063175 | 4374.490588 | 2904.854174 | 3428.545293 | 3418.296917 | 3.41E-08 | -0.42059 | 8.06E-09 | -5.76725 | 0.747119 | 7.466794 |
| CNAG_011 | - | hypothetic | 11.1133843 | 14.2663259 | 7.721744962 | 71.57445263 | 46.82957722 | 64.91074716 | 3.45E-08 | 2.486112 | 8.15E-09 | 5.765358 | 5.60266 | 7.462149 |
| CNAG_038 | - | hypothetic | 35.74385754 | 53.04075282 | 54.59355372 | 152.2898852 | 136.8722875 | 113.0265659 | 3.48E-08 | 1.483309 | 8.22E-09 | 5.763914 | 2.795893 | 7.458656 |
| CNAG_000 | - | vacuolar p | 1705.12394 | 1629.201211 | 1517.70401 | 2054.02592 | 2352.494946 | 2347.760343 | 3.53E-08 | 0.461857 | 8.35E-09 | 5.761145 | 1.377314 | 7.451754 |
| CNAG_022 | CRG1 | xap5-dom | 2158.995472 | 2281.86902 | 2143.506053 | 1638.733043 | 1757.764228 | 1645.427017 | 3.56E-08 | -0.40017 | 8.41E-09 | -5.75995 | 0.757769 | 7.448896 |
| CNAG_043 | - | hypothetic | 0.44683804 | 7.787378998 | 5.554625618 | 53.6774883 | 46.39517519 | 36.37560866 | 3.62E-08 | 3.479192 | 8.57E-09 | 5.756917 | 11.15171 | 7.441772 |
| CNAG_066 | - | hypothetic | 1226.400576 | 1290.852345 | 1146.574085 | 1720.421015 | 1694.291879 | 1620.714626 | 3.62E-08 | 0.444103 | 8.59E-09 | 5.756508 | 1.360468 | 7.440944 |
| CNAG_041 | - | histone H1 | 1162.440679 | 1479.43385 | 1554.964335 | 2237.430254 | 2106.652285 | 1926.024086 | 3.76E-08 | 0.565027 | 8.93E-09 | 5.749905 | 1.479415 | 7.424422 |
| CNAG_029 | - | flavoprote | 348.3070864 | 341.7028031 | 306.2527531 | 546.9130816 | 553.2649344 | 506.0642173 | 3.76E-08 | 0.674516 | 8.93E-09 | 5.749982 | 1.596061 | 7.424422 |
| CNAG_020 | - | acetoaceta | 116.1479395 | 133.7963849 | 116.457319 | 283.5953049 | 265.3373192 | 208.6076406 | 3.84E-08 | 1.035569 | 9.12E-09 | 5.746245 | 2.049922 | 7.415247 |
| CNAG_000 | MLR1 | hypothetic | 694.3915291 | 638.8025383 | 671.7267519 | 460.3888212 | 455.1982026 | 386.6418559 | 3.86E-08 | -0.6383 | 9.17E-09 | -5.74535 | 0.642472 | 7.413387 |
| CNAG_005 | - | hypothetic | 38.14425548 | 63.82868917 | 79.07785217 | 165.1551458 | 172.3118388 | 150.6551681 | 3.86E-08 | 1.421446 | 9.18E-09 | 5.745134 | 2.678538 | 7.413065 |
| CNAG_128 | - | unspecifie | 127.4270881 | 122.3995627 | 85.23797064 | 41.311686 | 29.62646557 | 38.12158106 | 3.96E-08 | -1.64354 | 9.42E-09 | -5.74079 | 0.320071 | 7.402377 |
| CNAG_033 | - | NifU-like p | 2829.235817 | 2641.818975 | 2936.89243 | 2123.631878 | 2162.260939 | 2162.73739 | 3.99E-08 | -0.39843 | 9.51E-09 | -5.73923 | 0.758682 | 7.398797 |
| CNAG_061 | - | hypothetic | 888.9534951 | 835.1805813 | 925.777299 | 1183.193693 | 1275.103785 | 1369.647351 | 4.00E-08 | 0.51581 | 9.55E-09 | 5.738475 | 1.429797 | 7.397542 |
| CNAG_079 | - | hypothetic | 12.22368794 | 5.677270233 | 9.832538175 | 65.33455931 | 43.40390467 | 58.27757081 | 4.00E-08 | 2.658101 | 9.55E-09 | 5.738542 | 6.312018 | 7.397542 |
| CNAG_057 | - | hypothetic | 1375.348648 | 1334.84431 | 1375.620305 | 956.1014312 | 1001.222818 | 1042.573973 | 4.00E-08 | -0.46086 | 9.54E-09 | -5.73864 | 0.726555 | 7.397542 |
| CNAG_051 | - | U3 small r | 1193.116423 | 1083.833205 | 1223.071897 | 762.2251388 | 840.7857334 | 854.9786419 | 4.03E-08 | -0.52638 | 9.62E-09 | -5.73736 | 0.694296 | 7.394908 |
| CNAG_012 | - | arginine b | 1989.73516 | 2103.04687 | 2218.093324 | 1609.989371 | 1628.346394 | 1537.758592 | 4.06E-08 | -0.41763 | 9.69E-09 | -5.73607 | 0.748656 | 7.391824 |
| CNAG_034 | - | BET3 fami | 1564.502418 | 1429.701031 | 1642.18466 | 1142.213487 | 1152.194959 | 1043.782939 | 4.16E-08 | -0.48987 | 9.94E-09 | -5.73169 | 0.712088 | 7.38105 |
| CNAG_033 | ZIP2 | solute car | 12.30028103 | 14.28482043 | 15.16402735 | 59.10914442 | 79.48931157 | 57.40599898 | 4.17E-08 | 2.234324 | 9.98E-09 | 5.731098 | 4.705422 | 7.37975 |
| CNAG_005 | - | hypothetic | 33.39367541 | 54.08404401 | 34.40047153 | 94.3721217 | 132.6447546 | 155.7198326 | 4.23E-08 | 1.641504 | 1.01E-08 | 5.72881 | 3.119909 | 7.374112 |
| CNAG_015 | - | splicing fa | 2526.758698 | 2315.360318 | 2602.02056 | 1837.191588 | 1932.837854 | 1858.574152 | 4.24E-08 | -0.4193 | 1.02E-08 | -5.72807 | 0.747787 | 7.372431 |
| CNAG_060 | SAC1 | inositol/ph | 4478.115094 | 4346.289954 | 4337.205265 | 3416.387577 | 3564.82376 | 3539.313198 | 4.24E-08 | -0.33866 | 1.02E-08 | -5.7279 | 0.790776 | 7.372221 |
| CNAG_052 | - | hypothetic | 889.4625898 | 801.5366819 | 895.6704472 | 577.048818 | 622.5728451 | 569.7418362 | 4.38E-08 | -0.56389 | 1.05E-08 | -5.72225 | 0.676473 | 7.358211 |
| CNAG_051 | - | ubiquinol- | 1912.443423 | 1943.51775 | 1645.518911 | 1364.025481 | 1348.836581 | 1309.072601 | 4.41E-08 | -0.46689 | 1.06E-08 | -5.72124 | 0.723523 | 7.355853 |
| CNAG_036 | - | hypothetic | 57.89991422 | 96.04241374 | 61.04842003 | 175.1435841 | 169.5014513 | 184.9181711 | 4.50E-08 | 1.289791 | 1.08E-08 | 5.717497 | 2.444926 | 7.346508 |

| | | | | | | | | | | | | | | |
|---|---|---|---|---|---|---|---|---|---|---|---|---|---|---|
| CNAG_042 | - | bromodor | 2364.555882 | 2440.291097 | 2492.205019 | 1886.283301 | 1925.078354 | 1871.883504 | 4.59E-08 | -0.3759 | 1.10E-08 | -5.71401 | 0.770622 | 7.337834 |
| CNAG_014 | - | hypothetic | 14.7373313 | 32.55653579 | 25.83604017 | 74.39696943 | 91.17900727 | 109.1344826 | 4.63E-08 | 1.935307 | 1.11E-08 | 5.712447 | 3.824596 | 7.334054 |
| CNAG_069 | - | hypothetic | 2182.256852 | 1952.279057 | 2359.887374 | 1504.00681 | 1638.692311 | 1567.725529 | 4.64E-08 | -0.47945 | 1.11E-08 | -5.71231 | 0.717251 | 7.333936 |
| CNAG_037 | - | pantetheir | 1114.80067 | 1001.926479 | 1033.296144 | 774.0961241 | 726.0213451 | 718.4866894 | 4.64E-08 | -0.52082 | 1.12E-08 | -5.712 | 0.696974 | 7.333344 |
| CNAG_013 | - | Nucleolar | 1232.885321 | 1158.174798 | 1321.174941 | 902.4854684 | 908.9418294 | 833.9599318 | 4.81E-08 | -0.50535 | 1.16E-08 | -5.7058 | 0.704491 | 7.317965 |
| CNAG_062 | MIZ1 | E3 SUMO- | 1348.534961 | 1361.76471 | 1380.951776 | 996.6371357 | 1060.800897 | 957.1672968 | 4.90E-08 | -0.45591 | 1.18E-08 | -5.70261 | 0.729051 | 7.310055 |
| CNAG_042 | - | hypothetic | 177.9205307 | 182.2702504 | 165.4944059 | 299.2893047 | 310.8488062 | 347.0581333 | 4.93E-08 | 0.852509 | 1.19E-08 | 5.701411 | 1.805638 | 7.307227 |
| CNAG_060 | ERG8 | phosphom | 564.6965185 | 512.7235901 | 525.626318 | 325.5875619 | 296.3234562 | 370.9479251 | 5.04E-08 | -0.70736 | 1.22E-08 | -5.69734 | 0.612438 | 7.297295 |
| CNAG_031 | - | hypothetic | 13.381729 | 8.901574998 | 9.836027825 | 57.08065898 | 51.36555326 | 59.39041495 | 5.34E-08 | 2.460842 | 1.29E-08 | 5.687249 | 5.505378 | 7.272282 |
| CNAG_065 | - | hypothetic | 2116.568189 | 1287.388861 | 1323.323084 | 601.5104236 | 570.4368694 | 585.50661 | 5.54E-08 | -1.44575 | 1.34E-08 | -5.6808 | 0.367101 | 7.256122 |
| CNAG_031 | - | hypothetic | 62.38679303 | 55.21508552 | 56.74402838 | 143.2440982 | 137.5806545 | 146.1117556 | 5.64E-08 | 1.284633 | 1.36E-08 | 5.677735 | 2.436201 | 7.248553 |
| CNAG_052 | RAD53 | CAMK pro | 1409.777698 | 1547.274138 | 1433.424151 | 1973.278045 | 2060.061163 | 1888.284582 | 5.66E-08 | 0.417007 | 1.37E-08 | 5.67722 | 1.335155 | 7.247464 |
| CNAG_047 | - | ATP-depe | 1682.972001 | 1664.751126 | 1696.818476 | 2131.134681 | 2332.619398 | 2246.812169 | 5.74E-08 | 0.396718 | 1.39E-08 | 5.674461 | 1.31651 | 7.240901 |
| CNAG_028 | - | nuclear po | 6243.732747 | 5893.496704 | 6082.976012 | 4658.519571 | 5104.541884 | 4650.627959 | 5.79E-08 | -0.35378 | 1.40E-08 | -5.67283 | 0.782533 | 7.236977 |
| CNAG_056 | - | YbgI/famil | 1430.215766 | 1319.796639 | 1489.690488 | 1076.889243 | 1008.961461 | 980.4837547 | 5.83E-08 | -0.48365 | 1.41E-08 | -5.6716 | 0.715166 | 7.234072 |
| CNAG_062 | - | MIF4G/MA | 1571.466069 | 1541.672462 | 1454.573303 | 1104.046467 | 988.1696848 | 1185.76004 | 6.14E-08 | -0.49355 | 1.49E-08 | -5.66276 | 0.710274 | 7.212123 |
| CNAG_049 | - | hypothetic | 1130.645141 | 1159.422373 | 1123.082314 | 1536.658428 | 1537.278691 | 1536.307625 | 6.39E-08 | 0.418719 | 1.55E-08 | 5.655612 | 1.33674 | 7.194256 |
| CNAG_041 | - | hypothetic | 2.849076744 | 4.551694925 | 4.474251104 | 44.10591015 | 34.05260015 | 39.48310516 | 6.58E-08 | 3.534217 | 1.60E-08 | 5.650614 | 11.58525 | 7.181841 |
| CNAG_036 | - | heterokary | 1959.711771 | 2011.652731 | 1852.547497 | 2540.607419 | 2666.42171 | 2478.887385 | 6.71E-08 | 0.385184 | 1.63E-08 | 5.646837 | 1.306027 | 7.172954 |
| CNAG_021 | - | hypothetic | 350.8883977 | 464.4991163 | 459.7544101 | 720.5563673 | 701.6087463 | 640.4090164 | 6.71E-08 | 0.680686 | 1.63E-08 | 5.646858 | 1.602902 | 7.172954 |
| CNAG_020 | - | DNA-direc | 1081.086028 | 1113.92056 | 1096.202972 | 826.2839743 | 799.3960134 | 679.7033138 | 6.87E-08 | -0.52923 | 1.67E-08 | -5.64274 | 0.692925 | 7.162832 |
| CNAG_033 | - | hypothetic | 2.932438941 | 13.1713418 | 8.768309419 | 67.04616358 | 54.10121596 | 43.06669868 | 6.94E-08 | 2.821776 | 1.69E-08 | 5.641028 | 7.070323 | 7.158729 |
| CNAG_123 | - | unspecifie | 15.85257229 | 22.89586806 | 24.74851491 | 65.03002079 | 100.2366808 | 76.20394222 | 6.97E-08 | 1.967146 | 1.70E-08 | 5.640275 | 3.909939 | 7.157047 |
| CNAG_010 | - | hypothetic | 609.0445919 | 540.746894 | 554.4293234 | 402.8735565 | 320.4848947 | 341.0882486 | 7.53E-08 | -0.69549 | 1.84E-08 | -5.62656 | 0.617498 | 7.123361 |
| CNAG_065 | - | allantoate | 12.31574702 | 17.50240184 | 14.11100732 | 62.25336953 | 51.09199816 | 92.26166109 | 7.54E-08 | 2.239666 | 1.84E-08 | 5.626287 | 4.722877 | 7.122893 |
| CNAG_061 | FZC15 | hypothetic | 82.4559776 | 131.624655 | 127.0740475 | 227.0333297 | 250.6620827 | 248.2767976 | 7.61E-08 | 1.07848 | 1.86E-08 | 5.624485 | 2.11181 | 7.118792 |
| CNAG_042 | - | transcripti | 1087.088973 | 1256.115159 | 1243.346444 | 805.8474102 | 864.9348075 | 886.0588091 | 7.75E-08 | -0.50377 | 1.90E-08 | -5.62128 | 0.705262 | 7.110953 |
| CNAG_051 | - | V-type pro | 698.0181998 | 760.4452031 | 654.7336417 | 499.3310855 | 452.5417857 | 462.0535534 | 7.91E-08 | -0.59456 | 1.94E-08 | -5.61754 | 0.662248 | 7.101761 |
| CNAG_003 | - | gamma-tu | 2139.575354 | 2321.930659 | 2011.462656 | 2907.248015 | 2781.89583 | 2914.00386 | 7.95E-08 | 0.396003 | 1.95E-08 | 5.616512 | 1.315857 | 7.099614 |
| CNAG_058 | - | hypothetic | 758.6543783 | 697.0004121 | 744.2480491 | 526.5905852 | 500.9025714 | 449.8917661 | 8.08E-08 | -0.59183 | 1.98E-08 | -5.61353 | 0.663501 | 7.092339 |
| CNAG_064 | - | cytochrom | 2539.473905 | 2266.845365 | 2391.951277 | 1910.52966 | 1736.118144 | 1501.224995 | 8.18E-08 | -0.49936 | 2.01E-08 | -5.61132 | 0.707421 | 7.087227 |
| CNAG_028 | - | hypothetic | 91.7027765 | 135.8884449 | 84.5143146 | 200.4014861 | 236.8662702 | 254.8023963 | 8.27E-08 | 1.140888 | 2.03E-08 | 5.609294 | 2.205167 | 7.082357 |
| CNAG_131 | - | unspecifie | 322.6236222 | 166.9043749 | 160.9752873 | 97.44754149 | 75.37675023 | 49.37743454 | 8.35E-08 | -1.57022 | 2.05E-08 | -5.6075 | 0.336756 | 7.078298 |
| CNAG_062 | - | Amidase | 831.237935 | 867.2015819 | 894.616105 | 609.8046859 | 633.7910214 | 577.5128567 | 8.47E-08 | -0.52616 | 2.09E-08 | -5.60477 | 0.694399 | 7.072314 |
| CNAG_007 | - | phosphori | 3568.419067 | 3361.53487 | 3524.546304 | 2765.782516 | 2812.669432 | 2672.318368 | 8.56E-08 | -0.35723 | 2.11E-08 | -5.60287 | 0.780663 | 7.067757 |
| CNAG_004 | - | hypothetic | 29.81102844 | 39.01433657 | 26.92195835 | 92.53880448 | 113.8123658 | 92.90678923 | 8.59E-08 | 1.645295 | 2.12E-08 | 5.602157 | 3.128117 | 7.066183 |
| CNAG_074 | - | hypothetic | 31.10079665 | 60.52987016 | 48.21695555 | 136.4442908 | 121.1750419 | 133.8581686 | 8.59E-08 | 1.475191 | 2.12E-08 | 5.602056 | 2.780205 | 7.066145 |
| CNAG_019 | FZC36 | nuclear pr | 1007.527227 | 1036.351674 | 994.9144778 | 731.2334061 | 748.5052131 | 721.7937722 | 8.60E-08 | -0.48028 | 2.12E-08 | -5.6017 | 0.716836 | 7.065475 |
| CNAG_049 | - | hypothetic | 72.89649231 | 67.05721557 | 67.40538215 | 169.0225756 | 176.6123188 | 138.5891674 | 8.66E-08 | 1.213694 | 2.14E-08 | 5.60047 | 2.319307 | 7.062601 |
| CNAG_053 | ITR3C | myo-inosi | 13.43755063 | 17.48352046 | 9.853871266 | 61.64294233 | 69.80383238 | 56.27765494 | 9.01E-08 | 2.239297 | 2.23E-08 | 5.593385 | 4.72167 | 7.045504 |
| CNAG_000 | - | cation trar | 1100.864148 | 1089.143771 | 1018.394756 | 817.7372337 | 742.408739 | 727.3768783 | 9.12E-08 | -0.50359 | 2.26E-08 | -5.59105 | 0.705351 | 7.040088 |
| CNAG_060 | SFB2 | copII vesic | 5084.113546 | 4736.327975 | 4738.196301 | 3649.824012 | 3857.21336 | 3966.70674 | 9.17E-08 | -0.35921 | 2.27E-08 | -5.58987 | 0.779589 | 7.037518 |
| CNAG_013 | GGT1 | Glycosylin | 511.7360128 | 505.443409 | 400.1285735 | 721.8726628 | 783.5957889 | 719.1922894 | 9.31E-08 | 0.635519 | 2.31E-08 | 5.587183 | 1.553497 | 7.031063 |
| CNAG_131 | - | unspecifie | 25.11146342 | 27.19512403 | 10.9585872 | 101.0830556 | 104.4819107 | 60.98884776 | 9.31E-08 | 2.080755 | 2.31E-08 | 5.58705 | 4.230286 | 7.030944 |
| CNAG_018 | - | hypothetic | 99.82733416 | 119.7902033 | 116.4211957 | 228.7570706 | 241.1982169 | 206.2766675 | 9.32E-08 | 1.000142 | 2.31E-08 | 5.58678 | 2.000197 | 7.030483 |
| CNAG_042 | LPI6 | hypothetic | 355.4924221 | 460.1578934 | 394.7424479 | 597.2371106 | 674.9094494 | 656.9450729 | 9.36E-08 | 0.658248 | 2.32E-08 | 5.586047 | 1.578165 | 7.028863 |
| CNAG_048 | - | atypical/A | 1068.823188 | 1133.576962 | 1180.643385 | 1521.849633 | 1511.376997 | 1595.094195 | 9.38E-08 | 0.437605 | 2.33E-08 | 5.585514 | 1.354354 | 7.027746 |
| CNAG_063 | - | hypothetic | 444.6552635 | 463.1390303 | 505.2893886 | 287.4576013 | 267.0164409 | 318.7898467 | 9.48E-08 | -0.71034 | 2.36E-08 | -5.58345 | 0.611174 | 7.023235 |
| CNAG_044 | - | hypothetic | 2.860040545 | 5.630568156 | 4.479499656 | 33.23217013 | 56.41896175 | 33.04986099 | 9.49E-08 | 3.45647 | 2.36E-08 | 5.583215 | 10.97744 | 7.022853 |
| CNAG_080 | - | hypothetic | 1440.785174 | 1339.209342 | 1663.457428 | 1016.797123 | 1057.274599 | 1067.037301 | 9.60E-08 | -0.51622 | 2.39E-08 | -5.58099 | 0.699199 | 7.017518 |
| CNAG_040 | - | Haloacid c | 266.5763533 | 271.6548822 | 191.1448697 | 422.366268 | 440.3104708 | 427.132721 | 9.87E-08 | 0.809369 | 2.46E-08 | 5.57622 | 1.752445 | 7.005815 |
| CNAG_024 | - | rRNA biog | 1017.768897 | 868.327809 | 903.1848749 | 573.8235867 | 693.3455862 | 610.8177212 | 1.02E-07 | -0.58766 | 2.53E-08 | -5.57112 | 0.665419 | 6.993511 |
| CNAG_035 | LYS2 | L-aminoac | 1814.368026 | 1920.857823 | 1591.121798 | 1277.599997 | 1371.333075 | 1127.086866 | 1.02E-07 | -0.51135 | 2.53E-08 | -5.5712 | 0.701563 | 6.993511 |
| CNAG_031 | - | trehalose | 1341.027741 | 1299.605335 | 850.2778794 | 2706.379773 | 2579.601366 | 2367.672758 | 1.03E-07 | 1.11764 | 2.57E-08 | 5.568254 | 2.169917 | 6.986589 |
| CNAG_037 | - | tRNA ade | 813.2709204 | 647.4617467 | 672.8387179 | 475.9115496 | 433.5317542 | 465.3743354 | 1.04E-07 | -0.65089 | 2.60E-08 | -5.56672 | 0.636885 | 6.982975 |
| CNAG_037 | ENV7 | NAK prote | 1617.059823 | 1567.564843 | 1535.623288 | 1167.870394 | 1235.037212 | 1154.747504 | 1.05E-07 | -0.42366 | 2.62E-08 | -5.56483 | 0.745533 | 6.978494 |
| CNAG_048 | - | vacuolar p | 437.1514431 | 517.2646489 | 421.4275227 | 792.2684403 | 723.1217508 | 651.5628047 | 1.06E-07 | 0.641092 | 2.64E-08 | 5.563627 | 1.559509 | 6.975701 |
| CNAG_051 | - | hypothetic | 2784.795084 | 2479.138982 | 2797.187143 | 1929.756753 | 2102.768091 | 2057.27323 | 1.06E-07 | -0.42071 | 2.65E-08 | -5.5632 | 0.747057 | 6.974843 |
| CNAG_073 | - | hypothetic | 3347.746946 | 3339.950953 | 3312.333645 | 2455.116737 | 2749.731867 | 2641.201841 | 1.08E-07 | -0.36521 | 2.69E-08 | -5.56032 | 0.776357 | 6.968113 |
| CNAG_024 | - | peptidyl-p | 1220.11168 | 1215.259842 | 1386.209395 | 963.2616105 | 832.9216617 | 909.4022502 | 1.16E-07 | -0.51413 | 2.91E-08 | -5.54683 | 0.700217 | 6.934998 |
| CNAG_006 | URO3 | hypothetic | 833.152266 | 766.9328128 | 651.5601384 | 493.812464 | 381.649826 | 511.9537408 | 1.18E-07 | -0.71502 | 2.97E-08 | -5.54334 | 0.609199 | 6.926974 |
| CNAG_017 | - | ATP-depe | 2832.760122 | 2715.071435 | 2871.866387 | 2160.999523 | 2251.137312 | 2203.807152 | 1.19E-07 | -0.36347 | 2.98E-08 | -5.5425 | 0.777294 | 6.925101 |
| CNAG_028 | - | potassium | 1296.494663 | 1455.671231 | 1205.242094 | 1803.616612 | 1830.576043 | 1811.619154 | 1.19E-07 | 0.446018 | 2.99E-08 | 5.541864 | 1.362275 | 6.923732 |
| CNAG_033 | - | mitochonc | 1388.076665 | 1269.123878 | 1273.2643 | 953.8422229 | 1009.047493 | 865.0633717 | 1.21E-07 | -0.49103 | 3.04E-08 | -5.53937 | 0.711515 | 6.917975 |
| CNAG_025 | - | transketol | 927.9789916 | 877.9830638 | 860.5368288 | 555.1485437 | 667.4660212 | 615.2244402 | 1.22E-07 | -0.55182 | 3.07E-08 | -5.53738 | 0.68216 | 6.913446 |
| CNAG_024 | - | hypothetic | 142.9580044 | 155.343113 | 159.0579001 | 278.4451718 | 282.4512165 | 275.0431339 | 1.24E-07 | 0.857699 | 3.13E-08 | 5.534259 | 1.812146 | 6.905933 |
| CNAG_068 | - | hypothetic | 1143.596652 | 961.9694387 | 807.2862263 | 703.3529068 | 537.8401036 | 514.3505805 | 1.24E-07 | -0.74658 | 3.13E-08 | -5.53403 | 0.596016 | 6.905569 |
| CNAG_057 | - | transcripti | 4922.246438 | 4723.569555 | 4812.984563 | 6168.230044 | 5920.728323 | 6123.51864 | 1.24E-07 | 0.31767 | 3.13E-08 | 5.53392 | 1.246316 | 6.905513 |

| | | | | | | | | | | | | | | |
|---|---|---|---|---|---|---|---|---|---|---|---|---|---|---|
| CNAG_005 | - | nitrilase | 12.27119124 | 9.987913127 | 16.20880633 | 65.81760407 | 53.90027254 | 61.67041735 | 1.25E-07 | 2.258527 | 3.16E-08 | 5.532463 | 4.785028 | 6.902114 |
| CNAG_079 | - | cell divisic | 1197.966796 | 1374.607077 | 1227.411487 | 938.3013269 | 871.776516 | 944.8980258 | 1.27E-07 | -0.47872 | 3.20E-08 | -5.53001 | 0.717613 | 6.89625 |
| CNAG_039 | - | F-actin ca | 2880.627593 | 2790.478002 | 2840.956492 | 2312.092346 | 2174.316795 | 2210.475803 | 1.30E-07 | -0.3612 | 3.27E-08 | -5.52639 | 0.778515 | 6.887497 |
| CNAG_067 | - | efflux prot | 231.5452382 | 171.5907227 | 216.6494109 | 424.3520101 | 333.1135899 | 382.6897906 | 1.34E-07 | 0.865956 | 3.39E-08 | 5.520226 | 1.822547 | 6.872469 |
| CNAG_028 | CAP64 | capsular a | 2006.087182 | 2180.597968 | 2142.411361 | 1687.108271 | 1622.292812 | 1561.067814 | 1.41E-07 | -0.39278 | 3.58E-08 | -5.51061 | 0.761662 | 6.849557 |
| CNAG_079 | - | hypothetic | 771.5568416 | 721.7894724 | 786.8964867 | 510.1200797 | 500.8588414 | 553.0109901 | 1.44E-07 | -0.5589 | 3.66E-08 | -5.50664 | 0.678818 | 6.84018 |
| CNAG_039 | IDP1 | isocitrate | 7730.454197 | 8040.815077 | 8995.390769 | 6461.078472 | 6588.0857 | 6482.369963 | 1.50E-07 | -0.35782 | 3.81E-08 | -5.49928 | 0.780341 | 6.822878 |
| CNAG_010 | - | hypothetic | 41.56400722 | 57.35128855 | 65.23001879 | 160.0150198 | 137.6221813 | 124.0974993 | 1.56E-07 | 1.354336 | 3.95E-08 | 5.493047 | 2.556794 | 6.808172 |
| CNAG_055 | - | hypothetic | 338.3340327 | 403.7100456 | 329.4301039 | 199.4368463 | 185.8987504 | 229.9913795 | 1.62E-07 | -0.81779 | 4.13E-08 | -5.48517 | 0.56731 | 6.789222 |
| CNAG_042 | - | hypothetic | 438.3193741 | 503.2764507 | 450.1953176 | 802.417579 | 713.6061711 | 644.9174484 | 1.63E-07 | 0.619525 | 4.16E-08 | 5.48413 | 1.536369 | 6.786877 |
| CNAG_014 | APG9 | autophagy | 446.45234 | 489.2750836 | 454.4482502 | 643.0811269 | 734.3910852 | 718.0248925 | 1.64E-07 | 0.577969 | 4.18E-08 | 5.482904 | 1.492746 | 6.784074 |
| CNAG_026 | - | hypothetic | 627.4728668 | 681.0727932 | 728.4690782 | 995.8663294 | 949.8698648 | 983.4049651 | 1.75E-07 | 0.508693 | 4.46E-08 | 5.471789 | 1.422761 | 6.757429 |
| CNAG_038 | - | galactose- | 2120.270085 | 1942.523031 | 1861.983724 | 1525.953731 | 1490.309514 | 1455.598567 | 1.75E-07 | -0.4216 | 4.47E-08 | -5.47122 | 0.746595 | 6.756245 |
| CNAG_027 | LIV13 | endoplasn | 1361.812689 | 1375.973231 | 1157.266036 | 1767.005156 | 1923.832483 | 1739.50018 | 1.75E-07 | 0.464641 | 4.47E-08 | 5.471077 | 1.379974 | 6.756099 |
| CNAG_064 | - | hypothetic | 1081.560629 | 1096.919829 | 947.1639129 | 1421.45923 | 1477.817903 | 1430.839609 | 1.77E-07 | 0.455226 | 4.52E-08 | 5.46921 | 1.370998 | 6.751942 |
| CNAG_078 | - | solute car | 11.25866555 | 31.49431853 | 33.26271915 | 89.59734239 | 100.8155742 | 89.49962268 | 1.77E-07 | 1.876678 | 4.52E-08 | 5.46923 | 3.672285 | 6.751942 |
| CNAG_063 | - | cytoplasm | 671.808372 | 760.7560763 | 683.7274488 | 1024.649095 | 1039.670636 | 955.7311125 | 1.84E-07 | 0.498812 | 4.70E-08 | 5.462308 | 1.413049 | 6.735455 |
| CNAG_065 | SKS1 | RAN prote | 113.8351888 | 157.4259513 | 98.37509155 | 272.5361371 | 267.9252204 | 225.1807892 | 1.87E-07 | 1.040017 | 4.80E-08 | 5.45871 | 2.056252 | 6.727275 |
| CNAG_076 | - | hypothetic | 548.3715838 | 486.8947908 | 519.2221923 | 322.4883764 | 362.9425617 | 306.6876488 | 1.87E-07 | -0.66549 | 4.80E-08 | -5.45874 | 0.630473 | 6.727275 |
| CNAG_030 | PTC2 | PP2Cc prc | 2797.5701 | 2497.425071 | 2554.084776 | 2126.114583 | 1871.505828 | 1901.894021 | 1.88E-07 | -0.42775 | 4.80E-08 | -5.4584 | 0.743421 | 6.726732 |
| CNAG_036 | - | lipoate-pr | 1316.799641 | 1121.558976 | 1226.312639 | 833.1046999 | 916.7276889 | 867.2236513 | 1.88E-07 | -0.50169 | 4.83E-08 | -5.45756 | 0.706277 | 6.725286 |
| CNAG_047 | - | Dihydrofo | 77.73415547 | 107.9439563 | 113.1990313 | 209.2859128 | 224.835982 | 195.1375138 | 1.95E-07 | 1.065693 | 5.00E-08 | 5.451148 | 2.093175 | 6.710036 |
| CNAG_010 | - | BEM46 far | 711.4883254 | 630.5107535 | 799.90908 | 1125.208976 | 1020.601633 | 1043.368973 | 1.96E-07 | 0.559199 | 5.02E-08 | 5.450523 | 1.473451 | 6.708715 |
| CNAG_130 | - | unspecifie | 57.89905965 | 86.39033048 | 80.17620322 | 152.1935958 | 171.2474977 | 195.9232549 | 1.97E-07 | 1.201959 | 5.07E-08 | 5.448943 | 2.300518 | 6.705064 |
| CNAG_126 | - | unspecifie | 29.81185706 | 36.87481678 | 33.2883727 | 83.72567038 | 108.5370048 | 103.8554578 | 1.98E-07 | 1.570186 | 5.09E-08 | 5.448215 | 2.96943 | 6.703492 |
| CNAG_022 | - | hypothetic | 306.2293043 | 292.1384616 | 206.0817425 | 498.854252 | 456.5978685 | 459.3562593 | 2.03E-07 | 0.798479 | 5.22E-08 | 5.443713 | 1.739267 | 6.692713 |
| CNAG_128 | - | unspecifie | 34.4382797 | 31.52969913 | 39.65812174 | 98.06709114 | 108.484752 | 95.13299366 | 2.07E-07 | 1.517402 | 5.33E-08 | 5.439971 | 2.862751 | 6.683795 |
| CNAG_050 | HRB1 | RNP doma | 10188.52716 | 10420.85605 | 10642.06192 | 8227.00184 | 8894.633447 | 8562.786236 | 2.14E-07 | -0.29832 | 5.51E-08 | -5.4339 | 0.8132 | 6.669405 |
| CNAG_075 | - | hypothetic | 763.5348529 | 810.0951306 | 839.1548608 | 580.266583 | 551.7780877 | 566.3699048 | 2.14E-07 | -0.52228 | 5.52E-08 | -5.43354 | 0.696271 | 6.668741 |
| CNAG_046 | - | histone de | 2304.197033 | 2274.570482 | 2372.930381 | 3003.708296 | 2965.641925 | 2941.811101 | 2.20E-07 | 0.342905 | 5.69E-08 | 5.428335 | 1.268308 | 6.656898 |
| CNAG_011 | - | endoplasn | 1600.098677 | 1750.89892 | 1621.112628 | 2191.968769 | 2135.847925 | 2195.74568 | 2.26E-07 | 0.376769 | 5.85E-08 | 5.423264 | 1.298431 | 6.644977 |
| CNAG_015 | - | hypothetic | 37.93250197 | 41.18410156 | 35.43096077 | 111.5034364 | 92.59217174 | 114.9291075 | 2.37E-07 | 1.473625 | 6.14E-08 | 5.414635 | 2.777189 | 6.625043 |
| CNAG_056 | - | neutral an | 6918.822276 | 7491.266452 | 6911.64177 | 5615.068179 | 5826.459256 | 5921.696018 | 2.38E-07 | -0.31109 | 6.17E-08 | -5.41369 | 0.806032 | 6.623163 |
| CNAG_033 | BCH1 | ChAPs fan | 1527.196192 | 1513.697271 | 1521.743107 | 1196.766315 | 1115.062014 | 1154.72104 | 2.38E-07 | -0.41171 | 6.18E-08 | -5.4135 | 0.75173 | 6.622903 |
| CNAG_007 | - | dihydroor | 593.9471165 | 496.6552567 | 619.3980016 | 373.1508679 | 355.0634181 | 364.3630714 | 2.42E-07 | -0.66138 | 6.27E-08 | -5.41095 | 0.632272 | 6.616915 |
| CNAG_011 | - | hypothetic | 558.0762298 | 663.4892031 | 670.6102035 | 423.7730887 | 361.0249353 | 436.464906 | 2.43E-07 | -0.64668 | 6.32E-08 | -5.4095 | 0.638748 | 6.613618 |
| CNAG_045 | - | mitochonc | 992.3270633 | 937.2957219 | 1049.240443 | 728.9364438 | 681.1539491 | 721.7726405 | 2.44E-07 | -0.49897 | 6.34E-08 | -5.40902 | 0.70761 | 6.612659 |
| CNAG_047 | - | hypothetic | 71.82423464 | 99.2792386 | 67.43636607 | 177.3995178 | 199.0603313 | 160.7474818 | 2.46E-07 | 1.163192 | 6.40E-08 | 5.407371 | 2.239523 | 6.608855 |
| CNAG_059 | - | hypothetic | 34.37769739 | 34.71372091 | 15.22614183 | 86.29637223 | 92.83865088 | 105.945901 | 2.47E-07 | 1.755115 | 6.42E-08 | 5.406606 | 3.375532 | 6.607205 |
| CNAG_040 | - | hypothetic | 833.5531107 | 837.0587215 | 885.0205181 | 596.5504718 | 626.8871647 | 600.7887591 | 2.50E-07 | -0.50315 | 6.52E-08 | -5.40395 | 0.705565 | 6.601586 |
| CNAG_078 | - | hypothetic | 0.308519602 | 0.147976978 | 0.127604829 | 19.09914892 | 29.21939365 | 27.24629781 | 2.65E-07 | 7.071083 | 6.90E-08 | 5.393819 | 134.4647 | 6.577266 |
| CNAG_038 | - | acyltransfe | 667.2498657 | 804.9556374 | 834.0450382 | 1072.771351 | 1154.407173 | 1119.953573 | 2.68E-07 | 0.522504 | 6.98E-08 | 5.391612 | 1.436446 | 6.572134 |
| CNAG_078 | - | hypothetic | 302.9319368 | 378.2888728 | 341.4084475 | 576.5887647 | 507.3723496 | 544.8462245 | 2.71E-07 | 0.657938 | 7.06E-08 | 5.389608 | 1.577825 | 6.567697 |
| CNAG_055 | - | hypothetic | 2421.652399 | 2273.316439 | 2302.406754 | 1803.775585 | 1836.225952 | 1837.449297 | 2.76E-07 | -0.36891 | 7.20E-08 | -5.38617 | 0.774368 | 6.559791 |
| CNAG_064 | - | acetolacta | 581.5078976 | 730.2789006 | 733.526884 | 461.9087999 | 448.2571825 | 436.5329018 | 2.79E-07 | -0.61828 | 7.29E-08 | -5.384 | 0.651448 | 6.554753 |
| CNAG_033 | - | hypothetic | 961.9656631 | 990.0123867 | 933.0664234 | 666.5755768 | 745.1132098 | 657.4341133 | 2.81E-07 | -0.49508 | 7.35E-08 | -5.38247 | 0.709523 | 6.551275 |
| CNAG_001 | - | hypothetic | 271.2114186 | 254.4429901 | 225.2149204 | 408.2999748 | 428.2080607 | 404.9844451 | 2.82E-07 | 0.709981 | 7.37E-08 | 5.381968 | 1.635783 | 6.550263 |
| CNAG_056 | - | hypothetic | 2.81572764 | 4.532816386 | 0.158696761 | 27.74569835 | 42.31567951 | 35.05332237 | 2.85E-07 | 4.101566 | 7.46E-08 | 5.379754 | 17.16699 | 6.545125 |
| CNAG_019 | TAL1 | transaldol | 23401.57913 | 23411.44833 | 22074.49206 | 19392.46674 | 19282.42652 | 18867.30838 | 3.00E-07 | -0.27482 | 7.85E-08 | -5.37058 | 0.826553 | 6.523412 |
| CNAG_029 | - | calcium/pr | 1302.330309 | 1380.287603 | 1371.545472 | 1849.597387 | 1773.592787 | 1745.059913 | 3.05E-07 | 0.39009 | 7.99E-08 | 5.367443 | 1.310475 | 6.516267 |
| CNAG_055 | - | DNA repa | 1037.928518 | 1095.605265 | 1036.513191 | 784.1905678 | 773.5005515 | 782.8267462 | 3.17E-07 | -0.45265 | 8.32E-08 | -5.36005 | 0.730699 | 6.498881 |
| CNAG_001 | - | alternative | 1011.491677 | 541.8276771 | 567.2127962 | 225.5039349 | 223.6562551 | 216.9789525 | 3.23E-07 | -1.69223 | 8.48E-08 | -5.35671 | 0.309448 | 6.491063 |
| CNAG_037 | ISP1 | short-chai | 5.161746024 | 3.469637815 | 3.403711874 | 37.57881071 | 38.60690765 | 34.04540619 | 3.27E-07 | 3.292715 | 8.59E-08 | 5.354212 | 9.799549 | 6.485265 |
| CNAG_002 | - | oligoribon | 1162.771492 | 1112.885255 | 1185.757225 | 878.4599252 | 841.6687665 | 670.8606676 | 3.33E-07 | -0.54967 | 8.75E-08 | -5.35088 | 0.683177 | 6.477473 |
| CNAG_079 | - | hypothetic | 36.88018329 | 59.45212846 | 49.27387262 | 116.5801871 | 125.5778614 | 134.9315045 | 3.35E-07 | 1.367077 | 8.80E-08 | 5.349857 | 2.579474 | 6.475214 |
| CNAG_044 | - | hypothetic | 67.13985998 | 78.86829411 | 78.0395995 | 168.9169387 | 176.5441193 | 147.4377902 | 3.38E-07 | 1.123638 | 8.88E-08 | 5.348307 | 2.178958 | 6.471697 |
| CNAG_058 | - | mitochonc | 818.3303431 | 804.7353323 | 876.4634845 | 610.6681872 | 500.7773311 | 597.4293173 | 3.40E-07 | -0.56446 | 8.96E-08 | -5.34656 | 0.676208 | 6.46791 |
| CNAG_065 | - | monocarb | 567.2575354 | 617.1627386 | 551.271295 | 377.7846708 | 404.3202231 | 385.4406126 | 3.56E-07 | -0.58826 | 9.39E-08 | -5.33805 | 0.665144 | 6.44873 |
| CNAG_063 | - | hypothetic | 2307.64392 | 2277.777269 | 1935.768218 | 2898.646185 | 2869.906399 | 2976.161418 | 3.63E-07 | 0.408427 | 9.59E-08 | 5.334238 | 1.327238 | 6.439796 |
| CNAG_068 | - | translation | 1060.068593 | 1035.318221 | 1104.713173 | 692.9369853 | 824.4660281 | 776.1704426 | 3.76E-07 | -0.49655 | 9.94E-08 | -5.32777 | 0.7088 | 6.424736 |
| CNAG_069 | - | hypothetic | 490.0777564 | 564.2500465 | 440.3784396 | 357.7967024 | 259.2012672 | 263.4419194 | 3.79E-07 | -0.77951 | 1.00E-07 | -5.3262 | 0.582564 | 6.42118 |
| CNAG_006 | CEL1 | glycosyl h | 48.44419028 | 57.31683876 | 46.09375182 | 143.6139744 | 117.6758998 | 118.4870495 | 3.99E-07 | 1.309343 | 1.06E-07 | 5.316824 | 2.478286 | 6.39939 |
| CNAG_055 | - | hypothetic | 8.824906427 | 17.48831146 | 18.33768764 | 67.28655671 | 61.79225576 | 59.56223432 | 4.17E-07 | 2.106709 | 1.10E-07 | 5.308646 | 4.307078 | 6.380093 |
| CNAG_013 | STE24 | STE24 enc | 1478.016282 | 1326.253429 | 1368.179845 | 1074.558353 | 1034.869875 | 992.6819843 | 4.21E-07 | -0.44418 | 1.12E-07 | -5.30679 | 0.735003 | 6.375862 |
| CNAG_018 | - | hypothetic | 1332.735751 | 1497.712523 | 1412.086217 | 1759.042519 | 1985.872972 | 1999.184667 | 4.21E-07 | 0.422611 | 1.12E-07 | 5.306636 | 1.340351 | 6.375705 |
| CNAG_023 | - | hypothetic | 1467.663531 | 1497.520115 | 1453.503676 | 1119.645552 | 1116.819112 | 1151.367812 | 4.27E-07 | -0.39858 | 1.13E-07 | -5.30419 | 0.758602 | 6.37007 |
| CNAG_030 | ERG28 | putative E | 527.4781676 | 483.6856114 | 566.0890313 | 366.239731 | 324.8629336 | 329.9645571 | 4.30E-07 | -0.64477 | 1.14E-07 | -5.30264 | 0.639597 | 6.366596 |
| CNAG_060 | GAL7 | galactose- | 152.3119191 | 173.6433088 | 162.2787825 | 335.6092737 | 320.503835 | 247.5086973 | 4.32E-07 | 0.874174 | 1.15E-07 | 5.301684 | 1.832958 | 6.364516 |

| | | | | | | | | | | | | | | |
|---|---|---|---|---|---|---|---|---|---|---|---|---|---|---|
| CNAG_063 | - | glycine cle | 668.341443 | 706.9443337 | 780.7204068 | 1129.142645 | 980.8781606 | 1007.874938 | 4.35E-07 | 0.517589 | 1.16E-07 | 5.300478 | 1.431561 | 6.361846 |
| CNAG_017 | - | hypothetic | 73.15568846 | 125.1731919 | 134.5088935 | 252.3087176 | 233.3136196 | 227.2835956 | 4.58E-07 | 1.086824 | 1.22E-07 | 5.290887 | 2.124059 | 6.339449 |
| CNAG_044 | - | hypothetic | 7.539158302 | 5.647536987 | 8.738508111 | 36.31137812 | 43.69125659 | 52.6004628 | 4.72E-07 | 2.695707 | 1.26E-07 | 5.284856 | 6.478711 | 6.325728 |
| CNAG_075 | - | DNA repa | 3287.388007 | 3236.693812 | 2883.824666 | 3896.269616 | 4351.028037 | 4087.479752 | 4.73E-07 | 0.37569 | 1.26E-07 | 5.28459 | 1.29746 | 6.3253 |
| CNAG_038 | - | hypothetic | 666.0815395 | 837.2525124 | 691.2299753 | 1350.538577 | 1139.725373 | 954.7311368 | 4.73E-07 | 0.635836 | 1.26E-07 | 5.284503 | 1.553838 | 6.325292 |
| CNAG_073 | - | hypothetic | 1019.192522 | 1033.127305 | 1001.31237 | 745.2659881 | 790.8264794 | 678.5504248 | 4.79E-07 | -0.47957 | 1.28E-07 | -5.28219 | 0.717193 | 6.320006 |
| CNAG_045 | - | cytochrom | 483.3566994 | 581.6025401 | 608.7226647 | 388.050998 | 304.069302 | 355.4779804 | 4.84E-07 | -0.68974 | 1.29E-07 | -5.28013 | 0.619965 | 6.315311 |
| CNAG_120 | - | unspecifie | 540.3821752 | 548.2669934 | 560.8105788 | 325.5108409 | 385.3541373 | 380.9631199 | 5.05E-07 | -0.61167 | 1.35E-07 | -5.27204 | 0.654438 | 6.296516 |
| CNAG_019 | NDX1 | hypothetic | 1370.642486 | 1332.672181 | 1269.020797 | 944.4411754 | 980.5213244 | 1042.554736 | 5.18E-07 | -0.43622 | 1.39E-07 | -5.26704 | 0.739071 | 6.285327 |
| CNAG_052 | TLK1 | phosphati | 122.1026599 | 177.9366478 | 167.595223 | 340.3554518 | 246.7727428 | 324.8485911 | 5.23E-07 | 0.951912 | 1.40E-07 | 5.265478 | 1.934434 | 6.281826 |
| CNAG_044 | - | hypothetic | 2883.436097 | 2987.815901 | 3115.168726 | 3765.595968 | 3859.278291 | 3738.918149 | 5.38E-07 | 0.323407 | 1.44E-07 | 5.260064 | 1.251282 | 6.269429 |
| CNAG_073 | - | hypothetic | 526.9577876 | 520.546017 | 541.8640538 | 748.2671401 | 754.9599202 | 781.3323289 | 5.40E-07 | 0.509358 | 1.45E-07 | 5.259193 | 1.423417 | 6.267571 |
| CNAG_000 | - | pre-mRNA | 4961.557114 | 4860.213809 | 4682.752436 | 3644.366093 | 3990.082916 | 4003.337226 | 5.46E-07 | -0.33295 | 1.47E-07 | -5.25689 | 0.793912 | 6.262717 |
| CNAG_066 | - | ubiquitin | 2049.076934 | 1814.353778 | 1936.594601 | 1493.254288 | 1471.33616 | 1472.227575 | 5.47E-07 | -0.40231 | 1.47E-07 | -5.2566 | 0.756645 | 6.262234 |
| CNAG_004 | - | mitochonc | 4793.891471 | 2456.486389 | 2963.467503 | 1139.213273 | 1102.591216 | 1129.524592 | 5.64E-07 | -1.61775 | 1.52E-07 | -5.25072 | 0.325843 | 6.248958 |
| CNAG_039 | - | nit proteir | 288.0125873 | 300.3665393 | 286.7121241 | 194.2749953 | 156.6112179 | 131.3906902 | 5.88E-07 | -0.87672 | 1.58E-07 | -5.24292 | 0.544605 | 6.230986 |
| CNAG_077 | - | hypothetic | 95.11224983 | 112.2306867 | 99.37508475 | 206.1695437 | 186.5786987 | 216.0674768 | 5.88E-07 | 0.976083 | 1.58E-07 | 5.242691 | 1.967118 | 6.23083 |
| CNAG_039 | - | hypothetic | 987.7061703 | 1039.57713 | 983.1894089 | 756.2043731 | 699.272869 | 752.8388268 | 5.94E-07 | -0.46209 | 1.60E-07 | -5.24054 | 0.725933 | 6.226163 |
| CNAG_075 | - | hypothetic | 181.3102208 | 165.0356757 | 139.9244356 | 286.1693293 | 297.9867277 | 288.3664735 | 6.01E-07 | 0.829284 | 1.62E-07 | 5.23827 | 1.776803 | 6.221014 |
| CNAG_040 | - | hypothetic | 10.01182669 | 17.50931524 | 24.70819253 | 71.90771742 | 64.30848384 | 68.37415395 | 6.15E-07 | 1.978422 | 1.66E-07 | 5.234029 | 3.940617 | 6.211239 |
| CNAG_047 | - | hypothetic | 34.51858016 | 43.35229065 | 50.30548975 | 94.51276758 | 119.6244527 | 130.3957875 | 6.17E-07 | 1.418598 | 1.67E-07 | 5.233251 | 2.673255 | 6.209998 |
| CNAG_050 | - | zinc finger | 988.9203243 | 1050.365889 | 1084.444746 | 686.7356648 | 827.9517262 | 716.2667586 | 6.17E-07 | -0.50138 | 1.67E-07 | -5.23328 | 0.706432 | 6.209998 |
| CNAG_021 | - | nuclear pr | 953.5736689 | 775.719568 | 889.291929 | 592.6294859 | 598.3619493 | 621.8629085 | 6.17E-07 | -0.54722 | 1.66E-07 | -5.23338 | 0.684337 | 6.209998 |
| CNAG_076 | - | voltage-ga | 1524.188057 | 1667.93311 | 1360.956037 | 1908.461444 | 2175.643893 | 2257.803123 | 6.35E-07 | 0.463656 | 1.72E-07 | 5.227545 | 1.379032 | 6.196984 |
| CNAG_025 | - | spermine | 648.2189292 | 419.0923633 | 474.4566897 | 256.8973208 | 263.5180026 | 339.8709364 | 6.35E-07 | -0.86159 | 1.72E-07 | -5.22746 | 0.550345 | 6.196978 |
| CNAG_010 | - | short-chai | 1536.598024 | 1677.405955 | 1535.628405 | 1282.438955 | 1155.600537 | 1146.987812 | 6.47E-07 | -0.42093 | 1.75E-07 | -5.22391 | 0.746942 | 6.188835 |
| CNAG_031 | - | hypothetic | 439.4945902 | 495.7489429 | 508.7967639 | 727.3823059 | 666.0389155 | 750.1981315 | 6.48E-07 | 0.556838 | 1.75E-07 | 5.223621 | 1.471042 | 6.188361 |
| CNAG_031 | - | Cullin 3 | 207.0379311 | 180.1547314 | 200.6437636 | 318.7159388 | 324.597399 | 368.1743616 | 6.62E-07 | 0.76733 | 1.79E-07 | 5.219547 | 1.702117 | 6.179197 |
| CNAG_050 | CLR5 | hypothetic | 12.28852652 | 21.75500159 | 6.66381673 | 65.69399944 | 48.52845986 | 75.85006629 | 6.62E-07 | 2.257029 | 1.79E-07 | 5.219577 | 4.780061 | 6.179197 |
| CNAG_075 | - | hypothetic | 126.5408384 | 136.9939213 | 106.8685288 | 230.9679363 | 235.0143897 | 232.8300083 | 6.63E-07 | 0.906661 | 1.80E-07 | 5.21904 | 1.874702 | 6.178204 |
| CNAG_018 | - | electron tı | 2537.324521 | 2565.263957 | 2424.002979 | 1965.671635 | 1959.555725 | 2053.900759 | 6.67E-07 | -0.34754 | 1.81E-07 | -5.21786 | 0.785923 | 6.175629 |
| CNAG_046 | - | hypothetic | 79.77628827 | 67.03640444 | 42.94913948 | 151.7496588 | 140.069447 | 170.4141408 | 6.75E-07 | 1.278899 | 1.83E-07 | 5.215545 | 2.426536 | 6.170599 |
| CNAG_051 | - | small subu | 942.021346 | 864.0166888 | 911.70384 | 703.3658875 | 596.5898743 | 618.5738874 | 6.93E-07 | -0.51868 | 1.88E-07 | -5.21041 | 0.698012 | 6.158974 |
| CNAG_034 | LAC2 | Laccase 2 | 100.8902962 | 104.7107721 | 95.11736007 | 214.1137247 | 210.9700482 | 174.1479718 | 7.22E-07 | 0.984779 | 1.96E-07 | 5.202727 | 1.979011 | 6.141576 |
| CNAG_041 | - | vacuolar p | 1704.55226 | 1541.730034 | 1523.905453 | 1142.131874 | 1220.348449 | 1229.082296 | 7.27E-07 | -0.42462 | 1.98E-07 | -5.20137 | 0.745034 | 6.138604 |
| CNAG_009 | - | hypothetic | 999.2362328 | 935.1074512 | 924.5346058 | 701.7009038 | 627.6359553 | 719.5193527 | 7.28E-07 | -0.49677 | 1.98E-07 | -5.20074 | 0.708689 | 6.137708 |
| CNAG_051 | - | hypothetic | 278.6939982 | 275.6487099 | 295.2003248 | 177.0413638 | 168.7660423 | 132.4858419 | 7.28E-07 | -0.84563 | 1.98E-07 | -5.20077 | 0.556467 | 6.137708 |
| CNAG_063 | - | hypothetic | 1849.919743 | 276.1098427 | 577.8797455 | 37.62700995 | 66.46927765 | 42.82347801 | 7.33E-07 | -4.2439 | 2.00E-07 | -5.19937 | 0.052779 | 6.134892 |
| CNAG_018 | CID14 | DNA polyı | 1730.301253 | 1612.838568 | 1606.0152 | 1284.689617 | 1204.761786 | 1289.026698 | 7.36E-07 | -0.405 | 2.01E-07 | -5.19842 | 0.755235 | 6.132884 |
| CNAG_054 | - | pi-transpc | 858.0067617 | 828.443148 | 891.4082066 | 552.0769949 | 680.4674277 | 543.1329185 | 7.39E-07 | -0.55367 | 2.02E-07 | -5.1977 | 0.681286 | 6.131388 |
| CNAG_079 | - | hypothetic | 44.86313352 | 43.33391029 | 25.88035812 | 137.0245151 | 89.92517945 | 104.0064116 | 7.43E-07 | 1.544428 | 2.03E-07 | 5.19665 | 2.916885 | 6.129129 |
| CNAG_066 | - | hypothetic | 189.8583022 | 305.0302603 | 277.422362 | 401.8186771 | 472.2261224 | 520.1903534 | 7.66E-07 | 0.839879 | 2.10E-07 | 5.190636 | 1.789901 | 6.115673 |
| CNAG_040 | - | cytochrom | 532.140498 | 504.1247086 | 536.2790473 | 324.0157796 | 334.3727445 | 379.8139069 | 7.68E-07 | -0.61555 | 2.10E-07 | -5.19017 | 0.65268 | 6.114771 |
| CNAG_076 | - | nicotinate | 610.6252114 | 723.8280461 | 696.2500283 | 474.3979297 | 485.4235828 | 416.5866761 | 7.84E-07 | -0.57612 | 2.15E-07 | -5.18601 | 0.670767 | 6.105457 |
| CNAG_043 | - | peptidyl-tl | 789.2340128 | 812.2819384 | 924.4195761 | 570.8372189 | 640.7412698 | 532.0320211 | 7.98E-07 | -0.55109 | 2.19E-07 | -5.18268 | 0.682503 | 6.098102 |
| CNAG_021 | - | vacuolar p | 110.3258493 | 127.3408286 | 134.5275523 | 273.4428628 | 202.8288812 | 246.105874 | 8.32E-07 | 0.944015 | 2.29E-07 | 5.174504 | 1.923875 | 6.079647 |
| CNAG_120 | - | unspecifie | 7.663536046 | 15.35059596 | 20.45383749 | 77.87580589 | 52.05263243 | 59.56163841 | 8.37E-07 | 2.148357 | 2.30E-07 | 5.173308 | 4.433227 | 6.077059 |
| CNAG_126 | - | unspecifie | 352.0239856 | 305.8382446 | 312.3230534 | 222.3392153 | 152.2021188 | 165.744599 | 8.43E-07 | -0.86179 | 2.32E-07 | -5.17203 | 0.55027 | 6.074291 |
| CNAG_064 | - | AP-2 com | 285.7185989 | 298.2310364 | 279.2779281 | 166.0058486 | 173.9370994 | 171.2149407 | 8.58E-07 | -0.77223 | 2.36E-07 | -5.16845 | 0.585511 | 6.066341 |
| CNAG_029 | - | CMGC/DY | 2622.64596 | 2775.358088 | 2663.939326 | 2094.850539 | 2180.410038 | 2192.678328 | 8.72E-07 | -0.3329 | 2.40E-07 | -5.16537 | 0.79394 | 6.059375 |
| CNAG_131 | - | unspecifie | 2930.247119 | 2393.904117 | 3085.937292 | 1447.811979 | 853.2568552 | 1262.630331 | 8.85E-07 | -1.25479 | 2.44E-07 | -5.16262 | 0.419056 | 6.053182 |
| CNAG_061 | - | carbamoyl | 7719.6794 | 7299.505016 | 6564.012307 | 5827.61596 | 5816.109495 | 5357.823146 | 8.94E-07 | -0.35975 | 2.46E-07 | -5.16053 | 0.779297 | 6.048739 |
| CNAG_063 | - | hypothetic | 106.592919 | 72.47813144 | 79.14533142 | 207.2752042 | 155.406732 | 203.7908165 | 9.05E-07 | 1.120423 | 2.50E-07 | 5.157997 | 2.174107 | 6.043431 |
| CNAG_051 | UGT1 | UDP-galac | 598.2117863 | 660.5560864 | 589.8832405 | 905.6115846 | 883.5056191 | 832.5026632 | 9.05E-07 | 0.489539 | 2.50E-07 | 5.158 | 1.403996 | 6.043431 |
| CNAG_056 | - | hypothetic | 1035.601926 | 1022.404872 | 1130.293482 | 816.9477574 | 777.816693 | 740.6885297 | 9.05E-07 | -0.46545 | 2.50E-07 | -5.15801 | 0.724244 | 6.043431 |
| CNAG_048 | - | tyrosine-tl | 1950.400931 | 1989.045707 | 2068.988163 | 2457.981062 | 2644.813631 | 2648.680928 | 9.09E-07 | 0.352119 | 2.51E-07 | 5.157092 | 1.276434 | 6.041526 |
| CNAG_052 | TPS1 | alpha,alph | 6220.293112 | 5805.121178 | 5356.780763 | 4513.00326 | 4754.310079 | 4547.362706 | 9.10E-07 | -0.3469 | 2.51E-07 | -5.15677 | 0.786273 | 6.040963 |
| CNAG_052 | - | cytoplasm | 352.2910567 | 311.3156429 | 411.3641367 | 228.4057357 | 205.8054412 | 199.0168019 | 9.18E-07 | -0.78095 | 2.53E-07 | -5.15514 | 0.581983 | 6.037389 |
| CNAG_051 | - | DNA polyı | 442.9901195 | 536.6434388 | 473.6390638 | 698.4746955 | 734.3392751 | 703.6598543 | 9.26E-07 | 0.542644 | 2.56E-07 | 5.153307 | 1.45664 | 6.033329 |
| CNAG_120 | - | unspecifie | 30.9202907 | 32.57123605 | 32.20902435 | 90.33478635 | 85.82133197 | 93.90476316 | 9.61E-07 | 1.496555 | 2.66E-07 | 5.146257 | 2.821681 | 6.017387 |
| CNAG_078 | - | hypothetic | 57.5536694 | 30.48671437 | 33.32170294 | 123.3064159 | 100.3850753 | 117.2092038 | 9.72E-07 | 1.485653 | 2.69E-07 | 5.143952 | 2.800439 | 6.012247 |
| CNAG_042 | - | hypothetic | 3671.328916 | 3963.768495 | 4035.354912 | 2834.126529 | 3023.09417 | 3286.14277 | 9.74E-07 | -0.36684 | 2.70E-07 | -5.14358 | 0.775481 | 6.011583 |
| CNAG_065 | - | MFS trans | 20.44024555 | 28.22486993 | 13.07012268 | 61.92699602 | 90.63464974 | 73.95528345 | 9.85E-07 | 1.862523 | 2.73E-07 | 5.141338 | 3.63643 | 6.006776 |
| CNAG_004 | - | hypothetic | 804.8348077 | 788.8096108 | 806.3382803 | 1123.467345 | 1118.115575 | 1043.450322 | 1.00E-06 | 0.438222 | 2.78E-07 | 5.137959 | 1.354933 | 5.999542 |
| CNAG_129 | - | unspecifie | 119.7674754 | 185.4481928 | 169.7160208 | 283.8165926 | 302.3290214 | 294.993202 | 1.01E-06 | 0.880117 | 2.79E-07 | 5.136965 | 1.840524 | 5.997437 |
| CNAG_062 | - | hypothetic | 282.9635004 | 321.1745832 | 233.7714229 | 481.6307254 | 428.8957522 | 467.1016065 | 1.01E-06 | 0.704592 | 2.81E-07 | 5.136031 | 1.629683 | 5.995469 |
| CNAG_043 | - | hypothetic | 1628.145611 | 1831.698564 | 1626.460427 | 2159.157391 | 2469.847115 | 2173.597704 | 1.02E-06 | 0.404624 | 2.84E-07 | 5.133458 | 1.323744 | 5.989718 |
| CNAG_070 | - | hypothetic | 1242.213954 | 1252.919176 | 1145.323812 | 861.9352892 | 922.7584588 | 938.216717 | 1.06E-06 | -0.43452 | 2.95E-07 | -5.1265 | 0.739941 | 5.974045 |

| | | | | | | | | | | | | | | |
|---|---|---|---|---|---|---|---|---|---|---|---|---|---|---|
| CNAG_004 | - | RING zinc | 120.8052397 | 164.9479412 | 106.8892933 | 255.9690574 | 241.8447018 | 257.2102132 | 1.07E-06 | 0.935491 | 2.97E-07 | 5.125657 | 1.912541 | 5.972294 |
| CNAG_079 | - | uracil tran | 951.5891632 | 1034.201642 | 1045.001326 | 700.0251029 | 768.3696122 | 757.2806478 | 1.09E-06 | -0.46053 | 3.05E-07 | -5.12055 | 0.726721 | 5.96092 |
| CNAG_034 | - | hypothetic | 21.64117137 | 19.69502237 | 33.23403955 | 65.73092101 | 86.03067268 | 94.82135571 | 1.10E-06 | 1.730949 | 3.07E-07 | 5.119203 | 3.31946 | 5.958301 |
| CNAG_041 | YAK1 | CMGC/DY | 1060.563382 | 1061.383321 | 1065.481337 | 1412.110457 | 1476.095906 | 1357.631282 | 1.14E-06 | 0.398666 | 3.17E-07 | 5.112927 | 1.318288 | 5.944124 |
| CNAG_057 | - | mitogen-a | 991.2703107 | 1010.545069 | 1123.881784 | 795.9246254 | 753.6538515 | 734.017921 | 1.14E-06 | -0.46841 | 3.18E-07 | -5.11241 | 0.722761 | 5.943133 |
| CNAG_035 | - | Phenylacry | 33.38354422 | 31.57253244 | 61.98548716 | 116.7334407 | 151.0867308 | 97.53938239 | 1.16E-06 | 1.525332 | 3.23E-07 | 5.109574 | 2.87853 | 5.937176 |
| CNAG_074 | - | asparagine | 1881.182925 | 2111.635886 | 2062.426644 | 1613.159756 | 1590.39985 | 1512.214489 | 1.18E-06 | -0.37548 | 3.30E-07 | -5.10559 | 0.770849 | 5.928218 |
| CNAG_018 | - | CCR4-NOT | 2656.55628 | 2840.017564 | 2896.391086 | 2256.023197 | 2291.694376 | 2151.654451 | 1.19E-06 | -0.34039 | 3.33E-07 | -5.10398 | 0.789827 | 5.924707 |
| CNAG_075 | - | hypothetic | 1091.974521 | 703.5449842 | 739.0023168 | 510.8072154 | 537.0940227 | 505.3932629 | 1.19E-06 | -0.7239 | 3.34E-07 | -5.10342 | 0.605458 | 5.923751 |
| CNAG_007 | - | dityrosine | 309.8980575 | 393.3371288 | 290.2759723 | 481.9812589 | 551.5342087 | 575.8241824 | 1.23E-06 | 0.681901 | 3.45E-07 | 5.09698 | 1.604252 | 5.910156 |
| CNAG_066 | - | hypothetic | 1366.060703 | 1374.701446 | 1343.660846 | 1058.976128 | 1040.920379 | 1049.24789 | 1.23E-06 | -0.39083 | 3.45E-07 | -5.097 | 0.76269 | 5.910156 |
| CNAG_043 | - | DNA-direc | 1629.919687 | 1502.968508 | 1633.70076 | 1255.92104 | 1165.088527 | 1216.884156 | 1.23E-06 | -0.40538 | 3.46E-07 | -5.09658 | 0.755037 | 5.909417 |
| CNAG_033 | - | AP-1 com | 2732.423252 | 2776.475765 | 2926.249766 | 2264.576954 | 2230.411805 | 2279.277533 | 1.24E-06 | -0.33164 | 3.49E-07 | -5.09482 | 0.79463 | 5.905581 |
| CNAG_065 | SPT8 | transcripti | 2172.966944 | 2187.080435 | 2061.406838 | 1593.570023 | 1773.319154 | 1664.276336 | 1.26E-06 | -0.36728 | 3.54E-07 | -5.09215 | 0.775243 | 5.899644 |
| CNAG_033 | - | hypothetic | 1540.972735 | 1202.423576 | 1489.682695 | 987.2502515 | 977.8866062 | 1041.484064 | 1.27E-06 | -0.50997 | 3.57E-07 | -5.09036 | 0.702237 | 5.895728 |
| CNAG_064 | - | rRNA-proc | 1311.200127 | 1243.316503 | 1541.881897 | 921.0315653 | 960.6473854 | 1051.440103 | 1.30E-06 | -0.49759 | 3.66E-07 | -5.08598 | 0.708288 | 5.886094 |
| CNAG_012 | - | hypothetic | 3441.173166 | 3234.416366 | 3568.248308 | 2775.938187 | 2777.308839 | 2604.606698 | 1.36E-06 | -0.34439 | 3.84E-07 | -5.07676 | 0.78764 | 5.865572 |
| CNAG_012 | - | hypothetic | 2336.483207 | 2311.015003 | 2309.871828 | 1786.630556 | 1888.871505 | 1880.726791 | 1.37E-06 | -0.34004 | 3.86E-07 | -5.0759 | 0.79002 | 5.8638 |
| CNAG_068 | PMT1 | dolichyl-p | 2255.161323 | 2366.117459 | 2262.037841 | 2814.456218 | 2896.622442 | 3043.883327 | 1.37E-06 | 0.331779 | 3.87E-07 | 5.075037 | 1.258564 | 5.86201 |
| CNAG_048 | - | hypothetic | 281.8769415 | 356.7104025 | 278.5269199 | 507.2544845 | 509.2820355 | 449.4850748 | 1.40E-06 | 0.664729 | 3.94E-07 | 5.071904 | 1.585271 | 5.855233 |
| CNAG_032 | - | dioxygena | 49.56797696 | 44.44567996 | 51.38877402 | 139.7707622 | 126.499763 | 96.47321342 | 1.40E-06 | 1.310073 | 3.96E-07 | 5.070944 | 2.47954 | 5.853229 |
| CNAG_023 | - | L-serine a | 1041.375075 | 1009.462248 | 1021.574311 | 785.7967075 | 752.7977219 | 759.5171388 | 1.47E-06 | -0.43492 | 4.15E-07 | -5.06213 | 0.739734 | 5.833873 |
| CNAG_029 | APH1 | phosphate | 143.1378386 | 250.0303157 | 167.6568493 | 318.6562158 | 347.098755 | 410.2210984 | 1.47E-06 | 0.928198 | 4.15E-07 | 5.062011 | 1.902898 | 5.833796 |
| CNAG_063 | - | protein CV | 783.1323162 | 714.2274235 | 730.4083485 | 539.0647499 | 414.4706822 | 537.4644889 | 1.47E-06 | -0.59503 | 4.16E-07 | -5.0617 | 0.662031 | 5.833286 |
| CNAG_055 | KIN4 | CAMK/CA | 315.5516836 | 283.5415275 | 274.2441339 | 476.1037446 | 478.2186858 | 422.8462511 | 1.48E-06 | 0.642409 | 4.18E-07 | 5.060593 | 1.560933 | 5.830942 |
| CNAG_059 | - | tRNA-spec | 851.0067357 | 904.8523298 | 777.3998989 | 590.3251429 | 565.5518686 | 642.9067756 | 1.48E-06 | -0.50938 | 4.20E-07 | -5.05963 | 0.702526 | 5.82894 |
| CNAG_064 | - | hypothetic | 2075.62455 | 1134.472648 | 1366.984806 | 631.1726193 | 529.8787199 | 501.1812456 | 1.50E-06 | -1.47976 | 4.25E-07 | -5.05749 | 0.358549 | 5.824238 |
| CNAG_002 | NAB2 | putative n | 1376.39492 | 1190.517137 | 1312.690956 | 923.4515578 | 859.6676797 | 1025.882519 | 1.56E-06 | -0.48195 | 4.44E-07 | -5.04926 | 0.71601 | 5.806086 |
| CNAG_060 | URO2 | hydroxyiso | 195.4344014 | 225.3054584 | 155.9439834 | 404.2270281 | 302.9045354 | 336.1305664 | 1.59E-06 | 0.843606 | 4.50E-07 | 5.046362 | 1.79453 | 5.799874 |
| CNAG_023 | - | Uracil pho | 19.17411225 | 8.920453188 | 15.15645714 | 69.79636489 | 68.92285284 | 45.39245755 | 1.59E-06 | 2.097696 | 4.52E-07 | 5.045817 | 4.280252 | 5.798823 |
| CNAG_033 | - | solute car | 2421.919053 | 2985.379691 | 2748.159557 | 2043.452772 | 2177.829972 | 2042.864108 | 1.62E-06 | -0.39532 | 4.60E-07 | -5.04244 | 0.760319 | 5.791344 |
| CNAG_037 | - | hypothetic | 2963.694905 | 3293.591335 | 3221.670415 | 2559.566825 | 2396.84045 | 2592.328002 | 1.62E-06 | -0.34322 | 4.62E-07 | -5.04143 | 0.788277 | 5.789787 |
| CNAG_055 | - | glycogen | 350.7387187 | 369.7236612 | 416.0003873 | 565.4314171 | 601.5546019 | 550.4801876 | 1.62E-06 | 0.58103 | 4.62E-07 | 5.04144 | 1.495917 | 5.789787 |
| CNAG_050 | - | NADH-ubi | 2438.195976 | 2774.241057 | 2573.284884 | 2131.542583 | 1981.097942 | 2026.187908 | 1.63E-06 | -0.35753 | 4.63E-07 | -5.04105 | 0.780497 | 5.789125 |
| CNAG_052 | - | hypothetic | 1699.793548 | 1442.62651 | 1452.456931 | 1160.913248 | 1157.370199 | 1098.143868 | 1.68E-06 | -0.44349 | 4.80E-07 | -5.03418 | 0.735353 | 5.774103 |
| CNAG_014 | - | ribosome | 3301.030007 | 3066.3411 | 3375.226152 | 2589.886989 | 2563.356627 | 2638.965797 | 1.71E-06 | -0.3382 | 4.87E-07 | -5.03125 | 0.791027 | 5.768191 |
| CNAG_019 | - | hypothetic | 32.19744215 | 58.33086131 | 34.3835235 | 112.1209755 | 99.50858563 | 125.979394 | 1.80E-06 | 1.425412 | 5.14E-07 | 5.020939 | 2.685912 | 5.745229 |
| CNAG_011 | NEM1 | nuclear er | 361.2669038 | 333.1621252 | 459.6958456 | 570.733489 | 611.853974 | 653.5516353 | 1.80E-06 | 0.653809 | 5.15E-07 | 5.020475 | 1.573316 | 5.744365 |
| CNAG_040 | - | hypothetic | 887.3182434 | 916.7859856 | 999.1026812 | 729.0579956 | 589.6501123 | 661.8409988 | 1.82E-06 | -0.51672 | 5.20E-07 | -5.01872 | 0.698957 | 5.740572 |
| CNAG_049 | - | hypothetic | 2774.364019 | 2755.978808 | 2614.902644 | 2235.047046 | 2211.468294 | 2069.511183 | 1.84E-06 | -0.33708 | 5.26E-07 | -5.01656 | 0.791643 | 5.736061 |
| CNAG_049 | - | protein FR | 1253.840158 | 1122.633265 | 1214.587054 | 878.3373914 | 942.6397221 | 851.6939281 | 1.88E-06 | -0.44207 | 5.40E-07 | -5.01137 | 0.736078 | 5.724901 |
| CNAG_054 | - | hypothetic | 671.3353747 | 729.2927473 | 783.6726997 | 521.8856037 | 482.7415776 | 530.814624 | 1.89E-06 | -0.52488 | 5.44E-07 | -5.01028 | 0.695018 | 5.722634 |
| CNAG_007 | - | coiled-coil | 1069.455826 | 1034.269462 | 1158.023131 | 806.7597946 | 765.6969994 | 838.2998192 | 1.90E-06 | -0.45225 | 5.47E-07 | -5.00904 | 0.730901 | 5.720208 |
| CNAG_013 | - | hypothetic | 636.2027092 | 677.5440943 | 679.19069 | 450.9997806 | 510.5172521 | 384.4314278 | 1.92E-06 | -0.58306 | 5.51E-07 | -5.00747 | 0.667544 | 5.716848 |
| CNAG_020 | - | hypothetic | 1203.834066 | 1243.274723 | 1367.046773 | 945.2830082 | 961.5636542 | 969.3171376 | 1.93E-06 | -0.42261 | 5.55E-07 | -5.00628 | 0.746075 | 5.714355 |
| CNAG_030 | PER1 | Protein pr | 696.4265357 | 580.5942084 | 499.0888559 | 361.3686571 | 399.9915731 | 379.9044424 | 2.02E-06 | -0.65594 | 5.83E-07 | -4.99683 | 0.634663 | 5.693602 |
| CNAG_046 | - | hypothetic | 302.8283414 | 330.90038 | 312.6078215 | 518.9797872 | 477.245793 | 453.920384 | 2.10E-06 | 0.602275 | 6.05E-07 | 4.989483 | 1.518109 | 5.677263 |
| CNAG_005 | - | hypothetic | 1053.489772 | 984.8972757 | 960.9933348 | 1291.389378 | 1404.512755 | 1337.597844 | 2.13E-06 | 0.41244 | 6.13E-07 | 4.986985 | 1.330935 | 5.6722 |
| CNAG_040 | - | potassium | 1041.398079 | 1099.897758 | 987.4819146 | 817.7586375 | 716.5081341 | 770.6163645 | 2.16E-06 | -0.45606 | 6.25E-07 | -4.98344 | 0.728975 | 5.664794 |
| CNAG_052 | - | hypothetic | 59.16507354 | 121.8949467 | 110.0094147 | 225.868978 | 203.957978 | 205.0590362 | 2.22E-06 | 1.113084 | 6.42E-07 | 4.978143 | 2.163076 | 5.653085 |
| CNAG_075 | - | hypothetic | 1007.502792 | 921.159417 | 1095.071848 | 656.365847 | 720.0125732 | 776.1171833 | 2.22E-06 | -0.50601 | 6.42E-07 | -4.97805 | 0.704166 | 5.653054 |
| CNAG_044 | HFM1 | ATP-depe | 84.65837308 | 106.8496758 | 104.6777668 | 167.5810028 | 218.8358359 | 202.7915839 | 2.26E-06 | 0.982789 | 6.53E-07 | 4.974987 | 1.976282 | 5.646557 |
| CNAG_064 | - | hypothetic | 2111.011258 | 1896.244134 | 2059.228466 | 1546.928146 | 1518.756017 | 1638.699883 | 2.26E-06 | -0.38312 | 6.53E-07 | -4.97473 | 0.76678 | 5.64617 |
| CNAG_069 | - | L-serine a | 67.153763 | 86.38367059 | 80.17098847 | 171.2342312 | 138.1543361 | 189.2949076 | 2.28E-06 | 1.081762 | 6.59E-07 | 4.972971 | 2.116619 | 5.642405 |
| CNAG_067 | - | Acyl-CoA- | 240.8835836 | 243.6446909 | 215.5993077 | 393.6552676 | 368.5829714 | 361.7219529 | 2.29E-06 | 0.669705 | 6.64E-07 | 4.971598 | 1.590748 | 5.639511 |
| CNAG_035 | - | hypothetic | 74.19703769 | 102.5365133 | 102.5344369 | 172.4309395 | 184.9680695 | 208.2338148 | 2.32E-06 | 1.006358 | 6.73E-07 | 4.9689 | 2.008834 | 5.63365 |
| CNAG_069 | - | hypothetic | 1152.262623 | 1099.958788 | 1057.865395 | 815.2999515 | 829.5866402 | 847.1920743 | 2.36E-06 | -0.42458 | 6.83E-07 | -4.9661 | 0.745055 | 5.627564 |
| CNAG_014 | HPI1 | E3 ubiquit | 1194.278925 | 1066.606456 | 1167.653444 | 824.6162163 | 790.6957175 | 904.8826781 | 2.38E-06 | -0.46025 | 6.89E-07 | -4.96435 | 0.726858 | 5.624004 |
| CNAG_041 | - | hypothetic | 189.47563 | 183.3191721 | 121.8420635 | 288.4305948 | 293.5870535 | 331.5084346 | 2.40E-06 | 0.872636 | 6.98E-07 | 4.962028 | 1.831005 | 5.619003 |
| CNAG_079 | - | hypothetic | 458.0223102 | 430.0525832 | 441.6340225 | 684.6658212 | 612.5736583 | 642.5981582 | 2.44E-06 | 0.528866 | 7.08E-07 | 4.959088 | 1.442795 | 5.612709 |
| CNAG_025 | - | hypothetic | 3484.429914 | 3373.402437 | 3857.224968 | 2659.824979 | 2799.699557 | 2955.336461 | 2.45E-06 | -0.3643 | 7.12E-07 | -4.95801 | 0.776844 | 5.610566 |
| CNAG_010 | - | pre-mRNA | 1732.715878 | 1635.48752 | 1728.632272 | 1344.626935 | 1343.709042 | 1304.594851 | 2.45E-06 | -0.36788 | 7.13E-07 | -4.95778 | 0.774921 | 5.610236 |
| CNAG_076 | - | hypothetic | 36.77437316 | 49.74192987 | 34.36779512 | 100.3300664 | 106.6374446 | 101.7698507 | 2.47E-06 | 1.35312 | 7.18E-07 | 4.956423 | 2.55464 | 5.607569 |
| CNAG_120 | - | unspecifie | 18.12090718 | 27.14505133 | 19.43036429 | 73.33098811 | 74.80023774 | 64.09709423 | 2.47E-06 | 1.706976 | 7.18E-07 | 4.956424 | 3.264758 | 5.607569 |
| CNAG_005 | - | hypothetic | 321.5364264 | 313.7315323 | 354.1796635 | 465.5578567 | 497.0203869 | 647.7902347 | 2.55E-06 | 0.688438 | 7.43E-07 | 4.94967 | 1.611538 | 5.593039 |
| CNAG_061 | HAD8 | glycerol-1 | 31.04702677 | 49.77391511 | 49.24187096 | 133.6578209 | 99.49168937 | 104.0650354 | 2.56E-06 | 1.367142 | 7.45E-07 | 4.949273 | 2.57959 | 5.592335 |
| CNAG_051 | - | membrane | 4318.330315 | 5029.313825 | 4853.337233 | 3553.3098 | 3692.440768 | 3988.88158 | 2.57E-06 | -0.35266 | 7.48E-07 | -4.94837 | 0.783136 | 5.5905 |
| CNAG_022 | - | RNA polyr | 2318.894249 | 2222.660718 | 2159.518767 | 1806.189973 | 1822.464774 | 1638.794179 | 2.58E-06 | -0.36274 | 7.53E-07 | -4.94727 | 0.777687 | 5.588227 |

| | | | | | | | | | | | | | | |
|---|---|---|---|---|---|---|---|---|---|---|---|---|---|---|
| CNAG_02 | - | hypothetic | 1094.398786 | 981.7040234 | 1119.848566 | 1422.221947 | 1406.133488 | 1480.760149 | 2.69E-06 | 0.415962 | 7.85E-07 | 4.939126 | 1.334188 | 5.57099 |
| CNAG_03 | - | cytoplasm | 649.8954832 | 510.6750135 | 609.852545 | 344.9552837 | 413.8230085 | 400.9590556 | 2.72E-06 | -0.62812 | 7.96E-07 | -4.93629 | 0.647019 | 5.565049 |
| CNAG_03 | - | monolyso | 1140.604304 | 1045.053415 | 1136.727268 | 798.9228416 | 803.6768209 | 872.6978626 | 2.73E-06 | -0.44091 | 7.98E-07 | -4.93584 | 0.736671 | 5.564222 |
| CNAG_04 | - | DNA-direc | 1831.322604 | 1867.311315 | 2020.989315 | 2726.086241 | 2397.16372 | 2402.300519 | 2.76E-06 | 0.380963 | 8.08E-07 | 4.933455 | 1.30221 | 5.559273 |
| CNAG_05 | - | hypothetic | 13.72061151 | 39.17393402 | 56.81274115 | 208.1284663 | 274.8032636 | 213.6739662 | 2.84E-06 | 2.672351 | 8.31E-07 | 4.927869 | 6.37467 | 5.547396 |
| CNAG_12 | - | unspecifie | 166.6465968 | 196.8475881 | 174.7253463 | 107.0035273 | 77.15544453 | 86.91393418 | 2.91E-06 | -1.00362 | 8.53E-07 | -4.92294 | 0.498747 | 5.536633 |
| CNAG_00 | - | hypothetic | 518.3887065 | 591.3514902 | 625.8234735 | 415.2869117 | 387.0203342 | 379.9047009 | 2.97E-06 | -0.56896 | 8.74E-07 | -4.91813 | 0.6741 | 5.526682 |
| CNAG_06 | - | diphthami | 1315.959868 | 1426.405552 | 1438.540208 | 1034.767416 | 1095.292992 | 1091.419302 | 2.97E-06 | -0.39109 | 8.74E-07 | -4.91813 | 0.762553 | 5.526682 |
| CNAG_03 | - | transcripti | 519.9126459 | 482.8525903 | 508.8132545 | 776.5074999 | 660.8032815 | 758.0003729 | 3.05E-06 | 0.524656 | 8.95E-07 | 4.913384 | 1.43859 | 5.516337 |
| CNAG_07 | CAS1 | O-acetyltr | 1654.508561 | 1679.613011 | 1661.453379 | 1389.110167 | 1270.361664 | 1168.112404 | 3.07E-06 | -0.39922 | 9.04E-07 | -4.91144 | 0.75827 | 5.51221 |
| CNAG_06 | ALO1 | d-arabino | 2249.989931 | 2048.15775 | 2187.211771 | 1639.520259 | 1674.908035 | 1773.022727 | 3.08E-06 | -0.36596 | 9.07E-07 | -4.91077 | 0.77595 | 5.5109 |
| CNAG_05 | - | hypothetic | 347.9963987 | 449.0702005 | 487.1059884 | 271.9520126 | 251.5207264 | 271.1039573 | 3.13E-06 | -0.70852 | 9.21E-07 | -4.90789 | 0.611946 | 5.50471 |
| CNAG_05 | - | beta-cater | 860.8777716 | 823.2976807 | 882.0408635 | 1177.913686 | 1196.621864 | 1096.750435 | 3.16E-06 | 0.419998 | 9.31E-07 | 4.905629 | 1.337926 | 5.500063 |
| CNAG_12 | - | unspecifie | 23.90486665 | 25.02813361 | 25.80194338 | 68.31222707 | 77.33664253 | 77.2641804 | 3.16E-06 | 1.589526 | 9.33E-07 | 4.905311 | 3.009505 | 5.499722 |
| CNAG_00 | PDR5 | ATP-bindi | 1391.302857 | 1171.044382 | 1014.194596 | 742.7166787 | 778.5948122 | 913.764621 | 3.16E-06 | -0.57122 | 9.33E-07 | -4.90536 | 0.673048 | 5.499722 |
| CNAG_06 | - | nuclear pr | 4382.408991 | 4311.822523 | 4298.82253 | 3527.734627 | 3659.732223 | 3598.13995 | 3.20E-06 | -0.28396 | 9.43E-07 | -4.90319 | 0.821332 | 5.495505 |
| CNAG_04 | - | hypothetic | 21.66589956 | 32.548947 | 32.19008777 | 84.10165603 | 71.86015742 | 97.08320094 | 3.23E-06 | 1.554032 | 9.54E-07 | 4.900983 | 2.936367 | 5.490866 |
| CNAG_01 | - | vesicle tra | 1551.759477 | 1557.87924 | 1582.529199 | 1132.784756 | 1276.471527 | 1230.188345 | 3.30E-06 | -0.38178 | 9.75E-07 | -4.89656 | 0.767491 | 5.48146 |
| CNAG_03 | - | ATP(GTP)- | 667.7469562 | 676.5202171 | 711.1925873 | 483.7092665 | 506.1308074 | 487.5461664 | 3.36E-06 | -0.49205 | 9.93E-07 | -4.89309 | 0.711014 | 5.474157 |
| CNAG_00 | - | hypothetic | 75.18839842 | 75.62765614 | 70.58488146 | 145.3084642 | 153.0985625 | 156.1672391 | 3.36E-06 | 1.030032 | 9.94E-07 | 4.892862 | 2.04207 | 5.473823 |
| CNAG_01 | - | hypothetic | 496.5725808 | 531.2615624 | 437.4137886 | 694.573297 | 726.5609701 | 697.0089416 | 3.49E-06 | 0.517244 | 1.03E-06 | 4.884945 | 1.431218 | 5.456719 |
| CNAG_07 | - | Xaa-Pro a | 869.648942 | 865.0325416 | 776.3352695 | 639.4935006 | 592.3355942 | 594.1288199 | 3.53E-06 | -0.4749 | 1.05E-06 | -4.88265 | 0.719519 | 5.451836 |
| CNAG_03 | - | aspartyl/g | 1835.536864 | 1810.020312 | 1883.269894 | 1473.050119 | 1485.182647 | 1441.134757 | 3.63E-06 | -0.34496 | 1.08E-06 | -4.87687 | 0.787331 | 5.439653 |
| CNAG_00 | - | protein di | 2290.901393 | 2160.198088 | 2228.818166 | 1824.100935 | 1773.261368 | 1745.319094 | 3.64E-06 | -0.33754 | 1.08E-06 | -4.87632 | 0.791389 | 5.438621 |
| CNAG_03 | - | Mannosyl- | 2931.332203 | 3244.210026 | 3110.921096 | 3731.254262 | 3886.843421 | 4071.895663 | 3.67E-06 | 0.317289 | 1.09E-06 | 4.874749 | 1.245987 | 5.435339 |
| CNAG_07 | - | hypothetic | 488.5457723 | 581.9136835 | 579.1726675 | 808.3142516 | 787.7530462 | 758.1016677 | 3.69E-06 | 0.499072 | 1.10E-06 | 4.873641 | 1.413304 | 5.43308 |
| CNAG_03 | YCH1 | hypothetic | 852.342664 | 928.6175456 | 942.6218254 | 660.4072603 | 659.6342007 | 696.2042109 | 3.69E-06 | -0.4488 | 1.10E-06 | -4.87332 | 0.732654 | 5.432556 |
| CNAG_07 | - | HMG box | 0.362084733 | 4.516377287 | 2.317120719 | 27.93803829 | 19.93687702 | 45.66873396 | 3.70E-06 | 3.911574 | 1.10E-06 | 4.87295 | 15.04877 | 5.431917 |
| CNAG_04 | CHI22 | chitinase, | 1616.692011 | 1347.757275 | 1159.246855 | 978.7247598 | 949.4225386 | 974.9054847 | 3.77E-06 | -0.52258 | 1.12E-06 | -4.86931 | 0.696126 | 5.424104 |
| CNAG_06 | HAD7 | magnesiu | 783.1114019 | 675.4802618 | 760.2283594 | 409.4761048 | 583.0140355 | 472.0617264 | 3.77E-06 | -0.61567 | 1.12E-06 | -4.86898 | 0.652626 | 5.423557 |
| CNAG_00 | - | hypothetic | 657.6078264 | 612.1018986 | 481.1777745 | 899.4414235 | 833.4255256 | 867.9392973 | 3.78E-06 | 0.555431 | 1.12E-06 | 4.868422 | 1.469607 | 5.422499 |
| CNAG_12 | - | unspecifie | 8.681743718 | 8.855715611 | 7.67653134 | 49.54814737 | 35.67684197 | 40.69876771 | 3.78E-06 | 2.415775 | 1.13E-06 | 4.868177 | 5.336059 | 5.422139 |
| CNAG_02 | - | hypothetic | 339.2688495 | 357.398165 | 277.2211777 | 226.2580661 | 174.7226403 | 138.0812126 | 3.89E-06 | -0.86969 | 1.16E-06 | -4.86272 | 0.547263 | 5.410509 |
| CNAG_01 | - | hypothetic | 412.5977187 | 381.6247658 | 490.6339779 | 594.8044017 | 650.6311998 | 723.4643722 | 4.09E-06 | 0.60081 | 1.22E-06 | 4.8523 | 1.516567 | 5.388198 |
| CNAG_04 | - | hypothetic | 9.947580209 | 12.12099562 | 19.37113262 | 61.87045709 | 61.07238767 | 44.23296899 | 4.28E-06 | 2.037661 | 1.28E-06 | 4.843368 | 4.105795 | 5.369005 |
| CNAG_13 | - | unspecifie | 25.23467369 | 52.95647847 | 38.614945 | 125.8951961 | 102.2297338 | 89.72830191 | 4.36E-06 | 1.445249 | 1.30E-06 | 4.839199 | 2.723098 | 5.360247 |
| CNAG_02 | - | DNA repli | 205.8194099 | 188.7307558 | 175.0818393 | 321.9902569 | 289.1278308 | 340.4645593 | 4.38E-06 | 0.726827 | 1.31E-06 | 4.838297 | 1.654995 | 5.358454 |
| CNAG_03 | - | hypothetic | 712.9797033 | 606.5108132 | 611.0005094 | 462.760629 | 462.9963132 | 396.6052287 | 4.39E-06 | -0.56331 | 1.32E-06 | -4.83746 | 0.676747 | 5.357163 |
| CNAG_02 | - | hypothetic | 5.214333492 | 9.917840828 | 8.731391136 | 33.10176775 | 46.43722222 | 43.91365511 | 4.39E-06 | 2.456997 | 1.31E-06 | 4.837504 | 5.490728 | 5.357163 |
| CNAG_01 | - | hypothetic | 264.3248207 | 272.7659454 | 300.8457424 | 385.3821776 | 476.5604569 | 461.5329001 | 4.40E-06 | 0.645787 | 1.32E-06 | 4.837266 | 1.564593 | 5.356911 |
| CNAG_07 | - | DNA poly | 1623.438007 | 1807.982392 | 1615.783327 | 2192.741688 | 2156.560767 | 2126.9592 | 4.43E-06 | 0.345075 | 1.33E-06 | 4.8356 | 1.270217 | 5.353449 |
| CNAG_04 | CDC24 | cell divisi | 1734.946057 | 1719.436754 | 1474.894757 | 1329.419486 | 1175.419495 | 1206.924163 | 4.45E-06 | -0.42431 | 1.33E-06 | -4.83483 | 0.745196 | 5.352126 |
| CNAG_05 | - | hypothetic | 54.2798942 | 57.34946421 | 70.53472952 | 125.8447373 | 143.7490068 | 131.7846594 | 4.48E-06 | 1.124976 | 1.34E-06 | 4.83327 | 2.180979 | 5.348895 |
| CNAG_02 | MEC1 | serine/thr | 2893.836826 | 2849.886952 | 2657.707141 | 3545.374631 | 3484.019329 | 3468.040416 | 4.50E-06 | 0.30647 | 1.35E-06 | 4.832386 | 1.236678 | 5.347143 |
| CNAG_06 | - | PEK/GCN2 | 4664.980299 | 4498.224534 | 4530.240818 | 3699.73297 | 3680.381441 | 3914.501864 | 4.57E-06 | -0.29322 | 1.37E-06 | -4.82897 | 0.816076 | 5.340033 |
| CNAG_04 | PDR16 | pleiotropi | 2082.032202 | 2182.806072 | 2413.24238 | 1693.230712 | 1825.061722 | 1697.597247 | 4.60E-06 | -0.37186 | 1.38E-06 | -4.82769 | 0.772786 | 5.33743 |
| CNAG_00 | - | MFS trans | 73.01075918 | 100.3715814 | 95.08041239 | 166.2163415 | 169.3640388 | 205.9659259 | 4.61E-06 | 0.996626 | 1.38E-06 | 4.827202 | 1.995328 | 5.336541 |
| CNAG_02 | - | Sugar trar | 387.4194825 | 379.148634 | 433.8434783 | 260.2912668 | 235.1204123 | 277.7050735 | 4.61E-06 | -0.65042 | 1.39E-06 | -4.8268 | 0.637094 | 5.336022 |
| CNAG_02 | - | hypothetic | 58.92570758 | 78.79443361 | 43.9978826 | 159.1220686 | 144.5596239 | 118.6340271 | 4.61E-06 | 1.219969 | 1.39E-06 | 4.82686 | 2.329417 | 5.336022 |
| CNAG_00 | - | hypothetic | 617.3558689 | 556.949357 | 583.243513 | 424.6237273 | 403.4262073 | 402.0827165 | 4.70E-06 | -0.53043 | 1.42E-06 | -4.82286 | 0.692346 | 5.327791 |
| CNAG_06 | - | scavenger | 203.6947659 | 284.5024678 | 201.7637207 | 397.5426571 | 362.4996783 | 394.9229309 | 4.74E-06 | 0.731316 | 1.43E-06 | 4.821131 | 1.660152 | 5.324197 |
| CNAG_03 | LAC1 | laccase | 393.7456051 | 360.0124852 | 293.4856349 | 492.0547176 | 548.0049267 | 620.1758553 | 4.80E-06 | 0.651078 | 1.45E-06 | 4.818534 | 1.570341 | 5.318719 |
| CNAG_01 | - | hypothetic | 3.935449359 | 3.438970187 | 0.145259058 | 33.0571198 | 34.4052314 | 19.8827657 | 4.86E-06 | 3.826072 | 1.46E-06 | 4.816117 | 14.18281 | 5.313636 |
| CNAG_00 | - | hypothetic | 123.041252 | 138.0559686 | 107.9248859 | 244.4308476 | 210.7179659 | 220.6608023 | 4.91E-06 | 0.858508 | 1.48E-06 | 4.813681 | 1.813162 | 5.308513 |
| CNAG_02 | PXA2 | ATP-bindi | 312.0341279 | 263.0951624 | 290.2068644 | 452.6753441 | 435.8326265 | 447.1697394 | 4.93E-06 | 0.609193 | 1.49E-06 | 4.813092 | 1.525405 | 5.307411 |
| CNAG_07 | - | hypothetic | 827.6539507 | 806.8897054 | 799.7548553 | 598.9560726 | 557.8015272 | 626.2537518 | 4.93E-06 | -0.46463 | 1.49E-06 | -4.81283 | 0.724658 | 5.307019 |
| CNAG_07 | - | glycylpept | 2582.852705 | 2368.171301 | 2676.684412 | 2094.189259 | 1983.709352 | 1942.945336 | 5.08E-06 | -0.35713 | 1.53E-06 | -4.80677 | 0.780717 | 5.294027 |
| CNAG_08 | - | hypothetic | 2.820897899 | 4.535761152 | 4.460636953 | 56.11773611 | 26.10388462 | 19.98205819 | 5.10E-06 | 3.322136 | 1.54E-06 | 4.805934 | 10.00144 | 5.292386 |
| CNAG_03 | - | translatior | 1043.769267 | 1089.152513 | 1027.992778 | 750.6391165 | 797.6782486 | 838.2791484 | 5.10E-06 | -0.42046 | 1.54E-06 | -4.80571 | 0.747184 | 5.29208 |
| CNAG_02 | LPI3 | hypothetic | 665.3101258 | 668.9378418 | 649.3669972 | 433.8152222 | 401.5933178 | 524.0670479 | 5.17E-06 | -0.56034 | 1.56E-06 | -4.80286 | 0.67814 | 5.286247 |
| CNAG_06 | - | G protein | 1286.63857 | 1233.593513 | 1303.107186 | 959.2997918 | 950.3263859 | 1027.001852 | 5.18E-06 | -0.39584 | 1.57E-06 | -4.80264 | 0.760047 | 5.285937 |
| CNAG_06 | - | glycoprote | 217.6089556 | 236.1052344 | 223.0401613 | 405.4964087 | 374.6927656 | 319.6540862 | 5.23E-06 | 0.684729 | 1.59E-06 | 4.80018 | 1.6074 | 5.281129 |
| CNAG_05 | - | TDG/mug | 263.0001747 | 235.0579011 | 219.8685062 | 360.6560881 | 370.2707806 | 427.0255959 | 5.39E-06 | 0.674981 | 1.63E-06 | 4.794153 | 1.596576 | 5.268766 |
| CNAG_04 | - | drug trans | 152.3901319 | 218.8247358 | 191.047469 | 295.2826107 | 318.6002558 | 354.8433898 | 5.41E-06 | 0.771626 | 1.64E-06 | 4.793139 | 1.707193 | 5.266743 |
| CNAG_07 | - | hypothetic | 18.06159575 | 21.77021486 | 15.16785528 | 74.49341665 | 55.53828369 | 58.5173607 | 5.44E-06 | 1.777588 | 1.65E-06 | 4.791929 | 3.428525 | 5.264648 |
| CNAG_04 | CPS1 | polysacch | 692.8682727 | 838.3107978 | 731.717065 | 1210.957181 | 1009.326247 | 1002.379924 | 5.53E-06 | 0.496222 | 1.68E-06 | 4.788483 | 1.410515 | 5.257363 |
| CNAG_01 | - | hypothetic | 295.6311855 | 237.2395801 | 219.8927259 | 471.7610913 | 393.5545777 | 382.859233 | 5.53E-06 | 0.714555 | 1.68E-06 | 4.788383 | 1.640977 | 5.257322 |
| CNAG_00 | - | peptidyl-p | 1187.453289 | 1210.947449 | 1308.402476 | 964.8334525 | 977.143683 | 835.0842112 | 5.55E-06 | -0.43192 | 1.69E-06 | -4.78753 | 0.741275 | 5.255817 |

| | | | | | | | | | | | | | | |
|---|---|---|---|---|---|---|---|---|---|---|---|---|---|---|
| CNAG_004 | ACT1 | actin | 3468.413405 | 3516.817703 | 3307.172975 | 4676.409452 | 4084.284765 | 4268.469609 | 5.55E-06 | 0.325228 | 1.69E-06 | 4.787316 | 1.252862 | 5.255535 |
| CNAG_035 | - | hypothetic | 67.19644634 | 105.7174081 | 84.44277666 | 158.3975094 | 186.8452757 | 179.4819641 | 5.73E-06 | 1.016837 | 1.75E-06 | 4.780616 | 2.023478 | 5.241575 |
| CNAG_003 | - | gly-X carb | 362.153179 | 279.2656403 | 283.8510435 | 497.8670206 | 464.2746304 | 491.552107 | 5.87E-06 | 0.636056 | 1.79E-06 | 4.775628 | 1.554075 | 5.231673 |
| CNAG_029 | - | hypothetic | 319.0615906 | 270.6422926 | 310.4563371 | 483.8670441 | 473.8615321 | 429.507841 | 6.15E-06 | 0.607522 | 1.88E-06 | 4.766006 | 1.52364 | 5.211272 |
| CNAG_040 | - | 26S protea | 4812.437513 | 4341.102471 | 4671.135043 | 5557.13551 | 5872.48997 | 5831.545179 | 6.38E-06 | 0.30448 | 1.95E-06 | 4.758439 | 1.234974 | 5.195501 |
| CNAG_059 | - | MFS trans | 275.8219818 | 242.5987437 | 226.2674741 | 386.4448146 | 371.9582235 | 427.0680717 | 6.38E-06 | 0.656837 | 1.95E-06 | 4.758218 | 1.576623 | 5.1952 |
| CNAG_004 | - | charged n | 812.1702968 | 631.3428468 | 700.5556111 | 482.896784 | 444.7439559 | 530.7862808 | 6.48E-06 | -0.57368 | 1.99E-06 | -4.75493 | 0.671901 | 5.188474 |
| CNAG_056 | - | ATP-depe | 1547.145133 | 1562.210355 | 1749.902825 | 1180.283143 | 1186.652164 | 1336.710765 | 6.55E-06 | -0.40727 | 2.01E-06 | -4.75261 | 0.754049 | 5.18367 |
| CNAG_131 | - | unspecifie | 5.109777188 | 0.175928197 | 4.447816883 | 22.05071044 | 37.01882975 | 34.97924178 | 6.66E-06 | 3.350648 | 2.04E-06 | 4.749065 | 10.20107 | 5.176393 |
| CNAG_017 | - | E3 ubiquit | 639.1742692 | 745.6885767 | 775.3807071 | 979.3722231 | 967.0864775 | 1062.175144 | 6.68E-06 | 0.463775 | 2.05E-06 | 4.748283 | 1.379145 | 5.175062 |
| CNAG_068 | - | vacuolar p | 73.00737365 | 86.42249802 | 106.7648689 | 160.6966324 | 171.9847509 | 207.0593765 | 6.70E-06 | 1.0068 | 2.06E-06 | 4.747805 | 2.009449 | 5.174208 |
| CNAG_007 | - | hypothetic | 242.1019169 | 256.5734391 | 253.9446523 | 366.0859311 | 406.6188827 | 392.7632217 | 6.86E-06 | 0.61708 | 2.11E-06 | 4.742633 | 1.533767 | 5.163628 |
| CNAG_045 | VPS4 | vacuolar p | 3286.87871 | 2886.40021 | 2917.773283 | 2413.997116 | 2341.658083 | 2465.77645 | 6.96E-06 | -0.34802 | 2.14E-06 | -4.73964 | 0.785662 | 5.157377 |
| CNAG_030 | - | polyamine | 404.4552726 | 423.598402 | 494.9002398 | 607.2846516 | 718.0696654 | 635.9565387 | 6.97E-06 | 0.553741 | 2.15E-06 | 4.739175 | 1.467887 | 5.15656 |
| CNAG_016 | - | peptidyl-p | 1508.522118 | 1453.390313 | 1504.682991 | 1170.276374 | 1128.883838 | 1205.742249 | 7.01E-06 | -0.3654 | 2.16E-06 | -4.73796 | 0.776256 | 5.154135 |
| CNAG_024 | YRM101 | putative tr | 156.8495845 | 162.8455318 | 140.9593446 | 271.4676937 | 239.1075748 | 277.2032085 | 7.10E-06 | 0.766242 | 2.19E-06 | 4.735445 | 1.700834 | 5.148914 |
| CNAG_017 | HAD1 | phosphata | 2050.133261 | 1918.769906 | 1594.364159 | 1397.497124 | 1423.052534 | 1344.593231 | 7.17E-06 | -0.43301 | 2.21E-06 | -4.73301 | 0.740714 | 5.144227 |
| CNAG_024 | - | GC-rich se | 2727.759156 | 2840.026298 | 2853.75211 | 2237.310352 | 2182.081445 | 2400.236699 | 7.25E-06 | -0.31924 | 2.24E-06 | -4.73086 | 0.801494 | 5.139792 |
| CNAG_078 | - | hypothetic | 1844.921258 | 1986.660731 | 1863.036493 | 1519.753745 | 1425.589146 | 1568.750061 | 7.34E-06 | -0.35018 | 2.26E-06 | -4.72826 | 0.784486 | 5.134563 |
| CNAG_013 | - | co-chaper | 3533.378221 | 3286.121884 | 3323.02029 | 2771.260794 | 2730.707963 | 2799.930942 | 7.36E-06 | -0.30466 | 2.27E-06 | -4.72747 | 0.809632 | 5.13306 |
| CNAG_058 | - | ubiquitin- | 1667.234719 | 1453.431325 | 1603.852034 | 1230.205213 | 1203.942673 | 1217.990525 | 7.48E-06 | -0.38764 | 2.31E-06 | -4.72402 | 0.764376 | 5.126032 |
| CNAG_025 | - | DNA repa | 21.47828721 | 19.62498619 | 4.531530614 | 52.64901898 | 72.42403337 | 63.91568331 | 7.53E-06 | 2.072461 | 2.33E-06 | 4.722365 | 4.206035 | 5.123172 |
| CNAG_017 | - | hypothetic | 1187.724158 | 1052.783986 | 1032.45028 | 1446.35188 | 1437.195421 | 1467.470961 | 7.61E-06 | 0.395661 | 2.36E-06 | 4.720061 | 1.315545 | 5.118422 |
| CNAG_047 | - | MFS trans | 0.35539212 | 3.440808875 | 3.379057055 | 13.00152232 | 44.34129813 | 32.77982835 | 7.83E-06 | 3.876627 | 2.42E-06 | 4.714369 | 14.68862 | 5.106448 |
| CNAG_025 | CPK2 | Calcium-d | 48.40772639 | 49.79965331 | 56.69034231 | 119.8642076 | 105.5111975 | 123.9064483 | 8.03E-06 | 1.166331 | 2.49E-06 | 4.708739 | 2.244401 | 5.095132 |
| CNAG_078 | - | hypothetic | 0.334307757 | 3.42731199 | 0.137978055 | 18.93285015 | 31.79601062 | 32.69296337 | 8.06E-06 | 4.742537 | 2.50E-06 | 4.707989 | 26.76985 | 5.093705 |
| CNAG_003 | - | 20S protea | 2235.270451 | 2248.686557 | 2243.89213 | 2828.570129 | 2758.594284 | 2755.321773 | 8.17E-06 | 0.295261 | 2.54E-06 | 4.705239 | 1.227107 | 5.08802 |
| CNAG_031 | - | THO comp | 1967.519142 | 1998.557423 | 2073.088727 | 1560.170603 | 1487.679652 | 1706.381928 | 8.17E-06 | -0.36028 | 2.54E-06 | -4.70515 | 0.779011 | 5.087993 |
| CNAG_015 | - | vacuolar r | 609.9789689 | 724.1283165 | 725.2678812 | 893.6509816 | 982.7159147 | 992.2636055 | 8.45E-06 | 0.463603 | 2.63E-06 | 4.698065 | 1.378982 | 5.073271 |
| CNAG_060 | - | cytoplasm | 963.2534565 | 1010.524009 | 1022.627123 | 742.9210454 | 777.002195 | 766.1560441 | 8.45E-06 | -0.40549 | 2.63E-06 | -4.6981 | 0.754982 | 5.073271 |
| CNAG_045 | - | hypothetic | 182.6845413 | 238.2233107 | 218.7577613 | 358.5996022 | 325.3867879 | 354.9761763 | 8.47E-06 | 0.686575 | 2.63E-06 | 4.697536 | 1.609458 | 5.072317 |
| CNAG_124 | - | unspecifie | 13437.19832 | 15878.98142 | 18965.43854 | 9301.58117 | 7234.632141 | 9655.312596 | 8.56E-06 | -0.8968 | 2.67E-06 | -4.69493 | 0.537076 | 5.067287 |
| CNAG_040 | - | RNA bindi | 14089.97077 | 13557.3058 | 14673.25112 | 11747.28476 | 11934.27209 | 12046.44644 | 8.56E-06 | -0.2598 | 2.67E-06 | -4.69499 | 0.835203 | 5.067287 |
| CNAG_079 | - | protein pł | 6083.841915 | 5993.701587 | 5997.686104 | 4962.129748 | 5139.026417 | 5155.674654 | 8.62E-06 | -0.25978 | 2.69E-06 | -4.69348 | 0.835216 | 5.064385 |
| CNAG_010 | - | nitric oxid | 792.8491604 | 883.3772247 | 976.6864528 | 683.8843539 | 612.1554434 | 617.4530533 | 8.65E-06 | -0.4864 | 2.70E-06 | -4.69267 | 0.713802 | 5.062833 |
| CNAG_064 | PMA1 | plasma-m | 20701.47272 | 18893.74282 | 15846.45928 | 14740.57703 | 14722.04143 | 13649.61842 | 8.81E-06 | -0.37841 | 2.75E-06 | -4.68888 | 0.769285 | 5.05496 |
| CNAG_015 | - | large subu | 206.0282256 | 172.2548866 | 179.0197217 | 102.1882836 | 97.09471284 | 104.6092337 | 9.23E-06 | -0.89416 | 2.88E-06 | -4.67911 | 0.53806 | 5.03475 |
| CNAG_053 | GLO3 | ADP-ribos | 2177.567823 | 2019.042672 | 2065.646745 | 1429.227816 | 1758.686338 | 1632.069483 | 9.35E-06 | -0.3934 | 2.92E-06 | -4.67627 | 0.761332 | 5.029089 |
| CNAG_060 | - | Chorismat | 1114.832341 | 1009.47481 | 982.1555045 | 779.5303291 | 767.4664094 | 795.0149205 | 1.00E-05 | -0.42299 | 3.14E-06 | -4.66141 | 0.745876 | 4.998006 |
| CNAG_027 | - | cytoplasm | 1014.68878 | 1083.800173 | 1182.543047 | 779.4427362 | 880.5593088 | 793.950414 | 1.01E-05 | -0.43459 | 3.17E-06 | -4.65931 | 0.739906 | 4.993908 |
| CNAG_045 | - | zinc metal | 643.687187 | 650.8805925 | 634.6483276 | 840.7679623 | 931.8667518 | 856.9144906 | 1.04E-05 | 0.431753 | 3.27E-06 | 4.652931 | 1.348872 | 4.981316 |
| CNAG_027 | - | hypothetic | 155.9168532 | 211.3306833 | 228.3171248 | 298.2946916 | 365.3321198 | 353.8147741 | 1.06E-05 | 0.759826 | 3.31E-06 | 4.650349 | 1.693286 | 4.976045 |
| CNAG_061 | - | cytoplasm | 902.4924841 | 1015.824353 | 861.6565618 | 704.8433753 | 669.9773416 | 694.0068981 | 1.06E-05 | -0.44107 | 3.33E-06 | -4.64917 | 0.736588 | 4.973736 |
| CNAG_050 | - | hypothetic | 33.44327257 | 56.26833818 | 76.88992129 | 135.4581427 | 123.6945868 | 141.6362429 | 1.06E-05 | 1.259756 | 3.34E-06 | 4.648707 | 2.394552 | 4.972927 |
| CNAG_064 | - | Cystathior | 74.00716798 | 71.32727239 | 69.51324244 | 139.8364163 | 132.2081288 | 169.3234432 | 1.08E-05 | 1.024877 | 3.39E-06 | 4.645948 | 2.034785 | 4.967457 |
| CNAG_023 | - | hypothetic | 1007.528505 | 1039.581644 | 938.4361328 | 775.6990902 | 772.6914733 | 709.5964385 | 1.09E-05 | -0.41858 | 3.42E-06 | -4.6441 | 0.74816 | 4.963904 |
| CNAG_013 | - | pumilio do | 1044.205682 | 1142.127913 | 869.3386204 | 1470.651615 | 1379.414227 | 1363.149413 | 1.10E-05 | 0.448774 | 3.45E-06 | 4.642027 | 1.36488 | 4.959815 |
| CNAG_033 | PCL103 | g1/s-speci | 128.9105447 | 163.8644098 | 103.6929387 | 271.7697097 | 240.1358101 | 219.6523766 | 1.11E-05 | 0.8757 | 3.51E-06 | 4.638529 | 1.834898 | 4.952871 |
| CNAG_012 | - | hypothetic | 113.6197712 | 109.0003843 | 87.67343544 | 214.1351147 | 195.3176128 | 174.1399166 | 1.12E-05 | 0.900672 | 3.53E-06 | 4.637506 | 1.866935 | 4.950892 |
| CNAG_049 | - | ADP-ribos | 2728.921526 | 2883.107829 | 2763.125915 | 2396.998366 | 2259.754458 | 2170.525913 | 1.14E-05 | -0.3101 | 3.58E-06 | -4.63461 | 0.806588 | 4.944977 |
| CNAG_044 | - | ubiquitin | 623.9103528 | 718.7140261 | 617.6123518 | 830.5517413 | 943.9151334 | 957.8300184 | 1.14E-05 | 0.464917 | 3.58E-06 | 4.634471 | 1.380238 | 4.944853 |
| CNAG_066 | - | COP9 sigr | 994.7013672 | 1013.743456 | 974.66528 | 689.1137288 | 745.9169327 | 807.1734938 | 1.14E-05 | -0.42672 | 3.58E-06 | -4.63416 | 0.743953 | 4.944378 |
| CNAG_035 | - | hypothetic | 78.89991767 | 98.28668715 | 138.7099001 | 197.4227625 | 216.9869915 | 207.2804863 | 1.14E-05 | 0.966384 | 3.61E-06 | 4.632717 | 1.953936 | 4.941678 |
| CNAG_032 | ARF2 | ADP-ribos | 12521.60625 | 11799.9989 | 12050.87269 | 10488.87513 | 10368.74348 | 9503.142766 | 1.14E-05 | -0.27626 | 3.61E-06 | -4.63273 | 0.825728 | 4.941678 |
| CNAG_069 | - | hypothetic | 13.42961667 | 16.41079437 | 20.44459671 | 55.99386766 | 67.17547124 | 50.81977906 | 1.15E-05 | 1.796168 | 3.64E-06 | 4.631157 | 3.472966 | 4.938572 |
| CNAG_059 | - | hypothetic | 39.0527177 | 31.53267738 | 47.08158908 | 90.03295279 | 92.66495995 | 112.6688448 | 1.16E-05 | 1.313427 | 3.65E-06 | 4.630229 | 2.485312 | 4.936794 |
| CNAG_029 | - | hypothetic | 5871.657373 | 6275.013132 | 5797.326649 | 6993.821339 | 7500.14082 | 7313.634566 | 1.20E-05 | 0.266361 | 3.79E-06 | 4.622701 | 1.20277 | 4.921183 |
| CNAG_030 | - | compass | 1096.396669 | 1150.599379 | 1186.85381 | 881.5097186 | 845.9462294 | 917.1022435 | 1.20E-05 | -0.39188 | 3.80E-06 | -4.6218 | 0.762135 | 4.919475 |
| CNAG_061 | - | N-acetylgl | 1096.294684 | 1076.26767 | 1096.225095 | 865.2315352 | 864.1438959 | 758.4651462 | 1.21E-05 | -0.40947 | 3.83E-06 | -4.62045 | 0.7529 | 4.916804 |
| CNAG_024 | - | Monosacc | 125.4789355 | 155.3041976 | 153.7057976 | 225.9628632 | 283.4678368 | 249.5343852 | 1.22E-05 | 0.791091 | 3.85E-06 | 4.619259 | 1.730382 | 4.91448 |
| CNAG_014 | - | hypothetic | 2064.427291 | 1981.359742 | 2250.085884 | 1613.082395 | 1762.985157 | 1561.070349 | 1.22E-05 | -0.36661 | 3.86E-06 | -4.61876 | 0.775605 | 4.913672 |
| CNAG_032 | MSL1 | histone ac | 1547.069461 | 1506.180651 | 1544.145817 | 1259.080452 | 1148.707852 | 1219.083139 | 1.22E-05 | -0.35758 | 3.88E-06 | -4.61793 | 0.780472 | 4.912262 |
| CNAG_006 | - | GTPase | 1805.019066 | 1665.633883 | 1591.115852 | 1325.155944 | 1277.243258 | 1361.167734 | 1.23E-05 | -0.36824 | 3.89E-06 | -4.61698 | 0.774728 | 4.910551 |
| CNAG_065 | - | nucleopor | 2061.945583 | 1910.21651 | 1832.135956 | 1522.833112 | 1567.132452 | 1523.280132 | 1.24E-05 | -0.34694 | 3.94E-06 | -4.61475 | 0.786253 | 4.906217 |
| CNAG_019 | - | large subu | 310.3725004 | 297.2926582 | 374.0673708 | 209.6999029 | 202.3986742 | 204.504275 | 1.25E-05 | -0.68928 | 3.98E-06 | -4.61248 | 0.620162 | 4.901813 |
| CNAG_012 | OCA101 | protein-ty | 68.17055392 | 71.29798568 | 57.80963621 | 151.9456283 | 135.812204 | 124.1411246 | 1.27E-05 | 1.049817 | 4.04E-06 | 4.609068 | 2.070267 | 4.895014 |
| CNAG_048 | - | vacuolar p | 24.05212977 | 24.02786894 | 52.37511335 | 131.733891 | 104.0938252 | 69.9291471 | 1.29E-05 | 1.587718 | 4.11E-06 | 4.605924 | 3.005735 | 4.888783 |
| CNAG_060 | - | mRNA exp | 2025.770318 | 1852.056779 | 1904.622728 | 1437.128779 | 1580.950438 | 1559.888342 | 1.30E-05 | -0.3529 | 4.12E-06 | -4.60501 | 0.783011 | 4.887036 |

| | | | | | | | | | | | | | | |
|---|---|---|---|---|---|---|---|---|---|---|---|---|---|---|
| CNAG_074 | - | SCF-assoc | 737.0889304 | 785.5170752 | 625.1131116 | 1066.785357 | 961.0460502 | 969.0377401 | 1.30E-05 | 0.466015 | 4.14E-06 | 4.60445 | 1.381289 | 4.88604 |
| CNAG_057 | - | GTPase in | 1610.207932 | 1776.541034 | 1615.625955 | 1342.321642 | 1366.175685 | 1221.37411 | 1.30E-05 | -0.36295 | 4.15E-06 | -4.60397 | 0.777571 | 4.88537 |
| CNAG_040 | - | hypothetic | 955.1501855 | 1063.300071 | 1097.247062 | 802.1514792 | 816.7012333 | 715.1716969 | 1.30E-05 | -0.43211 | 4.14E-06 | -4.604 | 0.741178 | 4.88537 |
| CNAG_057 | - | hypothetic | 414.1415435 | 413.5457469 | 367.856236 | 293.1759529 | 255.0294353 | 228.9962571 | 1.32E-05 | -0.63685 | 4.19E-06 | -4.60157 | 0.643116 | 4.880703 |
| CNAG_078 | - | centromer | 3163.602808 | 2970.596718 | 3183.423954 | 3721.920413 | 3957.609822 | 3877.674518 | 1.32E-05 | 0.295116 | 4.22E-06 | 4.600303 | 1.226983 | 4.878224 |
| CNAG_039 | - | hypothetic | 36.77717845 | 52.95279765 | 36.4909881 | 109.9158231 | 87.33911851 | 108.3499675 | 1.33E-05 | 1.280168 | 4.23E-06 | 4.599724 | 2.428672 | 4.877349 |
| CNAG_074 | - | AP-2 com | 832.3148953 | 793.9754283 | 790.1663782 | 591.1496033 | 587.1795641 | 625.1457806 | 1.33E-05 | -0.43695 | 4.24E-06 | -4.59918 | 0.738692 | 4.876386 |
| CNAG_056 | PPS1 | hypothetic | 1238.181656 | 1456.758492 | 1416.326046 | 1879.18 | 1672.564052 | 1883.747759 | 1.35E-05 | 0.388438 | 4.29E-06 | 4.596637 | 1.308976 | 4.871248 |
| CNAG_045 | - | translocati | 2358.633002 | 2173.147357 | 2322.657636 | 1817.811214 | 1849.180296 | 1869.621461 | 1.35E-05 | -0.32377 | 4.31E-06 | -4.59605 | 0.798978 | 4.87019 |
| CNAG_122 | - | unspecifie | 336.5716505 | 325.5231649 | 297.6992494 | 519.758485 | 440.0266417 | 480.4931768 | 1.35E-05 | 0.571875 | 4.31E-06 | 4.595733 | 1.486454 | 4.869698 |
| CNAG_064 | APT1 | Phospholi | 2818.677058 | 2845.362376 | 2406.99086 | 2211.70706 | 2080.305948 | 2080.594912 | 1.38E-05 | -0.35585 | 4.42E-06 | -4.59058 | 0.781411 | 4.859145 |
| CNAG_066 | - | hypothetic | 1026.170058 | 943.7737066 | 1000.240846 | 748.3998582 | 782.1970449 | 727.3362872 | 1.43E-05 | -0.41126 | 4.58E-06 | -4.58324 | 0.751964 | 4.84454 |
| CNAG_078 | - | hypothetic | 173.2708754 | 218.8032903 | 165.492947 | 297.7294513 | 318.6620243 | 302.8430058 | 1.44E-05 | 0.707534 | 4.62E-06 | 4.581392 | 1.633011 | 4.841029 |
| CNAG_124 | - | unspecifie | 61.61171565 | 100.6578725 | 61.74157666 | 29.74781458 | 20.19094993 | 25.87264804 | 1.44E-05 | -1.5975 | 4.63E-06 | -4.58096 | 0.330448 | 4.84029 |
| CNAG_015 | - | hypothetic | 51.02110865 | 105.7814993 | 124.8735302 | 211.7618686 | 209.239868 | 191.7828147 | 1.45E-05 | 1.112061 | 4.65E-06 | 4.580042 | 2.161542 | 4.838556 |
| CNAG_048 | - | peptidyl-p | 1197.121199 | 1214.337752 | 1014.350263 | 1464.168904 | 1532.107017 | 1556.266168 | 1.46E-05 | 0.395417 | 4.68E-06 | 4.578494 | 1.315322 | 4.835837 |
| CNAG_013 | SDP101 | hypothetic | 96.14907173 | 82.12669225 | 96.13723254 | 156.7689067 | 185.0671985 | 181.7116753 | 1.47E-05 | 0.914005 | 4.72E-06 | 4.577049 | 1.884269 | 4.833004 |
| CNAG_128 | - | unspecifie | 18.14851264 | 29.30076648 | 27.91847249 | 73.1670347 | 60.59167155 | 87.0726107 | 1.48E-05 | 1.558336 | 4.76E-06 | 4.575013 | 2.94514 | 4.82911 |
| CNAG_028 | - | hypothetic | 83.59402715 | 131.6052006 | 137.6893051 | 249.2199981 | 202.0562085 | 220.6358711 | 1.48E-05 | 0.920994 | 4.76E-06 | 4.574911 | 1.89342 | 4.829063 |
| CNAG_064 | - | Adenosine | 335.2541864 | 241.2149176 | 211.1385422 | 155.9719154 | 152.3659551 | 122.487747 | 1.49E-05 | -0.892 | 4.79E-06 | -4.57372 | 0.538866 | 4.826758 |
| CNAG_126 | - | unspecifie | 2.823352522 | 8.806821958 | 4.461829755 | 36.01581209 | 39.57089628 | 24.30677055 | 1.51E-05 | 2.808981 | 4.85E-06 | 4.571336 | 7.007892 | 4.82198 |
| CNAG_034 | TSA1 | Thiol pero | 11796.45518 | 10819.56702 | 12098.82376 | 9784.478814 | 9450.061724 | 9670.716723 | 1.51E-05 | -0.27991 | 4.86E-06 | -4.57067 | 0.823642 | 4.82076 |
| CNAG_039 | - | hypothetic | 0.330020538 | 1.2686367 | 3.365108902 | 40.90474151 | 14.65465027 | 24.11163913 | 1.53E-05 | 4.271318 | 4.92E-06 | 4.568364 | 19.31056 | 4.816151 |
| CNAG_074 | - | hypothetic | 103.3044207 | 132.6715548 | 111.1040295 | 223.2441679 | 171.6597363 | 269.1505568 | 1.54E-05 | 0.924572 | 4.96E-06 | 4.566301 | 1.898121 | 4.812206 |
| CNAG_067 | - | fructosyl a | 102.2031274 | 157.4044762 | 126.0196558 | 235.6327412 | 210.6490651 | 261.5541185 | 1.54E-05 | 0.861412 | 4.97E-06 | 4.565906 | 1.816816 | 4.811554 |
| CNAG_002 | PTP4 | serine/thre | 702.0532271 | 657.3804964 | 710.3445303 | 937.374297 | 885.9565874 | 996.6873433 | 1.56E-05 | 0.43036 | 5.03E-06 | 4.563402 | 1.34757 | 4.806699 |
| CNAG_065 | - | clathrin-cc | 2478.765145 | 2235.596592 | 2089.165713 | 1740.720496 | 1721.464529 | 1865.162667 | 1.56E-05 | -0.36838 | 5.03E-06 | -4.56345 | 0.774652 | 4.806699 |
| CNAG_058 | KRE64 | glucosidas | 138.252113 | 171.419067 | 135.6279185 | 247.1762431 | 270.4207463 | 242.9217187 | 1.56E-05 | 0.75904 | 5.04E-06 | 4.563161 | 1.692365 | 4.806364 |
| CNAG_129 | - | unspecifie | 197.7111615 | 209.1599167 | 183.6030083 | 296.0370094 | 301.2380537 | 362.5995678 | 1.59E-05 | 0.688557 | 5.14E-06 | 4.559142 | 1.611671 | 4.798545 |
| CNAG_069 | - | arsenical-r | 1288.999697 | 1272.367365 | 1282.868281 | 998.2581151 | 1053.933999 | 966.0119058 | 1.64E-05 | -0.36415 | 5.30E-06 | -4.55264 | 0.776926 | 4.785273 |
| CNAG_042 | - | hypothetic | 146.4926883 | 172.5616821 | 186.7430778 | 246.0754205 | 288.424263 | 332.5734947 | 1.65E-05 | 0.766224 | 5.32E-06 | 4.551563 | 1.700813 | 4.783214 |
| CNAG_032 | - | hypothetic | 21.66320989 | 38.95882596 | 29.00973988 | 83.31317025 | 67.47113767 | 97.07453242 | 1.66E-05 | 1.475957 | 5.38E-06 | 4.549248 | 2.78168 | 4.778598 |
| CNAG_042 | MPK2 | CMGC/MA | 238.6522189 | 304.977871 | 232.6675685 | 384.0147838 | 424.7454383 | 404.9828231 | 1.67E-05 | 0.632746 | 5.39E-06 | 4.549163 | 1.550514 | 4.778586 |
| CNAG_027 | - | hypothetic | 845.0786893 | 722.9234502 | 826.3717731 | 549.0406093 | 633.8703813 | 541.9829279 | 1.68E-05 | -0.49008 | 5.43E-06 | -4.54764 | 0.711986 | 4.775611 |
| CNAG_034 | - | hypothetic | 1652.510797 | 1596.843035 | 1480.353189 | 2051.098139 | 2022.871426 | 1952.684392 | 1.71E-05 | 0.334413 | 5.54E-06 | 4.543317 | 1.260865 | 4.767023 |
| CNAG_006 | - | transcripti | 7307.584923 | 6759.781919 | 7053.467002 | 5696.032165 | 5899.788976 | 6063.775618 | 1.75E-05 | -0.27382 | 5.66E-06 | -4.5388 | 0.827128 | 4.758048 |
| CNAG_022 | - | phosphok | 2491.073691 | 2862.7634 | 2278.090894 | 3565.815246 | 3190.721505 | 3226.027808 | 1.75E-05 | 0.372989 | 5.68E-06 | 4.538024 | 1.295033 | 4.75661 |
| CNAG_000 | FPP1 | fructose-1 | 2607.818863 | 2102.293438 | 1651.141922 | 3966.104233 | 3949.239587 | 3755.401967 | 1.78E-05 | 0.859686 | 5.77E-06 | 4.534553 | 1.814643 | 4.749628 |
| CNAG_038 | - | lysine dec | 533.9688664 | 541.0050354 | 577.033046 | 799.7682881 | 714.3071748 | 773.5970348 | 1.78E-05 | 0.454357 | 5.79E-06 | 4.533905 | 1.370172 | 4.748458 |
| CNAG_075 | - | proteasom | 825.798761 | 830.7879566 | 747.7131997 | 1043.15051 | 1060.262754 | 1090.002098 | 1.85E-05 | 0.395602 | 5.99E-06 | 4.526679 | 1.315491 | 4.733932 |
| CNAG_001 | - | hypothetic | 500.8832353 | 579.4923002 | 583.2019117 | 402.8307285 | 399.155681 | 371.0197549 | 1.85E-05 | -0.51878 | 6.02E-06 | -4.52562 | 0.697961 | 4.731917 |
| CNAG_063 | - | hypothetic | 127.9319896 | 182.2491738 | 208.0367242 | 292.3347425 | 297.0423883 | 312.7368712 | 1.87E-05 | 0.787035 | 6.07E-06 | 4.523916 | 1.725525 | 4.728583 |
| CNAG_005 | - | hypothetic | 769.3808654 | 826.2488732 | 782.6974346 | 560.733192 | 603.6135024 | 616.2642234 | 1.91E-05 | -0.43301 | 6.22E-06 | -4.51866 | 0.740715 | 4.717957 |
| CNAG_028 | - | hypothetic | 50.7481719 | 64.81068052 | 51.39968869 | 115.7560905 | 136.0590443 | 109.6932177 | 1.92E-05 | 1.110885 | 6.26E-06 | 4.517557 | 2.159782 | 4.715863 |
| CNAG_056 | - | hypothetic | 3436.439397 | 3265.625702 | 3104.420558 | 2779.897696 | 2673.795895 | 2540.210144 | 1.93E-05 | -0.31039 | 6.27E-06 | -4.51709 | 0.806421 | 4.715108 |
| CNAG_023 | - | cyclin binc | 1910.6157 | 1860.822966 | 1896.226997 | 2338.975809 | 2346.303271 | 2354.537272 | 1.93E-05 | 0.297481 | 6.28E-06 | 4.516836 | 1.228997 | 4.714874 |
| CNAG_006 | - | hypothetic | 205.7639856 | 189.7823668 | 150.5880142 | 289.9197633 | 295.2704865 | 321.609336 | 1.93E-05 | 0.719122 | 6.30E-06 | 4.516037 | 1.64618 | 4.713399 |
| CNAG_019 | - | endopepti | 1573.208444 | 1666.864543 | 1605.099144 | 1906.874016 | 2176.485844 | 2103.604589 | 1.96E-05 | 0.337489 | 6.40E-06 | 4.512723 | 1.263556 | 4.706772 |
| CNAG_120 | - | unspecifie | 1.584641371 | 1.274291761 | 5.48867537 | 36.54193875 | 20.91549179 | 24.14834283 | 1.97E-05 | 3.499013 | 6.42E-06 | 4.512218 | 11.30597 | 4.705899 |
| CNAG_013 | - | hypothetic | 1272.768254 | 1361.779142 | 1411.874011 | 1017.659219 | 1076.325206 | 1080.304163 | 1.98E-05 | -0.365 | 6.47E-06 | -4.51048 | 0.776467 | 4.702665 |
| CNAG_129 | - | unspecifie | 269818.0282 | 236555.5946 | 313610.5491 | 181750.6765 | 130649.8996 | 147604.0059 | 1.99E-05 | -0.84981 | 6.51E-06 | -4.50916 | 0.554857 | 4.700289 |
| CNAG_069 | - | tRNA-dihy | 221.5315762 | 211.0614485 | 267.3786075 | 108.2116564 | 111.6891899 | 163.2936636 | 2.00E-05 | -0.88698 | 6.52E-06 | -4.50894 | 0.540743 | 4.699998 |
| CNAG_048 | - | hypothetic | 43.90732908 | 88.44909653 | 61.00118281 | 150.2982003 | 144.5119351 | 132.9618474 | 2.00E-05 | 1.136898 | 6.54E-06 | 4.508251 | 2.199077 | 4.698752 |
| CNAG_054 | - | hypothetic | 1.665705843 | 9.897316215 | 8.71519522 | 42.39875795 | 38.51036054 | 30.86944137 | 2.03E-05 | 2.598731 | 6.63E-06 | 4.505115 | 6.057534 | 4.692499 |
| CNAG_043 | - | cation diff | 10.08310355 | 43.23455031 | 27.95185477 | 83.29053775 | 92.09761812 | 82.88315991 | 2.05E-05 | 1.672346 | 6.69E-06 | 4.503264 | 3.187325 | 4.688876 |
| CNAG_056 | UBP5 | ubiquitin | 11178.66762 | 10755.93893 | 10099.29938 | 8543.774187 | 9182.727099 | 9101.187432 | 2.05E-05 | -0.27126 | 6.71E-06 | -4.50276 | 0.828597 | 4.688013 |
| CNAG_043 | - | U3 small r | 2141.29827 | 2027.614023 | 1761.784722 | 1518.915407 | 1609.415443 | 1483.348368 | 2.08E-05 | -0.37844 | 6.81E-06 | -4.49968 | 0.769267 | 4.682029 |
| CNAG_013 | - | xeroderma | 1633.823132 | 1563.447366 | 1461.156629 | 1914.758911 | 2068.645972 | 1967.090443 | 2.08E-05 | 0.337828 | 6.81E-06 | 4.499557 | 1.263853 | 4.681948 |
| CNAG_079 | - | WD repea | 796.6390095 | 790.9469567 | 778.6148648 | 1149.287971 | 1006.720679 | 1014.588185 | 2.11E-05 | 0.407612 | 6.93E-06 | 4.495933 | 1.326488 | 4.674873 |
| CNAG_014 | - | hypothetic | 1555.046038 | 1283.208919 | 1432.156039 | 1150.901075 | 1044.334012 | 1041.515433 | 2.13E-05 | -0.41653 | 6.99E-06 | -4.49405 | 0.749226 | 4.671358 |
| CNAG_037 | - | hypothetic | 1199.085582 | 1108.653418 | 1326.508423 | 881.4219261 | 929.6621451 | 947.0935863 | 2.15E-05 | -0.4138 | 7.04E-06 | -4.49244 | 0.750646 | 4.668222 |
| CNAG_040 | - | cytoplasm | 1701.19184 | 1706.547111 | 1582.581978 | 1257.366445 | 1405.886448 | 1294.596259 | 2.16E-05 | -0.34999 | 7.10E-06 | -4.49075 | 0.784589 | 4.664936 |
| CNAG_041 | - | hypothetic | 1341.522635 | 1312.228634 | 1292.480656 | 1069.940177 | 977.9143369 | 1053.660925 | 2.18E-05 | -0.36341 | 7.17E-06 | -4.48866 | 0.777325 | 4.660851 |
| CNAG_011 | - | hypothetic | 3155.339237 | 2856.374948 | 2505.251711 | 3547.615771 | 3911.11398 | 3656.749127 | 2.20E-05 | 0.368495 | 7.21E-06 | 4.487313 | 1.291005 | 4.658263 |
| CNAG_036 | - | hypothetic | 1481.969543 | 1323.20433 | 983.4974246 | 2061.635956 | 1826.271125 | 1659.627266 | 2.24E-05 | 0.534937 | 7.35E-06 | 4.48327 | 1.448878 | 4.650188 |
| CNAG_060 | - | protein DO | 204.6944111 | 202.7183714 | 200.6359185 | 308.5613542 | 301.2181426 | 352.670268 | 2.26E-05 | 0.650154 | 7.42E-06 | 4.481366 | 1.569336 | 4.646795 |
| CNAG_052 | - | bud site s | 1670.895702 | 1713.022848 | 1688.125544 | 1382.035845 | 1412.772942 | 1226.935908 | 2.27E-05 | -0.35038 | 7.46E-06 | -4.48002 | 0.784378 | 4.644217 |
| CNAG_011 | - | cation-trai | 3534.535516 | 3209.6396 | 3335.81132 | 2674.695728 | 2796.29644 | 2803.255549 | 2.32E-05 | -0.30041 | 7.65E-06 | -4.47485 | 0.812022 | 4.634191 |

| | | | | | | | | | | | | | | |
|---|---|---|---|---|---|---|---|---|---|---|---|---|---|---|
| CNAG_052 | - | tRNA pseu | 2088.84528 | 1859.627172 | 2116.794884 | 1433.947703 | 1579.178579 | 1691.956209 | 2.35E-05 | -0.38224 | 7.74E-06 | -4.47229 | 0.767244 | 4.629309 |
| CNAG_060 | - | vesicle-fus | 3741.334634 | 3779.557957 | 3648.298557 | 3017.937346 | 3199.139825 | 3146.292182 | 2.36E-05 | -0.26969 | 7.77E-06 | -4.47149 | 0.829497 | 4.627836 |
| CNAG_046 | - | prolyl olig | 23.75555649 | 16.41726701 | 15.15875363 | 55.11564093 | 57.37906247 | 63.89935052 | 2.37E-05 | 1.680367 | 7.81E-06 | 4.470255 | 3.205094 | 4.625497 |
| CNAG_130 | - | unspecifie | 60.09493425 | 77.73205653 | 66.30785912 | 139.9982189 | 129.6818769 | 142.8474665 | 2.38E-05 | 0.999866 | 7.87E-06 | 4.468804 | 1.999814 | 4.62271 |
| CNAG_065 | - | hypothetic | 49.47985136 | 47.62089259 | 42.86797032 | 93.01560484 | 109.1823699 | 110.602467 | 2.45E-05 | 1.161599 | 8.07E-06 | 4.463298 | 2.237052 | 4.611699 |
| CNAG_123 | - | unspecifie | 14.63387329 | 28.18773556 | 20.47433298 | 62.99394697 | 64.31170241 | 66.19380905 | 2.46E-05 | 1.617505 | 8.12E-06 | 4.461946 | 3.068438 | 4.609281 |
| CNAG_035 | - | zinc finger | 416.2777306 | 314.5862596 | 374.1769867 | 252.5966176 | 223.9388919 | 231.1525131 | 2.48E-05 | -0.66131 | 8.20E-06 | -4.46002 | 0.632302 | 4.605541 |
| CNAG_043 | HSV2 | WD40 dor | 588.9079339 | 636.8852929 | 698.58216 | 854.0450489 | 878.2931839 | 889.0581964 | 2.49E-05 | 0.432419 | 8.21E-06 | 4.459512 | 1.349494 | 4.604669 |
| CNAG_015 | - | hypothetic | 46.15436492 | 58.39998359 | 68.39011145 | 107.7088374 | 143.8603488 | 125.1166844 | 2.54E-05 | 1.10941 | 8.41E-06 | 4.454413 | 2.157574 | 4.594504 |
| CNAG_031 | SET202 | histone-ly | 2326.123578 | 2639.581968 | 2494.369074 | 1947.751325 | 2050.175033 | 2070.547028 | 2.65E-05 | -0.31271 | 8.78E-06 | -4.44523 | 0.805129 | 4.576567 |
| CNAG_070 | RPN11 | 26S protea | 4307.981819 | 3763.622612 | 4350.114624 | 5087.859614 | 5146.165914 | 5419.638278 | 2.72E-05 | 0.317789 | 9.00E-06 | 4.439826 | 1.246419 | 4.565824 |
| CNAG_051 | - | transcripti | 900.246256 | 1006.182129 | 973.5781398 | 772.6383098 | 726.9434316 | 699.5904747 | 2.76E-05 | -0.40487 | 9.14E-06 | -4.43648 | 0.755306 | 4.559398 |
| CNAG_075 | - | hypothetic | 269.0882392 | 305.0894895 | 364.7910453 | 533.8286677 | 529.1669023 | 412.9626882 | 2.79E-05 | 0.636734 | 9.26E-06 | 4.433665 | 1.554805 | 4.553878 |
| CNAG_014 | - | PH and SE | 1021.979174 | 1010.729201 | 992.9613786 | 1310.927383 | 1273.272068 | 1304.303489 | 2.81E-05 | 0.347462 | 9.33E-06 | 4.432225 | 1.27232 | 4.551137 |
| CNAG_032 | - | AMME syr | 44.98037333 | 74.44909305 | 42.90845934 | 110.9625332 | 143.9210005 | 115.1933981 | 2.82E-05 | 1.188215 | 9.35E-06 | 4.431668 | 2.278707 | 4.550334 |
| CNAG_043 | - | hypothetic | 873.5396313 | 712.3400259 | 554.7891763 | 1206.32686 | 1085.375826 | 963.5518185 | 2.83E-05 | 0.589357 | 9.39E-06 | 4.430816 | 1.504576 | 4.548779 |
| CNAG_056 | - | hypothetic | 807.9165034 | 797.2337606 | 893.5417596 | 658.9934368 | 613.065412 | 596.351812 | 2.85E-05 | -0.43489 | 9.47E-06 | -4.42894 | 0.739748 | 4.545312 |
| CNAG_007 | SPS1 | STE/STE20 | 557.4307554 | 701.4646472 | 613.3308212 | 890.7549542 | 793.6195827 | 953.3398329 | 2.89E-05 | 0.480524 | 9.62E-06 | 4.42561 | 1.39525 | 4.538776 |
| CNAG_055 | - | hypothetic | 2300.240261 | 2159.123158 | 2176.58137 | 1758.651355 | 1821.5933 | 1836.30731 | 2.90E-05 | -0.30863 | 9.64E-06 | -4.42521 | 0.807409 | 4.538124 |
| CNAG_031 | - | 4'-phosph | 417.6633006 | 423.2367832 | 363.6109661 | 293.9277032 | 240.2991615 | 268.8632624 | 2.90E-05 | -0.60161 | 9.66E-06 | -4.42469 | 0.659018 | 4.537238 |
| CNAG_001 | VPS29 | retrograde | 1177.957834 | 1117.20553 | 1063.211595 | 917.4274096 | 847.6972822 | 821.7059494 | 2.92E-05 | -0.39239 | 9.71E-06 | -4.42354 | 0.761867 | 4.535088 |
| CNAG_066 | - | allantoate | 91.76626278 | 174.589698 | 119.6436333 | 234.0181145 | 246.2469308 | 242.8128987 | 2.94E-05 | 0.898143 | 9.79E-06 | 4.42186 | 1.863666 | 4.531871 |
| CNAG_003 | - | anon-23da | 5.076755615 | 2.360662049 | 3.37470178 | 30.5823758 | 24.49656521 | 27.3870622 | 2.95E-05 | 3.001443 | 9.82E-06 | 4.421042 | 8.008006 | 4.530386 |
| CNAG_010 | - | superkiller | 5870.042223 | 5558.408261 | 5258.664896 | 4407.151987 | 4691.353211 | 4743.800352 | 2.98E-05 | -0.2851 | 9.93E-06 | -4.4186 | 0.820682 | 4.525638 |
| CNAG_121 | - | unspecifie | 23.80742193 | 22.84570259 | 17.2924818 | 54.13159622 | 65.22880961 | 68.34569658 | 2.99E-05 | 1.577553 | 9.98E-06 | 4.417673 | 2.984632 | 4.523936 |
| CNAG_043 | - | thioredoxi | 1173.324634 | 1062.311726 | 1161.270334 | 865.9253692 | 881.3638532 | 889.3697503 | 3.00E-05 | -0.38172 | 1.00E-05 | -4.4169 | 0.767522 | 4.522541 |
| CNAG_027 | - | hypothetic | 564.3342124 | 573.3282088 | 628.2081194 | 777.7506588 | 790.2786902 | 841.2817461 | 3.01E-05 | 0.432949 | 1.00E-05 | 4.41614 | 1.34999 | 4.521174 |
| CNAG_010 | - | COP9 sigr | 2278.14889 | 2336.866197 | 2277.890766 | 1794.415916 | 1925.981416 | 1937.319335 | 3.02E-05 | -0.30007 | 1.01E-05 | -4.41557 | 0.812212 | 4.520196 |
| CNAG_065 | - | endoribor | 533.4115399 | 547.1962655 | 522.4783086 | 384.9027368 | 365.4568807 | 402.0150799 | 3.07E-05 | -0.49197 | 1.02E-05 | -4.41201 | 0.711053 | 4.513522 |
| CNAG_017 | - | ribosome | 2990.530027 | 3109.398522 | 3125.709603 | 2574.373228 | 2584.969648 | 2573.458377 | 3.12E-05 | -0.26985 | 1.04E-05 | -4.40823 | 0.829404 | 4.506103 |
| CNAG_061 | - | hypothetic | 509.9915137 | 537.4419154 | 454.2700333 | 349.0703576 | 374.1879959 | 323.3041045 | 3.13E-05 | -0.53512 | 1.05E-05 | -4.4073 | 0.690103 | 4.504391 |
| CNAG_022 | - | 26S protea | 6794.116375 | 6497.009606 | 6721.915134 | 7848.417778 | 8207.423275 | 7915.368986 | 3.15E-05 | 0.244827 | 1.05E-05 | 4.405783 | 1.18495 | 4.50167 |
| CNAG_058 | - | hypothetic | 592.8520732 | 537.558847 | 554.463985 | 418.427279 | 412.1064235 | 378.7905772 | 3.18E-05 | -0.4941 | 1.07E-05 | -4.40333 | 0.710005 | 4.497065 |
| CNAG_027 | PMU101 | hypothetic | 1006.280601 | 898.5307513 | 972.5053481 | 775.7771233 | 657.8459928 | 721.7593325 | 3.20E-05 | -0.43329 | 1.07E-05 | -4.40219 | 0.740569 | 4.494941 |
| CNAG_056 | - | hypothetic | 57.91968925 | 101.4220408 | 88.68871845 | 222.53417 | 161.5980005 | 139.8101952 | 3.25E-05 | 1.071979 | 1.09E-05 | 4.398621 | 2.102316 | 4.488121 |
| CNAG_067 | CSR3 | chitin synt | 62.23375364 | 54.08243136 | 40.77148093 | 110.3097338 | 108.1414604 | 127.1933391 | 3.29E-05 | 1.129847 | 1.10E-05 | 4.395948 | 2.188355 | 4.483091 |
| CNAG_025 | - | hypothetic | 211.7960343 | 240.4047043 | 249.6498895 | 346.6848862 | 361.6959715 | 365.0049299 | 3.29E-05 | 0.599977 | 1.10E-05 | 4.395742 | 1.515693 | 4.482837 |
| CNAG_056 | - | ABC trans | 1283.519708 | 1199.299542 | 1224.37813 | 1651.895734 | 1600.224461 | 1504.205001 | 3.31E-05 | 0.344023 | 1.11E-05 | 4.394517 | 1.269291 | 4.480546 |
| CNAG_001 | - | pre-mRNA | 2326.454919 | 2533.125427 | 2644.849253 | 3005.888619 | 3283.127645 | 3097.24764 | 3.33E-05 | 0.307787 | 1.12E-05 | 4.393155 | 1.237807 | 4.477981 |
| CNAG_043 | - | hypothetic | 311.0531075 | 363.2125402 | 375.4756672 | 504.6895331 | 490.9748668 | 534.8428624 | 3.35E-05 | 0.527934 | 1.12E-05 | 4.39188 | 1.441863 | 4.475592 |
| CNAG_067 | - | hypothetic | 97.53140595 | 146.6447856 | 133.4457182 | 231.8159141 | 222.8830854 | 214.0416604 | 3.38E-05 | 0.81296 | 1.14E-05 | 4.389375 | 1.756812 | 4.470746 |
| CNAG_017 | - | RTA1 dom | 1472.67909 | 1340.452301 | 1385.414065 | 1910.341226 | 1794.282633 | 1697.379038 | 3.43E-05 | 0.34821 | 1.15E-05 | 4.386269 | 1.27298 | 4.46486 |
| CNAG_021 | - | hypothetic | 18.14133364 | 33.56568922 | 14.13348066 | 44.19613563 | 72.07782965 | 109.9288419 | 3.47E-05 | 1.77939 | 1.17E-05 | 4.383586 | 3.43281 | 4.459665 |
| CNAG_125 | - | unspecifie | 32.15723432 | 60.43230773 | 29.05921268 | 92.35273591 | 102.2632014 | 110.5249854 | 3.51E-05 | 1.316545 | 1.18E-05 | 4.38096 | 2.49069 | 4.454587 |
| CNAG_129 | - | unspecifie | 2.80353452 | 3.453425681 | 8.679835999 | 33.67251809 | 27.9673629 | 30.71207037 | 3.56E-05 | 2.776139 | 1.20E-05 | 4.377949 | 6.850165 | 4.448744 |
| CNAG_003 | - | glucan 1,3 | 144.018401 | 164.9563161 | 121.7872572 | 245.6947434 | 265.2272604 | 223.0022171 | 3.56E-05 | 0.758096 | 1.20E-05 | 4.377796 | 1.691257 | 4.448596 |
| CNAG_050 | - | ubiquinol- | 1195.590747 | 1150.644127 | 1291.339559 | 1021.785601 | 856.2561194 | 886.0943423 | 3.62E-05 | -0.41153 | 1.22E-05 | -4.37402 | 0.751823 | 4.441241 |
| CNAG_064 | - | RNA exon | 3089.625046 | 2935.934858 | 2789.819875 | 2414.804682 | 2364.112577 | 2472.420153 | 3.63E-05 | -0.29697 | 1.22E-05 | -4.37362 | 0.813958 | 4.4406 |
| CNAG_046 | MRPL37 | large subu | 1556.71982 | 1245.696066 | 1431.270488 | 1964.03573 | 1961.715543 | 1738.462665 | 3.67E-05 | 0.40384 | 1.24E-05 | 4.370725 | 1.323024 | 4.434993 |
| CNAG_010 | - | hypothetic | 674.0188751 | 714.4045538 | 571.7865062 | 850.8581414 | 956.0227026 | 899.0690125 | 3.70E-05 | 0.45061 | 1.25E-05 | 4.368901 | 1.366618 | 4.431522 |
| CNAG_078 | - | hypothetic | 12.37793864 | 43.18421564 | 13.07626909 | 73.10908477 | 72.89926824 | 88.19528357 | 3.75E-05 | 1.766888 | 1.27E-05 | 4.365787 | 3.403192 | 4.425491 |
| CNAG_017 | - | mitochonc | 731.9451712 | 687.3473897 | 784.7600789 | 525.7456962 | 589.8850075 | 497.5831271 | 3.77E-05 | -0.46669 | 1.27E-05 | -4.36452 | 0.723622 | 4.423293 |
| CNAG_052 | APT4 | Aminopho | 2132.556753 | 2313.305265 | 2214.032269 | 2629.177054 | 2768.969328 | 2834.092591 | 3.83E-05 | 0.290879 | 1.30E-05 | 4.360739 | 1.223386 | 4.41625 |
| CNAG_029 | - | membrane | 151.1642043 | 172.576334 | 208.0346948 | 284.4900369 | 326.5479353 | 271.8420369 | 3.83E-05 | 0.715091 | 1.30E-05 | 4.360741 | 1.641587 | 4.41625 |
| CNAG_029 | - | hydrolase | 860.3547185 | 773.55993 | 878.6301559 | 623.0805398 | 597.5067826 | 659.5485632 | 3.91E-05 | -0.43393 | 1.32E-05 | -4.35627 | 0.740245 | 4.40755 |
| CNAG_129 | - | unspecifie | 3.914548174 | 4.498988975 | 0.139356905 | 34.04645702 | 30.87085025 | 14.46279645 | 3.94E-05 | 3.463635 | 1.33E-05 | 4.354731 | 11.03209 | 4.404646 |
| CNAG_074 | - | hypothetic | 1121.908109 | 1062.258944 | 1061.045221 | 862.1411128 | 871.935997 | 688.5844767 | 3.98E-05 | -0.43788 | 1.35E-05 | -4.35202 | 0.738216 | 4.399735 |
| CNAG_062 | - | hypothetic | 145.3065647 | 171.4728973 | 187.7930642 | 251.6418255 | 286.7381675 | 290.550179 | 3.99E-05 | 0.702438 | 1.35E-05 | 4.351806 | 1.627252 | 4.399475 |
| CNAG_079 | - | hypothetic | 1306.890785 | 1365.158573 | 1070.887365 | 1699.369393 | 1659.753065 | 1611.842258 | 3.99E-05 | 0.394723 | 1.35E-05 | 4.351529 | 1.31469 | 4.399082 |
| CNAG_007 | - | hypothetic | 1301.015887 | 1208.991374 | 1162.55063 | 1505.343033 | 1733.193339 | 1550.794862 | 4.00E-05 | 0.367584 | 1.36E-05 | 4.350781 | 1.290191 | 4.397757 |
| CNAG_127 | - | unspecifie | 185.1168355 | 157.1971342 | 176.848953 | 101.4824473 | 90.18838229 | 99.05230736 | 4.00E-05 | -0.85267 | 1.36E-05 | -4.35051 | 0.55376 | 4.397409 |
| CNAG_001 | CRG1 | Putative re | 70.67392623 | 104.6470825 | 97.19356809 | 167.0852939 | 164.1866429 | 181.6903473 | 4.05E-05 | 0.903888 | 1.37E-05 | 4.347941 | 1.871102 | 4.392759 |
| CNAG_061 | - | nuclear DI | 3453.87127 | 3098.638046 | 2968.987096 | 2445.837101 | 2510.740449 | 2696.652759 | 4.13E-05 | -0.33071 | 1.40E-05 | -4.34356 | 0.795143 | 4.384403 |
| CNAG_030 | - | Maltose p | 2.852062506 | 8.830399826 | 10.82586012 | 34.94930954 | 34.93426483 | 39.49479158 | 4.22E-05 | 2.406086 | 1.43E-05 | 4.338537 | 5.300344 | 4.374942 |
| CNAG_063 | - | hypothetic | 4668.335951 | 4429.21883 | 3799.791552 | 3387.609163 | 3407.806756 | 3554.833542 | 4.25E-05 | -0.3328 | 1.45E-05 | -4.33656 | 0.793995 | 4.371372 |
| CNAG_124 | - | unspecifie | 39.07427553 | 41.1788129 | 50.27200587 | 89.15659075 | 103.1411179 | 103.951627 | 4.30E-05 | 1.166876 | 1.46E-05 | 4.333983 | 2.245251 | 4.366727 |
| CNAG_131 | - | unspecifie | 88.31217315 | 41.97549859 | 48.93569091 | 18.77468013 | 13.29764189 | 20.28921634 | 4.31E-05 | -1.81297 | 1.47E-05 | -4.33351 | 0.284605 | 4.365947 |
| CNAG_052 | - | hypothetic | 77.54676141 | 98.16570207 | 59.97555769 | 191.119863 | 140.8036831 | 146.3315994 | 4.31E-05 | 1.013708 | 1.47E-05 | 4.33332 | 2.019093 | 4.36573 |

| | | | | | | | | | | | | | |
|---|---|---|---|---|---|---|---|---|---|---|---|---|---|
| CNAG_022 | - | copper ch | 1092.94258 | 1196.915317 | 1208.188071 | 879.906774 | 967.6921154 | 882.7436892 | 4.38E-05 | -0.37294 | 1.49E-05 | -4.32958 | 0.772208 | 4.358653 |
| CNAG_129 | - | unspecifie | 74.11707137 | 97.11940295 | 90.80329714 | 162.451601 | 174.7142084 | 152.999641 | 4.39E-05 | 0.88824 | 1.50E-05 | 4.329048 | 1.850916 | 4.357769 |
| CNAG_033 | - | hypothetic | 528.8207219 | 533.2428952 | 600.2446806 | 414.5466482 | 408.66375 | 364.3757126 | 4.39E-05 | -0.5014 | 1.50E-05 | -4.32896 | 0.706419 | 4.357748 |
| CNAG_036 | - | ATP-deper | 2759.189958 | 2623.495669 | 2626.628927 | 2144.691219 | 2195.936287 | 2281.459383 | 4.39E-05 | -0.29001 | 1.50E-05 | -4.32862 | 0.817896 | 4.357244 |
| CNAG_004 | - | hypothetic | 12.24851892 | 12.11951384 | 21.48228394 | 69.250168 | 47.76780082 | 44.22617499 | 4.43E-05 | 1.814477 | 1.51E-05 | 4.326724 | 3.517322 | 4.353653 |
| CNAG_016 | CSF1 | Cold sensi | 5820.218467 | 5788.036187 | 5495.523014 | 6504.99943 | 7068.843268 | 7189.256568 | 4.50E-05 | 0.264409 | 1.54E-05 | 4.323185 | 1.201144 | 4.346836 |
| CNAG_021 | - | origin recc | 428.8146498 | 407.4140527 | 427.7480377 | 586.3850417 | 618.7514576 | 560.5228924 | 4.58E-05 | 0.467577 | 1.57E-05 | 4.319181 | 1.382785 | 4.339111 |
| CNAG_078 | - | MIPC synt | 610.3035679 | 584.8884977 | 496.9525258 | 422.3373944 | 389.6237608 | 386.5451978 | 4.59E-05 | -0.5128 | 1.57E-05 | -4.31835 | 0.700861 | 4.337784 |
| CNAG_023 | - | hypothetic | 418.9553727 | 412.5466606 | 434.9770697 | 318.0784999 | 284.3799295 | 273.3407064 | 4.62E-05 | -0.54598 | 1.58E-05 | -4.31708 | 0.684924 | 4.335443 |
| CNAG_003 | - | hypothetic | 239.5773619 | 209.1839826 | 167.6539619 | 300.6345691 | 337.5777286 | 368.1959618 | 4.64E-05 | 0.693123 | 1.59E-05 | 4.315965 | 1.616779 | 4.333404 |
| CNAG_014 | - | eclair-PA | 2239.600395 | 2234.528349 | 2232.030667 | 1888.72964 | 1797.406691 | 1861.843711 | 4.64E-05 | -0.28898 | 1.59E-05 | -4.31582 | 0.818479 | 4.333276 |
| CNAG_034 | - | lipoyl(octa | 532.2757898 | 527.8479244 | 558.6968323 | 400.5111277 | 393.1171617 | 383.1949327 | 4.70E-05 | -0.47478 | 1.61E-05 | -4.31291 | 0.719576 | 4.327865 |
| CNAG_026 | - | hypothetic | 2364.457435 | 2258.228213 | 2152.07734 | 1847.425336 | 1837.965482 | 1871.836098 | 4.73E-05 | -0.30156 | 1.62E-05 | -4.31161 | 0.811375 | 4.325466 |
| CNAG_017 | - | nuclear pr | 292.0717462 | 245.8160273 | 170.8946346 | 409.219691 | 424.8313307 | 355.1329077 | 4.74E-05 | 0.732968 | 1.63E-05 | 4.310982 | 1.662055 | 4.324385 |
| CNAG_016 | - | Ras family | 12.05638118 | 3.48069886 | 7.660035671 | 33.25275378 | 40.28349845 | 38.4369108 | 4.74E-05 | 2.316775 | 1.63E-05 | 4.310687 | 4.982171 | 4.323959 |
| CNAG_012 | - | hypothetic | 7476.935198 | 7430.972907 | 6846.621993 | 6054.870484 | 6155.976963 | 6253.618759 | 4.78E-05 | -0.25175 | 1.64E-05 | -4.30876 | 0.839879 | 4.320335 |
| CNAG_042 | - | glycosyl-h | 764.1794407 | 1017.149187 | 968.4134266 | 1189.422128 | 1267.323279 | 1299.787581 | 4.93E-05 | 0.435734 | 1.69E-05 | 4.301726 | 1.352599 | 4.307148 |
| CNAG_063 | - | hypothetic | 410.5980492 | 405.9949534 | 341.2282665 | 268.9487595 | 277.5672218 | 214.5862645 | 4.96E-05 | -0.62234 | 1.71E-05 | -4.30038 | 0.649618 | 4.304663 |
| CNAG_062 | - | hypothetic | 1103.32737 | 1069.825504 | 1109.028933 | 875.3335426 | 861.5297683 | 853.8569474 | 5.01E-05 | -0.35713 | 1.72E-05 | -4.29789 | 0.780714 | 4.30009 |
| CNAG_021 | - | cytoplasm | 1071.828111 | 1118.259307 | 1067.45612 | 808.282268 | 823.5418524 | 907.075532 | 5.03E-05 | -0.37434 | 1.73E-05 | -4.29683 | 0.771458 | 4.29833 |
| CNAG_065 | - | rhamnoga | 451.0256762 | 419.2876672 | 483.1758214 | 658.8951504 | 613.4500297 | 619.3226699 | 5.05E-05 | 0.466654 | 1.74E-05 | 4.295909 | 1.381901 | 4.296675 |
| CNAG_009 | - | glutathion | 112.4947919 | 127.2586131 | 90.87272161 | 209.3013927 | 208.3137141 | 175.2796729 | 5.05E-05 | 0.832824 | 1.74E-05 | 4.29571 | 1.781168 | 4.29644 |
| CNAG_058 | - | hypothetic | 16.99023232 | 34.63444327 | 25.79760491 | 64.32020834 | 81.77975386 | 68.51495751 | 5.08E-05 | 1.492514 | 1.75E-05 | 4.294314 | 2.813789 | 4.29386 |
| CNAG_023 | - | hypothetic | 1243.319703 | 1090.323183 | 1180.472272 | 986.7636778 | 859.7468551 | 818.4100473 | 5.09E-05 | -0.41547 | 1.75E-05 | -4.29414 | 0.749774 | 4.29367 |
| CNAG_024 | - | hypothetic | 687.9900477 | 618.5949435 | 649.5736144 | 904.7419082 | 953.4478847 | 804.8273526 | 5.19E-05 | 0.429674 | 1.79E-05 | 4.28962 | 1.346929 | 4.284986 |
| CNAG_024 | BWC2 | white colla | 3259.104625 | 3315.202907 | 3420.044168 | 2719.863299 | 2878.272402 | 2803.257743 | 5.37E-05 | -0.26595 | 1.85E-05 | -4.28194 | 0.83165 | 4.270281 |
| CNAG_061 | - | hypothetic | 1576.057831 | 1306.905813 | 1387.396551 | 1098.650154 | 1131.5304 | 1081.457514 | 5.42E-05 | -0.38315 | 1.87E-05 | -4.27971 | 0.766761 | 4.266087 |
| CNAG_020 | - | hypothetic | 78.80092717 | 63.46270439 | 87.12236968 | 27.29505793 | 28.02980783 | 35.77422352 | 5.47E-05 | -1.3571 | 1.89E-05 | -4.27772 | 0.390366 | 4.262356 |
| CNAG_075 | TAF9 | transcripti | 741.4124833 | 788.5766559 | 827.4405511 | 573.2309189 | 634.7295927 | 570.8091931 | 5.51E-05 | -0.42214 | 1.90E-05 | -4.27582 | 0.746318 | 4.258963 |
| CNAG_010 | - | hypothetic | 863.9739922 | 947.9806573 | 853.1074954 | 623.7534407 | 733.9489186 | 661.8233194 | 5.52E-05 | -0.4158 | 1.91E-05 | -4.27493 | 0.749602 | 4.257681 |
| CNAG_055 | - | exocyst cc | 476.7222628 | 506.4858275 | 470.4232315 | 639.9847284 | 654.8620201 | 696.956924 | 5.52E-05 | 0.439103 | 1.91E-05 | 4.275002 | 1.355762 | 4.257681 |
| CNAG_027 | BIM1 | BCS-induc | 24382.32506 | 20174.93039 | 23873.50808 | 14916.92169 | 19027.14256 | 18986.0496 | 5.53E-05 | -0.38691 | 1.91E-05 | -4.27471 | 0.764764 | 4.257412 |
| CNAG_043 | - | hypothetic | 318.4162761 | 265.0281787 | 352.755796 | 191.7307019 | 199.8368734 | 206.6862263 | 5.56E-05 | -0.66577 | 1.93E-05 | -4.27321 | 0.630353 | 4.254798 |
| CNAG_015 | HDA1 | histone de | 1829.739256 | 1852.036689 | 1913.133031 | 1399.766425 | 1561.124952 | 1562.084527 | 5.58E-05 | -0.3222 | 1.93E-05 | -4.27232 | 0.79985 | 4.253208 |
| CNAG_029 | - | hypothetic | 524.6158142 | 561.4380698 | 543.9900205 | 702.9934652 | 780.0451244 | 723.6915936 | 5.63E-05 | 0.422571 | 1.95E-05 | 4.270404 | 1.340314 | 4.249635 |
| CNAG_036 | ROM21 | Predicted | 929.8969041 | 1101.203059 | 1082.5176 | 1283.538155 | 1385.476849 | 1407.50021 | 5.73E-05 | 0.374312 | 1.99E-05 | 4.266399 | 1.296221 | 4.242144 |
| CNAG_064 | - | nucleolar | 2684.376233 | 2347.709561 | 2600.991574 | 1837.892575 | 2082.956962 | 2138.247042 | 5.76E-05 | -0.34918 | 2.00E-05 | -4.26491 | 0.785033 | 4.239406 |
| CNAG_064 | - | 26S protea | 2631.996161 | 2609.494427 | 2768.428616 | 2235.048973 | 2282.237716 | 2119.446514 | 5.77E-05 | -0.28658 | 2.00E-05 | -4.26468 | 0.819845 | 4.239097 |
| CNAG_052 | - | solute car | 1120.993556 | 1205.56376 | 1316.928239 | 918.0474887 | 983.1894698 | 936.0150252 | 5.83E-05 | -0.37515 | 2.02E-05 | -4.26219 | 0.771025 | 4.234421 |
| CNAG_056 | - | GPI inosit | 1028.016356 | 1305.883185 | 1182.784241 | 1521.028233 | 1620.147203 | 1485.301274 | 5.84E-05 | 0.381846 | 2.03E-05 | 4.261438 | 1.303008 | 4.23326 |
| CNAG_055 | - | DNA excis | 1963.141764 | 1959.920116 | 1806.677324 | 2292.980163 | 2375.631883 | 2455.522951 | 5.84E-05 | 0.299124 | 2.03E-05 | 4.261481 | 1.230397 | 4.23326 |
| CNAG_031 | - | sulfite red | 1934.312948 | 1400.667039 | 1511.116392 | 1150.700137 | 1282.500107 | 1108.165116 | 5.90E-05 | -0.46962 | 2.05E-05 | -4.259 | 0.722156 | 4.228825 |
| CNAG_012 | - | hypothetic | 181.5157437 | 255.4153533 | 205.9825839 | 347.6366202 | 367.0060453 | 308.5196955 | 5.91E-05 | 0.658879 | 2.05E-05 | 4.258847 | 1.578856 | 4.228684 |
| CNAG_121 | - | unspecifie | 19.18354885 | 21.7568735 | 18.33629686 | 55.10559058 | 55.60154024 | 62.81737615 | 6.01E-05 | 1.553563 | 2.09E-05 | 4.254683 | 2.935412 | 4.221207 |
| CNAG_006 | - | TFIIH basa | 49.63657666 | 46.61850387 | 79.00878245 | 101.2968071 | 151.6815484 | 136.1352876 | 6.09E-05 | 1.140752 | 2.12E-05 | 4.251546 | 2.204959 | 4.215274 |
| CNAG_020 | PBP1 | poly(A)-bi | 1189.76611 | 1222.781475 | 1183.69835 | 995.2657694 | 867.4798694 | 959.2973 | 6.09E-05 | -0.36433 | 2.12E-05 | -4.25142 | 0.776829 | 4.215174 |
| CNAG_027 | - | hypothetic | 2099.39887 | 2117.04574 | 1894.009001 | 1638.822196 | 1601.590423 | 1715.28294 | 6.26E-05 | -0.31757 | 2.19E-05 | -4.24507 | 0.802423 | 4.203334 |
| CNAG_016 | - | 3-methylc | 87.09303607 | 112.2907905 | 166.4047305 | 205.7945863 | 241.9911646 | 241.6395899 | 6.52E-05 | 0.897306 | 2.28E-05 | 4.235717 | 1.862585 | 4.185991 |
| CNAG_040 | - | U3 small r | 1989.66736 | 1827.31225 | 2122.119059 | 1531.357614 | 1578.323862 | 1642.021215 | 6.54E-05 | -0.33745 | 2.29E-05 | -4.23484 | 0.79144 | 4.184454 |
| CNAG_025 | - | cobalamin | 225.0054202 | 245.3981671 | 201.4941581 | 146.7189492 | 129.9087445 | 134.5760231 | 6.54E-05 | -0.72887 | 2.29E-05 | -4.23474 | 0.603377 | 4.184407 |
| CNAG_025 | - | DNA polyı | 1653.430529 | 1773.343347 | 1663.620623 | 1399.148677 | 1378.22026 | 1384.488559 | 6.56E-05 | -0.30536 | 2.30E-05 | -4.23391 | 0.809243 | 4.183113 |
| CNAG_030 | - | enolase | 36812.13192 | 37271.56911 | 37387.67661 | 30992.82309 | 33577.71977 | 32394.58111 | 6.61E-05 | -0.21644 | 2.32E-05 | -4.23203 | 0.860684 | 4.179631 |
| CNAG_052 | - | nuclear pr | 2809.397186 | 2703.212881 | 2604.250145 | 2206.979889 | 2207.138059 | 2320.313067 | 6.65E-05 | -0.28481 | 2.33E-05 | -4.23073 | 0.820852 | 4.177276 |
| CNAG_004 | - | hypothetic | 2016.402492 | 1805.733931 | 1831.053722 | 1571.189615 | 1538.673189 | 1420.076764 | 6.74E-05 | -0.33534 | 2.36E-05 | -4.22767 | 0.7926 | 4.171511 |
| CNAG_006 | - | hypothetic | 1920.808541 | 1936.07285 | 1952.606924 | 1688.788834 | 1538.633898 | 1496.661858 | 6.83E-05 | -0.31386 | 2.40E-05 | -4.22446 | 0.804485 | 4.165466 |
| CNAG_018 | - | hypothetic | 2390.18017 | 2236.710606 | 2334.401764 | 1875.420886 | 2008.827563 | 1826.367388 | 6.87E-05 | -0.30154 | 2.41E-05 | -4.22292 | 0.811387 | 4.162804 |
| CNAG_077 | - | cell divisic | 1650.898893 | 1355.4357 | 1785.055841 | 1071.988053 | 1178.046202 | 1304.51477 | 6.94E-05 | -0.44721 | 2.44E-05 | -4.22066 | 0.73346 | 4.158603 |
| CNAG_024 | - | G patch d | 1454.700178 | 1367.171691 | 1255.199786 | 1046.487274 | 1064.224978 | 1105.825354 | 6.95E-05 | -0.35767 | 2.44E-05 | -4.22025 | 0.780422 | 4.158034 |
| CNAG_063 | - | hypothetic | 362.8795769 | 361.8851044 | 361.4152353 | 265.896447 | 246.442797 | 232.2665863 | 7.10E-05 | -0.56021 | 2.50E-05 | -4.21502 | 0.678205 | 4.148495 |
| CNAG_057 | - | phosphati | 317.9466712 | 349.1809621 | 310.4749412 | 439.1649024 | 481.5847162 | 482.6966814 | 7.11E-05 | 0.507695 | 2.50E-05 | 4.214821 | 1.421777 | 4.148263 |
| CNAG_068 | - | hypothetic | 40.18220526 | 43.29710208 | 31.16897813 | 129.3965824 | 83.05805222 | 72.10497428 | 7.14E-05 | 1.302194 | 2.51E-05 | 4.213615 | 2.466036 | 4.146092 |
| CNAG_130 | - | unspecifie | 5.241847545 | 16.33110788 | 10.86350129 | 51.02168251 | 32.039367 | 47.25445385 | 7.16E-05 | 2.056381 | 2.52E-05 | 4.212904 | 4.159417 | 4.144876 |
| CNAG_022 | - | ribonucle | 2373.844129 | 2360.568921 | 2193.674309 | 1913.595976 | 1916.476225 | 1908.478482 | 7.17E-05 | -0.28691 | 2.52E-05 | -4.21264 | 0.819656 | 4.144525 |
| CNAG_012 | - | hypothetic | 6.296455153 | 8.807215285 | 4.462060494 | 26.86211282 | 29.68101765 | 41.5345046 | 7.43E-05 | 2.401288 | 2.62E-05 | 4.204455 | 5.282745 | 4.12909 |
| CNAG_050 | - | hypothetic | 368.0428573 | 277.1374328 | 355.2269958 | 513.3715644 | 543.8119282 | 450.6526358 | 7.46E-05 | 0.574079 | 2.63E-05 | 4.203309 | 1.488726 | 4.127043 |
| CNAG_010 | - | hypothetic | 160.4354516 | 185.4595227 | 189.9389512 | 306.5472596 | 284.0705937 | 258.5548298 | 7.82E-05 | 0.652316 | 2.76E-05 | 4.192658 | 1.571689 | 4.106922 |
| CNAG_041 | - | hypothetic | 68.12980856 | 73.42029361 | 47.17680581 | 133.0066841 | 150.7485275 | 108.6960877 | 7.82E-05 | 1.04142 | 2.76E-05 | 4.192526 | 2.058252 | 4.10682 |
| CNAG_019 | RCV1 | Protein re | 363.4834185 | 339.5711866 | 410.6458546 | 557.7567704 | 515.1282366 | 518.2942186 | 7.87E-05 | 0.500266 | 2.78E-05 | 4.191077 | 1.414475 | 4.104196 |

| | | | | | | | | | | | | | | |
|---|---|---|---|---|---|---|---|---|---|---|---|---|---|---|
| CNAG_130 | - | unspecifie | 83.42565902 | 111.0926435 | 90.82686711 | 154.3845639 | 183.3148253 | 179.516232 | 7.96E-05 | 0.847217 | 2.81E-05 | 4.188344 | 1.799028 | 4.099114 |
| CNAG_027 | - | mitochonc | 1850.630538 | 1679.682943 | 1760.651171 | 1369.471534 | 1471.414166 | 1439.99162 | 8.12E-05 | -0.32143 | 2.87E-05 | -4.18371 | 0.800276 | 4.090706 |
| CNAG_032 | - | hypothetic | 712.6678329 | 774.7818928 | 840.41246 | 999.5525112 | 1043.044982 | 1022.317144 | 8.23E-05 | 0.382461 | 2.91E-05 | 4.180316 | 1.303564 | 4.08437 |
| CNAG_056 | - | hypothetic | 499.5742147 | 538.543298 | 490.4935377 | 380.3013427 | 320.5321415 | 387.5694573 | 8.32E-05 | -0.50497 | 2.94E-05 | -4.1778 | 0.704675 | 4.080007 |
| CNAG_021 | - | Alpha-amy | 2044.352375 | 1962.945281 | 1614.633662 | 1554.089648 | 1383.31802 | 1400.086938 | 8.33E-05 | -0.38899 | 2.95E-05 | -4.1773 | 0.763663 | 4.079206 |
| CNAG_064 | - | hypothetic | 129.2207459 | 131.2475512 | 132.0643235 | 68.6998592 | 75.62427199 | 66.90149197 | 8.38E-05 | -0.92269 | 2.97E-05 | -4.17581 | 0.527526 | 4.076517 |
| CNAG_016 | - | polyadeny | 442.856872 | 435.4211237 | 466.1244505 | 637.0485092 | 646.3347207 | 572.7738847 | 8.41E-05 | 0.450467 | 2.98E-05 | 4.17503 | 1.366483 | 4.075178 |
| CNAG_024 | - | AP-3 com | 1536.598304 | 1609.566985 | 1462.07863 | 1264.510722 | 1273.880755 | 1189.143266 | 8.58E-05 | -0.32124 | 3.04E-05 | -4.17047 | 0.800382 | 4.066634 |
| CNAG_064 | - | atypical/R | 1542.363655 | 1372.638125 | 1588.897893 | 1140.636951 | 1254.928054 | 1154.723797 | 8.58E-05 | -0.35954 | 3.04E-05 | -4.17016 | 0.779414 | 4.066341 |
| CNAG_007 | - | hypothetic | 64.69774578 | 47.70851693 | 84.34220427 | 128.1470644 | 152.4289257 | 125.227892 | 8.58E-05 | 1.037623 | 3.04E-05 | 4.17023 | 2.052843 | 4.066341 |
| CNAG_122 | - | unspecifie | 3.933076785 | 5.575633353 | 4.437248852 | 28.87127951 | 29.89577411 | 22.02917647 | 8.67E-05 | 2.704272 | 3.07E-05 | 4.167864 | 6.51729 | 4.062116 |
| CNAG_045 | - | hypothetic | 1670.798939 | 1614.984429 | 1519.665924 | 1247.294311 | 1227.217569 | 1370.00113 | 8.78E-05 | -0.33649 | 3.12E-05 | -4.16475 | 0.791965 | 4.056495 |
| CNAG_013 | - | large subu | 1488.77623 | 1513.72876 | 1636.890878 | 1295.692577 | 1131.428214 | 1257.920809 | 8.96E-05 | -0.34742 | 3.19E-05 | -4.15977 | 0.785987 | 4.047453 |
| CNAG_075 | - | xylitol deh | 120.7629424 | 164.9186091 | 118.5747869 | 215.9475507 | 221.9446551 | 250.5170133 | 9.09E-05 | 0.757243 | 3.23E-05 | 4.156479 | 1.690258 | 4.041353 |
| CNAG_001 | - | endoplasn | 396.5584863 | 327.4915898 | 375.2488571 | 277.6486715 | 231.7230452 | 221.1978635 | 9.32E-05 | -0.60691 | 3.32E-05 | -4.1506 | 0.656603 | 4.030494 |
| CNAG_060 | - | hypothetic | 72.91070056 | 92.79923545 | 78.03362038 | 171.3465095 | 126.0057407 | 163.9256453 | 9.35E-05 | 0.912646 | 3.33E-05 | 4.149853 | 1.882495 | 4.029221 |
| CNAG_075 | - | glucose-6 | 1814.353376 | 1635.482756 | 1622.030359 | 1438.163971 | 1187.460508 | 1341.200047 | 9.52E-05 | -0.37001 | 3.39E-05 | -4.14544 | 0.773778 | 4.021302 |
| CNAG_062 | - | transporti | 860.1016994 | 674.4516221 | 728.3151283 | 521.0246247 | 597.6366502 | 523.0957761 | 9.70E-05 | -0.47946 | 3.46E-05 | -4.14095 | 0.717248 | 4.013081 |
| CNAG_004 | ISP6 | Identified | 810.0683079 | 907.9939639 | 622.8543868 | 510.0625313 | 572.5491733 | 566.344637 | 9.72E-05 | -0.51956 | 3.46E-05 | -4.14058 | 0.697585 | 4.012529 |
| CNAG_053 | - | regucalcin | 1812.084228 | 1613.970566 | 1738.245534 | 1376.533213 | 1343.680328 | 1438.856945 | 9.80E-05 | -0.32834 | 3.49E-05 | -4.1386 | 0.796454 | 4.008942 |
| CNAG_073 | - | hypothetic | 58.97097563 | 76.68084253 | 84.37166977 | 158.9191943 | 135.7211184 | 130.8064739 | 9.97E-05 | 0.941717 | 3.56E-05 | 4.134474 | 1.920812 | 4.001429 |
| CNAG_063 | - | Guanine c | 55.24446923 | 29.41242748 | 53.47661283 | 86.60580116 | 113.5436536 | 120.4850668 | 9.98E-05 | 1.202014 | 3.56E-05 | 4.134126 | 2.300606 | 4.000921 |
| CNAG_010 | - | hypothetic | 685.2103384 | 650.6945761 | 682.4301733 | 519.6364771 | 532.9295911 | 456.512006 | 9.98E-05 | -0.43617 | 3.57E-05 | -4.134 | 0.739093 | 4.000833 |
| CNAG_018 | UPF1 | ATP-deper | 958.4316172 | 862.9801111 | 924.52731 | 718.883928 | 663.0636962 | 732.8156879 | 0.0001 | -0.39254 | 3.59E-05 | -4.13257 | 0.761787 | 3.998277 |
| CNAG_030 | FZC20 | hypothetic | 1002.927868 | 951.3379871 | 1095.110395 | 758.4798457 | 790.7950695 | 816.0783483 | 0.000102 | -0.3822 | 3.63E-05 | -4.12985 | 0.767268 | 3.993284 |
| CNAG_057 | - | endoplasn | 7.492442848 | 9.900330687 | 8.717574623 | 41.5362632 | 41.18105524 | 25.4749891 | 0.000102 | 2.141393 | 3.64E-05 | 4.129209 | 4.411877 | 3.992223 |
| CNAG_051 | - | myosin-1 | 2791.065937 | 2710.903476 | 2588.379444 | 3081.276994 | 3564.34253 | 3385.865973 | 0.000103 | 0.294855 | 3.67E-05 | 4.127339 | 1.226762 | 3.988841 |
| CNAG_050 | - | hypothetic | 16.93748015 | 22.85244995 | 28.93825683 | 55.70045664 | 60.77217383 | 72.72909504 | 0.000104 | 1.487918 | 3.74E-05 | 4.123173 | 2.80484 | 3.981424 |
| CNAG_079 | - | hypothetic | 632.3929469 | 511.7336138 | 542.7420758 | 449.7100497 | 362.8250475 | 367.7039589 | 0.000105 | -0.5324 | 3.77E-05 | -4.12142 | 0.691405 | 3.978262 |
| CNAG_018 | - | hypothetic | 189.6876347 | 233.9419495 | 248.5737869 | 395.3752932 | 370.402388 | 295.2904004 | 0.000105 | 0.646939 | 3.77E-05 | 4.121025 | 1.565842 | 3.977816 |
| CNAG_024 | - | pre-mRN/ | 2950.665852 | 2808.797739 | 2641.589883 | 2248.978901 | 2384.014338 | 2334.772628 | 0.000106 | -0.2853 | 3.81E-05 | -4.119 | 0.82057 | 3.974292 |
| CNAG_042 | - | hypothetic | 3497.072826 | 3096.492177 | 2980.722216 | 2448.934936 | 2636.719937 | 2686.674927 | 0.000106 | -0.31676 | 3.81E-05 | -4.11883 | 0.802872 | 3.974127 |
| CNAG_020 | - | DASH con | 399.4344494 | 564.2957754 | 510.6807312 | 352.2479052 | 298.9573762 | 348.7561278 | 0.000106 | -0.57532 | 3.81E-05 | -4.11844 | 0.671136 | 3.973542 |
| CNAG_062 | NGL3 | RNA exon | 1724.578909 | 1707.650867 | 1631.638594 | 1410.8453 | 1364.411379 | 1383.376125 | 0.000107 | -0.29936 | 3.85E-05 | -4.11613 | 0.812612 | 3.969332 |
| CNAG_033 | - | hypothetic | 176.8516504 | 233.9098265 | 231.5173049 | 322.5769464 | 318.4907086 | 354.9422244 | 0.000109 | 0.620904 | 3.90E-05 | 4.113238 | 1.537839 | 3.964034 |
| CNAG_029 | - | hypothetic | 95.02132011 | 116.4687445 | 85.52204312 | 168.546128 | 181.5289175 | 176.2443019 | 0.000109 | 0.812334 | 3.91E-05 | 4.112649 | 1.756051 | 3.963074 |
| CNAG_000 | - | solute car | 2428.570241 | 2167.734663 | 2052.910127 | 1716.589853 | 1879.429579 | 1741.987666 | 0.00011 | -0.33271 | 3.94E-05 | -4.11111 | 0.794044 | 3.960329 |
| CNAG_060 | - | Proliferati | 5138.010394 | 5270.731939 | 5086.950451 | 4466.432404 | 4461.055544 | 4462.935249 | 0.000112 | -0.22577 | 4.02E-05 | -4.10631 | 0.855141 | 3.951454 |
| CNAG_026 | - | hypothetic | 54.27087686 | 62.70317612 | 76.89594038 | 138.5984773 | 123.6723448 | 119.6790767 | 0.000112 | 0.969821 | 4.05E-05 | 4.104809 | 1.958598 | 3.948924 |
| CNAG_045 | - | hypothetic | 1.677219231 | 10.97393221 | 12.9478685 | 38.96589463 | 40.2333983 | 35.23227518 | 0.000113 | 2.28365 | 4.06E-05 | 4.104211 | 4.869083 | 3.947947 |
| CNAG_037 | - | dTDP-4-de | 8094.621288 | 7159.533661 | 7724.263939 | 6588.842497 | 6326.735218 | 6447.917418 | 0.000113 | -0.26284 | 4.07E-05 | -4.10334 | 0.833445 | 3.946455 |
| CNAG_052 | BUD16 | pyridoxal | 512.3985869 | 551.4634238 | 475.5957177 | 349.0047019 | 391.4433351 | 375.397967 | 0.000115 | -0.47891 | 4.16E-05 | -4.09842 | 0.717519 | 3.937662 |
| CNAG_018 | - | myo-inosi | 1422.098864 | 1298.280527 | 1427.889424 | 1118.945048 | 1088.37391 | 1121.37791 | 0.000115 | -0.33348 | 4.16E-05 | -4.09833 | 0.793621 | 3.937639 |
| CNAG_010 | - | hypothetic | 81.0617851 | 85.32377433 | 102.4877963 | 159.2784082 | 152.9229137 | 173.9269439 | 0.000116 | 0.837655 | 4.20E-05 | 4.096248 | 1.787143 | 3.933887 |
| CNAG_020 | - | hypothetic | 149.8287818 | 130.5940044 | 143.0647252 | 222.8472889 | 199.2571557 | 333.4151803 | 0.000116 | 0.822221 | 4.20E-05 | 4.096072 | 1.768126 | 3.933704 |
| CNAG_011 | RRP40 | exosome | 1123.114162 | 1008.447899 | 1127.133969 | 902.662543 | 845.1369312 | 776.2130867 | 0.000118 | -0.38467 | 4.27E-05 | -4.09245 | 0.765953 | 3.927058 |
| CNAG_064 | - | hypothetic | 457.9484551 | 405.2731717 | 438.4104531 | 580.8615635 | 601.4191952 | 608.1824694 | 0.00012 | 0.444956 | 4.32E-05 | 4.089603 | 1.361272 | 3.922025 |
| CNAG_029 | - | hypothetic | 328.9404986 | 303.738554 | 319.8133549 | 215.1882672 | 204.9983379 | 224.4162939 | 0.00012 | -0.57855 | 4.32E-05 | -4.08961 | 0.669636 | 3.922025 |
| CNAG_038 | SSK1 | osomolarit | 483.6093726 | 441.8764644 | 418.1867563 | 597.9743138 | 607.4205653 | 646.9992839 | 0.00012 | 0.447729 | 4.32E-05 | 4.08949 | 1.363892 | 3.92196 |
| CNAG_047 | - | hypothetic | 4.274632586 | 26.21631683 | 31.21908858 | 175.5739908 | 157.9644701 | 125.0538062 | 0.00012 | 2.893527 | 4.34E-05 | 4.088578 | 7.43085 | 3.9204 |
| CNAG_074 | - | heat shocl | 736.7132619 | 827.2988102 | 737.9296872 | 621.6702811 | 540.5603564 | 581.8757698 | 0.000121 | -0.41546 | 4.36E-05 | -4.08753 | 0.749782 | 3.918729 |
| CNAG_003 | - | 2-oxoisov | 170.9919874 | 217.7569166 | 215.5277416 | 330.5749731 | 280.4309835 | 333.85811 | 0.000121 | 0.633242 | 4.39E-05 | 4.085903 | 1.551047 | 3.915833 |
| CNAG_004 | - | hypothetic | 1979.503533 | 1947.0069 | 1900.507534 | 2237.624207 | 2437.749545 | 2576.489668 | 0.000125 | 0.300238 | 4.52E-05 | 4.079399 | 1.231348 | 3.904115 |
| CNAG_042 | - | hypothetic | 2096.019353 | 2191.415255 | 2150.979733 | 1858.419292 | 1773.283031 | 1730.877658 | 0.000126 | -0.27896 | 4.58E-05 | -4.07619 | 0.824187 | 3.898699 |
| CNAG_055 | - | hypothetic | 2911.467418 | 3154.790888 | 3247.392248 | 3911.977351 | 3716.012854 | 3687.884091 | 0.000126 | 0.26597 | 4.58E-05 | 4.076199 | 1.202444 | 3.898699 |
| CNAG_024 | - | vacuolar p | 1868.339668 | 2052.398274 | 1986.732852 | 1704.339335 | 1490.277339 | 1607.628177 | 0.000126 | -0.31356 | 4.58E-05 | -4.07628 | 0.804651 | 3.898699 |
| CNAG_015 | - | nucleolar | 4333.293607 | 3781.793947 | 4465.123215 | 3046.577 | 3513.104629 | 3542.600905 | 0.000126 | -0.33257 | 4.58E-05 | -4.07632 | 0.794123 | 3.898699 |
| CNAG_002 | LYS9 | alpha-ami | 824.9091779 | 1152.875064 | 991.9068183 | 1202.485873 | 1466.684962 | 1426.331138 | 0.000128 | 0.450592 | 4.63E-05 | 4.073738 | 1.366601 | 3.894424 |
| CNAG_036 | - | mRNA 3'- | 1356.72093 | 1299.325405 | 1321.273602 | 1033.25861 | 1081.510766 | 1100.266005 | 0.000128 | -0.32236 | 4.63E-05 | -4.07354 | 0.799763 | 3.894202 |
| CNAG_003 | - | hypothetic | 212.0332041 | 397.5719664 | 294.4975164 | 477.5360081 | 506.7018032 | 472.7223814 | 0.00013 | 0.675454 | 4.73E-05 | 4.068666 | 1.597099 | 3.885551 |
| CNAG_120 | - | unspecifie | 47.23514846 | 59.42583246 | 50.31663514 | 131.1129232 | 116.0579017 | 85.45458988 | 0.000132 | 1.074949 | 4.81E-05 | 4.064838 | 2.106647 | 3.879149 |
| CNAG_057 | - | hypothetic | 859.0148897 | 788.5557153 | 689.9850725 | 644.3135611 | 535.3766993 | 524.2310949 | 0.000132 | -0.47134 | 4.82E-05 | -4.06432 | 0.721296 | 3.878337 |
| CNAG_017 | IRK6 | serine/thre | 4514.716877 | 4896.959986 | 4370.418007 | 5434.952018 | 5460.974886 | 5622.848702 | 0.000133 | 0.246546 | 4.83E-05 | 4.063601 | 1.186364 | 3.877138 |
| CNAG_038 | UBP7 | putative u | 486.0617882 | 532.3234006 | 475.7584736 | 649.3174373 | 681.6379524 | 680.3594171 | 0.000135 | 0.41414 | 4.93E-05 | 4.058985 | 1.332504 | 3.868986 |
| CNAG_076 | - | hypothetic | 13.22797695 | 8.84761181 | 7.670171938 | 31.48375394 | 46.46136467 | 37.41763515 | 0.000135 | 2.008847 | 4.93E-05 | 4.059055 | 4.024605 | 3.868986 |
| CNAG_077 | FRE1 | ferric-chel | 2210.679172 | 2181.867101 | 2003.970005 | 2573.19895 | 2616.279269 | 2636.517431 | 0.000136 | 0.276183 | 4.96E-05 | 4.057547 | 1.210986 | 3.866603 |
| CNAG_043 | UTR1 | NAD kinas | 2056.41806 | 2477.935192 | 2262.949361 | 1725.895885 | 1836.243883 | 1923.98293 | 0.000137 | -0.32363 | 5.00E-05 | -4.05545 | 0.799058 | 3.863144 |
| CNAG_003 | - | hypothetic | 687.4948556 | 677.5748111 | 623.8256545 | 529.0507773 | 424.9032347 | 507.4743985 | 0.000137 | -0.45956 | 5.00E-05 | -4.0555 | 0.727208 | 3.863144 |

| | | | | | | | | | | | | | | |
|---|---|---|---|---|---|---|---|---|---|---|---|---|---|---|
| CNAG_057 | - | hypothetic | 1472.208219 | 1298.266919 | 1328.750734 | 1138.468003 | 1084.075699 | 1030.405206 | 0.000138 | -0.34937 | 5.03E-05 | -4.05409 | 0.784928 | 3.860906 |
| CNAG_006 | - | ARF guany | 2763.071792 | 2764.769747 | 2811.228486 | 3285.311552 | 3413.30472 | 3295.990535 | 0.000138 | 0.245894 | 5.05E-05 | 4.053229 | 1.185827 | 3.85945 |
| CNAG_004 | CDC14 | cell divisio | 207.1397422 | 240.4009484 | 257.0968187 | 373.3643866 | 332.2447302 | 352.8292175 | 0.00014 | 0.571557 | 5.13E-05 | 4.049496 | 1.486126 | 3.85281 |
| CNAG_061 | - | long-chair | 7622.908533 | 6892.336803 | 7745.568237 | 6318.751885 | 6245.674682 | 6271.399778 | 0.000143 | -0.25684 | 5.25E-05 | -4.04425 | 0.836918 | 3.843223 |
| CNAG_023 | - | hypothetic | 212.905428 | 222.1029233 | 232.5959226 | 288.7818167 | 361.7828992 | 369.3586118 | 0.000148 | 0.596954 | 5.43E-05 | 4.036224 | 1.51252 | 3.828798 |
| CNAG_040 | - | hypothetic | 515.3932608 | 595.9320736 | 649.5186737 | 770.7211226 | 887.0799779 | 760.370699 | 0.000149 | 0.443342 | 5.46E-05 | 4.034882 | 1.359751 | 3.826461 |
| CNAG_056 | - | cysteine sy | 1570.417671 | 1535.264686 | 1469.538818 | 1216.967094 | 1283.38035 | 1243.498946 | 0.000154 | -0.30498 | 5.65E-05 | -4.02696 | 0.809456 | 3.8124 |
| CNAG_056 | FZC37 | nuclear pr | 2392.615346 | 2458.62728 | 2411.202349 | 2009.314842 | 2001.856891 | 2101.605603 | 0.000154 | -0.26382 | 5.66E-05 | -4.0265 | 0.832879 | 3.811687 |
| CNAG_014 | SIT4 | serine/thre | 1026.39734 | 1098.897766 | 1168.713283 | 894.8219635 | 858.9325501 | 857.1888425 | 0.000155 | -0.35054 | 5.69E-05 | -4.02537 | 0.784291 | 3.80989 |
| CNAG_025 | - | hypothetic | 630.4923642 | 710.9564983 | 677.1086068 | 525.0924168 | 524.278317 | 480.9000505 | 0.000156 | -0.41449 | 5.72E-05 | -4.02412 | 0.750283 | 3.807776 |
| CNAG_075 | - | glutamate | 1407.972185 | 1232.545691 | 1286.080399 | 934.2921214 | 1034.049698 | 1103.566685 | 0.000163 | -0.36973 | 6.01E-05 | -4.01246 | 0.773926 | 3.786962 |
| CNAG_035 | - | hypothetic | 263.5657181 | 223.0068935 | 300.4893891 | 171.5930292 | 167.0594379 | 162.3246305 | 0.000163 | -0.66871 | 6.02E-05 | -4.01217 | 0.62907 | 3.786575 |
| CNAG_006 | - | hypothetic | 922.2032573 | 843.5616686 | 837.1131732 | 643.2867414 | 721.0077668 | 645.1792563 | 0.000165 | -0.38886 | 6.07E-05 | -4.01001 | 0.763733 | 3.782898 |
| CNAG_016 | - | tRNA (ade | 1203.856445 | 1261.589159 | 1287.121665 | 999.8412894 | 965.8676743 | 1055.841301 | 0.00017 | -0.3282 | 6.27E-05 | -4.00239 | 0.796532 | 3.769188 |
| CNAG_121 | - | unspecifie | 7.589917257 | 20.61664888 | 12.99722505 | 45.85531409 | 51.50673881 | 41.93597796 | 0.000172 | 1.799961 | 6.35E-05 | 3.999411 | 3.482108 | 3.764001 |
| CNAG_055 | - | hypothetic | 254.9445287 | 258.749461 | 302.9380863 | 370.6692158 | 398.741908 | 427.1281555 | 0.000173 | 0.538614 | 6.36E-05 | 3.99886 | 1.452576 | 3.763132 |
| CNAG_005 | - | lipid partic | 873.8285104 | 1038.700072 | 998.2706803 | 1212.759345 | 1320.830866 | 1221.078001 | 0.000174 | 0.352792 | 6.40E-05 | 3.997399 | 1.27703 | 3.760597 |
| CNAG_123 | - | unspecifie | 13.45010345 | 32.42041209 | 9.857866836 | 54.25473779 | 50.26873988 | 76.97034709 | 0.000175 | 1.726299 | 6.45E-05 | 3.995626 | 3.308778 | 3.757489 |
| CNAG_030 | - | hypothetic | 454.0524903 | 494.3697155 | 451.0341504 | 374.2026642 | 320.6107771 | 308.862047 | 0.000176 | -0.49634 | 6.49E-05 | -3.99419 | 0.708902 | 3.755009 |
| CNAG_048 | - | mRNA bin | 2398.443903 | 2472.62617 | 2577.515834 | 2081.77121 | 1671.311671 | 2131.56236 | 0.000177 | -0.35491 | 6.54E-05 | -3.9924 | 0.781918 | 3.75187 |
| CNAG_007 | RAD23 | UV excisio | 636.6834384 | 646.5695513 | 643.1686273 | 793.1741965 | 984.5987972 | 814.7791449 | 0.000177 | 0.413154 | 6.55E-05 | 3.991908 | 1.331594 | 3.751107 |
| CNAG_039 | - | transcripti | 4674.765003 | 4970.273074 | 5055.045465 | 5385.710455 | 6321.911597 | 6500.903931 | 0.000178 | 0.293636 | 6.59E-05 | 3.990708 | 1.225726 | 3.749196 |
| CNAG_061 | HAD6 | glycerol-1 | 2033.991155 | 1862.848813 | 1945.152566 | 1559.416262 | 1579.191789 | 1666.426206 | 0.00018 | -0.29729 | 6.65E-05 | -3.98859 | 0.813778 | 3.745746 |
| CNAG_129 | - | unspecifie | 43.69564626 | 45.46722434 | 50.27936364 | 97.89612792 | 116.2632382 | 82.05244059 | 0.00018 | 1.080399 | 6.65E-05 | 3.98865 | 2.114621 | 3.745746 |
| CNAG_011 | - | xenobiotic | 1621.107212 | 1847.833388 | 1675.48664 | 2262.865658 | 2072.828184 | 2068.157431 | 0.000182 | 0.301304 | 6.74E-05 | 3.985458 | 1.232257 | 3.740161 |
| CNAG_011 | MAK3201 | hypothetic | 1191.002582 | 1238.965336 | 1265.793521 | 931.2533961 | 1028.925215 | 1013.679545 | 0.000183 | -0.32893 | 6.77E-05 | -3.98423 | 0.796124 | 3.73806 |
| CNAG_019 | - | hypothetic | 9.807628287 | 10.97952362 | 10.84226029 | 44.70097645 | 35.73017196 | 32.00985711 | 0.000188 | 1.921444 | 6.96E-05 | 3.977641 | 3.788021 | 3.72659 |
| CNAG_075 | - | hypothetic | 0.270400832 | 2.314903552 | 2.27092483 | 20.21708966 | 18.48677121 | 18.58113239 | 0.000188 | 3.793818 | 6.99E-05 | 3.976629 | 13.86925 | 3.724885 |
| CNAG_026 | - | pre-mRNA | 9708.512028 | 9115.049176 | 9348.487259 | 7676.001215 | 8004.445261 | 8441.70788 | 0.000191 | -0.23941 | 7.08E-05 | -3.97355 | 0.847092 | 3.719552 |
| CNAG_038 | ARK1 | NAK prote | 260.5511096 | 219.9585044 | 193.2185944 | 316.9875216 | 345.2965137 | 369.3735206 | 0.000192 | 0.600475 | 7.13E-05 | 3.97181 | 1.516216 | 3.716522 |
| CNAG_041 | - | NEK prote | 1534.558168 | 1446.014462 | 1466.451626 | 1822.963577 | 1817.540328 | 1837.203746 | 0.000194 | 0.285167 | 7.21E-05 | 3.969411 | 1.218551 | 3.712576 |
| CNAG_043 | - | phosphati | 2335.223773 | 2295.894791 | 1960.15664 | 1773.471664 | 1751.691197 | 1824.092964 | 0.000198 | -0.31635 | 7.35E-05 | -3.96452 | 0.8031 | 3.703953 |
| CNAG_018 | - | hypothetic | 669.2733916 | 636.8643625 | 554.71003 | 939.233719 | 801.4358168 | 790.3579668 | 0.000199 | 0.429387 | 7.42E-05 | 3.962258 | 1.346661 | 3.700122 |
| CNAG_120 | - | unspecifie | 337.2513597 | 377.9662066 | 350.7422678 | 222.8788561 | 182.4060515 | 286.4525264 | 0.000203 | -0.63903 | 7.55E-05 | -3.95823 | 0.642146 | 3.693512 |
| CNAG_067 | - | hypothetic | 104.2753069 | 97.15964898 | 101.4609094 | 157.499155 | 184.1449261 | 177.3435885 | 0.000203 | 0.762854 | 7.56E-05 | 3.958035 | 1.696844 | 3.693295 |
| CNAG_056 | - | metal iron | 1562.909618 | 2097.796789 | 2031.64395 | 2433.952113 | 2455.87324 | 2511.025919 | 0.000204 | 0.364816 | 7.62E-05 | 3.956003 | 1.287718 | 3.689886 |
| CNAG_045 | - | hypothetic | 1053.206358 | 1040.7605 | 1204.945547 | 881.5669709 | 886.5731758 | 825.0254969 | 0.000205 | -0.36263 | 7.66E-05 | -3.95466 | 0.777744 | 3.68759 |
| CNAG_056 | - | cleft lip ar | 1145.44912 | 1196.936122 | 1175.167045 | 943.8103612 | 957.3007056 | 955.9637034 | 0.000205 | -0.31552 | 7.67E-05 | -3.95442 | 0.80356 | 3.687444 |
| CNAG_016 | - | aldo-keto | 891.8732173 | 826.3249354 | 848.8206072 | 688.5823039 | 580.1970901 | 693.9447513 | 0.000207 | -0.40266 | 7.72E-05 | -3.95278 | 0.756462 | 3.684601 |
| CNAG_131 | - | unspecifie | 7.377299316 | 6.647778471 | 0.147951007 | 25.45183706 | 36.17241502 | 23.12944023 | 0.000209 | 2.676431 | 7.80E-05 | 3.950613 | 6.392727 | 3.680809 |
| CNAG_013 | - | cyanate hy | 1166.783057 | 1263.865591 | 1049.523257 | 1519.545151 | 1469.087594 | 1437.568202 | 0.000214 | 0.332678 | 7.98E-05 | 3.944857 | 1.259349 | 3.670512 |
| CNAG_131 | - | unspecifie | 0.271293397 | 0.130475854 | 2.27140743 | 16.80656743 | 31.26487312 | 11.09902424 | 0.000214 | 4.840533 | 7.99E-05 | 3.94475 | 28.65138 | 3.670461 |
| CNAG_030 | - | hypothetic | 1278.519977 | 1205.618119 | 1370.271631 | 1094.928431 | 995.204879 | 970.4551861 | 0.000216 | -0.34852 | 8.08E-05 | -3.94214 | 0.785389 | 3.665864 |
| CNAG_034 | - | DNA repa | 370.166777 | 458.7964184 | 463.7351817 | 269.529176 | 290.4176365 | 335.3596314 | 0.000216 | -0.54455 | 8.09E-05 | -3.9416 | 0.685607 | 3.665044 |
| CNAG_027 | CLC-A | putative v | 2323.818428 | 2580.359384 | 2585.003268 | 2048.991226 | 2115.747341 | 2103.8527 | 0.000216 | -0.27192 | 8.10E-05 | -3.94149 | 0.828217 | 3.66498 |
| CNAG_007 | AFR1 | ATP-bindi | 10413.81363 | 10101.94083 | 9390.117386 | 8163.993226 | 8893.803357 | 8533.900567 | 0.000217 | -0.24019 | 8.14E-05 | -3.94013 | 0.846632 | 3.662809 |
| CNAG_057 | - | hypothetic | 2.716119997 | 2.336615425 | 4.408613997 | 25.88405818 | 19.25071231 | 20.82293584 | 0.000224 | 2.984589 | 8.40E-05 | 3.932799 | 7.915001 | 3.649822 |
| CNAG_034 | RDP1 | RNA-depe | 1780.635627 | 1799.216833 | 1647.65255 | 1425.58676 | 1420.48452 | 1472.162485 | 0.000225 | -0.29073 | 8.42E-05 | -3.93213 | 0.817486 | 3.64875 |
| CNAG_001 | - | hypothetic | 669.5538446 | 831.8535069 | 859.6073372 | 1095.468141 | 1009.321321 | 1037.866032 | 0.000225 | 0.398556 | 8.45E-05 | 3.931213 | 1.318188 | 3.647522 |
| CNAG_076 | - | COP9 sign | 1877.631652 | 1926.374706 | 1906.762307 | 1534.51296 | 1561.947763 | 1658.642534 | 0.000225 | -0.27947 | 8.45E-05 | -3.93126 | 0.823892 | 3.647522 |
| CNAG_001 | - | hypothetic | 134.7595506 | 188.5922249 | 127.1133117 | 263.7353385 | 232.2338324 | 244.0098092 | 0.000225 | 0.705259 | 8.45E-05 | 3.931364 | 1.630438 | 3.647522 |
| CNAG_027 | - | 20S protea | 1988.846982 | 1922.237928 | 2032.712799 | 2401.218703 | 2373.870378 | 2434.470192 | 0.000226 | 0.263258 | 8.48E-05 | 3.930377 | 1.200186 | 3.646153 |
| CNAG_129 | - | unspecifie | 0.305964016 | 5.538448517 | 2.28962511 | 17.43124528 | 31.05308247 | 18.68230581 | 0.000228 | 3.224911 | 8.56E-05 | 3.92814 | 9.349577 | 3.642395 |
| CNAG_030 | - | transcripti | 2122.827398 | 2040.622717 | 2213.867439 | 1667.560134 | 1719.785746 | 1864.010754 | 0.00023 | -0.29551 | 8.64E-05 | -3.92598 | 0.814783 | 3.638912 |
| CNAG_065 | SET302 | histone de | 3458.844926 | 3633.04455 | 3779.431715 | 3082.612726 | 3156.88193 | 3027.523295 | 0.00023 | -0.24567 | 8.64E-05 | -3.92589 | 0.843427 | 3.638904 |
| CNAG_060 | - | hypothetic | 448.2464069 | 508.3536126 | 439.3256022 | 331.9544989 | 352.6182088 | 334.3385297 | 0.000232 | -0.47066 | 8.74E-05 | -3.92322 | 0.721632 | 3.634229 |
| CNAG_061 | - | S-formylg | 1572.815502 | 1631.132951 | 1510.069064 | 1317.451069 | 1292.836165 | 1285.686074 | 0.000233 | -0.29041 | 8.76E-05 | -3.92265 | 0.817669 | 3.633333 |
| CNAG_003 | - | tRNA dim | 1077.635554 | 1033.20174 | 1069.579256 | 827.8004718 | 886.6061039 | 837.2003657 | 0.000233 | -0.33359 | 8.79E-05 | -3.9219 | 0.793562 | 3.632126 |
| CNAG_040 | - | hypothetic | 600.1166338 | 115.4805727 | 158.9227718 | 26.01461622 | 40.67377445 | 43.86654381 | 0.000235 | -3.02415 | 8.85E-05 | -3.9202 | 0.122925 | 3.629338 |
| CNAG_029 | - | RNA-bind | 3320.608969 | 2584.766683 | 2773.802028 | 2130.567981 | 2251.964815 | 2444.642036 | 0.000238 | -0.36263 | 8.96E-05 | -3.91724 | 0.777743 | 3.624285 |
| CNAG_006 | - | pre-rRNA- | 1502.703339 | 1337.100113 | 1648.585264 | 1130.51038 | 1159.958676 | 1244.573591 | 0.000239 | -0.3607 | 9.00E-05 | -3.91612 | 0.778786 | 3.622421 |
| CNAG_006 | - | histone de | 113.7172326 | 119.7790971 | 136.6094977 | 220.1542413 | 174.2959923 | 223.8952182 | 0.00024 | 0.73069 | 9.06E-05 | 3.91444 | 1.659432 | 3.619672 |
| CNAG_057 | - | V-type AT | 1533.196534 | 1669.918611 | 1641.197803 | 1410.16096 | 1289.358059 | 1252.424257 | 0.000241 | -0.3086 | 9.11E-05 | -3.91326 | 0.807426 | 3.617697 |
| CNAG_039 | - | f-actin-cap | 1538.9766 | 1648.355301 | 1498.338154 | 1252.772567 | 1298.888728 | 1306.756434 | 0.000242 | -0.29552 | 9.15E-05 | -3.91199 | 0.814779 | 3.615539 |
| CNAG_063 | - | hypothetic | 781.1336861 | 841.3683469 | 869.0426959 | 584.0557548 | 709.8334409 | 616.314056 | 0.000244 | -0.39917 | 9.23E-05 | -3.91 | 0.758296 | 3.6121 |
| CNAG_009 | - | alkylbase | 918.8757613 | 966.3455565 | 918.159368 | 702.4382373 | 761.5094607 | 762.7846804 | 0.000249 | -0.34768 | 9.42E-05 | -3.90518 | 0.785845 | 3.603725 |
| CNAG_129 | - | unspecifie | 12.12944241 | 13.12913441 | 10.85759842 | 36.2773258 | 44.5599299 | 40.73242009 | 0.00025 | 1.762595 | 9.44E-05 | 3.904455 | 3.393078 | 3.602564 |
| CNAG_045 | - | Tartrate d | 43.63565552 | 44.36841025 | 40.71426642 | 100.5082395 | 76.91131254 | 96.21102783 | 0.000252 | 1.084035 | 9.53E-05 | 3.902276 | 2.119958 | 3.598932 |

| | | | | | | | | | | | | | | |
|---|---|---|---|---|---|---|---|---|---|---|---|---|---|---|
| CNAG_024 | - | carboxype | 728.9019802 | 700.4588436 | 750.8560768 | 859.263285 | 1025.01531 | 975.6636697 | 0.000256 | 0.376933 | 9.70E-05 | 3.897969 | 1.298579 | 3.591346 |
| CNAG_065 | - | oxysterol l | 5714.744898 | 5171.653546 | 5266.115135 | 4612.744762 | 4658.591873 | 4501.816737 | 0.00026 | -0.24571 | 9.84E-05 | -3.89457 | 0.843399 | 3.585395 |
| CNAG_021 | - | DNA repli | 259.5623324 | 264.1101413 | 275.2446105 | 421.6758792 | 355.4824875 | 378.4038724 | 0.00026 | 0.517095 | 9.86E-05 | 3.894109 | 1.431071 | 3.584709 |
| CNAG_126 | - | unspecifie | 7.391492452 | 2.371309975 | 7.617932995 | 25.39590935 | 30.71460677 | 29.60335474 | 0.000261 | 2.374599 | 9.89E-05 | 3.893284 | 5.185915 | 3.583372 |
| CNAG_041 | - | transcripti | 766.2726253 | 840.4411613 | 701.8603398 | 1037.826491 | 976.5489333 | 984.5988575 | 0.000261 | 0.362824 | 9.91E-05 | 3.8927 | 1.28594 | 3.582605 |
| CNAG_017 | - | cytoplasm | 1170.976487 | 1073.07093 | 1082.399525 | 850.3423025 | 934.0581155 | 872.726876 | 0.000271 | -0.33977 | 0.000103 | -3.88376 | 0.790167 | 3.567313 |
| CNAG_129 | - | unspecifie | 602.5401673 | 707.726544 | 736.759981 | 528.2202994 | 425.7447967 | 540.7450609 | 0.000273 | -0.46829 | 0.000104 | -3.88173 | 0.722823 | 3.563971 |
| CNAG_011 | LAG1 | LAG1 fam | 814.0343359 | 756.4572207 | 681.6014161 | 1026.178733 | 954.1192776 | 963.5063399 | 0.000276 | 0.371372 | 0.000105 | 3.87915 | 1.293583 | 3.559634 |
| CNAG_054 | LIV11 | virulence | 2191.821773 | 2303.486661 | 2382.37671 | 1936.188312 | 1950.136603 | 1907.369238 | 0.000276 | -0.2624 | 0.000105 | -3.87891 | 0.833703 | 3.559342 |
| CNAG_009 | - | hypothetic | 805.408875 | 722.8971333 | 735.7884759 | 624.0405458 | 576.0062377 | 524.2209873 | 0.000277 | -0.40756 | 0.000105 | -3.87782 | 0.753899 | 3.557814 |
| CNAG_009 | - | aflatoxin e | 337.7475959 | 243.4177057 | 227.1308432 | 174.7175906 | 136.7076441 | 182.2303311 | 0.000277 | -0.72995 | 0.000106 | -3.87744 | 0.602923 | 3.557281 |
| CNAG_063 | - | pre-rRNA- | 2569.078247 | 2865.891749 | 3090.45481 | 2252.857483 | 2484.975237 | 2240.455338 | 0.00028 | -0.30425 | 0.000107 | -3.8745 | 0.809863 | 3.552169 |
| CNAG_061 | - | hypothetic | 802.6169435 | 1042.958757 | 845.8199295 | 1127.130365 | 1189.658243 | 1229.865441 | 0.000281 | 0.384715 | 0.000107 | 3.873943 | 1.305602 | 3.551321 |
| CNAG_001 | - | hypothetic | 177.8405648 | 183.3063458 | 185.679693 | 265.7114502 | 266.7318369 | 295.0107186 | 0.000281 | 0.584856 | 0.000107 | 3.873712 | 1.499889 | 3.551048 |
| CNAG_011 | - | glutamine | 5268.909651 | 4518.863757 | 4336.317162 | 5831.174004 | 5743.909167 | 5874.851847 | 0.000284 | 0.289314 | 0.000108 | 3.870793 | 1.222059 | 3.545983 |
| CNAG_052 | - | hypothetic | 5.098677351 | 7.713333814 | 4.443562675 | 29.6194964 | 28.93932709 | 24.21112364 | 0.000286 | 2.32555 | 0.000109 | 3.869124 | 5.012569 | 3.543428 |
| CNAG_047 | RAS2 | Ras family | 170.8348681 | 170.3942531 | 182.470311 | 246.1616499 | 276.3383118 | 276.1767086 | 0.000287 | 0.597008 | 0.00011 | 3.868267 | 1.512576 | 3.542041 |
| CNAG_079 | - | hypothetic | 6.303531412 | 7.746190138 | 8.697788581 | 44.309435 | 29.66007414 | 22.16092951 | 0.000287 | 2.162453 | 0.00011 | 3.86812 | 4.476754 | 3.541918 |
| CNAG_024 | - | methionyl | 61.2879374 | 70.25175959 | 91.80425593 | 126.3800222 | 126.1143763 | 174.8075384 | 0.00029 | 0.925532 | 0.000111 | 3.865752 | 1.899384 | 3.538119 |
| CNAG_043 | ATM1 | iron-sulfur | 1045.994854 | 936.2432474 | 961.8824497 | 726.5467 | 772.6880962 | 816.0505141 | 0.00029 | -0.36203 | 0.000111 | -3.86577 | 0.778069 | 3.538119 |
| CNAG_042 | - | YEATS dor | 1408.121358 | 1315.515684 | 1442.818286 | 1062.79985 | 1139.307826 | 1172.410815 | 0.00029 | -0.31987 | 0.000111 | -3.86588 | 0.801142 | 3.538119 |
| CNAG_009 | - | ATP-depe | 4460.615661 | 4228.877219 | 4142.080918 | 3597.81693 | 3810.730413 | 3553.745904 | 0.000301 | -0.24282 | 0.000115 | -3.85571 | 0.845092 | 3.520961 |
| CNAG_063 | - | solute car | 45851.72997 | 47263.67577 | 48146.04547 | 38058.08593 | 42833.87264 | 41684.39201 | 0.000306 | -0.21994 | 0.000117 | -3.85173 | 0.858604 | 3.514031 |
| CNAG_042 | - | nuclear RI | 1083.799219 | 996.7372816 | 1011.086567 | 1282.050292 | 1348.390539 | 1258.842429 | 0.000313 | 0.315779 | 0.00012 | 3.846017 | 1.244683 | 3.504324 |
| CNAG_010 | - | alpha/beta | 327.1921888 | 321.1882398 | 289.1553007 | 465.9051814 | 462.6226134 | 400.7142295 | 0.000314 | 0.486709 | 0.00012 | 3.84537 | 1.401245 | 3.503316 |
| CNAG_042 | - | hypothetic | 4.978422368 | 2.326352808 | 0.119645562 | 26.03256327 | 14.78699111 | 20.77116289 | 0.000319 | 3.306355 | 0.000122 | 3.841339 | 9.892639 | 3.496457 |
| CNAG_056 | - | hypothetic | 1873.987625 | 1812.154145 | 1641.267468 | 1431.790713 | 1440.320569 | 1503.242526 | 0.000319 | -0.2993 | 0.000123 | -3.84101 | 0.812644 | 3.496017 |
| CNAG_042 | - | phytanoyl | 708.6844365 | 735.7887396 | 734.7037208 | 579.5868734 | 571.7141778 | 555.2373143 | 0.00032 | -0.36771 | 0.000123 | -3.84024 | 0.775014 | 3.494927 |
| CNAG_032 | - | THO comp | 2065.596137 | 2094.443795 | 1969.711303 | 1652.817251 | 1667.172324 | 1796.294562 | 0.000325 | -0.27571 | 0.000125 | -3.83611 | 0.826041 | 3.487768 |
| CNAG_130 | - | unspecifie | 23.9319474 | 39.9978088 | 30.05085771 | 69.00179645 | 80.78197315 | 69.68040158 | 0.000326 | 1.230785 | 0.000125 | 3.835393 | 2.346947 | 3.486633 |
| CNAG_013 | - | ATP-depe | 181.4131897 | 229.5555746 | 184.6527682 | 286.7061188 | 316.9057764 | 296.2266199 | 0.00033 | 0.581872 | 0.000127 | 3.8326 | 1.49679 | 3.481836 |
| CNAG_058 | - | hypothetic | 1455.487021 | 1078.481038 | 1029.134068 | 904.0942868 | 918.4725668 | 799.5560212 | 0.000335 | -0.45918 | 0.000129 | -3.82882 | 0.727402 | 3.47531 |
| CNAG_074 | - | hypothetic | 209.4351551 | 104.8977858 | 70.75273796 | 490.3648383 | 399.8957264 | 435.7268112 | 0.000336 | 1.771487 | 0.000129 | 3.827632 | 3.414057 | 3.473344 |
| CNAG_019 | - | hypothetic | 40.70024349 | 51.47307039 | 63.61010225 | 16.44251444 | 13.33201295 | 24.62046673 | 0.000339 | -1.55308 | 0.000131 | -3.8254 | 0.340781 | 3.469686 |
| CNAG_060 | PTP1 | Putative p | 812.4185252 | 784.2483048 | 684.6559713 | 609.186862 | 548.341748 | 590.7400717 | 0.000346 | -0.39926 | 0.000134 | -3.8199 | 0.75825 | 3.460402 |
| CNAG_034 | SPP101 | compass c | 702.8998404 | 777.7783408 | 746.4393996 | 577.9849268 | 584.6527692 | 582.9680609 | 0.000347 | -0.36709 | 0.000134 | -3.81913 | 0.775343 | 3.459183 |
| CNAG_031 | - | hypothetic | 126.7128012 | 198.3181664 | 177.1473354 | 283.9137243 | 248.5827797 | 275.0597422 | 0.000348 | 0.672393 | 0.000134 | 3.818683 | 1.593715 | 3.458676 |
| CNAG_037 | - | mRNA 3'- | 1429.14222 | 1353.215465 | 1434.303171 | 1214.788449 | 1135.835881 | 1091.44979 | 0.000353 | -0.30899 | 0.000136 | -3.81472 | 0.807207 | 3.452117 |
| CNAG_067 | RPN14 | 19S protea | 784.8829239 | 671.4048307 | 768.9852446 | 985.6211683 | 1048.303995 | 896.9723233 | 0.000356 | 0.381772 | 0.000138 | 3.812502 | 1.302941 | 3.448492 |
| CNAG_021 | ASP1 | Inositol he | 2976.475458 | 2939.188662 | 2972.15716 | 2459.930502 | 2556.525321 | 2592.2968 | 0.000357 | -0.23967 | 0.000138 | -3.81202 | 0.846937 | 3.447774 |
| CNAG_065 | HMG1 | hydroxyme | 1572.630778 | 1450.14557 | 1323.465711 | 1222.537183 | 1134.945977 | 1135.841677 | 0.000358 | -0.33113 | 0.000138 | -3.81085 | 0.794911 | 3.44586 |
| CNAG_024 | - | Phospholi | 1362.899944 | 1393.166308 | 1095.411435 | 1735.193434 | 1591.526125 | 1645.137106 | 0.000363 | 0.353884 | 0.00014 | 3.807449 | 1.277997 | 3.440026 |
| CNAG_050 | - | GabA perr | 191.3949148 | 214.2613962 | 250.3577306 | 129.4543609 | 161.1839339 | 111.3586401 | 0.000364 | -0.72342 | 0.000141 | -3.80646 | 0.605661 | 3.43843 |
| CNAG_121 | - | unspecifie | 149.0764902 | 136.7246143 | 156.5719987 | 81.15882166 | 92.04474264 | 85.72584566 | 0.000367 | -0.78919 | 0.000142 | -3.80469 | 0.57867 | 3.43573 |
| CNAG_066 | - | hypothetic | 964.5727062 | 1167.776289 | 1106.884137 | 916.7065875 | 830.4560664 | 785.0836676 | 0.000367 | -0.36946 | 0.000142 | -3.80451 | 0.77407 | 3.435692 |
| CNAG_019 | - | carbohydr | 1707.252532 | 1971.578523 | 1873.692593 | 1508.075549 | 1571.479746 | 1518.81827 | 0.000367 | -0.28677 | 0.000142 | -3.80458 | 0.819736 | 3.435692 |
| CNAG_001 | - | hypothetic | 933.2468812 | 979.452813 | 891.6633519 | 1197.265268 | 1298.446454 | 1092.38048 | 0.00037 | 0.340608 | 0.000143 | 3.802426 | 1.26629 | 3.432309 |
| CNAG_122 | - | unspecifie | 12.0139789 | 6.685599924 | 5.530106808 | 40.888914 | 26.04170069 | 32.97505628 | 0.000373 | 2.07472 | 0.000145 | 3.80009 | 4.212626 | 3.428487 |
| CNAG_054 | - | SHO1 osm | 3466.982996 | 3450.986047 | 3762.357766 | 2943.245121 | 3100.815965 | 3048.591984 | 0.000376 | -0.24755 | 0.000146 | -3.79809 | 0.842325 | 3.425249 |
| CNAG_003 | - | hypothetic | 186.5557058 | 183.0479597 | 208.8218557 | 131.2101144 | 102.2439216 | 124.5345997 | 0.000382 | -0.70783 | 0.000148 | -3.79382 | 0.612242 | 3.418045 |
| CNAG_053 | - | hypothetic | 44.70172573 | 38.98589926 | 30.08713074 | 76.70751073 | 99.02244329 | 74.20517808 | 0.000382 | 1.136759 | 0.000149 | 3.793291 | 2.198865 | 3.417401 |
| CNAG_064 | - | ATP-depe | 1598.292717 | 1284.32075 | 1521.726536 | 1107.1611 | 1245.460792 | 1103.674449 | 0.000384 | -0.36626 | 0.000149 | -3.79196 | 0.77579 | 3.415341 |
| CNAG_058 | - | hypothetic | 394.8930332 | 390.1271383 | 338.2169501 | 533.53913 | 540.2382064 | 477.3037732 | 0.000389 | 0.450934 | 0.000151 | 3.788637 | 1.366925 | 3.409807 |
| CNAG_122 | - | unspecifie | 21.35987028 | 15.30977716 | 13.00990325 | 47.34382526 | 54.07615823 | 43.08156906 | 0.00039 | 1.538924 | 0.000152 | 3.787992 | 2.905778 | 3.408952 |
| CNAG_031 | - | cytoplasm | 626.2132874 | 675.6420195 | 702.8495261 | 853.2424886 | 876.5508017 | 851.3790483 | 0.000393 | 0.350126 | 0.000153 | 3.78589 | 1.274672 | 3.405689 |
| CNAG_010 | - | hypothetic | 141.6382 | 146.6708072 | 141.9724186 | 216.7205379 | 232.3614308 | 219.6201349 | 0.000399 | 0.624897 | 0.000156 | 3.781765 | 1.542101 | 3.398622 |
| CNAG_025 | - | hypothetic | 34.10146599 | 21.78278594 | 13.05383975 | 65.43544022 | 59.8970298 | 61.83296871 | 0.000403 | 1.427572 | 0.000157 | 3.779666 | 2.689937 | 3.395097 |
| CNAG_034 | - | NAD epim | 566.5011869 | 487.1555945 | 532.2504907 | 689.8092718 | 691.0654156 | 726.9675593 | 0.000403 | 0.394267 | 0.000157 | 3.779205 | 1.314275 | 3.394429 |
| CNAG_060 | RHO104 | rho family | 686.7559036 | 639.0187954 | 567.4985251 | 879.109173 | 857.6043801 | 773.7333653 | 0.000407 | 0.391731 | 0.000159 | 3.776788 | 1.311966 | 3.390486 |
| CNAG_039 | - | Transmem | 395.4075599 | 336.0940268 | 346.5197783 | 261.9722705 | 262.0258804 | 236.6968971 | 0.00041 | -0.52077 | 0.00016 | -3.77449 | 0.697001 | 3.386755 |
| CNAG_128 | - | unspecifie | 48.46348321 | 80.88664208 | 56.71722844 | 133.1695121 | 124.6237751 | 104.2392648 | 0.000411 | 0.958779 | 0.000161 | 3.773852 | 1.943664 | 3.385779 |
| CNAG_039 | - | wd-repeat | 1215.546475 | 1261.603632 | 1320.174974 | 1012.285885 | 991.7551196 | 1086.913628 | 0.000413 | -0.3124 | 0.000162 | -3.77258 | 0.805303 | 3.383699 |
| CNAG_040 | - | ATP-depe | 2134.432884 | 2010.440267 | 2007.024728 | 1588.924666 | 1726.729165 | 1791.857656 | 0.000414 | -0.28436 | 0.000162 | -3.77232 | 0.821103 | 3.383377 |
| CNAG_023 | - | hypothetic | 1116.188029 | 1073.071794 | 1114.372449 | 867.4981475 | 923.6925074 | 894.9081631 | 0.000415 | -0.31487 | 0.000162 | -3.77115 | 0.803925 | 3.381486 |
| CNAG_046 | - | hypothetic | 13.60232997 | 51.83153891 | 41.75971554 | 74.19169676 | 88.38748618 | 113.6539479 | 0.000416 | 1.377727 | 0.000163 | 3.771019 | 2.598586 | 3.381388 |
| CNAG_059 | - | O-sialogly | 15.77715088 | 30.31312579 | 22.58664005 | 45.23029942 | 62.572574 | 75.96463431 | 0.000417 | 1.428703 | 0.000163 | 3.770336 | 2.692045 | 3.380335 |
| CNAG_074 | PAN6 | pantoate-l | 112.4997624 | 87.54638751 | 141.8925651 | 211.6546187 | 191.7760329 | 183.0059606 | 0.000417 | 0.766506 | 0.000163 | 3.769827 | 1.701145 | 3.379586 |
| CNAG_028 | ADE12 | adenylosu | 17604.84381 | 15233.84796 | 18884.62562 | 13348.11215 | 15125.80625 | 14237.92712 | 0.000418 | -0.29244 | 0.000164 | -3.76948 | 0.81652 | 3.379259 |

| | | | | | | | | | | | | | | |
|---|---|---|---|---|---|---|---|---|---|---|---|---|---|---|
| CNAG_048 | - | mitochonc | 801.306874 | 849.0851013 | 808.4576226 | 1007.281185 | 1044.725002 | 1045.639675 | 0.000423 | 0.317919 | 0.000166 | 3.766176 | 1.246531 | 3.374047 |
| CNAG_064 | - | hypothetic | 238.141049 | 272.4406664 | 298.4028593 | 184.0826606 | 193.8977188 | 160.1436442 | 0.000423 | -0.60456 | 0.000166 | -3.7662 | 0.657674 | 3.374047 |
| CNAG_054 | - | hypothetic | 641.3958712 | 724.09771 | 674.0921964 | 806.3519291 | 951.6883473 | 905.7306182 | 0.000425 | 0.369782 | 0.000167 | 3.764615 | 1.292158 | 3.371469 |
| CNAG_055 | - | hypothetic | 1060.528893 | 1129.204475 | 1037.75399 | 1334.981861 | 1401.871991 | 1282.178197 | 0.000431 | 0.301326 | 0.000169 | 3.760836 | 1.232277 | 3.365174 |
| CNAG_013 | - | cell divisic | 2613.508519 | 2320.905463 | 2366.549172 | 2821.364825 | 2965.5972 | 3239.245757 | 0.000432 | 0.290592 | 0.000169 | 3.76065 | 1.223142 | 3.364986 |
| CNAG_007 | - | hypothetic | 10.82069671 | 5.598006613 | 4.454432322 | 32.8004689 | 30.64644734 | 25.34521337 | 0.000435 | 2.178145 | 0.000171 | 3.758697 | 4.525712 | 3.361731 |
| CNAG_007 | - | hypothetic | 356.5648992 | 421.3683085 | 431.9792487 | 516.1595789 | 585.1096164 | 556.0239016 | 0.000438 | 0.441106 | 0.000172 | 3.756801 | 1.357645 | 3.358577 |
| CNAG_073 | - | hypothetic | 138.1716638 | 104.4054093 | 113.9582579 | 43.522989 | 54.79587009 | 81.12788443 | 0.00044 | -1.01189 | 0.000173 | -3.75547 | 0.495897 | 3.356538 |
| CNAG_010 | - | geranylge | 598.7164037 | 523.5923949 | 564.0658679 | 444.9595425 | 418.1359925 | 409.8308829 | 0.000444 | -0.42256 | 0.000175 | -3.7532 | 0.7461 | 3.352738 |
| CNAG_015 | - | hypothetic | 92.55931047 | 86.36771244 | 73.7853138 | 179.267284 | 149.5387818 | 124.2966881 | 0.000445 | 0.833238 | 0.000175 | 3.752309 | 1.78168 | 3.351597 |
| CNAG_000 | - | D-3-phosp | 61.25015292 | 83.08866453 | 71.61588747 | 133.6475626 | 111.3533216 | 156.0385273 | 0.000449 | 0.881911 | 0.000177 | 3.749563 | 1.842815 | 3.347514 |
| CNAG_053 | - | hypothetic | 61.19175471 | 61.62849996 | 76.89242914 | 125.9000708 | 136.7949125 | 108.6776575 | 0.000462 | 0.88241 | 0.000183 | 3.742 | 1.843452 | 3.334966 |
| CNAG_131 | - | unspecifie | 59.98831795 | 49.82015847 | 71.56332498 | 94.24508255 | 145.6545369 | 116.3017513 | 0.000464 | 0.971361 | 0.000183 | 3.741266 | 1.96069 | 3.333832 |
| CNAG_075 | - | small subu | 642.1641666 | 653.9405034 | 747.4268516 | 482.8962299 | 553.6508919 | 523.0285412 | 0.000467 | -0.40574 | 0.000185 | -3.73921 | 0.754848 | 3.330683 |
| CNAG_127 | - | unspecifie | 20.63986676 | 72.2712823 | 54.56289114 | 95.20833935 | 118.6684745 | 133.7545678 | 0.00047 | 1.233471 | 0.000186 | 3.737708 | 2.351321 | 3.328221 |
| CNAG_063 | - | hypothetic | 435.4155832 | 440.5878131 | 480.8324981 | 339.8123019 | 335.3360983 | 332.1061638 | 0.00047 | -0.44496 | 0.000186 | -3.73756 | 0.734603 | 3.328104 |
| CNAG_007 | - | ATP-depe | 1684.804239 | 1507.303096 | 1650.781982 | 1141.290159 | 1445.638598 | 1259.069631 | 0.000471 | -0.34837 | 0.000186 | -3.73682 | 0.785471 | 3.32696 |
| CNAG_058 | GMT1 | GDP-manı | 6820.835676 | 7336.175522 | 7472.571961 | 6265.841771 | 6412.200514 | 6165.936112 | 0.000472 | -0.214 | 0.000187 | -3.73619 | 0.862143 | 3.326135 |
| CNAG_063 | - | hypothetic | 364.2018763 | 426.4565623 | 375.3157549 | 306.4651943 | 272.3309301 | 252.2539218 | 0.000472 | -0.50373 | 0.000187 | -3.73619 | 0.705279 | 3.326135 |
| CNAG_056 | - | hypothetic | 273.4824308 | 256.5704964 | 259.2648234 | 386.4756415 | 353.7856688 | 378.373382 | 0.000479 | 0.488032 | 0.00019 | 3.732247 | 1.402531 | 3.319735 |
| CNAG_121 | - | unspecifie | 4.048793121 | 16.30102946 | 10.84468555 | 36.41117878 | 33.92909482 | 44.98586671 | 0.000481 | 1.901723 | 0.000191 | 3.730804 | 3.736593 | 3.317515 |
| CNAG_039 | - | glucose-6 | 6938.678876 | 6822.26795 | 7067.331691 | 5720.939908 | 6313.037247 | 6036.029205 | 0.000482 | -0.2204 | 0.000191 | -3.73019 | 0.858324 | 3.316719 |
| CNAG_061 | APH4 | Acid phos | 224.6412497 | 258.7126413 | 286.933381 | 361.4341085 | 363.3367058 | 384.9922386 | 0.000488 | 0.513907 | 0.000194 | 3.727261 | 1.427912 | 3.31181 |
| CNAG_128 | - | unspecifie | 0.361671647 | 10.89717968 | 4.443821384 | 25.4331112 | 24.43325482 | 34.95071586 | 0.000489 | 2.558368 | 0.000194 | 3.726389 | 5.890411 | 3.310442 |
| CNAG_057 | - | hypothetic | 545.6663075 | 602.3768947 | 630.3276526 | 774.6779413 | 780.8031323 | 758.1284759 | 0.000491 | 0.364689 | 0.000195 | 3.725578 | 1.287604 | 3.309312 |
| CNAG_050 | - | histone de | 968.7745437 | 813.4113031 | 790.2190052 | 710.4294627 | 631.1682164 | 628.5358576 | 0.000497 | -0.4007 | 0.000197 | -3.7223 | 0.757493 | 3.303937 |
| CNAG_003 | - | methylglu | 268.9642507 | 317.9254237 | 298.7101422 | 381.496208 | 451.4702436 | 416.1396041 | 0.000502 | 0.484331 | 0.0002 | 3.719361 | 1.398937 | 3.299016 |
| CNAG_007 | - | pre-mRNA | 2267.69012 | 2304.575458 | 2416.502928 | 1900.324964 | 2099.435671 | 1859.668443 | 0.000507 | -0.2697 | 0.000202 | -3.71707 | 0.829492 | 3.295348 |
| CNAG_056 | - | hypothetic | 326.1069374 | 345.9624188 | 337.1038378 | 500.1755509 | 490.2180237 | 412.9613668 | 0.000508 | 0.460783 | 0.000202 | 3.716455 | 1.376288 | 3.294421 |
| CNAG_058 | - | translatior | 10016.87319 | 9759.356043 | 10168.5804 | 8359.366577 | 8952.450948 | 8891.342992 | 0.000508 | -0.20799 | 0.000202 | -3.71633 | 0.865741 | 3.294344 |
| CNAG_055 | - | hypothetic | 44.94778474 | 58.36765659 | 68.35669129 | 127.033585 | 101.1339767 | 105.2499303 | 0.000511 | 0.955414 | 0.000204 | 3.714484 | 1.939136 | 3.291302 |
| CNAG_046 | - | stearoyl-C | 1724.82921 | 1491.290746 | 1524.057132 | 1937.350515 | 2067.777647 | 1896.088573 | 0.000517 | 0.300186 | 0.000206 | 3.711577 | 1.231303 | 3.286578 |
| CNAG_023 | - | MRP famil | 163.2184474 | 159.3563063 | 192.7878082 | 111.7055789 | 84.96120037 | 113.4065151 | 0.000517 | -0.75401 | 0.000206 | -3.71134 | 0.592952 | 3.286296 |
| CNAG_051 | - | Nicotinate | 3186.612028 | 3151.414835 | 2911.409729 | 2520.613521 | 2714.402703 | 2633.39384 | 0.000518 | -0.24859 | 0.000207 | -3.71048 | 0.84172 | 3.285356 |
| CNAG_018 | PYK1 | pyruvate k | 11830.23217 | 10695.63805 | 10340.3264 | 9415.575478 | 9669.231054 | 9077.911374 | 0.000519 | -0.23859 | 0.000207 | -3.71013 | 0.847576 | 3.284889 |
| CNAG_026 | - | hypothetic | 464.9190975 | 430.0149532 | 403.2528485 | 594.9589119 | 538.3003678 | 628.1076754 | 0.00052 | 0.425185 | 0.000208 | 3.709314 | 1.342745 | 3.283626 |
| CNAG_045 | - | hypothetic | 616.4386329 | 639.9007415 | 667.4863716 | 518.1479744 | 515.6861419 | 433.2079106 | 0.000522 | -0.40607 | 0.000208 | -3.70871 | 0.754678 | 3.282727 |
| CNAG_044 | - | monocarb | 576.6797673 | 607.537484 | 556.6390832 | 465.2946287 | 432.7942628 | 438.6712691 | 0.000522 | -0.39634 | 0.000209 | -3.70846 | 0.759782 | 3.282421 |
| CNAG_008 | SIP401 | pathway-s | 119.4557548 | 89.69657545 | 138.7088562 | 175.3421825 | 196.1124215 | 217.2016512 | 0.000527 | 0.74867 | 0.000211 | 3.706041 | 1.680243 | 3.278413 |
| CNAG_014 | - | hypothetic | 1435.028734 | 1482.454652 | 1423.666083 | 1136.776195 | 1121.124609 | 1290.036551 | 0.000533 | -0.306 | 0.000213 | -3.70275 | 0.80888 | 3.273168 |
| CNAG_064 | OTU1 | ubiquitin t | 815.0706758 | 917.8636147 | 937.3026461 | 729.0553151 | 704.5473148 | 686.2349811 | 0.000534 | -0.3478 | 0.000214 | -3.70228 | 0.785782 | 3.272505 |
| CNAG_065 | - | hypothetic | 503.5079358 | 415.0125965 | 535.4072551 | 622.7993374 | 680.8036124 | 683.6593287 | 0.000535 | 0.433948 | 0.000214 | 3.701649 | 1.350926 | 3.271553 |
| CNAG_128 | - | unspecifie | 1.46971149 | 1.221351094 | 1.190451484 | 13.59476338 | 16.78652422 | 18.49100705 | 0.000536 | 3.954521 | 0.000215 | 3.701042 | 15.50349 | 3.270646 |
| CNAG_053 | FZC44 | hypothetic | 618.6281108 | 555.9174308 | 616.3017812 | 480.7882079 | 462.1667408 | 416.5290963 | 0.000541 | -0.41348 | 0.000217 | -3.69865 | 0.750809 | 3.266685 |
| CNAG_038 | - | hypothetic | 395.0162017 | 456.8887721 | 403.2348222 | 540.3314199 | 599.7695748 | 550.5192803 | 0.000544 | 0.414859 | 0.000218 | 3.697243 | 1.333168 | 3.264543 |
| CNAG_004 | - | hypothetic | 151.9911914 | 80.05773278 | 140.88071 | 194.0362171 | 216.8246917 | 247.1208222 | 0.000544 | 0.806956 | 0.000218 | 3.69728 | 1.749516 | 3.264543 |
| CNAG_074 | - | hypothetic | 410.1340275 | 453.6521771 | 359.5558924 | 574.7666554 | 596.3324052 | 511.7338866 | 0.000544 | 0.445248 | 0.000218 | 3.697012 | 1.361548 | 3.264281 |
| CNAG_034 | - | hypothetic | 349.3981915 | 609.4499533 | 520.2569479 | 326.4526518 | 363.8471353 | 286.7246653 | 0.000553 | -0.61352 | 0.000222 | -3.69264 | 0.653602 | 3.257077 |
| CNAG_062 | - | allergen | 2100.25652 | 3214.063475 | 2258.977383 | 5313.729568 | 4760.073131 | 3803.35067 | 0.000556 | 0.860732 | 0.000223 | 3.691358 | 1.81596 | 3.255016 |
| CNAG_033 | - | pre-mRNA | 693.2274167 | 621.5878612 | 567.3397838 | 483.0422121 | 450.0112638 | 491.9218168 | 0.000556 | -0.41697 | 0.000223 | -3.69117 | 0.748994 | 3.254823 |
| CNAG_004 | - | hypothetic | 98.64636343 | 138.0244308 | 137.6707473 | 194.9328579 | 199.5104912 | 216.1691381 | 0.000558 | 0.693087 | 0.000224 | 3.690081 | 1.616739 | 3.253099 |
| CNAG_028 | - | hypothetic | 1662.728046 | 1675.331138 | 1595.384156 | 1372.685972 | 1396.371701 | 1378.923928 | 0.000563 | -0.26563 | 0.000226 | -3.68778 | 0.831835 | 3.249302 |
| CNAG_035 | - | kynurenin | 228.1200164 | 267.3082413 | 272.0267776 | 359.8888968 | 384.143391 | 351.7915029 | 0.00058 | 0.497404 | 0.000233 | 3.680107 | 1.411672 | 3.236355 |
| CNAG_057 | CDA1 | Chitin dea | 2288.817698 | 2077.369453 | 1686.234577 | 2619.236567 | 2530.879748 | 2608.73969 | 0.000587 | 0.342972 | 0.000236 | 3.67731 | 1.268367 | 3.231725 |
| CNAG_031 | - | hypothetic | 796.1836479 | 740.1551179 | 825.3181517 | 630.9760579 | 658.0497936 | 528.6906067 | 0.000593 | -0.39452 | 0.000239 | -3.67427 | 0.760744 | 3.226817 |
| CNAG_120 | - | unspecifie | 2.836272378 | 14.13045981 | 6.588745508 | 34.24596691 | 38.60237891 | 26.49946043 | 0.000593 | 2.144187 | 0.000238 | 3.674327 | 4.420429 | 3.226817 |
| CNAG_039 | - | mitochonc | 637.4797087 | 615.1903394 | 763.39227 | 511.8153233 | 508.7187956 | 513.0408289 | 0.000595 | -0.41096 | 0.000239 | -3.6735 | 0.75212 | 3.225636 |
| CNAG_013 | - | hypothetic | 243.1158235 | 229.6285738 | 221.9540322 | 328.7713733 | 350.5104128 | 317.3847326 | 0.000603 | 0.506339 | 0.000243 | 3.669863 | 1.420441 | 3.219722 |
| CNAG_030 | - | hypothetic | 117.1099784 | 119.739225 | 113.1906102 | 183.2997309 | 194.4195551 | 181.8785037 | 0.00061 | 0.661817 | 0.000246 | 3.666893 | 1.582074 | 3.214941 |
| CNAG_014 | - | hypothetic | 3695.763379 | 3561.943772 | 3474.481653 | 2977.49265 | 3103.392097 | 3160.692266 | 0.00061 | -0.23104 | 0.000246 | -3.66673 | 0.852021 | 3.214791 |
| CNAG_050 | - | hypothetic | 704.9981005 | 621.6390362 | 660.0575841 | 508.7083715 | 545.0359013 | 475.35096 | 0.000617 | -0.39341 | 0.000249 | -3.66378 | 0.76133 | 3.209918 |
| CNAG_050 | - | hypothetic | 20.49259553 | 29.32780843 | 44.87753181 | 66.51567477 | 62.24320349 | 101.382957 | 0.000617 | 1.285804 | 0.000249 | 3.66355 | 2.438178 | 3.209663 |
| CNAG_007 | - | hypothetic | 37.81719942 | 37.91797783 | 38.56651855 | 98.37545142 | 93.73904826 | 57.78195826 | 0.000628 | 1.131128 | 0.000253 | 3.659129 | 2.1903 | 3.202299 |
| CNAG_012 | - | hypothetic | 483.5131849 | 403.1034988 | 362.7639967 | 528.5674613 | 626.5572356 | 589.3089291 | 0.000629 | 0.466037 | 0.000254 | 3.658532 | 1.38131 | 3.20155 |
| CNAG_050 | CKY1 | YjeF famil | 1130.004672 | 1085.915152 | 896.9067738 | 863.7599862 | 824.4644265 | 699.6530964 | 0.000629 | -0.39814 | 0.000254 | -3.65855 | 0.758838 | 3.20155 |
| CNAG_058 | - | cytoplasm | 2845.757876 | 2912.255905 | 3069.176289 | 2490.317864 | 2576.380745 | 2511.280934 | 0.000632 | -0.23541 | 0.000255 | -3.65719 | 0.849447 | 3.199405 |
| CNAG_017 | - | hypothetic | 375.133758 | 409.5033251 | 332.9000367 | 653.1674846 | 467.5328052 | 490.6173158 | 0.000634 | 0.514225 | 0.000256 | 3.656184 | 1.428227 | 3.197968 |
| CNAG_035 | - | hypothetic | 3537.94148 | 3168.669432 | 2881.571437 | 2594.560879 | 2628.078071 | 2716.648573 | 0.000638 | -0.2877 | 0.000258 | -3.65443 | 0.819205 | 3.195396 |

| | | | | | | | | | | | | | |
|---|---|---|---|---|---|---|---|---|---|---|---|---|---|
| CNAG_06( | GAL1 | galactokin | 1041.898105 | 1221.822744 | 999.3920026 | 1472.158913 | 1334.480762 | 1337.662082 | 0.00064 | 0.330998 | 0.000259 | 3.653445 | 1.257883 | 3.193855 |
| CNAG_05( | - | hypothetic | 921.0189197 | 889.8357591 | 753.9944219 | 650.3260416 | 676.0897097 | 678.4318225 | 0.000647 | -0.37015 | 0.000262 | -3.65049 | 0.773702 | 3.188993 |
| CNAG_078 | - | hypothetic | 2.622434311 | 0.114690511 | 0.09896651 | 20.47260817 | 15.88243102 | 12.09711099 | 0.000648 | 4.537 | 0.000262 | 3.65025 | 23.21523 | 3.188714 |
| CNAG_053 | ITR3 | myo-inosi | 30.8126711 | 27.175487 | 37.45476194 | 75.48790112 | 71.99382032 | 65.29407112 | 0.000648 | 1.150704 | 0.000262 | 3.649958 | 2.220222 | 3.188352 |
| CNAG_042 | - | hypothetic | 1300.90512 | 1593.361643 | 1459.906037 | 1159.384606 | 1134.077003 | 1231.244358 | 0.000651 | -0.31874 | 0.000263 | -3.64882 | 0.801767 | 3.186687 |
| CNAG_056 | - | hypothetic | 928.6214275 | 1066.685346 | 923.6538183 | 1215.910344 | 1260.395309 | 1185.573051 | 0.000654 | 0.312687 | 0.000265 | 3.647483 | 1.242019 | 3.184561 |
| CNAG_05( | - | ribonuclea | 3719.152924 | 3701.998189 | 3622.706594 | 2930.720894 | 3160.313149 | 3337.177065 | 0.000659 | -0.24333 | 0.000267 | -3.64535 | 0.844793 | 3.181087 |
| CNAG_038 | - | vacuolar r | 840.7095344 | 881.2673334 | 954.3519357 | 757.9267224 | 702.8186292 | 656.2988571 | 0.000661 | -0.3541 | 0.000268 | -3.64456 | 0.782357 | 3.179889 |
| CNAG_01( | - | hypothetic | 533.7074095 | 629.1080607 | 723.9064642 | 501.8151671 | 474.2313282 | 402.1491633 | 0.000662 | -0.46806 | 0.000268 | -3.64404 | 0.722935 | 3.179135 |
| CNAG_037 | - | IQ domair | 4156.061896 | 3717.234936 | 3541.797621 | 4482.303534 | 4700.248428 | 4611.509956 | 0.000664 | 0.257443 | 0.000269 | 3.64313 | 1.195358 | 3.177732 |
| CNAG_01( | - | hypothetic | 75.37731273 | 115.4271768 | 124.861928 | 214.997542 | 194.4870647 | 147.6487761 | 0.000669 | 0.807351 | 0.000272 | 3.641003 | 1.749995 | 3.174405 |
| CNAG_125 | - | unspecifie | 11.10397486 | 22.79487408 | 20.4301103 | 61.8468212 | 61.05831187 | 33.35702133 | 0.000677 | 1.534081 | 0.000275 | 3.637971 | 2.896038 | 3.169553 |
| CNAG_073 | - | hypothetic | 8.789383522 | 13.18865751 | 29.92482028 | 56.17569998 | 42.4485786 | 57.26086539 | 0.000683 | 1.617439 | 0.000277 | 3.635564 | 3.068298 | 3.165627 |
| CNAG_04( | - | WD40 rep | 1744.773062 | 1718.584718 | 1709.596395 | 2121.074176 | 2179.016575 | 1959.412606 | 0.000687 | 0.26006 | 0.000279 | 3.633781 | 1.197529 | 3.162754 |
| CNAG_075 | - | hypothetic | 425.3021465 | 409.5573444 | 440.5220474 | 599.715974 | 586.7731573 | 521.7330736 | 0.000701 | 0.405705 | 0.000285 | 3.628515 | 1.324736 | 3.154282 |
| CNAG_02( | - | hypothetic | 1918.492783 | 1982.390042 | 1855.603423 | 1653.703831 | 1585.235532 | 1638.687553 | 0.000708 | -0.25411 | 0.000288 | -3.62592 | 0.838505 | 3.150179 |
| CNAG_008 | - | hypothetic | 329.214535 | 370.493726 | 374.1944702 | 257.2601172 | 268.9459357 | 252.1972791 | 0.000719 | -0.48061 | 0.000293 | -3.62152 | 0.716673 | 3.143174 |
| CNAG_125 | - | unspecifie | 21.8180942 | 63.73834208 | 76.85704909 | 114.9694948 | 128.1903897 | 120.6703358 | 0.000721 | 1.164846 | 0.000294 | 3.620758 | 2.242093 | 3.14216 |
| CNAG_03( | - | hypothetic | 1705.588613 | 2723.832547 | 2532.937593 | 3730.069089 | 3728.29926 | 3783.159058 | 0.000723 | 0.678317 | 0.000295 | 3.619821 | 1.600272 | 3.140718 |
| CNAG_054 | - | hypothetic | 120.8139933 | 196.0902118 | 122.8496707 | 249.6412167 | 223.5891944 | 247.2888527 | 0.000724 | 0.702202 | 0.000295 | 3.619612 | 1.626986 | 3.140498 |
| CNAG_04( | - | hypothetic | 236.2278387 | 261.9170448 | 254.982443 | 362.3098691 | 364.2601973 | 334.0586918 | 0.000725 | 0.482183 | 0.000296 | 3.619107 | 1.396855 | 3.13978 |
| CNAG_057 | - | hypothetic | 286.99721 | 298.28903 | 272.9418438 | 203.5690082 | 203.3510234 | 195.5885673 | 0.000726 | -0.52529 | 0.000296 | -3.61866 | 0.694821 | 3.139167 |
| CNAG_018 | - | hypothetic | 254.8286697 | 262.994767 | 237.9535904 | 370.1376811 | 331.3299725 | 365.0389633 | 0.000729 | 0.486594 | 0.000298 | 3.617378 | 1.401133 | 3.13714 |
| CNAG_043 | - | transcripti | 803.1161151 | 729.3692645 | 741.1285774 | 634.1544309 | 569.0733406 | 584.0960009 | 0.000737 | -0.36259 | 0.000301 | -3.61425 | 0.777769 | 3.13242 |
| CNAG_027 | - | hypothetic | 2459.588073 | 1264.214502 | 1144.652174 | 3811.888453 | 4688.83221 | 4617.647878 | 0.000742 | 1.411859 | 0.000303 | 3.612463 | 2.660798 | 3.129808 |
| CNAG_005 | PKR1 | cAMP-dep | 3531.18494 | 3489.768332 | 3610.95621 | 3017.999869 | 3156.899131 | 3059.694662 | 0.00075 | -0.21882 | 0.000307 | -3.60963 | 0.859269 | 3.125196 |
| CNAG_128 | - | unspecifie | 75.30177427 | 66.6253238 | 72.30986399 | 24.9437372 | 20.18356289 | 46.69527523 | 0.000761 | -1.2578 | 0.000312 | -3.60539 | 0.418181 | 3.11849 |
| CNAG_125 | - | unspecifie | 61.89560549 | 38.65573235 | 34.07709491 | 15.75021679 | 18.67629978 | 12.54218918 | 0.000769 | -1.58072 | 0.000315 | -3.60252 | 0.334315 | 3.113954 |
| CNAG_024 | - | hypothetic | 10.84808086 | 9.868937304 | 2.330552174 | 38.53233542 | 27.8946721 | 26.4595059 | 0.00077 | 2.10079 | 0.000316 | 3.602053 | 4.289443 | 3.113427 |
| CNAG_12( | - | unspecifie | 7.263348044 | 0.145383486 | 0.125374863 | 31.00767253 | 13.8377082 | 21.88403459 | 0.000787 | 3.195046 | 0.000323 | 3.596319 | 9.158083 | 3.104239 |
| CNAG_025 | - | cardiolipir | 118.3357565 | 149.8212047 | 105.774168 | 202.062102 | 198.6615782 | 197.3998062 | 0.000787 | 0.668286 | 0.000323 | 3.596363 | 1.589184 | 3.104239 |
| CNAG_065 | - | hypothetic | 61.27012223 | 92.74409623 | 71.62554619 | 117.7161861 | 157.5766883 | 132.984618 | 0.000794 | 0.843922 | 0.000326 | 3.593815 | 1.794923 | 3.100192 |
| CNAG_128 | - | unspecifie | 37.95979611 | 57.25278442 | 58.77336041 | 94.60397834 | 99.53661208 | 106.2215895 | 0.000795 | 0.961981 | 0.000326 | 3.593474 | 1.947982 | 3.099752 |
| CNAG_063 | - | transcripti | 1067.570238 | 1147.531944 | 1128.374349 | 1341.140795 | 1486.446945 | 1313.272344 | 0.00081 | 0.293759 | 0.000333 | 3.588375 | 1.22583 | 3.09177 |
| CNAG_03( | - | glycerate- | 103.1533452 | 125.0675809 | 94.0354319 | 175.5723234 | 164.0660315 | 189.5113601 | 0.000813 | 0.699296 | 0.000334 | 3.587219 | 1.623713 | 3.089975 |
| CNAG_017 | - | hypothetic | 401.9273241 | 401.9841134 | 374.4478377 | 490.4298999 | 601.6030769 | 508.3591566 | 0.000823 | 0.427848 | 0.000339 | 3.58375 | 1.345226 | 3.08446 |
| CNAG_008 | - | E3 ubiquit | 3660.797492 | 3622.287902 | 3709.073494 | 3279.608592 | 3036.917403 | 3197.342467 | 0.000831 | -0.22373 | 0.000342 | -3.58109 | 0.856351 | 3.08017 |
| CNAG_041 | - | O-methylt | 1342.660428 | 1268.071145 | 1224.256337 | 1111.293353 | 1059.100479 | 941.619966 | 0.000832 | -0.317 | 0.000343 | -3.58076 | 0.802738 | 3.079744 |
| CNAG_015 | - | hypothetic | 0.20211901 | 1.199526586 | 0.084333653 | 18.16909629 | 14.19161687 | 6.687975839 | 0.000832 | 5.128018 | 0.000343 | 3.580644 | 34.96932 | 3.079681 |
| CNAG_018 | - | hypothetic | 0.273935613 | 1.239650376 | 4.386833327 | 17.63847761 | 30.33363674 | 8.955910418 | 0.000838 | 3.442513 | 0.000345 | 3.578716 | 10.87175 | 3.076734 |
| CNAG_016 | - | CMGC/CD | 1157.30604 | 955.8343595 | 1024.956792 | 1454.318264 | 1318.124688 | 1249.999162 | 0.000842 | 0.342581 | 0.000347 | 3.577408 | 1.268023 | 3.074819 |
| CNAG_001 | AOX1 | alternative | 5957.090751 | 1873.788755 | 2752.375288 | 1029.379722 | 992.1379705 | 978.5926112 | 0.000846 | -1.83955 | 0.000349 | -3.57604 | 0.279409 | 3.072675 |
| CNAG_028 | - | 20S protea | 1211.172902 | 1157.289176 | 1339.491406 | 1523.320045 | 1555.357599 | 1510.837001 | 0.000846 | 0.292002 | 0.000349 | 3.57594 | 1.224338 | 3.072639 |
| CNAG_038 | - | hypothetic | 10.81115756 | 4.525803733 | 6.569207674 | 27.82753671 | 30.6669272 | 27.48529943 | 0.000849 | 2.052945 | 0.00035 | 3.574991 | 4.14952 | 3.071192 |
| CNAG_017 | - | DASH con | 686.5669195 | 739.0238125 | 775.1929156 | 538.9755268 | 635.6619809 | 539.7252891 | 0.000858 | -0.37711 | 0.000355 | -3.57166 | 0.769978 | 3.06643 |
| CNAG_007 | - | hypothetic | 9.638330373 | 3.442070397 | 5.499594812 | 33.00960979 | 23.55156979 | 23.12401139 | 0.000858 | 2.198837 | 0.000355 | 3.571719 | 4.591091 | 3.06643 |
| CNAG_13( | - | unspecifie | 53.18043305 | 57.85914463 | 43.63367014 | 25.26738157 | 17.70641727 | 18.07478773 | 0.000868 | -1.36503 | 0.000359 | -3.56868 | 0.388227 | 3.061628 |
| CNAG_003 | - | hypothetic | 700.9025181 | 796.2544862 | 658.1325119 | 851.50263 | 962.8372644 | 975.638686 | 0.00087 | 0.357624 | 0.00036 | 3.567791 | 1.281313 | 3.060277 |
| CNAG_053 | - | hypothetic | 651.7210219 | 909.1192039 | 811.4667577 | 596.6321083 | 611.3865268 | 594.0967469 | 0.000871 | -0.41064 | 0.00036 | -3.56765 | 0.75229 | 3.060172 |
| CNAG_012 | - | hypothetic | 667.6356111 | 586.0893392 | 626.9971926 | 459.5922236 | 496.6849528 | 500.7952724 | 0.000872 | -0.38471 | 0.000361 | -3.56712 | 0.76593 | 3.059426 |
| CNAG_018 | - | hypothetic | 362.7394145 | 315.6141254 | 319.8632989 | 243.2944782 | 251.7117455 | 214.5098748 | 0.000882 | -0.50829 | 0.000365 | -3.56391 | 0.703057 | 3.054321 |
| CNAG_028 | - | hypothetic | 296.889395 | 523.2855084 | 494.5921743 | 314.1518003 | 304.2784539 | 224.6202673 | 0.000888 | -0.65448 | 0.000368 | -3.56203 | 0.635304 | 3.051381 |
| CNAG_065 | - | hypothetic | 164.8487797 | 140.2408247 | 141.9832423 | 224.5290489 | 225.3702565 | 230.6911704 | 0.00089 | 0.593929 | 0.000369 | 3.561609 | 1.509352 | 3.050811 |
| CNAG_03( | - | hypothetic | 1207.147091 | 1065.547719 | 1080.279377 | 908.0254894 | 927.9929982 | 890.4856886 | 0.000896 | -0.31453 | 0.000371 | -3.55957 | 0.804114 | 3.047572 |
| CNAG_01( | - | 2,4-dienoy | 176.4068923 | 141.3041966 | 93.04795998 | 204.1149646 | 247.125815 | 241.701481 | 0.000899 | 0.739353 | 0.000373 | 3.558675 | 1.669427 | 3.046214 |
| CNAG_067 | - | hypothetic | 0.294107249 | 3.400168611 | 5.45443709 | 11.61923781 | 25.65862097 | 25.05868049 | 0.000906 | 2.916833 | 0.000376 | 3.556613 | 7.551865 | 3.042933 |
| CNAG_021 | - | hypothetic | 999.784243 | 1048.396943 | 958.838847 | 1204.158699 | 1214.587802 | 1302.05018 | 0.000928 | 0.293046 | 0.000385 | 3.550103 | 1.225225 | 3.032442 |
| CNAG_121 | - | unspecifie | 17.05445737 | 53.92111819 | 31.13707496 | 96.09778657 | 85.0175604 | 64.314886 | 0.000937 | 1.266786 | 0.000389 | 3.547557 | 2.406249 | 3.028371 |
| CNAG_04( | YFH701 | hypothetic | 58.51200369 | 29.37259597 | 33.28060943 | 92.61896791 | 75.21602771 | 95.05798634 | 0.000937 | 1.109471 | 0.000389 | 3.547138 | 2.157665 | 3.028062 |
| CNAG_068 | - | hypothetic | 255.0397128 | 358.8100957 | 300.856213 | 408.8193622 | 436.6930588 | 448.280967 | 0.000937 | 0.486997 | 0.000389 | 3.547163 | 1.401525 | 3.028062 |
| CNAG_001 | - | pre-mRNA | 27485.14151 | 25474.76783 | 23891.76107 | 21419.85997 | 22655.08299 | 22589.62614 | 0.000937 | -0.22069 | 0.000389 | -3.54729 | 0.858157 | 3.028062 |
| CNAG_036 | NIC1 | high-affini | 1679.118742 | 1618.283357 | 1812.869708 | 1453.675807 | 1434.317144 | 1385.6111 | 0.000938 | -0.27368 | 0.00039 | -3.5469 | 0.827208 | 3.027798 |
| CNAG_038 | SDP102 | MAP kinas | 238.4574476 | 239.2970519 | 203.8561176 | 309.9705497 | 305.4558851 | 379.3005791 | 0.000941 | 0.531462 | 0.000391 | 3.54584 | 1.445394 | 3.026305 |
| CNAG_062 | - | MFS trans | 44.94372409 | 80.83683946 | 42.8945665 | 117.4843 | 97.6437413 | 118.4140423 | 0.000943 | 0.987669 | 0.000392 | 3.545364 | 1.982978 | 3.025649 |
| CNAG_062 | HXT2 | Low-affini | 313.2888334 | 221.9013031 | 216.4635624 | 174.0166055 | 164.4969693 | 144.6082987 | 0.000943 | -0.65448 | 0.000392 | -3.54526 | 0.635305 | 3.0256 |
| CNAG_041 | - | ATP-deper | 1530.77841 | 1438.356721 | 1616.644364 | 1230.202543 | 1254.879148 | 1317.831746 | 0.000947 | -0.28578 | 0.000394 | -3.54401 | 0.820298 | 3.023672 |
| CNAG_068 | ETF1alpha | electron tr | 3127.01275 | 2961.804495 | 2795.165834 | 2655.442817 | 2433.988311 | 2449.136819 | 0.000956 | -0.25241 | 0.000398 | -3.54148 | 0.839495 | 3.019758 |
| CNAG_06( | - | tRNA (5-m | 972.5399408 | 884.5594177 | 1029.004825 | 729.6938775 | 734.6971792 | 827.1231719 | 0.000956 | -0.34884 | 0.000398 | -3.54123 | 0.785216 | 3.019485 |

| | | | | | | | | | | | | | | |
|---|---|---|---|---|---|---|---|---|---|---|---|---|---|---|
| CNAG_05! | FHL1 | Pre-rRNA- | 1032.019904 | 891.0345487 | 1015.168373 | 755.4010503 | 830.5587979 | 762.8301862 | 0.000964 | -0.34006 | 0.000402 | -3.5388 | 0.79001 | 3.01574 |
| CNAG_05! | - | hypothetic | 33.42777591 | 56.25494471 | 89.59785322 | 118.0614861 | 129.0027383 | 119.6167436 | 0.000987 | 1.025238 | 0.000412 | 3.532138 | 2.035296 | 3.005546 |
| CNAG_043 | MAN1 | mannose- | 1473.426665 | 1351.049238 | 1307.451055 | 1182.855768 | 1123.765443 | 1103.638029 | 0.000988 | -0.29309 | 0.000413 | -3.53187 | 0.816151 | 3.005228 |
| CNAG_063 | - | hypothetic | 1187.605541 | 1368.220704 | 1342.577914 | 990.4080777 | 1050.444221 | 1133.524668 | 0.000988 | -0.31116 | 0.000413 | -3.53171 | 0.805994 | 3.005093 |
| CNAG_049 | - | nuclear pr | 1553.20623 | 1466.469067 | 1419.539078 | 1785.583259 | 1782.163115 | 1802.797749 | 0.000995 | 0.259348 | 0.000416 | 3.529647 | 1.196937 | 3.00209 |
| CNAG_008 | - | 2-hydroxy | 441.1511537 | 431.9492963 | 420.1162975 | 306.2444244 | 336.2474467 | 330.9628936 | 0.000997 | -0.42591 | 0.000417 | -3.52914 | 0.744369 | 3.001378 |
| CNAG_052 | - | U3 small r | 704.1572784 | 727.2432111 | 989.3919378 | 632.5297398 | 629.5275939 | 513.1723868 | 0.001008 | -0.46379 | 0.000422 | -3.526 | 0.725081 | 2.996613 |
| CNAG_015 | - | plasma m | 4278.400774 | 4234.206695 | 3629.161773 | 3647.787437 | 3291.36893 | 3218.508668 | 0.001018 | -0.27264 | 0.000426 | -3.52332 | 0.827805 | 2.992355 |
| CNAG_021 | NHP6B02 | Non-histo | 740.2148633 | 757.3507374 | 754.979006 | 612.2946015 | 611.4179178 | 585.204646 | 0.001018 | -0.33137 | 0.000426 | -3.52318 | 0.79478 | 2.992236 |
| CNAG_043 | - | hypothetic | 469.2985232 | 467.5450634 | 525.6199181 | 373.2926957 | 364.6608405 | 374.2639413 | 0.001025 | -0.41044 | 0.00043 | -3.5212 | 0.752392 | 2.989255 |
| CNAG_052 | - | ste/ste20/ | 1112.852562 | 1189.424433 | 1321.201599 | 929.7223838 | 984.0385791 | 1023.645593 | 0.001027 | -0.31805 | 0.00043 | -3.52066 | 0.802151 | 2.988497 |
| CNAG_128 | - | unspecifie | 16.80430148 | 20.64100972 | 17.24196827 | 43.24442152 | 48.73081471 | 49.60223471 | 0.001031 | 1.38681 | 0.000432 | 3.519451 | 2.614999 | 2.986643 |
| CNAG_041 | FZC47 | hypothetic | 644.7886331 | 606.7193064 | 659.1310054 | 783.1173277 | 849.8476212 | 803.6475058 | 0.001038 | 0.335493 | 0.000435 | 3.517618 | 1.261809 | 2.983769 |
| CNAG_014 | MAT3 | mat3 pher | 46.02431537 | 57.25050634 | 52.41147667 | 108.9916717 | 90.78339901 | 96.34663812 | 0.001046 | 0.909114 | 0.000439 | 3.515603 | 1.877891 | 2.980599 |
| CNAG_039 | PHO13 | 4-nitrophe | 1827.424262 | 1770.19773 | 1960.045858 | 1480.789534 | 1631.897842 | 1548.780493 | 0.001056 | -0.26987 | 0.000443 | -3.51281 | 0.829394 | 2.97629 |
| CNAG_073 | - | cactin | 2301.529831 | 2325.039937 | 2284.305051 | 1956.415393 | 1951.844978 | 2040.536806 | 0.001064 | -0.23165 | 0.000447 | -3.51069 | 0.851658 | 2.973077 |
| CNAG_036 | - | hypothetic | 972.6889343 | 970.7457392 | 1196.379555 | 815.3553381 | 808.0282778 | 859.3614811 | 0.001064 | -0.35378 | 0.000447 | -3.51073 | 0.78253 | 2.973077 |
| CNAG_079 | - | hypothetic | 1112.933254 | 1038.722879 | 833.0747745 | 1282.081434 | 1430.456381 | 1196.717891 | 0.001074 | 0.374125 | 0.000451 | 3.508096 | 1.296053 | 2.969089 |
| CNAG_017 | - | pre-mRNA | 5535.049603 | 5514.258121 | 5520.993041 | 4712.327482 | 4992.438726 | 4903.651431 | 0.001083 | -0.19714 | 0.000455 | -3.50575 | 0.872278 | 2.965394 |
| CNAG_012 | - | thiosulfate | 103.009402 | 72.44428568 | 98.23224161 | 173.5732465 | 145.978306 | 148.5775463 | 0.001087 | 0.755011 | 0.000457 | 3.504574 | 1.687644 | 2.963595 |
| CNAG_028 | - | sterol 3-b | 738.2251614 | 786.57878 | 706.0971294 | 938.0805715 | 904.0572725 | 961.2493039 | 0.001089 | 0.315104 | 0.000458 | 3.503952 | 1.244102 | 2.962833 |
| CNAG_036 | - | L-lactate c | 2090.105496 | 1987.826448 | 2061.396962 | 1768.124976 | 1812.178633 | 1578.82828 | 0.001089 | -0.26661 | 0.000458 | -3.50401 | 0.831271 | 2.962833 |
| CNAG_044 | - | hypothetic | 414.9516646 | 520.4517376 | 491.711547 | 616.6300347 | 681.7251408 | 601.6257694 | 0.00109 | 0.398903 | 0.000459 | 3.50375 | 1.318505 | 2.962629 |
| CNAG_064 | - | mitochonc | 759.8451059 | 657.1955863 | 677.1440161 | 563.269489 | 569.1707648 | 496.4553407 | 0.001094 | -0.37841 | 0.000461 | -3.50266 | 0.769286 | 2.960983 |
| CNAG_013 | ISP3 | mannose- | 695.0845149 | 801.6423387 | 640.0192919 | 1095.756803 | 910.9843 | 839.2798505 | 0.001107 | 0.39856 | 0.000466 | 3.499523 | 1.318192 | 2.955991 |
| CNAG_056 | LCB3 | sphingosir | 1845.087508 | 2128.906755 | 2158.405291 | 1610.743488 | 1729.320512 | 1769.665512 | 0.001109 | -0.27811 | 0.000467 | -3.49879 | 0.824673 | 2.954929 |
| CNAG_029 | - | hypothetic | 1098.719509 | 1070.926697 | 1154.877929 | 955.6101594 | 864.9515298 | 903.7894463 | 0.001111 | -0.30272 | 0.000468 | -3.49841 | 0.810723 | 2.954433 |
| CNAG_129 | - | unspecifie | 0.140781169 | 0.068356703 | 0.059039394 | 9.880513031 | 6.120516952 | 9.746541068 | 0.001115 | 5.428056 | 0.00047 | 3.497312 | 43.05343 | 2.952769 |
| CNAG_040 | - | translatior | 712.4420462 | 690.7073768 | 563.2509232 | 786.9118134 | 889.5240049 | 912.3396084 | 0.001122 | 0.381604 | 0.000473 | 3.495633 | 1.30279 | 2.95016 |
| CNAG_061 | - | hypothetic | 207.1835373 | 287.7277755 | 264.5710956 | 366.1830221 | 367.6970247 | 352.8867813 | 0.001128 | 0.50324 | 0.000476 | 3.494124 | 1.417393 | 2.947831 |
| CNAG_031 | CMT1 | alpha-1,3- | 959.8880723 | 1161.304332 | 1031.209301 | 849.6514454 | 818.3785891 | 876.0080539 | 0.001129 | -0.3236 | 0.000476 | -3.4938 | 0.799075 | 2.947423 |
| CNAG_013 | - | hypothetic | 304.9564065 | 273.810644 | 272.071132 | 401.188097 | 432.4983257 | 354.1072654 | 0.001138 | 0.467667 | 0.00048 | 3.491467 | 1.382871 | 2.943762 |
| CNAG_003 | - | survival fa | 4800.484267 | 4534.893602 | 4716.878764 | 4203.435576 | 4028.038571 | 4042.190286 | 0.001138 | -0.21058 | 0.000481 | -3.49134 | 0.864189 | 2.943684 |
| CNAG_062 | DRP1 | Putative tr | 202.1489411 | 184.3549054 | 127.1437537 | 257.9706845 | 275.4846939 | 264.0131597 | 0.00115 | 0.618522 | 0.000486 | 3.488383 | 1.535301 | 2.939126 |
| CNAG_120 | - | unspecifie | 34.14246873 | 22.86368783 | 20.4835381 | 64.51007306 | 43.95523363 | 83.67027624 | 0.001156 | 1.306323 | 0.000489 | 3.486918 | 2.473104 | 2.936873 |
| CNAG_078 | - | hypothetic | 5.023222562 | 5.54599319 | 1.224407074 | 31.73947477 | 14.70222771 | 20.84978984 | 0.001184 | 2.549052 | 0.000501 | 3.480078 | 5.852498 | 2.926527 |
| CNAG_059 | - | Poly(3-hyc | 104.2750095 | 112.1773711 | 85.52130206 | 213.6162615 | 132.7851556 | 172.9356171 | 0.001185 | 0.766135 | 0.000502 | 3.479797 | 1.700708 | 2.926198 |
| CNAG_017 | - | hypothetic | 408.9771047 | 454.7318448 | 390.4490011 | 563.8054445 | 591.9974543 | 512.8479805 | 0.001217 | 0.397257 | 0.000516 | 3.47246 | 1.317002 | 2.914691 |
| CNAG_028 | KCS1 | inositol he | 1543.883044 | 1421.241019 | 1432.335086 | 1656.976818 | 1798.556107 | 2019.137535 | 0.00122 | 0.300698 | 0.000517 | 3.471692 | 1.23174 | 2.913575 |
| CNAG_000 | CSE4 | Histone H | 1022.786216 | 995.4944516 | 1020.532236 | 818.5019286 | 866.780959 | 817.2086534 | 0.001231 | -0.29558 | 0.000522 | -3.46927 | 0.814742 | 2.909784 |
| CNAG_075 | - | hypothetic | 274.2214823 | 261.7614315 | 318.6702611 | 213.7827376 | 204.226369 | 175.6746929 | 0.001245 | -0.5427 | 0.000528 | -3.46593 | 0.686486 | 2.90489 |
| CNAG_029 | - | RNA exon | 441.9733863 | 321.0824828 | 372.097645 | 241.5356979 | 253.3184518 | 300.9283434 | 0.001245 | -0.53056 | 0.000528 | -3.46598 | 0.692287 | 2.90489 |
| CNAG_021 | TUP1 | glucose re | 1399.251016 | 1645.26303 | 1512.30173 | 1886.046417 | 1881.39616 | 1776.206676 | 0.001248 | 0.268276 | 0.00053 | 3.465139 | 1.204368 | 2.903733 |
| CNAG_030 | - | hypothetic | 333.1513753 | 340.6148083 | 402.0912257 | 474.2135417 | 462.4406078 | 532.6092091 | 0.001264 | 0.43433 | 0.000537 | 3.46169 | 1.351284 | 2.898415 |
| CNAG_021 | - | hypothetic | 347.1015663 | 594.4239934 | 555.4131603 | 352.2190396 | 364.6855154 | 302.2487109 | 0.001265 | -0.56888 | 0.000538 | -3.46118 | 0.674142 | 2.897839 |
| CNAG_014 | - | hypothetic | 476.3199016 | 521.3417716 | 495.8230083 | 414.6937214 | 368.0964449 | 358.7763333 | 0.001285 | -0.40308 | 0.000546 | -3.45693 | 0.756243 | 2.891116 |
| CNAG_026 | - | mitochonc | 435.3139453 | 406.1324304 | 441.3987304 | 321.9134931 | 286.0693517 | 347.559432 | 0.001286 | -0.44166 | 0.000547 | -3.45654 | 0.736288 | 2.890617 |
| CNAG_039 | SOG2 | ram signa | 786.0838568 | 832.9025942 | 717.8443817 | 941.0830776 | 966.1881631 | 1042.248873 | 0.001287 | 0.322012 | 0.000547 | 3.456424 | 1.250073 | 2.890548 |
| CNAG_079 | FZC29 | hypothetic | 1594.199424 | 1688.399207 | 1694.645955 | 1945.074802 | 1941.701345 | 2082.514481 | 0.001287 | 0.247303 | 0.000548 | 3.456165 | 1.186986 | 2.890255 |
| CNAG_033 | - | hypothetic | 929.7955425 | 1063.459688 | 946.0400798 | 1144.148228 | 1212.884551 | 1325.318549 | 0.001288 | 0.31073 | 0.000548 | 3.455956 | 1.240336 | 2.890043 |
| CNAG_003 | YMR1 | myotubula | 677.6489385 | 828.5834705 | 756.2004505 | 883.3870681 | 1081.988583 | 961.2872474 | 0.001289 | 0.356824 | 0.000549 | 3.455741 | 1.280603 | 2.889821 |
| CNAG_065 | CET1 | mRNA-cap | 715.6204592 | 624.9228326 | 804.9918976 | 545.2828599 | 494.8258585 | 592.8965161 | 0.001296 | -0.40964 | 0.000552 | -3.45417 | 0.752809 | 2.887413 |
| CNAG_005 | - | hypothetic | 95.07565861 | 143.3183761 | 87.66718285 | 192.8716186 | 158.8052355 | 198.3674922 | 0.001299 | 0.740745 | 0.000553 | 3.453552 | 1.671039 | 2.886544 |
| CNAG_000 | - | 26S protea | 5669.295641 | 5609.057207 | 5326.922986 | 5049.468395 | 4781.05789 | 4719.401692 | 0.001305 | -0.20583 | 0.000556 | -3.45204 | 0.867039 | 2.884233 |
| CNAG_075 | - | ribonuclea | 566.3998545 | 631.3088556 | 769.758933 | 469.6886503 | 507.0198127 | 510.8002887 | 0.001325 | -0.41899 | 0.000565 | -3.44772 | 0.747949 | 2.877657 |
| CNAG_012 | - | hypothetic | 4.029674621 | 9.901502533 | 16.10038 | 36.55279163 | 27.75027914 | 41.67671942 | 0.001325 | 1.828471 | 0.000565 | 3.447725 | 3.551605 | 2.877657 |
| CNAG_038 | HRT1 | RING-box | 896.7920876 | 976.0639835 | 990.6468019 | 802.2385953 | 776.155294 | 762.8162179 | 0.001333 | -0.30549 | 0.000569 | -3.44606 | 0.809167 | 2.87524 |
| CNAG_054 | - | hypothetic | 908.4529753 | 1041.730607 | 934.1706054 | 778.0472534 | 743.3261403 | 816.0435354 | 0.001338 | -0.31742 | 0.000571 | -3.44484 | 0.802504 | 2.873402 |
| CNAG_024 | - | hypothetic | 491.8831886 | 489.2773253 | 521.5676648 | 758.7075273 | 644.4627897 | 592.8179218 | 0.001341 | 0.394393 | 0.000573 | 3.444175 | 1.314389 | 2.872459 |
| CNAG_075 | - | hypothetic | 510.6828927 | 652.945412 | 568.5168639 | 806.7575094 | 739.3518204 | 718.1993891 | 0.001365 | 0.373226 | 0.000584 | 3.439167 | 1.295246 | 2.864791 |
| CNAG_002 | YAP1 | hypothetic | 87.87328641 | 74.54739746 | 83.32026894 | 144.5686301 | 120.856042 | 151.7396486 | 0.001371 | 0.750753 | 0.000586 | 3.437889 | 1.682671 | 2.862865 |
| CNAG_014 | - | hypothetic | 1178.401225 | 1017.251151 | 1210.463801 | 1356.678373 | 1460.502384 | 1438.638597 | 0.001382 | 0.305164 | 0.000591 | 3.435619 | 1.235559 | 2.85935 |
| CNAG_029 | - | transcripti | 3327.034949 | 3320.69841 | 3090.667652 | 3709.424603 | 3781.553771 | 3957.599062 | 0.001385 | 0.218305 | 0.000592 | 3.435035 | 1.163366 | 2.858538 |
| CNAG_041 | - | hypothetic | 233.9965513 | 93.8166032 | 101.343547 | 71.00541369 | 68.61399717 | 70.23820416 | 0.001391 | -1.0497 | 0.000595 | -3.43374 | 0.483067 | 2.856708 |
| CNAG_007 | MSH2 | DNA misn | 2036.675314 | 2006.242955 | 1920.762174 | 2310.867321 | 2364.390656 | 2388.965112 | 0.001391 | 0.229039 | 0.000595 | 3.433769 | 1.172054 | 2.856708 |
| CNAG_051 | - | DNA-3-me | 553.3316488 | 552.6394125 | 557.6767813 | 452.7723494 | 409.487255 | 437.5302017 | 0.001396 | -0.37178 | 0.000598 | -3.43272 | 0.772827 | 2.855198 |
| CNAG_066 | - | mitofusin | 2204.469139 | 2064.304534 | 1942.006132 | 1713.555926 | 1825.107952 | 1710.878388 | 0.001401 | -0.25835 | 0.0006 | -3.43162 | 0.836041 | 2.853559 |
| CNAG_004 | OST2 | oligosacch | 935.2172314 | 1053.548435 | 845.7062774 | 725.0391977 | 782.2263137 | 758.362342 | 0.001408 | -0.33742 | 0.000603 | -3.43005 | 0.791457 | 2.851302 |

| | | | | | | | | | | | | | | |
|---|---|---|---|---|---|---|---|---|---|---|---|---|---|---|
| CNAG_043 | - | dolichyl-d | 2524.484435 | 2524.333848 | 2241.691636 | 2106.679361 | 2121.814697 | 1954.030388 | 0.001408 | -0.25308 | 0.000603 | -3.43008 | 0.839101 | 2.851302 |
| CNAG_079 | - | hypothetic | 3.768266454 | 0.117365824 | 2.256057622 | 16.13802675 | 13.12478376 | 18.50082018 | 0.001413 | 3.214785 | 0.000606 | 3.429115 | 9.28425 | 2.849925 |
| CNAG_029 | - | ubiquitin- | 2078.340568 | 1787.468088 | 1977.1407 | 1639.657886 | 1633.562396 | 1626.492776 | 0.001418 | -0.2702 | 0.000608 | -3.42804 | 0.829205 | 2.84832 |
| CNAG_022 | - | ubiquitin- | 518.2511543 | 474.0382555 | 535.2429781 | 393.5337089 | 401.8023237 | 382.0599293 | 0.00142 | -0.39253 | 0.000609 | -3.42759 | 0.761795 | 2.847737 |
| CNAG_044 | RSM18 | small subu | 927.003836 | 858.6891792 | 959.7054524 | 741.4775458 | 788.3103082 | 675.1748925 | 0.001431 | -0.33199 | 0.000614 | -3.42543 | 0.79444 | 2.844399 |
| CNAG_063 | - | hypothetic | 918.6939105 | 752.0654409 | 885.0565927 | 664.3684129 | 692.5031003 | 668.4560314 | 0.001456 | -0.35229 | 0.000625 | -3.42055 | 0.783338 | 2.836733 |
| CNAG_048 | - | hypothetic | 24.89011535 | 21.75256967 | 21.50639935 | 43.77299888 | 52.07935547 | 66.06076064 | 0.001467 | 1.271188 | 0.00063 | 3.418354 | 2.413602 | 2.833469 |
| CNAG_024 | - | hypothetic | 871.3572782 | 1350.82375 | 1213.483188 | 747.4037581 | 826.96385 | 966.9906393 | 0.001483 | -0.44891 | 0.000637 | -3.41546 | 0.732598 | 2.828975 |
| CNAG_074 | - | hypothetic | 93.78244965 | 74.59608919 | 108.8595445 | 196.4837329 | 148.566752 | 132.0730342 | 0.001494 | 0.775283 | 0.000642 | 3.413147 | 1.711526 | 2.825537 |
| CNAG_041 | - | hypothetic | 9.821558305 | 12.05208705 | 14.01534407 | 41.32111121 | 48.20875539 | 23.36646625 | 0.001503 | 1.662516 | 0.000646 | 3.41136 | 3.165681 | 2.822937 |
| CNAG_129 | - | unspecifie | 53.17403966 | 55.22972773 | 107.7058387 | 114.5526856 | 141.8665137 | 159.3714549 | 0.001503 | 0.929461 | 0.000646 | 3.411424 | 1.904564 | 2.822937 |
| CNAG_128 | - | unspecifie | 72.66096528 | 61.60352411 | 57.77105092 | 132.4417332 | 98.43383579 | 113.0030256 | 0.001507 | 0.836804 | 0.000648 | 3.410626 | 1.78609 | 2.82189 |
| CNAG_068 | - | hypothetic | 2.808675658 | 3.45605008 | 13.94498874 | 19.48870966 | 46.86896517 | 23.19145057 | 0.001509 | 2.272048 | 0.00065 | 3.409984 | 4.830084 | 2.821238 |
| CNAG_002 | - | hypothetic | 158.0716231 | 134.4952415 | 105.5804578 | 72.60111121 | 86.04460125 | 73.53687641 | 0.001509 | -0.79707 | 0.00065 | -3.40999 | 0.575517 | 2.821238 |
| CNAG_073 | - | cystinosin | 104.5165881 | 174.5738039 | 129.2061793 | 182.904438 | 218.4833527 | 273.676693 | 0.001527 | 0.712633 | 0.000658 | 3.406665 | 1.638792 | 2.816201 |
| CNAG_127 | - | unspecifie | 112.5644524 | 167.0045312 | 96.21141335 | 200.4254 | 229.9270146 | 188.6011862 | 0.001537 | 0.704869 | 0.000662 | 3.404763 | 1.629997 | 2.813299 |
| CNAG_062 | - | YeeE/YedE | 1432.758203 | 1441.565541 | 1565.458744 | 1192.83762 | 1278.222945 | 1254.574394 | 0.001537 | -0.26843 | 0.000663 | -3.40462 | 0.830225 | 2.813198 |
| CNAG_022 | - | exportin-T | 2540.81169 | 2370.311219 | 2309.921569 | 2042.028637 | 2115.778838 | 2033.920624 | 0.001547 | -0.2374 | 0.000667 | -3.40271 | 0.848273 | 2.81041 |
| CNAG_023 | - | casein kin | 1109.342166 | 1130.201569 | 1361.69477 | 986.6594533 | 980.5929077 | 943.7831612 | 0.00156 | -0.32293 | 0.000673 | -3.40047 | 0.799443 | 2.806971 |
| CNAG_046 | - | hypothetic | 210.4036463 | 203.7495143 | 166.5500213 | 282.8590432 | 338.6496831 | 246.4449377 | 0.001561 | 0.565223 | 0.000674 | 3.4 | 1.479616 | 2.806468 |
| CNAG_076 | - | zinc finger | 547.7946202 | 510.8031912 | 477.8908579 | 664.9341043 | 652.2362313 | 663.7426985 | 0.001561 | 0.352103 | 0.000674 | 3.400046 | 1.27642 | 2.806468 |
| CNAG_000 | ZFC6 | hypothetic | 573.2536866 | 605.4185616 | 607.7909492 | 461.2313082 | 501.9254517 | 442.0260933 | 0.001564 | -0.3616 | 0.000675 | -3.39952 | 0.778299 | 2.805831 |
| CNAG_027 | - | aryl-alcoh | 1536.72853 | 1715.163658 | 1599.638921 | 1368.03466 | 1374.80454 | 1370.036399 | 0.001571 | -0.25291 | 0.000678 | -3.39826 | 0.839201 | 2.803952 |
| CNAG_041 | PTP3 | hypothetic | 3250.907041 | 3197.78012 | 3206.789557 | 2822.699137 | 2867.936504 | 2768.83982 | 0.001585 | -0.20619 | 0.000684 | -3.39573 | 0.866823 | 2.800054 |
| CNAG_053 | - | hypothetic | 1174.552528 | 1118.321966 | 1141.04511 | 833.8982426 | 944.373769 | 1015.829314 | 0.001587 | -0.31357 | 0.000685 | -3.39532 | 0.804647 | 2.799534 |
| CNAG_005 | - | DNA ligas | 1424.513545 | 1473.831225 | 1344.781342 | 1150.046885 | 1185.892277 | 1229.016847 | 0.00159 | -0.26589 | 0.000687 | -3.39472 | 0.831687 | 2.798702 |
| CNAG_026 | - | hypothetic | 16.82410202 | 25.97542671 | 14.07665539 | 60.31497731 | 46.02046058 | 39.85932342 | 0.001596 | 1.385036 | 0.00069 | 3.393438 | 2.611785 | 2.796913 |
| CNAG_020 | - | hypothetic | 699.0847236 | 607.6150516 | 582.2642082 | 494.7286091 | 490.6244436 | 489.7208668 | 0.001606 | -0.3738 | 0.000695 | -3.39162 | 0.771748 | 2.794269 |
| CNAG_007 | - | hypothetic | 249.0766261 | 269.4749764 | 281.6216023 | 414.6789261 | 339.920417 | 361.7829129 | 0.00162 | 0.464575 | 0.000701 | 3.389081 | 1.379911 | 2.790424 |
| CNAG_001 | - | hypothetic | 321.4038472 | 342.7109671 | 327.4997621 | 418.0988183 | 447.0049099 | 460.5200799 | 0.001625 | 0.40551 | 0.000704 | 3.388204 | 1.324557 | 2.789231 |
| CNAG_126 | - | unspecifie | 0.310885427 | 2.339653457 | 8.621025352 | 17.40083701 | 31.02413701 | 17.6238367 | 0.001637 | 2.684206 | 0.000709 | 3.385985 | 6.427269 | 2.785842 |
| CNAG_005 | - | dynein lig | 1011.25404 | 1125.818461 | 1119.695741 | 889.3488812 | 876.1981396 | 924.8443652 | 0.001647 | -0.29039 | 0.000714 | -3.38433 | 0.81768 | 2.783341 |
| CNAG_120 | - | unspecifie | 2.814776281 | 9.862689628 | 11.84794662 | 33.5835844 | 27.0244801 | 27.51756254 | 0.001651 | 1.970346 | 0.000715 | 3.383573 | 3.91862 | 2.78227 |
| CNAG_126 | - | unspecifie | 34.55980357 | 73.36174842 | 67.30415331 | 109.451511 | 100.2331045 | 130.5201154 | 0.001651 | 0.949454 | 0.000716 | 3.383474 | 1.931142 | 2.782237 |
| CNAG_050 | - | hypothetic | 271.1034596 | 269.4473414 | 221.9878011 | 370.1263677 | 335.655499 | 361.7257249 | 0.001663 | 0.471294 | 0.000721 | 3.381385 | 1.386353 | 2.779055 |
| CNAG_122 | - | unspecifie | 21.58180645 | 32.48954304 | 34.25471932 | 73.27302323 | 67.70220693 | 54.29760113 | 0.001667 | 1.142504 | 0.000723 | 3.380673 | 2.207638 | 2.778053 |
| CNAG_052 | - | sentrin/su | 761.280544 | 806.9041721 | 799.7671331 | 602.8304099 | 664.932114 | 650.6570737 | 0.001681 | -0.31944 | 0.00073 | -3.37822 | 0.801378 | 2.774305 |
| CNAG_066 | - | hypothetic | 3.928459865 | 11.93809287 | 2.3102008 | 22.19212128 | 27.20312483 | 26.31474353 | 0.001688 | 2.217966 | 0.000733 | 3.377077 | 4.652371 | 2.772616 |
| CNAG_125 | - | unspecifie | 34.53254473 | 71.19202516 | 60.91957085 | 101.6227588 | 111.6934581 | 106.3037328 | 0.001691 | 0.933419 | 0.000734 | 3.376519 | 1.909797 | 2.771911 |
| CNAG_014 | - | vacuolar p | 2414.750873 | 2397.219307 | 2238.4759 | 2072.454188 | 2025.182396 | 1978.418363 | 0.001699 | -0.22974 | 0.000738 | -3.37498 | 0.852787 | 2.76968 |
| CNAG_026 | - | ribonucleo | 976.3769121 | 899.7715085 | 949.2136015 | 1181.688779 | 1246.63841 | 1082.389381 | 0.001734 | 0.297551 | 0.000754 | 3.369162 | 1.229056 | 2.760864 |
| CNAG_017 | STE7 | STE/STE7/ | 432.5386032 | 615.2241852 | 542.9067795 | 692.1419338 | 716.1291688 | 710.3506752 | 0.001741 | 0.400678 | 0.000757 | 3.368104 | 1.320128 | 2.759319 |
| CNAG_040 | - | AFG3 fam | 8455.541555 | 7980.520317 | 8193.524146 | 6760.725939 | 7326.516221 | 7491.405511 | 0.001743 | -0.20633 | 0.000758 | -3.36758 | 0.866739 | 2.758615 |
| CNAG_018 | - | vacuolar p | 3800.854424 | 3684.763191 | 3463.836017 | 3150.334515 | 3110.265303 | 3260.600081 | 0.001764 | -0.21673 | 0.000768 | -3.36424 | 0.860513 | 2.753598 |
| CNAG_012 | - | hypothetic | 695.5913354 | 533.3710061 | 677.0661624 | 490.8236832 | 487.1668802 | 478.6387444 | 0.001764 | -0.40522 | 0.000768 | -3.36413 | 0.755123 | 2.753544 |
| CNAG_003 | - | hypothetic | 334.2684876 | 362.1046341 | 356.2789901 | 474.3121067 | 453.8452276 | 466.1246271 | 0.001784 | 0.390136 | 0.000777 | 3.360732 | 1.310517 | 2.748571 |
| CNAG_066 | - | protein ph | 5433.448384 | 5398.971907 | 5317.311285 | 4718.600045 | 4897.561008 | 4726.04029 | 0.00181 | -0.18653 | 0.000789 | -3.35671 | 0.878719 | 2.742374 |
| CNAG_007 | - | hypothetic | 525.3642559 | 565.5308458 | 535.2969063 | 433.2701392 | 429.398161 | 416.4629189 | 0.001813 | -0.36151 | 0.000791 | -3.35601 | 0.778347 | 2.741642 |
| CNAG_080 | - | hypothetic | 3299.905467 | 3175.151319 | 3111.889025 | 2742.49877 | 2718.660364 | 2879.806097 | 0.001813 | -0.2162 | 0.000791 | -3.35602 | 0.860831 | 2.741642 |
| CNAG_010 | - | cohesin co | 768.0275758 | 674.4254003 | 687.8078524 | 623.370779 | 542.3600489 | 482.0567991 | 0.001813 | -0.38615 | 0.00079 | -3.35615 | 0.765171 | 2.741642 |
| CNAG_043 | - | hypothetic | 728.6541777 | 989.858726 | 748.6228239 | 511.52385 | 740.9823038 | 555.3148936 | 0.001816 | -0.46297 | 0.000792 | -3.35548 | 0.725492 | 2.740933 |
| CNAG_024 | - | response | 231.462429 | 205.9653835 | 241.0961331 | 336.6929514 | 339.2913219 | 287.4898025 | 0.001821 | 0.490953 | 0.000795 | 3.354615 | 1.405373 | 2.73969 |
| CNAG_057 | - | hypothetic | 318.9430679 | 293.1807465 | 257.1765284 | 368.2307978 | 472.292081 | 381.8049744 | 0.001825 | 0.476562 | 0.000797 | 3.353871 | 1.391424 | 2.738766 |
| CNAG_027 | - | galactinol | 792.1692772 | 566.714354 | 547.1004251 | 451.7972785 | 455.1986455 | 493.0295466 | 0.001825 | -0.46184 | 0.000797 | -3.35392 | 0.726059 | 2.738766 |
| CNAG_000 | - | hypothetic | 89.22073312 | 101.4443902 | 118.4517413 | 155.9426705 | 192.0086713 | 153.0834624 | 0.001831 | 0.683617 | 0.0008 | 3.352828 | 1.606161 | 2.73725 |
| CNAG_000 | - | hypothetic | 4863.581536 | 4815.004851 | 4776.615451 | 4387.130403 | 4177.26452 | 4216.477497 | 0.00184 | -0.19283 | 0.000804 | -3.35139 | 0.87489 | 2.735108 |
| CNAG_073 | - | hypothetic | 387.9174514 | 414.8722227 | 353.1305647 | 522.5742665 | 477.9350051 | 532.6833558 | 0.001841 | 0.393138 | 0.000804 | 3.351256 | 1.313247 | 2.735027 |
| CNAG_010 | - | exosome | 901.2779526 | 836.0471199 | 861.6381299 | 706.4514864 | 728.7544354 | 690.6577895 | 0.001849 | -0.30621 | 0.000808 | -3.3499 | 0.808762 | 2.733025 |
| CNAG_023 | - | pre-mRNA | 4749.096367 | 4488.560693 | 4617.703553 | 3835.15202 | 4057.392333 | 4201.999584 | 0.001849 | -0.21161 | 0.000809 | -3.34978 | 0.86357 | 2.732961 |
| CNAG_049 | CAT1 | Catalase 1 | 4827.024077 | 5042.122561 | 3624.907866 | 3443.721973 | 3045.435813 | 2715.753707 | 0.001884 | -0.56674 | 0.000825 | -3.34441 | 0.675139 | 2.724901 |
| CNAG_019 | - | hypothetic | 51.7843877 | 54.04310795 | 54.53343689 | 85.03043284 | 93.40433248 | 116.0875184 | 0.001927 | 0.871947 | 0.000844 | 3.338006 | 1.830132 | 2.71513 |
| CNAG_032 | - | tRNA (ura | 310.8190507 | 283.5087626 | 299.770131 | 394.8302431 | 450.6208177 | 375.1766655 | 0.001931 | 0.43476 | 0.000846 | 3.337279 | 1.351686 | 2.714115 |
| CNAG_053 | - | hypothetic | 8.51937899 | 9.838552733 | 1.244932033 | 30.44227055 | 29.83225251 | 20.98779355 | 0.001936 | 2.119639 | 0.000848 | 3.336533 | 4.345853 | 2.713071 |
| CNAG_059 | - | hypothetic | 13.38418126 | 23.84402498 | 19.36185192 | 43.19014675 | 38.93562035 | 63.7349706 | 0.001937 | 1.375563 | 0.000849 | 3.336271 | 2.594691 | 2.712782 |
| CNAG_061 | - | sterility pr | 2243.452492 | 2417.799886 | 2341.986457 | 2772.459801 | 2682.633261 | 2750.894277 | 0.00196 | 0.213876 | 0.000859 | 3.333016 | 1.1598 | 2.70782 |
| CNAG_035 | - | ribosomal | 1729.348716 | 1684.00587 | 1912.038183 | 1395.933691 | 1428.228918 | 1605.315621 | 0.001967 | -0.28149 | 0.000862 | -3.33196 | 0.82274 | 2.706288 |
| CNAG_045 | - | hypothetic | 28.69656838 | 52.9503147 | 61.92924467 | 79.55762574 | 100.4992254 | 106.1510446 | 0.001971 | 1.001352 | 0.000865 | 3.331173 | 2.001875 | 2.705262 |
| CNAG_023 | - | hypothetic | 660.7982338 | 713.1234316 | 631.3132812 | 507.8831348 | 543.2770199 | 549.6280418 | 0.001975 | -0.34056 | 0.000867 | -3.33053 | 0.789736 | 2.704429 |

| | | | | | | | | | | | | | | |
|---|---|---|---|---|---|---|---|---|---|---|---|---|---|---|
| CNAG_00( | - | hypothetic | 429.62272 | 442.7489935 | 482.9718094 | 334.3155436 | 343.1101266 | 360.9090285 | 0.001987 | -0.39894 | 0.000872 | -3.32882 | 0.758414 | 2.701879 |
| CNAG_00! | - | hypothetic | 883.65364 | 753.1094038 | 796.5880559 | 656.6424084 | 644.165951 | 655.1134773 | 0.00199 | -0.33081 | 0.000874 | -3.32833 | 0.79509 | 2.701232 |
| CNAG_00( | - | hypothetic | 769.6455794 | 746.7261678 | 606.9721368 | 860.9037246 | 899.7880214 | 983.3776196 | 0.002001 | 0.355454 | 0.000879 | 3.326553 | 1.279389 | 2.698706 |
| CNAG_02( | - | peroxin-1( | 76.10534643 | 64.8123017 | 49.2778513 | 123.7077755 | 110.6843949 | 106.4021103 | 0.002001 | 0.827361 | 0.000879 | 3.326601 | 1.774437 | 2.698706 |
| CNAG_03( | - | transketol | 2931.084628 | 2755.052069 | 2218.384279 | 3420.14384 | 3255.435075 | 3141.69489 | 0.002009 | 0.29728 | 0.000883 | 3.325345 | 1.228825 | 2.696946 |
| CNAG_057 | UBC6 | ubiquitin- | 2489.338008 | 2104.227431 | 2228.849688 | 1912.834599 | 1835.355872 | 1980.593693 | 0.002029 | -0.26806 | 0.000892 | -3.32249 | 0.830436 | 2.692615 |
| CNAG_00 | - | hypothetic | 60.13959627 | 115.2597663 | 55.70489001 | 127.1457188 | 138.3107446 | 161.6295179 | 0.00203 | 0.877277 | 0.000893 | 3.322359 | 1.836904 | 2.692536 |
| CNAG_03( | - | hypothetic | 196.4789571 | 186.5595226 | 215.4962622 | 265.5740801 | 311.748544 | 279.6125416 | 0.002043 | 0.50249 | 0.000899 | 3.320488 | 1.416657 | 2.689746 |
| CNAG_01( | - | MFS trans | 415.8760329 | 409.5064228 | 358.4651149 | 540.5118113 | 505.5904486 | 512.7736977 | 0.002052 | 0.382433 | 0.000903 | 3.319158 | 1.303539 | 2.687917 |
| CNAG_13( | - | unspecifie | 26.01427217 | 15.3413065 | 23.61438536 | 64.9631176 | 49.4514135 | 44.28675986 | 0.002062 | 1.277739 | 0.000908 | 3.317616 | 2.424588 | 2.685641 |
| CNAG_12( | - | unspecifie | 0.205563092 | 0.099339495 | 2.23379342 | 8.65991467 | 15.09647793 | 16.26367751 | 0.002073 | 4.269195 | 0.000913 | 3.316143 | 19.28216 | 2.683471 |
| CNAG_05 | - | amino aci | 7.479946805 | 10.95744488 | 13.98505587 | 33.31649752 | 34.94877168 | 29.77521693 | 0.002074 | 1.663133 | 0.000914 | 3.315878 | 3.167035 | 2.683179 |
| CNAG_07! | - | hypothetic | 270.0634736 | 296.3850597 | 290.1645284 | 396.5148004 | 364.0971111 | 400.5744833 | 0.002079 | 0.423638 | 0.000916 | 3.315084 | 1.341306 | 2.682066 |
| CNAG_03 | - | yippee-lik | 56.34704412 | 32.61695787 | 52.39868243 | 94.7032636 | 132.0629695 | 66.6900371 | 0.002089 | 1.042359 | 0.000921 | 3.313549 | 2.059593 | 2.680042 |
| CNAG_04! | - | hypothetic | 1152.329796 | 1017.089578 | 1122.899545 | 917.4097285 | 861.5059552 | 928.1798678 | 0.00209 | -0.29805 | 0.000922 | -3.31341 | 0.813351 | 2.679951 |
| CNAG_05 | - | DNA excis | 500.0476707 | 517.2622426 | 531.1605464 | 660.2375192 | 624.5647931 | 696.9893224 | 0.002095 | 0.340079 | 0.000924 | 3.312665 | 1.265826 | 2.678911 |
| CNAG_04 | - | TPR repea | 1224.846098 | 1209.908722 | 1265.805922 | 963.9730172 | 1111.808397 | 1002.603423 | 0.002141 | -0.2812 | 0.000945 | -3.3065 | 0.822904 | 2.669468 |
| CNAG_04 | - | mitochonc | 1507.532171 | 1580.538184 | 1675.307148 | 1346.251571 | 1407.636992 | 1263.516567 | 0.002168 | -0.2611 | 0.000957 | -3.30281 | 0.834451 | 2.663987 |
| CNAG_08( | - | hypothetic | 54.11973208 | 40.13834099 | 70.44741965 | 103.2869941 | 103.8570384 | 100.7844442 | 0.002168 | 0.88333 | 0.000957 | 3.302845 | 1.844628 | 2.663987 |
| CNAG_07 | - | 2-hydroxy | 1.443913806 | 2.281120478 | 1.180207904 | 10.29889362 | 14.12546417 | 17.36464254 | 0.00217 | 3.343795 | 0.000959 | 3.302382 | 10.15272 | 2.663446 |
| CNAG_03( | YKU80 | ATP-deper | 1274.250884 | 1449.166703 | 1261.71203 | 1633.850375 | 1520.725083 | 1713.918776 | 0.002181 | 0.274092 | 0.000964 | 3.300813 | 1.209233 | 2.661258 |
| CNAG_04 | - | pre-mRNA | 657.1794795 | 609.769518 | 612.092402 | 524.4069895 | 485.4340736 | 485.290851 | 0.002182 | -0.34549 | 0.000964 | -3.30069 | 0.78704 | 2.661184 |
| CNAG_00 | - | hypothetic | 483.4281122 | 561.2069932 | 560.8388668 | 415.3353027 | 370.6168387 | 447.4603801 | 0.002186 | -0.39517 | 0.000967 | -3.29996 | 0.760401 | 2.660291 |
| CNAG_02 | - | hypothetic | 560.4749858 | 620.4948729 | 632.3010948 | 442.4550538 | 493.2599522 | 494.1200065 | 0.002186 | -0.35754 | 0.000967 | -3.29996 | 0.780496 | 2.660291 |
| CNAG_12 | - | unspecifie | 0.222463055 | 0.107373724 | 3.298364871 | 18.87210322 | 14.11309071 | 9.929941517 | 0.002207 | 3.755864 | 0.000977 | 3.297157 | 13.50914 | 2.656201 |
| CNAG_13 | - | unspecifie | 166.2389252 | 215.5389402 | 179.2993712 | 260.982228 | 251.9745887 | 309.383601 | 0.002229 | 0.536227 | 0.000987 | 3.294276 | 1.450175 | 2.651868 |
| CNAG_06 | - | hypothetic | 408.8097778 | 536.3819588 | 530.9483117 | 359.9906928 | 379.3609594 | 373.1610741 | 0.002235 | -0.42223 | 0.000989 | -3.29352 | 0.746269 | 2.650813 |
| CNAG_05 | - | endoplasn | 1007.581082 | 917.9620854 | 1034.364733 | 804.5126694 | 843.4927498 | 780.5893145 | 0.002237 | -0.30125 | 0.000991 | -3.29316 | 0.811548 | 2.650382 |
| CNAG_00 | - | actin-2 | 1391.779395 | 1359.656241 | 1305.313467 | 1155.586083 | 1181.627748 | 1084.773192 | 0.002248 | -0.26109 | 0.000996 | -3.29171 | 0.834458 | 2.648268 |
| CNAG_07( | - | ATP-deper | 1677.788458 | 1464.221797 | 1550.575703 | 1230.154953 | 1341.183353 | 1348.919317 | 0.002257 | -0.27525 | 0.001 | -3.29052 | 0.826306 | 2.646539 |
| CNAG_02! | - | hypothetic | 541.1059838 | 759.5830794 | 644.2293977 | 838.4707608 | 846.3432289 | 848.0238378 | 0.002291 | 0.367401 | 0.001016 | 3.286099 | 1.290026 | 2.639965 |
| CNAG_00 | - | ubiquitin- | 6582.51322 | 6799.599306 | 6192.895209 | 5598.030225 | 5791.145457 | 5911.65809 | 0.002304 | -0.19308 | 0.001023 | -3.28422 | 0.874738 | 2.637455 |
| CNAG_06 | - | regulatory | 2765.236617 | 2952.1117 | 3100.097738 | 2565.08681 | 2622.978446 | 2442.479851 | 0.002307 | -0.22396 | 0.001024 | -3.28373 | 0.856212 | 2.63697 |
| CNAG_06( | - | aspartate | 2704.38655 | 2660.135702 | 2634.10824 | 2266.94452 | 2284.80037 | 2415.753684 | 0.002325 | -0.21475 | 0.001033 | -3.2814 | 0.861698 | 2.633557 |
| CNAG_03 | FZC46 | hypothetic | 575.6502918 | 608.6565665 | 622.7117785 | 434.6501005 | 473.3788243 | 515.1688592 | 0.002334 | -0.35909 | 0.001037 | -3.28024 | 0.779656 | 2.631886 |
| CNAG_07 | - | peptidyl-p | 749.8118205 | 688.5877968 | 697.5467467 | 875.7655689 | 929.1881148 | 858.0850471 | 0.002348 | 0.303306 | 0.001044 | 3.278518 | 1.233969 | 2.629362 |
| CNAG_06 | - | hypothetic | 696.800239 | 602.252066 | 622.7573349 | 483.7564132 | 488.0081178 | 541.8238669 | 0.002355 | -0.36032 | 0.001047 | -3.27747 | 0.778991 | 2.627995 |
| CNAG_04 | - | GTPase in | 895.5842266 | 960.9720556 | 924.5600374 | 801.5070582 | 744.2207352 | 751.7055984 | 0.002355 | -0.29036 | 0.001047 | -3.27755 | 0.817696 | 2.627995 |
| CNAG_04( | - | ATP-bindi | 5623.987932 | 5073.741522 | 5641.573465 | 6506.714372 | 6243.283335 | 6363.367092 | 0.002361 | 0.210588 | 0.001051 | 3.276634 | 1.15716 | 2.626822 |
| CNAG_02( | - | vacuolar p | 1626.650862 | 1822.90602 | 1709.479019 | 1349.219412 | 1522.354488 | 1488.801121 | 0.002372 | -0.25735 | 0.001056 | -3.2753 | 0.836621 | 2.62492 |
| CNAG_06( | - | Allantoin | 852.3704409 | 901.7223815 | 891.4875483 | 711.8789208 | 686.3991618 | 770.5030549 | 0.002377 | -0.30192 | 0.001058 | -3.27454 | 0.811172 | 2.623962 |
| CNAG_02 | - | methionyl | 889.7536996 | 1015.868063 | 925.6243207 | 867.8342373 | 711.3962194 | 658.5608519 | 0.002388 | -0.35401 | 0.001064 | -3.27307 | 0.782406 | 2.621937 |
| CNAG_00 | - | 8-Amino- | 491.6522444 | 587.0676053 | 600.2811289 | 416.8149778 | 415.5353107 | 466.3394881 | 0.002388 | -0.38611 | 0.001064 | -3.27298 | 0.765191 | 2.621922 |
| CNAG_05( | GPI13 | phosphati | 402.2237798 | 331.7392706 | 287.944897 | 219.7767295 | 288.090961 | 190.1670389 | 0.00241 | -0.56577 | 0.001075 | -3.27007 | 0.675595 | 2.617929 |
| CNAG_13 | - | unspecifie | 30.32849613 | 47.10639582 | 48.77284817 | 18.20345945 | 18.69621933 | 11.43625289 | 0.002421 | -1.42546 | 0.00108 | -3.26874 | 0.372301 | 2.616007 |
| CNAG_02 | - | hypothetic | 70.64139909 | 103.5502632 | 108.8557447 | 139.5214295 | 171.2362541 | 155.1964917 | 0.00244 | 0.712244 | 0.001089 | 3.266422 | 1.63835 | 2.612566 |
| CNAG_03 | - | hypothetic | 924.7108749 | 931.9036631 | 920.2937527 | 747.6816176 | 707.0806461 | 820.4453905 | 0.002449 | -0.30227 | 0.001094 | -3.26513 | 0.810974 | 2.610943 |
| CNAG_02 | SAV1 | GTP-bindi | 4196.692 | 3985.371033 | 3722.99169 | 3393.921766 | 3388.875324 | 3563.668438 | 0.002451 | -0.21784 | 0.001095 | -3.26485 | 0.859853 | 2.610624 |
| CNAG_05 | - | 2-dehydrc | 375.085757 | 365.3745916 | 396.7848361 | 478.8297191 | 489.2164161 | 522.6867401 | 0.002464 | 0.375049 | 0.001101 | 3.263342 | 1.296883 | 2.608438 |
| CNAG_07 | - | Urea carb | 688.866512 | 742.2369692 | 695.2784937 | 557.7426354 | 563.9465928 | 600.6729969 | 0.002468 | -0.3199 | 0.001103 | -3.26278 | 0.801126 | 2.607691 |
| CNAG_03( | - | hypothetic | 224.4810503 | 247.8766626 | 196.3934733 | 282.5881211 | 307.2335261 | 368.1992359 | 0.002477 | 0.506359 | 0.001109 | 3.261387 | 1.420461 | 2.606035 |
| CNAG_05 | - | hypothetic | 2544.446266 | 2530.853603 | 2634.08077 | 2131.47732 | 2312.464787 | 2254.811908 | 0.002479 | -0.21822 | 0.00111 | -3.26114 | 0.859625 | 2.605771 |
| CNAG_01 | SEC6 | exocyst cc | 1690.671346 | 1634.387226 | 1481.306027 | 1409.370355 | 1277.265527 | 1351.163986 | 0.00248 | -0.26641 | 0.001111 | -3.26089 | 0.831387 | 2.605514 |
| CNAG_12 | - | unspecifie | 548.8137402 | 673.201169 | 577.9705168 | 497.1625209 | 468.1993898 | 440.9217569 | 0.002501 | -0.3702 | 0.00112 | -3.2584 | 0.773674 | 2.601927 |
| CNAG_06 | - | hypothetic | 48.43521344 | 70.16036736 | 70.4933205 | 122.9405814 | 114.1985661 | 98.70106761 | 0.002501 | 0.814757 | 0.00112 | 3.258414 | 1.759002 | 2.601927 |
| CNAG_05( | - | meiotic Si | 584.9552875 | 664.6228029 | 589.7079673 | 499.4500589 | 496.6972648 | 466.4333861 | 0.002508 | -0.34512 | 0.001124 | -3.2575 | 0.787242 | 2.600678 |
| CNAG_03 | - | hypothetic | 505.2247563 | 478.2421646 | 403.1224385 | 350.7297476 | 287.7346138 | 385.2770428 | 0.002512 | -0.4544 | 0.001126 | -3.25696 | 0.729811 | 2.599963 |
| CNAG_06 | - | high-affini | 454.0648095 | 469.6427479 | 441.4564026 | 364.020432 | 369.9212922 | 319.9424564 | 0.002524 | -0.38992 | 0.001132 | -3.2555 | 0.763172 | 2.597978 |
| CNAG_07 | - | AP-1 com | 2287.234792 | 1914.576623 | 1857.764179 | 1714.39798 | 1684.451681 | 1635.399083 | 0.002533 | -0.28338 | 0.001137 | -3.25426 | 0.821666 | 2.596309 |
| CNAG_03 | - | DNA poly | 2428.963351 | 2764.603689 | 2764.171818 | 2214.776354 | 2268.416777 | 2355.819178 | 0.002533 | -0.23341 | 0.001137 | -3.25429 | 0.850623 | 2.596309 |
| CNAG_12( | - | unspecifie | 38.88588745 | 45.3554437 | 24.76429361 | 74.5235088 | 71.02136301 | 75.19550463 | 0.002536 | 1.025817 | 0.001138 | 3.253892 | 2.036112 | 2.595867 |
| CNAG_02 | PTP5 | hypothetic | 1617.426027 | 1535.418349 | 1479.257812 | 1844.724464 | 1921.959204 | 1772.886461 | 0.002557 | 0.242279 | 0.001148 | 3.251486 | 1.18286 | 2.592308 |
| CNAG_01 | - | hypothetic | 494.3008934 | 564.653632 | 592.9877353 | 683.5176634 | 722.159298 | 701.5017119 | 0.002566 | 0.337111 | 0.001153 | 3.250348 | 1.263224 | 2.590688 |
| CNAG_03( | - | hypothetic | 8.660219368 | 17.35641319 | 8.726848153 | 35.62658745 | 50.06288298 | 22.25899798 | 0.002577 | 1.683546 | 0.001158 | 3.249101 | 3.212165 | 2.588902 |
| CNAG_04 | - | hypothetic | 7.336792929 | 6.629619999 | 4.429721087 | 23.93257114 | 25.43960111 | 21.99310249 | 0.002579 | 2.005515 | 0.001159 | 3.248775 | 4.01532 | 2.588521 |
| CNAG_02 | - | Aldehyde | 2441.693359 | 2415.576965 | 2580.746454 | 2101.944124 | 2154.572402 | 2203.731262 | 0.002586 | -0.21879 | 0.001162 | -3.24798 | 0.859287 | 2.58743 |
| CNAG_06( | - | kinetochoi | 1686.436799 | 1695.968558 | 1810.876757 | 1900.556533 | 2214.404841 | 2145.813255 | 0.002597 | 0.254511 | 0.001168 | 3.246614 | 1.192932 | 2.585578 |
| CNAG_07 | - | hypothetic | 0.265178112 | 4.442212067 | 3.326484445 | 24.5343345 | 9.433362635 | 18.5657871 | 0.002608 | 2.854149 | 0.001173 | 3.245249 | 7.230769 | 2.583614 |

| | | | | | | | | | | | | | |
|---|---|---|---|---|---|---|---|---|---|---|---|---|---|
| CNAG_055 | - | hypothetic | 19.15759135 | 31.32890313 | 14.0918013 | 59.28332532 | 45.03531296 | 52.98147896 | 0.002618 | 1.27262 | 0.001178 | 3.244049 | 2.415999 | 2.582019 |
| CNAG_076 | - | hypothetic | 605.4975391 | 492.3224623 | 446.8537811 | 357.5771202 | 422.5646042 | 375.411953 | 0.002645 | -0.43522 | 0.001191 | -3.24101 | 0.739582 | 2.577507 |
| CNAG_039 | RDS2 | Regulator | 1347.352754 | 1335.9162 | 1175.236727 | 1147.910731 | 1047.857873 | 992.6552938 | 0.002655 | -0.29089 | 0.001196 | -3.23984 | 0.8174 | 2.575953 |
| CNAG_129 | - | unspecifie | 127.4882454 | 108.9931833 | 100.4243358 | 218.9235506 | 183.1601354 | 141.0382869 | 0.00266 | 0.678255 | 0.001199 | 3.239097 | 1.600203 | 2.575062 |
| CNAG_073 | - | amino aci | 29.70849672 | 40.01827904 | 38.53526933 | 65.67558863 | 93.91489511 | 62.0783431 | 0.00266 | 1.024964 | 0.001199 | 3.239165 | 2.034909 | 2.575062 |
| CNAG_130 | - | unspecifie | 34.40289688 | 22.58205593 | 34.93068103 | 7.959158091 | 14.47120006 | 6.957722561 | 0.002682 | -1.75491 | 0.001209 | -3.23671 | 0.296291 | 2.571551 |
| CNAG_056 | DAM1 | pre-mRNA | 1718.766778 | 1590.278605 | 1739.311358 | 1508.244854 | 1407.563321 | 1356.759927 | 0.002686 | -0.2565 | 0.001211 | -3.23619 | 0.837117 | 2.570874 |
| CNAG_026 | - | hypothetic | 518.7420062 | 578.6315987 | 522.6614066 | 746.0538547 | 685.8797542 | 638.3117147 | 0.00269 | 0.339919 | 0.001214 | 3.235571 | 1.265685 | 2.570168 |
| CNAG_003 | - | mitochond | 1009.911569 | 936.2626499 | 1005.591844 | 774.089971 | 865.9526924 | 795.0046865 | 0.0027 | -0.2931 | 0.001219 | -3.23437 | 0.816145 | 2.56858 |
| CNAG_003 | - | hypothetic | 577.9325142 | 575.3041948 | 648.2753016 | 493.2405912 | 466.4614099 | 470.847364 | 0.002709 | -0.34835 | 0.001224 | -3.23323 | 0.785483 | 2.567192 |
| CNAG_042 | - | ATP-deper | 1159.766801 | 1122.792396 | 1246.716482 | 1454.088763 | 1367.217112 | 1454.190516 | 0.002709 | 0.262107 | 0.001224 | 3.233294 | 1.199229 | 2.567192 |
| CNAG_036 | - | mRNA gua | 1082.668463 | 1076.433717 | 1044.142324 | 1198.560356 | 1458.890983 | 1291.035748 | 0.002726 | 0.286327 | 0.001232 | 3.231298 | 1.219532 | 2.564491 |
| CNAG_014 | - | hypothetic | 106.7258865 | 133.7098582 | 128.0867874 | 204.4997589 | 219.5810409 | 153.2401867 | 0.002754 | 0.635362 | 0.001246 | 3.228031 | 1.553327 | 2.559997 |
| CNAG_010 | UBC4 | ubiquitin-c | 117.0666858 | 103.6243693 | 118.4869166 | 211.8373165 | 160.5486985 | 161.9844843 | 0.002763 | 0.638782 | 0.001251 | 3.227077 | 1.557014 | 2.558666 |
| CNAG_010 | - | nuclear po | 3388.564906 | 3030.797414 | 3031.909532 | 2584.430088 | 2829.14216 | 2716.651616 | 0.002775 | -0.23286 | 0.001257 | -3.22558 | 0.850945 | 2.556744 |
| CNAG_120 | - | unspecifie | 5.006152917 | 3.407528015 | 5.462407518 | 21.65822481 | 20.1606897 | 19.74959258 | 0.002796 | 2.190026 | 0.001267 | 3.223292 | 4.563137 | 2.553511 |
| CNAG_055 | - | hypothetic | 268.9846629 | 311.4849742 | 339.1780161 | 414.3368161 | 461.8278378 | 379.6356832 | 0.002872 | 0.434828 | 0.001303 | 3.215337 | 1.35175 | 2.541808 |
| CNAG_036 | - | hypothetic | 1094.166769 | 1203.405476 | 1206.082812 | 1017.872992 | 957.2881409 | 964.8464638 | 0.002883 | -0.26864 | 0.001308 | -3.21411 | 0.8301 | 2.540189 |
| CNAG_006 | - | pre-mRNA | 1019.628408 | 995.6463241 | 1007.876524 | 1093.395775 | 1436.520434 | 1271.014688 | 0.002889 | 0.31536 | 0.001312 | 3.213368 | 1.244322 | 2.539184 |
| CNAG_065 | - | pirin | 209.1750308 | 166.1158538 | 186.7441174 | 261.7804332 | 302.3039129 | 249.7054962 | 0.002924 | 0.517161 | 0.001329 | 3.209707 | 1.431136 | 2.534001 |
| CNAG_058 | SHB17 | Phosphog | 2820.001533 | 2722.645442 | 2880.427407 | 2218.566192 | 2487.550564 | 2541.197034 | 0.002925 | -0.23245 | 0.001329 | -3.20956 | 0.851189 | 2.533898 |
| CNAG_045 | - | oxidoredu | 6.328780019 | 14.14256114 | 11.87594712 | 32.50575341 | 30.48061523 | 34.08422924 | 0.002933 | 1.617751 | 0.001334 | 3.208587 | 3.068961 | 2.532659 |
| CNAG_034 | - | hypothetic | 869.7971459 | 856.4897224 | 832.8608894 | 691.6437902 | 705.4389024 | 720.5832733 | 0.002934 | -0.28835 | 0.001334 | -3.20848 | 0.818835 | 2.532609 |
| CNAG_073 | - | hypothetic | 1739.844201 | 1664.619772 | 1853.406479 | 1464.514766 | 1563.762933 | 1449.992579 | 0.002945 | -0.2474 | 0.00134 | -3.2072 | 0.842411 | 2.530914 |
| CNAG_030 | - | hypothetic | 6.556491781 | 37.85314283 | 40.62498658 | 77.92881866 | 80.82275541 | 60.91638013 | 0.002969 | 1.37436 | 0.001353 | 3.20456 | 2.592528 | 2.527396 |
| CNAG_010 | - | class II alc | 29.79400383 | 46.48807124 | 43.87843041 | 103.0519453 | 113.8761946 | 44.65215028 | 0.002969 | 1.124917 | 0.001353 | 3.204596 | 2.180891 | 2.527396 |
| CNAG_034 | - | hypothetic | 220.9723247 | 253.2367427 | 186.8066964 | 302.2465253 | 328.9379533 | 301.8322054 | 0.002973 | 0.483927 | 0.001355 | 3.204046 | 1.398545 | 2.526737 |
| CNAG_012 | - | hypothetic | 747.1364417 | 747.6336003 | 680.3756092 | 578.79088 | 637.3897029 | 526.4240258 | 0.002996 | -0.33582 | 0.001366 | -3.20176 | 0.792336 | 2.523525 |
| CNAG_131 | - | unspecifie | 1.592458139 | 4.506872546 | 10.75857615 | 43.21586037 | 15.49524846 | 17.71985676 | 0.003012 | 2.30772 | 0.001374 | 3.200126 | 4.951001 | 2.521176 |
| CNAG_020 | - | glyoxal ox | 2017.76964 | 1537.637496 | 1379.09564 | 2207.600268 | 2107.390072 | 2044.830959 | 0.003034 | 0.349814 | 0.001385 | 3.19782 | 1.274397 | 2.517934 |
| CNAG_055 | - | ino eighty | 626.0105543 | 827.2769435 | 760.2862915 | 599.8252485 | 599.3492461 | 544.1680187 | 0.003043 | -0.35952 | 0.001389 | -3.19692 | 0.779422 | 2.516702 |
| CNAG_005 | - | Carnitine | 264.2113956 | 279.1622067 | 289.0828196 | 360.557897 | 394.4610388 | 365.1244733 | 0.003051 | 0.412337 | 0.001393 | 3.196047 | 1.33084 | 2.515498 |
| CNAG_057 | - | mitochond | 1363.86386 | 1383.378964 | 1469.490172 | 1216.31436 | 1127.182932 | 1225.686247 | 0.003067 | -0.25556 | 0.001401 | -3.19449 | 0.837662 | 2.513268 |
| CNAG_013 | - | arf/Sar far | 562.7404175 | 568.8217668 | 609.9054048 | 451.1108381 | 493.3038213 | 440.9008737 | 0.003115 | -0.34485 | 0.001423 | -3.18985 | 0.787392 | 2.506531 |
| CNAG_002 | - | ESCRT-II c | 332.0455612 | 426.7030484 | 397.860954 | 512.4131347 | 498.7030817 | 513.8534635 | 0.003116 | 0.384698 | 0.001424 | 3.18969 | 1.305587 | 2.506401 |
| CNAG_122 | - | unspecifie | 51.57237302 | 30.42735294 | 40.68157613 | 80.73659146 | 76.17391119 | 81.85660678 | 0.003138 | 0.953089 | 0.001436 | 3.187354 | 1.936014 | 2.503355 |
| CNAG_010 | - | C2 domair | 720.7341613 | 807.028207 | 690.1092122 | 1009.891432 | 928.2567261 | 851.4811972 | 0.003138 | 0.315904 | 0.001436 | 3.187388 | 1.244792 | 2.503355 |
| CNAG_051 | FRK101 | CAMK prc | 32.03693036 | 40.03693006 | 45.96075836 | 67.18362709 | 84.14724691 | 77.43657929 | 0.003173 | 0.947564 | 0.001453 | 3.183909 | 1.928613 | 2.498533 |
| CNAG_122 | - | unspecifie | 9.828363187 | 15.24856566 | 16.12799708 | 37.16568334 | 37.45990531 | 36.37530196 | 0.00318 | 1.445205 | 0.001457 | 3.183161 | 2.723016 | 2.497526 |
| CNAG_028 | - | hypothetic | 661.3045158 | 807.0387787 | 787.0934382 | 988.7803058 | 929.0980741 | 909.1518976 | 0.003185 | 0.311091 | 0.001459 | 3.182637 | 1.240646 | 2.496857 |
| CNAG_019 | - | hypothetic | 455.2194018 | 423.3966131 | 477.6532841 | 339.7803027 | 344.8340003 | 367.5588873 | 0.003186 | -0.38344 | 0.00146 | -3.18252 | 0.766606 | 2.496798 |
| CNAG_063 | - | SUMO act | 922.3097001 | 884.5092492 | 855.2655523 | 762.5786427 | 729.5834726 | 711.7503859 | 0.003189 | -0.28851 | 0.001462 | -3.18212 | 0.818745 | 2.496317 |
| CNAG_077 | - | hypothetic | 296.8962997 | 297.5029553 | 308.3004595 | 340.752754 | 513.7911506 | 411.7320294 | 0.003212 | 0.472688 | 0.001474 | 3.179812 | 1.387692 | 2.493199 |
| CNAG_012 | - | agmatinas | 711.4943171 | 844.7539592 | 811.6346677 | 916.0848498 | 963.5761727 | 1111.00924 | 0.003212 | 0.323259 | 0.001473 | 3.179868 | 1.251153 | 2.493199 |
| CNAG_101 | - | unspecifie | 2.788049852 | 13.01628093 | 4.444554495 | 27.09507051 | 33.44098054 | 19.91054139 | 0.003276 | 2.045175 | 0.001504 | 3.173852 | 4.127234 | 2.484621 |
| CNAG_127 | - | unspecifie | 189.0428601 | 220.6784851 | 205.712869 | 145.2182165 | 118.6629276 | 155.525462 | 0.003278 | -0.56907 | 0.001506 | -3.17349 | 0.674051 | 2.484403 |
| CNAG_129 | - | unspecifie | 29.56318842 | 29.2654357 | 24.71577625 | 58.87318945 | 77.52591386 | 45.5100456 | 0.003278 | 1.118543 | 0.001506 | 3.173557 | 2.171275 | 2.484403 |
| CNAG_028 | ERG130 | hydroxym | 4131.330216 | 3621.275542 | 4223.068364 | 3292.684325 | 3505.361472 | 3500.408583 | 0.003289 | -0.23378 | 0.001512 | -3.17241 | 0.850401 | 2.482927 |
| CNAG_036 | - | ATP-deper | 1235.674744 | 1271.440498 | 1295.790019 | 1508.496486 | 1462.10025 | 1565.197167 | 0.003331 | 0.239652 | 0.001532 | 3.168537 | 1.180708 | 2.477367 |
| CNAG_008 | CNB1 | calcineuri | 1606.657349 | 1632.21788 | 1469.568541 | 1386.803333 | 1234.119295 | 1358.912125 | 0.003349 | -0.25767 | 0.00154 | -3.16696 | 0.836439 | 2.475126 |
| CNAG_012 | - | hypothetic | 329.3557532 | 395.2950686 | 444.5082847 | 303.3093982 | 308.6781938 | 241.1914346 | 0.003363 | -0.47017 | 0.001547 | -3.16563 | 0.721879 | 2.473263 |
| CNAG_129 | - | unspecifie | 23.72874544 | 21.74394247 | 23.61255013 | 53.57829584 | 56.54920082 | 43.19270193 | 0.003365 | 1.166026 | 0.001549 | 3.165418 | 2.243927 | 2.473055 |
| CNAG_125 | - | unspecifie | 1.503337066 | 2.313862463 | 5.434724652 | 15.97367572 | 9.424201463 | 28.15729424 | 0.003367 | 2.686652 | 0.00155 | 3.165125 | 6.438176 | 2.472733 |
| CNAG_065 | - | monocarb | 470.6502075 | 596.7181976 | 553.3959361 | 405.1479166 | 443.2447934 | 413.1324534 | 0.003414 | -0.37499 | 0.001572 | -3.16096 | 0.771112 | 2.466756 |
| CNAG_000 | - | COPII-coa | 1997.875524 | 1956.566511 | 1904.661164 | 1757.284337 | 1643.036955 | 1669.780416 | 0.003427 | -0.22371 | 0.001579 | -3.15975 | 0.856363 | 2.465065 |
| CNAG_123 | - | unspecifie | 114.0260677 | 127.9134677 | 91.69445327 | 49.86219938 | 65.27282567 | 77.81650919 | 0.003439 | -0.81199 | 0.001585 | -3.15861 | 0.569598 | 2.463598 |
| CNAG_068 | - | allantoate | 236.7788626 | 266.9625762 | 211.1286404 | 141.0727529 | 142.8384266 | 200.9421745 | 0.003447 | -0.57604 | 0.00159 | -3.15778 | 0.670802 | 2.462582 |
| CNAG_044 | - | hypothetic | 555.502635 | 515.9969501 | 477.7458118 | 451.3696598 | 389.6809044 | 344.4033412 | 0.003449 | -0.4013 | 0.001591 | -3.1575 | 0.757175 | 2.462291 |
| CNAG_038 | - | Ras family | 862.0638931 | 946.0475705 | 943.8657119 | 1211.414428 | 1053.216819 | 1114.512108 | 0.003451 | 0.28141 | 0.001593 | 3.157234 | 1.215382 | 2.462003 |
| CNAG_063 | - | hypothetic | 210.7004305 | 310.3236401 | 270.9736081 | 406.0746942 | 354.654216 | 351.8175265 | 0.003453 | 0.478159 | 0.001594 | 3.156974 | 1.392965 | 2.461838 |
| CNAG_125 | - | unspecifie | 26.11360078 | 27.12280819 | 28.93921008 | 70.27643688 | 46.63856393 | 59.66406825 | 0.003455 | 1.096593 | 0.001595 | 3.156741 | 2.138491 | 2.461613 |
| CNAG_077 | - | hypothetic | 4.936633267 | 4.442582186 | 0.110250144 | 19.39987073 | 16.68568328 | 15.36622334 | 0.003468 | 2.629569 | 0.001602 | 3.155546 | 6.188413 | 2.459949 |
| CNAG_014 | - | endoplasn | 957.43789 | 929.7932742 | 1014.10722 | 785.8147674 | 801.1814357 | 829.3674808 | 0.003474 | -0.27944 | 0.001606 | -3.15479 | 0.823911 | 2.459162 |
| CNAG_007 | CCD3 | hypothetic | 527.4972742 | 500.9023179 | 445.7703085 | 384.9887941 | 363.7765354 | 399.7550803 | 0.003494 | -0.37635 | 0.001616 | -3.15307 | 0.770384 | 2.456726 |
| CNAG_060 | - | chromatin | 1993.477058 | 1994.384398 | 1927.150155 | 2239.227579 | 2337.657926 | 2332.357002 | 0.003496 | 0.208915 | 0.001617 | 3.152777 | 1.155818 | 2.456401 |
| CNAG_004 | - | hypothetic | 32.10795327 | 50.77053518 | 52.35768917 | 78.15877785 | 93.63127531 | 83.03512156 | 0.003498 | 0.908259 | 0.001619 | 3.152531 | 1.876779 | 2.45615 |
| CNAG_079 | - | hypothetic | 5324.899993 | 5877.285743 | 5163.768338 | 4732.596716 | 4970.885945 | 4707.180605 | 0.003541 | -0.19832 | 0.001639 | -3.14881 | 0.871562 | 2.450845 |
| CNAG_013 | - | thioredoxi | 1775.926566 | 1654.893667 | 1602.865094 | 1389.779601 | 1431.733603 | 1479.90621 | 0.003561 | -0.24228 | 0.001649 | -3.14702 | 0.845405 | 2.44842 |

| | | | | | | | | | | | | | | |
|---|---|---|---|---|---|---|---|---|---|---|---|---|---|---|
| CNAG_01 | RPA32 | replicatior | 1253.134585 | 1217.580953 | 1243.543212 | 1479.713884 | 1491.49422 | 1446.475652 | 0.003576 | 0.234756 | 0.001658 | 3.145526 | 1.176708 | 2.446606 |
| CNAG_02 | - | quinone c | 471.6326146 | 479.3794034 | 506.4569214 | 363.1203678 | 426.9445345 | 337.7150518 | 0.003596 | -0.38636 | 0.001668 | -3.14374 | 0.765058 | 2.444122 |
| CNAG_12 | - | unspecifie | 15.63934245 | 19.56334926 | 20.399562 | 44.96310656 | 39.01063287 | 48.47526295 | 0.003598 | 1.264084 | 0.001669 | 3.143501 | 2.401747 | 2.443993 |
| CNAG_03 | CFT3 | iron ion tr | 1015.904147 | 1149.495975 | 1044.023079 | 880.7841264 | 911.6187597 | 899.3301715 | 0.00361 | -0.26879 | 0.001676 | -3.14234 | 0.830017 | 2.442495 |
| CNAG_00 | - | arp2/3 cor | 1929.068624 | 1988.888348 | 2013.410515 | 1718.2937 | 1766.453482 | 1678.671045 | 0.00361 | -0.21532 | 0.001676 | -3.1424 | 0.861354 | 2.442495 |
| CNAG_06 | - | hypothetic | 404.2022607 | 386.8995904 | 371.2337358 | 479.5714372 | 490.0559493 | 541.5348169 | 0.003631 | 0.363947 | 0.001687 | 3.140452 | 1.286942 | 2.439928 |
| CNAG_01 | - | 6-phospho | 785.50332 | 713.1685528 | 616.4311168 | 574.1673845 | 577.79266 | 524.1804307 | 0.003634 | -0.35137 | 0.001689 | -3.14013 | 0.783842 | 2.43956 |
| CNAG_05 | - | hypothetic | 584.2661918 | 802.663034 | 637.8527284 | 900.8341583 | 834.1950713 | 868.0187359 | 0.003727 | 0.349396 | 0.001733 | 3.132462 | 1.274027 | 2.428664 |
| CNAG_00 | - | large subu | 1569.350986 | 1656.994183 | 1518.60989 | 1426.534355 | 1249.648217 | 1353.375072 | 0.003727 | -0.25053 | 0.001734 | -3.13232 | 0.840586 | 2.42859 |
| CNAG_06 | - | 26S protea | 5598.243199 | 4970.296625 | 5204.352925 | 6117.512337 | 6258.871373 | 5961.520391 | 0.003736 | 0.201456 | 0.001739 | 3.131514 | 1.149858 | 2.427603 |
| CNAG_05 | - | transcripti | 8670.434431 | 8536.452382 | 8190.352479 | 7201.28295 | 7744.016597 | 7716.7783 | 0.003766 | -0.17969 | 0.001754 | -3.12907 | 0.882895 | 2.424109 |
| CNAG_04 | - | cleavage s | 434.3108504 | 504.0638702 | 438.2691909 | 364.7840342 | 377.6968569 | 324.3803867 | 0.003794 | -0.38436 | 0.001767 | -3.12685 | 0.766119 | 2.420939 |
| CNAG_01 | - | lysophosp | 3486.936109 | 3173.118591 | 3004.298051 | 3642.458699 | 3796.22916 | 3998.656645 | 0.003797 | 0.227532 | 0.001769 | 3.126535 | 1.17083 | 2.420586 |
| CNAG_06 | - | hypothetic | 1591.889464 | 1742.261085 | 1773.5465 | 2030.749118 | 1970.161782 | 2029.281028 | 0.003803 | 0.224826 | 0.001772 | 3.125953 | 1.168636 | 2.41984 |
| CNAG_13 | - | unspecifie | 54.23698124 | 75.53458893 | 73.69348913 | 101.3535487 | 122.8737993 | 121.8196348 | 0.003829 | 0.753668 | 0.001785 | 3.123877 | 1.686075 | 2.41689 |
| CNAG_00 | - | tRNA-dihy | 500.5370516 | 397.5736381 | 443.5706578 | 339.8089067 | 335.3340364 | 352.0406814 | 0.00385 | -0.40129 | 0.001795 | -3.12219 | 0.757182 | 2.414514 |
| CNAG_06 | - | hypothetic | 251.4548107 | 273.803052 | 321.0343986 | 346.3900663 | 384.8864608 | 433.7655721 | 0.003882 | 0.445772 | 0.00181 | 3.119712 | 1.362042 | 2.410973 |
| CNAG_12 | - | unspecifie | 6.304517985 | 9.875895608 | 13.96577803 | 30.97431731 | 31.45156519 | 27.55956976 | 0.003901 | 1.637114 | 0.00182 | 3.118095 | 3.11043 | 2.408818 |
| CNAG_03 | - | hypothetic | 38.94174994 | 53.92841039 | 27.96015899 | 79.17996665 | 75.32315118 | 78.55222422 | 0.003924 | 0.961859 | 0.001831 | 3.116307 | 1.947818 | 2.406297 |
| CNAG_07 | - | U3 small r | 409.4511551 | 345.8078278 | 351.8763845 | 310.4455038 | 256.7804674 | 257.7642776 | 0.003925 | -0.44086 | 0.001833 | -3.11611 | 0.736695 | 2.406121 |
| CNAG_03 | - | phosphati | 739.3055749 | 710.1095873 | 668.7684283 | 847.6970642 | 933.5278041 | 843.6590942 | 0.003932 | 0.293736 | 0.001836 | 3.115515 | 1.22581 | 2.405358 |
| CNAG_03 | - | hypothetic | 199.9575512 | 224.1638398 | 175.0632747 | 302.4903261 | 249.3077527 | 299.5032334 | 0.003944 | 0.492993 | 0.001843 | 3.114457 | 1.407361 | 2.404027 |
| CNAG_02 | - | peptide al | 1474.520581 | 1251.958951 | 1246.671816 | 1083.132071 | 1129.851496 | 1096.95034 | 0.003981 | -0.27945 | 0.001862 | -3.11147 | 0.823903 | 2.399973 |
| CNAG_03 | - | hypothetic | 628.3791681 | 764.8840707 | 883.8899878 | 591.2166997 | 593.26996 | 599.6150891 | 0.004012 | -0.36633 | 0.001877 | -3.10908 | 0.775755 | 2.396686 |
| CNAG_13 | - | unspecifie | 157.1847626 | 134.5981293 | 142.7944618 | 100.0858446 | 97.24939211 | 83.53362269 | 0.004016 | -0.64488 | 0.001879 | -3.10867 | 0.639546 | 2.396196 |
| CNAG_13 | - | unspecifie | 213.8489165 | 329.3999158 | 286.7431567 | 191.8564867 | 190.3827229 | 198.8788781 | 0.004066 | -0.52962 | 0.001904 | -3.10478 | 0.692736 | 2.390819 |
| CNAG_12 | - | unspecifie | 14.7587884 | 38.99696674 | 61.86790869 | 96.68783438 | 94.57055924 | 67.6655041 | 0.004075 | 1.171231 | 0.001909 | 3.104068 | 2.252037 | 2.389888 |
| CNAG_02 | - | ATP-bindi | 3442.350146 | 3447.697785 | 3002.088894 | 2872.503014 | 2892.935467 | 2838.775979 | 0.004169 | -0.21654 | 0.001955 | -3.09702 | 0.860626 | 2.380001 |
| CNAG_07 | - | hypothetic | 216.8224995 | 194.9319684 | 207.8405014 | 168.0369337 | 112.5856438 | 137.860014 | 0.00421 | -0.58058 | 0.001974 | -3.09405 | 0.668695 | 2.375764 |
| CNAG_12 | - | unspecifie | 11.11605041 | 29.19533787 | 23.6083466 | 53.61228664 | 61.00664716 | 39.91470511 | 0.004216 | 1.266946 | 0.001979 | 3.093423 | 2.406516 | 2.375077 |
| CNAG_06 | - | hypothetic | 1448.995699 | 1380.147151 | 1388.480432 | 1128.228223 | 1128.911728 | 1297.785395 | 0.004254 | -0.2621 | 0.001997 | -3.09068 | 0.833875 | 2.37118 |
| CNAG_06 | - | cytoplasm | 718.1181422 | 756.2924744 | 818.9196099 | 563.0789961 | 620.0353065 | 671.6866161 | 0.004265 | -0.32107 | 0.002002 | -3.08987 | 0.800477 | 2.370105 |
| CNAG_01 | - | tyrosyl-DN | 2525.533838 | 2224.865428 | 2123.312461 | 1842.695362 | 1951.859453 | 2038.305311 | 0.004285 | -0.25265 | 0.002012 | -3.08841 | 0.839352 | 2.368089 |
| CNAG_00 | FZC28 | hypothetic | 747.5504959 | 787.6555938 | 719.9512179 | 922.4720964 | 866.9062075 | 992.292872 | 0.004293 | 0.288459 | 0.002017 | 3.087776 | 1.221335 | 2.367271 |
| CNAG_08 | - | hypothetic | 252.2467541 | 181.2042111 | 187.8493637 | 294.531305 | 281.3343695 | 313.9261247 | 0.0043 | 0.502068 | 0.002021 | 3.08709 | 1.416242 | 2.366494 |
| CNAG_06 | - | hypothetic | 114.6793644 | 97.15784899 | 112.083045 | 172.5131035 | 162.382704 | 159.7070233 | 0.004352 | 0.594248 | 0.002047 | 3.083355 | 1.509686 | 2.361265 |
| CNAG_02 | POS5 | NADH kin | 399.5901681 | 406.2829248 | 403.2014503 | 484.1904035 | 528.0746793 | 538.2540317 | 0.004355 | 0.344469 | 0.002049 | 3.083084 | 1.269684 | 2.360983 |
| CNAG_03 | - | solute car | 633.968065 | 647.4631321 | 663.2528102 | 558.6968754 | 475.8829118 | 534.088226 | 0.004371 | -0.3253 | 0.002057 | -3.08192 | 0.798133 | 2.35939 |
| CNAG_12 | - | unspecifie | 47.13000375 | 43.31863158 | 61.92548949 | 99.55590847 | 86.48230202 | 87.51092378 | 0.004401 | 0.833348 | 0.002071 | 3.079851 | 1.781816 | 2.356492 |
| CNAG_00 | - | hypothetic | 814.9077259 | 753.1058224 | 827.4835992 | 658.2120461 | 638.1226307 | 677.2809782 | 0.00441 | -0.29456 | 0.002076 | -3.07911 | 0.81532 | 2.355522 |
| CNAG_13 | - | unspecifie | 17.95901058 | 20.6479168 | 22.52833749 | 37.50071655 | 48.70120392 | 53.96609406 | 0.004444 | 1.210999 | 0.002093 | 3.076748 | 2.314979 | 2.352195 |
| CNAG_00 | PTC1 | Protein ph | 554.8578114 | 591.5482961 | 490.7007227 | 712.4533658 | 662.5327451 | 693.72674 | 0.00445 | 0.323515 | 0.002096 | 3.076313 | 1.251376 | 2.351673 |
| CNAG_10 | - | unspecifie | 0.18521499 | 2.256896189 | 0.077388533 | 7.045775462 | 10.5795531 | 17.25796708 | 0.004483 | 4.07696 | 0.002112 | 3.07398 | 16.87669 | 2.348389 |
| CNAG_08 | - | hypothetic | 244.1767786 | 209.1665345 | 205.9565619 | 313.2718471 | 311.6231057 | 285.2279978 | 0.004504 | 0.450406 | 0.002122 | 3.07255 | 1.366425 | 2.34642 |
| CNAG_00 | - | hypothetic | 25.9705567 | 24.92387082 | 11.96429935 | 40.66839196 | 64.61658881 | 43.15049082 | 0.004513 | 1.277347 | 0.002128 | 3.071772 | 2.423928 | 2.345513 |
| CNAG_03 | - | GTP-bindi | 794.33107 | 919.0728816 | 822.3201707 | 1065.757376 | 1053.340882 | 982.4525028 | 0.004513 | 0.275599 | 0.002128 | 3.071786 | 1.210497 | 2.345513 |
| CNAG_02 | MRE11 | DNA repa | 966.8564356 | 1055.790558 | 1034.395134 | 827.0598135 | 811.4858336 | 913.6958634 | 0.004533 | -0.27488 | 0.002138 | -3.07038 | 0.826519 | 2.343602 |
| CNAG_05 | - | Sugar trar | 2.716440367 | 7.65848995 | 4.40877613 | 26.73013463 | 16.53613453 | 17.61103233 | 0.004539 | 2.170098 | 0.002141 | 3.069902 | 4.500541 | 2.343018 |
| CNAG_02 | ALG12 | alpha-1,6- | 754.619095 | 810.2999649 | 834.0104546 | 944.9681577 | 1005.053837 | 973.5144787 | 0.004577 | 0.270027 | 0.00216 | 3.067271 | 1.20583 | 2.339418 |
| CNAG_12 | - | unspecifie | 0.118430101 | 0.05759837 | 0.049758096 | 10.91307181 | 6.184103792 | 3.354926199 | 0.00458 | 5.090114 | 0.002162 | 3.066975 | 34.06254 | 2.339101 |
| CNAG_12 | - | unspecifie | 87.02145233 | 56.07566127 | 96.73471946 | 53.48989325 | 36.70352887 | 34.74034562 | 0.004595 | -0.97542 | 0.00217 | -3.06596 | 0.508591 | 2.337737 |
| CNAG_12 | - | unspecifie | 33.10105933 | 32.51595042 | 38.51056066 | 62.60910441 | 68.47746974 | 71.84017622 | 0.004595 | 0.949121 | 0.002171 | 3.065836 | 1.930695 | 2.337671 |
| CNAG_02 | - | TatD DNa | 838.3367179 | 860.795753 | 848.8446248 | 706.4694657 | 719.2620071 | 706.1709779 | 0.004603 | -0.27186 | 0.002175 | -3.06526 | 0.82825 | 2.336941 |
| CNAG_00 | - | tRNA thre | 1228.478728 | 1315.491777 | 1421.478047 | 1109.628865 | 1153.158882 | 1095.846423 | 0.004649 | -0.25496 | 0.002198 | -3.06215 | 0.83801 | 2.332657 |
| CNAG_01 | - | prefoldin | 662.8846433 | 498.9025846 | 665.3057525 | 543.275994 | 388.6675694 | 429.8360894 | 0.004652 | -0.44065 | 0.0022 | -3.06186 | 0.736801 | 2.332346 |
| CNAG_05 | - | hypothetic | 6.15230542 | 4.479405358 | 5.467461613 | 24.14223347 | 19.22770593 | 18.69777026 | 0.004678 | 2.007724 | 0.002213 | 3.060016 | 4.021473 | 2.329898 |
| CNAG_00 | - | nuclease | 811.7999416 | 901.8376385 | 775.4197285 | 1058.717217 | 940.1994836 | 1072.252215 | 0.004678 | 0.289339 | 0.002213 | 3.060057 | 1.22208 | 2.329898 |
| CNAG_01 | - | hypothetic | 357.6106553 | 410.5438594 | 359.5026425 | 510.9396539 | 488.3739196 | 461.7622423 | 0.004711 | 0.35908 | 0.00223 | 3.057799 | 1.282608 | 2.326908 |
| CNAG_01 | PUB1 | nucleolysi | 5412.706143 | 5326.917177 | 5888.97358 | 6065.220193 | 6908.450519 | 6417.773816 | 0.004714 | 0.206266 | 0.002232 | 3.057535 | 1.153699 | 2.326638 |
| CNAG_12 | - | unspecifie | 19.29112381 | 38.89817666 | 36.37631632 | 69.99035857 | 63.26361879 | 61.95792465 | 0.004753 | 1.039818 | 0.002251 | 3.054959 | 2.055968 | 2.323019 |
| CNAG_12 | - | unspecifie | 101.5953203 | 140.8792948 | 137.3666668 | 52.89580054 | 82.59556165 | 90.0263997 | 0.004817 | -0.76905 | 0.002282 | -3.05082 | 0.586802 | 2.317252 |
| CNAG_03 | YOR1 | ATP-bindi | 1896.543696 | 1846.790191 | 1822.637337 | 2046.134626 | 2209.136002 | 2300.109162 | 0.004817 | 0.221169 | 0.002282 | 3.050827 | 1.165678 | 2.317252 |
| CNAG_07 | - | Sugar trar | 5.039981343 | 5.554433995 | 6.53092237 | 25.72447213 | 12.87351587 | 28.37600814 | 0.004818 | 2.008078 | 0.002284 | 3.050629 | 4.022459 | 2.317088 |
| CNAG_12 | - | unspecifie | 0.153548226 | 1.16987787 | 0.064325477 | 8.930538652 | 7.003910698 | 11.88833705 | 0.004832 | 4.581736 | 0.002291 | 3.049695 | 23.94639 | 2.315849 |
| CNAG_05 | - | hypothetic | 59.25558088 | 68.69793989 | 61.70860244 | 29.80275279 | 21.96326302 | 42.27037227 | 0.004848 | -1.04496 | 0.002299 | -3.04861 | 0.484657 | 2.314394 |
| CNAG_01 | - | hypothetic | 973.8928974 | 1074.113724 | 1084.501629 | 810.6354953 | 906.4681246 | 901.5230824 | 0.004849 | -0.27367 | 0.0023 | -3.04847 | 0.827214 | 2.314304 |
| CNAG_01 | - | hypothetic | 41.12557782 | 21.82949238 | 38.51520117 | 67.41177298 | 85.20581269 | 56.55883874 | 0.004882 | 1.0386 | 0.002316 | 3.046414 | 2.054233 | 2.311443 |
| CNAG_12 | - | unspecifie | 9.705727196 | 7.737382418 | 10.79744928 | 38.57225621 | 19.84634874 | 27.52465041 | 0.004894 | 1.676418 | 0.002323 | 3.045485 | 3.196334 | 2.310324 |

| | | | | | | | | | | | | | | |
|---|---|---|---|---|---|---|---|---|---|---|---|---|---|---|
| CNAG_043 | - | E3 ubiquit | 6422.664517 | 6992.480752 | 7160.102597 | 5891.443233 | 6105.118269 | 6284.662946 | 0.004906 | -0.18548 | 0.002329 | -3.04468 | 0.879354 | 2.309272 |
| CNAG_053 | CDC42 | cell divisio | 1138.68834 | 1083.989703 | 1121.969081 | 1410.577708 | 1342.262769 | 1260.012259 | 0.004921 | 0.248108 | 0.002337 | 3.043693 | 1.187649 | 2.307968 |
| CNAG_019 | - | aryl-alcoh | 577.8890129 | 549.4630818 | 619.4980225 | 460.4711208 | 471.676444 | 475.2575394 | 0.004925 | -0.32716 | 0.00234 | -3.04328 | 0.797104 | 2.307583 |
| CNAG_019 | - | cytoskelet | 669.3342678 | 702.5525915 | 693.2594822 | 833.7275812 | 834.2081116 | 854.7051635 | 0.004938 | 0.273513 | 0.002347 | 3.042407 | 1.208748 | 2.306439 |
| CNAG_035 | - | hypothetic | 951.7588066 | 1061.203259 | 1160.173923 | 845.7163451 | 904.7288093 | 882.6800791 | 0.004957 | -0.28482 | 0.002357 | -3.04115 | 0.820846 | 2.304738 |
| CNAG_126 | - | unspecifie | 35.53480648 | 46.47787397 | 50.22360886 | 82.25935688 | 84.03294765 | 75.32676077 | 0.005022 | 0.864717 | 0.00239 | 3.036871 | 1.820982 | 2.299127 |
| CNAG_057 | HPI4 | symplekin | 5013.088901 | 5127.449285 | 4937.661502 | 4433.74756 | 4524.051298 | 4590.566118 | 0.005126 | -0.16948 | 0.002442 | -3.03046 | 0.889164 | 2.290234 |
| CNAG_035 | - | hypothetic | 34.50123848 | 62.60396282 | 65.13441561 | 95.38724937 | 107.4132775 | 91.9717837 | 0.005128 | 0.849863 | 0.002443 | 3.030252 | 1.80233 | 2.290045 |
| CNAG_006 | - | hypothetic | 714.5088184 | 670.1434556 | 754.9456322 | 559.2864833 | 574.3065655 | 613.9831899 | 0.005154 | -0.30773 | 0.002459 | -3.02833 | 0.807913 | 2.287844 |
| CNAG_031 | FZC5 | hypothetic | 20.47929225 | 42.13342154 | 41.6928622 | 67.3852626 | 76.37085676 | 66.40594412 | 0.005181 | 1.007262 | 0.002472 | 3.026715 | 2.010093 | 2.285628 |
| CNAG_031 | - | hypothetic | 575.6118361 | 594.6742594 | 622.7156959 | 507.295351 | 482.8835243 | 463.094504 | 0.005207 | -0.31793 | 0.002486 | -3.025 | 0.80222 | 2.283381 |
| CNAG_076 | CSR2 | protoplast | 1564.942438 | 1609.735989 | 1554.953943 | 1931.982595 | 1804.539116 | 1811.729008 | 0.005211 | 0.215454 | 0.002489 | 3.024648 | 1.161069 | 2.283103 |
| CNAG_059 | - | hypothetic | 28.41660982 | 28.19707079 | 31.06033099 | 58.07372873 | 56.32155247 | 62.95630415 | 0.005211 | 1.001302 | 0.002489 | 3.024723 | 2.001805 | 2.283103 |
| CNAG_031 | - | hypothetic | 109.1802692 | 172.4465185 | 164.3081705 | 247.3883898 | 209.7503302 | 213.015068 | 0.005219 | 0.57446 | 0.002494 | 3.024074 | 1.48912 | 2.28239 |
| CNAG_010 | - | vacuolar r | 245.4543827 | 201.6850375 | 266.6590447 | 324.8204943 | 298.4937652 | 374.9081414 | 0.005222 | 0.468431 | 0.002496 | 3.023833 | 1.383604 | 2.282156 |
| CNAG_051 | - | ENTH don | 1374.417249 | 1451.244716 | 1534.535782 | 1132.843686 | 1336.953999 | 1216.844139 | 0.00527 | -0.25802 | 0.00252 | -3.02098 | 0.836233 | 2.278166 |
| CNAG_121 | - | unspecifie | 11.89228956 | 3.442157364 | 7.610649951 | 21.28945215 | 24.45132351 | 30.64305805 | 0.005298 | 1.818449 | 0.002535 | 3.019095 | 3.527018 | 2.275914 |
| CNAG_036 | - | hypothetic | 534.5627047 | 496.6364302 | 511.8244056 | 375.5332349 | 353.3553649 | 466.2535854 | 0.005332 | -0.38327 | 0.002552 | -3.01707 | 0.7667 | 2.273123 |
| CNAG_038 | - | delta8-fatt | 869.8745511 | 907.1160803 | 877.6448452 | 750.8684894 | 707.9822519 | 766.0816661 | 0.005345 | -0.27007 | 0.002559 | -3.01624 | 0.829282 | 2.272049 |
| CNAG_122 | - | unspecifie | 127.8161759 | 210.0814608 | 149.4595183 | 236.9272314 | 222.6516743 | 270.5415415 | 0.005352 | 0.570077 | 0.002563 | 3.015788 | 1.484603 | 2.271507 |
| CNAG_080 | - | hypothetic | 452.1675016 | 490.2950316 | 463.9918786 | 543.2754094 | 627.3150282 | 611.5451932 | 0.005403 | 0.327124 | 0.002588 | 3.012826 | 1.25451 | 2.267377 |
| CNAG_016 | - | DNA misn | 2985.975468 | 2652.736396 | 2722.724764 | 3182.528689 | 3296.822839 | 3284.879189 | 0.005519 | 0.208095 | 0.002645 | 3.006189 | 1.155162 | 2.258109 |
| CNAG_065 | - | hypothetic | 730.9060634 | 712.1545179 | 796.5340572 | 615.4063941 | 634.7445699 | 595.1863211 | 0.005552 | -0.2949 | 0.002663 | -3.00417 | 0.81513 | 2.255559 |
| CNAG_021 | - | hypothetic | 28.19958724 | 15.31599861 | 15.13118721 | 54.63957295 | 49.59543083 | 36.58270282 | 0.005626 | 1.243679 | 0.0027 | 2.99999 | 2.368016 | 2.249816 |
| CNAG_071 | - | hypothetic | 14.3591746 | 15.23356463 | 9.784842708 | 38.9227706 | 35.73542919 | 29.84300434 | 0.005636 | 1.409696 | 0.002705 | 2.999366 | 2.656811 | 2.249038 |
| CNAG_053 | - | hypothetic | 1111.87219 | 1219.667932 | 986.599967 | 1370.019018 | 1246.404321 | 1460.7757 | 0.005696 | 0.283309 | 0.002735 | 2.996029 | 1.216983 | 2.244394 |
| CNAG_054 | MTN1 | Copper-de | 2552.670537 | 1847.988026 | 2247.083114 | 2794.328164 | 2747.395234 | 2682.067809 | 0.005754 | 0.29041 | 0.002764 | 2.99283 | 1.222987 | 2.240059 |
| CNAG_044 | YAK103 | CMGC/DY | 565.1676245 | 444.048712 | 417.137379 | 564.2895244 | 681.7334626 | 639.2906624 | 0.005754 | 0.385606 | 0.002765 | 2.992749 | 1.306408 | 2.240055 |
| CNAG_015 | - | hypothetic | 98.32634001 | 96.01246381 | 80.15611087 | 140.4630626 | 136.4716066 | 150.7127078 | 0.005802 | 0.621671 | 0.002791 | 2.98982 | 1.538656 | 2.23644 |
| CNAG_068 | - | cytoplasm | 538.4693113 | 503.2667636 | 515.1753646 | 656.3545764 | 651.3803674 | 632.7094299 | 0.005841 | 0.301972 | 0.002812 | 2.987596 | 1.232828 | 2.2335 |
| CNAG_076 | - | hypothetic | 15.58810928 | 14.21250672 | 19.31911598 | 29.60006108 | 40.89723811 | 51.62535907 | 0.005846 | 1.306493 | 0.002815 | 2.987203 | 2.473395 | 2.233161 |
| CNAG_054 | - | hypothetic | 2.755594352 | 7.689541614 | 9.695971684 | 26.46990962 | 19.13195908 | 23.06015518 | 0.005941 | 1.904263 | 0.002864 | 2.982023 | 3.743175 | 2.226138 |
| CNAG_048 | - | hypothetic | 9.872838089 | 11.01290377 | 26.69346484 | 40.2389827 | 41.7872789 | 40.79987889 | 0.005941 | 1.379526 | 0.002863 | 2.982043 | 2.601828 | 2.226138 |
| CNAG_011 | - | hypothetic | 1111.467185 | 993.3797247 | 1046.143716 | 889.3928212 | 890.9080374 | 880.463025 | 0.005967 | -0.26017 | 0.002878 | -2.98046 | 0.834988 | 2.224257 |
| CNAG_051 | - | peroxisom | 93.88255536 | 134.7123918 | 104.6575928 | 156.647903 | 168.4359306 | 182.8829891 | 0.005969 | 0.598522 | 0.00288 | 2.980293 | 1.514164 | 2.224124 |
| CNAG_071 | - | V-type pro | 1637.092318 | 1725.956955 | 1569.810266 | 1449.036348 | 1405.850613 | 1425.532521 | 0.005975 | -0.21952 | 0.002884 | -2.97989 | 0.858852 | 2.223665 |
| CNAG_013 | - | phosphati | 91.08357897 | 46.60555557 | 57.7728996 | 113.3224162 | 122.9039944 | 107.5168452 | 0.006024 | 0.800234 | 0.002908 | 2.977325 | 1.741384 | 2.220137 |
| CNAG_128 | - | unspecifie | 166.6097564 | 201.1120696 | 117.3776901 | 95.20224851 | 95.42002395 | 117.7865347 | 0.006045 | -0.67043 | 0.002919 | -2.97609 | 0.62832 | 2.218609 |
| CNAG_128 | - | unspecifie | 4.044286566 | 12.04344661 | 20.32492967 | 36.44445991 | 29.48371964 | 41.73030328 | 0.006058 | 1.544418 | 0.002927 | 2.975349 | 2.916865 | 2.217669 |
| CNAG_032 | - | hypothetic | 380.8777017 | 549.2830724 | 546.9162639 | 355.3067776 | 330.9247366 | 414.1363541 | 0.006122 | -0.43821 | 0.002958 | -2.97206 | 0.738051 | 2.213126 |
| CNAG_125 | - | unspecifie | 16.766047 | 22.74648671 | 15.11336066 | 38.47444904 | 35.48643306 | 54.949274 | 0.006134 | 1.247275 | 0.002967 | 2.971102 | 2.373925 | 2.212256 |
| CNAG_020 | NAP1 | Nucleosor | 11071.3825 | 10872.35039 | 11188.12486 | 9805.514253 | 9783.925134 | 10347.83715 | 0.006134 | -0.16164 | 0.002967 | -2.97113 | 0.89401 | 2.212256 |
| CNAG_052 | - | hypothetic | 3429.521393 | 3315.211662 | 3124.708222 | 2854.580195 | 2873.944864 | 2989.710689 | 0.006134 | -0.19435 | 0.002966 | -2.97127 | 0.873966 | 2.212256 |
| CNAG_058 | - | cytochrom | 2003.831322 | 2109.556097 | 2128.585804 | 1764.939278 | 1799.187169 | 1897.303883 | 0.006162 | -0.20766 | 0.002983 | -2.96949 | 0.865942 | 2.210264 |
| CNAG_043 | - | hypothetic | 1563.779644 | 1689.432982 | 1431.291641 | 1931.196487 | 1976.32453 | 1712.990617 | 0.006189 | 0.247952 | 0.003 | 2.967781 | 1.18752 | 2.20839 |
| CNAG_013 | - | protein tra | 1038.332731 | 1118.417583 | 1045.200343 | 1188.455425 | 1295.695652 | 1364.219258 | 0.006189 | 0.250342 | 0.003 | 2.967784 | 1.189489 | 2.20839 |
| CNAG_026 | - | hypothetic | 352.536832 | 375.9121749 | 376.3612237 | 246.2203637 | 319.1153926 | 276.5870535 | 0.006189 | -0.40653 | 0.002999 | -2.96779 | 0.754434 | 2.20839 |
| CNAG_012 | - | ATP-deper | 1512.278496 | 1408.27301 | 1263.864706 | 1596.340967 | 1689.019812 | 1746.141253 | 0.006189 | 0.250937 | 0.002999 | 2.967891 | 1.18998 | 2.20839 |
| CNAG_123 | - | unspecifie | 9.74145456 | 10.94547634 | 12.920132 | 30.92385573 | 28.73211466 | 30.81543474 | 0.006196 | 1.490196 | 0.003004 | 2.967278 | 2.809271 | 2.207899 |
| CNAG_123 | - | unspecifie | 90.7909736 | 106.3905552 | 84.18173631 | 68.34059541 | 44.44598007 | 50.22308691 | 0.006228 | -0.8036 | 0.003021 | -2.96561 | 0.572916 | 2.205653 |
| CNAG_126 | - | unspecifie | 15.85388908 | 44.25623761 | 39.56871453 | 69.05022612 | 68.47568811 | 69.65785828 | 0.006235 | 1.059345 | 0.003025 | 2.965174 | 2.083985 | 2.205145 |
| CNAG_013 | - | hypothetic | 3287.144073 | 3261.358256 | 3339.023331 | 2838.213821 | 3018.910246 | 2948.654728 | 0.006302 | -0.18273 | 0.003059 | -2.96173 | 0.881032 | 2.200505 |
| CNAG_051 | - | hypothetic | 84.53704942 | 76.7439999 | 130.0979938 | 148.9800209 | 152.0428114 | 167.3278248 | 0.006381 | 0.671403 | 0.003099 | 2.957744 | 1.592621 | 2.195103 |
| CNAG_010 | - | hypothetic | 1234.275403 | 1299.323643 | 1340.457414 | 1097.973285 | 1038.338606 | 1166.814767 | 0.006386 | -0.2455 | 0.003103 | -2.95735 | 0.843526 | 2.194761 |
| CNAG_018 | TUB1 | tubulin be | 11028.06663 | 9989.950702 | 10350.96972 | 9792.41158 | 9017.947813 | 9060.122206 | 0.006386 | -0.18625 | 0.003103 | -2.95737 | 0.878886 | 2.194761 |
| CNAG_003 | - | RAD51-lik | 142.9002642 | 172.4973341 | 186.6819104 | 256.5962341 | 222.6537753 | 234.0955965 | 0.006415 | 0.495286 | 0.003118 | 2.955875 | 1.409601 | 2.192799 |
| CNAG_122 | - | unspecifie | 9.52016751 | 3.409923414 | 2.290781421 | 22.47737675 | 16.52833772 | 22.97016692 | 0.006434 | 2.082144 | 0.003128 | 2.954911 | 4.234361 | 2.19155 |
| CNAG_010 | - | glutaredo: | 1108.983047 | 984.7064512 | 906.4853158 | 944.9380198 | 731.19926 | 716.2724216 | 0.006503 | -0.34192 | 0.003163 | -2.95144 | 0.788988 | 2.18689 |
| CNAG_032 | CCD4 | hypothetic | 322.5843728 | 386.8139213 | 301.9496854 | 400.0676065 | 454.7732916 | 483.7858456 | 0.006587 | 0.391766 | 0.003205 | 2.947405 | 1.311998 | 2.181324 |
| CNAG_057 | - | oligopepti | 1055.416846 | 1096.683936 | 874.5293112 | 830.9889418 | 870.2402477 | 795.027052 | 0.006618 | -0.29285 | 0.003221 | -2.94581 | 0.816288 | 2.179298 |
| CNAG_027 | RRP6 | exosome c | 4067.201828 | 3998.329188 | 4227.361866 | 3334.666609 | 3794.373695 | 3700.219063 | 0.006704 | -0.19843 | 0.003266 | -2.94157 | 0.8715 | 2.173675 |
| CNAG_028 | - | bloom syr | 1617.546875 | 1792.873878 | 1673.329549 | 1900.637415 | 1982.258993 | 2079.197604 | 0.006733 | 0.215414 | 0.003281 | 2.940123 | 1.161037 | 2.171759 |
| CNAG_052 | - | hypothetic | 91.43145652 | 85.30721568 | 103.5317529 | 125.3526746 | 186.989055 | 133.1345088 | 0.006786 | 0.654127 | 0.003308 | 2.937559 | 1.573663 | 2.168385 |
| CNAG_049 | - | hypothetic | 443.6473862 | 486.8744659 | 455.3183118 | 345.199664 | 385.4624146 | 372.0186271 | 0.006787 | -0.34365 | 0.003309 | -2.93746 | 0.788045 | 2.16835 |
| CNAG_129 | - | unspecifie | 12.13870415 | 6.725408101 | 24.56873538 | 47.75905079 | 32.05287559 | 37.50285728 | 0.006793 | 1.448418 | 0.003314 | 2.936997 | 2.729086 | 2.167924 |
| CNAG_068 | - | hypothetic | 1083.413822 | 959.9752024 | 989.6331221 | 827.8472243 | 863.3158297 | 862.6821306 | 0.006829 | -0.26389 | 0.003332 | -2.9353 | 0.832841 | 2.165661 |
| CNAG_011 | PBX1 | parallel be | 1843.770123 | 1807.896825 | 1896.089784 | 1417.659072 | 1632.751601 | 1697.457019 | 0.006837 | -0.24012 | 0.003337 | -2.93484 | 0.846676 | 2.165116 |
| CNAG_032 | - | hypothetic | 585.1950316 | 366.3304109 | 333.8578001 | 309.4313721 | 321.5731229 | 273.3563533 | 0.006846 | -0.52515 | 0.003343 | -2.93426 | 0.694889 | 2.164533 |

| | | | | | | | | | | | | | |
|---|---|---|---|---|---|---|---|---|---|---|---|---|---|
| CNAG_067 | SEC24 | protein tra | 1214.194552 | 1044.035456 | 1147.441126 | 852.6102132 | 1005.690791 | 975.9062136 | 0.006866 | -0.28184 | 0.003354 | -2.93332 | 0.822543 | 2.163326 |
| CNAG_015 | - | hypothetic | 772.7517325 | 665.8415513 | 708.0725676 | 576.4564531 | 558.7460769 | 619.5286831 | 0.006877 | -0.30685 | 0.00336 | -2.93272 | 0.808402 | 2.162597 |
| CNAG_047 | EZA1 | hypothetic | 610.9235974 | 598.0757714 | 629.2659357 | 815.2655683 | 774.7426467 | 690.5233238 | 0.006917 | 0.295586 | 0.003384 | 2.930543 | 1.227384 | 2.160088 |
| CNAG_008 | - | phosphati | 1715.699821 | 1329.420744 | 1045.178003 | 942.1410762 | 880.378376 | 868.3934551 | 0.006937 | -0.62027 | 0.003394 | -2.92959 | 0.650549 | 2.158858 |
| CNAG_000 | - | hypothetic | 94.98744284 | 118.5929792 | 104.6338519 | 167.0582513 | 154.5908067 | 154.1627577 | 0.007033 | 0.573585 | 0.003446 | 2.924883 | 1.488217 | 2.152833 |
| CNAG_043 | - | exocyst cc | 2157.921769 | 2333.611826 | 2046.52683 | 1969.761293 | 1819.84998 | 1900.677215 | 0.007042 | -0.21519 | 0.003452 | -2.92434 | 0.861435 | 2.152285 |
| CNAG_055 | JJJ1 | DnaJ-like | 2407.760679 | 2379.991518 | 2241.678211 | 1991.421828 | 2040.710731 | 2141.538801 | 0.007063 | -0.20234 | 0.003463 | -2.92336 | 0.869141 | 2.151034 |
| CNAG_064 | - | jumonji dc | 555.6826115 | 530.0583857 | 570.4666316 | 462.9081469 | 455.3046551 | 426.461963 | 0.007085 | -0.3169 | 0.003476 | -2.92221 | 0.802793 | 2.149646 |
| CNAG_079 | - | hypothetic | 570.7354252 | 625.7611328 | 443.6981102 | 478.6217629 | 359.3610433 | 405.3776848 | 0.007112 | -0.4135 | 0.00349 | -2.92089 | 0.7508 | 2.148015 |
| CNAG_008 | - | hypothetic | 623.6655713 | 588.3571694 | 521.6253694 | 782.6024808 | 738.5187342 | 661.650887 | 0.007129 | 0.317138 | 0.003499 | 2.920074 | 1.245856 | 2.146992 |
| CNAG_033 | - | hypothetic | 608.1270955 | 615.0778681 | 505.5200857 | 487.0641973 | 461.3164937 | 433.1400889 | 0.007139 | -0.33842 | 0.003507 | -2.9194 | 0.79091 | 2.146381 |
| CNAG_067 | - | hypothetic | 1577.690392 | 1820.771209 | 1828.887249 | 1441.138472 | 1551.678958 | 1502.135093 | 0.007163 | -0.23223 | 0.003521 | -2.9182 | 0.851316 | 2.144918 |
| CNAG_044 | - | multidrug | 493.5343267 | 431.9761881 | 390.3266094 | 335.9192579 | 325.829156 | 361.9950487 | 0.007167 | -0.37948 | 0.003524 | -2.91794 | 0.768717 | 2.144659 |
| CNAG_074 | ERG27 | 3-keto ste | 818.445759 | 755.2763082 | 871.1860641 | 692.5097866 | 695.9884089 | 637.3918502 | 0.007174 | -0.28781 | 0.003528 | -2.91754 | 0.819147 | 2.14421 |
| CNAG_010 | - | hypothetic | 287.5267822 | 295.3209208 | 310.4012747 | 383.935548 | 396.1111151 | 381.7842635 | 0.007241 | 0.363758 | 0.003563 | 2.914431 | 1.286773 | 2.140213 |
| CNAG_020 | - | hypothetic | 878.34332 | 976.1701903 | 831.9392546 | 1051.589614 | 1131.002792 | 1065.679705 | 0.007243 | 0.259406 | 0.003565 | 2.914252 | 1.196986 | 2.140072 |
| CNAG_024 | - | endoplasn | 225.3754044 | 178.9949326 | 145.2396419 | 261.8487508 | 272.8473065 | 254.0941215 | 0.007248 | 0.506228 | 0.003569 | 2.913954 | 1.420332 | 2.139765 |
| CNAG_008 | - | Flavin-con | 131.0872435 | 138.0227345 | 132.3547934 | 184.6930826 | 181.2585733 | 215.0619457 | 0.007263 | 0.517612 | 0.003577 | 2.913265 | 1.431584 | 2.138914 |
| CNAG_031 | CHO2 | Phosphati | 1826.50245 | 1881.240088 | 1787.443099 | 2074.233581 | 2191.042957 | 2100.365377 | 0.007273 | 0.197156 | 0.003583 | 2.912738 | 1.146436 | 2.138289 |
| CNAG_032 | - | 28 kda go | 1070.864184 | 1219.564324 | 1230.603784 | 966.3852588 | 998.7287027 | 1024.747127 | 0.007295 | -0.25077 | 0.003595 | -2.91165 | 0.840448 | 2.136988 |
| CNAG_058 | - | CDK-activ | 669.0493615 | 712.0972624 | 704.8598031 | 611.6383017 | 557.042989 | 557.4373909 | 0.007305 | -0.28844 | 0.003601 | -2.91113 | 0.818789 | 2.136382 |
| CNAG_052 | - | hypothetic | 125.2431931 | 131.5570116 | 125.9557159 | 176.1341153 | 185.6773379 | 190.7340205 | 0.007312 | 0.517227 | 0.003605 | 2.910768 | 1.431202 | 2.135981 |
| CNAG_021 | - | ubiquitin- | 616.3679484 | 566.7044053 | 612.0681153 | 478.378908 | 492.3880481 | 497.450074 | 0.007321 | -0.30591 | 0.003611 | -2.9103 | 0.808932 | 2.135443 |
| CNAG_006 | EGCrP1 | fungus-sp | 1048.458348 | 968.5930282 | 1028.003085 | 893.3577063 | 837.3948846 | 849.3815767 | 0.007326 | -0.2547 | 0.003614 | -2.91 | 0.838162 | 2.135121 |
| CNAG_049 | - | peroxin-12 | 118.4055066 | 158.4518956 | 151.5184324 | 212.1180974 | 210.7161132 | 195.2793109 | 0.007336 | 0.516618 | 0.00362 | 2.90948 | 1.430598 | 2.134513 |
| CNAG_054 | - | pyridoxam | 1059.124846 | 1124.778373 | 1186.869777 | 883.0332693 | 916.7419581 | 1032.460296 | 0.007356 | -0.26598 | 0.003631 | -2.90857 | 0.831633 | 2.133362 |
| CNAG_122 | - | unspecifie | 7.270772618 | 3.40873372 | 5.463711869 | 25.03034403 | 18.34284211 | 16.54016547 | 0.007428 | 1.958573 | 0.003667 | 2.905461 | 3.886774 | 2.129148 |
| CNAG_006 | - | hypothetic | 1488.862697 | 1508.38527 | 1711.546488 | 1305.729533 | 1388.648343 | 1343.387719 | 0.007429 | -0.2373 | 0.003669 | -2.90534 | 0.848331 | 2.129082 |
| CNAG_063 | - | hypothetic | 120.4130001 | 81.04914396 | 107.8172932 | 141.7819459 | 165.9237452 | 171.7813503 | 0.007461 | 0.614416 | 0.003685 | 2.903914 | 1.530938 | 2.127216 |
| CNAG_064 | - | hypothetic | 8.874798633 | 38.88758551 | 38.48188853 | 74.09310453 | 64.17872253 | 57.56862351 | 0.007477 | 1.20304 | 0.003694 | 2.903143 | 2.302244 | 2.126254 |
| CNAG_056 | - | hypothetic | 835.0490119 | 764.006937 | 806.3012906 | 991.7874639 | 976.5474738 | 939.1399681 | 0.007528 | 0.257372 | 0.00372 | 2.900955 | 1.195299 | 2.123328 |
| CNAG_021 | - | anaphase- | 847.7557561 | 936.1910416 | 901.0976859 | 745.3801758 | 793.4948575 | 722.8529544 | 0.007572 | -0.26304 | 0.003744 | -2.89896 | 0.833329 | 2.120778 |
| CNAG_024 | - | hypothetic | 579.2817051 | 517.277115 | 515.1930982 | 682.8321529 | 663.4334683 | 658.2357401 | 0.007575 | 0.298605 | 0.003746 | 2.898784 | 1.229955 | 2.120642 |
| CNAG_128 | - | unspecifie | 82.9167117 | 35.64509231 | 60.594615 | 34.72935141 | 31.6725451 | 11.5337473 | 0.007614 | -1.23909 | 0.003766 | -2.89708 | 0.42364 | 2.118394 |
| CNAG_128 | - | unspecifie | 128.7328645 | 144.441383 | 111.0811251 | 191.8721045 | 203.9189451 | 168.6895308 | 0.007647 | 0.538401 | 0.003785 | 2.895576 | 1.452362 | 2.116522 |
| CNAG_012 | - | protein M | 2243.204979 | 2283.043364 | 2360.006611 | 2007.814385 | 2071.80279 | 2018.354239 | 0.007685 | -0.19107 | 0.003804 | -2.89394 | 0.875958 | 2.114369 |
| CNAG_036 | - | hypothetic | 107.6301298 | 94.96794807 | 93.98408935 | 147.4535878 | 148.5924402 | 149.6787715 | 0.007693 | 0.579159 | 0.00381 | 2.89351 | 1.493978 | 2.113878 |
| CNAG_034 | - | hypothetic | 1256.299763 | 1164.675432 | 1178.397924 | 996.7489326 | 1011.652204 | 1070.245468 | 0.007722 | -0.24114 | 0.003825 | -2.89228 | 0.846079 | 2.112284 |
| CNAG_036 | - | hypothetic | 761.1332705 | 669.0812729 | 740.0444291 | 619.3709336 | 608.8556559 | 559.6870677 | 0.00774 | -0.29684 | 0.003834 | -2.89147 | 0.814035 | 2.111268 |
| CNAG_069 | - | hypothetic | 11.95303098 | 8.795216238 | 8.684011706 | 25.29171397 | 21.66130935 | 35.03099738 | 0.007759 | 1.569592 | 0.003845 | 2.890626 | 2.968207 | 2.110213 |
| CNAG_000 | - | palmitoyl- | 380.8783181 | 395.4818607 | 354.1754037 | 458.5382867 | 466.748761 | 521.5536152 | 0.007901 | 0.341151 | 0.003917 | 2.88474 | 1.266767 | 2.1023 |
| CNAG_006 | - | hypothetic | 295.7883832 | 352.3776805 | 341.3319778 | 413.4473782 | 416.7371267 | 453.8534379 | 0.007931 | 0.361251 | 0.003934 | 2.883394 | 1.284539 | 2.100658 |
| CNAG_074 | - | pantothen | 370.5410088 | 447.1911796 | 446.8930335 | 553.6785704 | 502.9243038 | 559.3525815 | 0.007956 | 0.339303 | 0.003948 | 2.882326 | 1.265145 | 2.099293 |
| CNAG_004 | UBP16 | ubiquitin | 764.8239329 | 835.987981 | 827.4901057 | 606.6856709 | 764.272227 | 607.4458208 | 0.007958 | -0.31054 | 0.003949 | -2.88219 | 0.806337 | 2.099208 |
| CNAG_034 | NDX3 | cleavage a | 1.425214049 | 3.331694782 | 0.085194742 | 18.15212638 | 5.963632561 | 13.07953741 | 0.007987 | 3.159501 | 0.003965 | 2.880941 | 8.935203 | 2.097598 |
| CNAG_029 | - | hypothetic | 283.9982963 | 289.9253946 | 282.7054667 | 427.9813784 | 371.0455264 | 334.1603969 | 0.007997 | 0.39137 | 0.003972 | 2.880414 | 1.311639 | 2.097085 |
| CNAG_017 | - | hypothetic | 461.5459143 | 539.8244418 | 488.5190772 | 610.3636865 | 605.6203392 | 642.6285942 | 0.007997 | 0.305048 | 0.003971 | 2.880444 | 1.23546 | 2.097085 |
| CNAG_044 | - | hypothetic | 11.09284784 | 35.547379 | 11.96002356 | 45.60057529 | 61.99462737 | 38.77869584 | 0.008018 | 1.326022 | 0.003983 | 2.879518 | 2.507105 | 2.095957 |
| CNAG_047 | - | hypothetic | 1891.880273 | 1731.548195 | 1962.300066 | 2211.307812 | 2103.827722 | 2251.292527 | 0.00813 | 0.217886 | 0.004041 | 2.874969 | 1.163028 | 2.08991 |
| CNAG_025 | - | translatior | 293.9436525 | 258.5399312 | 292.0822738 | 255.3783559 | 171.2906559 | 178.9937948 | 0.008136 | -0.49662 | 0.004044 | -2.87467 | 0.708765 | 2.089605 |
| CNAG_050 | - | hypothetic | 285.4026319 | 412.6610919 | 362.6729637 | 507.9386304 | 452.1013849 | 447.3120342 | 0.008193 | 0.395062 | 0.004074 | 2.872393 | 1.315 | 2.086582 |
| CNAG_011 | - | omega-6 | 3946.635131 | 3399.265077 | 3099.145369 | 3104.497043 | 2901.508279 | 2925.384407 | 0.008206 | -0.24167 | 0.004081 | -2.8718 | 0.845767 | 2.085873 |
| CNAG_005 | - | hypothetic | 1298.823191 | 1386.745521 | 1583.679567 | 1671.125351 | 1691.584743 | 1737.287255 | 0.008282 | 0.241835 | 0.004122 | 2.868638 | 1.182496 | 2.08185 |
| CNAG_038 | PRE3 | 20S protea | 2650.876781 | 2509.415064 | 2516.892565 | 2792.512419 | 3093.299447 | 2999.56869 | 0.008299 | 0.195506 | 0.004132 | 2.867935 | 1.145126 | 2.080991 |
| CNAG_027 | - | transcripti | 793.8809511 | 752.0055608 | 769.9263398 | 644.2191596 | 690.870512 | 588.5670172 | 0.008303 | -0.28343 | 0.004136 | -2.86763 | 0.821637 | 2.080781 |
| CNAG_077 | CTR1 | putative c | 1682.815514 | 1936.074372 | 2105.051735 | 1470.603125 | 1738.896429 | 1618.709055 | 0.008303 | -0.26095 | 0.004135 | -2.86768 | 0.834537 | 2.080781 |
| CNAG_004 | MAK32 | Maintenar | 496.0773134 | 510.57677 | 451.0908854 | 438.1773804 | 354.2677532 | 356.552293 | 0.008321 | -0.3588 | 0.004146 | -2.86686 | 0.779814 | 2.079841 |
| CNAG_044 | - | ribonuclea | 1482.991014 | 1477.13797 | 1623.054559 | 1269.919536 | 1361.050311 | 1332.271561 | 0.008357 | -0.22517 | 0.004165 | -2.86541 | 0.855494 | 2.077956 |
| CNAG_078 | - | hypothetic | 139.3887176 | 198.2471071 | 145.191301 | 211.8379263 | 242.6729467 | 239.5777674 | 0.008362 | 0.507932 | 0.004168 | 2.86513 | 1.42201 | 2.077674 |
| CNAG_128 | - | unspecifie | 3.825398665 | 5.514105183 | 4.390098114 | 14.22137371 | 18.45311709 | 19.67237309 | 0.008387 | 2.073406 | 0.004183 | 2.864002 | 4.208792 | 2.076416 |
| CNAG_049 | - | condensin | 1958.170482 | 1922.086082 | 1824.692231 | 1643.561086 | 1627.52441 | 1727.453872 | 0.008423 | -0.20609 | 0.004203 | -2.86249 | 0.866885 | 2.074524 |
| CNAG_022 | - | 2-oxoisova | 210.356518 | 175.7939367 | 194.1985153 | 318.3089512 | 271.0434637 | 227.615942 | 0.00847 | 0.47897 | 0.004229 | 2.860543 | 1.393749 | 2.072132 |
| CNAG_056 | - | hypothetic | 2.747264283 | 5.558741721 | 9.689383656 | 35.81985877 | 12.85905204 | 18.74739258 | 0.008493 | 2.00862 | 0.004242 | 2.859604 | 4.02397 | 2.070952 |
| CNAG_005 | - | saccharop | 7745.787661 | 7953.627064 | 7058.94373 | 8456.979247 | 8809.444507 | 8617.009574 | 0.008521 | 0.17067 | 0.004259 | 2.858307 | 1.125581 | 2.069495 |
| CNAG_120 | - | unspecifie | 27.16371574 | 13.20523961 | 31.01012124 | 47.03624782 | 53.86363245 | 55.17863093 | 0.00856 | 1.108839 | 0.00428 | 2.856778 | 2.15672 | 2.067508 |
| CNAG_077 | - | hypothetic | 35.60711748 | 48.66261226 | 65.09601365 | 76.41006223 | 82.99536175 | 111.5962951 | 0.008622 | 0.847566 | 0.004312 | 2.854418 | 1.799462 | 2.064387 |
| CNAG_001 | - | large subu | 375.6617099 | 312.4465265 | 349.7115711 | 280.7756056 | 256.8311199 | 261.0438894 | 0.008637 | -0.39428 | 0.004321 | -2.85372 | 0.760868 | 2.063644 |
| CNAG_036 | - | hypothetic | 13.27770724 | 12.06717691 | 21.40942561 | 38.71354818 | 30.27678656 | 45.07344633 | 0.008639 | 1.281247 | 0.004323 | 2.853564 | 2.43049 | 2.063537 |

| | | | | | | | | | | | | | | |
|---|---|---|---|---|---|---|---|---|---|---|---|---|---|---|
| CNAG_052 | - | 20S protea | 3865.222141 | 3816.258416 | 4222.009854 | 3541.081187 | 3568.338313 | 3484.880755 | 0.008698 | -0.18368 | 0.004354 | -2.85131 | 0.880457 | 2.060559 |
| CNAG_014 | - | cysteine d | 1328.77002 | 1315.492517 | 1283.981962 | 1124.440227 | 1105.665965 | 1172.382521 | 0.008702 | -0.22228 | 0.004357 | -2.85109 | 0.857212 | 2.060365 |
| CNAG_130 | - | unspecifie | 101.5952733 | 126.969204 | 139.4867971 | 75.81751186 | 86.083146 | 74.61333234 | 0.008719 | -0.65406 | 0.004366 | -2.8504 | 0.635491 | 2.059529 |
| CNAG_060 | - | trans-acor | 431.0252308 | 348.1919908 | 431.9618009 | 477.1360874 | 534.1187123 | 558.1989257 | 0.00874 | 0.3587 | 0.004378 | 2.84957 | 1.28227 | 2.058505 |
| CNAG_001 | PBY101 | 5'/3'-nucle | 324.8704655 | 324.4167435 | 347.7209172 | 425.9698434 | 447.0339666 | 405.1464131 | 0.008904 | 0.344565 | 0.004463 | 2.843415 | 1.269768 | 2.050427 |
| CNAG_023 | - | hypothetic | 276.5119773 | 268.193231 | 266.5426835 | 198.9020271 | 196.4410455 | 218.7890317 | 0.008966 | -0.4196 | 0.004497 | -2.84102 | 0.74763 | 2.04738 |
| CNAG_007 | - | protein sy | 1492.492026 | 1351.214165 | 1502.670184 | 1735.786762 | 1635.451697 | 1763.926954 | 0.009095 | 0.224843 | 0.004564 | 2.836314 | 1.16865 | 2.041183 |
| CNAG_074 | - | hypothetic | 45.06417358 | 104.5123922 | 81.17112043 | 139.1872417 | 125.3077719 | 125.2611232 | 0.009125 | 0.745712 | 0.004581 | 2.835131 | 1.676802 | 2.039786 |
| CNAG_127 | - | unspecifie | 3.897856692 | 5.555653076 | 9.685635824 | 29.93429303 | 18.26881659 | 16.58902496 | 0.009125 | 1.874956 | 0.00458 | 2.835159 | 3.667903 | 2.039786 |
| CNAG_130 | - | unspecifie | 32.01895229 | 48.56775634 | 36.42435289 | 90.56202594 | 54.29157398 | 75.21974312 | 0.009234 | 0.903556 | 0.004639 | 2.831094 | 1.870671 | 2.034618 |
| CNAG_003 | - | vacuolar p | 2678.985653 | 2863.843229 | 2693.922258 | 3015.908068 | 3164.826943 | 3261.54387 | 0.009275 | 0.182295 | 0.004662 | 2.829509 | 1.134687 | 2.032676 |
| CNAG_079 | - | hypothetic | 43.41014038 | 37.85829742 | 25.81100396 | 81.12126875 | 64.05901681 | 57.64964269 | 0.009305 | 0.928558 | 0.004678 | 2.828401 | 1.903372 | 2.031279 |
| CNAG_065 | - | ribosome | 348.8930111 | 309.2002171 | 348.6257295 | 276.1293641 | 246.4686493 | 254.3799361 | 0.009397 | -0.38874 | 0.00473 | -2.82485 | 0.763795 | 2.026989 |
| CNAG_130 | - | unspecifie | 1.3785892 | 0.081532092 | 1.154186329 | 9.71571142 | 12.48329194 | 7.691513955 | 0.009443 | 3.789326 | 0.004754 | 2.823218 | 13.82613 | 2.024885 |
| CNAG_074 | - | hypothetic | 1677.95397 | 1746.434781 | 1617.793339 | 1470.040285 | 1329.859828 | 1550.914189 | 0.00945 | -0.22752 | 0.004759 | -2.8229 | 0.854102 | 2.024559 |
| CNAG_023 | CHI4 | endochitir | 830.6085754 | 1095.718125 | 873.5340639 | 1077.202706 | 1257.885079 | 1133.391209 | 0.009577 | 0.295184 | 0.004828 | 2.818309 | 1.227042 | 2.018766 |
| CNAG_023 | - | GTP-bindi | 4906.946548 | 5625.187134 | 5378.082636 | 4759.112279 | 4832.004244 | 4631.684805 | 0.009676 | -0.17643 | 0.00488 | -2.81486 | 0.884887 | 2.014313 |
| CNAG_020 | URE6 | urease acc | 932.9005893 | 867.3300899 | 993.8339172 | 790.5601939 | 738.1604169 | 817.1366521 | 0.009686 | -0.26722 | 0.004886 | -2.81446 | 0.83092 | 2.013877 |
| CNAG_015 | - | Myo-inosi | 8265.10282 | 7504.286968 | 7025.813443 | 6680.676986 | 6855.564591 | 6681.031806 | 0.009699 | -0.18878 | 0.004895 | -2.81386 | 0.877349 | 2.013275 |
| CNAG_014 | HSP12 | heat shocl | 7665.563512 | 10534.05943 | 6634.635783 | 14150.10841 | 11517.0406 | 12394.75177 | 0.009705 | 0.603109 | 0.004899 | 2.813579 | 1.518986 | 2.013002 |
| CNAG_040 | FZC18 | hypothetic | 455.3135873 | 496.5710769 | 452.1375429 | 392.8702082 | 382.8439441 | 357.637766 | 0.009747 | -0.32582 | 0.004923 | -2.81205 | 0.797844 | 2.011147 |
| CNAG_075 | TRS130 | Trafficking | 2498.870268 | 2528.66868 | 2336.598696 | 2138.558621 | 2149.394888 | 2237.020009 | 0.009806 | -0.1894 | 0.004956 | -2.80987 | 0.876971 | 2.008522 |
| CNAG_078 | - | hypothetic | 3.808516795 | 4.444576256 | 4.382627963 | 12.5900807 | 17.58630451 | 19.64022655 | 0.009892 | 2.110895 | 0.005001 | 2.806983 | 4.319592 | 2.004732 |
| CNAG_054 | - | NADPH de | 706.450212 | 814.3982313 | 715.5657326 | 619.3011747 | 622.6403707 | 625.1199289 | 0.009897 | -0.27492 | 0.005005 | -2.80674 | 0.826497 | 2.004506 |
| CNAG_027 | RPP1 | hypothetic | 457.9569689 | 446.164594 | 478.8913314 | 570.6851344 | 620.4364761 | 537.289138 | 0.009928 | 0.30698 | 0.005025 | 2.805426 | 1.237116 | 2.003129 |
| CNAG_000 | NDX6 | A/G-speci | 1151.189722 | 1160.288853 | 984.3603217 | 936.8820843 | 971.177099 | 874.9590559 | 0.009928 | -0.25983 | 0.005023 | -2.80555 | 0.835185 | 2.003129 |
| CNAG_025 | - | hypothetic | 559.5932598 | 628.1780601 | 572.7683 | 751.3615043 | 727.2808913 | 679.3849103 | 0.009954 | 0.279459 | 0.00504 | 2.804495 | 1.213739 | 2.002007 |
| CNAG_063 | SBH1 | protein tra | 775.3420188 | 773.5772866 | 903.1538398 | 665.1856465 | 657.1012603 | 714.9998437 | 0.009968 | -0.28297 | 0.005048 | -2.80398 | 0.821897 | 2.001412 |
| CNAG_027 | - | hypothetic | 441.474176 | 363.2452529 | 391.4809828 | 491.2497645 | 483.1022239 | 563.7135215 | 0.009981 | 0.347333 | 0.005056 | 2.803455 | 1.272207 | 2.000816 |
| CNAG_075 | - | prenylatec | 54.20832415 | 77.65532486 | 69.43507347 | 96.66476284 | 117.6885484 | 112.9864669 | 0.009982 | 0.686197 | 0.005057 | 2.803362 | 1.609037 | 2.000795 |
| CNAG_016 | - | mRNA cap | 378.4178697 | 335.1903599 | 312.5981522 | 457.2014615 | 432.2702303 | 431.7525111 | 0.010015 | 0.349372 | 0.005077 | 2.802134 | 1.274006 | 1.999365 |
| CNAG_040 | - | Cytochron | 110.1613899 | 131.5405256 | 129.1301417 | 166.7364075 | 178.7632376 | 191.7998768 | 0.010038 | 0.517352 | 0.005092 | 2.801149 | 1.431326 | 1.998338 |
| CNAG_027 | MAL13 | Fungal Zn | 821.2081372 | 949.2520782 | 920.3971765 | 1055.4986 | 1061.906473 | 1104.48727 | 0.010111 | 0.244781 | 0.005131 | 2.798671 | 1.184913 | 1.995213 |
| CNAG_079 | - | hypothetic | 11.92034345 | 11.96446302 | 1.247089807 | 21.22074129 | 31.60783902 | 26.38412114 | 0.010123 | 1.747362 | 0.005139 | 2.798206 | 3.357442 | 1.994693 |
| CNAG_035 | - | AP-2 com | 2802.49701 | 2926.219175 | 2609.617594 | 2332.287016 | 2410.743304 | 2592.243519 | 0.010129 | -0.19978 | 0.005143 | -2.79792 | 0.870682 | 1.994418 |
| CNAG_064 | DPP101 | hypothetic | 433.4997551 | 489.199278 | 440.541629 | 569.1335038 | 653.3114014 | 510.6945866 | 0.010135 | 0.331667 | 0.005147 | 2.797669 | 1.258467 | 1.99418 |
| CNAG_015 | STE14 | protein-S- | 133.4469817 | 129.4560528 | 150.4432258 | 207.4530273 | 241.1832342 | 158.827905 | 0.010211 | 0.540666 | 0.005188 | 2.795104 | 1.454644 | 1.990941 |
| CNAG_069 | APH2 | Phytase | 2434.62493 | 2277.682963 | 2392.012948 | 2040.473462 | 2108.012993 | 2140.444019 | 0.010215 | -0.19137 | 0.005192 | -2.79488 | 0.875773 | 1.990746 |
| CNAG_033 | - | hypothetic | 240.9082363 | 294.1841503 | 268.8338257 | 336.3753849 | 336.4946266 | 388.3000223 | 0.010229 | 0.388054 | 0.0052 | 2.794369 | 1.308627 | 1.990163 |
| CNAG_077 | - | 3-ketodihy | 505.5901616 | 525.7343478 | 594.941471 | 377.773421 | 451.8761019 | 468.5373828 | 0.010233 | -0.34076 | 0.005203 | -2.79417 | 0.789624 | 1.990005 |
| CNAG_041 | - | hypothetic | 45.05358969 | 92.72070892 | 90.7118608 | 117.0147592 | 125.3383213 | 137.3317436 | 0.01024 | 0.726342 | 0.005208 | 2.793864 | 1.654438 | 1.989694 |
| CNAG_129 | - | unspecifie | 35.40020006 | 42.1268758 | 30.04764956 | 80.3150656 | 68.46301007 | 53.2763999 | 0.010345 | 0.895034 | 0.005263 | 2.790482 | 1.859653 | 1.985258 |
| CNAG_126 | - | unspecifie | 44.55280284 | 83.68033154 | 74.4385326 | 36.89833787 | 35.92827206 | 36.8585872 | 0.010414 | -0.92268 | 0.005299 | -2.78827 | 0.527528 | 1.982389 |
| CNAG_020 | - | hypothetic | 919.842692 | 469.8024671 | 527.8554348 | 366.815488 | 319.5681321 | 359.9713036 | 0.010429 | -0.89331 | 0.005308 | -2.78771 | 0.538378 | 1.981752 |
| CNAG_009 | KRE6 | glucosidas | 3336.443735 | 3689.13809 | 3364.725196 | 3887.651286 | 4023.092284 | 3921.033357 | 0.01043 | 0.172726 | 0.00531 | 2.787618 | 1.127186 | 1.98173 |
| CNAG_079 | - | hypothetic | 287.7080359 | 187.512841 | 220.6876775 | 183.5225371 | 141.9825378 | 161.1567915 | 0.01046 | -0.53365 | 0.005326 | -2.78661 | 0.690803 | 1.980486 |
| CNAG_034 | - | efflux prot | 21.50206057 | 22.83824494 | 36.31541932 | 63.92208675 | 43.16353523 | 57.43118872 | 0.010512 | 1.029759 | 0.005356 | 2.784824 | 2.041683 | 1.978298 |
| CNAG_126 | - | unspecifie | 43.92901679 | 109.8979029 | 87.55939775 | 155.7210711 | 130.4740255 | 123.1130533 | 0.010614 | 0.759362 | 0.00541 | 2.781555 | 1.692742 | 1.974131 |
| CNAG_032 | - | universal s | 877.5574355 | 418.0856435 | 451.0983769 | 295.8645828 | 302.3643599 | 293.4160788 | 0.010614 | -0.99083 | 0.005412 | -2.78145 | 0.503187 | 1.974101 |
| CNAG_063 | - | hypothetic | 15.51162133 | 14.17888282 | 12.95882611 | 40.51053886 | 43.7448804 | 21.19205469 | 0.010633 | 1.32687 | 0.005423 | 2.7808 | 2.508579 | 1.973329 |
| CNAG_078 | - | hypothetic | 825.5554323 | 871.5775415 | 891.4823257 | 744.6661063 | 760.7075241 | 686.2332172 | 0.010677 | -0.25555 | 0.005447 | -2.77932 | 0.837667 | 1.971564 |
| CNAG_066 | - | hypothetic | 685.6578943 | 713.3206815 | 696.4596086 | 852.4462724 | 812.5968787 | 854.7124667 | 0.010677 | 0.251011 | 0.005447 | 2.779346 | 1.19004 | 1.971564 |
| CNAG_025 | GPX2 | glutathion | 1722.276827 | 1649.512608 | 1645.507992 | 1347.688075 | 1459.353788 | 1549.802637 | 0.010701 | -0.21922 | 0.005461 | -2.77851 | 0.859028 | 1.970579 |
| CNAG_018 | - | hypothetic | 603.9408363 | 659.4287077 | 609.024781 | 770.742639 | 765.2264012 | 734.8669155 | 0.010715 | 0.263344 | 0.005469 | 2.778017 | 1.200258 | 1.970024 |
| CNAG_028 | - | hypothetic | 920.1073375 | 912.5543352 | 982.1238351 | 815.4964169 | 793.4290587 | 794.9752121 | 0.010822 | -0.24373 | 0.005529 | -2.77447 | 0.84456 | 1.965704 |
| CNAG_076 | - | Triacylglyc | 1023.13972 | 1078.561588 | 1060.109249 | 1290.662249 | 1204.190307 | 1229.989648 | 0.010868 | 0.221091 | 0.005554 | 2.772999 | 1.165615 | 1.963841 |
| CNAG_063 | - | hypothetic | 627.0234575 | 653.9398258 | 683.5110213 | 543.809552 | 571.8048828 | 519.7029277 | 0.010896 | -0.28081 | 0.005571 | -2.77202 | 0.823129 | 1.962745 |
| CNAG_003 | - | hypothetic | 254.8922495 | 303.879504 | 275.2359992 | 359.0292876 | 341.6491256 | 387.2336382 | 0.010958 | 0.371272 | 0.005604 | 2.770089 | 1.293492 | 1.96027 |
| CNAG_000 | - | hypothetic | 42.61714963 | 56.21210306 | 81.05894509 | 88.07461258 | 108.1876783 | 111.792818 | 0.010998 | 0.7633 | 0.005626 | 2.768829 | 1.697368 | 1.958694 |
| CNAG_078 | - | hypothetic | 25.8723172 | 15.29947912 | 17.23027141 | 35.99420512 | 46.13032184 | 46.29812697 | 0.011038 | 1.139024 | 0.005649 | 2.767472 | 2.20232 | 1.957094 |
| CNAG_034 | - | hypothetic | 268.903843 | 287.7848414 | 322.1024344 | 404.3420117 | 385.7432386 | 354.0988021 | 0.011135 | 0.366952 | 0.0057 | 2.76456 | 1.289625 | 1.953318 |
| CNAG_039 | - | hydantoin | 1315.931006 | 1347.789414 | 1261.594236 | 1018.423535 | 1125.533731 | 1215.63413 | 0.011155 | -0.23937 | 0.005712 | -2.76389 | 0.847113 | 1.952532 |
| CNAG_027 | - | hypothetic | 1243.386697 | 1117.26575 | 1090.970862 | 926.6516027 | 1050.570875 | 955.9651995 | 0.011169 | -0.25097 | 0.005721 | -2.76333 | 0.840329 | 1.951996 |
| CNAG_047 | - | hypothetic | 124.1227417 | 157.3292514 | 116.4039361 | 191.0340216 | 196.0598119 | 183.0533021 | 0.011169 | 0.507562 | 0.00572 | 2.763397 | 1.421645 | 1.951996 |
| CNAG_066 | - | hypothetic | 27.92474089 | 3.490279184 | 4.487311907 | 34.75483239 | 38.39131892 | 38.50992775 | 0.011181 | 1.672265 | 0.005729 | 2.762893 | 3.187145 | 1.951514 |
| CNAG_025 | DAK102 | dihydroxy | 118.4066005 | 85.07153406 | 88.45306585 | 68.24484401 | 54.88224367 | 56.85350188 | 0.01125 | -0.72543 | 0.005766 | -2.7608 | 0.604817 | 1.948832 |
| CNAG_123 | - | unspecifie | 151.754843 | 191.563163 | 168.4098495 | 124.2340597 | 114.464158 | 121.1705167 | 0.011286 | -0.52076 | 0.00579 | -2.75945 | 0.697002 | 1.947447 |
| CNAG_002 | - | hypothetic | 783.6449037 | 756.4204691 | 675.180018 | 849.1770641 | 964.5766409 | 882.5027484 | 0.011286 | 0.268237 | 0.005789 | 2.759506 | 1.204335 | 1.947447 |

| | | | | | | | | | | | | | | |
|---|---|---|---|---|---|---|---|---|---|---|---|---|---|---|
| CNAG_004 | - | cyclin | 711.3854781 | 723.050591 | 788.1394314 | 829.6646933 | 938.6530378 | 927.9598346 | 0.011286 | 0.262551 | 0.005788 | 2.759538 | 1.199598 | 1.947447 |
| CNAG_001 | - | hypothetic | 92.46134451 | 98.11065821 | 59.9496989 | 147.0160302 | 124.3788087 | 124.2047671 | 0.011358 | 0.651234 | 0.005831 | 2.757149 | 1.570511 | 1.944706 |
| CNAG_044 | - | membrane | 25.97218432 | 21.72912603 | 22.54273693 | 48.80023313 | 48.63485522 | 44.23976628 | 0.011358 | 1.026024 | 0.005831 | 2.757163 | 2.036404 | 1.944706 |
| CNAG_007 | - | DNA repa | 1089.839908 | 1342.496132 | 1284.050747 | 1442.289194 | 1456.953206 | 1534.109094 | 0.011421 | 0.240209 | 0.005866 | 2.755176 | 1.181164 | 1.942292 |
| CNAG_023 | - | hypothetic | 742.72756 | 888.7497003 | 833.9021525 | 661.270437 | 738.3140012 | 656.2489571 | 0.011421 | -0.27599 | 0.005866 | -2.75519 | 0.825882 | 1.942292 |
| CNAG_032 | CAC1 | Adenylate | 3013.782582 | 2850.847991 | 2705.595182 | 2449.842951 | 2585.028895 | 2565.643197 | 0.011526 | -0.18858 | 0.005921 | -2.75211 | 0.877469 | 1.938323 |
| CNAG_031 | AGS1 | alpha-1,3- | 10710.59412 | 9791.759865 | 8264.08515 | 11375.67116 | 11782.75101 | 10745.2262 | 0.011546 | 0.221481 | 0.005933 | 2.751449 | 1.165929 | 1.937554 |
| CNAG_123 | - | unspecifie | 2.756624556 | 9.809697765 | 8.645924841 | 30.67448837 | 18.22838065 | 18.76705896 | 0.011656 | 1.784206 | 0.005992 | 2.748199 | 3.444289 | 1.933453 |
| CNAG_131 | - | unspecifie | 0.271793585 | 4.447718863 | 6.486875447 | 15.95457465 | 8.515833925 | 28.16959459 | 0.011662 | 2.339103 | 0.005997 | 2.747948 | 5.059881 | 1.933224 |
| CNAG_01( | - | DNA misn | 385.6454736 | 428.8859847 | 429.8377214 | 506.8248048 | 491.7142504 | 567.0730304 | 0.011681 | 0.317844 | 0.006008 | 2.74735 | 1.246467 | 1.932535 |
| CNAG_015 | - | S-adenosy | 382.9071255 | 408.239425 | 406.2302061 | 365.8473665 | 300.83227 | 261.1552779 | 0.011696 | -0.38421 | 0.006017 | -2.74684 | 0.766201 | 1.931961 |
| CNAG_02( | WSP1 | wiskott-Al | 312.7960914 | 343.5438688 | 296.4467045 | 266.8278018 | 241.3273928 | 228.8868963 | 0.011847 | -0.38648 | 0.006101 | -2.74233 | 0.764996 | 1.926403 |
| CNAG_011 | - | nuclear pc | 1643.000213 | 1314.606485 | 1343.839362 | 1579.09968 | 2038.602394 | 1699.599997 | 0.011847 | 0.289485 | 0.006098 | 2.742457 | 1.222204 | 1.926403 |
| CNAG_121 | - | unspecifie | 7.207310639 | 3.388204742 | 2.274615252 | 14.2288762 | 13.92005903 | 24.99805421 | 0.011967 | 2.071043 | 0.006166 | 2.738845 | 4.201905 | 1.922008 |
| CNAG_015 | - | hypothetic | 746.0935363 | 796.1208242 | 739.0191568 | 594.2856238 | 612.2475929 | 698.2993758 | 0.012027 | -0.27542 | 0.006201 | -2.73696 | 0.826209 | 1.919825 |
| CNAG_128 | - | unspecifie | 0.132601705 | 0.064423749 | 1.134551593 | 6.430262608 | 8.915892552 | 7.617600397 | 0.01204 | 4.272763 | 0.006209 | 2.736539 | 19.32991 | 1.919373 |
| CNAG_029 | PLC2 | Phosphati | 325.1436744 | 257.425857 | 215.4664815 | 191.8934182 | 206.0126905 | 183.381124 | 0.012068 | -0.47463 | 0.006226 | -2.73563 | 0.719652 | 1.918378 |
| CNAG_073 | - | ubiquitin | 2711.575786 | 2608.51734 | 2425.202637 | 2830.662346 | 3024.248067 | 3086.135345 | 0.012068 | 0.191887 | 0.006225 | 2.735667 | 1.142257 | 1.918378 |
| CNAG_053 | - | transposas | 262.7030961 | 296.1777384 | 301.6925532 | 214.4811503 | 218.9090912 | 228.8001515 | 0.01217 | -0.39502 | 0.006286 | -2.73246 | 0.760476 | 1.914711 |
| CNAG_008 | - | hypothetic | 331.4936817 | 333.929614 | 348.6348089 | 263.570571 | 240.3945986 | 287.5856891 | 0.01224 | -0.37234 | 0.006327 | -2.73033 | 0.772528 | 1.912218 |
| CNAG_027 | - | peroxin-13 | 229.1012809 | 210.2492337 | 245.336245 | 302.2479792 | 317.6681337 | 288.562178 | 0.012294 | 0.39135 | 0.006358 | 2.728728 | 1.31162 | 1.910309 |
| CNAG_003 | - | chromatin | 2268.049193 | 2576.183909 | 2664.01634 | 2761.399986 | 2961.2905 | 3007.312264 | 0.012431 | 0.202557 | 0.00643 | 2.725005 | 1.150736 | 1.90551 |
| CNAG_08( | - | hypothetic | 26.04783912 | 18.55690417 | 33.13301872 | 46.97310629 | 56.47556526 | 54.12408061 | 0.01244 | 1.00192 | 0.006436 | 2.724681 | 2.002663 | 1.905187 |
| CNAG_058 | - | hypothetic | 921.40248 | 126.3929826 | 252.7803618 | 99.30037592 | 89.86341739 | 97.13024212 | 0.012463 | -2.21158 | 0.00645 | -2.72399 | 0.215898 | 1.904378 |
| CNAG_027 | - | sorting ne | 413.6217898 | 462.2562768 | 405.3567005 | 508.3181462 | 508.9714088 | 598.1257096 | 0.012633 | 0.319971 | 0.006541 | 2.719367 | 1.248305 | 1.898511 |
| CNAG_059 | - | ATP synth | 772.7468421 | 618.4870436 | 765.6000293 | 619.404924 | 501.7111959 | 613.9843895 | 0.012654 | -0.33039 | 0.006553 | -2.71873 | 0.79532 | 1.897771 |
| CNAG_057 | - | hypothetic | 1.363983309 | 1.172321606 | 0.065918203 | 8.040881213 | 9.754138872 | 9.786600956 | 0.012664 | 3.685758 | 0.00656 | 2.718392 | 12.86837 | 1.897436 |
| CNAG_062 | - | taurine ca | 96.22348285 | 111.1310402 | 139.7300062 | 160.5296197 | 152.7311963 | 206.0656716 | 0.012721 | 0.567472 | 0.006591 | 2.716826 | 1.481924 | 1.895483 |
| CNAG_039 | - | hypothetic | 99.33826565 | 119.5051266 | 162.8386793 | 93.18156405 | 75.59363756 | 75.73706964 | 0.012757 | -0.66006 | 0.006613 | -2.71574 | 0.632854 | 1.894262 |
| CNAG_13( | - | unspecifie | 65.81465206 | 79.8419179 | 80.07884306 | 113.9586737 | 107.9485264 | 129.5708402 | 0.012815 | 0.625242 | 0.006644 | 2.714163 | 1.542469 | 1.892297 |
| CNAG_01( | - | hypothetic | 321.4822278 | 373.9432611 | 393.5557727 | 435.9444029 | 464.2130155 | 486.0686702 | 0.012825 | 0.334731 | 0.006651 | 2.713814 | 1.261142 | 1.891942 |
| CNAG_018 | GLN3 | GATA fam | 32.96858263 | 31.38870769 | 24.70897676 | 61.36133901 | 61.65197132 | 46.58055935 | 0.012839 | 0.933989 | 0.00666 | 2.71337 | 1.910552 | 1.891462 |
| CNAG_023 | - | hypothetic | 238.3334827 | 204.8531261 | 205.9440814 | 269.3897372 | 336.8167473 | 271.9292068 | 0.012846 | 0.421104 | 0.006665 | 2.713122 | 1.338951 | 1.891239 |
| CNAG_00( | - | Sugar trar | 50.39052037 | 34.69505417 | 40.67000922 | 70.38282069 | 64.79656064 | 88.37527238 | 0.012872 | 0.824154 | 0.006681 | 2.712362 | 1.770497 | 1.890345 |
| CNAG_016 | - | CTD kinas | 761.676888 | 843.7063699 | 928.8886383 | 992.4432529 | 1068.039209 | 996.858351 | 0.012978 | 0.256237 | 0.006739 | 2.709485 | 1.194359 | 1.886781 |
| CNAG_122 | - | unspecifie | 13.42365954 | 21.74296774 | 33.11203376 | 54.39941728 | 68.08752208 | 30.11364328 | 0.012978 | 1.166158 | 0.006738 | 2.709516 | 2.244132 | 1.886781 |
| CNAG_028 | - | glucan en | 7852.88253 | 6962.3561 | 6757.058078 | 6383.385303 | 6356.113168 | 6477.849933 | 0.01299 | -0.18247 | 0.006748 | -2.70904 | 0.88119 | 1.886407 |
| CNAG_045 | - | hypothetic | 258.0056426 | 346.627744 | 243.1617889 | 219.2471833 | 225.0144717 | 179.0035053 | 0.013178 | -0.45711 | 0.006849 | -2.70411 | 0.728443 | 1.880157 |
| CNAG_008 | - | hypothetic | 2.601143298 | 2.284036419 | 3.299028063 | 17.13871377 | 13.19496246 | 8.864078127 | 0.013178 | 2.424781 | 0.006849 | 2.704116 | 5.369474 | 1.880157 |
| CNAG_028 | - | signal pep | 181.283461 | 203.6972148 | 179.2811315 | 247.7228315 | 279.8023545 | 236.4053998 | 0.013196 | 0.421682 | 0.006861 | 2.703499 | 1.339488 | 1.879564 |
| CNAG_01( | - | hypothetic | 140.3474352 | 142.3137284 | 124.9127121 | 191.794711 | 184.7401378 | 195.2019178 | 0.013196 | 0.4745 | 0.006861 | 2.70351 | 1.389437 | 1.879564 |
| CNAG_005 | VPS17 | vacuolar p | 284.1256454 | 379.2425442 | 299.792569 | 433.0846449 | 416.7768777 | 400.6872418 | 0.013328 | 0.362633 | 0.006933 | 2.700031 | 1.28577 | 1.875238 |
| CNAG_003 | - | transforme | 810.7419512 | 995.56301 | 937.4626428 | 1159.969553 | 1089.513785 | 1051.284728 | 0.013371 | 0.252327 | 0.006959 | 2.698798 | 1.191127 | 1.873834 |
| CNAG_124 | - | unspecifie | 47.58429465 | 62.13723836 | 41.54756081 | 22.80736535 | 15.93254463 | 35.56676017 | 0.01342 | -1.07177 | 0.006991 | -2.69726 | 0.475735 | 1.872233 |
| CNAG_00( | - | membrane | 1639.406765 | 1549.33948 | 1673.203935 | 1461.527463 | 1449.028616 | 1346.755285 | 0.013548 | -0.20737 | 0.007061 | -2.69394 | 0.866113 | 1.868109 |
| CNAG_015 | SCP1 | sterol regu | 1450.603635 | 1716.35236 | 1551.753889 | 1752.749267 | 1861.51911 | 1911.574492 | 0.013576 | 0.213353 | 0.007079 | 2.693117 | 1.15938 | 1.86724 |
| CNAG_078 | - | hypothetic | 243.1444339 | 268.3369551 | 240.0581809 | 338.9091622 | 319.2750327 | 320.729526 | 0.013613 | 0.364751 | 0.0071 | 2.692133 | 1.28766 | 1.866061 |
| CNAG_126 | - | unspecifie | 46.98612259 | 26.14504824 | 54.44186684 | 82.34784187 | 85.84660078 | 64.34321998 | 0.013628 | 0.856862 | 0.007109 | 2.691672 | 1.811095 | 1.865562 |
| CNAG_121 | - | unspecifie | 43.61234282 | 32.19019082 | 27.68446999 | 14.29809351 | 19.68472701 | 10.29323158 | 0.013673 | -1.26295 | 0.007134 | -2.69051 | 0.416692 | 1.864152 |
| CNAG_002 | - | hypothetic | 92.57704658 | 109.9588551 | 86.53720217 | 143.5728661 | 157.3582066 | 128.7190536 | 0.013809 | 0.561831 | 0.007209 | 2.687048 | 1.476142 | 1.859851 |
| CNAG_014 | - | monooxyg | 406.1308364 | 368.4488988 | 398.7693726 | 331.4248881 | 269.6864832 | 324.2505786 | 0.013854 | -0.3584 | 0.007234 | -2.68586 | 0.780028 | 1.858413 |
| CNAG_025 | - | nuclear pc | 8959.946621 | 8388.846627 | 7723.275122 | 6966.982549 | 7599.963297 | 7730.083196 | 0.013872 | -0.18468 | 0.007245 | -2.68537 | 0.879844 | 1.857872 |
| CNAG_12( | - | unspecifie | 196.100708 | 188.541712 | 259.9621364 | 163.1965514 | 138.5675797 | 161.1045802 | 0.01394 | -0.49245 | 0.007286 | -2.6835 | 0.710815 | 1.855744 |
| CNAG_027 | - | dephosph | 1037.904944 | 916.8933343 | 1015.190605 | 709.330169 | 884.9435162 | 879.2895 | 0.01394 | -0.27972 | 0.007284 | -2.68355 | 0.823752 | 1.855744 |
| CNAG_078 | - | hypothetic | 7.177596361 | 2.310891856 | 2.267486531 | 12.61097584 | 22.16234745 | 14.29415396 | 0.014198 | 2.106605 | 0.007424 | 2.677198 | 4.306766 | 1.847773 |
| CNAG_057 | - | hypothetic | 374.9774684 | 351.3438237 | 329.6523606 | 475.1157202 | 517.0402949 | 378.640475 | 0.014361 | 0.362527 | 0.007513 | 2.673214 | 1.285676 | 1.842815 |
| CNAG_034 | CTS1 | calcineurir | 92.57740929 | 96.02472271 | 98.21855126 | 113.5092979 | 160.8451195 | 158.4486012 | 0.014399 | 0.576419 | 0.007535 | 2.672246 | 1.491143 | 1.841663 |
| CNAG_06( | - | hypothetic | 1146.704469 | 1208.823374 | 1233.823702 | 1073.93693 | 999.5568717 | 1045.849695 | 0.0144 | -0.21763 | 0.007537 | -2.67214 | 0.859978 | 1.841624 |
| CNAG_001 | - | etoposide | 369.1950925 | 393.3070333 | 340.3136397 | 459.4037266 | 452.0897854 | 473.8957293 | 0.014411 | 0.313895 | 0.007544 | 2.671819 | 1.243059 | 1.841314 |
| CNAG_055 | PRY1 | septum fo | 600.0418322 | 602.197177 | 566.2555932 | 533.8813985 | 421.5107021 | 490.7876333 | 0.014413 | -0.30697 | 0.007547 | -2.67169 | 0.80834 | 1.841246 |
| CNAG_129 | - | unspecifie | 75.3855572 | 64.52896204 | 62.83640173 | 40.07173116 | 41.17572588 | 35.77473723 | 0.014444 | -0.80894 | 0.007565 | -2.6709 | 0.5708 | 1.840324 |
| CNAG_041 | - | actin bind | 5097.086819 | 4824.71854 | 4705.188928 | 4394.870291 | 4392.939308 | 4441.8104 | 0.014465 | -0.16047 | 0.007578 | -2.67031 | 0.894733 | 1.839671 |
| CNAG_055 | - | septum-pr | 941.35414 | 842.6887179 | 1007.814927 | 1146.664854 | 1054.076208 | 1157.765193 | 0.014515 | 0.251225 | 0.007608 | 2.669001 | 1.190217 | 1.838175 |
| CNAG_042 | - | peroxisom | 244.3396212 | 232.8802688 | 292.2226423 | 330.2170451 | 344.3631219 | 334.0351374 | 0.014555 | 0.376649 | 0.00763 | 2.668015 | 1.298323 | 1.837 |
| CNAG_037 | - | hypothetic | 196.7083784 | 271.5956834 | 307.1384237 | 366.2135978 | 330.461374 | 352.8725063 | 0.014619 | 0.422928 | 0.007666 | 2.666459 | 1.340646 | 1.835091 |
| CNAG_048 | - | cytoplasm | 1700.41653 | 1661.494171 | 1850.312567 | 1973.835183 | 1956.33211 | 2139.136903 | 0.014633 | 0.204317 | 0.007675 | 2.666063 | 1.152141 | 1.834681 |
| CNAG_066 | - | Haloacid | 181.2105773 | 182.1796208 | 164.362169 | 228.9992337 | 275.5657829 | 218.6627287 | 0.0147 | 0.436762 | 0.007713 | 2.664368 | 1.353563 | 1.832694 |
| CNAG_031 | - | FAD depe | 28.22894423 | 58.89836109 | 54.144889 | 25.2965625 | 25.64679127 | 16.97035981 | 0.014714 | -1.09822 | 0.007726 | -2.6638 | 0.467094 | 1.832259 |

| | | | | | | | | | | | | | | |
|---|---|---|---|---|---|---|---|---|---|---|---|---|---|---|
| CNAG_041 | PKA2 | AGC/PKA | 473.164964 | 495.6936777 | 502.3561801 | 632.2950826 | 621.2162928 | 555.073659 | 0.014714 | 0.283359 | 0.007726 | 2.663816 | 1.217025 | 1.832259 |
| CNAG_042 | - | hypothetic | 823.0932891 | 787.5607058 | 789.1377074 | 698.7676937 | 645.0171525 | 696.1426158 | 0.014724 | -0.24986 | 0.007733 | -2.66351 | 0.84098 | 1.831986 |
| CNAG_021 | - | hypothetic | 190.372138 | 223.9942515 | 249.3699633 | 158.4107504 | 170.6388513 | 161.1379555 | 0.014769 | -0.45195 | 0.007759 | -2.66239 | 0.731054 | 1.830642 |
| CNAG_006 | - | transcripti | 1061.661179 | 1402.675463 | 1382.0141 | 1094.882806 | 1050.443928 | 1082.503177 | 0.01482 | -0.26741 | 0.007787 | -2.66115 | 0.830811 | 1.829149 |
| CNAG_056 | - | glutaminy | 1452.6094 | 1525.568791 | 1447.150408 | 1254.3914 | 1207.40962 | 1388.813152 | 0.014832 | -0.21569 | 0.007795 | -2.6608 | 0.861134 | 1.828798 |
| CNAG_012 | IPK1 | inositol-pe | 440.5294243 | 503.2101484 | 488.4949813 | 612.0350964 | 618.6585664 | 541.7529714 | 0.014838 | 0.292618 | 0.007802 | 2.660514 | 1.224861 | 1.828629 |
| CNAG_049 | - | hypothetic | 168.3470624 | 167.0945932 | 146.2440716 | 218.1915049 | 247.0588201 | 199.7938967 | 0.014877 | 0.449751 | 0.007828 | 2.659401 | 1.365804 | 1.827496 |
| CNAG_037 | - | hypothetic | 271.2879067 | 340.5048932 | 317.8730489 | 433.9703219 | 395.1906106 | 370.7569917 | 0.014895 | 0.353953 | 0.007839 | 2.658906 | 1.278058 | 1.826958 |
| CNAG_029 | - | hypothetic | 1255.472372 | 1226.198013 | 1288.315295 | 1407.206215 | 1470.766415 | 1511.925703 | 0.014927 | 0.204023 | 0.007858 | 2.658113 | 1.151906 | 1.826037 |
| CNAG_012 | - | charged n | 671.6543731 | 672.4080429 | 716.6976347 | 774.4247942 | 816.0732308 | 901.253656 | 0.014939 | 0.259576 | 0.007866 | 2.657749 | 1.197127 | 1.825669 |
| CNAG_127 | - | unspecifie | 5.1311185 | 14.10142431 | 10.79256458 | 31.94151956 | 26.13979178 | 23.19803479 | 0.014983 | 1.451313 | 0.007893 | 2.656606 | 2.734568 | 1.824398 |
| CNAG_022 | - | niemann-I | 2151.147593 | 2315.41797 | 2072.198886 | 2526.484544 | 2392.761331 | 2604.323959 | 0.015048 | 0.187563 | 0.007931 | 2.654983 | 1.138839 | 1.82251 |
| CNAG_123 | - | unspecifie | 35.35226543 | 41.03361642 | 24.73789627 | 65.13316291 | 76.48798833 | 48.85691745 | 0.01506 | 0.903599 | 0.007939 | 2.654652 | 1.870727 | 1.822184 |
| CNAG_122 | - | unspecifie | 5.117000308 | 8.787628507 | 14.9864954 | 21.18921127 | 25.28260637 | 31.77659738 | 0.015179 | 1.497162 | 0.00801 | 2.651665 | 2.822869 | 1.818743 |
| CNAG_073 | - | hypothetic | 19.0419461 | 24.87110458 | 14.05566312 | 41.75130021 | 40.82101177 | 41.94045866 | 0.015246 | 1.083796 | 0.00805 | 2.649956 | 2.119605 | 1.816847 |
| CNAG_011 | - | hypothetic | 422.7789839 | 466.4776998 | 574.6080753 | 338.0922389 | 448.5665877 | 349.9042466 | 0.015449 | -0.38084 | 0.008161 | -2.64533 | 0.767988 | 1.811107 |
| CNAG_066 | - | hypothetic | 4040.504656 | 3733.395624 | 4000.30534 | 4369.38417 | 4304.197196 | 4840.145515 | 0.015588 | 0.183382 | 0.008236 | 2.642226 | 1.135543 | 1.807223 |
| CNAG_126 | - | unspecifie | 2.565026982 | 2.267419187 | 1.170491985 | 16.45124826 | 12.36457677 | 5.613566321 | 0.015592 | 2.70702 | 0.008242 | 2.641972 | 6.529716 | 1.807099 |
| CNAG_006 | CDC7 | CDC7 prot | 1398.870551 | 1395.232557 | 1397.016598 | 1200.702038 | 1192.79021 | 1285.597052 | 0.015592 | -0.20358 | 0.008241 | -2.64205 | 0.868391 | 1.807099 |
| CNAG_010 | - | hypothetic | 156.7038451 | 132.71134 | 167.4889956 | 247.4364783 | 199.350881 | 194.2274276 | 0.015625 | 0.476303 | 0.008262 | 2.641184 | 1.391174 | 1.806188 |
| CNAG_056 | CML1 | EF-hand c | 700.5266867 | 730.4058789 | 676.1067945 | 538.2116566 | 623.5744462 | 607.3301638 | 0.015679 | -0.26747 | 0.008292 | -2.63992 | 0.830773 | 1.804673 |
| CNAG_032 | - | hypothetic | 659.880728 | 632.5279603 | 600.5059707 | 774.6220375 | 807.5608976 | 709.3839646 | 0.015684 | 0.260716 | 0.008297 | 2.639747 | 1.198073 | 1.804547 |
| CNAG_020 | - | Short-chai | 691.3972135 | 736.9574732 | 480.1117116 | 807.3184208 | 790.2168212 | 841.3351839 | 0.015757 | 0.339989 | 0.008338 | 2.638082 | 1.265747 | 1.802515 |
| CNAG_126 | - | unspecifie | 169.4108367 | 232.5113319 | 219.5520972 | 150.6707185 | 129.9187831 | 167.7006144 | 0.015781 | -0.48905 | 0.008352 | -2.6375 | 0.712493 | 1.80187 |
| CNAG_080 | PHO85 | CMGC/CD | 1500.556361 | 1596.702454 | 1632.678868 | 1414.033882 | 1409.346482 | 1346.732542 | 0.015922 | -0.19663 | 0.008434 | -2.63417 | 0.872585 | 1.79801 |
| CNAG_031 | FZC24 | nuclear pr | 1572.896262 | 1580.558984 | 1652.940955 | 1303.337778 | 1389.475544 | 1505.384178 | 0.01606 | -0.21008 | 0.008516 | -2.63091 | 0.864487 | 1.794246 |
| CNAG_122 | - | unspecifie | 84.57097405 | 133.5999937 | 100.383054 | 143.3553976 | 169.4028109 | 159.6671496 | 0.016176 | 0.557619 | 0.008581 | 2.62832 | 1.471838 | 1.791133 |
| CNAG_038 | ASG1 | nuclear pr | 1298.648861 | 1280.049807 | 1241.421005 | 1472.656935 | 1572.631517 | 1405.451941 | 0.016205 | 0.204874 | 0.0086 | 2.627558 | 1.152586 | 1.79036 |
| CNAG_067 | LMP1 | Low matir | 1089.691784 | 1095.82774 | 1172.060124 | 1260.11664 | 1318.962549 | 1346.514672 | 0.016337 | 0.210292 | 0.008674 | 2.624633 | 1.156922 | 1.786828 |
| CNAG_059 | PAK1 | ste/ste20/ | 777.8733106 | 828.570944 | 719.9576997 | 910.7370796 | 956.7400289 | 905.8295787 | 0.01636 | 0.238854 | 0.008688 | 2.624084 | 1.180055 | 1.786226 |
| CNAG_051 | - | GABA peri | 228.0691649 | 133.6787419 | 174.8030435 | 102.1339433 | 143.1219338 | 116.7785166 | 0.016403 | -0.5853 | 0.008713 | -2.62311 | 0.666509 | 1.785089 |
| CNAG_049 | - | hypothetic | 885.106978 | 935.1434615 | 930.9588025 | 796.0315125 | 827.1452911 | 739.5147771 | 0.016434 | -0.23461 | 0.008732 | -2.62237 | 0.849918 | 1.784245 |
| CNAG_003 | - | 1-phospha | 433.3954294 | 440.7324722 | 398.960308 | 582.6258682 | 529.7474936 | 478.4864146 | 0.016473 | 0.307009 | 0.008754 | 2.621505 | 1.23714 | 1.783239 |
| CNAG_018 | - | hypothetic | 15.24399113 | 5.577396084 | 0.145472468 | 16.32447425 | 34.40205426 | 25.25956258 | 0.016603 | 1.887503 | 0.008829 | 2.618586 | 3.699942 | 1.779822 |
| CNAG_006 | - | cytoplasm | 2383.474206 | 2310.079543 | 2338.780206 | 2614.357276 | 2713.721784 | 2653.227821 | 0.016637 | 0.167246 | 0.008852 | 2.617715 | 1.122913 | 1.778912 |
| CNAG_063 | - | Alcohol de | 825.6359641 | 928.6688433 | 962.9130235 | 739.1110923 | 737.3229759 | 825.9824804 | 0.01668 | -0.25344 | 0.008877 | -2.61676 | 0.838891 | 1.777796 |
| CNAG_061 | - | hypothetic | 13.34792672 | 27.00427936 | 18.28609677 | 42.52873045 | 33.69467917 | 51.7179517 | 0.01675 | 1.107007 | 0.008918 | 2.615187 | 2.153983 | 1.775995 |
| CNAG_016 | GCN3 | translatior | 644.5005675 | 643.1832548 | 674.9925392 | 572.6968328 | 529.442993 | 552.9559882 | 0.016891 | -0.26173 | 0.008997 | -2.61216 | 0.834089 | 1.772341 |
| CNAG_052 | - | hypothetic | 420.5268134 | 367.5486384 | 421.3019716 | 546.7771433 | 523.7633258 | 454.0691254 | 0.016983 | 0.318346 | 0.009048 | 2.610228 | 1.2469 | 1.769993 |
| CNAG_079 | - | hypothetic | 0.087003557 | 0.042411767 | 0.036649734 | 4.026313072 | 7.217606137 | 3.306550816 | 0.017092 | 4.648603 | 0.009108 | 2.607956 | 25.08239 | 1.767208 |
| CNAG_044 | - | thiamine p | 382.9415759 | 423.3089944 | 391.3412981 | 321.9694056 | 321.5836014 | 327.6020821 | 0.017119 | -0.31845 | 0.009125 | -2.60734 | 0.801932 | 1.766525 |
| CNAG_078 | - | hypothetic | 75.1537513 | 108.8253803 | 69.51013695 | 143.8269708 | 148.8027203 | 103.2982477 | 0.017133 | 0.630835 | 0.009134 | 2.606981 | 1.548461 | 1.766171 |
| CNAG_001 | - | auxin-indu | 459.3145623 | 319.9834383 | 290.1224711 | 254.896741 | 257.6796919 | 286.5112959 | 0.017233 | -0.43872 | 0.00919 | -2.6049 | 0.737791 | 1.76363 |
| CNAG_042 | - | hypothetic | 0.153126723 | 2.232386182 | 0.064151137 | 10.67796433 | 6.087473946 | 9.776451206 | 0.017339 | 3.634065 | 0.009248 | 2.602733 | 12.41545 | 1.760985 |
| CNAG_061 | - | hypothetic | 312.1980611 | 416.9823755 | 381.8555257 | 453.1013879 | 529.0811363 | 440.6977322 | 0.017371 | 0.344195 | 0.009268 | 2.602022 | 1.269443 | 1.760184 |
| CNAG_002 | - | hypothetic | 210.3222412 | 202.6390441 | 167.5900303 | 261.8036632 | 284.0979984 | 236.432887 | 0.017383 | 0.416354 | 0.009276 | 2.601704 | 1.33455 | 1.759881 |
| CNAG_057 | - | acyl-prote | 1143.232721 | 1235.753891 | 1261.54668 | 1084.041539 | 1070.349955 | 1020.354771 | 0.01743 | -0.21229 | 0.009304 | -2.60069 | 0.863167 | 1.758702 |
| CNAG_039 | NUP75 | nuclear pc | 2289.846946 | 2197.942633 | 2243.786815 | 1996.163729 | 2069.236397 | 1959.527354 | 0.017432 | -0.17517 | 0.009307 | -2.60058 | 0.885664 | 1.758654 |
| CNAG_039 | - | hypothetic | 43.66016398 | 51.86686643 | 59.79661224 | 109.228758 | 68.98534481 | 82.00494051 | 0.017539 | 0.744848 | 0.009366 | 2.598404 | 1.675797 | 1.756003 |
| CNAG_044 | - | hypothetic | 273.1691542 | 282.222225 | 310.2121813 | 234.067107 | 218.8959064 | 223.2812681 | 0.017668 | -0.37301 | 0.009437 | -2.59581 | 0.772172 | 1.752819 |
| CNAG_031 | - | hypothetic | 3351.826113 | 4434.702441 | 3138.750674 | 5571.629596 | 5132.480566 | 4440.629277 | 0.017698 | 0.457612 | 0.009455 | 2.595137 | 1.373267 | 1.752071 |
| CNAG_122 | - | unspecifie | 19.1328976 | 22.79559753 | 29.9318155 | 45.53758896 | 41.54132036 | 54.0241795 | 0.017865 | 0.985186 | 0.009551 | 2.59168 | 1.979568 | 1.748002 |
| CNAG_020 | - | hypothetic | 102.7678014 | 155.8721526 | 110.8689475 | 83.7121076 | 87.81253081 | 72.41965847 | 0.01791 | -0.61436 | 0.00958 | -2.59063 | 0.65322 | 1.746904 |
| CNAG_015 | - | hypothetic | 71.46230626 | 70.14832819 | 55.63462665 | 122.2122251 | 111.6123138 | 81.0974709 | 0.017939 | 0.664587 | 0.009599 | 2.589938 | 1.585114 | 1.746201 |
| CNAG_016 | - | hypothetic | 1392.170052 | 1505.217175 | 1508.001773 | 1611.858841 | 1723.506102 | 1763.929987 | 0.017986 | 0.195538 | 0.009626 | 2.588966 | 1.145151 | 1.745074 |
| CNAG_040 | - | pumilio 2 | 502.4238513 | 571.0932163 | 575.9262354 | 714.8278128 | 670.3306288 | 627.2209177 | 0.018045 | 0.272318 | 0.00966 | 2.587762 | 1.207747 | 1.743655 |
| CNAG_131 | - | unspecifie | 10.90462457 | 12.02172493 | 16.09106323 | 27.52122687 | 33.15266912 | 31.93718115 | 0.018184 | 1.244642 | 0.009737 | 2.585025 | 2.369598 | 1.740305 |
| CNAG_030 | - | hypothetic | 157.7102833 | 45.39870153 | 45.97912923 | 48.68699451 | 43.7012506 | 25.94586397 | 0.018212 | -1.11132 | 0.009754 | -2.58442 | 0.462872 | 1.73964 |
| CNAG_000 | - | hypothetic | 486.0861644 | 575.3809594 | 548.2131127 | 621.929381 | 656.5344447 | 687.0329383 | 0.01826 | 0.273027 | 0.009784 | 2.583355 | 1.208341 | 1.738499 |
| CNAG_040 | - | hypothetic | 262.0558711 | 178.8713703 | 192.9965867 | 162.4353667 | 150.7320846 | 144.5242388 | 0.018421 | -0.48759 | 0.009878 | -2.58008 | 0.713218 | 1.734676 |
| CNAG_057 | RIB4 | 6,7-dimetl | 190.711017 | 193.0243617 | 254.8786042 | 251.371786 | 323.8329048 | 291.8128781 | 0.018446 | 0.425689 | 0.009893 | 2.579549 | 1.343214 | 1.734105 |
| CNAG_056 | - | hypothetic | 468.2656703 | 425.6514982 | 315.8644671 | 569.4140375 | 511.6263895 | 488.4227359 | 0.018527 | 0.360998 | 0.009941 | 2.577579 | 1.284314 | 1.732204 |
| CNAG_124 | - | unspecifie | 28.9181423 | 32.12969661 | 30.78452538 | 14.39393894 | 14.42322153 | 11.35038112 | 0.018552 | -1.22137 | 0.009957 | -2.57732 | 0.428875 | 1.731604 |
| CNAG_077 | - | hypothetic | 59.99108083 | 37.69898352 | 53.12111451 | 25.24705799 | 8.949264152 | 35.55074861 | 0.018576 | -1.14749 | 0.009974 | -2.57672 | 0.451411 | 1.731042 |
| CNAG_010 | ZFC4 | hypothetic | 470.6310792 | 425.6669108 | 368.0681367 | 544.3262071 | 520.2416668 | 517.248085 | 0.01863 | 0.308554 | 0.010006 | 2.575636 | 1.238466 | 1.729782 |
| CNAG_078 | - | insulysin | 1322.895797 | 1265.942948 | 1228.540523 | 1153.335301 | 1090.151059 | 1099.152521 | 0.018719 | -0.20631 | 0.010058 | -2.57383 | 0.866754 | 1.727717 |
| CNAG_036 | - | hypothetic | 216.3010035 | 219.9102894 | 238.9405902 | 325.0603534 | 292.5580783 | 267.5293995 | 0.01888 | 0.378112 | 0.010147 | 2.570787 | 1.29964 | 1.723993 |
| CNAG_126 | - | unspecifie | 150.7156724 | 209.8668788 | 193.9588941 | 112.3013582 | 119.6057863 | 158.7535001 | 0.018881 | -0.51865 | 0.01015 | -2.57069 | 0.698026 | 1.723966 |

| | | | | | | | | | | | | | | |
|---|---|---|---|---|---|---|---|---|---|---|---|---|---|---|
| CNAG_015 | - | hypothetic | 4.893580577 | 3.362465944 | 2.255579876 | 12.72866791 | 21.35192673 | 8.900171714 | 0.01893 | 2.170653 | 0.01018 | 2.569641 | 4.502272 | 1.722851 |
| CNAG_077 | - | hypothetic | 585.2800707 | 647.5799154 | 623.9339282 | 727.8318158 | 720.3042582 | 781.4021954 | 0.01895 | 0.249558 | 0.010194 | 2.569193 | 1.188843 | 1.722389 |
| CNAG_060 | - | hypothetic | 61.25296132 | 87.37546549 | 96.0253578 | 134.4315169 | 119.2021203 | 120.8584684 | 0.018962 | 0.597958 | 0.010202 | 2.568899 | 1.513573 | 1.722119 |
| CNAG_126 | - | unspecifie | 182.2450238 | 145.6153415 | 150.4950089 | 219.769677 | 247.069714 | 194.2655666 | 0.018976 | 0.452185 | 0.010214 | 2.56849 | 1.368111 | 1.721803 |
| CNAG_127 | - | unspecifie | 27.30011435 | 42.07363305 | 26.84149484 | 72.54101705 | 47.45510908 | 59.72695783 | 0.018976 | 0.892836 | 0.010213 | 2.568519 | 1.856823 | 1.721803 |
| CNAG_067 | - | kinesin | 2580.665921 | 2678.435335 | 2523.217885 | 2360.445965 | 2234.760829 | 2395.76592 | 0.019117 | -0.1698 | 0.010293 | -2.56583 | 0.888968 | 1.718575 |
| CNAG_045 | - | hypothetic | 144.1173599 | 182.1981984 | 216.4927272 | 239.9294527 | 232.1156787 | 277.2501883 | 0.019145 | 0.450202 | 0.01031 | 2.565243 | 1.366231 | 1.717934 |
| CNAG_056 | - | solute car | 162.735689 | 198.3386533 | 211.2025996 | 306.6133115 | 250.2689519 | 225.3746814 | 0.019228 | 0.437102 | 0.010357 | 2.563664 | 1.353882 | 1.716056 |
| CNAG_127 | - | unspecifie | 179.2262612 | 133.5743204 | 137.5309237 | 101.5654032 | 94.58110958 | 117.7551814 | 0.019848 | -0.54121 | 0.010696 | -2.55249 | 0.687193 | 1.702294 |
| CNAG_024 | - | nuclear pr | 751.9675034 | 770.3146516 | 797.6393832 | 631.7018386 | 665.7910537 | 687.2433418 | 0.019852 | -0.24037 | 0.010701 | -2.55233 | 0.846527 | 1.702191 |
| CNAG_015 | - | biotin-[ace | 1905.702413 | 1988.872585 | 1916.385405 | 1657.552998 | 1889.915981 | 1532.195646 | 0.019872 | -0.20938 | 0.010713 | -2.55191 | 0.864908 | 1.701769 |
| CNAG_128 | - | unspecifie | 48.39083533 | 41.82899211 | 30.916441 | 23.85356725 | 14.27359246 | 20.18522025 | 0.020038 | -1.07703 | 0.01081 | -2.54877 | 0.474003 | 1.698151 |
| CNAG_053 | ZAP104 | specific RI | 91.55208745 | 118.6154898 | 128.0207007 | 145.631846 | 163.2394275 | 179.5540963 | 0.020148 | 0.516649 | 0.010875 | 2.546694 | 1.430629 | 1.695768 |
| CNAG_031 | - | hypothetic | 54.00031565 | 56.14122374 | 44.96240984 | 108.4411261 | 82.13746589 | 67.74676151 | 0.020335 | 0.723867 | 0.010981 | 2.543313 | 1.651603 | 1.691759 |
| CNAG_034 | - | hypothetic | 712.5627178 | 798.409812 | 759.3724496 | 863.2003664 | 936.9111663 | 891.3842364 | 0.020355 | 0.23066 | 0.010997 | 2.542809 | 1.173372 | 1.691329 |
| CNAG_067 | - | hypothetic | 1538.001412 | 1485.836923 | 1425.921389 | 1714.721854 | 1674.287978 | 1726.218751 | 0.020355 | 0.186022 | 0.010994 | 2.542879 | 1.137622 | 1.691329 |
| CNAG_001 | - | hypothetic | 455.5810885 | 449.3710396 | 430.9396189 | 577.0238415 | 620.494684 | 472.996605 | 0.020447 | 0.307655 | 0.011049 | 2.541161 | 1.237694 | 1.689379 |
| CNAG_025 | RIB3 | 3,4-dihydr | 693.457653 | 618.4521264 | 706.9698104 | 563.2769613 | 606.3440139 | 519.7284188 | 0.020644 | -0.27287 | 0.01116 | -2.53764 | 0.827671 | 1.685209 |
| CNAG_062 | - | amyloid b | 1421.126207 | 1621.409612 | 1388.526536 | 1291.797395 | 1323.960007 | 1267.903651 | 0.020655 | -0.2055 | 0.011169 | -2.53738 | 0.867237 | 1.684977 |
| CNAG_019 | - | NAD synth | 8024.581528 | 6938.687161 | 7419.290361 | 6553.800568 | 6834.011477 | 6746.509766 | 0.020783 | -0.16868 | 0.011243 | -2.53505 | 0.889658 | 1.682291 |
| CNAG_034 | - | 1-alkyl-2-a | 233.8469143 | 235.0146145 | 280.4988055 | 360.0896315 | 321.0109458 | 296.3922232 | 0.020856 | 0.369881 | 0.011285 | 2.533741 | 1.292246 | 1.680761 |
| CNAG_066 | - | ribonuclea | 157.7694306 | 136.970428 | 136.6243256 | 200.3609818 | 205.5394735 | 184.2155754 | 0.021041 | 0.441509 | 0.011387 | 2.530578 | 1.358024 | 1.676941 |
| CNAG_067 | - | hypothetic | 313.5033532 | 474.0991198 | 480.9863605 | 521.579302 | 507.2238943 | 635.8013627 | 0.021131 | 0.37879 | 0.011439 | 2.528988 | 1.30025 | 1.675072 |
| CNAG_059 | - | hypothetic | 1276.534208 | 1372.682469 | 1310.72781 | 1435.166219 | 1602.800153 | 1544.150508 | 0.021161 | 0.195873 | 0.01146 | 2.52834 | 1.145417 | 1.674466 |
| CNAG_016 | - | hypothetic | 488.3388901 | 447.2862641 | 569.4823545 | 590.0356759 | 638.4694668 | 633.7717977 | 0.021168 | 0.29145 | 0.011467 | 2.528145 | 1.22387 | 1.674322 |
| CNAG_033 | URK1 | Uridine ki | 1941.745779 | 1817.58161 | 1679.677368 | 1476.117529 | 1683.69482 | 1600.923875 | 0.021187 | -0.20768 | 0.01148 | -2.52775 | 0.865931 | 1.67393 |
| CNAG_035 | RIM20 | pH-respor | 1714.20626 | 1719.554742 | 1790.522931 | 1539.298695 | 1607.770344 | 1509.916977 | 0.021221 | -0.18147 | 0.011503 | -2.52702 | 0.881806 | 1.673225 |
| CNAG_069 | TUB4 | tubulin ga | 1275.120442 | 1339.177662 | 1296.769781 | 1163.430326 | 1015.006053 | 1213.418463 | 0.021221 | -0.22023 | 0.011503 | -2.52702 | 0.858429 | 1.673225 |
| CNAG_000 | - | hypothetic | 22.65404086 | 34.56890143 | 29.9832251 | 48.49314291 | 61.71088647 | 50.90716676 | 0.02126 | 0.889267 | 0.01153 | 2.526222 | 1.852235 | 1.672434 |
| CNAG_004 | - | hypothetic | 22.6103078 | 18.55262687 | 37.34686889 | 48.62844407 | 52.0716188 | 51.93089502 | 0.02126 | 0.955231 | 0.011527 | 2.526292 | 1.93889 | 1.672434 |
| CNAG_008 | - | ligase | 739.1287852 | 777.8373331 | 773.123993 | 609.8710043 | 694.3249043 | 652.8654872 | 0.021276 | -0.24273 | 0.011541 | -2.52588 | 0.845146 | 1.672105 |
| CNAG_051 | GAT5 | hypothetic | 414.6905569 | 443.9190744 | 340.3514478 | 482.6755282 | 504.7484223 | 514.9698997 | 0.021288 | 0.310831 | 0.01155 | 2.525603 | 1.240422 | 1.671864 |
| CNAG_033 | - | N-acetyltr | 649.192925 | 638.8944174 | 711.2269786 | 568.7534898 | 538.0648085 | 582.8982753 | 0.021393 | -0.25856 | 0.011609 | -2.5238 | 0.835924 | 1.669734 |
| CNAG_010 | - | hypothetic | 281.451691 | 333.8781305 | 336.8842657 | 252.7075997 | 224.8634867 | 269.8261 | 0.021512 | -0.36516 | 0.011682 | -2.52161 | 0.776381 | 1.667319 |
| CNAG_024 | - | hypothetic | 1375.731464 | 1363.013563 | 1355.518296 | 1474.067866 | 1617.427164 | 1639.590746 | 0.021512 | 0.193399 | 0.01168 | 2.521664 | 1.143454 | 1.667319 |
| CNAG_050 | PIP201 | hypothetic | 818.7655955 | 703.732049 | 929.9370293 | 932.4322304 | 1065.490521 | 995.7181216 | 0.021512 | 0.272103 | 0.011677 | 2.521746 | 1.207567 | 1.667319 |
| CNAG_122 | - | unspecifie | 28.26888707 | 17.47052207 | 27.82943167 | 39.80797482 | 53.92121007 | 50.78712499 | 0.021566 | 0.962813 | 0.011714 | 2.520639 | 1.949107 | 1.666221 |
| CNAG_077 | CLR6 | transcripti | 1138.812037 | 1604.148721 | 1644.323184 | 1198.323351 | 1255.794018 | 1195.766343 | 0.02158 | -0.27992 | 0.011724 | -2.52033 | 0.823636 | 1.665939 |
| CNAG_031 | - | hypothetic | 750.7387986 | 800.4183665 | 690.0026208 | 644.2597346 | 639.9101564 | 626.2393777 | 0.021593 | -0.24555 | 0.011734 | -2.52005 | 0.843496 | 1.665693 |
| CNAG_029 | - | anaphase- | 831.4093503 | 853.2919294 | 927.7235728 | 774.2870971 | 715.7625932 | 754.989931 | 0.021647 | -0.23434 | 0.011768 | -2.51902 | 0.850075 | 1.66461 |
| CNAG_027 | - | hypothetic | 203.5700539 | 241.428053 | 260.2444523 | 304.5534609 | 295.966359 | 319.5479199 | 0.021647 | 0.368202 | 0.011766 | 2.519077 | 1.290743 | 1.66461 |
| CNAG_069 | AKP1 | alkaline pl | 12.14582574 | 18.45834128 | 19.30813842 | 34.57933205 | 55.22924583 | 23.41368747 | 0.021753 | 1.177204 | 0.011832 | 2.517127 | 2.261381 | 1.662475 |
| CNAG_019 | - | hypothetic | 667.5306205 | 569.9099886 | 522.575351 | 510.4659479 | 512.3085191 | 391.0470815 | 0.021768 | -0.33123 | 0.011842 | -2.51681 | 0.794858 | 1.662186 |
| CNAG_051 | - | hypothetic | 652.598786 | 633.4746665 | 614.2508689 | 529.0165772 | 530.3411487 | 552.9275362 | 0.022127 | -0.25254 | 0.012046 | -2.5108 | 0.839415 | 1.655078 |
| CNAG_051 | - | homoserir | 3071.310935 | 3583.47718 | 3849.787188 | 3163.64804 | 3160.359202 | 2907.644823 | 0.022178 | -0.20128 | 0.012076 | -2.50992 | 0.869781 | 1.654086 |
| CNAG_058 | - | hypothetic | 212.3120033 | 226.1046883 | 194.0684138 | 142.7545249 | 183.7097249 | 147.8486673 | 0.022247 | -0.4363 | 0.012117 | -2.50873 | 0.739025 | 1.652726 |
| CNAG_060 | - | peroxin-14 | 378.269673 | 403.9451608 | 399.8485178 | 320.4274644 | 295.6315909 | 343.0884579 | 0.02233 | -0.31553 | 0.012164 | -2.50734 | 0.803557 | 1.651117 |
| CNAG_059 | CDC420 | cell divisic | 114.2094037 | 125.8713913 | 107.6585175 | 81.40835304 | 70.41985552 | 85.5985682 | 0.022361 | -0.56653 | 0.012184 | -2.50676 | 0.675239 | 1.650502 |
| CNAG_040 | - | hypothetic | 18.79860847 | 9.886967034 | 8.707024487 | 39.18440411 | 26.91987604 | 23.27334934 | 0.022456 | 1.313035 | 0.012239 | 2.50519 | 2.484638 | 1.648666 |
| CNAG_038 | - | hypothetic | 596.9942414 | 682.0550052 | 668.7126181 | 839.3643073 | 752.212621 | 747.0986399 | 0.022478 | 0.250027 | 0.012256 | 2.504688 | 1.18923 | 1.648244 |
| CNAG_059 | - | multidrug | 1863.643094 | 1276.629924 | 978.008261 | 946.8843333 | 808.7421155 | 660.9426941 | 0.022489 | -0.78709 | 0.012265 | -2.50443 | 0.579511 | 1.648025 |
| CNAG_004 | - | dolichyl-p | 431.60759 | 340.4625982 | 347.6450997 | 298.6240383 | 285.3104425 | 305.3926013 | 0.022542 | -0.34829 | 0.012296 | -2.50352 | 0.785514 | 1.647008 |
| CNAG_066 | - | cathepsin | 3380.469399 | 3119.154141 | 3133.226026 | 2711.310075 | 2900.71814 | 3015.216601 | 0.02256 | -0.17469 | 0.012309 | -2.50315 | 0.885959 | 1.646654 |
| CNAG_016 | - | hypothetic | 528.1105968 | 629.2389471 | 581.2800689 | 720.1759652 | 673.7204425 | 703.7427976 | 0.022607 | 0.255711 | 0.01234 | 2.50226 | 1.193924 | 1.64575 |
| CNAG_031 | - | hypothetic | 6.131860124 | 6.594520932 | 5.459558347 | 24.22875496 | 11.14160696 | 20.80821812 | 0.022608 | 1.679619 | 0.012344 | 2.502164 | 3.203433 | 1.645729 |
| CNAG_059 | - | hypothetic | 70.17103947 | 56.18056247 | 44.98874194 | 115.3863047 | 87.28059376 | 77.69583332 | 0.022732 | 0.698964 | 0.012414 | 2.50016 | 1.623339 | 1.643369 |
| CNAG_077 | - | hypothetic | 48.42538883 | 85.13360803 | 67.30591525 | 107.8484903 | 98.47843681 | 110.766694 | 0.022788 | 0.642197 | 0.01245 | 2.499117 | 1.560704 | 1.642284 |
| CNAG_029 | - | CCR4-NOT | 3564.740773 | 2935.990553 | 2738.686392 | 2801.778752 | 2734.245824 | 2410.336382 | 0.023131 | -0.23351 | 0.012643 | -2.49367 | 0.85056 | 1.635808 |
| CNAG_022 | - | hypothetic | 296.3558122 | 328.4035833 | 235.7342921 | 229.3845604 | 227.5767143 | 204.4681016 | 0.023226 | -0.39479 | 0.012704 | -2.49197 | 0.760599 | 1.634024 |
| CNAG_052 | - | chaperone | 1108.157932 | 1189.408898 | 1142.135281 | 1067.003947 | 996.1506796 | 939.3370761 | 0.023227 | -0.21105 | 0.012707 | -2.49188 | 0.863909 | 1.634002 |
| CNAG_128 | - | unspecifie | 7.304403188 | 5.552632384 | 8.632204244 | 15.65606266 | 27.32825886 | 18.72774316 | 0.023328 | 1.567988 | 0.012765 | 2.490254 | 2.964909 | 1.632117 |
| CNAG_004 | - | hypothetic | 435.1631369 | 341.5616138 | 376.4076478 | 318.1367442 | 294.7981809 | 309.8499597 | 0.02341 | -0.33896 | 0.012813 | -2.48893 | 0.790611 | 1.630591 |
| CNAG_052 | - | DNA misn | 196.4233324 | 211.2449044 | 191.0041102 | 284.5384721 | 230.284426 | 276.2401325 | 0.023516 | 0.38707 | 0.01288 | 2.487082 | 1.307735 | 1.628631 |
| CNAG_033 | - | hypothetic | 13.18866827 | 16.2762182 | 11.88403989 | 34.91695869 | 34.0196689 | 24.39025692 | 0.023567 | 1.186949 | 0.01291 | 2.486236 | 2.276707 | 1.627694 |
| CNAG_079 | - | hypothetic | 628.2032103 | 641.0343019 | 718.6717439 | 607.0541199 | 569.200553 | 467.6177732 | 0.023586 | -0.28976 | 0.012924 | -2.48587 | 0.81804 | 1.627344 |
| CNAG_016 | - | hypothetic | 1504.791351 | 1167.955975 | 1183.771117 | 1106.56303 | 1084.111802 | 1081.401282 | 0.023589 | -0.25288 | 0.012928 | -2.48574 | 0.839221 | 1.627288 |
| CNAG_009 | - | protein tra | 372.4154628 | 404.99888 | 379.6054749 | 350.2251794 | 282.6735424 | 298.7853492 | 0.023705 | -0.32735 | 0.012995 | -2.48392 | 0.796999 | 1.625154 |
| CNAG_027 | - | transcripti | 651.656077 | 715.3663067 | 803.9749717 | 636.5266174 | 589.8301559 | 602.9266654 | 0.023706 | -0.26257 | 0.012998 | -2.48383 | 0.833599 | 1.625144 |

| | | | | | | | | | | | | | |
|---|---|---|---|---|---|---|---|---|---|---|---|---|---|
| CNAG_028 | - | GPI mann | 148.7778595 | 200.4693536 | 209.0591953 | 264.2481746 | 219.0761534 | 275.0661165 | 0.023863 | 0.430452 | 0.01309 | 2.48131 | 1.347656 | 1.622267 |
| CNAG_035 | - | hypothetic | 17.56180748 | 5.595359177 | 6.569539926 | 27.82383052 | 29.76455912 | 21.02403829 | 0.023879 | 1.4404 | 0.013101 | 2.481 | 2.713962 | 1.621986 |
| CNAG_021 | - | hypothetic | 1672.301247 | 1646.364898 | 1551.765438 | 1772.964838 | 1931.4119 | 1880.53509 | 0.023948 | 0.182119 | 0.013149 | 2.479699 | 1.134549 | 1.620733 |
| CNAG_002 | - | hypothetic | 122.8917417 | 128.3202007 | 129.128691 | 169.8987669 | 187.4675663 | 167.5362003 | 0.023948 | 0.449362 | 0.013148 | 2.479728 | 1.365437 | 1.620733 |
| CNAG_020 | CAS4 | putative s | 456.6304873 | 529.9958436 | 535.2654589 | 413.0251372 | 424.2577158 | 425.2872464 | 0.023968 | -0.28314 | 0.013165 | -2.47927 | 0.821802 | 1.620364 |
| CNAG_059 | - | serine carl | 54.29319607 | 90.56744036 | 90.70200695 | 113.0918963 | 129.7358332 | 117.5174279 | 0.02402 | 0.603484 | 0.013196 | 2.478424 | 1.519382 | 1.619429 |
| CNAG_032 | - | hypothetic | 2684.646946 | 2466.298561 | 2110.643137 | 3100.306009 | 2854.315811 | 2612.228813 | 0.02412 | 0.223164 | 0.013257 | 2.476781 | 1.167291 | 1.617622 |
| CNAG_023 | - | hypothetic | 245.5796244 | 311.3971287 | 274.1659546 | 391.1883047 | 356.3898457 | 321.9397093 | 0.024273 | 0.349326 | 0.013347 | 2.474361 | 1.273965 | 1.614872 |
| CNAG_045 | - | hypothetic | 567.398074 | 518.2560211 | 622.686994 | 479.2382749 | 495.0322841 | 446.4417619 | 0.024305 | -0.28144 | 0.013368 | -2.47382 | 0.822771 | 1.614312 |
| CNAG_014 | - | myo-inosi | 769.8027612 | 808.1613217 | 909.688327 | 933.9946028 | 1005.019544 | 1014.566715 | 0.024334 | 0.232633 | 0.013387 | 2.473315 | 1.174977 | 1.613793 |
| CNAG_057 | - | multidrug | 234.9289173 | 241.4257869 | 232.5755396 | 308.4849708 | 280.370163 | 321.7548427 | 0.024439 | 0.347644 | 0.013447 | 2.471697 | 1.272481 | 1.611924 |
| CNAG_054 | APN1 | AP endon | 943.4857993 | 980.4053036 | 973.6269557 | 848.9660753 | 878.9013118 | 798.3376423 | 0.024526 | -0.21357 | 0.013505 | -2.47018 | 0.862403 | 1.61037 |
| CNAG_080 | - | hypothetic | 121.8827837 | 172.4040976 | 146.2029609 | 215.270718 | 186.3862675 | 204.1082604 | 0.024717 | 0.448883 | 0.013616 | 2.467242 | 1.364983 | 1.606998 |
| CNAG_014 | - | hypothetic | 1073.176149 | 1016.064232 | 836.2417339 | 1152.795326 | 1236.249739 | 1115.676688 | 0.024768 | 0.244766 | 0.013647 | 2.466427 | 1.1849 | 1.606105 |
| CNAG_026 | - | hypothetic | 285.1740151 | 286.7059547 | 310.3891575 | 430.9979181 | 364.1021504 | 333.064292 | 0.024835 | 0.339935 | 0.013696 | 2.465141 | 1.265699 | 1.604931 |
| CNAG_126 | - | unspecifie | 80.91530924 | 88.46547188 | 92.85962908 | 127.9954639 | 127.8774971 | 124.1979769 | 0.024835 | 0.524842 | 0.013695 | 2.465153 | 1.438776 | 1.604931 |
| CNAG_018 | - | hypothetic | 212.9037172 | 47.58858223 | 59.79592203 | 31.8494589 | 30.52170784 | 36.9970612 | 0.025003 | -1.73452 | 0.013795 | -2.46257 | 0.300508 | 1.602008 |
| CNAG_122 | - | unspecifie | 10.98747064 | 24.81616204 | 14.02673025 | 29.71082086 | 45.45223757 | 33.15602868 | 0.025238 | 1.143679 | 0.013937 | 2.458895 | 2.209437 | 1.597944 |
| CNAG_041 | - | hypothetic | 368.0254054 | 330.918636 | 394.6210121 | 396.0456775 | 505.7533045 | 481.6406487 | 0.025373 | 0.32386 | 0.014014 | 2.456899 | 1.251675 | 1.595627 |
| CNAG_017 | - | FACT com | 6231.238758 | 6458.156295 | 6062.8486 | 6537.58881 | 7292.217955 | 7202.638343 | 0.02538 | 0.150464 | 0.014021 | 2.456727 | 1.109926 | 1.595516 |
| CNAG_051 | CAN2 | Beta-type | 2892.338285 | 2676.322639 | 3329.27726 | 1853.2811 | 2654.986214 | 2128.375119 | 0.025548 | -0.43957 | 0.014117 | -2.45427 | 0.737354 | 1.592638 |
| CNAG_062 | - | hypothetic | 288.4823388 | 439.2220897 | 291.1550448 | 198.5913647 | 312.3124853 | 239.9880717 | 0.025562 | -0.45627 | 0.014128 | -2.454 | 0.72887 | 1.592412 |
| CNAG_122 | - | unspecifie | 14.54003622 | 29.17103082 | 25.7020221 | 35.02185575 | 50.44320946 | 51.82170311 | 0.025568 | 0.98668 | 0.014138 | 2.453749 | 1.98162 | 1.592301 |
| CNAG_029 | - | metal resi | 2955.364955 | 2753.878703 | 2546.710146 | 2545.721537 | 2466.8431 | 2344.775074 | 0.025568 | -0.18185 | 0.014135 | -2.4538 | 0.88157 | 1.592301 |
| CNAG_068 | - | Methylene | 1181.774405 | 1304.694452 | 1283.956256 | 1119.834232 | 1162.702998 | 1017.060597 | 0.025612 | -0.20725 | 0.014165 | -2.45305 | 0.866189 | 1.591557 |
| CNAG_027 | - | kynurenin | 1756.310785 | 1927.456336 | 1871.586755 | 1695.025029 | 1681.927567 | 1589.859414 | 0.025663 | -0.17667 | 0.014196 | -2.45226 | 0.884744 | 1.59069 |
| CNAG_030 | - | RNA exon | 424.0099855 | 366.4696958 | 427.6887454 | 494.3908862 | 500.4127198 | 506.1180345 | 0.025733 | 0.285793 | 0.014238 | 2.451193 | 1.21908 | 1.589503 |
| CNAG_040 | - | hypothetic | 257.0073824 | 214.5620343 | 242.1578441 | 314.7375817 | 308.9594113 | 297.4380915 | 0.025795 | 0.350227 | 0.014279 | 2.450169 | 1.274761 | 1.58846 |
| CNAG_000 | - | protein PI | 3124.725358 | 3038.300929 | 2716.282632 | 2701.347504 | 2707.521354 | 2533.499794 | 0.025795 | -0.17622 | 0.014276 | -2.45024 | 0.885019 | 1.58846 |
| CNAG_004 | - | cytoplasm | 295.3153314 | 279.0405934 | 333.6587009 | 226.1316588 | 244.8339212 | 248.7683075 | 0.025808 | -0.35273 | 0.014289 | -2.44991 | 0.783101 | 1.588248 |
| CNAG_012 | HOC2 | alpha-1,6- | 182.3312712 | 142.4228589 | 191.9979657 | 242.4404442 | 217.4474666 | 237.4072432 | 0.025864 | 0.4184 | 0.014327 | 2.448968 | 1.336444 | 1.587298 |
| CNAG_010 | - | dihydroce | 1438.836137 | 983.6420989 | 854.2750965 | 757.6820132 | 687.1351902 | 750.6614222 | 0.026008 | -0.59459 | 0.014409 | -2.44689 | 0.662232 | 1.584892 |
| CNAG_022 | ZIP3 | solute car | 105.3551585 | 112.1388989 | 98.24518758 | 162.4082313 | 157.2586036 | 129.8823408 | 0.026087 | 0.493057 | 0.014456 | 2.445719 | 1.407424 | 1.583575 |
| CNAG_075 | YAP4 | hypothetic | 1707.070934 | 1557.958785 | 1595.390787 | 1370.329732 | 1476.647871 | 1461.028215 | 0.026399 | -0.18962 | 0.014645 | -2.44104 | 0.876834 | 1.57842 |
| CNAG_127 | - | unspecifie | 2.576458377 | 2.272684363 | 3.287646668 | 7.800854452 | 10.50787049 | 17.32741997 | 0.026399 | 2.265225 | 0.014643 | 2.44108 | 4.807294 | 1.57842 |
| CNAG_018 | - | splicing fa | 1333.589572 | 1372.634272 | 1564.378074 | 1241.981886 | 1278.246405 | 1223.496066 | 0.026452 | -0.20546 | 0.014681 | -2.44015 | 0.867262 | 1.577539 |
| CNAG_059 | - | hypothetic | 838.6780699 | 990.1523136 | 845.7849769 | 1070.354575 | 1085.250385 | 998.0145209 | 0.026497 | 0.223701 | 0.014709 | 2.439458 | 1.167725 | 1.576806 |
| CNAG_079 | - | hypothetic | 5.977472668 | 2.285987722 | 2.24613557 | 11.11468983 | 12.27166766 | 15.26201118 | 0.026617 | 2.060996 | 0.014779 | 2.437742 | 4.172742 | 1.574839 |
| CNAG_023 | - | hypothetic | 680.5943248 | 608.7457908 | 670.7287335 | 582.8576347 | 576.1254341 | 485.3459087 | 0.026827 | -0.26896 | 0.014899 | -2.43482 | 0.829915 | 1.57143 |
| CNAG_009 | - | glucan 1,3 | 2.661889574 | 7.614469834 | 2.268328682 | 11.75816111 | 5.836639332 | 32.39318074 | 0.026828 | 2.11695 | 0.014903 | 2.434724 | 4.33776 | 1.571407 |
| CNAG_016 | - | hypothetic | 837.2153196 | 904.9486633 | 832.8786634 | 770.4104146 | 692.4502226 | 760.5214394 | 0.026961 | -0.22633 | 0.014987 | -2.4327 | 0.854807 | 1.569269 |
| CNAG_044 | - | F-box prot | 253.5830016 | 243.6072891 | 251.7585093 | 284.8367657 | 332.2943251 | 341.7275999 | 0.02705 | 0.342002 | 0.015043 | 2.431344 | 1.267514 | 1.56783 |
| CNAG_052 | - | telomere | 1203.772569 | 1162.496349 | 1081.387516 | 980.4331506 | 999.599711 | 1043.5952 | 0.02745 | -0.20466 | 0.015275 | -2.42579 | 0.867743 | 1.561463 |
| CNAG_033 | - | hypothetic | 138.2185762 | 204.6808254 | 146.2505022 | 227.5968859 | 253.112845 | 194.2871833 | 0.027478 | 0.451985 | 0.015298 | 2.425255 | 1.367921 | 1.561015 |
| CNAG_120 | - | unspecifie | 6.245753928 | 9.838700788 | 11.82438271 | 17.09992742 | 39.76295209 | 16.67609692 | 0.027478 | 1.45354 | 0.015297 | 2.42526 | 2.738792 | 1.561015 |
| CNAG_077 | PIK1 | 1-phospha | 1738.572705 | 1537.500857 | 1604.98748 | 1468.522634 | 1434.340633 | 1421.094051 | 0.027798 | -0.1903 | 0.015486 | -2.42081 | 0.876424 | 1.555981 |
| CNAG_073 | - | hypothetic | 1519.170068 | 1446.99531 | 1623.063749 | 1376.679723 | 1388.662619 | 1294.565211 | 0.027856 | -0.19274 | 0.015521 | -2.41998 | 0.874944 | 1.555088 |
| CNAG_034 | - | hypothetic | 478.7365053 | 497.7235685 | 534.1982845 | 422.4115203 | 411.2872313 | 431.9307577 | 0.02787 | -0.27073 | 0.015533 | -2.41971 | 0.8289 | 1.554859 |
| CNAG_002 | YFH7 | hypothetic | 386.5179675 | 472.817086 | 403.0885671 | 367.257961 | 309.4112334 | 348.6872554 | 0.027895 | -0.31453 | 0.01555 | -2.4193 | 0.804112 | 1.554466 |
| CNAG_075 | - | hypothetic | 297.9091353 | 290.9829794 | 266.7231716 | 383.3043713 | 365.8915128 | 329.7041444 | 0.028077 | 0.321089 | 0.015662 | 2.416697 | 1.249273 | 1.551646 |
| CNAG_019 | - | hypothetic | 331.6333095 | 376.9941793 | 383.8175865 | 283.0053002 | 335.5587963 | 251.1342787 | 0.028148 | -0.34211 | 0.015705 | -2.41571 | 0.788886 | 1.550559 |
| CNAG_130 | - | unspecifie | 205.4871568 | 239.0504212 | 228.135886 | 174.8595524 | 168.0003354 | 177.7381802 | 0.028166 | -0.38736 | 0.015719 | -2.41538 | 0.764525 | 1.55027 |
| CNAG_017 | - | vacuolar p | 14038.69792 | 13084.38384 | 11600.98644 | 14511.32281 | 14441.9152 | 14957.11321 | 0.028167 | 0.16601 | 0.015723 | 2.415294 | 1.121951 | 1.550258 |
| CNAG_074 | - | hypothetic | 250.1277501 | 257.594527 | 264.5419415 | 332.611682 | 318.3945237 | 324.059111 | 0.02817 | 0.322051 | 0.015728 | 2.415173 | 1.250106 | 1.550209 |
| CNAG_051 | - | DNA poly | 1090.671454 | 1196.946943 | 1151.728668 | 1022.54888 | 1056.602187 | 934.9022194 | 0.028197 | -0.20545 | 0.015746 | -2.41475 | 0.867269 | 1.549798 |
| CNAG_009 | - | phosphati | 3650.399991 | 3373.501747 | 3169.587202 | 3661.85367 | 3926.477186 | 4025.333805 | 0.028201 | 0.172694 | 0.015752 | 2.414622 | 1.127161 | 1.549743 |
| CNAG_020 | - | hypothetic | 1127.896319 | 1011.725568 | 1159.153975 | 1019.529345 | 891.7056473 | 942.6214967 | 0.02826 | -0.22485 | 0.015788 | -2.41378 | 0.855682 | 1.548827 |
| CNAG_001 | - | oxidoredu | 332.7946806 | 278.0892937 | 232.6615462 | 386.4045655 | 425.63908 | 304.2637239 | 0.028282 | 0.389009 | 0.015804 | 2.413412 | 1.309494 | 1.548489 |
| CNAG_131 | - | unspecifie | 24.52119213 | 12.0416202 | 6.608260548 | 33.98353291 | 32.16950283 | 34.16279247 | 0.028418 | 1.219927 | 0.015884 | 2.411579 | 2.32935 | 1.546401 |
| CNAG_009 | - | importin s | 1394.046801 | 1338.092597 | 1159.265108 | 1157.992486 | 1201.531742 | 981.5848691 | 0.028476 | -0.23622 | 0.015923 | -2.41068 | 0.848968 | 1.545515 |
| CNAG_014 | - | hypothetic | 384.0781088 | 384.5903613 | 407.2983542 | 301.6473681 | 356.2273456 | 302.1253944 | 0.028636 | -0.30859 | 0.016016 | -2.40856 | 0.80743 | 1.543087 |
| CNAG_038 | - | hypothetic | 1481.637391 | 1286.46125 | 1343.708027 | 1165.658202 | 1230.804451 | 1215.694985 | 0.028687 | -0.20298 | 0.016051 | -2.40775 | 0.868752 | 1.542315 |
| CNAG_069 | - | hexose tra | 220.9995963 | 238.2060374 | 254.9260398 | 288.0724182 | 290.7609397 | 338.3502084 | 0.028895 | 0.348573 | 0.016175 | 2.404945 | 1.2733 | 1.539172 |
| CNAG_025 | - | 2,4-dihydr | 77.35623303 | 53.06058284 | 99.17506955 | 119.4963329 | 119.2889193 | 114.1942467 | 0.029073 | 0.603368 | 0.016278 | 2.402622 | 1.51926 | 1.536507 |
| CNAG_018 | - | large subu | 1596.228553 | 1535.331496 | 1717.966694 | 1192.667306 | 1580.269952 | 1398.886477 | 0.029127 | -0.23316 | 0.016312 | -2.40186 | 0.850772 | 1.5357 |
| CNAG_131 | - | unspecifie | 27.3291765 | 31.42761813 | 45.88662427 | 47.42707618 | 62.38839747 | 75.04421565 | 0.029246 | 0.821928 | 0.016385 | 2.400216 | 1.767767 | 1.533935 |
| CNAG_034 | - | ATP-bindi | 3326.790205 | 3264.573581 | 3039.401024 | 2941.837855 | 3014.62142 | 2743.326284 | 0.029246 | -0.16206 | 0.016382 | -2.40028 | 0.893746 | 1.533935 |
| CNAG_045 | - | D-aminoa | 204.4347504 | 144.5945224 | 196.280973 | 259.6045486 | 244.2783608 | 230.8403566 | 0.02927 | 0.413577 | 0.016402 | 2.399834 | 1.331985 | 1.533577 |

| | | | | | | | | | | | | | | |
|---|---|---|---|---|---|---|---|---|---|---|---|---|---|---|
| CNAG_043 | CLR1 | hypothetic | 296.8144533 | 292.088743 | 310.3953631 | 424.698309 | 357.1587766 | 351.8906913 | 0.029308 | 0.318795 | 0.016427 | 2.399286 | 1.247288 | 1.533021 |
| CNAG_022 | - | cytosolic F | 680.4093805 | 573.1657057 | 543.9012027 | 451.8066078 | 507.0776451 | 530.6988415 | 0.029314 | -0.28722 | 0.016434 | -2.39913 | 0.819482 | 1.532927 |
| CNAG_040 | YRM103 | putative tr | 8.470669049 | 10.87116144 | 4.430718458 | 18.89639314 | 35.38539136 | 12.30991546 | 0.029426 | 1.547613 | 0.016504 | 2.397566 | 2.923331 | 1.531265 |
| CNAG_032 | - | hypothetic | 557.1949289 | 525.9046434 | 610.0282264 | 677.2908475 | 701.4259336 | 653.8375357 | 0.029818 | 0.247782 | 0.016735 | 2.392481 | 1.18738 | 1.525526 |
| CNAG_030 | NPP1 | type I pho | 974.066666 | 956.8462126 | 1041.94865 | 1141.876079 | 1108.425323 | 1208.834478 | 0.029883 | 0.203285 | 0.016786 | 2.391357 | 1.151317 | 1.524574 |
| CNAG_059 | - | hypothetic | 20.3097697 | 26.010952 | 32.062732 | 59.24729354 | 42.36475481 | 45.37977076 | 0.029883 | 0.88981 | 0.016785 | 2.39138 | 1.852932 | 1.524574 |
| CNAG_042 | - | cytoplasm | 950.628343 | 979.4003507 | 789.3072877 | 1043.008275 | 1073.99301 | 1101.165681 | 0.029903 | 0.228309 | 0.016805 | 2.390949 | 1.171461 | 1.52428 |
| CNAG_039 | - | THO comp | 8001.334017 | 7784.424043 | 7737.101817 | 6546.690852 | 7199.734994 | 7604.593822 | 0.029903 | -0.15516 | 0.016803 | -2.39099 | 0.898031 | 1.52428 |
| CNAG_001 | - | hypothetic | 1781.902454 | 1750.801697 | 1795.867317 | 1467.567435 | 1667.309027 | 1615.334788 | 0.030116 | -0.18085 | 0.016928 | -2.38827 | 0.88218 | 1.521203 |
| CNAG_042 | - | cytosolic F | 1909.539393 | 1170.148848 | 1360.744523 | 1001.997325 | 1093.546436 | 996.0377505 | 0.030127 | -0.5403 | 0.016938 | -2.38805 | 0.687627 | 1.521047 |
| CNAG_011 | - | hypothetic | 24.68625696 | 12.08475555 | 19.32503033 | 25.4945797 | 38.2016306 | 55.97802713 | 0.030298 | 1.084224 | 0.017041 | 2.385815 | 2.120235 | 1.51859 |
| CNAG_040 | - | hypothetic | 70.45951436 | 65.93108251 | 100.2488583 | 129.763033 | 133.2278215 | 96.62069654 | 0.0303 | 0.591058 | 0.017047 | 2.385703 | 1.506351 | 1.518551 |
| CNAG_069 | NOP1 | rRNA 2'-C | 920.1416193 | 920.105569 | 1003.45159 | 866.9478602 | 802.907846 | 809.4090679 | 0.030563 | -0.21412 | 0.017201 | -2.38237 | 0.86207 | 1.51481 |
| CNAG_041 | - | hypothetic | 978.4899776 | 916.9059342 | 1087.668744 | 934.7371394 | 779.5420204 | 837.1681305 | 0.030624 | -0.24083 | 0.01724 | -2.38155 | 0.846258 | 1.513933 |
| CNAG_032 | YOX101 | specific tra | 4.845321532 | 2.280375747 | 2.241322703 | 12.01520306 | 15.96267775 | 8.853865803 | 0.030803 | 2.111814 | 0.017348 | 2.379247 | 4.322344 | 1.511408 |
| CNAG_130 | - | unspecifie | 40.96067866 | 71.83233744 | 58.50319539 | 43.55336622 | 20.24281468 | 29.10711555 | 0.030913 | -0.88988 | 0.017414 | -2.37785 | 0.539658 | 1.509857 |
| CNAG_008 | - | hypothetic | 19.94309347 | 13.08224745 | 5.535694118 | 28.35987316 | 42.12962756 | 21.1236979 | 0.031166 | 1.281405 | 0.017564 | 2.37468 | 2.430756 | 1.506313 |
| CNAG_073 | - | hypothetic | 440.7225827 | 675.5098629 | 583.407895 | 712.3790375 | 722.1355763 | 681.5712698 | 0.031239 | 0.303194 | 0.017613 | 2.373659 | 1.233873 | 1.5053 |
| CNAG_124 | - | unspecifie | 0.198987573 | 5.435196907 | 1.170133913 | 12.12879082 | 6.8834034 | 15.18531168 | 0.031247 | 2.451908 | 0.017621 | 2.373491 | 5.471394 | 1.505197 |
| CNAG_041 | - | hypothetic | 157.4780695 | 193.6737281 | 136.5266979 | 125.8693023 | 108.4103198 | 120.0405281 | 0.031349 | -0.47526 | 0.017682 | -2.3722 | 0.719338 | 1.503779 |
| CNAG_126 | - | unspecifie | 7.394959593 | 9.84457548 | 13.93097053 | 22.05045571 | 19.01702917 | 32.83239516 | 0.031507 | 1.31743 | 0.017787 | 2.370024 | 2.492217 | 1.501592 |
| CNAG_013 | PPP1 | vacuolar p | 1477.197889 | 1387.775374 | 1159.368613 | 1587.112361 | 1575.135705 | 1556.380569 | 0.031507 | 0.214607 | 0.017787 | 2.370029 | 1.160388 | 1.501592 |
| CNAG_042 | - | hypothetic | 1773.798988 | 1831.60189 | 1911.025389 | 1610.101242 | 1585.256105 | 1738.52334 | 0.031962 | -0.17596 | 0.018056 | -2.36448 | 0.885178 | 1.495364 |
| CNAG_009 | - | transcripti | 2125.093541 | 1911.354985 | 1923.868775 | 1595.16315 | 1873.473055 | 1796.292675 | 0.032044 | -0.19469 | 0.018106 | -2.36345 | 0.873759 | 1.494252 |
| CNAG_078 | - | hypothetic | 1002.265786 | 1153.997963 | 1294.660437 | 1439.362556 | 1401.808131 | 1250.056645 | 0.032316 | 0.230887 | 0.018271 | 2.360072 | 1.173556 | 1.490578 |
| CNAG_031 | - | hypothetic | 364.6026955 | 474.0095949 | 301.9931598 | 542.9951289 | 479.7096252 | 445.1590469 | 0.032346 | 0.350406 | 0.018292 | 2.359652 | 1.274919 | 1.490181 |
| CNAG_050 | - | hypothetic | 1.377962516 | 1.178823669 | 2.209939726 | 11.45702372 | 11.56202078 | 4.494800317 | 0.032489 | 2.683898 | 0.018381 | 2.357858 | 6.425897 | 1.488269 |
| CNAG_045 | - | hypothetic | 13.30098709 | 15.27453724 | 25.63612931 | 50.12674683 | 34.67983988 | 28.86114925 | 0.032522 | 1.06398 | 0.018404 | 2.357394 | 2.090691 | 1.48782 |
| CNAG_024 | - | Phosphoa | 1352.415058 | 1368.404205 | 1435.473134 | 1539.521249 | 1562.158074 | 1647.367207 | 0.032541 | 0.177273 | 0.018418 | 2.357105 | 1.130745 | 1.487575 |
| CNAG_014 | - | hypothetic | 77.45668143 | 113.1044013 | 67.38394449 | 138.2933974 | 104.3218164 | 145.1049678 | 0.032585 | 0.577673 | 0.018447 | 2.356514 | 1.49244 | 1.486978 |
| CNAG_121 | - | unspecifie | 6.202541717 | 7.691640161 | 11.79739254 | 23.09258808 | 16.4247459 | 25.2122627 | 0.032607 | 1.400298 | 0.018464 | 2.356186 | 2.639561 | 1.486688 |
| CNAG_067 | - | hypothetic | 0.189051731 | 3.318870426 | 2.223598338 | 18.27695047 | 7.820479356 | 5.597064712 | 0.032637 | 2.573828 | 0.018485 | 2.355761 | 5.953873 | 1.486285 |
| CNAG_129 | - | unspecifie | 6.223612087 | 10.88372281 | 10.76044749 | 26.35726108 | 17.28411064 | 24.17724976 | 0.03269 | 1.352045 | 0.01852 | 2.355047 | 2.552737 | 1.485586 |
| CNAG_003 | - | NADH deh | 490.4712812 | 487.0117548 | 642.8712615 | 412.925885 | 480.4070443 | 431.9854272 | 0.032772 | -0.30708 | 0.018573 | -2.35399 | 0.808277 | 1.484495 |
| CNAG_074 | - | F-type H - | 11411.23791 | 11092.17235 | 12100.99015 | 10449.18228 | 10619.78383 | 10808.56372 | 0.032895 | -0.13389 | 0.018647 | -2.35252 | 0.911368 | 1.482877 |
| CNAG_004 | - | solute car | 865.0061147 | 707.9009825 | 786.996081 | 659.7712227 | 655.4030507 | 698.3447715 | 0.032959 | -0.24493 | 0.018687 | -2.35172 | 0.843858 | 1.482029 |
| CNAG_122 | - | unspecifie | 22.26591784 | 15.23068119 | 6.60787514 | 33.16383549 | 28.60152816 | 37.40153398 | 0.032977 | 1.172823 | 0.018702 | 2.351426 | 2.254525 | 1.481783 |
| CNAG_028 | FZC51 | hypothetic | 488.9665157 | 434.1599247 | 410.5827206 | 367.1587831 | 388.084302 | 346.5279269 | 0.03308 | -0.29051 | 0.018765 | -2.35016 | 0.817616 | 1.480435 |
| CNAG_005 | - | large subu | 137.0720854 | 104.4220404 | 92.77613638 | 98.19355637 | 45.20210596 | 69.0444086 | 0.033114 | -0.667 | 0.018795 | -2.34957 | 0.629813 | 1.479989 |
| CNAG_034 | - | hypothetic | 383.1498776 | 380.3971592 | 349.8957924 | 453.9378397 | 431.3278034 | 478.318992 | 0.03327 | 0.277529 | 0.018896 | 2.347581 | 1.212117 | 1.477951 |
| CNAG_016 | - | ER to Golg | 451.8372951 | 443.87786 | 500.0597454 | 388.944833 | 362.0726786 | 410.8062962 | 0.03327 | -0.28038 | 0.018892 | -2.34765 | 0.823373 | 1.477951 |
| CNAG_001 | CAS32 | Capsule st | 336.7436549 | 443.9419253 | 444.7441118 | 429.4373746 | 615.4393746 | 519.4394931 | 0.033331 | 0.339219 | 0.018935 | 2.34681 | 1.265071 | 1.477146 |
| CNAG_063 | - | vacuolar r | 2658.921273 | 2730.168835 | 2671.442475 | 2313.631066 | 2409.905231 | 2576.685891 | 0.033338 | -0.15815 | 0.018943 | -2.34665 | 0.896174 | 1.477058 |
| CNAG_057 | - | hypothetic | 728.7339761 | 873.6752946 | 811.5215361 | 722.1596971 | 695.1188667 | 659.5689211 | 0.033649 | -0.23099 | 0.019124 | -2.34311 | 0.852048 | 1.473028 |
| CNAG_013 | - | hypothetic | 26.07632339 | 34.56328341 | 27.86585571 | 56.62112688 | 56.42110018 | 43.26601578 | 0.033794 | 0.818658 | 0.019223 | 2.341184 | 1.763764 | 1.471154 |
| CNAG_125 | - | unspecifie | 7.603673798 | 31.26083671 | 22.5122251 | 41.67629159 | 37.22571772 | 49.56866654 | 0.034372 | 1.070914 | 0.019568 | 2.334532 | 2.100764 | 1.463798 |
| CNAG_067 | FZC49 | hypothetic | 412.5352093 | 498.8791579 | 479.9510562 | 519.904612 | 596.2350198 | 574.9448138 | 0.034476 | 0.267449 | 0.019631 | 2.33332 | 1.203677 | 1.462484 |
| CNAG_065 | - | hypothetic | 4.848070919 | 0.105956863 | 4.34624572 | 8.594706317 | 15.95699482 | 13.11821322 | 0.034723 | 2.150019 | 0.019781 | 2.33048 | 4.438336 | 1.459378 |
| CNAG_124 | - | unspecifie | 148.0820327 | 151.7945772 | 141.7791986 | 114.192426 | 111.1091908 | 99.01578176 | 0.034723 | -0.4567 | 0.019777 | -2.33056 | 0.728649 | 1.459378 |
| CNAG_022 | - | autophagy | 195.3127492 | 225.2375429 | 211.2383542 | 238.1084896 | 282.2645499 | 297.2914627 | 0.0349 | 0.357978 | 0.019903 | 2.328174 | 1.281628 | 1.45717 |
| CNAG_033 | - | hypothetic | 1839.068843 | 1688.340851 | 1805.463155 | 1531.462767 | 1673.348853 | 1563.192381 | 0.034902 | -0.17735 | 0.019908 | -2.32807 | 0.884328 | 1.457144 |
| CNAG_050 | - | hypothetic | 11.17306238 | 42.04537762 | 34.22520091 | 54.87327344 | 74.02354335 | 43.31110988 | 0.035676 | 0.962538 | 0.02038 | 2.319278 | 1.948735 | 1.447625 |
| CNAG_031 | - | hypothetic | 49.24421842 | 44.31133706 | 39.61078599 | 67.97810269 | 79.74273552 | 67.59592099 | 0.035747 | 0.674649 | 0.020425 | 2.318451 | 1.596208 | 1.446763 |
| CNAG_066 | - | CCR4-NOT | 1955.81974 | 1821.921007 | 1855.601458 | 1483.840994 | 1833.844608 | 1653.108253 | 0.036326 | -0.1962 | 0.020773 | -2.31208 | 0.872849 | 1.439786 |
| CNAG_011 | - | co-chaper | 780.043427 | 841.4065253 | 833.9144928 | 724.4529377 | 691.6345264 | 727.2188101 | 0.036445 | -0.21108 | 0.020846 | -2.31076 | 0.863889 | 1.438365 |
| CNAG_063 | - | hypothetic | 1973.495923 | 1712.137891 | 1775.708269 | 2109.32807 | 2054.681706 | 2079.268589 | 0.036581 | 0.177316 | 0.020928 | 2.309274 | 1.130779 | 1.436744 |
| CNAG_125 | - | unspecifie | 20.43573278 | 44.22283665 | 36.3718367 | 63.56130365 | 80.93430356 | 40.09699094 | 0.036917 | 0.858191 | 0.021134 | 2.305579 | 1.812764 | 1.432772 |
| CNAG_042 | - | hypothetic | 2954.150826 | 2377.94839 | 2779.120626 | 2237.309582 | 2363.311902 | 2535.619374 | 0.037156 | -0.201 | 0.021285 | -2.3029 | 0.869945 | 1.429969 |
| CNAG_014 | - | hypothetic | 670.8598319 | 317.8821023 | 306.1275541 | 237.57291 | 247.2266557 | 221.2542697 | 0.03718 | -0.8938 | 0.021303 | -2.30257 | 0.538194 | 1.42969 |
| CNAG_125 | - | unspecifie | 22.53922565 | 49.35188418 | 67.81673437 | 16.44188517 | 32.71576559 | 20.2519768 | 0.037434 | -1.03544 | 0.021453 | -2.29992 | 0.487867 | 1.426734 |
| CNAG_027 | - | hypothetic | 81.96156021 | 95.90402578 | 53.54805846 | 102.0379275 | 136.7707441 | 109.7915234 | 0.037508 | 0.585124 | 0.021505 | 2.299007 | 1.500168 | 1.425875 |
| CNAG_016 | - | anaphase- | 865.3954222 | 803.8562056 | 848.9444592 | 927.7491591 | 984.2837662 | 1021.221816 | 0.037508 | 0.205229 | 0.021504 | 2.299014 | 1.152869 | 1.425875 |
| CNAG_001 | - | mitotic sp | 206.4187062 | 191.7173738 | 195.0927496 | 139.6970537 | 179.4410765 | 131.2370827 | 0.037541 | -0.4148 | 0.021528 | -2.29859 | 0.750122 | 1.42549 |
| CNAG_078 | - | hypothetic | 0.082979192 | 0.040461953 | 0.034966173 | 0.612840207 | 6.297215407 | 7.501616998 | 0.037608 | 4.542679 | 0.021576 | 2.297757 | 23.3068 | 1.424719 |
| CNAG_123 | - | unspecifie | 51.55619084 | 46.45652052 | 38.55844885 | 63.92742126 | 70.92243029 | 85.11971205 | 0.037722 | 0.676185 | 0.021645 | 2.296533 | 1.597909 | 1.423408 |
| CNAG_120 | - | unspecifie | 21.47534628 | 30.28526571 | 31.02038098 | 56.72017298 | 56.48080505 | 37.78174679 | 0.037772 | 0.846859 | 0.021679 | 2.295943 | 1.798581 | 1.422825 |
| CNAG_077 | - | siderophо | 186.8267822 | 217.515331 | 216.3857203 | 163.9925169 | 166.3493189 | 150.0543541 | 0.038017 | -0.38477 | 0.021829 | -2.29334 | 0.765902 | 1.420025 |
| CNAG_050 | - | GTP-bindi | 5202.2853 | 5108.098343 | 5333.295753 | 4960.003751 | 4773.337702 | 4678.299126 | 0.038071 | -0.13373 | 0.021869 | -2.29263 | 0.911471 | 1.419404 |

| | | | | | | | | | | | | | | |
|---|---|---|---|---|---|---|---|---|---|---|---|---|---|---|
| CNAG_12( | - | unspecifie | 80.63739407 | 105.4276219 | 112.8548251 | 71.32556495 | 63.56319899 | 68.988641 | 0.03809 | -0.5705 | 0.021885 | -2.29236 | 0.673385 | 1.419187 |
| CNAG_00 | - | hypothetic | 420.5061976 | 425.6308877 | 362.7173427 | 445.9256004 | 551.4734191 | 489.4952341 | 0.038185 | 0.283794 | 0.021944 | 2.291334 | 1.217392 | 1.418104 |
| CNAG_04 | - | glyceralde | 12.11594857 | 20.56175373 | 17.18359502 | 32.21837517 | 44.59820769 | 26.63392927 | 0.038242 | 1.043264 | 0.021986 | 2.29061 | 2.060886 | 1.41746 |
| CNAG_12( | - | unspecifie | 4.48440354 | 3.196023636 | 4.200288783 | 0.647333892 | 0 | 0.046501109 | 0.038276 | -4.33369 | 0.02201 | -2.29019 | 0.049594 | 1.417073 |
| CNAG_00 | - | cellulase | 4952.412068 | 4461.713282 | 4195.518723 | 5167.200877 | 5376.485353 | 4885.817531 | 0.038328 | 0.165347 | 0.022045 | 2.289592 | 1.121435 | 1.41648 |
| CNAG_02 | - | aldose rec | 3175.098413 | 3495.101109 | 3123.645032 | 2830.427718 | 3135.408131 | 2913.140649 | 0.039058 | -0.15638 | 0.022489 | -2.28201 | 0.897275 | 1.408288 |
| CNAG_12 | - | unspecifie | 8.637602209 | 17.3383148 | 16.09892104 | 24.18124092 | 44.76821201 | 24.39291988 | 0.039228 | 1.15168 | 0.022596 | 2.280195 | 2.221724 | 1.406401 |
| CNAG_02( | - | ARP2/3 cc | 1907.894001 | 1671.108799 | 1751.09469 | 1643.681755 | 1555.94904 | 1558.75332 | 0.039263 | -0.17988 | 0.022621 | -2.27978 | 0.882774 | 1.406021 |
| CNAG_06( | - | pre-rRNA- | 43.64255453 | 42.23227295 | 70.37303607 | 75.67641266 | 93.56859515 | 80.88568554 | 0.039308 | 0.662002 | 0.022657 | 2.279176 | 1.582277 | 1.405516 |
| CNAG_01( | FRK102 | serine/thre | 200.906585 | 170.3589146 | 164.3569822 | 214.8747636 | 251.27595 | 235.2201995 | 0.039308 | 0.374676 | 0.022652 | 2.279248 | 1.296548 | 1.405516 |
| CNAG_02 | - | hypothetic | 2.557183855 | 4.378688818 | 0.081276517 | 7.004355716 | 17.90689236 | 7.742396301 | 0.039655 | 2.35598 | 0.022871 | 2.27558 | 5.11942 | 1.401699 |
| CNAG_00 | - | phosphop | 216.8263833 | 187.4224077 | 185.525886 | 142.0626643 | 138.6242797 | 169.8826612 | 0.039742 | -0.40459 | 0.022926 | -2.27467 | 0.755451 | 1.40075 |
| CNAG_02 | - | alfa-L-rhar | 734.6837782 | 937.2546132 | 966.0869005 | 694.6897212 | 743.4038989 | 800.4513857 | 0.03981 | -0.2517 | 0.02297 | -2.27393 | 0.839905 | 1.400011 |
| CNAG_02 | CHI21 | Chitinase | 114.7666965 | 115.4338323 | 145.0539954 | 160.4795468 | 176.1902544 | 174.1282881 | 0.039847 | 0.431971 | 0.023001 | 2.273411 | 1.349075 | 1.3996 |
| CNAG_01 | - | vacuolar-s | 1116.268684 | 1119.402678 | 1103.746614 | 1082.666276 | 960.7771002 | 874.984224 | 0.039997 | -0.21027 | 0.023093 | -2.2719 | 0.864378 | 1.397971 |
| CNAG_00 | - | hypothetic | 1051.151908 | 1073.189649 | 1136.861973 | 1212.641759 | 1233.527354 | 1298.774498 | 0.040042 | 0.184142 | 0.023123 | 2.271389 | 1.136141 | 1.397486 |
| CNAG_07 | - | mitochonc | 735.6841253 | 776.7861246 | 827.4850724 | 690.210117 | 670.9555226 | 676.1742752 | 0.040204 | -0.21465 | 0.023227 | -2.26968 | 0.861754 | 1.395728 |
| CNAG_07 | - | amidohyd | 43.54147764 | 43.25508243 | 50.20685888 | 83.96670381 | 80.57935368 | 56.67205891 | 0.040427 | 0.671614 | 0.023371 | 2.267319 | 1.592854 | 1.393325 |
| CNAG_00( | - | hypothetic | 2265.296031 | 2136.530815 | 2145.686087 | 1810.793929 | 2001.084403 | 2069.350439 | 0.04047 | -0.17027 | 0.023401 | -2.26683 | 0.888678 | 1.392862 |
| CNAG_12( | - | unspecifie | 8.716051966 | 25.89942086 | 20.36775763 | 42.74236306 | 32.90778225 | 36.44871344 | 0.04068 | 1.041395 | 0.023527 | 2.264772 | 2.058217 | 1.390621 |
| CNAG_03 | OPS1 | opsin 1 | 800.6823469 | 681.9761243 | 614.3115995 | 641.2851641 | 564.8070436 | 558.5572105 | 0.040865 | -0.26462 | 0.023644 | -2.26287 | 0.832415 | 1.388652 |
| CNAG_05 | - | hypothetic | 935.2088975 | 856.5559181 | 913.9036326 | 792.9250539 | 762.3624091 | 816.0154085 | 0.040994 | -0.20573 | 0.023739 | -2.26133 | 0.867099 | 1.387277 |
| CNAG_01( | - | hypothetic | 266.5334232 | 274.8600233 | 321.0151664 | 330.0104938 | 405.7263419 | 349.6299314 | 0.04107 | 0.316713 | 0.023792 | 2.260466 | 1.245489 | 1.386477 |
| CNAG_07 | - | hypothetic | 139.4431625 | 196.1390341 | 199.4571096 | 251.0218335 | 251.2619295 | 209.8212582 | 0.041172 | 0.398586 | 0.023856 | 2.259436 | 1.318215 | 1.385403 |
| CNAG_00 | - | peptidase | 876.9104689 | 884.5320277 | 907.4992731 | 725.8538315 | 770.1551272 | 837.0661546 | 0.041564 | -0.20881 | 0.024099 | -2.25555 | 0.865249 | 1.381282 |
| CNAG_00 | - | phytanoyl | 1.394088896 | 0.087144259 | 4.313716034 | 8.787268148 | 13.35940021 | 7.714490684 | 0.041649 | 2.466553 | 0.024164 | 2.254517 | 5.527215 | 1.380391 |
| CNAG_00 | - | hypothetic | 663.2265495 | 694.8797066 | 689.9453433 | 566.3476669 | 560.4995499 | 639.4555748 | 0.041959 | -0.22807 | 0.024349 | -2.25158 | 0.853779 | 1.37717 |
| CNAG_00 | - | YjeF famil | 726.2141926 | 725.0543844 | 693.1764434 | 606.8530208 | 613.1608526 | 647.2667803 | 0.042323 | -0.21549 | 0.02457 | -2.2481 | 0.861251 | 1.373423 |
| CNAG_01 | - | alternative | 204.6198238 | 210.2057953 | 229.3331555 | 315.7989685 | 267.4968692 | 245.3560475 | 0.042426 | 0.347806 | 0.024635 | 2.247081 | 1.272624 | 1.37237 |
| CNAG_05 | - | glucan 1,3 | 3631.510428 | 3472.511885 | 3100.210974 | 3066.337373 | 3257.032128 | 2828.830749 | 0.042436 | -0.17249 | 0.024646 | -2.2469 | 0.887309 | 1.372261 |
| CNAG_04 | - | hypothetic | 1114.097015 | 1268.063444 | 1265.817514 | 1098.836007 | 1087.608459 | 1050.311401 | 0.04245 | -0.18777 | 0.024659 | -2.2467 | 0.877961 | 1.372123 |
| CNAG_04 | FZC30 | hypothetic | 17.90996702 | 22.74903608 | 22.50394035 | 42.54824333 | 51.48389971 | 26.74053834 | 0.042499 | 0.948765 | 0.024696 | 2.246125 | 1.93022 | 1.371619 |
| CNAG_03( | - | hypothetic | 204.416682 | 257.3525058 | 247.3155149 | 172.4052431 | 201.7743946 | 182.2079178 | 0.042675 | -0.36587 | 0.024811 | -2.24433 | 0.775999 | 1.369822 |
| CNAG_13( | - | unspecifie | 0.077058031 | 0.037591038 | 0.032487053 | 6.741802662 | 2.596374344 | 3.290062118 | 0.043022 | 4.302676 | 0.025018 | 2.241126 | 19.73488 | 1.366308 |
| CNAG_00 | - | lipid parti | 162.3586414 | 148.7598694 | 107.9072703 | 189.4003164 | 171.6851974 | 211.7649791 | 0.043064 | 0.438472 | 0.025047 | 2.24067 | 1.355169 | 1.365887 |
| CNAG_03( | - | long-chair | 1627.729546 | 1601.024332 | 1551.676774 | 1317.356005 | 1455.080795 | 1503.171616 | 0.043132 | -0.17605 | 0.025092 | -2.23998 | 0.885125 | 1.365197 |
| CNAG_01 | - | DNA helic | 899.2816827 | 874.9457621 | 891.6046857 | 997.0611393 | 1093.047354 | 989.1271423 | 0.043166 | 0.193365 | 0.025118 | 2.23959 | 1.143427 | 1.364854 |
| CNAG_04( | - | HAT1-inte | 792.7752466 | 733.7337357 | 813.6340031 | 692.551416 | 696.8784591 | 648.4595774 | 0.043402 | -0.21523 | 0.02526 | -2.23741 | 0.861409 | 1.362493 |
| CNAG_02 | RUB1 | ubiquitin-l | 1049.809269 | 1110.788011 | 1177.28331 | 885.360542 | 1091.193606 | 931.5452542 | 0.043575 | -0.21398 | 0.025371 | -2.2357 | 0.862155 | 1.360761 |
| CNAG_01 | - | DNA-direc | 11649.4145 | 11287.17025 | 11325.72237 | 10344.88071 | 10769.89403 | 10788.57773 | 0.043619 | -0.11837 | 0.025402 | -2.23523 | 0.92123 | 1.360326 |
| CNAG_06 | - | hypothetic | 2313.132779 | 2032.057822 | 2194.72658 | 1876.238271 | 1976.918185 | 2024.966276 | 0.043908 | -0.17005 | 0.025576 | -2.23259 | 0.888811 | 1.357451 |
| CNAG_01 | - | hypothetic | 892.4071761 | 1110.789803 | 857.545762 | 1233.979252 | 1133.511648 | 1030.252006 | 0.043932 | 0.23417 | 0.025595 | 2.232299 | 1.17623 | 1.35722 |
| CNAG_00( | PKP1 | pyruvate c | 529.9623231 | 479.4477768 | 523.5593965 | 481.8002217 | 424.2446694 | 376.5536333 | 0.043943 | -0.27296 | 0.025607 | -2.23212 | 0.827622 | 1.357109 |
| CNAG_06 | - | hypothetic | 55.01605285 | 46.46632096 | 39.62462959 | 71.89754655 | 81.43684131 | 69.82618468 | 0.044088 | 0.642776 | 0.025697 | 2.230761 | 1.56133 | 1.355679 |
| CNAG_01 | - | hypothetic | 1283.296291 | 1263.811189 | 1338.344004 | 1098.705764 | 1192.010713 | 1180.150475 | 0.044591 | -0.17801 | 0.026023 | -2.22587 | 0.88392 | 1.350751 |
| CNAG_06 | BLP3 | pr4/barwir | 191.2584941 | 192.7368211 | 175.9219615 | 178.4038376 | 125.6560962 | 112.4174691 | 0.044704 | -0.44224 | 0.026094 | -2.22481 | 0.735991 | 1.349657 |
| CNAG_00( | - | hypothetic | 1551.93233 | 1550.421255 | 1736.101506 | 1428.780534 | 1450.747294 | 1458.810386 | 0.044757 | -0.17313 | 0.026136 | -2.22419 | 0.886915 | 1.349144 |
| CNAG_12 | - | unspecifie | 27.18831923 | 22.82587985 | 35.24352033 | 47.78512232 | 55.59112477 | 46.50246544 | 0.044785 | 0.800751 | 0.026158 | 2.223859 | 1.742007 | 1.348866 |
| CNAG_05 | - | peptide al | 609.4867258 | 547.3790934 | 721.8268471 | 502.4898062 | 518.2569004 | 558.452703 | 0.044802 | -0.26618 | 0.026173 | -2.22363 | 0.831518 | 1.348703 |
| CNAG_02 | - | hypothetic | 586.1356353 | 534.4408012 | 686.6452461 | 507.2445686 | 533.8629381 | 484.171874 | 0.044824 | -0.2609 | 0.026191 | -2.22336 | 0.834566 | 1.348493 |
| CNAG_07 | - | hypothetic | 643.1290497 | 512.9011918 | 604.6067948 | 533.8646534 | 496.7194126 | 448.6872543 | 0.045051 | -0.26838 | 0.026335 | -2.22123 | 0.830249 | 1.346297 |
| CNAG_12 | - | unspecifie | 13.33607105 | 24.8693112 | 24.60748135 | 44.21630056 | 39.93738135 | 34.33989597 | 0.04513 | 0.925781 | 0.026387 | 2.220474 | 1.899712 | 1.345539 |
| CNAG_05 | - | replicatior | 742.4825 | 720.7320194 | 655.8731485 | 562.3907918 | 566.5165965 | 680.4956133 | 0.04538 | -0.24266 | 0.026552 | -2.21805 | 0.845186 | 1.343132 |
| CNAG_05 | - | hypothetic | 1059.389766 | 1243.35664 | 1135.828631 | 1251.50628 | 1374.206346 | 1328.78637 | 0.04538 | 0.187156 | 0.02655 | 2.218064 | 1.138517 | 1.343132 |
| CNAG_03 | - | solute car | 3839.270785 | 3650.250034 | 2869.906086 | 3109.938355 | 3047.334531 | 2956.463575 | 0.04538 | -0.20011 | 0.026549 | -2.21809 | 0.870485 | 1.343132 |
| CNAG_07( | - | hypothetic | 11.98382126 | 13.05558404 | 12.90809505 | 25.20496254 | 32.37601077 | 22.14856194 | 0.045435 | 1.118633 | 0.026593 | 2.217438 | 2.171412 | 1.342607 |
| CNAG_12( | - | unspecifie | 17.83230913 | 22.70367994 | 15.0874636 | 37.87412004 | 35.6095071 | 33.17407401 | 0.045544 | 0.945161 | 0.026669 | 2.216336 | 1.925403 | 1.341567 |
| CNAG_00( | - | hypothetic | 76.13048901 | 53.03843743 | 89.59472817 | 102.1712024 | 108.9031589 | 114.1172121 | 0.045624 | 0.555031 | 0.026726 | 2.215498 | 1.4692 | 1.340807 |
| CNAG_02 | - | hypothetic | 928.1564544 | 839.3080654 | 852.0759584 | 775.0426443 | 790.9106503 | 731.7177487 | 0.045656 | -0.20559 | 0.02675 | -2.21514 | 0.867182 | 1.340501 |
| CNAG_06 | SNF102 | CAMK/CA | 1253.981169 | 1180.83144 | 1129.374658 | 1022.46747 | 1028.051651 | 1110.180374 | 0.045843 | -0.18809 | 0.026871 | -2.21339 | 0.87777 | 1.338729 |
| CNAG_01 | - | hypothetic | 1111.736069 | 1243.283955 | 1228.500369 | 1001.414096 | 1064.314871 | 1112.404376 | 0.045843 | -0.18791 | 0.02687 | -2.2134 | 0.877879 | 1.338729 |
| CNAG_03 | - | DIL and A | 1943.06909 | 1944.732269 | 1931.32334 | 1680.892697 | 1751.779279 | 1843.989764 | 0.045861 | -0.15681 | 0.026893 | -2.21307 | 0.897004 | 1.338556 |
| CNAG_06 | - | hypothetic | 126.21526 | 120.7442851 | 90.82714197 | 167.8188368 | 153.7033735 | 144.2571687 | 0.045861 | 0.450511 | 0.02689 | 2.213105 | 1.366524 | 1.338556 |
| CNAG_12 | - | unspecifie | 1.470319213 | 8.636973037 | 2.254525874 | 13.5917471 | 14.04548616 | 13.1705406 | 0.045878 | 1.852873 | 0.026914 | 2.212768 | 3.612187 | 1.3384 |
| CNAG_05( | CAS91 | maltose C | 935.5806445 | 1026.827783 | 1015.30029 | 1184.783242 | 1156.797732 | 1081.291484 | 0.045878 | 0.186064 | 0.026913 | 2.212775 | 1.137655 | 1.3384 |
| CNAG_02( | - | hypothetic | 956.2994546 | 977.1719322 | 928.8654173 | 823.2313762 | 840.8988628 | 872.6279862 | 0.045968 | -0.18961 | 0.026972 | -2.21192 | 0.876845 | 1.337548 |
| CNAG_03 | CAP4 | beta-1,2-x | 809.0286154 | 785.3709728 | 685.754884 | 705.1046875 | 622.6009274 | 638.4542432 | 0.046083 | -0.229 | 0.027051 | -2.21078 | 0.853228 | 1.336461 |
| CNAG_12 | - | unspecifie | 34.06892457 | 19.31227355 | 12.84797681 | 8.944416922 | 9.256759293 | 9.064112871 | 0.046093 | -1.33918 | 0.027062 | -2.21062 | 0.395246 | 1.336366 |

| | | | | | | | | | | | | |
|---|---|---|---|---|---|---|---|---|---|---|---|---|
| CNAG_078 | - | hypothetic | 33565.93088 | 31436.32119 | 33858.84746 | 36593.76206 | 35455.5949 | 36648.57927 | 0.046169 | 0.121287 | 0.027113 | 2.209892 | 1.087705 | 1.335652 |
| CNAG_131 | - | unspecifie | 58.61627271 | 59.35896447 | 49.20929006 | 80.38645881 | 104.9100106 | 71.05644036 | 0.046195 | 0.603801 | 0.027134 | 2.20959 | 1.519715 | 1.335407 |
| CNAG_019 | - | hypothetic | 4.958400591 | 2.319409347 | 8.587837969 | 21.01455635 | 22.09788893 | 5.723839541 | 0.046213 | 1.758956 | 0.02715 | 2.209354 | 3.384531 | 1.335235 |
| CNAG_063 | - | phosphog | 7912.707203 | 7550.689761 | 7132.511214 | 8537.995153 | 8221.10745 | 8254.012163 | 0.046227 | 0.131331 | 0.027164 | 2.209152 | 1.095304 | 1.3351 |
| CNAG_130 | - | unspecifie | 5.003257612 | 0.145409756 | 12.80158834 | 20.82509664 | 17.45108887 | 18.67421758 | 0.046246 | 1.674576 | 0.027186 | 2.208837 | 3.192256 | 1.33493 |
| CNAG_060 | - | hypothetic | 66.32844859 | 60.29186166 | 75.53828667 | 44.82038686 | 39.39060577 | 46.75655472 | 0.046246 | -0.65539 | 0.027185 | -2.20885 | 0.634902 | 1.33493 |
| CNAG_020 | - | stromal m | 330.4117912 | 356.5397068 | 352.9258508 | 309.704757 | 249.0086121 | 293.1628761 | 0.046455 | -0.30132 | 0.027315 | -2.20699 | 0.81151 | 1.332969 |
| CNAG_014 | SWI6 | Cell-cycle | 195.2783391 | 240.2571265 | 173.9870028 | 304.1451368 | 265.8283799 | 227.6225846 | 0.0469 | 0.374435 | 0.027599 | 2.202938 | 1.296332 | 1.32883 |
| CNAG_131 | - | unspecifie | 38.23074292 | 43.93434625 | 33.01221887 | 26.31834688 | 23.11558589 | 10.34134775 | 0.046975 | -0.96532 | 0.027649 | -2.20223 | 0.512166 | 1.328131 |
| CNAG_127 | - | unspecifie | 15.4589673 | 13.09314028 | 16.09998525 | 33.24690052 | 28.64498565 | 29.80272602 | 0.047198 | 1.016775 | 0.027792 | 2.200214 | 2.02339 | 1.32608 |
| CNAG_121 | - | unspecifie | 9.721469271 | 21.52585176 | 3.399505955 | 33.49769073 | 19.82136304 | 30.7816669 | 0.04782 | 1.303288 | 0.02817 | 2.194908 | 2.467906 | 1.320387 |
| CNAG_068 | - | mitochond | 1470.07973 | 1341.420457 | 1512.162452 | 1229.473982 | 1235.916945 | 1374.377523 | 0.048029 | -0.18727 | 0.028305 | -2.19304 | 0.878267 | 1.3185 |
| CNAG_071 | - | origin rec | 326.0868092 | 333.0457714 | 404.1778351 | 470.4786533 | 433.1194825 | 406.3000699 | 0.048071 | 0.284896 | 0.028336 | 2.192609 | 1.218322 | 1.318115 |
| CNAG_031 | - | hypothetic | 1533.162093 | 1468.5305 | 1538.851336 | 1307.314661 | 1361.038507 | 1417.699864 | 0.048115 | -0.16746 | 0.028373 | -2.19209 | 0.890412 | 1.317718 |
| CNAG_122 | - | unspecifie | 28.78718425 | 25.73422709 | 25.4925681 | 11.23111979 | 8.277780644 | 17.81210155 | 0.048158 | -1.13038 | 0.028405 | -2.19165 | 0.456795 | 1.317328 |
| CNAG_004 | - | glyoxal ox | 3042.395276 | 1385.632301 | 1320.34371 | 1095.276667 | 1080.432383 | 955.1029682 | 0.048272 | -0.89566 | 0.028483 | -2.19057 | 0.537503 | 1.316308 |
| CNAG_022 | - | cell cycle | 705.0913976 | 581.8588078 | 703.7691161 | 611.7092897 | 532.8799659 | 549.648234 | 0.048737 | -0.24935 | 0.028764 | -2.18671 | 0.841276 | 1.312139 |
| CNAG_038 | - | hypothetic | 871.0294584 | 814.5396916 | 901.0833216 | 807.0648518 | 740.8183293 | 716.1826344 | 0.048819 | -0.20808 | 0.028824 | -2.18588 | 0.865691 | 1.311408 |
| CNAG_077 | - | ABC trans | 800.2796759 | 1108.630183 | 661.4233186 | 1136.500747 | 1178.436855 | 1325.227596 | 0.048819 | 0.489415 | 0.028822 | 2.185916 | 1.403875 | 1.311408 |
| CNAG_056 | PUT2 | 1-pyrrolin | 16933.38204 | 15440.67324 | 14898.36811 | 13707.80727 | 14965.41016 | 14686.35969 | 0.048863 | -0.14032 | 0.028856 | -2.18545 | 0.907316 | 1.311016 |
| CNAG_060 | - | cytoplasm | 47.05412794 | 58.24795063 | 48.12023695 | 75.76472038 | 85.73288743 | 74.2692939 | 0.048906 | 0.598347 | 0.028887 | 2.185023 | 1.513981 | 1.31064 |
| CNAG_127 | - | unspecifie | 114.2921806 | 73.44398045 | 55.68737441 | 128.8962025 | 96.52842497 | 146.1391265 | 0.048938 | 0.596429 | 0.028912 | 2.184683 | 1.51197 | 1.310355 |
| CNAG_001 | UBP16 | ubiquitin | 1787.762134 | 1859.584882 | 1723.387253 | 1563.387564 | 1609.452116 | 1688.574756 | 0.048963 | -0.15866 | 0.028939 | -2.18432 | 0.895858 | 1.310135 |
| CNAG_050 | - | DNA/RNA | 841.8966662 | 787.629345 | 970.3454328 | 749.3179327 | 731.3141342 | 773.837593 | 0.049184 | -0.2212 | 0.029082 | -2.18238 | 0.857854 | 1.308177 |
| CNAG_041 | - | oxysterol- | 3899.034889 | 3956.27828 | 3623.822782 | 3501.407925 | 3457.054194 | 3596.953277 | 0.049239 | -0.13611 | 0.02912 | -2.18186 | 0.909971 | 1.307693 |
| CNAG_052 | - | solute car | 1170.074769 | 1215.307693 | 1264.754119 | 1088.692665 | 1033.201173 | 1135.718445 | 0.049459 | -0.17943 | 0.029256 | -2.18002 | 0.88305 | 1.305754 |
| CNAG_007 | DPH1 | diphthami | 907.1495497 | 749.951278 | 938.3760111 | 759.4689626 | 759.8286718 | 722.8326312 | 0.049481 | -0.2279 | 0.029275 | -2.17976 | 0.853877 | 1.30556 |

| Gene_ID | gene_nam | Function | WT(H99)_1_DESeq2_normalized_coun | WT(H99)_2_DESeq2_normalized_coun | WT(H99)_3_DESeq2_normalized_coun | *hht2*Δ mutant_1_DESeq2_normalized_cou | *hht2*Δ mutant_2_DESeq2_normalized_cou | *hht2*Δ mutant_3_DESeq2_normalized_cou | padj | log2FoldC | pvalue | stat | foldChang | log10padj |
|---|---|---|---|---|---|---|---|---|---|---|---|---|---|---|
| CNAG_058 | - | endoribon | 289.4359451 | 284.9050809 | 307.5334983 | 2631.762714 | 2566.375465 | 2642.331183 | 1.00E-50 | 3.139844 | 2.38E-292 | 36.54215 | 8.814289 | 288.188 |
| CNAG_105 | - | unspecifie | 212236.7795 | 232456.525 | 238929.797 | 46437.80732 | 35447.86589 | 47200.20725 | 1.00E-50 | -2.41931 | 3.24E-168 | -27.6446 | 0.186945 | 164.5298 |
| CNAG_034 | GRE202 | oxidoredu | 311.5311983 | 351.6450886 | 298.9717957 | 1507.394863 | 1615.762301 | 1533.512183 | 1.00E-50 | 2.262824 | 1.95E-124 | 23.7259 | 4.7993 | 120.9731 |
| CNAG_067 | H3 | Histone H | 4976.280707 | 5347.972113 | 5278.613174 | 1891.861074 | 1945.370038 | 1964.325258 | 1.00E-50 | -1.44247 | 7.29E-122 | -23.4753 | 0.367937 | 118.4546 |
| CNAG_032 | - | large subu | 48568.51403 | 48515.41322 | 50756.48573 | 21236.3777 | 22640.89556 | 21286.63808 | 1.00E-50 | -1.19736 | 4.39E-115 | -22.8019 | 0.436072 | 111.7235 |
| CNAG_058 | - | small subu | 47368.08381 | 47359.33595 | 51714.12206 | 21259.74634 | 21190.89638 | 20432.93646 | 1.00E-50 | -1.23505 | 1.59E-108 | -22.1311 | 0.424827 | 105.2482 |
| CNAG_074 | - | U3 small r | 442315.4359 | 435261.7619 | 363444.9676 | 152310.4886 | 135643.9974 | 139878.5612 | 1.00E-50 | -1.55141 | 7.23E-99 | -21.1045 | 0.341177 | 95.74017 |
| CNAG_037 | - | large subu | 56265.24962 | 56311.74948 | 57218.18768 | 27901.76113 | 28956.61768 | 28409.36895 | 1.00E-50 | -1.00906 | 2.68E-98 | -21.0425 | 0.49687 | 95.19007 |
| CNAG_055 | - | large subu | 72627.72158 | 69521.08798 | 79484.56113 | 32744.98446 | 33375.45463 | 32084.06231 | 1.00E-50 | -1.19001 | 2.04E-86 | -19.7028 | 0.438299 | 83.43027 |
| CNAG_052 | RPL2 | large subu | 69678.04811 | 68077.33661 | 74129.95884 | 34605.17827 | 34369.15618 | 34674.01597 | 1.00E-50 | -1.04704 | 4.31E-85 | -19.5478 | 0.483961 | 82.15996 |
| CNAG_054 | - | Dihydroxy | 21456.91824 | 22419.99822 | 21339.52808 | 10624.48203 | 11247.38365 | 11163.00458 | 1.00E-50 | -0.99627 | 9.54E-83 | -19.2703 | 0.501295 | 79.88288 |
| CNAG_026 | ISP4 | Identified | 229.7664583 | 243.9006142 | 225.3757555 | 959.874231 | 1115.417395 | 1179.190549 | 1.00E-50 | 2.207563 | 1.98E-82 | 19.23256 | 4.618943 | 79.58327 |
| CNAG_032 | - | large subu | 14569.48096 | 15151.62758 | 17881.84365 | 6128.14049 | 6118.452172 | 5883.206813 | 1.00E-50 | -1.40808 | 2.49E-82 | -19.2205 | 0.376812 | 79.48924 |
| CNAG_059 | - | cytochrom | 13804.60551 | 14109.77155 | 14867.13009 | 6166.33604 | 6854.328893 | 6534.791586 | 1.00E-50 | -1.14482 | 3.91E-82 | -19.1972 | 0.452246 | 79.30132 |
| CNAG_052 | - | alpha-amy | 868.4258059 | 700.7376649 | 902.5226748 | 3117.061388 | 2851.515126 | 2771.522838 | 1.00E-50 | 1.806726 | 2.00E-81 | 19.11217 | 3.498474 | 78.62467 |
| CNAG_037 | - | large subu | 84544.66302 | 81775.80647 | 95900.45156 | 38365.79014 | 38470.58728 | 37934.60929 | 1.00E-50 | -1.20768 | 2.86E-79 | -18.8514 | 0.432966 | 76.50498 |
| CNAG_036 | CAS3 | capsule re | 83.86696317 | 98.48012665 | 120.8662458 | 720.3111906 | 681.6164918 | 628.7802638 | 1.00E-50 | 2.737817 | 1.63E-76 | 18.51261 | 6.670603 | 73.7874 |
| CNAG_037 | - | small subu | 75723.52114 | 73726.32753 | 82057.96281 | 38067.01791 | 38048.84925 | 38193.22336 | 1.00E-50 | -1.03363 | 3.25E-75 | -18.3509 | 0.488478 | 72.50488 |
| CNAG_059 | - | pyruvate c | 11351.00915 | 11020.76348 | 10356.13682 | 5160.092842 | 5053.265356 | 5347.996734 | 1.00E-50 | -1.08785 | 5.61E-75 | -18.3212 | 0.470461 | 72.27247 |
| CNAG_051 | - | chaperone | 22577.81113 | 22519.10567 | 22901.80344 | 9014.034158 | 9457.452046 | 10845.5107 | 1.00E-50 | -1.2289 | 7.00E-73 | -18.0566 | 0.426642 | 70.21461 |
| CNAG_057 | - | large subu | 34703.939 | 35264.12082 | 38974.29685 | 16895.90466 | 18208.23806 | 17644.14994 | 1.00E-50 | -1.0618 | 4.11E-69 | -17.571 | 0.479035 | 66.50264 |
| CNAG_031 | URA5 | Orotate pl | 3462.879507 | 3307.459077 | 3508.351519 | 1555.806614 | 1534.006692 | 1549.061302 | 1.00E-50 | -1.16333 | 1.44E-68 | -17.4996 | 0.446482 | 65.96888 |
| CNAG_035 | - | 3-beta hyc | 239.1217192 | 296.6746295 | 311.7223567 | 1048.639242 | 1239.702301 | 1137.239508 | 1.00E-50 | 2.002075 | 2.00E-68 | 17.48098 | 4.005759 | 65.83493 |
| CNAG_055 | PLR1 | pyridoxal | 805.302745 | 791.1736418 | 840.6400724 | 2078.007281 | 2314.080969 | 2499.382177 | 1.00E-50 | 1.4849 | 1.79E-66 | 17.22267 | 2.798978 | 63.91189 |
| CNAG_009 | - | hypothetic | 66.3678462 | 131.8588762 | 88.89586282 | 681.7437303 | 678.0369707 | 797.1079597 | 1.00E-50 | 2.906404 | 6.02E-65 | 17.0182 | 7.497471 | 62.41784 |
| CNAG_035 | - | mandelate | 466.7421248 | 429.2087954 | 572.960177 | 1592.729219 | 1634.447396 | 1696.758902 | 1.00E-50 | 1.730629 | 2.03E-63 | 16.81081 | 3.318724 | 60.91203 |
| CNAG_033 | - | Succinyl-C | 12189.64105 | 11856.93174 | 13253.59228 | 6052.028906 | 6378.224302 | 6163.96783 | 1.00E-50 | -1.01992 | 2.97E-62 | -16.6511 | 0.493142 | 59.76319 |
| CNAG_037 | - | 3-phosphc | 8996.889947 | 8384.361703 | 8676.446202 | 4495.517025 | 4270.949856 | 4493.152765 | 1.00E-50 | -0.99025 | 1.56E-61 | -16.5514 | 0.50339 | 59.05402 |
| CNAG_049 | - | zuotin | 6810.914678 | 6448.223134 | 7060.720929 | 3301.554081 | 3391.156961 | 3435.177193 | 1.00E-50 | -1.02013 | 7.52E-61 | -16.4566 | 0.493071 | 58.38062 |
| CNAG_055 | - | hypothetic | 38649.66603 | 35674.67673 | 44428.0635 | 18031.43276 | 17910.59549 | 17464.3456 | 1.00E-50 | -1.1688 | 1.46E-57 | -15.9915 | 0.44479 | 55.13051 |
| CNAG_068 | UAP1 | uric acid x | 213.4009751 | 284.7860177 | 264.7906518 | 889.9702189 | 972.1752834 | 893.0969932 | 1.00E-50 | 1.841044 | 1.35E-55 | 15.70698 | 3.582691 | 53.18533 |
| CNAG_036 | - | Threonine | 3119.788162 | 3417.30931 | 3394.264945 | 1629.031084 | 1641.882987 | 1575.690141 | 1.00E-50 | -1.05024 | 4.85E-55 | -15.626 | 0.482887 | 52.6395 |
| CNAG_078 | - | Histone H | 8952.698623 | 11298.19763 | 10990.01934 | 23772.27384 | 23278.48224 | 22388.24271 | 1.00E-50 | 1.138004 | 3.08E-53 | 15.35907 | 2.200763 | 50.85114 |
| CNAG_055 | HIS3 | Imidazoleg | 5513.455727 | 5021.706808 | 5181.691721 | 2683.640548 | 2685.59261 | 2715.785292 | 1.02E-50 | -0.97483 | 2.31E-52 | -15.2278 | 0.5088 | 49.98984 |
| CNAG_031 | - | small subu | 111752.2731 | 110867.601 | 129792.3853 | 60518.39933 | 59631.00413 | 59163.05046 | 1.21E-50 | -0.99031 | 2.75E-52 | -15.2164 | 0.503371 | 49.9188 |
| CNAG_034 | - | hypothetic | 70.98152194 | 95.20938128 | 88.86231184 | 448.0997714 | 510.953151 | 443.4112643 | 5.90E-50 | 2.454512 | 1.40E-51 | 15.10974 | 5.481277 | 49.22938 |
| CNAG_033 | - | small subu | 18007.39082 | 17601.80007 | 19054.06362 | 10162.34309 | 9151.331917 | 8908.301617 | 7.18E-50 | -0.96922 | 1.72E-51 | -15.0961 | 0.510784 | 49.14401 |
| CNAG_054 | - | 2-isopropy | 9872.989895 | 11886.94041 | 10678.1869 | 5255.832024 | 5667.488578 | 5430.145637 | 3.37E-48 | -1.00255 | 8.39E-50 | -14.8375 | 0.499115 | 47.47287 |
| CNAG_049 | - | Nascent p | 8722.771596 | 9820.515294 | 10622.61982 | 4735.1697 | 4926.528977 | 4633.075779 | 9.54E-48 | -1.04378 | 2.41E-49 | -14.7665 | 0.485055 | 47.02065 |
| CNAG_075 | - | calcium/pr | 1119.304913 | 851.5328035 | 780.9708922 | 2676.057689 | 2784.240567 | 2710.457071 | 1.19E-47 | 1.554626 | 3.01E-49 | 14.75143 | 2.937575 | 46.92593 |
| CNAG_027 | - | T-complex | 7017.70758 | 7081.73064 | 7402.011024 | 3629.241228 | 3580.906554 | 4022.394936 | 2.11E-47 | -0.95197 | 5.47E-49 | -14.7111 | 0.516926 | 46.67522 |
| CNAG_058 | - | large subu | 15891.41574 | 15160.35994 | 17006.45819 | 8826.779434 | 8206.84287 | 7797.033127 | 3.57E-47 | -0.96836 | 9.33E-49 | -14.6749 | 0.511088 | 46.44711 |
| CNAG_126 | - | unspecifie | 1638.977883 | 1765.574157 | 1876.602818 | 742.4540239 | 641.1560906 | 756.4132265 | 7.71E-46 | -1.31816 | 2.09E-47 | -14.4626 | 0.401047 | 45.11322 |
| CNAG_052 | XKS1 | xylulokina | 885.7037464 | 784.6695065 | 802.2226459 | 1844.562287 | 1893.002904 | 1958.940659 | 7.91E-46 | 1.188977 | 2.16E-47 | 14.4604 | 2.279911 | 45.10156 |
| CNAG_051 | - | ATP-bindi | 3537.761961 | 3519.733939 | 3448.724838 | 1928.774162 | 1896.381021 | 1847.638006 | 2.42E-45 | -0.90453 | 6.68E-47 | -14.3824 | 0.534207 | 44.61628 |
| CNAG_079 | - | hypothetic | 3705.719922 | 3588.06973 | 3298.760897 | 6681.544116 | 6542.056773 | 6724.99744 | 5.20E-45 | 0.898108 | 1.46E-46 | 14.32834 | 1.86362 | 44.28358 |
| CNAG_037 | - | conidiatio | 9.031092404 | 6.856382731 | 1.37144751 | 274.3570513 | 267.6743983 | 295.2824843 | 8.20E-44 | 5.69222 | 2.36E-45 | 14.13343 | 51.70456 | 43.08611 |
| CNAG_059 | - | small subu | 38279.6471 | 37092.59669 | 42857.28016 | 20729.08622 | 22028.5792 | 21387.56706 | 1.01E-43 | -0.89784 | 2.94E-45 | -14.1181 | 0.53669 | 42.9951 |
| CNAG_050 | - | hypothetic | 32.43412052 | 38.1244306 | 13.1524087 | 331.2637547 | 276.829659 | 329.8282574 | 3.11E-43 | 3.481303 | 9.24E-45 | 14.03711 | 11.16803 | 42.50668 |
| CNAG_055 | - | small subu | 40361.82215 | 40838.8189 | 46208.08761 | 23773.82613 | 24012.46981 | 23559.03481 | 1.82E-42 | -0.852 | 5.61E-44 | -13.9087 | 0.554018 | 41.73952 |
| CNAG_058 | PRM1 | plasma m | 140.9306578 | 152.2722424 | 127.2409325 | 509.3197503 | 588.4274013 | 516.7671738 | 1.80E-41 | 1.929653 | 5.72E-43 | 13.74154 | 3.809636 | 40.74401 |
| CNAG_127 | - | unspecifie | 862.4122226 | 938.0898454 | 1050.027127 | 342.3064139 | 329.7721127 | 352.3640132 | 4.71E-41 | -1.49318 | 1.51E-42 | -13.6711 | 0.35523 | 40.3273 |
| CNAG_031 | - | saccharop | 508.1686135 | 478.0080884 | 534.9149226 | 136.7094439 | 145.1468321 | 125.986899 | 7.05E-41 | -1.92023 | 2.31E-42 | -13.6403 | 0.264212 | 40.15184 |
| CNAG_057 | - | ketol-acid | 35210.79862 | 34675.90129 | 34627.58452 | 20548.62235 | 22396.98513 | 20600.43219 | 7.36E-41 | -0.73331 | 2.43E-42 | -13.6366 | 0.601521 | 40.13317 |
| CNAG_058 | TRR1 | Thioredox | 2141.106407 | 2156.757228 | 2116.702332 | 1040.616091 | 1123.519902 | 1103.829528 | 1.05E-40 | -0.98872 | 3.50E-42 | -13.6098 | 0.503925 | 39.97887 |
| CNAG_067 | - | succinate | 6523.738975 | 6192.908282 | 6880.511401 | 3740.681211 | 3480.887748 | 3485.122146 | 2.24E-40 | -0.88777 | 7.48E-42 | -13.5542 | 0.54045 | 39.65045 |
| CNAG_057 | - | alanine-tR | 5101.35073 | 4787.919516 | 5236.039558 | 2450.86119 | 2672.676676 | 2787.90622 | 2.48E-40 | -0.95087 | 8.31E-42 | -13.5465 | 0.51732 | 39.60642 |
| CNAG_032 | - | Adenylosu | 7826.861999 | 7203.533679 | 7673.987206 | 3816.787479 | 4249.475342 | 4265.526877 | 8.39E-40 | -0.89636 | 2.85E-41 | -13.4558 | 0.537242 | 39.07624 |
| CNAG_026 | - | translation | 10449.6554 | 9374.5944 | 10406.23465 | 5636.548127 | 5777.941102 | 5569.988141 | 1.13E-39 | -0.84781 | 3.89E-41 | -13.4327 | 0.555626 | 38.94719 |
| CNAG_035 | - | large subu | 8583.825937 | 9100.864843 | 9980.67161 | 5189.042826 | 5178.44494 | 5077.077716 | 7.01E-39 | -0.85609 | 2.51E-40 | -13.2939 | 0.552448 | 38.15398 |
| CNAG_034 | - | Spermidin | 10633.17538 | 10566.21261 | 11491.88225 | 6249.070635 | 6687.12542 | 6410.349928 | 2.97E-37 | -0.77229 | 1.12E-38 | -13.0067 | 0.585486 | 36.52721 |
| CNAG_034 | - | endo alph | 2852.393845 | 2922.852591 | 2986.92571 | 1674.315468 | 1577.22302 | 1622.24785 | 9.82E-37 | -0.86125 | 3.75E-38 | -12.914 | 0.550475 | 36.00809 |
| CNAG_058 | CHS5 | Chitin syn | 3555.071128 | 3484.626456 | 2991.638889 | 6170.893489 | 6379.079404 | 6482.961979 | 3.94E-36 | 0.908896 | 1.55E-37 | 12.80411 | 1.877608 | 35.40427 |
| CNAG_051 | - | hypothetic | 3778.460041 | 3937.794582 | 3865.720524 | 2208.968525 | 2287.17233 | 2312.729354 | 4.52E-36 | -0.78161 | 1.79E-37 | -12.7931 | 0.581716 | 35.34529 |
| CNAG_050 | - | hypothetic | 4.301300246 | 2.515281294 | 6.730717124 | 225.2218604 | 245.4885605 | 225.5587371 | 7.18E-36 | 5.8403 | 2.88E-37 | 12.75606 | 57.29352 | 35.14392 |
| CNAG_080 | - | hypothetic | 308.9620203 | 309.5346102 | 329.7838471 | 861.1631918 | 769.8827666 | 834.3259421 | 3.40E-35 | 1.365951 | 1.41E-36 | 12.63203 | 2.577463 | 34.46897 |
| CNAG_036 | - | translation | 8791.574189 | 8367.190594 | 8963.352076 | 5108.895709 | 5432.969421 | 5280.19 | 5.93E-35 | -0.73906 | 2.47E-36 | -12.5877 | 0.599128 | 34.22702 |

| | | | | | | | | | | | | | | |
|---|---|---|---|---|---|---|---|---|---|---|---|---|---|---|
| CNAG_037 | - | efflux prot | 79.08521441 | 95.17709805 | 95.22593483 | 374.1266476 | 368.2349956 | 378.9961667 | 6.94E-35 | 2.043043 | 2.91E-36 | 12.57458 | 4.121138 | 34.15891 |
| CNAG_025 | - | homoacor | 1918.098608 | 1946.665004 | 1885.300034 | 1013.463303 | 1055.406304 | 973.9582029 | 1.05E-34 | -0.93402 | 4.43E-36 | -12.5414 | 0.523398 | 33.97815 |
| CNAG_048 | - | hypothetic | 325.2907615 | 306.3121983 | 303.1511952 | 904.8512301 | 752.5471724 | 896.4049153 | 1.48E-34 | 1.435011 | 6.24E-36 | 12.51422 | 2.703842 | 33.83054 |
| CNAG_026 | - | translatior | 2582.378291 | 2383.052501 | 2524.044783 | 1155.771079 | 1343.512195 | 1057.301349 | 5.19E-34 | -1.09091 | 2.24E-35 | -12.4125 | 0.469466 | 33.28505 |
| CNAG_127 | - | unspecifie | 44.05547146 | 29.50493565 | 41.93290538 | 300.1519809 | 269.2470413 | 231.5072725 | 5.80E-34 | 2.794687 | 2.52E-35 | 12.40291 | 6.938804 | 33.23683 |
| CNAG_037 | - | AdoMet-d | 5994.700041 | 5545.377176 | 5795.991469 | 3128.065052 | 3340.344372 | 3486.158398 | 1.14E-33 | -0.81585 | 5.01E-35 | -12.3478 | 0.568075 | 32.94334 |
| CNAG_033 | BZP4 | bZip trans | 60.41444927 | 79.01440694 | 61.12841866 | 338.5533277 | 296.5167315 | 344.4588107 | 1.99E-33 | 2.275659 | 8.86E-35 | 12.30174 | 4.842187 | 32.70142 |
| CNAG_058 | - | Pyrroline-! | 2496.298719 | 2759.034963 | 2841.83797 | 1309.111445 | 1502.225484 | 1408.01278 | 3.43E-33 | -0.95535 | 1.54E-34 | -12.2569 | 0.515716 | 32.4641 |
| CNAG_068 | KRE61 | glucosidas | 235.5253853 | 300.9144395 | 315.9235719 | 760.437483 | 910.0536718 | 791.0706783 | 1.74E-32 | 1.518874 | 8.05E-34 | 12.12228 | 2.865673 | 31.75966 |
| CNAG_055 | - | small subu | 2218.052846 | 1932.749768 | 2196.635686 | 1053.881098 | 1024.266155 | 1107.14794 | 2.75E-32 | -1.01083 | 1.28E-33 | -12.084 | 0.496261 | 31.5607 |
| CNAG_051 | - | sarcosine | 3875.163383 | 3561.802704 | 3482.893965 | 2124.957426 | 2039.578056 | 2068.52381 | 1.28E-31 | -0.8247 | 6.18E-33 | -11.9541 | 0.564601 | 30.89214 |
| CNAG_052 | - | hypothetic | 34.73459662 | 44.59933376 | 44.05629008 | 238.7097652 | 275.4727373 | 232.5250391 | 1.54E-31 | 2.595367 | 7.49E-33 | 11.93816 | 6.043426 | 30.81129 |
| CNAG_050 | - | hypothetic | 2433.255984 | 2619.00605 | 2861.008943 | 1418.973719 | 1327.039402 | 1435.741945 | 3.04E-31 | -0.9357 | 1.50E-32 | -11.8801 | 0.522789 | 30.5174 |
| CNAG_052 | - | SET doma | 4509.50216 | 4584.293815 | 4648.502601 | 2737.460884 | 2882.377148 | 2889.989013 | 5.71E-31 | -0.70682 | 2.87E-32 | -11.8259 | 0.612671 | 30.24322 |
| CNAG_051 | - | hypothetic | 105.9183783 | 139.311084 | 111.229922 | 466.9403384 | 390.4205717 | 444.5166952 | 1.04E-30 | 1.858546 | 5.23E-32 | 11.77535 | 3.62642 | 29.98423 |
| CNAG_058 | - | cytochrom | 7245.481292 | 6848.022288 | 7470.311574 | 4459.101495 | 4498.81595 | 4211.131906 | 1.26E-30 | -0.7272 | 6.41E-32 | -11.7582 | 0.604076 | 29.89886 |
| CNAG_075 | GPM1 | hypothetic | 732.9697258 | 973.4885469 | 897.6087014 | 350.1905021 | 322.0418924 | 286.912107 | 1.38E-30 | -1.45571 | 7.05E-32 | -11.7502 | 0.364576 | 29.85996 |
| CNAG_033 | - | Mannose- | 218.9837612 | 158.7510942 | 156.0214374 | 589.6559953 | 569.998898 | 557.8823724 | 4.88E-30 | 1.672534 | 2.55E-31 | 11.64105 | 3.187741 | 29.31191 |
| CNAG_036 | - | large subu | 2671.332434 | 2645.960424 | 2606.242229 | 1476.583313 | 1436.624907 | 1598.878386 | 6.06E-30 | -0.82787 | 3.20E-31 | -11.6216 | 0.563361 | 29.21782 |
| CNAG_036 | - | NADH del | 16494.15152 | 16405.90825 | 16293.1472 | 10465.93976 | 11241.51489 | 10890.90078 | 6.34E-30 | -0.60907 | 3.36E-31 | -11.6174 | 0.655619 | 29.1976 |
| CNAG_031 | - | hypothetic | 2690.101551 | 2752.631791 | 2912.253252 | 1538.841367 | 1667.019649 | 1445.776063 | 1.24E-29 | -0.86067 | 6.76E-31 | -11.5576 | 0.550695 | 28.90721 |
| CNAG_032 | - | succinate | 2714.14325 | 2203.179894 | 2311.883187 | 1236.821575 | 1146.739946 | 1197.10159 | 1.38E-29 | -1.0301 | 7.56E-31 | -11.5479 | 0.489677 | 28.86039 |
| CNAG_076 | - | ATP-depe | 8100.156543 | 7559.089376 | 7636.712234 | 4144.486176 | 4589.347373 | 4846.084034 | 1.61E-29 | -0.79416 | 8.88E-31 | -11.5341 | 0.576678 | 28.79263 |
| CNAG_033 | ERG13 | hydroxym | 10463.80372 | 9719.395169 | 10493.73092 | 6414.939893 | 6631.087446 | 6392.561598 | 2.16E-29 | -0.67397 | 1.20E-30 | -11.5081 | 0.626779 | 28.66517 |
| CNAG_033 | - | flavonol sy | 45.20343989 | 47.78542516 | 30.21254278 | 258.4606427 | 277.20648 | 213.7968261 | 3.03E-29 | 2.601004 | 1.70E-30 | 11.47787 | 6.067089 | 28.51798 |
| CNAG_036 | - | DNA-direc | 1163.714417 | 1176.297645 | 1213.372208 | 622.7956109 | 573.1575191 | 537.7101202 | 3.34E-29 | -1.05147 | 1.88E-30 | -11.4694 | 0.482476 | 28.47629 |
| CNAG_056 | - | hypothetic | 123.3761778 | 132.8600411 | 144.2413809 | 421.3907911 | 419.0195569 | 427.9443573 | 8.33E-29 | 1.651857 | 4.77E-30 | 11.38848 | 3.142378 | 28.07923 |
| CNAG_068 | - | hypothetic | 1255.940945 | 1257.103836 | 1258.191187 | 599.3172302 | 594.7003466 | 680.8182861 | 1.04E-28 | -1.02431 | 6.00E-30 | -11.3685 | 0.491646 | 27.98248 |
| CNAG_037 | - | staphyloc | 5428.40618 | 5145.653462 | 5316.101207 | 3223.158446 | 2961.638008 | 3341.81511 | 2.13E-28 | -0.75356 | 1.27E-29 | -11.3032 | 0.593138 | 27.67126 |
| CNAG_026 | - | hypothetic | 109.4115728 | 144.6891672 | 114.4255476 | 466.1400844 | 453.7079709 | 375.9755139 | 2.99E-28 | 1.803526 | 1.79E-29 | 11.27274 | 3.490724 | 27.52431 |
| CNAG_037 | - | ribosomal | 5954.935732 | 5112.297136 | 5655.213119 | 3160.810014 | 3336.059303 | 3217.534774 | 3.05E-28 | -0.79973 | 1.83E-29 | -11.2707 | 0.574456 | 27.51534 |
| CNAG_034 | - | hypothetic | 66.18531757 | 70.38955725 | 58.98314443 | 294.0989946 | 298.5240298 | 255.9699844 | 1.11E-27 | 2.111188 | 6.76E-29 | 11.15511 | 4.320468 | 26.95486 |
| CNAG_027 | - | hypothetic | 35.82896201 | 29.47598399 | 19.53961005 | 201.5998976 | 201.0842357 | 191.3786237 | 1.88E-27 | 2.821424 | 1.16E-28 | 11.10728 | 7.068598 | 26.72697 |
| CNAG_067 | - | hypothetic | 2192.649722 | 2192.395568 | 2474.971152 | 1309.30054 | 1222.70542 | 1257.009396 | 1.95E-27 | -0.8717 | 1.21E-28 | -11.1035 | 0.546503 | 26.70957 |
| CNAG_034 | - | hypothetic | 1942.551489 | 1751.742175 | 1905.537412 | 1001.795804 | 990.6757616 | 1054.936662 | 2.06E-27 | -0.89412 | 1.28E-28 | -11.0984 | 0.538074 | 26.68556 |
| CNAG_036 | STE6 | ATP-bindi | 79.01917738 | 88.68827864 | 71.7753963 | 317.3044328 | 287.8376599 | 331.2067698 | 3.60E-27 | 1.959415 | 2.28E-28 | 11.04634 | 3.889043 | 26.44335 |
| CNAG_124 | - | unspecifie | 8.990126283 | 13.31389342 | 15.26015864 | 204.2065337 | 128.1622635 | 162.4041692 | 3.80E-27 | 3.764173 | 2.42E-28 | 11.04098 | 13.58717 | 26.42076 |
| CNAG_037 | - | hypothetic | 13.68811807 | 28.38488373 | 18.46535514 | 198.1178358 | 172.535329 | 166.9890046 | 3.81E-27 | 3.16977 | 2.44E-28 | 11.04055 | 8.999035 | 26.41954 |
| CNAG_051 | - | hypothetic | 72.03737136 | 86.534338 | 80.28830021 | 295.3155125 | 317.4160531 | 303.5826461 | 3.88E-27 | 1.927973 | 2.49E-28 | 11.03857 | 3.805203 | 26.41078 |
| CNAG_033 | - | hypothetic | 2182.09076 | 2108.367796 | 2316.108167 | 1203.374541 | 1265.885637 | 1288.057926 | 4.72E-27 | -0.83007 | 3.04E-28 | -11.0205 | 0.562503 | 26.32586 |
| CNAG_052 | - | Glutamate | 3177.836238 | 3282.018191 | 2880.684033 | 5317.913339 | 5156.726076 | 5014.368086 | 1.47E-26 | 0.714739 | 9.76E-28 | 10.91511 | 1.641186 | 25.83366 |
| CNAG_049 | - | hypothetic | 1365.88245 | 1443.523194 | 1552.483152 | 775.3282619 | 709.4307864 | 782.968139 | 1.56E-26 | -0.95919 | 1.04E-27 | -10.9092 | 0.514347 | 25.80606 |
| CNAG_034 | - | hypothetic | 1140.25609 | 1322.183423 | 1063.488386 | 2281.903784 | 2148.118574 | 2342.033388 | 1.89E-26 | 0.927452 | 1.27E-27 | 10.8912 | 1.901914 | 25.72267 |
| CNAG_025 | - | Sugar trar | 31.21462034 | 45.61444296 | 30.20219817 | 259.7889391 | 197.3351038 | 202.5818437 | 3.54E-26 | 2.620041 | 2.41E-27 | 10.83271 | 6.147674 | 25.45077 |
| CNAG_058 | - | protein FA | 4038.799968 | 3859.199331 | 3931.857151 | 2553.136972 | 2287.140606 | 2390.442698 | 4.50E-26 | -0.72549 | 3.08E-27 | -10.8103 | 0.60479 | 25.34692 |
| CNAG_034 | - | protein GI | 2802.207305 | 2717.131495 | 2903.763217 | 1689.902903 | 1755.010122 | 1691.044624 | 6.32E-26 | -0.7293 | 4.37E-27 | -10.7781 | 0.603197 | 25.19911 |
| CNAG_068 | - | elongatior | 109713.4274 | 107226.9944 | 105211.9045 | 65427.95577 | 73166.49511 | 72996.61743 | 7.06E-26 | -0.62185 | 4.89E-27 | -10.7676 | 0.649838 | 25.1514 |
| CNAG_075 | - | ATP-bindi | 1986.58715 | 1606.26915 | 1609.119898 | 855.4135013 | 862.9929327 | 851.8462078 | 7.94E-26 | -1.03426 | 5.52E-27 | -10.7565 | 0.488267 | 25.10035 |
| CNAG_055 | - | ATPase | 753.7380215 | 757.6811856 | 750.9976592 | 1354.607341 | 1569.506516 | 1702.441913 | 9.21E-26 | 1.017973 | 6.44E-27 | 10.74238 | 2.025072 | 25.03551 |
| CNAG_027 | - | hypothetic | 1919.458909 | 2016.753756 | 2254.205239 | 1116.208368 | 1179.635121 | 1113.809648 | 9.42E-26 | -0.87608 | 6.59E-27 | -10.7402 | 0.544845 | 25.02584 |
| CNAG_050 | GCS2 | ADP-ribos | 2203.10546 | 2078.221003 | 2360.887894 | 1289.071052 | 1252.931791 | 1256.995932 | 1.82E-25 | -0.82215 | 1.30E-26 | -10.6776 | 0.5656 | 24.73924 |
| CNAG_055 | - | hypothetic | 715.1678047 | 764.101452 | 620.9236279 | 1460.166494 | 1386.573108 | 1357.409039 | 1.84E-25 | 0.986935 | 1.31E-26 | 10.67682 | 1.98197 | 24.73626 |
| CNAG_027 | - | mitochonc | 192.1568622 | 195.3085475 | 182.6327144 | 501.4191485 | 477.5030711 | 525.6509758 | 2.29E-25 | 1.386353 | 1.65E-26 | 10.65525 | 2.614171 | 24.64067 |
| CNAG_068 | - | oxidoredu | 150.1497124 | 138.2461525 | 148.508538 | 433.0495425 | 462.2978642 | 391.5079616 | 3.19E-25 | 1.544788 | 2.32E-26 | 10.62337 | 2.917611 | 24.49574 |
| CNAG_032 | - | hypothetic | 6401.20635 | 5941.922822 | 7382.756336 | 3790.486475 | 3823.390464 | 3479.58321 | 3.79E-25 | -0.8464 | 2.76E-26 | -10.6071 | 0.556172 | 24.42104 |
| CNAG_123 | - | unspecifie | 28.83747879 | 30.54056408 | 13.13696646 | 191.5691248 | 199.5216606 | 163.7654274 | 4.61E-25 | 2.946534 | 3.37E-26 | 10.58861 | 7.708946 | 24.33642 |
| CNAG_074 | - | hypothetic | 44.04926407 | 66.05799335 | 45.12451939 | 229.9125294 | 265.8161061 | 246.9023328 | 4.89E-25 | 2.241132 | 3.59E-26 | 10.58253 | 4.727677 | 24.31039 |
| CNAG_057 | - | protein M | 2669.014767 | 2445.643208 | 2864.242098 | 1460.214827 | 1550.542276 | 1532.305016 | 6.26E-25 | -0.82851 | 4.63E-26 | -10.5588 | 0.563109 | 24.20339 |
| CNAG_035 | - | 3-beta hy | 14.82350581 | 17.61249364 | 9.922835446 | 127.088968 | 178.2488589 | 145.8208505 | 6.63E-25 | 3.476876 | 4.90E-26 | 10.5533 | 11.13381 | 24.17865 |
| CNAG_068 | SXI1alpha | Alpha cell | 144.4157968 | 200.6743558 | 166.6486765 | 545.434304 | 487.1018767 | 458.1025053 | 1.70E-24 | 1.532606 | 1.30E-25 | 10.46125 | 2.893079 | 23.76828 |
| CNAG_026 | - | hypothetic | 1096.93289 | 1063.632026 | 1014.393161 | 1868.48679 | 1999.049697 | 1793.786004 | 2.03E-24 | 0.819504 | 1.56E-25 | 10.44378 | 1.764799 | 23.69197 |
| CNAG_027 | DCR1 | dicer-like | 311.2504793 | 328.8877037 | 336.1582041 | 782.4191867 | 669.6734443 | 793.2380694 | 3.73E-24 | 1.187755 | 2.91E-25 | 10.38475 | 2.27798 | 23.42848 |
| CNAG_035 | - | hypothetic | 58.0361672 | 81.12652261 | 45.13700687 | 281.6162736 | 260.339501 | 280.1745976 | 3.76E-24 | 2.143934 | 2.94E-25 | 10.38371 | 4.419657 | 23.42444 |
| CNAG_027 | - | hypothetic | 13.62802171 | 6.827437705 | 10.98022394 | 136.3709115 | 124.1977417 | 133.5250815 | 5.07E-24 | 3.744342 | 3.97E-25 | 10.35502 | 13.40169 | 23.29541 |
| CNAG_052 | TIM54 | mitochonc | 1241.920824 | 1123.603644 | 1291.212349 | 639.1056248 | 637.8915623 | 632.010518 | 5.14E-24 | -0.95358 | 4.03E-25 | -10.3535 | 0.51635 | 23.28914 |
| CNAG_067 | - | DNA-direc | 1370.204286 | 1157.005228 | 1289.110713 | 630.488994 | 688.820414 | 603.1923616 | 8.49E-24 | -1.00638 | 6.71E-25 | -10.3046 | 0.497794 | 23.0713 |
| CNAG_051 | - | hypothetic | 187.6095451 | 249.1611015 | 258.3057632 | 601.8380534 | 593.1670687 | 592.3585254 | 2.04E-23 | 1.349246 | 1.64E-24 | 10.21843 | 2.547789 | 22.69098 |
| CNAG_075 | - | 6-phosphc | 72679.45768 | 73413.95664 | 73880.57034 | 49850.60231 | 50824.42833 | 53821.87312 | 3.11E-23 | -0.52491 | 2.52E-24 | -10.1765 | 0.695002 | 22.50673 |

| | | | | | | | | | | | | | | |
|---|---|---|---|---|---|---|---|---|---|---|---|---|---|---|
| CNAG_05 | - | hypothetic | 16.06577054 | 52.07850601 | 45.11684739 | 225.3762518 | 269.4525933 | 235.7787336 | 6.46E-23 | 2.676593 | 5.34E-24 | 10.10341 | 6.393443 | 22.18956 |
| CNAG_05 | MMS2 | ubiquitin- | 2826.802173 | 2874.422895 | 2894.205433 | 1883.019863 | 1831.778385 | 1835.336052 | 7.32E-23 | -0.64628 | 6.06E-24 | -10.091 | 0.638925 | 22.13575 |
| CNAG_05 | - | hypothetic | 67.34764944 | 78.98729404 | 79.20442846 | 282.2663087 | 287.216669 | 250.4551818 | 8.51E-23 | 1.855322 | 7.10E-24 | 10.07541 | 3.618325 | 22.07017 |
| CNAG_03 | - | biotin trar | 141.9898263 | 164.046418 | 137.8537262 | 411.1771234 | 393.8700464 | 425.738775 | 9.64E-23 | 1.460814 | 8.08E-24 | 10.06267 | 2.752637 | 22.0158 |
| CNAG_05 | - | U3 small r | 6832.088278 | 6173.587869 | 7044.792986 | 3996.727719 | 4269.384831 | 4305.432139 | 1.13E-22 | -0.68939 | 9.54E-24 | -10.0463 | 0.620117 | 21.94794 |
| CNAG_05 | - | hypothetic | 2264.988056 | 2170.861113 | 2155.166591 | 1371.625976 | 1361.667334 | 1371.297132 | 3.04E-22 | -0.69876 | 2.64E-23 | -9.94555 | 0.6161 | 21.51752 |
| CNAG_06 | - | pre-60S fa | 3700.278628 | 3631.891224 | 3809.21681 | 2443.375116 | 2452.844633 | 2465.886947 | 3.20E-22 | -0.61335 | 2.78E-23 | -9.94023 | 0.653679 | 21.49552 |
| CNAG_07 | DST1 | transcripti | 2232.354347 | 2140.720201 | 2365.188161 | 1381.733458 | 1373.738248 | 1353.554683 | 3.27E-22 | -0.72956 | 2.85E-23 | -9.93776 | 0.603088 | 21.48535 |
| CNAG_03 | - | large subu | 10881.86363 | 10098.65417 | 10122.67059 | 6927.88912 | 7048.586927 | 7147.415149 | 3.84E-22 | -0.57377 | 3.37E-23 | -9.92125 | 0.67186 | 21.41529 |
| CNAG_03 | - | cytoplasm | 7.789091275 | 16.52267266 | 14.17825594 | 151.5336163 | 119.7766206 | 122.6011246 | 9.26E-22 | 3.39131 | 8.32E-23 | 9.830563 | 10.49267 | 21.03318 |
| CNAG_03 | - | cytoplasm | 3262.78359 | 2469.306485 | 2379.1437 | 1263.169324 | 1344.298399 | 1409.112342 | 1.58E-21 | -1.0303 | 1.44E-22 | -9.77493 | 0.489609 | 20.80038 |
| CNAG_03 | CHI2 | Chitinase | 932.2692599 | 934.3188931 | 824.577466 | 1582.089236 | 1629.807804 | 1541.751888 | 1.68E-21 | 0.805757 | 1.53E-22 | 9.768656 | 1.748063 | 20.77522 |
| CNAG_00 | RRP44 | exosome | 3500.505917 | 3420.679608 | 3295.227592 | 2090.718294 | 2282.081658 | 2222.767848 | 1.70E-21 | -0.64682 | 1.56E-22 | -9.76721 | 0.638686 | 20.76958 |
| CNAG_03 | TUF1 | translatior | 15538.82135 | 13478.6057 | 15696.93708 | 9602.073134 | 9352.43147 | 9679.722215 | 3.81E-21 | -0.65896 | 3.59E-22 | -9.68224 | 0.633335 | 20.41956 |
| CNAG_03 | - | hypothetic | 5047.901344 | 5020.705765 | 5196.687694 | 3425.591825 | 3597.519066 | 3409.527777 | 7.57E-21 | -0.56456 | 7.25E-22 | -9.61013 | 0.67616 | 20.12098 |
| CNAG_03 | - | hypothetic | 1764.104116 | 1765.740255 | 1868.229249 | 1010.369166 | 1121.03142 | 1111.516287 | 1.29E-20 | -0.75063 | 1.25E-21 | -9.55381 | 0.594342 | 19.88856 |
| CNAG_07 | - | DNA-direc | 1051.593187 | 998.5797623 | 1210.062235 | 428.7122735 | 463.5437182 | 588.6533059 | 1.31E-20 | -1.15523 | 1.27E-21 | -9.55255 | 0.448995 | 19.88382 |
| CNAG_06 | - | hypothetic | 213.1360947 | 232.9725336 | 208.2052802 | 495.7915636 | 516.3878707 | 502.4865001 | 1.36E-20 | 1.197325 | 1.32E-21 | 9.548438 | 2.29314 | 19.86712 |
| CNAG_07 | - | hypothetic | 373.0642246 | 410.1068285 | 395.3530672 | 130.5769165 | 140.0397341 | 169.1029537 | 1.57E-20 | -1.43876 | 1.52E-21 | -9.53332 | 0.368884 | 19.80488 |
| CNAG_03 | - | pescadillo | 6816.920987 | 6158.509479 | 6781.41673 | 4159.44357 | 4439.344676 | 4390.901835 | 1.94E-20 | -0.62086 | 1.89E-21 | -9.51064 | 0.650283 | 19.71173 |
| CNAG_07 | - | protein m | 1108.958552 | 1105.261523 | 1235.75817 | 653.9823583 | 591.2879527 | 637.5275498 | 2.00E-20 | -0.88954 | 1.95E-21 | -9.50745 | 0.539787 | 19.69894 |
| CNAG_05 | - | small subu | 1330.985226 | 1571.686987 | 1480.047518 | 857.9124037 | 844.0971441 | 810.7253975 | 2.31E-20 | -0.8176 | 2.27E-21 | -9.49194 | 0.567384 | 19.63591 |
| CNAG_02 | - | T-complex | 10507.01643 | 9595.508603 | 9915.767379 | 6673.339659 | 7066.739659 | 6696.714802 | 2.48E-20 | -0.5705 | 2.43E-21 | -9.48454 | 0.673383 | 19.60615 |
| CNAG_03 | - | ribonucleo | 9746.778762 | 8585.974507 | 9471.020056 | 6253.145873 | 6011.753061 | 6092.791593 | 2.56E-20 | -0.6148 | 2.52E-21 | -9.48088 | 0.653022 | 19.59198 |
| CNAG_03 | - | hypothetic | 1737.133138 | 1677.375911 | 1684.832515 | 904.4651738 | 1075.340027 | 1007.184718 | 4.94E-20 | -0.78724 | 4.97E-21 | -9.40992 | 0.579452 | 19.30622 |
| CNAG_05 | LIV12 | eukaryotic | 13244.26778 | 12028.37253 | 13347.57423 | 8471.215761 | 8827.197612 | 8861.504227 | 6.23E-20 | -0.5778 | 6.28E-21 | -9.38519 | 0.669985 | 19.20575 |
| CNAG_05 | - | NADH del | 4475.649063 | 4209.427245 | 4830.829333 | 2988.208379 | 2837.51099 | 2864.460552 | 7.11E-20 | -0.65283 | 7.18E-21 | -9.37103 | 0.636033 | 19.14796 |
| CNAG_03 | - | Sugar trar | 3701.323199 | 3430.412306 | 3439.196305 | 2293.932477 | 2392.49715 | 2236.115967 | 8.31E-20 | -0.62657 | 8.42E-21 | -9.35431 | 0.647714 | 19.08027 |
| CNAG_03 | - | hypothetic | 28.81737384 | 36.97307454 | 23.78857699 | 169.549039 | 189.1945309 | 138.39025 | 9.40E-20 | 2.4965 | 9.57E-21 | 9.340752 | 5.643148 | 19.02672 |
| CNAG_04 | - | hypothetic | 7.775490981 | 11.13517494 | 17.3617971 | 117.3755033 | 105.0141955 | 137.7769274 | 1.01E-19 | 3.343972 | 1.03E-20 | 9.332623 | 10.15397 | 18.99547 |
| CNAG_03 | - | hormone- | 366.0809082 | 417.1768902 | 404.3786053 | 786.030984 | 761.17403 | 792.2779 | 1.13E-19 | 0.963815 | 1.16E-20 | 9.320242 | 1.950461 | 18.94683 |
| CNAG_05 | - | minichron | 4556.26821 | 4612.337562 | 4633.606772 | 3079.25375 | 3283.564118 | 2922.20332 | 1.17E-19 | -0.58733 | 1.20E-20 | -9.31644 | 0.665572 | 18.93177 |
| CNAG_03 | - | protein LT | 2840.830143 | 2864.741187 | 3200.197184 | 1650.880272 | 1858.52004 | 1903.036999 | 1.18E-19 | -0.73394 | 1.21E-20 | -9.31565 | 0.60126 | 18.92907 |
| CNAG_07 | - | hypothetic | 21.76229714 | 15.44806565 | 13.11340389 | 121.0842809 | 145.2239093 | 118.2255332 | 1.39E-19 | 2.954214 | 1.43E-20 | 9.297995 | 7.750093 | 18.85793 |
| CNAG_05 | - | U6 snRNA | 1673.04927 | 1709.707048 | 1720.031151 | 1064.200759 | 1094.300637 | 1048.251533 | 2.02E-19 | -0.68552 | 2.10E-20 | -9.25728 | 0.621782 | 18.69509 |
| CNAG_03 | - | U3 small r | 6866.112182 | 6964.384794 | 6850.791096 | 4705.878166 | 4920.682393 | 5008.104184 | 2.55E-19 | -0.51418 | 2.67E-20 | -9.23144 | 0.700191 | 18.59422 |
| CNAG_05 | - | glycine-tR | 18471.21468 | 18318.38052 | 18945.40859 | 11855.16276 | 13344.47121 | 13258.82778 | 3.32E-19 | -0.55071 | 3.51E-20 | -9.20204 | 0.682685 | 18.47822 |
| CNAG_03 | - | hypothetic | 1702.264908 | 1835.721934 | 1720.054739 | 1132.743911 | 1089.10007 | 1063.803418 | 3.61E-19 | -0.69369 | 3.84E-20 | -9.19258 | 0.618271 | 18.44197 |
| CNAG_03 | TSR2 | pre-rRNA- | 2681.869384 | 2340.103058 | 3006.040457 | 1559.136108 | 1563.480107 | 1483.486335 | 5.73E-19 | -0.81781 | 6.15E-20 | -9.14174 | 0.567304 | 18.24171 |
| CNAG_05 | - | D-asparta | 143.2239351 | 194.2009577 | 193.2554274 | 493.1113183 | 442.1615855 | 429.2549635 | 6.68E-19 | 1.349396 | 7.18E-20 | 9.124835 | 2.548054 | 18.17534 |
| CNAG_03 | - | hypothetic | 273.7995222 | 279.2992036 | 272.1639892 | 585.2918575 | 551.5571912 | 606.8060927 | 7.79E-19 | 1.065018 | 8.39E-20 | 9.108066 | 2.092196 | 18.10866 |
| CNAG_03 | - | ATP-depe | 3340.475471 | 3004.849274 | 3311.162011 | 1905.45045 | 1975.798953 | 2162.785567 | 9.98E-19 | -0.69193 | 1.08E-19 | -9.08032 | 0.619024 | 18.00073 |
| CNAG_07 | - | nuclear po | 6973.505879 | 7025.786467 | 7009.670899 | 4389.790443 | 4707.587594 | 5082.468663 | 1.60E-18 | -0.58236 | 1.75E-19 | -9.02779 | 0.66787 | 17.79611 |
| CNAG_05 | - | nuclear tra | 5790.597714 | 5632.710808 | 5892.027562 | 4047.489296 | 4127.991065 | 4076.703139 | 1.82E-18 | -0.5146 | 2.00E-19 | -9.01329 | 0.699988 | 17.73907 |
| CNAG_02 | - | 3-demeth | 1599.245753 | 1376.844677 | 1533.371248 | 855.5215302 | 888.9635089 | 899.4916772 | 2.62E-18 | -0.78678 | 2.90E-19 | -8.97251 | 0.579636 | 17.58174 |
| CNAG_03 | - | cell wall s | 90.58891979 | 89.74530967 | 72.82541191 | 237.2437989 | 274.9792324 | 313.3891711 | 2.69E-18 | 1.699998 | 2.98E-19 | 8.969282 | 3.249004 | 17.57046 |
| CNAG_00 | - | ATP-depe | 2137.730554 | 2090.046108 | 1982.432238 | 1307.803398 | 1336.679046 | 1363.494837 | 3.26E-18 | -0.64745 | 3.63E-19 | -8.94773 | 0.638408 | 17.48653 |
| CNAG_05 | - | hypothetic | 1564.563197 | 1842.141123 | 1798.898976 | 1099.291113 | 986.4012917 | 930.6482335 | 3.59E-18 | -0.80209 | 4.00E-19 | -8.93684 | 0.573516 | 17.44515 |
| CNAG_03 | - | hypothetic | 1282.536895 | 1227.378344 | 1029.353467 | 2174.527474 | 2016.998402 | 1995.819815 | 3.72E-18 | 0.790648 | 4.17E-19 | 8.932313 | 1.729851 | 17.42928 |
| CNAG_03 | - | arginine-t | 7718.448855 | 7159.429747 | 7427.72469 | 4723.692386 | 5243.284745 | 5199.070774 | 4.28E-18 | -0.57222 | 4.83E-19 | -8.91619 | 0.672583 | 17.36839 |
| CNAG_03 | - | malate de | 8601.207868 | 7880.217939 | 7959.894431 | 5766.794754 | 5011.131562 | 5405.595373 | 4.66E-18 | -0.61033 | 5.26E-19 | -8.90664 | 0.655048 | 17.33145 |
| CNAG_02 | - | nonsense- | 6045.063022 | 5392.476745 | 6458.227236 | 3750.07034 | 3772.531304 | 3964.594816 | 5.68E-18 | -0.65555 | 6.44E-19 | -8.88419 | 0.634832 | 17.24552 |
| CNAG_07 | - | valine-tRN | 10656.54441 | 10048.02418 | 10107.74545 | 6888.957578 | 7324.641433 | 7463.778169 | 6.02E-18 | -0.52292 | 6.85E-19 | -8.87737 | 0.695964 | 17.22028 |
| CNAG_05 | - | hypothetic | 1484.983109 | 1513.589491 | 1487.543378 | 958.4198027 | 919.1808991 | 811.8605581 | 6.07E-18 | -0.75352 | 6.91E-19 | -8.87637 | 0.593153 | 17.21686 |
| CNAG_03 | - | zinc finger | 1531.696366 | 1529.773153 | 1539.797775 | 1004.361895 | 881.1615758 | 926.1404119 | 6.19E-18 | -0.72539 | 7.06E-19 | -8.87395 | 0.604832 | 17.20834 |
| CNAG_12 | - | unspecifie | 21.80118933 | 32.64599417 | 18.44914462 | 153.4334712 | 134.4313619 | 132.6586016 | 7.56E-18 | 2.547164 | 8.63E-19 | 8.851506 | 5.84484 | 17.12134 |
| CNAG_03 | - | 2-oxoglut | 10021.11107 | 8452.377306 | 9039.136912 | 5954.999121 | 6110.974419 | 6091.673013 | 7.67E-18 | -0.61575 | 8.77E-19 | -8.84979 | 0.65259 | 17.11511 |
| CNAG_05 | - | hypothetic | 1463.999226 | 1416.688188 | 1617.576299 | 897.6093327 | 931.2669059 | 865.1081556 | 9.51E-18 | -0.75505 | 1.10E-18 | -8.82456 | 0.592527 | 17.02169 |
| CNAG_05 | SPR1 | endogluca | 125.5161516 | 104.826481 | 98.39777611 | 313.1594215 | 299.0103085 | 308.1261895 | 9.57E-18 | 1.476731 | 1.11E-18 | 8.823752 | 2.783174 | 17.01902 |
| CNAG_03 | - | phosphori | 1630.902077 | 1596.5699 | 1538.764687 | 979.3560385 | 926.0237421 | 1034.881244 | 9.85E-18 | -0.71199 | 1.14E-18 | -8.82026 | 0.610479 | 17.00638 |
| CNAG_04 | PRO2 | glutamate | 3531.022877 | 3588.792233 | 3648.207425 | 2484.674193 | 2566.763949 | 2431.472194 | 1.04E-17 | -0.5406 | 1.21E-18 | -8.81356 | 0.687485 | 16.98224 |
| CNAG_05 | - | GTP-bindi | 2033.790813 | 1896.150043 | 1967.463444 | 1296.200562 | 1220.198723 | 1263.597276 | 1.25E-17 | -0.65722 | 1.47E-18 | -8.79226 | 0.6341 | 16.90207 |
| CNAG_02 | - | nuclear G | 2268.404008 | 1930.66959 | 2225.492541 | 1255.584551 | 1362.565466 | 1271.411393 | 1.32E-17 | -0.74036 | 1.55E-18 | -8.78628 | 0.598588 | 16.8803 |
| CNAG_03 | - | hypothetic | 54.56138495 | 85.43460392 | 94.09891744 | 293.3018913 | 281.9964028 | 237.2170832 | 2.44E-17 | 1.784937 | 2.91E-18 | 8.714987 | 3.446034 | 16.61308 |
| CNAG_06 | ERG1 | Squalene | 2053.497617 | 1833.640588 | 1833.105374 | 1086.657748 | 1124.428665 | 1248.022629 | 3.21E-17 | -0.7414 | 3.87E-18 | -8.68261 | 0.59816 | 16.49319 |
| CNAG_02 | APH3 | Phytase | 200.2903477 | 199.6057082 | 229.4833237 | 499.1212784 | 468.8720978 | 444.8536067 | 3.71E-17 | 1.152478 | 4.49E-18 | 8.665736 | 2.222953 | 16.43054 |
| CNAG_03 | - | AFG1 fam | 2104.951143 | 1873.537558 | 1999.445444 | 1180.089548 | 1228.828454 | 1315.744734 | 5.03E-17 | -0.69841 | 6.13E-18 | -8.6301 | 0.616251 | 16.29845 |
| CNAG_03 | - | hypothetic | 1110.340652 | 1334.667246 | 1391.44764 | 699.0671258 | 762.198686 | 634.2674703 | 6.00E-17 | -0.88716 | 7.35E-18 | -8.60929 | 0.540677 | 16.22215 |

| | | | | | | | | | | | | | |
|---|---|---|---|---|---|---|---|---|---|---|---|---|---|
| CNAG_033 | - | Phosphog | 15270.98766 | 14394.66597 | 15705.76015 | 21432.12487 | 21126.03597 | 22817.99088 | 6.45E-17 | 0.511533 | 7.92E-18 | 8.600733 | 1.425564 | 16.19016 |
| CNAG_067 | - | cysteine-t | 3383.870348 | 3358.236835 | 3533.015059 | 2343.011728 | 2451.185495 | 2284.940219 | 6.82E-17 | -0.55303 | 8.40E-18 | -8.59397 | 0.681588 | 16.1663 |
| CNAG_057 | - | signal rec | 1340.062327 | 1270.109727 | 1268.923173 | 801.1702167 | 790.6654678 | 666.4534097 | 7.66E-17 | -0.79653 | 9.48E-18 | -8.58007 | 0.575733 | 16.11592 |
| CNAG_033 | - | DNA supe | 290.1958452 | 300.8644011 | 344.6389614 | 671.7642941 | 585.9677604 | 684.5035306 | 7.72E-17 | 1.039518 | 9.57E-18 | 8.578949 | 2.055541 | 16.11212 |
| CNAG_009 | - | carboxype | 1635.435791 | 1583.590806 | 1353.258779 | 878.1026198 | 922.6205575 | 911.7069526 | 8.94E-17 | -0.7689 | 1.11E-17 | -8.56182 | 0.586863 | 16.0488 |
| CNAG_036 | - | hypothetic | 1027.181433 | 1061.03354 | 972.4774276 | 626.0366643 | 581.8618389 | 591.9976067 | 9.68E-17 | -0.78212 | 1.21E-17 | -8.55208 | 0.581512 | 16.01427 |
| CNAG_058 | - | allantoate | 40.41358663 | 34.82814852 | 37.61806971 | 172.6900064 | 169.0570009 | 139.5046847 | 1.54E-16 | 2.09666 | 1.94E-17 | 8.497263 | 4.277182 | 15.81323 |
| CNAG_050 | - | sulfite red | 4782.452242 | 4007.928698 | 4054.543483 | 2546.778896 | 2891.078401 | 2434.894902 | 1.72E-16 | -0.72264 | 2.18E-17 | -8.48393 | 0.605986 | 15.76551 |
| CNAG_049 | - | hypothetic | 3334.793381 | 3234.33929 | 3426.370415 | 2276.080315 | 2198.374029 | 2373.696865 | 2.27E-16 | -0.56092 | 2.90E-17 | -8.45037 | 0.677869 | 15.64334 |
| CNAG_057 | MFE2 | multifunct | 1796.343503 | 1846.860695 | 1732.072249 | 2619.284339 | 2906.360829 | 2660.867368 | 2.31E-16 | 0.591765 | 2.95E-17 | 8.448479 | 1.507089 | 15.63671 |
| CNAG_032 | ZWF1 | glucose-6 | 14130.63142 | 13650.9462 | 12934.94971 | 9777.285827 | 9848.497558 | 10062.6607 | 2.97E-16 | -0.471 | 3.83E-17 | -8.41813 | 0.721467 | 15.52692 |
| CNAG_057 | ATP1 | ATP synth | 55366.40877 | 54237.86314 | 57593.6937 | 40234.88557 | 41741.71747 | 42441.6011 | 3.29E-16 | -0.44179 | 4.25E-17 | -8.40588 | 0.736223 | 15.4832 |
| CNAG_078 | BOT1 | Eukaryotic | 6337.156912 | 6275.944674 | 6575.628913 | 4706.761912 | 4650.706921 | 4642.872998 | 4.03E-16 | -0.47026 | 5.23E-17 | -8.38142 | 0.721836 | 15.39484 |
| CNAG_059 | - | hypothetic | 483.9241606 | 496.9034257 | 523.7842279 | 858.9655367 | 919.933091 | 877.8528058 | 4.08E-16 | 0.805999 | 5.30E-17 | 8.379811 | 1.748356 | 15.38933 |
| CNAG_055 | - | aldose rec | 1659.65961 | 1526.899437 | 1324.745715 | 2591.591301 | 2478.466155 | 2392.184023 | 5.22E-16 | 0.711128 | 6.80E-17 | 8.350388 | 1.637084 | 15.28218 |
| CNAG_036 | - | pumilio do | 5301.1303 | 4488.5271 | 5333.143811 | 3128.191917 | 3343.882021 | 3324.041016 | 5.33E-16 | -0.64269 | 6.95E-17 | -8.34792 | 0.640516 | 15.2735 |
| CNAG_056 | - | tyrosine a | 174.5708283 | 180.1859972 | 162.3502084 | 364.0105291 | 399.8793733 | 445.6926672 | 5.36E-16 | 1.212247 | 7.01E-17 | 8.346912 | 2.316982 | 15.27062 |
| CNAG_033 | STU2 | microtubu | 2591.497911 | 2378.061195 | 2406.057343 | 3565.711308 | 3720.524228 | 3627.802298 | 5.61E-16 | 0.549572 | 7.36E-17 | 8.341154 | 1.463651 | 15.25067 |
| CNAG_025 | - | hypothetic | 2702.487618 | 2754.048867 | 2489.263271 | 3728.272994 | 4149.264034 | 4049.625749 | 6.23E-16 | 0.570867 | 8.18E-17 | 8.328544 | 1.485416 | 15.2056 |
| CNAG_009 | - | nucleolar | 5768.151293 | 4999.17929 | 5026.098547 | 3269.05307 | 3629.422412 | 3512.764479 | 6.23E-16 | -0.61725 | 8.20E-17 | -8.32837 | 0.651912 | 15.20536 |
| CNAG_031 | - | DNA cros | 438.4914961 | 547.5062697 | 519.5319186 | 890.132884 | 953.5977849 | 910.0287018 | 6.63E-16 | 0.857639 | 8.75E-17 | 8.32062 | 1.812071 | 15.17816 |
| CNAG_036 | FRR1 | FK506-bin | 9434.048791 | 9429.587011 | 9567.054629 | 7114.806706 | 7202.19829 | 6997.519974 | 7.29E-16 | -0.431 | 9.67E-17 | -8.30873 | 0.741749 | 15.13741 |
| CNAG_008 | - | hypothetic | 8255.827225 | 8445.888901 | 9107.360503 | 6232.969372 | 6222.292073 | 6177.121221 | 7.37E-16 | -0.48533 | 9.80E-17 | -8.30723 | 0.714332 | 15.13232 |
| CNAG_056 | - | Nucleolar | 2024.411283 | 1855.208148 | 1880.034477 | 1296.235668 | 1183.963152 | 1233.624959 | 7.77E-16 | -0.64913 | 1.04E-16 | -8.30058 | 0.637664 | 15.10961 |
| CNAG_055 | - | GDSL Lipa | 1682.368096 | 1606.328587 | 1812.760565 | 1039.268093 | 994.1665621 | 1130.340471 | 9.80E-16 | -0.7046 | 1.32E-16 | -8.27192 | 0.613611 | 15.00896 |
| CNAG_031 | - | peptide-m | 2129.656835 | 2041.624972 | 2263.899716 | 1441.755285 | 1363.396233 | 1425.662619 | 1.03E-15 | -0.62063 | 1.38E-16 | -8.26607 | 0.650386 | 14.98847 |
| CNAG_051 | - | nuclear ca | 1051.717137 | 1084.744848 | 990.6160952 | 619.7516178 | 603.4314089 | 656.3333776 | 1.44E-15 | -0.74959 | 1.96E-16 | -8.22462 | 0.594774 | 14.84064 |
| CNAG_037 | - | hypothetic | 43.90171422 | 33.75960616 | 46.12646432 | 173.3768039 | 152.3927578 | 163.7276606 | 1.51E-15 | 1.98414 | 2.05E-16 | 8.219169 | 3.956266 | 14.82129 |
| CNAG_050 | CAT4 | catalase 4 | 52.05869796 | 41.29522331 | 53.58726223 | 174.6293578 | 175.7749352 | 181.4769511 | 1.66E-15 | 1.844157 | 2.26E-16 | 8.207645 | 3.590432 | 14.78115 |
| CNAG_059 | - | Alpha-glu | 608.9052333 | 524.5588673 | 607.6182076 | 270.0892313 | 284.212509 | 314.4585191 | 2.04E-15 | -1.01832 | 2.81E-16 | -8.18119 | 0.49369 | 14.68952 |
| CNAG_036 | - | ATP-bindi | 5272.018756 | 4947.454651 | 4932.250574 | 3305.678042 | 3595.792829 | 3659.255187 | 2.13E-15 | -0.53645 | 2.93E-16 | -8.17598 | 0.689464 | 14.67229 |
| CNAG_037 | HXS1 | high-affini | 409.4661778 | 262.2434025 | 241.3925557 | 1219.800305 | 1355.080538 | 1579.850283 | 2.13E-15 | 2.170151 | 2.93E-16 | 8.176059 | 4.500704 | 14.67229 |
| CNAG_057 | - | elongator | 967.2364326 | 903.0804629 | 923.703602 | 1413.717594 | 1565.940006 | 1525.086592 | 2.20E-15 | 0.67351 | 3.04E-16 | 8.171546 | 1.594949 | 14.65708 |
| CNAG_031 | - | hypothetic | 1245.300563 | 1509.599831 | 1381.201427 | 2147.759258 | 2179.137558 | 2180.108661 | 2.22E-15 | 0.639428 | 3.08E-16 | 8.170274 | 1.557712 | 14.65289 |
| CNAG_057 | - | hypothetic | 675.6379507 | 633.3684734 | 730.2527552 | 391.7411131 | 348.921629 | 337.8402307 | 2.52E-15 | -0.9371 | 3.50E-16 | -8.15481 | 0.522283 | 14.59879 |
| CNAG_026 | - | Cystathior | 3984.257315 | 4770.64707 | 4478.967681 | 2929.803173 | 3086.01168 | 2977.667221 | 3.34E-15 | -0.57184 | 4.66E-16 | -8.12003 | 0.672757 | 14.47674 |
| CNAG_036 | - | hypothetic | 1699.824262 | 1608.465912 | 1605.959304 | 1068.133101 | 1034.758303 | 1105.920585 | 4.18E-15 | -0.63044 | 5.87E-16 | -8.092 | 0.645977 | 14.37849 |
| CNAG_034 | - | hypothetic | 347.3174043 | 353.6186853 | 364.9043954 | 662.1840468 | 625.6503885 | 712.2947422 | 4.40E-15 | 0.894742 | 6.18E-16 | 8.085712 | 1.859277 | 14.35646 |
| CNAG_032 | UVE1 | UV damag | 1587.447516 | 1983.647828 | 1638.246531 | 2754.080761 | 2753.598145 | 2776.258117 | 4.50E-15 | 0.655286 | 6.32E-16 | 8.08289 | 1.574929 | 14.34678 |
| CNAG_034 | - | T-complex | 8755.54296 | 8451.29103 | 9041.261233 | 6087.360008 | 6660.529808 | 6215.986114 | 4.81E-15 | -0.48466 | 6.77E-16 | -8.07452 | 0.714667 | 14.31774 |
| CNAG_057 | EMC3 | hypothetic | 1621.723058 | 1675.244747 | 1759.465804 | 1043.164085 | 1009.705529 | 1161.402768 | 4.97E-15 | -0.6688 | 7.00E-16 | -8.07057 | 0.629028 | 14.30405 |
| CNAG_026 | - | hypothetic | 3770.270097 | 3431.527001 | 3583.167206 | 2428.602711 | 2414.891837 | 2591.280516 | 6.87E-15 | -0.55246 | 9.81E-16 | -8.02917 | 0.681859 | 14.16275 |
| CNAG_077 | - | tricarboxyl | 43.855062 | 28.37507999 | 37.60771249 | 156.9779514 | 150.8157546 | 142.719406 | 8.66E-15 | 2.036124 | 1.25E-15 | 7.999717 | 4.101421 | 14.0624 |
| CNAG_051 | - | hypothetic | 625.1275735 | 617.550634 | 543.0322358 | 995.1202006 | 1131.346943 | 992.2572479 | 9.41E-15 | 0.789343 | 1.36E-15 | 7.989174 | 1.728288 | 14.02634 |
| CNAG_050 | - | hypothetic | 67.2143546 | 57.43949931 | 62.12911416 | 189.9495966 | 243.4323643 | 183.9279559 | 9.86E-15 | 1.709171 | 1.42E-15 | 7.983389 | 3.26973 | 14.00634 |
| CNAG_008 | - | Glutamate | 8317.183173 | 7431.924793 | 6076.634893 | 4378.107828 | 4460.840061 | 4628.494073 | 1.11E-14 | -0.71206 | 1.61E-15 | -7.9679 | 0.61045 | 13.95407 |
| CNAG_027 | - | hypothetic | 69.68479852 | 82.2187989 | 116.4440779 | 274.2742309 | 291.494029 | 244.98387 | 1.12E-14 | 1.583899 | 1.63E-15 | 7.966771 | 2.997789 | 13.95046 |
| CNAG_033 | - | Ribose-ph | 3413.055055 | 3454.105587 | 3394.412796 | 2242.536022 | 2560.774905 | 2361.518164 | 1.76E-14 | -0.53387 | 2.59E-15 | -7.90912 | 0.690699 | 13.75444 |
| CNAG_034 | - | ribonuclea | 785.4709806 | 766.9587822 | 849.7225183 | 446.9280332 | 398.0491872 | 488.6987766 | 1.81E-14 | -0.86464 | 2.67E-15 | -7.90531 | 0.549182 | 13.74223 |
| CNAG_051 | - | L-lactate c | 735.1509018 | 774.3701193 | 640.8430349 | 413.5143416 | 390.3594391 | 377.7798851 | 2.15E-14 | -0.87972 | 3.19E-15 | -7.8834 | 0.543471 | 13.66846 |
| CNAG_057 | - | acetyl-CoA | 42781.29801 | 38009.56048 | 36806.44366 | 26661.43935 | 28384.08296 | 28198.26707 | 2.20E-14 | -0.5142 | 3.26E-15 | -7.88034 | 0.700181 | 13.65856 |
| CNAG_033 | NPY1 | NAD diph | 1384.688038 | 1433.906543 | 1497.129658 | 970.1255081 | 938.183625 | 900.5980166 | 2.29E-14 | -0.63469 | 3.41E-15 | -7.87488 | 0.644079 | 13.6403 |
| CNAG_068 | - | Histidinol | 1474.35067 | 1418.778671 | 1286.052571 | 819.7427338 | 940.8298257 | 806.2736412 | 2.89E-14 | -0.71936 | 4.32E-15 | -7.84525 | 0.607366 | 13.53968 |
| CNAG_075 | - | hypothetic | 206.0380436 | 209.2530844 | 164.5030736 | 485.2984438 | 404.0534026 | 411.5438032 | 2.99E-14 | 1.152013 | 4.49E-15 | 7.840509 | 2.222238 | 13.52363 |
| CNAG_026 | - | cytoplasm | 5351.516901 | 5100.475021 | 5414.25878 | 3604.586382 | 3822.649868 | 3951.210739 | 3.44E-14 | -0.49522 | 5.17E-15 | -7.8227 | 0.709451 | 13.46352 |
| CNAG_074 | - | hypothetic | 49.77026738 | 65.99709146 | 46.15639659 | 213.2913723 | 174.7936445 | 168.3579303 | 3.48E-14 | 1.780261 | 5.23E-15 | 7.821194 | 3.434884 | 13.45867 |
| CNAG_052 | - | Stomatin f | 1008.029351 | 737.2358358 | 780.8575939 | 1434.049244 | 1605.706953 | 1579.400114 | 3.75E-14 | 0.85443 | 5.66E-15 | 7.811237 | 1.808045 | 13.4261 |
| CNAG_050 | SSK2 | ste/ste11/ | 3506.494329 | 3550.005668 | 3444.553386 | 2439.519298 | 2514.136073 | 2617.908081 | 4.71E-14 | -0.48715 | 7.14E-15 | -7.78194 | 0.713434 | 13.32673 |
| CNAG_067 | - | ATP-deper | 2883.938963 | 2546.978525 | 3066.901895 | 1669.616027 | 1765.343834 | 1952.947272 | 4.96E-14 | -0.67325 | 7.55E-15 | -7.77496 | 0.627091 | 13.30416 |
| CNAG_033 | - | hypothetic | 1432.360782 | 1372.474969 | 1330.810298 | 883.6855667 | 919.2495105 | 756.3530395 | 5.22E-14 | -0.70812 | 7.95E-15 | -7.76839 | 0.612116 | 13.28267 |
| CNAG_078 | - | mitochonc | 1264.214655 | 1239.931813 | 1155.928391 | 758.3323944 | 819.1987678 | 695.2699511 | 7.62E-14 | -0.70254 | 1.18E-14 | -7.71868 | 0.614491 | 13.11798 |
| CNAG_058 | KAP123 | importin b | 2294.113363 | 2029.765463 | 2051.747074 | 1432.428694 | 1441.958738 | 1264.767701 | 9.30E-14 | -0.63951 | 1.45E-14 | -7.69226 | 0.641931 | 13.03156 |
| CNAG_008 | - | rab family | 924.324978 | 826.2564027 | 876.4602936 | 524.0379341 | 499.052471 | 543.1050582 | 1.24E-13 | -0.76138 | 1.94E-14 | -7.65486 | 0.589931 | 12.90792 |
| CNAG_033 | - | ribosomal | 2823.167124 | 2414.463239 | 2826.98536 | 1679.804949 | 1879.308463 | 1675.494919 | 1.32E-13 | -0.63991 | 2.07E-14 | -7.64651 | 0.641751 | 12.88039 |
| CNAG_037 | - | mRNA tur | 1657.823511 | 1504.009631 | 1778.628637 | 1111.015363 | 989.8717761 | 1020.494869 | 1.74E-13 | -0.6784 | 2.75E-14 | -7.60936 | 0.624858 | 12.76 |
| CNAG_075 | ELP3 | elongator | 1598.246375 | 1480.278471 | 1534.499847 | 1012.097152 | 1053.805966 | 1014.912521 | 2.02E-13 | -0.59855 | 3.21E-14 | -7.5894 | 0.660418 | 12.69533 |
| CNAG_033 | - | nicotinam | 995.6182197 | 898.460111 | 977.7657516 | 590.997588 | 580.1751244 | 596.3991087 | 2.14E-13 | -0.71623 | 3.42E-14 | -7.58139 | 0.608688 | 12.66983 |
| CNAG_034 | - | cytoplasm | 2360.928256 | 2350.838439 | 2434.582285 | 1621.518256 | 1632.563018 | 1755.315277 | 3.25E-13 | -0.52772 | 5.24E-14 | -7.52583 | 0.69365 | 12.48746 |

| | | | | | | | | | | | | | | |
|---|---|---|---|---|---|---|---|---|---|---|---|---|---|---|
| CNAG_032 | - | lactamase | 1028.38239 | 1042.752399 | 968.2283465 | 657.9982626 | 658.7280261 | 584.2466009 | 5.44E-13 | -0.69311 | 8.86E-14 | -7.45688 | 0.618517 | 12.26419 |
| CNAG_033 | - | cytochrom | 1591.301003 | 1526.601827 | 1608.066043 | 1099.361586 | 989.028864 | 1067.069413 | 6.30E-13 | -0.59801 | 1.03E-13 | -7.43708 | 0.660664 | 12.20067 |
| CNAG_078 | - | alanine-gl | 2154.050087 | 2023.271873 | 1957.909462 | 1262.618304 | 1492.061457 | 1273.61981 | 6.59E-13 | -0.62284 | 1.08E-13 | -7.43078 | 0.649393 | 12.18092 |
| CNAG_052 | - | 1,2-dihydr | 2093.406063 | 1899.431921 | 2094.366006 | 1336.630724 | 1420.406961 | 1393.453141 | 7.59E-13 | -0.5685 | 1.25E-13 | -7.41162 | 0.674319 | 12.11973 |
| CNAG_057 | TEL1 | serine/thr | 1295.318452 | 1256.433771 | 1167.926914 | 1836.372843 | 1985.993408 | 1790.531601 | 1.07E-12 | 0.578306 | 1.77E-13 | 7.365166 | 1.493095 | 11.96995 |
| CNAG_033 | - | hypothetic | 701.5700646 | 760.4752864 | 815.6088702 | 464.9244731 | 446.4648788 | 444.3410609 | 1.27E-12 | -0.76432 | 2.12E-13 | -7.34117 | 0.588731 | 11.8945 |
| CNAG_057 | - | hypothetic | 42.72604294 | 42.34021055 | 50.36778616 | 140.9032565 | 164.6983152 | 150.4859691 | 1.38E-12 | 1.744952 | 2.30E-13 | 7.329876 | 3.351836 | 11.85917 |
| CNAG_033 | TIM13 | mitochonc | 1605.356259 | 1532.012207 | 1728.537485 | 1119.578576 | 1042.529395 | 1085.951547 | 1.54E-12 | -0.59926 | 2.57E-13 | -7.31525 | 0.660091 | 11.81244 |
| CNAG_009 | - | hypothetic | 674.0801583 | 570.1807765 | 447.1117916 | 1147.331898 | 1052.718898 | 998.9050469 | 2.27E-12 | 0.903171 | 3.80E-13 | 7.262332 | 1.870171 | 11.64475 |
| CNAG_032 | - | hypothetic | 30.02069162 | 65.97867175 | 49.33566043 | 151.6330023 | 208.0723164 | 173.777794 | 2.38E-12 | 1.872001 | 4.01E-13 | 7.255212 | 3.6604 | 11.62284 |
| CNAG_027 | BUD32 | BUD32 pro | 1820.337944 | 1988.770627 | 1931.226017 | 1381.907094 | 1347.947913 | 1282.464564 | 2.92E-12 | -0.53213 | 4.95E-13 | -7.22659 | 0.691532 | 11.53399 |
| CNAG_036 | - | hypothetic | 1382.20543 | 1346.625785 | 1262.591483 | 915.6692275 | 901.1253342 | 821.7904829 | 3.96E-12 | -0.61285 | 6.75E-13 | -7.18437 | 0.653903 | 11.40253 |
| CNAG_057 | - | hypothetic | 1632.357095 | 1882.064765 | 1795.768289 | 1273.7485 | 1127.912724 | 1159.241283 | 4.84E-12 | -0.59155 | 8.29E-13 | -7.1563 | 0.66363 | 11.31505 |
| CNAG_037 | RINT1 | hypothetic | 2454.180705 | 2266.795316 | 2212.804617 | 1580.292394 | 1476.381158 | 1687.594713 | 5.44E-12 | -0.56281 | 9.35E-13 | -7.13974 | 0.67698 | 11.26452 |
| CNAG_008 | - | adenylate | 1117.224054 | 1258.173264 | 1054.643104 | 688.2271999 | 730.2967518 | 760.6623639 | 5.67E-12 | -0.66934 | 9.76E-13 | -7.13378 | 0.628794 | 11.24663 |
| CNAG_033 | CHS2 | Chitin syn | 234.0939078 | 289.983309 | 209.2746348 | 491.829792 | 457.4771106 | 519.1099796 | 5.88E-12 | 0.988926 | 1.02E-12 | 7.128437 | 1.984707 | 11.23038 |
| CNAG_050 | - | translatior | 1841.381368 | 1986.640285 | 2083.684797 | 1304.721347 | 1353.102273 | 1421.16974 | 6.09E-12 | -0.55028 | 1.05E-12 | -7.12369 | 0.682885 | 11.21571 |
| CNAG_037 | - | hypothetic | 17.1900956 | 42.3428624 | 54.61911531 | 144.8433626 | 179.5347055 | 159.2936693 | 7.85E-12 | 2.078744 | 1.36E-12 | 7.087588 | 4.224392 | 11.10494 |
| CNAG_034 | - | alginate ly | 653.2258733 | 780.1743463 | 650.7174406 | 1057.249838 | 1134.593487 | 1200.849588 | 9.48E-12 | 0.688683 | 1.66E-12 | 7.06059 | 1.611812 | 11.02314 |
| CNAG_009 | - | DNA-bind | 2522.000817 | 2474.721849 | 2306.673307 | 1659.64547 | 1799.972687 | 1747.573198 | 9.71E-12 | -0.50369 | 1.70E-12 | -7.05684 | 0.705298 | 11.01262 |
| CNAG_055 | - | hypothetic | 1117.368297 | 1243.178859 | 1276.368844 | 819.9125371 | 781.1816265 | 798.4280616 | 9.92E-12 | -0.61538 | 1.74E-12 | -7.05365 | 0.652756 | 11.00356 |
| CNAG_049 | - | U6 snRNA | 1598.391873 | 1631.093225 | 1620.894536 | 1126.56611 | 1184.965952 | 1085.964552 | 1.34E-11 | -0.52955 | 2.38E-12 | -7.01017 | 0.69277 | 10.87151 |
| CNAG_034 | - | small subu | 1714.986973 | 1500.786493 | 1685.903053 | 1081.38235 | 999.3609836 | 1146.96658 | 1.51E-11 | -0.61843 | 2.68E-12 | -6.99377 | 0.651378 | 10.82127 |
| CNAG_048 | - | derlin-2/3 | 270.1247082 | 238.3426008 | 227.3682928 | 464.5176377 | 447.1447114 | 481.431197 | 1.64E-11 | 0.905876 | 2.92E-12 | 6.981307 | 1.873682 | 10.78418 |
| CNAG_037 | - | hypothetic | 5.30367251 | 2.442265342 | 6.653322381 | 70.06180052 | 81.53499279 | 39.8781144 | 1.86E-11 | 3.852108 | 3.32E-12 | 6.963384 | 14.44109 | 10.73092 |
| CNAG_052 | - | signal rec | 3462.134023 | 3291.485465 | 3579.957256 | 2522.083871 | 2559.01448 | 2555.755387 | 1.94E-11 | -0.45178 | 3.47E-12 | -6.95729 | 0.731139 | 10.71301 |
| CNAG_055 | - | solute car | 851.6657772 | 899.7988011 | 879.9572808 | 1336.773074 | 1427.935322 | 1238.817195 | 2.19E-11 | 0.590624 | 3.93E-12 | 6.939685 | 1.505898 | 10.66031 |
| CNAG_036 | LDP1 | hypothetic | 745.3107766 | 698.3440463 | 678.4213399 | 1085.379209 | 1079.328554 | 1108.801663 | 2.31E-11 | 0.610045 | 4.16E-12 | 6.931568 | 1.526306 | 10.63615 |
| CNAG_055 | - | U3 small r | 5129.545112 | 4707.204418 | 4637.933161 | 3253.556176 | 3521.62361 | 3676.985372 | 2.37E-11 | -0.48542 | 4.28E-12 | -6.92759 | 0.714291 | 10.62463 |
| CNAG_077 | - | U3 small r | 2305.778185 | 1975.917996 | 1976.058173 | 1400.482514 | 1465.265046 | 1387.926821 | 2.44E-11 | -0.57303 | 4.41E-12 | -6.92341 | 0.672201 | 10.61211 |
| CNAG_034 | - | hypothetic | 181.4646199 | 185.5229077 | 151.6776567 | 356.4307039 | 354.1032063 | 346.042566 | 2.46E-11 | 1.015375 | 4.44E-12 | 6.922321 | 2.021429 | 10.60906 |
| CNAG_074 | - | mitochonc | 1336.792319 | 1385.412681 | 1324.430009 | 974.099443 | 885.5545083 | 910.5470848 | 2.81E-11 | -0.56177 | 5.10E-12 | -6.90285 | 0.67747 | 10.55177 |
| CNAG_055 | - | hypothetic | 14.73789217 | 25.06703088 | 20.53102566 | 92.07299759 | 88.53570552 | 106.9528795 | 2.87E-11 | 2.263564 | 5.22E-12 | 6.899526 | 4.801764 | 10.5419 |
| CNAG_003 | - | hypothetic | 291.3442376 | 376.172995 | 331.8484962 | 632.7209338 | 638.7777571 | 550.4659 | 2.91E-11 | 0.852366 | 5.30E-12 | 6.897358 | 1.80546 | 10.53585 |
| CNAG_055 | - | methylthic | 1559.787301 | 1437.217777 | 1627.241108 | 1039.361252 | 1076.242476 | 1042.654317 | 2.91E-11 | -0.56601 | 5.30E-12 | -6.89737 | 0.675484 | 10.53585 |
| CNAG_037 | ECM22 | hypothetic | 1489.73613 | 1413.494276 | 1546.200955 | 1064.372755 | 1005.483269 | 962.7539814 | 2.99E-11 | -0.56893 | 5.45E-12 | -6.89345 | 0.674117 | 10.52451 |
| CNAG_025 | SEN54 | tRNA-splic | 1026.243709 | 1177.402477 | 1081.258989 | 752.2354504 | 676.7780776 | 714.0595478 | 3.05E-11 | -0.63105 | 5.56E-12 | -6.89056 | 0.645707 | 10.51598 |
| CNAG_008 | - | multisite-s | 4114.905665 | 3968.099802 | 4221.96786 | 2883.924358 | 3225.842227 | 2927.690451 | 3.05E-11 | -0.46105 | 5.56E-12 | -6.89042 | 0.726456 | 10.51582 |
| CNAG_077 | - | ubiquitin c | 7045.827562 | 6259.81379 | 6427.435597 | 4634.299822 | 4965.578709 | 4932.599919 | 3.19E-11 | -0.45714 | 5.84E-12 | -6.88353 | 0.728431 | 10.49596 |
| CNAG_055 | - | hypothetic | 279.6210643 | 323.4085405 | 324.3540545 | 562.5964044 | 529.0551538 | 552.5546336 | 3.52E-11 | 0.812509 | 6.46E-12 | 6.869201 | 1.756263 | 10.45288 |
| CNAG_033 | ZNF2 | C2H2 type | 82.37154538 | 95.06155989 | 78.1129975 | 181.6746376 | 237.0229915 | 252.4642538 | 3.68E-11 | 1.379023 | 6.75E-12 | 6.86288 | 2.600922 | 10.43422 |
| CNAG_068 | - | hypothetic | 224.6780534 | 221.0975915 | 223.0777317 | 413.8428622 | 473.2954415 | 394.9989982 | 3.90E-11 | 0.922692 | 7.16E-12 | 6.85439 | 1.895649 | 10.409 |
| CNAG_050 | - | L-xylulose | 115.099976 | 148.9038994 | 148.4254778 | 309.062645 | 282.4225925 | 310.413913 | 4.12E-11 | 1.117973 | 7.58E-12 | 6.846312 | 2.170418 | 10.38505 |
| CNAG_052 | SMG1 | glucose-m | 123.5920427 | 42.46626496 | 31.3028438 | 1254.094621 | 1119.136296 | 1020.546081 | 4.34E-11 | 4.093599 | 8.02E-12 | 6.838262 | 17.07245 | 10.36208 |
| CNAG_058 | ALG9 | Alpha-1,2- | 2914.358622 | 2764.579911 | 2644.742525 | 2039.558827 | 2065.636942 | 1984.050121 | 5.19E-11 | -0.46641 | 9.64E-12 | -6.81183 | 0.723762 | 10.28442 |
| CNAG_059 | - | hypothetic | 419.5738597 | 394.5334964 | 345.7466464 | 754.28803 | 693.0254793 | 612.6658327 | 5.20E-11 | 0.814896 | 9.65E-12 | 6.811701 | 1.759172 | 10.28431 |
| CNAG_074 | - | hypothetic | 909.2499051 | 866.1054451 | 859.4387916 | 588.7680701 | 578.5145368 | 519.8448821 | 5.73E-11 | -0.66001 | 1.07E-11 | -6.79709 | 0.632872 | 10.24221 |
| CNAG_050 | - | hypothetic | 387.6734333 | 574.9521364 | 558.511549 | 240.5402354 | 268.7496522 | 247.9099328 | 5.79E-11 | -1.02255 | 1.08E-11 | -6.79543 | 0.492247 | 10.2375 |
| CNAG_050 | - | endopepti | 21.63765156 | 17.55060831 | 15.20790596 | 94.75537933 | 78.11242608 | 90.44505907 | 6.81E-11 | 2.290155 | 1.28E-11 | 6.771068 | 4.891086 | 10.16679 |
| CNAG_052 | - | hypothetic | 756.3041523 | 716.3603186 | 791.1105398 | 478.9756862 | 491.4053518 | 406.6543011 | 8.23E-11 | -0.73456 | 1.56E-11 | -6.74261 | 0.601002 | 10.08436 |
| CNAG_067 | - | endoplasn | 1771.180924 | 1751.777411 | 1695.576941 | 1279.184266 | 1247.03458 | 1225.815265 | 9.60E-11 | -0.4912 | 1.82E-11 | -6.7197 | 0.711432 | 10.01792 |
| CNAG_127 | - | unspecifie | 1448.031731 | 780.9722897 | 875.3129863 | 269.763765 | 251.1739032 | 208.1948443 | 9.60E-11 | -2.10993 | 1.82E-11 | -6.71949 | 0.231658 | 10.01756 |
| CNAG_026 | - | flavonol s | 93.91419217 | 70.3464876 | 84.47671916 | 207.1013627 | 216.2532063 | 199.4641285 | 9.72E-11 | 1.316207 | 1.85E-11 | 6.71758 | 2.490106 | 10.01215 |
| CNAG_058 | - | splicing fa | 5234.835075 | 5252.388708 | 5254.33451 | 3918.335594 | 4192.765179 | 4069.993422 | 1.07E-10 | -0.38534 | 2.04E-11 | -6.70325 | 0.765597 | 9.970884 |
| CNAG_057 | ARG5 | N-acetyl-g | 5742.670179 | 5251.333694 | 5304.469764 | 4177.575836 | 4086.612129 | 4070.015902 | 1.20E-10 | -0.4177 | 2.28E-11 | -6.68661 | 0.748617 | 9.922036 |
| CNAG_034 | GMT2 | GDP-mani | 979.5990809 | 1070.8205 | 1282.650428 | 623.5560464 | 734.6598482 | 673.0190102 | 1.29E-10 | -0.72982 | 2.47E-11 | -6.67517 | 0.602979 | 9.889246 |
| CNAG_055 | - | hypothetic | 1059.469693 | 1134.642628 | 1005.816114 | 1550.744155 | 1514.860084 | 1601.724387 | 1.31E-10 | 0.52992 | 2.50E-11 | 6.673257 | 1.443849 | 9.883848 |
| CNAG_027 | - | Monosacc | 54.47091468 | 97.16084071 | 60.0069064 | 181.188832 | 229.4368652 | 200.4840788 | 1.37E-10 | 1.518289 | 2.64E-11 | 6.665371 | 2.86451 | 9.861898 |
| CNAG_049 | URA3 | orotidine | 1215.362713 | 1221.680111 | 1265.74103 | 805.0415662 | 853.6902028 | 886.0735107 | 1.69E-10 | -0.55606 | 3.26E-11 | -6.63438 | 0.680157 | 9.772647 |
| CNAG_075 | - | hypothetic | 12807.64669 | 11888.3362 | 12674.70405 | 9311.110261 | 9650.153024 | 9936.058343 | 2.07E-10 | -0.38654 | 4.03E-11 | -6.60281 | 0.764962 | 9.683762 |
| CNAG_037 | - | signal rec | 2687.922302 | 2551.294151 | 2845.154618 | 1939.90095 | 2023.380519 | 1956.283621 | 2.21E-10 | -0.46541 | 4.32E-11 | -6.59276 | 0.724263 | 9.655135 |
| CNAG_032 | - | rRNA biog | 1726.740901 | 1603.121179 | 1681.674578 | 1056.377191 | 1180.619621 | 1227.985243 | 2.31E-10 | -0.54836 | 4.51E-11 | -6.58619 | 0.683798 | 9.637019 |
| CNAG_050 | - | hypothetic | 40.45182715 | 61.67386091 | 59.95163734 | 136.6855355 | 188.090457 | 162.7130647 | 2.49E-10 | 1.588415 | 4.87E-11 | 6.574722 | 3.007187 | 9.604592 |
| CNAG_075 | - | threonine | 334.3368112 | 324.4884727 | 275.3626046 | 547.7058659 | 517.7916267 | 593.5343851 | 2.73E-10 | 0.813636 | 5.36E-11 | 6.560542 | 1.757636 | 9.564329 |
| CNAG_067 | - | hypothetic | 1955.692587 | 1820.790521 | 2010.126058 | 1411.485843 | 1369.508598 | 1372.348208 | 2.85E-10 | -0.49412 | 5.63E-11 | -6.55337 | 0.709994 | 9.545072 |
| CNAG_033 | - | pre-mRNA | 5568.830471 | 5838.48017 | 5639.324643 | 4018.657328 | 4373.006449 | 4549.545545 | 3.17E-10 | -0.41251 | 6.26E-11 | -6.53738 | 0.751315 | 9.499414 |
| CNAG_070 | - | hypothetic | 19.3723723 | 23.99891687 | 26.89952674 | 92.84240925 | 92.0348539 | 97.13736855 | 3.30E-10 | 2.032092 | 6.53E-11 | 6.531068 | 4.089976 | 9.481891 |
| CNAG_036 | - | malonic se | 1594.941855 | 1670.958859 | 1634.771492 | 1200.569643 | 1180.622208 | 1173.624524 | 3.33E-10 | -0.47821 | 6.60E-11 | -6.52938 | 0.717869 | 9.477533 |

| | | | | | | | | | | | | | | |
|---|---|---|---|---|---|---|---|---|---|---|---|---|---|---|
| CNAG_050 | URM1 | ubiquitin- | 866.115417 | 785.373003 | 975.5560199 | 540.4045115 | 577.6656729 | 512.0735987 | 3.49E-10 | -0.70488 | 6.93E-11 | -6.52208 | 0.613494 | 9.457438 |
| CNAG_031 | - | U4/U6.U5 | 502.8661151 | 494.3573912 | 498.9343524 | 299.134586 | 270.4468022 | 307.7450203 | 3.60E-10 | -0.78533 | 7.18E-11 | -6.51693 | 0.580221 | 9.443337 |
| CNAG_073 | - | hypothetic | 51.01973644 | 101.4870035 | 78.10340519 | 184.1376673 | 214.46925 | 262.3138286 | 3.82E-10 | 1.508035 | 7.64E-11 | 6.507594 | 2.844224 | 9.417665 |
| CNAG_057 | XRN2 | 5'-3' exori | 6804.317012 | 6889.011178 | 6976.649418 | 5065.50502 | 5527.149397 | 5472.106836 | 3.94E-10 | -0.37891 | 7.88E-11 | -6.50285 | 0.769021 | 9.405012 |
| CNAG_033 | - | alpha-keto | 3331.317843 | 3193.419678 | 3227.002575 | 2389.77199 | 2351.102592 | 2561.256456 | 3.95E-10 | -0.43277 | 7.91E-11 | -6.50234 | 0.740839 | 9.403803 |
| CNAG_032 | - | hypothetic | 1809.630876 | 1636.536535 | 1691.298166 | 1212.971189 | 1215.1128 | 1234.674339 | 4.09E-10 | -0.50404 | 8.21E-11 | -6.49676 | 0.705128 | 9.388487 |
| CNAG_037 | HXK2 | Hexokinas | 12925.42786 | 11361.45324 | 11117.73094 | 8816.139211 | 9015.306739 | 8747.13103 | 5.48E-10 | -0.42946 | 1.11E-10 | -6.45125 | 0.742538 | 9.261029 |
| CNAG_036 | - | peptidylpr | 2240.4979 | 1918.857636 | 2130.650767 | 1527.454345 | 1476.464137 | 1441.201385 | 7.01E-10 | -0.51671 | 1.43E-10 | -6.41306 | 0.698962 | 9.153984 |
| CNAG_077 | HAD2 | hypothetic | 7127.818936 | 7426.640524 | 7315.794961 | 5959.938551 | 5669.426452 | 5691.942149 | 7.03E-10 | -0.35151 | 1.43E-10 | -6.41254 | 0.783764 | 9.153025 |
| CNAG_055 | - | hypothetic | 130.1674737 | 140.2911501 | 139.8994355 | 265.9317821 | 314.6216172 | 270.5916739 | 7.07E-10 | 1.038416 | 1.44E-10 | 6.411602 | 2.053971 | 9.150608 |
| CNAG_050 | - | hypothetic | 2.918470113 | 0.227157895 | 9.819147809 | 48.34569775 | 52.41820136 | 63.58073789 | 7.47E-10 | 3.872573 | 1.52E-10 | 6.402864 | 14.64741 | 9.126507 |
| CNAG_026 | - | hypothetic | 475.5827465 | 438.6997657 | 437.4057317 | 755.5673193 | 717.0825909 | 691.4439777 | 7.77E-10 | 0.664571 | 1.59E-10 | 6.396749 | 1.585097 | 9.109632 |
| CNAG_055 | PBX2 | parallel be | 2184.696617 | 2261.42237 | 2326.885509 | 1659.738685 | 1737.028626 | 1660.976233 | 8.88E-10 | -0.4366 | 1.82E-10 | -6.37579 | 0.738875 | 9.051697 |
| CNAG_068 | - | hypothetic | 27.71096868 | 69.21920509 | 73.79460942 | 180.7587767 | 193.0882653 | 181.5807793 | 8.96E-10 | 1.699641 | 1.84E-10 | 6.374348 | 3.248202 | 9.047868 |
| CNAG_055 | URE1 | urease | 9708.455851 | 9368.198259 | 9283.417649 | 7242.44005 | 7668.891984 | 7543.670637 | 8.96E-10 | -0.35233 | 1.84E-10 | -6.37419 | 0.783318 | 9.047668 |
| CNAG_008 | FRE7 | ferric-chel | 1165.614346 | 871.5192416 | 848.8113439 | 436.5679457 | 619.9520855 | 533.1734415 | 9.07E-10 | -0.8776 | 1.86E-10 | -6.3723 | 0.544271 | 9.042577 |
| CNAG_008 | SKN1 | putative g | 3765.043983 | 3763.569115 | 3498.089311 | 4930.789913 | 5080.73372 | 4665.917341 | 1.08E-09 | 0.39717 | 2.23E-10 | 6.344326 | 1.316922 | 8.965826 |
| CNAG_068 | - | hypothetic | 475.0483965 | 514.8331984 | 591.5999207 | 302.1970059 | 273.0065276 | 327.7155418 | 1.10E-09 | -0.82366 | 2.27E-10 | -6.34219 | 0.565007 | 8.960055 |
| CNAG_032 | - | 18S rRNA | 2150.46047 | 1755.084088 | 1905.643689 | 1239.300013 | 1395.42357 | 1327.954979 | 1.14E-09 | -0.56892 | 2.37E-10 | -6.33546 | 0.674123 | 8.941871 |
| CNAG_074 | - | hypothetic | 19.43790168 | 42.2507224 | 29.06002817 | 114.7396194 | 113.6617288 | 108.3399993 | 1.36E-09 | 1.883078 | 2.83E-10 | 6.307835 | 3.688613 | 8.866249 |
| CNAG_032 | - | hypothetic | 937.985041 | 901.9627264 | 926.8649558 | 1286.811036 | 1313.034881 | 1391.888383 | 1.53E-09 | 0.514297 | 3.18E-10 | 6.289497 | 1.428298 | 8.815898 |
| CNAG_034 | - | alanine tra | 1776.088516 | 2273.11808 | 1976.033806 | 1337.417483 | 1429.044116 | 1396.777681 | 1.64E-09 | -0.54785 | 3.43E-10 | -6.27783 | 0.684039 | 8.784299 |
| CNAG_032 | - | hypothetic | 164.201522 | 271.6484437 | 211.3734937 | 410.645778 | 524.3637242 | 412.7322675 | 1.77E-09 | 1.04538 | 3.71E-10 | 6.26556 | 2.063911 | 8.751088 |
| CNAG_009 | - | hypothetic | 9.922912043 | 6.752061922 | 4.51547054 | 76.005946 | 54.06894468 | 41.99981916 | 1.79E-09 | 3.154025 | 3.75E-10 | 6.263928 | 8.901358 | 8.746794 |
| CNAG_056 | RPD304 | histone de | 4390.49531 | 4423.813754 | 4067.411016 | 3312.890145 | 3190.391086 | 3389.445357 | 1.87E-09 | -0.39597 | 3.93E-10 | -6.25695 | 0.75998 | 8.728114 |
| CNAG_031 | - | vacuolar p | 299.3228007 | 259.8999714 | 278.5237697 | 429.0076011 | 562.9698097 | 534.716436 | 1.91E-09 | 0.851313 | 4.02E-10 | 6.253386 | 1.804142 | 8.718453 |
| CNAG_056 | CKA1 | CMGC/CK | 6653.70963 | 7091.537139 | 6639.683934 | 5524.084311 | 5299.390491 | 5337.79561 | 2.15E-09 | -0.34982 | 4.54E-10 | -6.23421 | 0.784681 | 8.667454 |
| CNAG_026 | - | D-lactate | 1461.815021 | 1576.115479 | 1454.562717 | 1115.764072 | 1052.073456 | 1075.928439 | 2.23E-09 | -0.48498 | 4.71E-10 | -6.22861 | 0.714507 | 8.652664 |
| CNAG_027 | DCR2 | dicer-like | 2157.724856 | 2016.884557 | 2191.43762 | 1475.204492 | 1596.415262 | 1591.012172 | 2.28E-09 | -0.46461 | 4.84E-10 | -6.2243 | 0.724665 | 8.641484 |
| CNAG_025 | - | thioredoxi | 51.89905389 | 43.3832605 | 34.40819644 | 127.7120404 | 118.5677297 | 150.3057326 | 2.30E-09 | 1.610506 | 4.87E-10 | 6.223314 | 3.053589 | 8.639003 |
| CNAG_049 | - | signal rec | 1210.537173 | 1079.500922 | 1129.265027 | 802.8274268 | 769.1328154 | 789.5209337 | 2.54E-09 | -0.55003 | 5.41E-10 | -6.20672 | 0.683008 | 8.594367 |
| CNAG_036 | - | ATP-deper | 1239.602387 | 1013.80349 | 1116.450434 | 735.8061883 | 712.1617194 | 777.294892 | 2.56E-09 | -0.61472 | 5.46E-10 | -6.20531 | 0.653055 | 8.590953 |
| CNAG_033 | - | tRNA (gua | 1850.426946 | 1600.990255 | 1658.24444 | 1129.598159 | 1229.801538 | 1229.117344 | 2.62E-09 | -0.52585 | 5.58E-10 | -6.20188 | 0.694549 | 8.581744 |
| CNAG_049 | - | hypothetic | 1058.652295 | 934.0262202 | 930.9239397 | 673.621894 | 634.5518344 | 649.6605878 | 2.69E-09 | -0.59469 | 5.75E-10 | -6.19713 | 0.662185 | 8.569621 |
| CNAG_057 | SLA2 | cell wall o | 425.4551201 | 451.5946334 | 444.8471745 | 665.0286277 | 703.3107764 | 700.2589668 | 2.78E-09 | 0.632639 | 5.94E-10 | 6.192077 | 1.550399 | 8.556191 |
| CNAG_027 | - | NAD bind | 1002.19193 | 1001.055725 | 922.6258486 | 1422.365411 | 1344.906182 | 1418.576542 | 2.98E-09 | 0.50138 | 6.38E-10 | 6.180654 | 1.415567 | 8.525976 |
| CNAG_032 | - | hypothetic | 490.7462965 | 444.0894949 | 283.975956 | 864.9074066 | 733.4808491 | 759.058277 | 3.08E-09 | 0.937946 | 6.62E-10 | 6.174863 | 1.915798 | 8.510794 |
| CNAG_037 | - | hypothetic | 1014.530947 | 1013.751145 | 1117.465218 | 732.7841764 | 743.3163867 | 647.5000861 | 3.19E-09 | -0.5827 | 6.84E-10 | -6.1697 | 0.667715 | 8.496847 |
| CNAG_049 | - | hypothetic | 744.2906938 | 926.6661639 | 727.5020516 | 1152.918549 | 1285.568455 | 1279.740074 | 3.23E-09 | 0.618326 | 6.94E-10 | 6.167537 | 1.535093 | 8.49116 |
| CNAG_123 | - | unspecifie | 35.75837138 | 53.05155821 | 51.41797281 | 136.2655201 | 118.4578889 | 170.1818039 | 3.24E-09 | 1.595277 | 6.97E-10 | 6.166874 | 3.021524 | 8.489589 |
| CNAG_033 | - | hypothetic | 318.0738219 | 287.9228539 | 373.3640436 | 568.7408835 | 537.642544 | 588.0445946 | 3.29E-09 | 0.775993 | 7.07E-10 | 6.164542 | 1.712368 | 8.483432 |
| CNAG_037 | - | hypothetic | 5.244303133 | 6.728790644 | 4.499014559 | 47.70515672 | 48.96526695 | 50.50813888 | 3.41E-09 | 3.258316 | 7.35E-10 | 6.158394 | 9.568653 | 8.467061 |
| CNAG_076 | - | hypothetic | 116.0842265 | 113.3601374 | 102.6066453 | 234.1897461 | 256.7920614 | 219.5550324 | 3.42E-09 | 1.083545 | 7.37E-10 | 6.157879 | 2.119237 | 8.46614 |
| CNAG_075 | - | hypothetic | 409.3864114 | 337.1845556 | 483.809314 | 229.050874 | 212.6563088 | 203.501347 | 3.95E-09 | -0.94867 | 8.55E-10 | -6.13433 | 0.51811 | 8.403874 |
| CNAG_051 | CXT1 | beta-1,2-x | 1108.892299 | 1075.091208 | 929.90155 | 730.4750642 | 639.6806446 | 690.7345191 | 4.00E-09 | -0.61096 | 8.67E-10 | -6.13215 | 0.654762 | 8.398434 |
| CNAG_034 | - | exportin-1 | 4002.739594 | 3839.86288 | 3578.979108 | 2845.863976 | 2986.890346 | 2888.796433 | 4.10E-09 | -0.40459 | 8.91E-10 | -6.12787 | 0.755451 | 8.387489 |
| CNAG_026 | VPS15 | VPS15 pro | 2402.974276 | 2418.709803 | 2250.163982 | 1742.23785 | 1831.052887 | 1769.758851 | 4.29E-09 | -0.41952 | 9.33E-10 | -6.12047 | 0.747672 | 8.368031 |
| CNAG_033 | - | Phosphati | 2050.317666 | 2053.466978 | 1963.273083 | 1499.42325 | 1575.742568 | 1472.258972 | 4.47E-09 | -0.43143 | 9.76E-10 | -6.11324 | 0.741527 | 8.349563 |
| CNAG_051 | - | hypothetic | 21.75266567 | 36.89507224 | 36.4869207 | 96.36293986 | 105.7809761 | 126.9438029 | 5.07E-09 | 1.800127 | 1.11E-09 | 6.092238 | 3.482508 | 8.294896 |
| CNAG_067 | - | DNA repa | 260.8954401 | 271.7200192 | 305.1342729 | 438.4692684 | 498.0491064 | 523.6036875 | 5.29E-09 | 0.788186 | 1.16E-09 | 6.085102 | 1.726901 | 8.276265 |
| CNAG_055 | - | phosphati | 389.2908192 | 441.8794117 | 416.0585051 | 644.0731976 | 655.8418143 | 664.7315003 | 5.59E-09 | 0.640976 | 1.23E-09 | 6.075953 | 1.559384 | 8.252207 |
| CNAG_037 | ALF1 | tubulin-fo | 916.462656 | 972.7630219 | 961.8142831 | 679.8872225 | 619.8792433 | 664.0639133 | 5.95E-09 | -0.55305 | 1.31E-09 | -6.06574 | 0.681578 | 8.225773 |
| CNAG_033 | - | phosphol | 340.0572971 | 229.7816309 | 266.8131025 | 490.0250911 | 555.1555842 | 509.272151 | 5.95E-09 | 0.879294 | 1.32E-09 | 6.065503 | 1.839475 | 8.225386 |
| CNAG_037 | - | microtubu | 2122.096847 | 2259.471528 | 2255.631989 | 2945.363948 | 2901.825703 | 2907.372842 | 6.64E-09 | 0.384454 | 1.47E-09 | 6.047608 | 1.305366 | 8.178054 |
| CNAG_052 | CAT2 | catalase 2 | 1250.248395 | 553.1842254 | 566.6817243 | 4324.300301 | 4325.057944 | 4152.365327 | 6.69E-09 | 2.41488 | 1.48E-09 | 6.046274 | 5.33275 | 8.174698 |
| CNAG_078 | - | hypothetic | 272.5059929 | 258.8011904 | 297.6689081 | 487.0780654 | 447.0578808 | 478.1861787 | 7.53E-09 | 0.756367 | 1.68E-09 | 6.026475 | 1.689232 | 8.12335 |
| CNAG_051 | - | prefoldin | 1555.035503 | 1341.350424 | 1556.858272 | 1037.86991 | 1084.921016 | 895.0854472 | 7.60E-09 | -0.57804 | 1.70E-09 | -6.02456 | 0.669875 | 8.118915 |
| CNAG_058 | - | protein RA | 1235.013918 | 1070.887939 | 1109.016234 | 794.2492311 | 786.4068736 | 781.7573483 | 7.69E-09 | -0.54787 | 1.72E-09 | -6.02266 | 0.684028 | 8.114281 |
| CNAG_050 | - | iron dono | 994.6953067 | 1010.511231 | 1084.421375 | 751.5269825 | 715.6869101 | 651.9275857 | 8.23E-09 | -0.56 | 1.84E-09 | -6.01124 | 0.6783 | 8.08485 |
| CNAG_032 | - | splicing fa | 3105.946193 | 3088.877165 | 2858.049886 | 2173.361578 | 2367.574108 | 2340.355109 | 9.29E-09 | -0.41089 | 2.09E-09 | -5.99101 | 0.75216 | 8.032147 |
| CNAG_052 | - | clathrin bi | 3115.283369 | 3015.634393 | 2876.175456 | 2281.626625 | 2302.849694 | 2352.562675 | 9.37E-09 | -0.39228 | 2.11E-09 | -5.98924 | 0.761923 | 8.028133 |
| CNAG_033 | - | SIT4-assoc | 5961.136646 | 5609.057189 | 5819.560428 | 4593.980625 | 4492.889833 | 4678.334056 | 1.12E-08 | -0.35286 | 2.53E-09 | -5.95931 | 0.783032 | 7.951238 |
| CNAG_036 | - | UDP-N-ac | 616.0595952 | 554.7440375 | 551.2466623 | 370.8012723 | 349.8721152 | 368.7970227 | 1.21E-08 | -0.67842 | 2.74E-09 | -5.94669 | 0.624847 | 7.918423 |
| CNAG_126 | - | unspecifie | 43.80842038 | 46.59392242 | 41.84012641 | 119.792021 | 156.2369927 | 108.5714442 | 1.25E-08 | 1.543356 | 2.83E-09 | 5.941291 | 2.914717 | 7.904356 |
| CNAG_051 | - | hypothetic | 1235.858739 | 1410.45601 | 1270.282581 | 1670.280891 | 2029.966239 | 2015.774313 | 1.25E-08 | 0.530783 | 2.84E-09 | 5.940768 | 1.444713 | 7.903205 |
| CNAG_033 | - | hypothetic | 31.06884273 | 53.00391163 | 37.58158644 | 126.3146148 | 130.0649139 | 112.9017451 | 1.29E-08 | 1.590991 | 2.94E-09 | 5.93501 | 3.012563 | 7.888423 |
| CNAG_051 | - | exoribonu | 683.9236444 | 620.5156618 | 721.7916201 | 422.1096565 | 465.5676905 | 407.7024002 | 1.31E-08 | -0.66129 | 2.97E-09 | -5.93309 | 0.632313 | 7.883562 |
| CNAG_057 | - | OPT famil | 1249.762696 | 1301.636918 | 1218.008941 | 1677.484321 | 1725.334125 | 1740.551066 | 1.35E-08 | 0.433507 | 3.07E-09 | 5.927862 | 1.350513 | 7.870211 |

| | | | | | | | | | | | | | | |
|---|---|---|---|---|---|---|---|---|---|---|---|---|---|---|
| CNAG_034 | - | mitochond | 1037.759609 | 994.339314 | 975.7075348 | 620.5141038 | 731.2652772 | 708.4593081 | 1.49E-08 | -0.56137 | 3.42E-09 | -5.91011 | 0.677659 | 7.825426 |
| CNAG_075 | - | pyridoxal | 634.0249133 | 795.9823941 | 802.8253797 | 446.1872959 | 512.1522761 | 438.7982455 | 1.53E-08 | -0.69109 | 3.50E-09 | -5.90602 | 0.619387 | 7.814887 |
| CNAG_055 | - | hypothetic | 3786.775721 | 3522.077558 | 3784.746559 | 2867.707612 | 2866.108749 | 2894.326406 | 1.57E-08 | -0.37819 | 3.59E-09 | -5.90212 | 0.769402 | 7.805317 |
| CNAG_037 | - | ankyrin re | 1529.42612 | 1379.055121 | 1563.271756 | 1037.054632 | 1135.84002 | 987.1736422 | 1.66E-08 | -0.5171 | 3.81E-09 | -5.89228 | 0.698775 | 7.780579 |
| CNAG_056 | - | ubiquitin- | 2491.609383 | 2290.528347 | 2379.14779 | 1861.471699 | 1452.176695 | 1713.152827 | 1.72E-08 | -0.52588 | 3.95E-09 | -5.88617 | 0.694533 | 7.765441 |
| CNAG_074 | MBS1 | Mbp1 anc | 2815.226511 | 2808.765019 | 2705.533711 | 2205.427126 | 2159.675393 | 2116.12734 | 1.77E-08 | -0.3772 | 4.08E-09 | -5.88107 | 0.76993 | 7.752525 |
| CNAG_033 | - | hypothetic | 964.2106292 | 873.7248985 | 972.4585286 | 655.7217882 | 649.264796 | 635.2246755 | 2.01E-08 | -0.55047 | 4.66E-09 | -5.85902 | 0.682799 | 7.69612 |
| CNAG_057 | - | WSC dom | 225.7584645 | 152.2397797 | 227.3033981 | 414.0846566 | 427.5344004 | 351.7489818 | 2.03E-08 | 0.963722 | 4.70E-09 | 5.857585 | 1.950335 | 7.692608 |
| CNAG_031 | SWE1 | WEE prote | 1242.838259 | 1331.82824 | 1379.010224 | 1715.526215 | 1956.613277 | 1840.471307 | 2.14E-08 | 0.464557 | 4.97E-09 | 5.848295 | 1.379894 | 7.669026 |
| CNAG_033 | - | hypothetic | 2092.46421 | 2211.850985 | 2171.212557 | 1629.410047 | 1697.361687 | 1649.852003 | 2.42E-08 | -0.39503 | 5.63E-09 | -5.82726 | 0.760473 | 7.615359 |
| CNAG_031 | - | rhomboid | 1403.196045 | 1244.3452 | 1254.066638 | 975.7053592 | 862.2646426 | 903.8711961 | 2.47E-08 | -0.52527 | 5.74E-09 | -5.82424 | 0.694827 | 7.607954 |
| CNAG_025 | - | avenacina | 258.5920387 | 305.0771994 | 310.4706812 | 465.7983555 | 549.9888054 | 468.2925801 | 3.04E-08 | 0.748492 | 7.14E-09 | 5.787605 | 1.680036 | 7.51752 |
| CNAG_078 | - | aldo-keto | 2812.716091 | 2419.867621 | 2572.200937 | 2004.608945 | 1896.54514 | 1897.446337 | 3.15E-08 | -0.4446 | 7.42E-09 | -5.78116 | 0.734786 | 7.501781 |
| CNAG_057 | - | hypothetic | 2058.408673 | 1800.353254 | 2069.847253 | 1456.630755 | 1494.643272 | 1353.510448 | 3.15E-08 | -0.478 | 7.44E-09 | -5.78078 | 0.717972 | 7.501011 |
| CNAG_025 | - | hypothetic | 2847.978254 | 2876.657575 | 3046.737141 | 2115.778874 | 2141.508611 | 2384.712094 | 3.22E-08 | -0.41638 | 7.60E-09 | -5.77713 | 0.749301 | 7.491828 |
| CNAG_036 | - | NADPH2:c | 807.2133379 | 815.7471965 | 827.6727554 | 1127.271398 | 1143.970115 | 1174.34396 | 3.57E-08 | 0.4767 | 8.45E-09 | 5.759221 | 1.391557 | 7.447251 |
| CNAG_036 | - | dynactin 6 | 539.2600101 | 546.1352347 | 622.5918283 | 373.9338311 | 355.928985 | 362.1466221 | 3.86E-08 | -0.66245 | 9.17E-09 | -5.74537 | 0.631805 | 7.413387 |
| CNAG_033 | - | NifU-like p | 2829.235817 | 2641.818975 | 2936.89243 | 2123.631878 | 2162.260939 | 2162.73739 | 3.99E-08 | -0.39843 | 9.51E-09 | -5.73923 | 0.758682 | 7.398797 |
| CNAG_079 | - | hypothetic | 12.22368794 | 5.677270233 | 9.832538175 | 65.33455931 | 43.40390467 | 58.27757081 | 4.00E-08 | 2.658101 | 9.55E-09 | 5.738542 | 6.312018 | 7.397542 |
| CNAG_057 | - | hypothetic | 1375.348648 | 1334.84431 | 1375.620305 | 956.1014312 | 1001.222818 | 1042.573973 | 4.00E-08 | -0.46086 | 9.54E-09 | -5.73864 | 0.726555 | 7.397542 |
| CNAG_051 | - | U3 small r | 1193.116423 | 1083.833205 | 1223.071897 | 762.2251388 | 840.7857334 | 854.9786419 | 4.03E-08 | -0.52638 | 9.62E-09 | -5.73736 | 0.694296 | 7.394908 |
| CNAG_034 | - | BET3 fami | 1564.502418 | 1429.701031 | 1642.18466 | 1142.213487 | 1152.194959 | 1043.782939 | 4.16E-08 | -0.48987 | 9.94E-09 | -5.73169 | 0.712088 | 7.38105 |
| CNAG_033 | ZIP2 | solute car | 12.30028103 | 14.28482043 | 15.16402735 | 59.10914442 | 79.48931157 | 57.40599898 | 4.17E-08 | 2.234324 | 9.98E-09 | 5.731098 | 4.705422 | 7.37975 |
| CNAG_052 | - | hypothetic | 889.4625898 | 801.5366819 | 895.6704472 | 577.048818 | 622.5728451 | 569.7418362 | 4.38E-08 | -0.56389 | 1.05E-08 | -5.72225 | 0.676473 | 7.358211 |
| CNAG_055 | - | hypothetic | 5.225325734 | 4.572280125 | 7.678141072 | 40.44226354 | 44.59453842 | 50.43282628 | 4.38E-08 | 3.050466 | 1.05E-08 | 5.722303 | 8.284796 | 7.358211 |
| CNAG_051 | - | ubiquinol- | 1912.443423 | 1943.51775 | 1645.518911 | 1364.025481 | 1348.836581 | 1309.072601 | 4.41E-08 | -0.46689 | 1.06E-08 | -5.72124 | 0.723523 | 7.355853 |
| CNAG_036 | - | hypothetic | 57.89991422 | 96.04241374 | 61.04842003 | 175.1435841 | 169.5014513 | 184.9181711 | 4.50E-08 | 1.289791 | 1.08E-08 | 5.717497 | 2.444926 | 7.346508 |
| CNAG_037 | - | pantethein | 1114.80067 | 1001.926479 | 1033.296144 | 774.0961241 | 726.0213451 | 718.4866894 | 4.64E-08 | -0.52082 | 1.12E-08 | -5.712 | 0.696974 | 7.333344 |
| CNAG_032 | - | hypothetic | 1175.068231 | 1302.695134 | 1157.232029 | 1598.828237 | 1767.689794 | 1658.424971 | 5.31E-08 | 0.452137 | 1.28E-08 | 5.688255 | 1.368065 | 7.274619 |
| CNAG_031 | - | hypothetic | 13.381729 | 8.901574998 | 9.836027825 | 57.08065898 | 51.36555326 | 59.39041495 | 5.34E-08 | 2.460842 | 1.29E-08 | 5.687249 | 5.505378 | 7.272282 |
| CNAG_031 | - | hypothetic | 62.38679303 | 55.21508552 | 56.74402838 | 143.2440982 | 137.5806545 | 146.1117556 | 5.64E-08 | 1.284633 | 1.36E-08 | 5.677735 | 2.436201 | 7.248553 |
| CNAG_052 | RAD53 | CAMK pro | 1409.777698 | 1547.274138 | 1433.424151 | 1973.278045 | 2060.061163 | 1888.284582 | 5.66E-08 | 0.417007 | 1.37E-08 | 5.67722 | 1.335155 | 7.247464 |
| CNAG_049 | - | hypothetic | 1130.645141 | 1159.422373 | 1123.082314 | 1536.658428 | 1537.278691 | 1536.307625 | 6.39E-08 | 0.418719 | 1.55E-08 | 5.655612 | 1.33674 | 7.194256 |
| CNAG_036 | - | heterokary | 1959.711771 | 2011.652731 | 1852.547497 | 2540.607419 | 2666.42171 | 2478.887385 | 6.71E-08 | 0.385184 | 1.63E-08 | 5.646837 | 1.306027 | 7.172954 |
| CNAG_033 | - | hypothetic | 2.932438941 | 13.1713418 | 8.768309419 | 67.04616358 | 54.10121596 | 43.06669868 | 6.94E-08 | 2.821776 | 1.69E-08 | 5.641028 | 7.070323 | 7.158729 |
| CNAG_123 | - | unspecifie | 15.85257229 | 22.89586806 | 24.74851491 | 65.03002079 | 100.2366808 | 76.20394222 | 6.97E-08 | 1.967146 | 1.70E-08 | 5.640275 | 3.909939 | 7.157047 |
| CNAG_051 | - | V-type pro | 698.0181998 | 760.4452031 | 654.7336417 | 499.3310855 | 452.5417857 | 462.0535534 | 7.91E-08 | -0.59456 | 1.94E-08 | -5.61754 | 0.662248 | 7.101761 |
| CNAG_058 | - | hypothetic | 758.6543783 | 697.0004121 | 744.2480491 | 526.5905852 | 500.9025714 | 449.8917661 | 8.08E-08 | -0.59183 | 1.98E-08 | -5.61353 | 0.663501 | 7.092339 |
| CNAG_127 | - | unspecifie | 28.59327181 | 29.3491655 | 18.40777759 | 114.5993237 | 76.21546133 | 81.82238533 | 8.45E-08 | 1.838785 | 2.08E-08 | 5.605161 | 3.577087 | 7.073067 |
| CNAG_074 | - | hypothetic | 31.10079665 | 60.52987016 | 48.21695555 | 136.4442908 | 121.1750419 | 133.8581686 | 8.59E-08 | 1.475191 | 2.12E-08 | 5.602056 | 2.780205 | 7.066145 |
| CNAG_049 | - | hypothetic | 72.89649231 | 67.05721557 | 67.40538215 | 169.0225756 | 176.6123188 | 138.5891674 | 8.66E-08 | 1.213694 | 2.14E-08 | 5.60047 | 2.319307 | 7.062601 |
| CNAG_037 | - | tRNA ader | 813.2709204 | 647.4617467 | 672.8387179 | 475.9115496 | 433.5313542 | 465.3743354 | 1.04E-07 | -0.65089 | 2.60E-08 | -5.56672 | 0.636885 | 6.982975 |
| CNAG_037 | ENV7 | NAK prote | 1617.059823 | 1567.564843 | 1535.623288 | 1167.870394 | 1235.037212 | 1154.747504 | 1.05E-07 | -0.42366 | 2.62E-08 | -5.56483 | 0.745533 | 6.978494 |
| CNAG_051 | - | hypothetic | 2784.795823 | 2479.138982 | 2797.187143 | 1929.756753 | 2102.768091 | 2057.27323 | 1.06E-07 | -0.42071 | 2.65E-08 | -5.5632 | 0.747057 | 6.974843 |
| CNAG_033 | - | mitochond | 1388.076665 | 1269.123878 | 1273.2643 | 953.8422229 | 1009.047493 | 865.0633717 | 1.21E-07 | -0.49103 | 3.04E-08 | -5.53937 | 0.711515 | 6.917975 |
| CNAG_057 | - | transcripti | 4922.246438 | 4723.565995 | 4812.984563 | 6168.230044 | 5920.728323 | 6123.51864 | 1.24E-07 | 0.31767 | 3.13E-08 | 5.53392 | 1.246316 | 6.905513 |
| CNAG_051 | - | ornithine- | 3698.518623 | 4014.573586 | 3688.961661 | 4684.297454 | 4886.578489 | 4958.933946 | 1.38E-07 | 0.334983 | 3.49E-08 | 5.51498 | 1.261362 | 6.860098 |
| CNAG_055 | - | hypothetic | 48.64152135 | 62.79773254 | 103.559812 | 160.8024269 | 196.4565893 | 199.284384 | 1.45E-07 | 1.366365 | 3.68E-08 | 5.505717 | 2.578202 | 6.838121 |
| CNAG_031 | - | solute car | 899.0067514 | 925.4140597 | 1032.150817 | 670.5029157 | 665.6522192 | 682.9238111 | 1.50E-07 | -0.51703 | 3.80E-08 | -5.49974 | 0.698808 | 6.823818 |
| CNAG_123 | - | unspecifie | 89.17251167 | 58.50169297 | 78.06092549 | 202.765331 | 192.1114188 | 145.3106196 | 1.59E-07 | 1.243421 | 4.04E-08 | 5.488965 | 2.367592 | 6.798342 |
| CNAG_055 | - | hypothetic | 338.3340327 | 403.7100456 | 329.4301039 | 199.4368463 | 185.8987504 | 229.9913795 | 1.62E-07 | -0.81779 | 4.13E-08 | -5.48517 | 0.56731 | 6.789222 |
| CNAG_026 | - | hypothetic | 627.4728668 | 681.0727932 | 728.4690782 | 995.8663294 | 949.8698648 | 983.4049651 | 1.75E-07 | 0.508693 | 4.46E-08 | 5.471789 | 1.422761 | 6.757429 |
| CNAG_078 | - | solute car | 11.25866555 | 31.49431853 | 33.26271915 | 89.59734239 | 100.8155742 | 89.49962268 | 1.77E-07 | 1.876678 | 4.52E-08 | 5.46923 | 3.672285 | 6.751942 |
| CNAG_055 | - | peroxin-10 | 548.0396308 | 564.7265035 | 613.2997879 | 818.3096942 | 880.1378856 | 819.1356441 | 1.88E-07 | 0.529764 | 4.82E-08 | 5.457842 | 1.443693 | 6.725773 |
| CNAG_036 | - | lipoate-pr | 1316.799641 | 1121.558976 | 1226.312639 | 833.1046999 | 916.7276889 | 867.2236513 | 1.88E-07 | -0.50169 | 4.83E-08 | -5.45756 | 0.706277 | 6.725286 |
| CNAG_050 | HRB1 | RNP doma | 10188.52716 | 10420.85605 | 10642.06192 | 8227.00184 | 8894.633447 | 8562.786236 | 2.14E-07 | -0.29832 | 5.51E-08 | -5.4339 | 0.8132 | 6.669405 |
| CNAG_075 | - | hypothetic | 763.5348529 | 810.0951306 | 839.1548608 | 580.266583 | 551.7780877 | 566.3699048 | 2.14E-07 | -0.52228 | 5.52E-08 | -5.43354 | 0.696271 | 6.668741 |
| CNAG_008 | - | ATP-bindi | 10519.92285 | 9200.15317 | 9357.013371 | 7493.8495 | 7957.867972 | 7418.25779 | 2.29E-07 | -0.36245 | 5.91E-08 | -5.42138 | 0.777841 | 6.6406 |
| CNAG_131 | - | unspecifie | 27.2977453 | 6.791255725 | 13.06352696 | 62.84358802 | 88.96362587 | 74.98525306 | 2.31E-07 | 2.264543 | 5.98E-08 | 5.419476 | 4.80502 | 6.636182 |
| CNAG_058 | - | translation | 430.1161367 | 491.4116725 | 438.4558256 | 694.7176645 | 691.2016918 | 650.4026398 | 2.37E-07 | 0.567725 | 6.15E-08 | 5.414345 | 1.482185 | 6.624544 |
| CNAG_056 | - | neutral an | 6918.822276 | 7491.266452 | 6911.64177 | 5615.068179 | 5826.459256 | 5921.696018 | 2.38E-07 | -0.31109 | 6.17E-08 | -5.41369 | 0.806032 | 6.623163 |
| CNAG_033 | BCH1 | ChAPs fan | 1527.196192 | 1513.697271 | 1521.743107 | 1196.766315 | 1115.062014 | 1154.72104 | 2.38E-07 | -0.41171 | 6.18E-08 | -5.4135 | 0.75173 | 6.622903 |
| CNAG_032 | - | hypothetic | 27.30722682 | 14.31226408 | 10.9400859 | 81.41653255 | 56.25818909 | 88.0768822 | 2.48E-07 | 2.125147 | 6.45E-08 | 5.405852 | 4.362475 | 6.605786 |
| CNAG_078 | - | hypothetic | 302.9319368 | 378.2888728 | 341.4084475 | 576.5887647 | 507.3723496 | 544.8462245 | 2.71E-07 | 0.657938 | 7.06E-08 | 5.389608 | 1.577825 | 6.567697 |
| CNAG_055 | - | hypothetic | 2421.652399 | 2273.316439 | 2302.406754 | 1803.775585 | 1836.225952 | 1837.449297 | 2.76E-07 | -0.36891 | 7.20E-08 | -5.38617 | 0.774368 | 6.559791 |
| CNAG_033 | - | hypothetic | 961.9656631 | 990.0123867 | 933.0664234 | 666.5755768 | 745.1132098 | 657.4341133 | 2.81E-07 | -0.49508 | 7.35E-08 | -5.38247 | 0.709523 | 6.551275 |
| CNAG_055 | - | DNA repa | 1037.928518 | 1095.605265 | 1036.513191 | 784.1905678 | 773.5005515 | 782.8267462 | 3.17E-07 | -0.45265 | 8.32E-08 | -5.36005 | 0.730699 | 6.498881 |
| CNAG_037 | ISP1 | short-chai | 5.161746024 | 3.469637815 | 3.403711874 | 37.57881071 | 38.60690765 | 34.04540619 | 3.27E-07 | 3.292715 | 8.59E-08 | 5.354212 | 9.799549 | 6.485265 |

| | | | | | | | | | | | | | | |
|---|---|---|---|---|---|---|---|---|---|---|---|---|---|---|
| CNAG_123 | - | unspecifie | 21.62461941 | 21.82898974 | 27.92980479 | 90.80499337 | 72.86740958 | 74.03623705 | 3.39E-07 | 1.758822 | 8.92E-08 | 5.347534 | 3.384217 | 6.470043 |
| CNAG_058 | - | mitochonc | 818.3303431 | 804.7353323 | 876.4634845 | 610.6681872 | 500.7773311 | 597.4293173 | 3.40E-07 | -0.56446 | 8.96E-08 | -5.34656 | 0.676208 | 6.46791 |
| CNAG_031 | ATG2602 | UDP-gluc | 429.0300163 | 521.5891969 | 475.7788622 | 684.3640436 | 692.8085496 | 856.6004521 | 3.46E-07 | 0.633164 | 9.11E-08 | 5.343578 | 1.550962 | 6.461363 |
| CNAG_050 | - | hypothetic | 1663.092495 | 1714.273773 | 1515.56252 | 2107.059215 | 2243.764215 | 2139.148379 | 3.51E-07 | 0.392503 | 9.25E-08 | 5.340777 | 1.312669 | 6.455053 |
| CNAG_036 | - | translatior | 9060.368751 | 8510.591828 | 9070.097836 | 6488.97168 | 7417.058236 | 7242.797361 | 3.90E-07 | -0.34882 | 1.03E-07 | -5.32118 | 0.785225 | 6.409377 |
| CNAG_055 | - | hypothetic | 8.824906427 | 17.48831146 | 18.33768764 | 67.28655671 | 61.79225576 | 59.56223432 | 4.17E-07 | 2.106709 | 1.10E-07 | 5.308646 | 4.307078 | 6.380093 |
| CNAG_075 | - | DNA repa | 3287.388007 | 3236.693812 | 2883.824666 | 3896.269616 | 4351.028037 | 4087.479752 | 4.73E-07 | 0.37569 | 1.26E-07 | 5.28459 | 1.29746 | 6.3253 |
| CNAG_077 | - | glucan en | 227.2966468 | 217.4901147 | 250.3677594 | 128.6531754 | 117.7698711 | 129.044172 | 5.14E-07 | -0.90732 | 1.37E-07 | -5.26892 | 0.533175 | 6.28936 |
| CNAG_052 | TLK1 | phosphati | 122.1026599 | 177.9366478 | 167.595223 | 340.3554518 | 246.7727428 | 324.8485911 | 5.23E-07 | 0.951912 | 1.40E-07 | 5.265478 | 1.934434 | 6.281826 |
| CNAG_033 | RDI1 | rho gdp-c | 856.2479029 | 941.7483031 | 798.9198998 | 1203.584877 | 1183.622303 | 1214.331725 | 5.41E-07 | 0.457577 | 1.45E-07 | 5.258861 | 1.373234 | 6.266985 |
| CNAG_049 | - | cytoplasm | 1674.369888 | 1659.167726 | 1747.809619 | 1260.478888 | 1373.076646 | 1259.097932 | 5.43E-07 | -0.39979 | 1.46E-07 | -5.25799 | 0.75797 | 6.265114 |
| CNAG_031 | CHK1 | CAMK/CA | 1292.518047 | 1338.055175 | 1316.982841 | 957.6961566 | 1039.241186 | 991.532284 | 5.50E-07 | -0.41687 | 1.48E-07 | -5.25534 | 0.749045 | 6.259465 |
| CNAG_032 | - | methionin | 4986.151945 | 4885.027 | 5181.808197 | 4021.937561 | 4235.934145 | 3962.310774 | 5.88E-07 | -0.31646 | 1.58E-07 | -5.24273 | 0.803039 | 6.23083 |
| CNAG_075 | - | hypothetic | 181.3102208 | 165.0356757 | 139.9244356 | 286.1693293 | 297.9867277 | 288.3664735 | 6.01E-07 | 0.829284 | 1.62E-07 | 5.23827 | 1.776803 | 6.221014 |
| CNAG_050 | - | zinc finge | 988.9203243 | 1050.365889 | 1084.444746 | 686.7356648 | 827.9517262 | 716.2667586 | 6.17E-07 | -0.50138 | 1.67E-07 | -5.23328 | 0.706432 | 6.209998 |
| CNAG_009 | - | hypothetic | 360.5130187 | 348.9685736 | 401.8325001 | 231.4670395 | 238.6709058 | 209.0079917 | 6.25E-07 | -0.72554 | 1.69E-07 | -5.23069 | 0.604769 | 6.204183 |
| CNAG_025 | - | spermine | 648.2189292 | 419.0923633 | 474.4566897 | 256.8973208 | 263.5180026 | 339.8709364 | 6.35E-07 | -0.86159 | 1.72E-07 | -5.22746 | 0.550345 | 6.196978 |
| CNAG_050 | CLR5 | hypothetic | 12.28852652 | 21.75500159 | 6.66381673 | 65.69399944 | 48.52845986 | 75.85006629 | 6.62E-07 | 2.257029 | 1.79E-07 | 5.219577 | 4.780061 | 6.179197 |
| CNAG_075 | - | hypothetic | 126.5408384 | 136.9939213 | 106.8685288 | 230.9679363 | 235.0143897 | 232.8300083 | 6.63E-07 | 0.906661 | 1.80E-07 | 5.21904 | 1.874702 | 6.178204 |
| CNAG_075 | - | sterol este | 23.99675717 | 32.56511256 | 31.14385572 | 72.72912438 | 106.969856 | 90.5975535 | 6.73E-07 | 1.625061 | 1.83E-07 | 5.216326 | 3.084552 | 6.172233 |
| CNAG_051 | - | small subu | 942.021346 | 864.0166888 | 911.70384 | 703.3658875 | 596.5898743 | 618.5738874 | 6.93E-07 | -0.51868 | 1.88E-07 | -5.21041 | 0.698012 | 6.158974 |
| CNAG_034 | LAC2 | Laccase 2 | 100.8902962 | 104.7107721 | 95.11736007 | 214.1137247 | 210.9700482 | 174.1479718 | 7.22E-07 | 0.984779 | 1.96E-07 | 5.202727 | 1.979011 | 6.141576 |
| CNAG_009 | - | hypothetic | 999.2362328 | 935.1074512 | 924.5346058 | 701.7009038 | 627.6359553 | 719.5193527 | 7.28E-07 | -0.49677 | 1.98E-07 | -5.20074 | 0.708689 | 6.137708 |
| CNAG_051 | - | hypothetic | 278.6939982 | 275.6487099 | 295.2003248 | 177.0413638 | 168.7660423 | 132.4858419 | 7.28E-07 | -0.84563 | 1.98E-07 | -5.20077 | 0.556467 | 6.137708 |
| CNAG_058 | - | hypothetic | 800.686956 | 770.2211095 | 712.3282495 | 551.4867272 | 553.569341 | 482.0743251 | 7.31E-07 | -0.54034 | 1.99E-07 | -5.19987 | 0.687608 | 6.135864 |
| CNAG_032 | - | solute car | 188.3247875 | 158.6077769 | 159.0938142 | 282.9092901 | 348.2332051 | 282.8986835 | 7.55E-07 | 0.836661 | 2.06E-07 | 5.193563 | 1.785912 | 6.122309 |
| CNAG_008 | CLR3 | hypothetic | 65.83143011 | 63.7824252 | 36.55273062 | 159.2144212 | 119.2144807 | 142.7830864 | 7.95E-07 | 1.342127 | 2.18E-07 | 5.183428 | 2.535248 | 6.099645 |
| CNAG_051 | UGT1 | UDP-galac | 598.2117863 | 660.5560864 | 589.8832405 | 905.6115846 | 883.5056191 | 832.5026632 | 9.05E-07 | 0.489539 | 2.50E-07 | 5.158 | 1.403996 | 6.043431 |
| CNAG_056 | - | hypothetic | 1035.601926 | 1022.404872 | 1130.293482 | 816.9477574 | 777.816693 | 740.6885297 | 9.05E-07 | -0.46545 | 2.50E-07 | -5.15801 | 0.724244 | 6.043431 |
| CNAG_052 | - | cytoplasm | 352.2910567 | 311.3156429 | 411.3641367 | 228.4057357 | 205.8054412 | 199.0168019 | 9.18E-07 | -0.78095 | 2.53E-07 | -5.15514 | 0.581983 | 6.037389 |
| CNAG_051 | - | DNA poly | 442.9901195 | 536.6434388 | 473.6390638 | 698.4746955 | 734.3392751 | 703.6598543 | 9.26E-07 | 0.542644 | 2.56E-07 | 5.153307 | 1.45664 | 6.033329 |
| CNAG_027 | - | elongatior | 42531.70758 | 39949.01753 | 45849.90638 | 32884.76416 | 36358.58482 | 33775.68125 | 9.33E-07 | -0.33282 | 2.58E-07 | -5.15191 | 0.793985 | 6.030292 |
| CNAG_052 | - | NADH del | 1522.591853 | 1556.797883 | 1631.55779 | 1276.213908 | 1182.370373 | 1119.249344 | 1.10E-06 | -0.41193 | 3.07E-07 | -5.11917 | 0.75162 | 5.958301 |
| CNAG_034 | - | hypothetic | 21.64117137 | 19.69502237 | 33.23403955 | 65.73092101 | 86.03067268 | 94.82135571 | 1.10E-06 | 1.730949 | 3.07E-07 | 5.119203 | 3.31946 | 5.958301 |
| CNAG_057 | - | mitogen-a | 991.2703107 | 1010.545069 | 1123.881784 | 795.9246254 | 753.6538515 | 734.017921 | 1.14E-06 | -0.46841 | 3.18E-07 | -5.11241 | 0.722761 | 5.943133 |
| CNAG_026 | - | hypothetic | 20.48573518 | 29.32321126 | 25.8176114 | 93.9450576 | 85.16035661 | 63.14143754 | 1.15E-06 | 1.688949 | 3.20E-07 | 5.111011 | 3.224218 | 5.940099 |
| CNAG_033 | - | hypothetic | 476.7803226 | 486.0673225 | 511.9966225 | 674.2407994 | 754.2135705 | 710.31706 | 1.15E-06 | 0.522173 | 3.21E-07 | 5.110627 | 1.436117 | 5.939406 |
| CNAG_035 | - | Phenylacr | 33.38354422 | 31.57253244 | 61.98548716 | 116.7334407 | 151.0867308 | 97.53938239 | 1.16E-06 | 1.525332 | 3.23E-07 | 5.109574 | 2.87853 | 5.937176 |
| CNAG_076 | SAT4 | HAL prote | 5234.829499 | 4861.344681 | 5259.663769 | 4113.000725 | 4200.533832 | 4173.212492 | 1.20E-06 | -0.31396 | 3.35E-07 | -5.10248 | 0.804432 | 5.921817 |
| CNAG_033 | - | AP-1 com | 2732.423252 | 2776.475765 | 2926.249766 | 2264.576954 | 2230.411805 | 2279.277533 | 1.24E-06 | -0.33164 | 3.49E-07 | -5.09482 | 0.79463 | 5.905581 |
| CNAG_033 | - | hypothetic | 1540.972735 | 1202.423576 | 1489.682695 | 987.2502515 | 977.8866062 | 1041.484064 | 1.27E-06 | -0.50997 | 3.57E-07 | -5.09036 | 0.702237 | 5.895728 |
| CNAG_058 | - | nuclear pc | 4202.465189 | 3904.554285 | 3846.655993 | 3210.977679 | 3120.559463 | 3277.299093 | 1.32E-06 | -0.33054 | 3.70E-07 | -5.08355 | 0.795237 | 5.880713 |
| CNAG_068 | PMT1 | dolichyl-p | 2255.161323 | 2366.117459 | 2262.037841 | 2814.456218 | 2896.622442 | 3043.883327 | 1.37E-06 | 0.331779 | 3.87E-07 | 5.075037 | 1.258564 | 5.86201 |
| CNAG_050 | ATG1 | ULK/ULK | 154.7594449 | 205.9599904 | 229.3847199 | 348.4930135 | 347.9791986 | 337.239406 | 1.39E-06 | 0.79697 | 3.93E-07 | 5.072308 | 1.737449 | 5.855967 |
| CNAG_032 | - | dioxygena | 49.56797696 | 44.44567996 | 51.38877402 | 139.7707622 | 126.499763 | 96.47321342 | 1.40E-06 | 1.310073 | 3.96E-07 | 5.070944 | 2.47954 | 5.853229 |
| CNAG_032 | - | coatomer | 9276.367574 | 8378.085507 | 8794.980936 | 7047.140348 | 7297.997848 | 7277.213454 | 1.46E-06 | -0.30647 | 4.13E-07 | -5.06308 | 0.808616 | 5.83586 |
| CNAG_055 | KIN4 | CAMK/CA | 315.5516836 | 283.5415275 | 274.2441339 | 476.1037446 | 478.2186858 | 422.8462511 | 1.48E-06 | 0.642409 | 4.18E-07 | 5.060593 | 1.560933 | 5.830942 |
| CNAG_050 | - | Cysteine s | 4619.227204 | 3884.116959 | 4106.848541 | 3062.955229 | 3413.030824 | 3254.025544 | 1.53E-06 | -0.39022 | 4.33E-07 | -5.054 | 0.763013 | 5.816485 |
| CNAG_033 | - | solute car | 2421.919053 | 2985.379691 | 2748.159557 | 2043.452777 | 2177.829972 | 2042.864108 | 1.62E-06 | -0.39532 | 4.60E-07 | -5.04244 | 0.760319 | 5.791344 |
| CNAG_037 | - | hypothetic | 2963.694905 | 3293.591335 | 3221.670415 | 2559.566825 | 2396.84045 | 2592.328002 | 1.62E-06 | -0.34322 | 4.62E-07 | -5.04143 | 0.788277 | 5.789787 |
| CNAG_055 | - | glycogen | 350.7387187 | 369.7236612 | 416.0003873 | 565.4314171 | 601.5546019 | 550.4801876 | 1.62E-06 | 0.58103 | 4.62E-07 | 5.04144 | 1.495917 | 5.789787 |
| CNAG_050 | - | NADH-ubi | 2438.195976 | 2774.241057 | 2573.284884 | 2131.542583 | 1981.097942 | 2026.187908 | 1.63E-06 | -0.35753 | 4.63E-07 | -5.04105 | 0.780497 | 5.789125 |
| CNAG_026 | - | pirin | 350.523866 | 284.6453082 | 289.1769209 | 470.4653765 | 511.8778615 | 478.2707081 | 1.64E-06 | 0.644551 | 4.68E-07 | 5.038857 | 1.563253 | 5.784327 |
| CNAG_052 | - | hypothetic | 1699.793548 | 1442.62651 | 1452.456931 | 1160.913248 | 1157.370199 | 1098.143868 | 1.68E-06 | -0.44349 | 4.80E-07 | -5.03418 | 0.735353 | 5.774103 |
| CNAG_105 | - | unspecifie | 30958.3911 | 23918.42878 | 12531.97345 | 6676.692076 | 4450.041858 | 6425.031121 | 1.79E-06 | -1.95658 | 5.11E-07 | -5.02203 | 0.257639 | 5.747512 |
| CNAG_049 | - | protein FF | 1253.840158 | 1122.633265 | 1214.587054 | 878.3373914 | 942.6397221 | 851.6939281 | 1.88E-06 | -0.44207 | 5.40E-07 | -5.01137 | 0.736078 | 5.724901 |
| CNAG_033 | - | YTH doma | 260.8285836 | 300.7236014 | 261.4466778 | 443.3453801 | 491.2941017 | 389.5833961 | 1.90E-06 | 0.673426 | 5.45E-07 | 5.009771 | 1.594856 | 5.721667 |
| CNAG_026 | - | hypothetic | 96.30735524 | 119.7650689 | 114.2743296 | 191.8450505 | 216.0776796 | 220.5440231 | 2.12E-06 | 0.915758 | 6.11E-07 | 4.987791 | 1.88656 | 5.673827 |
| CNAG_058 | UBA2 | ubiquitin- | 1717.554126 | 1612.873839 | 1802.185046 | 1285.394189 | 1335.93828 | 1344.534325 | 2.16E-06 | -0.38789 | 6.25E-07 | -4.98346 | 0.764245 | 5.664794 |
| CNAG_052 | - | hypothetic | 59.16507354 | 121.8949467 | 110.0094147 | 225.868978 | 203.957978 | 205.0590362 | 2.22E-06 | 1.113084 | 6.42E-07 | 4.978143 | 2.163076 | 5.653085 |
| CNAG_075 | - | hypothetic | 1007.502792 | 921.159417 | 1095.071848 | 656.365847 | 720.0125732 | 776.1171833 | 2.22E-06 | -0.50601 | 6.42E-07 | -4.97805 | 0.704166 | 5.653054 |
| CNAG_067 | - | Acyl-CoA- | 240.8835836 | 243.6446909 | 215.5993077 | 393.6552676 | 368.5829714 | 361.7219529 | 2.29E-06 | 0.669705 | 6.64E-07 | 4.971598 | 1.590748 | 5.639511 |
| CNAG_035 | - | hypothetic | 74.19703769 | 102.5365133 | 102.5344369 | 172.4309395 | 184.9680695 | 208.2338148 | 2.32E-06 | 1.006358 | 6.73E-07 | 4.9689 | 2.008834 | 5.63365 |
| CNAG_051 | - | membran | 4318.330315 | 5029.313825 | 4853.337233 | 3553.3098 | 3692.440768 | 3988.88158 | 2.57E-06 | -0.35266 | 7.48E-07 | -4.94837 | 0.783136 | 5.5905 |
| CNAG_076 | - | hypothetic | 310.9818674 | 345.9629672 | 281.727014 | 467.3025601 | 552.5394188 | 456.143913 | 2.60E-06 | 0.641185 | 7.58E-07 | 4.945753 | 1.559609 | 5.58503 |
| CNAG_075 | - | peptide cl | 1367.347533 | 1475.974555 | 1524.909733 | 1127.460988 | 1158.27594 | 1065.943063 | 2.63E-06 | -0.39727 | 7.67E-07 | -4.94369 | 0.759293 | 5.580617 |
| CNAG_031 | - | cytoplasm | 649.8954832 | 510.6750135 | 609.852545 | 344.9552837 | 413.8230085 | 400.9590556 | 2.72E-06 | -0.62812 | 7.96E-07 | -4.93629 | 0.647019 | 5.565049 |
| CNAG_051 | - | hypothetic | 13.72061151 | 39.17393402 | 56.81274115 | 208.1284663 | 274.8032636 | 213.6739662 | 2.84E-06 | 2.672351 | 8.31E-07 | 4.927869 | 6.37467 | 5.547396 |

| | | | | | | | | | | | | | |
|---|---|---|---|---|---|---|---|---|---|---|---|---|---|
| CNAG_034 | - | transcripti | 519.9126459 | 482.8525903 | 508.8132545 | 776.5074999 | 660.8032815 | 758.0003729 | 3.05E-06 | 0.524656 | 8.95E-07 | 4.913384 | 1.43859 | 5.516337 |
| CNAG_057 | - | hypothetic | 347.9963987 | 449.0702005 | 487.1059884 | 271.9520126 | 251.5207264 | 271.1039573 | 3.13E-06 | -0.70852 | 9.21E-07 | -4.90789 | 0.611946 | 5.50471 |
| CNAG_126 | - | unspecifie | 23.90486665 | 25.02813361 | 25.80194338 | 68.31222707 | 77.33664253 | 77.2641804 | 3.16E-06 | 1.589526 | 9.33E-07 | 4.905311 | 3.009505 | 5.499722 |
| CNAG_008 | PDR5 | ATP-bindi | 1391.302857 | 1171.044382 | 1014.194596 | 742.7166787 | 778.5948122 | 913.764621 | 3.16E-06 | -0.57122 | 9.33E-07 | -4.90536 | 0.673048 | 5.499722 |
| CNAG_105 | - | unspecifie | 433646.2318 | 319514.2973 | 204568.166 | 118586.4337 | 75544.08427 | 93314.00798 | 3.20E-06 | -1.75214 | 9.43E-07 | -4.90316 | 0.296861 | 5.495505 |
| CNAG_031 | - | hypothetic | 368.3271891 | 447.2602257 | 475.7141032 | 641.7255729 | 646.326917 | 635.9290813 | 3.54E-06 | 0.560505 | 1.05E-06 | 4.882005 | 1.474786 | 5.450598 |
| CNAG_032 | - | Mannosyl- | 2931.332203 | 3244.210026 | 3110.921096 | 3731.254262 | 3886.843421 | 4071.895663 | 3.67E-06 | 0.317289 | 1.09E-06 | 4.874749 | 1.245987 | 5.435339 |
| CNAG_077 | - | HMG box | 0.362084733 | 4.516377287 | 2.317120719 | 27.93803829 | 19.93687702 | 45.66873396 | 3.70E-06 | 3.911574 | 1.10E-06 | 4.87295 | 15.04877 | 5.431917 |
| CNAG_126 | - | unspecifie | 88.10665557 | 86.44419086 | 113.1644735 | 202.4235477 | 175.3747421 | 188.41437 | 3.93E-06 | 0.961783 | 1.17E-06 | 4.860615 | 1.947716 | 5.406072 |
| CNAG_031 | - | hypothetic | 712.9797033 | 606.5108132 | 611.0005094 | 462.760629 | 462.9963132 | 396.6052287 | 4.39E-06 | -0.56331 | 1.32E-06 | -4.83746 | 0.676747 | 5.357163 |
| CNAG_032 | KIC102 | ste/ste20/ | 853.8186517 | 771.5910129 | 821.2660313 | 1090.666141 | 1144.00854 | 1080.05942 | 4.39E-06 | 0.423016 | 1.31E-06 | 4.837564 | 1.340728 | 5.357163 |
| CNAG_027 | - | prenylcyst | 412.7083848 | 557.0957239 | 489.6242505 | 692.2029707 | 730.8663148 | 738.0220642 | 4.54E-06 | 0.55295 | 1.36E-06 | 4.830566 | 1.467083 | 5.343348 |
| CNAG_049 | PDR16 | pleiotropic | 2082.032202 | 2182.806072 | 2413.24238 | 1693.230712 | 1825.061722 | 1697.597247 | 4.60E-06 | -0.37186 | 1.38E-06 | -4.82769 | 0.772786 | 5.33743 |
| CNAG_009 | - | MFS trans | 73.01075918 | 100.3715814 | 95.08041239 | 166.2163415 | 169.3640388 | 205.9659259 | 4.61E-06 | 0.996626 | 1.38E-06 | 4.827202 | 1.995328 | 5.336541 |
| CNAG_034 | LAC1 | laccase | 393.7456051 | 360.0124852 | 293.4856349 | 492.0547176 | 548.0049267 | 620.1758553 | 4.80E-06 | 0.651078 | 1.45E-06 | 4.818534 | 1.570341 | 5.318719 |
| CNAG_075 | - | hypothetic | 827.6539507 | 806.8897054 | 799.7548553 | 598.9560726 | 557.8015272 | 626.2537518 | 4.93E-06 | -0.46463 | 1.49E-06 | -4.81283 | 0.724658 | 5.307019 |
| CNAG_034 | - | translation | 1043.769267 | 1089.152513 | 1027.992778 | 750.6391165 | 797.6782486 | 838.2791484 | 5.10E-06 | -0.42046 | 1.54E-06 | -4.80571 | 0.747184 | 5.29208 |
| CNAG_050 | - | deoxyhypu | 609.3880352 | 614.0473796 | 696.2157883 | 440.096994 | 446.5638957 | 466.4256929 | 5.17E-06 | -0.52058 | 1.56E-06 | -4.8032 | 0.697093 | 5.286794 |
| CNAG_025 | LPI3 | hypothetic | 665.3101258 | 668.9378418 | 649.3669972 | 433.8152222 | 401.5933178 | 524.0670479 | 5.17E-06 | -0.56034 | 1.56E-06 | -4.80286 | 0.67814 | 5.286247 |
| CNAG_068 | FRE2 | ferric redu | 602.2217008 | 609.6585984 | 551.2905749 | 376.182651 | 451.8539408 | 388.7846257 | 5.25E-06 | -0.55157 | 1.59E-06 | -4.79963 | 0.682278 | 5.280106 |
| CNAG_058 | - | TDG/mug | 263.0001747 | 235.0579011 | 219.8685062 | 360.6560881 | 370.2707806 | 427.0255959 | 5.39E-06 | 0.674981 | 1.63E-06 | 4.794153 | 1.596576 | 5.268766 |
| CNAG_032 | - | pre-mRNA | 4782.824466 | 4403.407687 | 4289.24407 | 3529.245808 | 3582.909475 | 3793.501105 | 5.79E-06 | -0.3208 | 1.77E-06 | -4.77843 | 0.800624 | 5.237028 |
| CNAG_037 | - | hypothetic | 71.81393001 | 78.89247585 | 101.4315616 | 158.4531511 | 160.7436132 | 193.7649725 | 6.81E-06 | 1.016362 | 2.09E-06 | 4.744222 | 2.022812 | 5.166865 |
| CNAG_078 | - | hypothetic | 1844.921258 | 1986.660731 | 1863.036493 | 1519.753745 | 1425.589146 | 1568.750061 | 7.34E-06 | -0.35018 | 2.26E-06 | -4.72826 | 0.784486 | 5.134563 |
| CNAG_058 | - | ubiquitin- | 1667.234719 | 1453.431325 | 1603.852034 | 1230.205213 | 1203.942673 | 1217.990525 | 7.48E-06 | -0.38764 | 2.31E-06 | -4.72402 | 0.764376 | 5.126032 |
| CNAG_124 | - | unspecifie | 105.6137169 | 109.5585182 | 87.33424589 | 46.10839677 | 40.10364327 | 40.30825016 | 7.53E-06 | -1.27117 | 2.33E-06 | -4.72241 | 0.414324 | 5.123172 |
| CNAG_127 | - | unspecifie | 26.43989074 | 61.57051519 | 50.31943364 | 103.1746419 | 119.5445807 | 133.7541151 | 7.87E-06 | 1.360917 | 2.44E-06 | 4.713124 | 2.568484 | 5.103964 |
| CNAG_031 | - | THO comp | 1967.519142 | 1998.557423 | 2073.088727 | 1560.170603 | 1487.679652 | 1706.381928 | 8.17E-06 | -0.36028 | 2.54E-06 | -4.70515 | 0.779011 | 5.087993 |
| CNAG_037 | NTH1 | alpha,alph | 1750.575448 | 1633.486862 | 1479.306668 | 2103.205428 | 2146.248792 | 2081.433459 | 8.93E-06 | 0.365217 | 2.79E-06 | 4.686059 | 1.288076 | 5.049309 |
| CNAG_032 | - | hypothetic | 133.5360834 | 155.2672355 | 106.8804919 | 229.2850539 | 224.5213802 | 263.7880925 | 1.03E-05 | 0.847597 | 3.21E-06 | 4.656681 | 1.799501 | 4.988711 |
| CNAG_050 | - | hypothetic | 33.44327257 | 56.26833818 | 76.88992129 | 135.4581427 | 123.6945868 | 141.6362429 | 1.06E-05 | 1.259756 | 3.34E-06 | 4.648707 | 2.394552 | 4.972927 |
| CNAG_026 | - | hypothetic | 2017.699623 | 2028.719135 | 2024.056339 | 1465.877229 | 1700.876251 | 1639.818761 | 1.07E-05 | -0.35255 | 3.35E-06 | -4.64832 | 0.7832 | 4.97229 |
| CNAG_033 | PCL103 | g1/s-speci | 128.9105447 | 163.8644098 | 103.6929387 | 271.7697097 | 240.1358101 | 219.6523766 | 1.11E-05 | 0.8757 | 3.51E-06 | 4.638529 | 1.834898 | 4.952871 |
| CNAG_032 | ARF2 | ADP-ribos | 12521.60625 | 11799.9989 | 12050.87269 | 10488.87513 | 10368.74348 | 9503.142766 | 1.14E-05 | -0.27626 | 3.61E-06 | -4.63273 | 0.825728 | 4.941678 |
| CNAG_057 | - | cytoplasm | 252.6464143 | 272.7425288 | 285.9179805 | 424.6534894 | 456.7252863 | 372.9368386 | 1.22E-05 | 0.614718 | 3.88E-06 | 4.617878 | 1.531259 | 4.912262 |
| CNAG_032 | MSL1 | histone ac | 1547.069461 | 1506.180651 | 1544.145817 | 1259.080452 | 1148.707852 | 1219.083139 | 1.22E-05 | -0.35758 | 3.88E-06 | -4.61793 | 0.780472 | 4.912262 |
| CNAG_031 | CSN4 | COP9 sigr | 335.4475027 | 338.4486684 | 328.5972989 | 522.0137624 | 492.7701226 | 460.6039646 | 1.26E-05 | 0.543073 | 4.00E-06 | 4.61145 | 1.457073 | 4.899823 |
| CNAG_057 | - | GTPase in | 1610.207932 | 1776.541034 | 1615.625955 | 1342.321642 | 1366.175685 | 1221.37411 | 1.30E-05 | -0.36295 | 4.15E-06 | -4.60397 | 0.777571 | 4.88537 |
| CNAG_057 | - | hypothetic | 414.1415435 | 413.5457469 | 367.856236 | 293.1759529 | 255.0294353 | 228.9962571 | 1.32E-05 | -0.63685 | 4.19E-06 | -4.60157 | 0.643116 | 4.880703 |
| CNAG_078 | - | centromer | 3163.602808 | 2970.596718 | 3183.423954 | 3721.920413 | 3957.609822 | 3877.674518 | 1.32E-05 | 0.295116 | 4.22E-06 | 4.600303 | 1.226983 | 4.878224 |
| CNAG_027 | - | small subu | 350.9773946 | 311.2545868 | 317.6814324 | 229.3093826 | 207.6024749 | 171.306332 | 1.32E-05 | -0.70395 | 4.22E-06 | -4.60007 | 0.613888 | 4.877907 |
| CNAG_074 | - | AP-2 com | 832.3148953 | 793.9754283 | 790.1663782 | 591.1496033 | 587.1795641 | 625.1457806 | 1.33E-05 | -0.43695 | 4.24E-06 | -4.59918 | 0.738692 | 4.876386 |
| CNAG_078 | - | hypothetic | 173.2708754 | 218.8032903 | 165.492947 | 297.7294513 | 318.6620243 | 302.8430058 | 1.44E-05 | 0.707534 | 4.62E-06 | 4.581392 | 1.633011 | 4.841029 |
| CNAG_032 | - | hypothetic | 45.06013263 | 89.51490891 | 52.50154796 | 140.7968538 | 158.5047936 | 128.5512523 | 1.46E-05 | 1.182697 | 4.68E-06 | 4.578553 | 2.270008 | 4.835837 |
| CNAG_126 | - | unspecifie | 2.823352522 | 8.806821958 | 4.461829755 | 36.01581209 | 39.57089628 | 24.30677055 | 1.51E-05 | 2.808981 | 4.85E-06 | 4.571336 | 7.007892 | 4.82198 |
| CNAG_034 | TSA1 | Thiol pero | 11796.45518 | 10819.56702 | 12098.82293 | 9784.478814 | 9450.061724 | 9670.716723 | 1.51E-05 | -0.27991 | 4.86E-06 | -4.57067 | 0.823642 | 4.82076 |
| CNAG_058 | KRE64 | glucosidas | 138.252113 | 171.419067 | 135.6279185 | 247.1762431 | 270.4207463 | 242.9217187 | 1.56E-05 | 0.75904 | 5.04E-06 | 4.563161 | 1.692365 | 4.806364 |
| CNAG_003 | - | DNA topo | 5297.782521 | 4690.029232 | 4740.348399 | 4087.382448 | 3900.323321 | 4024.447008 | 1.58E-05 | -0.3098 | 5.10E-06 | -4.56054 | 0.806751 | 4.801097 |
| CNAG_032 | - | hypothetic | 21.66320989 | 38.95882596 | 29.00973988 | 83.31317025 | 67.47113767 | 97.07453242 | 1.66E-05 | 1.475957 | 5.38E-06 | 4.549248 | 2.78168 | 4.778598 |
| CNAG_034 | - | hypothetic | 1652.510797 | 1596.843035 | 1480.353189 | 2051.098139 | 2022.871426 | 1952.684392 | 1.71E-05 | 0.334413 | 5.54E-06 | 4.543317 | 1.260865 | 4.767023 |
| CNAG_035 | - | mitochonc | 1551.453378 | 1408.075007 | 1240.268413 | 1144.748546 | 1011.567947 | 839.5949106 | 1.82E-05 | -0.50362 | 5.90E-06 | -4.53 | 0.705332 | 4.74059 |
| CNAG_075 | - | proteasom | 825.798761 | 830.7879566 | 747.7131997 | 1043.15051 | 1060.262754 | 1090.002098 | 1.85E-05 | 0.395602 | 5.99E-06 | 4.526679 | 1.315491 | 4.733932 |
| CNAG_052 | - | hypothetic | 10.02512056 | 32.47128704 | 19.42507707 | 66.92625758 | 61.59229336 | 79.32142398 | 1.99E-05 | 1.744209 | 6.49E-06 | 4.509845 | 3.350111 | 4.701529 |
| CNAG_057 | - | Swi5-depe | 28.63763174 | 30.44114825 | 47.0535332 | 80.61576122 | 101.5810718 | 91.7644827 | 2.10E-05 | 1.357517 | 6.89E-06 | 4.497068 | 2.562438 | 4.677027 |
| CNAG_037 | - | hypothetic | 1199.085582 | 1108.653418 | 1326.508423 | 881.4219261 | 929.6621451 | 947.0935863 | 2.15E-05 | -0.4138 | 7.04E-06 | -4.49244 | 0.750646 | 4.668222 |
| CNAG_036 | - | hypothetic | 1481.969543 | 1323.20433 | 983.4974246 | 2061.635956 | 1826.271125 | 1659.627266 | 2.24E-05 | 0.534937 | 7.35E-06 | 4.48327 | 1.448878 | 4.650188 |
| CNAG_052 | - | hypothetic | 605.6668441 | 493.4585968 | 639.6668141 | 414.5072415 | 414.6932207 | 346.6634554 | 2.25E-05 | -0.5816 | 7.41E-06 | -4.48151 | 0.668221 | 4.646932 |
| CNAG_052 | - | bud site s | 1670.895702 | 1713.022848 | 1688.125544 | 1382.035845 | 1412.772942 | 1226.935908 | 2.27E-05 | -0.35038 | 7.46E-06 | -4.48002 | 0.784378 | 4.644217 |
| CNAG_079 | - | hypothetic | 42.45754832 | 33.65750937 | 37.52384812 | 81.37486223 | 95.39793668 | 97.25928747 | 2.34E-05 | 1.26946 | 7.72E-06 | 4.472668 | 2.410713 | 4.629915 |
| CNAG_052 | - | tRNA pseu | 2088.84528 | 1859.627172 | 2116.794884 | 1433.947703 | 1579.178579 | 1691.956209 | 2.35E-05 | -0.38224 | 7.74E-06 | -4.47229 | 0.767244 | 4.629309 |
| CNAG_123 | - | unspecifie | 14.63387329 | 28.18773556 | 20.47433298 | 62.99394697 | 64.31170241 | 66.19380905 | 2.46E-05 | 1.617505 | 8.12E-06 | 4.461946 | 3.068438 | 4.609281 |
| CNAG_078 | - | class III ar | 29.68419399 | 27.18342859 | 22.63628057 | 98.80490239 | 88.69595326 | 48.91884971 | 2.60E-05 | 1.569161 | 8.61E-06 | 4.449498 | 2.967321 | 4.584719 |
| CNAG_068 | CAP1alpha | O-glucosy | 2342.411791 | 2397.221839 | 2487.951608 | 1971.951325 | 2045.029659 | 1920.72018 | 2.63E-05 | -0.299 | 8.70E-06 | -4.4471 | 0.812815 | 4.580193 |
| CNAG_031 | SET202 | histone-ly | 2326.123578 | 2639.581968 | 2494.369074 | 1947.751325 | 2050.175033 | 2070.547028 | 2.65E-05 | -0.31271 | 8.78E-06 | -4.44523 | 0.805129 | 4.576567 |
| CNAG_051 | - | transcripti | 900.246256 | 1006.182129 | 973.5781398 | 772.6383098 | 726.9434316 | 699.5904747 | 2.76E-05 | -0.40487 | 9.14E-06 | -4.43648 | 0.755306 | 4.559398 |
| CNAG_075 | - | hypothetic | 269.0882392 | 305.0894895 | 364.7910453 | 533.8286677 | 529.1669023 | 412.9626882 | 2.79E-05 | 0.636734 | 9.26E-06 | 4.433665 | 1.554805 | 4.553878 |
| CNAG_032 | - | AMME syr | 44.98037333 | 74.44909305 | 42.90845934 | 110.9625332 | 143.9210005 | 115.1933981 | 2.82E-05 | 1.188215 | 9.35E-06 | 4.431668 | 2.278707 | 4.550334 |
| CNAG_056 | - | hypothetic | 807.9165034 | 797.2337606 | 893.5417596 | 658.9934368 | 613.065412 | 596.351812 | 2.85E-05 | -0.43489 | 9.47E-06 | -4.42894 | 0.739748 | 4.545312 |

| | | | | | | | | | | | | | | |
|---|---|---|---|---|---|---|---|---|---|---|---|---|---|---|
| CNAG_05 | - | hypothetic | 2300.240261 | 2159.123158 | 2176.58137 | 1758.651355 | 1821.5933 | 1836.30731 | 2.90E-05 | -0.30863 | 9.64E-06 | -4.42521 | 0.807409 | 4.538124 |
| CNAG_03 | - | 4'-phosph | 417.6633006 | 423.2367832 | 363.6109661 | 293.9277032 | 240.2991615 | 268.8632624 | 2.90E-05 | -0.60161 | 9.66E-06 | -4.42469 | 0.659018 | 4.537238 |
| CNAG_02 | - | hypothetic | 564.3342124 | 573.3282088 | 628.2081194 | 777.7506588 | 790.2786902 | 841.2817461 | 3.01E-05 | 0.432949 | 1.00E-05 | 4.41614 | 1.34999 | 4.521174 |
| CNAG_05 | - | hypothetic | 592.8520732 | 537.558847 | 554.463985 | 418.427279 | 412.1064235 | 378.7905772 | 3.18E-05 | -0.4941 | 1.07E-05 | -4.40333 | 0.710005 | 4.497065 |
| CNAG_06 | FAO1 | putative fa | 30.96146775 | 45.43001881 | 38.58839779 | 109.3874953 | 84.83519844 | 84.14143013 | 3.22E-05 | 1.279639 | 1.08E-05 | 4.400713 | 2.427782 | 4.492148 |
| CNAG_06 | CSR3 | chitin synt | 62.23375364 | 54.08243136 | 40.77148093 | 110.3097338 | 108.1414604 | 127.1933391 | 3.29E-05 | 1.129847 | 1.10E-05 | 4.395948 | 2.188355 | 4.483091 |
| CNAG_06 | - | hypothetic | 97.53140595 | 146.6447856 | 133.4457182 | 231.8159141 | 222.8830854 | 214.0416604 | 3.38E-05 | 0.81296 | 1.14E-05 | 4.389375 | 1.756812 | 4.470746 |
| CNAG_00 | - | glucan 1,3 | 144.018401 | 164.9563161 | 121.7872572 | 245.6947434 | 265.2722604 | 223.0022171 | 3.56E-05 | 0.758096 | 1.20E-05 | 4.377796 | 1.691257 | 4.448596 |
| CNAG_05 | - | ubiquinol- | 1195.590747 | 1150.644127 | 1291.339559 | 1021.785601 | 856.2561194 | 886.0943423 | 3.62E-05 | -0.41153 | 1.22E-05 | -4.37402 | 0.751823 | 4.441241 |
| CNAG_05 | APT4 | Aminopho | 2132.556753 | 2313.305265 | 2214.032269 | 2629.177054 | 2768.969328 | 2834.092591 | 3.83E-05 | 0.290879 | 1.30E-05 | 4.360739 | 1.223386 | 4.41625 |
| CNAG_03 | - | hypothetic | 821.7222574 | 754.1093038 | 719.8099811 | 545.9650793 | 557.8546055 | 602.9269829 | 3.94E-05 | -0.44435 | 1.33E-05 | -4.35465 | 0.734913 | 4.404642 |
| CNAG_07 | - | hypothetic | 1121.908109 | 1062.258944 | 1061.045221 | 862.1411128 | 871.935997 | 688.5844767 | 3.98E-05 | -0.43788 | 1.35E-05 | -4.35202 | 0.738216 | 4.399735 |
| CNAG_12 | - | unspecifie | 1523.82079 | 1553.597102 | 1737.11308 | 1283.942555 | 1162.477371 | 1316.744313 | 4.02E-05 | -0.37096 | 1.36E-05 | -4.3497 | 0.77327 | 4.39609 |
| CNAG_05 | - | hypothetic | 9.860865112 | 11.00678831 | 11.92303761 | 58.41242018 | 29.36081788 | 44.02037777 | 4.14E-05 | 2.065079 | 1.41E-05 | 4.342663 | 4.184569 | 4.382785 |
| CNAG_04 | NDX4 | 7,8-dihydr | 672.8188962 | 664.8727607 | 605.8757157 | 889.1705223 | 886.0693857 | 830.3067738 | 4.15E-05 | 0.409235 | 1.41E-05 | 4.342185 | 1.327982 | 4.381995 |
| CNAG_05 | - | lipoyl synt | 802.2563981 | 934.0118296 | 1028.941216 | 640.0982835 | 702.8161514 | 696.2163205 | 4.25E-05 | -0.45449 | 1.45E-05 | -4.33649 | 0.729766 | 4.371372 |
| CNAG_05 | - | hypothetic | 77.54676141 | 98.16570207 | 59.97555769 | 191.119863 | 140.8036831 | 146.3315994 | 4.31E-05 | 1.013708 | 1.47E-05 | 4.33332 | 2.019093 | 4.36573 |
| CNAG_03 | - | hypothetic | 528.8207219 | 533.2428952 | 600.2446806 | 414.5466482 | 408.66375 | 364.3757126 | 4.39E-05 | -0.5014 | 1.50E-05 | -4.32896 | 0.706419 | 4.357748 |
| CNAG_03 | - | ATP-depe | 2759.189958 | 2623.495669 | 2626.628927 | 2144.691219 | 2195.936287 | 2281.459383 | 4.39E-05 | -0.29001 | 1.50E-05 | -4.32862 | 0.817896 | 4.357244 |
| CNAG_04 | - | pre-mRNA | 1111.92453 | 1162.624403 | 1094.278629 | 1355.114116 | 1485.540854 | 1468.589284 | 4.59E-05 | 0.340492 | 1.57E-05 | 4.31862 | 1.266188 | 4.33816 |
| CNAG_03 | - | lipoyl(octa | 532.2757898 | 527.8479244 | 558.6968323 | 400.5111277 | 393.1171617 | 383.1949327 | 4.70E-05 | -0.47478 | 1.61E-05 | -4.31291 | 0.719576 | 4.327865 |
| CNAG_02 | - | hypothetic | 2364.457435 | 2258.228213 | 2152.07734 | 1847.425336 | 1837.965482 | 1871.836098 | 4.73E-05 | -0.30156 | 1.62E-05 | -4.31161 | 0.811375 | 4.325466 |
| CNAG_02 | - | oxidoredu | 1106.372253 | 796.2285473 | 892.5469878 | 656.4933951 | 619.8871032 | 703.9720348 | 4.97E-05 | -0.51475 | 1.71E-05 | -4.29998 | 0.699913 | 4.304033 |
| CNAG_00 | - | glutathion | 112.4947919 | 127.2586131 | 90.87272161 | 209.3013927 | 208.3137141 | 175.2796729 | 5.05E-05 | 0.832824 | 1.74E-05 | 4.29571 | 1.781168 | 4.29644 |
| CNAG_05 | - | hypothetic | 16.99023232 | 34.63444327 | 25.79760491 | 64.32020834 | 81.77975386 | 68.51495751 | 5.08E-05 | 1.492514 | 1.75E-05 | 4.294314 | 2.813789 | 4.29386 |
| CNAG_07 | TAF9 | transcripti | 741.4124833 | 788.5766559 | 827.4405511 | 573.2309189 | 634.7295927 | 570.8091931 | 5.51E-05 | -0.42214 | 1.90E-05 | -4.27582 | 0.746318 | 4.258963 |
| CNAG_03 | GAT203 | hypothetic | 1265.641462 | 1198.058699 | 1287.11623 | 982.7001471 | 976.2379247 | 1017.015009 | 5.52E-05 | -0.34947 | 1.91E-05 | -4.27535 | 0.784875 | 4.258203 |
| CNAG_05 | - | exocyst cc | 476.7222628 | 506.4858275 | 470.4232315 | 639.9847284 | 654.8620201 | 696.956924 | 5.52E-05 | 0.439103 | 1.91E-05 | 4.275002 | 1.355762 | 4.257681 |
| CNAG_03 | - | hypothetic | 1047.775323 | 1182.006892 | 1058.03642 | 1571.863864 | 1545.98767 | 1270.041409 | 5.67E-05 | 0.401249 | 1.97E-05 | 4.268803 | 1.320651 | 4.24667 |
| CNAG_03 | ROM21 | Predicted | 929.8969041 | 1101.203059 | 1082.5176 | 1283.538155 | 1385.476849 | 1407.50021 | 5.73E-05 | 0.374312 | 1.99E-05 | 4.266399 | 1.296221 | 4.242144 |
| CNAG_05 | - | solute car | 1120.993556 | 1205.56376 | 1316.928239 | 918.0474887 | 983.1894698 | 936.0150252 | 5.83E-05 | -0.37516 | 2.02E-05 | -4.26219 | 0.771025 | 4.234421 |
| CNAG_05 | - | GPI inosito | 1028.016356 | 1305.883185 | 1182.784241 | 1521.028233 | 1620.147203 | 1485.301274 | 5.84E-05 | 0.381846 | 2.03E-05 | 4.261438 | 1.303008 | 4.23326 |
| CNAG_05 | - | DNA excis | 1963.141764 | 1959.920116 | 1806.677324 | 2292.980163 | 2375.631883 | 2455.522951 | 5.84E-05 | 0.299124 | 2.03E-05 | 4.261481 | 1.230397 | 4.23326 |
| CNAG_03 | - | sulfite red | 1934.312948 | 1400.667039 | 1511.116392 | 1150.700377 | 1282.507102 | 1108.165116 | 5.90E-05 | -0.46962 | 2.05E-05 | -4.259 | 0.722156 | 4.228825 |
| CNAG_05 | - | glycosyl tr | 0.367896414 | 9.843985308 | 2.319838834 | 29.56229835 | 26.20522075 | 34.9771185 | 5.94E-05 | 3.007098 | 2.07E-05 | 4.257334 | 8.039456 | 4.225898 |
| CNAG_05 | - | anaphase- | 399.3117929 | 519.0519772 | 444.6104739 | 328.9145665 | 322.3896522 | 276.7100569 | 6.14E-05 | -0.57092 | 2.14E-05 | -4.2497 | 0.673186 | 4.21199 |
| CNAG_03 | - | polyamine | 1148.122864 | 1186.334778 | 1116.679546 | 1464.968079 | 1626.240805 | 1373.234063 | 6.22E-05 | 0.356116 | 2.17E-05 | 4.246625 | 1.279976 | 4.20619 |
| CNAG_02 | - | hypothetic | 2099.39887 | 2117.04574 | 1894.009001 | 1638.822196 | 1601.590423 | 1715.28294 | 6.26E-05 | -0.31757 | 2.19E-05 | -4.24507 | 0.802423 | 4.203334 |
| CNAG_03 | - | sterol 3be | 1641.971662 | 1541.899814 | 1452.621798 | 1985.704958 | 1908.100327 | 1950.433472 | 6.27E-05 | 0.319088 | 2.19E-05 | 4.24478 | 1.247542 | 4.202919 |
| CNAG_05 | - | DNA-direc | 1365.027065 | 1411.373956 | 1579.27541 | 1214.758686 | 1103.872726 | 1098.118283 | 6.48E-05 | -0.36603 | 2.26E-05 | -4.23728 | 0.775917 | 4.188713 |
| CNAG_03 | - | RAN prote | 2963.602405 | 2870.236713 | 2970.010579 | 2403.107651 | 2533.248755 | 2441.354723 | 6.54E-05 | -0.27043 | 2.29E-05 | -4.23452 | 0.82907 | 4.184134 |
| CNAG_02 | - | DNA poly | 1653.430529 | 1773.343347 | 1663.620623 | 1399.148677 | 1378.22026 | 1384.488559 | 6.56E-05 | -0.30536 | 2.30E-05 | -4.23391 | 0.809243 | 4.183113 |
| CNAG_05 | - | nuclear pr | 2809.397186 | 2703.212881 | 2604.250145 | 2206.979889 | 2207.138059 | 2320.313067 | 6.65E-05 | -0.28481 | 2.33E-05 | -4.23073 | 0.820852 | 4.177276 |
| CNAG_00 | ISP7 | Identified | 96.2572084 | 122.9531715 | 109.9983043 | 186.4812423 | 190.955196 | 193.9951981 | 6.95E-05 | 0.7847 | 2.44E-05 | 4.22021 | 1.722735 | 4.158034 |
| CNAG_05 | - | phosphati | 317.9466712 | 349.1809621 | 310.4749412 | 439.1649024 | 481.5847162 | 482.6966814 | 7.11E-05 | 0.507695 | 2.50E-05 | 4.214821 | 1.421777 | 4.148263 |
| CNAG_06 | - | hypothetic | 40.18220526 | 43.29710208 | 31.16897813 | 129.3965813 | 83.05805222 | 72.10497428 | 7.14E-05 | 1.302194 | 2.51E-05 | 4.213615 | 2.466036 | 4.146092 |
| CNAG_05 | - | hypothetic | 368.0428573 | 277.1374328 | 355.2269958 | 513.3715644 | 543.8119282 | 450.6526358 | 7.46E-05 | 0.574079 | 2.63E-05 | 4.203309 | 1.488726 | 4.127043 |
| CNAG_03 | - | hypothetic | 2.803033815 | 8.789909101 | 4.451917976 | 25.33082764 | 42.38935163 | 23.17753333 | 7.82E-05 | 2.671278 | 2.76E-05 | 4.192772 | 6.369931 | 4.10699 |
| CNAG_05 | GCS1 | ceramide | 1113.950009 | 1119.408765 | 1372.323382 | 808.1465513 | 945.2229362 | 945.971742 | 8.05E-05 | -0.43326 | 2.84E-05 | -4.18582 | 0.740587 | 4.094446 |
| CNAG_03 | - | hypothetic | 712.6678329 | 774.7818928 | 840.41246 | 999.5525112 | 1043.044982 | 1022.317144 | 8.23E-05 | 0.382461 | 2.91E-05 | 4.180316 | 1.303564 | 4.08437 |
| CNAG_03 | - | hypothetic | 617.4714221 | 575.2862524 | 619.4999762 | 451.0911711 | 449.1933382 | 449.7748532 | 8.70E-05 | -0.44044 | 3.09E-05 | -4.16688 | 0.736907 | 4.060388 |
| CNAG_07 | - | glucose-6 | 1814.353376 | 1635.482756 | 1622.030359 | 1438.163971 | 1187.460508 | 1341.200047 | 9.52E-05 | -0.37001 | 3.39E-05 | -4.14544 | 0.773778 | 4.021302 |
| CNAG_05 | - | hypothetic | 1234.384242 | 1020.482236 | 1066.56353 | 1496.236017 | 1413.007935 | 1438.639839 | 9.52E-05 | 0.372895 | 3.39E-05 | 4.145363 | 1.294949 | 4.021302 |
| CNAG_05 | - | endoplasn | 7.492442848 | 9.900330687 | 8.717574623 | 41.5362632 | 41.18105524 | 25.4749891 | 0.000102 | 2.141393 | 3.64E-05 | 4.129209 | 4.411877 | 3.992223 |
| CNAG_05 | - | myosin-1 | 2791.065937 | 2710.903476 | 2588.379444 | 3081.276994 | 3564.34253 | 3385.865973 | 0.000103 | 0.294855 | 3.67E-05 | 4.127339 | 1.226762 | 3.988841 |
| CNAG_05 | - | hypothetic | 16.93748015 | 22.85244995 | 28.93825683 | 55.70045664 | 60.77217383 | 72.72909504 | 0.000104 | 1.487918 | 3.74E-05 | 4.123173 | 2.80484 | 3.981424 |
| CNAG_03 | - | hypothetic | 176.8516504 | 233.9098265 | 231.5173049 | 322.5769464 | 318.4907086 | 354.9422244 | 0.000109 | 0.620904 | 3.90E-05 | 4.113238 | 1.537839 | 3.964034 |
| CNAG_02 | - | hypothetic | 54.27087686 | 62.70317612 | 76.89594038 | 138.5984773 | 123.6723448 | 119.6790767 | 0.000112 | 0.969821 | 4.05E-05 | 4.104809 | 1.958598 | 3.948924 |
| CNAG_05 | LPI11 | glycosyl h | 3700.420055 | 3789.222303 | 3323.071296 | 3007.877241 | 3020.560819 | 3008.642743 | 0.000112 | -0.27403 | 4.05E-05 | -4.10482 | 0.827005 | 3.948924 |
| CNAG_03 | - | dTDP-4-de | 8094.621288 | 7159.533661 | 7724.263939 | 6588.842497 | 6326.735218 | 6447.917418 | 0.000113 | -0.26284 | 4.07E-05 | -4.10334 | 0.833445 | 3.946455 |
| CNAG_02 | - | hypothetic | 372.8765532 | 417.0715659 | 425.5945444 | 550.5351473 | 553.0847249 | 579.2976878 | 0.000115 | 0.455552 | 4.13E-05 | 4.099947 | 1.371308 | 3.940384 |
| CNAG_05 | BUD16 | pyridoxal | 512.3985869 | 551.4634238 | 475.5957177 | 349.0047019 | 391.4433351 | 375.397967 | 0.000115 | -0.47891 | 4.16E-05 | -4.09842 | 0.717519 | 3.937662 |
| CNAG_02 | - | 3-hydroxy | 293.4647149 | 339.4742972 | 300.8710653 | 475.2853861 | 459.1507022 | 410.6838102 | 0.000124 | 0.513378 | 4.47E-05 | 4.081568 | 1.427389 | 3.90802 |
| CNAG_05 | - | hypothetic | 2911.467418 | 3154.790888 | 3247.392248 | 3911.977351 | 3716.012854 | 3687.884091 | 0.000126 | 0.26597 | 4.58E-05 | 4.076199 | 1.202444 | 3.898699 |
| CNAG_03 | - | mRNA 3'- | 1356.72093 | 1299.325405 | 1321.273602 | 1033.25861 | 1081.510766 | 1100.266005 | 0.000128 | -0.32236 | 4.63E-05 | -4.07354 | 0.799763 | 3.894202 |
| CNAG_12 | - | unspecifie | 54.40960613 | 113.179816 | 72.73121451 | 146.7157456 | 181.7564365 | 156.258653 | 0.000128 | 1.002343 | 4.67E-05 | 4.071759 | 2.003251 | 3.89117 |
| CNAG_05 | - | hypothetic | 859.0148897 | 788.5557153 | 689.9850725 | 644.3135611 | 535.3766993 | 524.2310949 | 0.000132 | -0.47134 | 4.82E-05 | -4.06432 | 0.721296 | 3.878337 |
| CNAG_05 | - | hypothetic | 1472.208219 | 1298.266919 | 1328.750734 | 1138.468003 | 1084.075699 | 1030.405206 | 0.000138 | -0.34937 | 5.03E-05 | -4.05409 | 0.784928 | 3.860906 |

| | | | | | | | | | | | | | | |
|---|---|---|---|---|---|---|---|---|---|---|---|---|---|---|
| CNAG_025 | - | hypothetic | 630.4923642 | 710.9564983 | 677.1086068 | 525.0924168 | 524.278317 | 480.9000505 | 0.000156 | -0.41449 | 5.72E-05 | -4.02412 | 0.750283 | 3.807776 |
| CNAG_037 | - | peroxin-5, | 274.6807248 | 236.1551183 | 279.5048745 | 406.7742423 | 413.508971 | 351.8519272 | 0.000159 | 0.552379 | 5.86E-05 | 4.018373 | 1.466502 | 3.797572 |
| CNAG_075 | - | glutamate | 1407.972185 | 1232.545691 | 1286.080399 | 934.2921214 | 1034.049698 | 1103.566685 | 0.000163 | -0.36973 | 6.01E-05 | -4.01246 | 0.773926 | 3.786962 |
| CNAG_035 | - | hypothetic | 263.5657181 | 223.0068935 | 300.4893891 | 171.5930922 | 167.0594379 | 162.3246305 | 0.000163 | -0.66871 | 6.02E-05 | -4.01217 | 0.62907 | 3.786575 |
| CNAG_027 | - | hypothetic | 104.5496746 | 177.8243705 | 136.6683826 | 247.3432554 | 227.9666369 | 239.53067 | 0.000165 | 0.760661 | 6.08E-05 | 4.009635 | 1.694266 | 3.78235 |
| CNAG_055 | - | hypothetic | 254.9445287 | 258.749461 | 302.9380863 | 370.6692158 | 398.741908 | 427.1281555 | 0.000173 | 0.538614 | 6.36E-05 | 3.99886 | 1.452576 | 3.763132 |
| CNAG_123 | - | unspecifie | 13.45010345 | 32.42041209 | 9.857866836 | 54.25473779 | 50.26873988 | 76.97034709 | 0.000175 | 1.726299 | 6.45E-05 | 3.995626 | 3.308778 | 3.757489 |
| CNAG_026 | - | pre-mRNA | 9708.512028 | 9115.049176 | 9348.487259 | 7676.001215 | 8004.445261 | 8441.70788 | 0.000191 | -0.23941 | 7.08E-05 | -3.97355 | 0.847092 | 3.719552 |
| CNAG_075 | - | DNA ligas | 926.1842848 | 877.1373605 | 876.7170769 | 1121.726422 | 1111.958098 | 1183.259652 | 0.000193 | 0.334849 | 7.16E-05 | 3.970942 | 1.261246 | 3.715224 |
| CNAG_067 | - | hypothetic | 104.2753069 | 97.15964898 | 101.4609094 | 157.490155 | 184.1449261 | 177.3435885 | 0.000203 | 0.762854 | 7.56E-05 | 3.958035 | 1.696844 | 3.693295 |
| CNAG_131 | - | unspecifie | 0.271293397 | 0.130475854 | 2.27140743 | 16.80656743 | 31.26487312 | 11.09902424 | 0.000214 | 4.840533 | 7.99E-05 | 3.94475 | 28.65138 | 3.670461 |
| CNAG_034 | - | DNA repa | 370.166777 | 458.7964184 | 463.7351817 | 269.529176 | 290.4176365 | 335.3596314 | 0.000216 | -0.54455 | 8.09E-05 | -3.9416 | 0.685607 | 3.665044 |
| CNAG_027 | CLC-A | putative v | 2323.818428 | 2580.359384 | 2585.003268 | 2048.991226 | 2115.747341 | 2103.8527 | 0.000216 | -0.27192 | 8.10E-05 | -3.94149 | 0.828217 | 3.66498 |
| CNAG_051 | - | hypothetic | 612.1534272 | 651.9231877 | 597.3259263 | 890.091872 | 772.0773042 | 798.0980504 | 0.000221 | 0.388512 | 8.27E-05 | 3.936481 | 1.309042 | 3.656338 |
| CNAG_057 | - | hypothetic | 2.716119997 | 2.336615425 | 4.408613997 | 25.88405818 | 19.25071231 | 20.82293584 | 0.000224 | 2.984589 | 8.40E-05 | 3.932799 | 7.915001 | 3.649822 |
| CNAG_034 | RDP1 | RNA-depe | 1780.635627 | 1799.216833 | 1647.65255 | 1425.58676 | 1420.48452 | 1472.162485 | 0.000225 | -0.29073 | 8.42E-05 | -3.93213 | 0.817486 | 3.64875 |
| CNAG_027 | - | 20S prote | 1988.846982 | 1922.237928 | 2032.712799 | 2401.218703 | 2373.870378 | 2434.470192 | 0.000226 | 0.263258 | 8.48E-05 | 3.930377 | 1.200186 | 3.646153 |
| CNAG_075 | - | hypothetic | 51.79789714 | 46.55717819 | 53.4789653 | 88.18624377 | 118.7909415 | 102.9513847 | 0.000236 | 1.022084 | 8.91E-05 | 3.918401 | 2.030851 | 3.626241 |
| CNAG_127 | - | unspecifie | 7.626123401 | 18.52181984 | 22.53347809 | 57.85196113 | 54.8930612 | 42.03703887 | 0.00024 | 1.688879 | 9.04E-05 | 3.915077 | 3.224061 | 3.620678 |
| CNAG_057 | - | V-type AT | 1533.196534 | 1669.918611 | 1641.197803 | 1410.16096 | 1289.358059 | 1252.424257 | 0.000241 | -0.3086 | 9.11E-05 | -3.91326 | 0.807426 | 3.617697 |
| CNAG_037 | - | Putative tı | 204.8657504 | 288.8083648 | 243.2858439 | 330.892725 | 421.4018023 | 372.8106749 | 0.000245 | 0.597376 | 9.25E-05 | 3.909354 | 1.512962 | 3.611086 |
| CNAG_009 | - | alkylbase | 918.8757613 | 966.3455565 | 918.159368 | 702.4382373 | 761.5094607 | 762.7846804 | 0.000249 | -0.34768 | 9.42E-05 | -3.90518 | 0.785845 | 3.603725 |
| CNAG_057 | - | hypothetic | 332.853072 | 440.4498918 | 429.6134151 | 283.7254305 | 265.3870526 | 284.3727565 | 0.000265 | -0.54376 | 0.0001 | -3.88953 | 0.685979 | 3.577489 |
| CNAG_032 | - | large subı | 305.4846694 | 259.5922535 | 283.5524002 | 193.4261189 | 204.2539534 | 164.596972 | 0.000271 | -0.61047 | 0.000103 | -3.88351 | 0.654983 | 3.56701 |
| CNAG_009 | - | aflatoxin ε | 337.7475959 | 243.4177057 | 227.1308432 | 174.7175906 | 136.7076441 | 182.2303311 | 0.000277 | -0.72995 | 0.000106 | -3.87744 | 0.602923 | 3.557281 |
| CNAG_127 | - | unspecifie | 12.15507441 | 13.14201024 | 14.03826838 | 40.27958547 | 46.27088955 | 40.78163305 | 0.000286 | 1.676577 | 0.000109 | 3.869491 | 3.196685 | 3.543803 |
| CNAG_052 | - | hypothetic | 5.098677351 | 7.713333814 | 4.443562675 | 29.6194964 | 28.93932709 | 24.21112364 | 0.000286 | 2.32555 | 0.000109 | 3.869124 | 5.012569 | 3.543428 |
| CNAG_079 | - | hypothetic | 6.303531412 | 7.746190138 | 8.697788581 | 44.309435 | 29.66007414 | 22.16092951 | 0.000287 | 2.162453 | 0.00011 | 3.86812 | 4.476754 | 3.541918 |
| CNAG_034 | - | peptidyl-p | 2014.309521 | 2153.701658 | 2118.981468 | 1834.309119 | 1711.161921 | 1719.762317 | 0.000294 | -0.2707 | 0.000113 | -3.86184 | 0.828916 | 3.531305 |
| CNAG_009 | - | ATP-depe | 4460.615661 | 4228.877219 | 4142.080918 | 3597.81693 | 3810.730413 | 3553.745904 | 0.000301 | -0.24282 | 0.000115 | -3.85571 | 0.845092 | 3.520961 |
| CNAG_032 | - | THO comp | 2065.596137 | 2094.443795 | 1969.711303 | 1652.817251 | 1667.172324 | 1796.294562 | 0.000325 | -0.27571 | 0.000125 | -3.83611 | 0.826041 | 3.487768 |
| CNAG_058 | - | hypothetic | 1455.487021 | 1078.481038 | 1029.134068 | 904.0942868 | 918.4725668 | 799.5560212 | 0.000335 | -0.45918 | 0.000129 | -3.82882 | 0.727402 | 3.47531 |
| CNAG_055 | - | hypothetic | 674.7690879 | 643.1791037 | 719.7297143 | 547.714756 | 492.2810935 | 521.9193978 | 0.000337 | -0.39962 | 0.00013 | -3.82673 | 0.758059 | 3.47189 |
| CNAG_034 | SPP101 | compass ( | 702.8998404 | 777.7783408 | 746.4393996 | 577.9849268 | 584.6527692 | 582.9680609 | 0.000347 | -0.36709 | 0.000134 | -3.81913 | 0.775343 | 3.459183 |
| CNAG_031 | - | hypothetic | 126.7128012 | 198.3181664 | 177.1473354 | 283.9137243 | 248.5827797 | 275.0597422 | 0.000348 | 0.672393 | 0.000134 | 3.818683 | 1.593715 | 3.458676 |
| CNAG_037 | - | mRNA 3'- | 1429.14222 | 1353.215465 | 1434.303171 | 1214.788449 | 1135.835881 | 1091.44979 | 0.000353 | -0.30899 | 0.000136 | -3.81472 | 0.807207 | 3.452117 |
| CNAG_057 | - | Alcohol de | 267.8045508 | 352.3365329 | 247.6027854 | 423.7743115 | 411.6332544 | 438.2788023 | 0.000354 | 0.540864 | 0.000137 | 3.813716 | 1.454843 | 3.450488 |
| CNAG_067 | RPN14 | 19S prote | 784.8829239 | 671.4048307 | 768.9852446 | 985.6211683 | 1048.303995 | 896.9723233 | 0.000356 | 0.381772 | 0.000138 | 3.812502 | 1.302941 | 3.448492 |
| CNAG_050 | - | GabA per | 191.3949148 | 214.2613962 | 250.3577306 | 129.4543609 | 161.1839339 | 111.3586401 | 0.000364 | -0.72342 | 0.000141 | -3.80646 | 0.605661 | 3.43843 |
| CNAG_032 | - | Aldehyde | 1338.10576 | 1402.711764 | 1302.10437 | 1051.149424 | 1146.245384 | 1116.92577 | 0.000365 | -0.30198 | 0.000141 | -3.80587 | 0.811138 | 3.437528 |
| CNAG_121 | - | unspecifie | 149.0764902 | 136.7246143 | 156.5719987 | 81.15882166 | 92.04474264 | 85.72584566 | 0.000367 | -0.78919 | 0.000142 | -3.80469 | 0.57867 | 3.43573 |
| CNAG_058 | - | hypothetic | 2242.986611 | 1928.609743 | 2220.258901 | 1820.283849 | 1764.676791 | 1577.737554 | 0.000372 | -0.32428 | 0.000144 | -3.8005 | 0.798698 | 3.429062 |
| CNAG_054 | - | SHO1 osn | 3466.982996 | 3450.986047 | 3762.357766 | 2943.245121 | 3100.815965 | 3048.591984 | 0.000376 | -0.24755 | 0.000146 | -3.79809 | 0.842325 | 3.425249 |
| CNAG_058 | - | hypothetic | 394.8930332 | 390.1271383 | 338.2169501 | 533.53913 | 540.2382064 | 477.3037732 | 0.000389 | 0.450934 | 0.000151 | 3.788637 | 1.366925 | 3.409807 |
| CNAG_068 | - | hypothetic | 99.64986428 | 108.9607976 | 94.02379961 | 177.2387276 | 142.3540861 | 201.5847528 | 0.000392 | 0.77437 | 0.000153 | 3.786612 | 1.710443 | 3.406813 |
| CNAG_034 | - | NAD epim | 566.5011869 | 487.1555945 | 532.2504907 | 689.8092158 | 691.0654156 | 726.9675593 | 0.000403 | 0.394267 | 0.000157 | 3.779205 | 1.314275 | 3.394429 |
| CNAG_031 | SEC14-1 | SEC14 cyt | 7677.868914 | 7567.850463 | 7684.822125 | 6743.73273 | 6664.90016 | 6773.163768 | 0.000429 | -0.19949 | 0.000168 | -3.76251 | 0.87086 | 3.36794 |
| CNAG_055 | - | hypothetic | 1060.528893 | 1129.204475 | 1037.75399 | 1334.981861 | 1401.871991 | 1282.178197 | 0.000431 | 0.301326 | 0.000169 | 3.760836 | 1.232277 | 3.365174 |
| CNAG_034 | - | hypothetic | 1149.336739 | 1225.12467 | 1284.054695 | 1503.859214 | 1586.459999 | 1454.25074 | 0.000447 | 0.297387 | 0.000176 | 3.751148 | 1.228916 | 3.34999 |
| CNAG_050 | - | carnitine a | 553.7084125 | 571.1086995 | 485.3786356 | 747.5745022 | 674.61879 | 714.7962109 | 0.00046 | 0.393223 | 0.000181 | 3.743644 | 1.313324 | 3.33754 |
| CNAG_075 | - | small subı | 642.1641666 | 653.9405034 | 747.4268516 | 482.8962299 | 553.6508919 | 523.0285412 | 0.000467 | -0.40574 | 0.000185 | -3.73921 | 0.754848 | 3.330683 |
| CNAG_058 | GMT1 | GDP-man | 6820.835676 | 7336.175522 | 7472.571961 | 6265.841771 | 6412.200514 | 6165.936112 | 0.000472 | -0.214 | 0.000187 | -3.73619 | 0.862143 | 3.326135 |
| CNAG_057 | - | cytoplasm | 190.5800979 | 163.9558741 | 178.2226905 | 297.9911035 | 264.1633705 | 248.5738759 | 0.000473 | 0.59164 | 0.000187 | 3.735395 | 1.506959 | 3.324898 |
| CNAG_056 | - | hypothetic | 273.4824308 | 256.5704964 | 259.2648234 | 386.4756415 | 353.7856688 | 378.373382 | 0.000479 | 0.488032 | 0.00019 | 3.732247 | 1.402531 | 3.319735 |
| CNAG_057 | - | hypothetic | 545.6663075 | 602.3768947 | 630.3276526 | 774.6779413 | 780.8031323 | 758.1284759 | 0.000491 | 0.364689 | 0.000195 | 3.725578 | 1.287604 | 3.309312 |
| CNAG_050 | - | histone de | 968.7745437 | 813.4113031 | 790.2190052 | 710.4294627 | 631.1682164 | 628.5358576 | 0.000497 | -0.4007 | 0.000197 | -3.7223 | 0.757493 | 3.303937 |
| CNAG_058 | - | translatior | 10016.87319 | 9759.356043 | 10168.5804 | 8359.366577 | 8952.450948 | 8891.342992 | 0.000508 | -0.20799 | 0.000202 | -3.71633 | 0.865741 | 3.294344 |
| CNAG_055 | - | hypothetic | 44.94778474 | 58.36765659 | 68.35669129 | 127.033585 | 101.1339767 | 105.2499303 | 0.000511 | 0.955414 | 0.000204 | 3.714484 | 1.939136 | 3.291302 |
| CNAG_074 | - | hypothetic | 25.10875621 | 31.47103873 | 44.8841897 | 84.17763293 | 71.01917396 | 73.01148135 | 0.000518 | 1.174037 | 0.000206 | 3.711066 | 2.256422 | 3.286011 |
| CNAG_051 | - | Nicotinate | 3186.612028 | 3151.414835 | 2911.409729 | 2520.613521 | 2714.402703 | 2633.39384 | 0.000518 | -0.24859 | 0.000207 | -3.71048 | 0.84172 | 3.285356 |
| CNAG_026 | - | hypothetic | 464.9190975 | 430.0149532 | 403.2528485 | 594.9589119 | 538.3003678 | 628.1076754 | 0.00052 | 0.425185 | 0.000208 | 3.709314 | 1.342745 | 3.283626 |
| CNAG_035 | - | Ubiquitin | 1377.770782 | 1339.19127 | 1363.93134 | 1140.009018 | 1033.121556 | 1166.848812 | 0.000531 | -0.30399 | 0.000212 | -3.7037 | 0.810008 | 3.274661 |
| CNAG_074 | - | hypothetic | 410.1340275 | 453.6521771 | 359.5558924 | 574.7666554 | 596.3324052 | 511.7338866 | 0.000544 | 0.445248 | 0.000218 | 3.697012 | 1.361548 | 3.264281 |
| CNAG_034 | - | hypothetic | 349.3981915 | 609.4499533 | 520.2569479 | 326.4526518 | 363.8471353 | 286.7246653 | 0.000553 | -0.61352 | 0.000222 | -3.69264 | 0.653602 | 3.257077 |
| CNAG_033 | - | pre-mRNA | 693.2274167 | 621.5878612 | 567.3397838 | 483.0422121 | 450.0112628 | 491.9218168 | 0.000556 | -0.41697 | 0.000223 | -3.69117 | 0.748994 | 3.254823 |
| CNAG_057 | CDA1 | Chitin dea | 2288.817698 | 2077.369453 | 1686.234577 | 2619.236567 | 2530.879748 | 2608.73969 | 0.000587 | 0.342972 | 0.000236 | 3.67731 | 1.268367 | 3.231725 |
| CNAG_050 | - | hypothetic | 704.9981005 | 621.6390362 | 660.0575841 | 508.7083715 | 545.0359013 | 475.35096 | 0.000617 | -0.39341 | 0.000249 | -3.66378 | 0.76133 | 3.209918 |
| CNAG_050 | - | hypothetic | 20.49259553 | 29.32780843 | 44.87753181 | 66.51567477 | 62.24320349 | 101.382957 | 0.000617 | 1.285804 | 0.000249 | 3.66355 | 2.438178 | 3.209663 |

| | | | | | | | | | | | | | | |
|---|---|---|---|---|---|---|---|---|---|---|---|---|---|---|
| CNAG_05( | CKY1 | YjeF family | 1130.004672 | 1085.915152 | 896.9067738 | 863.7599862 | 824.4644265 | 699.6530964 | 0.000629 | -0.39814 | 0.000254 | -3.65855 | 0.758838 | 3.20155 |
| CNAG_058 | - | cytoplasm | 2845.757876 | 2912.255905 | 3069.176289 | 2490.317864 | 2576.380745 | 2511.280934 | 0.000632 | -0.23541 | 0.000255 | -3.65719 | 0.849447 | 3.199405 |
| CNAG_034 | - | hypothetic | 0.347644799 | 7.700371541 | 6.548576993 | 33.10416505 | 26.29972975 | 18.79385638 | 0.000637 | 2.554349 | 0.000257 | 3.654697 | 5.874025 | 3.195713 |
| CNAG_035 | - | hypothetic | 3537.94148 | 3168.669432 | 2881.571437 | 2594.560879 | 2628.078071 | 2716.648573 | 0.000638 | -0.2877 | 0.000258 | -3.65443 | 0.819205 | 3.195396 |
| CNAG_037 | - | hypothetic | 806.7233798 | 830.5850464 | 758.2164831 | 619.9951663 | 665.7918132 | 620.7288039 | 0.00065 | -0.34553 | 0.000263 | -3.64914 | 0.787022 | 3.187097 |
| CNAG_056 | - | hypothetic | 928.6214275 | 1066.685346 | 923.6538183 | 1215.910344 | 1260.395309 | 1185.573051 | 0.000654 | 0.312687 | 0.000265 | 3.647483 | 1.242019 | 3.184561 |
| CNAG_037 | - | IQ domain | 4156.061896 | 3717.234936 | 3541.797621 | 4482.303534 | 4700.248428 | 4611.509956 | 0.000664 | 0.257443 | 0.000269 | 3.64313 | 1.195358 | 3.177732 |
| CNAG_068 | - | hypothetic | 2.743953737 | 2.34927525 | 8.636639942 | 23.17655325 | 24.58817882 | 23.03128344 | 0.000665 | 2.528108 | 0.00027 | 3.642824 | 5.768145 | 3.177347 |
| CNAG_075 | - | hypothetic | 425.3021465 | 409.5573444 | 440.5220474 | 599.715974 | 586.7731573 | 521.7330736 | 0.000701 | 0.405705 | 0.000285 | 3.628515 | 1.324736 | 3.154282 |
| CNAG_008 | - | hypothetic | 329.214535 | 370.493726 | 374.1944702 | 257.2601172 | 268.9459357 | 252.1972791 | 0.000719 | -0.48061 | 0.000293 | -3.62152 | 0.716673 | 3.143174 |
| CNAG_057 | - | hypothetic | 286.99721 | 298.28903 | 272.9418438 | 203.5690082 | 203.3510234 | 195.5885673 | 0.000726 | -0.52529 | 0.000296 | -3.61866 | 0.694821 | 3.139167 |
| CNAG_027 | - | hypothetic | 2459.588073 | 1264.214502 | 1144.652174 | 3811.888453 | 4688.83221 | 4617.647878 | 0.000742 | 1.411859 | 0.000303 | 3.612463 | 2.660798 | 3.129808 |
| CNAG_025 | - | hypothetic | 143.8429099 | 139.1040955 | 90.89419056 | 230.3826065 | 205.5779814 | 186.3960089 | 0.000765 | 0.724309 | 0.000313 | 3.60402 | 1.652109 | 3.116326 |
| CNAG_058 | - | hypothetic | 295.6802642 | 282.4237865 | 290.1794367 | 398.789298 | 440.2593794 | 370.7298659 | 0.000809 | 0.462894 | 0.000333 | 3.58848 | 1.378303 | 3.091817 |
| CNAG_036 | - | hypothetic | 374.0039182 | 401.9904084 | 413.8613995 | 473.2051932 | 555.7309996 | 589.2222657 | 0.000856 | 0.429755 | 0.000353 | 3.57261 | 1.347005 | 3.067498 |
| CNAG_067 | - | septation | 1003.314683 | 1017.187 | 1049.450429 | 1292.222543 | 1185.191748 | 1337.570233 | 0.000882 | 0.298481 | 0.000365 | 3.563885 | 1.229849 | 3.054321 |
| CNAG_067 | - | hypothetic | 0.294107249 | 3.400168611 | 5.45443709 | 11.61923781 | 25.65862097 | 25.05868049 | 0.000906 | 2.916833 | 0.000376 | 3.556613 | 7.551865 | 3.042933 |
| CNAG_036 | NIC1 | high-affini | 1679.118742 | 1618.283357 | 1812.869708 | 1453.675807 | 1434.317144 | 1385.6111 | 0.000938 | -0.27368 | 0.00039 | -3.5469 | 0.827208 | 3.027798 |
| CNAG_055 | FHL1 | Pre-rRNA- | 1032.019904 | 891.0345487 | 1015.168373 | 755.4010503 | 830.5587979 | 762.8301862 | 0.000964 | -0.34006 | 0.000402 | -3.5388 | 0.79001 | 3.01574 |
| CNAG_055 | - | hypothetic | 33.42777591 | 56.25494471 | 89.59785322 | 118.0614861 | 129.0027383 | 119.6167436 | 0.000987 | 1.025238 | 0.000412 | 3.532138 | 2.035296 | 3.005546 |
| CNAG_026 | - | hypothetic | 0.22541254 | 0.108773863 | 2.245624187 | 12.82895952 | 12.27460484 | 20.56531945 | 0.000992 | 4.444067 | 0.000415 | 3.530609 | 21.76694 | 3.003415 |
| CNAG_123 | - | unspecifie | 82.17386854 | 73.03303711 | 57.54109994 | 36.09930529 | 32.42164663 | 31.37682645 | 0.000993 | -1.11397 | 0.000415 | -3.53025 | 0.462022 | 3.002945 |
| CNAG_050 | - | hypothetic | 7.448528706 | 10.93815453 | 8.697789929 | 27.65911653 | 32.35228598 | 32.94595256 | 0.001003 | 1.849612 | 0.00042 | 3.527483 | 3.604032 | 2.998793 |
| CNAG_036 | CPA2 | Cyclophilin | 17923.74977 | 17140.87218 | 19394.43113 | 16119.03837 | 15814.95158 | 15584.53939 | 0.001003 | -0.21229 | 0.00042 | -3.52726 | 0.863166 | 2.998562 |
| CNAG_052 | - | U3 small r | 704.1572784 | 727.2432111 | 989.3919378 | 632.5297398 | 629.5275939 | 513.1723868 | 0.001008 | -0.46379 | 0.000422 | -3.526 | 0.725081 | 2.996613 |
| CNAG_052 | - | ste/ste20/ | 1112.852562 | 1189.424433 | 1321.201599 | 929.7223838 | 984.0385791 | 1023.645593 | 0.001027 | -0.31805 | 0.00043 | -3.52066 | 0.802151 | 2.988497 |
| CNAG_036 | - | syntaxin 7 | 1181.501221 | 1074.161058 | 1102.66059 | 954.0391342 | 839.9000553 | 930.411193 | 0.001047 | -0.31743 | 0.000439 | -3.51519 | 0.802498 | 2.980048 |
| CNAG_036 | - | hypothetic | 972.6889343 | 970.7457392 | 1196.379555 | 815.3553381 | 808.0282778 | 859.3614811 | 0.001064 | -0.35378 | 0.000447 | -3.51073 | 0.78253 | 2.973077 |
| CNAG_079 | - | hypothetic | 1112.933254 | 1038.722879 | 833.0747745 | 1282.081434 | 1430.456381 | 1196.717891 | 0.001074 | 0.374125 | 0.000451 | 3.508096 | 1.296053 | 2.969089 |
| CNAG_036 | - | L-lactate c | 2090.105496 | 1987.826448 | 2061.396962 | 1768.124976 | 1812.178633 | 1578.82828 | 0.001089 | -0.26661 | 0.000458 | -3.50401 | 0.831271 | 2.962833 |
| CNAG_056 | LCB3 | sphingosir | 1845.087508 | 2128.906755 | 2158.405291 | 1610.743488 | 1729.320512 | 1769.665512 | 0.001109 | -0.27811 | 0.000467 | -3.49879 | 0.824673 | 2.954929 |
| CNAG_031 | CMT1 | alpha-1,3- | 959.8880723 | 1161.304332 | 1031.209301 | 849.6514454 | 818.3785891 | 876.0080539 | 0.001129 | -0.3236 | 0.000476 | -3.4938 | 0.799075 | 2.947423 |
| CNAG_033 | - | AAA famil | 1576.322731 | 1522.375268 | 1594.282859 | 1261.324114 | 1364.488867 | 1328.955053 | 0.001235 | -0.26274 | 0.000524 | -3.46817 | 0.833506 | 2.908252 |
| CNAG_074 | - | hypothetic | 692.3061038 | 689.481294 | 700.5832569 | 594.5001405 | 526.8153143 | 530.8181358 | 0.001265 | -0.35003 | 0.000537 | -3.46139 | 0.784567 | 2.898052 |
| CNAG_026 | - | mitochonc | 435.3139453 | 406.1324304 | 441.3987304 | 321.9134931 | 286.0693517 | 347.559432 | 0.001286 | -0.44166 | 0.000547 | -3.45654 | 0.736288 | 2.890617 |
| CNAG_033 | - | hypothetic | 929.7955425 | 1063.459688 | 946.0400798 | 1144.148228 | 1212.884551 | 1325.318549 | 0.001288 | 0.31073 | 0.000548 | 3.455956 | 1.240336 | 2.890043 |
| CNAG_031 | - | hypothetic | 375.0966473 | 412.7110776 | 347.7985074 | 520.2656346 | 489.2036847 | 504.9792667 | 0.00132 | 0.401299 | 0.000563 | 3.448937 | 1.320696 | 2.879366 |
| CNAG_057 | - | hypothetic | 1268.998385 | 1133.400659 | 1060.050941 | 934.4827798 | 976.3201323 | 898.2768041 | 0.00136 | -0.31735 | 0.000581 | -3.44035 | 0.802542 | 2.866564 |
| CNAG_075 | - | hypothetic | 510.6828927 | 652.945412 | 568.5168639 | 806.7575094 | 739.3518204 | 718.1993891 | 0.001365 | 0.373226 | 0.000584 | 3.439167 | 1.295246 | 2.864791 |
| CNAG_051 | - | DNA-3-me | 553.3316488 | 552.6394125 | 557.6767813 | 452.7723494 | 409.487255 | 437.5302017 | 0.001396 | -0.37178 | 0.000598 | -3.43272 | 0.772827 | 2.855198 |
| CNAG_036 | - | hypothetic | 402.7040491 | 402.8805745 | 428.5996794 | 319.6220306 | 303.4081342 | 310.9950192 | 0.001509 | -0.41787 | 0.000649 | -3.41027 | 0.748531 | 2.82144 |
| CNAG_068 | - | hypothetic | 2.808675658 | 3.45605008 | 13.94498874 | 19.48870966 | 46.86896517 | 23.19145057 | 0.001509 | 2.272048 | 0.00065 | 3.409984 | 4.830084 | 2.821238 |
| CNAG_027 | - | aryl-alcoh | 1536.72853 | 1715.163658 | 1599.638921 | 1368.03466 | 1374.80454 | 1370.036399 | 0.001571 | -0.25291 | 0.000678 | -3.39826 | 0.839201 | 2.803952 |
| CNAG_034 | - | hypothetic | 1353.517201 | 1320.991854 | 1162.554643 | 1534.945457 | 1582.080412 | 1576.318336 | 0.00159 | 0.275772 | 0.000687 | 3.394634 | 1.210642 | 2.798686 |
| CNAG_026 | - | hypothetic | 16.82410202 | 25.97542671 | 14.07665539 | 60.31497731 | 46.02046058 | 39.85932342 | 0.001596 | 1.385036 | 0.00069 | 3.393438 | 2.611785 | 2.796913 |
| CNAG_126 | - | unspecifie | 34.55980357 | 73.36174842 | 67.30415331 | 109.4515511 | 100.2331045 | 130.5201154 | 0.001651 | 0.949454 | 0.000716 | 3.383474 | 1.931142 | 2.782237 |
| CNAG_050 | - | hypothetic | 271.1034596 | 269.4473414 | 221.9878011 | 370.1263677 | 335.655499 | 361.7257249 | 0.001663 | 0.471294 | 0.000721 | 3.381385 | 1.386353 | 2.779055 |
| CNAG_052 | - | sentrin/su | 761.280544 | 806.9041721 | 799.7671331 | 602.8304099 | 664.932114 | 650.6570737 | 0.001681 | -0.31944 | 0.00073 | -3.37822 | 0.801378 | 2.774305 |
| CNAG_026 | - | ribonuclec | 976.3769121 | 899.7715085 | 949.2136015 | 1181.688779 | 1246.63841 | 1082.389381 | 0.001734 | 0.297551 | 0.000754 | 3.369162 | 1.229056 | 2.760864 |
| CNAG_057 | - | hypothetic | 318.9430679 | 293.1807465 | 257.1765284 | 368.2307978 | 472.292081 | 381.8049744 | 0.001825 | 0.476562 | 0.000797 | 3.353871 | 1.391424 | 2.738766 |
| CNAG_075 | - | alpha-1,6- | 9.796537124 | 13.10205886 | 9.781127937 | 37.31317286 | 17.03877274 | 54.6468735 | 0.00186 | 1.788191 | 0.000814 | 3.348084 | 3.453817 | 2.730417 |
| CNAG_049 | CAT1 | Catalase 1 | 4827.024077 | 5042.122561 | 3624.907866 | 3443.721973 | 3045.435813 | 2715.753707 | 0.001884 | -0.56674 | 0.000825 | -3.34441 | 0.675139 | 2.724901 |
| CNAG_032 | - | tRNA (ura | 310.8190507 | 283.5087626 | 299.770131 | 394.8302431 | 450.6208177 | 375.1766655 | 0.001931 | 0.43476 | 0.000846 | 3.337279 | 1.351686 | 2.714115 |
| CNAG_035 | - | ribosomal | 1729.348716 | 1684.00587 | 1912.038183 | 1395.933691 | 1428.228918 | 1605.315621 | 0.001967 | -0.28149 | 0.000862 | -3.33196 | 0.82274 | 2.706288 |
| CNAG_057 | UBC6 | ubiquitin- | 2489.338008 | 2104.227431 | 2228.849688 | 1912.834599 | 1835.355872 | 1980.593693 | 0.002029 | -0.26806 | 0.000892 | -3.32249 | 0.830436 | 2.692615 |
| CNAG_036 | - | hypothetic | 196.4789571 | 186.5595226 | 215.4962622 | 265.5740801 | 311.748544 | 279.6125416 | 0.002043 | 0.50249 | 0.000899 | 3.320488 | 1.416657 | 2.689746 |
| CNAG_126 | - | unspecifie | 0.205563092 | 0.099339495 | 2.23379342 | 8.65991467 | 15.09647793 | 16.26367751 | 0.002073 | 4.269195 | 0.000913 | 3.316143 | 19.28216 | 2.683471 |
| CNAG_032 | - | yippee-like | 56.34704412 | 32.61695787 | 52.39868243 | 94.7032636 | 132.0629695 | 66.6900371 | 0.002089 | 1.042359 | 0.000921 | 3.313549 | 2.059593 | 2.680042 |
| CNAG_058 | - | DNA excis | 500.0476707 | 517.2622426 | 531.1605464 | 660.2375192 | 624.5647931 | 696.9893224 | 0.002095 | 0.340079 | 0.000924 | 3.312665 | 1.265826 | 2.678911 |
| CNAG_078 | - | 2-hydroxy | 1.443913806 | 2.281120478 | 1.180207904 | 10.29889362 | 14.12546417 | 17.36464254 | 0.00217 | 3.343795 | 0.000959 | 3.302382 | 10.15272 | 2.663446 |
| CNAG_036 | YKU80 | ATP-deper | 1274.250884 | 1449.166703 | 1261.71203 | 1633.855035 | 1520.725083 | 1713.918776 | 0.002181 | 0.274092 | 0.000964 | 3.300813 | 1.209233 | 2.661258 |
| CNAG_027 | - | hypothetic | 560.4749858 | 620.4948729 | 632.3010948 | 442.4550538 | 493.2599522 | 494.1200065 | 0.002186 | -0.35754 | 0.000967 | -3.29996 | 0.780496 | 2.660291 |
| CNAG_050 | - | sulfite red | 79.66309993 | 73.42123425 | 63.10477565 | 113.139551 | 128.896355 | 125.1883166 | 0.002187 | 0.752113 | 0.000967 | 3.299834 | 1.684258 | 2.660222 |
| CNAG_057 | - | endoplasn | 1007.581082 | 917.9620854 | 1034.364733 | 804.5126694 | 843.4927498 | 780.5893145 | 0.002237 | -0.30125 | 0.000991 | -3.29316 | 0.811548 | 2.650382 |
| CNAG_033 | - | hypothetic | 924.7108749 | 931.9036631 | 920.2937527 | 747.6816176 | 707.0806461 | 820.4453905 | 0.002449 | -0.30227 | 0.001094 | -3.26513 | 0.810974 | 2.610943 |
| CNAG_068 | - | SBDS fam | 1879.798016 | 1738.915046 | 1611.411683 | 1554.164717 | 1456.735414 | 1355.676255 | 0.002473 | -0.2758 | 0.001106 | -3.26213 | 0.825995 | 2.606818 |
| CNAG_127 | - | unspecifie | 548.8137402 | 673.201169 | 577.9705168 | 497.1625209 | 468.1993898 | 440.9217569 | 0.002501 | -0.3702 | 0.00112 | -3.2584 | 0.773674 | 2.601927 |
| CNAG_05( | - | meiotic Si | 584.9552875 | 664.6228029 | 589.7079673 | 499.4500589 | 496.6972648 | 466.4333861 | 0.002508 | -0.34512 | 0.001124 | -3.2575 | 0.787242 | 2.600678 |

| | | | | | | | | | | | | | | |
|---|---|---|---|---|---|---|---|---|---|---|---|---|---|---|
| CNAG_02! | - | hypothetic | 292.5293207 | 220.8566324 | 245.1989261 | 179.4506651 | 185.2759965 | 160.1111693 | 0.002521 | -0.5472 | 0.001131 | -3.25583 | 0.684349 | 2.598359 |
| CNAG_03 | - | DNA poly | 2428.963351 | 2764.603689 | 2764.171818 | 2214.776354 | 2268.416777 | 2355.819178 | 0.002533 | -0.23341 | 0.001137 | -3.25429 | 0.850623 | 2.596309 |
| CNAG_03 | - | hypothetic | 8.660219368 | 17.35641319 | 8.726848153 | 35.62658745 | 50.06288298 | 22.25899798 | 0.002577 | 1.683546 | 0.001158 | 3.249101 | 3.212165 | 2.588902 |
| CNAG_04 | - | hypothetic | 347.567057 | 308.0597472 | 291.1073262 | 230.8205849 | 230.0953267 | 234.3952912 | 0.002588 | -0.46247 | 0.001164 | -3.24761 | 0.725743 | 2.586982 |
| CNAG_05! | - | hypothetic | 19.15759135 | 31.32890313 | 14.0918013 | 59.28332532 | 45.03531296 | 52.98147896 | 0.002618 | 1.27262 | 0.001178 | 3.244049 | 2.415999 | 2.582019 |
| CNAG_05( | DAM1 | pre-mRNA | 1718.766778 | 1590.278605 | 1739.311358 | 1508.244854 | 1407.563321 | 1356.759927 | 0.002686 | -0.2565 | 0.001211 | -3.23619 | 0.837117 | 2.570874 |
| CNAG_02( | - | hypothetic | 518.7420062 | 578.6315987 | 522.6614066 | 746.0538547 | 685.8797542 | 638.3117147 | 0.00269 | 0.339919 | 0.001214 | 3.235571 | 1.265685 | 2.570168 |
| CNAG_03 | - | mRNA gu | 1082.668463 | 1076.433717 | 1044.142324 | 1198.560356 | 1458.890983 | 1291.035748 | 0.002726 | 0.286327 | 0.001232 | 3.231298 | 1.219532 | 2.564491 |
| CNAG_03 | - | hypothetic | 1573.124037 | 1648.513969 | 1536.836655 | 1837.661095 | 1903.798286 | 1883.855187 | 0.002728 | 0.226269 | 0.001234 | 3.230918 | 1.169805 | 2.564149 |
| CNAG_05 | - | hypothetic | 99.71459136 | 120.7969039 | 125.9293976 | 203.1163369 | 170.1040381 | 163.098537 | 0.002767 | 0.625116 | 0.001253 | 3.22646 | 1.542335 | 2.557965 |
| CNAG_05! | - | hypothetic | 268.9846629 | 311.4849742 | 339.1780161 | 414.3368161 | 461.8278378 | 379.6356832 | 0.002872 | 0.434828 | 0.001303 | 3.215337 | 1.35175 | 2.541808 |
| CNAG_03 | - | hypothetic | 1094.166769 | 1203.405476 | 1206.082812 | 1017.872992 | 957.2881409 | 964.8464638 | 0.002883 | -0.26864 | 0.001308 | -3.21411 | 0.8301 | 2.540189 |
| CNAG_05( | - | hypothetic | 5.155955835 | 13.06353886 | 9.754360513 | 26.81710362 | 40.42938629 | 25.40543969 | 0.002894 | 1.739411 | 0.001314 | 3.21281 | 3.338987 | 2.538573 |
| CNAG_05 | SHB17 | Phosphog | 2820.001533 | 2722.645442 | 2880.427407 | 2218.566192 | 2487.550564 | 2541.197034 | 0.002925 | -0.23245 | 0.001329 | -3.20956 | 0.851189 | 2.533898 |
| CNAG_03 | - | hypothetic | 869.7971459 | 856.4897224 | 832.8608894 | 691.6437902 | 705.4389024 | 720.5832733 | 0.002934 | -0.28835 | 0.001334 | -3.20848 | 0.818835 | 2.532609 |
| CNAG_03 | - | hypothetic | 220.9723247 | 253.2367427 | 186.8066964 | 302.2465332 | 328.9379533 | 301.8322054 | 0.002973 | 0.483927 | 0.001355 | 3.204046 | 1.398545 | 2.526737 |
| CNAG_05! | - | ino eighty | 626.0105543 | 827.2769435 | 760.2862915 | 599.8524485 | 599.3492461 | 544.1680187 | 0.003043 | -0.35952 | 0.001389 | -3.19692 | 0.779422 | 2.516702 |
| CNAG_05 | - | mitochon | 1363.86386 | 1383.378964 | 1469.490172 | 1216.31436 | 1127.182932 | 1225.686247 | 0.003067 | -0.25556 | 0.001401 | -3.19449 | 0.837662 | 2.513268 |
| CNAG_03 | - | hypothetic | 823.1369431 | 815.5647644 | 859.4785701 | 689.3554777 | 639.8071062 | 719.4470047 | 0.00313 | -0.30186 | 0.001431 | -3.1883 | 0.811203 | 2.504431 |
| CNAG_12 | - | unspecifie | 51.57237302 | 30.42735294 | 40.68157613 | 80.73659146 | 76.17391119 | 81.85660678 | 0.003138 | 0.953089 | 0.001436 | 3.187354 | 1.936014 | 2.503355 |
| CNAG_05 | FRK101 | CAMK pro | 32.03693036 | 40.03693006 | 45.96075836 | 67.18362709 | 84.14724961 | 77.43657929 | 0.003173 | 0.947564 | 0.001453 | 3.183909 | 1.928613 | 2.498533 |
| CNAG_12 | - | unspecifie | 189.0428601 | 220.6784851 | 205.712869 | 145.2182165 | 118.6629276 | 155.525462 | 0.003278 | -0.56907 | 0.001506 | -3.17349 | 0.674051 | 2.484403 |
| CNAG_03 | - | ATP-depe | 1235.674744 | 1271.440498 | 1295.790019 | 1508.496486 | 1462.10025 | 1565.197167 | 0.003331 | 0.239652 | 0.001532 | 3.168537 | 1.180708 | 2.477367 |
| CNAG_00 | CNB1 | calcineuri | 1606.657349 | 1632.21788 | 1469.568541 | 1386.803333 | 1234.119295 | 1358.912125 | 0.003349 | -0.25767 | 0.00154 | -3.16696 | 0.836439 | 2.475126 |
| CNAG_12 | - | unspecifie | 114.0260677 | 127.9134677 | 91.69445327 | 49.86219938 | 65.27282567 | 77.81650919 | 0.003439 | -0.81199 | 0.001585 | -3.15861 | 0.569598 | 2.463598 |
| CNAG_07 | - | hypothetic | 2102.841961 | 2154.712422 | 1747.938796 | 1416.029212 | 1412.588114 | 1921.618867 | 0.003453 | -0.35252 | 0.001594 | -3.15697 | 0.783215 | 2.461838 |
| CNAG_05 | STP1 | putative s | 729.6868017 | 703.5206727 | 747.5031025 | 570.1819179 | 604.5327496 | 608.46018 | 0.003532 | -0.30561 | 0.001635 | -3.14966 | 0.809102 | 2.451999 |
| CNAG_03 | SAF2 | hypothetic | 854.676356 | 837.1274251 | 909.5914826 | 736.8760836 | 677.7749823 | 731.6846997 | 0.003543 | -0.2935 | 0.001641 | -3.14858 | 0.815922 | 2.450613 |
| CNAG_12 | - | unspecifie | 15.63934245 | 19.56334926 | 20.399562 | 44.96310656 | 39.01063287 | 48.47526295 | 0.003598 | 1.264084 | 0.001669 | 3.143501 | 2.401747 | 2.443993 |
| CNAG_03 | CFT3 | iron ion tr | 1015.904147 | 1149.495975 | 1044.023079 | 880.7841264 | 911.6187597 | 899.3301715 | 0.00361 | -0.26879 | 0.001676 | -3.14234 | 0.830017 | 2.442495 |
| CNAG_05 | - | hypothetic | 584.2661918 | 802.663034 | 637.8527284 | 900.8341583 | 834.1950713 | 868.0187359 | 0.003727 | 0.349396 | 0.001733 | 3.132462 | 1.274027 | 2.428664 |
| CNAG_05! | - | transcripti | 8670.434431 | 8536.452382 | 8190.352479 | 7201.28295 | 7744.016597 | 7716.7783 | 0.003766 | -0.17969 | 0.001754 | -3.12907 | 0.882895 | 2.424109 |
| CNAG_03 | - | hypothetic | 199.9575512 | 224.1638398 | 175.0632747 | 302.4903261 | 249.3077527 | 299.5032334 | 0.003944 | 0.492993 | 0.001843 | 3.114457 | 1.407361 | 2.404027 |
| CNAG_05 | - | helicase s | 7710.508827 | 7522.581146 | 7072.71714 | 6141.985079 | 6752.068929 | 6783.126368 | 0.003981 | -0.19617 | 0.001861 | -3.11159 | 0.872863 | 2.400029 |
| CNAG_03 | - | hypothetic | 628.3791681 | 764.8840707 | 883.8899878 | 591.2166997 | 593.26996 | 599.6150891 | 0.004012 | -0.36633 | 0.001877 | -3.10908 | 0.775755 | 2.396686 |
| CNAG_03 | - | phosphati | 831.4529565 | 978.1783121 | 934.137089 | 740.6834524 | 695.8641273 | 807.1270597 | 0.004096 | -0.30539 | 0.001919 | -3.10242 | 0.809224 | 2.387587 |
| CNAG_07 | - | hypothetic | 216.8224995 | 194.9319684 | 207.8405014 | 168.0369337 | 112.5856438 | 137.860014 | 0.00421 | -0.58058 | 0.001974 | -3.09405 | 0.668695 | 2.375764 |
| CNAG_05 | - | ESCRT-II c | 213.6504344 | 253.034186 | 256.8638158 | 178.7330642 | 143.6834555 | 186.6054583 | 0.004215 | -0.52438 | 0.001977 | -3.0936 | 0.695256 | 2.37523 |
| CNAG_08( | - | hypothetic | 252.2467541 | 181.2042111 | 187.8493637 | 294.531305 | 281.3343695 | 313.9261247 | 0.0043 | 0.502068 | 0.002021 | 3.08709 | 1.416242 | 2.366494 |
| CNAG_12 | - | unspecifie | 29.82642741 | 65.75160516 | 44.9581365 | 82.06188545 | 97.94918067 | 87.46515108 | 0.0043 | 0.934091 | 0.002021 | 3.087155 | 1.910686 | 2.366494 |
| CNAG_03! | - | solute car | 633.968065 | 647.4631321 | 663.2528102 | 558.6968754 | 475.8829118 | 534.088226 | 0.004371 | -0.3253 | 0.002057 | -3.08192 | 0.798133 | 2.35939 |
| CNAG_00 | - | hypothetic | 814.9077259 | 753.1058224 | 827.4835992 | 658.2120461 | 638.1226307 | 677.2809782 | 0.00441 | -0.29456 | 0.002076 | -3.07911 | 0.81532 | 2.355522 |
| CNAG_02 | ALG12 | alpha-1,6- | 754.619095 | 810.2999649 | 834.0104546 | 944.9681577 | 1005.053837 | 973.5144787 | 0.004577 | 0.270027 | 0.00216 | 3.067271 | 1.20583 | 2.339418 |
| CNAG_12( | - | unspecifie | 19.29112381 | 38.89817666 | 36.37631632 | 69.99035857 | 63.26361879 | 61.95792465 | 0.004753 | 1.039818 | 0.002251 | 3.054959 | 2.055968 | 2.323019 |
| CNAG_03! | YOR1 | ATP-bindi | 1896.543696 | 1846.790191 | 1822.637337 | 2046.134626 | 2209.136002 | 2300.109162 | 0.004817 | 0.221169 | 0.002282 | 3.050827 | 1.165678 | 2.317252 |
| CNAG_05( | - | hypothetic | 59.25558088 | 68.69793989 | 61.70860244 | 29.80267279 | 21.96326302 | 42.27037227 | 0.004848 | -1.04496 | 0.002299 | -3.04861 | 0.484657 | 2.314394 |
| CNAG_03 | - | DNA-dire | 971.3794842 | 922.238321 | 935.2320483 | 838.1331505 | 778.7499242 | 745.0721496 | 0.004891 | -0.27584 | 0.002321 | -3.04577 | 0.82597 | 2.310628 |
| CNAG_05! | - | transcripti | 768.1119125 | 622.8002802 | 830.595557 | 644.366824 | 580.3413839 | 492.0637878 | 0.004967 | -0.38798 | 0.002362 | -3.04051 | 0.764198 | 2.303932 |
| CNAG_12( | - | unspecifie | 35.53480648 | 46.47787397 | 50.22360886 | 82.25935688 | 84.03294765 | 75.32676077 | 0.005022 | 0.864717 | 0.00239 | 3.036871 | 1.820982 | 2.299127 |
| CNAG_05 | HPI4 | symplekin | 5013.088901 | 5127.449285 | 4937.661502 | 4433.74756 | 4524.051283 | 4590.566118 | 0.005126 | -0.16948 | 0.002442 | -3.03046 | 0.889164 | 2.290234 |
| CNAG_12 | - | unspecifie | 6.311354641 | 18.36615527 | 1.260935752 | 21.83591285 | 35.91761926 | 39.42556074 | 0.00515 | 1.9165 | 0.002455 | 3.028789 | 3.775061 | 2.288163 |
| CNAG_03! | - | hypothetic | 639.7726548 | 649.606823 | 622.7663763 | 523.5670759 | 533.8102407 | 509.6984028 | 0.005151 | -0.3026 | 0.002456 | -3.0287 | 0.810789 | 2.288144 |
| CNAG_03 | FZC5 | hypothetic | 20.47929225 | 42.13342154 | 41.6928622 | 67.3852626 | 76.37085676 | 66.40594412 | 0.005181 | 1.007262 | 0.002472 | 3.026715 | 2.010093 | 2.285628 |
| CNAG_03 | - | hypothetic | 575.6118361 | 594.6742594 | 622.7156959 | 507.295351 | 482.8835243 | 463.094504 | 0.005207 | -0.31793 | 0.002486 | -3.025 | 0.80222 | 2.283381 |
| CNAG_03 | - | hypothetic | 109.1802692 | 172.4465185 | 164.3081705 | 247.3883898 | 209.7503302 | 213.015068 | 0.005219 | 0.57446 | 0.002494 | 3.024074 | 1.48912 | 2.28239 |
| CNAG_05 | - | ENTH dor | 1374.417249 | 1451.244716 | 1534.535782 | 1132.843686 | 1336.953999 | 1216.844139 | 0.00527 | -0.25802 | 0.00252 | -3.02098 | 0.836233 | 2.278166 |
| CNAG_12 | - | unspecifie | 11.89228956 | 3.442157364 | 7.610649951 | 21.28945215 | 24.45132351 | 30.64305805 | 0.005298 | 1.818449 | 0.002535 | 3.019095 | 3.527018 | 2.275914 |
| CNAG_03( | - | hypothetic | 534.5627047 | 496.6364302 | 511.8244056 | 375.5332349 | 353.3553649 | 466.2535854 | 0.005332 | -0.38327 | 0.002552 | -3.01707 | 0.7667 | 2.273123 |
| CNAG_12 | - | unspecifie | 127.8161759 | 210.0814608 | 149.4595183 | 236.9272314 | 222.6516743 | 270.5415415 | 0.005352 | 0.570077 | 0.002563 | 3.015788 | 1.484603 | 2.271507 |
| CNAG_05 | - | AP-3 com | 383.9966417 | 386.7026236 | 375.3311767 | 297.0301495 | 308.6680367 | 288.8024945 | 0.005801 | -0.37261 | 0.00279 | -2.98993 | 0.772383 | 2.236482 |
| CNAG_02! | - | hypothetic | 12.9710791 | 1.274593519 | 4.431836474 | 16.38396239 | 25.42364198 | 31.65402502 | 0.005949 | 2.062273 | 0.002869 | 2.981434 | 4.176437 | 2.225523 |
| CNAG_05 | - | peroxisom | 93.88255536 | 134.7123918 | 104.6575928 | 156.647903 | 168.4359306 | 182.8829891 | 0.005969 | 0.598522 | 0.00288 | 2.980293 | 1.514164 | 2.224124 |
| CNAG_03 | - | hypothetic | 380.8777017 | 549.2830724 | 546.9162639 | 355.3067776 | 330.9247366 | 414.1363541 | 0.006122 | -0.43821 | 0.002958 | -2.97206 | 0.738051 | 2.213126 |
| CNAG_03 | - | RecQ-mec | 298.0831588 | 336.2289803 | 321.0873062 | 391.5923167 | 402.9264711 | 462.6719734 | 0.006134 | 0.379455 | 0.002968 | 2.971059 | 1.300851 | 2.212256 |
| CNAG_05 | - | hypothetic | 3429.521393 | 3315.211662 | 3124.708222 | 2854.580195 | 2873.944864 | 2989.710689 | 0.006134 | -0.19435 | 0.002966 | -2.97127 | 0.873966 | 2.212256 |
| CNAG_05 | - | cytochron | 2003.831322 | 2109.556097 | 2128.585804 | 1764.939278 | 1799.187169 | 1897.303883 | 0.006162 | -0.20766 | 0.002983 | -2.96949 | 0.865942 | 2.210264 |
| CNAG_02( | - | hypothetic | 352.536832 | 375.9121749 | 376.3612237 | 246.2203637 | 319.1153926 | 276.5870535 | 0.006189 | -0.40653 | 0.002999 | -2.96779 | 0.754434 | 2.20839 |
| CNAG_03 | - | hypothetic | 42.23136557 | 34.6419628 | 21.5640982 | 58.63608492 | 59.70155824 | 82.72057363 | 0.006192 | 1.021552 | 0.003002 | 2.967527 | 2.030101 | 2.208141 |

| | | | | | | | | | | | | | | |
|---|---|---|---|---|---|---|---|---|---|---|---|---|---|---|
| CNAG_123 | - | unspecifie | 9.74145456 | 10.94547634 | 12.920132 | 30.92385573 | 28.73211466 | 30.81543474 | 0.006196 | 1.490196 | 0.003004 | 2.967278 | 2.809271 | 2.207899 |
| CNAG_123 | - | unspecifie | 90.7909736 | 106.3905552 | 84.18173631 | 68.34059541 | 44.44598007 | 50.22308691 | 0.006228 | -0.8036 | 0.003021 | -2.96561 | 0.572916 | 2.205653 |
| CNAG_051 | - | hypothetic | 84.53704942 | 76.7439999 | 130.0979938 | 148.9800209 | 152.0428114 | 167.3278248 | 0.006381 | 0.671403 | 0.003099 | 2.957744 | 1.592621 | 2.195103 |
| CNAG_122 | - | unspecifie | 9.52016751 | 3.409923414 | 2.290781421 | 22.47737672 | 16.52833772 | 22.97016692 | 0.006434 | 2.082144 | 0.003128 | 2.954911 | 4.234361 | 2.19155 |
| CNAG_124 | - | unspecifie | 26.20332917 | 38.91071534 | 35.32861279 | 73.95367645 | 51.77607321 | 72.90303188 | 0.006442 | 0.971678 | 0.003133 | 2.954407 | 1.96112 | 2.190951 |
| CNAG_032 | CCD4 | hypothetic | 322.5843728 | 386.8139213 | 301.9496854 | 400.0676065 | 454.7732916 | 483.7858456 | 0.006587 | 0.391766 | 0.003205 | 2.947405 | 1.311998 | 2.181324 |
| CNAG_121 | - | unspecifie | 0.117830818 | 0.057309428 | 0.049508769 | 13.58403411 | 5.262154913 | 1.205988879 | 0.006611 | 5.090161 | 0.003217 | 2.946199 | 34.06364 | 2.179739 |
| CNAG_057 | - | oligopepti | 1055.416846 | 1096.683936 | 874.5293112 | 830.9889418 | 870.2402477 | 795.027052 | 0.006618 | -0.29285 | 0.003221 | -2.94581 | 0.816288 | 2.179298 |
| CNAG_052 | - | hypothetic | 91.43145652 | 85.30721568 | 103.5317529 | 125.3526746 | 186.989055 | 133.1345088 | 0.006786 | 0.654127 | 0.003308 | 2.937559 | 1.573663 | 2.168385 |
| CNAG_049 | - | hypothetic | 443.6473862 | 486.8744659 | 455.3183118 | 345.199664 | 385.4624146 | 372.0186271 | 0.006787 | -0.34365 | 0.003309 | -2.93746 | 0.788045 | 2.16835 |
| CNAG_032 | - | hypothetic | 585.1950316 | 366.3304109 | 333.8578001 | 309.4313721 | 321.5731229 | 273.3563533 | 0.006846 | -0.52515 | 0.003343 | -2.93426 | 0.694889 | 2.164533 |
| CNAG_055 | - | bis(5'-ade | 401.5387164 | 378.1496502 | 437.1144199 | 338.4096181 | 297.3514387 | 312.100814 | 0.006898 | -0.3762 | 0.003373 | -2.93157 | 0.770464 | 2.161303 |
| CNAG_055 | - | hypothetic | 1053.378108 | 1376.756102 | 1141.078615 | 988.9944029 | 954.6826348 | 1012.549679 | 0.006902 | -0.28717 | 0.003376 | -2.9313 | 0.819511 | 2.161032 |
| CNAG_032 | - | THO comp | 3186.684474 | 3096.513187 | 3158.797161 | 2916.198639 | 2803.238191 | 2632.32098 | 0.007014 | -0.19228 | 0.003435 | -2.92589 | 0.875222 | 2.154027 |
| CNAG_055 | JJJ1 | DnaJ-like | 2407.760679 | 2379.991518 | 2241.678211 | 1991.421828 | 2040.710731 | 2141.538801 | 0.007063 | -0.20234 | 0.003463 | -2.92336 | 0.869141 | 2.151034 |
| CNAG_079 | - | hypothetic | 570.7354252 | 625.7611328 | 443.6981102 | 478.6217629 | 359.3610433 | 405.3776848 | 0.007112 | -0.4135 | 0.00349 | -2.92089 | 0.7508 | 2.148015 |
| CNAG_033 | - | hypothetic | 608.1270955 | 615.0778681 | 505.5200857 | 487.0641973 | 461.3164937 | 433.1400889 | 0.007139 | -0.33842 | 0.003507 | -2.9194 | 0.79091 | 2.146381 |
| CNAG_027 | - | hypothetic | 1008.209442 | 1386.662228 | 1322.437883 | 1515.517296 | 1502.683873 | 1586.270588 | 0.007139 | 0.294867 | 0.003507 | 2.919415 | 1.226772 | 2.146381 |
| CNAG_075 | - | hypothetic | 22.76520248 | 34.64818251 | 40.62234751 | 98.95160144 | 47.36166193 | 58.7247082 | 0.007159 | 1.066545 | 0.003518 | 2.918433 | 2.094412 | 2.145137 |
| CNAG_031 | CHO2 | Phosphati | 1826.50245 | 1881.240088 | 1787.443099 | 2074.233581 | 2191.042957 | 2100.365377 | 0.007273 | 0.197156 | 0.003583 | 2.912738 | 1.146436 | 2.138289 |
| CNAG_032 | - | 28 kda go | 1070.864184 | 1219.564324 | 1230.603784 | 966.3852588 | 998.7287027 | 1024.747127 | 0.007295 | -0.25077 | 0.003595 | -2.91165 | 0.840448 | 2.136988 |
| CNAG_058 | - | CDK-activ | 669.0493615 | 712.0972624 | 704.8598031 | 611.6383017 | 557.042989 | 557.4373909 | 0.007305 | -0.28844 | 0.003601 | -2.91113 | 0.818789 | 2.136382 |
| CNAG_052 | - | hypothetic | 125.2431931 | 131.5570116 | 125.9557159 | 176.1341153 | 185.6773379 | 190.7340205 | 0.007312 | 0.517227 | 0.003605 | 2.910768 | 1.431202 | 2.135981 |
| CNAG_056 | - | hypothetic | 835.0490119 | 764.006937 | 806.3012906 | 991.7874639 | 976.5474738 | 939.1399681 | 0.007528 | 0.257372 | 0.00372 | 2.900955 | 1.195299 | 2.123328 |
| CNAG_036 | - | hypothetic | 107.6301298 | 94.96794807 | 93.98408935 | 147.4535878 | 148.5924402 | 149.6787715 | 0.007693 | 0.579159 | 0.00381 | 2.89351 | 1.493978 | 2.113878 |
| CNAG_034 | - | hypothetic | 1256.299763 | 1164.675432 | 1178.397924 | 996.7489326 | 1011.652204 | 1070.245468 | 0.007722 | -0.24114 | 0.003825 | -2.89228 | 0.846079 | 2.112284 |
| CNAG_036 | - | hypothetic | 761.1332705 | 669.0812729 | 740.0444291 | 619.3709336 | 608.8556559 | 559.6870677 | 0.00774 | -0.29684 | 0.003834 | -2.89147 | 0.814035 | 2.111268 |
| CNAG_034 | NDX3 | cleavage a | 1.425214049 | 3.331694782 | 0.085194742 | 18.15212638 | 5.963632561 | 13.07953741 | 0.007987 | 3.159501 | 0.003965 | 2.880941 | 8.935203 | 2.097598 |
| CNAG_123 | - | unspecifie | 13.44435071 | 36.67163589 | 22.56825553 | 59.97433578 | 58.25388802 | 42.14110539 | 0.00804 | 1.150876 | 0.003995 | 2.878552 | 2.220487 | 2.094734 |
| CNAG_050 | - | hypothetic | 285.4026319 | 412.6610919 | 362.6729637 | 507.9386304 | 452.1013849 | 447.3120342 | 0.008193 | 0.395062 | 0.004074 | 2.872393 | 1.315 | 2.086582 |
| CNAG_078 | - | hypothetic | 0.140649945 | 1.161561372 | 0.058985004 | 9.005305348 | 7.963857874 | 7.634873014 | 0.008225 | 4.404376 | 0.004092 | 2.870983 | 21.17627 | 2.084858 |
| CNAG_051 | - | cation-tra | 3671.087532 | 3229.029622 | 2975.429221 | 2754.133221 | 2989.592517 | 2761.085315 | 0.008378 | -0.23146 | 0.004177 | -2.86446 | 0.851773 | 2.076868 |
| CNAG_027 | - | hypothetic | 12.21886695 | 18.50688984 | 27.79089072 | 55.53412967 | 37.20481202 | 41.99166239 | 0.008511 | 1.207843 | 0.004252 | 2.858832 | 2.30992 | 2.070001 |
| CNAG_077 | - | hypothetic | 35.60711748 | 48.66261226 | 65.09601365 | 76.41006223 | 82.99536175 | 111.5962951 | 0.008622 | 0.847566 | 0.004312 | 2.854418 | 1.799462 | 2.064387 |
| CNAG_036 | - | hypothetic | 13.27770724 | 12.06717691 | 21.40942561 | 38.71354818 | 30.27678656 | 45.07344633 | 0.008639 | 1.281247 | 0.004323 | 2.853564 | 2.43049 | 2.063537 |
| CNAG_052 | - | 20S protea | 3865.222141 | 3816.258416 | 4222.009854 | 3541.081187 | 3568.338313 | 3484.880755 | 0.008698 | -0.18368 | 0.004354 | -2.85131 | 0.880457 | 2.060559 |
| CNAG_057 | STB4 | putative tr | 6.209730561 | 3.432724689 | 11.80188904 | 20.55216318 | 22.71319067 | 27.37050307 | 0.008762 | 1.769627 | 0.00439 | 2.848695 | 3.409657 | 2.057416 |
| CNAG_051 | PMR1 | Ca2 -trans | 815.2794316 | 870.6080952 | 786.0705146 | 923.0529749 | 1118.175384 | 975.7686458 | 0.008935 | 0.272409 | 0.00448 | 2.842211 | 1.207823 | 2.048892 |
| CNAG_127 | - | unspecifie | 6.223725948 | 9.824700619 | 7.604916746 | 25.51866833 | 27.18844818 | 18.80126142 | 0.009185 | 1.65532 | 0.004613 | 2.832855 | 3.14993 | 2.036904 |
| CNAG_008 | - | hypothetic | 420.4091415 | 301.8845346 | 366.9453592 | 495.3540348 | 503.1223452 | 440.6953061 | 0.009313 | 0.386999 | 0.004683 | 2.828037 | 1.30767 | 2.03089 |
| CNAG_120 | - | unspecifie | 8.369022501 | 4.466822692 | 2.284190833 | 20.03380464 | 19.29247551 | 17.58278921 | 0.009378 | 1.983157 | 0.004719 | 2.82559 | 3.953573 | 2.027889 |
| CNAG_124 | - | unspecifie | 183.3304058 | 256.1299837 | 195.1189356 | 157.725025 | 154.2098928 | 138.9969401 | 0.009527 | -0.51163 | 0.004799 | -2.82024 | 0.701431 | 2.021059 |
| CNAG_075 | TRS130 | Trafficking | 2498.870268 | 2528.66868 | 2336.598696 | 2138.558621 | 2149.394888 | 2237.020009 | 0.009806 | -0.1894 | 0.004956 | -2.80987 | 0.876971 | 2.008522 |
| CNAG_027 | RPP1 | hypothetic | 457.9569689 | 446.164594 | 478.8913314 | 570.6851344 | 620.4364761 | 537.289138 | 0.009928 | 0.30698 | 0.005025 | 2.805426 | 1.237116 | 2.003129 |
| CNAG_058 | - | protein CF | 3513.632561 | 3359.409907 | 3399.824074 | 2886.430873 | 2992.113803 | 3230.5777 | 0.009928 | -0.18872 | 0.005022 | -2.8056 | 0.877382 | 2.003129 |
| CNAG_027 | - | hypothetic | 441.474176 | 363.2452529 | 391.4809828 | 491.2497645 | 483.1022239 | 563.7135215 | 0.009981 | 0.347333 | 0.005056 | 2.803455 | 1.272207 | 2.000816 |
| CNAG_075 | - | prenylatec | 54.20832415 | 77.65532486 | 69.43507347 | 96.66476284 | 117.6885484 | 112.9864669 | 0.009982 | 0.686197 | 0.005057 | 2.803362 | 1.609037 | 2.000795 |
| CNAG_035 | - | AP-2 com | 2802.49701 | 2926.219175 | 2609.617594 | 2332.287016 | 2410.743304 | 2592.243519 | 0.010129 | -0.19978 | 0.005143 | -2.79792 | 0.870682 | 1.994418 |
| CNAG_033 | - | hypothetic | 240.9082363 | 294.1841503 | 268.8338257 | 336.3753849 | 336.4946266 | 388.3000223 | 0.010229 | 0.388054 | 0.0052 | 2.794369 | 1.308627 | 1.990163 |
| CNAG_126 | - | unspecifie | 44.55280284 | 83.68033154 | 74.4385326 | 36.89833787 | 35.92872206 | 36.8585872 | 0.010414 | -0.92268 | 0.005299 | -2.78827 | 0.527528 | 1.982389 |
| CNAG_009 | KRE6 | glucosidas | 3336.443735 | 3689.13809 | 3364.725196 | 3887.651286 | 4023.092284 | 3921.033357 | 0.01043 | 0.172726 | 0.00531 | 2.787618 | 1.127186 | 1.98173 |
| CNAG_034 | - | efflux prot | 21.50206057 | 22.83824494 | 36.31541932 | 63.92208675 | 43.16353523 | 57.43118872 | 0.010512 | 1.029759 | 0.005356 | 2.784824 | 2.041683 | 1.978298 |
| CNAG_126 | - | unspecifie | 43.92901679 | 109.8979029 | 87.55939775 | 155.7210711 | 130.4740255 | 123.1130533 | 0.010614 | 0.759362 | 0.00541 | 2.781555 | 1.692742 | 1.974131 |
| CNAG_032 | - | universal s | 877.5574355 | 418.0856435 | 451.0983769 | 295.8645828 | 302.3643599 | 293.4160788 | 0.010614 | -0.99083 | 0.005412 | -2.78145 | 0.503187 | 1.974101 |
| CNAG_078 | - | hypothetic | 825.5554323 | 871.5775415 | 891.4823257 | 744.6661063 | 760.7075241 | 686.2332172 | 0.010677 | -0.25555 | 0.005447 | -2.77932 | 0.837667 | 1.971564 |
| CNAG_122 | - | unspecifie | 0.230678295 | 3.353109391 | 4.354967716 | 11.09130078 | 19.57621888 | 11.01423516 | 0.010724 | 2.536274 | 0.005475 | 2.777646 | 5.800888 | 1.969632 |
| CNAG_032 | INM101 | myo-inosi | 2045.658826 | 1745.45995 | 1915.30373 | 1615.518103 | 1669.826797 | 1663.098462 | 0.010822 | -0.22164 | 0.005528 | -2.77455 | 0.857591 | 1.965709 |
| CNAG_078 | - | hypothetic | 25.8723172 | 15.29947912 | 17.23027141 | 35.99420512 | 46.13032184 | 46.29812697 | 0.011038 | 1.139024 | 0.005649 | 2.767472 | 2.20232 | 1.957094 |
| CNAG_034 | - | hypothetic | 268.903843 | 287.7848414 | 322.1024344 | 404.3420117 | 385.7432386 | 354.0988021 | 0.011135 | 0.366952 | 0.0057 | 2.76456 | 1.289625 | 1.953318 |
| CNAG_032 | CAC1 | Adenylate | 3013.782582 | 2850.847991 | 2705.595182 | 2449.842951 | 2585.028895 | 2565.643197 | 0.011526 | -0.18858 | 0.005921 | -2.75211 | 0.877469 | 1.938323 |
| CNAG_121 | - | unspecifie | 7.207310639 | 3.388204742 | 2.274615252 | 14.2288792 | 13.92005903 | 24.99805421 | 0.011967 | 2.071043 | 0.006166 | 2.738845 | 4.201905 | 1.922008 |
| CNAG_003 | - | hypothetic | 107.3711313 | 134.4699987 | 127.8337366 | 82.92504824 | 93.04759394 | 65.80735897 | 0.011976 | -0.6315 | 0.006171 | -2.73853 | 0.645503 | 1.921696 |
| CNAG_057 | UBP14 | ubiquitin | 1084.528014 | 885.6741096 | 998.1403557 | 774.0705479 | 822.7484856 | 881.5078223 | 0.012171 | -0.27647 | 0.006288 | -2.73235 | 0.825611 | 1.914672 |
| CNAG_073 | - | hypothetic | 6.35309728 | 18.40851576 | 13.99981527 | 33.20096051 | 44.72602208 | 22.24391106 | 0.012257 | 1.406737 | 0.006337 | 2.729798 | 2.651368 | 1.911615 |
| CNAG_027 | - | peroxin-13 | 229.1012809 | 210.2492337 | 245.336245 | 302.2479792 | 317.6681337 | 288.562178 | 0.012294 | 0.39135 | 0.006358 | 2.728728 | 1.31162 | 1.910309 |
| CNAG_003 | - | chromatin | 2268.049193 | 2576.183909 | 2664.01634 | 2761.399986 | 2961.2905 | 3007.312264 | 0.012431 | 0.202557 | 0.00643 | 2.725005 | 1.150736 | 1.90551 |
| CNAG_058 | - | hypothetic | 921.40248 | 126.3929826 | 252.7803618 | 99.30037592 | 89.86341739 | 97.13024212 | 0.012463 | -2.21158 | 0.00645 | -2.72399 | 0.215898 | 1.904378 |
| CNAG_027 | - | sorting ne | 413.6217898 | 462.2562768 | 405.3567005 | 508.3181462 | 508.9714088 | 598.1257096 | 0.012633 | 0.319971 | 0.006541 | 2.719367 | 1.248305 | 1.898511 |

| | | | | | | | | | | | | | | |
|---|---|---|---|---|---|---|---|---|---|---|---|---|---|---|
| CNAG_057 | - | hypothetic | 1.363983309 | 1.172321606 | 0.065918203 | 8.040881213 | 9.754138872 | 9.786600956 | 0.012664 | 3.685758 | 0.00656 | 2.718392 | 12.86837 | 1.897436 |
| CNAG_122 | - | unspecifie | 13.42365954 | 21.74296774 | 33.11203376 | 54.39941728 | 68.08752208 | 30.11364328 | 0.012978 | 1.166158 | 0.006738 | 2.709516 | 2.244132 | 1.886781 |
| CNAG_008 | - | hypothetic | 2.601143298 | 2.284036419 | 3.299028063 | 17.13871377 | 13.19496246 | 8.864078127 | 0.013178 | 2.424781 | 0.006849 | 2.704116 | 5.369474 | 1.880157 |
| CNAG_124 | - | unspecifie | 47.58429465 | 62.13723836 | 41.54756081 | 22.80730535 | 15.93254463 | 35.56676017 | 0.01342 | -1.07177 | 0.006991 | -2.69726 | 0.475735 | 1.872233 |
| CNAG_057 | - | hypothetic | 374.9774684 | 351.3438237 | 329.6523606 | 475.1157202 | 517.0402949 | 378.640475 | 0.014361 | 0.362527 | 0.007513 | 2.673214 | 1.285676 | 1.842815 |
| CNAG_034 | CTS1 | calcineurir | 92.57740929 | 96.02472271 | 98.21855126 | 113.5092979 | 160.8451195 | 158.4486012 | 0.014399 | 0.576419 | 0.007535 | 2.672246 | 1.491143 | 1.841663 |
| CNAG_060 | - | hypothetic | 1146.704469 | 1208.823374 | 1233.823702 | 1073.93693 | 999.5568717 | 1045.849695 | 0.0144 | -0.21763 | 0.007537 | -2.67214 | 0.859978 | 1.841624 |
| CNAG_055 | PRY1 | septum fo | 600.0418322 | 602.197177 | 566.2555932 | 533.8813985 | 421.5107021 | 490.7876333 | 0.014413 | -0.30697 | 0.007547 | -2.67169 | 0.80834 | 1.841246 |
| CNAG_055 | - | septum-pı | 941.35414 | 842.6887179 | 1007.814927 | 1146.664854 | 1054.076208 | 1157.765193 | 0.014515 | 0.251225 | 0.007608 | 2.669001 | 1.190217 | 1.838175 |
| CNAG_037 | - | hypothetic | 196.7083784 | 271.5956834 | 307.1384237 | 366.2135978 | 330.461374 | 352.8725063 | 0.014619 | 0.422928 | 0.007666 | 2.666459 | 1.340646 | 1.835091 |
| CNAG_123 | - | unspecifie | 7.267112446 | 7.657170475 | 2.289009764 | 24.19726617 | 16.54059055 | 17.60770441 | 0.014665 | 1.818601 | 0.007693 | 2.665249 | 3.52739 | 1.83373 |
| CNAG_031 | - | FAD depe | 28.22894423 | 58.89836109 | 54.144889 | 25.2965625 | 25.64679127 | 16.97035981 | 0.014714 | -1.09822 | 0.007726 | -2.6638 | 0.467094 | 1.832259 |
| CNAG_131 | - | unspecifie | 11.97866435 | 7.740391909 | 13.9575073 | 25.21945236 | 26.10272721 | 31.84417873 | 0.014853 | 1.386922 | 0.007812 | 2.660081 | 2.615201 | 1.828172 |
| CNAG_037 | - | hypothetic | 271.2879067 | 340.5048932 | 317.8730489 | 433.9703219 | 395.1906106 | 370.7569917 | 0.014895 | 0.353953 | 0.007839 | 2.658906 | 1.278058 | 1.826958 |
| CNAG_127 | - | unspecifie | 5.1311185 | 14.10142431 | 10.79256458 | 31.94151956 | 26.13979178 | 23.19803479 | 0.014983 | 1.451313 | 0.007893 | 2.656606 | 2.734568 | 1.824398 |
| CNAG_058 | - | Cytochron | 207.0813025 | 282.2817339 | 225.1396412 | 302.1431459 | 316.7325674 | 331.7423934 | 0.015155 | 0.396538 | 0.007992 | 2.652426 | 1.316345 | 1.819449 |
| CNAG_122 | - | unspecifie | 5.117000308 | 8.787628507 | 14.9864954 | 21.18921127 | 25.28260637 | 31.77659738 | 0.015179 | 1.497162 | 0.00801 | 2.651665 | 2.822869 | 1.818743 |
| CNAG_032 | - | hypothetic | 659.880728 | 632.5279603 | 600.5059707 | 774.6220375 | 807.5608976 | 709.3839646 | 0.015684 | 0.260716 | 0.008297 | 2.639747 | 1.198073 | 1.804547 |
| CNAG_126 | - | unspecifie | 169.4108367 | 232.5113319 | 219.5520972 | 150.6707185 | 129.9187831 | 167.7006144 | 0.015781 | -0.48905 | 0.008352 | -2.6375 | 0.712493 | 1.80187 |
| CNAG_127 | - | unspecifie | 2.650180135 | 7.604915926 | 1.198446113 | 14.34524152 | 10.35841481 | 22.79848303 | 0.015944 | 2.191996 | 0.00845 | 2.633548 | 4.569372 | 1.797413 |
| CNAG_031 | FZC24 | nuclear pı | 1572.896262 | 1580.558984 | 1652.940955 | 1303.337778 | 1389.475544 | 1505.384178 | 0.01606 | -0.21008 | 0.008516 | -2.63091 | 0.864487 | 1.794246 |
| CNAG_025 | - | hypothetic | 8.495610644 | 7.70429452 | 8.658879791 | 17.99380724 | 34.40972442 | 18.80437395 | 0.016107 | 1.568295 | 0.008542 | 2.629845 | 2.965541 | 1.79298 |
| CNAG_051 | - | GABA pen | 228.0691649 | 133.6787419 | 174.8030435 | 102.1339433 | 143.1219338 | 116.7785166 | 0.016403 | -0.5853 | 0.008713 | -2.62311 | 0.666509 | 1.785089 |
| CNAG_003 | APT3 | Aminophc | 3007.944262 | 2924.090662 | 2616.033748 | 2514.495565 | 2500.458228 | 2580.067797 | 0.016829 | -0.18574 | 0.008962 | -2.6135 | 0.879199 | 1.773951 |
| CNAG_052 | - | hypothetic | 420.5268134 | 367.5486384 | 421.3019716 | 546.7771433 | 523.7633258 | 454.0691254 | 0.016983 | 0.318346 | 0.009048 | 2.610228 | 1.2469 | 1.769993 |
| CNAG_078 | - | hypothetic | 75.1537513 | 108.8253803 | 69.51013695 | 143.8269708 | 148.8027203 | 103.2982477 | 0.017133 | 0.630835 | 0.009134 | 2.606981 | 1.548461 | 1.766171 |
| CNAG_057 | - | acyl-prote | 1143.232721 | 1235.753891 | 1261.54668 | 1084.041539 | 1070.349955 | 1020.354771 | 0.01743 | -0.21229 | 0.009304 | -2.60069 | 0.863167 | 1.758702 |
| CNAG_031 | - | hypothetic | 3351.826113 | 4434.702441 | 3138.750674 | 5571.629596 | 5132.480566 | 4440.629277 | 0.017698 | 0.457612 | 0.009455 | 2.595137 | 1.373267 | 1.752071 |
| CNAG_122 | - | unspecifie | 19.1328976 | 22.79559753 | 29.9318155 | 45.53758896 | 41.54132036 | 54.0241795 | 0.017865 | 0.985186 | 0.009551 | 2.59168 | 1.979568 | 1.748002 |
| CNAG_057 | RIB4 | 6,7-dimetl | 190.711017 | 193.0243617 | 254.8786042 | 251.371786 | 323.8329048 | 291.8128781 | 0.018446 | 0.425689 | 0.009893 | 2.579549 | 1.343214 | 1.734105 |
| CNAG_127 | - | unspecifie | 1.310753354 | 0.058166456 | 0.050248287 | 8.253229814 | 4.337013421 | 7.588904414 | 0.01849 | 4.129429 | 0.009919 | 2.578641 | 17.50177 | 1.733063 |
| CNAG_051 | HOB3 | hypothetic | 456.3159593 | 395.4092074 | 442.4925226 | 375.8325816 | 363.0453308 | 292.225448 | 0.018576 | -0.34414 | 0.009972 | -2.57681 | 0.787775 | 1.731058 |
| CNAG_058 | - | hypothetic | 2.6048389 | 3.349339101 | 3.300726543 | 8.565456464 | 16.84676024 | 14.1979779 | 0.018668 | 2.233046 | 0.010028 | 2.574848 | 4.701256 | 1.728891 |
| CNAG_078 | - | insulysin | 1322.895797 | 1265.942948 | 1228.540523 | 1153.335301 | 1090.151059 | 1099.152521 | 0.018719 | -0.20631 | 0.010058 | -2.57383 | 0.866754 | 1.727717 |
| CNAG_036 | - | hypothetic | 216.3010035 | 219.9105894 | 238.9405902 | 325.0603534 | 292.5580783 | 267.5293995 | 0.01888 | 0.378112 | 0.010147 | 2.570787 | 1.29964 | 1.723993 |
| CNAG_127 | - | unspecifie | 27.30011435 | 42.07363305 | 26.84149484 | 72.54101705 | 47.45510908 | 59.72695783 | 0.018976 | 0.892836 | 0.010213 | 2.568519 | 1.856823 | 1.721803 |
| CNAG_067 | - | kinesin | 2580.665921 | 2678.435335 | 2523.217885 | 2360.445965 | 2234.760829 | 2395.76592 | 0.019117 | -0.1698 | 0.010293 | -2.56583 | 0.888968 | 1.718575 |
| CNAG_056 | - | solute car | 162.735689 | 198.3386533 | 211.2025996 | 306.6133115 | 250.2689519 | 225.3746814 | 0.019228 | 0.437102 | 0.010357 | 2.563664 | 1.353882 | 1.716056 |
| CNAG_127 | - | unspecifie | 179.2262612 | 133.5743204 | 137.5309237 | 101.5654032 | 94.58110958 | 117.7551814 | 0.019848 | -0.54121 | 0.010696 | -2.55249 | 0.687193 | 1.702294 |
| CNAG_055 | - | hypothetic | 101.9530093 | 103.5922679 | 130.1374538 | 155.9302484 | 157.1657135 | 166.3137585 | 0.020144 | 0.501807 | 0.01087 | 2.546836 | 1.415986 | 1.695846 |
| CNAG_031 | - | hypothetic | 54.00031565 | 56.14122374 | 44.96240984 | 108.4411261 | 82.13746589 | 67.74676151 | 0.020335 | 0.723867 | 0.010981 | 2.543313 | 1.651603 | 1.691759 |
| CNAG_034 | - | hypothetic | 712.5627178 | 798.409812 | 759.3724496 | 863.2003664 | 936.9111663 | 891.3842364 | 0.020355 | 0.23066 | 0.010997 | 2.542809 | 1.173372 | 1.691329 |
| CNAG_067 | - | hypothetic | 1538.001412 | 1485.836923 | 1425.921389 | 1714.721854 | 1674.287978 | 1726.218751 | 0.020355 | 0.186022 | 0.010994 | 2.542879 | 1.137622 | 1.691329 |
| CNAG_034 | - | 1-alkyl-2-a | 233.8469143 | 235.0146145 | 280.4988055 | 360.0896315 | 321.0109458 | 296.3922232 | 0.020856 | 0.369881 | 0.011285 | 2.533741 | 1.292246 | 1.680761 |
| CNAG_033 | URK1 | Uridine kir | 1941.745779 | 1817.58161 | 1679.677368 | 1476.117529 | 1683.69482 | 1600.923875 | 0.021187 | -0.20768 | 0.01148 | -2.52775 | 0.865931 | 1.67393 |
| CNAG_051 | GAT5 | hypothetic | 414.6905569 | 443.9190744 | 340.3514478 | 482.6755282 | 504.7484223 | 514.9698997 | 0.021288 | 0.310831 | 0.01155 | 2.525603 | 1.240422 | 1.671864 |
| CNAG_033 | - | N-acetyltr | 649.192925 | 638.8944174 | 711.2269786 | 568.7534898 | 538.0648085 | 582.8982753 | 0.021393 | -0.25856 | 0.011609 | -2.5238 | 0.835924 | 1.669734 |
| CNAG_050 | PIP201 | hypothetic | 818.7655955 | 703.732049 | 929.9370293 | 932.4322304 | 1065.490521 | 995.7181216 | 0.021512 | 0.272103 | 0.011677 | 2.521746 | 1.207567 | 1.667319 |
| CNAG_077 | CLR6 | transcripti | 1138.812037 | 1604.148721 | 1644.323184 | 1198.323351 | 1255.794018 | 1195.766343 | 0.02158 | -0.27992 | 0.011724 | -2.52033 | 0.823636 | 1.665939 |
| CNAG_031 | - | hypothetic | 750.7387986 | 800.4183665 | 690.0026208 | 644.2597346 | 639.9101564 | 626.2393777 | 0.021593 | -0.24555 | 0.011734 | -2.52005 | 0.843496 | 1.665693 |
| CNAG_051 | - | hypothetic | 652.598786 | 633.4746665 | 614.2508689 | 529.0165772 | 530.3411487 | 552.9275362 | 0.022127 | -0.25254 | 0.012046 | -2.5108 | 0.839415 | 1.655078 |
| CNAG_051 | - | homoserir | 3071.310935 | 3583.47718 | 3849.787188 | 3163.64804 | 3160.359202 | 2907.644823 | 0.022178 | -0.20128 | 0.012076 | -2.50992 | 0.869781 | 1.654086 |
| CNAG_058 | - | hypothetic | 212.3120033 | 226.1046883 | 194.0684138 | 142.7545313 | 183.7097249 | 147.8486673 | 0.022247 | -0.4363 | 0.012117 | -2.50873 | 0.739025 | 1.652726 |
| CNAG_031 | - | hypothetic | 6.131860124 | 6.594520932 | 5.459558347 | 24.22875496 | 11.14160696 | 20.80821812 | 0.022608 | 1.679619 | 0.012344 | 2.502164 | 3.203433 | 1.645729 |
| CNAG_077 | - | hypothetic | 48.42538883 | 85.13360803 | 67.30591525 | 107.8484903 | 98.47843681 | 110.766694 | 0.022788 | 0.642197 | 0.01245 | 2.499117 | 1.560704 | 1.642284 |
| CNAG_077 | CIN1 | Intersectir | 1882.421358 | 1734.727905 | 1549.671437 | 2069.6899 | 2140.200872 | 1855.098123 | 0.02315 | 0.215871 | 0.012656 | 2.493302 | 1.161405 | 1.635455 |
| CNAG_052 | - | chaperone | 1108.157932 | 1189.408898 | 1142.135281 | 1067.003947 | 996.1506796 | 939.3370761 | 0.023227 | -0.21105 | 0.012707 | -2.49188 | 0.863909 | 1.634002 |
| CNAG_009 | - | membrane | 288.7326977 | 359.858739 | 290.1911064 | 412.8259861 | 378.7408853 | 398.4274597 | 0.02345 | 0.328225 | 0.012838 | 2.48824 | 1.255467 | 1.62985 |
| CNAG_052 | - | DNA misn | 196.4233324 | 211.2449044 | 191.0041102 | 284.5384721 | 230.284426 | 276.2401325 | 0.023516 | 0.38707 | 0.01288 | 2.487082 | 1.307735 | 1.628631 |
| CNAG_033 | - | hypothetic | 13.18866827 | 16.2762182 | 11.88403989 | 34.91695869 | 34.0196689 | 24.39025692 | 0.023567 | 1.186949 | 0.01291 | 2.486236 | 2.276707 | 1.627694 |
| CNAG_079 | - | hypothetic | 628.2032103 | 641.0343019 | 718.6717439 | 607.0541199 | 569.200553 | 467.6177732 | 0.023586 | -0.28976 | 0.012924 | -2.48587 | 0.81804 | 1.627344 |
| CNAG_009 | - | protein tra | 372.4154628 | 404.99888 | 379.6054749 | 350.2251794 | 282.6735424 | 298.7853492 | 0.023705 | -0.32735 | 0.012995 | -2.48392 | 0.796999 | 1.625154 |
| CNAG_032 | - | hypothetic | 2684.646946 | 2466.298561 | 2110.643137 | 3100.306009 | 2854.315811 | 2612.228813 | 0.02412 | 0.223164 | 0.013257 | 2.476781 | 1.167291 | 1.617622 |
| CNAG_027 | - | UTP-gluco | 13095.24279 | 12799.94603 | 12803.88093 | 14237.3572 | 14305.63996 | 14087.90932 | 0.024255 | 0.124244 | 0.013335 | 2.474702 | 1.089937 | 1.61519 |
| CNAG_057 | - | multidrug | 234.9289173 | 241.4257869 | 232.5755396 | 308.4849708 | 280.370163 | 321.7548427 | 0.024439 | 0.347644 | 0.013447 | 2.471697 | 1.272481 | 1.611924 |
| CNAG_037 | - | tubulin alı | 8493.020656 | 8398.575803 | 8810.995647 | 7675.334709 | 8089.063652 | 7879.977694 | 0.02445 | -0.13583 | 0.013457 | -2.47145 | 0.910147 | 1.611716 |
| CNAG_057 | - | 2-dehydrc | 331.7680152 | 308.2501839 | 314.6827333 | 405.7594408 | 389.1198205 | 395.1157455 | 0.024642 | 0.303761 | 0.013571 | 2.468412 | 1.234358 | 1.608321 |
| CNAG_026 | - | hypothetic | 285.1740151 | 286.7059547 | 310.3891575 | 430.9979181 | 364.1021504 | 333.064292 | 0.024835 | 0.339935 | 0.013696 | 2.465141 | 1.265699 | 1.604931 |

| | | | | | | | | | | | | | | |
|---|---|---|---|---|---|---|---|---|---|---|---|---|---|---|
| CNAG_122 | - | unspecifie | 10.98747064 | 24.81616204 | 14.02673025 | 29.71082086 | 45.45223757 | 33.15602868 | 0.025238 | 1.143679 | 0.013937 | 2.458895 | 2.209437 | 1.597944 |
| CNAG_051 | CAN2 | Beta-type | 2892.338285 | 2676.322639 | 3329.27726 | 1853.2811 | 2654.986214 | 2128.375119 | 0.025548 | -0.43957 | 0.014117 | -2.45427 | 0.737354 | 1.592638 |
| CNAG_122 | - | unspecifie | 14.54003622 | 29.17103082 | 25.7020221 | 35.02185575 | 50.44320946 | 51.82170311 | 0.025568 | 0.98668 | 0.014138 | 2.453749 | 1.98162 | 1.592301 |
| CNAG_068 | - | Methylene | 1181.774405 | 1304.694452 | 1283.956256 | 1119.834232 | 1162.702998 | 1017.060597 | 0.025612 | -0.20725 | 0.014165 | -2.45305 | 0.866189 | 1.591557 |
| CNAG_127 | - | unspecifie | 2.576458377 | 2.272684363 | 3.287646668 | 7.800854452 | 10.50787049 | 17.32741997 | 0.026399 | 2.265225 | 0.014643 | 2.44108 | 4.807294 | 1.57842 |
| CNAG_059 | - | hypothetic | 838.6780699 | 990.1523136 | 845.7849769 | 1070.354575 | 1085.250385 | 998.0145209 | 0.026497 | 0.223701 | 0.014709 | 2.439458 | 1.167725 | 1.576806 |
| CNAG_009 | - | glucan 1,3 | 2.661889574 | 7.614469834 | 2.268328682 | 11.75816111 | 5.836639332 | 32.39318074 | 0.026828 | 2.11695 | 0.014903 | 2.434724 | 4.33776 | 1.571407 |
| CNAG_051 | - | hypothetic | 180.1972132 | 206.9606102 | 228.2481727 | 275.0577153 | 270.152982 | 258.5820173 | 0.026865 | 0.372168 | 0.014927 | 2.434145 | 1.294296 | 1.570808 |
| CNAG_055 | - | serine/thr | 831.1212232 | 702.482431 | 716.6404246 | 647.3783944 | 609.6621098 | 652.8453785 | 0.027069 | -0.25299 | 0.015057 | -2.43101 | 0.839156 | 1.567529 |
| CNAG_037 | LYSO1 | lysophosp | 384.4218826 | 450.3674939 | 366.9749174 | 482.6933245 | 501.2992735 | 495.0261545 | 0.027416 | 0.286103 | 0.015253 | 2.426319 | 1.219342 | 1.562001 |
| CNAG_052 | - | telomere | 1203.772569 | 1162.496349 | 1081.387516 | 980.4331506 | 999.599711 | 1043.5952 | 0.02745 | -0.20466 | 0.015275 | -2.42579 | 0.867743 | 1.561463 |
| CNAG_033 | - | hypothetic | 138.2185762 | 204.6808254 | 146.2505022 | 227.5968859 | 253.112845 | 194.2871833 | 0.027478 | 0.451985 | 0.015298 | 2.425255 | 1.367921 | 1.561015 |
| CNAG_077 | PIK1 | 1-phospha | 1738.572705 | 1537.500857 | 1604.98748 | 1468.522634 | 1434.340633 | 1421.094051 | 0.027798 | -0.1903 | 0.015486 | -2.42081 | 0.876424 | 1.555981 |
| CNAG_034 | - | hypothetic | 478.7365053 | 497.7235685 | 534.1982845 | 422.4115203 | 411.2872313 | 431.9307577 | 0.02787 | -0.27073 | 0.015533 | -2.41971 | 0.8289 | 1.554859 |
| CNAG_031 | - | calcium-bi | 728.8457805 | 670.292598 | 808.3758532 | 902.2803718 | 850.5470451 | 879.1592478 | 0.027901 | 0.238517 | 0.015557 | 2.419153 | 1.179779 | 1.554388 |
| CNAG_051 | - | DNA poly | 1090.671454 | 1196.946943 | 1151.728668 | 1022.54888 | 1056.602187 | 934.9022194 | 0.028197 | -0.20545 | 0.015746 | -2.41475 | 0.867269 | 1.549798 |
| CNAG_025 | - | Rab small | 2546.55823 | 2101.013663 | 2217.138437 | 2141.058756 | 2036.429882 | 1799.748062 | 0.028435 | -0.21608 | 0.015896 | -2.41129 | 0.860904 | 1.546146 |
| CNAG_026 | - | early grov | 33.01599368 | 13.23764118 | 46.92008724 | 62.84800675 | 49.22337398 | 69.53415008 | 0.028644 | 0.950419 | 0.016023 | 2.408381 | 1.932433 | 1.542974 |
| CNAG_025 | - | 2,4-dihydr | 77.35623303 | 53.06058284 | 99.17506955 | 119.4963329 | 119.2889193 | 114.1942467 | 0.029073 | 0.603368 | 0.016278 | 2.402622 | 1.51926 | 1.536507 |
| CNAG_034 | - | ATP-bindi | 3326.790205 | 3264.573581 | 3039.401024 | 2941.837855 | 3014.62142 | 2743.326284 | 0.029246 | -0.16206 | 0.016382 | -2.40028 | 0.893746 | 1.533935 |
| CNAG_009 | - | hypothetic | 217.9098472 | 147.6922259 | 215.2511164 | 121.6638657 | 144.7397233 | 155.497824 | 0.029736 | -0.48004 | 0.016686 | -2.39356 | 0.716957 | 1.526712 |
| CNAG_032 | - | hypothetic | 557.1949289 | 525.9046434 | 610.0282264 | 677.2908475 | 701.4259336 | 653.8375357 | 0.029818 | 0.247782 | 0.016735 | 2.392481 | 1.18738 | 1.525526 |
| CNAG_051 | - | hypothetic | 34.99500417 | 15.31433733 | 16.1872962 | 44.85468484 | 43.38926812 | 44.18443783 | 0.030317 | 0.993365 | 0.01706 | 2.38542 | 1.990823 | 1.518312 |
| CNAG_032 | YOX101 | specific tr | 4.845321532 | 2.280375747 | 2.241322703 | 12.01520306 | 15.96267775 | 8.853865803 | 0.030803 | 2.111814 | 0.017348 | 2.379247 | 4.322344 | 1.511408 |
| CNAG_008 | - | hypothetic | 19.94309347 | 13.08224745 | 5.535694118 | 28.35987316 | 42.12962756 | 21.1236979 | 0.031166 | 1.281405 | 0.017564 | 2.37468 | 2.430756 | 1.506313 |
| CNAG_126 | - | unspecifie | 7.394959593 | 9.84457548 | 13.93097053 | 22.05045571 | 19.01702917 | 32.83239516 | 0.031507 | 1.31743 | 0.017787 | 2.370024 | 2.492217 | 1.501592 |
| CNAG_009 | - | transcripti | 2125.093541 | 1911.354985 | 1923.868775 | 1595.16315 | 1873.473055 | 1796.292675 | 0.032044 | -0.19469 | 0.018106 | -2.36345 | 0.873759 | 1.494252 |
| CNAG_032 | - | glutaryl-C | 910.6630856 | 844.6847277 | 854.2023606 | 735.2612192 | 783.1439843 | 754.9949962 | 0.032166 | -0.21456 | 0.018179 | -2.36196 | 0.861812 | 1.492599 |
| CNAG_078 | - | hypothetic | 1002.265786 | 1153.997963 | 1294.660437 | 1439.362556 | 1401.808131 | 1250.056645 | 0.032316 | 0.230887 | 0.018271 | 2.360072 | 1.173556 | 1.490578 |
| CNAG_050 | - | hypothetic | 1.377962516 | 1.178823669 | 2.209939726 | 11.45702372 | 11.56202078 | 4.494800317 | 0.032489 | 2.683898 | 0.018381 | 2.357858 | 6.425897 | 1.488269 |
| CNAG_121 | - | unspecifie | 6.202541717 | 7.691640161 | 11.79739254 | 23.09258808 | 16.4247459 | 25.2122627 | 0.032607 | 1.400298 | 0.018464 | 2.356186 | 2.639561 | 1.486688 |
| CNAG_067 | - | hypothetic | 0.189051731 | 3.318870426 | 2.223598338 | 18.27695047 | 7.820479356 | 5.597064712 | 0.032637 | 2.573828 | 0.018485 | 2.355761 | 5.953873 | 1.486285 |
| CNAG_058 | - | hypothetic | 144.6301879 | 91.76716032 | 72.7481597 | 162.395144 | 149.4110883 | 148.6183522 | 0.03269 | 0.563554 | 0.018523 | 2.355002 | 1.477906 | 1.485586 |
| CNAG_122 | - | unspecifie | 2.674121485 | 8.679021829 | 4.387209649 | 19.33523357 | 14.83605037 | 14.31924564 | 0.03308 | 1.735747 | 0.018768 | 2.350112 | 3.33052 | 1.480435 |
| CNAG_034 | - | hypothetic | 383.1498776 | 380.3971592 | 349.8957924 | 453.9378397 | 431.3278034 | 478.318992 | 0.03327 | 0.277529 | 0.018896 | 2.347581 | 1.212117 | 1.477951 |
| CNAG_057 | - | hypothetic | 728.7339761 | 873.6752946 | 811.5215361 | 722.1596971 | 695.1188667 | 659.5689211 | 0.033649 | -0.23099 | 0.019124 | -2.34311 | 0.852048 | 1.473028 |
| CNAG_057 | TPM1 | tropomyo | 5666.051467 | 5551.00054 | 5951.892277 | 6621.108385 | 6332.11372 | 6166.932208 | 0.034079 | 0.139849 | 0.019388 | 2.337977 | 1.101789 | 1.467519 |
| CNAG_067 | FZC49 | hypothetic | 412.5352093 | 498.8791579 | 479.9510562 | 519.904612 | 596.2350198 | 574.9448138 | 0.034476 | 0.267449 | 0.019631 | 2.33332 | 1.203677 | 1.462484 |
| CNAG_075 | - | hypothetic | 78.24246879 | 112.8421299 | 89.50526817 | 65.08033308 | 54.01733565 | 66.73696847 | 0.034728 | -0.60628 | 0.019789 | -2.33033 | 0.656888 | 1.459318 |
| CNAG_036 | - | hypothetic | 949.3683046 | 803.8730453 | 813.790473 | 971.3766318 | 995.4742776 | 1065.608231 | 0.034835 | 0.225055 | 0.019857 | 2.329041 | 1.168822 | 1.457988 |
| CNAG_033 | - | hypothetic | 1839.068843 | 1688.340851 | 1805.463155 | 1531.462767 | 1673.348853 | 1563.192381 | 0.034902 | -0.17735 | 0.019908 | -2.32807 | 0.884328 | 1.457144 |
| CNAG_050 | - | hypothetic | 11.17306238 | 42.04537762 | 34.22520091 | 54.87237344 | 74.02354335 | 43.31110988 | 0.035676 | 0.962538 | 0.02038 | 2.319278 | 1.948735 | 1.447625 |
| CNAG_125 | - | unspecifie | 22.53922565 | 49.35188418 | 67.81673437 | 16.44188517 | 32.71576559 | 20.2519768 | 0.037434 | -1.03544 | 0.021453 | -2.29992 | 0.487867 | 1.426734 |
| CNAG_050 | - | GTP-bindi | 5202.2853 | 5108.098343 | 5333.295753 | 4960.003751 | 4773.337702 | 4678.299126 | 0.038071 | -0.13373 | 0.021869 | -2.29263 | 0.911471 | 1.419404 |
| CNAG_052 | - | hypothetic | 454.6171945 | 603.3474737 | 533.2895797 | 672.7448934 | 641.8602294 | 613.8789237 | 0.038223 | 0.262417 | 0.02197 | 2.290878 | 1.199486 | 1.417674 |
| CNAG_056 | - | pre-mRN/ | 2176.547092 | 2025.565893 | 2065.700794 | 1898.909987 | 1894.107557 | 1881.790843 | 0.038536 | -0.15897 | 0.022169 | -2.28746 | 0.895664 | 1.414136 |
| CNAG_131 | - | unspecifie | 3.821928874 | 8.681347142 | 4.388564415 | 14.23377915 | 15.73656435 | 18.59858989 | 0.038816 | 1.635587 | 0.02234 | 2.284543 | 3.10714 | 1.410991 |
| CNAG_055 | - | hypothetic | 60.94754854 | 51.8882558 | 60.87766581 | 85.90650521 | 99.59355252 | 78.76465261 | 0.038816 | 0.599444 | 0.022337 | 2.284596 | 1.515133 | 1.410991 |
| CNAG_027 | - | aldose rec | 3175.098413 | 3495.101109 | 3123.645032 | 2830.427718 | 3135.408131 | 2913.140649 | 0.039058 | -0.15638 | 0.022489 | -2.28201 | 0.897275 | 1.408288 |
| CNAG_009 | - | phosphop | 216.8263833 | 187.4224077 | 185.525886 | 142.0626643 | 138.6242797 | 169.8826612 | 0.039742 | -0.40459 | 0.022926 | -2.27467 | 0.755451 | 1.40075 |
| CNAG_025 | - | alfa-L-rha | 734.6837782 | 937.2546132 | 966.0869005 | 694.6897212 | 743.4038989 | 800.4513857 | 0.03981 | -0.2517 | 0.02297 | -2.27393 | 0.839905 | 1.400011 |
| CNAG_025 | CHI21 | Chitinase | 114.7666965 | 115.4338323 | 145.0539954 | 160.4795468 | 176.1902544 | 174.1282881 | 0.039847 | 0.431971 | 0.023001 | 2.273411 | 1.349075 | 1.3996 |
| CNAG_075 | - | mitochon | 735.6841253 | 776.7861246 | 827.4850724 | 690.210117 | 670.9555226 | 676.1742752 | 0.040204 | -0.21465 | 0.023227 | -2.26968 | 0.861754 | 1.395728 |
| CNAG_122 | - | unspecifie | 68.14499927 | 100.1992589 | 71.60098894 | 132.1659906 | 116.6453815 | 103.214294 | 0.040375 | 0.540726 | 0.023336 | 2.267895 | 1.454705 | 1.393887 |
| CNAG_077 | - | amidohyd | 43.54147764 | 43.25508243 | 50.20685888 | 83.96670381 | 80.57935368 | 56.67205891 | 0.040427 | 0.671614 | 0.023371 | 2.267319 | 1.592854 | 1.393325 |
| CNAG_051 | - | ER lumen | 107.7901557 | 115.417822 | 139.7249425 | 175.5523945 | 167.5356699 | 152.0414848 | 0.040951 | 0.437204 | 0.023708 | 2.261823 | 1.353977 | 1.387737 |
| CNAG_058 | - | hypothetic | 935.2088975 | 856.5559181 | 913.9036326 | 792.9250539 | 762.3624091 | 816.0154085 | 0.040994 | -0.20573 | 0.023739 | -2.26133 | 0.867099 | 1.387277 |
| CNAG_009 | - | peptidase | 876.9104689 | 884.5320277 | 907.4992731 | 725.8538315 | 770.1551272 | 837.0661546 | 0.041564 | -0.20881 | 0.024099 | -2.25555 | 0.865249 | 1.381282 |
| CNAG_051 | BWC1 | white coll | 549.9884279 | 485.9986994 | 412.864421 | 584.6663803 | 648.0515005 | 543.975775 | 0.041564 | 0.279976 | 0.024104 | 2.255466 | 1.214175 | 1.381281 |
| CNAG_009 | - | phytanoyl | 1.394088896 | 0.087144259 | 4.313716034 | 8.787268148 | 13.35940021 | 7.714490684 | 0.041649 | 2.466553 | 0.024164 | 2.254517 | 5.527215 | 1.380391 |
| CNAG_055 | - | arf/Sar fa | 672.5816394 | 694.8976423 | 718.7256673 | 626.4291654 | 585.5387463 | 599.5769089 | 0.042276 | -0.21849 | 0.024537 | -2.24861 | 0.859466 | 1.37391 |
| CNAG_009 | - | YjeF famil | 726.2141926 | 725.0543844 | 693.1764434 | 606.8530208 | 613.1608526 | 647.2667803 | 0.042323 | -0.21549 | 0.02457 | -2.2481 | 0.861251 | 1.373423 |
| CNAG_051 | - | glucan 1,3 | 3631.510428 | 3472.511885 | 3100.210974 | 3066.337373 | 3257.032128 | 2828.830749 | 0.042436 | -0.17249 | 0.024646 | -2.2469 | 0.887309 | 1.372261 |
| CNAG_121 | - | unspecifie | 7.160443072 | 4.43497418 | 1.198141117 | 20.32297459 | 15.80987889 | 8.925409007 | 0.042599 | 1.824479 | 0.024761 | 2.245108 | 3.541791 | 1.370605 |
| CNAG_036 | - | hypothetic | 204.416682 | 257.3525058 | 247.3155149 | 172.4052431 | 201.7743946 | 182.2079178 | 0.042675 | -0.36587 | 0.024811 | -2.24433 | 0.775999 | 1.369822 |
| CNAG_009 | - | lipid parti | 162.3586414 | 148.7598694 | 107.9072703 | 189.4003164 | 171.6851974 | 211.7649791 | 0.043064 | 0.438472 | 0.025047 | 2.24067 | 1.355169 | 1.365887 |
| CNAG_075 | - | cytoplasm | 639.856537 | 660.4071881 | 661.1497451 | 564.0896419 | 613.2812295 | 508.6309692 | 0.044192 | -0.23347 | 0.025763 | -2.22977 | 0.850587 | 1.35466 |
| CNAG_026 | - | hypothetic | 1292.594113 | 1262.720316 | 1272.247477 | 1226.60698 | 1065.962265 | 1106.925868 | 0.044205 | -0.18682 | 0.025776 | -2.22957 | 0.878542 | 1.354532 |

| | | | | | | | | | | | | | | |
|---|---|---|---|---|---|---|---|---|---|---|---|---|---|---|
| CNAG_027 | - | hypothetic | 202.422776 | 260.7813949 | 258.1242471 | 264.5235254 | 342.7531876 | 319.5651091 | 0.044738 | 0.346841 | 0.02612 | 2.224428 | 1.271772 | 1.34932 |
| CNAG_051 | - | hypothetic | 1059.389766 | 1243.35664 | 1135.828631 | 1251.50628 | 1374.206346 | 1328.78637 | 0.04538 | 0.187156 | 0.02655 | 2.218064 | 1.138517 | 1.343132 |
| CNAG_076 | - | hypothetic | 11.98382126 | 13.05558404 | 12.90809505 | 25.20496254 | 32.37601077 | 22.14856194 | 0.045435 | 1.118633 | 0.026593 | 2.217438 | 2.171412 | 1.342607 |
| CNAG_100 | - | unspecifie | 13.28589111 | 9.936569964 | 29.83290928 | 48.5536427 | 30.26175336 | 31.01096163 | 0.045544 | 1.078931 | 0.026674 | 2.216261 | 2.112471 | 1.341567 |
| CNAG_031 | - | DIL and A | 1943.06909 | 1944.732269 | 1931.32334 | 1680.892697 | 1751.779279 | 1843.989764 | 0.045861 | -0.15681 | 0.026893 | -2.21307 | 0.897004 | 1.338556 |
| CNAG_050 | CAS91 | maltose C | 935.5806445 | 1026.827783 | 1015.30029 | 1184.783242 | 1156.797732 | 1081.291484 | 0.045878 | 0.186064 | 0.026913 | 2.212775 | 1.137655 | 1.3384 |
| CNAG_037 | CAP4 | beta-1,2-x | 809.0286154 | 785.3709728 | 685.754884 | 705.1046875 | 622.6009274 | 638.4542432 | 0.046083 | -0.229 | 0.027051 | -2.21078 | 0.853228 | 1.336461 |
| CNAG_127 | - | unspecifie | 15.4589673 | 13.09314028 | 16.09998525 | 33.24690052 | 28.64498565 | 29.80272602 | 0.047198 | 1.016775 | 0.027792 | 2.200214 | 2.02339 | 1.32608 |
| CNAG_127 | - | unspecifie | 155.7061422 | 198.2980439 | 199.4668493 | 225.0364616 | 251.2347361 | 239.669145 | 0.047275 | 0.358664 | 0.027843 | 2.199494 | 1.282238 | 1.325373 |
| CNAG_078 | - | syntaxin-b | 630.4392087 | 638.8382763 | 590.8019277 | 542.3330342 | 463.8013285 | 578.3949962 | 0.04784 | -0.2459 | 0.028187 | -2.19467 | 0.843289 | 1.320212 |
| CNAG_068 | - | mitochonc | 1470.07973 | 1341.420457 | 1512.162452 | 1229.473982 | 1235.916945 | 1374.377523 | 0.048029 | -0.18727 | 0.028305 | -2.19304 | 0.878267 | 1.3185 |
| CNAG_071 | - | origin recc | 326.0868092 | 333.0457714 | 404.1778351 | 470.4786533 | 433.1194825 | 406.3000699 | 0.048071 | 0.284896 | 0.028336 | 2.192609 | 1.218322 | 1.318115 |
| CNAG_127 | - | unspecifie | 0.176648753 | 1.184280722 | 4.310824601 | 11.40113931 | 6.940577915 | 10.89064626 | 0.048087 | 2.418468 | 0.028351 | 2.192395 | 5.346029 | 1.317968 |
| CNAG_031 | - | hypothetic | 1533.162093 | 1468.5305 | 1538.851336 | 1307.314661 | 1361.038507 | 1417.699864 | 0.048115 | -0.16746 | 0.028373 | -2.19209 | 0.890412 | 1.317718 |
| CNAG_077 | - | ABC transp | 800.2796759 | 1108.630183 | 661.4233186 | 1136.500747 | 1178.436855 | 1325.227596 | 0.048819 | 0.489415 | 0.028822 | 2.185916 | 1.403875 | 1.311408 |
| CNAG_127 | - | unspecifie | 114.2921806 | 73.44398045 | 55.68737441 | 128.8962025 | 96.52842497 | 146.1391265 | 0.048938 | 0.596429 | 0.028912 | 2.184683 | 1.51197 | 1.310355 |
| CNAG_059 | - | hypothetic | 187.3596195 | 299.5042781 | 266.6644765 | 341.2648831 | 335.7417455 | 297.4991843 | 0.049055 | 0.357999 | 0.028999 | 2.183498 | 1.281647 | 1.309319 |
| CNAG_050 | - | DNA/RNA | 841.8966662 | 787.629345 | 970.3454328 | 749.3179327 | 731.3141342 | 773.837593 | 0.049184 | -0.2212 | 0.029082 | -2.18238 | 0.857854 | 1.308177 |
| CNAG_052 | - | solute car | 1170.074769 | 1215.307693 | 1264.754119 | 1088.692665 | 1033.201173 | 1135.718445 | 0.049459 | -0.17943 | 0.029256 | -2.18002 | 0.88305 | 1.305754 |
| CNAG_036 | - | hypothetic | 1241.30672 | 1428.549277 | 1223.226206 | 1104.951021 | 1079.762657 | 1246.696967 | 0.049711 | -0.19706 | 0.029417 | -2.17785 | 0.872324 | 1.303551 |
