## Supplementary Dataset 3 for "Systematic histone mutagenesis reveals nucleosome-dependent maintenance of three-dimensional chromosome architecture and virulence in *Cryptococcus neoformans*"

Supplementary Dataset 3. Validated chromatin interaction pairs identified by Hi-C analysis.

| | WT | *hht2*Δ |
|---|---|---|
| valid interaction | 30995979 | 27153926 |
| valid interaction rmdup | 30995979 | 27153926 |
| trans interaction | 16806517 | 20190288 |
| cis interaction | 14189462 | 6963638 |
| cis_1kb+ | 1512025 | 1701370 |
| cis_2kb+ | 1434613 | 1668095 |
| cis_4kb+ | 1388041 | 1652091 |
| cis_10kb+ | 1343742 | 1627562 |
| cis_20kb+ | 1303024 | 1596177 |
| cis_40kb+ | 1260264 | 1548167 |
